# An integrative taxonomic treatment of the Mycetophilidae (Diptera: Bibionomorpha) from Singapore reveals 115 new species on 730km^2^

**DOI:** 10.1101/2023.09.02.555672

**Authors:** Dalton de Souza Amorim, Sarah Siqueira Oliveira, Maria Isabel P. A. Balbi, Yuchen Ang, Ambrosio Torres, Darren Yeo, Amrita Srivathsan, Rudolf Meier

**Affiliations:** Departamento de Biologia, FFCLRP, Universidade de São Paulo; Departamento de Ecologia, Instituto de Ciências Biológicas, Universidade Federal de Goiás; Lee Kong Chian Natural History Museum, National University of Singapore; Centre for Wildlife Forensics, National Parks Board Singapore; Center for Integrative Biodiversity Discovery, Leibniz Institute for Evolution and Biodiversity Science, Museum für Naturkunde Berlin

**Keywords:** Mycetophilidae, Integrative Taxonomy, Reverse Taxonomic Workflow, Oriental Insect Fauna, NGS barcodes

## Abstract

“Open ended” or “dark taxa” are species-rich clades that are so abundant and diverse that conventional taxonomic methods tend to struggle with the onslaught of specimens and species. New approaches based on presorting specimens to putative species with low-cost DNA barcodes may make tackling these taxa manageable. However, this will still require limiting the geographic scope of taxonomic revisions, given that most countries and biogeographic regions will have too many specimens and species for comprehensive coverage. We demonstrate the power of this approach by carrying out a revision of the Mycetophilidae fungus gnats (Diptera) of Singapore. The material revised here was obtained from 496 samples collected with 71 Malaise traps placed at 107 sites in different habitats in Singapore: mangroves, swamp forests, freshwater swamps, primary rainforests, and different types of secondary forests (old, maturing, young, urban). Based on molecular and morphological data for 1,454 specimens, we delimit 120 species of 23 genera. Of these, only five were species previously described. The remaining 115 species are new to science and described here. We name, however, only 98 of these species since 14 species are currently only known from females and we cannot prepare a fully satisfying morphological diagnosis (*Manota* spp. A–G and *Neoempheria* spp. A–G), and three species lack molecular data (*Epicypta* sp. A, *Epicypta* sp. B, and *Neoempheria* sp. H). To assess congruence between species delimited with DNA barcodes (3% clusters) and morphology, we determined a match ratio and found it to be overall high (95%) with even higher match ratios (99%) observed for MOTUs clustered at 5% with Objective Clustering and MOTUs obtained with ABGD set to (P=0.060). Overall, the ratio of undescribed to described is an astonishing 20:1. Only revising the Singapore fauna increases the number of described species of Oriental Mycetophilidae by about 25%, highlighting the size of the taxonomic impediment for fungus gnats. Most of the Singapore Mycetophilidae diversity belongs to three genera—*Neoempheria* Osten-Sacken (31 species), *Epicypta* Winnertz (29 species) and *Manota* Williston (14 species), but we also describe a new genus— *Integricypta*, **gen. n.**, which is the putative sistergroup of *Aspidionia* Colless—belonging to the Mycetophilinae Mycetophilini based on three species. The species sequenced, illustrated, and named are: *Leptomorphus rafflesi*, **sp. n.**; *Monoclona simhapura*, **sp. n.**; *Azana demeijeri*, **sp. n.**; *Azana leekongchiani*, **sp. n.** (**Sciophilinae**); *Tetragoneura crawfurdi*, **sp. n.**; *Tetragoneura chola*, **sp. n.**; *Tetragoneura dayuan*, **sp. n.**; *Tetragoneura farquhari*, **sp. n.**; *Ectrepesthoneura johor*, **sp. n.** (**Tetragoneurinae**); *Mohelia zubirsaidi*, **sp. n.**; *Allactoneura tumasik*, **sp. n.**; *Allactoneura limbosengi*, **sp. n.**; *Manota banzu*, **sp. n.**; *Manota tantocksengi*, **sp. n.**; *Manota bukittimah*, **sp. n.**; *Manota chiamassie*, **sp. n.**; *Manota danmaxi*, **sp. n.**; *Manota mahuan*, **sp. n.**; *Manota temenggong*, **sp. n.**; *Clastobasis sritribuana*, **sp. n.**; *Clastobasis bugis*, **sp. n.**; *Clastobasis oranglaut*, **sp. n.** (**Leiinae**); *Parempheriella mait*, **sp. n.**; *Parempheriella longyamen*, **sp. n.**; *Parempheriella peranakan*, **sp. n.**; *Mycomya sachmatich*, **sp. n.**; *Neoempheria merlio*, **sp. n.**; *Neoempheria sabana*, **sp. n.**; *Neoempheria sangabo*, **sp. n.**; *Neoempheria shicheng*, **sp. n.**; *Neoempheria ujong*, **sp. n.**; *Neoempheria subaraji*, **sp. n.**; *Neoempheria kokoiyeeae*, **sp. n.**; *Neoempheria mandai*, **sp. n.**; *Neoempheria malacca*, **sp. n.**; *Neoempheria sinkapho*, **sp. n.**; *Neoempheria singapura*, **sp. n.**; *Neoempheria xinjiapo*, **sp. n.**; *Neoempheria puluochung*, **sp. n.**; *Neoempheria merdeka*, **sp. n.**; *Neoempheria neesoon*, **sp. n.**; *Neoempheria pulau*, **sp. n.**; *Neoempheria cinkappur*, **sp. n.**; *Neoempheria temasek*, **sp. n.**; *Neoempheria polunini*, **sp. n.**; *Neoempheria fajar*, **sp. n.**; *Neoempheria riatanae*, **sp. n.** (**Mycomyinae**); *Brachycampta glorialimae*, **sp. n.**; *Brachycampta murphyi*, **sp. n.**; *Brachycampta limtzepengi*, **sp. n.**; *Rymosia teopohlengi*, **sp. n.**; *Exechia tanswiehiani*, **sp. n.**; *Exechia alinewongae*, **sp. n.**; *Mycetophila chngseoktinae*, **sp. n.**; *Mycetophila georgettechenae*, **sp. n.**; *Mycetophila aishae*, **sp. n.**; *Platyprosthiogyne phanwaithongae*, **sp. n.**; *Platyprosthiogyne gohsookhimae*, **sp. n.**; *Platyprosthiogyne rahimahae*, **sp. n.**; *Platyprosthiogyne lynetteseahae*, **sp. n.**; *Platyprosthiogyne neilaae*, **sp. n.**; *Platyprosthiogyne snehalethaae*, **sp. n.**; *Platurocypta adeleneweeae*, **sp. n.**; *Platurocypta tanhoweliangi*, **sp. n.**; *Epicypta constancesingamae*, **sp. n.**; *Epicypta jennylauae*, **sp. n.**; *Epicypta limchiumeiae*, **sp. n.**; *Epicypta janetyeeae*, **sp. n.**; *Epicypta kohkhenglianae*, **sp. n.**; *Epicypta daintoni*, **sp. n.**; *Epicypta holltumi*, **sp.n.**; *Epicypta ridleyi*, **sp. n.**; *Epicypta chezaharaae*, **sp. n.**; *Epicypta tanjiakkimi*, **sp. n.**; *Epicypta gehminae*, **sp. n.**; *Epicypta jackieyingae*, **sp. n.**; *Epicypta khatijunae*, **sp. n.**; *Epicypta purchoni*, **sp. n.**; *Epicypta foomaoshengi*, **sp. n.**; *Epicypta ganengsengi*, **sp. n.**; *Epicypta nanyangu*, **sp. n.**; *Epicypta nus*, **sp. n.**; *Epicypta peterngi*, **sp. n.**; *Epicypta maggielimae*, **sp. n.**; *Epicypta yupeigaoae*, **sp. n.**; *Epicypta annwee*, **sp. n.**; *Epicypta wallacei*, **sp. n.**; *Epicypta lamtoongjini*, **sp. n.**; *Epicypta catherinelimae*, **sp. n.**; *Epicypta grootaerti*, **sp. n.**; *Epicypta joaquimae*, **sp. n.**; *Aspidionia cheesweeleeae*, **sp. n.**; *Aspidionia janetjesudasonae*, **sp. n.**; *Aspidionia fatimahae*, **sp. n.**; *Integricypta fergusondavie*, **sp. n.**; *Integricypta teosoonkimae*, **sp. n.**; *Integricypta shirinae*, **sp. n.**; *Integricypta hoyuenhoeae*, **sp. n.** (**Mycetophilinae**). The previously described species are *Metanepsia malaysiana* Kallweit, *Eumanota racola* Søli, *Parempheriella defectiva* (Edwards), *Neoempheria dizonalis* (Edwards) (all known from Sumatra and/or the Malaysian peninsula), and *Chalastonepsia hokkaidensis* Kallweit a species spread in east Asia. The mycomyine genus *Vecella* Wu & Yang is here proposed as a new synonym to *Parempheriella*, with *P. guadunana* (Wu & Yang), **n**. **comb**. corresponding to an additional Palearctic species of *Parempheriella*. Barcodes for a second set of 1,493 Singapore mycetophilid specimens suggest the presence of an additional 18 MOTUs. We thus estimate that approximately 85% of all the Singapore species that routinely enter Malaise traps are identified or described here. The revision concludes with a discussion of the biogeography and generic composition of the mycetophilid fauna at the southern end of the Malay Peninsula.

**Zusammenfassung:** “Open-ended” oder “Dark Taxa” sind artenreiche Klade, die so abundant und arteinreich sind, dass die herkömmlichen taxonomischen Methoden angesichts der großen Anzahl von Exemplaren und Arten nicht gut funktionieren. Neue Ansätze, die auf das Vorsortierung von Exemplaren mit DNA Barcodes bis zum Artniveau basieren, erlauben es jetzt aber solche Taxa zu revidieren. Allerdings kann auch mit DNA Barcodes, nur die Fauna eines vergleichsweise kleinen Gebietes bearbeiten werden, weil für die meisten Länder und biogeografischen Regionen zu viele Exemplare und Arten abgedeckt werden müssten. Wir demonstrieren hier wie eine solche Revision durchgeführt werden kann. Wir revidieren hier die Pilzmücken (Diptera: Mycetophilidae) von Singapur, die mit Malaisefallen gefangen werden können. Das hier überarbeitete Material stammt aus 496 Malaise-Fallenproben, die mit 71 Malaise-Fallen an 107 Sammelstellen in verschiedenen Habitaten gefangen wurden: Mangroven, Sumpfwälder, Süßwassersümpfe, Primärregenwälder und Sekundärwälder. Basierend auf molekularen und morphologischen Daten für mehr als 1454 Tiere grenzen wir 120 Arten mit molekularen und morphologischen Daten ab, wobei nur 5 dieser Arten bereits beschrieben sind. Die verbleibenden 115 werden hier beschrieben. Allerdings benennen wir nur 98 Arten, da für zwei Arten molekulare Daten fehlen und für 14 weitere Arten derzeit nur Weibchen bekannt sind. Daher können wir derzeit keine zufriedenstellende morphologische Diagnose erstellen. Darüber hinaus fehlen molekulare Daten für drei Arten (*Epicypta* sp. A, *Epicypta* sp. B und *Neoempheria* sp. H). Was die Artgrenzen betrifft, so stimmen die molekularen und morphologischen Daten in den meisten Fällen überein (match ratio: 95% für 3% MOTUs). Eine noch höhere „match ratio“ von 99% wird für 5% und ABGD MOTUs (P=0,060) beobachtet. Insgesamt ist das Verhältnis zwischen unbeschrieben und beschrieben erstaunlich hoch (20:1), und die Überarbeitung der singapurer Fauna erhöht die Anzahl der beschriebenen Arten in der Orientalischen Region um über 25%. Dies unterstreicht den Ausmaß des „taxonomic impediments“ für Pilzmücken. Die meisten der Mycetophilidenarten in Singapur gehören zu drei von 22 Gattungen - *Neoempheria* Osten-Sacken (31 Arten), *Epicypta* Winnertz (29 Arten) und *Manota* Williston (14 Arten), aber wir beschreiben hier auch eine neue Gattung, *Integricypta*, gen. n. basierend auf drei Arten. Die Gattung gehört zu den Mycetophilinae Mycetophilini und ist die mutmaßliche Schwestergruppe von *Aspidionia* Colless. Die sequenzierten, illustrierten und benannten Arten sind: *Leptomorphus rafflesi*, **sp. n.**; *Monoclona simhapura*, **sp. n.**; *Azana demeijeri*, **sp. n.**; *Azana leekongchiani*, **sp. n.** (**Sciophilinae**); *Tetragoneura crawfurdi*, **sp. n.**; *Tetragoneura chola*, **sp. n.**; *Tetragoneura dayuan*, **sp. n.**; *Tetragoneura farquhari*, **sp. n.**; *Ectrepesthoneura johor*, **sp. n.** (**Tetragoneurinae**); *Mohelia zubirsaidi*, **sp. n.**; *Allactoneura tumasik*, **sp. n.**; *Allactoneura limbosengi*, **sp. n.**; *Manota banzu*, **sp. n.**; *Manota tantocksengi*, **sp. n.**; *Manota bukittimah*, **sp. n.**; *Manota chiamassie*, **sp. n.**; *Manota danmaxi*, **sp. n.**; *Manota mahuan*, **sp. n.**; *Manota temenggong*, **sp. n.**; *Clastobasis sritribuana*, **sp. n.**; *Clastobasis bugis*, **sp. n.**; *Clastobasis oranglaut*, **sp. n.** (**Leiinae**); *Parempheriella mait*, **sp. n.**; *Parempheriella longyamen*, **sp. n.**; *Parempheriella peranakan*, **sp. n.**; *Mycomya sachmatich*, **sp. n.**; *Neoempheria merlio*, **sp. n.**; *Neoempheria sabana*, **sp. n.**; *Neoempheria sangabo*, **sp. n.**; *Neoempheria shicheng*, **sp. n.**; *Neoempheria ujong*, **sp. n.**; *Neoempheria subaraji*, **sp. n.**; *Neoempheria kokoiyeeae*, **sp. n.**; *Neoempheria mandai*, **sp. n.**; *Neoempheria malacca*, **sp. n.**; *Neoempheria sinkapho*, **sp. n.**; *Neoempheria singapura*, **sp. n.**; *Neoempheria xinjiapo*, **sp. n.**; *Neoempheria puluochung*, **sp. n.**; *Neoempheria merdeka*, **sp. n.**; *Neoempheria neesoon*, **sp. n.**; *Neoempheria pulau*, **sp. n.**; *Neoempheria cinkappur*, **sp. n.**; *Neoempheria temasek*, **sp. n.**; *Neoempheria polunini*, **sp. n.**; *Neoempheria fajar*, **sp. n.**; *Neoempheria riatanae*, **sp. n.** (**Mycomyinae**); *Brachycampta glorialimae*, **sp. n.**; *Brachycampta murphyi*, **sp. n.**; *Brachycampta limtzepengi*, **sp. n.**; *Rymosia teopohlengi*, **sp. n.**; *Exechia tanswiehiani*, **sp. n.**; *Exechia alinewongae*, **sp. n.**; *Mycetophila chngseoktinae*, **sp. n.**; *Mycetophila georgettechenae*, **sp. n.**; *Mycetophila aishae*, **sp. n.**; *Platyprosthiogyne phanwaithongae*, **sp. n.**; *Platyprosthiogyne gohsookhimae*, **sp. n.**; *Platyprosthiogyne rahimahae*, **sp. n.**; *Platyprosthiogyne lynetteseahae*, **sp. n.**; *Platyprosthiogyne neilaae*, **sp. n.**; *Platyprosthiogyne snehalethaae*, **sp. n.**; *Platurocypta adeleneweeae*, **sp. n.**; *Platurocypta tanhoweliangi*, **sp. n.**; *Epicypta constancesingamae*, **sp. n.**; *Epicypta jennylauae*, **sp. n.**; *Epicypta limchiumeiae*, **sp. n.**; *Epicypta janetyeeae*, **sp. n.**; *Epicypta kohkhenglianae*, **sp. n.**; *Epicypta daintoni*, **sp. n.**; *Epicypta holltumi*, **sp.n.**; *Epicypta ridleyi*, **sp. n.**; *Epicypta chezaharaae*, **sp. n.**; *Epicypta tanjiakkimi*, **sp. n.**; *Epicypta gehminae*, **sp. n.**; *Epicypta jackieyingae*, **sp. n.**; *Epicypta khatijunae*, **sp. n.**; *Epicypta purchoni*, **sp. n.**; *Epicypta foomaoshengi*, **sp. n.**; *Epicypta ganengsengi*, **sp. n.**; *Epicypta nanyangu*, **sp. n.**; *Epicypta nus*, **sp. n.**; *Epicypta peterngi*, **sp. n.**; *Epicypta maggielimae*, **sp. n.**; *Epicypta yupeigaoae*, **sp. n.**; *Epicypta annwee*, **sp. n.**; *Epicypta wallacei*, **sp. n.**; *Epicypta lamtoongjini*, **sp. n.**; *Epicypta catherinelimae*, **sp. n.**; *Epicypta grootaerti*, **sp. n.**; *Epicypta joaquimae*, **sp. n.**; *Aspidionia cheesweeleeae*, **sp. n.**; *Aspidionia janetjesudasonae*, **sp. n.**; *Aspidionia fatimahae*, **sp. n.**; *Integricypta fergusondavie*, **sp. n.**; *Integricypta teosoonkimae*, **sp. n.**; *Integricypta shirinae*, **sp. n.**; *Integricypta hoyuenhoeae*, **sp. n.** (**Mycetophilinae**). Die Tiere, die zu den bereits beschriebenen Arten gehören, gehören zu den folgenden Arten: *Metanepsia malaysiana* Kallweit, *Eumanota racola* Søli, *Parempheriella defectiva* (Edwards), *Neoempheria dizonalis* (Edwards) (alle Arten sind derzeit aus Sumatra und/oder der malaysischen Halbinsel bekannt), und *Chalastonepsia hokkaidensis* Kallweit, eine in Ostasien verbreitete Art. Die Gattung *Vecella* Wu & Yang wird hier als neues Synonym für *Parempheriella* vorgeschlagen, wobei *P. guadunana* (Wu & Yang), n. comb., wobei die Gattung damit eine zusätzlichen paläarktischen Art erhält. Die DNA Barcodes für eine zweite Stichprobe mit 1.493 Pilzmücken deuten darauf hin, dass in Singapur 18 zusätzliche Arten mit Malaisefallen gefangen werden können. Es sieht aber dennoch so aus, als würden wir bereits hier circa 85% aller Arten beschreiben, die regelmäßig in Malaise-Fallen geraten. Die Revision endet mit einer Diskussion der Biogeographie und taxonomische Zusammensetzung der Mycetophilidenfauna im südlichen Teil der Malaiischen Halbinsel.

## Introduction

Much of our planet’s species-level diversity belongs to hyperdiverse taxa that are nowadays often called “open-ended” (Bickel 2009) or “dark taxa” (Hausmann et al. 2020, Hartop et al. 2022). Hartop et al. (2022) proposed a precise definition of “dark taxa” by suggesting that the term should be restricted to clades with an estimated diversity of more than 1,000 species worldwide for which the undescribed fauna exceeds the described fauna by at least one order of magnitude. Dark taxa often dominate specimen samples obtained with Malaise traps and other mass sampling techniques. Most “dark” species are small in size, lack showy color patterns, and belong to genera that challenge taxonomists by being abundant and species-rich (Srivathsan et al. 2023). This means that comprehensive taxonomic revisions of these clades are missing for many parts of the world.

We here use a new approach to tackling the taxonomy of dark taxa that was called “dark taxonomy” in Meier et al. (2023). The process starts with the “reverse workflow”; i.e., sorting of specimens into MOTUs with DNA barcodes (Wang et al., 2018, Hartop et al., 2022). This is followed by the application of “Large-scale Integrative Taxonomy” (LIT) *sensu* Hartop et al., (2022). LIT is a formalized algorithm that allows for testing the congruence between morphospecies and MOTUs. LIT also provides guidance regarding how conflict between MOTUs and morphology can be resolved. A third element is the recommendation that dark taxa should be studied at geographic scales that are largely determined by the number of specimens and species that can be effectively managed during a revision. To increase the probability that the species are correctly delimited despite covering only a limited geographic range, we propose that dark taxonomy has to be integrative and delimits species for which molecular and morphological data are available. We envision that species for which only morphological or molecular data are available should be treated only later. The last element of dark taxonomy is the recommendation that revisions should initially be based on the kind of fresh samples obtained with methods that are also used for biomonitoring. This ensures that the results of the taxonomic revisions are immediately relevant for monitoring biodiversity.

Revising a fauna at a regional scale means that in the future it will become more important to combine information from multiple faunistic revisions covering neighboring areas. This is fairly straightforward for barcodes but also needs careful documentation of morphology. This is why we consider it indispensable that faunistic revisions are well illustrated (“digitized”) (Ang et al. 2013). Otherwise, new studies would have to physically examine type specimens. Generating morphological information is also important because specimen identification with barcodes is likely to be partially replaced with identification with artificial intelligence models (e.g., Convolutional Neural Networks; Wührl et al. 2022; Høye et al. 2021). In the process of training of CNNs, image sets will be needed and should ideally come from specimens identified by the author who described the species.

The known world fauna of Mycetophilidae may be over 5,000 described species (Kjærandsen, pers. comm.), but the total species richness is estimated to be at least 10 times larger. Evidence comes from at least three studies covering temperate and tropical habitats. Firstly, in Hebert et al.’s (2016) study of Canadian insects from Malaise trap material, Mycetophilidae ranked fifth highest in terms of number of BINs while the family was the third richest in Great Britain (Duff et al. 2012). Note that the 1,199 mycetophilid BINs from Canada are about 2.5 times the number of species currently known for the country (Savage et al. 2019). Secondly, in Brown et al.’s (2018) single-site inventory of the Diptera fauna of a patch of cloud forest in Costa Rica (see also Borkent et al. 2018), Mycetophilidae had the fourth highest number of species, Thirdly, considering ground level and canopy elements of the fauna of an Amazon tropical forest, mycetophilids were second among 38 fly families (Amorim et al. 2022). Overall, there is thus no doubt that the Mycetophilidae fit Hartop et al.’s criteria for a dark taxon.

The most recent classification for the Mycetophilidae includes six subfamilies—Sciophilinae, Leiinae, Tetragoneurinae, Gnoristine, Mycomyinae and Mycetophilinae (Oliveira & Amorim 2021). Some of the groups in the family are more species-rich in temperate areas, while some others are particularly diverse in tropical areas. The known Oriental fauna of the family consists so far of about 450 described species in 50 genera (of the overall slightly less than 150 extant genera in the family worldwide). Southeast Asian mycetophilids have been described mostly from Borneo, Sumatra, Malaysia, and Thailand—basically nothing is known about the Singaporean fauna of the family.

Unfortunately, we currently also do have little information on the natural history of tropical fungus gnats. This makes it difficult to predict how and whether they could be used as indicator taxa for habitat quality or fungus richness given that mycetophilids are critically dependent on fungi. It would indeed be very useful to have indirect ways to assess fungal diversity in a tropical forest. Økland (1994, 1996) and Økland et al. (2005) demonstrated that high mycetophilid species-richness and woodland types are correlated in boreonemoral forests, but more evidence is needed, which could be obtained through matching larvae and adults with barcodes (see Yeo et al. 2018). Larvae could be collected from fruiting bodies and mycelia and sequenced to understand to what extent the presence of adult mycetophilids may be indicative of fungal diversity in a forest or of forest health.

This revision of Singapore mycetophilids is part of a larger project on the insect diversity of Singapore as sampled by Malaise traps (Yeo et al. 2021, Srivathsan et al. 2023). However, this revision is the first to cover the taxonomy of an entire family using the format of a monographic treatment. Previous analyses of the material collected and sequenced concentrated on detecting diversity patterns across habitats, testing the effectiveness of the reverse workflow, starting with barcodes for all available specimens (Wang et al. 2018, Yeo et al. 2021) and revising genera of Dolichopodidae (e.g., Grootaert & Puniamoorthy, 2014; Grootaert 2018).

One of the main challenges in taxonomy is distinguishing poorly described species from those that are new to science. Therefore, we extensively reviewed the literature on the Oriental fauna of the family before describing new species. For the Oriental mycetophilid species described by Edwards (e.g., 1925, 1926, 1927, 1928, 1929, 1931, 1933, 1935), we studied and photographed primary types deposited at the Natural History Museum, London. For more recently described Oriental species of Mycetophilidae, we were able to resolve species identity based on published careful descriptions with good illustrations that are here cited following recommendations in Meier (2017): Colless (1966), Zaitzev (1982), Sivec & Plassmann (1982), Wu & Yang (1986), Bechev (1995), Søli (1996, 2002), Kallweit (1998), Matile (1999), Xu & Wu (2002), Wu et al. (2003), Papp (2004), Hippa et al. (2005), Hippa (2006, 2007, 2008, 2009, 2011), Ševčík & Hippa (2010), Hippa & Ševčík (2010, 2013), Ševčík et al. (2011), Ševčík & Kjaerandsen (2012), Borkent & Wheeler (2012), Kurina & Hippa (2015), Hippa & Saigusa (2016), Hippa & Kurina (2018), Magnussen et al. (2019), Fitzgerald (2017), Kaspřák et al. (2017). This unfortunately leaves some Oriental species—especially from Sri Lanka—that cannot be clarified in limbo because the descriptions are insufficient and the types are lost or unavailable. The names of these species are ignored in this treatment.

The results of this study are published in two papers. One describes the rationale for the approach—“dark taxonomy” (Meier et al., 2023), while this paper describes the species and addresses the implication of the investigation of the Singaporean fauna for the Mycetophilidae systematics. This monograph only provides scant information on the rationale of the approach and the reader is referred to the companion paper for more information on why we only treated only fresh specimens from Singapore. Note that all species descriptions presented here also include proper molecular diagnoses as recommended by Rheindt et al. (2023).

## Material and Methods

This revision of the Mycetophilidae fauna of Singapore proceeded through five phases as described in Meier et al. (2023): i) collecting, ii) molecular processing (barcoding and establishing haplotype networks), iii) integrating morphological and molecular data, iv) formally describing, and v) assessing the completeness of the fauna.

All Singapore samples were obtained with Gressit-type Malaise traps between 2012 and 2019. Under “Material examined” for each species, we refer to the collectors as “MIP leg.”; the sampling period is indicated by the collecting end date (in almost all cases, with samples recovered every week) in the format dd/mm/yy.

All specimens studied in this paper (that have a ZRCBDP code) were collected in Singapore (see map of the collecting sites in Figs. 1A-B). The collecting effort varied considerably between sites and environments. The project covered the following habitats: (1) mangrove (Fig. 2A); (2) primary rainforest (Fig. 2B); (3) maturing secondary rainforest; (4) old secondary rainforest; (5) coastal forest; (6) degraded urban forest (Fig. 2C); (7) old mangrove; (8) replanted mangrove; (9) swamp forest (Fig. 2D); and (10) freshwater swamp (Fig. 2E).

The techniques used for i) molecular processing (barcoding and establishing haplotype networks) and ii) integrating morphological and molecular data are described in Meier et al. (2023), but we here also follow recent recommendations regarding the provision of formal molecular diagnoses (Rheindt et al, 2023). For this purpose, we derived Diagnostic Molecular Combinations (DMCs) using UITOTO (Torres et al., 2024) for all 117 species with molecular data, encompassing 1,454 sequences. We conducted 50,000 search iterations, varying the minimum number of exclusive character states for the DMCs (3, 4, and 5), without defining a minimum length. An additional search was performed, setting the minimum length for the final DMCs to ten nucleotides for a minimum number of exclusive character states of 4. The resulting DMCs from these searches were tested against 60,349 unaligned Mycetophilidae sequences from The Barcode of Life Data System (BOLD; Ratnasingham & Hebert, 2007) using UITOTO. The DMCs with the best performance, in terms of discrimination power, were selected and visualized (also in UITOTO) on one sequence per species, typically the holotype sequence. Detailed molecular diagnosis analyses are available in the Supplementary Material (files ScriptMolecularDiagnosis.R, Mycetos_dataset1_27mar24.fasta, MycetophilidaeTesting.fasta, MycetoSpeciesList.csv, DMCsMycetoFinal.csv, and the functions ALnID2.r and ConfuClass.R).

All holotypes, most paratypes and most identified specimens are housed in the Zoological Research Collection (ZRC) within the Lee Kong Chian Natural History Museum, National University of Singapore (LKCHNM). All specimens in the list of examined material are deposited in the ZRC unless stated otherwise (some of the paratypes and identified specimens are deposited in the Museu de Zoologia da Universidade de São Paulo: MZUSP). The types of some of the Oriental species of mycetophilids referred to in the paper were examined at the Natural History Museum (NHM), London. All specimens were originally in ethanol and most are still kept in their original vials with ethanol. The exception are most holotypes and some paratypes that were slide-mounted in Canada balsam. Specimens that were too damaged, though recognized as part of a species were not designated as paratypes. Slide-mounted specimens are indicated in the list of examined material— otherwise kept in ethanol in 1 ml glass tubes.

A set of over 13,000 photographs of habitus and individual photos that were later stacked to illustrate details of the morphology are available from the Lee Kong Chian Natural History Museum. Photographs of the habitus and important features are also published here and were acquired with the Dun Inc. Passport II Imaging system, using a Canon 6D MkII chassis fitted with a MP-E 65 mm lens. Specimens were imaged at different depths of field, usually with 30 to 40 image ‘slices’, then focus-stacked using Zerene Stacker LLC to achieve a fully-focused image of the specimen. The images were then processed using TopazLab©’s ‘Mask AI’, ‘Sharpen AI’ and ‘DeNoise AI’ plugins, as well as Adobe Photoshop© and Adobe Illustrator© CC for manual image clean-up and figure plating. Slide mounted specimens were imaged using a Leica DC500 camera attached to a Leica MZ-16 stereomicroscope or to a compound microscope model Leica DM2500. The photos were later stacked with Helicon Focus 6. The high-resolution stack images of the habitus of most species are presented on the open-access *Biodiversity of Singapore*online platform (https://singapore.biodiversity.online/taxon/A-Arth-Hexa-Dipt-Mycetophilidae).

Due to the description of a large number of species in a single paper, the textual description of morphological structures is reduced compared to what is found in most papers describing few species. Measurements, for example, are provided only of wing length and wing width (always in millimeters). We opted instead for presenting more photographic evidence summarized in plates. Some may feel that some information is missing, and minimalists will complain that there is too much information: this paper tries to find a balance that allows for large-scale revision of a dark taxon. Future taxonomic studies dealing with the Singapore species of Mycetophilidae may further concentrate on the relationships between species, add drawings of the terminalia, have keys to the species of each genus etc.— sections that we see in regular taxonomic revisions.

Based on a principle of integrative taxonomy, we describe but do not formally name species known only from females for which there is insufficient information to propose a satisfying morphological diagnosis. The same applies to species known only from morphology, i.e., without a barcode sequence. These species will be named when males become available or when they have a barcode.

The standards of species descriptions in the literature vary between genera of Mycetophilidae. This is partly due to differences in the morphology of different genera but also to idiosyncrasies in the way particular communities of taxonomists describe species. We tried as much as possible to follow the standards of species descriptions present in the literature for each genus, what accounts for some differences in the way different genera are described in this paper. We provide a more detailed description for at least one species of each genus.

With few exceptions, we follow the morphological nomenclature for Diptera in Cumming & Wood (2017) and Søli’s (1997, 2017) monographs for structures typical of the Mycetophilidae. However, there are some few inconsistencies between both systems, e.g., we use here prosternum (Cumming & Wood 2017) for Søli’s (1997) basisternum. For the particularly complex morphology (and, hence, nomenclature) of the male terminalia of *Manota*, we follow Hippa & Ševčík (2013). The term “macroseta” is used here for flat, capitulate setae, as typically seen in *Manota* Williston, but also present in species of *Platurocypta* Enderlein.

Strict homology criteria are applied to name morphological structures. For example, the maxillary palpomere 3 in the Mycetophilidae ground-plan has a sensory pit. In some groups, however, the palpomere with the sensory pit is the second most basal palpomere—it is still referred to here as palpomere 3 even if there is no evidence of loss of palpomere 1 or fusion of palpomeres 2 and 3.

Regarding some details of the wing venation, we use the homology interpretation of the radial sector in Bibionomorpha as discussed in Amorim (1993) and Amorim & Rindal (2007): the Anisopodiformia preserve R_2+3_ (branching more basally than the insertion bibionomorphs of r-m, with R_4_ lost), while the remaining members of the suborder (in the sense of Amorim & Yeates 2006) preserve R_4_ (branching beyond or much beyond the base of r-m, with R_2+3_ lost). About the homology of the cubital and anal veins, we follow Cumming & Wood (2017). There are some minor additional problems of interpretation in the literature for this area of the wing. In sciophilines, there is a well-defined CuP (formerly referred to as A_1_) and a sclerotized cubital fold (“pseudovein”) adjacent to CuA—with no evidence of any anal vein—but two further changes during the evolution of mycetophilids obscure this original pattern. The anal lobe has a very weak anal fold in, e.g., tetragoneurines and leiines, that becomes more evident in mycomyines and is well sclerotized in mycetophilines. In the literature, this fold is often referred to as the anal vein, what leads to an erroneous interpretation of homology; it is referred to here as an “anal fold” (af). Additionally, CuP and the cubital pseudovein seem to be fused in some genera and it is hard to distinguish between them.

The list of abbreviation used in the illustration plates is as follows:

ae: aedeagus;
aep: aedeagal apodeme;
af: sclerotized anal fold;
allgc: apicolateral lobe of gonocoxite;
anp: antepronotum;
ans: anepisternum;
asp: anterior spiracle;
avlgc: apicoventral lobe of gonocoxite;
car: cardo;
ce: cercus;
ce1–2: cercomere 1–2; **ces**, cervical sclerite;
cly: clypeus;
cpv: cubital pseudovein;
cxI–III: coxae I–III; **es**, tergite 9 megaseta;
fc: face;
flg: flagellomere;
fmv: false medial vein;
frf: frontal furrow;
fr: frons;
gc: gonocoxite;
gca: gonocoxal apodeme;
gcd: gonocoxal distal setae on lateral projection;
gcpb: gonocoxite posterior ventral border;
gdl: gonocoxite dorsal lobe
gfk: genital fork;
gil: gonocoxite internal lobe
gll: gonocoxite lateral lobe
gvl: gonocoxite ventral lobe;
gci: gonocoxal inner seta on latera projection;
gfk: genital fork;
glp: gonocoxite lateral projection
gmp: gonocoxite medial projection
gn: gena;
gop: genital opening;
gon ap: gonocoxal apodeme;
gpb: gonocoxite posterior border;
gpl: gonocoxal posterior lobe;
gs: gonostylus;
gsbl: gonostylus basal lobe
gsdl: gonostylus dorsal lobe;
gsil: gonostylus inner lobe;
gsl: gonostylus main lobe;
gsml: gonostylus medial lobe;
gsol: gonostylus outer lobe;
gspl: gonostylus posterior lobe;
gsvl: gonostylus ventral lobe;
hal: halter;
hypd: hypandrium;
ibr: internal branch of gonostylus;
jgs: juxtagonostylar lobe;
kts: katepisternum;
lbl: labellum;
loc: lateral ocellus;
lt: laterotergite;
mdt: mediotergite;
mk: medial keel of scutum;
moc: mid ocellus;
mem: mesepimerom;
mtem: metepimeron;
mtp: metepisternum;
mxp1–5: maxillary palpomeres 1–5;
oc s: ocellar seta;
ocf: occipital foramen;
ocp: occiput;
pa: paramere;
pal: parastylar lobe;
pap: parameral apodeme;
pda: parameral distal arm;
ped: antennal pedicel;
ppm: postpronotum;
pem: proepimeron;
pep: proepisternum;
prs: prosternum;
ps: parameral spine;
psl: parastylar lobe;
psp: posterior spiracle
pst: pseudotrachaea;
S1–10: sternites 1–10;
sai: scutum anterior incision;
scl: scutellum;
scp: scape;
sc: scutum;
spd: spermathecal duct;
sep: maxillary palpus sensory pit;
sgms: syngonocoxite medial suture;
sgvp: syngonocoxite medioventral process;
sti: stipes;
syngcxm: syngonocoxite medial sclerite;
T1–9: tergites 1–9;
tr: trochanter;
vgf: vaginal furca
vrt: vertex;
wf: wing medial fold.

This article conforms to the requirements of the amended International Code of Zoological Nomenclature (ICZN) and the new names were registered in ZooBank, the online registration system for the ICZN.

## Results

We found 120 species in the first batch of 1,456 specimens of the project. Of these, 115 are new to science. The five previously described species with confirmed specimens in our material are: *Metanepsia malaysiana* Kallweit, *Eumanota racola* Søli, *Parempheriella defectiva* (Edwards), *Neoempheria dizonalis* (Edwards) (all known from Sumatra and/or the Malaysian peninsula), and *Chalastonepsia hokkaidensis* Kallweit (distributed from the Seychelles Islands to Hokkaido). The remaining 115 are new species. This implies a 20:1 ratio of undescribed to described species in the Oriental fauna of mycetophilids. Overall, this revision of the Singapore fauna thus increases the number of described species of Mycetophilidae in the entire Oriental Region by over 25%—a good measure of the taxonomic impediment for fungus gnats. Note that 25% and 9% of the species in the first batch are known respectively from singletons or doubletons.

A second batch of 1,567 specimens was sequenced after the cut-off date for the taxonomic revision. The clustering of the barcodes suggests the presence of an additional 28 MOTUs (Meier et al., 2023), meaning that the first batch of specimens covered ca. 85% of the sampled diversity.

As mentioned above, principles of integrative taxonomy guided our decisions on which new species should be formally named in this monograph. Species known only from barcodes and from females are described only if the females have a sufficiently large number of diagnostic features. This was not the case for seven putative species of *Manota* and seven of *Neoempheria* Osten-Sacken. These species clearly diverge from other species based on DNA barcodes and are likely different species, but without their respective males, there is not enough information to prepare morphological diagnoses. The females of these species were described and illustrated in the paper, but they were referred by letters, “sp. A”, “sp. B” etc. This applies to all species of *Manota* and most species of *Neoempheria* without males in our samples. Species of *Epicypta* Winnertz and of some other genera known only from females have enough features to build a diagnosis. In addition, one of the species of *Epicypta* and one of the species of *Neoempheria* are only known of a single specimen that did not barcode successfully. This species is similarly described but not named; it will be formally named when a barcoded male specimen that matches the morphology becomes available. This approach to the fauna of the family allows for the actual characterization of the known local diversity of mycetophilids but avoids generating unnecessary burden to future taxonomists using barcode or morphological information. As more mycetophilid specimens are collected in Singapore, these species will eventually be named.

Besides the addition of species to the world fauna of the family, this paper has a substantial amount of new systematic information on Mycetophilidae. The paper has the first color photographs of the habitus of some mycetophilid genera (e.g., *Aspidionia* Colless). Many of the details of body parts of different genera are described and/or illustrated in detail for the first time. Various issues related to the delimitation of subfamilies and genera are also addressed, for example the question of the limits of *Dziedzickia* Johannsen and *Neoempheria*, the relationships among the Mycetophilini, diagnoses of genera mostly with Holarctic distribution (e.g., *Brachycampta* Winnertz), the position of the Manotini among the Leiinae etc.

Some of the species-rich genera of the family, such as *Epicypta*, have had almost no species described worldwide in the last 20 years. With this faunistic survey, it was possible to identify diversity patterns within the family that have been masked by the lack of research on a dark genus like *Epicypta*: our results suggest that *Epicypta* and *Neoempheria* may each be two or three times more diverse than *Manota,* a genus that received much more attention in the last 20 years (now with over 300 described species). Indeed, most of the diversity of Mycetophilidae found in Singapore belong to three of 21 genera— *Neoempheria* (31 species), *Epicypta* (29 species) and *Manota* (14 species).

We also describe a new genus in the mycetophiline tribe Mycetophilini based on three species. As discussed below, the genus may be sister to *Aspidionia*. Finding a new genus in this study was quite unexpected, given that *Brachyradia* Ševčík & Kjærandsen, 2012 was the only mycetophiline genus described since Tuomikoski (1966) and Colless (1966). This finding thus reveals the size of knowledge gaps for the biodiversity of tropical Mycetophilidae.

A key to the genera of Mycetophilidae expected to be found in the Malay Peninsula, including Singapore, is provided below. This key is largely based on Søli et al. (2000) and Søli (2017).

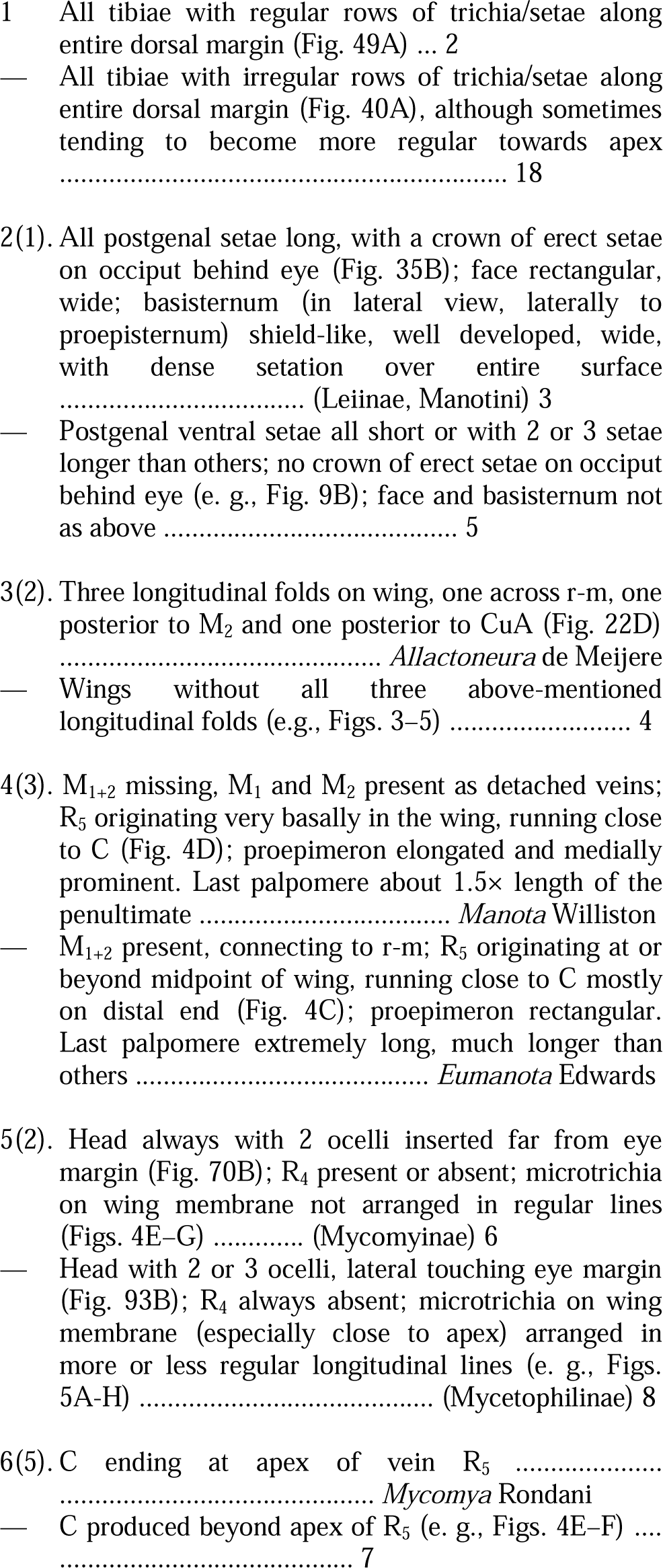
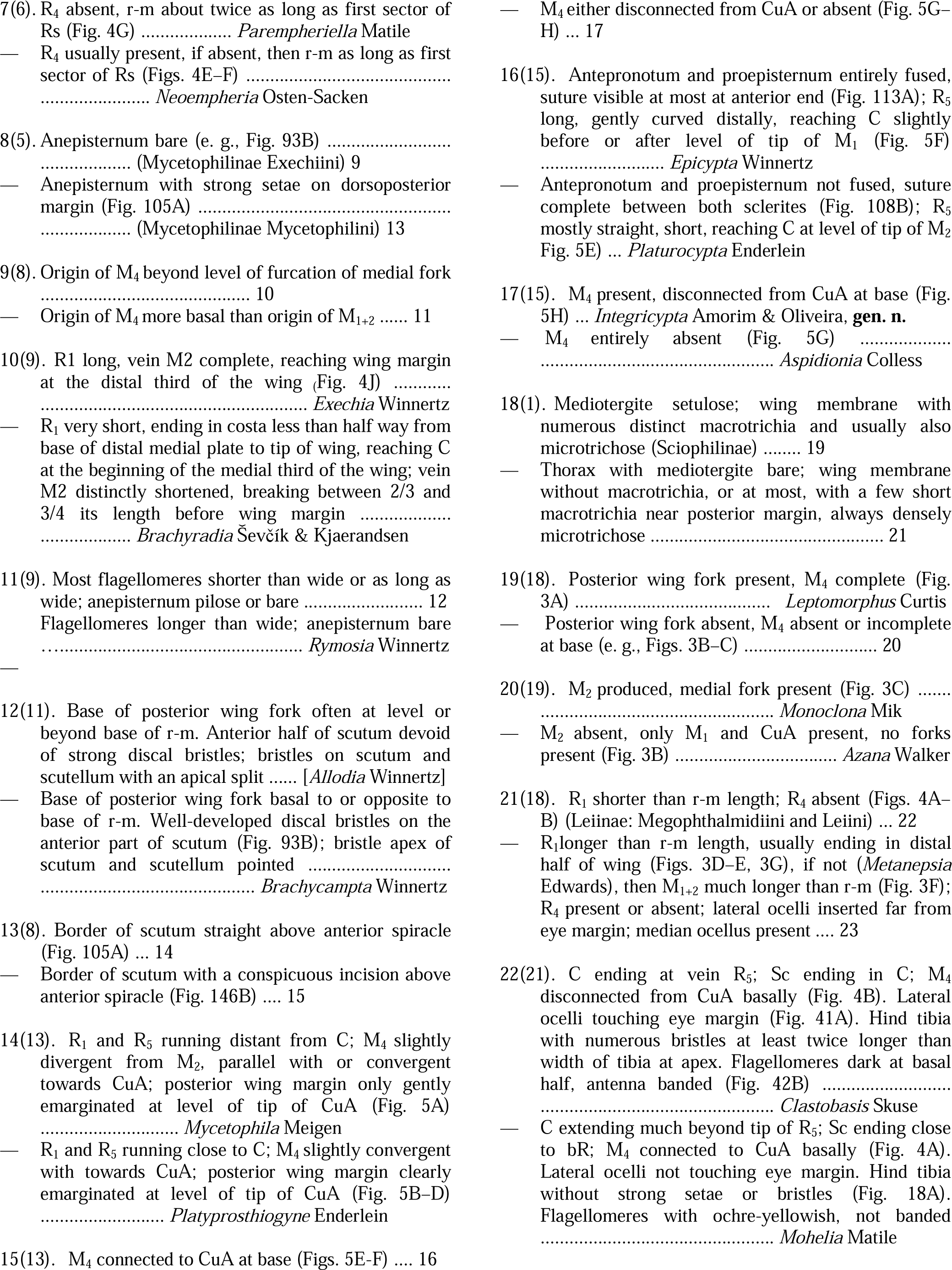
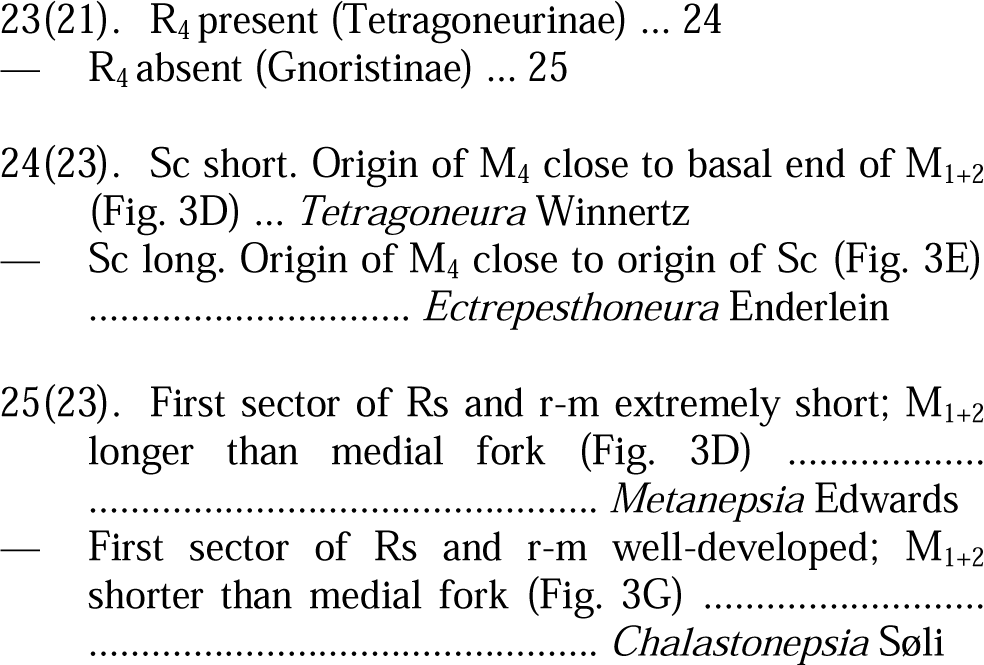

## Taxonomy

### Sciophilinae

Søli’s (1997) phylogenetic study of the Mycetophilidae showed that the traditional “Sciophilinae s.l.” (which includes all mycetophilids except those in the subfamily Mycetophilinae) was a large paraphyletic group. This has been corroborated by all phylogenies of the family published more recently. We use here Sciophilinae in the sense of Borkent & Wheeler (2013), with about 37 genera. The subfamily consists mostly of species with macrotrichia on the wing membrane. Søli’s (1997) phylogeny had a small clade with *Drepanocercus* Vockeroth and *Paratinia* Mik sister to the remaining mycetophilids, meaning that even in this stricter sense the sciophilines could be paraphyletic. This differs from Borkent & Wheeler’s (2013) phylogenetic analysis of the Sciophilinae (ranked as a tribe), in which *Drepanocercus*, *Paratinia*, *Loicia* Vockeroth and *Acomoptera* Vockeroth are present in a clade sister to the set of remaining genera of the subfamily.

The Oriental sciophiline fauna is composed of eight genera—*Acnemia* Winnertz, *Anaclileia* Meunier, *Azana*, *Leptomorphus*, *Monoclona*, *Neuratelia* Rondani, *Polylepta* Winnertz, and *Sciophila* Meigen. Of these genera, only *Azana*, *Leptomorphus*, *Monoclona* are present in our Singapore samples.

### Leptomorphus Curtis

*Leptomorphus* Curtis, 1831: 365. Type species, *Leptomorphus walkeri* Curtis, by monotypy.

*Diomonus* Walker, 1848: 87 (Type species: *Diomonus nebulosus* Walker, by monotypy). Edwards (1925: 556), synonymization of *Diomonus*.

**Diagnosis** (modified from Borkent & Wheeler 2012, 2013). Flagellomere 1 with a distinctly offset basal stalk, flagellomeres slightly compressed laterally; katepisternum and anepimeron bare, antepronotum and proepisternum setose, acrostichal bristles absent. Wing membrane without microtrichia; sc-r (when present) placed before or at origin of Rs, C not produced beyond tip of R_5_, R_4_ present or absent, anterior and posterior forks present, M_1+2_shorter than medial fork, base of M_1_ complete, anterior end of medial fork beyond anterior end of posterior fork. Tibial bristles short, no more than half the width of tibia; male abdominal segment 7 more than half length of segment 6.

Eight Oriental species of *Leptomorphus* were described by Papp & Ševčík (2011) and one by Kaspřák et al. (2017). The worldwide diversity of *Leptomorphus* was revised by Borkent & Wheeler (2012). Their phylogeny of the genus demonstrates that the subgeneric arrangement proposed for *Leptomorphus* rendered paraphyletic taxa. The genus now includes 46 species, of which ten are from the Oriental region, while other five species occur east of the Wallace line, into the Australasian region. We carefully considered the knowledge of the Oriental species of the genus and the species in our samples does not fit into any of the described species—although it certainly belongs into one of the small groups of species (see below).

*Leptomorphus rafflesi* Amorim & Oliveira, sp. n. (Figs. 6A-D, **7A-D**)

urn:lsid:zoobank.org:act:06855A29-80E5-4642-8158-B4BA4A4AE583

**Diagnosis**. Antennal scape yellowish, pedicel and flagellum brown. Thorax mostly brown, metepisternum light brown on anterior half, yellowish on posterior half. Scutum with scarce fine yellowish setation, especially along dorsocentral line. Tibiae with short, sparse darker setae, but not densely distributed anteroventrally. R_4_ absent. Abdominal tergites 1–2 and 6–7 brown, tergites 3– 5 brown with a yellowish band on anterior third or half. Terminalia longer than wide, gonocoxite with a short distal pointed beak; gonostylus articulated distally on gonocoxite, folded backwards, “phallic organ” present, wide.

***Leptomorphus_rafflesi_ZRCBDP0066807_hapZRCBDP 0066806_SMH_holotype* [43: T, 67: G, 130: G, 214: T, 2 29: C, 280: G, 301: C]**

cttatcttctacaatcgcccatgccggagcatctgtagatctTgcaatcttctctcttcatttggcGggtatttcttcaattttgggggccgttaattttattactactattattaatatacgagccccGggaatttctttcgaccgaataccactatttgtatgatcagttttaattacagcagtacttctccttctatctttaccagttctTgctggagcaatcacCatacttttaacagatcgaaatttaaatacatcattttttgacccagctggGggaggagacccaattctttaCcaacatttattc

#### Description

**Male** (Fig. 6A). Wing length, 4.08; width, 1.24. Body mostly dark brown. **Head** (Fig. 6B). Head yellow, palpus and antennal scape light brownish-yellow, pedicel brown, flagellum light brown, flagellomeres slightly flattened (flagellomeres 11–14 missing on holotype). Ocellar triangle black, ocelli large. Anterior ocellus slightly anterior to lateral ocelli. Lateral ocelli far from eye margins. Prefrons ventrally with four stronger setae and some other smaller setae. Eyes bare. Palpomere 3 short, with a deep sensorial pit, distal palpomere slender, twice longer than penultimate palpomere. **Thorax** (Fig. 6C). Scutum dark brown, scutellum cream-yellow. Scattered yellowish thin setae over scutum, with a more or less regular row of dorsocentrals. Scutellum small, weakly sclerotized, entirely bare. Pleural sclerites brown, except for metepisternum light brown on anterior half, yellowish on distal half, laterotergite and mediotergite dark brown; pleural membrane around anterior spiracle yellowish. Some few thin setae on antepronotum, one single thin fine seta on proepisternum; anepisternum, katepisternum, mesepimeron, and metepisternum entirely devoid of setation, three stronger setae dorsoposteriorly on laterotergite, about 15 fine setae and three pairs of stronger setae on mediotergite laterally on ventral half. Coxae yellow, femora yellow with brown proximal and distal fifth, tarsi light brown. Unaligned microsetae on tibiae, short scattered macrosetae on all tibiae. Empodia not developed, claws minute with a basal tooth. **Wing** (Fig. 6D). Membrane fumose light brownish, with dense macrotrichia, veins brown. All veins with dorsal setae, all veins but M_4_ and second sector of CuA with ventral setae. Sc reaching C before mid of wing; sc-r close to apex of Sc. R_1_ long, reaching C at distal fifth of wing, M_1_, M_2_, and M_4_ weakly sclerotized close to wing margin. M_1+2_ about half of length of M_1_, M_4_ with a gentle depression on distal fourth. CuA gradually and gently divergent from M_4_. CuP sclerotized, reaching margin at about level of mid of second sector of CuA. Haltere brown. **Abdomen**. Abdominal tergites 1–2 and 6–7 brown, tergites 3–5 yellow on anterior half, brown on distal half; sternites 1–2 light brown, sternites 3–5 yellow on anterior half, brown on distal half, sternites 6–7 brown, tergite and sternite 8 short, light brown. Distribution of setae on tergite 8 slightly asymmetrical, sternite 8 rhomboid, larger. **Terminalia** (Figs. 7A–C). Whitish-yellow. Gonocoxites longer than wide and fused to each other anteromedially, without sclerotized suture medially; laterodistal lobe of gonocoxites short, with a short spine projecting at distal borders. Gonostylus folded backwards, articulated distally on gonocoxal lobe, more sclerotized basally, bare, with a condylus-like projection on internal margin midway to apex and a short digitiform process distally at dorsal margin. “Phallic organ” wide, present medio-distally between gonocoxites in ventral view, ejaculatory apodeme long. Cercus weakly sclerotized, with fine setae.

**Female**. Unknown.

#### Material examined

**Holotype**: male, ZRCBDP0066807, Bukit Timah, maturing secondary forest (BT08), 16.aug.2016, MIP leg. (slide-mounted). **Additional sequenced specimens**: male, ZRCBDP0066806, Bukit Timah (BT08), maturing secondary forest, 16.aug. 16, Mangrove Insect Project leg.

**Etymology**. This species is named after Sir Thomas Stamford Raffles (1781-1826), knighted in 1816, British East Indian administrator, largely responsible for the creation of Britain’s Far Eastern empire. The Raffles Museum of Biodiversity Research, National University of Singapore, had its origins in the Raffles Museum, founded in 1849 due to an initiative by Raffles. Considered the founder of modern Singapore (in 1819) by the Singaporean government, Raffles was also a naturalist collector who inspired the establishment of a natural history collection of Southeast Asian biodiversity— resulting in the Zoological Research Collection (ZRC) now housed in the Lee Kong Chian Natural History Museum, National University of Singapore.

**Remarks**. In Borkent & Wheeler’s (2012) key for the species of the genus, *Leptomorphus rafflesi* Amorim & Oliveira, **sp. n.** runs to *L. tagbanua* Borkent, from the Philippines, from which it is clearly different, as can be seen by different aspects of the male terminalia. It is also similar to *L. ascutellatus* Ševčik, from Thailand, but the differences in the male gonostylus are striking. The shape of the “phallic organ” shows that *L. rafflesi* Amorim & Oliveira, **sp. n.** clearly belongs in the clade of the genus including, e. g., *L. hyalinus* Coquillet, *L. titiwangsensis* Borkent and *L. tagbanua* Borkent. This species was collected only in Bukit (=hill) Timah and there is a single haplotype in our material for this species. (Fig. 7D).

### Monoclona Mik

*Staegeria* van der Wulp, 1876: xlix (preoccupied Rondani, 1856). Type species: *Sciophila halterata* Staeger, 1840: 275 (monotypy) [= *rufilatera (*Walker)].

*Monoclona* Mik, 1886: 279 (nom. n. for *Staegeria* Wulp).

**Diagnosis** (modified from Borkent & Wheeler 2013). Two or three ocelli. Metepisternum setose. Wing membrane macrotrichia reflexed towards wing base; sc-r placed before origin of Rs; R4 present or absent; anterior fork present, posterior fork absent (M4 missing), M1+2shorter than medial fork.

Of 19 species of *Monoclona* described worldwide (Borkent & Wheeler 2013; Amaral et al. 2022), only one is known so far from the Oriental region—*M. laosilvatica* Ševčík, from Laos. The remaining species are from the Palearctic (six), Nearctic (four), and Neotropical (eight) regions. The species from Singapore, described here based on females, clearly diverges from *M. laosilvatica*, as discussed below. There are no worldwide revisions of the genus or a phylogeny for the species of *Monoclona*.

*Monoclona simhapura* Amorim & Oliveira, sp. n. (Figs. 8A-F)

(https://singapore.biodiversity.online/species/A-Arth-Hexa-Diptera-000769)

urn:lsid:zoobank.org:act:08741C27-4ACE-45F6-8AB5-C189762E858F

**Diagnosis**. Head brownish, scutum and abdominal tergites with ochreous-yellow thoracic pleura, wing approximately 2 mm long.

Monoclona_simhapura_ZRCBDP0048567_hapZRCBDP 0048567_SMH_holotype [40: C, 70: G, 103: C, 106: C, 1 09: A, 138: T, 139: T, 151: G]

attagcttctacaatcgcccatgcaggggcatcagttgaCttagctattttttcattacatttagccggGatttcttcaattttaggagcagttaattttatCacCacAattattaatatacgagccccaggtattaTTtttgatcgaatGccattatttgtatgatcagttttaattacagctattttacttcttctatcattaccagttttagccggagctattactatacttttaacagatcgaaatttaaatacttcctttttcgatcctgcgggagggggagacccaattttataccaacacctattt

#### Description

**Female** (Fig. 8A). Wing length, 2.04 mm, width, 0.92 mm. **Head** (Fig. 8B). Elliptical, higher than long. Vertex dark brown, frons brown, lighter ventrally, setose, ocellar triangle dark brown. Three ocelli, lateral ocelli close to each other, away from eye margin, mid ocellus small. Occiput light yellowish-brown on ventral half laterally, ochre-brown towards vertex, brown around ocelli. Eye with small interommatidial setae. Antennal scape and pedicel light ochre-brown, with small setae, scape slightly longer than pedicel; 14 light brown flagellomeres, darker towards apex, almost as long as wide, with scattered setae. Fronto-clypeus light ochre-brown, covered with short setae; labella cream, elongate; five palpomeres, basal one hardly visible, palpomeres 2 and 3 light brown, distal two segment whitish-brown, palpomeres gradually longer, no evidence of sensorial pit on third palpomere. **Thorax** (Fig. 8B). Scutum and scutellum mostly light brown, yellowish only at anterior margin of scutum. Two parallel rows of irregular acrostichal setae and a pair of irregular rows of dorsocentral larger setae, separated anteriorly and coming close together near scutoscutellar suture. Some stronger setae along entire lateral margin of scutum, with some supra-alars and some prescutellars. Transverse suture short but well-marked. Pronotum and proepisternum light brown, anepisternum brownish-ochre, katepisternum cream-yellow, mesepimeron slightly darker, laterotergite and mediotergite brown, pleural membrane whitish-yellow. Scutellum with some setulae on disc, at least four longer and many shorter setae along distal margin. Prosternum with seven setae. Antepronotum with four stronger and 10 smaller setae, proepisternum quite small, subquadrate, with two larger and 12 smaller setae, proepimeron elongate, bare. Anepisternum, katepisternum, mesepimeron, and metepisternum entirely bare, mesepimeron reaching ventral margin of thorax. Laterotergite bulging, with 11-14 setae of different sizes. Mediotergite only slightly curved in profile, ventral half with three delicate setae. Haltere pedicel whitish-yellow, knob brown, with setulae on both parts. **Legs**. Fore coxa brownish ochre, mid and hind coxae light ochre with brownish tinge on distal third of wing; femora brownish-ochre, slightly darker on distal half, tibiae yellowish-brown, strongly setose, trochanters light brown. Length of tarsomere 1 more than twice tarsomere 2 in all legs. Tibial spurs brownish, length more than twice tibia width at apex, spurs subequal in length. Tarsal claws with a conspicuous tooth midway to apex. **Wing** (Fig. 8C). Membrane homogenously fumose, densely covered with microtrichia and macrotrichia, macrotrichia reflexed, directed towards wing base, all wing veins setose except sc-r, first sector of Rs, r-m, and M_1+2_. Sc complete, ending at C slightly beyond base of Rs. C extending slightly beyond R_5_. First sector of Rs oblique. R_1_ long, reaching C close to level of tip of M_2_; R_4_ absent; R_5_ gently sinuous on distal fifth, reaching C clearly before wing apex; r-m oblique, distal end more basal than basal end, well sclerotized. M_1+2_ present, shorter than r-m, M_1_ and M_2_ produced, base of M_2_ less sclerotized. No trace of M_4_, CuA well-sclerotized, CuP absent. **Abdomen**. Tergite 1 yellowish-brown, tergites 2-6 light brown, tergite 7 yellowish-brown, sternites 1-7 light brownish-yellow, sternite 6 yellowish-brown. **Terminalia** (Figs. 8D–E). Terminalia yellowish, sternite 8 slightly darker, with some few darker setae at distal margin. Sternite 8 with a pair of medial wide projections with setae along distal margin. Tergite 8 and 9 very short, partially fused to each other, bare. Tergite 10 slender, wide, extending laterodistally towards ventral margin of terminalia, with 6–7 pairs of setae, most of which originating at tip of small projections. Two cercomeres, distal one short, nearly fused to cercomere 1.

**Male**. Unknown.

#### Material examined

**Holotype**: female, ZRCBDP0048567, Nee Soon (NS1), swamp forest, 25.apr. 12, MIP leg. (slide-mounted). **Paratype**, 1 female: ZRCBDP0048568, Nee Soon (NS2), swamp forest, 30.may. 12, MIP leg.

**Etymology**. The specific epithet of this species name comes from the Sanskrit original form of the name of Singapore, gathering the words *simha*, for lion, and *pura*, for place or town. The noun is used in apposition.

**Remarks**. We have only females from this species in our samples. *M. laosilvatica* is mostly yellowish, including the abdomen (Ševčík 2001, p. 159), while *M. simhapura* Amorim & Oliveira, **sp. n.** is mostly brownish. Also, *M. simhapura* Amorim & Oliveira, **sp. n.** is smaller, with the wing about 2 mm long, while *M. laosilvatica* is larger, the wing length being 3.15 mm. Since we can establish a morphological diagnosis for this species, we formally name it. This species was collected only in the swamp forest and both specimens share the same haplotype (Fig. 8F).

### *Azana* Walker

*Azana* Walker, 1856: 26 (type-species: *Azana scatopsoides* Walker 1856: 26, by monotypy; = *Azana anomala* Staeger 1840).

**Diagnosis** (modified from Borkent & Wheeler 2013). No interommatidial setae present; frons bare. Mediotergite and laterotergite bare. Wing sc-r placed before origin of Rs, R_4_ present or absent, when present forming a cell that is three times longer than high; anterior and posterior forks present; M_1+2_longer than medial fork, origin of anterior fork beyond level of origin of posterior fork. Male abdominal segment 7 less than half length of segment 6.

There are no revisions available for the Oriental species of *Azana*. Recent publications on the genus include a new species for the Neotropical region (Amorim et al. 2008a,b) and two new species for the Nearctic region (Kerr 2010). The Oriental region so far includes three species—*A. asiatica* Senior-White (from Sri Lanka) (Senior-White 1922), and *A. grandispinosa* Xu & Wu and *A. sinensis* Xu & Wu (from China, Zhejiang) (Xu & Wu 2002). There are consistent differences in color and shape of the terminalia sclerites of the species from China that allow to safely describe the species from Singapore as new. We examined an “allotype” of *Azana asiatica* at the NHM, which is basically brown in color—possibly not conspecific with the holotype, referred to as yellowish in the original description (Senior-White 1922).

*Azana demeijeri* Amorim & Oliveira, sp. n. (Figs. 9A-I)

https://singapore.biodiversity.online/species/A-Arth-Hexa-Diptera-000762

urn:lsid:zoobank.org:act:7FB79B40-B007-4AAF-8792-9DBD7328BCD9

**Diagnosis**. Abdomen cream-yellow to ochre-yellow, with brown transverse marks on posterior half of tergites 3–6. Male gonostylus with a bifid end at tip, outer branch strongly sclerotized. Female gonocoxite 8 with short medial incision on posterior border, lobes more rounded. ***Azana_demeijeri_ZRCBDP0048550_hapZRCBDP00485 48_SMH_holotype* [44: A, 55: A, 148: C, 173: A, 184: A, 202: C, 205: C, 229: C]**

tttatcatctactatcgctcatgcaggagcatcagtagatttaActattttttcAttacatttagcaggaatttcttcaattttaggagctgtaaattttattacaactattattaatatacgatcccccggaatttcattcgatcgCatacctttatttgtttgatcagtaAtaattacagcAattttactccttttatcCctCccagttttagcaggagctattacCatattactaactgatcgaaatttaaatacttcattttttgaccctgcaggaggaggagatccaattctatatcaacatttattt

#### Description

**Male**. **Head** (Figs. 9B–C). Elliptical, higher than long. Vertex and frons cream-yellow, setose, ocellar triangle dark brown, epicranial suture evident, from mid ocellus to ventral margin of vertex. Three ocelli, lateral ocelli close to mid ocellus, mid ocellus about half width of lateral ocelli. Occiput cream-yellow. Eye with interommatidial setulae. Antennal scape and pedicel brownish-yellow, rounded, with small setulae, pedicel with some slightly longer setae dorsally; flagellomeres light brown, slightly shorter than wide, with scattered setae. Face cream-yellow, covered with short setae. Labella light yellow, well-developed, projected backwards. Five palpomeres, first three light brown, distal segment whitish-brown, palpomeres gradually longer; first palpomere hardly recognizable, third palpomere with sensorial setulae, no pit. **Thorax** (Figs. 9B–D). Scutum and scutellum homogenously dark cream-yellow. Pleural sclerites light cream-yellow, pleural membrane whitish-yellow. Thorax short, scutum arched, covered with scattered small setae, an irregular band of longer and shorter dorsocentrals. Scutellum with about four pairs of slightly stronger setae and additional smaller setae along posterior margin, setulae on disc. Prosternum present, setose. Antepronotum with some 6–7 setae, proepisternum large, setose, proepimeron bare. Anepisternum with 9–11 setulae, katepisternum small, bare. Mesepimeron wide, extending to ventral margin of thorax, bare. Laterotergite only slightly bulging, with 19 larger and smaller setae, suture at contact with mediotergite incomplete dorsally. Mediotergite slightly curved in profile, with two pairs of longer setae. Haltere whitish-yellow, knob darker, covered with setulae. **Legs**. Fore coxa brownish-yellow, mid and hind coxae light yellow; trochanters dark brown, femora, tibiae, and tarsi brownish-yellow. Tarsomere 1 more than twice length of second one in all legs; mid and hind tibiae with erect darker bristles along almost entire outer length, mid and hind tarsi with some slightly longer, darker setae. Fore tibia with a wide anteroapical depressed area covered with some setulae irregularly distributed and a regular comb of setae. Mid and hind tibial spurs about twice tibia width at apex, mid and hind internal spurs shorter than outer spur. Tarsal claws with a long median tooth. **Wing** (Fig. 9E). Membrane homogenously fumose, densely covered with microtrichia and macrotrichia. Sc short, ending free slightly beyond humeral vein. C ending almost halfway between R_5_ and M_1_. First sector of Rs transverse, short. R_1_ short, reaching C slightly before middle of wing; R_4_ absent; R_5_ reaching C well before wing apex; r-m perfectly longitudinal, well sclerotized. No trace of M_1_, connection on M_1+2_ to tip of r-m very basal at wing, basal third of medial vein (supposedly M_1+2_) hardly sclerotized, recognizable by a well-defined row of setae. No trace of M_4_. CuA well sclerotized, basal fifth of CuP visible, weak. All veins with dorsal macrotrichia, except Sc and first sector of Rs. **Abdomen**. Tergites 2–6 yellow with a brown medial transverse band, setose, slender, T1 and T7 yellow. Sternites 1–2 whitish-yellow, sternites 3–7 yellowish. Tergite and sternite 8 produced. **Terminalia** (Fig. 9F). Terminalia brownish-yellow. Gonocoxites well-developed, ending in a short peak slightly more projected than insertion of gonostylus. Gonostylus well-sclerotized, elongate, bifid distally. Parameres separate, elongate, with a comb of small spines distally.

**Female** (Fig. 9A). As male, except as follows. Wing length, 2.42; width, 0.98. **Terminalia** (Figs. 9G-H). Sternite 8 setose, trapezoid, posterior margin with a short medial wide incision. Tergite 8 large, tergite 9 short and wide. Cercomere 1 elongate, 1.6× length of cercomere 2.

#### Material examined

**Holotype**: male, ZRCBDP0048550, Nee Soon (NS1), swamp forest, Malaise trap, 04.apr. 12, MIP leg. (slide-mounted). **Paratypes**: 9 females, ZRCBDP0048548, Nee Soon (NS1), swamp forest, 23.may. 12, MIP leg.; ZRCBDP0048549, Nee Soon (NS1), swamp forest, 09.may. 12, MIP leg.; ZRCBDP0048691, Nee Soon (NS1), swamp forest, 04.apr. 12, MIP leg.; ZRCBDP0047941, Nee Soon (NS2), swamp forest, 20.nov.13, MIP leg. (slide-mounted); ZRCBDP0048551, Nee Soon (NS2), swamp forest, 13.jun. 12, MIP leg.; ZRCBDP0048554, Nee Soon (NS2), swamp forest, 23.jan.13, MIP leg.; ZRCBDP0048553, Nee Soon (NS2), swamp forest, 16.may. 12, MIP leg.; ZRCBDP0048555, Nee Soon (NS2), swamp forest, 18.apr. 12, MIP leg.; ZRCBDP0078992, Singapore, date range 2012-2018, MIP leg.

**Etymology**. The species epithet of this species honors the Dutch naturalist Johannes Cornelis Hendrik de Meijere (1866–1947), specialized in Diptera and Coleoptera systematics, who gave relevant contribution to the understanding of the Oriental diversity of flies.

**Remarks**. We have three haplotypes for the species, and different delimitation algorithms generate no conflict for the species delimitation (Fig. 9I). *Azana demeijeri* Amorim & Oliveira, **sp. n.** was found in the mangrove and in swamp forest samples.

***Azana leekongchiani* Amorim & Oliveira, sp. n. (**Figs. 10A-F) https://singapore.biodiversity.online/species/A-Arth-Hexa-Diptera-000800

urn:lsid:zoobank.org:act:A120091C-0DC6-43F0-99C4-F6A149B1036A

**Diagnosis**. Abdominal segments 2–6 ochre-brown to brown, with a slender ochre-yellowish transverse band along posterior margin. Male gonostylus complex, with three branches, inner branch more sclerotized towards tip, outer branch weakly sclerotized and more setulose. Female sternite 8 with deeper distal medial incision, genital fork with a pair of well characterized anterior flaps. ***Azana_leekongchiani_ZRCBDP0048268_hapZRCBDP0 048079_SMH_holotype* [44: A, 46: A, 76: A, 124: T, 142 : A, 173: A, 203: C, 205: T, 209: G]**

actatcttcaacaattgctcatgcaggagcatcagttgatttaAcAattttttctttacatttagcaggaatttcAtcaattttaggagctgtaaattttattacaactattattaatatacgTtctcctggaatttatttAgatcgaatacctttatttgtatgatcagtaAtaattacagctattttattattattatcaCtTcctGtattagcaggagctattactatattattaacagatcgaaatttaaatacttctttttttgatcctgcaggaggaggagatcctattttatatcaacatttattt

#### Description

**Male**. Wing length, 1. 76, width, 0.79. **Head** (Fig. 10B). Vertex and frons light brownish-yellow, **Thorax** (Fig. 10B). Scutum and scutellum dark cream-yellow, posterior fourth of scutum darker. Pleural sclerites light brownish-yellow. **Wing** (Fig. 10C). **Abdomen**. Tergite 1 light brownish-yellow, tergites 2-6 ochre-brown, with a slender ochre-yellowish transverse band along posterior margin, T7 brownish-yellow. **Terminalia** (Fig. 10D). Terminalia brownish-yellow. Sternite 9 present as an independent setose plate. Gonocoxite large, not projected beyond base of gonostylus. Gonostylus large, complex, well sclerotized, with three branches, ventral branch wider, with a group of setae along margin, medial branch weakly sclerotized, dorsal branch digitiform, more sclerotized. Aedeagus tubular, projected first dorsally and then distally, with a pair of digitiform basal projections ending as small hooks. Parameres well-developed, digitiform, widened distally, with a long comb of short spinules on inner margin distally. Cercus small, lobose.

**Female** (Fig. 10A). As males except as for the following. Wing length, 1,84; width, 0,84. **Terminalia** (Figs. 10E–F). Terminalia brownish-yellow. Sternite 8 wide, with a pair of projections with a posterior incision in between, a row of longer setae along distal margin, tergite 9 partially fused to tergite 8, slightly projecting distally at sides; tergite 10 produced, with a row of setae along posterior margin, sternite 10 triangular, with a number of distal setulae. Cercus 2-segmented, clearly separated, distal cercomere about one fourth of basal cercomere.

#### Material examined

**Holotype**: male, ZRCBDP0048268, Pulau Semakau (SMO2), old mangrove, 10.oct.13, MIP leg. (slide-mounted). **Paratypes** (7 males, 4 females). **Males**: ZRCBDP0048146, Sungei Buloh (SB1), mangrove, 02.oct.13, MIP leg.; ZRCBDP0048258, Sungei Buloh (SB1), mangrove, 09.oct.13, MIP leg.; ZRCBDP0048312, Sungei Buloh (SB1), mangrove, 19.jun.13, MIP leg.; ZRCBDP0049277, National University of Singapore (Uhall), 15.apr.15, MIP leg.; ZRCBDP0049296, National University of Singapore (Icube), 13.may.15, MIP leg.; ZRCBDP0049320, National University of Singapore (Uhall), 01.apr.15, MIP leg.; ZRCBDP0278186, Pulau Ubin (PU18), mangrove, 31.may.18, MIP leg. **Females**: ZRCBDP0048079, Sungei Buloh (SB1), mangrove, 04.set.13, MIP leg.; ZRCBDP0049121, Nee Soon (NS1), 24.dec.14, MIP leg. (slide-mounted); ZRCBDP0279131, Singapore, 26.apr.18, MIP leg.; ZRCBDP0284192, Singapore, date range 2012-2018, MIP leg.

**Etymology**. This species proudly honors Lee Kong Chian [1893–1967], a prominent Chinese businessman who founded the Lee Foundation. He poured his wealth into education and other philanthropic work in Singapore, investing between 1952 and 1993 over US$ 200 million to various causes with no conditions attached. He was the first Chancellor of the National University of Singapore (1962-1965). The Lee Kong Chian Museum of Natural History, of the National University of Singapore, is named after him.

**Remarks**. *A. leekongchiani* Amorim & Oliveira, **sp. n.** was collected in different kinds of environment in Singapore, including urban forests. There are two haplotypes (Fig. 9I). *A. leekongchiani* Amorim & Oliveira, **sp. n.** can be clearly separated from *A. demeijeri* based on the shape of the gonostylus and in the abdomen color pattern (Figs. 10A, D).

### Tetragoneurinae

There is a group of mycetophilid genera about which there has been scarce agreement in the literature about its placement within the family—*Tetragoneura* Winnertz, *Ectrepesthoneura* Enderlein, *Novakia* Strobl and *Docosia* Winnertz. They have been alternately placed in the Gnoristinae, in the Leiinae or in a higher taxon of their own. These genera are connected to a number of fossils, known from the Valanginian to the Campanian, along the Cretaceous (Blagoderov & Grimaldi 2004; Oliveira & Amorim 2021). Oliveira & Amorim’s (2021) extensive study of the Leiinae showed that they do not belong in the subfamily and that they need indeed a taxon of subfamily rank, the Tetragoneurinae, a name that was available in the literature. Of the four extant genera belonging to the subfamily, we found in the Singapore samples species of *Tetragoneura* and *Ectrepesthoneura*.

### Tetragoneura Winnertz

*Tetragoneura* Winnertz, 1846: 18. Type species: *Tetragoneura distincta* Winnertz, 1846 [*Sciophila sylvatica* Curtis, 1837], designated Johannsen 1909: 34.

**Diagnosis**. Flagellomeres about as long as wide. Three ocelli, lateral ocelli remote from eye margin. Laterotergite and mediotergite bare. C extending beyond tip of R_5_ at wing margin; R_1_ at most slightly longer than r-m; Sc incomplete, short; R_4_, when present, forming a small cell; M_4_ originating beyond level of origin of M_1+2_. Male terminalia flexed, ventral face often directed posteriorly.

*Tetragoneura* is a quite large genus, with nearly 140 described species, particularly diversified in the Neotropical region, with 79 described species, and the Australasian region, with 26 species. There are no previous records for the genus in the Oriental region. No comprehensive studies have been made so far for the genus worldwide and it is not possible to group the species described here with the species found elsewhere.

We found four species of *Tetragoneura* in the samples from Singapore. The haplotypes for these species are gathered in two species (mPTP) or four species (all other method and parameters) (Fig. 11). The morphology of the male terminalia of the four species allows a clear recognition of each of the species. Some of the New Zealand species of the genus also show long, curved blade-like parameres, with tips extending beyond the base of the gonostylus, also present in three species from Singapore described here, but absent in *Tetragoneura crawfurdi* Amorim & Oliveira, **sp. n.**

*Tetragoneura crawfurdi* Amorim & Oliveira, sp. n. (Figs. 12A–D)

https://singapore.biodiversity.online/species/A-Arth-Hexa-Diptera-000749

urn:lsid:zoobank.org:act:19F3266B-F025-41E5-AA77-9130C8D2A5A0

**Diagnosis**. Scutum yellowish. Male with anterior half of abdomen yellowish, brown marks only medially on tergites 2–4, tergites 5–6 blackish-brown; female with tergite and sternite 1 yellowish, segments 2–6 dark brown. Male syngonocoxite medially with a pair of slightly divergent posterior projections, gonocoxites laterally not projecting beyond base of gonostylus. Gonostylus with a pointed short projection on outer face, no long curved blades. Parameres short, not curved.

***Tetragoneura_crawfurdi_ZRCBDP0048499_hapZRCBD P0048499_SMH_holotype* [10: C, 45: G, 122: T, 126: T, 208: C, 212: C]**

tctttcagcCactattgcccactcaggagcatctgttgatttatGtattttctcattacatttagcaggaatttcctcaattttaggagccgttaattttatcacaacaattattaatataTattTaccagaaataaatatagataaaataccattatttgtttgatcagtatttattacagctattttacttctattatcattaccCgtaCtagctggagcaattacaatattattaacagatcgtaatttaaatacctcattttttgatcctgctggaggaggagatccaattctataccaacatctattt

#### Description

**Male** (Fig. 12A). Wing length, 1.81; width, 0.74. **Head**. Blackish-brown. Three ocelli, lateral ocelli separated from eye margin by distance much larger than its own diameter, mid ocellus at posterior end of frontal furrow. Clypeus brownish, with scattered short setae, palpus and labella whitish. Eyes densely covered by interommatidial setulae. Antennal scape and pedicel whitish-yellow, with a crown of brownish short setae and an additional conspicuous dorsal bristle; flagellomere 1 light yellowish-brown, other flagellomeres brown. Flagellomeres about as long as wide except first and last, slightly longer than wide. Mouthparts slightly projected. Maxillary palpus 5-segmented, second segment very short, third segment with a well-developed, deep sensorial pit, fourth segment slightly longer than third, distal segment almost twice longer than fourth, thin, weakly sclerotized. **Thorax**. Scutum and scutellum yellowish, scutellum yellowish-brown laterally. Pleural sclerites cream-yellowish, except for anepisternum, only slightly more brownish, dorsal half of mesepimeron, dorsal half of laterotergite and mediotergite, light brownish-yellow. Scutum with scattered brownish short setae and a line of nine conspicuous dorsocentrals, some longer setae along acrostichal line; some longer supra-alars; scutellum with one pair of strong setae and one pair of smaller setae closer to lateral margins. Antepronotum and proepisternum with bristles and setae, other pleural sclerites bare. Haltere mostly yellowish, light brown at base of knob; some setulae along pedicel. **Wing** (Fig. 12B). Wing largely greyish-brown, membrane brown on distal third of wing, at tip of costal cell, on cell r1, and along second sector of CuA on cell cua. C extending over almost ¾ of distance between R_5_ and M_1_. Sc short, ending free. R_4_ present. R_1_ longer than r-m; r-m longitudinal on most of its length, slightly curved basally towards posterior margin, aligned to second sector of Rs and to bM. M_1+2_ much shorter than medial fork. M_4_ originating slightly beyond level of origin of M_1+2_, with very gentle depression on distal half. CuA slightly sinuous beyond origin of M_4_. All veins with dorsal setae, except for Sc, first sector of Rs, R_4_, M_1+2_ and bM. No trace of CuP. **Legs**. Coxae whitish-yellow; femora light brownish-yellow, hind femur brownish at distal fifth; tibiae and tarsi greyish-brown. Fore tibia without bristles except at tip, mid and hind tibiae and tarsi with a long regular row of small dark dorsal bristles, scattered laterally. Anteroapical depressed area on inner face of fore tibia wide, lined with setulae. Tibial spurs dark brown, more than 4× width of tibiae at apex. Tarsal claw with a strong basal tooth. **Abdomen**. Tergites 1–3 yellowish, tergites 4–7 dark brown, sternites 1–2 yellowish, sternite 3 light brown, sternites 4–7 brown. **Terminalia** (Figs. 12C–D). Dark brown. Gonocoxites with deep V-shaped median cleft, reaching anterior third of terminalia; medial area of syngonocoxite more sclerotized, with a pair of projections densely covered with microtrichia. Gonostylus at tip of gonocoxites, well-developed, with three basal branches in addition to main branch densely covered with setae: a long, slender, bare blade, a pointed basal branch covered with setulae, and a short digitiform projection densely covered with setulae. Aedeagus short, hardly detectable. Aedeagal-parameral complex well sclerotized medially, with a pair of short lateral horns, no long, curved blades inside terminalia, each branch crossing towards the other side of terminalia but not extending beyond tip of aedeagus. Gonocoxal bridge wide, gonocoxal apodemes short, well separated. Tergite 9 weakly sclerotized, wide and short, not reaching posteriorly level of base of gonostylus. Cerci small, lobose, covered with microtrichia and some elongate fine setae.

**Female**. Unknown.

#### Material examined

**Holotype**: male, ZRCBDP0048499, Nee Soon (NS1), swamp forest, 27.jun. 12, MIP leg. (slide-mounted).

**Etymology**. This species is named after John Crawfurd FRS (1783–1868), a Scottish physician, with an important role in founding Singapore, the second and last British Resident of Singapore (as this position was later replaced by the Governor of the Straits Settlements). He was an important and efficient administrator, laying out many crucial foundations that ensured the growth of modern Singapore. He was also an author, known for his work on Asian languages and his History of the Indian Archipelago.

**Remarks**. This is the only species of *Tetragoneura* in Singapore that does not have the long, curved paramere blades crossing sides in the male terminalia. It is known only from the holotype.

*Tetragoneura chola* Amorim & Oliveira, sp. n. (Figs. 13A–D)

https://singapore.biodiversity.online/species/A-Arth-Hexa-Diptera-000809

urn:lsid:zoobank.org:act:2CFAC9C8-8441-4DF0-803C-ADB6E74C8C32

**Diagnosis**. Scutum yellowish. Males with anterior half of abdomen yellowish, with brown marks only medially on tergites 2–4, tergites 5–6 blackish-brown (female unknown). Male syngonocoxite medially without a pair of conspicuous posterior projections, gonocoxites laterally not projecting beyond base of gonostylus. Gonostylus with a long, curved slender blade that extends distally. Parameres long, curved and slender, crossing inside terminalia, ending with a brush-like concentration of setae. ***Tetragoneura_chola_ZRCBDP0047776_hapZRCBDP00 47776_SMH_holotype* [19: A, 43: T, 142: A, 146: A, 199: G, 208: A, 220: G, 251: C] -**

ctttcatcatcaattgcAcattcaggagcatcagtagatttTgctattttttctttacatttagcaggtatttcttcaattttaggagccgtaaattttattacaacaattattaatatacgatctcctggaataacaatAgatAaaataccattatttgtatgatcagtatttattacagctattttattacttctGtcattaccAgtattagcaggGgctattacaatattattaacagatcgaaacCtaaatacttcattttttgacccagctggaggaggagatccaattttatatcaacatttattt

#### Description

**Male** (Fig. 13A). Wing length, 1. 91–2.22; width, 0.79 (n=2). **Head**. Blackish-brown. Three ocelli, lateral ones separated from eye margin by distance of its own diameter, mid ocellus at posterior end of frontal furrow. Clypeus brownish with scattered short setae, palpus and labella whitish-yellow. Eyes densely covered by interommatidial setulae. Antennal scape and pedicel dirty-yellow, with a crown of blackish-brown short setae and one additional dorsal bristle, flagellomere 1 brownish-yellow, other flagellomeres dark brown. Flagellomeres about as long as wide, except last one, slightly longer. Mouthparts slightly projected. Maxillary palpus 5-segmented, second segment very short, third segment with a well-developed, deep sensorial pit, fourth segment slightly longer than third, distal segment almost twice longer than fourth, thin, weakly sclerotized. **Thorax**. Scutum yellowish, scutellum yellowish-brown. Pleural sclerites cream-yellowish, except for anepisternum, only slightly more brownish, dorsal half of mesepimeron and dorsal half of laterotergite and mediotergite, brownish. Scutum with scattered brownish short setae and a line of nine conspicuous dorsocentrals, some longer setae along acrostichal line; some longer supra-alars; scutellum with one pair of strong setae and one pair of smaller setae closer to lateral margins. Antepronotum and proepisternum with setae and bristles, other pleural sclerites bare, including laterotergite and mediotergite. Haltere mostly yellowish, light brown at base of knob. **Wing** (Fig. 13B). Wing largely greyish-brown, membrane brown on distal third of wing, at tip of costal cell, over cell m4, and along CuA on cell cua; veins dark brown except for M_1+2_, basal fourth of M_1_, bM and first section of CuA, which are weakly sclerotized. C extending over almost ¾ of distance between R_5_ and M_1_. Sc short, ending free. R_4_ present. R_1_ longer than r-m; r-m longitudinal on most its length, slightly curved basally towards posterior margin, aligned to second sector of Rs and to bM. M_1+2_ much shorter than medial fork. M_4_ originating slightly beyond level of origin of M_1+2_, with very gentle depression on distal half. CuA slightly sinuous beyond origin of M_4_. All veins with dorsal setae, except for Sc, first sector of Rs, R_4_, M_1+2_ and bM. No trace of CuP. **Legs**. Coxae whitish-yellow, femora light brownish-yellow, tibiae and tarsi greyish-brown, fore tibia without bristles except at tip, mid and hind tibiae and tarsi with some scattered small dark bristles. Anteroapical depressed area on inner face of fore tibia wide, lined with setulae. Tibial spurs dark brown, more than 4× width of tibiae at apex. Tarsal claw with long, strong basal tooth. **Abdomen**. Tergite 1 yellowish, tergites 2–3 yellowish with a medial longitudinal brown band, tergite 4 yellowish with a medial brown band that expands laterally at distal margin, tergites 5–7 dark brown, sternite 1–4 yellowish, sternite 5 light brownish, sternite 6–7 dark brown. **Terminalia** (Figs. 13C–D). Dark brown. Gonocoxites with deep median V-shaped cleft, reaching anterior third of terminalia; a small area bulging on each gonocoxite with a group of concentrated setulae. Gonostylus relatively small, bifid from base, one branch straight with setulae, other branch long, curved projecting medially on basal half and posteriorly on distal half. Aedeagus short, hardly detectable. Parameres extremely long, with a pair of curved slender blades inside the terminalia, each branch crossing towards the other side of terminalia, tip of blades not reaching level of tip of syngonocoxite projections. Gonocoxal bridge wide, gonocoxal apodemes short, well separated. Tergite 9 weakly sclerotized, wide and short, not reaching posteriorly level of base of gonostylus. Cerci small, lobose, covered with microtrichia and some elongate fine setae.

**Female**. Unknown.

#### Material examined

**Holotype**: male, ZRCBDP0047776, Nee Soon (NS1), swamp forest, 28.aug.13, MIP leg. (slide-mounted). **Paratypes**: 5 males: ZRCBDP0047848, Nee Soon (NS1), swamp forest, 26.jun.13, MIP leg.; ZRCBDP0048501, Nee Soon (NS1), swamp forest, 20.jun. 12, MIP leg.; ZRCBDP0049175, Nee Soon (NS2), 13.may.15, MIP leg. (extracted); ZRCBDP0049186, Nee Soon (NS2), 13.may.15, MIP leg.; ZRCBDP0155113, Singapore, NSM2, 25.feb.15, MIP leg. (slide-mounted).

**Etymology**. This species is named after Rajendra Chola I, a Tamil Chola emperor of South India, who succeeded his father to the throne in 1014 CE. He greatly extended the influence of the Chola empire – North through the banks of the river Ganga, South to Sri Lanka, and East through South East Asia, conquering much of the Malay Peninsula (including Singapore) and parts of Indonesia. There are no records, however, of his visit to the island of Singapore itself. The noun is used in aposition.

*Tetragoneura dayuan* Amorim & Oliveira, sp. n. (Figs. 14A–F)

https://singapore.biodiversity.online/species/A-Arth-Hexa-Diptera-000751

urn:lsid:zoobank.org:act:C0B23FEA-A178-4EFA-8767-2E7747A01A70

**Diagnosis**. Scutum yellowish. Male with anterior half of abdomen yellowish, brown marks only medially on tergites 2–4, tergites 5–6 blackish-brown; female with tergite and sternite 1 yellowish, segments 2–6 dark brown. Male syngonocoxite medially with a pair of parallel posterior projections, gonocoxites laterodistally with a short projection beyond base of gonostylus. Gonostylus simple, without long, curved blade or pointed projections. Parameres long, curved inside the terminalia, crossing medially, without a distal brush.

***Tetragoneura_dayuan_ZRCBDP0048504_hapZRCBDP 0048504_SMH_holotype* [31: C, 83: C, 140: A, 142: A, 1 62: C, 178: C, 211: G]**

actgtcttcatctattgcccattcaggagcCtcagttgatttagctattttttccctccatttagccggaatttcttctattCtaggagctgtaaattttattacaacaatcattaatatacgatctcctggaataacaAtAgataaaataccactatttgCatgatcagtatttatCacagcaattttattacttttatctttacctgtGctagccggagctattacaatattattaacagaccgtaatttaaatacatccttctttgatcctgctggagggggagatccaattttatatcaacatttattt

#### Description

**Male** (Fig. 14A). Wing length, 2.17; width, 0.89. **Head** (Fig. 14B). Blackish-brown. Three ocelli, lateral ones separated from eye margin by distance much larger than its own diameter. Clypeus light brown, with scattered short setae, palpus whitish-yellow, labella whitish. Eyes densely covered by interommatidial setulae. Antennal scape and pedicel whitish-yellow, with a crown of brownish short setae and an additional conspicuous dorsal bristle; flagellomere 1 light yellowish-brown, other flagellomeres brown. Flagellomeres about as long as wide, first flagellomere 1 1. 5× length of flagellomere 2. Clypeus extended into a short proboscis. Maxillary palpus 5-segmented, second segment very short, third segment with a well-developed sensorial pit, fourth segment slightly longer than third, distal segment almost twice longer than fourth, thin, weakly sclerotized. **Thorax** (Fig. 14C). Scutum yellowish, slightly darker posterior to transverse suture, with scattered blackish short setae, some supra-alars and scattered stronger setae, scutellum yellowish-brown laterally. Pleural sclerites cream-yellowish, except for a dirty-yellow anepisternum, laterotergite mostly brownish, yellowish only ventrally, mediotergite brownish on dorsal half. Antepronotum and proepisternum with some scattered thin setae, antepronotum with three bristles, proepisternum with two bristles along ventral margin. Other pleural sclerites bare, including laterotergite and mediotergite. **Legs**. Coxae whitish-yellow; trochanters with brownish marks, fore and mid femora whitish-yellow, hind femora cream-yellowish with a brown mark close to tip, fore tibia and tarsus light greyish-brown, mid tibia and tarsus greyish-brown, hind tibia and tarsus brownish. Fore coxa with internal face covered with fine setae, mid coxa covered with fine setae on distal half of internal face, hind coxa with a regular row of fine setae along mid of external face. Fore tibia without bristles except at tip, mid and hind tibiae and tarsi with long regular rows of small dark dorsal and lateral bristles. Anteroapical depressed area on inner face of fore tibia wide, lined with setulae. Tibial spurs dark brown, about 3× width of tibiae at apex. Tarsal claw with strong basal tooth. Haltere mostly yellowish, light brown at base of knob. **Wing** (Fig. 14D). Wing largely greyish-brown, membrane brown on distal fourth of wing, at tip of costal cell, over cell m4, and along CuA on cell cua; veins dark brown except for basal fourth of M_1_, M_1+2_ and bM. R_4_ present. C extending over ¾ of distance between R_5_ and M_1_. Sc short, ending free. R_1_ longer than r-m; r-m longitudinal on most its length, slightly curved basally towards posterior margin, aligned to second sector of Rs and to bM. M_1+2_ much shorter than medial fork, M_2_ slightly curved anteriorly on distal half. M_4_ originating slightly beyond level of origin of M_1+2_, with very gentle depression on distal half. CuA slightly sinuous beyond origin of M_4_. All veins with dorsal setae, except for Sc, first sector of Rs, R_4_, M_1+2_, and bM, three setae on a weakly sclerotized CuP. **Abdomen**. Tergite 1–3 yellowish, tergite 4 brown with yellowish longitudinal lateral bands, tergites 5–7 dark brown, sternites 1–3 yellowish, sternite 4 light brown, sternites 4–7 brown. **Terminalia** (Figs. 14E– F). Dark brown. Gonocoxites with V-shaped deep median cleft, reaching anterior third of terminalia; medial part of posterior margin of syngonocoxite with a pair of distal projections with a number of fine setulae; part of posterior margin of syngonocoxite at each side of these projections slightly bulging, with a group of concentrated setulae; gonocoxite with a distal-posterior extension beyond base of gonostylus. Aedeagus short, weakly sclerotized, triangular distally. Aedeagal-parameral complex, with a pair of elongate, curved slender blades inside terminalia, each branch curved basally and then crossing towards other side of terminalia, extending almost to level of tip of medial syngonocoxite projections. Gonocoxal bridge wide, gonocoxal apodemes short, well separated. Tergite 9 weakly sclerotized, wide and short, not reaching posteriorly level of base of gonostylus. Cerci small, lobose, covered with microtrichia and some elongate fine setae.

**Female**. Unknown.

#### Material examined

**Holotype**: male, ZRCBDP0048504, Nee Soon (NS2), swamp forest, 26.Sep.12, MIP leg. (slide-mounted).

**Etymology**. This species is named after Wang Dayuan [=汪大渊] (1311-1350), a 14th Century Yuan Dynasty Chinese traveler who visited Singapore around 1330. His book, Dao Yi Zhi Lue [=岛夷志略; A Brief Account of Island Barbarians], is one of the few records documenting life in Dan Ma Xi [=淡马锡; transcription of Malay Temasek], a small settlement in early Singapore. The noun is used in apposition.

*Tetragoneura farquhari* Amorim & Oliveira, sp. n. (Figs. 15A–D, 16A–D)

https://singapore.biodiversity.online/species/A-Arth-Hexa-Diptera-000810

urn:lsid:zoobank.org:act:7B3540D1-8C5F-4124-A3DC-51717EE7A471

**Diagnosis**. Scutum yellowish, with a brownish mark above wing base. Male with anterior half of abdomen yellowish, brown marks only medially on tergites 2–4, tergites 5–6 blackish-brown; female with tergite and sternite 1 yellowish, segments 2–6 dark brown. Male syngonocoxite medially with a pair of posterior projections distally rounded, gonocoxites laterodistally with a short, pointed projection beyond base of gonostylus. Gonostylus simple, with a pointed basal short projection on outer face. Parameres not crossing medially, without a distal brush.

***Tetragoneura_farquhari_ZRCBDP0048710_hapZRCBD P0048500_SMH_holotype* [7: A, 11: T, 46: T, 61: C, 67: C, 115: C, 140: A, 142: A, 184: C, 202: T]**

actttcAtcaTctatcgcccattcaggagcttctgttgatttggcTattttttctcttcaCttagcCggaatttcttctattttaggagccgtaaattttattacaacaattatCaatatacgatctccaggaataacaAtAgataaaatacctttatttgtatgatctgtatttattacagcCattttattacttttatcTttacctgtattagctggagctattacaatattattaacagatcgtaatttaaatacctcattttttgacccagctggaggaggagaccctattttatatcaacatttattt

#### Description

**Male**. Wing length, 2.14–2.30; width, 0.89 (n=2). **Head** (Fig. 15B). Blackish-brown. Three ocelli, lateral ocelli separated from eye margin by distance much larger than its own diameter. Clypeus light brown, with scattered short setae, palpus whitish-yellow, labella whitish. Eyes densely covered by interommatidial setulae. Antennal scape and pedicel whitish-yellow, with a crown of brownish short setae and an additional conspicuous dorsal bristle; flagellomere 1 light yellowish-brown, other flagellomeres brown. Flagellomeres about as long as wide, except first and last, slightly longer than wide. Clypeus extended into a short proboscis. Maxillary palpus 5-segmented, second segment very short, third segment with a deep sensorial pit, fourth segment slightly longer than third, distal segment almost twice longer than fourth, thin, weakly sclerotized. **Thorax** (Fig. 15C). Scutum yellowish except for a brownish mark above wing base, scutellum brownish-yellow. Pleural sclerites cream-yellowish, except for a dirty-yellowish anepisternum, laterotergite brownish dorsoposteriorly, mediotergite brownish on dorsal half, dirty-yellowish on ventral half. Antepronotum and proepisternum with some scattered thin setae, antepronotum with three bristles in line, proepisternum with two bristles along ventral margin. Other pleural sclerites bare, including laterotergite and mediotergite. **Legs**. Coxae whitish-yellow; trochanters with brownish marks; fore and mid femora yellowish, hind femur yellowish with a brown mark close to tip, fore tibia and tarsus light brown, mid tibia and tarsus brown, hind tibia and tarsus dark brown. Fore coxa with internal face covered with fine setae, mid coxa covered with fine setae on distal half of internal face, hind coxa with a regular row of fine setae along mid of external face. Fore tibia without bristles except at tip, mid and hind tibiae and tarsi with long regular row of small dark dorsal and lateral bristles. Anteroapical depressed area on inner face of fore tibia wide, lined with setulae. Tibial spurs dark brown, about 3× width of tibiae at apex. Tarsal claws with a strong basal tooth. Haltere mostly yellowish, light brown at base of knob. **Wing** (Fig. 15D). Wing membrane brown on distal fourth of wing, at distal third of costal cell, over cell m4, and along CuA on cell cua; veins brown except for basal fourth of M_1_, M_1+2_ and bM. R_4_ present. C extending over ⅔ of distance between R_5_ and M_1_. Sc short, ending free. R_1_ longer than r-m; r-m longitudinal on most its length, gently curved basally towards posterior margin, aligned to second sector of Rs and to bM. M_1+2_ much shorter than medial fork, M_2_ gently curved anteriorly on distal half. M_4_ originating slightly beyond level of origin of M_1+2_, with a very gentle depression on distal half. CuA slightly sinuous beyond origin of M_4_. All veins with dorsal setae, except for Sc, first sector of Rs, R_4_, M_1+2_ and bM. CuP absent. **Abdomen**. Tergite 1 yellowish, tergites 2–3 yellowish with a brownish medial area, tergite 4 brown with yellow areas along anterior margin and along lateral margin, tergites 5–7 dark brown, sternites 1–3 yellowish, sternite 4 light brown, sternites 4–7 brown. **Terminalia** (Figs. 16A– B). Dark brown. Gonocoxites connected on anterior half of terminalia; medial part of posterior margin of syngonocoxite with a pair of distal projections with fine setulae; gonocoxite with a distal pointed short projection beyond base of gonostylus. Aedeagus short, weakly sclerotized. Aedeagal-parameral complex with a pair of elongate, curved slender blades inside terminalia, each branch curved basally and then extending distally almost to level of tip of medial syngonocoxite projections, not crossing to other half of terminalia. Gonocoxal bridge wide, gonocoxal apodemes short, well separated. Tergite 9 weakly sclerotized, wide and short, not reaching posteriorly level of base of gonostylus. Cerci small, lobose, covered with microtrichia and some elongate fine setae.

**Female** (Fig. 15A). As male, except for the following. **Wing l**ength, 2.14; width, 0.82. **Abdomen**. Tergites brownish, sternites brownish-yellow, distal segments darker. **Terminalia** (Figs. 16C–D). Sternite 8 with a pair of elongated lobes distally, connected along anterior half; no chambers on lobes, scattered microtrichia on lobes, some setulae on distal half of lobes, longer setae along posterior margin. Sternite 9 (vaginal furca) weakly sclerotized, hardly visible, without typical Y-shape. Sternite 10 weakly sclerotized, hardly visible. Tergite 8 with a row of fine setae along posterior margin. Tergite 9 short, slender, with a row of elongate fine setae. Tergite 10 short, slender, with a single row of fine elongate setae. Cercus 2-segmented, cercomere 1 about 3× length of cercomere 2, tip of cercomere 1 projecting slightly beyond insertion of cercomere 2.

#### Material examined

**Holotype**: male, ZRCBDP0048710, Nee Soon (NS2), 28.jan.15, MIP leg. (extracted, slide-mounted). **Paratypes**: 4 males, 2 females. **Males**: ZRCBDP0048500, Nee Soon (NS1), swamp forest, 25.apr. 12, MIP leg.; ZRCBDP0048716, Nee Soon (NS2), 28.jan.15, MIP leg.; ZRCBDP0048739, Nee Soon (NS1), 25.feb.15, MIP leg.; **Females**: ZRCBDP0048502, Nee Soon (NS1), swamp forest, 23.may. 12, MIP leg.; ZRCBDP0048503, Nee Soon (NS1), swamp forest, 04.apr. 12, MIP leg. (slide-mounted). **Sequence failure**: male, ZRCBDP0154852, Singapore, MIP leg. (slide-mounted, sequence failure) (MZUSP).

**Etymology**. The species name honors William Farquhar, a Scottish employee of the East India Company, who was the first British Resident and Commandant of Singapore. He was designated by Raffles to manage the colony of Singapore according to specific plans set in the period from 1819 to 1823.

**Remarks**. Specimens of this species came from both, the swamp forest and the mangroves. This species is closely related to *Tetragoneura dayuan* Amorim & Oliveira, **sp. n.** as can be seen in different details of the male terminalia. We have two haplotypes for this species, but no delimitation conflicts.

### Ectrepesthoneura Enderlein

*Ectrepesthoneura* Enderlein 1911: 155. Type species *Tetragoneura hirta* Winnertz 1846: 19, by original designation and monotypy.

*Ectrepesthoneura* has about 20 described species worldwide, most of which from the Nearctic and Palearctic regions. The genus has two Cretaceous and two Cenozoic fossil species described (Blagoderov & Grimaldi 2004). This is the first non-Holarctic species of the genus. The long Sc ending free is a quite unique feature of this species within *Ectrepesthoneura*, the Holarctic species having Sc fused to R_1_. Only samples of the Nee Soon swamp forest had specimens of this species.

**Diagnosis**. Three ocelli, lateral ocelli remote from eye margin. Maxillary palpomeres 2 and 3 more or less swollen. Laterotergite and mediotergite setose. C produced beyond tip of R_5_;Sc long, ending free or fused to R_1_ distally; R_4_ present; first section of CuA very short, M_4_ originating close to wing base. Male terminalia flexed, ventral face directed posteriorly.

*Ectrepesthoneura johor* Amorim & Oliveira, sp. n. (Figs. 17A–J)

https://singapore.biodiversity.online/species/A-Arth-Hexa-Diptera-000810

urn:lsid:zoobank.org:act:FFFFE4DD-25CA-4ED3-8034-4BC10E2079D8

**Diagnosis**. Head dark brown. Male mid tibia dorsally with a sensory organ at basal fourth. Scutum ochre-yellowish, thoracic pleural sclerites greyish brown. Abdominal tergite 1 cream-yellowish, tergites 2–7 greyish-brown. Male gonocoxites with a short pointed blade directed inwards medially on posterior margin of ventral face; gonostylus small, spine-like distally; tergite 9 largely developed. ***Ectrepesthoneura_johor_ZRCBDP0047876_hapZRCBD P0047876_SMH_holotype* [104: T, 170: A, 172: C, 182: A, 187: C, 238: T]**

cctttcatctacaattgcccattcaggggcatctgttgatttagcaattttttcacttcatttagcaggtatctcctctattttaggagcagtaaattttattTcaacaattattaatatacgtgcccccggaatttcatttgaccgaatacctttatttgtttgatctAtCttaattaccAcaatCttacttttgctttctttacctgttttagccggagctatcactatacttctTacagatcgaaacttaaatacttccttttttgatccagctggaggaggtgatcctattctataccaacacctattt

#### Description

**Male** (Fig. 17A). Wing length, 1.89; width, 0.82. **Head** (Fig. 17C). Blackish-brown. Three ocelli, lateral ocelli separated from eye margin by distance twice its own diameter. Clypeus light brown, with scattered setae, palpus dirty whitish-yellow, labella whitish. Antennal scape and pedicel whitish-yellow, flagellomere 1 dirty-yellowish, remaining flagellomeres light ochre-brownish; scape and pedicel with a crown of short setae around distal margin, one strong dorsal bristle at distal margin of pedicel. Flagellomeres slightly wider than long, except first and last, slightly longer than wide. Clypeus bulging, covered with setulae and some few longer setae, not projecting into a proboscis. Maxillary palpus 5-segmented, basal palpomere slightly elongate, second very short, third well-developed, widening towards apex, projecting beyond base of next flagellomere, a shallow sensorial pit on basal half at inner face and a shallow depression on distal half, tapering end; second palpomere slightly longer than first, third palpomere about twice longer than second. **Thorax** (Fig. 17D). Scutum ochre-yellowish, with a blackish pre-sutural mark along anterior margin medially, dark marks posteriorly to transverse suture and above wing. Small setae scattered over scutum, some scattered stronger supra-alars and a pair of regular rows of strong dorsocentrals. Scutellum brown, yellowish laterally, a pair of strong scutellars and some additional fine setae. Antepronotum blackish on anterior half, cream-yellowish on posterior half, anterior margin of proepisternum blackish, remaining sclerite cream-yellowish, anepisternum, katepisternum and mesepimeron with cream-yellowish diffuse marks over a brownish background, laterotergite and metepisternum brown, mediotergite mostly brownish yellowish-brown laterally. Antepronotum with a row of three bristles and some fine setae, proepisternum with fine setae and one strong seta close to ventral margin and another one more dorsally; other pleural sclerites bare. **Legs**. Coxae whitish-yellow, some light brown tinge at anterior tip; trochanters with brownish marks, femora whitish-yellow, tibiae and tarsi light greyish-brown, mid and hind tibiae and tarsi slightly darker. Fore coxa with scattered setae frontally and on external face, mid coxa with setae on distal half of external face and along entire anterior face, hind coxa with a regular row of setae medially on external face. Hind femur slightly wider medially. Fore tibia without bristles except at tip, mid tibia with irregular rows of small dark bristles on external, dorsal and internal faces, hind tibia with irregular rows of small bristles on external and dorsal faces. Tarsomeres 1–2 with a row of ventral stronger setae and a group of setae at tip, tarsomeres 3–4 with a pair of distal setae, only setulae on distal tarsomere. Anteroapical depressed area on inner face of fore tibia wide, lined with setulae. Tibial spurs dark brown, about 3× width of tibiae at apex. A long basal tooth on tarsal claw. Haltere mostly yellowish, light brown at base of knob. **Wing** (Fig. 17E). Wing light greyish; anterior veins brown, posterior veins with a light brownish tinge, M_1+2_ and basal fourth of M_1_ weakly sclerotized. C extending over ¾ of distance between R_5_ and M_1_. Sc long, ending free, bare. R_1_ slightly longer than r-m; R_4_ present; r-m longitudinal on most its. R_5_ short, ending at level of tip of M_2_. M_1+2_ less than a third medial fork length, M_4_ originating from CuA slightly beyond level of humeral vein, CuP weakly sclerotized but produced on basal third. Dorsal macrotrichia on R_1_, R_5_, r-m, distal ⅔ of M_1_, M_2,_ M_4_, CuA and CuP. **Abdomen**. Tergite 1 yellowish-brown, tergites 2–7 brownish, sternites 1–3 yellowish with light brown diffuse areas, sternites 4–7 light brown. **Terminalia** (Figs. 17F–G). Brownish-yellow. Gonocoxites in contact medially only at anterior end of terminalia, elongate, posterior margin with a pointed blade-like extension at each side crossing medially, two other pointed projections medially and a longer projection external to base of gonostylus. Gonostylus small, strongly sclerotized on distal half, falciform, distal end curved outwards, a few fine setulae on basal half, articulating at inner end of distal margin of gonocoxite. Gonocoxal bridge conspicuous, with well-developed apodemes. Aedeagal-parameral complex well-developed, aedeagus with an elongate apodeme anteriorly, a pair of distal projections and a short medial projection with a rounded tip. Tergite 9 wide, covering entirely dorsal face of terminalia, with a V-shapes medial incision on posterior margin. Cerci delicate, short, weakly sclerotized, with lateral apodemes extending anteriorly.

**Female** (Fig. 17B). As male, except for the following. **Terminalia** (Figs. 17H–I). Sternite 8 wide, without lobose distal projections, medially along posterior margin with an inverted triangular depression densely covered with microtrichia and fine setulae. Sternite 9 without Y-shape sclerotized area, gonopore on a more sclerotized area. Tergite 8 wide, short, weakly sclerotized, only with few fine setulae. Tergites 9 and 10 separate, weakly sclerotized except for band along anterior margin, with microtrichia and setulae. Cercus 2-segmented, basal cercomere much longer than second.

#### Material examined

**Holotype**: male, ZRCBDP0047876, Nee Soon (NS1), swamp forest, 19.jun. 2013, MIP leg. (extracted, slide-mounted). **Paratypes** (1 male, 1 female). **Male**: ZRCBDP0048505, Nee Soon (NS1), swamp forest, 25.apr. 12, MIP leg. (slide-mounted) (MZUSP). **Female**: ZRCBDP0048506, Nee Soon (NS2), swamp forest, 15.aug. 2012, MIP leg. (slide-mounted).

**Etymology**. The species name refers to Johor [sometimes Johor-Riau or Johor-Riau-Lingga], the name of the Sultanate founded in 1528 by Sultan Alauddin Riayat Shah II (son of the Malaccan Sultan Mahmud Shah), originally part of the Malaccan Sultanate, before Portugal’s 1511 conquer of Malacca’s capital. At its height, the sultanate included areas of modern-day Malaysia and Indonesia, as Johor, Riau, Muar, Batu Pahat, etc., besides Singapore. The noun is used in aposition.

**Remarks**. There is a single haplotype for *Ectrepesthoneura johor* Amorim & Oliveira, **sp. n.** in our samples (Fig. 17J).

### Leiinae

Oliveira & Amorim (2021) published the only phylogenetic study of the Leiinae including all genera of the subfamily as well as a large number of genera from other subfamilies. *Allactoneura* de Meijere joins *Sticholeia* Søli and the manotine genera in a clade deeply nested within the Leiinae. Our Singapore samples of the leiines include the genera *Mohelia* Matile, *Allactoneura*, *Eumanota* Edwards, *Manota* and *Clastobasis*. Comments are added individually to these genera. There is a record of *Indoleia* Edwards for Java (Ulimah et al. 2021) associated to citrus plantation and it is possible that the genus comes to be found in Singapore.

### Mohelia Matile

*Mohelia* Matile 1979a: 270. Type-species, *M. nigricauda* Matile (orig. desig.).

**Diagnosis**. Labrum elongated, almost twice length of clypeus, triangular, labella also elongated, almost as long as head height. R_1_ about as long as r-m, r-m almost longitudinal, laterotergite with setae, mediotergite bare.

*Mohelia* was created by Matile (1979a) for a single Afrotropical species, *M. nigricauda* Matile, while three additional Afrotropical species were added more recently by Oliveira (2015). *Mohelia* was considered by Matile (1979a) to be related to the strictly Neotropical genus *Aphrastomyia* Coher & Lane, and the more cosmopolitan *Megophthalmidia* Dziedzicki. This latter genus has seven species described from the Neotropical region, nine from the Nearctic and ten from the Palearctic (Bechev 1999; Chandler et al. 2005; Oliveira & Amorim 2014; Kerr 2014). Jaschhof & Kallweit’s (2004) understanding about this group was similar to Matile’s (1979a) position and reinforced the possible relationship between *Aphrastomyia* and *Mohelia*. Kerr (2014) made a careful revision of the Nearctic species of *Megophthalmidia* and also mentioned its similarities with *Aphrastomyia*. A key for including these three genera, *Aphrastomyia*, *Mohelia,* and *Megophthalmidia*, was published by Oliveira & Amorim (2021).

The species from Singapore described here unquestionably belongs to the clade (*Megophthalmidia* + (*Mohelia* + *Aphrastomyia*)). All species of *Aphrastomyia* have an obvious sclerotization of the posterior border of the syngonocoxite ventrally, not seen in the Singaporean species. *Megophthalmidia* has a flat and elongate syngonocoxite and a largely modified tergite 9 (Kerr 2014), both features absent in *Mohelia zubirsaidi* Amorim & Oliveira, **sp. n.** The Afrotropical species of *Mohelia* have simpler terminalia, but with a general shape and details on most sclerites that differ from the species from Singapore, except for the simple, band-like tergite 9 (Oliveira 2015). The problem of the limits between *Aphrastomyia* and *Mohelia* was already in need of solution, which is beyond the scope of this paper. Some features of this Singaporean species suggest that its inclusion in *Mohelia* may be the best solution for the time being—as the presence of a short proboscis, the shape of some of the wing veins etc.

In Oliveira & Amorim’s (2021) phylogeny, the clade with *Aphrastomyia*, *Mohelia* and *Megophthalmidia* belongs in the Leiinae, but too many nodes distant from the base of the subfamily. The presence of *Mohelia* in Southeast Asia results in another case of an Afro-oriental distribution pattern (see, e. g., Amorim & Tozoni 1995; Matile 1999), as also seen, e. g., in the genus *Platyprosthiogyne* Enderlein.

***Mohelia zubirsaidi* Amorim & Oliveira, sp. n.** (Figs. 18A–F, 19) https://singapore.biodiversity.online/species/A-Arth-Hexa-Diptera-000815

urn:lsid:zoobank.org:act:2A8346FE-F3AC-4195-B9CC-4E96538BC7CE

**Diagnosis**. Scutum yellowish-brown, darker on anterior half, pleural sclerites mostly dark ochre-yellowish. Tergite 1 yellowish with anteromedial brown mark, tergites 2–6 brownish with a cream-yellow band at posterior fourth. Gonocoxites with a deep V-shaped medial cleft; gonostylus small, with a pair of distal spines on apical projection; aedeagus long, coiled. Tergite 9 wide, short, weakly sclerotized.

***Mohelia_zubirsaidi_ZRCBDP0048984_hapZRCBDP004 8725_SMH_paratype* [12: A, 24: T, 34: C, 67: G, 113: T, 122: A, 123: A, 127: C, 262: G, 275: T]**

tttagcatcaaAtattgctcatgTtggagcttcCgttgatttagctattttttctcttcatttagcGggaatttcttcaattttaggagcagtaaattttattactacaattTttaatataAAaacCcctggaatttcatttgaccaaataccattatttgtttgatcagttttaatcacagcaattttattattactctctttaccagttttagctggagcaattactatactattaacagaccgaaatttaaatacatcGttctttgaccctTcaggaggaggagatccaattttatatcaacatttattc

#### Description

**Male** (Fig. 18A). Wing length, 1.48–1.63; width, 0.64–0.74 (n=2). **Head**. Dark ochre-yellowish, area across ocelli dark brown, face yellowish-brown, brown along laterals, clypeus cream-yellowish. Three ocelli, lateral ocelli separated from eye margin by about 1. 5× its own diameter, mid ocellus at posterior end of frontal furrow. Face with scattered short setae, clypeus cream-yellowish, palpus and labella whitish-yellow. Eyes densely covered by interommatidial setulae. Maxillary palpus with four palpomeres, third palpomere with sensorial pit, distal palpomere about 2× length of preceding palpomere. Antennal scape and pedicel whitish-yellow, with a crown of blackish-brown setae along distal border, with one stronger and longer dorsal setae. Flagellum light brownish-yellow, flagellomeres wider than long, last flagellomere slightly more projected ventrally than dorsally. Mouthparts slightly projected as a short proboscis. Maxillary palpus 5-segmented, second segment very short, third segment with a well-developed, deep sensorial pit, fourth segment slightly longer than third, distal segment almost twice longer than fourth, thin, weakly sclerotized. Labella well developed, directed backwards with pseudotrachaea. **Thorax**. Scutum yellowish-brown, darker on anterior half, with scattered brownish setae, supra-alars only slightly stronger. Scutellum yellowish-brown laterally. Pleural sclerites dark ochre-yellowish, with light brown maculae on anepisternum, katepisternum, laterotergite, and mediotergite. Antepronotum with some small setulae and three larger setae. Proepisternum well-developed, with 11– 13 smaller setae, 2–3 stronger setae along ventral margin.

Anepisternum, katepisternum, mesepimeron, metepisternum, and mediotergite bare, laterotergite with 4–8 setae. Mesepimeron not reaching ventral margin of thoracic pleura, katepisternum dorsoposteriorly in contact with laterotergite. Haltere pedicel yellowish-brown, knob brown. **Legs**. Fore coxa dirty-yellowish, mid and hind coxae whitish-yellow, femora dirty-yellowish, tibiae and tarsi greyish brown. Fore tibia without bristles except at tip, mid and hind tibiae with some few slightly longer setae dorsally and laterally, tibia without regularly arranged trichia; tarsomeres with some setae ventrally, particularly basal ones. Anteroapical depressed area on inner face of fore tibia wide, lined with setulae. Tibial spurs dark brown, more than 4× width of tibiae at apex. Tarsal claw with strong basal tooth. **Wing** (Fig. 19). Wing light brownish fumose, veins brownish. C extending over 4/5 of distance between R_5_ and M_1_. Sc short, ending free close to bR. R_1_ shorter than r-m; r-m longitudinal, aligned to second sector of Rs and to bM; R_4_ absent; R_5_ short, reaching C midway between level of tips of M_2_ and M_4_. M_1+2_ shorter than medial fork, M_1_ and M_2_ quite parallel along most their length, M_1+2_ shorter than medial fork, M_1_ slightly less sclerotized, especially at basal half, but not interrupted. M_4_ originating slightly beyond level of origin of M_1+2_, M_4_ and CuA unsclerotized close to margin; CuA unsclerotized at tip, not reaching wing margin. Cubital pseudovein present, no trace of CuP, anal fold absent. Macrotrichia dorsally on bR, R_1_, second sector of Rs, r-m, distal half of M_1_ and M_2_ and distal fourth of M_4_. **Abdomen**. Tergite 1 yellowish, brown anteromedially, tergites 2–6 brownish with a cream-yellow band at posterior fourth, wider at laterally. Sternites 1–7 dirty-yellowish, darker towards tip of abdomen. **Terminalia** (Figs. 18B–C). Brown. Gonocoxites large, occupying most of anterior face of terminalia, setose, fused medially on anterior half, with a deep V-shaped median cleft, which is slender anteriorly, gonocoxites with a pair of pointed projections at distal border ventrally and a lateroposterior projection reaching level of tip of gonostylus, from which a short blade projects inwards dorsally to gonostylus. Gonostylus small, complex, more or less rounded on basal two-thirds, with an apical slender projection bearing a pair of distal spines and a pair of additional short beaks. Gonocoxal bridge with a pair of long arms extending anteriorly, gonocoxal apodemes close together. Aedeagus long, partially coiled inside terminalia, projecting beyond tip of gonostylus and then curved ventrally. Tergite 9 wide, weakly sclerotized, not reaching level of base of gonostylus distally. Cerci lobose, covered with microtrichia and some elongate fine setae.

**Female**. As males, except for the following. **Wing l**ength, 1. 56; width, 0.64. **Abdomen**. Tergites yellowish-brown, darker laterally. Sternites yellowish, sternites 5–7 brownish medially on distal half. **Terminalia** (Figs. 18D– E). Sternite 8 with a pair of distal elongated lobes, connected along their anterior half; lobes with a small chamber on inner face densely covered by microtrichia; setulae scattered on ventral face of lobes, some longer setae along lobe posterior margin and on inner face. Sternite 9 (vaginal furca) weakly sclerotized, hardly visible, not with typical Y-shape. Sternite 10 weakly sclerotized, hardly visible. Tergite 8 separated from tergite 9, anterior border with sclerotized band extending laterally into apodemes directed ventrally, with scattered elongate fine setae. Tergite 9 short, slender, also with sclerotized band along anterior margin projecting laterally, with elongate fine setae. Tergite 10 short, slender, not fused to tergite 9, with a single row of fine elongate setae. Cercus 2-segmented, cercomere 1 about 1. 5× length of cercomere 2.

#### Material examined

**Holotype**: male, ZRCBDP0048725, Nee Soon (NS2), 15.apr. 2015, MIP leg. **Paratypes**: 8 males, 2 females. Males: ZRCBDP0048940, Nee Soon (NS2), 03.dec. 2014, MIP leg. (extracted, slide-mounted) (MZUSP); ZRCBDP0048951, Nee Soon (NS2), 03.dec. 2014, MIP leg.; ZRCBDP0048984, Nee Soon (NS2), 17.dec. 2014, MIP leg.; ZRCBDP0049011, Nee Soon (NS2), 17.dec. 2014, MIP leg.; ZRCBDP0049195, Nee Soon (NS2), 13.may. 2015, MIP leg.; ZRCBDP0072459, Bukit Timah, old secondary forest (BT07), 08.dec. 2016, MIP leg.; ZRCBDP0074038, Bukit Timah, primary forest (BT05), 08.dec. 2016, MIP leg.; ZRCBDP0074039, Bukit Timah, primary forest (BT05), 08.dec. 2016, MIP leg. Females: ZRCBDP0048999, Nee Soon (NS2), 17.dec. 2014, MIP leg. (slide-mounted); ZRCBDP0066748, Bukit Timah, maturing secondary forest (BT06), 28.Sep.2016, MIP leg. **Sequence failures**: male, ZRCBDP0133972, Singapore, date range 2012-2018, MIP leg. (slide-mounted); male, ZRCBDP0078990, Singapore, MIP leg.; male, ZRCBDP0133980, Singapore, MIP leg.; female, ZRCBDP0078997, Singapore, date range 2012-2018, MIP leg. female, ZRCBDP0134012, date range 2012-2018, Singapore, MIP leg.

**Etymology**. This species honors Zubir bin Said (1907-1987), a prolific Singaporean composer who wrote the national anthem of Singapore, “*Majulah Singapura*” in 1959. He was awarded the *Bintang Bakti Masyarakat* (Public Service Star) in 1963 by the Republic in recognition of his contributions in song and music and to Singapore.

**Remarks**. Specimens of this species are present in samples from the swamp forest and the tropical forest, with a single haplotype. Four of the specimens lack DNA barcodes due to sequencing failures.

### Allactoneura de Meijere

*Allactoneura* de Meijere 1907: 201. Type-species:

*Allactoneura cincta* de Meijere 1907: 202.

*Scottella* Enderlein 1910: 60.

**Diagnosis.** Hind margin of head with 30 or more long setae in arow on occiput behind eyes; 3 ocelli, median ocellus very small; palpus 4-segmented; flagellar segments cylindrical, closely appressed to each other. Thorax strongly flattened dorsoventrally; prosternum broad, reaching lateral surface of thorax; dorsal part of pronotum forming a slender strip sharply expanded laterally; mesonotum strongly flattened, covered with dense small setae and lanceolate scales; mediotergite bare. Wing membrane without microtrichia; C not extending beyond tip of R_5_; Sc ending at C; sc-r present; three longitudinal folds on wing, one across r-m, one posterior to M2, and one posterior to CuA. Male genitalia withdrawn into abdomen. Gonostylus elongate, attached to dorsolateral surface of gonocoxites. Aedeagus large, digitate. Female genitalia with long, stiff setae, cercus 1-segmented.

The genus *Allactoneura* was revised by Zaitzev (1982), who provided a key for the six species of the genus known to him. More recently, a species was added to the genus by Bechev (1995). *Allactoneura* is basically Afro-oriental in distribution, extending into New Guinea and northern Australia. In Oliveira & Amorim’s (2021) study of the phylogeny of the Leiinae, *Allactoneura* is sister of a clade with *Manota*, *Eumanota*, *Promanota* Tuomikoski, and *Paramanota* Tuomikoski.

*Allactoneura* has a particularly puzzling wing venation, with M_4_ detached from CuA, originating close to the wing base and a stump arising from Rs directed towards the wing base, which results in an intricate problem of homology. The stump, however puzzling, is only a secondary sclerotization of a fold going across r-m. In different manotine and leiine genera, for example, r-m originates longitudinally at the tip of the first sector of Rs, extending basally, becoming oblique at its proximal end. There is a careful discussion of the homology of the male terminalia sclerites of the genus in Oliveira & Amorim (2021).

Both species of *Allactoneura* in our samples would run into *A. nigrofemorata* de Meijere in Zaitzev’s (1982) key, which type locality (of a female) is Java and with a supposedly conspecific female known from Sumatra—no males are known of that species. The species from Singapore have clearly “abdomen with yellow transverse bands or yellow blotches on tergites”, so they do not run into *A. argenteosquamosa* Enderlein. The details of the male terminalia of *A. tumasik* Amorim & Oliveira, **sp. n.** are clearly different from those of the males of *A. cincta* de Meijere, *A. formosana* Enderlein, and *A. ussuriensis* Zaitzev. The antenna of *A. nigrofemorata* has the scape, pedicel, and first two flagellomeres yellow, while in our *A. tumasik* the antenna is darker, ochre-yellow. Also, in *A. nigrofemorata* the fore coxa is entirely whitish, with black femur. In *A. limbosengi* Amorim & Oliveira, **sp. n.** the base of the fore coxa is brown and the femora are entirely black; in *A. tumasik* Amorim & Oliveira, **sp. n.** the fore coxa is entirely yellow, with the mid femur black only at the distal third. The abdomen color pattern of *A. tumasik* Amorim & Oliveira, **sp. n.** concurs with *A. nigrofemorata*. The differences between both species from Singapore and the description of *A. nigrofemorata* are enough to accept that they are three separate species. Indeed, the situation of the genus is complex. Species with different body color patterns share a relatively similar male terminalia pattern. It would be useful to sequence specimens from different parts of Asia to check the molecular and morphological delimitation of the species of the genus. There are records in the literature of *A. argenteosquamosa* Enderlein, besides the type-series from Seychelles, for Sumatra (Fort de Kock), Thailand, Mauritius, Diego Garsia Island, India (Assam), and Sri Lanka. It is likely that some of the older records for *A. argenteosquamosa* correspond to misidentifications. Indeed, Bechev (pers.comm.) clarified that larvae of *A. akasakana* have been reported from shiitake mushroom indoor facilities in Japan and China (Sueyosh et al. 2019) and that *Allactoneura* sp. larvae were found on rotting wood, what suggests that some of the species of the genus may have a synanthropic association and could be widespread in Asia.

The haplotype network of *Allactoneura* (Fig. 21F) shows four haplotypes, with specimens from the subclusters in the swamp forest separated from a subcluster in the mangrove. PTP, mPTP etc. split them into separate species.

*Allactoneura tumasik* Amorim & Oliveira, sp. n. (Figs. 20A–B, 21A–E)

(https://singapore.biodiversity.online/species/A-Arth-Hexa-Diptera-000789)

urn:lsid:zoobank.org:act:5FC8095D-3B88-48AF-88FE-DE15DCF260DB

**Diagnosis**. Fore coxa entirely whitish, mid and hind coxae whitish with brown basal band. Fore femur and tibia whitish, mid femur whitish basally with a blackish-brown distal fourth, tibiae greyish-yellow. Male abdominal tergites 3–5 with cream-yellow marks on anterolateral corners. Gonostylus with a strong tooth on inner margin midway to apex, distally with a pair of short projections.

***Allactoneura_tumasik_ZRCBDP0048240_hapZRCBDP0 048240_SMH_holotype* [5: G, 12: A, 67: C, 70: T, 106: A, 140: T, 175: T, 184: A, 265: C, 286: A]**

tttaGctgctaAtattgctcatactggagcatcagttgatttagctattttttctttacatttagcCggTatttcttctattttaggtgcagtaaattttattacAacaattattaatatacgagctcctggaattagaTttgataaaatacctttatttgtttgatcagttctTattacagcAattttattattattatctttaccagtattagcaggagctattactatattattaacagatcgtaatttaaatacttcattCtttgatccagctggaggaggAgatcctattttataccaacatttattt

#### Description

**Male** (Fig. 20A). Wing length, 3.27, width, 1.02. **Head**. Vertex dark brown, with scattered setulae and some strong, black setae. Three ocelli in line, mid ocellus much smaller than lateral ocelli, lateral ocelli not close to eye margin. Occiput dark brown, with a crown of longer setae posteriorly to eye margin, at ventral end three stronger bristles at postgena, projecting laterally on posterior border. Ommatidia small, no interommatidial setulae. Scape and pedicel light brownish-yellow; scape elongate, with strong blackish setae on distal half; pedicel about as long as wide, with smaller, black setae on distal half; flagellomeres 1–5 ochre-yellow, flagellomeres 6–14 brownish, flagellomeres about as long as wide, with scattered setae. Frons blackish-brown, face dark brown, both with scattered setulae; clypeus longer than face, yellowish-brown, ventral margin lighter, densely covered with dark brown setae; labella whitish; palpomeres whitish, apical ones increasingly longer, last one about twice length of penultimate. Labella longer than length of head capsule, a row of 6–8 dark, long setae at anterior end. **Thorax** (Fig. 20B). Cervical sclerite robust. Scutum shinning dark brown, dorsally compressed, entirely covered with scattered setae, no bristles except for a pair of black, strong prescutellar bristles. Scutellum dark brown, with a pair of long bristles. Pleural sclerites brown. Pleural membrane yellowish. Antepronotum largely developed, projected towards proepisternum, covered with yellowish setulae. Prosternum shield-like, projecting above fore coxae anteriorly, with longer setae directed downwards at ventral margin in additional to more dorsal setulae. Proepisternum relatively small with some setulae. Proepimeron large, extending ventroposteriorly to articulate with anterodorsal corner of katepisternum and extending dorsally beyond anterior spiracle, ventral end of proepimeron with a fold. Anepisternum and katepisternum bare, anapleural suture incomplete; katepisternum slightly compressed dorsoventrally. Anepisternum wide, extending dorsoposteriorly over mesepimeron, posterior anepisternal sclerite well-developed. Mesepimeron bare, inclinate, not reaching ventral margin of thorax, dorsoposterior end of katepisternum fused to anteroventral end of laterotergite. Metepisternum short and wide, bare. Laterotergite bulging, with five bristles along outer margin dorsally in addition to smaller setae at dorsal surface. Mediotergite strongly curved in profile. Haltere whitish. **Legs**. Fore, mid, and hind coxae whitish-yellow, with a dark brown band basally and some brownish tinge at tip. Trochanters whitish-yellow with brownish areas; fore femur whitish-yellow, darker at tip, fore tibia whitish-yellow, darker basally at dorsal side; tarsi whitish-yellow densely covered by blackish setulae, giving a greyish appearance. Mid femur whitish-yellow at basal 4/5, dark brown at tip; hind femur entirely dark brown; mid and hind tibiae and tarsi light yellowish-brown, with dense coverage of blackish setulae. Mid and hind coxae twice wider than forecoxa. Fore leg tarsomere 1 slightly longer than fore tibia; tibiae and tarsi with erect dark short bristles along almost entire length, those on hind tibia more or less aligned dorsally and laterally. Tibial spurs very long, fore tibia spur yellowish-brown, mid and hind tibial spurs whitish, internal spurs slightly longer than inner spur. Mid and hind tibiae and tarsomeres 1-2 with short, spiny black setulae, tarsomeres 3-4 with a pair only with black setulae at distal margin. Tarsal claws with a large basal tooth. **Wing** (Fig. 21A). Membrane homogenously brownish, with distal fourth slightly darker; membrane densely covered with irregularly arranged microtrichia on all cells. Macrotrichia absent on membrane, present on all veins dorsally except on first sector of Rs, base of r-m, M_2_, basal ⅔ of M_4_, and bM; ventral setae only on Sc and R_1_; dorsal setae on R_1_ and Rs in two rows, setae slightly flattened, as slender scales. Sc well sclerotized, complete, reaching C way before base of Rs. C ending at tip of R_5_, before wing apex. First sector of Rs nearly transverse, bare, less than fourth length of r-m. R_1_ relatively long, reaching C on distal fifth of wing; R_4_ absent; R_5_ reaching C before wing apex; r-m strictly longitudinal at origin on Rs, then transverse across wing fold; a well sclerotized, setose wing fold from r-m towards base. M_1+2_ almost one fifth of medial fork; M_1_ and M_2_ running more or less parallel along most of their length; M_4_ long, disconnected from CuP at base; M_4_ and CuA reaching wing margin, CuA also detached from other veins at base. A conspicuous fold along the anal lobe. **Abdomen**. Tergites and sternites 1–7 dark brown, except for tergite 2, sternite 2, and tergite 5 light brown, tergites 3–4 whitish-yellow on ventral half and brown on distal half, sternites 3–4 yellowish-brown, sternite 5 yellowish-brown, lighter medially. Abdominal sclerites with flattened setae, especially along posterior margin. **Terminalia** (Figs. 21B–C). Terminalia dark brown with yellowish cercus. Gonocoxite very long, distal end almost reaching level of tip of gonostylus; gonocoxites close together medially at anterior end, no evidence of sternite 9, distal end with an apical spine; a small, rounded lobe on internal face midway to apex, bearing 5–6 small setae and a long seta. Gonostylus very long, displaced to a dorsal position, partially articulated to tergite 9, internal face covered with setation, a long tooth midway to apex, distally with a short subapical tooth and a pair of short digitiform projections apically. Gonocoxal bridge present at anterior end of terminalia, apodemes more or less close to each other. Aedeagus large, projected distally, strongly sclerotized at tip. Tergite 9 short, trapezoid, with four elongate setae medially, typically a pair of sutures extending anteriorly from posterior margin. Sternite 10 small, weakly sclerotized, with a pair of short lobes on posterior margin. Cerci small, elongate, with microtrichia all over surface and some fine setae along posterior border.

**Female**. As male, except as following. **Wing**. Membrane homogenously brownish, with distal fourth slightly darker; membrane densely covered with irregularly arranged microtrichia on all cells. Macrotrichia absent on membrane, present on all veins dorsally except on first sector of Rs, base of r-m, M_2_, basal ⅔ of M_4_, and bM; ventral setae only on Sc and R_1_; dorsal setae on R_1_ and Rs in two rows, setae slightly flattened, as slender scales. Sc well sclerotized, complete, reaching C way before base of Rs. C ending at tip of R_5_, before wing apex. First sector of Rs nearly transverse, bare, less than ¼length of r-m. R_1_ relatively long, reaching C on distal fifth of wing; R_4_ absent; R_5_ reaching C before wing apex; r-m strictly longitudinal at origin on Rs, then transverse crossing wing fold; a well sclerotized, setose wing fold from r-m towards base. M_1+2_ almost one fifth of medial fork; M_1_ and M_2_ running more or less parallel along most of their length; M_4_ long, disconnected from CuP at base; M_4_ and CuA reaching wing margin, CuA also detached from other veins at base. A conspicuous fold along anal lobe. **Abdomen**. Entirely brown except for anterolateral corners of tergite 4 and across sternite 4. **Terminalia** (Figs. 21D– E). Sternite 8 well-developed, trapezoid, medial posterior projection rounded. Sternite 9 slender, with a short anterior extension, membranous area between posterior arms slightly more sclerotized around genital opening. Tergite 8 wide, entirely bare, sclerotized medially along posterior margin. Tergite 8 short, largely membranous, with a sclerotized slender band along anterior margin and a row of setae and setulae. Tergite 9+10 slender, with a large setose area. Cerci 1-segmented, elongated, fused together along anterior half.

#### Material examined

**Holotype**: male: ZRCBDP0048240, Sungei Buloh (SB1), mangrove, 10.jul.13, MIP leg. (imaged, slide-mounted). **Paratypes** (6 females): ZRCBDP0048284, Pulau Semakau (SMO2), old mangrove, 17.oct.13, MIP leg. (slide-mounted); ZRCBDP0048772, National University of Singapore (PGP), urban forest, 20.may.15, MIP leg.; ZRCBDP0278288, Singapore, 17.may.18, MIP leg.; ZRCBDP0284273, Singapore, 02.may.18, MIP leg.; ZRCBDP0278334, Singapore, 03.may.18, MIP leg.; ZRCBDP0278211, Pulau Ubin (PU18), mangrove, 31.may.18, MIP leg. **Sequence failures specimens**: ZRCBDP0069322, Pulau Ubin (PU01), mangrove, 23.jun. 16, MIP leg.

**Etymology**. The specific epithet of this species refers to the name Tumasik used in the Old Javanese [=Kawi] epic poem Nagarakretagama, written in the 14th century, to refer to the settlement (also referred to elsewhere as Temasek, meaning “sea town”) in the island that is now Singapore. The noun is used in apposition.

**Remarks**. In the type-series of *A. tumasik*, the fore femur is whitish, the mid femur is whitish on its basal third, and tergites and sternite 3–5 have cream-yellowish anterolateral marks. In *A. tumasik* Amorim & Oliveira, **sp. n.,** there is variation on the extent of light areas on these sternites in some females. The gonostylus in *A. tumasik* Amorim & Oliveira, **sp. n.** has typically a pair of small projections at the tip and the gonocoxites are more slender; in *A. limbosengi* Amorim & Oliveira, **sp. n.** the gonocoxites are larger and the gonostylus is blunt distally, with the posterior end hardly sclerotized.

*Allactoneura limbosengi* Amorim & Oliveira, sp. n. (Figs. 22A–D, 23A)

https://singapore.biodiversity.online/species/A-Arth-Hexa-Diptera-000789,-002117

urn:lsid:zoobank.org:act:3D6EB176-E9EC-4073-AFC0-FD37EDE8D364

**Diagnosis**. Gonocoxite large, with a long seta and a pair of small spines distally on inner face, Gonostylus with a strong tooth on inner margin midway to apex, blunt distally with a well-sclerotized tip.

***Allactoneura_limbosengi_ZRCBDP0278244_holotype* [8 : G, 11: A, 12: A, 25: T, 67: T, 76: A, 106: T, 146: A, 17 3: C, 184: T, 194: T, 229: A]**

ttagctGctAAtattgctcatacTggagcgtcagttgatttagctattttttcattacatttagcTggtatttcAtctattttaggagcagtaaattttattacTacaattattaatatacgagctcctggaattagatttgatAaaatacctttatttgtatgatcagttCttattacagcTattttattaTtactttctttaccagtattagcaggagctattacAatattattaacagatcgtaatttaaatacttcattttttgatccagctggaggaggagatccgattttataccaacatttattt

#### Description

**Female** (Fig. 22A). Wing length, 3.65; width, 1.15. **Head** (Fig. 22B). Vertex dark brown, occiput dark brown. Scape and pedicel light brownish-yellow; flagellomeres ochre-brownish, darker towards apex. Frons blackish-brown, face dark brown; clypeus yellowish-brown, ventral margin lighter; labella whitish; palpomeres whitish. Labella whitish-yellow. **Thorax**. Scutum dark shinning brown, dorsally compressed, scutellum dark brown. Pleural sclerites brown. Pleural membrane ochre-brown. Anepisternum, katepisternum, mesepimeron and metepisternum bare. Laterotergite bulging, with five bristles along outer margin dorsally. Mediotergite strongly curved in profile. Haltere whitish. **Legs**. All coxae whitish-yellow, with a dark brown band basally and some brownish tinge at tip. Trochanters brown. All femora brown, tibiae and tarsi brown, tarsi densely covered by blackish setulae giving a greyish tinge. Fore leg tarsomere 1 slightly longer than fore tibia; tibiae and tarsi with erect darker short bristles along almost entire length, those on hind tibia more or less aligned dorsally and laterally. Tibial spurs very long, fore tibia spur yellowish-brown, mid and hind tibial spurs whitish, internal spurs slightly longer than inner spur. Mid and hind tibiae and tarsomeres 1-2 with short, spiny black setulae, tarsomeres 3-4 with a pair only with black setulae at distal margin. Tarsal claws with a large basal tooth. **Wing** (Fig. 22D). Membrane homogenously light brownish, distal fourth slightly darker; microtrichia irregularly arranged on all wing cells. Macrotrichia slightly flattened, absent on wing membrane, present dorsally on all veins except on first sector of Rs, base of r-m, M_2_, basal ⅔ of M_4_, and bM, R_1_ and Rs with two rows; ventral setae on Sc and R_1_. Sc complete, reaching C much more basally than origin of Rs. C ending at tip of R_5_, before wing apex. First sector of Rs nearly transverse, bare, less than ¼ of length of r-m. R_1_ reaching C on distal fifth of wing; R_4_ absent; r-m strictly longitudinal at anterior half, transverse on posterior half, crossing wing fold; a well sclerotized, setose wing fold from r-m towards wing base. M_1+2_ almost one fifth of medial fork; M1 and M2 running more or less parallel along most of their length; M4 long, incomplete at base; M4 and CuA reaching wing margin, CuA also detached from other veins at base. A conspicuous fold along anal lobe. Abdomen (Fig. 22C). Entirely brown except for whitish anterolateral corners of tergite 4 and whitish sternite 4. **Terminalia** (Fig. 22C). Sternite 8 and tergite 8 light caramel-brown, rest of terminalia distally whitish.

**Male**. As females, except for the following. **Terminalia** (Fig. 23) Dark brown, cercus lighter. Gonocoxites fused together at anterior end, no evidence of sternite 9, gonocoxite wide at base, extending ventrally way beyond insertion of gonostylus, distal end almost at level of tip of gonostylus, inner face of distal third with a row of one very long seta, some smaller setae and 2–3 small spines. Gonostylus very long, displaced to a more dorsal position basally, also articulated to tergite 9, outer face densely setose, inner face also with setae, a long tooth along inner margin on distal third, more slender at distal end, strongly sclerotized at tip. Paramere elongate, with some fine setae along its length; aedeagus large, elongate, more sclerotized at tip. Tergite 9 short, trapezoid, bare. Sternite 10 elongate, weakly sclerotized. Cerci elongate, densely covered with microtrichia and with some fine setae distally.

#### Material examined

**Holotype**: male, ZRCBDP0278244, Pulau Ubin (PU18), mangrove, date range 2012-2018, MIP leg. (slide-mounted). **Paratypes** (21 males, 12 female). **Males**: ZRCBDP0278169, Pulau Ubin (PU18), mangrove, 10.may. 18 (slide-mounted); ZRCBDP0278173, Pulau Ubin (PU18), mangrove, 10.may. 18 (slide-mounted) (MZUSP); ZRCBDP0278176, Pulau Ubin (PU18), mangrove, 10.may. 18 (slide-mounted); ZRCBDP0278202, Pulau Ubin (PU18), mangrove, 10.may. 18 (slide-mounted); ZRCBDP0278210, Pulau Ubin (PU18), mangrove, 31.may. 18 (slide-mounted); ZRCBDP0278213, Pulau Ubin (PU18), mangrove, 31.may. 18 (slide-mounted); ZRCBDP0278221, Pulau Ubin (PU18), mangrove, 31.may. 18 (slide-mounted); ZRCBDP0278264, Singapore, 07.jun.18, MIP leg. (slide-mounted); ZRCBDP0278278, Singapore, 07.jun.18, MIP leg. (slide-mounted); ZRCBDP0278298, Singapore, 07.may.18, MIP leg.; ZRCBDP0278317, Singapore, 03.may.18, MIP leg. (slide-mounted); ZRCBDP0278323, Singapore, 03.may.18, MIP leg. (slide-mounted); ZRCBDP0284167, Pulau Ubin (PU18), mangrove, 2018, MIP leg.; ZRCBDP0284168, Pulau Ubin (PU18), mangrove, 2018, MIP leg. (slide-mounted); ZRCBDP0284170, Pulau Ubin (PU18), mangrove, 2018, MIP leg.; ZRCBDP0284171, Pulau Ubin (PU18), mangrove, 2018, MIP leg.; ZRCBDP0284172, Pulau Ubin (PU18), mangrove, 2018, MIP leg.; ZRCBDP0284174, Pulau Ubin (PU18), mangrove, 2018, MIP leg. (slide-mounted); ZRCBDP0284175, Pulau Ubin (PU18), mangrove, 2018, MIP leg.; ZRCBDP0284176, Pulau Ubin (PU18), mangrove, 2018, MIP leg. (slide-mounted) (MZUSP); ZRCBDP0284179, Pulau Ubin (PU18), mangrove, 2018, MIP leg. (slide-mounted). **Females**: ZRCBDP0048563, Nee Soon (NS1), swamp forest, 04.apr. 12, MIP leg. (website photo specimen, extracted); ZRCBDP0048564, Nee Soon (NS2), swamp forest, 04.apr. 12, MIP leg.; ZRCBDP0048707, Nee Soon (NS2), 28.jan.15, MIP from r-m towards wing base. M 1+2 almost one fifth of leg.; ZRCBDP0047786, Nee Soon (NS2), swamp forest, 26.feb.14, MIP leg. (website photo specimen, slide-mounted); medial fork; M_1_ and M_2_ running more or less parallel along most of their length; M_4_ long, incomplete at base; M_4_ and CuA reaching wing margin, CuA also detached from other veins at base. A conspicuous fold along anal lobe. **Abdomen** (Fig. 22C). Entirely brown except for whitish anterolateral corners of tergite 4 and whitish ZRCBDP0047846, Nee Soon (NS1), swamp forest, 26.jun.13, MIP leg.; ZRCBDP0047874, Nee Soon (NS1), swamp forest, 19.jun.13, MIP leg.; ZRCBDP0047880, Nee Soon (NS1), swamp forest, 25.set.13, MIP leg.; ZRCBDP0284243, Singapore, date range 2012-2108, MIP leg.; ZRCBDP0048981, Nee Soon (NS2), 17.dec.14, MIP leg.; ZRCBDP0134000, Singapore, date range 2012-2108, MIP leg.; ZRCBDP0134057, Singapore, date range 2012-2108, MIP leg. (slide-mounted); ZRCBDP0154977, Nee Soon (NSM2a), swamp forest, 14.jan.15, MIP leg. (website photo specimen).

**Etymology**. The specific epithet of this species honors Lim Bo Seng (1909–1944), a Chinese resistance fighter based in Singapore and Malaya during World War II. Before the outbreak of War World II, he was a prominent businessman among the Chinese community in Singapore. When the Second Sino-Japanese War broke, he participated in anti-Japanese activities in Malaya and Singapore. He was captured and died while interred. After the war, he is remembered as a war hero in Singapore.

**Remarks**. Most specimens of *Allactoneura limbosengi* Amorim & Oliveira, **sp. n.** have dark fore and mid femora and a small whitish-yellow mark on the tergite 4 and only the sternite 4 with a posterior whitish-yellow band. We have a larger number of specimens of this species and there are males with the fore femur and the mid femur with the basal ⅔ whitish, but with the terminalia strictly similar to that of the holotype. This show intraspecific variation and the color pattern cannot be used alone as a diagnosis to discriminate between the species. We have eight haplotypes for *A. limbosengi*, that lump with *Allactoneura tumasik* Amorim & Oliveira, **sp.n.** at OC 4-5%.

### Eumanota Edwards

*Eumanota* Edwards 1933: 231. Type species: *Eumanota leucura* Edwards 1933: 232, by original designation.

Ten species of *Eumanota* have been described to date—with an additional female not formally described. The known distribution of the genus includes Taiwan, Myanmar, Thailand, Malaysia (Pahang and Borneo), Indonesia (Sumatra and Maluku Utara), and Papua New Guinea (Edwards 1933, Søli 2002, Papp 2004, Hippa et al. 2005), with an additional Neotropical species described from the Colombian highlands (Amorim et al. 2018). The descriptions of all species include careful illustrations of the male terminalia. The shape of the gonostylus leaves no question that the species from Singapore fits into a small group also including *Eumanota racola* Søli, *E. suthepensis* Søli, *E. kambaiti* Hippa et al. and *E. humeralis* Edwards. As discussed below, the shape of the posterior margin of the gonocoxites ventrally and dorsally is characteristic of each species of *Eumanota*, and the specimens from Singapore should be conspecific with the holotype of *E. racola*.

***Eumanota racola*** Søli https://singapore.biodiversity.online/species/A-Arth-Hexa-Diptera-000765 (Figs. 24A–H) *Eumanota racola* Søli 2002: 50, male terminalia (figs. 14– 16). Type locality: Thailand, Phang Nga Province, Koh Ra.

**Diagnosis** (modified from Søli 2002). Posterior, ventromedial corner of gonocoxites less pronounced, gonostylus not produced into a small hook distally, subtriangular, with a blunt, well sclerotized distal end.

***Eumanota_racola_ZRCBDP0048561_hapZRCBDP0047 931_describedearlier_type* [1: A, 8: A, 12: T, 13: T, 24: A, 104: A, 113: C, 115: A, 125: G, 127: A]**

AttatctAataTTattgctcatgAaggagcttctgtagattttgcaattttttcccttcatttagcaggaatttcttctattttaggagcagtaaattttattActacaattCtAaatatacgaGcAccaggcatttcttttgaccaaatacctttattcatttgatcagtattaattactgcaattcttttattactttcattaccagtattagcaggtgctattacaatattacttacagatcgaaatttaaatacatctttttttgatcctataggaggaggagatccaattctttatcaacatctattt

**Redescription**. **Male**. Wing length, 2.60, width, 0.92. **Head** (Fig. 24B). Vertex brown, with scattered setulae. Three ocelli, lateral ocelli separated from eye margin by less than their own diameter, mid ocellus about half size of lateral ocelli. Occiput dark brown dorsally, brown towards ventral margin. Occiput and postgena with several strong bristles projecting along lateral border, posteriorly to eye. Dense interommatidial setulae. Scape and pedicel yellow, rounded, with setae distally, no strong seta dorsally on pedicel; flagellomeres 1–8 brown with basal yellow band, flagellomeres 9–14 brown, flagellomeres longer than wide, with scattered setae. A crown of setae on occiput around eyes. Frons, face and clypeus light brown; frons with some strong setae, face densely covered with setae medially and some dark brown setae along laterals, clypeus very short, with a line of setae and setulae. Palpomeres yellow, with four palpomeres, palpomere 3 widely developed, with a conspicuous apical sensory pit, palpomere 4 inserted subapically on previous palpomere, about as long as palpomere 3, palpomere 5 about 5× length of penultimate; a strong seta on palpomere 2, microtrichia all over surface of palpomeres, elongate setulae on distal face and a group of stronger dorsal setae on palpomere 3, a number of stronger setae dorsally along entire palpomere 4 length, fine setulae along entire dorsal side of palpomere 5. Labella yellow, well-developed. **Thorax**. Scutum compressed, light brown, with yellowish anterior corners and lateral margins. Scutellum darker than scutum. Scutum covered with small setae and some few scattered slightly longer setae. Scutellum with eight scutellar bristles close to posterior margin, plus a large number of small setae covering most of disk. Pleural sclerites yellowish, except for light brown laterotergite, a distal light brown band on mesepimeron, and light brown borders of anepisternum. Pleural membrane yellow. Basisternum largely developed, projected between fore coxa and antepronotum, covered with short setae and some longer setae along posterior margin. Antepronotum largely developed, covered with small setae and a row of longer setae along rounded posterior border. Proepisternum reduced in size, triangular, extending from ventroposterior corner of antepronotum. Anepisternum larger than katepisternum, entirely covered with setulae and some few longer setae close to posterior margin. Mesepimeron wide, reaching ventral margin of pleura, bare. Laterotergite bulging, with over 30 setulae and about 12 longer setae, suture separating from mediotergite incomplete dorsoposteriorly. Mediotergite curved in profile, bare. Haltere whitish, some few setae on pedicel, knob densely covered with setulae. **Legs**. Legs whitish, tips of mid and hind coxae with a brown macula, trochanters brown. Mid and hind coxae twice wider than fore coxa. Fore coxa with small setae on frontal and lateral faces, some stronger setae on distal end; mid coxa covered with small setae on frontal face; hind coxa with a line of fine setae along entire length. Fore tibia shorter than femur; mid and hind tibiae longer than femora. Anteroapical depressed area on inner face of fore tibia wide, lined with setulae. Tibiae and tarsi with aligned trichia, tibia and first two tarsomeres of fore leg with rows of setae ventrally, mid and hind tibiae with a pair of dorsolateral rows of setae and a ventrolateral row of setae. Mid and hind tarsomeres 1 and 2 with rows of ventrolateral setae. Tarsomere 1 almost 1. 5 length of second one. Tibial spurs yellowish, about 2× tibia width at apex, internal spur almost half of length of outer tibial spur. Tarsal claws with a large basal tooth. **Wing** (Fig. 24C). Membrane homogenously light greyish-brown, lighter close to wing base; membrane densely covered with microtrichia on all cells, macrotrichia on cells m1, m2, m4 and cua, anal lobe extensively setose. Sc short, ending in bR. C ending much beyond apex of R_5_, covering about ¾ of extension of M_1_. R_1_ short, reaching C on distal fourth of wing. First sector of Rs nearly transverse, devoid of setae, much less than half length of r-m, less sclerotized than R_1_ and Rs. R_4_ absent. R_5_ gradually curved on distal half, reaching C before wing apex; r-m almost transverse, well sclerotized, setose, more than 2× length of R_1_; bM about twice r-m length, cell br conspicuously wider at level of insertion of M_1+2_. M_1+2_ very short, less than ¼ of length of r-m; M_1_ and M_2_ gradually diverging along most of their length, M_1+2_ weakly sclerotized, M_1_ not sclerotized at base. M_4_ originating at base of wing, not directly in contact with CuA, slightly sinuous on distal third. CuP extending almost to level of origin of M_1+2_, with a row of setae along its distal fifth. All veins with dorsal macrotrichia except first sector of Rs. **Abdomen**. Abdomen greyish-brown, except for yellowish sternites 1–4. **Terminalia** (Figs. 24D–E). Terminalia brown, with yellow cercomere. Gonocoxites fused along anterior fourth of ventral face of terminalia, incision deep, wide on distal half, almost closed on anterior end; no posterior gonocoxite lobes projecting beyond tip of gonostylus, with a short dorsoposterior projection at corners of tergite 9 and a short inner projection at ventral face, to which tip of gonostylus fits. Gonostylus subtriangular, rounded on dorsal margin, ending into a sclerotized acute projection.

Gonocoxal bridge with a pair of long apodemes directed towards mid of terminalia anteriorly. Aedeagal-parameral complex with a pair of wide lateral plates that extend at each side into a digitiform posterior projection reaching level of base of gonostylus, bearing around 20 setae placed very close to each other; aedeagus protruding beyond tip of parameres, with a slender sclerotized axis, opening distally into a membranous area that extends almost to level of tip of gonostylus. Tergite 9 about as long as broad, but wider midway to apex. Cercus fused to each other on anterior half, extending way beyond tip of gonostylus, densely covered with microtrichia and elongate setae; sternite 10 membranous, with a pair of distal lobes, almost as long as cerci, densely covered with microtrichia and elongate setae.

**Female**. As male, except for the following. Wing length, 2.70; width, 0.92. **Terminalia** (Figs. 24F–G). Sternite 8 with a pair of elongate lobes entirely separated from each other, microtrichia and setulae all over ventral face and some darker setae, especially at distal end. Tergite 8 well sclerotized, wide, with a sclerotized band along entire anterior and posterior margins, setose on posterior third. Sternite 9 (vaginal furca) with elongate, curved anterior arm. Tergite 9+10 weakly sclerotized, with microtrichia, some scattered setulae, and a row of short digitiform bases from which longer setae emerge. Cerci long, 1-segmented, with microtrichia and setae.

**Material examined** (6 males, 3 females). **Males**: ZRCBDP0047923, Nee Soon (NS1), swamp forest, 31. jul.13, MIP leg.; ZRCBDP0047931, Nee Soon (NS1), swamp forest, 17.jul.13, MIP leg.; ZRCBDP0048560, Nee Soon (NS1), swamp forest, 04.apr. 12, MIP leg. (slide-mounted); ZRCBDP0048561, Nee Soon (NS2), swamp forest, 04.apr. 12, MIP leg.; ZRCBDP0048562, Nee Soon (NS2), swamp forest, 05.Sep.12, MIP leg.; ZRCBDP0048954, Nee Soon (NS1), 15.apr.15, MIP leg. **Females**: ZRCBDP0047833, Nee Soon (NS1), swamp forest, 24.jul.13, MIP leg.; ZRCBDP0048927, Nee Soon (NS1), 31. dec.14, MIP leg. (slide-mounted) (MZUSP); ZRCBDP0133453, Singapore, date range 2012-2018, Nee Soon Swamp Forest, MIP leg.

**Remarks**. This species is close to *Eumanota humeralis*, *E. suthepensis*, *Eumanota parahumeralis* Papp, and *E. kambaiti*, which have a triangular gonostylus with a distal, sclerotized short projection. There are differences between these three species in the shape of the posterior border of the gonocoxites, the shape of the tergite 9 and the parameres, and in details of the gonostylus itself. Our specimens fit in all details with *E. racola* for which the type locality is Koh Ra, in the Phang Nga Province, an island with lowland tropical forest along the Malaysian peninsula. There are two haplotypes among our specimens (Fig. 24H) and no delimitation conflicts.

### Manota

*Manota* Williston, 1896: 260. Type-species, *M. defecta* Williston (mon.).

**Diagnosis**. Head with a row of strong bristles around posterior margin of eye, directed posteriorly; three ocelli. Sc short, incomplete; R_4_ missing; r-m and bM perfectly aligned, almost longitudinal; M_1+2_ missing, M_1_and M_2_ present as detached veins reaching wing margin.

*Manota* is presently one of the largest genera of Mycetophilidae. Of the almost 320 described species, 68 are Afrotropical (see Kurina & Hippa 2014, Hippa et al. 2019), 17 Palaearctic (review in Hippa & Saigusa 2016), one Nearctic (Jaschhof et al. 2011), 39 Australasian (Kurina & Hippa 2015; Kurina et al. 2019), 109 Oriental (Hippa & Kurina 2018), and 96 Neotropical (Kurina & Hippa 2021). The known Oriental diversity of the genus is concentrated in Thailand (e.g., Hippa 2009, 2011) and in the Peninsular Malaysia (Hippa 2006), with some species also known from Nepal and China (Hippa & Saigusa 2016), Sulawesi (Hippa & Kurina 2018) etc.

The species of the genus *Manota* are ochre-yellowish in color and there are only few features with the exception of terminalia that can be used to separate groups of species. A key for the Oriental species of *Manota* was provided by Hippa (2011) and subsequent papers indicate where they fit in the key. The complex male terminalia features are mostly very well described and illustrated in published papers and allow good comparisons.

Of the 14 species of *Manota* we found in the Singapore samples, seven are known only from females. As stated above, by protocol in this paper we describe but do not formally name the species of *Manota* for which we cannot present a morphological differential diagnosis—which is the case of these seven species with only females available in our sample. We carefully compared the male terminalia of the species from Singapore with those in published papers on Oriental *Manota*. None of the described species fit the species in our samples that have males available. This only shows the high level of species turnover in the genus between rather close localities. A revision of the genus is needed to better understand the diversity of the genus in Singapore and SE Asia. It is worth noting that, despite the number of species and specimens of *Manota*, there is a delimitation conflict in our haplotype networks for only one species, *Manota chiamassie* Amorim & Oliveira, **sp. n.** (Fig. 25A).

*Manota banzu* Amorim & Oliveira, sp. n. (Figs. 25B–C, 26A–D)

https://singapore.biodiversity.online/species/A-Arth-Hexa-Diptera-000756

urn:lsid:zoobank.org:act:88A6E1F3-E3A0-4BBE-BF35-351DF8EBA160

**Diagnosis**. Thorax dirty ochre-yellowish. Laterotergite bare. Anal lobe devoid of setae. Abdomen with segments 1–3 mostly greyish-yellow, segments 4–6 with tergites caramel-brown. Gonocoxites directed lateroposteriorly, posterior margin ventrally not extending beyond base of gonostylus; parastylar lobe short, two long setae directed ventrally; short juxtagonostylar lobe with two long, strong, curved setae; gonostylus relatively small, slightly elongate, flat, with long setae on outer face, distally with seven long, slightly curved setae. Tergite 9 with wide posterior incision separating a pair of lateral lobes, internal posterior margin dorsally projected much beyond base of gonostylus, six strong setae concentrated on distal short beak.

***Manota_banzu_ZRCBDP0047877_hapZRCBDP004787 7_SMH_holotype* [85: T, 133: T, 138: G, 144: C, 181: G, 260: A]**

tttagctgcttcaattgctcattctggagcttctgttgatttatcaattttttctcttcatcttgcaggaatttcttcaattctTggagcagtaaattttattacaacaattattaatatacgaactccaaaTataaGttttaCtcaaataccactatttgtatgatcagttttaattacGgcaattttacttttattatcattacctgtattagctggagctattactatacttttaactgatcgaaacttaaatacaAcattttttgatccagctggtgggggtgatcctattttataccaacatttattt

#### Description

**Male** (Fig. 25B). Wing length, 1. 89; width, 0.71. **Head** (Fig. 25C). Brownish-yellow, brown along ocelli line, occiput darker dorsally to condyle medially. Antenna light brown, scape and pedicel paler. Face light brown, clypeus, palpus, and labella whitish-yellow. Mid ocellus present, slightly smaller than lateral ocelli, at posterior end of frontal furrow, lateral ocelli separated from eyes by a distance larger than ocellus width. About 10–12 strong postocular setae. Eyes densely covered by inter-ommatidial setulae. Scape only slightly longer than pedicel, flagellomeres 1 not longer than following flagellomeres, flagellomeres 1–5 about as long as wide, flagellomeres 6–13 slightly longer than wide, flagellomere 14 about twice as long as wide. Scape with fine setae on external face and around distal end, four ventral setae stronger; internal face with a concentrated group of dorsal setae distally. Pedicel with fine setae on external face and around distal end, four ventral stronger setae. Frons with a pair of sutures laterally extending from internal dorsal corner of eye straight backwards ending behind level of ocellar line; frons projected medially as an inverted triangle, reaching level of insertion of antennae, frontal furrow reaching anterior end of triangular extension; frons almost entirely devoid of setae, except for 10–11 setae on a slender ventral extension of frons that connects with face and small setae dorsolaterally, externally to lateral ocelli. Face wide, clypeus rectangular, elongate, slightly bulging, entirely covered with elongate setae; clypeus short, trapezoid, covered with setae, some stronger setae on ventral half. Maxillary palpomere 1 pretty much reduced; palpomere 2 short, with microtrichia and small setae; palpomere 3 well-developed, sensorial pit elongate, with opening laterally on internal face, apicomedial digitiform projection extending well beyond base of palpomere 4, setae on dorsal and external faces, palpomere 4 with small subdistal parasegment, 1. 6× length of palpomere 3; palpomere 5 weakly sclerotized, elongate, almost twice longer than palpomere 4. Labella short, weakly sclerotized. **Thorax**. Mostly cream-yellow, medial part of scutum and scutellum somewhat darker, anepisternum dark ochre-yellowish, dorsoposterior corner of katepisternum and ventral half of mesepimeron dark ochre-brown, metepisternum greyish, cream-yellowish on anteroventral corner, laterotergite and mediotergite brownish-yellow. Scutum densely covered with scattered small setae, 8–9 larger setae restricted to more sclerotized area along margin above wing and a row of long prescutellars. Scutellum well-developed, with scattered small setae and two pairs of stronger marginal setae, inner pair stronger. Cervical sclerite small. Basisternum dorsoposterior arms elongate, bare of setae. Antepronotum largely developed, lateral lobes connected medially by a slender, bare stripe, posterior end covered with short setae; proepisternum well-developed, extending dorsally to level of spiracular sclerite, covered with small setae and a row of about 10 longer setae along ventral margin. Anepisternum densely covered with small setae, no longer setae along posterior margin; katepisternum bare except for a medial line of small setae along anteroventral margin; anterior basalare with no setae; katepisternum well-developed; mesepimeron reaching ventral margin of thorax, bare, suture separating from katepisternum absent; laterotergite bulging, bare; metepisternum with 17 fine small setae distributed along sclerite length. Mediotergite strongly curved medially, bare. Haltere pedicel yellow, knob brown, small setae on pedicel and on knob. **Legs**. Coxae whitish, femora whitish-yellow, tibiae and tarsi light yellowish. Fore coxa largely developed, covered with small setae on entire face, some stronger setae along anterior end of ventral face and at distal margin posteriorly. Mid coxa covered with small setae anteriorly, less densely on external face, and some strong setae along distal margin of anterior face; hind coxa relatively smaller, a band of small setae medially along entire external face and some few stronger setae on distal margin anteriorly. Femora laterally compressed, mid and hind femora larger, densely covered with small setae, a sequence of longer setae on ventral edge distally. Tibiae and tarsi with trichia more or less aligned. Fore tibia with one subapical small bristle and a row of setae lateroventrally on external face, besides distal setae. Fore tibia with a wide anteroapical depressed area lined with setulae; mid tibia with an irregular row of stronger setae along dorsal edge, some additional lateral and lateroventral setae; hind tibia with an irregular row of stronger setae along dorsal and ventral edges, some additional lateral setae on external face. Fore tibia slightly shorter than femur, first fore tarsomere about as long as tibia. Mid and hind tibial organs absent. Tarsal claws with a longer, more distal tooth coming out on inner face and a more basal, blunt sclerotized tooth. **Wing** (Fig. 26A). Membrane slightly fumose. Sc present, incomplete, ending free. R_1_ short, meeting C before mid of wing. First sector of Rs weakly sclerotized. R_5_ short, reaching C at level of tip of M_4_; r-m long, slightly longer than R_1_. Tip of medial and cubital veins hardly sclerotized. M_1_sclerotized only distally, a line of dorsal setae present on non-sclerotized part of vein more basally. M_2_ originating beyond level of tip of R_1_. M_4_ continuous with first sector of CuA, second sector of CuA disconnected from first sector by a short unsclerotized part. Cubital pseudovein present, CuP absent. Sclerotized anal fold present, long, curved, not reaching posterior wing margin. Dorsal macrotrichia on Sc, bR, R_1_, second sector of Rs, r-m, M_1_, M_2_, M_4_, CuA and anal fold; ventral setae on bR, R_1_, second sector of Rs, and r-m. **Abdomen**. Tergites 1–3 brownish-yellow, darker medially, tergites 4–6 light brown, tergite 7 yellowish; sternites 1–4 whitish-yellow, sternites 5–6 light brown. Tergite and sternite 8 trapezoid, wide basally, rounded posteriorly. Abdominal setosity pale brown. **Terminalia** (Figs. 26B–D). Cream-yellowish. Sternite 9 wide anteriorly, not fused to gonocoxites, posterior margin broadly convex, lateral ends anteriorly with a small apodeme, no medioventral process, mostly covered with fine setae, 4–5 long setae on each side along posterior margin. Medial margins of gonocoxites widely separated, posterior margin short ventrally, gonocoxites directed laterally, posterior margin ventrally not extending beyond base of gonostylus, parastylar lobe short, with two long setae directed ventrally; a short juxtagonostylar lobe with two long, strong, curved setae. Gonostylus relatively small, subquadrate, slightly elongate, flat, with long setae on outer face, distally with seven long, slightly curved setae. Aedeagus subtriangular, elongated distally, apex curved ventrally. Tergite 9 with wide posterior incision separating a pair of lateral lobes, internal posterior margin dorsally projected much beyond base of gonostylus, six strong setae concentrated on distal short beak and a line of smaller setae along margin inwards, long setae more laterally on ventral face, laterally and on dorsal face. Cerci medially separate, elongate, reaching level of tip of gonocoxites dorsally.

**Female**. As male, except for the following. **Wing l**ength, 1. 71; width, 0.64. **Terminalia**. Sternite 8 wide at base, trapezoid, distal end with a pair of short lobes bearing a shallow medial incision, setation restricted to lobes distally from base of incision. Sternite 9 present as a pair of long sclerotized bands connected distally and diverging towards anterior end, extending anteriorly to distal end of segment 7, distal medial end of sternite 9 reaching level of tip of lobes of sternite 8. Tergite 8 wide, not overlapping laterally with lateral border of sternite 8, covered with microtrichia and fine setae. Tergite 9 covered with microtrichia and scattered fine setae. Tergite 10 reduced to an inconspicuous slender band and two pairs of short digitiform projections each bearing a long seta. Cercomere 1 slender, almost 10× longer than wide, over 3× longer than cercomeres 2, both covered with microtrichia and fine setae.

#### Material examined

**Holotype**: male, ZRCBDP0047877, Nee Soon (NS1), swamp forest, 19.jun.13, MIP leg. (slide-mounted). **Paratypes** (7 males, 4 females). **Males**: ZRCBDP0048517, Nee Soon (NS1), swamp forest, 09.may. 12, MIP leg. (website photo specimen); ZRCBDP0048675, Nee Soon (NS2), swamp forest, 02. may. 12, MIP leg.; ZRCBDP0048699, Nee Soon (NS1), 13.may.15, MIP leg.; ZRCBDP0048708, Nee Soon (NS2), 28.jan.15, MIP leg.; ZRCBDP0048822, Nee Soon (NS1), 14.jan.15, MIP leg.; ZRCBDP0048973, Nee Soon (NS1), 13.may.15, MIP leg.; ZRCBDP0049185, Nee Soon (NS2), 13.may.15, MIP leg.

**Females**: ZRCBDP0048677, Nee Soon (NS2), swamp forest, 13.jun. 12, MIP leg. (slide-mounted);ZRCBDP0048997, Nee Soon (NS2), 17.dec.14, MIP leg.; ZRCBDP0072693, Bukit Timah, maturing secondary forest (BT06), 13.oct. 16, MIP leg.; ZRCBDP0155093, Singapore, date range 2012-2018, MIP leg. **Additional sequenced specimens**: male, ZRCBDP0078993; female, ZRCBDP0137281.

**Etymology**. The species epithet refers to the locality of Ban Zu [=班卒, from the Malay word *pancur*, meaning “spring of water”] as recorded by the Chinese traveler Wang Dayuan—thought to be, in present days, Fort Canning Hill—an important 14th century settlement in the island of Singapore. The noun is used in apposition.

**Remarks**. Specimens come the swamp forest and from the Bukit Timah forest. To some extent, *Manota banzu* Amorim & Oliveira, **sp. n.** is similar to *M. stricta* Hippa & Ševčík, from Brunei (Hippa & Ševčík 2010) and to *M. aconcinna* Hippa and *M. inflata* Hippa and *M. submirifica* Hippa (Hippa 2008), from Thailand, to *M. biunculata* Hippa, from Papua-New Guinea, and to *M. gemela* Hippa, from Maluku Utara (Hippa 2007). We have two haplotypes for this species and no delimitation conflicts.

*Manota tantocksengi* Amorim & Oliveira, sp. n. (Figs. 27A–E)

https://singapore.biodiversity.online/species/A-Arth-Hexa-Diptera-000758,-002093

urn:lsid:zoobank.org:act:D8E73AB1-4634-4AED-BA5F-39F101AC3B73

**Diagnosis**. Scutum ochre-yellow on anterior third, brownish yellow on posterior third, darker towards posterior end. Pleural sclerites brownish cream-yellow. Laterotergite setose. Anal lobe devoid of setae. Abdomen with tergites 1–6 brownish. Gonocoxites large, ventromedial margin lobes setose, approximate to each other, setae on posterior margin longer; parastylar lobe small, devoid of setae; juxtagonostylar lobe digitiform, extending distally, with one juxtagonostylar long seta distally; an apicolateral digitiform lobe with a distal megaseta; gonostylus ovoid, elongate, evenly covered with fine setae on ventral face, some stronger setae on distal margin and a group of concentrated small setae on dorsal face. Tergite 9 with wide posterior incision separating a pair of lateral lobes projecting beyond level of base of gonostylus, with a projection on inner margin bearing distally a group of concentrated strong setae and more anteriorly at inner margin a group of longer setae, dorsal face of gonocoxite with fine setae.

***Manota_tantocksengi_ZRCBDP0072684_hapZRCBDP0 047933_SMH_holotype* [82: C, 94: C, 97: C, 122: A, 123 : A, 144: G, 197: A]**

tttagctgcatcaattgctcactctggggcttctgtagatttatcaattttttctttacatttagcaggtatttcttcaatCctaggagcaatCaaCttcattactacaattattaacataAAaacccctgaaataaattttaGacaaatacccttatttgtctgatcagtattaattacagctattcttcttctaAtctcattacctgttttagcaggagcaattacaatattattaactgatcgtaatttaaatacatcattttttgatccagccggaggaggagatccaattctataccaacatttattt

#### Description

**Male** (Fig. 27A). Wing length, 1. 51; width, 0.61. **Head**. Brownish-yellow, darker dorsally. Antenna light brown, scape and pedicel light brownish-yellow. Face, clypeus, palpus and labella dirty-yellowish. Occiput with line of 9–10 strong postocular setae. **Thorax**. Basically cream-yellow with brownish tinge, as well as antepronotum; proepisternum, anepisternum and mesepimeron dark ochre-yellowish, katepisternum and metepisternum lighter; laterotergite brown, mediotergite light brown. Anepisternum covered with fine setae except for anteroventral corner; anterior basalare setose; katepisternum with some fine setae close to posterior border; laterotergite with 23 setae; metepisternum bare. Haltere pedicel yellow, knob brown. **Legs**. Coxae whitish, fore femur whitish-yellow, mid and hind femora light brownish-yellow; tibiae and tarsi light greyish-brown. Mid-and hind tibial organs absent. **Wing** (Fig. 27B). Membrane light greyish-brown; R_1_ meeting C before mid of wing; sclerotized part of M_2_ originating slightly beyond level of tip of R_1_; CuA not connected basally to M_4_. **Abdomen**. Tergites 1–2 brownish-yellow, tergites 3–7 light brown; sternites 1–3 light brown, sternite 4–6 light yellowish-brown. **Terminalia** (Figs. 27C–D). Cream-yellowish. Sternite 9 trapezoid with deep anterior medial incision, on anterior third of gonocoxite, fused to gonocoxites at lateral ends, with two groups of elongate setae separate from each other, no incision at posterior margin medially. Gonocoxites large, lobes of ventromedial margin setose, approximate to each other, setae on posterior margin longer; parastylar lobe small, devoid of setae; juxtagonostylar lobe digitiform, extending distally, with one juxtagonostylar long distal seta; an apicolateral digitiform lobe with a distal megaseta. Gonostylus ovoid, elongate, evenly covered with fine setae on ventral face, some stronger setae on distal margin and a group of concentrated small setae on dorsal face. Gonocoxal bridge well-marked, no apodeme directed anteriorly. Tegmen with a pair of distal digitiform extensions separated by a deep posterior incision, extending anteriorly into a pair of lateral apodemes. Sternite 10 well-developed, wider and projected more distally than tegmen, distal margin rounded, setose, medially with stronger setae. Tergite 9 with wide posterior incision separating a pair of lateral lobes, projecting beyond level of base of gonostylus, with a projection on inner margin bearing distally a group of concentrated strong setae and more anteriorly at inner margin a group of longer setae, dorsal face of gonocoxite with fine setae. Cerci elongate, entirely separate, with fine setae and microtrichia, placed behind tergite 9.

**Females**. As males, except for the following. Wing length, 1. 73; width, 0.66. **Terminalia** (Fig. 27E). Sternite 8 trapezoid, wide at base, distal end with a pair of short lobes bearing a shallow medial incision, setation restricted to lobes distally from base of incision. Sternite 9 present as a pair of long sclerotized bands connected distally and diverging towards anterior end, extending anteriorly to distal end of segment 7, medial end of sternite 9 posteriorly reaching level of tip of sternite 8 lobes, with setulae along distal margin. Tergite 8 wide, not overlapping laterally with lateral border of sternite 8, covered with microtrichia and fine setae. Tergite 9 covered with microtrichia and scattered fine setae. Tergite 10 reduced to an inconspicuous slender band and two pairs of short digitiform projections each bearing a long seta. Cercomere 1 slender, 3.5× longer than wide, covered with microtrichia and fine setae [both cercomeres 2 broken].

#### Material examined

**Holotype**: male, ZRCBDP0072684, Bukit Timah, primary forest (BT05), 22.dec. 16, MIP leg. (slide-mounted). **Paratypes** (3 males, 3 females). **Males**: ZRCBDP0048525, Nee Soon (NS2), swamp forest, 23.may. 12, MIP leg. (website photo specimen, extracted, abdomen separated from thorax); ZRCBDP0072735, Bukit Timah, primary forest (BT05), 14.dec. 16, MIP leg.; ZRCBDP0074028, Bukit Timah, primary forest (BT02), 22.dec. 16, MIP leg. (slide-mounted). **Females**: ZRCBDP0047933, Nee Soon (NS1), swamp forest, 17.apr.13, MIP leg.; ZRCBDP0072664, Bukit Timah, primary forest (BT05), 28.dec. 16, MIP leg.; ZRCBDP0072744, Bukit Timah, primary forest (BT05), 14.dec. 16, MIP leg. (slide-mounted). **Additional sequenced specimens**: male, ZRCBDP0072720, Singapore, date range 2012-2108, MIP leg.; male, ZRCBDP0137247, Bukit Timah (BT05), 19.jan.17.

**Etymology**. The species name honors Tan Tock Seng (1798–1850), merchant and philanthropist, acting Kapitan China of Singapore (government-appointed head of the Chinese community), a successful businessman and the city’s first Asian Justice of Peace. He offered $5,000 for the construction of Singapore’s first privately funded hospital. Many of these immigrants were poor and destitute, and malnutrition was common; it was estimated that about 100 immigrants died each year from starvation. The British government set up a pauper’s hospital in the 1820s, closed in the 1830s due to insufficient funds, suggesting that better-off members of each community take care of their own poor.

**Remarks**. This species was collected from the mangrove to the swamp forest and the primary tropical forest. There are three haplotypes for this species, that group together using any of the delimitation approaches.

*Manota bukittimah* Amorim & Oliveira, sp. n. (Figs. 28A–G)

https://singapore.biodiversity.online/species/A-Arth-Hexa-Diptera-000757

urn:lsid:zoobank.org:act:046F3268-7C33-405A-892B-F6A215943E8B

**Diagnosis**. Thorax ochre-yellowish. Laterotergite bare. Membrane with macrotrichia on cell cua and on anal lobe membrane. Abdomen basically ochre-yellowish, slightly darker medially. Gonocoxites large, ventromedial lobes setose, setae on posterior margin longer; parastylar lobe large, with microtrichia and a pair of long setae directed inwards; juxtagonostylar lobe slender, digitiform, with a pair of long, hook-like juxtagonostylar setae distally directed inwards; no apicolateral lobe; gonostylus with a short neck basally, subquadrate distally, flat, inner face with elongate setae, distal margin with long, curved setae, outer face with microtrichia medially and some setae close to margin; tergite 9 with wide posterior incision separating a pair of lateral lobes, posterior margin extending straight inwards and then curved anteriorly, an internal short lobe on dorsoposterior corner with a group of concentrated setae, inner margin with a long row of setae, posterior margin with five stronger, dorsal face with long setae.

***Manota_bukittimah_ZRCBDP0047832_hapZRCBDP00 47832_SMH_holotype* [1: A, 22: C, 28: T, 143: A, 145: A, 199: G, 211: A, 286: T]**

AttagcagcttcaattgctcaCtctggTgcctcagttgatttatctattttttctcttcatttagcaggtatttcttctattttaggagcagtaaattttattactacaattattaatatacgagcccctaatataaattttAcAcaaatacctttatttgtatgatcagtattaattactgctattttacttttattGtctttacctgtAttagcaggagctattacaatattattaactgatcgaaatttaaatacttcattttttgacccagcaggtggaggTgacccaattctatatcaacatcttttt

#### Description

**Male**. Wing length, 1. 81; width, 0.74. **Head**. Light brownish-yellow, face and clypeus dirty-yellow, vertex with brown mark along line of ocelli, occiput darker posteriorly to eye margin and dorsally to occiput condyle medially. Antennal scape and pedicel light brownish-yellow. Maxillary palpus and labella whitish. Flagellum light brown; flagellomeres 1–8 about as long as wide, flagellomeres 9–13 slightly longer than wide. Maxillary palpomere 4 with small subdistal parasegment, 1. 3× length of palpomere 3; palpomere 5 slightly over twice longer than palpomere 4. About 8 strong postocular setae on each side. **Thorax**. Scutum mostly ochre-yellowish, darker medio-posteriorly, scutellum light brown, antepronotum, proepisternum, anepisternum, mesepimeron, laterotergite and mediotergite ochre-yellowish, katepisternum whitish-brown, metepisternum whitish-brown on anterior fourth, brownish on posterior ¾. Anepisternum with over 50 small scattered setae, except ventromedially; anterior basalare bare; laterotergite and metepisternum bare. **Legs**. Coxae whitish, femora whitish-yellow, tibiae and tarsi light yellowish. Haltere stem light brown, knob dark brown. **Wing** (Fig. 180B).

Membrane greyish infuscate. C ending at about ¾ of distance to M_1_; R_1_ meeting C before mid of wing; sclerotized part of M_2_ originating slightly beyond level of tip of R_1_. Membrane with macrotrichia on cell cua and on anal lobe membrane. **Abdomen**. Tergites 1–6 yellowish-brown, darker medially, tergite 7 light yellowish-brown; sternites 1–3 light brown, sternite 4–6 light yellowish-brown, sternite 7 yellowish. **Terminalia** (Figs. 28C–D). Cream-yellowish. Sternite 9 trapezoid, with rounded posterior end and a deep anteromedial incision, almost half of gonocoxite length, with a slender fusion to gonocoxites at lateral ends, medial depression separating two elongate, setose pair of lobes. Gonocoxites large, margin of ventromedial lobes setose, approximate to each other, setae on posterior margin longer; parastylar lobe large, with microtrichia and a pair of long setae directed inwards; juxtagonostylar lobe slender, digitiform, with a pair of long, hook-like juxtagonostylar setae distally directed inwards; no apicolateral lobe. Gonostylus with a short neck, subquadrate distally, flat, inner face with elongate setae, distal margin with long, curved setae, outer face with microtrichia medially and some setae close to margin. Gonocoxal bridge with no anteriorly directed apodeme. Tegmen triangular, with an elongate distal, curved projection ending pointed. Sternite 10 well-developed, wide, projected to level of tip of tegmen, distal margin rounded, with two pairs of setae medially and a stronger additional external pair. Tergite 9 with a wide posterior incision separating a pair of lateral lobes, posterior margin extending straight inwards and then curved anteriorly, an internal short lobe on dorsoposterior corner with a group of concentrated setae, inner margin with a long row of setae, posterior margin with five stronger setae, dorsal face with long setae. Cerci elongate, flat, entirely separate from each other, with microtrichia and fine setae, long setae distally.

**Female**. As male, except as follow. Wing length, 1.86; width, 0.69. Wing membrane with few dorsal macrotrichia on anal lobe and cell cua. **Terminalia** (Figs. 28E–F). Sternite 8 trapezoid, wide at base, distal end with a pair of lobes and a medial incision posteriorly, fine setae entirely covering sclerite, especially on lobes, two longer setae at tip of each lobe. Sternite 9 present as a pair of long sclerotized bands connected distally and diverging towards anterior end, extending anteriorly to distal end of segment 7, distal medial end of sternite 9 almost reaching level of tip of lobes of sternite 8, gonopore connected to a pair of gonoducts. Tergite 8 wide, laterally barely overlapped to lateral border of sternite 8, covered with microtrichia and fine setae. Tergite 9 about as long as tergite 8 and more slender, covered with microtrichia and scattered fine setae. Sternite 10 with one pair of outer long setae and one pair of inner short setae on distal margin. Tergite 10 reduced to an inconspicuous slender band and with two pairs of setae at tip of long digitiform projection, one pair at lateral corners and one pair sub-medially. Cercomere 1 slender, about 4× longer than wide, cercomere 2 ovoid, covered with microtrichia and fine setae.

#### Material examined

**Holotype**: male, ZRCBDP0047832, Nee Soon (NS1), swamp forest, 24.jul.13, MIP leg. (slide-mounted). **Paratypes** (2 males, 4 females). **Males**: ZRCBDP0048530, Nee Soon (NS2), swamp forest, 04.apr. 12, MIP leg. (website photo specimen); ZRCBDP0048522, Nee Soon (NS1), swamp forest, 06.jun. 12, MIP leg. **Females**: ZRCBDP0048527, Nee Soon (NS2), swamp forest, 08.aug. 12, MIP leg. (slide-mounted); ZRCBDP0048528, Nee Soon (NS2), swamp forest, 04.jul. 12, MIP leg.; ZRCBDP0048673, Nee Soon (NS1), swamp forest, 02.may. 12, MIP leg.; ZRCBDP0048678, Nee Soon (NS2), swamp forest, 04.apr. 12, MIP leg. **Additional sequenced specimens**: female, ZRCBDP0137035, Bukit Timah (BT08), 05.jul. 17, MIP leg. (website photo specimen).

**Etymology**. The species epithet refers to the Bukit Timah Nature Reserve, a 1.64km² reserve of primary tropical forest on the slopes of Bukit [=hill in Malay] Timah, near the geographic center of Singapore. Together with the neighboring Central Catchment Nature Reserve, it houses over 840 species of flowering plants. Its status as a biological reserve goes back to 1882, when Nathaniel Cantley, then Superintendent of the Singapore Botanic Gardens, was commissioned by the Government of the Straits Settlements to prepare a report on the forests of the settlements and Bukit Timah was one of the first forest reserves established, in 1883. By 1937, the forest reserves in Singapore were depleted under economic pressures for development. Three areas, however, including the Bukit Timah Reserve, were retained for the protection of flora and fauna, under the management of the Singapore Botanic Gardens. In 1951, a Nature Reserves Board was established for the administration of the reserves, which total some 28km² in area. The noun is used in apposition.

**Remarks**. This species was sampled in the Bukit Timah tropical forest, as well as in the swamp forest and in mangroves. We have two haplotypes for this species and no delimitation conflicts.

*Manota chiamassie* Amorim & Oliveira, sp. n. (Figs. 29A–D)

https://singapore.biodiversity.online/species/A-Arth-Hexa-Diptera-000781

urn:lsid:zoobank.org:act:26208BB8-358C-4255-8880-B71AC5879BDB

**Diagnosis**. Thorax ochre-yellowish. Laterotergite bare. Macrotrichia on anal lobe membrane. Abdomen light brownish-yellow. Gonocoxites large, medial lobes of ventral margin setose; parastylar lobe with a pair of elongate fine setae directed inwards; juxtagonostylar lobe slender, digitiform, with a pair of long, hook-like juxtagonostylar setae directed inwards distally; no apicolateral lobe; gonostylus with a short neck, subquadrate distally, flat, inner face with elongate setae, distal margin with long, curved setae, outer face with microtrichia medially and some setae close to margin; tergite 9 with wide posterior incision separating a pair of lateral lobes, posterior margin extending straight inwards, dorsoposterior corner truncate, with a line of longer setae. ***Manota_chiamassie_ZRCBDP0048953_hapZRCBDP00 48953_SMH_holotype* [8: G, 55: A, 94: T, 131: A, 145: C, 226: C, 244: C]**

tttagcaGcatctattgctcattctggagcttctgtagatttatctattttttcAttacatttagcaggaatttcttcaattttaggagcagtTaattttattacaacaattattaatatacgagcccctAatataaattttacCcaaatacctttatttgtatgatcagttctaattacagctattcttttattactttctttaccagtattagctggagcaatCactatattattaacagaCcgaaatttaaatacatcattttttgaccctgctggaggaggagacccaattttataccaacacttattt

#### Description

**Male** (Fig. 29A). Wing length, 1.79; width, 0.66. **Head**. Vertex brown, face light yellowish-brown, clypeus whitish-yellow, occiput light brown, a brown band along line of ocelli and along posterior margin of eye; nine strong postocular setae on each side. Antennal scape and pedicel whitish-yellow, darker dorsally, flagellum light brown. Maxillary palpus and labella whitish. Flagellomeres slightly longer than wide, except distal flagellomere, twice longer than wide. **Thorax**. Scutum mostly ochre-yellowish, scutellum light brown. Antepronotum and proepisternum ochre-yellowish, anepisternum, mesepimeron and laterotergite ochre with brownish tinge, katepisternum whitish, mediotergite light brown on dorsal half, ochre on ventral half, metepisternum light brown. Three pairs of long prescutellars. Anepisternum covered with scattered small setae except at ventral fourth; anterior basalare bare; katepisternum with a band of fine setae dorsoposteriorly; laterotergite bare; metepisternum with about 30 fine setae along its length. Haltere stem light brown, knob dark brown. **Legs**. Coxae whitish, femora, tibiae and tarsi whitish-yellow, tarsi with a brownish tinge. **Wing** (Fig. 29B). Membrane with light brown infuscation. C ending at about 4/5 of distance to M_1_; R_1_ meeting C before mid of wing; sclerotized part of M_2_ originating at level of tip of R_1_. Membrane with macrotrichia on anal lobe membrane. **Abdomen**. Tergites 1–6 yellowish-brown, darker medially, tergite 7 light yellowish-brown; sternites 1–3 light brown, sternite 4–6 light yellowish-brown, sternite 7 yellowish. **Terminalia** (Figs. 29C–D). Sternite 9 trapezoid with rounded posterior end and a deep anterior medial incision, almost half of gonocoxite length, with a slender fusion to gonocoxites at lateral ends, medial depression separating two elongate, setose pair of lobes. Gonocoxites large, ventromedial margin lobes setose; parastylar lobe with a pair of elongate fine setae directed inwards; juxtagonostylar lobe slender, digitiform, with a pair of long, hook-like juxtagonostylar setae directed inwards distally; no apicolateral lobe. Gonostylus with a short neck, subquadrate distally, flat, inner face with elongate setae, distal margin with long, curved setae, outer face with microtrichia medially and some setae close to margin. Gonocoxal bridge with oblique apodemes. Tegmen triangular, with an elongate distal, curved projection with pointed end, apodemes anteriorly directed obliquely outwards. Sternite 10 well-developed, wide, projected to level of tip of tegmen, distal margin rounded, with two pairs of setae medially and a stronger additional pair externally. Tergite 9 with wide posterior incision separating a pair of lateral lobes, posterior margin straight extending inwards, dorsoposterior corner truncate, with a line of longer setae. Cerci elongate, flat, entirely separate from each other, with microtrichia and fine setae, long setae distally.

**Female**. As males, except for the following. **Terminalia**. Light yellowish-brown. Sternite 8 wide at base, trapezoid, distal end with a pair of lobes and a medial incision posteriorly, fine setae entirely covering sclerite, especially on lobes, two longer setae at tip of each lobe. Sternite 9 present as a pair of long sclerotized bands connected distally and diverging towards anterior end, extending anteriorly to distal end of segment 7, anterior end very thin, distal end of sternite 9 medially almost reaching level of tip of lobes of sternite 8, gonopore connected to one single gonoduct. Tergite 8 apparently fused to tergite 9, slightly longer lateroposteriorly than medially, covered with microtrichia and scattered fine setae. Sternite 10 with one pair of outer long setae and one pair of inner short setae on distal margin. Tergite 10 reduced to an inconspicuous slender band with two pairs of setae at tip of long digitiform projections, one pair at lateral corners and one pair sub-medially [tip of cercomeres 1 broken, cercomeres 2 missing].

#### Material examined

**Holotype**: male, ZRCBDP0048953, Nee Soon (NS1), 15.apr. 2015, MIP leg. (website photo specimen, slide-mounted). **Paratype** (17 males, 4 females). **Males**: ZRCBDP0066795, Bukit Timah, primary forest (BT05), 22.Sep.2016, MIP leg.; ZRCBDP0066801, Bukit Timah, primary forest (BT05), 14.dec. 2016, MIP leg.; ZRCBDP0066822, Bukit Timah, primary forest (BT09), 05.oct. 2016, MIP leg.; ZRCBDP0071106, Bukit Timah, primary forest (BT01), 09.nov. 2016, MIP leg.; ZRCBDP0078968, Singapore, date range 2012-2018, MIP leg.; ZRCBDP0078998, Singapore, date range 2012-2018, MIP leg.; ZRCBDP0079013, Singapore, date range 2012-2018, MIP leg.; ZRCBDP0136941, Bukit Timah, primary forest (BT08), 12.jul. 2017, MIP leg.; ZRCBDP0136970, Bukit Timah, primary forest (BT08), 12.jul. 2017, MIP leg.; ZRCBDP0136989, Bukit Timah, primary forest (BT06), 28.jun. 2017, MIP leg.; ZRCBDP0136990, Bukit Timah, primary forest (BT06), 28.jun. 2017, MIP leg.; ZRCBDP0136993, Bukit Timah, primary forest (BT06), 28.jun. 2017, MIP leg.; ZRCBDP0137013, Bukit Timah, primary forest (BT06), 19.apr. 2017, MIP leg.; ZRCBDP0137015, Bukit Timah, primary forest (BT06), 19.apr. 2017, MIP leg.; ZRCBDP0137043, Bukit Timah, primary forest (BT06), 07.jun. 2017, MIP leg.; ZRCBDP0278178, Pulau Ubin, mangrove (PU18), 10.may. 2018, MIP leg.; ZRCBDP0137089, Bukit Timah, primary forest (BT05), 15.Mar. 17, MIP leg. **Females**: ZRCBDP0137009, Bukit Timah, maturing secondary forest (BT06), 19.apr.17, MIP leg.; ZRCBDP0137077, Bukit Timah, maturing secondary forest (BT06), 17.oct. 17, MIP leg.; ZRCBDP0137109, Bukit Timah, old secondary forest (BT07), 17.may. 17, MIP leg.; ZRCBDP0049262, Nee Soon (NS1), 10.dec. 2014, MIP leg. (slide-mounted). **Additional sequenced specimens:** ZRCBDP0066821, Bukit Timah, primary forest (BT09), 05.oct.16; ZRCBDP0079014, Singapore; ZRCBDP0082314, Bukit Timah, primary forest (BT05), 09.nov.16; ZRCBDP0136983, Bukit Timah, primary forest (BT08), 12.apr.17; ZRCBDP0137007, Bukit Timah, primary forest (BT06), 19.apr.17; ZRCBDP0154886, Singapore.

**Etymology**. The species epithet refers to the transcription of the name “Temasek” by Marco Polo, as Chiamassie. Temasek is an early name of a settlement corresponding to modern Singapore. The noun is used in apposition.

**Remarks**. The sequences we have for this species come from specimens from both, Bukit Timah Forest and from Nee Soon Swamp Forest. The haplotype network shows the one haplotype that is separated from a couple of other haplotypes but there are no delimitation conflicts

*Manota danmaxi* Amorim & Oliveira, sp. n. (Figs. 30A-G)

https://singapore.biodiversity.online/species/A-Arth-Hexa-Diptera-000783

urn:lsid:zoobank.org:act:F1C163EA-4BF2-404A-9B6A-5407A0C7D2D2

**Diagnosis**. Thorax ochre-yellowish on anterior half, more brownish towards posterior end, katepisternum lighter. Laterotergite bare. Macrotrichia on wing anal lobe membrane. Abdominal tergites more brownish. Gonocoxites ventromedial margin lobes well separate, setose; parastylar lobe long, with a pair of long setae distally, a small basal projection with a single small fine seta distally directed ventrally; juxtagonostylar lobe with a pair of long, hook-like juxtagonostylar setae directed inwards distally; no apicolateral lobe; gonostylus rectangular, flat, elongate, one megaseta and nine smaller setae distally, a basal digitiform projection with three elongate setae at apex; tergite 9 with wide posterior incision separating a pair of lateral lobes, dorsal face covered with elongate setae, longer closer to posterior margin, an internal lobe near posterior margin with five elongate setae directed inwards and a concentrated group of elongate small setae slightly more anteriorly.

***Manota_danmaxi_ZRCBDP0049202_hapZRCBDP0049 193_SMH_holotype* [124: T, 139: C, 145: A, 154: C, 155 : C, 157: T, 169: T, 196: C]**

tttagctgcttcaattgctcattcaggagcttcagttgacttatctattttttcccttcatttagctggaatttcttctattttaggagctgtaaattttattactacaattattaatatacgTgccccaaatataaaCtttacAcaaataccCCtTtttgtatgatcTgttttaattacagcaattttacttctCttatctcttcctgtattagcaggagctattactatacttttaacagatcgaaatttaaatacttctttttttgatcctgcaggaggtggagatcctattttatatcaacatttattt

#### Description

**Male** (Fig. 30A). Wing length, 1.84; width, 0.74. **Head**. Greyish-brown. Antennal scape and pedicel light greyish-brown, flagellum light brown. Face light yellowish-brown, clypeus, palpus and labella whitish. Occiput with ten strong postocular setae. Flagellomeres 1–5 about as long as wide, flagellomeres 6–13 slightly longer than wide, flagellomere 14 more than twice as long as wide. Maxillary palpomere 4 only slightly longer than palpomere 3; palpomere 5 only slightly longer than palpomere 4. **Thorax** (Fig. 30C). Scutum light brown, darker medially on posterior end, scutellum light brown; antepronotum and proepisternum light ochre-brown; anepisternum and mesepimeron ochre with brownish tinge, katepisternum cream-yellowish with light brownish tinge, laterotergite greyish-ochre, mediotergite light brown on dorsal half, greyish-ochre on ventral half; metepisternum greyish-brown. Anepisternum with small scattered setae over dorsal ¾; anterior basalare bare; katepisternum with a medial vertical line with 10 small setae near posterior end; mesepimeron bare; laterotergite bare; metepisternum with 12 small fine setae on anterior half. Haltere stem light brown, knob dark brown. **Legs**. Coxae whitish, fore coxa with light brownish tinge, femora brownish-yellow, tibiae and tarsi light-brown. **Wing** (Fig. 30B). Membrane brownish fumose; C extending for 4/5 of distance to M_1_; R_1_ meeting C before mid of wing; sclerotized part of M_2_ originating slightly beyond level of tip of R_1_. Membrane of anal lobe, cell cua and cell m4 distally with dorsal macrotrichia. **Abdomen**. Tergites 1–6 brownish; sternites 1–6 light brown, tergite and sternite 7 light brown. Abdominal setosity brownish. **Terminalia** (Figs. 30D–E). Light brown, with yellowish cerci. Sternite 9 trapezoid with rounded posterior end, a deep anterior medial incision almost reaching mid of gonocoxite length, a slender fusion laterally to gonocoxites, medial depression separating a pair of elongate, setose lobes. Gonocoxites ventromedial margin lobes well separate, setose; parastylar lobe long, with a pair of long setae distally, a small basal projection with a single small fine seta distally directed ventrally; juxtagonostylar lobe with a pair of long, hook-like juxtagonostylar setae directed inwards distally; no apicolateral lobe. Gonostylus rectangular, flat, elongate, one megaseta and nine smaller setae distally, a basal digitiform projection with three elongate setae at apex. Gonocoxal bridge with apodemes directed obliquely inwards. Tegmen triangular, aedeagus elongate, curved forwards and sinuous distally, anteriorly with a pair of oblique apodemes directed outwards. Sternite 10 well-developed, projected to level of tip of tegmen, distal margin rounded, with two pairs of setae medially and a stronger additional external seta, a pair of wide digitiform projections with a number of fine setae distally laterad to opening of aedeagus. Tergite 9 with wide posterior incision separating a pair of lateral lobes, dorsal face covered with elongate setae, longer closer to posterior margin, an internal lobe near posterior margin with five elongate setae directed inwards and a concentrated group of elongate small setae slightly more anteriorly. Cerci elongate, flat, entirely separate from each other, with microtrichia and fine setae, long setae distally.

**Female**. Wing length, 1.76; width, 0.66. **Head**. Flagellomeres 1–7 about as long as wide, flagellomeres 8– 13 slightly longer than wide, flagellomere 14 more than twice as long as wide. **Thorax**. Metepisternum with 21 small fine setae along sclerite. **Wing**. Macrotrichia dorsally on membrane of anal lobe. **Terminalia** (Figs. 30F–G). Sternite 8 wide at base, trapezoid, distal end with a pair of elongate lobes and a medial incision posteriorly, fine setae entirely covering sclerite, especially on lobes, three longer setae at tip of each lobe. Sternite 9 present as a pair of long sclerotized bands connected distally and diverging towards anterior end, extending anteriorly to distal end of segment 7, anterior arm weakly sclerotized, distal medial end of sternite 9 almost reaching level of tip of lobes of sternite 8, gonopore connected to two gonoducts. Tergite 8 apparently fused to tergite 9, slightly longer posterior at laterals than medially, covered with microtrichia and scattered fine setae. Sternite 10 with one pair of outer long setae and one pair of inner short setae on distal margin. Tergite 10 reduced to an inconspicuous slender band with two pairs of setae at tip of long digitiform projection, one pair at lateral corners and one pair sub-medially. Cercomere 1 slender, 3.5× longer than wide, cercomere 2 rounded, covered with microtrichia and fine setae.

#### Material examined

**Holotype**: male, ZRCBDP0049202, Nee Soon (NS2), may.15, MIP leg. (website photo specimen, slide-mounted). **Paratypes** (2 males, 1 female). **Male**: ZRCBDP0049193, Nee Soon (NS2), 13.may.15, MIP leg.; ZRCBDP0137040, Bukit Timah (BT06), 01.feb.17. **Female**: ZRCBDP0074033, Bukit Timah, primary forest (BT05), 08.dec. 16, MIP leg. (slide-mounted).

**Etymology**. The species epithet refers to the transcription of the name “Temasek” as recorded in Yuan and Ming Chinese documents as Dan Ma Xi [=淡馬錫]. Temasek is an early name of a settlement corresponding to modern Singapore. The noun is used in apposition.

**Remarks**. Bukit Timah and Nee Soon samples are the source of specimens examined of this species.

*Manota mahuan* Amorim & Oliveira, sp. n. (Figs. 31A–E)

https://singapore.biodiversity.online/species/A-Arth-Hexa-Diptera-000817

urn:lsid:zoobank.org:act:B3D67417-A320-4819-81D7-BFD518D724D6

**Diagnosis**. Thorax bright ochre-yellowish, more brownish on posterior fourth medially, pleura very light brown, katepisternum lighter. Laterotergite setose. Macrotrichia on anal lobe membrane. Anterior abdominal segments more dirty-yellowish, segments 4–6 brown. Gonocoxite ventromedial margin lobe setose; parastylar lobe with a digitiform projection ventrad bearing three elongate setae at tip; juxtagonostylar lobe with a wide ventral area from which a slender digitiform is projected, distally with a pair of long juxtagonostylar megasetae directed inwards, one of them pointed, the other one capitate at tip; dorsomedial border of gonocoxite approximate to each other medially, posterior margin with a distal extension with a number of long setae at dorsal face and a strong curved setae distally, inner posterior corner with 4–5 setae directed ventrally; gonostylus more or less flattened, inner and outer faces with microtrichia and elongate setae, distal margin with long, curved setae, a subdistal beak directed dorsally with three bristles directed anteriorly.

***Manota_mahuan_ZRCBDP0047061_hapZRCBDP00470 61_SMH_holotype* [10: C, 125: A, 127: A, 133: T, 144: C, 172: C, 190: G, 196: T, 205: T]**

tttagctgcCtcaattgctcattctggagcatcagtagatttatcaattttttccttacatttagcaggaatttcttcaattttaggggcagtaaattttattacaactattattaatatacgtAcAcctaaTataaattttaCtcaaatacctttatttgtttgatcagtCttaattacagcaattctGttactTttatcactTccagttttagcaggagcaattactatacttttaacagatcgaaatttaaatacgtctttttttgatcctgcggggggtggagacccaattttatatcaacatttattt

#### Description

**Male** (Fig. 31A). Wing length, 1.84; width, 0.66. **Head**. Brown, a dark brown band at level of line of ocelli. Occiput with 10 strong postocular setae on each side. Antennal scape light brownish-yellow, pedicel and basal two flagellomeres light brown, flagellomeres 3–14 brownish. Face light brown, clypeus, palpus and labella whitish-brown. Flagellomeres 1–2 about as long as wide, flagellomeres 3–13 slightly longer than wide, flagellomere 14 more than twice as long as wide. Maxillary palpomere 4 1.4× palpomere 3 length, palpomere 4 with parasegment; palpomere 5 1.4× palpomere 4 length. **Thorax**. Scutum brownish-yellowish, darker on posterior fourth, a brown band medio-posteriorly; scutellum light brown. Antepronotum and proepisternum ochre-yellowish, anepisternum brownish-yellow, katepisternum whitish with light brown tinge, mesepimeron light brownish-yellow, laterotergite ochreish-brown, mediotergite light brown on dorsal half, ochreish-brown on ventral half, metepisternum light brown. Anepisternum covered with small setae on dorsal ¾, katepisternum with a vertical band of small fine setae at level of ventral end of anepisternum; anterior basalare bare; laterotergite setose; metepisternum with 13 fine small setae on anterior ¾. Haltere stem light brown, knob dark brown. **Legs**. Coxae whitish, femora light brownish-yellow, tibiae and tarsi light brown, darker towards tip. **Wing** (Fig. 31B). Membrane brownish fumose. C extending to 4/5 of distance to M_1_; R_1_ meeting C before mid of wing; sclerotized part of M_2_ originating slightly beyond level of tip of R_1_. Macrotrichia on membrane of anal lobe and on cell cua. **Abdomen**. Tergites 1–3 light brownish-yellow with brown mark medio-posteriorly, tergite 4 mostly brownish, with brownish-yellow anterolateral corners, tergites 5–6 brownish, tergite 7, brownish-yellow; sternites 1–5 light brown, sternite 5–6 brown, sternite 7 brownish-yellow. **Terminalia** (Figs. 31C–D). Terminalia brownish-yellow with whitish gonostylus. Sternite 9 trapezoid, a pair of lobes with a deep anterior medial incision on posterior end and on anterior end, about half of gonocoxite length, slender fusion to gonocoxites at lateral ends. Gonocoxite lobe at ventromedial margin setose, oblique on distal half; parastylar lobe with digitiform projection ventrally bearing three elongate setae at tip; juxtagonostylar lobe with a wide ventral area from which a slender digitiform is projected, distally with a pair of long juxtagonostylar megasetae directed inwards, one of them pointed, the other capitate at tip; dorsomedial border of gonocoxite approximate to each other medially, posterior margin with a distal extension bearing a number of long setae at dorsal face and a strong distal curved setae, inner posterior corner with 4–5 setae directed ventrally. Gonostylus about half of gonocoxite length, elongate, more or less flattened, inner and outer faces with microtrichia and elongate setae, distal margin with long, curved setae, a subdistal beak directed dorsally with three strong setae directed anteriorly. Gonocoxal bridge with apodemes directed obliquely. Tegmen triangular, with curved distal end tubular ending pointed, reaching level of mid of gonostylus, apodemes anteriorly directed obliquely outwards. Sternite 10 well-developed, triangular, projected to level of tip of aedeagus opening, with three long setae at distal end, a row of long curved setae on each side directed ventrolaterally close to tip. Tergite 9 arched distally, with scattered microtrichia dorsally and a band of setae along distal margin, four strong setae at margin medially. Cerci elongate, flat, entirely separate from each other, with microtrichia and fine setae, long setae distally.

**Female**. Wing length, 1.76; width, 0.64. **Head**. Flagellomeres 1–4 about as long as wide, flagellomeres 5– 13 slightly longer than wide, flagellomere 14 more than twice as long as wide. **Thorax**. Metepisternum with 21 small fine setae along sclerite. **Wing**. Macrotrichia dorsally on membrane of anal lobe. **Terminalia** (Figs. 31E). Sternite 8 wide at base, trapezoid, distal end with a pair of elongate lobes and a medial incision posteriorly, fine setae entirely covering sclerite, especially on lobes, three longer setae at tip of each lobe. Sternite 9 present as a pair of long sclerotized bands connected distally and diverging towards anterior end, extending anteriorly to distal end of segment 7, anterior arm weakly sclerotized, distal medial end of sternite 9 almost reaching level of tip of lobes of sternite 8, gonopore connected to two gonoducts. Tergite 8 apparently fused to tergite 9, slightly wider at posterior margin than medially, covered with microtrichia and scattered fine setae. Sternite 10 with one pair of long outer setae and one pair of inner short setae on distal margin. Tergite 10 reduced to an inconspicuous slender band and with two pairs of setae, at tip of long digitiform projections, one pair at lateral corners and one pair sub-medially. Cercomere 1 over 3× longer than cercomere 2 rounded, both covered with microtrichia and fine setae.

#### Material examined

**Holotype**: male, ZRCBDP0047061, National University of Singapore (PGP), 08.jul.15, MIP leg. (slide-mounted). **Paratypes** (7 males, 3 females). **Males**: ZRCBDP0047100, National University of Singapore (PGP), 08.jul.15, MIP leg.; ZRCBDP0047101, National University of Singapore (PGP), 08.jul.15, MIP leg.; ZRCBDP0049301, National University of Singapore (Icube), 01.apr.15, MIP leg.; ZRCBDP0066723, Bukit Timah, maturing secondary forest (BT06), 16.aug. 16, MIP leg.; ZRCBDP0048305, Pulau Semakau (SMO2), old mangrove, 26.dec.13, MIP leg.; ZRCBDP0048762, Nee Soon (NS1), 25.feb.15, MIP leg. (extracted); ZRCBDP0049338, National University of Singapore (PGP), 08.apr.15, MIP leg. (extracted, website photo specimen). **Females**: ZRCBDP0048936, Nee Soon (NS1), 31.dec.14, MIP leg.; ZRCBDP0049137, National University of Singapore (PGP), 22.apr.15, MIP leg. (slide-mounted); ZRCBDP0072694, Bukit Timah, primary forest (BT05), 01.dec. 16, MIP leg. **Additional sequenced specimens:** ZRCBDP0066834, Singapore; ZRCBDP0132821, Singapore; ZRCBDP0132822, Singapore; ZRCBDP0133477, Singapore; ZRCBDP0133516, Singapore; ZRCBDP0133523, Singapore; ZRCBDP0133532, Singapore; ZRCBDP0133533, Singapore; ZRCBDP0278038, Singapore Botanical Gardens (CUGE), 17.nov.17; ZRCBDP0278057, Singapore Botanical Gardens (CUGE), 15.dec.17; ZRCBDP0278263, Singapore, 07.jun.18; ZRCBDP0278269, Singapore, 07.jun.18; ZRCBDP0278474, Pulau Ubin (PU17), 17.may.18; ZRCBDP0279125, Singapore, 31.may.18; ZRCBDP0279176, Singapore, 31.may.18; ZRCBDP0284245, Singapore; ZRCBDP0284246, Singapore; ZRCBDP0284247, Singapore; ZRCBDP0284250, Singapore; ZRCBDP0284251, Singapore; ZRCBDP0284254, Singapore; ZRCBDP0284256, Singapore.

**Etymology**. This species is named after Ma Huan [=马欢] (1380–1460), a Chinese Muslim voyager and scribe who followed the Chinese Admiral Zheng. He explored Southeast Asia and much of the rest of the old world. His book Ying Ya Sheng Lan [=瀛涯胜览, The Overall Survey of the Ocean’s Shores] was one of the first extensive natural and cultural history records of Southeast Asia, long before Wallace and other Western naturalists.

His records mention the Long Ya Men [=龙牙门; Dragon’s Teeth Gate], a navigational rock off the coast of Singapore. The noun is used in apposition.

**Remarks**. This is a widespread species in Singapore, occurring in different environments, from the mangroves to swamp forest, primary tropical forest, as well as degraded secondary forest in urbanized areas. This species is similar to, e. g., *Manota anceps* Hippa & Ševčík and *Manota capillata* Hippa &Ševčík, from Sumatra, *Manota dolichothrix* Hippa & Ševčík (Malaysia) and *Manota hyboloma* Hippa & Ševčík, from Brunei (Hippa & Ševčík 2010). We have four haplotypes for this species and no delimitation conflicts.

*Manota temenggong* Amorim & Oliveira, sp. n. (Figs. 32A–E)

https://singapore.biodiversity.online/species/A-Arth-Hexa-Diptera-000814

urn:lsid:zoobank.org:act:124E3C8B-9896-495F-BADD-74AB1EDCAAD4

**Diagnosis**. Small specimens, < 1.50 mm. Head, thorax and abdomen brown, distal third of hind femur brownish. Laterotergite setose. Anal lobe devoid of setation. Gonocoxite ventromedial margin lobe large; parastylar lobe with digitiform projection bearing three elongate fine setae at tip directed inwards; juxtagonostylar lobe slender, digitiform, distally with a pair of fine long juxtagonostylar setae directed inwards; dorsomedial border of gonocoxite with a distal extension, almost reaching tip of gonostylus, inner face with a dense group of strong curved setae. Gonostylus about half of gonocoxite length, distal margin with a pair of short, more or less pointed projections with microtrichia and elongate setae on external face, distal margin with long, curved setae, long setae directed inwards on distal half of inner face. Tergite 9 arched distally, with scattered microtrichia dorsally and a band of setae along distal margin, four strong setae at margin medially.

***Manota_temenggong_ZRCBDP0049118_hapZRCBDP0 049118_SMH_holotype* [11: T, 100: C, 130: A, 140: C, 1 42: A, 145: T, 160: C, 275: A]**

tttagcttctTcaattgctcattctggagcttcagtagatttatcaattttttctcttcacttagcaggaatttcatcaattttaggagcagtaaatttCattacaacaattgttaatatacgagctccAgaaataaatCtAacTcaaatacctttattCgtttgatcagttttaattacagctattttattattattatcattaccagtattggcaggagcaattactatattattaacagatcgaaacttaaatacatcatttttcgacccaAcaggaggaggagatccaattttatatcaacatttattc

#### Description

**Male** (Fig. 32A). Wing length, 1.40; width, 0.54. **Head**. Brown, a darker band at level of line of ocelli and on occiput posteriorly to eye. Antennal scape brownish-yellow, pedicel and flagellum light brown. Flagellomere 1 about as long as wide, flagellomeres 2–13 slightly longer than wide, flagellomere 14 more than twice as long as wide. Face, clypeus, palpus and labella whitish-brown. Maxillary palpomere 4 only slightly longer than palpomere 3, palpomere 5 slightly over 2.0× palpomere 4 length. Eight strong postocular setae on each side. **Thorax** (Fig. 32C). Scutum brown, darker medially, scutellum brown. Antepronotum, proepisternum, anepisternum, mesepimeron, laterotergite and metepisternum light brown, mediotergite light brown, brown medially, katepisternum whitish with a light brown tinge. Anepisternum covered with small setae on dorsal ¾; anterior basalare bare; katepisternum bare; laterotergite with 19 small setae; metepisternum with eight small fine setae on anterior half. Haltere stem light brown, knob brown. **Legs**. Coxae whitish, femora yellowish, hind femur brown on distal fifth; tibiae and tarsi brownish, darker towards tip. **Wing** (Fig. 32B). Membrane brownish fumose, posterior margin slightly emarginated at tip of CuA. C extending to 4/5 of distance to M_1_; R_1_ meeting C before mid of wing; sclerotized part of M_2_ originating well beyond level of tip of R_1_. M_2_ gently curved at basal fourth. CuA not disconnected from M_4_ at base. **Abdomen**. Tergites 1–6 brownish with lighter lateral margins, tergite 7 light brown; sternites 1–7 light brown. **Terminalia** (Figs. 32D–E). Yellowish-brown. Sternite 9 trapezoid, a pair of lobes with a deep anterior medial incision on posterior end and on anterior end, about half of gonocoxite length. No fusion of sternite 9 laterally to gonocoxites. Gonocoxite ventromedial margin lobe large, ventral face bare only at inner corner anteriorly; parastylar lobe with digitiform projection bearing three elongate fine setae at tip directed inwards; juxtagonostylar lobe present as a slender digitiform distally with a pair of fine long juxtagonostylar setae directed inwards; dorsomedial border of gonocoxite with a distal extension almost reaching tip of gonostylus, dorsal face with long fine setae, inner face with a group of concentrated strong curved setae. Gonostylus about half of gonocoxite length, subquadrate elongate, more or less flattened, distal margin with a pair of short, more or less pointed projections, with microtrichia and elongate setae on external face, distal margin with long, curved setae, long setae directed inwards on distal half of inner face. Gonocoxal bridge with apodemes directed obliquely, slightly curved. Tegmen triangular, with curved tubular distal end pointed at tip, reaching level of base of gonostylus, pointed apodemes at anterior end directed obliquely outwards. Sternite 10 well-developed, triangular, projected to level of tip of aedeagus opening, with three long setae at distal end, close to tip laterally with a row of long curved setae directed ventrally on each side. Tergite 9 arched distally, with scattered microtrichia dorsally and a band of setae along distal margin, four strong setae at margin medially. Cerci elongate, flat, entirely separate from each other, with microtrichia and fine setae, long setae distally.

**Female**. Unknown.

#### Material examined

**Holotype**: male, ZRCBDP0049118, Nee Soon (NS1), 24.dec.14, MIP leg. (website photo specimen, slide-mounted).

**Etymology**. The species epithet is a reference to “Temenggong”, an old Malay title of nobility, usually given to the chief of public security, responsible for the safety of the Sultan and overseeing the state’s security forces. The Temenggong Abdul Rahman, of the Johor Sultanate, helped Raffles set a formal treaty with the Sultan Hussein of Johor, signed on February 6, 1819, resulting in Singapore’s colony status. The noun is used in apposition.

***Manota* sp**. **A** (Figs. 33A–B)

https://singapore.biodiversity.online/species/A-Arth-Hexa-Diptera-000759

***Manota_spA_ZRCBDP0047870_hapZRCBDP0047870_ SMH_unnamed_type* [7: G, 10: C, 28: C, 127: T, 138: A, 145: A, 199: C, 205: C]**

tttagcGgcCtctattgctcattcaggCgcatctgtagatttatcaattttttctttacatttagcaggaatttcatcaattttaggagcagtaaattttattacaactattattaatatacgaacTcctaatataaAttttacAcaaataccattatttgtttgatcagttttaattacagcaattttacttctcctCtcactCccagttttagcaggagctattactatacttttaacagatcgaaatttaaatacatctttttttgatcccgcaggtggaggagatccaattttatatcaacatttattt

#### Description

**Female**. Wing length, 1.81; width, 0.71. **Head**. Brownish-yellow, a dark brown band at level of line of ocelli and on occiput. Antenna light yellowish-brown, scape and pedicel lighter. Face, clypeus, palpus and labella dirty-yellowish. Occiput with line of 9–10 strong postocular setae. **Thorax**. Scutum brownish cream-yellow, as well as antepronotum; proepisternum, anepisternum, and mesepimeron dark ochre-yellowish, katepisternum and metepisternum lighter; laterotergite brown, mediotergite light brown. Five supra-alars, two pairs of prescutellars. Anepisternum covered with fine setae except for anterior margin, including anteroventral corner; anterior basalare bare; katepisternum with some fine setae close to posterior border; laterotergite bare; metepisternum with 18–20 setulae. Haltere brownish-yellow, setae on pedicel, knob densely setose. **Legs**. Coxae whitish, fore femur whitish-yellow, mid and hind femora light brownish-yellow; tibiae and tarsi light greyish-brown. Mid and hind tibial organs absent. **Wing** (Fig. 33A). Membrane light greyish-brown. C almost reaching tip of M_1_, R_1_ meeting C before mid of wing. First sector of Rs not produced. Sclerotized part of M_2_ originating at level of tip of R_1_; CuA not connected basally to M_4_. **Abdomen**. Tergites 1–2 brownish-yellow, tergites 3–7 light brown; sternites 1–3 light brown, sternite 4–6 light yellowish-brown. **Terminalia** (Fig. 33B). Sternite 8 wide at base, trapezoid, distal end with a pair of short lobes bearing a medial incision posteriorly, fine setae entirely covering sclerite, larger setae on lobes, setae on distal margin of lobes longer. Sternite 9 present as a pair of long sclerotized bands connected distally and diverging towards anterior end, extending anteriorly to distal end of segment 7, distal medial end of sternite 9 reaching level of tip of lobes of sternite 8, with setulae along posterior margin, gonopore connected to a pair of gonoducts. Tergite 8 wide and short, almost not overlapping laterally with lateral border of sternite 8, covered with microtrichia and fine setae. Tergite 9 longer than tergite 8, covered with microtrichia and scattered fine setae. Sternite 10 with two pairs of longer setae. Tergite 10 reduced to an inconspicuous slender band, two pairs of setae, at tip of long digitiform projections, one pair at lateral corners and one pair sub-medially. Cercomere 1 slender, over 3× longer than wide, cercomere 2 ovoid, covered with microtrichia and fine setae.

**Male**. Unknown.

**Material examined** (3 females): ZRCBDP0047870, Nee Soon (NS1), swamp forest, 26.mar.14, MIP leg. (slide-mounted); ZRCBDP0048529, Nee Soon (NS2), swamp forest, 30.may. 12, MIP leg. (website photo specimen); ZRCBDP0048672, Nee Soon (NS1), swamp forest, 04.apr. 12, MIP leg.

**Remarks**. Only specimens from the swamp forest were found for this species. We have two haplotypes and no delimitation conflicts.

***Manota* sp**. **B** (Figs. 34A–C)

https://singapore.biodiversity.online/species/A-Arth-Hexa-Diptera-000820,and-000759

***Manota_spB_ZRCBDP0047826_hapZRCBDP0047826_ SMH_unnamed_type* [1: A, 7: C, 8: G, 11: G, 37: G, 64: T, 67: G, 73: C, 76: C, 145: A]**

AttagcCGcaGctattgctcattcagggtcttcagtGgatttatctattttttctctgcatctTgcGggtatCtcCtcaattttaggggcagtaaattttattacaactattataaatatgcgaacacctgaaatatcattaacAcaaatacctttatttgtatgatcagttctaattacagcaattctacttttattatctcttcctgttttagctggtgctattactatattattaacagatcgaaatttaaatacaactttttttgatcctataggaggaggagatccaattttatatcaacatttattt

#### Description

**Female** (Fig. 34A). Wing length, 1.76; width, 0.66. **Head**. Light brown. Antennal scape light brownish-yellow, pedicel and flagellum light brown. Flagellomeres 1–5 about as long as wide, flagellomeres 6–13 slightly longer than wide, flagellomere 14 more than twice as long as wide. Face light brown, clypeus, palpus and labella whitish. Maxillary palpomere 4 2.0× length of palpomere 3; palpomere 5 1.1× palpomere 4 length. Occiput with ten strong postocular setae. **Thorax**. Scutum yellowish-brown, darker medially; antepronotum and proepisternum light ochre-brown; anepisternum, mesepimeron, laterotergite and mediotergite yellowish-brown, katepisternum whitish-brown, metepisternum whitish-brown on anterior fourth, brownish on posterior ¾, laterotergite with blackish-brown mark on ventral end. Anepisternum with small scattered setae over dorsal half; anterior basalare bare; katepisternum with a medial vertical line with 10 small setae; mesepimeron bare; laterotergite with about 30 scattered fine setae; metepisternum with nine small setae on anterior half. Haltere stem light brown, knob dark brown. **Legs**. Coxae whitish, trochanters brownish, femora brownish-yellow, tibiae and tarsi yellowish-brown. **Wing** (Fig. 34B). Membrane brownish; R_1_ meeting C before mid of wing; sclerotized part of M_2_ originating at level of tip of R_1_. No dorsal macrotrichia on wing membrane. **Abdomen**. Tergites 1–6 brownish; sternites 1–6 light brown, tergite and sternite 7 light brown. Abdominal setosity brownish. **Terminalia** (Fig. 34C). Light brown, with yellowish cerci, brownish tinges ventrally. Sternite 8 wide at base, trapezoid, distal end with a pair of lobes and a medial incision posteriorly, fine setae entirely covering sclerite, especially on lobes, two longer setae at tip of each lobe. Sternite 9 present as a pair of long sclerotized bands connected distally and diverging towards anterior end, extending anteriorly to distal end of segment 7, anterior arm ending very thin, distal medial end of sternite 9 almost reaching level of tip of lobes of sternite 8, gonopore connected to a single gonoduct. Tergite 8 apparently fused to tergite 9, slightly longer posterior laterally than medially, covered with microtrichia and scattered fine setae. Sternite 10 with one pair of outer long setae and one pair of inner short setae on distal margin. Tergite 10 reduced to an inconspicuous slender band and with two pairs of setae at tip of long digitiform projection, one pair at lateral corners and one pair sub-medially. Cercomere 1 slender, about 3× longer than wide, cercomere 2 ovoid, covered with microtrichia and fine setae.

**Male**. Unknown.

**Material examined** (1 female): ZRCBDP0047826, Nee Soon (NS1), swamp forest, 06.mar.13, MIP leg. (website photo specimen, slide-mounted).

**Remarks**. The specimen of this species was collected in the swamp forest.

***Manota* sp**. **C** (Figs. 35A–F)

https://singapore.biodiversity.online/species/A-Arth-Hexa-Diptera-000773

***Manota_spC_ZRCBDP0048300_hapZRCBDP0048300_ SMH_unnamed_type* [10: A, 31: C, 40: C, 55: A, 88: T, 125: A, 145: A, 154: A, 229: C]**

tttagcttcAtctattgctcattctggagcCtcagtagaCttatcaattttttcAttacatttagcaggaatttcttcaattttaggTgcagtaaattttattacaacaattattaatatacgaActccaaatataaattttacAcaaataccAttatttgtatgatcagttttaattacagctattcttttactattatctttaccagtattagcaggagctattacCatattattaactgatcgaaatttaaatacatcattttttgatcctgctggaggaggagatccaattttatatcaacatttattt

#### Description

**Female** (Fig. 35A). Wing length, 1.79; width, 0.66. **Head** (Fig. 35B). Brown, ocellar line darker. Antennal scape light brownish-yellow, pedicel and flagellum light brown. Flagellomeres 1–10 about as long as wide, flagellomeres 11–13 slightly longer than wide, flagellomere 14 more than twice as long as wide. Face densely setose. Face and clypeus light brown, labella yellowish-brown [both maxillary palpi missing]. Nine strong postocular setae on each side. **Thorax** (Fig. 35C). Scutum and scutellum light brown. Antepronotum and proepisternum, anepisternum, mesepimeron, laterotergite, and mediotergite light brown, katepisternum whitish with brown tinge, metepisternum light brown dorsally, whitish ventrally. Anepisternum covered with small setae except for bare anteroventral corner; anterior basalare bare; laterotergite bare, metepisternum with some setulae on anterior half. Haltere stem light brown, knob brown. **Legs**. Coxae whitish, femora brownish-yellow, tibiae and tarsi brownish, darker towards tip, covered with regular rows of brownish setulae and regular rows of short brown bristles. Fore tibia shorter than femur and shorter than first tarsomere. **Wing** (Fig. 35D). Membrane brownish fumose. Wing margin slightly emarginated at level of tip of CuA. C extending over 4/5 of distance to M_1_. R_1_ meeting C before mid of wing; sclerotized part of M_2_ originating beyond level of tip of R_1_. **Abdomen**. Tergites 1–6 brownish; sternites 1–6 light brown, tergite and sternite 7 light brown. Abdominal setosity brownish. **Terminalia** (Figs. 35E–F). Light brown, cerci yellowish with brownish tinge. Sternite 8 wide at base, trapezoid, distal end with a pair of lobes and a medial incision posteriorly, fine setae entirely covering sclerite, especially on lobes, two longer setae at tip of each lobe. Sternite 9 present as a pair of long sclerotized bands connected distally and diverging towards anterior end, extending anteriorly to distal end of segment 7, anterior arm weakly sclerotized, distal medial end of sternite 9 almost reaching level of tip of lobes of sternite 8, gonopore connected to a pair of gonoducts. Tergite 8 apparently fused to tergite 9, slightly longer laterally than medially, covered with microtrichia and scattered fine setae. Sternite 10 with one pair of outer long setae and one pair of inner short setae on distal margin. Tergite 10 reduced to an inconspicuous slender band and with two pairs of setae at tip of long digitiform projection, one pair at lateral corners and one pair sub-medially. Cercomere 1 slender, slightly over 3× longer than wide, cercomere 2 ovoid, short, covered with microtrichia and fine setae.

**Male**. Unknown.

**Material examined** (1 female): ZRCBDP0048300, Pulau Semakau (SMO1), old mangrove, 20.dec. 12, MIP leg. (website photo specimen, slide-mounted).

***Manota* sp**. **D (**Figs. 36A–D)

https://singapore.biodiversity.online/species/A-Arth-Hexa-Diptera-000825

***Manota_spD_ZRCBDP0048676_hapZRCBDP0048676_ SMH_unnamed_type* [49: C, 76: A, 82: C, 139: C, 202: C, 238: G, 250: C, 259: C]**

attagcagcttctattgctcattcagggtcttcagttgatttatctatCttttctcttcatttagctggtatttcAtctatCttaggggctgtaaattttattactacaattattaatatacgagccccaaatataaaCtttactcaaatacctttatttgtttgatcagttttaattactgctattttacttcttttatcCttacctgtattagctggagctattactatattattGactgatcgaaaCttaaatacCtcattttttgacccagctggaggaggagatccaatcttatatcaacacttattt

#### Description

**Female** (Fig. 36A). Wing length, 1.99; width, 0.69. **Head**. Vertex brown, face light brown, clypeus whitish-yellow, occiput yellowish with light brownish tinge, brown along posterior margin of eye; 11 strong postocular setae on each side. Antennal scape and pedicel whitish-yellow, flagellum light brown. Maxillary palpus and labella whitish. Flagellomeres slightly shorter than long, except for distal flagellomere. **Thorax**. Scutum mostly ochre-yellowish, scutellum light brown. Antepronotum and proepisternum ochre-yellowish, anepisternum, mesepimeron, and laterotergite ochre with brownish tinge, katepisternum whitish, mediotergite light brown on dorsal half, ochre on ventral half, metepisternum light brown. Three pairs of long prescutellars. Anepisternum covered with scattered small setae except at ventral fourth; anterior basalare bare; katepisternum with a band of fine setae dorsoposteriorly at level of ventral end of anepisternum; laterotergite bare; metepisternum with about 30 fine setae along its length. Haltere stem light brown, knob dark brown. **Legs**. Coxae whitish, femora and tibiae whitish-yellow, tarsi with brownish tinge. **Wing** (Fig. 36B). Membrane with light brown infuscation. C ending at about 4/5 of distance to M_1_; R_1_ meeting C before mid of wing; sclerotized part of M_2_ originating at level of tip of R_1_. Membrane with no macrotrichia. **Abdomen**. Tergites 1–6 yellowish-brown, darker medially, tergite 7 light yellowish-brown; sternites 1–3 light brown, sternites 4–6 light yellowish-brown, sternite 7 yellowish. **Terminalia** (Figs. 36C–D). Sternite 8 wide at base, trapezoid, distal end with a pair of lobes and a medial incision posteriorly, fine setae entirely covering sclerite, especially on lobes, two longer setae at tip of each lobe. Sternite 9 present as a pair of long sclerotized bands connected distally and diverging towards anterior end, extending anteriorly to distal end of segment 7, distal medial end of sternite 9 almost reaching level of tip of lobes of sternite 8, gonopore connected to a pair of gonoducts. Tergite 8 wide, barely overlapping laterally with lateral border of sternite 8, covered with microtrichia and fine setae. Tergite 9 about as long as tergite 8, covered with microtrichia and scattered fine setae. Sternite 10 with one pair of outer long setae and one pair of inner short setae on distal margin. Tergite 10 reduced to an inconspicuous slender band and with two pairs of setae, each at tip of a long digitiform projection, one pair at lateral corners and one pair sub-medially. Cercomere 1 slender, about 2.0× longer than wide, cercomere 2 ovoid, slightly elongate, covered with microtrichia and fine setae.

**Male**. Unknown.

**Material examined** (1 female): ZRCBDP0048676, Nee Soon (NS2), swamp forest, 23.may. 12, MIP leg. (website photo specimen, slide-mounted).

***Manota* sp**. **E (**Figs. 37A–C)

https://singapore.biodiversity.online/species/A-Arth-Hexa-Diptera-002076

***Manota_spE_ZRCBDP0133494_unnamed_type* [13: T, 6 7: C, 73: C, 100: C, 143: A, 144: C, 172: A, 211: A, 235:T]**

cttagctgcttcTattgctcattctggagcttcagtagatttatctattttttcacttcacttagcCggaatCtcttcaattttaggagccgtaaatttCattactacaattattaacatacgagcccctgaaataaactttACtcaaatacctctatttgtttgatcagtAttaattacagcaattttattattactttctttaccagtAttagccggagcaattacaatactTctaacagatcgaaatttaaatacatcattttttgaccctgcaggaggaggagatccaattttatatcaacacttattt

#### Description

**Female** (Fig. 37A). Wing length, 1.40; width, 0.51. **Head**. Brown, a darker band at level of line of ocelli and on occiput posteriorly to eyes. Antennal scape and pedicel yellowish-brown, flagellum light brown. Flagellomere 1–6 wider than long, flagellomeres 7–13 slightly longer than wide, flagellomere 14 more than twice as long as wide. Face light brown, clypeus yellowish-brown, palpus and labella whitish. Maxillary palpomere 4 only slightly longer than palpomere 3, palpomere 5 1.7× palpomere 4 length. Occiput with eight strong postocular setae. **Thorax**. Scutum light brown, lighter along margin on anterior half, scutellum light brown. Antepronotum, proepisternum, anepisternum, mesepimeron, laterotergite, and metepisternum light brown, mediotergite light brown on ventral half, brown on dorsal half, katepisternum whitish with a light brown tinge. Anepisternum covered with small setae on dorsal ¾; anterior basalare bare; katepisternum with a vertical band of small, fine setae at level of ventral end of anepisternum; laterotergite with 19 small setae; metepisternum with 13 small fine setae on its anterior ¾. Haltere stem light brown, knob brown. **Legs**. Coxae whitish with a yellowish tinge, hind coxa with a brown band on basal end, femora whitish, hind femur brown on distal half; tibiae and tarsi light brown, darker towards tip. **Wing** (Fig. 37B). Membrane brownish fumose, posterior margin slightly emarginated at tip of CuA. C extending for 4/5 of distance to M_1_; R_1_ meeting C before mid of wing. Sclerotized part of M_2_ originating beyond level of tip of R_1_. CuA not disconnected from M_4_ at base. Anal fold weakly sclerotized. One single macrotrichium on membrane of anal lobe. **Abdomen**. Tergites 1–6 light brown, darker medio-anteriorly, tergite 7 light yellowish-brown; sternites 1–7 light yellowish-brown. **Terminalia** (Fig. 37C). Yellowish-brown. Sternite 8 wide at base, trapezoid, distal end with a pair of elongate lobes and a medial incision posteriorly, fine setae entirely covering sclerite, especially on lobes, three longer setae at tip of each lobe. Sternite 9 present as a pair of long sclerotized bands connected distally, diverging towards anterior end, anterior arm weakly sclerotized, distal end of sternite 9 almost reaching level of mid of cercomere 1, gonopore connected to two gonoducts. Tergite 8 apparently fused to tergite 9, covered with microtrichia and scattered fine setae. Sternite 10 with one pair of outer long setae and one pair of inner short setae on distal margin. Tergite 10 reduced to an inconspicuous slender band and with two pairs of setae at tip of short digitiform projections, one pair at lateral corners and one pair sub-medially. Cercomere 1 slender, flat, over 8× longer than cercomere 2, cercomere 2 very short, both covered with microtrichia and fine setae. **Male**. Unknown.

**Material examined** (35 females): ZRCBDP0047060, National University of Singapore (PGP), 08.jul.15, MIP leg. (slide-mounted); ZRCBDP0047062, National University of Singapore (PGP), 08.jul.15, MIP leg.; ZRCBDP0047079, National University of Singapore (PGP), 22.jul.15, MIP leg.; ZRCBDP0278303, Singapore, 10.may.18, MIP leg.; ZRCBDP0279123, Singapore, 31.may.18, MIP leg.; ZRCBDP0279127, Singapore, 31.may.18, MIP leg.; ZRCBDP0284252, Singapore, date range 2012-2018, MIP leg.; ZRCBDP0132837, Singapore, date range 2012-2018, MIP leg.; ZRCBDP0133108, Singapore, date range 2012-2018, MIP leg.; ZRCBDP0133113, Singapore, date range 2012-2018, MIP leg.; ZRCBDP0133122, Singapore, date range 2012-2018, MIP leg.; ZRCBDP0133175, Singapore, date range 2012-2018, MIP leg.; ZRCBDP0069329, Singapore, PU01, 11.jun. 16, MIP leg.; ZRCBDP0132841, National University of Singapore (PGP), 24.may. 17, MIP leg.; ZRCBDP0132853, National University of Singapore (PGP), 24.may. 17, MIP leg.; ZRCBDP0132885, National University of Singapore (Uhall), 05.jul. 17, MIP leg.; ZRCBDP0133097, National University of Singapore (PGP), 17.may. 17, MIP leg.; ZRCBDP0133105, National University of Singapore (PGP), 17.may. 17, MIP leg.; ZRCBDP0133109, National University of Singapore (PGP), 17.may. 17, MIP leg.; ZRCBDP0133110, National University of Singapore (PGP), 17.may. 17, MIP leg.; ZRCBDP0133117, National University of Singapore (PGP), 17.may. 17, MIP leg.; ZRCBDP0133126, National University of Singapore (PGP), 17.may. 17, MIP leg.; ZRCBDP0133131, National University of Singapore (PGP), 17.may. 17, MIP leg.; ZRCBDP0133134, National University of Singapore (PGP), 17.may. 17, MIP leg.; ZRCBDP0133144, National University of Singapore (PGP), 17.may. 17, MIP leg.; ZRCBDP0133380, National University of Singapore (PGP), 14.apr. 17, MIP leg.; ZRCBDP0133392, National University of Singapore (PGP), 14.apr. 17, MIP leg.; ZRCBDP0133393, National University of Singapore (PGP), 14.apr. 17, MIP leg.; ZRCBDP0133394, National University of Singapore (PGP), 14.apr. 17, MIP leg.; ZRCBDP0133486, National University of Singapore (PGP), 09.may. 17, MIP leg.; ZRCBDP0133488, National University of Singapore (PGP), 09.may. 17, MIP leg.; ZRCBDP0133494, National University of Singapore (PGP), 31.may. 17, MIP leg. (website photo specimen); ZRCBDP0133498, National University of Singapore (PGP), 31.may. 17, MIP leg.; ZRCBDP0133515, National University of Singapore (PGP), 03.may. 17, MIP leg.; ZRCBDP0133526, National University of Singapore (PGP), 03.may. 17, MIP leg.

**Remarks**. This species, together with *Manota temenggong* Amorim & Oliveira, **sp. n.** and *Manota* sp. D, may form a small clade. They are pretty distinctive from other species of *Manota* being small and entirely brownish. We have five haplotypes and no delimitation conflicts.

***Manota* sp**. **F (**Figs. 38A–E)

https://singapore.biodiversity.online/species/A-Arth-Hexa-Diptera-002075

***Manota_spF_ZRCBDP0133440_unnamed_type* [10: C, 22: C, 103: C, 126: C, 132: A, 133: C, 140: T, 268: C, 30 4: G]**

gttagccgcCtcaattgctcaCtctggggcctcagttgatttatcaattttttctttacatttagctggtatttcttcaattttaggagcagtaaattttatCacaactattattaacatacgagCccctaACataaatTttactcaaatacctttatttgtttgatcagttttaattacagctattttacttcttttatctttaccagttttagcaggagcgattactatacttttaacagatcgaaatttaaatacatcattcttCgatcccgcaggaggaggagatcctatcttatatcaGcatctattt

#### Description

**Female** (Fig. 38A). Wing length, 1.86; width, 0.66. **Head** (Fig. 38B). Brown, a dark brown band at level of line of ocelli and on occiput posteriorly to eye. Antennal scape, pedicel and flagellum light brown. Flagellomere 1 and flagellomeres 6–13 slightly longer than wide, flagellomeres 2–5 as long as wide, flagellomere 14 more than twice as long as wide. Face, clypeus, palpus and labella light brown. Maxillary palpomere 4 about as long as palpomere 3, palpomere 5 1.5× palpomere 4 length. Occiput with eight strong postocular setae. **Thorax**. Scutum light ochre-brown, darker medio-posteriorly, scutellum light brown. Antepronotum, proepisternum, anepisternum, mesepimeron, laterotergite, and metepisternum light ochre-brown, mediotergite light brown dorsally, brown medially, katepisternum whitish with a light brown tinge. Anepisternum almost entirely covered with small setae; anterior basalare bare; katepisternum with a vertical group of small, fine setae medially at level of ventral end of anepisternum; laterotergite bare; metepisternum with three small fine setae along its length. Haltere stem light brown, knob brown. **Legs**. Coxae light yellowish, fore and mid femora whitish; fore and mid tibiae and tarsi light brown, darker towards tip [hind femora, tibia and tarsi missing]. **Wing** (Fig. 38C). Membrane brownish fumose, posterior margin slightly emarginated at tip of CuA. C extending for 4/5 of distance to M_1_; R_1_ meeting C before mid of wing. A row of macrotrichia along part of non-sclerotized M_1_, sclerotized part of M_2_ originating slightly beyond level of tip of R_1_. CuA not disconnected from M_4_ at base. Macrotrichia on membrane of anal lobe and cell cua. **Abdomen**. Tergites 1–6 brownish with lighter lateral margins, tergite 7 light brown; sternites 1–7 light brown. **Terminalia** (Figs. 38D–E). Yellowish-brown. Sternite 8 wide at base, trapezoid, distal end with a pair of elongate lobes, a medial incision posteriorly, fine setae entirely covering sclerite, especially on lobes, three longer setae at tip of each lobe. Sternite 9 present as a pair of long sclerotized bands connected distally, diverging towards anterior end, extending anteriorly to distal end of segment 7, anterior arm weakly sclerotized, distal end of sternite 9 medially almost reaching level of tip of lobes of sternite 8, gonopore connected to two gonoducts. Tergite 8 apparently fused to tergite 9, slightly longer laterally, covered with microtrichia and scattered fine setae. Sternite 10 with one pair of outer long setae and one pair of inner short setae on distal margin. Tergite 10 reduced to an inconspicuous slender band and with two pairs of setae, at tip of long digitiform projections, one pair at lateral corners and one pair sub-medially. Cercomere 1 over 3× longer than cercomere 2, which is rounded, both covered with microtrichia and fine setae.

**Male**. Unknown.

**Material examined** (2 females): ZRCBDP0278396, Singapore Botanical Gardens (CUGE), 03.nov. 17, MIP leg.; ZRCBDP0058640, Bukit Timah Forest (BT08), 14.sep. 16, MIP leg.; Two specimens with abdomen missing: ZRCBDP0072692, Bukit Timah Forest, (BT06), 13.oct. 16, MIP leg.; ZRCBDP0133440, Nee Soon Swamp Forest (NSM1), 28.jan.15, MIP leg. (website photo specimen).

**Remarks**. We have two haplotypes for this species and no delimitation conflicts.

***Manota* sp**. **G (**Figs. 39A–F)

https://singapore.biodiversity.online/species/A-Arth-Hexa-Diptera-002068

***Manota_spG_ZRCBDP0132831_unnamed_type* [2: T, 7: T, 8: G, 40: C, 130: T, 136: A, 193: T, 196: T, 202: T, 2 77: T]**

aTtagcTGcttcaattgctcattctggagcctcagttgaCttatcaattttttctttacatttagcaggtatttcttcaattttaggagcagtaaattttattacaactattattaatatacgagccccTaatatAaattttactcaaatacctttatttgtttgatcagttttaattacagctattttactTctTttatcTttaccagttttagcaggagctattactatacttctaacagatcgaaatttaaatacatcattttttgatcctgcTggtggaggggaccctattttatatcaacatttattt

#### Description

**Female** (Fig. 39A). Wing length, 1.79–1.99; width, 0.66–0.74 (n=2). **Head** (Figs. 39B–C). Brown, a band at level of line of ocelli and occiput posteriorly to eye darker. Antennal scape, pedicel and flagellomere 1 yellowish-brown, flagellum light brown. Flagellomere 2–4 wider than long, flagellomeres 1 and 7–13 slightly longer than wide, flagellomere 14 more than twice as long as wide. Face brown, clypeus light brown, palpus and labella yellowish-brown. Maxillary palpomere 4 about as long as palpomere 3, palpomere 5 1.7× palpomere 4 length. Occiput with nine strong postocular setae. **Thorax**. Scutum light brown, darker medio-posteriorly, scutellum brown. Antepronotum, proepisternum, anepisternum, mesepimeron, laterotergite, and metepisternum light brown, mediotergite light brown on ventral half, brown on dorsal half, katepisternum whitish with a light brown tinge. Anepisternum covered with small setae on dorsal four-fifth; anterior basalare bare; katepisternum with a group of small, fine setae medially at level of ventral end of anepisternum; laterotergite bare; metepisternum with 15 small setae along its entire length. Haltere stem light brown, knob brown. **Legs**. Coxae whitish with a yellowish tinge, hind coxa with a brownish tinge on basal end, femora light yellowish-brown; tibiae and tarsi light brown, darker towards tip. **Wing** (Fig. 39D). Membrane brownish fumose. Posterior margin slightly emarginated at tip of CuA. C extending for 4/5 of distance to M_1_; R_1_ meeting C before mid of wing. A row of setae along part of the unsclerotized line of M_1_; sclerotized part of M_2_ originating beyond level of tip of R_1_; distal end of M_4_ gently sinuous. CuA not disconnected from M_4_ at base. Anal fold weakly sclerotized. Macrotrichia on membrane of anal lobe and cell cua. **Abdomen**. Tergites 1–6 light brown, darker medially, tergite 7 yellowish-brown; sternites 1–7 light yellowish-brown. **Terminalia** (Figs. 39E–F). Yellowish-brown. Sternite 8 wide at base, trapezoid, distal end with a pair of elongate lobes and a medial incision posteriorly, fine setae entirely covering sclerite, especially on lobes, three longer setae at tip of each lobe. Sternite 9 present as a pair of long sclerotized bands connected distally, diverging towards anterior end, anterior arm weakly sclerotized, distal medial end of sternite 9 almost reaching level of mid of cercomere 1, gonopore connected to two gonoducts. Tergite 8 apparently fused to tergite 9, covered with microtrichia and scattered fine setae. Sternite 10 with one pair of outer long setae and one pair of inner short setae on distal margin. Tergite 10 reduced to an inconspicuous slender band and with two pairs of long setae, at tip of long digitiform projections, one pair at lateral corners and one pair sub-medially [distal end of both cercomere 1 broken, cercomere 2 missing].

**Male**. Unknown.

**Material examined** (25 females): ZRCBDP0278331, Singapore, 03.may.18, MIP leg. (slide-mounted); ZRCBDP0058745, Bukit Timah Forest (BT08), 11.aug. 16, MIP leg.; ZRCBDP0071045, Bukit Timah Forest (BT04), 17.nov. 16, MIP leg.; ZRCBDP0132805, National University of Singapore (PGP), 7.jun. 17, MIP leg.; ZRCBDP0132813, National University of Singapore (PGP), 7.jun. 17, MIP leg.; ZRCBDP0132830, National University of Singapore (PGP), 24.may. 17, MIP leg.; ZRCBDP0132831, National University of Singapore (PGP), 24.may. 17, MIP leg. (extracted; website photo specimen); ZRCBDP0132835, National University of Singapore (PGP), 24.may. 17, MIP leg.; ZRCBDP0132846, National University of Singapore (PGP), 24.may. 17, MIP leg.; ZRCBDP0132848, National University of Singapore (PGP), 24.may. 17, MIP leg.; ZRCBDP0133093, National University of Singapore (PGP), 17.may. 17, MIP leg.; ZRCBDP0133096, National University of Singapore (PGP), 17.may. 17, MIP leg.; ZRCBDP0133107, National University of Singapore (PGP), 17.may. 17, MIP leg.; ZRCBDP0133120, National University of Singapore (PGP), 17.may. 17, MIP leg.; ZRCBDP0133123, National University of Singapore (PGP), 17.may. 17, MIP leg.; ZRCBDP0133145, National University of Singapore (PGP), 14.jun. 17, MIP leg.; ZRCBDP0133146, National University of Singapore (PGP), 14.jun. 17, MIP leg.; ZRCBDP0133148, National University of Singapore (PGP), 14.jun. 17, MIP leg.; ZRCBDP0133150, National University of Singapore (PGP), 14.jun. 17, MIP leg.; ZRCBDP0133154, National University of Singapore (RVR), 03.may. 17, MIP leg.; ZRCBDP0133469, National University of Singapore (PGP), 09.may. 17, MIP leg.; ZRCBDP0133474, National University of Singapore (PGP), 09.may. 17, MIP leg.; ZRCBDP0133482, National University of Singapore (PGP), 09.may. 17, MIP leg.; ZRCBDP0133528, National University of Singapore (PGP), 03.may. 17, MIP leg. (slide-mounted); ZRCBDP0136975, Bukit Timah Forest (BT08), 17.apr. 17, MIP leg.

### Clastobasis Skuse

*Clastobasis* Skuse, 1890: 617. Type-species, *Clastobasis tryoni* Skuse 1890: 619, by monotypy.

**Diagnosis**. Sc complete, reaching C; sc-r weak or absent; R_1_ shorter than r-m; R_5_ often gently sinuous; M_4_ incomplete basally, not connected to CuA, origin at level of tip of Sc. Lateral ocelli touching eye margins, most often antenna dark-ringed.

*Clastobasis* and *Leia* Meigen are both paraphyletic, in their current definition (Oliveira & Amorim 2021) and need revision. There is indeed a core clade of *Clastobasis* species, including, e.g., *C. villiersi* Matile and a number of Afrotropical species assigned to the genus of the *maculicoxa*-group of species (Olavi Kurina, pers. com.). Two of the three species of the genus we found in Singapore belong to this group. All three species from Singapore also share some features of the wing venation and the lateral ocelli touching the eye margins with those species of *Clastobasis*.

There are seven described Oriental species of *Clastobasis*, four are described from India (Brunetti 1912) and three from Sri Lanka (Sivec & Plassmann 1982). Some of Brunetti’s (1912) types of mycetophilids are badly damaged (Väisänen 1996). There are, however, about 20 Oriental species assigned to *Leia* that may actually correspond to species of *Clastobasis* —most of which described from India, Pakistan, Nepal and China (see, e. g., Colless & Liepa 1973). Four species of *Leia* are known from the Malay Peninsula and Indonesia: *L. albicincta* de Meijere, *L. major* Edwards, *L. nigriventris* Edwards, and *L. nigripalpis* Edwards. The original description of all these species has enough information on the color patterns of the thorax and abdomen to conclude that our material of *Clastobasis* does not belong to any of the described species. There is one species described by Colless (1966) from Micronesia in *Leia*, later moved to *Clastobasis* by Matile (1989), but the color pattern also differs from the species from Singapore.

*Clastobasis sritribuana* Amorim & Oliveira, **sp. n.** was collected in the swamp-and rainforest. The other two species of *Clastobasis* had specimens collected in the mangrove, but also in impacted urban forests. The haplotype network (Fig. 40E) for the genus is complex, with a number of conflicts depending on what species delimitation method was used. *Clastobasis sritribuana* Amorim & Oliveira, **sp. n.** and *C. bugis* Amorim & Oliveira, **sp. n.** are well defined, mixed with *Clastobasis oranglaut* Amorim & Oliveira, **sp. n.** in a single cluster using mPTP. There are five small groups of haplotypes that are separated from the main haplotype of *Clastobasis oranglaut* Amorim & Oliveira, **sp. n.** only by PTP.

*Clastobasis sritribuana* Amorim & Oliveira, sp. n. (Figs. 40A–D)

https://singapore.biodiversity.online/species/A-Arth-Hexa-Diptera-000801

urn:lsid:zoobank.org:act:9446F6A2-1A82-4336-BB48-0FAAB44C26E8

**Diagnosis**. Head and most of thorax dark brown, only shoulders whitish-yellow. Flagellomeres 1–8 annelate, yellow on anterior half, brown on distal half. Coxae whitish, no marks, femora and tibiae cream-yellowish. Abdominal tergites 1–4 and 6 brown, tergites 3–5 cream-yellow, sternite 1–2 and 6 brown, sternites 3–5 cream-yellow with slender brownish lateral marks.

***Clastobasis_sritribuana_ZRCBDP0137106_hapZRCBD P0137106_SMH_holotype* [19: A, 28: T, 52: C, 70: G, 11 0: G, 112: A, 139: C, 145: C]**

tttagcttcaacaattgcAcacgcaggTgcttctgttgatttagcaattttCtctttacatttagctggGatttcctcaattttaggagctgtaaattttattactactGtAattaatatacgagctccaggaatttcCtttgaCcgaatacctttatttgtttgatcagttctaattactgctgttttattattactttcattaccagtattagcaggagctattactatacttttaacagatcgaaatttaaatacttctttctttgatccagcaggaggaggagatcctattttatatcaacatttattt

#### Description

**Female** (Fig. 40A). Wing length, 2.76–2.98; width, 0.94–0.99 (n=2). **Head**. Vertex brown, with scattered setae posteriorly to ocelli, an irregular crown of longer setae on occiput around eyes. Mid ocellus present, placed at posterior end of frontal furrow, less than half of diameter of lateral ocelli, lateral ocelli nearly touching eye margin. Frons, face, and clypeus yellow. Frons devoid of setae, frontal furrow complete, reaching mid ocellus; face slender, bare, entirely separated from clypeus; clypeus large, truncate ventrally, entirely covered with setae. Scape twice length of pedicel, yellowish. Scape with a subapical crown of setulae and a distal crown of stronger setae; pedicel with a single distal crown of setulae and larger setae, dorsally a conspicuous seta. Flagellomeres 1– 8 yellowish with a brown distal band, flagellomeres 9–14 light brown. Maxillary palpus yellowish, five palpomeres, basal one weakly sclerotized, bare, palpomere 2 short, with dorsal setae, palpomere 3 well-developed, with a well-defined sensorial pit opening laterally on basal third and scattered setulae on dorsal face, about 4× length of palpomere 2; palpomere 4 elongate, weakly sclerotized, slightly longer than palpomere 2, with scattered setulae dorsally; palpomere 5 more than twice palpomere 4 length, slender, weakly sclerotized, with scattered dorsal setulae. Labella yellow, large, with a pair of pseudotracheae. **Thorax**. Scutum dark brown, scutellum brown. Scutum covered with scattered short setae, some stronger supra-alars and prescutellars; a pair of strong scutellar bristles, a pair of strong setae more externally and 2–4 setulae laterally. Pleural sclerites brown, membrane whitish. Basisternum with wide dorsoposterior arms. Antepronotum wide, with five bristles and additional small setae; proepisternum well-developed, with three bristles and two setulae on ventral half. All remaining pleural sclerites bare except for laterotergite, with about 35 setae and setulae. Mesepisternum wide on ventral half, reaching ventral margin of pleura, mediotergite long, only gently curved. Haltere whitish, setose. **Legs**. Coxae whitish-yellow, distal margin of mid and hind coxae, and mid and hind trochanters with dark brown markings, hind coxa with a dark brown mark on basal fifth. Mid and hind femora, tibiae and tarsi light brownish yellow [both fore legs broken]. Fore coxa with fine setae covering entire frontal and lateral faces, and some few strong bristles on distal margin; mid coxa covered with setae on frontal face and on distal half of lateral face, some few strong bristles on distal margin; hind coxa with setae restricted to basal fifth of frontal face, some few strong bristles on distal margin. Mid and hind tibiae and tarsi with irregularly distributed trichia; tibia with two irregular rows of short bristles dorsolaterally, hind tibia with a pair of irregular dorsolateral rows and a dorsal row of short bristles. Mid and hind tarsomeres 1 and 2 with some few short bristles. Tibial spurs yellow, outer spurs about 5× tibial diameter at apex, outer spur over 1.5× longer than inner spur. Tarsal claw with a small proximal tooth and a large, blunt medial tooth. **Wing** (Fig. 40B). Membrane without macrotrichia, translucent, with a light brownish-yellow tinge. C ending at apex of R_5_. Sc complete, ending at C in wing basal fourth, weakly sclerotized distally; sc-r not sclerotized. R_1_ short, 0.88× r-m length, reaching C at mesal third of wing; first sector of Rs nearly transverse, short; R_5_ short, reaching C before level of tip of M_2_, straight except very distally; r-m curved on basal third, more than 6× length of first sector of Rs. M_1+2_ shorter than length of r-m; M_1_ and M_2_ more than 5× length of M_1+2_; M_1_ gently curved posteriorly on distal half, M_2_ curved anteriorly on distal half, fading on distal fourth, line of setae reaching margin. M_4_ gradually curved posteriorly, interrupted at its very base, not connecting to CuA. CuA gently sinuous midway to apex. Veins bR, R_1_, R_1_ and R_5_ with ventral and dorsal setae, M_1+2_, M_1_, M_2_, M_4_ and CuA with only dorsal setulae; Sc, first sector of Rs and CuP bare. **Abdomen**. Tergites 1–4 brown, tergite 5 yellowish laterally, brown medially, tergites 6–7 brown; sternites 1–2 brownish, sternite 3–5 cream-yellow with brown lateral margins (extension of brown area varying between individuals), sternites 6–7 brown. Sternite 1 strongly modified, U-shaped, displaced posteriorly, fit into a large incision on anterior half of sternite 2. **Terminalia** (Figs. 40C–D). Terminalia brownish. Sternite 8 wide, with a pair of overlapped lobes at each side, widely covered with microtrichia, setae restricted to lobes and posterior margin. Sternite 9 complex, with two pairs of weakly sclerotized elongated lobes, curved sclerotized bands beneath posterior lobe, medial anterior apodeme of vaginal furca not visible. Tergite 8 large, with a pair of projections on each side of posterior margin, mostly covered only by microtrichia, setation restricted to lateroposterior lobes. No evidence of tergites 9 and 10, possibly fused indistinctly to tergite 8. Cerci very modified into a pair of lobes connected to each other medially on anterior end, entirely devoid of microtrichia, short, strong spine-like setae at apex of each lobe. A pair of long apodemes extending anteriorly from each side of anus may correspond to a modified sternum 10.

**Male**. Unknown.

#### Material examined

**Holotype**: female, ZRCBDP0137106, Bukit Timah Forest (BT07), old secondary forest, 15.nov. 16, MIP leg. (slide-mounted). **Paratype** (female): ZRCBDP0047849, Nee Soon (NS1), swamp forest, 26.jun.13, MIP leg. (website photo specimen, slide-mounted). **Additional sequenced specimen**: ZRCBDP0138569, Bukit Timah Forest (BT06), maturing secondary forest, 15.nov. 16, MIP leg.

**Etymology**. The species epithet of this species refers to Sri Tri Buana, the official title of the Srivijayan (a Buddhist thalassocratic empire in Southeast Asia during the 7th to 12th century) prince Sang Nila Utama. He is the founder of the Kingdom of Singapura (Kerajaan Singapura; est. 1299). Sang Nila Utama is said to have landed on Temasek (the island of Singapore) on a hunting trip, seeing an animal supposed to be a lion. Taken as an auspicious sign, he founded the Kingdom based on the settlement hence called Singapura, in one of the standing interpretations considered to come from the Sanskrit “Lion City”. The noun is used in apposition.

**Remarks**. This species clearly differs in the color pattern from the other two species of *Clastobasis* described here. Most species of *Clastobasis* fit in the yellowish color pattern seen in *C. bugis* Amorim & Oliveira, **sp. n.** and *C. oranglaut* Amorim & Oliveira, **sp. n.,** but there are some species of *Clastobasis* in Japan that share this general brown color pattern of *C. sritribuana* Amorim & Oliveira, **sp. n.** (Kurina, 2020). We have two haplotypes for this species and no delimitation conflicts.

*Clastobasis bugis* Amorim & Oliveira, sp. n. (Figs. 41A–F)

https://singapore.biodiversity.online/species/A-Arth-Hexa-Diptera-000808

urn:lsid:zoobank.org:act:C145CE97-C965-4D7D-8B1E-9774B38C1B40

**Diagnosis**. Thorax mostly ochreous-yellowish, some parts of pleura with a light greyish-brown tinge. Abdomen cream-yellow, tergites 1–5 with brownish triangular wide marks on posterior half, sternites 3–5 with slender diagonal greyish-brown marks. Male terminalia with a pair of strongly sclerotized structures in aedeagus, long rounded setose projections of gonocoxites with a short ventral digitiform projection bearing spines at apex, gonostylus wider at base, with a long distal flattened projection, slightly widening towards apex.

***Clastobasis_bugis_ZRCBDP0048253_hapZRCBDP0048 245_SMH_holotype* [103: C, 109: C, 110: G, 112: A, 124 : C, 184: C, 208: T]**

tttagcatctactattgctcatgcgggggcttcagttgatttagctattttttcattacatttagcaggtatttcttcaattttaggagctgttaattttatCactacCGtAattaatatacgCgccccaggaatttcttttgatcgaatacctttatttgtttgatcagtattaattactgcCgttttacttttactatcacttccTgtattagcaggagctattactatattattaacagatcgaaatttaaatacatcattttttgacccagcaggaggtggagatccaatcttatatcaacatttattt

#### Description

**Male**. Wing length, 2.14; width, 0.82. **Head**. Vertex light ochre-yellowish, except for blackish-brown area around ocelli, occiput dark ochre-yellowish. Frons, face, and clypeus light ochre-yellowish. Scape and pedicel yellowish, flagellomeres with yellowish basal half, brown on distal half. Maxillary palpus whitish-yellow, darker on basal three palpomeres. Labella yellow. **Thorax**. Scutum and scutellum ochre-yellowish, lighter laterally. Anepisternum, katepisternum, and mediotergite light ochre-yellowish with some areas with a brownish tinge; mesepimeron ochre-yellowish on ventral half, light brownish on dorsal half; laterotergite ochre-yellowish with brownish ventral border. Laterotergite with about 18 setae of different sizes, other pleural sclerites bare except for antepronotum and proepisternum. **Legs**. Coxae and femora whitish-yellow, coxae with brown marks at tip, hind femur light ochre-brown on distal half and a brown mark distally; trochanters with a brown mark; tibiae and tarsi with light brown tinge. **Wing** (Fig. 41B). Sc complete, ending at C on basal fourth of wing. R_1_ short, 0.78× r-m length; R_5_ reaching C more basally than level of tip of M_1_, straight except distally, gently curved. M_2_ not reaching wing margin. Veins bR, R_1_ and second sector of Rs with dorsal and ventral setae; M_1_, M_2_, M_4_, second sector of CuA and CuP with setulae dorsally, distal fifth of M_2_ with no setulae. **Abdomen**. Tergites 1–6 cream-yellow with a greyish-brown posterior triangle pointed anteriorly, tergite 7 cream-yellow; sternites 1–2 cream-yellow, sternite 3–5 yellow with a pair of slender diagonal brownish bands, light brown medio-posteriorly; sternite 6 yellowish with a faint brownish diagonal band; sternite 7 yellow with brown anterolateral corners. **Terminalia** (Fig. 41C). Terminalia light yellowish-brown. Gonocoxites well developed, largely fused to each other ventrally, a pair of posterior incisions on posterior border isolating a medial lobe that does not project beyond level of base of gonostylus, a pair of spines at posterior margin externally to incisions, dorsally a pair of flat, oval large projections densely covered with strong, spine-like curved setae on ventral face and a sub-basal digitiform projection with a group of distal spines; an additional short digitiform lobe on gonocoxite dorsal to insertion of gonostylus. Gonostylus well sclerotized, simple, basal third wider, entirely bare of setae. Aedeagal sclerite complex, with a pair of hardly sclerotized small sclerites close to each other. Tergite 9 apparently fused indistinctly to gonocoxites. Cerci small, inconspicuous.

**Female** (Fig. 41A). Wing length, 2.73; width, 0.94. **Terminalia** (Figs. 41D–F). Sternite 8 large, with a pair of short lobes with a short medial posterior incision, most of sternite 8 with only microtrichia, setation restricted to posterior margin. Sternite 9 present as a wide transversal plate with gonoducts reaching independently, two short groups of setae, anterior medial apodeme short, extending anteriorly. Tergite 8 wide, rectangular, no medio-posterior incision. Tergite 9+10 with a pair of short lateral lobes with a slender medial connection, entirely bare of setae. Cerci not cylindrical, flat and elongate, fused to each other medially, covered with microtrichia and fine setae.

#### Material examined

**Holotype**: male, ZRCBDP0048253, Pulau Semakau (SMN2), planted mangrove, 17.oct.13, MIP leg. (slide-mounted). **Paratypes** (3 females): ZRCBDP0048242, Pulau Semakau (SMO2), old mangrove, 24.oct.13, MIP leg. (extracted, slide-mounted); ZRCBDP0048245, Pulau Semakau (SMO2), old mangrove, 24.oct.13, MIP leg. (website photo specimen); ZRCBDP0048306, Pulau Semakau (SMN1), planted mangrove, 09.may.13, MIP leg. **Additional sequenced specimens**: ZRCBDP0278014, National University of Singapore (SDE), urban forest, 19.oct. 17, MIP leg.; ZRCBDP0278015, National University of Singapore (SDE), urban forest, 19.oct. 17, MIP leg.; ZRCBDP0278021, National University of Singapore (SDE), urban forest, 19.oct. 17, MIP leg.; ZRCBDP0278024, National University of Singapore (SDE), urban forest, 19.oct. 17, MIP leg.; ZRCBDP0278026, National University of Singapore (SDE), urban forest, 19.oct. 17, MIP leg.; ZRCBDP0278030, National University of Singapore (SDE), urban forest, 19.oct. 17, MIP leg.; ZRCBDP0278020, National University of Singapore (SDE), urban forest, 19.oct. 17.

**Etymology**. The species epithet refers to the Bugis, an ethnic group from South Sulawesi (Indonesia) and one of the first groups of people to arrive in Singapore when the British established the colonial port-city of Singapore. The Bugis were leading maritime traders in the region at that point in time and made significant contributions to Singapore in its development as a trading port. The noun is used in apposition.

**Remarks**. *Clastobasis bugis* Amorim & Oliveira, **sp. n.** and *C. oranglaut* Amorim & Oliveira, **sp. n.** belong to a group in the genus clearly apart from *C. sritribuana* Amorim & Oliveira**, sp. n.,** as can be seen by the body color patterns. A species of *Clastobasis* collected in Java in citrus plantation (Ulimah et al. 2021) has a color pattern that in some extension agrees with *C. bugis* Amorim & Oliveira, **sp. n.,** especially the oblique brown marks on sternites 2–4. We have three haplotypes for this species and no delimitation conflicts.

*Clastobasis oranglaut* Amorim & Oliveira, sp. n. (Figs. 42A–F, 43A–B)

https://singapore.biodiversity.online/species/A-Arth-Hexa-Diptera-000829,and-002108

urn:lsid:zoobank.org:act:1E5623E9-B833-49CB-B74E-F2DD03214BCF

**Diagnosis**. Most of thorax ochreous-yellow, some parts of the pleura with a light greyish-brown tinge. Abdomen cream-yellow, more yellowish towards apex of abdomen, tergites 1–5 with brownish transverse marks along posterior half, light brown lateral marks on sternites 3–6. Male terminalia with a single strongly sclerotized aedeagus structure, a pair of long rounded setose projections of gonocoxites, gonostylus slender, simple.

***Clastobasis_oranglaut_ZRCBDP0047070_hapZRCBDP 0047070_SMH_holotype* [34: A, 46: A, 110: G, 112: A, 1 37: A, 139: A, 209: G, 217: T, 220: T, 241: A, 292: A]**

tttagcatctacaattgctcatgcaggagcttcAgttgatttagcAattttttctttacatttagcaggaatttcttcaattttaggagctgtaaattttattactactGtAattaatatacgagccccaggaattAcAtttgatcgaatacctttatttgtttgatcagttttaattactgctgttttacttttattatcattaccaGtattagcTggTgctattactatattattaacAgatcgaaatttaaatacttctttttttgatccagcaggaggaggagatccAattttataccaacatttattt

#### Description

**Male** (Fig. 42A). Wing length, 2.50; width, 0.87. **Head** (Fig. 42B). Vertex light yellowish-brown, except for dark brown area over ocellar line. Mid ocellus present, small. Occiput light brown. Frons, face, and clypeus light yellowish-brown. Maxillary palpus first two palpomeres yellowish-brown, last two palpomeres whitish-yellow. Scape and pedicel light yellowish-brown, flagellomeres with a yellowish band on basal half and a brownish band on distal half. **Thorax** (Fig. 42C). Scutum and scutellum ochre-yellowish, some dark brown sclerites on wing articulation. Antepronotum and proepisternum ochre-yellowish, with a light brown mark at ventral end of proepisternum; anepisternum, katepisternum, mediotergite, and mesepimerom ochre-yellowish with areas with greyish-brown tinge; laterotergite ochre-yellowish with brownish ventral border. Two strong bristles and additional small setae aligned on scutellum; 2– 3 larger setae and almost 30 small setae on laterotergite. Pleural membrane yellow. **Legs**. Legs whitish-yellow, with dark brown marks at tip; trochanters with a brown spot distally; femora light brownish-yellow with a dark brown mark at distal end; tibiae and tarsi light yellowish-brown. Fore tibia slightly shorter than fore femur, mid and hind tibiae slightly longer than femora. Fore tibia with a wide anteroapical depressed area lined with setulae. **Wing** (Fig. 42D). Sc weakly sclerotized, complete, ending at C on basal fourth of wing. R_1_ short, 0.87× r-m length; R_5_ reaching C more basally than level of tip of M_1_, straight except distally, gently curved. M_2_ not reaching wing margin. Veins bR, R_1_, and second sector of Rs with dorsal and ventral setae; M_1_, M_2_, M_4_, second sector of CuA, and CuP with setulae dorsally, distal fifth of M_2_ with no setulae. **Abdomen**. Tergites 1–5 ochre-yellow with a triangular posterior light brown mark, tergites 6–7 brownish-yellow; sternites 1–2 yellow; sternites 3–4 ochre-yellow with brownish bands laterally; sternites 5–6 yellow medially and brown at lateral margins; sternite 7 ochre-yellow with dark brown on lateral margins. **Terminalia** (Figs. 42E–F). Terminalia yellowish, with a dark brown distal projection of gonocoxites and dark brown sclerites internally. Gonocoxites complex, fused medially along anterior third, a medial large conical distal projection bearing ventrally 4 long fine setae and a number of strong short setae directed outwards distally; a pair of short lateroposterior projections extending beyond level of base of gonostylus; a short lobe internally to base of gonostylus, bearing a group of short black setae; a short, round bare lobe on posterior margin on dorsal face of terminalia; and a pair of large, well-sclerotized rounded lobes extending beyond tip of gonostylus covered with regular setae on dorsal face and with dense, darker and stronger setae on ventral face on distal two-thirds, part of setae sinuous. Gonostylus simple, well-sclerotized, digitiform, slightly curved at tip, entirely bare. Gonocoxal bridge with long apodemes, extending anteriorly more or less close to each other. Aedeagal-parameral complex with two main components: ejaculatory apodeme at anterior end and a strongly sclerotized median structure bearing a pair of pointed lobes anteriorly, with some other smaller, well-sclerotized pieces; second element capsular, placed more dorsally, with rounded anterior end, a pair of sclerotized slender arms laterally and with a thin medial bifid projection at distal end. Area corresponding to tergite 9 medially with a medial incision on posterior margin. Cerci small, well-defined, placed between bases of large gonocoxite rounded dorsal lobes.

**Female**. As male, except for the following. **Wing l**ength, 2.86; width, 1.02. **Abdomen**. Tergites 1–6 yellowish with brown band on posterior margin, more rectangular on anterior segments and more triangular on posterior segments, tergite 7 yellowish; sternites 1–2 whitish-yellow, sternites 3–6 yellowish with a pair of oblique brown bands, sternite 7 yellowish with brown marks at lateroanterior corners. Fine setae on all tergites and on sternites 1–6, sternite 7 with slightly stronger setae on posterior half. **Terminalia** (Figs. C42A–B). Sternite 8 large, with two pairs of short lobes along posterior margin, a short medial posterior incision, most of tergite only with microtrichia, setation restricted to margins of posterior lobes. Sternite 9 present as a wide transversal plate, a pair of lateroposterior setose lobes, gonoduct sclerotized medially, sided by a pair of apodemes directed anteriorly. Tergite 8 wide, rectangular, no medio-posterior incision, long setae restricted to posterior margin. Tergite 9 with a pair of large lateral lobes separated by deep medial incision. Sternite 10 wide, weakly sclerotized but well-defined, with a pair of posterior lobes separated by a medial incision, covered with fine setae. Cerci apparently fused to tergite 10 to form a distal pair of large lobes covered with microtrichia and fine setae.

#### Material examined

**Holotype**: male, ZRCBDP0047070, National University of Singapore (Uhall), 22.jul.15, MIP leg. (slide-mounted). **Paratypes** (5 males, 14 females). **Males**: ZRCBDP0047071, National University of Singapore (Uhall), 22.jul.15, MIP leg.; ZRCBDP0048790, National University of Singapore (Utown), 08.apr.15, MIP leg.; ZRCBDP0049283, National University of Singapore (Utown), 06.may.15, MIP leg.; ZRCBDP0049312, National University of Singapore (Utown), 10.jun.15, MIP leg. (website photo); ZRCBDP0066825, Bukit Timah, primary forest (BT05), 28.Sep.16, MIP leg. **Females**: ZRCBDP0047065, National University of Singapore (PGP), 01.jul.15, MIP leg.; ZRCBDP0048250, Sungei Buloh (SB1), mangrove, 18.set.13, MIP leg.; ZRCBDP0048255, Sungei Buloh (SB1), mangrove, 09.oct.13, MIP leg.; ZRCBDP0048256, Sungei Buloh (SB1), mangrove, 09.oct.13, MIP leg.; ZRCBDP0048769, National University of Singapore (PGP), 20.may.15, MIP leg.; ZRCBDP0049074, National University of Singapore (PGP), 22.apr.15, MIP leg.; ZRCBDP0049138, National University of Singapore (Utown), 20.may.15, MIP leg.; ZRCBDP0049139, National University of Singapore (Utown), 20.may.15, MIP leg.; ZRCBDP0049140, National University of Singapore (Icube), 03.jun.15, MIP leg.; ZRCBDP0049303, National University of Singapore (Utown), 29.apr.15, MIP leg.; ZRCBDP0049304, National University of Singapore (Utown), 29.apr.15, MIP leg.; ZRCBDP0049311, National University of Singapore (Utown), 10.jun.15, MIP leg.; ZRCBDP0049319, National University of Singapore (Utown), 27.may.15, MIP leg.; ZRCBDP0049336, National University of Singapore (PGP), 08.apr.15, MIP leg. (extracted, slide-mounted). **Additional sequenced specimens**: ZRCBDP0278233, Pulau Ubin (PU18), mangrove, 31.may.18, MIP leg.; ZRCBDP0278261, Singapore, 07.jun.18, MIP leg.; ZRCBDP0278301, Singapore, 10.may.18, MIP leg.; ZRCBDP0278305, Singapore, 10.may.18, MIP leg.; ZRCBDP0278307, Singapore, 10.may.18, MIP leg.; ZRCBDP0278312, Singapore, 10.may.18, MIP leg.; ZRCBDP0278313, Singapore, 10.may.18, MIP leg.; ZRCBDP0278327, Singapore, 03.may.18, MIP leg.; ZRCBDP0278346, Singapore, 26.apr.18, MIP leg.; ZRCBDP0278473, Pulau Ubin (PU17), mangrove, 17.may.18, MIP leg.; ZRCBDP0279148, Singapore, 16.may.18, MIP leg.; ZRCBDP0279149, Singapore, 16.may.18, MIP leg.; ZRCBDP0279164, Singapore, 26.apr.18, MIP leg.; ZRCBDP0279175, Singapore, 31.may.18, MIP leg.; ZRCBDP0279185, Singapore, 07.jun.18, MIP leg.; ZRCBDP0279192, Singapore, 07.jun.18, MIP leg.; ZRCBDP0284181, Kranji Marshes (KM03), freshwater swamp, date range 2018-2019,MIP leg.; ZRCBDP0284188, Kranji Marshes (KM03), freshwater swamp, date range 2018-2019,MIP leg.; ZRCBDP0284203, Pulau Ubin (PU20), mangrove, 2018, MIP leg.; ZRCBDP0284207, Pulau Ubin (PU20), mangrove, 2018, MIP leg.; ZRCBDP0284257, Singapore, 03.may.18, MIP leg.; ZRCBDP0284262, Singapore, 03.may.18, MIP leg.; leg.; ZRCBDP0284262, Singapore, 03.may.18, MIP leg.; ZRCBDP0284265, Singapore, 03.may.18, MIP leg.; ZRCBDP0284266, Singapore, 03.may.18, MIP leg.; ZRCBDP0284269, Singapore, 02.may.18, MIP leg.; ZRCBDP0284270, Singapore, 02.may.18, MIP leg.; ZRCBDP0284271, Singapore, 02.may.18, MIP leg.; ZRCBDP0284272, Singapore, 02.may.18, MIP leg.; ZRCBDP0284274, Singapore, 03.may.18, MIP leg.; ZRCBDP0284275, Singapore, 03.may.18, MIP leg.; ZRCBDP0284276, Singapore, 03.may.18, MIP leg.; ZRCBDP0284277, Singapore, 03.may.18, MIP leg.; ZRCBDP0284278, Singapore, 03.may.18, MIP leg.; ZRCBDP0284280, Singapore, 26.apr.18, MIP leg.; ZRCBDP0284282, Singapore, 26.apr.18, MIP leg.; ZRCBDP0284283, Singapore, 26.apr.18, MIP leg.; ZRCBDP0314192, Pulau Ubin (PU11), mangrove, 31.may.18, MIP leg.; ZRCBDP0326310, Pulau Ubin (PU14), coastal forest, 03.may.18, MIP leg.; ZRCBDP0326313, Pulau Ubin (PU14), coastal forest, 03.may.18, MIP leg.; ZRCBDP0326314, Pulau Ubin (PU14), coastal forest, 03.may.18, MIP leg.; ZRCBDP0326315, Pulau Ubin (PU14), coastal forest, 03.may.18, MIP leg.; ZRCBDP0326316, Pulau Ubin (PU14), coastal forest, 03.may.18, MIP leg.; ZRCBDP0326317, Pulau Ubin (PU14), coastal forest, 03.may.18, MIP leg.; ZRCBDP0326318, Pulau Ubin (PU14), coastal forest, 03.may.18, MIP leg.; ZRCBDP0326319, Pulau Ubin (PU14), coastal forest, 03.may.18, MIP leg.; ZRCBDP0278306, Singapore, 10.may.18; ZRCBDP0278308, Singapore, 10.may.18; ZRCBDP0278471, Singapore; ZRCBDP0284267, Singapore, 03.may.18; ZRCBDP0326311, Pulau Ubin (PU14), coastal forest, 03.may.18; ZRCBDP0326312, Pulau Ubin (PU14), coastal forest, 03.may.18; ZRC_MIS0000001, L01-MT, urban forest, 18.apr.19; ZRC_MIS0000002, L01-MT, urban forest, 18.apr.19; ZRC_MIS0000003, L01-MT, urban forest, 18.apr.19; ZRC_MIS0000004, L01-MT, urban forest, 18.apr.19; ZRC_MIS0000005, L01-MT, urban forest, 18.apr.19; ZRC_MIS0000006, L01-MT, urban forest, 18.apr.19; ZRC_MIS0000007, L01-MT, urban forest, 18.apr.19; ZRC_MIS0000008, L01-MT, urban forest, 18.apr.19; ZRC_MIS0000009, L01-MT, urban forest, 18.apr.19; ZRC_MIS0000010, L01-MT, urban forest, 18.apr.19; ZRC_MIS0000011, L01-MT, urban forest, 18.apr.19; ZRC_MIS0000012, L01-MT, urban forest, 18.apr.19; ZRC_MIS0000013, L01-MT, urban forest, 18.apr.19; ZRC_MIS0000014, L01-MT, urban forest, 18.apr.19; ZRC_MIS0000015, L01-MT, urban forest, 18.apr.19; ZRC_MIS0000016, L01-MT, urban forest, 18.apr.19; ZRC_MIS0000017, L01-MT, urban forest, 18.apr.19; ZRC_MIS0000018, L01-MT, urban forest, 18.apr.19; ZRC_MIS0000019, L01-MT, urban forest, 18.apr.19; ZRC_MIS0000020, L01-MT, urban forest, 18.apr.19; ZRC_MIS0000021, L01-MT, urban forest, 18.apr.19;ZRC_MIS0000022, L01-MT, urban forest, 18.apr.19; ZRC_MIS0000023, L01-MT, urban forest, 18.apr.19; ZRC_MIS0000025, L01-MT, urban forest, 18.apr.19; ZRC_MIS0000026, L01-MT, urban forest, 18.apr.19; ZRC_MIS0000027, L01-MT, urban forest, 18.apr.19; ZRC_MIS0000028, L01-MT, urban forest, 18.apr.19; ZRC_MIS0000029, L01-MT, urban forest, 18.apr.19; ZRC_MIS0000030, L01-MT, urban forest, 18.apr.19; **Sequence failure specimens**: ZRCBDP0040844; ZRCBDP0040847; ZRCBDP0067281.

**Etymology**. The species epithet of this species refers to the Orang [=people] Laut [=sea], a name that applies to several seafaring ethnic groups and tribes living around Singapore, Peninsular Malaysia and the Riau Islands. The name is used in apposition.

**Remarks**. This is the most abundant species of *Clastobasis*, collected mostly on urban areas in Singapore. We have eleven haplotypes in our material and no delimitation conflicts.

### Gnoristinae

The Gnoristinae constitute a complex problem of mycetophilid systematics. The subfamily includes over 340 species in 28 genera worldwide. There is evidence that, in its present composition, the subfamily is not monophyletic (Søli 1996; Rindal et al. 2009; Kasprak et al. 2019; Oliveira & Amorim 2021). The subfamily is basically defined by plesiomorphic features as the long R_1_, tibiae and tarsi trichia not organized in regular rows, microtrichia on the wings not arranged in regular rows, absence of macrotrichia on the wing membrane etc.

The Singapore fauna belong to only two genera, *Chalastonepsia* and *Metanepsia*, each with a single species. Two additional genera, *Pectinepsia* Ševčík & Hippa and *Dziedzickia*, are known from other tropical areas of Southeast Asia and may thus eventually be found in Singapore. Some additional gnoristine genera of Palearctic affinities are known to occur along the northern range of the Oriental region (e. g., *Aglaomyia*, *Boletina*, *Coelosia*, *Deimyia* Kallweit, *Hemisphaeronotus* Saigusa etc.) and are thus less likely to reach Singapore.

Besides the problem of the monophyly of Gnoristinae, there is an additional issue with the “tribe” Metanepsini, as discussed by Kallweit (1998). The Metanepsini were originally proposed by Matile (1971a) for *Metanepsia* Edwards as a tribe in the paraphyletic sciophiline grade. Later Väisänen (1986) proposed a subfamily rank for Metanepsini, which was followed by Ševčík & Hippa (2010). There is no question that *Metanepsia* and related genera compose a clade. The question is, as already posed by Søli (1996), whether the subfamily (or tribal) status for the Metanepsinae renders any other group paraphyletic. Note that a number of Neotropical species of *Dziedzickia*—e. g., *D. intermedia* Lane and *D. laticornis* (Enderlein)—have reduced mouthparts and other features shared with *Metanepsia* and its two related genera, *Chalastonepsia* and *Pectinepsia*. These Neotropical species of *Dziedzickia*, however, do not share the autapomorphies of any of these genera. This means that the entire “Metanepsini” may be a subclade of *Dziedzickia* in the current delimitation of the genus. The Neotropical genus *Schnusea* Edwards differs from *Dziedzickia* basically by r-m reaching M_1_, instead of reaching M_1+2_ and is also a subclade of *Dziedzickia* (Oliveira & Amorim 2014). Even *Hadroneura* Lundstrom, *Palaeodocosia* Meunier and *Syntemna* Winnertz may be subclades of *Dziedzickia*. In Ševčik et al. (2013), *Schnusea*, one species of *Metanepsia*, one species of *Dziedzickia, Syntemna*, *Palaeodocosia* and *Chalastonepsia* compose a clade within the Gnoristinae, but a second species of *Metanepsia* is sister to *Paratinia* and a second species of *Dziedzickia* sister to (*Ectrepesthoneura* + *Metanepsia* + *Paratinia*).

As discussed by Kallweit (1998) and by Ševčík et al. (2011), a thorough review of the “Gnoristinae” (or at least of *Dziedzickia*) is necessary before a proper solution can be found. It may be the case that the name Metanepsinae or Metanepsini will come to be applied to a clade including *Metanepsia*, *Chalastonepsia*, and *Pectinepsia* and part of the genus *Dziedzickia*. We presently do not have a phylogeny for the subfamily and we keep both, *Metanepsia* and *Chalastonepsia*, in the gnoristine grade without using tribal ranks.

### Chalastonepsia Søli

***Chalastonepsia*** Søli 1996: 79. Type species. *Chalastonepsia orientalis* Søli (orig. des.).

**Diagnosis**. Palpus 1-segmented. Male flagellum bead-like, each flagellomere bulbous with a long stalk-like bare apical portion, basal part with numerous long setae. Sc ending in R_1_ via sc-r at tip; r-m reaching Rs well apart from R_1_; medial fork longer than M_1+2_; R_4_ absent; M_4_ originating slightly more basal than level of origin of M_1+2_; first sector of CuA at most slightly shorter than second sector.

There are no keys including all genera of gnoristines. In the Afrotropical manual of Diptera (Søli 2017), the key does not include *Chalastonepsia* and our species of this genus runs into couplet 26, with part of the features of *Metanepsia* (e. g., reduced mouthparts) and part of the features of *Dziedzickia* (e. g., long r-m). All four species of the *Clalastonepsia* described so far included only on males—which have either a strongly pectinate flagellum or a flagellum with a bulbous base with a long distal neck. We have only females of the species found in Singapore. They have laterally compressed flagellomeres, but no pectination or flagellomeres with a distal neck. The type-species of *Chalastonepsia*—*C. orientalis* Søli— is known from Pahang, Malaysia, while *C. nigricoxa* Ševčík & Papp is known from Thailand, *C. montana* Ševčik & Papp from the type-locality in Sumatra and Pahang, Malaysia, and *C. hokkaidensis* Kallweit from Hokkaido (Japan).

A barcode comparison of the sequence of the Singapore species of *Chalastonepsia* on GenBank has a 100% hit against “*Chalastonepsia* cf. *hokkaidensis*” (https://www.gbif.org/species/10652424) and is consistent with the photo of a damaged female collected in Kuala Lumpur, Malaysia. The species is also known from Sumatra in the Bold system. Our specimens agree with the description of the female in Ševčík & Hippa (2010). Conspecificity between Singapore and Malay Peninsula mycetophilid populations is seen, as mentioned above, in *Eumanota racola*. Malay Peninsula populations conspecific with a Hokkaido population would be acceptable for introduced species—as is the case of some species of *Leia*, *Clastobasis* and *Allactoneura*. Finding males of the Singapore populations would help confirming its conspecificity with *C. hokkaidensis*.

*Chalastonepsia hokkaidensis* Kallweit (Figs. 44A–D, 45A–C)

https://singapore.biodiversity.online/species/A-Arth-Hexa-Diptera-002101

***Chalastonepsia hokkaidensis_sp_ZRCBDP0143086_ unnamed_type*[11: G, 12: G, 18: T, 125: C, 132: T, 171: C, 129: T]**

actttcatcaGGaatatTccatacaagatcatctgtagatctatcaattttttctcttcatctagctggaatttcatctattcttggtgcaattaatttcattactacaattataaatataaaaCcaaTaaTtataacatttgatcaaattccactatttgtttgatcaaCtttaattacagctattttactacttttatctttacctgtattagcaggagctattactatattattaacagatcgaaatttaaatacttccttttttgatcctgcaggaggaggggatcccattttatatcaacatttattc

#### Description

**Female** (Fig. 44A). Wing length, 2.32; width, 1.02. **Head** (Fig. 45A). Vertex and frons brown, occiput brown dorsally, lighter laterally towards ventral margin, with few scattered setae anteriorly. Face and clypeus greyish-brown. Mid ocellus present, lateral ocelli distant from eye margin, lateral ocelli large. Frontal furrow present. Frons bare, face with long setae, clypeus bare. Inter-ommatidial setulae on part of eye. Antenna ochre-yellowish, greyish towards apex of flagellum. Scape and pedicel of similar length, about as long as wide; pedicel flat on distal face, extended dorsally and with a group of elongate setae at dorsal tip. Flagellomeres in some degree laterally compressed, slightly projected ventrally, densely covered with setae, distal six flagellomeres less modified. Mouthparts reduced, maxillary palpus greyish-brown, 1-segmented, with setae and sensorial setulae; labella whitish, reduced. **Thorax** (Fig. 45B). Scutum and scutellum dark brown, strongly arched. Scutum with scattered short setae, about three rows of slightly stronger dorsocentrals at each side. Scutellum with an irregular row of setae along distal margin, four stronger scutellar bristles. Pleural sclerites brown, antepronotum lighter towards anterior end, metepisternum mostly yellowish, brownish-yellow on dorsoposterior corner. Pleural membrane whitish-yellow. Antepronotum with three longer and additional smaller setae; proepisternum with 6-7 regular, thin setae; proepimeron wide. Anepisternum, katepisternum, mesepimeron, metepisternum, and mediotergite bare; laterotergite slightly bulging, with 9–11 thin setae medially. Mesepimeron reaching ventral margin of thoracic pleura. Mediotergite with a gentle fold medially. Haltere with pedicel whitish-yellow, knob mostly whitish-yellow, dark brown on proximal end, thin elongate setulae present. **Legs**. Coxae mostly cream-yellowish, brownish on distal end, hind coxa darker. Femora cream-yellowish with darker tinge on distal half, tibiae and tarsi greyish. Tibiae and tarsi covered with by irregularly distributed short setae, some few longer dark setae on hind tibiae. Tibial spurs cream-yellowish, spurs short, slightly more than twice tibial diameter at apex. Tarsal claws with three small teeth on basal half. **Wing** (Fig. 44B). Membrane without macrotrichia, very light brown fumose. C ending beyond apex of R_5_, at about a third of distance between R_5_ and M_1_; Sc fused to bR more basally than origin of M_4_. R_1_ almost straight, reaching C at distal third of wing, more than 4× longer than r-m; first sector of Rs oblique, slightly over half of r-m length; R_5_ reaching C before wing tip, slightly arched posteriorly close to margin; r-m oblique, distal end directed towards wing base. Medial fork about 2.9× M_1+2_ length; M_1_ and M_2_ reaching wing margin. Base of M_4_ arched, originating at level of distal end of r-m, clearly more basally than origin of medial fork. CuA gently curved towards posterior margin. Sc, r-m, first sector of Rs, M_1+2_, M_1_, M_2_, M_4_, CuA and CuP devoid of macrotrichia; bR, R_1_ and R_5_ with dorsal setae; false cubital vein sclerotized; CuP present, produced to slightly beyond origin of M_4_. **Abdomen**. Tergite and sternite 1 cream-yellowish, tergite and sternites 2-7 greyish-brown, tergites and sternites covered with scattered brown setae. **Terminalia** (Figs. 44C–D). Brown. Sternite 8 with a sclerotized band along anterior margin, a weakly sclerotized area medially, and a pair of large, well-separated setose projections; sternite 9 (vaginal furca) more sclerotized medially on a tubular structure around base of spermathecal ducts; sternite 10 present as a thin blade with some few setulae along distal lateral margins. Tergite 8 wide, short; tergite 9 present as a slender band; tergite 10 divided into a pair of sclerites basal to cerci. Cercus with a short second cercomere, largely fused to first cercomere.

**Material examined** (3 females): ZRCBDP0048536, Nee Soon (NS2), swamp forest, 16.jan.13, MIP leg. (website photo specimen, slide-mounted); ZRCBDP0047804, Nee Soon (NS1), swamp forest, 01.may.13, MIP leg. (slide-mounted) (MZUSP); ZRCBDP0143086, Nee Soon (MSM1a), swamp forest, 21.jan. 2015, MIP leg. (website photo specimen).

**Remarks**. The barcodes of the *Chalastonepsia* species from Singapore have a 100% Genbank match to a *Chalastonepsia hokkaidensis* specimen from Thailand (Sequence ID: MH114427.1) and four additional 100% hits in BOLD for specimens identified as “*Chalastonepsia* cf. *hokkaidensis*” (BOLD:ACP9308) (Sumatra and Malaysia). In addition, the image of one damaged female in belonging to BOLD:ACP9308 is consistent with the morphology of the Singapore specimens. Three females of this species have been collected, all in the Nee Soon swamp forest, with a single haplotype.

### Metanepsia Edwards

*Metanepsia* Edwards 1927: 361. Type-species, *M. javana* Edwards (orig. des.).

**Diagnosis**. Mid ocellus present, smaller than lateral ocelli, lateral ocelli well separated from eye margin. Flagellomeres strongly enlarged, flattened, pedunculate. Mouthparts reduced, maxillary palpus very small, 1-segmented. Thorax strongly arched, thoracic bristles small. Laterotergite protruding, rather small, with some few setae. Tibiae with irregularly arranged microtrichia, devoid of larger tibial bristles. Fore tibia distal comb reduced to a few undifferentiated setulae, no posterior tibial comb. Wing microtrichia irregularly arranged, macrotrichia only on veins C, R_1_ and R_5_. Sc fairly long, free at the apex; sc-r missing (or weakly sclerotized); C extending to about half of distance between R_5_ and M_1_; first sector of Rs very short, evanescent, R_1_ very close to Rs at its base; R_1_ relatively long, r-m short, almost longitudinal; R_5_ reaching wing margin before level of tip of M_1_; medial fork sub-equal to length of petiole; posterior fork short, wide open, origin of M_4_ well before medial fork. CuP short, not even reaching level of origin of M_4_. Male terminalia simple, gonocoxites separated anteriorly, aedeagus reduced to a single more or less curved tube.

The genus *Metanepsia* was established by Edwards (1927) for a species from Java; in the same paper, a second species, from western Africa, was assigned to the same genus but was not formally described. Matile (1971a,b, 1973b, 1980a) added five Afrotropical species to the genus, while Kallweit (1998) added a second Oriental species.

*Metanepsia* has clearly derived features compared to other gnoristines. Some of them are shared with *Chalastonepsia* and *Pectinepsia* (e. g., the reduced mouthparts), while other features are exclusive of the genus, as the incomplete Sc, the displacement of the origin of Rs and of r-m towards the wing base, the shorter R_1_ and R_5_, and the longer first sector of CuA, with a shorter posterior fork. Apparently, the pectinate antenna varies within the genus—the type-species, *M. javana* Edwards, has slightly modified flagellomeres, which are not pectinate.

*Metanepsia malaysiana* Kallweit (Figs. 46A–D, 47A–E)

https://singapore.biodiversity.online/species/A-Arth-Hexa-Diptera-000760

*Metanepsia malaysiana* Kallweit 1998: 350, Fig. 2 (antenna), Fig. 6 (wing), figs. 16–18 (male terminalia). Type locality: Malaysia, B. Camp. 5°30’07” N, 101°26’21” E.

**Diagnosis**. Flagellomeres with ventral half more extended than dorsal half, a naked neck present distally on flagellomeres. Maxillary palpus with a single, small palpomere. Sc incomplete, sc-r absent; first sector of Rs very short, almost missing, M_1+2_ forking beyond level of tip of R_1_. Aedeagus elongated, heavily sclerotized, gonostylus rounded distally with a short row of curved spines along outer margin.

***Metanepsia_malaysiana_ZRCBDP0048531_hapZRCBD P0047793_SMH_describedearlier_type* [10: C, 11: T, 14: T, 16: A, 17: T, 18: T, 23: C, 35: A, 40: C, 44: G, 49: C, 132: C, 260: A]**

actatcttcCTctTtATTccatCcgggggcttctAttgaCctgGcaatCttttctctacatttagccggaatttcctctattttaggggcaattaattttattacaacaattattaatatacgagccccaaCaataacatttgatcaaatacctttattcgtatgatctgtatttatcacagctattttattactattatctttaccagtattagcgggggctattacaatattattaactgatcgaaatttaaatactAcattctttgaccctgcgggagggggagaccctattttataccaacatttattt

**Redescription**. **Male** (Fig. 46A). Wing length, 1.56 mm, width, 0.82 mm. **Head** (Fig. 46B). Shinning blackish-brown, frons elongate, flat, occiput blackish-brown, slightly lighter towards ventral margin, frontal furrow complete. Eyes rounded, interommatidial setulae present. Antenna light brownish-yellow, slightly pectinate, ventral half more developed and projected than dorsal half, densely pilose, each flagellomere with a distal neck almost as long as body of flagellomere. Scape elongate, pedicel very short. Mouthparts reduced, palpus with one visible palpomere, labella very small. **Thorax** (Fig. 46C). Mesonotum dark ochre-brown, slightly lighter laterally. No prescutum suture. Scutum covered by scattered, not too dense thin setae, a pair of clear lines of dorsocentrals, setae more dense over lateral margins. Scutellum dark brown, with a row of short setae along distal margin. Pleural sclerites ochre-brown, laterotergite and mediotergite darker, metepisternum light ochre-yellowish. Antepronotum well-developed, almost bulging, with three thin setae, almost entirely fused to proepisternum; proepisternum with three thin setae, proepimeron triangular, not fusing to katepisternum. Prosternum produced, bare, connected to proepisternum. Anepisternum, katepisternum, mesepimeron, mediotergite, and metepisternum bare, laterotergite with 16 thin setae concentrated dorsoposteriorly. Mesepimeron slender, reaching ventral margin of thoracic pleura, mediotergite slightly curved. Haltere pedicel light ochre-yellowish, pedicel brown, with few thin setulae. **Wing** (Fig. 46D). Membrane only slightly fumose, unmarked, membrane only with microtrichia, without macrotrichia. C produced beyond tip of R_5_ to about half distance to tip of M_1_. Sc long, incomplete, bare, ending well beyond first sector of Rs. R_1_ short, ending slightly over mid of wing. R_4_ absent, R_5_ ending before wing tip, at about level of tip of M_2_. First sector of Rs barely sclerotized, perfectly transverse, very short, R_1_ and R_5_ almost in contact at anterior end. Crossvein r-m more than 3× as long as first sector of Rs. Medial fork weakly sclerotized, stem of M_1+2_ very long, longer than medial fork. First sector of CuA long, origin of M_4_ more basal than base of medial fork. CuP weak, visible only half way to wing margin. **Legs**. Coxae light ochre-yellowish, only very distal tip of coxae with a brown marking, fore and mid femora, tibiae, and tarsi rather whitish-yellow, hind femur, tibia, and tarsus slightly darker. Legs covered with small, irregularly distributed trichia, distinctive setae entirely absent. Tibial spurs short, spurs of fore and mid tibiae less than twice tibiae apex width, hind spurs slightly over twice tibia width at apex. **Abdomen**. Tergite and sternite 1, whitish ochre-yellow, tergites and sternites 2-8, light ochre-brown; posterior segments with tergite gradually wider than sternite.

Setation on abdominal sclerites quite scattered and regularly distributed. **Terminalia** (Figs. 47A–B). Gonocoxites close to each other medially but in contact only at anterior fifth, not projected beyond base of gonostylus, each gonocoxite with a scale-like tooth at distal end of inner margin ventrally. Gonostylus curved, rather flattened, about half as long as gonocoxite, with five curved short spines at outer margin on distal half, with a distal rounded dorsal extension bearing a subdistal tooth. Gonocoxal bridge conspicuous, apodemes quite close to each other medially. Parameres medially fused to aedeagus ventrally, aedeagus elongate, curved ventrally at distal end. Tergite 9 rectangular, length less than half width, no medial protuberance. Sternite 10 triangular, weakly sclerotized, with some thin distal setae. Tergite 10 basically produced laterally to cerci. Cerci longer than tergite 9, with scattered thin setae.

**Female**. As males, except for the following. **Terminalia** (Figs. 47C–D). Sternite 8 with a pair of large lobes well separated by a deep medial incision, lobes densely covered by microtrichia and some scattered fine setae. Tergite 8 larger than tergite 9+10, both weakly sclerotized. Cercomere 1 1.6× length of cercomere 2, cercomere 2 less sclerotized.

**Material examined** (28 males, 4 females). **Males**: ZRCBDP0047791, Pulau Ubin (PU4), mangrove, 20.apr.13, MIP leg.; ZRCBDP0047922, Nee Soon (NS1), swamp forest, 31.jul.13, MIP leg.; ZRCBDP0048545, Nee Soon (NS2), swamp forest, 25.apr. 12, MIP leg.; ZRCBDP0048685, Nee Soon (NS2), swamp forest, 13.jun. 12, MIP leg.; ZRCBDP0048944, Nee Soon (NS2), 03.dec.14, MIP leg.; ZRCBDP0047792, Pulau Ubin (PU4), mangrove, 20.apr.13, MIP leg.; ZRCBDP0047957, Nee Soon (NS2), swamp forest, 18.set.13, MIP leg.; ZRCBDP0048532, Nee Soon (NS1), swamp forest, 25.apr. 12, MIP leg.; ZRCBDP0048535, Nee Soon (NS2), swamp forest, 05.Sep.12, MIP leg.; ZRCBDP0048537, Nee Soon (NS2), swamp forest, 21.nov. 12, MIP leg.; ZRCBDP0048538, Nee Soon (NS2), swamp forest, 17.oct. 12, leg.; ZRCBDP0048544, Nee Soon (NS2), swamp forest, 23.may. 12, MIP leg.; ZRCBDP0048546, Nee Soon (NS2), swamp forest, 16.may. 12, MIP leg.; ZRCBDP0048680, Nee Soon (NS1), swamp forest, 02.may. 12, MIP leg. (slide-mounted); ZRCBDP0048684, Nee Soon (NS2), swamp forest, 05.dec. 12, MIP leg.; ZRCBDP0048995, Nee Soon (NS2), 17.dec.14, MIP leg.; ZRCBDP0047793, Pulau Ubin (PU4), mangrove, 20.apr.13, MIP leg.; ZRCBDP0047824, Nee Soon (NS1), swamp forest, 29.may.13, MIP leg.; ZRCBDP0048070, Nee Soon (NS1), swamp forest, 12.jun.13, MIP leg.; ZRCBDP0048531, Nee Soon (NS1), swamp forest, 04.apr. 12, MIP leg. (website photo specimen); ZRCBDP0048539, Nee Soon (NS2), swamp forest, 30.may. 12, MIP leg.; ZRCBDP0048540, Nee Soon (NS2), swamp forest, 05.dec. 12, MIP leg.; ZRCBDP0048542, Nee Soon (NS2), swamp forest, 11.apr. 12, MIP leg.; ZRCBDP0048547, Nee Soon (NS2), swamp forest, 18.apr. 12, MIP leg.; ZRCBDP0048679, Nee Soon (NS1), swamp forest, 04.apr. 12, MIP leg.; ZRCBDP0048681, Nee Soon (NS2), swamp forest, 04.apr. 12, MIP leg.; ZRCBDP0048686, Nee Soon (NS2), swamp forest, 11.apr. 12, MIP leg.; ZRCBDP0048690, Nee Soon (NS2), swamp forest, 18.apr. 12, MIP leg. **Females**: ZRCBDP0048683, Nee Soon (NS1), swamp forest, 07.nov. 12, MIP leg.; ZRCBDP0048687, Nee Soon (NS2), swamp forest, 09.may. 12, MIP leg.; ZRCBDP0048688, Nee Soon (NS2), swamp forest, 02.may. 12, MIP leg. (slide-mounted) (MZUSP); ZRCBDP0048689, Nee Soon (NS2), swamp forest, 16.may. 12, MIP leg. (slide-mounted).

**Remarks**. There are six haplotypes among the specimens of *Metanepsia malaysiana* collected covered by this project (Fig. 47E) and there is no conflict between different criteria to delimit the species. There are specimens from the Nee Soon swamp forest and from the Pulau Ubin mangrove. This species fits in Matile’s (1971a, figs. 1-3) original description of *Metanepsia* in details of the antenna, mouthparts, wings and male terminalia, including the light color of the tergite and sternite 1, the light color of the metepisternum, and the short pedicel. The gonostylus is similar to that of *M. africana* Matile. Edwards (1927) illustrated the male gonostylus of *M. javana* Edwards and it clearly differs from the species in Singapore. In all details except the antenna, our specimens fit *M. malaysiana* Kallweit (Kallweit 1998, Fig. 6) including a nearly identical male terminalia. The antenna of *M. malaysiana* in the original description, however, has a rather long pectination (Kallweit 1998, Fig. 2), while the antenna of the specimens from Singapore is almost identical to that attributed to *Chalastonepsia* sp. (Kallweit 1998, Fig. 3), a species with a very different wing venation (Kallweit 1998, Fig. 5). The male terminalia of our Singapore specimens perfectly fits *M. malaysiana* and for the time being we accept them as conspecific with the holotype of *M. malaysiana*.

### Mycomyinae

The Mycomyinae are the second smallest subfamily of mycetophilid in terms of number of genera (with the Manotini as a subclade of Leiinae and having the Tetragoneurinae as a separate subfamily), but the third most species-rich, with over 600 described species. The subfamily in the past included only the genera *Mycomya* Rondani (which has several subgenera, see e. g., Väisänen 1984) and *Neoempheria*. More recently, eight additional genera were added: *Echinopodium* Freeman, with 36 species known from temperate South America (Freeman 1951), *Viridivora* Matile with two Afrotropical species (Matile 1973,b), *Mycomyiella* Matile with nine Afrotropical species (Matile 1973), *Moriniola* Matile with one Afrotropical species (Matile 1976b), *Dinempheria* Matile with seven Afrotropical species (Matile 1979b), *Parempheriella* with 36 Afrotropical and two Oriental species (see, e.g., Matile 1973b,c, 1980b) *Syndocosia* Speiser with nine species from Africa (see, e.g., Matile 1976a), and *Vecella* Wu & Yang with one species from southeast China (Wu & Yang 1996).

The phylogenetic position of these genera within the Mycomyinae is not clear. *Echinopodium* seems to be a subclade of *Mycomya* (Väisänen, 1984). *Vecella* is actually a species of *Parempheriella*, a genus overlooked in Wu & Yang’s (1986) original publication—formal synonymy is proposed here. *Viridivora* and *Dinempheria* are obviously associated to the *ferruginea*-group of *Neoempheria*. *Mycomyiella* and *Moriniola* be offshoots of *Neoempheria*, with a small cell r1, while *Parempheriella* and *Syndocosia* subclades of *Neoempheria* that lost R_4_.

What is needed is a global review of *Neoempheria* as was discussed above for *Dziedzickia*. A partial solution would be the recognition of smaller clades within *Neoempheria* and *Mycomya*, and then gradually assigning species from these genera to the newly created genera or subgenera. This would restrict, at the end, *Neoempheria*, for example, to the “*ferruginea*-group” or part of it, to which *N. striata* (Meigen), the type-species of the genus, probably belongs. Some of Edwards (1940) informal groups of species already have generic names available, as *Neurocompsa* Enderlein and *Pleonazoneura* Enderlein (Enderlein 1910).

It was rather surprising that no species of *Mycomya* showed up in our samples from Singapore. There are over 40 described *Mycomya* species from the Oriental region, some of them known from India and Sri Lanka, Myanmar, Borneo and Sumatra, and Guangxi-Zhuang, in southern China (see, e.g., Colless & Liepa 1973, Väisänen 1984a, 1996, 2013a, 2013c, Wu & Yang 1994, Wu et al. 2001).

And there are also tropical species of *Mycomya* in the Neotropical region. This is one of the genera that are still expected to be collected in Singapore. Our Singapore samples so far included four species of *Parempheriella* and 31 species of *Neoempheria*.

### Parempheriella Matile

*Parempheriella* Matile 1974a: 227. Type-species, *P. lobayensis* Matile (orig. des.).

*Vecella* Wu & Yang 1986: 86. Type-species, *V. guadunana* Wu & Yang (orig. des.). **new synonym**

**Diagnosis** (modified from Matile 1974b). Lateral ocelli large, close to small mid ocellus. Maxillary palpus with 3-palpomeres. Scutum with a pair of bare bands, setation sparse. Laterotergite and mediotergite bare. C extending beyond tip of R_5_, Sc reaching C at level of origin of Rs or more basally, sc-r present; R_4_ absent; medial fold present, either sclerotized or not; basal end of r-m apart from origin of Rs; M_1_rather unsclerotized at basal end, M_4_ originating way more basal than origin of medial fork. Trichia regularly distributed on tibiae.

*Parempheriella* was originally described for the Afrotropical region and now has 36 known species. Besides the Afrotropical species, we have *P. septentrionalis* Matile, from South Korea (a northern extension of an Oriental clade into the Palearctic region), *P. guadunana* (Wu & Yang), **n**. **comb**., from the Fujian Province, in southeast China, and *Parempheriella defectiva* (Edwards), from Sumatra. Matile (1999) referred to undescribed species he was aware of from Sulawesi and from Sri Lanka. We found in our samples in Singapore four species of *Parempheriella*. One of these species fits in great detail with the description of *P. defectiva*—for which there is a quite good illustration of the male terminalia (Edwards 1931, Fig. 4). There are no species delimitation conflicts for the haplotype network of the four species (Fig. 48I). Wu & Wang (1986) may have missed Matile’s (1974a) original description of *Parempheriella*. The wing of *Vecella* perfectly fits the wing of different *Parempheriella* species—shape of R_1_ and R_5_, extension of r-m, shape of the medial fork, extension of R_1_ and position of sc-r etc. Also, the details of the male terminalia of *Vecella*, as the shape and sclerotization of the gonostylus and the gonocoxite lobes, also agrees with the species of *Parempheriella*.

*Parempheriella defectiva* (**Edwards) (**Figs. 48A–H)

https://singapore.biodiversity.online/species/A-Arth-Hexa-Diptera-000763

*Mycomya (Neoempheria*) *defectiva* Edwards 1931: 266, Figs. 4v, 4d (male terminalia, ventral and dorsal views). Type locality: Indonesia, Sumatra, Fort de Kock [=Bukittinggi] (920 m). Matile 1974b:611 (new combination).

**Diagnosis**. Scutum dark brown, thoracic pleura greyish brown, head and abdominal sternites ochre-brown, hind femur entirely cream-yellow with a greyish tinge, tergites 1–6 dark brown, tergite and sternite 7 in males and females dark ochre-yellowish. Gonostylus with a basal peduncle, clavate distally, inner face distally with a regular row of six elongate setae directed inwards.

***Parempheriella_defectiva_ZRCBDP0155005_hapZRCB DP0048717_describedearlier_type* [43: G, 64: T, 73: C, 1 00: C, 169: C, 205: T, 247: G]**

tctttcttctacaattgcccatacaggagcatcagtagatttGgctattttttctttacacctTgcaggtatCtcttcaattttaggagctgtaaatttCattacaacaattattaatatacgagccccaggaattcaatttgatcgaatacctttatttgtttgatcCgttttaattactgcggttctattactattatctctTcctgttttagcaggggctattactatattattaacagatcgGaatttaaatacaagcttctttgaccccgctggaggtggtgatcctattttatatcaacatttattt

### Redescription

**Male** (Figs. 48A). Wing length, 2.04–2.45; width, 0.84–0.99 (n=2). **Head**. Occiput and frons light brown. Ocelli in contact with each other medially. Antennae brownish, scape and pedicel light brown, with a crown of setulae around distal margin, pedicel with a strong dorsal seta; flagellomeres slightly wider than long, except first and last, longer than wide. Eyes densely covered with interommatidial setulae. Face and clypeus slightly bulging, yellowish-brown, palpi dirty-yellow, palpomeres darker towards tip, proboscis dirty-yellow. Maxillary palpus 3-segmented (palpomeres 1 and 2 largely membranous, not forming complete segments), palpomere 3 wider, with a sensorial pit, palpomere 4 longer than 3, projecting slightly beyond insertion of palpomere 5, last palpomere almost twice as long as second, with setulae on distal half. Labella small. **Thorax** (Figs. 48C). Scutum and scutellum shining blackish-brown, lighter areas over sutures and laterally on scutellum. Scattered smaller setae on scutum, long supra-alars; a more or less regular row of dorsocentrals and acrostichals; scutellum with two long, diverging lateroapical bristles and an additional pair of smaller setae. Pleural sclerites brown, mesepimeron lighter dorsally, metepisternum lighter anteriorly, laterotergite and mediotergite dark brown. Dorso-lateral branches of basisternite 1 wide, not fused to proepisternum. Antepronotum wide, three stronger and some additional smaller setae, proepisternum reduced, devoid of any setae or setulae; all other pleural sclerites bare. Haltere light brown, darker at base of knob. **Legs**. Anterior coxae cream-yellow, more yellowish towards tip, mid and posterior coxae light cream-yellow. Femora cream-yellowish, tibiae and tarsi greyish-yellow, with regularly distributed setulae, a row of stronger setae along tibiae laterally and dorsally. Fore coxa covered with fine setae on anterior and external faces, mid coxa with fine setae restricted to distal half of external face, hind coxa with a regular line of fine setae along its length. Femora slightly enlarged medially, especially hind femur. Fore tibia long and slender, with trichia arranged in more or less regular rows, only some few setae dorsally and laterally along its length and some distal setae; anteroapical depressed area at internal face small, with a more or less irregular comb of setae. Mid tibia with two dorsolateral rows with few setae. Hind tibia with two long dorsolateral rows of setae and a long regular comb of setulae distally. Tibial spurs greyish-brown, mid and hind spurs subequal. Tarsi with few setae, besides rows of trichia, tarsal claw with two minute teeth. **Wings** (Figs. 48D). Membrane light greyish. C extending beyond apex of Rs for about a ¼ of distance to tip of M_1_. Humeral vein strongly inclinate. Sc bare, long, ending at C before level of basal end of r-m; sc-r weak but present. First section of Rs strictly transverse, straight, both R_1_ and R_5_ mostly straight; r-m oblique, very slightly curved. Medial fork longer than M_1+2_; M_4_ originating at level of anterior end of M_1+2_. M_4_ gently sinuous on distal half. CuP weakly sclerotized; R_1_, R_5_, distal 4/5 of M_1_, distal half of M_2_ and M_4_, and distal ⅔ of CuA setose dorsally. Abdomen. Tergites 1-6 shining brown, sternites 1-6 ochre-yellowish with a brownish tinge, tergite and sternite 7 yellowish. Terminalia (Figs. 48E-F). Yellowish-brown. Gonocoxites fused medially on anterior fifth of terminalia, suture of fusion evident, a sclerotized membranous area between medial projections; ventral face of gonocoxite with three posterior lobes: medial lobe long, blade-like, bare; second lobe long, digitiform, with a pair of long, fine, curved setae; third lobe short, digitiform, with a long subapical seta and two fine apical setae; gonocoxite dorsally with a short lateral projection with setae, from which a long falciform blade projects at apex. Gonostylus club-shaped, a wider base bearing a strong seta, a long neck with a strong seta directed inwards and a widened, truncate distal end covered with a group of setulae, with a row of fine setae on distal margin and a modified area with some setulae and a strong seta. Aedeagal-parameral complex large, present as a wide plate with sclerotized lateral margins and a medial sclerotized axis, ventrally with a pair of short digitiform projections with short denticules on distal end, and projected into a trapezoid membrane distally between gonostyli; anteriorly with a pair of slender apodemes. Gonocoxal bridge wide, without evident apodemes. Tergite 9 with a pair of very long projections with setae on basal half, extending way beyond rest of terminalia into a pair of digitiform projections, each bearing a group of five short, curved setulae. Sternite 10 trapezoid, weakly sclerotized, cerci quite elongate, weakly sclerotized medio-distally.

**Female** (Figs. 48B). As male, except for the following. **Abdomen**. Tergites 1–6 light brown, sternites lighter, tergite and sternite 7 yellowish-brown. **Terminalia** (Figs. 48G–H). Sternite 8 trapezoid, with a medio-posterior incision separating a pair of distal setose lobes. Sternite 9 wide, with a short medial apodeme directed anteriorly. Tergite 8 rectangular, wide and short, with a single row of setae, tergite 9 with a sclerotized band along anterior border and a pair of long setae on dorsoposterior corners, tergite 10 partially fused to tergite 9, setose. Cerci elongate, cercomere 1 2.5× length of cercomere 2.

#### Material examined

*Parempheriella defectiva* (Edwards), syntype male, “Sumatra, Fort de Kock, 2005” (NHM). **Sequenced specimens** (17 males, 2 females). **Males**: ZRCBDP0047858, Nee Soon (NS1), swamp forest, 10.jul.13, MIP leg.; ZRCBDP0047865, Nee Soon (NS2), swamp forest, 02.oct.13, MIP leg.; ZRCBDP0048717, Nee Soon (NS2), 28.jan.15, MIP leg.; ZRCBDP0048818, Nee Soon (NS1), 14.jan.15, MIP leg.; ZRCBDP0048828, Nee Soon (NS1), jan.15, MIP leg. (slide-mounted) (MZUSP); ZRCBDP0048843, Nee Soon (NS1), 14.jan.15, MIP leg.; ZRCBDP0048914, Nee Soon (NS1), 31.dec.14, MIP leg.; ZRCBDP0049009, Nee Soon (NS2), 17.dec.14, MIP leg. (extracted; ZRCBDP0049025, Nee Soon (NS2), 07.jan.15, MIP leg.; ZRCBDP0049029, Nee Soon (NS2), 07.jan.15, MIP leg.; ZRCBDP0049030, Nee Soon (NS2), 07.jan.15, MIP leg.; ZRCBDP0049032, Nee Soon (NS2), 07.jan.15, MIP leg.; ZRCBDP0049040, Nee Soon (NS2), 07.jan.15, MIP leg.; distal 4/5 of M_1_, distal half of M_2_ and M_4_, and distal ⅔ of CuA setose dorsally. **Abdomen**. Tergites 1–6 shining brown, sternites 1–6 ochre-yellowish with a brownish tinge, tergite and sternite 7 yellowish. **Terminalia** (Figs. 48E–F). Yellowish-brown. Gonocoxites fused medially on anterior fifth of terminalia, suture of fusion evident, a sclerotized membranous area between medial projections; ventral face of gonocoxite with three posterior lobes: medial lobe long, blade-like, bare; second lobe long, digitiform, with a pair of long, fine, curved setae; third lobe short, digitiform, with a long subapical seta and two fine apical setae; gonocoxite dorsally with a short lateral projection with setae, from which a long falciform blade

ZRCBDP0049041, Nee Soon (NS2), 07.jan.15, MIP leg.; ZRCBDP0049048, Nee Soon (NS2), 07.jan.15, MIP leg.; ZRCBDP0049049, Nee Soon (NS2), 07.jan.15, MIP leg.; ZRCBDP0155005, Singapore, NSM,14.jan.15, MIP leg.

**Females**: ZRCBDP0048556, Nee Soon (NS1), swamp forest, 09.may. 12, MIP leg. (website photo specimen, slide-mounted); ZRCBDP0049054, Nee Soon (NS2), 07.jan.15, MIP leg.

**Additional sequenced specimens:** ZRCBDP0128631, Bukit Timah (BT06), 08.feb.17; ZRCBDP0140219, Nee Soon (NS2), 21.jan.15; ZRCBDP0140220, Singapore; ZRCBDP0140224, Nee Soon (NS2), 21.jan.15; ZRCBDP0140707, Nee Soon (NS2), 04.feb.15; ZRCBDP0140718, 04.feb.15; ZRCBDP0140719, Nee Soon (NS2), 04.feb.15; ZRCBDP0140766, Nee Soon (NS2), 31.dec.14; ZRCBDP0154972, Nee Soon (NS2), 14.jan.15; ZRCBDP0154974, Nee Soon (NS2), 14.jan.15; ZRCBDP0154975, Nee Soon (NS2), 14.jan.15; ZRCBDP0154979, Nee Soon (NS2), 14.jan.15, ZRCBDP0154999, Nee. Soon (NS2), 14.jan.15; ZRCBDP0155100, Nee Soon (NS2), 21.jan.15. **Sequence failure specimen**: male, ZRCBDP0154761 (slide-mounted).

**Remarks**. The type locality of *P. defectiva* is “Fort De Kock” (now Bukittinggi), in West Sumatra. The minimum width of the Strait of Malacca, separating Sumatra from the Malay Peninsula, is 2.8 Km, with a minimum depth of 25 m. It is highly probable that part of the Singapore fauna of mycetophilids is shared with Sumatra at the species level. The only disagreement between our specimens and Edwards’ (1931) original description is with regard to the color of tergite 4. In his description the tergite 4 is referred to as “yellowish” (like in our other species of *Parempheriella*), our specimens of *P. defectiva* have tergite 4 concolor with the remaining tergites. We assume that our specimens are conspecific with the holotype. Having specimens of *P. defectiva* from Sumatra that can be sequenced would allow a better understanding of the genetic divergence between populations on both sides. In a more general sense, understanding the evolution of the tropical forests in Southeast Asia. All but one of the specimens of *P. defectiva* in our samples come from the Nee Soon swamp forest, with a single specimen from Bukit Timah. There are two different haplotypes. We have examined and photographed one of the syntypes of *P. defectiva* (NHM).

*Parempheriella mait* Amorim & Oliveira, sp. n. (Figs. 49A–D, 50A–D)

https://singapore.biodiversity.online/species/A-Arth-Hexa-Diptera-000764

urn:lsid:zoobank.org:act:484D5620-207F-41F4-873A-E137C32445CB

**Diagnosis**. Head brown, yellowish dorsolaterally, scutum and scutellum shining brown, lighter along laterals, pleural sclerites brown, antepronotum, laterotergite, and mediotergite darker, femora yellowish-brown. Tergites 1-3 and 5-6 uniformly brown, tergite 4 cream-yellow with a slender brown band along anterior margin, tergite 7 yellowish-brown. Male syngonocoxite with a pair of long, digitiform ventral lobes and a pair of well-developed setose lateral lobes with an inner small digitiform projection; gonostylus small, digitiform.

***Parempheriella_mait_ZRCBDP0048925_hapZRCBDP0 048925_SMH_holotype* [62: C, 64: T, 67: C, 70: G, 76: C, 196: G, 212: C, 265: C, 268: C, 298: T]**

tctttcttctacaattgctcatacaggggcatctgtagatttagctattttttctcttcatCtTgcCggGatttcCtcaattttaggggcagtaaattttattactacaattattaatatacgagccccaggtattcaatttgatcgaatacccttatttgtttgatcagttttaattactgctgttcttctattGctatctttacctgtcCtagcaggggctattacaatattattaacagatcgaaatttaaatactagtttCttCgacccagctggtgggggagaccctatcctTtatcaacacttattt

#### Description

**Male**. **Head** (Fig. 50A). Occiput and frons brown, more yellowish dorsolaterally. Antennae uniformly light brown. Face and clypeus light brown, palpus and labella dirty whitish-yellow. **Thorax** (Fig. 50B). Scutum and scutellum shining brown, lighter areas along lateral margins. Scutellum with two very long, diverging lateroapical bristles, no additional small setae. Pleural sclerites brown, antepronotum, laterotergite, and mediotergite darker. Halter pedicel light brown, knob slightly darker. **Legs**. Coxae whitish-yellow, fore coxa brownish on basal fifth; femora yellowish-brown, tibiae and tarsi light brownish, with regularly distributed setulae, a row of stronger, regularly distributed setae along tibiae laterally and dorsally. **Wings**. Membrane light greyish. C extending very shortly beyond tip of R_5_. **Abdomen**. Tergites 1-3 and 5-6 uniformly brown, tergite 4 cream-yellow with a slender brown band along anterior margin, tergite 7 yellowish-brown; sternites 1-4 and 7 brownish-yellow, sternites 5–6 darker. **Terminalia** (Figs. 50C–D). Gonocoxites fused medially on anterior fifth of terminalia, an evident suture between them, with only some few setae; syngonocoxite ventrally with a pair of long, digitiform lobes extending beyond tip of gonostylus and a pair of well-developed setose lateral lobes truncate distally, bearing an inner small digitiform lobe. Gonostylus small, digitiform, with sub-basal seta and a group of small distal setae. Aedeagal-parameral plate well-defined, with a pair of hook-like laterodistal projections. Gonocoxal bridge evident. Tergite 9 with a pair of lateral projections distally, with fine setae along most length. Cerci present as a pair of setose lobes medially.

**Female** (Fig. 49A). As males, except for the following. **Wing** (Fig. 49A). Length, 2.07; width, 0.79. **Terminalia** (Figs. 49C–D). Brownish-yellow, cerci more yellowish. Sternite 8 wide, trapezoid, with a pair of short projections on posterior margin with a few dark short setae medially. Sternite 9 (vaginal furca) wide, with a short anterior medial apodeme, distal part of ducts leading to gonopore composing a sclerotized slender cone. Tergite 8 wide, with a single row of setae along posterior margin. Tergite 9 slender, bare. Tergite 10 slender, with a row of setae along posterior margin. Cercomere 1 almost 3× longer than cercomere 2.

**Etymology**. The specific epithet of this species refer to Ma’it, a 9^th^ century Arab description of an island that fits Singapore. The noun is used in aposition.

#### Material examined

**Holotype**: male, ZRCBDP0048925, Nee Soon (NS1), 31.dec.14, MIP leg. (slide-mounted). **Paratypes** (2 females): ZRCBDP0048557, Nee Soon (NS1), swamp forest, 11.apr. 12, MIP leg.; ZRCBDP0048559, Nee Soon (NS2), swamp forest, 11.jul. 12, MIP leg. (slide-mounted).

**Remarks**. The color pattern of *Parempheriella mait* Amorim & Oliveira, **sp. n.** is similar to that of *P. longyamen* Amorim & Oliveira, **sp. n.,** but the male terminalia morphology reveals clear differences. We have three haplotypes for this species and no delimitation conflicts.

*Parempheriella longyamen* Amorim & Oliveira, sp. n. (Figs. 51A–G)

https://singapore.biodiversity.online/species/A-Arth-Hexa-Diptera-000790

urn:lsid:zoobank.org:act:97498B6E-3F4B-4AE9-B215-0F02B1D96E17

**Diagnosis**. Head dark greyish-brown, scutum and thoracic pleura dark brown, hind femur ochre-yellowish, abdominal sternites light brownish-yellow, tergites 1–3 and 5–6 brown, tergite 4 cream-yellow with a small medial mark along anterior margin, tergite and sternite 7 cream-yellow. Gonostylus strongly sclerotized, falciform. ***Parempheriella_longyamen_ZRCBDP0049091_hapZRC BDP0049091_SMH_holotype* [55: C, 137: C, 190: C, 19 3: T, 220: G, 226: C, 241: T]**

tttatcttcaactattgctcatacaggggcttcagttgatttagctattttttcCttacatcttgcaggtatttcttctattttaggggctgtaaatttcattactactattattaatatacgggccccaggtattCaatttgatcgaatacctttatttgtttgatctgttttaattacagcagttctCctTttattatctttaccagttttagccggGgctatCactatactattaacTgatcgaaatttaaatactagtttctttgacccagcaggaggaggagaccctattttataccaacatttattt

#### Description

**Male** (Fig. 51A). Wing length, 2.37–2.55; width, 0.92–0.99. **Head** (Fig. 51B). Occiput and frons brown. Antennae uniformly light brown; flagellomeres slightly longer than wide. Face and clypeus light brown, palpus and labella whitish. **Thorax** (Fig. 51C). Scutum and scutellum shining brown, lighter areas along lateral margin. Scutellum with two very long diverging lateroapical bristles, in addition to a row of four long fine setae slightly more dorsally. Pleural sclerites brown, antepronotum, laterotergite, and mediotergite darker. Halter light brown, pedicel lighter. **Legs**. Coxae whitish-yellow, darkened on distal fourth of wing; femora yellowish-brown, hind femur darkened towards tip, tibiae and tarsi light brownish, with regularly distributed setulae and a row of stronger and regularly distributed setae along tibiae laterally and dorsally. **Wings** (Fig. 51D). Membrane almost translucent. C extending only shortly beyond tip of R_5_. Sc reaching C at level of anterior end of r-m. **Abdomen**. Tergites 1-3 and 5-6 uniformly dark brown, tergite 4 mostly yellow, with a brown mark along anterior margin medially; sternites 1-5 whitish-yellow, darker towards apex. Tergite 7 light yellowish-brown, sternites 6-7 light yellowish-brown. **Terminalia** (Fig. 51E). Light brown on basal half, darker distally, gonostylus dark brown with strong black setae. Gonocoxites fused medially on anterior fifth of terminalia, with an evident suture between them, bearing few setae; syngonocoxite ventrally with a pair of long, well sclerotized digitiform bare projections laterodistally, extending beyond tip of gonostylus; a pair of posterior extensions of gonocoxites supporting basally tergite 9 and a pair of posterior extensions dorsally to insertion of gonostylus. Gonostylus with a basal elongate branch extending ventrally to main branch, both well-sclerotized, main branch with some fine, long setae on distal third. Aedeagal-parameral plate well-defined, wide with a posterior V-shaped extension. Gonocoxal bridge evident, apodeme closer to each other medially. Tergite 9 with a pair of lateral projections barely in contact medially, lateral projections with long, fine setae, extending to level of tip of gonostylus. Tergite 10 well-defined, lateral projections with a slender medial connection, each side with two digitiform projections, medial one slightly stronger, with three well developed setae on middle third, three additional strong setae and one spine, not reaching level of tip of gonostylus, outer digitiform projection longer, reaching level of tip of gonostylus, with a conspicuous distal seta. Sternite 10 visible, slightly elongate, rectangular. Cerci weakly sclerotized, fused to each other, lobose distally.

**Female**. As male, except as following. **Wing l**ength, 2.53; width, 0.97. **Terminalia** (Figs. 51F–G). Sternite 10 well-developed, laterally with a band with microtrichia and setae, medially with a transparent area. Tergite 9 more sclerotized, tergite 10 reduced to a band fused to tergite 9, with a row of three pairs of longer and some smaller setae. Cercomere 1 more than 3× longer than wide, 2.4× longer than cercomere 2.

#### Material examined

**Holotype**: male, ZRCBDP0049091, Nee Soon (NS1), 24.dec.14, MIP leg. (extracted, slide-mounted). **Paratypes** (4 males, 1 female). **Males**: ZRCBDP0278299, Singapore, 10.may.18, MIP leg. (slide-mounted) (MZUSP); ZRCBDP0134029, Singapore, date range 2012-2018, MIP leg.; ZRCBDP0278300, Singapore, 10.may.18, MIP leg.; male, ZRCBDP0278309, Singapore, 10.may.18, MIP leg. **Female**: ZRCBDP0072675, Bukit Timah (BT05), primary forest, 10.may.18, MIP leg. (slide-mounted).

**Etymology**. The specific epithet of this species refers to Long Ya Men [=龍牙門], or “Dragon’s Teeth Gate”, the Mandarin name of a place reached in 1320 by a Mongol Yuan Empire trade mission. It is believed to be the present day Keppel Harbour, at the southern part of Singapore. The noun is used in apposition.

**Remarks**. The specimens of the type-series come from Nee Soon swamp forest samples and from Bukit Timah primary forest samples. We have four haplotypes for this species and no delimitation conflicts.

*Parempheriella peranakan* Amorim & Oliveira, sp. n. (Figs. 52A–E)

https://singapore.biodiversity.online/species/A-Arth-Hexa-Diptera-000819,and-002062

urn:lsid:zoobank.org:act:37510D03-43B1-4F9F-806F-A30EDB8B73CF

**Diagnosis**. Scutum dark brown, head greyish-brown, thoracic pleura concolor with head, mid and hind femur whitish with brown distal fourth, abdominal sternites very light brown, tergites 1–3 and 5–6 greyish-brown, tergite 4 greyish-brown with a pair of cream-yellow marks on lateroposterior corners, tergite and sternite 7 cream-yellow. Gonostylus elongate, with two distal spines, more ventral one long and curved inwards.

***Parempheriella_peranakan_ZRCBDP0047777_hapZRC BDP0047777_SMH_holotype* [10: A, 19: C, 137: C, 217: T, 220: G, 244: C, 250: C, 262: C, 283: T, 298: T] -**

ttatcttcAacaattgcCcatacaggagcttcagttgatttagcaattttttctcttcatttagcaggtatttcttcaattttaggagctgttaattttattacaacaattattaacatacgagccccaggtattCaatttgatcgaatacctttatttgtttgatctgtattaattacagctattttattattattgtctttacctgttttagcTggGgccattactatattattaacagaCcgaaaCttaaatactagCttttttgacccagcaggaggTggagatcctattctTtatcaacatttattt

#### Description

**Male**. Wing length, 2.04; width, 0.82. **Head**. Occiput and frons light greyish-brown. Antennae light brownish, scape light ochre-yellow, pedicel light brown; flagellomeres only slightly longer than wide, except first and last, about 1.5 longer than wide. Face, clypeus, palpus and proboscis whitish. **Thorax**. Scutum and scutellum shining blackish-brown, lighter areas over sutures and laterally on scutellum. Scattered smaller setae on scutum, longer supra-alars; a more or less regular row of dorsocentrals and acrostichals. Scutellum with two long, diverging lateroapical bristles and an additional pair of small setae. Pleural sclerites brown, mesepimeron lighter dorsally and metepisternum anteriorly, laterotergite and mediotergite dark brown. Dorso-lateral branches of prosternum wide but not fused to proepisternum. Antepronotum wide, with three stronger and some additional smaller setae, proepisternum reduced, devoid of any setae or setulae; all other pleural sclerites bare. Haltere greyish. **Legs**. Coxae whitish, anterior coxa brownish on anterior fifth. Femora cream-yellowish, mid and hind femora brownish at distal fourth; tibiae and tarsi greyish, with regularly distributed setulae and a row of stronger and regularly distributed setae along tibiae laterally and dorsally. **Wing** (Fig. 52B). Membrane light greyish. C extending beyond apex of Rs for about one ¼ distance to tip of M_1_. Humeral vein strongly inclinate. Sc bare, ending at R before anterior end of r-m; sc-r weak but present. First section of Rs almost transverse, straight, both R_1_ and R_5_ mostly straight, slightly curved distally; r-m oblique, only slightly curved. Medial fork longer than M_1+2_; M_4_ originating slightly more basal than anterior end of M_1+2_. M_4_ slightly depressed on distal half, gently sinuous, posterior fork not too open.

CuP weakly sclerotized, ending at level of origin of M_4_. Veins R_1_, second sector of Rs, most of extension of M_1_ distal half of M_2_, distal 4/5 of M4, distal ⅔ of CuA with dorsal setation. Abdomen. Tergites 1-6 shining brown, sternites 1-6 ochre-yellowish with brownish tinge, tergite and sternite 7 yellowish. Terminalia (Figs. 52C-D). Yellowish-brown. Terminalia slightly asymmetric, Gonocoxites fused medially on anterior fifth of terminalia, with an evident suture between them, with very few setae; syngonocoxite ventrally with a pair of long digitiform projections at posterior margin bearing a strong seta basally at external face, and a strong seta, some setulae and a sclerotized tooth at internal margin midway to apex, with a long apical spine. Gonostylus elongate, articulated at limit between gonocoxite and tergite 9, bearing one fine seta at external face on distal third and two sclerotized distal spines, inner one straight and pointed, outer one long and curved. Aedeagal-parameral complex large, present as a wide plate with three sclerotized curved laterodistal teeth and a pair of blade-like extensions laterally, a pair of apodemes at anterior end. Tergite 9 fused to tergite 10, asymmetric, complex, mostly devoid of setation, a wide and deep incision separating two extensions; right projection with a short digitiform extension bearing a long spine, a long and slender digitiform projection with a small group of long and short setae only distally, and a laterodistal extension with some few long setae, to which articulates the gonostylus; left projection with a short digitiform projection with a long distal spine, a stronger digitiform projection with setulae along its length and three strong curved spines distally, no laterodistal extension. Cerci weakly sclerotized fused to each other, lobose distally.

**Female** (Fig. 52A). As male, except for the following. **Wing l**ength, 2.14–2.22; width, 0.87–0,92. **Terminalia** (Fig. 52E). Sternite 8 trapezoid, with a short medial incision on posterior margin separating small setose lobes, stronger setae along posterior margin. Sternite 9 wide, with a short medial apodeme anteriorly directed. Tergite 8 rectangular, wide and short, with a single row of setae, tergite 9 with a sclerotized band along anterior border and a pair of long setae on dorsoposterior corners, tergite 10 partially fused to tergite 9, setose. Cerci elongate, cercomere 1 2.5× length of cercomere 2.

#### Material examined

**Holotype**: male, ZRCBDP0047777, Nee Soon (NS1), swamp forest, 28.aug.13, MIP leg. (slide-mounted). **Paratypes** (8 males, 3 females). **Males**: ZRCBDP0048733, Nee Soon (NS2), 06.may.15, MIP leg.; ZRCBDP0048924, Nee Soon (NS1), 31.dec.14, MIP leg.; ZRCBDP0048931, Nee Soon (NS1), 31.dec.14, MIP leg.; ZRCBDP0048952, Nee Soon (NS1), apr.15, MIP leg.; ZRCBDP0048975, Nee Soon (NS1), 13.may.15, MIP leg.; ZRCBDP0048977, Nee Soon (NS1), 13.may.15, MIP leg.; ZRCBDP0049196, Nee Soon (NS2), 13.may.15, MIP leg.; ZRCBDP0049222, Nee Soon (NS1),10.dec.14, MIP leg. (slide-mounted) (MZUSP). **Females**: ZRCBDP0049227, Nee Soon (NS1), 10.dec.14, MIP leg.; ZRCBDP0133974, Singapore, NA, date range 2012-2018, MIP leg. (slide-mounted); ZRCBDP0072705, Bukit Timah Forest (BT05), 01.dec. 16, MIP leg. (slide-mounted). distal half of M_2_, distal 4/5 of M_4_, distal ⅔ of CuA with dorsal setation. **Abdomen**. Tergites 1–6 shining brown, sternites 1–6 ochre-yellowish with brownish tinge, tergite and sternite 7 yellowish. **Terminalia** (Figs. 52C–D). Yellowish-brown. Terminalia slightly asymmetric.

**Etymology**. The specific epithet refers to the Peranakan [= a Malay/Indonesian grammatical inflection roughly translating to ‘descendant’], an ethnic group of descendants from early 19th century Chinese settlers in the Malay Archipelago who intermarried with the indigenous populations of Malay, Javanese or Thai, resulting in a hybrid culture distinctive in the Malay Peninsula and Indonesian Archipelago. The noun is used in apposition.

**Remarks**. We found two haplotypes of *Parempheriella peranakan* Amorim & Oliveira, **sp. n.** and no delimitation conflicts. The specimens collected came from the Nee Soon swamp forest samples and in the Bukit Timah samples.

### Mycomya Rondani

*Mycomya* Rondani, 1856: 194. Type species: *Sciophila marginata* Meigen, 1818, by monotypy.

**Diagnosis** (modified from Väisänen 1984). Eyes slightly but distinctly emarginate above antennae, ocellar prominence seldom well developed, ocellar setae seldom reaching far beyond ocelli. Pronotum with several setae of different sizes, scutellum with 2 or 4 long setae. Suture between anepisternum and katepisternum more or less distinctly declining anteriorly, katepisternum not very wide, mediotergite bare or with small setae. Males of some groups with long spur on coxa 1. Wing hyaline, usually with no dark marks. C not distinctly produced beyond apex of R_5_, R_5_ and C usually reaching wing apex. Sc long, usually reaching level of cell r1, R_4_ present.

The only species of *Mycomya* in our Singapore samples belongs to the genus *M.* (*Cymomya*) Väisänen. The subgenus was described by Väisänen (1984) and includes species from the Palaearctic, Nearctic, and Neotropical, with further descriptions of species by Matile (1991), from New Caledonia, and by Väisänen (2013), from Myanmar and Nepal.

*Mycomya sachmatich* Amorim & Oliveira, sp. n. (Figs. 54A–D)

https://singapore.biodiversity.online/species/A-Arth-Hexa-Diptera-000745

urn:lsid:zoobank.org:act:8CBA50D0-3394-46C1-9CF8-0272A24BF977

**Diagnosis**. Head whitish-yellow, scutum yellowish with a pair of slender brown bands on posterior half, scutellum yellowish-brown; pleural sclerites whitish with an orange tinge. Tergites 1–7 brown with a cream-yellow longitudinal band on anterior half extending posteriorly on laterals. Wing with a brown mark at level of cell r1, not extending beyond M_1+2_, and a mild brownish tinge transverse band at level of distal half of R_1_, not extending beyond M_1_. Cell r1 slightly elongate, anterior margin of cell r1 4.0× length of R_4_; sc-r present at distal half of cell r1, tip of Sc not produced, Sc reaching R_1_ (through sc-r). Male syngonocoxite with a medio-posterior short extension towards aedeagus, gonocoxites entirely bare ventrally, a long laterodistal lobe extending to level of tip of gonostylus, with a dorsal short basal lobe bearing a pair of elongated spines directed inwards; gonostylus long, pedunculate, wider at apex, with a pair of short, curved spines; parameres with a pair of strongly sclerotized ventral extensions crossing each other medially; tergite 9 with a pair of elongate projections covered with microtrichia and setae.

***Mycomya_sachmatich_ZRCBDP0048490_hapZRCBDP 0047837_SMH_holotype* [4: C, 24: T, 31: C, 85: T, 91: T, 214: G]**

actCtcatctactattgctcataTaggagcCtcagtagatttagctattttttcccttcatttagcaggaatttcctcaattctTggagcTattaactttattactacaattattaatatacgatctccaggaattaaatttgatcgaatacctttatttgtttgatcagtattaattactgcaattttactattattatctctaccagtactGgcaggagctattactatattattaacagatcgaaatttaaatacttcattttttgacccagcaggaggaggagacccaattttatatcaacatttattt

#### Description

**Male** (Fig. 54A). Wing length, 2.43; width, 0.98. **Head**. Cream-yellow, lighter at frons and ventrally at occiput. Some scattered short setae over frons and occiput, some slightly longer setae around eye on occiput. Two ocelli placed medially on vertex over a blackish background. Face whitish, clypeus light brown. Antennal scape and pedicel light ochre-yellow, flagellum light yellowish with some light greyish tinge. Maxillary palpomeres light brown, last palpomere lighter. Frons with no setae anteriorly to line of ocelli, ocellar setae absent. Scape 1.2× longer than pedicel, flagellomere 1 1.3× longer than flagellomere 4, flagellomere 4 1.0× longer than wide.

Palpomere 4 1.3× palpomere 3 length, palpomere 5 1.9× palpomere 4 length. **Thorax**. Scutum background light yellowish, an even lighter area laterally, a pair of brown bands externally to dorsocentral line on posterior half of scutum; scutellum yellowish-brown. Pleural sclerites cream-yellow with an orangish tinge except for brownish tinge on laterotergite, mediotergite with a yellowish-brown mark medially. Scutum with a pair of prescutellar bristles at dorsocentral lines and two pairs of prescutellar bristles at lateroposterior corners; scutellum with two pairs of scutellar bristles, no additional setae. Antepronotum with 2–3 bristles and some additional smaller setae, proepisternum bare. **Legs**. Coxae whitish, fore coxa with a yellowish tinge, femora whitish-yellow, tibiae and tarsi slightly more greyish. Mid tibial inner spur 3.0× longer than tibia width at apex [both hind leg femora, tibiae and tarsi missing on holotype]. Foreleg tarsomere 1 0.8× tibia length, 1.6× tarsomere 2 length. **Wing** (Fig. 54B). Membrane background light greyish, with a dark greyish-brown mark on membrane at level of tip of Sc. C barely produced beyond tip of R_5_; Sc reaching sc-r, missing beyond sc-r (not visible even under phase contrast); sc-r regularly sclerotized; sc-r placed at distal half of cell r1; first sector of Rs short, R_4_ apart from base of r-m, anterior margin of cell r1 2.5× length of R_4_; first sector of Rs 0.67× r-m length; r-m oblique. False medial vein weak, not sclerotized. M_1+2_ 6.0× r-m length; bM 9.0× length of first sector of Rs. First sector of CuA 0.9× longer than second sector. M_1_, M_2_ and M_4_ gently curved. Cubital pseudovein sclerotized to distal third of second sector of CuA, CuP sclerotized to level of origin of M_4_, faint more distally. Wing margin gently emarginated at tip of CuA. No ventral macrotrichia on veins, dorsal macrotrichia on entire length of bR, R_1_, second sector of Rs; Sc, sc-r, first sector of Rs, R_4_, r-m, bM and all posterior wing veins entirely devoid of setation; no macrotrichia on wing membrane. **Abdomen**. Tergites 1–7 with a cream-yellow band anteriorly and a light brownish band posteriorly, tergites 2–7 with wider cream-yellow band, tergite 7 rectangular, no ornamentation. Sternites 1–7 cream-yellow. **Terminalia** (Fig. 54C). Cream-yellow. Gonocoxites large, fused medially without a suture, entirely bare ventrally, a large medial, weakly sclerotized plate connecting gonocoxites, with a medio-posterior short extension towards aedeagus, laterodistally a large setose projection extending to level of tip of gonostylus, inner margin, posteriorly to level of insertion of gonostylus, a pair of spines directed obliquely towards distal end. Gonostylus insertion displaced more dorsally, gonostylus elongate, capitate, bare at basal ¾, with a pair of short, curved spines on a small lateral projection, distal enlarged area covered with setae. Gonocoxal bridge wide, weakly sclerotized. Aedeagus with a pair of slender, elongate blades projecting almost to level of tip of gonostylus. Parameres with a pair of strongly sclerotized ventral extensions crossing each other medially, lateroanteriorly a pair of apodemes with a concentrate group of small setae, connected medially behind aedeagus. Tergite 9 with a pair of elongate projections covered with microtrichia and setae. Cerci small, with setae directed ventrally.

**Female** (Fig. 54A). As male, except for the following. **Wing l**ength, 2.52; width, 0.98. **Terminalia** (Fig. 54D). Sternite 8 as a pair of triangular, elongate lobes not fused together medially, deep posterior incision almost reaching anterior end of sclerite, entirely covered with microtrichia, setae on distal half of each lobe, stronger setae at tip. Sternite 9 weakly sclerotized, anterior apodeme extending towards anterior margin of sternite 8. Tergite 8 with a pair of lateral lobes with a slender medial connection, lateral end of anterior margin extending to ventral face of terminalia, covered only with microtrichia. Tergite 9+10 with a pair of lateral lobes with microtrichia and setae, a slender medial connection present, lateral extensions with longer setae. Sternite 10 with an elongate medial transparent area. Cercomeres dorsoventrally compressed, densely covered with microtrichia and short setae, cercomere 1 2.2× length of cercomere 2.

#### Material examined

**Holotype**: male, ZRCBDP0048490, Nee Soon (NS1), swamp forest, 09.may. 12, MIP leg. (slide-mounted). **Paratypes** (2 males, 3 females). **Males**: ZRCBDP0048721, Nee Soon (NS2), 28.jan.15, MIP leg.; male, ZRCBDP0155050, Singapore, date range 2012-2018, MIP leg. **Females**: ZRCBDP0047837, Nee Soon (NS2), swamp forest, 29.may.13, MIP leg.; ZRCBDP0047952, Nee Soon (NS1), swamp forest, 03.jul.13, MIP leg.; ZRCBDP0048498, Nee Soon (NS1), swamp forest, 25.apr. 12, MIP leg. (website photo specimen, slide-mounted).

**Etymology**. The specific epithet of this species refers to the 14th century Vietnamese transcription of Temasek, what is now Singapore, Sachmatich. The noun is used in apposition.

**Remarks**. This species of *Mycomya* belongs to *M.* (*Cymomyia*), as indicated by the vivid contrast of yellow and the pair of dark brown longitudinal marks, lack of the male mid-coxal spur, bare mediotergite, yellow abdominal tergites with posterior margins dark, and features of the male terminalia. There are two haplotypes for this species (Fig. 53), but there is no question about the species delimitation based on morphological features or molecular methods. It is present only in the swamp forest samples.

### *Neoempheria* Osten-Sacken

*Neoempheria* Osten-Sacken, 1878: 9 (*nom. nov.* for *Empheria* Winnertz). Type-species, *Sciophila striata* Meigen (auct.).

**Diagnosis**. Mid ocellus missing, lateral ocelli close together on vertex. Scutellum with bristles, laterotergite and mediotergite bare. Trichia regularly arranged along entire length of tibiae and tarsi, tibiae with bristles. Wing membrane with brownish marks; microtrichia on wing membrane not arranged regular longitudinal lines; C produced or not beyond tip of R_5_; R_4_ most usually present; false vein present between R_5_ and M_1_.

*Neoempheria* is a large genus of Mycetophilidae, with over 130 species described worldwide. *Neoempheria* is the genus with the largest number of species in our samples (31), slightly over *Epicypta* (30), though not as abundant. This number of congeneric species in sympatry is particularly impressive. It suggests that *Neoempheria* may be an open-ended or dark taxon nightmare even worse than *Manota*—except for the fact that *Neoempheria* species are large and quite colorful, while *Manota* species are small and more homogeneous in color patterns.

As mentioned above, it is likely that *Viridivora*, *Dinempheria*, *Mycomyiella*, *Moriniola*, *Syndocosia* and *Parempheriella* may be nested in *Neoempheria* to which generic status was given. Most species with large cell r1 (referred to here as the *ferruginea*-group) seem to belong to a clade deeply nested within the genus: considering the condition of cell r1 in most species of *Mycomya* and of other mycetophilid genera, the large cell r1 should be a secondary feature within the genus, not a retained plesiomorphy. In this scenario, *Parempheriella* seems to be a regular *Neoempheria* that misses R_4_. As mentioned above, for the time being we will not remove the generic status of *Parempheriella*.

There are 25 species of *Neoempheria* described from the Oriental region—14 of these species from China, three from India, two from Sri Lanka and India, three from Java and three from Sumatra. One of Edwards’ (1931) species from Sumatra originally described in *Neoempheria* was transferred to *Parempheriella* by Matile (1999). His other three *Neoempheria* species from Sumatra have small cell r1, with Sc ending beyond the origin of Rs—all with male terminalia illustrated. Two of his species do not fit any of the species of *Neoempheria* in our Singapore samples, while *N. zonalis* Edwards is present in our material. Colless (1966) described *Neoempheria vicina* Colless, from Micronesia, a species that also fits in the *ferruginea* species-group, Of the 31 species of *Neoempheria* in our samples, ten are known only from females and one species lacks a barcode. There are not enough features on color patterns and wing venation to differentiate the females of seven of these species from other species in the genus (even with female terminalia in most species being particularly complex, with lobes and digitiform projections on sternite 7, sternite 8, sternite 9, tergite 10 etc.). As happens with the genus *Manota*, we describe here these seven species of *Neoempheria* (species A–G), but we do not formally name them. The species lacking a barcode (species H) is described but not named. Three additional species known only from females are very distinctive, for which it is possible to prepare diagnoses, and are named here. The haplotype network for the genus (Fig. 53) has six species that have specimens split in smaller units, discussed below under the species descriptions.

Ten of the 31 species of *Neoempheria* found in our samples belong to the *ferruginea*-group, with a large cell r1 (see, e.g., Sueyoshi 2014). Edwards (1940) discriminated the Neotropical species of *Neoempheria* in eight informal groups and most these groups have species with large cell r1. Coher (1959) also has an extensive treatment of the genus in the New World, with a key for the Neotropical and Nearctic species of *Neoempheria*. Interestingly, few of the Neotropical species of *Neoempheria* have a small cell r1—as is the case of *N. neivai* Edwards. Just as an example, the Nearctic species *N. indulgens* Johannsen fits in the groups with small cell r1, while the Palearctic species *N. lineola* Meigen and the Nearctic *N. balioptera* Loew fit in the groups with a large cell r1. Different from what is seen in most other biogeographical regions, there is a large diversity of species of *Neoempheria* in Singapore with a small cell r1. We organized the diversity of species of the genus into four main groups and we use the name *ferruginea* for one of them; the others are *dizonalis*-group, *merlio*-group, and *puluochong*-group. This allows better visualizing similarities and patterns in the wing venation and in the male terminalia morphology between the species.

### Group *merlio*

This group includes seven of the species of *Neoempheria* described from Singapore, all of which with a short cell r4, around and a sclerotized medio-posterior process of the syngonocoxite in the male terminalia.

*Neoempheria merlio* Amorim & Oliveira, sp. n. (Figs. 55A–C)

https://singapore.biodiversity.online/species/A-Arth-Hexa-Diptera-000741

urn:lsid:zoobank.org:act:BB289C42-6544-43C5-A9B4-00FBD2B81C5B

**Diagnosis**. Head, scutum and scutellum ochre-yellowish with a brown tinge medially; pleural sclerites whitish-yellow, mediotergite light brown, dorsoposterior end of laterotergite with a light brown tinge. Tergite 1 light brown, tergites 2 and 4 with a brown medial longitudinal band, ochre-yellow laterally; tergites 3, 5–6 brown with a slender yellow band along anterior margin; tergite 7 yellowish; sternite 6–7 with brown lateral marks. Wing with a brown band distally across wing and a brown band more basally, from level of cell r1 to posterior margin on cell cua. Cell r1 small, anterior margin of cell r1 1.6× length of R_4_; sc-r not sclerotized; Sc reaching C before mid of cell r1. Male syngonocoxite with medial-posterior process extending towards aedeagus dorsally, gonocoxite with a long laterodorsal lobe, capitate distally, with a long spine directed inwards. Gonostylus long, slender, dorsoventrally compressed. Parameres with a pair of digitiform extensions laterally directed inwards.

***Neoempheria_merlio_ZRCBDP0047889_hapZRCBDP0 047889_SMH_holotype* [4: G, 7: C, 19: C, 55: C, 103: C, 127: C, 187: A, 202: A, 223: A, 244: C]**

tctGtcCtctacaattgcCcatacaggagcttcagtagatttagctattttttcCttacatttagcaggtatctcttcaattttaggtgctgttaattttatCacaactattattaatatacgagcCccaggaattcaatttgaccgaatgcctttatttgtttgatcagttttaattacagcagtAttattattactttcAttaccagttttagccggagcAattactatattattaacagaCcgaaatttaaatactagtttctttgaccctgctggaggaggagacccaattttataccaacatttattt

#### Description

**Male** (Fig. 55A). Wing length, 1.93; width, 0.72. **Head**. Light ochre, slightly darker at vertex. Antennal scape and pedicel dirty whitish, flagellum light ochre-yellowish. Face and clypeus light brown. Maxillary palpomeres brown, last palpomere lighter. Labella whitish-yellow. Ocellar setae absent, some setae on frons ahead of ocelli. Scape 1.5× pedicel length, flagellomere 1 2.3× longer than flagellomere 2; flagellomere 4 length 1.1× width. Clypeus projected beyond level of ventral margin of eye, a short proboscis present. Palpomere 4 1.5× palpomere 3 length, palpomere 5 1.5× palpomere 4 length. **Thorax**. Scutum ochre-yellowish, slightly darker close to posterior end, no brown stripes, scutellum ochre-yellowish. Pleural sclerites whitish. Halter whitish with light brown tinge, base of knob light brown. Scutum with irregular rows of long dorsocentrals, a pair of elongate bristles above level of anepisternum. Two pairs of prescutellars, one pair medially, one pair at lateroposterior corner; scutellum with one pair of scutellar bristles and one additional pair of fine setae placed more anteriorly. Two bristles on antepronotum. **Legs**. Coxae whitish, femora ochre-yellow, tibiae and tarsi light greyish-brown. Fore tibia with a small anteroapical depression lined with few setulae. Fore leg tarsomere 1 0.9× tibial length, 1.9× tarsomere 2 length. **Wing** (Fig. 55B). Membrane background light greyish, a dark brown band across wing at level of cell r1 and a lighter apical band beginning beyond basal end of medial fork. Sc weak, ending slightly beyond level of origin of Rs, sc-r absent; R_4_ weakly sclerotized, anterior margin of cell r1 1.7× length of R_4_. M_1+2_ 2.6× r-m length; bM 5.0× r-m length. False medial vein conspicuous, slightly sclerotized, almost reaching r-m. Medial fork wide open. M_4_ gently sinuous, first sector of CuA 0.70× length of second sector. Among posterior wing veins, only M_1_with macrotrichia on dorsal face, at distal fourth. **Abdomen**. Tergites 1–6 light brown, tergite 7 ochre-yellowish with a medial light brown mark along posterior margin. Sternites 1–7 light ochre-yellowish. **Terminalia** (Fig. 55C). Whitish-yellow. Gonocoxites fused medially, no suture of fusion, medial-posterior process of syngonocoxite extending towards aedeagus dorsally, a pair of very long laterodorsal lobes capitate distally, extending beyond tip of gonostylus, with setae on outer face, a long spine directed inwards and some strong setae at distal end. Gonostylus long, slender, dorsoventrally compressed, with setae mostly on outer face, long setae at distal end. Aedeagus with a hardly sclerotized structure with a pair of lateral winglets, a short anterior medial apodeme and a long pair of weakly sclerotized distal extensions. Parameres composed of a pair of long digitiform extensions laterally, almost as long as gonostylus, each with a subdistal seta and a pair of short and strong distal setae, a pair of lateral slender blades almost of same length, and a pair of valve-like, weakly sclerotized blades touching each other medially. Dorsally a subtriangular elongate membranous area that may correspond to tergite 9, with a pair of elongate lobes distally with microtrichia and setae.

**Female**. As male, except for the following. **Wing l**ength, 2.03; width, 0.66. **Abdomen**. Abdominal tergite 3 with medial brown band extending laterally until margin, tergite 7 light brown. **Terminalia**. Light brown. Sternite 8 trapezoid, with a medial U-shaped incision on posterior margin reaching posterior third of terminalia, entire sclerite with microtrichia and setae, long setae along tip of lobes. Sternite 9 with a sclerotized area around genital chamber, with a pair of anterior lobes close together directed anteriorly and a pair of lateral sclerotized extensions projecting dorsally bearing three strong setae. Tergite 8 wide, anterior margin laterally with sclerotized arms extending ventrally, posteriorly with a pair of lobes bearing a deep incision, strong setae on medio-posterior end of lobes. A pair of long digitiform extensions extending laterodistally from tergite 9+10. [Cerci broken].

#### Material examined

**Holotype**: male, ZRCBDP0047889, Nee Soon (NS2), swamp forest, 23.oct.13, MIP leg. (slide-mounted). **Paratypes** (7 males, 3 females). **Males**: ZRCBDP0048483, Nee Soon (NS1), swamp forest, 16.may. 12, MIP leg.; ZRCBDP0048726, Nee Soon (NS2), 15.apr.15, MIP leg.; ZRCBDP0048732, Nee Soon (NS2), 06.may.15, MIP leg.; ZRCBDP0048854, Nee Soon (NS1), 14.jan.15, MIP leg.; ZRCBDP0049184, Nee Soon (NS2), 13.may.15, MIP leg.; ZRCBDP0049199, Nee Soon (NS2), 13.may.15, MIP leg.; ZRCBDP0072731, Bukit Timah, primary forest (BT05), 14.dec. 16, MIP leg. **Females**: ZRCBDP0048481, Nee Soon (NS1), swamp forest, 25.apr. 12, MIP leg. (website photo specimen); ZRCBDP0049192, Nee Soon (NS2), 13.may.15, MIP leg. (slide-mounted); ZRCBDP0072702, Bukit Timah, primary forest (BT05), 01.dec. 16, MIP leg. **Additional sequenced specimens**: male, ZRCBDP0078969, Singapore, date range 2012-2018, MIP leg. ZRCBDP0048958, Nee Soon (NS1), 15.apr.15; ZRCBDP0155067, Nee Soon (NS2), 10.dec.14.

**Etymology**. The specific epithet of this species refers to “merlion”, the official mascot of Singapore (designed by the Singapore Tourism Board in 1964), corresponding to a mythical creature with the body of a fish and the head of a lion. The merlion is a prominent symbol of the nature of Singapore and of Singaporeans. In 2002 the statue and its ‘cub’ were relocated to its current position in the Merlion Park, that fronts Marina Bay, where it stands on a newly reclaimed promontory, just in front the historic The Fullerton Hotel. The noun is used in apposition.

**Remarks**. There are five haplotypes for *Neoempheria merlio* Amorim & Oliveira, **sp. n.** and there are no conflicts between the different delimitation approaches. The species was collected in the swamp forest, the Bukit Timah tropical forest and in the mangrove.

*Neoempheria sabana* Amorim & Oliveira, sp. n. (Figs. 56A–D)

https://singapore.biodiversity.online/species/A-Arth-Hexa-Diptera-000747

urn:lsid:zoobank.org:act:384A5842-FDF7-4227-BEE2-F48ED6C81413

**Diagnosis**. Head ochre-brownish with some whitish-yellow areas, scutum dark ochre-yellowish some areas with a light brown tinge, scutellum brownish; pleural sclerites whitish-yellow, mediotergite light brown on dorsal half, laterotergite dorsoposterior end brown. Wing with a brown band distally across wing and a brown band more basally, from level of tip of R_1_ to posterior margin on cell cua; cell r1 small, anterior margin of cell r1 1.2× length of R_4_; sc-r not sclerotized; Sc reaching C basal to origin of Rs. Tergites 1–2 brown, medially with large cream-yellowish lateral bands, tergites 3–4 brown with a slender cream-yellow band on lateral margin, tergites 5–6 brown, tergite 7 yellowish. Male syngonocoxite with a medio-posterior projection bearing a pair of strongly sclerotized arms, bare, with rounded, short lateral lobes; gonostylus displaced medially, long, dorsoventrally compressed, slender towards apex; parameres with a pair of short digitiform distal projections; tergite 9 with a pair of long, slightly curved projections widening towards apex as a valve.

***Neoempheria_sabana_ZRCBDP0048987_hapZRCBDP0 048495_SMH_holotype* [19: C, 37: A, 67: T, 127: C, 187 : A, 250: C, 251: C, 261: G, 262: T, 274: A]**

actatcttctacaattgcCcatacaggagcatctgtAgatttagctattttttctttacatttagcTggaatttcttctattttaggggctgttaattttattacaactattattaatatacgagcCccaggtattcaatttgatcgaatacctctatttgtttgatctgttttaattacagccgtAttattattattatcattaccagtattagctggagctattactatattattaacagatcgaaaCCtaaatacaaGTttttttgatccAgcaggaggaggagatccaattttatatcaacatttattt

#### Description

**Male**. Wing length, 2.30, width, 0.85. **Head**. Light ochre-brown at vertex, more ochre-yellow towards frons and ventral margin of occiput, ocelli with blackish background. Antennal scape and pedicel cream-yellow, flagellum whitish-yellow. Face cream-yellow, clypeus light brown. Maxillary palpus dark greyish-brown, labella small, ochre-yellowish. Setae on frons anteriorly to ocelli. Antennal scape as long as pedicel, flagellomere 1 2.0× longer than flagellomere 2, flagellomere 4 1.2× wider than long. Palpomere 4 0.8× palpomere 3 length, palpomere 5 2.0× palpomere 4 length. **Thorax**. Scutum dark ochre-yellow, more brownish towards posterior end; scutellum brownish. Pleural sclerites cream-yellow with an orangish tinge, laterotergite brown on dorsoposterior corner, mediotergite brownish dorsally, more yellowish-brown on ventral half. Scutum with one pair of medial pre-scutellar bristles and one pair on lateroposterior corner; one pair of scutellar bristles. Antepronotum with two strong bristles and some additional smaller setae, proepisternum with one bristle and three smaller setae. **Legs**. Coxae whitish with orangish tinge; femora ochre-yellow, hind femur darker; tibiae and tarsi light greyish-brown. Hind leg tibial inner spurs 3.0× tibia width at apex. Fore leg tarsomere 1 0.81× tibia length, 1.7× tarsomere 2 length. **Wing** (Fig. 56B). Membrane background light greyish, with a dark brown mark across distal third of wing (lighter on cell m2) and a dark brown band at level of tip of Sc to level of distal end of cubital pseudovein. Sc complete, reaching C slightly basal to origin of Rs, sc-r not clear (even under phase contrast); anterior margin of cell r1 1.2× length of first sector of Rs. False medial vein conspicuous, slightly sclerotized. Medial fork not wide open, M_2_ short. Origin of M_4_ before level of base of M_1+2_, M_4_ slightly sinuous on distal half. No ventral macrotrichia on veins, dorsal macrotrichia on entire length of bR, R_1_, second sector of Rs, anterior end of r-m, and on distal fourth of M_1_; Sc, sc-r, first sector of Rs, R_4_, anterior half of r-m, bM, M_1+2_, M_2_, M_4_, distal half of first sector, and CuA entirely devoid of setation; no macrotrichia on wing membrane. **Abdomen**. Tergites 1–2 brown medially, with large cream-yellow lateral bands, tergite 3 brown, with a slender cream-yellow band on lateral margin, tergite 4 brown laterally and along posterior margin, with a large cream-yellow area, tergites 5–6 brown, tergite 7 yellowish. Sternites 1–7 cream-yellow, sternite 6 and in some individuals sternite 5 with brown lateroanterior corner. **Terminalia** (Fig. 56C). Mostly ochre-yellow. Gonocoxites fused medially on anterior end of terminalia, entirely bare, a pair of elongate, strongly sclerotized medio-posterior projections connected to aedeagus, and a pair of short laterodistal lobes curved towards distal margin; ventrally to tergite 9 lateral projections, lobes extending to almost mid of gonostylus. Gonostylus displaced to a more medial position, elongate, slender towards apex, covered ventrally with microtrichia and long setae. Gonocoxal bridge wide, no clear apodemes. Aedeagus with a strongly sclerotized anterior portion, a medial ejaculatory apodeme directed anteriorly, weakly sclerotized on distal two-thirds, a pair of valves touching medially with a medial sclerotized suture, extending to level of tip of gonostylus, curved at distal end. Parameres with a pair of well-developed anterior apodemes, laterodistally with a pair of short digitiform projections with setae, medially with a membrane projection distally curved reaching slightly beyond tip of aedeagus. Tergite 9 with a slender median sclerite extending lateroposteriorly into a pair of long projections widening to apex and slightly curved as a valve, with setae along entire external face, bifid distally, dorsal end at apex with a hook-like projection, ventral end slightly capitate. Sternite 10 just dorsally to paramere, with a pair of short lobes densely covered with microtrichia and with setae close to tip. Cerci lobose, slightly elongate, not reaching level of tip of sternite 10, with microtrichia and fine setae, longer setae distally.

**Female** (Fig. 56A). As male, except for the following. **Wing l**ength, 2.20–2.43; width, 0.82–0.85 (n=2). **Abdomen**. Tergites 1–6 entirely brown, anterior segments lighter. **Terminalia** (Fig. 56D). Sternite 8 longer than wide, posterior margin with a V-shaped medial incision separating a pair of pointed lobes, anterior ¾ of sclerite only with macrotrichia, setae on distal fourth, lobes with longer and stronger setae, laterally a short projection with 3–4 long setae directed dorsally midway to apex. Sternite 9 wide, well sclerotized, anterior end with a pair of rounded lobes separated by a short medial incision, laterally with a pair of large arms, medially well sclerotized with a pair of winglets directed dorsally, distally with a pair of pointed, slender valves that open to a weakly sclerotized genital chamber with a rounded opening distally. Tergite 8 more slender on anterior end, wider medially, bare, with a wide and shallow medial depression along posterior margin. Tergite 9 wide, each lateral end with a pair long of digitiform projections, medially with two separate lobes close to each other, each with two short digitiform distal projections bearing a long seta at tip. Tergite 10 weakly sclerotized, U-shaped, densely covered with macrotrichia and setae, subapically on lateroventral extension a group of 3–4 long setae directed ventrally. Cercomere 1 elongate, wide at base and slender distally; cercomere 2 small, slender at base and widening to apex, rounded distally, both densely covered with microtrichia and short setae, cercomere 1 about twice cercomere 2 length.

#### Material examined

**Holotype**: male, ZRCBDP0048987, Nee Soon (NS2), 17.dec.14, MIP leg. (slide-mounted). **Paratypes**: 7 females, ZRCBDP0047068, National University of Singapore (Icube),01.jul.15, MIP leg. (slide-mounted); ZRCBDP0048495, Nee Soon (NS1), swamp forest, 12.dec. 12, MIP leg. (slide-mounted); ZRCBDP0048883, Nee Soon (NS1), 31.dec.14, MIP leg.; ZRCBDP0048949, Nee Soon (NS2), 03.dec.14, MIP leg.; ZRCBDP0048989, Nee Soon (NS2), 17.dec.14, MIP leg.; ZRCBDP0049105, Nee Soon (NS1), 24.dec.14, MIP leg. ZRCBDP0072724, Bukit Timah, old secondary forest (BT01), 08.dec. 16, MIP leg. (slide-mounted).

**Etymology**. The specific epithet of this species refers to the name given in the second century by Ptolemy [90–168 A. D. ] to a place at the Southern tip of the Golden Chersonese [= the Malay Peninsula], assumed to be Singapore, Sabana. The noun is used in apposition.

**Remarks**. Females have darker abdomen, nearly without any cream-yellow areas. Two of the females have the sclerotized areas of sternite 9 have slightly different shape. They cluster together at p-distances >2% and only diverge at 1.6%. We have specimens sequenced from the Bukit Timah tropical forest and the Nee Soon swamp forest, as well as from the campus of the National University of Singapore.

*Neoempheria* sp. A (Figs. 57A–D)

https://singapore.biodiversity.online/species/A-Arth-Hexa-Diptera-000748

***Neoempheria_spA_ZRCBDP0048497_hapZRCBDP0048 497_SMH_unnamed_type* [41: C, 55: C, 70: A, 169: T, 1 99: T, 205: C, 217: C, 277: C]**

actttcttctacaattgctcatacaggtgcttcagttgatCtagcaattttttcCcttcatttagccggAatctcttcaattttaggggcagttaattttattactactattattaatatacgagccccaggaattcaatttgaccgaatacctttatttgtatgatcTgttttaattacagctattttattactactTtctctCcctgtattagcCggagctatcactatattattaacagatcgaaatttaaatacaagtttttttgaccctgcCggaggaggagaccctattttataccaacacttattt

#### Description

**Female** (Fig. 57A). Wing length, 2.23; width, 0.79. **Head** (Fig. 57B). Ochre-brown at vertex, cream-yellow towards frons and ventral margin of occiput. Face and clypeus cream-yellow. Antennal scape and pedicel cream-yellow, flagellum whitish-yellow. Palpomeres dark greyish-brown, last palpomere lighter. Labella whitish-yellowish. Scape as long as pedicel, flagellomere 1 1.4× flagellomere 2 length, flagellomere 4 1.0× longer than wide. Palpomere 4 1.0× palpomere 3 length, palpomere 5 1.8× palpomere 4 length. **Thorax**. Scutum ochre-yellow, more brownish towards posterior end, scutellum ochre-yellow. Pleural sclerites cream-yellow except for brown laterotergite and dark brown mediotergite on dorsal half and yellowish-brown on ventral half. Scutum with one pair of prescutellar bristles on dorsocentral lines, two pairs on lateroposterior corner; scutellum with two pairs of bristles and smaller setae spread over disc. Antepronotum with three larger bristles and one small bristle, besides smaller setae, proepisternum bare. **Legs**. Coxae whitish, anterior coxa with a yellowish tinge, femora ochre-yellow, hind femur more greyish-brown; tibiae and tarsi light greyish-brown. First tarsomeres of hind leg with a row of longer setae ventrally, all tarsomeres with a pair of distal ventral setae. Hind tibia inner spur 3.1× tibia width at apex. Fore leg tarsomere 1 0.9× tibia length, 1.6× tarsomere 2 length. **Wing** (Fig. 57C). Membrane background light greyish, a dark brown mark across most of wing at level of R_4_ to level of cubital pseudovein and a dark brown mark at distal third of wing, beginning at level of base of medial fork. C produced beyond tip of R_5_ for about a fifth of distance to tip of M_1_; Sc scarcely visible, reaching C slightly beyond level of origin of Rs; sc-r very faint, separated from tip of Sc by about length of r-m; R_4_ close to origin of Rs, anterior margin of cell r1 1.2× length of first sector of Rs; first sector of Rs 1.1× r-m length. False medial vein conspicuous, sclerotized, anterior end across r-m. M_1+2_ 3.5× r-m length; bM 5.3× length of first sector of Rs. First sector of CuA 1.3× longer than second sector. Medial fork not wide open, M_2_ short. M_4_ barely depressed on distal half. Cubital pseudovein sclerotized to level of origin of M_4_. No ventral macrotrichia on veins, dorsal macrotrichia on entire length of bR, R_1_, second sector of Rs, anterior end of r-m, distal third of M_1_, M_4_, distal half of first sector, and entire second sector of CuA; Sc, sc-r, first sector of Rs, R_4_, anterior half of r-m, bM, M_1+2_, M_2_, and basal half of first sector of CuA entirely devoid of setation; no macrotrichia on wing membrane. **Abdomen**. Tergite 1 whitish, tergites 2–6 brown, tergite 4 with cream-yellow lateroposterior corners, tergite 7 cream-yellow; tergite 7 rectangular, no digitiform lateral projections. Sternites 1–5 and 7 cream-yellow, sternites 6 cream-yellow with light brown lateral marks. **Terminalia** (Fig. 57D). Mostly cream-yellow, cerci slightly more brownish. Sternite 8 trapezoid, wide at base, medially at distal end a short medial incision separating a pair of pointed lobes, microtrichia and fine setae all over sclerite, lobes with stronger setae at tip, on inner face of lobes a group of spines directed dorsally. Sternite 9 wide at base, short, sclerotized genital chamber short, not reaching level of tip of sternite 8, genital opening largely developed, elongate, projecting well beyond tip of sternite 8 lobes, basally at each side with a lateral projection densely covered with teeth. Tergite 8 wide at anterior end, projecting ventrally at lateral ends, apparently bare and medially separated into two plates. Tergite 9 wider at posterior margin, covered with microtrichia but no setae. Tergite 10 slender, with three pairs of short digitiform projections along posterior margin, each with a long fine seta. Cercomere 1 elongate, slender, tip on dorsal face projected beyond insertion of cercomere 2, 3.0× longer than cercomere 2.

**Male**. Unknown.

#### Material examined

Female, ZRCBDP0048497, Nee Soon (NS2), swamp forest, 30.may. 12, MIP leg. (slide-mounted).

*Neoempheria sangabo* Amorim & Oliveira, sp. n. (Figs. 58A–D)

https://singapore.biodiversity.online/species/A-Arth-Hexa-Diptera-000796

urn:lsid:zoobank.org:act:A0925185-6D52-4107-B6FB-36CACBD0C6C5

**Diagnosis**. Head light brown, scutum ochre-yellowish, some areas with a light brown tinge; pleural sclerites whitish-yellow, mediotergite light brown, dorsoposterior end of laterotergite with a very light brown tinge. Tergite 1 brown, tergites 2 and 4 brown with a cream-yellow lateral band; tergites 3, 5–6 brown; tergite 7 yellowish; sternites light cream-yellowish, sternite 6 with a brownish-yellow tinge. Wing with a brown band distally across wing and a brown band more basally, from level of cell r1 to posterior margin on cell cua. Cell r1 small, anterior margin of cell r1 1.3× length of R_4_; sc-r well before level of origin of Rs; Sc reaching C before mid of cell r1. Male syngonocoxite with a medial sclerotized projection, gonocoxite with a pair of blade-like lobes extending beyond insertion of gonostylus; gonostylus large, slender basally, dorsoventrally compressed, displaced medially; tergite 9 with a pair of long lateroposterior projections, trifid distally.

***Neoempheria_sangabo_ZRCBDP0047815_hapZRCBDP 0047815_SMH_holotype* [1: A, 7: C, 127: T, 143: A, 178 : C, 184: C, 205: T, 253: T]**

AttatcCtctacaattgcccatacaggagcttctgtagatttagctattttttccttacatttagccggaatttcttcaattttaggagcagttaattttattactacaattattaatatacgagcTccaggaattcaatttAatcgaatacctttatttgtttgatctgttttaatCactgcCattttattattattatctctTccagtattagcaggagctattactatattattaacagatcgaaatctTaatacaagtttttttgatccagcaggaggaggagacccaattttatatcaacacttattt

#### Description

**Male** (Fig. 58A). Wing length, 2.30; width, 0.85. **Head**. Light-brown at vertex, lighter towards frons and ventral margin of occiput. Antennal scape and pedicel ochre-yellow, flagellum light ochre-yellow. Face light brown, clypeus yellowish-brown. Maxillary palpus light brown, last palpomere lighter; labella yellowish-brown. Setae on frons anteriorly to line of ocelli and on occiput. Scape 1.2× longer than pedicel, flagellomere 1 1.8× longer than flagellomere 4, flagellomere 4 1.1× longer than wide.

Palpomere 4 0.8× palpomere 3 length, palpomere 5 1.8× palpomere 4 length. **Thorax**. Scutum mostly ochre-yellow, light brown on anterior end, a pair of light brown bands outside dorsocentrals on posterior half of scutum; scutellum ochre-yellow. Scutum with one pair of long prescutellar setae at dorsocentral line and one pair of small bristles on lateroposterior corners. Pleural sclerites whitish with an orangish tinge, antepronotum light brownish medially, laterotergite with a diffuse brown area on dorsoposterior end, mediotergite light brown. Antepronotum with two bristles and additional smaller setae. **Legs**. Coxae whitish, mid and hind coxae with a light brown ring at tip; femora ochre-yellow, hind femur slightly darker; tibiae and tarsi light brown. Mid tibial inner spur 3.3× longer than tibia width at apex [both hind leg femora, tibiae and tarsi missing]. Fore leg tarsomere 1 0.8× tibia length, 1.8× tarsomere 2 length. **Wing** (Fig. 58B). Membrane background light greyish, a dark brown band across wing from tip of Sc to base of second sector of CuA and a dark brown mark at distal third of wing, beginning at level of tip of R_1_. C produced beyond tip of R_5_ for about a third of distance to M_1_; Sc complete, reaching C at level of origin of Rs, sc-r not produced; first sector of Rs oblique, R_4_ not far from base of r-m, also oblique, anterior margin of cell r1 1.3× length of first sector of Rs; first sector of Rs 1.0× r-m length. False medial vein conspicuous, sclerotized. M_1+2_ 4.5× r-m length; bM 6.5× length of first sector of Rs. First sector of CuA 1.5× longer than second sector. M_4_ with a gentle depression on distal half. Cubital pseudovein sclerotized to level of origin of M_4_, CuP barely sclerotized, close to wing base. Wing margin not emarginated at tip of CuA. No ventral macrotrichia on veins, dorsal macrotrichia on entire length of bR, R_1_, second sector of Rs, and distal fifth of M_1_ and M_2_; Sc, sc-r, first sector of Rs, R_4_, r-m, bM, M_4_, CuA and CuP devoid of macrotrichia. **Abdomen**. Tergite 1 light brown, tergites 2–4 light brown medially with cream-yellow lateral bands, slender on tergites 3–4, tergites 5–6 light brown with a slender ochre-yellow band along anterior margin, tergite 7 ochre-yellowish. Sternites 1–7 cream-yellow, sternites 5–6 with a brownish tinge. **Terminalia** (Figs. 58C–D). Cream-yellow, gonocoxal lateroposterior extensions light brownish. Gonocoxites fused medially, bare, syngonocoxite with a pair of lateral rounded lobes extending to level of middle of gonostylus. Gonostylus large, slender basally, dorsoventrally compressed, insertion displaced slightly towards middle, a short rounded lobe on dorsal face with two long distal setae, ventral face and margin covered with microtrichia and fine, elongate setae. Gonocoxal bridge wide, sub-medial apodeme directed anteriorly. Aedeagus and parameres hard to identify, weakly sclerotized. Tergite 9 with a slender medial stripe connecting a pair of long lateroposterior projections, with fine setae along external margin, distally trifid, outer projection shorter, ventral projection long and curved, dorsal projection digitiform, setose. Tergite 10 large, bare medially, extending lateroposteriorly into a pair of setose lobes. Cerci weakly sclerotized, lobose, covered with microtrichia and fine setae.

**Female**. Unknown.

#### Material examined

**Holotype**: male, ZRCBDP0047815, Nee Soon (NS1), swamp forest, 18.dec.13, MIP leg. (slide-mounted).

**Etymology**. The specific epithet of this species refers to the Cantonese dialect transcription of Singapore, Xin Jia Po (新加坡). Cantonese is one of the largest Chinese dialect groups in Singapore. The noun is used in apposition.

**Remarks**. This is a singleton species and the holotype was collected in the swamp forest.

*Neoempheria shicheng* Amorim & Oliveira, sp. n. (Figs. 59A–D)

https://singapore.biodiversity.online/species/A-Arth-Hexa-Diptera-000796

urn:lsid:zoobank.org:act:D9AF8FD3-05A9-4AA5-9FB4-81BD9A75F0CE

**Diagnosis**. Head, scutum and scutellum light brown, yellowish above wing; pleural sclerites whitish, mediotergite light brown, dorsoposterior end of laterotergite light brown. Tergite 1 light brown, tergites 2 and 4 with a brown medial longitudinal band, ochre-yellow laterally; tergites 3, 5–6 brown, with a slender yellow band along anterior margin; tergite 7 yellowish; sternites cream-yellowish, sternite 6–7 with brown lateral marks. Wing with a brown band distally across wing and a brown band more basally, from level of cell r1 to posterior margin on cell cua. Cell r1 small, anterior margin of cell r1 1.6× length of R_4_; sc-r well before level of origin of Rs; Sc reaching C before mid of cell r1. Male syngonocoxite with medio-posterior area strongly sclerotized; gonostylus dorsoventrally compressed; tergite 9 with a pair of long lateroposterior bifid projections, more ventral one extending beyond tip of cerci, a short digitiform lobe distally projected backwards bearing a long fine seta at tip. ***Neoempheria_shicheng_ZRCBDP0047883_hapZRCBD P0047883_SMH_holotype* [1: T, 137: C, 187: A, 194: C, 196: T, 202: C, 229: C, 250: C, 251: C, 262: T]**

TctatcttctacaattgctcatacaggagcttctgttgatttagctattttttctttacatttagctggtatttcttctattttaggagctgttaattttattacaacaattattaatatacgagcacctggtattCaattcgatcgaatacctttatttgtttgatctgttttaattacagccgtAttattaCtTttatcCttaccagtattagcaggagctattacCatattattaacagatcgaaaCCttaatacaagTttctttgatcctgcaggaggaggagatcctattttatatcaacatttattt

#### Description

**Male** (Fig. 59A). Wing length, 2.26; width, 0.85. **Head**. Light-brown at vertex, ochre-yellowish above antennae and on ventral half of occiput. Face whitish-yellow, clypeus ochre with dark brown mark laterally. Antennal scape and pedicel light ochre-yellow, flagellum dark ochre-yellow. Maxillary palpus light brown, last palpomere lighter. Labella light brown. Scape 1.0× pedicel length; flagellomere 1.7× longer than flagellomere 2, flagellomere 4 1.4× longer than wide. Palpomere 4 0.81× palpomere 3 length, palpomere 5 1.6× palpomere 4 length. **Thorax**. Scutum mostly light brown, an ochre-yellowish band laterally on posterior half of scutum, medially brown; scutellum ochre-yellowish. Pleural sclerites whitish with an orangish tinge, antepronotum with a brown mark anteriorly, laterotergite with a light brown area at dorsoposterior end, mediotergite brown. Scutum with a pair of prescutellar bristles medially on dorsocentral line and a pair of prescutellar bristles on lateroposterior corner, scutellum with one pair of bristles and one additional pair of small setae. Antepronotum with two bristles and additional small setae, proepisternum with one small bristle and some few setae. **Legs**. Coxae whitish; femora light ochre-yellow; tibiae and tarsi light brown. Mid tibia inner spur 3.1× tibia width at apex [fore and hind femora, tibiae and tarsi missing]. **Wing** (Fig. 59B). Membrane fumose light brown, a dark brown band across wing from tip of Sc to base of second sector of CuA and a dark brown mark at tip of wing beginning at level of tip of R_1_ to M_2_. C extending beyond tip of R_5_ for a third of distance to M_1_. Sc complete, reaching C slightly beyond level of origin of Rs, sc-r barely sclerotized; first sector of Rs oblique, R_4_ not far from base of r-m, cell r1 short, anterior margin 1.6× length of R_4_; first sector of Rs 1.1× r-m length. False medial vein conspicuous, sclerotized. M_1+2_ 4.6× r-m length; bM 6.5× length of first sector of Rs. First sector of CuA 1.3× longer than second sector. M_4_ only gently depressed on distal half. Cubital pseudovein sclerotized to level of origin of M_4_, CuP sclerotized only close to wing base. Wing margin gently emarginated at tip of CuA. No ventral macrotrichia on veins, dorsal macrotrichia on entire length of bR, R_1_, second sector of Rs, and distal fourth of M_1_; Sc, sc-r, first sector of Rs, R_4_, r-m, bM, M_2_, M_4_, CuA, and CuP devoid of macrotrichia. **Abdomen**. Tergite 1 brown, slender cream-yellow band laterally; tergites 2 and 4 brown medially with wide cream-yellow lateral bands, tergites 3, 5 and 6 brown, tergite 6 with a slender yellowish band along posterior margin, tergite 7 ochre-yellowish. Sternites 1–4 whitish-yellow, sternites 5–6 light brown, with a yellowish band along posterior margin, sternite 7 ochre-yellowish. **Terminalia** (Fig. 59C). Ochre-yellow, with some more brownish sclerites. Gonocoxites fused medially, no suture of fusion, entirely bare, syngonocoxite with a strongly sclerotized medio-posterior projection bearing a pair of lateral arms. Gonostylus insertion displaced medially, dorsoventrally compressed, with microtrichia and long fine setae on ventral face. Gonocoxal bridge anteriorly with a pair of large, triangular apodemes. Aedeagus not distinguishable, no distal tubular extension. Parameres with two pairs of digitiform projections laterally on anterior half, each with some elongate setae distally, posteriorly with a wide rounded projection, a median sclerotized line. Tergite 9 with a slender medial connection between a pair of long lateroposterior projections extending beyond tip of cerci, covered on external face with elongate fine setae, distally bifid, dorsal branch shorter, ventral branch elongate, curved inwards, distal end directed anteriorly with a seta at tip. Cerci in a quite posterior position, elongate, close to each other along medial line, densely covered with microtrichia and short setae.

**Female**. As male, except for the following. **Wing l**ength, 2.10; width, 0.79. **Terminalia** (Fig. 59D). Sternite 8 trapezoid, elongate, slender posteriorly, a medial incision separating two distal lobes entirely covered by microtrichia, fine setae on posterior third, lobes densely setose distally, a short projection with a distal seta on posterior third of lateral margin. Sternite 9 wide, no anterior medial apodeme, an incision separating a pair of lobes, two gonoducts. Sternite 10 with a large, transparent area between lateral bands with microtrichia and setae. Tergite 9 large, reaching base of cercomere 1, devoid of setae, only with microtrichia, a pair of sub-medial projections, each distally with three digitiform short extensions with a terminal seta. Tergite 10 slender, with a pair of lateral digitiform projections with a terminal seta. Cercomere 1 with a wide base, tapering distally, covered with microtrichia and fine setae, cercomere 2 elongate, about 0.5× length of cercomere 1.

#### Material examined

**Holotype**: male, ZRCBDP0047883, Nee Soon (NS2), swamp forest, 23.oct.13, MIP leg. (slide-mounted). **Paratypes** (2 females): ZRCBDP0047906, Nee Soon (NS2), swamp forest, 30.oct.13, MIP leg. (slide-mounted); ZRCBDP0048257, Sungei Buloh (SB1), mangrove, 09.oct.13, MIP leg.

**Etymology**. The specific epithet of this species refers to Shi Cheng (獅城), a Mandarin local colloquial nickname for Singapore, meaning ‘Lion City’. The noun is used in apposition.

**Remarks**. We have two haplotypes for this species and no delimitation conflicts.

*Neoempheria ujong* Amorim & Oliveira, sp. n. (Figs. 60A–D)

https://singapore.biodiversity.online/species/A-Arth-Hexa-Diptera-000797

urn:lsid:zoobank.org:act:01B2ADF3-2413-4375-A5B4-66AF1FF10F05

**Diagnosis**. Head and scutum ochre-yellowish, scutellum light brown; pleural sclerites whitish-yellow, mediotergite light brown, dorsoposterior end of laterotergite with a light brown tinge. Wing with a brown band across distal third of wing and a brown band more basally, from level of cell r1 to posterior margin on cell cua. Cell r1 small, anterior margin of cell r1 1.4× length of R_4_; sc-r slightly before level of origin of Rs; Sc reaching C before mid of cell r1. Tergite 1 brown, tergites 2–4 brown with a cream-yellow lateral band; tergites 5–6 brown; tergite 7 yellowish; sternites cream-yellowish, sternite 6 brownish-yellow. Gonocoxites entirely bare, with a short rounded lobe extending to level of basal third of gonostylus and a medio-posterior extension; gonostylus displaced medially, large, obovoid, dorsoventrally compressed; parameres with a pair of digitiform lateral extensions posteriorly with some 3–4 fine setae at distal end. Tergite 9 with a pair of large bifid lobes, separated midway to apex, dorsal branch with tip curved inward.

***Neoempheria_ujong_ZRCBDP0048904_hapZRCBDP00 48904_SMH_holotype* [41: C, 43: T, 73: C, 199: C, 205: G, 212: C, 247: T, 251: C, 253: T] -**

ttatcttctacaattgctcatacaggggcttctgtagatCtTgctattttttcattacatttagccggaatCtcttctattttaggagctgttaattttattactactattattaatatacgagcaccaggaattcaatttgatcgaatacctttatttgtttgatctgttttaatcacagcagtattattattactCtcattGccagttCtagcaggagctattactatactactaacagatcgTaatCtTaatacaagtttttttgatcctgctggaggaggagatcctattttataccaacatttattt

#### Description

**Male** (Fig. 60A). Wing length, 2.10, width, 0.79. **Head**. Light ochre-brown. Antennal scape and pedicel whitish, flagellum whitish ochre-yellow. Face light brown, clypeus light brown. Maxillary palpus greyish-brown, last palpomere lighter, labella light brown. Two ocelli placed medially on vertex. Frons covered with microtrichia and setae. Scape 1.0× pedicel length, flagellomere 1 1.9× flagellomere 2 length, flagellomere 4 1.3× longer than wide. Palpomere 4 1.1× palpomere 3 length, palpomere 5 1.5× palpomere 4 length. **Thorax**. Scutum ochre-yellowish, slightly darker at anterior and posterior ends, scutellum light brown. Pleural sclerites light cream-yellow, a brownish mark on antepronotum lobe anteriorly, laterotergite light brown at dorsoposterior end, mediotergite brown, lighter laterally. Halter light brown. Scutum with a prescutellar bristle medially on dorsocentral line, two pairs of prescutellar bristles on lateroposterior corners; scutellum with one pair of bristles, no additional small setae. Antepronotum with two bristles and some small setae, proepisternum with some small setae. **Legs**. Coxae whitish, femora ochre-yellow, hind femur darker, tibiae and tarsi light greyish-brown, light brown towards tip of tarsi. Fore leg tarsomere 1 0.90× tibia length, 1.9× tarsomere 2 length. Hind tibia inner spur 2.9× tibia width at apex. **Wing** (Fig. 60B). Membrane with a brown band across wing at level of cell r1 and another band at distal third of wing beginning slightly beyond basal end of medial fork. C extending beyond tip of R_5_ for a ¼ of distance to M_1_; Sc reaching C slightly beyond origin of Rs, sc-r hardly visible (even under phase contrast), R_4_ present, inclined, cell r1 short, trapezoid, anterior margin of cell r1 1.4× longer than length of R_4_; first sector of Rs as long as r-m. False medial vein present, sclerotized, slightly curved. M_1+2_ 3.5× r-m length; bM 5.6× length of first sector of Rs. First sector of CuA 1.8× longer than second sector. M_4_ gently depressed on distal half. Cubital pseudovein sclerotized barely beyond level of origin of M_4_; CuP sclerotized only at very base of wing. No sign of anal fold. Wing margin gently emarginated at tip of CuA. No ventral macrotrichia on veins, dorsal macrotrichia on entire length of bR, R_1_, second sector of Rs, and distal fourth of M_1_; Sc, sc-r, first sector of Rs, R_4_, r-m, bM, M_2_, M_4_, and CuP entirely devoid of macrotrichia. **Abdomen**. Tergites 1, 5–6 light brown, tergites 2–4 light brown with a wide cream-yellow mark laterally, tergite 7 brownish ochre-yellow. Sternites 1–3 whitish, sternites 4–5 cream-yellow, sternites 6–7 ochre-yellow with light brownish tinge. **Terminalia** (Figs. 60C–D). Light ochre-yellow. Gonocoxites fused, suture present medially on anterior half of syngonocoxite, entirely bare, laterally a short rounded posterior lobe, extending to level of basal third of gonostylus, medially extending into aedeagal sclerite. Gonostylus displaced medially, large, obovoid, compressed dorsoventrally, slightly pointed distally, covered with scattered microtrichia and fine setae on ventral face, a concentrated row of long setae along distal half of internal margin and some long setae on external margin on basal half. Gonocoxal bridge weakly sclerotized. Aedeagus with a strongly sclerotized sclerite medially, with a short anterior extension and a pair of lateral extensions and a distal extension. Parameres wide, a pair of digitiform lateral extensions posteriorly with some 3–4 fine setae at distal end. Tergite 9 with a pair of large lobes connected medially through a slender, weakly sclerotized band, each lobe bifid, diverging midway to apex, dorsal branch an inverted L-shape with tip curved inward, more sclerotized, with some few fine setae, ventral branch longer, reaching level of tip of gonostylus, with a digitiform dorsal extension bearing a seta at tip directed inwards and a digitiform ventral extension with some setae at distal end. Sternite 10 wide, with a pair of short lobes laterally on distal margin bearing microtrichia and setae on both, ventral and dorsal faces. Tergite 10 membranous. Cerci weakly sclerotized, slightly elongate, with microtrichia and short setae

**Female**. Wing length, 2.07; width, 0.79. **Terminalia**. Sternite 8 anterior band weak, with lateroanterior extensions directed dorsoanteriorly articulating with lateroanterior arms of tergite 8, a subquadrate medial sclerite distally with a median short medial incision, covered with microtrichia, setae on distal third, especially along posterior margin. Sternite 9 wide, with a pair of plates laterally to genital chamber. Tergite 8 wide, bare. Tergite 9+10 rectangular, posterior margin with three pairs of elongate digitiform projections each with a seta at tip, median pair close to middle, lateral two pairs extending laterally to cerci. Sternite 10 slender, with median transparent window, with microtrichia and setae along posterior margin. Cercomere 1 large, entirely covered with microtrichia, setae restricted to distal third of inner border as a regular row of 9–10 elongate setae directed inwards, cercomere 2 slender basally, rounded distally.

#### Material examined

**Holotype**: male, ZRCBDP0048904, Nee Soon (NS1), 31.dec.14, MIP leg. (slide-mounted). **Paratypes** (1 male, 2 females). **Male**: ZRCBDP0049228, Nee Soon (NS1), 10.dec.14, MIP leg. **Females**: ZRCBDP0048978, Nee Soon (NS1), 13.may.15, MIP leg. (slide-mounted); ZRCBDP0049258, Nee Soon (NS1), 10.dec.14, MIP leg. **Additional sequenced specimens** (2 males): ZRCBDP0072698, Bukit Timah (BT05), primary forest, 01.dec. 16, MIP leg.; ZRCBDP0136953, Bukit Timah (BT08), maturing secondary forest, 09.may. 17, MIP leg.

**Etymology**. The species epithet refers to Ujong (in Malay, “at the end”), used in Pulau Ujong. It is the indigenous Malay name for pre-colonial Singapore, meaning the island at the end [of the Malay Peninsula]. The noun is used in apposition.

**Remarks**. This species includes specimens from the swamp forest and from the Bukit Timah tropical forest. There are two haplotypes and there are no conflicts between the delimitation approaches, despite considerable morphological variation on the abdominal tergite color between males and females.

*Neoempheria subaraji* Amorim & Oliveira, sp. n. (Figs. 61A–D)

https://singapore.biodiversity.online/species/A-Arth-Hexa-Diptera-000774

urn:lsid:zoobank.org:act:1B452FA3-FE3A-4EE2-B73B-9B2F579AFB98

**Diagnosis**. Head, scutum and scutellum dark brown; pleural sclerites whitish, mediotergite dark brown, laterotergite with a light brown tinge at dorsoposterior end. Wing with a brown band on distal third of wing and a brown band more basally, from level of cell r1 to slightly over CuA. Cell r1 small, R_4_ originating at level of insertion of r-m, cell r1 triangular, anterior margin 0.83× length of R_4_; sc-r reaching bR well basal to origin of Rs. Tergites 1–2 and 4 largely dark brown, with yellow marks laterally, larger on tergite 4; tergites 3, 5–6 dark brown with a slender yellow band along anterior margin; tergite 7 yellowish; sternites 1–5 whitish, sternites 6–7 with brown lateral marks. Gonocoxites bare, no lateroposterior projection beyond base of gonostylus, no medio-posterior process; gonostylus long, slender at base, dorsoventrally compressed; tergite 9 with a slender, bare band connecting a pair of long lateral projections gently curved inwards, with a group of 6–7 elongate spines.

***Neoempheria_subaraji_ZRCBDP0049198_hapZRCBDP 0049198_SMH_holotype* [4: C, 19: G, 25: G, 73: C, 94: C, 130: A, 238: G]**

actCtcatctactattgcGcatacGggagcttcagttgatttagctattttttcattacatttagctgggatCtcttcaattttaggggcagtCaattttattacaacaattattaatatacgtgccccAggaattcaatttgatcgaatacctttatttgtatgatctgttttaattacagctgttttattattactttcattacctgtattagccggagctattactatactattGacagatcgaaatctaaatacaagcttttttgaccctgctggaggaggagaccctattttatatcaacacttattt

#### Description

**Male** (Fig. 61A). Wing length, 2.33; width, 0.85. **Head**. Vertex, face, and clypeus brown. Antennal scape and pedicel ochre-yellowish, flagellum light ochre-yellow, brownish towards apex. Maxillary palpus brown, last palpomere lighter, labella brown. Ocellar setae present, well-developed, frons with setae. Scape 1.1× pedicel length, flagellomere 1 1.6× flagellomere 2 length, flagellomere 4 1.2× longer than wide. Palpomere 4 as long as palpomere 3, palpomere 5 1.4× palpomere 4 length. **Thorax**. Scutum dark ochre-yellow, brown on posterior half; scutellum brown. Pleural sclerites whitish except for a light brown mark on dorsoposterior end of laterotergite, and brown mediotergite. Halter light brown, knob darker. Scutum with a pair of prescutellar bristles medially on dorsocentral lines and one pair of prescutellar bristles on lateroposterior corner; scutellum with one pair of bristles, no additional small setae. Antepronotum with three bristles and some additional larger and smaller setae. **Legs**. Coxae whitish, hind coxa dark at tip; femora ochre-yellow, hind femur with a brown diffuse mark internally at basal half; tibia and tarsus light greyish-brown. Tarsomere 1 of fore leg 0.58× tibia length, 1.9× tarsomere 2 length. Hind tibia inner spur 2.9× longer than tibia width at apex. **Wing** (Fig. 61B). Membrane with a dark brown band across wing at level of tip of Sc and a wide band on distal third of wing, beginning slightly beyond basal end of medial fork. C extending beyond tip of R_5_ for a ¼ of distance to M_1_; Sc complete, reaching C at level of origin of Rs, sc-r basally to origin of Rs for a distance smaller than length of anterior margin of cell r1, R_4_ present, more of less transverse, originating at level of r-m; anterior margin of cell r1 short, 1.1× length of R_4_; first sector of Rs 1.4× r-m length. False medial vein present, slightly arched, not strongly sclerotized. M_1+2_ 5.2× r-m length; bM 7.5× length of first sector of Rs. First sector of CuA 1.5× longer than second sector. M_4_ not strongly diverging from CuA, straight on distal half. Cubital pseudovein weakly sclerotized, ending before level of origin of M_4_; CuP not sclerotized. Anal fold faint. Wing margin gently emarginated at tip of CuA. No ventral macrotrichia on veins, dorsal macrotrichia on entire length of bR, R_1_, and second sector of Rs, distal fourth of M_1_, and tip of M_2_. Sc, sc-r, first sector of Rs, R_4_, r-m, bM, M_4_, CuA, and CuP entirely devoid of macrotrichia. **Abdomen**. Tergite 1 dark brown medially with whitish lateral bands, tergites 2 and 4 with a dark brown medial band, slender medially at posterior margin and yellowish lateral bands, tergites 3, 5– 6 dark brown with a slender yellowish band along anterior margin, tergite 3 with additional small diffuse yellowish marks, tergite 7 yellow. Sternites 1–2 whitish, sternites 3–7 ochre-yellow. **Terminalia** (Figs. 61C–D). Brownish-yellow. Gonocoxites fused medially, bare, no medial suture, no lateroposterior projection beyond base of gonostylus, no medio-posterior process. Gonostylus long, slender at base, dorsoventrally compressed, with setae on both faces. Gonocoxal bridge with a pair of wide anterolateral apodemes. Aedeagal plate wide, with a median suture, gently rounded on posterior margin, not reaching level of tip of cerci. Parameres wide, posterior margin at level of tip of tergite 9, weakly sclerotized, a pair of digitiform projections laterally with 1–2 elongate setae at tip, reaching level of mid of gonostylus. Tergite 9 with a slender, bare band connecting a pair of long lateral projections almost reaching level of tip of gonostylus, gently curved inwards on distal half, with setae on lateral face and a group of 6–7 elongate spines at distal end. Sternite 10 trapezoid, with more or less rounded posterior margin, reaching beyond tip of aedeagus, with microtrichia and fine setae. Cerci close to each other medially, slightly elongate, with microtrichia and fine setae.

**Female**. Unknown.

#### Material examined

**Holotype**: male, ZRCBDP0049198, Nee Soon (NS2), 13.may.15, MIP leg. (slide-mounted).

**Etymology**. The species epithet honors Subaraj Rajathurai (1963-2019), veteran wildlife consultant and conservationist, who had a key role in drafting the masterplan for nature conservation in Singapore.

*Neoempheria kokoiyeeae* Amorim & Oliveira, sp. n. (Figs. 62A–D)

https://singapore.biodiversity.online/species/A-Arth-Hexa-Diptera-000770

urn:lsid:zoobank.org:act:081DE055-CAFE-437E-AF68-7DAEFA286A6A

**Diagnosis**. Head ochre-yellowish with brownish tinge dorsally, scutum dark ochre-yellowish, scutellum light brown; pleural sclerites whitish-yellow, mediotergite brown, laterotergite dorsoposterior end light brown. Wing with a brown band on distal third across wing and a brown band more basally, from level of cell r1 to posterior margin on cell cua. Cell r1 small, anterior margin of cell r1 1.2× length of R_4_; sc-r not sclerotized; Sc reaching C at level of origin of Rs. Tergite 1 dark brown medially, with large cream-yellow lateral band, tergite 2 dark brown, with large cream-yellow lateroanterior marks, tergites 3–6 dark brown, tergite 7 ochre-yellowish with a brown band along posterior margin. Male syngonocoxite with a short medio-posterior process, gonocoxite with a lateroventral projection extending beyond tip of gonostylus and a lateroposterior projection; gonostylus short, digitiform; tergite 9 with a pair of long projections with a hook-shaped distal end.

***Neoempheria_kokoiyeeae_ZRCBDP0048489_hapZRCB DP0048489_SMH_holotype* [22: C, 28: T, 133: C, 184: C, 187: A, 188: C, 261: G, 277: C]**

gctatcttctacaattgctcaCacaggTgcatctgtagatttagctattttttccctacatttagccggaatttcttctattttaggggctgttaattttattacaactattattaacatacgagccccaggCattcaatttgatcgaatacctttatttgtatgatccgttttaattacagcCgtACtattattattatcattaccagtattagctggagcaattactatattattaacagatcgaaacttaaatacaaGcttttttgacccagcCggaggaggagatcccattctataccaacatttattt

#### Description

**Male**. Wing length, 2.49; width, 0.95. **Head**. Light ochre-yellowish with a brown tinge, lighter towards frons and ventral half of occiput. Some few scattered short setae over frons and occiput. Ocellar setae present. Face light brown. Antennal scape and pedicel whitish-yellow, scape 1.3× longer than pedicel, flagellum light ochre-yellow except for flagellomere 1 cream-yellow, flagellomere 1 1.9× longer than flagellomere 2, flagellomere 4 1.1× longer than wide. Clypeus yellowish-brown, triangular, with scattered setae. Palpomeres brown, last palpomere lighter; palpomere 4 about as long as palpomere 3, a short distal projection over base of distal palpomere, palpomere 1.4× palpomere 4 length. Labella small, brownish. **Thorax**. Scutum ochre-yellowish; a row of longer dorsocentral setae, some long supra-alars, two pairs of pre-scutellar bristles on lateroposterior corners of scutum, one pair of prescutellars medially, on dorsocentral line. Scutellum light brown, with one pair of longer setae, no smaller setae on disc. Pleural sclerites whitish with an orangish tinge, antepronotum with a brown mark anteriorly, laterotergite with a light brown mark at dorsal margin, mediotergite brown. Antepronotum with one strong bristle and smaller setae of different sizes. Halter light brown. **Legs**. Coxae whitish, mid and hind coxae with a light brown ring at tip; femora ochre-yellow, fore coxa lighter; tibiae and tarsi light greyish-brown. **Wing** (Fig. 62B). Membrane background light greyish, a dark brown band across wing at level of origin of Rs and a dark brown band across distal third of wing beginning slightly beyond level of base of medial fork. Sc complete, reaching C at level of origin of Rs, sc-r not distinguishable; cell r1 short, anterior margin 1.2× length of first sector of Rs. False medial vein conspicuous, sclerotized, with a bend on basal third. Medial fork wide open, M_2_ short. M_1+2_ 4.0× r-m length; bM 6.4× length of first sector of Rs. First sector of CuA 1.8× longer than second sector. M_4_ gently depressed on distal half. Cubital pseudovein sclerotized, ending at first third of second sector of CuA; CuP short, limited to level of first third of first sector of CuA. Anal fold faint. Wing margin slightly emarginated at tip of CuA. No ventral macrotrichia on veins, dorsal macrotrichia on entire length of bR, R_1_, second sector of Rs, and on distal third of M_1_. **Abdomen**. Tergites 1–2 and 4 cream-yellow, with a medial brown band, tergite 3 and 5–6 brown, tergite 7 ochre-yellowish. Sternites 1–6 cream-yellow, sternite 5–6 with a slender lateral light brown band, sternite 7 ochre-yellow. **Terminalia** (Figs. 62C–D). Ochre-yellow, cerci lighter. Gonocoxites fused medially, a short medio-posterior sclerotized process, a pair of lateroventral projections extending beyond tip of gonostylus, no setae on ventral face, lateroposterior projections with microtrichia and elongate setae. Gonostylus short, digitiform, mostly bare, except for four fine setae distally. Aedeagus subquadrate, wide, posterior margin reaching almost level of tip of gonocoxite lobes. Parameres weakly sclerotized. Tergite 9 with a slender anterior blade medially connecting a pair of long projections bearing a hook-shaped distal curve. Sternite 10 weakly sclerotized, subquadrate, with a pair of short laterodistal lobes, covered with microtrichia and fine setae. Cerci small, close together, weakly sclerotized, with microtrichia and fine setae.

**Female** (Fig. 62A). As male, except for the following. **Wing l**ength, 2.62; width, 0.98. **Terminalia**. Ochre-yellowish, cerci lighter. Sternite 8 wide anteriorly, with a pair of weakly sclerotized distal lobes with a medial incision. Sternite 9 wide, complex, without a furca at anterior end. Tergite 8 wide, slightly projected medially covered with microtrichia, but no setae. Tergite 9 wide, short, with a row of elongate setae. Tergite 10 slender, a group of short protuberances each with a seta at tip, a pair of digitiform lobes, inner pair each with two setae, outer pair each with a single seta. Sternite 10 with a pair of short lateral lobes on posterior margin, transparent window wide midway to apex. Cerci laterally compressed, cercomere 1 1.4× longer than cercomere 2, both covered with microtrichia and short setae, distal cercomere clavate.

#### Material examined

**Holotype**: male, ZRCBDP0048489, Nee Soon (NS2), swamp forest, 04.apr. 12, MIP leg. (slide-mounted). **Paratypes** (2 females): ZRCBDP0049205, Nee Soon (NS1), 10.dec.14, MIP leg. (website photo specimen, slide-mounted); ZRCBDP0049263, Nee Soon (NS1), 10.dec.14, MIP leg.

**Etymology**. The species epithet honors Mdm. Kok Oi Yee (1942-), a veteran naturalist who was one of Singapore’s first women nature guides and first female biology lab technician in the National University of Singapore. She is also an Honorary Museum Associate of the Lee Kong Chian Natural History Museum.

**Remarks**. This species has specimens known from both, Bukit Timah and Nee Soon forests. We have two haplotypes for this species and no delimitation conflicts.

*Neoempheria* sp. G (Figs. 63A–D)

***Neoempheria_spG_ZRCBDP0137178_hapZRCBDP0137 178_SMH_unnamed_type* [7: A, 46: C, 55: C, 130: G, 13 3: T, 137: C, 283: T, 295: C] --**

tatcAtctacaattgctcatacaggagcttctgtagatttagcCattttttcCttacatttagccggaatttcttctattttaggagctgttaattttattacaactattattaatatacgagctccGggTattCaatttgatcgaatacctttatttgtttgatctgttttaattacagctgtattattattattatcattaccagtattagctggagctattactatattattaacagatcgaaatttaaatacaagtttttttgacccagcaggaggTggagaccctatCttatatcaacacttattt

#### Description

**Female**. Wing length, 2.36; width, 0.72. **Head** (Fig. 63A). Brown, lighter on occiput, face and clypeus light brown. Face and clypeus with scattered small setae and microtrichia, some setae laterally on frons anteriorly to ocelli. Post-ocellar setae present. Antennal scape and pedicel yellowish-brown, scape as long as pedicel, flagellum ochre-yellow, flagellomere 1 2.0× longer than flagellomere 2, flagellomere 4 1.3× longer than wide. Palpomeres dark brown, last palpomere lighter, palpomere about 0.85× palpomere 3 length, with a short distal projection over base of palpomere 5, distal palpomere 2.0× palpomere 4 length. Labella small, brownish. **Thorax**. Scutum homogenously brown; a row of longer dorsocentral setae, some long supra-alars, two pairs of pre-scutellar bristles on lateroposterior corners of scutum, one pair of prescutellars medially on dorsocentral lines. Scutellum brown, with one pair of longer setae, no smaller setae on disc. Antepronotum, proepisternum, proepimeron, anepisternum, katepisternum, mesepimeron, and metepisternum whitish, katepisternum, mesepimeron, laterotergite, and mediotergite light brown. Antepronotum with two stronger bristles and smaller setae of different sizes. Halter yellowish-brown. **Legs**. Coxa whitish, mid and hind coxae with a brownish mark distally; anterior and mid femora ochre-yellowish, hind femur brownish-yellow; tibiae and tarsi yellowish-brown, anterior tarsus lighter. Hind tibial spurs 2.7× length of tibia at apex. **Wing** (Figs. 63B–C). Membrane with light greyish-brown background, a brown band across wing at level of tip of Sc and a large brown mark at distal third of wing, beginning at level of medial fork. C extending beyond tip of R_5_ for a fifth of distance to M_1_. Sc complete, reaching C slightly beyond level of origin of Rs; sc-r present, reaching bR slightly more basally than level of origin of Rs; cell r1 short, anterior margin 1.4× length of R_4_. False medial vein present, not strongly sclerotized, gently sinuous on basal fourth. Medial fork not wide open. M_1+2_ 4.9× r-m length; bM 6.8× r-m length. First sector of CuA 1.6× longer than second sector. M_4_ very gently depressed on distal half. Cubital pseudovein weakly sclerotized, not even reaching level of origin of M_4_; CuP barely sclerotized. Anal fold faint. Wing margin slightly emarginated at tip of CuA. No ventral macrotrichia on veins, dorsal macrotrichia on entire length of bR, R_1_, second sector of Rs, and distal fifth of M_1_. **Abdomen**. Tergites 1–7 light brown. Sternites 1–4 and 7 ochre-yellow, sternites 5–6 light brown. **Terminalia** (Fig. 63D). Ochre-yellow. Sternite 8 wide, with a pair of triangular distal lobes separated by a wide V-shaped incision, a pair of smaller sub-lobes on external margin of lobes, sclerite mostly bare on anterior two-thirds, microtrichia and setation on posterior lobes, distal setae stronger. Sternite 9 wide, with median projection at anterior end. Tergite 8 rectangular, bare. Tergite 9+10 slender, with three pairs of long digitiform projections, each with a distal long seta, inner pair with a second, subdistal seta. Sternite 10 with a transparent window midway to apex, a pair of short digitiform lateral projections more basally, with two distal long setae and a pair of flat distal lateral lobes. Cercomere 1 wide at basal half, 2.0× longer than cercomere 2, both covered with microtrichia and short setae.

**Male**. Unknown.

**Material examined**: female, ZRCBDP0137178, Bukit Timah Forest (BT06), 09.may. 17, MIP leg. (partially broken, slide-mounted).

### Group ferruginea

This is possibly the best known group in the genus and includes all species with large cell r4 and the typical five-banded pattern of the scutum. There is more than one male terminalia pattern among the species. As mentioned above, there are many formal species groups with large cell r4 in Edwards’ (1940) study of the Neotropical species of the *Neoempheria*. A wide study of *Neoempheria* worldwide would allow to correlate the diversity of the genus in the Oriental and the Holarctic regions with the Neotropical species groups.

*Neoempheria mandai* Amorim & Oliveira, sp. n. (Figs. 64A–D)

https://singapore.biodiversity.online/species/A-Arth-Hexa-Diptera-00771,%20and%20-000740

urn:lsid:zoobank.org:act:0BFB28BB-372C-4FD7-8A21-8BDC36081F86

**Diagnosis**. Scutum with five longitudinal brown stripes, a greyish brown line over dorsal half of mediotergite extending over dorsal third of laterotergite. Wing with a brown mark across distal third of wing, with marks over sc-r, first section of Rs, R_4_, and basally on cell br. Vein sc-r at basal third of cell r1; anterior margin of cell r1 about 5× length of R_4_. Abdominal tergites and sternites ochre-yellow, dark brown marks over most of tergite 3 and 5, as well as medially and along posterior margin on tergites 1, 2, 4, 6 and 7. Gonostylus large, with a row of strong spines along distal third of inner margin directed inwards; parameres with a pair of small lobes projecting beyond tip of gonostylus with a group of short spines directed inwards.

***Neoempheria_mandai_ZRCBDP0048968_hapZRCBDP0 048968_SMH_holotype* [1: T, 10: A, 62: C, 64: T, 67: T, 70: T, 79: T, 106: T, 142: T, 169: A, 190: T, 262: A]**

TttatcatcAacaattgctcatacaggagcttctgtagatttagctatcttttctttacacCtTgcTggTatttcttcTattttaggagctgtaaattttattacTacaattattaatatacgtgctccaggaattcaattTgatcgaatacctttatttgtatgatcAgttttaattacagctattctTttattattatctttacctgttttagcaggagctattacaatattattaacagatcgaaatttaaatacaagAttttttgatcctgcaggtgggggagaccctattttataccaacatttattt

#### Description

**Male**. Wing length, 3.02; width, 1.11. **Head**. Light greyish-brown, with a pair of ochre-yellow bands running from behind ocelli towards base of antenna. Face yellowish, clypeus light brown. Antennal scape and pedicel ochre-brown, flagellum brown. Maxillary palpus dark brown, last palpomere lighter. Labella light greyish-brown. No setae on frons beyond line of ocelli. Flagellomere 4 slightly longer than flagellomere 4, flagellomere 4 1.5 wider than long. Maxillary palpomere 4 1.2× palpomere 3 length, palpomere 5 1.9× palpomere 4 length. **Thorax**. Scutum with ochre-yellow background and five brown longitudinal stripes, a slender additional light brown band at lateroanterior corner; scutellum ochre-yellow. Pleural sclerites whitish-yellow, except for a light brown band dorsally on laterotergite extending over dorsal half of mediotergite. Two pairs of prescutellars, inner pair over dorsocentral lines, outer pair on lateroposterior corner. Scutellum with one pair of bristles and smaller setae spread over disc. Fine setae and bristles over antepronotum, proepisternum bare. Halter pedicel ochre-yellow, knob light brown. **Legs**. Coxae whitish-yellow, femora dark ochre-yellow, tibiae and tarsi light greyish-brown. Fore coxa with scattered setae on anterior face and some few longer brown setae a distal end, mid coxa with some longer brown setae at distal end, hind coxa with a long row of greyish setae along most of its length. Anterior tibia with two setae dorsolaterally, mid tibia with two irregular dorsolateral rows of 6–8 setae, two lateral setae and two long irregular ventrolateral rows of 15–19 small bristles, hind tibia with a long dorsolateral row of 12 small bristles on outer face and a dorsolateral row of five small bristles on inner face. Fore leg tarsomere 1 0.9× tibia length, 1.5× tarsomere 2 length. **Wing** (Fig. 64B). Membrane background light greyish-brown, a dark brown over sc-r, first sector of Rs, R_4_, and base of M_4_, a greyish-brown large maculae on distal third of wing beginning at base of medial fork, a light brown mark at origin of M_4_ extending towards posterior margin. C extending beyond tip of R_5_ for a fifth of distance to M_1_. Sc complete, reaching C on anterior third of cell r1, setose at distal third until base of sc-r; sc-r close to origin of Rs. R_1_ reaching C at distal fifth of wing. First sector of Rs 2.5× r-m length. R_4_ present, transverse, length of cell r1 at anterior margin × length of R_4_. R_5_ reaching C at level of tip of M_1_, gently curved at distal half. False medial vein conspicuous, sclerotized. Length of M_1+2_ 5.4× r-m length; M_1_ and M_2_ gradually diverging; bM 4.6× length of first sector of Rs. M_4_ gently sinuous on distal half, origin basal to anterior end of M_1+2_. First sector of CuA only slightly longer than second sector. CuP produced to mid of second half of CuA, a gentle sign of sclerotized anal fold. No ventral macrotrichia on veins, dorsal macrotrichia on distal half of Sc (except on tip), entire length of bR, R_1_, second sector of Rs, on entire length of M_1_, distal third of M_2_ and M_4_, distal half of first sector of CuA, and entire second sector of CuA; sc-r, basal ⅔ of Sc, first sector of Rs, r-m, bM, M_1+2_ and CuP devoid of setation; membrane entirely devoid of setae. **Abdomen**. Tergites with light ochre-yellow background color, tergites 1–2 with medial brown longitudinal mark, tergites 3–4 and 6–7 with medial brown mark and a slender lateral band, tergite 5 mostly brown, with anterolateral corners light ochre-yellow. Sternites 1–4 whitish-yellow, sternites 5–7 more yellowish. **Terminalia** (Fig. 64C). Whitish-yellow. Gonocoxites fused extensively along medial line, suture of fusion evident, syngonocoxite entire ventral face bare, a short and wide medioventral projection of posterior margin that extends towards aedeagus, no laterodistal projection beyond base of gonostylus, dorsal border of gonocoxites separated from each other. Gonostylus large, longer than length of gonocoxites, setose on external face, a long sequence of spines along ventrodistal margin directed inwards, two much stronger distal spines, a laterodorsal setose sublobe. Gonocoxal bridge not visible. Aedeagus anteriorly with a median ejaculatory apodeme, medially a sclerotized inverted T, distally a medial plate with denticles. Parameres anteriorly with a pair of long lateral apodemes, laterodistally with a pair of triangular lobes bearing a concentration of strong spines distally on inner margin, a pointed triangle medially on distal margin. Tergite 9 apparently divided into a pair of rhomboid plates that touch each other medially, distal external margin with a sequence of setae, distal setae very long. Cerci not detected.

**Female** (Fig. 64A). As males, except for the following. **Wing l**ength, 3.70; width, 1.31. **Terminalia** (Fig. 64D). Sternite 8 with a single medial posterior subquadrate lobe, mostly with microtrichia, setae present along lateral margins and on ventral and dorsal faces at distal third, three long strong setae on each laterodistal corner. Sternite 10 with setae along distal margin and some shorter setae directed ventrally on ventral face. Tergite 8 wide, entirely bare. Tergite 9+10 with microtrichia and setae along posterior margin, much longer on lateroposterior corners. Cercomere 1 wide, long, cercomere 2 more or less rounded, both with dense microtrichia and short setae.

#### Material examined

**Holotype**: male, ZRCBDP0048968, Nee Soon (NS1), 15.apr.15, MIP leg. (website photo specimen, slide-mounted). **Paratypes** (1 female): ZRCBDP0048962, Nee Soon (NS1), 15.apr.15, MIP leg. (slide-mounted) (MZUSP). **Additional sequenced specimens:** ZRCBDP0066751, Bukit Timah, primary forest (BT09), 22.sep.16; ZRCBDP0136969, Bukit Timah, primary forest (BT08), 07.jun.17.

**Sequence failure specimens**: ZRCBDP0082303, Bukit Timah, primary forest (BT05), 09.nov.16.

**Possibly non-conspecific subcluster** (10 females): ZRCBDP0047780, Nee Soon (NS1), swamp forest, 28.aug.13, MIP leg.; ZRCBDP0047854, Nee Soon (NS1), swamp forest, 09.may.13, MIP leg.; ZRCBDP0047911, Nee Soon (NS1), swamp forest, 21.aug.13, MIP leg.; ZRCBDP0047913, Nee Soon (NS1), swamp forest, 21.aug.13, MIP leg.; ZRCBDP0047914, Nee Soon (NS1), swamp forest, 21.aug.13, MIP leg.; ZRCBDP0047915, Nee Soon (NS1), swamp forest, 21.aug.13, MIP leg.; ZRCBDP0048069, Nee Soon (NS1), swamp forest, 12.jun.13, MIP leg.; ZRCBDP0048474, Nee Soon (NS1), swamp forest, 04.apr. 12, MIP leg.; ZRCBDP0048476, Nee Soon (NS1), swamp forest, 18.apr. 12, MIP leg.; ZRCBDP0048478, Nee Soon (NS2), swamp forest, 30.may. 12, MIP leg. (website photo specimen, slide-mounted);

**Etymology**. The specific epithet of this species refers to a river (in the Franklin and Jackson’s 1828 Plan of Singapore) as well as a hill (Bukit Mandai) in the northern part of Singapore. It is now a planning area in Singapore.

The name is said to come from the Malay tree called *pokok* Mandai, but others suggest it might be a corruption of mandi (meaning “bathe” in Malay), as the river could have been used for this purpose. The noun is used in apposition.

**Remarks**. There are eight haplotypes for *Neoempheria mandai* Amorim & Oliveira, **sp. n.** The distances between some haplotypes is 4.15%. (e.g., ZRCBDP0047780 and ZRCBDP0048962). This is a case where most species delimitation methods point to two mOTUs. We have a single male from one of the subclusters and no males of the other subcluster, which would help with verifying conspecificity. This is another good example of how haplotype maps and different species delimitation approaches for barcode sequences guide the study of morphology in the quest of recognizing potentially hidden species diversity.

*Neoempheria malacca* Amorim & Oliveira, sp. n. (Figs. 65A–C, 66A-D)

https://singapore.biodiversity.online/species/A-Arth-Hexa-Diptera-00791,and-00742

urn:lsid:zoobank.org:act:2DAF4278-4B2C-4049-8469-65F027DA68F5

**Diagnosis**. Scutum with five longitudinal brown stripes. Wing with brownish marks, especially over distal end of radial sector—but no dark bands across the wing. Vein sc-r at basal third of cell r1; cell r1 long, anterior margin of over 6× length of R_4_. Abdominal tergites 3–6 ochre-yellow, with dark brown longitudinal marks along lateral margins; tergites 2–5 with a short brown mark medially along posterior margin. Gonocoxite with a large dorsal lobe extending to beyond tip of gonostylus; gonostylus long, petiolate, slightly wider at distal end, setae present only at distal third; aedeagus wide, cylindrical on distal half; tergite 9 with a pair of large lobes projecting posteriorly, close to each other, covered with strong setae on ventral face directed inwards.

***Neoempheria_malacca_ZRCBDP0048303_hapZRCBDP 0048303_SMH_holotype* [73: C, 130: T, 137: C, 139: A, 163: A, 202: A, 236: C, 238: T, 247: T, 295: C]**

tctctcttcaacaattgctcatacaggagcttctgttgatttagctattttttctcttcatttagcagggatCtcttcaattctaggggctgtaaattttattactacaattattaatatacgagccccTggtatcCaAtttgatcgaatacctttatttgtAtgatcagttttaattacagctattcttctattattgtcAttacctgtattagcaggagctattacaatattaCtTacagatcgTaatttaaatacaagcttttttgaccctgcaggtggaggagatcctatCctttatcaacatttattt

**Description** (Fig. 65A). **Male**. Wing length, 4.59, width, 1.64. **Head**. Light ochre-yellow background, with a light brown band over vertex. Antennal scape and pedicel light brown, flagellum brown. Face light brown, clypeus light ochre-yellowish, triangular, with scattered setae. Maxillary palpus brown, labella light yellowish-brown. Head elongate antero-posteriorly. Eyes small, rounded, covered with interommatidial setulae. Two ocelli placed medially on vertex at posterior end of frontal furrow, each ocellus over a black background; post-ocellar setae present. Frons bare, occiput with longer setae posteriorly to eye, small setae dorsally on occiput. Antennal scape with setulae around distal margin ventrally, dorsally with elongate setae; pedicel with setulae around distal margin, some setae and one long seta dorsally; scape 1.6× pedicel length; flagellomere 4 1.2× wider than long. Face slender, bare. Clypeus triangular, slightly bulging, projected ventrally beyond insertion of palpus, with scattered fine setae. Palpomere 1 not produced; palpomere 2 with small setae ventrodistally; palpomere 3 with conspicuous sensorial pit opening dorsally on proximal half, scattered setulae and dorsoapically with small setae; palpomere 4 with a short distal projection dorsally over base of palpomere 5, with some setulae and setae dorsally, palpomere 4 1.4× length of palpomere 3; palpomere 5 with annellation on most of its length, setulae and small setae on distal half, especially close to tip, palpomere 5 1.3× palpomere 4 length. Labella small, extending backwards, with a pair of pseudotrachea, small setae along its length on distal half. **Thorax** (Fig. 66A). Scutum with ochre-yellow background and five brown longitudinal stripes; scutellum ochre-yellow with a couple of small brownish marks over separation with scutum. Pleural sclerites whitish-yellow, except for yellowish laterotergite, a light brown mark on antepronotum medially, cervical sclerite brown, a light brown medial mark over dorsal half of mediotergite. Irregular rows of long brown setae on scutum along brown stripes, medial ochre-yellow stripes nearly bare, setae over ochre-yellow lateral bands, six fine bristles above anepisternum, a line of five brown supra-alar bristles; four pairs of prescutellars, two on dc rows, one sub-medially, one on lateroposterior corner. Scutellum short, one pair of bristles, smaller setae spread over disc. Bristles and smaller setae on antepronotum. Proepisternum subquadrate, bare, propepimeron elongate, directed almost ventrally. Anepisternum elongate from ventroanterior to dorsoposterior ends, a large membranous area around anterior spiracle, anterior basalare bare, katepisternum subquadrate, rounded ventrally, bare. Mesepimeron wider dorsoposteriorly, ventral extension slender, reaching ventral margin of pleura. Laterotergite slightly bulging, placed obliquely, bare. Metepisternum slender, elongate antero-posteriorly, bare; metepimeron produced, small. Mediotergite gently curved on dorsal two-thirds. Haltere yellowish, with small setae along pedicel and base of knob, extending distally at ventral face. **Legs**. Coxae yellowish, mid and hind coxae darker towards distal end. Femora dark ochre-yellowish; tibiae light greyish-brown, brown distally, tarsi brownish. Fore coxa with setation on anterior face, some few longer, curved setae at distal end; mid coxa with an irregular row of setae medially along distal half and some larger setae on distal third of anterior face; hind coxa medially with a row of long setae and some smaller fine setae along most of length of lateral face. Femora with scattered short setae along entire length, a short row of longer brownish setae close to distal end ventrally. Tibiae and tarsi with regular lines of brownish trichia. Fore tibia with additional scattered setae along most of its length, a stronger brown seta at tip; anteroapical depressed area on inner face of fore tibia wide, lined with setulae. Mid and hind tibia with long rows of dorsal, lateral and ventral brown setae, a regular comb of elongate setae distally at inner face and some additional strong setae at tip. Tarsomeres 1–3 with regular rows of setae dorsally and at both lateral faces, all tarsomeres with a pair of distal ventral setae. Tibial spurs on mid and hind tibiae subequal in length, spurs over 2.9× longer than tibia width at apex. Tarsal claw with a strong ventral tooth medially and two additional small teeth more basally. **Wing** (Fig. 65B). Membrane fumose light brown, lighter medially on distal wing cells, darker along most veins. C extending beyond tip of R_5_ for about a fifth of distance to M_1_. Sc complete, reaching C before mid of cell r1, sc-r reaching R_1_ at basal third of cell r1, anterior margin of cell r1 over 5× longer than length of R_4_. Origin of Rs before mid of wing, first sector of Rs oblique, long, twice length of r-m, bare; R_1_ reaching C on distal tenth of wing; R_4_ present, beyond level of medial fork, more or less transverse; R_5_ reaching C at level of tip of M_1_. False medial vein inconspicuous, not sclerotized. Posterior veins sclerotized until slightly before wing margin. M_1_and M_2_ gradually diverging, M_1+2_ about 5× r-m length; bM long, 6.5× length of first sector of Rs. M_4_ with a discrete sinuosity on distal half, first sector of CuA as long as second sector. Cubital pseudovein long, sclerotized, extending to distal fourth of second sector of CuA. CuP produced to mid of second sector of CuA. No anal pseudovein, only a mild fold produced. No ventral macrotrichia on veins, dorsal macrotrichia on distal half of Sc (except at distal end), entire length of bR, R_1_, and second sector of Rs, on entire length of M_1_, M_2_, M_4_, and CuA, and on basal fifth of CuP; R_4_, first sector of Rs, r-m, bM, and M_1+2_ devoid of setation; membrane entirely devoid of setae. **Abdomen**. Tergites mostly ochre-yellow, tergite 1 with a faint medial brownish mark, tergites 2–5 with a medial brown mark on anterior and posterior margins, tergites 3–6 with a slender brown band along lateral margins. Sternites 1–7 ochre-yellow. **Terminalia** (Figs. 66B–D). Ochre-yellow. Gonocoxites fused medially along anterior end of terminalia, syngonocoxite entire ventral face bare, medio-posterior margin ventrally extending towards aedeagus, a laterodistal large setose lobe on each side extending to beyond tip of gonostylus. Gonostylus simple, digitiform, slender, slightly capitate distally, bare along proximal two-thirds, setae close to tip longer, slightly curved. Gonocoxal bridge extending distally and connecting both gonocoxites almost at level of tip of laterodistal projections, gonocoxal apodemes directed medially on anterior end. Aedeagus long, apodeme elongate, reaching anterior end of terminalia, wide and strongly sclerotized medially, with two pairs of lateral expansions, tips curved anteriorly, distal end tubular, wide, opening at level of tip of gonostylus.

Parameres present at a posterior position, dorsal to tip of aedeagus, crown-shaped, anterior end straight, laterals projecting distally. Tergite 9 displaced posteriorly, present as a pair of long, sclerotized lobes curved inwards, connected medially at anterior end, closer to each other at tip than midway to apex, with setae on dorsal face, ventral face densely covered with strong setae. Cerci small, lobose, covered with microtrichia and setae, placed ventrally to base of tergite 9.

**Female**. As males, except for the following. **Wing l**ength, 4.95; width, 1.70. **Terminalia** (Fig. 65C). Sternite 8 composed of a wide slender anterior sclerotized band, with a medial subquadrate projection, a pair of short lobes medially along anterior margin, some elongate fine setae medially at anterior end, two pairs of slender spines medially at posterior margin, some additional setae directed inwards. Sternite 9 with a vaginal furca strongly sclerotized subquadrate medial plate. Sternite 10 as a pair of digitiform lobes with long setae emerging distally. Tergite 8 wide, short, as a transverse band covered with microtrichia, entirely devoid of setae. Tergite 9+10 rectangular, connecting laterally to sternite 9 laterodistal arms, covered with microtrichia, bearing long setae along posterolateral margin. Cercomere 1 elongate, 2.5× length of cercomere 2, densely covered with microtrichia and with small setae, distal border slightly beyond insertion of cercomere 2; cercomere 2 subspherical, with microtrichia and small setae.

#### Material examined

**Holotype**: male, ZRCBDP0048303, Sungei Buloh (SB1), mangrove, 16.oct. 2013, MIP leg. (website photo specimen, slide-mounted). **Paratypes** (10 males): ZRCBDP0279172, Singapore, 31.may. 2018, MIP leg; ZRCBDP0278314, Singapore, 31.may. 2018, MIP leg.; ZRCBDP0070102, Pulau Ubin (PU01), mangrove, 27.jul. 16, MIP leg.; ZRCBDP0041023, Singapore Botanical Gardens (CUGE), 20.Oct. 2017, MIP leg.; ZRCBDP0278001, Singapore Botanical Gardens (CUGE), 20.Oct. 17, MIP leg.; ZRCBDP0278438, National University of Singapore (SDE), 20.Dec. 2017, MIP leg. (MZUSP); ZRCBDP0284205, Pulau Ubin (PU20), no date, MIP leg.; ZRCBDP0314193, Sungei Buloh (SB1), mangrove, 20.Mar. 2013, MIP leg.; ZRCBDP0278351, Singapore Botanical Gardens (CUGE), 24.nov. 2017, MIP leg.; ZRCBDP0284286, Singapore, 03.may. 2018.

**Possibly non-conspecific subcluster** (4 females): ZRCBDP0048482, Nee Soon (NS2), swamp forest, 18.apr. 2012, MIP leg. (website photo specimen, slide-mounted); ZRCBDP0048942, Nee Soon (NS2), 03.dec. 2014, MIP leg.; ZRCBDP0069309, Pulau Ubin (PU01), mangrove, 11.may. 2016, MIP leg.; ZRCBDP0154957, Nee Soon (NS1), swamp forest, 04.Mar. 2015, MIP leg.

**Etymology**. The specific epithet of this species refers to the Strait of Malacca, a narrow stretch of water of 930 km in length, between the Malay Peninsula—with Singapore at its southern end— and the Indonesian island of Sumatra. The Strait of Malacca is historically an important maritime trade route between India and China. The noun is used in apposition.

**Remarks**. The haplotype of the holotype of *Neoempheria malacca* Amorim & Oliveira, **sp. n.** differs from another haplotype consisting only of females by 3.88%. PTP, OC 2% and 3%, and ABGD up to P=0.036 suggest two species. We cannot check conspecificity of the second subcluster based only on females and the specimens of the second subcluster were not include as paratypes. This species belongs to the group *ferruginea* of *Neoempheria* (see Sueyoshi 2014) and the male terminalia is similar to the male terminalia of *N. proxima* Zaitzev and *N. subproxima* Zaitzev (Zaitzev 2001).

*Neoempheria sinkapho* Amorim & Oliveira, sp. n. (Figs. 67A-D)

https://singapore.biodiversity.online/species/A-Arth-Hexa-Diptera-00788

urn:lsid:zoobank.org:act:2A828E5D-53FF-4CFD-9B4D-EC75F753B0BE

**Diagnosis**. Scutum with five longitudinal brown stripes, a brownish line over dorsal half of mediotergite extending to dorsal third of laterotergite. Wing with a brown mark across tip of wing, a second, light brown mark across wing between level of sc-r and anal lobe through level of origin of M_4_, with an additional dark brown mark over R_4_. C extending beyond tip of R_5_ for a ¼ of distance to M_1_; sc-r at basal third of cell r1; anterior margin of cell r1 slightly less than 3× length of R_4_. Abdominal tergites 1–2 and 7 cream-yellow with a caramel-brown transverse mark along posterior margin; tergites 3–4 and 6 cream-yellow with brown marks extending medially to the anterior margin and a slender brown mark along lateral margin; tergite 5 mostly caramel-brown with a large cream-yellow mark on lateroanterior corners. Gonocoxites with a dorsal setose extension reaching beyond tip of gonostylus; gonostylus small, flat, with a short outer lobe near tip; aedeagus with a medial crest with a row of spines; parameres with a line of short spines medially at posterior margin; tergite 9 with a pair of posterior setose lobes.

***Neoempheria_sinkapho_ZRCBDP0047884_hapZRCBD P0047884_SMH_holotype* [46: A, 55: A, 70: G, 130: C, 134: G, 147: A, 261: G, 274: C]**

tttatcttcaactattgctcatacaggagcttctgttgatttagcAattttttcAttacatttagctggGatttcttctattttaggagctgttaattttattacaacaatcattaatatacgagctccCggaGtacaatttgataAaatacctttatttgtatgatctgtattaattacagcaattttattattattatcattaccagtattagctggagctattacaatattactaacagatcgaaatttaaatacaaGattctttgacccCgcaggtggaggagaccctattctttaccaacatttattt

#### Description

**Male** (Fig. 67A). Wing length, 3.34; width, 1.31. **Head**. Ochre background, with a brownish band running from dorsal end of eye towards mid of vertex and a dark brown medial band above base of antennae. Face brownish, clypeus light brown. Antennal scape and pedicel brown, flagellum dark brown (Fig. 67B). Maxillary palpus brown, last palpomere lighter, labella whitish. No post-ocellar setae, no setae on frons anteriorly to line of ocelli. Scape 1.4× length of pedicel, flagellomere 1 1.5× flagellomere 2 length, flagellomere 4 0.7× longer than wide. Palpomere 4 1.1× palpomere 3 length, palpomere 5 1.7× palpomere 4 length. **Thorax**. Scutum ochre-yellow, a pair of brown stripes along dorsocentral lines, a slender light brown medial band, an additional pair of brownish bands along laterals; scutellum ochre-yellow with a brownish tinge medially. Pleural sclerites whitish with an orangish tinge, except for a brown mark anteriorly on antepronotum lobes, a brown mark on laterotergite dorsoposterior end, and a brown band along dorsal half of mediotergite. Scutum with a pair of strong prescutellar setae on dorsocentral line and two pairs of stronger prescutellar setae on lateroposterior corner; one pair of scutellar strong setae, besides an irregular row of small setae more anteriorly. Antepronotum with one bristle and some other large and some small setae, proepisternum bare. Halter pedicel ochre-yellow, knob light brown. **Legs**. Coxae whitish, fore femur whitish-yellow, mid and hind femur ochre-yellow, tibiae and tarsi light greyish-brown. Hind tibia inner spur 3.5× longer than tibia width at apex. **Wing** (Fig. 67C). Membrane background light greyish-brown, with brown marks over sc-r, first sector of Rs, R_4_, and base of M_4_, and a large greyish-brown macula on distal third of wing, beginning at base of medial fork. C produced beyond tip of R_5_ for about a fifth of distance to M_1_. Sc complete, reaching C almost at mid of cell r1; R_4_ present, transverse, far from origin of Rs, anterior margin of cell r1 3.2× length of R_4_; first sector of Rs 1.0× r-m length. False medial vein present, hardly sclerotized. M_1+2_ 3.3× r-m length; bM 7.7× length of first sector of Rs. First sector of CuA 0.90× longer than second sector. M_4_ gently depressed on distal half. Cubital pseudovein sclerotized to distal fourth of CuA; CuA unsclerotized at very tip; CuP sclerotized to basal third of second sector of CuA. Wing margin gently emarginated at tip of CuA. No macrotrichia on wing membrane, no ventral macrotrichia on veins, dorsal macrotrichia on distal ¾ of Sc, entire length of bR, R_1_, second sector of Rs, M_1_, M_2_, M_4_, and CuA; sc-r, first sector of Rs, R_4_, r-m, bM, M_2_, M_4_, and CuP devoid of macrotrichia. **Abdomen**. Tergites with light ochre-yellow background color, tergites 1–2 with a medial brown longitudinal mark, tergites 3–7 with medial brown mark extending laterally on posterior margin and a slender lateral band, tergite 5 mostly brown with medial and lateral brown bands connected, only anterolateral corners light ochre-yellow. Sternites 1–4 whitish-yellow, sternites 5–7 more yellowish. **Terminalia** (Fig. 67D). Whitish-yellow. Gonocoxites fused along a slender medial connection, no suture of fusion, short medio-posterior projection that extends dorsally, connecting to aedeagal sclerite, lateroposteriorly projecting only slightly beyond insertion of gonostylus. Gonostylus laterally at terminalia, a straight, bare inner lobe projecting posteriorly and a bifid outer lobe, ventral branch directed obliquely inwards with some short setae on distal half, dorsal branch directed posteriorly covered with setae on external face. Gonocoxal bridge with a pair of long arms that project towards anterior end of terminalia, a pair of apodemes anteriorly close to each other. Aedeagal sclerite with anterior end slender, widening mid-way to apex, a pair of sub-medial short lobes, a pair of longer slender lobes projected obliquely inwards reaching level of insertion of gonostylus. Paramere widening towards posterior third, then with a slender distal projection ending with a dorsoventral crest with a sequence of short teeth. Tergite 9 with a pair of long lobes connected medially, each lobe long, slendering towards apex, projecting way beyond tip of gonostylus, covered with long setae on dorsal face, a distal strong curved seta at tip of inner margin directed anteriorly. Sternite 10 triangular, with a group of five spines directed ventrally on distal end. Cerci slender, elongate.

**Female**. Unknown.

#### Material examined

**Holotype**: male, ZRCBDP0047884, Nee Soon (NS2), swamp forest, 23.oct.13, MIP leg. (website photo specimen, slide-mounted). **Paratypes** (2 males): ZRCBDP0048809, Nee Soon (NS1), 14.jan.15, MIP leg. (extracted); ZRCBDP0048955, Nee Soon (NS1), 15.apr.15, MIP leg.

**Etymology**. The specific epithet of this species refers to the Chinese Hokkien dialect transcription of Singapore, Xin Jia Po [新加坡]. Hokkien is one of the largest Chinese dialect groups in Singapore. The noun is used in apposition.

**Remarks**. This is another species of the group *ferruginea* of *Neoempheria*, with specimens from the swamp forest. There are two haplotypes for *Neoempheria sinkapho* Amorim & Oliveira, **sp. n.,** which are separated into two species by only objective clustering at 2% p-distance.

*Neoempheria singapura* Amorim & Oliveira, sp. n. (Figs. 68A–F)

https://singapore.biodiversity.online/species/A-Arth-Hexa-Diptera-00830

urn:lsid:zoobank.org:act:EC145915-5842-479B-BB11-92DCCB71743F

**Diagnosis**. Scutum with five longitudinal brown stripes, a brownish line over dorsal half of mediotergite extending on dorsal third of laterotergite. Wing with brown mark across tip of wing and a second, not as strong mark across wing from level of sc-r to anal lobe at level of origin of M_4_. C extending beyond R_5_ for about a ¼ of distance to M_1_; sc-r at basal third of cell r1; anterior margin of cell r1 slightly less than 4× length of R_4_. Abdominal tergites 1–2 cream-yellow, with a caramel-brown longitudinal mark medially; tergites 3–4 and 6 mostly caramel-brown, with cream-yellow marks close to lateroanterior corner, and slender lateral brown band; tergite 5 mostly caramel-brown, with a small cream-yellow mark on lateroanterior corners; tergite 7 mostly cream-yellow with a small caramel-brown mark medially along posterior margin. Gonostylus large, with a small ventral digitiform lobe; aedeagus with a medial crest with a row of spines; parameres with a group of short spines at distal end medially; tergite 9 with a pair of distal rounded lobes, with a subapical spine directed inwards.

***Neoempheria_singapura_ZRCBDP0047796_hapZRCBD P0047796_SMH_holotype* [4: T, 25: T, 40: C, 109: T, 17 2: A, 262: C, 265: C, 271: T, 298: T]**

tctTtcttcaacaattgctcatacTggagcttcagttgaCttagctattttttctttacatttagctggtatttcttcaattttaggagctgtaaattttattactacTattattaatatacgagctccaggaattcaatttgatcgaatacctttatttgtttgatctgtAttaattacagctattcttttactattatctttacctgtattagcaggagctattacaatattattaacagatcgaaatttaaatactagCttCtttgaTccagctggaggaggagatcctattctTtatcaacatttattt

#### Description

**Male**. Wing length, 2.79; width, 1.08. **Head**. Light ochre-brown, lighter towards base of antenna, whitish at ventral half of occiput. Face ochre. Antennal scape and pedicel light brown, clypeus light brown. Maxillary palpus 2–3 dark brown, last two palpomeres light brown, labella light brown. No setae on frons anteriorly to line of ocelli, post-ocellar setae absent. Scape 1.4× pedicel length, flagellomere 1 1.3× flagellomere 2 length, flagellomere 4 as long as wide. Palpomere 4 1.3× palpomere 3 length, palpomere 5 1.5× palpomere 4 length. **Thorax**. Scutum with ochre-yellow background and five brown longitudinal stripes, besides a slender light brown band along anterior margin; scutellum ochre-yellow. Pleural sclerites whitish, except for a brown mark on antepronotum internal margin anteriorly, a light brown band dorsally on laterotergite and a brown band at dorsal half of mediotergite. Scutum with a pair of prescutellar bristles on dorsocentral line and two pairs of bristles on lateroposterior corners; scutellum with one pair of bristles, no smaller setae. Antepronotum with two bristles, one strong seta and some additional small setae. Halter pedicel ochre-yellow, knob light brown. **Legs**. Coxae whitish, femora ochre-yellow, mid and hind femora darker, tibiae light greyish-brown, brown at tip, tarsi brownish. Hind tibia inner spurs 3.5× longer than tibia width at apex. **Wing** (Fig. 68B). Membrane background light greyish-brown, with dark brown marks over sc-r, first sector of Rs, R_4_, and base of M_4_, and a greyish-brown large macula on distal third of wing, beginning at base of medial fork. C produced beyond tip of R_5_ for about a ¼ of distance to M_1_; Sc complete, reaching C slightly before mid of cell r1, sc-r sclerotized, reaching R_1_ beyond level of origin of Rs. R_1_ reaching C at distal fourth of wing; R_5_ reaching C before level of tip of M_1_; R_4_ present, transverse, far from base of Rs, cell r1 long, anterior margin of cell r1 3.8× longer than R_4_ length; first sector of Rs 2.3× r-m length. False medial vein conspicuous, sclerotized; M_1+2_ 5.5× r-m length; bM 4.5× length of first sector of Rs; first sector of CuA 1.2× longer than second sector. M_4_ not sinuous on distal half.

Cubital pseudovein sclerotized to mid of second sector of CuA; CuP sclerotized to mid of second sector of CuA. Anal fold present. Wing margin gently emarginated at tip of CuA. No ventral macrotrichia on veins, dorsal macrotrichia on distal half of Sc, entire length of bR, R_1_, second sector of Rs, most of M_1_, distal third of M_2_ and M_4_, distal third of first sector, and entire second sector of CuA; sc-r, first sector of Rs, R_4_, r-m, bM and CuP entirely devoid of macrotrichia. **Abdomen**. Tergites 1–2 cream-yellow with medial light brown mark, on tergite 2 extending laterally along posterior margin, tergites 3–4 and 6 with light brown medial mark extending outwards on posterior margin, reaching or almost reaching a slender longitudinal light brown lateral mark, tergite 5 light brown with cream-yellow anterolateral corner, tergite 7 mostly ochre-yellow with a light brown mark along posterior margin. Sternites 1–5 whitish-yellow, sternites 6–7 more yellowish. **Terminalia** (Figs. 68C–E). Light ochre-brown. Gonocoxites fused medially, no suture, entirely bare, a medial extension on posterior margin connecting to a strongly sclerotized aedeagal sclerite inwards, no lateroposterior extension beyond insertion of gonostylus. Gonostylus placed laterally on terminalia, with a short digitiform ventral process basally directed inwards bearing fine setae concentrated at distal end and a large additional dorsal branch, wider midway to apex, covered with setae on most of external face, strong and concentrated setae midway to apex, and fine setae on inner margin at distal fourth. Gonocoxal bridge large, connected medially on anterior end. Aedeagus with anterior apodeme weakly sclerotized, a sclerotized structure with a pair of winglets midway to apex close to posterior margin of syngonocoxite medially, and a distal trapezoid plate posteriorly, with a sclerotized median ridge and a crest at tip with a sequence of short spines, laterally a pair of elongate pointed projections almost reaching level of medial crest, dorsally to genital opening a weakly sclerotized area with microtrichia and setae at posterior margin. Parameres with a large plate, distally with a pair of lateroposterior short projections with setae and a medial curved projection with 16 spines directed ventrally. Tergite 9 present as a pair of long projections barely connected medially, each lobe dorsally with no microtrichia and a sequence of three long setae, a rounded lobe midway to apex with a strong spine directed inwards and a ventral-posterior rounded lobe extending beyond tip of cerci, with a concentration of short setae on dorsal face. Cerci elongate, weakly sclerotized, partially covered by tergite 9 posterior projections, with microtrichia and setae. **Female** (Fig. 68A). As male, except for the following. **Wing l**ength, 2.75; width, 1.02. **Terminalia** (Fig. 68F). Sternite 8 small, subquadrate, slightly elongate, covered with microtrichia and fine setae, two pairs of long setae on posterior margin. Sternite 9 with short medial projection at anterior end, a pair of lateral arms rounded anteriorly meeting medially, with a slender genital chamber. Tergite 8 wide, lateral ends projecting ventrally to almost reach sternite 8 plate, covered only with microtrichia, bare of setae. Sternite 10 large, trapezoid, weakly sclerotized, medially with a circular area covered by a transparent area with two pairs of setae. Tergite 9+10 slender medially, with long setae along posterior margin, connected laterally to sternite 10. Cercomere 1 elongate, 2.6× length of cercomere 2, both covered with microtrichia and short setae.

#### Material examined

**Holotype**: male, ZRCBDP0047796, Nee Soon (NS2), swamp forest, 13.nov.2013, MIP leg. (slide-mounted). **Paratypes** (6 males, 12 females). **Males**: ZRCBDP0047807, Nee Soon (NS1), swamp forest, 01.may.13, MIP leg.; ZRCBDP0047944, Nee Soon (NS2), swamp forest, 20.nov.13, MIP leg.; ZRCBDP0048734, Nee Soon (NS2), 06.may.15, MIP leg.; ZRCBDP0048901, Nee Soon (NS1), 31.dec.14, MIP leg.; ZRCBDP0048993, Nee Soon (NS2), 17.dec.14, MIP leg.; ZRCBDP0049107, Nee Soon (NS1), 24.dec.14, MIP leg.

**Females**: ZRCBDP0047802, Nee Soon (NS2), swamp forest, 13.nov.13, MIP leg.; ZRCBDP0047844, Nee Soon (NS1), swamp forest, 24.apr.13, MIP leg. (slide-mounted); ZRCBDP0047928, Nee Soon (NS1), swamp forest, 17.jul.13, MIP leg.; ZRCBDP0047930, Nee Soon (NS1), swamp forest, 17.jul.13, MIP leg. (website photo specimen); ZRCBDP0047935, Nee Soon (NS1), swamp forest, 17.apr.13, MIP leg.; ZRCBDP0048479, Nee Soon (NS2), swamp forest, 18.jul. 12, MIP leg.; ZRCBDP0048692, Nee Soon (NS2), swamp forest, 02.may. 12, MIP leg.; ZRCBDP0048872, Nee Soon (NS1), 31.dec.14, MIP leg.; ZRCBDP0048890, Nee Soon (NS1), 31.dec.14, MIP leg.; ZRCBDP0049089, Nee Soon (NS1), 24.dec.14, MIP leg.; ZRCBDP0049177, Nee Soon (NS2), 13.may.15, MIP leg.; ZRCBDP0154903, Singapore, date range 2012-2018, MIP leg. **Additional sequenced specimens**: female, ZRCBDP0133978, Singapore, date range 2012-2018, MIP leg.

**Etymology**. The specific epithet of this species comes from the Malay (as the official language) name for Singapore. The noun is used in aposition.

**Remarks**. Most of the specimens of *Neoempheria singapura* Amorim & Oliveira, **sp. n.** are from the swamp forest.

*Neoempheria* sp. C (Figs. 69A–C)

https://singapore.biodiversity.online/species/A-Arth-Hexa-Diptera-00777

***Neoempheria_spC_ZRCBDP0048477_hapZRCBDP0048 477_SMH_unnamed_type* [1: A, 7: C, 10: A, 37: A, 73: C, 88: T, 178: C, 196: A, 296: C, 298: T]**

ActttcCtcAacaattgctcatacaggggcatctgtAgatttagctattttttctttacatttagctggtatCtcttcaattttaggTgctgttaattttattacaacaattattaatatacgagccccaggaattcaatttgatcgaatacctttatttgtttgatctgttttaatCacagctattttattattAttatctttaccagtattagctggagctattacaatattattaacagatcgaaatttaaatactagtttttttgacccagctggagggggagatccaattCtTtatcaacatttattt

#### Description

**Female** (Fig. 69A). Wing length, 2.79; width, 1.02. **Head**. Vertex and occiput light brown. Face light brown, clypeus light ochre-yellowish. Antennal scape and pedicel light brown, flagellum brown. Maxillary palpus brown, labella light brown. Scape 1.5× pedicel length, flagellomere 1 1.4× flagellomere 2 length, flagellomere 4 1.0× longer than wide. Palpomere 4 1.1× palpomere 3 length, palpomere 5 2.1× palpomere 4 length. **Thorax**. Scutum with ochre-yellow background and five brown longitudinal stripes; scutellum ochre-yellow. Pleural sclerites whitish, except for a light brown wide mark on antepronotum anteriorly, a brown cervical sclerite and a light brown mark medially on dorsal half of mediotergite. Halter whitish with light brown tinge. Scutum with a pair of prescutellar bristles at dorsocentral line and two pairs of bristles at lateroposterior corner; scutellum with one pair of bristles, no additional smaller setae. Antepronotum with one larger bristle, one smaller bristle, five long setae and some smaller setae, proepisternum bare. **Legs**. Coxae whitish-yellow, femora ochre-yellow, tibiae light greyish-brown with brown tips, tarsi brown. Hind tibia inner spur 3.0× longer than tibia width at apex. Tarsomere 1 of fore leg 1.0× tibia length, 1.9× tarsomere 2 length. **Wing** (Fig. 69B). Membrane with dark brown separate marks over sc-r, first sector of Rs, R_4_, and bM, a wide brown band on distal third of wing beginning at level of base of medial fork. C produced beyond tip of R_5_ for about a fifth of distance to M_1_; Sc complete, reaching C at basal fourth of cell r1, sc-r sclerotized, reaching R_1_ slightly beyond level of origin of Rs. R_1_ reaching C at distal fourth of wing; R_5_ reaching C before level of tip of M_1_; R_4_ present, transverse, far from base of Rs, cell r1 long, anterior margin of cell r1 4.0× longer than length of R_4_; first sector of Rs 2.2× r-m length. False medial vein conspicuous, sclerotized, curved on basal third. M_1+2_ 6.0× r-m length; bM 4.9× length of first sector of Rs. First sector of CuA 1.5× longer than second sector. M_4_ gently depressed on distal half. Cubital pseudovein sclerotized to beyond mid of second sector of CuA; CuP sclerotized to slightly beyond base of M_4_. Anal fold faint. Wing margin not emarginated at tip of CuA. No ventral macrotrichia on veins, dorsal macrotrichia on entire length of bR, R_1_, second sector of Rs, distal third of M_1_, M_2_ and M_4_, distal fourth of first sector, and entire second sector of CuA; sc-r, Sc, base of bR, first sector of Rs, R_4_, r-m, bM, and CuP entirely devoid of macrotrichia. **Abdomen**. Tergites 1–4 and 6–7 light cream-yellow with medial brown mark extending laterally along posterior margin on segments 2– 4 and 6–7, tergites 3–6 with a slender brown band along lateral margin, tergite 5 mostly brown with a ochre-yellow mark laterally along anterior ¾. Tergite 7 with no digitiform projections. Sternites 1–4 whitish, sternites 5–7 more ochre-yellow with brownish tinge. **Terminalia** (Fig. 69C). Ochre-yellow, tip of sternite 8 lobes and cerci brown. Sternite 8 with a wide weakly sclerotized anterior area covered with microtrichia and no setae, and a small, subquadrate, slightly elongate plate, with a short medial V-shaped incision distally separating short, pointed lobes, covered with microtrichia and fine setae, two pairs of long setae on posterior margin. Sternite 9 wide, weakly sclerotized. Tergite 8 wide, lateral ends projecting ventrally, covered only with microtrichia, bare of setae, anterior margin with a sclerotized band along its length. Sternite 10 trapezoid, weakly sclerotized, medially with a circular area covered by a transparent window with two pairs of setae laterally. Tergite 9+10 slender, with long setae along posterior margin, connected laterally to sternite 10. Cercomere 1 elongate, close to each other medially, 1.4× length of cercomere 2, both strongly sclerotized, covered with microtrichia and short setae.

**Male**. Unknown.

#### Material examined

Female, ZRCBDP0048477, Nee Soon (NS2), swamp forest, 09.may. 12, MIP leg. (slide-mounted).

*Neoempheria xinjiapo* Amorim & Oliveira, sp. n. (Figs. 70A–D, 71A–B)

https://singapore.biodiversity.online/species/A-Arth-Hexa-Diptera-000824

urn:lsid:zoobank.org:act:59C6E24D-154A-4EE4-8634-C89AECE2D5EA

**Diagnosis**. Scutum with five longitudinal brown stripes, a brownish line over dorsal half of mediotergite extending to dorsal third of laterotergite. Wing with a subapical brown band and a second band across wing at level of sc-r/first section of Rs and anal lobe at level of origin of M_4_, dark brown mark over R_4_; sc-r at basal third of cell r1; anterior margin of cell r1 long, more than 4× length of R_4_. Abdominal tergites 1–2 and 7 whitish with a caramel-brown transverse mark along posterior margin; tergites 3– 4 and 6 with a similar pattern but with a brown mark extending medially to anterior margin and a slender brown mark along lateral margin; tergite 5 mostly caramel-brown with a large cream-yellow mark on lateroanterior corners—tergites 5–7 with a more yellowish background. Gonocoxite with a short, hook-like projection extending beyond base of gonostylus. Gonostylus at lateroposterior end of gonocoxite, digitiform, with fine setae. Aedeagal plate with two pairs of sclerotized short projection directed inwards. Parameres with a short digitiform lobe directed outwards and a lobe directed inwards with a strongly sclerotized tooth. Tergite 9 with a pair of digitiform lobes laterodistally bearing fine setulae and two small and one strong spines distally.

***Neoempheria_xinjiapo_ZRCBDP0047820_hapZRCBDP 0047809_SMH_holotype* [46: C, 55: A, 76: A, 79: T, 97: C, 110: A, 190: T, 193: T, 298: T]**

tttatcctcaacaattgctcatacaggagcttctgttgatttagcCattttttcAttacatttagcaggaatttcAtcTattttaggagctgttaaCttcattacaacaAttattaatatacgagcccccggaattcaatttgatcgaatacctttgtttgtttgatcagttttaattacagctgtactTctTttactttctttaccagtattagctggagctattacaatattattaacagatcgaaatttaaatactagattctttgacccagctgggggaggagaccctattctTtatcaacatttattt

#### Description

**Male** (Fig. 70A). **Head** (Fig. 70B). Light brown, ochre-yellow posteriorly to vertex, more whitish towards ventral margin of occiput. Face light brown, clypeus light brown. Antennal scape and pedicel light brown, flagellum dark brown. Maxillary palpus brown, distal palpomere lighter, labella light brown. Ocellar setae present. Scape 1.2× pedicel length, flagellomere 1 1.5× flagellomere 2 length, flagellomere 4 1.2× longer than wide. Palpomere 4 1.5× palpomere 3 length, palpomere 5 2.0× palpomere 4 length. **Thorax**. Scutum with ochre-yellow background with five brown longitudinal stripes plus a pair of slender brown marks laterally, scutellum ochre-yellow. Pleural sclerites whitish, except for a brown mark on antepronotum anteriorly, a brown mark across laterotergite dorsoposterior end and a brown mark across mediotergite dorsal half. Halter light brown. Scutellum with a pair of prescutellar bristles on dorsocentral line, a pair of prescutellars at each lateroposterior corner. Antepronotum with two larger bristles, one smaller bristles and some additional setae, proepisternum bare. **Legs**. Coxae whitish-yellow, femora ochre-yellow, tibiae light greyish-brown with brown tips, tarsi brown. Fore leg tarsomere 1 0.90× tibia length, 1.9× tarsomere 2 length. Hind tibia inner spur 3.9× tibia width at apex. **Wing** (Fig. 70C). Membrane with a dark brown band across wing at level of sc-r, a brown mark over R_4_ and a brown band across wing at level of tip of R_1_ on distal end of wing. C produced beyond tip of R_5_ for about a fifth of distance to M_1_; Sc reaching C at level of basal third of cell r1; sc-r reaching R_1_ at level of anterior end of r-m. R_1_ reaching C at distal fifth of wing; R_5_ reaching C at level of tip of M_1_; R_4_ present, gently inclined, far from base of Rs, cell r1 trapezoid, long, anterior margin of cell r1 4.2× longer than length of R_4_; first sector of Rs 2.4× r-m length. False medial vein present, gently sclerotized. M_1+2_ 6.1× r-m length; bM 12.0× length of first sector of Rs. First sector of CuA 1.3× longer than second sector. M_4_ gently depressed on distal half. Cubital pseudovein sclerotized to distal third of second sector of CuA; CuP sclerotized to basal third of second sector of CuA. Anal fold faint. Wing margin gently emarginated at tip of CuA. No ventral macrotrichia on veins, dorsal macrotrichia on a short sector of Sc basally to sc-r, entire length of bR, R_1_, second sector of Rs, most of M_1_, distal third of M_2_ and M_4_, distal third of first sector of CuA, and entire second sector of CuA; most Sc, sc-r, first sector of Rs, R_4_, r-m, bM, and CuP entirely devoid of macrotrichia. **Abdomen**. Tergites 1–2 whitish, with a dark brown medial mark; tergites 3–4 cream-yellow with a medial brown mark and a pair of brown marks along laterals, tergite 5 dark brown with a pair of ochre-yellow marks on anterolateral corners, tergite 6 ochre-yellow, with anterior and posterior medial brown marks and a thin lateral brown mark along lateral margin, tergite 7 ochre-yellow with a brown mark along posterior margin; tergite 7 with no ornamentation. Sternites 1–4 whitish, sternites 5–7 more ochre-yellow. **Terminalia** (Figs. 71A–B).

Gonocoxites medially fused at anterior end of terminalia, a suture clearly present, a short, hook-like projection extending slightly beyond base of gonostylus. Gonostylus at lateroposterior end of gonocoxite, bifid, with both branches digitiform, with fine setae along most of its length. Aedeagal plate wide, rectangular, with two pairs of sclerotized short projections directed inwards. Parameres large, projecting beyond tip of gonostylus, with a short digitiform lobe directed outwards and a lobe directed inwards with a strong, sclerotized tooth. Tergite 9 with a pair of slender spines directed inwards close to apex.

**Female**. As male, except for the following. **Wing** (Fig. 70C). Length, 2.95; width, 1.05. **Terminalia** (Fig. 70D). Light ochre-yellow, with some brown sclerites. Sternite 8 with an anterior plate weakly sclerotized, and only with microtrichia, lateroposterior corner with a short rounded lobe, a pair of lateroanterior long, oblique extensions directed dorsally to meet lateroanterior extensions of tergite 8, medially with an elongate plate covered with microtrichia and short setae, four stronger setae on posterior margin. Sternite 9 weakly sclerotized, especially on anterior end, a pair of winglets medially, laterally to an elongate genital chamber. Tergite 8 wide, medially on anterior margin a shallow U-shaped incision, laterally an extension towards ventral face of terminalia, covered only with microtrichia, no setae. Tergite 9+10 slender medially, lateral ends slightly projected on posterior corners, covered with microtrichia and a row of elongate setae along posterior margin. Sternite 10 with microtrichia and setae along posterior margin, medially an elongate transparent window. Cerci elongate, dorsoventrally compressed, cercomere 1 2.5× length of cercomere 2, covered with microtrichia and short setae.

#### Material examined

**Holotype**: male, ZRCBDP0047820, Nee Soon (NS1), swamp forest, 15.may.2013, MIP leg. (slide-mounted). **Paratypes** (42 males, 11 females). **Males**: ZRCBDP0047953, Nee Soon (NS1), swamp forest, 03.jul.13, MIP leg.; ZRCBDP0048475, Nee Soon (NS1), swamp forest, 16.may. 12, MIP leg.; ZRCBDP0048807, Nee Soon (NS1), 14.jan.15, MIP leg.; ZRCBDP0048808, Nee Soon (NS1), 14.jan.15, MIP leg.; ZRCBDP0048813, Nee Soon (NS1), 14.jan.15, MIP leg.; ZRCBDP0048826, Nee Soon (NS1), 14.jan.15, MIP leg.; ZRCBDP0048829, Nee Soon (NS1), 14.jan.15, MIP leg.; ZRCBDP0048830, Nee Soon (NS1), 14.jan.15, MIP leg. (slide-mounted); ZRCBDP0048835, Nee Soon (NS1), 14.jan.15, MIP leg.; ZRCBDP0048838, Nee Soon (NS1), 14.jan.15, MIP leg.; ZRCBDP0048839, Nee Soon (NS1), 14.jan.15, MIP leg.; ZRCBDP0048848, Nee Soon (NS1), 14.jan.15, MIP leg.; ZRCBDP0048869, Nee Soon (NS1), 31.dec.14, MIP leg.; ZRCBDP0048870, Nee Soon (NS1), 31.dec.14, MIP leg.; ZRCBDP0048873, Nee Soon (NS1), 31.dec.14, MIP leg.; ZRCBDP0048879, Nee Soon (NS1), 31.dec.14, MIP leg.; ZRCBDP0048882, Nee Soon (NS1), 31.dec.14, MIP leg.; ZRCBDP0048885, Nee Soon (NS1), 31.dec.14, MIP leg.; ZRCBDP0048889, Nee Soon (NS1), 31.dec.14, MIP leg.; ZRCBDP0048892, Nee Soon (NS1), 31.dec.14, MIP leg.; ZRCBDP0048899, Nee Soon (NS1), 31.dec.14, MIP leg.; ZRCBDP0048919, Nee Soon (NS1), 31.dec.14, MIP leg.; ZRCBDP0048923, Nee Soon (NS1), 31.dec.14, MIP leg. (slide-mounted) (MZUSP); ZRCBDP0048930, Nee Soon (NS1), 31.dec.14, MIP leg.; ZRCBDP0048934, Nee Soon (NS1), 31.dec.14, MIP leg.; ZRCBDP0048959, Nee Soon (NS1), 15.apr.15, MIP leg.; ZRCBDP0048979, Nee Soon (NS1), 13.may.15, MIP leg.; ZRCBDP0048980, Nee Soon (NS1), 13.may.15, MIP leg.; ZRCBDP0049027, Nee Soon (NS2), 07.jan.15, MIP leg.; ZRCBDP0049033, Nee Soon (NS2), 07.jan.15, MIP leg.; ZRCBDP0049092, Nee Soon (NS1), 24.dec.14, MIP leg.; ZRCBDP0049095, Nee Soon (NS1), 24.dec.14, MIP leg.; ZRCBDP0049111, Nee Soon (NS1), 24.dec.14, MIP leg.; ZRCBDP0049207, Nee Soon (NS1), 10.dec.14, MIP leg.; ZRCBDP0049218, Nee Soon (NS1), 10.dec.14, MIP leg.; ZRCBDP0049219, Nee Soon (NS1), 10.dec.14, MIP leg.; ZRCBDP0049235, Nee Soon (NS1), 10.dec.14, MIP leg.; ZRCBDP0049246, Nee Soon (NS1), 10.dec.14, MIP leg.; ZRCBDP0049257, Nee Soon (NS1), 10.dec.14, MIP leg.; ZRCBDP0074027, Bukit Timah, maturing secondary forest (BT08), 28.dec. 16, MIP leg.; ZRCBDP0074034, Bukit Timah, primary forest (BT05), 08.dec. 16, MIP leg.; ZRCBDP0154998, Singapore, date range 2012-2018, MIP leg. **Females**: ZRCBDP0047794, Nee Soon (NS2), swamp forest, 13.nov.13, MIP leg.; ZRCBDP0047795, Nee Soon (NS2), swamp forest, 13.nov.13, MIP leg. (slide-mounted); ZRCBDP0047809, Nee Soon (NS1), swamp forest, 01.may.13, MIP leg.; ZRCBDP0047845, Nee Soon (NS1), swamp forest, 24.apr.13, MIP leg.; ZRCBDP0047937, Nee Soon (NS1), swamp forest, 30.may-05.jun.13, MIP leg.; ZRCBDP0048473, Nee Soon (NS1), swamp forest, 30.may. 12, MIP leg.; ZRCBDP0049039, Nee Soon (NS2), 07.jan.15, MIP leg.; ZRCBDP0048878, Nee Soon (NS1), 31.dec.14, MIP leg.; ZRCBDP0066814, Bukit Timah, maturing secondary forest (BT09), 05.oct. 16, MIP leg.; ZRCBDP0072689, Bukit Timah, maturing secondary forest (BT06), 28.dec. 16, MIP leg.; ZRCBDP0072696, Bukit Timah, primary forest (BT05), 01.dec. 16, MIP leg. **Additional sequenced specimens**: male, ZRCBDP0133907, Singapore, date range 2012-2018, MIP leg; ZRCBDP0133927, Singapore.

**Etymology**. The species epithet refers to Xin Jia Po (新加坡), the Mandarin transcription of Singapore. The noun is used in aposition.

**Remarks**. This is a species with specimens from the Bukit Timah forest and the Nee Soon swamp forest. This is one of the most abundant species of *Neoempheria* found in Singapore, with eleven different haplotypes, all of which come together using any of the delimitation approaches.

*Neoempheria* sp. D (Figs. 72A–D)

https://singapore.biodiversity.online/species/A-Arth-Hexa-Diptera-000780

***Neoempheria_spD_ZRCBDP0049022_hapZRCBDP0049 022_SMH_unnamed_type* [1: A, 16: C, 49: C, 55: A, 56: C, 109: C, 137: C, 139: A, 173: C]**

AttatcttcaactatCgctcacacaggggcttctgttgatttagcaatCttttcACttcatttagcaggtatttcctcaattttaggagctgtaaattttattactacCattattaatatacgagccccaggaattCaAtttgatcgaatacctttatttgtttgatcagttCtaattacagctgttttactattactttctttaccagttttagctggggctattacaatattattaacagatcgtaatttaaatactagtttttttgaccctgcaggaggaggagatcctatcctttaccaacatttattt

#### Description

**Female** (Fig. 72A). Wing length, 3.61; width, 1.31. **Head**. Light brown, ochre-yellow posteriorly to vertex, cream-yellow on ventral half of occiput. Antennal scape and pedicel ochre, with a light brown longitudinal band, flagellum dark brown. Face and clypeus light brown. Maxillary palpus brown, distal palpomere lighter, labella whitish. Post-ocellar setae present, no setae on frons beyond line of ocelli. Scape 1.5× pedicel length, flagellomere 1 1.3× flagellomere 2 length, flagellomere 4 1.0× longer than wide. Palpomere 4 1.2× palpomere 3 length, palpomere 5 1.9× palpomere 4 length. **Thorax**. Scutum with ochre-yellow background, five brown longitudinal stripes, an additional slender brown mark laterally above anepisternum. Pleural sclerites whitish with orangish tinge except for a brown mark on antepronotum anteriorly, brown cervical sclerite, a slender ochre-brown mark dorsoposteriorly on laterotergite, and a brown mark across dorsal half of mediotergite. Scutum with a prescutellar bristle medially on dorsocentral line and two prescutellar bristles on lateroposterior corner; scutellum with one pair of bristles, no additional small setae. Antepronotum with two bristles and some additional smaller setae, proepisternum bare. Halter light brown. **Legs**. Coxae whitish with an orangish tinge, femora ochre-yellow with some elongate light brown marks, tibiae light greyish-brown with brown tips, tarsi brown [fore tibiae and tarsi missing]. Hind tibia inner spur 3.5× longer than tibia width at apex. **Wing** (Figs. 72B–C). Membrane with a slender dark brown band across wing at level of sc-r, a brown mark over R_4_ and a brown band across distal third of wing beginning at level of base of medial fork. C extending beyond tip of R_5_ for a fifth of distance to M_1_; Sc reaching C almost at level of mid of cell r1; sc-r well sclerotized, at level of origin of Rs; R_4_ present, anterior end slightly inclined towards base of wing, cell r1 long, anterior margin of cell r1 3.4× longer than length of R_4_; first sector of Rs 1.1× r-m length. False medial vein present, weakly sclerotized. M_1+2_ 4.1× r-m length; bM 7.6× length of first sector of Rs. First sector of CuA 1.1× longer than second sector. M_4_ not depressed on distal half. Cubital pseudovein sclerotized almost to posterior margin; CuP sclerotized to level of basal third of second sector of CuA. Anal fold faint. Wing margin gently emarginated at tip of CuA. No ventral macrotrichia on veins, dorsal macrotrichia on entire length of bR, R_1_ and second sector of Rs, distal ⅔ of M_1_, distal third of M_2_, distal half of M_4_, distal third of first sector of CuA, and entire second sector of CuA; Sc, sc-r, first sector of Rs, R_4_, r-m, bM, and CuP entirely devoid of macrotrichia. **Abdomen**. Tergites 1–6 ochre-yellow with dark brown medial marks, brown mark of tergite 5 extending laterally, tergites 3–5 with a light brown mark along lateral margin; tergite 7 ochre-yellow anteriorly with a diffuse light brown mark along posterior half; no lateral projections on tergite 7. Sternites 1–2 whitish, sternites 3–6 cream-yellow, sternite 7 ochre-yellow. **Terminalia** (Fig. 72D). Dark ochre-yellow, tip of sternite 8 lobes light brown, cerci dark brown. Sternite 8 anterior plate weakly sclerotized, wide, lateroanterior corners extending anteriorly to articulate with tergite 8, medio-posteriorly a wide medial subquadrate plate with a medial incision separating a pair of short elongate lateroposterior projections. Sternite 9 with a pair of slender lateral arms. Tergite 8 large, wide, with microtrichia and no setae, lateroanterior corners extending ventrally to meet sternite 8. Tergite 9+10 slender, with a sequence of short protuberance along posterior margin bearing an elongate seta at tip. Sternite 10 extending to level of tip of cercomere 1, with a medial transparent elongate window. Cercomere 1 1.6× longer than cercomere 2, both laterally compressed, densely covered with microtrichia and short setae.

**Male**. Unknown.

**Material examined** (2 females): ZRCBDP0049022, Nee Soon (NS2), 07.jan.15, MIP leg. (website photo specimen, slide-mounted); ZRCBDP0049255, Nee Soon (NS1), 10.dec.14, MIP leg.

*Neoempheria* sp. E (Figs. 73A–C)

https://singapore.biodiversity.online/species/A-Arth-Hexa-Diptera-000811

***Neoempheria_spE_ZRCBDP0049180_hapZRCBDP0049 180_SMH_unnamed_type* [2: T, 40: C, 127: A, 133: T, 1 45: C, 147: G, 187: C, 196: A, 262: A]**

cTtatcctcaacaattgcccatacaggggcttctgtagaCttagctattttttctttacatttagctggtatctcttcaattttaggggctgttaattttattacaacaattattaatatacgagcAccaggTattcaatttgaCcGaatacctttatttgtatgatctgttttaattacagctatCcttttattAttatctttaccagttttagcaggagctattacaatattattaacagatcgtaacttaaatactagAttttttgaccctgctggaggaggagaccctattttataccaacatttattt

#### Description

**Female** (Fig. 73A). **Wing l**ength, 2.26; width, 0.89. **Head**. Light brown, including face and clypeus. Antennal scape and pedicel light brown, flagellum dark brown. Maxillary palpus brown, distal palpomere lighter, labella light brown. Two ocelli placed medially on vertex over black background. Post-ocellar setae present, well-developed, only some few setae on frons laterally, close to eyes. Scape 1.2× pedicel length, flagellomere 1 1.5× flagellomere 2 length, flagellomere 4 0.8× longer than wide. Palpomere 4 1.2× palpomere 3 length, palpomere 5 1.0× palpomere 4 length. **Thorax**. Scutum with ochre-yellow background, five brown longitudinal stripes, plus a pair of slender brown bands laterally; scutellum brown above, light brown laterally and posteriorly. Pleural sclerites whitish, except for a brown mark along antepronotum anteriorly, a brown cervical sclerite, a light brown mark on laterotergite on dorsoposterior end and a light brown mark across mediotergite dorsally. Scutum with a prescutellar bristle medially on dorsocentral line and two prescutellar bristles on lateroposterior corner; scutellum with one pair of bristles and some few additional small setae. Antepronotum with four larger and smaller bristles, and additional small setae. Halter light brown. **Legs**. Coxae whitish-yellow, femora ochre-yellow, tibiae light greyish-brown with brown tips, tarsi brown. Tarsomere 1 of fore leg 0.84× tibia length, 1.8× tarsomere 2 length. Hind tibia inner spur 3.3× longer than tibia width at apex. **Wing** (Fig. 73B). Membrane with dark brown marks over sc-r, first sector of Rs, and R_4_, a light brown mark on distal end of anal lobe and a band across distal third of wing at level of base of M_1+2_, besides some brownish areas on cell bR and cell bM. C extending beyond tip of R_5_ for a fifth of distance to M_1_; Sc complete, reaching C barely beyond origin of Rs, sc-r at level of origin of Rs, cell r1 large, over 3.4× longer than wide; first sector of Rs 1.2× r-m length. False medial vein present, slightly arched. M_1+2_ 4.1× r-m length; bM 6.7× length of first sector of Rs. First sector of CuA 1.6× longer than second sector. M_4_ with a strong basal curvature, gently depressed on distal half. Cubital pseudovein sclerotized to level of distal third of CuA; CuP sclerotized to level of basal third of second sector of CuA. Anal fold faint. Wing margin very gently emarginated at tip of CuA. No ventral macrotrichia on veins, dorsal macrotrichia on entire length of bR, R_1_, second sector of Rs, distal third of M_1_, M_2_ and M_4_, and entire second sector of CuA; Sc, sc-r, first sector of Rs, R_4_, r-m, bM, first sector of CuA, and CuP entirely devoid of macrotrichia. **Abdomen**. Tergites 1–2, 4 and 6 whitish with dark brown medial marks, tergite 3 and 5 cream-yellow with a brown mark medially and along posterior margin, tergites 3–5 with a pair of slender brown bands along laterals, tergite 5 brown with a pair of ochre-yellow marks on anterolateral corners, tergite 7 ochre-yellow with a brown mark medially and along posterior margin; no ornamentation on tergite 7. Sternites 1–6 whitish, sternite 7 ochre-yellow. **Terminalia** (Fig. 73C). Light ochre-yellow, with tip of sternite 8 lobes light brown, cerci dark brown. Sternite 8 anterior wide band with microtrichia but no setae, a pair of rounded lateroposterior lobes, medial posterior sclerite subquadrate, elongate, with a short incision medially separating a pair of pointed projections, lateroanterior end extending towards articulation with tergite 8. Sternite 9 with a wide genital chamber, short lateral arms. Tergite 8 wide, with elongate lateroanterior projections, covered with microtrichia but no setae. Tergite 9+10 slender, a group of short protuberances each with an elongate seta at tip. Sternite 10 with a pair of short lateral lobes on posterior margin, transparent window wide midway to apex. Cerci laterally compressed, cercomere 1 1.4× longer than cercomere 2, both covered with microtrichia and short setae.

**Male**. Unknown.

**Material examined** (female): ZRCBDP0049180, Nee Soon (NS2), swamp forest, 13.may.15, MIP leg. (slide-mounted).

*Neoempheria* sp. F (Figs. 74A–D)

https://singapore.biodiversity.online/species/A-Arth-Hexa-Diptera-000776

***Neoempheria_spF_ZRCBDP0047902_hapZRCBDP0047 836_SMH_unnamed_type* [19: A, 23: A, 25: A, 139: T, 1 52: C, 196: A, 197: C, 199: T, 223: C, 233: C]**

tttatcttcaacaattgcAcatAtAggagcttctgtagatttagcaattttttctttacatttagcaggtatttcttctattttaggagcagttaattttattacaacaattatcaatatacgagctccaggaattcaTtttgatcgtataCctctatttgtttgatctgttttaattacagctattttattattACtTtctttacctgttttagcaggagcCattacaataCtattaacagatcgaaacttaaatacaagtttttttgatccagcaggagggggtgatcctattctatatcaacatttattc

#### Description

**Female** (Fig. 74A). Wing length, 3.80; width, 1.44. **Head**. Light greyish-brown, ochre-yellowish on frons dorsally to base of antennae, cream-yellow along ventral margin of occiput. Face light brown, yellowish on ventral end, clypeus light brown, yellowish on dorsal end. Antennal scape and pedicel ochre, with a light brown ventral longitudinal band, flagellum dark brown. Maxillary palpus brown, labella cream-yellow. Post-ocellar setae present, only some few setae on frons laterally. Scape 1.2× pedicel length, flagellomere 1 1.0× flagellomere 2 length, flagellomere 4 1.5× longer than wide. Palpomere 4 1.2× palpomere 3 length, dorsal tip projecting slightly beyond base of palpomere 5, palpomere 5 2.1× palpomere 4 length. **Thorax**. Scutum with ochre-yellow background, two pairs of brown stripes, plus a slender brown medial band, scutellum brown, ochre-brown laterally. Pleural sclerites whitish with an orangish tinge except for a brown mark on antepronotum anteriorly, a brown cervical sclerite, and a slender ochre-brown band dorsally across mediotergite dorsally. Halter light brown. Scutum with a pair of prescutellar bristles medially on dorsocentral lines, one pair of prescutellar bristles on lateroposterior corner; scutellum with one pair of bristles and additional fine setae. Antepronotum with three larger bristles, additional larger and smaller setae. **Legs**. Coxae whitish with an orangish tinge, femora ochre-yellow with some longitudinal light brown marks, tibiae light greyish-brown, with brown tips, tarsi brown. Fore leg tarsomere 1 0.85× tibia length, 2.1× tarsomere 2 length. Hind tibia inner spur 3.8× longer than tibia width at apex. **Wing** (Figs. 74B–C). Membrane with dark brown marks over sc-r, first sector of Rs, and R_4_, and a greyish brown band across distal fourth of wing beyond level of base of medial fork and posteriorly to distal half of M_4_. C extending beyond tip of R_5_ for a fifth of distance to M_1_; Sc reaching C slightly before level of mid of cell r1, sc-r reaching R_1_ beyond origin of Rs for about half of length of first sector of Rs; R_4_ present, almost transverse, cell r1 large, anterior margin × length of R_4_; first sector of Rs 1.6× r-m length. False medial vein present, slightly sinuous on basal third; M_1+2_ 5.6× r-m length; bM 9.8× length of first sector of Rs. First sector of CuA 1.1× longer than second sector. M_4_ gently depressed on distal half. Cubital pseudovein weakly sclerotized, ending at level of distal third of second sector of CuA; CuP reaching level of mid of second sector of CuA. Anal fold faint. Wing margin gently emarginated at tip of CuA. No ventral macrotrichia on veins, dorsal macrotrichia on distal third of Sc, on entire length of bR, R_1_, second sector of Rs, distal two thirds of M_1_, distal fourth of M_2_, distal third of M_4_, distal third of first sector, and entire second sector of CuA; first sector of Rs, r-m, bM, and CuP entirely devoid of macrotrichia. **Abdomen**. Tergite 1 light ochre-yellow, tergites 2–4 light ochre-yellow, with brown short band medially on posterior margin, tergite 5 dark ochre-yellow on anterior half and brown on posterior half, tergites 6 mostly dark ochre-yellow, brownish along posterior margin, tergite 7 light brown; no ornamentation on tergite 7. Sternites 1–4 cream-yellow, sternites 5–6 light ochre-yellow, tergite 7 light brownish ochre-yellow. **Terminalia** (Fig. 74D). Brownish ochre-yellow, distal end of sternite 8 darker, cerci light brown. Sternite 8 with a wide anterior sclerite and a slender medial sclerite, a deep medial posterior incision separating a pair of elongate lobes, small setae covering entire sclerite, some longer setae on lobes. Sternite 9 wide, with short anterior arm of furca. Tergite 8 wide, with long lateroanterior extensions covered with microtrichia, but no setae. Tergite 9+10 slender, a group of short protuberances with an elongate seta at tip. Sternite 10 with a pair of short lateral lobes on posterior margin, transparent window wide midway to apex. Cerci laterally compressed, cercomere 1 1.9× longer than cercomere 2, both covered with microtrichia and short setae.

**Male**. Unknown.

**Material examined** (3 females): ZRCBDP0047836, Nee Soon (NS2), swamp forest, 29.may.13, MIP leg. (slide-mounted); ZRCBDP0047902, Nee Soon (NS2), swamp forest, 30.oct.13, MIP leg.; ZRCBDP0048480, Nee Soon (NS2), swamp forest, 11.apr. 12, MIP leg.

*Neoempheria* sp. H (Figs. 75A–E)

**Diagnosis**. Scutum with five longitudinal brown stripes, a brownish line over dorsal half of mediotergite extending on dorsal third of laterotergite. Abdominal tergites 1–2 and 7 whitish, with a caramel-brown transverse mark along posterior margin extending medially to anterior margin; tergites 3–4 and 6 with similar pattern, but with an additional slender brown mark along lateral margins; tergite 5 mostly caramel-brown with a small cream-yellow mark on lateroanterior corners. Wing with a brown mark at distal third of wing and a second band across wing at level of sc-r and first section of Rs, and an additional dark brown mark over R_4_. C extending beyond tip of R_5_ for about a ¼ of distance to M_1_; sc-r at basal third of cell r1; anterior margin of cell r1 long, more than 4× length of R_4_. Gonocoxites with no extension beyond base of gonostylus; gonostylus large, with a dorsal sub-basal digitiform lobe, a line of spines present at inner edge along distal third, three apical strong spines; parameres with three short, pointed projections distally; tergite 9 with a par of distal lobes with a group of short spines directed inwards.

#### Description

**Male**. Wing length, 3.05; width, 1.15. **Head** (Fig. 75A). Brown, lighter on face and clypeus. No setae on frons anteriorly to ocelli. Ocellar setae present. Antennal scape and pedicel light brown, scape 1.6× pedicel length, flagellum light ochre-yellow except for flagellomere 1 cream-yellow, flagellomere 1 1.6× flagellomere 2 length, flagellomere 4 1.6× wider than long. Palpomeres dark brown, last palpomere lighter, palpomere 4 about 1.2× palpomere 3 length, a short distal projection over base of distal one, last palpomere 2.0× length of penultimate palpomere. Labella small, brownish. **Thorax**. Scutum brown, darker along longitudinal lines; a row of longer dorsocentral setae, some long supra-alars, two pre-scutellar bristles at each lateroposterior corner of scutum, one pair of prescutellars medially at dorsocentral lines. Scutellum brown, with one pair of bristles and some smaller setae on disc. Pleural sclerites whitish with an orangish tinge, antepronotum with a brown mark anteriorly, laterotergite with a brown band along dorsal margin, mediotergite with a brown transverse band on dorsal third. Antepronotum with two strong bristles and smaller setae of different sizes. Halter light brown. **Legs**. Coxae whitish, mid and hind coxae with a light brown tinge, hind coxa slightly darker; femora yellowish-brown, fore femur lighter; tibiae and tarsi light greyish-brown. Hind tibial spurs 3.5× tibia length at apex. **Wing** (Fig. 75B). Membrane background light greyish fumose, a brown band across wing at level of origin of Rs and a brown band at distal third of wing beginning at level of base of medial fork, dark brown mark over first sector of Rs and over R_4_. Sc complete, reaching C at level of basal fourth of cell r1, sc-r just beyond level of origin of Rs; cell r1 long, anterior margin 4.4× length of R_4_. False medial vein conspicuous, sclerotized, sinuous on basal fourth. Medial fork not wide open. M_1+2_ 4.6× r-m length; bM 8.7× length of first sector of Rs. First sector of CuA 1.1× longer than second sector. M_4_ not depressed on distal half. Cubital pseudovein sclerotized, long, ending at distal third of second sector of CuA; CuP weakly sclerotized but produced to level of mid of second sector of CuA. Anal fold faint. Wing margin slightly emarginated at tip of CuA. No ventral macrotrichia on veins, dorsal macrotrichia on entire length of Sc, bR, R_1_, second sector of Rs, most of length of M_1_, distal third of M_2_, distal half of M_4_, long distal half of first sector of CuA, and entire length of second sector of CuA. **Abdomen** (Fig. 75C). Abdominal tergites 1–2 and 7 whitish, with a caramel-brown transverse mark along posterior margin extending medially to anterior margin; tergites 3–4 and 6 with similar pattern, but with an additional slender brown mark along lateral margin; tergite 5 mostly caramel-brown with a small cream-yellow mark on lateroanterior corners. Sternites 1–4 and sternites 6–7 cream-yellow, sternite 5 brown. **Terminalia** (Figs. 75D–E). Ochre-yellow, cerci lighter. Gonocoxites medially fused, with a short, weak medial suture, a weakly sclerotized medio-posterior process extending towards aedeagus, no lateral projections extending beyond base of gonostylus, entirely devoid of setae, some scattered microtrichia laterally. Gonostylus placed laterally, wide at base, with a dorsal digitiform branch bearing elongate setae distally, a large, elongate ventral branch with fine setae on outer face and a row of strong setae and fine spines distally on ventrodistal margin. Aedeagus tubular anterior half, bearing a pair of short apodemes laterally on anterior end, and a medial sub-rectangular plate extending posteriorly to slightly beyond level of mid of gonostylus. Parameres present as a pair of sub-medial weakly sclerotized laminar projections and a wide plate posteriorly to aedeagus with a medial short beak and a pair of short digitiform lobes on lateroposterior corners. Tergite 9 with a slender medial blade connecting a pair of long projections, each of which bearing a short row of long setae laterally and a group of setae and spines directed inwards at distal end. Sternite 10 weakly sclerotized, subquadrate, covered with microtrichia and fine setae. Cerci small, close together, with microtrichia and fine setae, laterally to sternite 10.

**Female**. Unknown.

**Material examined**: male, ZRCBDP0066766, Bukit Timah, maturing secondary forest (BT09), 22.Sep.2016, MIP leg. (slide-mounted).

**Remarks**. This is a species recognized based on the morphology of a sequence failure singleton. It is clearly divergent on the male terminalia morphology, but does not have a sequence and, hence, it is not formally named here.

### Group puluochung

This is one of the species groups in *Neoempheria* that, together with the group *merlio*, seems to be related to *Parempheriella*. *Neoempheria puluochung* Amorim & Oliveira, **sp. n.** has a more standard male terminalia pattern and a medium size cell r4, while *N. merdeka* Amorim & Oliveira, **sp. n.** has a large cell r4 and a scutum with five dark bands, similar to that seen in the group *ferruginea*—and a very unusual male terminalia.

*Neoempheria puluochung* Amorim & Oliveira, sp. n. (Figs. 76A–D, 77A–B)

https://singapore.biodiversity.online/species/A-Arth-Hexa-Diptera-000746

urn:lsid:zoobank.org:act:69DACA24-2425-4B52-804E-19966A2FC57C

**Diagnosis**. Head ochre with a brownish tinge; scutum brownish-ochre, darker on posterior margin, scutellum brown; mediotergite brown, with a brown band running across laterotergite, reaching ventral margin of thorax. Abdominal tergites 1–2 and 4 with cream-yellow background, a brown mark medially along tergites 1–2 and on anterior half medially on tergite 4; tergite 3 and 5 mostly brownish, tergite 3 with a yellowish tinge on lateroanterior corner; tergite 7 mostly yellowish, with a brown median tinge. Wing with oblique brown mark across distal third of wing extending from before tip of R_1_ to posterior margin more basally than tip of M_4_, irregular additional marks more basally, over tip of Sc, cell r1, base of cell m1+2, and on anal lobe close to CuA. Cell r1 rectangular, length of anterior margin 2.2× length of R_4_; sc-r reaching bR at level of origin of Rs. Male gonocoxite entirely bare; gonostylus bifid from base, both branches long, slightly widened at apex, inner lobe with a group of slender spines directed inwards; parameres with a pair of distal blade-like extensions turned inwards midway to apex; tergite 9 with a couple of long, digitiform branches with some short, blunt spines at apex. Female tergite 7 with a pair of digitiform lateroposterior projections reaching level of base of cerci. Female terminalia elongated, sternite 8 with a median short crest between distal lobes.

***Neoempheria_puluochung_ZRCBDP0047867_hapZRC BDP0047867_SMH_holotype* [2: T, 7: C, 25: T, 67: T, 7 3: T, 194: C, 199: T, 202: A, 205: G, 261: G, 274: A]**

tTtatcCtcaacaattgctcatacTggagcatctgttgatttagctattttttctttacatttagcTggaatTtcttctattttaggagctgttaattttattacaacaattattaatatacgagcccctggaattcaatttgatcgaatacctttatttgtttgatctgtattaattacagcagttttattaCttctTtcAttGcctgttttagcaggagctattacaatattattaacagatcgaaatttaaatactaGtttttttgatccAgctggaggaggagacccaattttatatcaacatttattt

#### Description

**Male**. Wing length, 2.95; width, 1.11. **Head**. Light ochre-brown at vertex, more ochre-yellow towards frons and on occiput towards ventral margin. Face cream-yellow, clypeus light brown. Antennal scape and pedicel dark ochre-yellow, flagellum brown, flagellomere 1 with ochre base. Maxillary palpus greyish-brown. Labella ochre-brownish. No post-ocellar setae, frons with only two pairs of setae at level of anterior end of ocelli. Scape as long as pedicel, flagellomere 1 1.5× flagellomere 2, flagellomere 4 1.1× longer than wide. Palpomere 4 1.6× palpomere 3 length, palpomere 5 1.7× palpomere 4 length. **Thorax**. Scutum dark ochre-yellow, more brownish towards posterior end, scutellum ochre-yellowish with a small a brownish mark at anterior margin. Pleural sclerites cream-yellow with an orangish tinge, except for laterotergite, light greyish-brown, darker dorsoposteriorly, and mediotergite brownish on dorsal third. Three pairs of prescutellars, one pair medially, two pairs on lateroposterior corner; one pair of scutellars and some few additional fine setae. Antepronotum with two bristles and some additional long and short setae, proepisternum with one bristle and some few additional setae. **Legs**. Coxae whitish with orangish tinge, anterior face of fore coxa with a brownish tinge on basal half. Femora ochre-yellow, fore and mid femora darker; tibiae and tarsi light greyish-brown. Tibiae with some few dorsal and lateral setae. Hind tibia inner spur 3.0× longer than tibia width at apex. Fore leg tarsomere 1 0.9× tibia length, 1.0× tarsomere 2 length. **Wing** (Fig. 76B). Membrane background light greyish, with dark greyish-brown marks over tip of Sc and sc-r, first sector of Rs, R_4_, from base of M_4_ to posterior margin, and across distal third of wing, beginning at level of base of medial fork. C extending beyond tip of R_5_ for a third of distance to M_1_. Sc complete, reaching C slightly beyond origin of Rs, sc-r at level of origin of Rs, cell r1 medium-sized, anterior margin of cell r1 1.4× longer than length of first sector of Rs. First sector of Rs 2.0× r-m length; R_4_ transverse. False medial vein conspicuous, sclerotized. M_1+2_ 2.7× r-m length; M_2_ slightly curved towards M_1_close to tip; bM 5.6× length of first sector of Rs; M_2_ relatively weak on distal third. Cubital pseudovein extending to level of mid of second sector of CuA. M_4_ gently sinuous medially, curved towards posterior margin at tip. First sector of CuA 1.2× length of second sector. No ventral macrotrichia on veins, dorsal macrotrichia on entire length of bR, R_1_, second sector of Rs, distal fifth of M_1_, and at tip of M_4_ and distal fourth of second sector of CuA; Sc, sc-r, first sector of Rs, R_4_, r-m, bM, M_1+2_, M_2_ and CuP entirely devoid of setation; some few macrotrichia on membrane of cell sc before tip of R_1._ **Abdomen**. Tergites 1–2 light brown medially with cream-yellow lateral bands, tergite 3 light brown with cream-yellow marks on anterolateral corners, tergite 4 largely cream-yellow with a slender, light brown medial band, tergite 5 light brown, with cream-yellow anterior band, tergite 6 cream-yellow with a pair of light brown marks sub-medially, tergite 7 cream-yellow on anterior half, posterior half light brown. Sternites 1–7 cream-yellow. **Terminalia** (Figs. 76C–D). Gonocoxites fused along anterior margin, entirely devoid of setae ventrally, a wide medial, weakly sclerotized triangular plate projecting to level of insertion of gonostylus, a pair of long lateroposterior projections slightly widening to apex, bearing small setae on inner face and large scales on ventral face. Gonostylus long, deeply bifid, outer branch curved at base, slightly clavate, bare on proximal ¾, with three long setae along inner margin distally and a dense group of elongate setulae on dorsal face, inner branch more sclerotized, also clavate, with a number of long setae at outer margin and a concentrated group of setae on inner margin close to apex. Gonocoxal apodemes projected anteriorly, aedeagus with a pair of short apodemes on anterior end, elongate, weakly sclerotized distally. Parameres with a pair of L-shaped digitiform lateroposterior projections, distal part directed inwards, with some setulae. Tergite 9 lateroanterior ends articulating to gonocoxites dorsally to insertion of gonostylus, medially along anterior margin with a wide, inverted V-shaped incision, extending posteriorly to almost tip of outer branch of gonostylus, a straight long seta and one very long, curved seta along lateral margin close to base at each side, ventral face of lateroposterior corner extending into a digitiform process almost reaching level of tip of cerci, bearing some long setae on distal half directed ventrally and three short spines directed dorsally. Sternite 10 weakly sclerotized, bare, with a medial rounded posterior incision separating a pair of short lobes. Tergite 10 wider than both cerci, bare, with a medial deep incision on posterior margin. Cerci slender, in contact with each other, placed very distally, entirely covered with dense microtrichia, short setae on dorsal face, setae on distal end longer.

**Female** (Fig. 76A). As male, except for the following. Tergite 7 highly modified, short medially, with a pair of digitiform expansions laterally reaching level of base of cerci. Sternites 1–8 ochre-yellow. **Terminalia** (Figs. 77A– B). Mostly ochre-yellow, tergite 8 light brown. Sternite 8 trapezoid, slender distal end with a medial incision separating a pair of slender lobes, microtrichia scattered over entire sclerite, setae restrict to tip of lobes. Sternite 9 with a wide anterior end, a pair of sclerotized arms medially. Sternite 10 rectangular, with a large transparent window. Tergite 8 bare, wide, more sclerotized laterally; tergite 9+10 wide, slender, with a row of setae along posterior margin, some few short digitiform projections laterally with a distal seta. Cerci elongate, dorsoventrally compressed, covered with short setae and microtrichia, cercomere 1 wider midway to apex, 3.2× cercomere 2 length, cercomere 2 slightly ovoid.

#### Material examined

**Holotype**: male, ZRCBDP0047867, Nee Soon (NS2), swamp forest, 02.oct.2013, MIP leg. (slide-mounted). **Paratypes**: 3 females, ZRCBDP0048493, Nee Soon (NS1), swamp forest, 11.apr.2012, MIP leg. (website photo specimen, slide-mounted); ZRCBDP0048494, Nee Soon (NS1), swamp forest, 23.may.2012, MIP leg. (website photo specimen, extracted); ZRCBDP0049005, Nee Soon (NS2), 17.dec.14, MIP leg. (extracted).

**Etymology**. The species epithet refers to the name Pu Luo Chung, the Mandarin transcription [蒲罗中] of Pulau Ujong—in the oldest known written document referring to Singapore, from the third century. The Malay name “Pulau Ujong” means literally “island at the end”. The noun is used in aposition.

**Remarks**. There are three haplotypes for *Neoempheria puluochung* Amorim & Oliveira, **sp. n.,** recognized as a single species by all of delimitation algorithms except PTP, that splits it into two separate species.

*Neoempheria merdeka* Amorim & Oliveira, sp. n. (Figs. 78A–D, 79A–B)

https://singapore.biodiversity.online/species/A-Arth-Hexa-Diptera-002126

urn:lsid:zoobank.org:act:376F6AEB-C10A-46A0-93B6-9E46D612FAE2

**Diagnosis**. Head ochre-yellowish with a brownish tinge; scutum brownish, with ochre-yellow longitudinal stripes, scutellum brown; mediotergite dark brown, a brown band extending over laterotergite, mesepimeron and katepisternum. Abdominal tergite 1 brown medially, cream-yellowish on laterals; tergites 2–3 brown with cream-yellow lateroanterior corners; tergite 4 background ochre-yellowish, with a wide medial brown band along posterior margin and a large brown mark on lateroposterior corner; tergites 5–7 mostly brownish, with an ochre-yellowish band along anterior half. Wing with a complex color pattern, an N-shaped brown mark over posterior ⅔ of wing (extending anteriorly over R_4_) and a basal band from wing margin to cubital pseudovein; cell r1 large, anterior margin of cell r1 3.6× R_4_ length; sc-r reaching R_1_ at basal third of cell r1. Male terminalia complex, gonocoxites largely reduced, bare; gonostylus bifid from base, both extending into long, sigmoid projections; parameres wide distally, projected beyond tip of cerci; tergite 9 with a pair of long, sigmoid branches.

***Neoempheria_merdeka_ZRCBDP0066818_hapZRCBDP 0066818_SMH_holotype* [19: A, 73: C, 76: C, 138: A, 15 4: C, 155: C, 220: T, 274: C, 283: T, 308: C] -**

ctttcatctactattgcAcatacaggagcttctgttgatttagcaattttttctttacatttagcaggtatCtcCtcaattttaggagctgtaaattttattacaacaattattaatatacgagccccaggaattcAatttgatcgaataccCCtatttgtttgatctgtattaattactgcaattttactgctattatcattacctgttttagctggTgctattacaatattattaacagatcgaaacttaaatacaagtttttttgatccCgcaggtggTggtgacccaattttatatcaacatCtattt

#### Description

**Male**. Wing length, 3.87; width, 1.38. **Head**. Brown vertex, ochre-yellowish on occiput, face and clypeus light brown, face with macrotrichia but no setae, clypeus setose. No setae on frons anteriorly to ocelli. Ocellar setae present. Antennal scape light brown, pedicel yellowish-brown, scape 1.7× pedicel length, flagellum light ochre-yellow except for flagellomere 1 cream-yellow, flagellomere 1 1.5× longer than flagellomere 2, flagellomere 4 as long as long wide. Palpomeres dark brown, last palpomere lighter, palpomere 4 about 0.85× palpomere 3 length, a short distal projection over base of distal palpomere, last palpomere 2.2× palpomere 4 length. Labella small, brownish. **Thorax**. Scutum ochre-yellowish, a brown band laterally on posterior fourth and a pair of light brown bands along dorsocentral line; a row of longer dorsocentral setae, some long supra-alars, two pre-scutellar bristles on each lateroposterior corner of scutum, a pair of prescutellars medially on dorsocentral lines. Scutellum ochre-yellowish, light brownish along anterior margin, with one pair of longer setae and some smaller setae on disc. Antepronotum, proepisternum, proepimeron, anepisternum, and metepisternum cream-yellow with an orangish tinge, a brown band across katepisternum, mesepimeron, laterotergite, and dorsal half of mediotergite; ventral half of mediotergite ochre-brown. Antepronotum with four stronger bristles and smaller setae of different sizes. Halter yellowish-brown. **Legs**. Anterior coxa with a brown band on dorsal half (in continuation of brown band on pleura), brownish-yellow ventrally with a brownish tinge on distal end, mid and hind coxae whitish, with a brown mark on distal end; anterior femur ochre-yellowish, mid and hind femora brownish-yellow; tibiae and tarsi yellowish-brown, anterior tarsus lighter. Hind tibial spurs 3.8× length of tibia at apex. **Wing** (Fig. 78B). Membrane with light greyish-brown background with a brown mark close to base of wing and a pair of large brown connected marks, one from tip of Sc to posterior margin across sc-r, origin of Rs, M_1+2_ and origin of M_4_, and another from distal end of R_1_ to posterior margin, with a light area over distal fourth of R_5_. Sc complete, reaching C close to level of origin of Rs; sc-r present, reaching R_1_ almost at same level than tip of Sc; anterior margin of cell r1 long, anterior margin 3.6× length of R_4_. False medial vein present, not strongly sclerotized, gently sinuous on basal fifth. Medial fork not wide open. M_1+2_ 5.0× r-m length; bM 8.8× r-m length. First sector of CuA 1.3× longer than second sector. M_4_ not depressed on distal half. Cubital pseudovein sclerotized, long, ending at distal third of second sector of CuA; CuP sclerotized, produced to level of mid of second sector of CuA. Anal fold faint. Wing margin slightly emarginated at tip of CuA. No ventral macrotrichia on veins, dorsal macrotrichia on distal half of Sc, on entire length of bR, R_1_, second sector of Rs, most length of M_1_ and of M_2_, distal ¾ of M_4_, and along entire length of CuA. **Abdomen**. Tergite 1 ochre-yellow, tergites 2–5 ochre-yellow on anterior half and brown on posterior half, tergite 6 ochre-yellow with a brown mark on lateroposterior corners, tergite 7 ochre-yellow on anterior half and brown on posterior half. Sternites 1–3 and 7 ochre-yellow, sternites 4–6 cream-yellow, with brown bands laterally. **Terminalia** (Figs. 78C–D). Ochre-yellow, cerci lighter. Gonocoxites largely modified, medially fused, with a conspicuous suture, anterior margin of syngonocoxite with a medial anterior incision, posterior margin also with a medial incision, a short digitiform lateral projection slightly beyond base of gonostylus, no setation or microtrichia at all. Gonostylus slightly displaced medially, bifid, with a pair of long, sinuous, digitiform sclerotized extensions, inner one almost reaching level of distal margin of parameres, outer one longer, extending beyond level of tergite 10 lateral projections, no setae at all except for a terminal short seta on outer branch. Gonocoxal bridge conspicuous, with an anterior medial acute extension. Aedeagus sclerite complex, with a pair of short sclerotized arms. Parameres present as a pair of weakly sclerotized laminar blades posteriorly to aedeagus. Tergite 9 with a blade medially connecting a pair of long sinuous projections widening distally, bifid at apex, with some long fine setae along inner face of distal half. Sternite 10 weakly sclerotized, with a pair of lateral lobes covered with microtrichia and fine setae. Cerci small, close together, with microtrichia and fine setae, laterally to sternite 10.

**Female** (Fig. 78A). As male, except as follows. **Terminalia** (Figs. 79A–B). Sternite 8 trapezoid, with slender distal end, a medial incision on posterior margin, small setae scattered over entire sclerite, slightly stronger distally. Sternite 9 with a wide anterior end, a pair of sclerotized arms on posterior margin. Sternite 10 rectangular, elongate. Tergite 8 bare, wide, tergite 9+10 slender, with a row of setae along posterior margin, some at tip of short digitiform projections. Cerci elongate, dorsoventrally compressed, covered with short setae and microtrichia, cercomere 1 3.0× cercomere 2 length, cercomere 2 rounded, slightly ovoid.

#### Material examined

**Holotype**: male, ZRCBDP0066818, Bukit Timah, maturing secondary forest (BT09), 05.oct. 2016, MIP leg. (slide-mounted). **Paratypes** (2 females): ZRCBDP0120496, Bukit Timah Forest (BTNR), date range 2012-2018, MIP leg., date range 2012-2018, MIP leg.; ZRCBDP0155081, Nee Soon (NS1), 03.dec.14, MIP leg. **Additional sequenced specimens**: female, ZRCBDP0133981. **Sequencing failures:** ZRCBDP0074035, Singapore, date range 2012-2018, MIP leg.

**Etymology**. The species epithet refers to the Malay word merdeka [=independence/freedom], the name of the movement after the post-war period that resulted in the gradual increase of self-governance for Singapore and a separate Legislative Council, elected in March 1948. The noun is used in apposition.

**Remarks**. There are three haplotypes for this species, both with specimens from Bukit Timah. OC separates them into two species at the level of 2%. All other approaches point to a single species. This is a very unusual species of *Neoempheria*. The large r1 cell and the longitudinal stripes in the wing suggest that it belongs in the *ferruginea*-group. Some other features, however, are shared with the species of *Parempheriella* and other *Neoempheria* species with a small cell r1. The male terminalia are also very unique.

### Group dizonalis

The group *dizonalis* includes seven species and may be the closest to *Parempheriella*. All species have a small cell r4, the first sector of Rs basically transverse, a quite similar pattern of abdomen color (with tergite 4 mostly or entirely yellow) and male terminalia with well-developed tergite 9, often with a distal spine. Some of the described Oriental species of *Neoempheria* possibly belong here.

*Neoempheria dizonalis* (**Edwards) (**Figs. 80A–D)

https://singapore.biodiversity.online/species/A-Arth-Hexa-Diptera-000798

*Mycomya* (*Neoempheria*) *dizonalis* Edwards 1931: 265 (fig. 2v, d, male terminalia). Type locality: Indonesia, Sumatra, Fort de Kock [= Bukittinggi]. Colless & Liepa 1973: 458), new combination.

**Diagnosis**. Head light ochre-yellow; scutum light ochre-yellow, no stripes; pleural sclerites whitish, mediotergite and laterotergite brown. Wing with brown band distally across wing and a brown band more basally, from level of cell r1 to posterior margin; cell r1 small, anterior margin 1.1× length of R_4_; sc-r reaching bR well basal to origin of Rs. Tergite 1 cream-yellowish; tergite 2 cream-yellow with a dark brown medial mark on posterior half medially; tergites 3, 5–6 mostly dark brown, with slender cream-yellow lateroposterior bands; tergite 4 cream-yellow; tergite 6 dark brown; tergite 7 ochre-yellowish; sternite 6 with a large brown mark laterally. Gonocoxites with no lateroposterior extensions beyond insertion of gonostylus; gonostylus with three branches, more ventral branch digitiform, long, extending beyond tip of cercus; tergite 9 large, with a pair of posterior strong arms, each with a distal spine directed inwards.

***Neoempheria_dizonalis_ZRCBDP0047918_hapZRCBD P0047800_SMH_describedearlier_type* [10: A, 55: C, 76: A, 124: G, 127: A, 154: C, 155: C, 262: C] -**

ttatcatcAacaattgctcatacaggggcatcagtagatttagctattttttcCctacatttagcaggtatttcAtctattttaggagcagttaattttattacaacaattattaatatacgGgcAcctggaattcaatttgatcgaataccCCtatttgtatgatcagttttaattacagcaattcttttacttctttctttacctgtattagcaggagcaattactatactattaacagatcgtaatttaaatacaagCttttttgatccagctggaggaggggatcctattttatatcaacatttattt

**Redescription**. **Male** (Fig. 80A). Wing length, 2.79; width, 1.05. **Head**. Dark ochre-yellow at vertex, lighter above antennae and on ventral half of occiput. Face whitish-yellow, clypeus whitish-yellow. Antennal scape and pedicel light ochre-yellow, flagellum ochre-yellow, slightly darker toward apex. Maxillary palpus light brown, last palpomere lighter, labella light ochre-yellowish. Ocellar setae present, no setae on frons anteriorly to line of ocelli. Scape 1.2× pedicel length, flagellomere 1 1.3× flagellomere 2 length, flagellomere 4 1.1× longer than wide. Palpomere 4 1.2× palpomere 3 length, palpomere 5 2.0× palpomere 4 length. **Thorax**. Scutum mostly ochre-yellow, slightly darker medio-posteriorly; scutellum ochre-yellow. Pleural sclerites whitish with an orangish tinge, laterotergite light brown, ventroanterior end lighter, mediotergite brown, ochreish laterally. Scutum with a pair of prescutellar bristles on each dorsocentral line and two prescutellar bristles at each lateroposterior corner; scutellum with one pair of strong setae and some additional small setae. Antepronotum with two bristles, one strong seta and some smaller setae, proepisternum with one bristle and one long seta. Halteres whitish-yellow, brownish at base of knob. **Legs**. Coxae whitish, Fore coxa with some orangish tinge; fore femur whitish-yellow, mid and hind femora darker; tibiae and tarsi light greyish-brown. Hind tibia inner spur 4.7× longer than tibia width at apex. Fore leg tarsomere 1 0.86× tibia length, 1.9× tarsomere 2 length. **Wing** (Fig. 80B). Membrane background light brown fumose, a dark brown band across wing at level of cell r1 reaching posterior margin and a brown mark across distal third of wing beginning slightly beyond basal end of medial fork. C produced beyond tip of R_5_ for about a third of distance to M_1_; Sc complete, reaching C slightly beyond level of origin of Rs, sc-r weakly sclerotized, reaching bR before level of origin of Rs. R_1_ reaching C at distal fourth of wing; R_5_ reaching C slightly before level of tip of M_1_; R_4_ present, close to base of Rs, cell r1 short, trapezoid, anterior margin of cell r1 1.1× longer than R_4_; first sector of Rs 0.83× r-m length. False medial vein conspicuous, sclerotized. M_1+2_ 3.6× r-m length; bM 7.0× length of first sector of Rs. First sector of CuA 1.1× longer than second sector. M_4_ gently depressed on distal half. Cubital pseudovein sclerotized to distal fourth of CuA; CuP sclerotized to basal fourth of second sector of CuA. Anal fold present. Wing margin gently emarginated at tip of CuA. No ventral macrotrichia on veins, dorsal macrotrichia on entire length of bR, R_1_, second sector of Rs, most of M_1_, M_2_, M_4_ and CuA; Sc, sc-r, first sector of Rs, R_4_, r-m, bM, M_2_, M_4_, and CuP devoid of macrotrichia. **Abdomen**. Tergite 1 whitish, tergite 2 brown medially on posterior half, cream-yellow on anterior half and laterally; tergites 3, 5–6 brown, tergites 5–6 with a slender cream-yellow band on lateroposterior corner, tergite 4 and 7 with a slender yellowish band along anterior margin. Sternites 1–5 whitish-yellow, sternite 6 light brown with a yellowish medial area posteriorly, sternite 7 ochre-yellowish. **Terminalia** (Figs. 80C–D). Ochre-yellow, some sclerites more brownish. Gonocoxites fused, a slender connection medially, an inverted deep U-shape incision on anterior margin with sclerotized border, no lateroposterior extensions beyond insertion of gonostylus, entirely bare. Gonostylus with three branches, a more ventral, long digitiform branch extending beyond tip of cercus, with long and short fine setae along entire length of external face, a medial, short digitiform process with a single setula at tip, and a dorsal, shorter ovoid branch with a concentration of fine setae and setulae at tip, base lobose, with setae and setulae. Gonocoxal apodeme large, lateral arms extending anteriorly, medial connection beyond anterior end of terminalia. Aedeagus wide on anterior end, not strongly sclerotized, distal end with a wide opening. Parameres more anteriorly with a pair of lateral digitiform projections almost reaching level of tip of aedeagus, laterodistally with a pair of slightly more sclerotized blades. Sternite 10 weakly sclerotized, with a pair of short lobes sub-medially with microtrichia and small setae. Tergite 9 large, with a pair of lateroanterior weakly sclerotized extensions that reach gonocoxite, a medial wide incision on anterior margin, posterior margin rounded, with a row of longer setae and a pair of posterior strong projections with a spine at tip directed inwards. Tergite 10 with a pair of long, large lobes with a slender medial connection, each lobe with a short ventral sublobe on ventral face with fine setae and a long dorsal, well sclerotized sublobe extending beyond tip of gonostylus, densely covered with long setae on dorsal face and a spine at tip directed inwards. Cerci large, elongate, almost reaching tip of tergite 10 projections, covered with microtrichia and setae.

**Female**. Unknown.

#### Material examined

**Holotype** (as *Mycomya (Neoempheria*) *dizonalis* Edwards, 1925): male, “Sumatra, Fort de Kock, 1925” (NHM). **Sequenced specimens** (3 males): ZRCBDP0047800, Nee Soon (NS2), swamp forest, 13.nov.13, MIP leg. (slide-mounted); ZRCBDP0047918, Nee Soon (NS2), swamp forest, 27.nov.13, MIP leg. (website photo specimen); ZRCBDP0049043, Nee Soon (NS2), 07.jan.15, MIP leg.

**Remarks**. We examined the type of *Neoempheria dizonalis* at the NHM and it largely agrees with our specimen.

*Neoempheria neesoon* Amorim & Oliveira, sp. n. (Figs. 81A–C)

https://singapore.biodiversity.online/species/A-Arth-Hexa-Diptera-000744,and-000793

urn:lsid:zoobank.org:act:4840D933-A430-4C04-BC62-EAD748E762DD

**Diagnosis**. Head and scutum dark brown, no stripes; pleural sclerites ochre-yellowish, mediotergite light brown, laterotergite with a light greyish brown tinge on dorsoposterior third. Tergite 1 ochre-yellowish; tergite 2 brownish medially; tergite 3 brownish with a slender anterior ochre-yellowish band; tergites 2–4 ochre-yellowish with a brown medial mark along posterior margin; tergites 5–6 brownish, with an ochre-yellowish slender band at lateroanterior corner. Wing with brown band distally across wing and a couple of brown marks more basally, one over cell r1, r-m, cell m1+2 close to injunction of bM, and M_1+2_, and another one at same level on cell cup; cell r1 short, anterior margin 0.94× length of R_4_; sc-r reaching bR well basal to origin of Rs. Female tergite 7 with a lateroposterior projection; terminalia sternite 8 with a pair of distal large posterior lobes widening towards apex, sternite 10 with transparent medial “window”.

***Neoempheria_neesoon_ZRCBDP0047823_hapZRCBDP 0047823_SMH_holotype* [34: C, 64: T, 67: T, 136: C, 14 2: C, 155: C, 184: C]**

tctatcttctacaattgctcatacaggggcatcCgttgatttagctattttttcattacatctTgcTggaatttcttcaattttaggagctgtaaattttattacaacaatcattaatatacgagccccaggaatCcaattCgaccgaatacctCtatttgtatgatccgttttaattacagcCattttattacttttatctttaccagttttagctggagctattacaatattattaacagatcgaaatttaaatacaagattttttgatcctgcaggagggggggaccctatcttatatcaacacttattt

#### Description

**Female** (Fig. 81A). Wing length, 2.43–3.02, width, 0.85–1.08 (n=3). **Head**. Cream-yellow, lighter at frons and ventrally at occiput. Some scattered short setae over frons and occiput, some slightly longer setae around eye on occiput. Two ocelli placed medially on vertex over a blackish background. Ocellar setae absent. Face whitish. Antennal scape and pedicel cream-yellow, scape with setulae around distal margin, pedicel with one longer seta besides crown of short setae distally. Flagellum light ochre-yellowish, flagellomere 1 slightly longer than flagellomere 4, flagellomere 4 1.1 wider than long. Mouthparts short. Clypeus light brown, triangular, with scattered setae. Four light brown palpomeres, last palpomere lighter on distal half, palpomere 4 not projected beyond base of distal one, last palpomere 2.1× length of penultimate palpomere. **Thorax**. Scutum ochre-yellow, an irregular row of longer dorsocentral and acrostichal setae, as well as some long supra-alars. Scutellum with one pair of longer setae and few smaller setae. Pleural sclerites cream-yellow with an orangish tinge except for light greyish-brown on most laterotergite and a greyish-brown mediotergite. One strong seta on proepisternum, besides smaller setae on both sclerites. **Legs**. Coxae whitish, femora ochre-yellow, tibiae and tarsi light greyish-brown. Fore coxa with scattered setae on anterior face, with some additional longer ones more laterally and ventrally, mid coxa with some longer greyish setae at distal end, hind coxa with a long row of grey setae along most of its length. Femora with scattered brownish short setae all over and some additional ventral longer setae close to distal end. Tibiae with some slightly longer setae dorsally, laterally and ventrally; tibiae and tarsi with dark setulae arranged in regular rows. Tibial spurs subequal, spur over 3× longer than tibia width at apex. First tarsomeres of hind leg with a row of longer setae ventrally, all tarsomeres with a pair of distal ventral setae. Tarsomere 1 of fore leg 0.8× length of tibia and twice tarsomere 2 length. **Wing** (Fig. 81B). Membrane background light greyish, with a dark greyish-brown band across wing at level of tip of R_1_ and a dark greyish-brown mark at tip of wing beginning at level of medial fork. Sc complete, reaching C at level of origin of Rs, sc-r well basal to origin of M_1+2_, cell r1 short, anterior margin 0.94× length of R_4_. False medial vein conspicuous, slightly sclerotized. Medial fork wide open. Origin of M_4_ at level of base of M_1+2_. **Abdomen**. Tergite 1 cream-yellow, tergites 2, 5 and 6 with a cream-yellow band anteriorly (wider on tergite 2), tergite 4 and 7–8 cream-yellow mostly cream-yellowish, tergite 4 with a median brown mark at posterior margin. Sternites cream-yellowish. Sternite 7 with a pair of short lateroposterior projections. **Terminalia** (Fig. 81C). Cream-yellow, with slightly darker sclerites. Posterior lobes of sternite 8 widening towards apex, a fringe of setae along posterior margin, a short incision medially. Sternite 9 with a pair of well sclerotized apodemes at anterior end. Sternite 10 with transparent window. Tergite 8 wide, with a pair of short lateroposterior projections. Tergite 9+10 short, slender, with a pair of small lateral lobes. Cercus elongate, cercomere 1 almost twice as long as cercomere 2.

**Male**. Unknown.

#### Material examined

**Holotype**: Female, ZRCBDP0047823, Nee Soon (NS1), swamp forest, 14.aug.2013, MIP leg. (slide-mounted). **Paratypes** (8 females): ZRCBDP0047955, Nee Soon (NS2), swamp forest, 18.set.13, MIP leg. (slide-mounted) (MZUSP); ZRCBDP0048486, Nee Soon (NS1), swamp forest, 18.apr. 12, MIP leg. (website photo, slide-mounted); ZRCBDP0048906, Nee Soon (NS1), 31.dec.14, MIP leg. (slide-mounted); ZRCBDP0049171, Nee Soon (NS2), 13.may.15, MIP leg. (slide-mounted); ZRCBDP0049176, Nee Soon (NS2), 13.may.15, MIP leg. (website photo specimen, slide-mounted); ZRCBDP0049243, Nee Soon (NS1), 10.dec.14, MIP leg. (slide-mounted); ZRCBDP0066792, Bukit Timah, primary forest (BT05), 11.apr.13, MIP leg; ZRCBDP0154986, Nee Soon (NS2), 14.jan.15.

**Etymology**. The specific epithet for this species refers to the Nee Soon Freshwater Swamp Forest – named after Lim Nee Soon (1879-1936), a prominent Chinese businessman and respected community leader in Singapore. Freshwater swamp forests (or flooded forests) are forests permanently or seasonally inundated with freshwater. They are known from different parts of the globe—not only in tropical areas—and are an important component of the forests in Southeast Asia. There was loss of the original areas of swamp forest in Singapore since the early 1800s due to land conversion for cultivation, e. g., of pepper, pineapple and rubber. Canalization, residential and industrial development wiped out fragments left from the previous use of these areas for agriculture. The Nee Soon Freshwater Swamp Forest is one of the most important protected fragments of this kind of habitat in Singapore. The noun is used in apposition.

**Remarks**. There are seven haplotypes for *Neoempheria neesoon* Amorim & Oliveira, **sp. n.** Two haplotypes differ by 3.5% (ZRCBDP0047823 and ZRCBDP0048486).

Several molecular species delimitation algorithms suggest two mOTUs within the limits of one morphological species. There are specimens from the swamp forest and from the Bukit Timah forests. There is enough information to prepare a diagnosis for the species, although only females in our samples.

*Neoempheria pulau* Amorim & Oliveira, sp. n. (Figs. 82A–G)

https://singapore.biodiversity.online/species/A-Arth-Hexa-Diptera-000743

urn:lsid:zoobank.org:act:F4107048-0B6E-4F8A-82DF-B0543A1C84C6

**Diagnosis**. Head light ochre-yellow, with a light brown diffuse mark over vertex; scutum light ochre-yellow, no stripes; pleural sclerites whitish, mediotergite light brown, laterotergite with a small light brown mark at dorsoposterior end. Tergites 1–2 and 4 cream-yellow; tergites 3, 5–6 dark brown with a slender cream-yellow lateroposterior band; tergite 7 whitish-yellow; sternite 6 with a large brown mark laterally. Wing with brown band distally across wing and a brown band more basally, from level of cell r1 to posterior margin. Cell r1 small, anterior margin 0.86× R_4_ length; sc-r reaching bR well basal to origin of Rs. Gonocoxites with no laterodistal projections; gonostylus large, with a basal peduncle expanding distally into a large, conch-like structure partially folded; tergite 9 with a pair of long, sclerotized bottle-like structures laterodistally, longer than gonostylus.

***Neoempheria_pulau_ZRCBDP0047805_hapZRCBDP00 47805_SMH_holotype* [40: C, 91: C, 155: C, 163: C, 259 : C, 283: T, 298: C] --**

tatcttctacaattgcccatactggagcatctgttgaCttagctattttttcattacatttagctggtatttcttcaattttaggagcCgttaattttattacaactattattaatatacgagccccaggaattcaatttgatcgaatacctCtatttgtCtgatctgtattaattactgcaattcttcttttattatctttaccagttttagctggagctattacaatattattaacagatcgtaacttaaatacCagattttttgaccctgctggaggTggagaccctattctCtatcaacacttattt

#### Description

**Male** (Fig. 82A). Wing length, 2.92; width, 1.11. **Head**. Light brown at vertex, more cream-yellow on occiput towards ventral margin. Antennal scape and pedicel cream-yellow, flagellum light ochre-yellowish. Face and clypeus whitish-yellow. Maxillary palpomeres 2–4 brown, palpomere 5 light brown. Labella whitish-yellow. Scape as long as pedicel, flagellomere 1 1.2× flagellomere 2 length, flagellomere 4 only slightly longer than wide. Palpomere 4 1.6× palpomere 3 length, palpomere 5 1.8× palpomere 4 length. **Thorax**. Scutum and scutellum dark ochre-yellow. Pleural sclerites dark cream-yellow except for light greyish-brown laterotergite dorsally and a greyish-brown mediotergite. Two pairs of prescutellars, one pair medially, one pair on lateroposterior corner; one pair of scutellars and some few additional fine setae. Antepronotum with three bristles and scattered smaller setae, proepisternum bare. **Legs**. Coxae whitish with orangish tinge, femora ochre-yellow, tibiae and tarsi light greyish-brown. Hind tibia inner spur 3.9× longer than tibia width at apex. Fore leg tarsomere 1 1.0× tibia length, 1.2× tarsomere 2 length. **Wing** (Fig. 82C). Membrane light greyish with a greyish-brown band across distal third of wing beginning before R_1_ and a band across wing at level of tip of Sc. C extending beyond tip of R_5_ for about a ¼ of distance to tip of M_1_. Sc weakly sclerotized at very tip, slightly beyond level of R_4_, sc-r way before level of origin of Rs. R_1_ reaching C at distal fifth of wing; R_5_ reaching C before level of tip of M_1_; cell r1 short, length of anterior margin 0.8× length of first sector of Rs, posterior end even shorter, cell almost triangular; first sector of Rs short, 0.76× r-m length. False medial vein conspicuous, slightly sclerotized. Medial fork wide open, M_1+2_ 3.4× r-m length; bM 8.2× length of first sector of Rs; M_4_ gently sinuous on distal half. First sector of CuA 1.1× length of second sector. Cubital pseudovein well sclerotized, ending close to posterior wing margin. CuP long, almost reaching level of second sector of CuA. No ventral macrotrichia on veins, dorsal macrotrichia on entire length of bR, R_1_, second sector of Rs, most length of M_1_, distal half of M_2_, distal ⅔ of M_4_, distal half of first sector of CuA, and entire second sector of CuA; sc-r, Sc, R_4_, first sector of Rs, r-m, bM, M_1+2_, and CuP devoid of setation; membrane entirely devoid of setae. **Abdomen**. Tergite 1 cream-yellow, tergites 2–3 and 5–6 with a cream-yellow band anteriorly, a brown band posteriorly, and a slender cream-yellow band along posterior margin, wider cream-yellow band on tergite 2, bands more slender on tergites 5–6, tergite 4 and 7–8 cream-yellow, with a median brown mark at posterior margin. Sternites 1–8 cream-yellow, darker towards distal segments. **Terminalia** (Figs. 82D–E). Dark cream-yellow, tip of some sclerites dark brown. Gonocoxites short, fused medially with no medial suture, bare ventrally, no laterodistal projections, a large rounded incision medially on anterior margin of syngonocoxite. Gonostylus large, complex, with a basal, slender neck expanding distally into a large conch-like, partially folded structure, with some few long setae on ventral and dorsal faces, internal area with a short peduncular projection densely surrounded by setulae, distally with a corona, apparently with a dorsal digitiform additional projection. Gonocoxal bridge wide, weakly sclerotized, with short apodemes directed obliquely inwards. Aedeagus composed by a large, weakly sclerotized rectangular plate, with a pair of semicircular lobes on anterior margin touching each other medially, at posterior margin with a pair of weakly sclerotized elongate valves laterally, and a tubular structure medially. Parameres with more or less digitiform structures anteriorly, distally with a pair of subapical setae and a pair of apical setae. Tergite 9 with a pair of long, sclerotized bottle-like structures laterodistally that extend beyond tip of gonostylus, connected medially by a slender bare stripe, lateral projections with setae on dorsal and lateral faces, and with a long, curved digitiform branch originating on ventral face midway to apex bearing some few setulae along its length. Sternite 10 weakly sclerotized, with microtrichia and small setae, between aedeagus and cerci. Cerci small, lobose, covered with microtrichia and small setae.

**Female** (Fig. 82B). As male, except as follows.

**Terminalia** (Figs. 82F–G). Sternite 8 trapezoid, a pair of short distal lobes, small setae restrict to lobes, slightly stronger at tip. Sternite 10 elongate, with elongate transparent window. Tergite 8 large, bare, tergite 9+10 slender, with a row of elongate setae on posterior margin. Cerci elongate, covered with short setae and microtrichia, cercomere 1 2.2× cercomere 2 length.

#### Material examined

**Holotype**: male, ZRCBDP0047805, Nee Soon (NS1), swamp forest, 01.may.2013, MIP leg. (slide-mounted). **Paratypes**: 7 males, 6 females. **Males**: ZRCBDP0047909, Nee Soon (NS2), swamp forest, 30.oct.13, MIP leg.; ZRCBDP0048484, Singapore, NS01, 16.may. 12, MIP leg. (MZUSP); ZRCBDP0048491, Nee Soon (NS1), swamp forest, 25.jul. 12, MIP leg. (website photo specimen); ZRCBDP0048492, Nee Soon (NS1), swamp forest, 25.apr. 12, MIP leg.; ZRCBDP0048812, Nee Soon (NS1), 14.jan.15, MIP leg.; ZRCBDP0048880, Nee Soon (NS1), 31.dec.14, MIP leg.; ZRCBDP0049172, Nee Soon (NS2), 13.may.15, MIP leg.

**Females**: ZRCBDP0048485, Nee Soon (NS1), swamp forest, 04.apr. 12, MIP leg. (slide-mounted); ZRCBDP0047932, Nee Soon (NS1), swamp forest, 17.apr.13, MIP leg.; ZRCBDP0047945, Nee Soon (NS2), swamp forest, 20.nov.13, MIP leg.; ZRCBDP0048974, Nee Soon (NS1), 13.may.15, MIP leg.; ZRCBDP0049082, Nee Soon (NS1), 24.dec.14, MIP leg; ZRCBDP0155089, Nee Soon (NS1), 03.dec.14. **Additional sequenced specimens**: ZRCBDP0133979, Singapore, date range 2012-2018, MIP leg. (abdomen missing).

**Etymology**. The specific epithet of this species, *pulau*, means island in Malay, as a reference to the island of Singapore. The noun is used in aposition.

**Remarks**. There are three haplotypes for *Neoempheria pulau* Amorim & Oliveira, **sp. n.,** but there are no conflicts between different delimitation approaches.

*Neoempheria cinkappur* Amorim & Oliveira, sp. n. (Figs. 83A–D)

https://singapore.biodiversity.online/species/A-Arth-Hexa-Diptera-000804

urn:lsid:zoobank.org:act:F841CDFF-9DF1-4731-A046-AFD8EA9382E9

**Diagnosis**. Head dark brown; scutum dark ochre-yellow, no stripes; pleural sclerites whitish, mediotergite and dorsoposterior end of laterotergite dark brown. Wing with brown band distally across wing and a couple of brown marks more basally, anterior one over cell r1, r-m and cell m1+2 close to injunction of bM and M_1+2_, and posterior one at same level on cell cup. Cell r1 small, anterior margin 0.94× length of R_4_; sc-r reaching bR well basal to origin of Rs. Tergite 1 cream-yellowish; tergite 2 cream-yellowish on anterior half, brown on posterior half; tergites 3, 5–6 dark brown, with an anterolateral cream-yellowish mark; tergite 4 cream-yellowish, with a slender brown band medially along posterior margin; tergite 7 ochre-yellowish. Female tergite 7 with lateroposterior digitiform extension; sternite 8 with a pair of distal, large posterior lobes covered with microtrichia and a number of setae, sternite 10 with transparent medial window; tergite 8 large, devoid of setation.

***Neoempheria_cinkappur_ZRCBDP0048487_hapZRCBD P0048487_SMH_holotype* [1: A, 10: A, 22: C, 61: C, 83: C, 145: C, 148: T, 151: G]**

ActatcatcAacaattgcccaCactggggcatcagttgatttagctattttttcattacaCcttgccggaatttcctcaattCtaggggctgtaaattttattactacaattattaatatacgagctccaggaattcaatttgaCcgTatGcctttatttgtctgatctgttttaattactgctattttattacttttatctttaccagtgttagcaggggctattacaatacttctaacagaccgaaatttaaatactagtttttttgatcccgcaggtgggggagaccctattttatatcaacatttattt

#### Description

**Female** (Fig. 83A). Wing length, 2.82; width, 1.11. **Head**. Light brown, slightly darker at vertex, ochre-brownish towards frons. Face brown, clypeus brown. Antennal scape and pedicel ochre-yellowish, flagellum light ochre-brown, brownish towards apex. Maxillary palpus brown, last palpomere lighter, labella light brown. Ocellar setae absent, no setae on frons anteriorly to line of ocelli. Scape 1.1× pedicel length, flagellomere 1 1.2× flagellomere 2 length, flagellomere 4 1.2× longer than wide. Palpomere 4 1.0× palpomere 3 length, palpomere 5 2.1× palpomere 4 length. **Thorax** (Fig. 83B). Scutum dark ochre, with ochre-yellow areas laterally; scutellum dark ochre. Pleural sclerites whitish with an orangish tinge, except for a brown mark on antepronotum anteriorly, a brown mark dorsally on laterotergite, mediotergite brown. Scutum with one pair of prescutellars medially along dorsocentral line, two prescutellars at each lateroposterior corner. Antepronotum with two long bristles and some additional long and short setae, proepisternum bare. Halter light brown, knob darker. **Legs**. Fore coxa light brown anteriorly and whitish posteriorly, mid coxa whitish with a brownish tinge antero-distally, hind coxa whitish; hind femur ochre-yellow, tibia and tarsus light greyish-brown, tarsomeres darker towards tip [holotype fore and mid femora, tibiae and tarsi missing]. Hind tibia inner spur 3.6× longer than tibia width at apex. **Wing** (Fig. 83C). Membrane with dark brown marks over sc-r, first sector of Rs, R_4_ and along bM, yellowish along anterior margin. C produced beyond tip of R_5_ for about a fifth of distance to M_1_; Sc complete (visible on phase contrast), reaching C slightly before level of origin of Rs, sc-r present, reaching bR before of origin of Rs for a distance equal to first sector of Rs. R_1_ reaching C at distal fourth of wing; R_5_ reaching C before level of tip of M_1_; R_4_ present, slightly oblique, very close to base of Rs, cell r1 trapezoid, short, anterior margin of cell r1 1.1× longer than length of R_4_; first sector of Rs 0.70× r-m length. False medial vein conspicuous, sclerotized. M_1+2_ 3.7× r-m length; bM 6.2× length of first sector of Rs. First sector of CuA 1.1× longer than second sector. M_4_ almost straight on distal half. Cubital pseudovein sclerotized to mid of second sector of CuA; CuP faint, not even reaching origin of M_4_. Anal fold faint. Wing margin gently emarginated at tip of CuA. No ventral macrotrichia on veins, dorsal macrotrichia on distal third of Sc before sc-r, entire length of bR, R_1_, second sector of Rs, most of M_1_ and M_4_, distal ⅔ of M_4_, and entire length of CuA; sc-r, most of Sc, base of bR, first sector of Rs, R_4_, r-m, bM, and CuP entirely devoid of macrotrichia. **Abdomen**. Tergite 1 light brown, tergites 2–3 and 5–6 with a dark brown medial band, a cream-yellow anterior band extending posteriorly along laterals on tergites 2–3 and with a cream-yellow mark on anterolateral corner, tergite 4 cream-yellow with a small median brown mark ochre-yellow. Sternites 1–2 whitish, sternites 3–5 cream-yellow, sternites 6–7 ochre-yellow. Tergite 7 with a digitiform projection on lateroposterior corners. **Terminalia** (Fig. 83D). Ochre-yellow, tergite 8 light brown. Sternite 8 trapezoid, lateroanterior corners extended dorsally, a wide incision on anterior margin, distally with a deep slender incision separating a pair of pointed lobes, microtrichia on entire sclerite, setae restricted to lobes. Sternite 9 with anterior apodeme considerably wide, genital chamber elongate, a pair of lateral winglets. Sternite 10 trapezoid, elongate, with a long transparent window with microtrichia and setae. Tergite 8 large, trapezoid, lateroanterior corners extending ventrally to articulate with sternite 8, distally more sclerotized, a pair of small pointed projections laterally to base of tergite 9+10. Tergite 9+10 with a slender medial connection between a pair of short lateral lobes with microtrichia and setae, laterally a pair of short digitiform projections, each with a long seta at tip. Cercomere 1 relatively short, widening to apex, 1.2× longer than cercomere 2, both covered with microtrichia and short setae.

**Male**. Unknown.

#### Material examined

**Holotype**: female, ZRCBDP0048487, Nee Soon (NS2), swamp forest, 13.jun.2012, MIP leg. (slide-mounted).

**Etymology**. The species epithet refers to CiDkappūr (LJLJLJLJLJLJLJLJLJLJLJ), the Tamil transcription for Singapore. The noun is used in aposition.

**Remarks**. There are features in the female morphology of this species that allow the presentation of a diagnosis and it is formally described and named here. This includes the color pattern of the head and thorax, the small size of cell r4 and the wing markings, and even the shape of the terminalia sclerites.

*Neoempheria temasek* Amorim & Oliveira, sp. n. (Figs. 84A–D)

https://singapore.biodiversity.online/species/A-Arth-Hexa-Diptera-000823

urn:lsid:zoobank.org:act:B1F0472E-E149-4713-AA0E-6A54AD0111B9

**Diagnosis**. Head light ochre-yellow; scutum light ochre-yellow, no stripes; pleural sclerites whitish, mediotergite and laterotergite brown. Wing with brown band distally across wing and a brown band more basally, over cell r1, r-m, cell m1+2 close to injunction of bM, and M_1+2_. Cell r1 small, anterior margin 0.94× length of R_4_; sc-r reaching bR well basal to origin of Rs. Tergite 1 cream-yellowish; tergite 2 cream-yellowish on anterior half, brown on posterior half; tergites 3, 5–6 dark brown with an anterolateral cream-yellowish mark; tergite 4 cream-yellowish with a slender brown medial band; tergite 7 ochre-yellowish. Gonocoxites with a ventral projection directed obliquely inwards; gonostylus small, displaced laterodistally, bifid from base, external branch capitate, slightly more dorsal in position; tergite 9 with a pair of large lobes more or less compressed dorsoventrally, extending much beyond tip of gonostylus.

***Neoempheria_temasek_ZRCBDP0047801_hapZRCBDP 0047801_SMH_holotype* [73: C, 145: C, 172: A, 202: C, 203: C, 238: T, 244: C, 247: T, 265: C] --**

tatcatcaacaattgctcatacaggagcttcagttgatttagctattttttctttacatttagcaggtatCtcttctattttaggagctgtaaattttattacaactattattaatatgcgagccccaggtatccaatttgaCcgaatacctttatttgtttgatctgtAttaattacagcaattttattacttttatcCCtacctgttttagcaggagcaattactatgttactTacagaCcgTaatttaaatactagattCtttgatcctgcaggaggaggagaccctattttataccaacatttattt

#### Description

**Male** (Fig. 84A). Wing length, 3.15; width, 1.21. **Head**. Ochre-yellow, darker on vertex. Face and clypeus whitish-yellow, with scattered setae. Antennal scape and pedicel light ochre-yellow, flagellum ochre-yellow. Maxillary palpus light brown, last palpomere lighter, labella light ochre-yellowish. Scape 1.4× pedicel length, flagellomere 1 1.3× flagellomere 2 length, flagellomere 4 as long as wide. Palpomere 4 1.5× palpomere 3 length, palpomere 5 2.4× palpomere 4 length. **Thorax**. Scutum mostly ochre-yellow, lighter above wing; scutellum light ochre-yellow. Pleural sclerites whitish with an orangish tinge, antepronotum slightly darker anteriorly, laterotergite light brown except for lighter ventral end, mediotergite brown with ochreish areas laterally. Halter whitish with brown tinge, base of knob slightly darker. Scutellum with a pair of prescutellar bristles on dorsocentral line, two pairs of prescutellars on lateroposterior corners. Antepronotum with three bristles and some additional smaller setae, proepisternum bare. **Legs**. Coxae whitish, fore coxa with some orangish tinge; femora light ochre-yellow; tibiae and tarsi light greyish-brown. Fore leg tarsomere 1 0.85× tibia length, 2.1× tarsomere 2 length. Hind tibia inner spur 4.2× tibia width at apex. **Wing** (Fig. 84B). Membrane with a light yellowish-brown background, with a dark brown band across distal third of wing, beginning at level of base of medial fork and a more basal band across wing from level of Sc to posterior margin. C produced beyond tip of R_5_ for about a fifth of distance to M_1_; Sc weakly sclerotized beyond sc-r (barely visible even on phase contrast), sc-r present, reaching bR well before origin of Rs. R_1_ reaching C at distal third of wing; R_5_ reaching C slightly before level of tip of M_1_; R_4_ present, slightly oblique, close to base of Rs, cell r1 trapezoid, short, anterior margin of cell r1 0.94× longer than length of R_4_; first sector of Rs 0.89× r-m length. False medial vein conspicuous, sclerotized. M_1+2_ 4.0× r-m length; bM 8.5× length of first sector of Rs; base of M_1_ weakly sclerotized. First sector of CuA 1.1× longer than second sector. M_4_ straight on distal half. Cubital pseudovein sclerotized to distal third of second sector of CuA; CuP sclerotized to basal third of second sector of CuA. Anal fold faint. Wing margin emarginated at tip of CuA. No ventral macrotrichia on veins, dorsal macrotrichia on entire length of bR, R_1_, second sector of Rs, most of M_1_ and of M_4_, distal ⅔ of M_2_, and entire length of CuA; sc-r, Sc, base of bR, first sector of Rs, R_4_, r-m, bM, and CuP entirely devoid of macrotrichia. **Abdomen**. Tergite 1 cream-yellow, tergites 2–3 brown with a cream-yellow band on anterior third, tergite 4 mostly ochre-yellow, with a medial slender longitudinal brown band; tergites 5–6 brown, with slender ochre-yellow slender bands anteriorly and posteriorly, tergite 7 light ochre-yellow. Sternites 1–5 whitish, sternite 6 brown with a whitish-yellow band medially, brown laterally, sternite 7 ochre-yellowish. **Terminalia** (Figs. 84C–D). Ochre-yellow, with some more brownish sclerites. Gonocoxites fused medially along a short extension, no medio-posterior process, an oblique projection inwards ventrally, with some few elongate fine setulae. Gonostylus placed laterodistally, bifid from base, external branch capitate, thin, directed outwards, with a concentration of elongate setulae distally, inner branch more sclerotized, projected inwards, with long, strong setae along posterior margin, distal setae curved. Gonocoxal apodeme wide, with a pair of short lateroanterior apodemes. Aedeagus wide, with a pair of short anterior apodemes, a pair of lateral winglets on basal third, a subquadrate distal plate with a U-shaped medial incision between a pair of lateral extensions distally. Parameres weakly sclerotized, wide, blade-like. Tergite 9 wide medially, bare, with a pair of large lobes more or less compressed dorsoventrally, extending to much beyond tip of gonostylus, densely covered with long setae on dorsal face, with some fine, long setae on ventral face. Sternite 10 small, trapezoid, slender at distal end, covered with microtrichia, setae along posterior margin. Cerci small, lobose, covered with microtrichia and small setae. **Female**. As male, except for the following. **Wing l**ength, 2.52; width, 0.95. **Terminalia**. Sternite 9 with a pair of elongated lobes barely in contact medially, no microtrichia, covered with elongate setae, distal setae on each lobe more concentrated and longer. Tergite 8 wide, covered with elongate setae, lateroposterior end at each side with a concentrated group of setae. Sternite 9 weakly sclerotized. Tergite 9+10 slender, with microtrichia and a sequence of elongate setae along posterior margin, lateroposterior corner with a short extension and a longer seta apically. Sternite 10 elongate, slightly arched, with transparent window between lateral margins. Cercomere 1 2.5× cercomere 2 length, both densely covered by microtrichia and short setae, distal tip on dorsal face of cercomere 1 slightly projected beyond insertion of cercomere 2.

#### Material examined

**Holotype**: male, ZRCBDP0047801, Nee Soon (NS2), swamp forest, 13.nov.2013, MIP leg. (slide-mounted). **Paratypes** (16 males, 8 females): **Males**: ZRCBDP0047851, Nee Soon (NS1), swamp forest, 09.may.13, MIP leg.; ZRCBDP0048695, Nee Soon (NS1), swamp forest, 18.apr. 12, MIP leg. (website photo specimen); ZRCBDP0048704, Nee Soon (NS2), 28.jan.15, MIP leg.; ZRCBDP0048810, Nee Soon (NS1), 14.jan.15, MIP leg.; ZRCBDP0048850, Nee Soon (NS1), 14.jan.15, MIP leg.; ZRCBDP0048891, Nee Soon (NS1), 31.dec.14, MIP leg.; ZRCBDP0048947, Nee Soon (NS2), 03.dec.14, MIP leg.; ZRCBDP0049240, Nee Soon (NS1), 10.dec.14, MIP leg.; ZRCBDP0066789, Bukit Timah, primary forest (BT05), 16.aug. 16, MIP leg.; ZRCBDP0072465, Bukit Timah, maturing secondary forest (BT08), 01.dec. 16, MIP leg. (slide-mounted) (MZUSP); ZRCBDP0072695, Bukit Timah, primary forest (BT05), 01.dec. 16, MIP leg.; ZRCBDP0072704, Bukit Timah, primary forest (BT05), 01.dec. 16, MIP leg.; ZRCBDP0072707, Bukit Timah, primary forest (BT05), 01.dec. 16, MIP leg.; ZRCBDP0072710, Bukit Timah, primary forest (BT05), 01.dec. 16, MIP leg.; ZRCBDP0072711, Bukit Timah, primary forest (BT05), 01.dec. 16, MIP leg.; ZRCBDP0072712, Bukit Timah, primary forest (BT05), 01.dec. 16, MIP leg. **Females**: ZRCBDP0048840, Nee Soon (NS1), 14.jan.15, MIP leg. (slide-mounted).; ZRCBDP0048876, Nee Soon (NS1), 31.dec.14, MIP leg. (slide-mounted); ZRCBDP0048960, Nee Soon (NS1), 15.apr.15, MIP leg.; ZRCBDP0072457, Bukit Timah, old secondary forest (BT07), 08.dec. 16, MIP leg. (slide-mounted); ZRCBDP0066783, Bukit Timah, primary forest (BT05), 16.aug. 16, MIP leg.; ZRCBDP0066802, Bukit Timah, primary forest (BT05), 23.aug. 16, MIP leg.; ZRCBDP0066816, Bukit Timah, maturing secondary forest (BT09), 05.oct. 16, MIP leg.; ZRCBDP0072464, Bukit Timah, maturing secondary forest (BT08), 01.dec. 16, MIP leg. **Additional sequenced specimens:** ZRCBDP0078949, Singapore; ZRCBDP0078950, Singapore; ZRCBDP0078952, Singapore; ZRCBDP0078954, Singapore; ZRCBDP0078994, Singapore; ZRCBDP0078995, Singapore; ZRCBDP0078999, Singapore; ZRCBDP0079000, Singapore; ZRCBDP0079009, Singapore.

**Etymology**. The species epithet refers to the early recorded, likely Malay/Javanese name for pre-colonial Singapore, Temasek. The noun is used in aposition.

**Remarks**. A large number of specimens of this species were collected in different environments in Singapore. There are three separate haplotypes, which cluster together under all delimitation methods.

*Neoempheria polunini* Amorim & Oliveira, sp. n. (Figs. 85A–D)

https://singapore.biodiversity.online/species/A-Arth-Hexa-Diptera-000812

urn:lsid:zoobank.org:act:77C82CD9-6371-4002-BF36-6541FA2FF920

**Diagnosis**. Head light ochre-yellow, with a light brown diffuse mark over vertex; scutum light ochre-yellow, no stripes; pleural sclerites whitish, mediotergite light brown, laterotergite with a small light brown mark at dorsoposterior end. Wing with brown band across distal third of wing and a brown band more basally, from level of cell r1 to posterior margin; R_4_ absent; sc-r reaching bR well basal to origin of Rs. Tergites 1–2 and 4 cream-yellow; tergites 3, 5–6 dark brown, tergite 3 with slender cream-yellow lateroposterior band; tergite 7 whitish-yellow; sternite 6 with a large brown mark laterally. Male syngonocoxite with a wide medio-posterior sclerotized process bearing short lateroposterior projections, a pair of sub-medial, bifid long blade-like projections internally to insertion of gonostylus; gonostylus long, digitiform, curved inwards distally; tergite 9 with a slender blade medially connecting a pair of long lateral setose projections with a group of 5–6 spines apically.

***Neoempheria_polunini_ZRCBDP0049204_hapZRCBDP 0048849_SMH_holotype* [43: G, 46: C, 49: C, 70: G, 73: C, 91: C, 127: A, 244: C]**

cctatcttctacaattgctcatacaggggcatctgtcgatttGgcCatCttttctttacatttagcaggGatCtcttctattttaggagcCgtaaattttattacaacaattattaatatacgagcAcctggaattcaatttgatcgaatacctttatttgtatgatcagttttaattacagctattttattacttttatctttacctgttttagctggagctattacaatattattaactgaCcgaaatttaaatacaagtttttttgaccctgctggaggaggagacccaattttatatcaacatttattt

#### Description

**Male** (Fig. 85A). Wing length, 2.49; width, 0.95. **Head**. Brownish ochre-yellow at vertex, lighter above antennae and laterally on occiput. Antennal scape and pedicel light ochre-yellow, flagellum ochre-yellow with brownish tinge. Face and clypeus whitish-yellow. Palpomeres brown, last palpomere lighter; labella light brownish. Ocellar setae present, small, only some few setae on frons laterally. Scape 1.2× pedicel length, flagellomere 1 1.76× flagellomere 2 length, flagellomere 4 1.4× longer than wide. Palpomere 4 1.0× palpomere 3 length, palpomere 5 2.0× palpomere 4 length. **Thorax**. Scutum mostly ochre-yellow, scutellum light brown. Pleural sclerites whitish with an orangish tinge, laterotergite light brown on dorsoposterior end, mediotergite light brown, ochreish laterally. Scutum with a prescutellar bristle medially on each dorsocentral line, one prescutellar bristle at each lateroposterior corner; scutellum with one pair of bristles and additional fine setae. Antepronotum with three bristles and additional larger and smaller setae. **Legs**. Coxae whitish, fore coxa with some orangish tinge; fore femur light ochre-yellow; tibiae and tarsi light greyish-brown. Hind tibia inner spur 3.6× longer than tibia width at apex. **Wing** (Fig. 85B). Membrane with a dark brown band across wing at level of origin of Rs and a brown mark before wing tip beginning slightly beyond basal end of medial fork, both marks connected briefly along posterior margin. C extending beyond tip of R_5_ for a fifth of distance to M_1_; Sc not sclerotized apically (not observable even under phase contrast), sc-r absent; R_4_ absent; first sector of Rs 1.3× r-m length. False medial vein present, curved along entire length. M_1+2_ 3.8× r-m length; bM 7.1× length of first sector of Rs. First sector of CuA 1.2× longer than second sector. M_4_ depressed on distal half. Cubital pseudovein weakly sclerotized, ending at level of distal fourth of second sector of CuA; CuP reaching level of basal third of second sector of CuA. Anal fold very faint. Wing margin emarginated at tip of CuA. No ventral macrotrichia on veins, dorsal macrotrichia on entire length of bR, R_1_, second sector of Rs, most M_1_, distal half of M_2_, most M_4_, and almost entire CuA. Sc, first sector of Rs, r-m, bM, and CuP entirely devoid of macrotrichia. **Abdomen**. Tergite 1 whitish, tergite 2 cream-yellow with a slender light brown medial band; tergites 3, 5–6 brown, tergite 3 with a slender cream-yellow band along anterior margin, tergite 4 and 7 ochre-yellow. Sternites 1–2 whitish, sternites 3–5 whitish-yellow, sternite 6 light brown laterally, light ochre-yellow medially, sternite 7 ochre-yellowish. **Terminalia** (Figs. 85C–D). Ochre-yellow, tip of gonostylus brownish. Gonocoxites fused medially for a short distance, gonocoxite with a wide medio-posterior sclerotized process bearing a pair short projections lateroposteriorly, sub-medially a pair of bifid blade-like projections internally to insertion of gonostylus, ventral branch capitate extending almost to level of tip of gonostylus, curved inwards on distal half, some few short setulae at caput, dorsal branch flat, bare projection reaching slightly beyond base of gonostylus. Gonostylus placed laterodistally, long, digitiform, setose on outer face, curved inwards distally, reaching beyond tip of tergite 9 projections, with a group of fine setae directed inwards distally. Aedeagus subquadrate, wide, posterior margin almost reaching tip of ventral lobe of gonocoxite. Parameres weakly sclerotized, with a pair of wide blades close together medially extending slightly beyond posterior margin of aedeagus. Tergite 9 with a slender blade medially connecting a pair of long lateral setose projections, a sub-medial projection branching inwards and a group of 5–6 spines apically. Sternite 10 weakly sclerotized, with a pair of short lobes, bearing microtrichia and fine setae. Cerci small, close together, weakly sclerotized lobes, with microtrichia and fine setae.

**Female**. Unknown.

#### Material examined

**Holotype**: male, ZRCBDP0049204, Nee Soon (NS1), 10.dec.2014, MIP leg. (website photo specimen, slide-mounted).

**Paratypes** (4 males): ZRCBDP0048849, Nee Soon (NS1), 14.jan.15, MIP leg.; ZRCBDP0049214, Nee Soon (NS1), 10.dec.14, MIP leg.; ZRCBDP0049231, Nee Soon (NS1), 10.dec.14, MIP leg.; ZRCBDP0049045, Nee Soon (NS2), 07.jan.15, MIP leg.

**Additional sequenced specimens:** ZRCBDP0154867, Nee Soon (NS1), 21.jun.15.

**Etymology**. The species epithet honors Ivan Polunin (1920–2010), a medical doctor and enthusiastic naturalist, who documented wildlife in Singapore using color films during the 1950s. He has as well documented lampyrid light synchronization.

**Remarks**. All specimens of this species come from the Nee Soon swamp forests. There are four haplotypes and all delimitation approaches point to a single species. This is the only species of *Neoempheria* we know of without R_4_; in Søli’s (2017) key for the Afrotropical genera of mycetophilids, this species would run into *Parempheriella*. As discussed above, *Neoempheria* is most certainly paraphyletic in relation to *Parempheriella*, but *N. polunini* Amorim & Oliveira, **sp. n.** has other regular features of *Neoempheria*: wing with maculae, a conspicuous false medial vein etc. Some additional relevant details reinforce the idea that this is a species of *Neoempheria*. In the wing of *Parempheriella* species, r-m is almost longitudinal, in such a way that the origin of M_1+2_ is much more basal than the origin of r-m in Rs. In *Neoempheria polunini* Amorim & Oliveira, **sp. n.,** r-m is almost transverse, as in all other *Neoempheria* species, in such a way that the origin of M_1+2_ is almost below the origin of r-m. Evidence strongly suggests that this is an independent case of loss of R_4_ in the evolution of the genus.

*Neoempheria fajar* Amorim & Oliveira, sp. n. (Figs. 86A–C)

urn:lsid:zoobank.org:act:FC916C3C-BE98-45C2-8BD8-D6CC99D1BBCE

**Diagnosis**. Head brown, scutum dark brown, no stripes, scutellum ochre-yellowish; pleural sclerites whitish-yellow, laterotergite and mediotergite brown, mediotergite lighter on ventral fourth. Wing with a single large, brown band across wing distal half, beginning at level of tip of Sc; cell r1 small, length of anterior margin 1.7× R_4_ length; sc-r reaching R_1_ at distal half of cell r1. Tergite 1 cream-yellow, tergites 2–6 brown with cream-yellow anterior band on tergites 2–4, cream-yellow lateroanterior corners on tergites 5–6; tergite 7 entirely cream-yellow. Female terminalia with unique pattern, wide, sternite 8 with a pair of distal setose lobes largely separate.

***Neoempheria_fajar_ZRCBDP0284230_hapZRCBDP028 4230_SMH_holotype* [7: A, 67: G, 169: T, 184: A, 233: C, 250: C, 259: C, 271: C, 283: G] -**

**ttatcAtctactattgctcatacaggagcttctgttgatttagctattttttcattacatttagcGggtatttcttcaattttaggagctgttaattttattacaactattattaatatacgagccccaggaattcaatttgatcgaatacctttatttgtatgatcTgttttaattacagcAattttattattactatctttacctgttttagcaggagctattacaataCtattaacagatcgaaaCttaaatacCannttntttgaCccagctggnggGggagatcctattttataccaacat------**

#### Description

**Female**. Wing length, 4.33; width, 1.64. **Head**. Brown, lighter ventrally on occiput, face and clypeus light brown, face with scattered small setae and macrotrichia, clypeus bare. No setae on frons anteriorly to ocelli. Post-ocellar setae present. Antennal scape and pedicel yellowish-brown, scape 1.4× pedicel length, flagellum light brown, flagellomere 1 2.0× flagellomere 2 length, flagellomere 4 1.4 longer than wide. Palpomeres light brown, last palpomere brownish-yellow, palpomere 4 about 1.2× palpomere 3 length, a short distal projection over base of distal palpomere, palpomere 5 2.1× palpomere 4 length. Labella small, brownish. **Thorax**. Scutum homogenously dark brown; a row of longer dorsocentral setae, some long supra-alars, two pairs of prescutellar bristles on lateroposterior corners of scutum, one pair of prescutellars medially on dorsocentral line. Scutellum ochre-yellow, with one pair of bristles and some smaller setae on disc. Pleural sclerites whitish-yellow, antepronotum light brown anteriorly, laterotergite and mediotergite brown, mediotergite lighter on ventral fourth. Antepronotum with one stronger bristle and smaller setae of different sizes. Halter light brown. **Legs**. Coxa whitish; anterior and mid femora ochre-yellowish, hind femur brownish-yellow; tibiae and tarsi yellowish-brown, anterior tarsus lighter. Hind tibial spurs 2.7× length of tibia at apex. **Wing** (Fig. 86A). Membrane with light greyish-brown with a single wide brown band across wing, at anterior margin between tip of Sc and tip of R_1_. Sc complete, reaching C slightly beyond level of tip of R_4_; sc-r present, reaching R_1_ slightly beyond mid of cell r1; cell r1 relatively short, anterior margin 1.7× length of R_4_. False medial vein present, gently sinuous on basal fourth. Medial fork not wide open. M_1+2_ 2.4× r-m length; bM 6.5× r-m length. First sector of CuA 1.2× longer than second sector. M_4_ gently depressed on distal half. Cubital pseudovein reaching level of distal third of second sector of CuA; CuP present to level of mid of second sector of CuA. Anal fold faint. Wing margin emarginated at tip of CuA. No ventral macrotrichia on veins, dorsal macrotrichia on distal half of Sc, entire length of bR, R_1_, second sector of Rs, most of M_1_, distal ⅔ of M_2_, entire M_4_, and on entire CuA. **Abdomen** (Fig. 86B). Tergite 1 cream-yellow, tergites 2–6 brown with cream-yellow anterior band on tergites 2–4 (extending more posteriorly along lateral margin on tergite 4) and with cream-yellow lateroanterior corners on tergites 5–6, tergite 7 cream-yellow. Sternites 1–5 and 7 cream-yellow, sternite 6 light brown. **Terminalia** (Fig. 86C). Ochre-yellow. Sternite 8 large, with a pair of large lobes densely covered by setation, each bearing a subdistal dorsal protuberance on ventral face, a pair of setose digitiform projections on lateral ends of sclerite. Sternite 9 wide, with a short median anterior projection of vaginal furca and a well-sclerotized triangular area distally to genital opening. Tergite 8 wide, with a pair of rounded lobes widely separated medially projecting outwards distally, sclerite mostly bare on anterior third, lobes covered with microtrichia and setation. Tergite 9+10 slender, with three pairs of long digitiform projections, each with a distal long seta, inner pair with a second, subdistal seta. Sternite 10 small, weakly sclerotized, covered with microtrichia and fine setae. Cerci small, weakly sclerotized.

**Male**. Unknown.

#### Material examined

**Holotype**: female, ZRCBDP0284230, Singapore, freshwater swamp (KM03), date range, 2012-2018 (slide-mounted).

**Etymology**. The species epithet refers to a magazine named Fajar [=dawn in Arabic], published by the University Socialist Club in Singapore in the early 1950s. The Club played an important role in the politics of colonial Malaya and post-colonial Malaysia and Singapore. The editorial board members were arrested, went to trial and were released three days after the trial. The Club’s victory stands as a landmark in the progress of decolonisation of this part of the world. The noun is used in apposition.

**Remarks**. This is a very unusual species of *Neoempheria*. The entire distal half of the wing is covered by a brownish macula and the abdomen color pattern is as well different from what is seen in other species of the genus. Even the female terminalia has a different general pattern.

### Unplaced species of Neoempheria

*Neoempheria riatanae* Amorim & Oliveira, sp. n. (Figs. 87A–D)

https://singapore.biodiversity.online/species/A-Arth-Hexa-Diptera-000802

urn:lsid:zoobank.org:act:3C612CAF-0310-4C0E-85A2-4D8FAE7760F1

**Diagnosis**. Head and scutum dark brown, no stripes, mediotergite dark brown, laterotergite only with a tinge of brown along posterior margin. Wing with oblique brown mark across tip of wing, on anterior margin beginning slightly before tip of R_1_, on posterior margin ending more basally than tip of CuA; a second oblique band more basally on the wing, beginning at tip of Sc and ending on basal lobe; anterior margin of cell r1 small, length of anterior margin 1.4 R_4_ length; sc-r reaching R_1_ on basal third of cell r1. Abdominal tergites 1–6 dark brown, tergites 1–4 with laterals yellowish, tergite 7 brownish-yellow. Sternite 8 yellowish, cerci whitish. Gonocoxites fused, bare, with a short suture on anterior end ventrally, a long dorsal setose projection. Gonostylus short, ear-like displaced laterally. Gonocoxal apodemes long, parallel on anterior end. Aedeagal plate not visualized. Parameres with a pair of setose lobes posteriorly. Tergite 9 with a short medial connection, with a pair of separate projections bearing some few denticles distally. Cerci small, elongate, separate from each. Female terminalia elongated, sternite 8 with a median short crest between the distal lobes.

***Neoempheria_riatanae_ZRCBDP0049212_hapZRCBDP 0047825_SMH_holotype* [22: C, 28: G, 110: A, 136: C, 1 55: C, 161: G, 166: G, 199: A, 241: T]**

tctttcttctacaatcgctcaCacaggGgcttcagtagatttagctattttttctcttcatctagcagggatttcttcaattttaggggctgtaaattttattactacaAttattaatatacgagccccaggaatCcaatttgatcgaatacccCtatttGtttgGtcagttttaattactgctgttttattacttttAtcattgccagtattagcaggagctattacaatactattaacTgatcgaaatttaaatactagcttttttgaccctgctggtggaggtgaccctattttataccaacatttattt

#### Description

**Male**. Wing length, 2.59–2.75; width, 0.95 (n=2). **Head**. Brown, lighter at ventral half of occiput. No setae on frons anteriorly to ocelli. Ocellar setae present. Face light brown, clypeus yellowish-brown. Antennal scape light brown, pedicel dark ochre-yellow, flagellum light brown. Palpomeres brown, last palpomere lighter, labella short, light brown. Scape 1.0× pedicel length, flagellomere 1 1.6× flagellomere 2 length, flagellomere 4 1.5× wider than long. Palpomere 4 1.0× palpomere 3 length, dorsally tip slightly projecting beyond base of palpomere 5, palpomere 5 1.7× palpomere 4 length. **Thorax**. Scutum light brown, with ochre-brown areas laterally and along anterior margin; scutellum light brown. Pleural sclerites mostly whitish, antepronotum ochre-brown, cervical sclerite dark brown, mediotergite brown. Scutum with some strong supra-alars, one pair of prescutellar bristles medially on dorsocentral line, one pair of prescutellar bristles on lateroposterior corner; scutellum with one pair of bristles and additional fine setae. Antepronotum with two strong bristles and additional smaller setae. Halter light brown, knob darker. Halter light brown. **Legs**. Coxae ochre-yellowish, fore coxa darker, hind coxa lighter; femora ochre-yellow; tibia and tarsus light greyish-brown, tarsomeres darker towards tip. Fore leg tarsomere 1 0.8× tibia length, 1.8× tarsomere 2 length. Hind tibia inner spur 3.0× longer than tibia width at apex. **Wing** (Fig. 87B). Membrane with a dark brown band across wing at level of base of Rs and a wide band distally at wing, beginning slightly beyond base of medial fork. Sc reaching C beyond level of base of second sector of Rs, sc-r complete, reaching R_1_ beyond origin of R_1_, anterior margin of cell r1short, about as wide as long; first sector of Rs 0.8× r-m length. False medial vein present, slightly sclerotized. Medial fork wide open. M_1+2_ 3.3× r-m length; bM 5.2× length of first sector of Rs. First sector of CuA 0.90× longer than second sector. M_4_ gently depressed on distal half. Cubital pseudovein weakly sclerotized, ending at level of second third of second sector of CuA; CuP reaching level of mid of second sector of CuA. Anal fold faint. Wing margin not emarginated at tip of CuA. No ventral macrotrichia on veins, dorsal macrotrichia on distal half of Sc, on entire length of bR, R_1_, and second sector of Rs, and on distal fourth of M_1_ and M_2_, distal fourth of M_2_. **Abdomen**. Tergites 1–2 and 4 dark brown medially, wide cream-yellow bands laterally, tergite 3 mostly dark brown, lighter along lateral margins, tergites 5–6 brown, tergite 7 with a light brownish-yellow medial band and an ochre-yellow band laterally. Sternites 1–4 whitish to light ochre-yellow, sternites 5–6 brown, sternites 7 ochre-yellow. Tergite 7 with no projection on lateroposterior corner. **Terminalia** (Fig. 87C). Gonocoxites fused, bare, with a short suture on anterior end ventrally, dorsally a long, flat projection covered with setae. Gonostylus short, ear-like displaced laterally. Gonocoxal apodemes long, parallel on anterior end. Aedeagal plate not visualized. Parameres with a pair of setose lobes posteriorly. Tergite 9 with a short medial connection, with a pair of separate projections bearing some few denticles distally. Cerci small, elongate, separate from each.

**Female** (Fig. 87A). As male, except as following. **Terminalia** (Fig. 87D). Ochre-yellow, cerci lighter. Sternite 8 wide anteriorly, extending posteriorly with a subtriangular shape, bearing a medio-distal short keel sided by a pair of subdistal lateral lobes. Sternite 9 wide, complex, without a furca at anterior end. Tergite 8 wide, slightly projected medially covered with microtrichia, but no setae. Tergite 9 wide, short, with a row of elongate setae. Tergite 10 slender, two pairs of short protuberances each with an elongate seta at tip, a pair of more developed lobes at lateral ends. Sternite 10 with a pair of short lateral lobes on posterior margin, transparent window wide midway to apex. Cerci laterally compressed, cercomere 1 2.0× longer than cercomere 2, both covered with microtrichia and short setae.

#### Material examined

**Holotype**: 1 male, ZRCBDP0049212, Nee Soon (NS1), 10.dec.2014, MIP leg. **Paratypes** (3 females): ZRCBDP0047840, Nee Soon (NS1), swamp forest, 24.apr.13, MIP leg. (slide-mounted); ZRCBDP0047825, Nee Soon (NS1), swamp forest, 20.feb.13, MIP leg. (slide-mounted); ZRCBDP0049247, Nee Soon (NS1), 10.dec.14, MIP leg. (website photo specimen).

**Etymology**. The species epithet honors Ms. Ria Tan (1961-), prominent naturalist and conservationist, with a key role in raising awareness and protecting Singapore’s littoral and mangrove habitats.

*Neoempheria* sp. B (Figs. 88A–D)

https://singapore.biodiversity.online/species/A-Arth-Hexa-Diptera-000834

***Neoempheria_spB_ZRCBDP0047779_hapZRCBDP0047 779_SMH_unnamed_type* [31: C, 76: C, 88: T, 100: C, 1 06: G, 127: A, 194: T, 199: T]**

tctatcctctacaattgctcatactggagcCtcagtagatttagctattttttctttacatttagccggaatctcCtcaattttaggTgcagttaatttCattacGactattattaatatacgtgcAccagggattcaatttgaccgaatacctttatttgtttgatcagttttaattacagcagtattattaTtactTtcattaccagttttagctggagcaattactatactattaacagaccgaaatttaaatactagtttctttgaccctgccggaggaggagaccctattttataccaacatttattt

#### Description

**Female** (Fig. 88A). Wing length, 2.20; width, 0.85. **Head**. Light-brown at vertex, lighter towards frons and ventral margin of occiput. Face light brown, clypeus yellowish-brown. Antennal scape and pedicel ochre-yellow, flagellum light ochre-yellow. Palpomeres brown, last palpomere lighter; labella yellowish-brown. Frons with some few scattered short setae anteriorly to line of ocelli. Scape 1.0× pedicel length, flagellomere 1 2.0× length of flagellomere 4, flagellomere 4 1.2× longer than wide.

Palpomere 4 1.0× palpomere 3 length, palpomere 5 1.7× palpomere 4 length. **Thorax**. Scutum mostly dark ochre-yellow, more brownish laterally along anterior half and medially on distal half; scutellum light brown. Pleural sclerites cream-yellow with an orangish tinge, antepronotum with a dark brown mark at inner margin anteriorly, dorsoposterior end of laterotergite with a light brown mark, mediotergite light brown. Scutum with a pair of prescutellar bristles at dorsocentral lines and one prescutellar bristle at each lateroposterior corner; scutellum with one pair of scutellar bristles and one additional pair of small setae. Antepronotum and proepisternum each with one strong bristle and some smaller setae. **Legs**. Coxae whitish, mid and hind coxae with a brown mark at tip; femora ochre-yellow; tibiae and tarsi light greyish-brown. Mid tibia inner spur 3.3× tibia width at apex [both hind leg femora, tibiae and tarsi missing on holotype]. Fore leg tarsomere 1 0.8× tibia length, 1.6× tarsomere 2 length. **Wing** (Figs. 88B–C). Membrane background light greyish fumose, a dark brown band across wing at level of tip of Sc and a dark brown mark at distal two-fifths of wing, beginning at level of base of medial fork. C produced beyond tip of R_5_ for less than a ¼ of distance to M_1_; Sc reaching C slightly beyond level of origin of Rs; sc-r not produced; first sector of Rs oblique, R_4_ not far from base of r-m, anterior margin of cell r1 1.4× length of first sector of Rs; first sector of Rs 1.0× r-m length. False medial vein conspicuous, sclerotized. M_1+2_ 3.3× r-m length; bM 7.4× length of first sector of Rs. First sector of CuA 1.3× longer than second sector. M_4_ gently curved on distal half. Cubital pseudovein sclerotized to beyond level of origin of M_4_, CuP barely sclerotized, reduced to wing base, anal fold weakly sclerotized. Wing margin not emarginated at tip of CuA. No ventral macrotrichia on veins, dorsal macrotrichia on entire length of bR, R_1_, anterior third of r-m, second sector of Rs, and distal third of M_1_; Sc, sc-r, first sector of Rs, R_4_, r-m posteriorly, bM, M_2_, M_4_, CuA, and CuP devoid of macrotrichia. **Abdomen**. Tergite 1 light brown, tergite 2 light brown with wide cream-yellow lateral bands, tergites 3–4 with slender cream-yellow lateral bands, tergite 5–6 light brown, tergite 7 ochre-yellowish on anterior half, light brownish on posterior half; tergite 7 wider on distal margin, no other lateroposterior projections. Sternite 1 whitish, sternites 2–4 cream-yellow, sternites 5–6 light brownish-yellow, sternite 7 whitish-yellow. **Terminalia** (Fig. 88D). Cream-yellow, tip of sternite 8, sternite 10, tergite 9+10 and base of cerci light yellowish-brown. Sternite 8 trapezoid, a pair of short lobes medially on posterior end with a shallow incision between them, microtrichia covering entire sclerite, setae on distal third. Sternite 9 wide, two gonoducts reaching genital opening, distally with a ventral slender medial blade apically bifid, genital chamber elongate, weakly sclerotized. Sternite 10 with wide medial transparent window. Tergite 8 slender medially. Tergite 9 medially slender, a pair of digitiform projections on each side each with a distal seta, and a number of short digitiform projections more medially along posterior margin. Cercomere 1 long, with a group of 3–4 large setae laterally near base and a group of three strong, short subapical setae; cercomere 2 elongate, with microtrichia and some fine setae, three stronger setae along inner margin.

**Male**. Unknown.

**Material examined** (female): ZRCBDP0047779, Nee Soon (NS1), swamp forest, 28.aug.13, MIP leg. (slide-mounted).

#### Mycetophilinae

There are 34 genera in the subfamily Mycetophilinae, 20 of which are in the Exechiini with about 700 described species, and 14 in the Mycetophilini with about 1,500 described species (Kjærandsen 2005; Magnussen 2020). The phylogeny of the subfamily has been addressed in different papers (Rindal and Søli 2006; Rindal et al. 2009; Burdíková et al. 2019), and the Exechiini emerged as a problematic taxon (Rindal et al. 2007; Burdíková et al. 2019).

The Exechiini are largely temperate in distribution—in both hemispheres. In the Mycetophilini, *Mycetophila* and some of the small genera also have a typically temperate distribution, while especially *Epicypta* is more speciose in tropical areas.

We found in Singapore only five species of three genera of Exechiini—*Allodia*, *Brachycampta*, and *Exechia*—and 53 species of Mycetophilini, quite expected for tropical environments. However, our study has the first records of the genera *Platyprosthiogyne* and *Aspidionia* for the Oriental region, and we describe a new genus belonging to the Mycetophilini.

The limited taxon sample for *Mycetophila* in all published molecular analyses does not allow to properly address the monophyly of the genus. For example, Rindal & Søli’s (2009) phylogeny only contained *Epicypta*, *Platurocypta*, *Zygomyia* Winnertz, *Sceptonia* Winnertz, *Platyprosthiogyne*, and *Aspidionia*. In the phylogenetic studies of Rindal and Søli (2006) and Rindal et al. (2009), *Platyprosthiogyne* and *Aspidionia* were not included, but *Mycetophila*, *Platurocypta* and *Epicypta* were forming a clade in Mycetophilini.

Our morphological information for Mycetophilini yields very interesting results. The group of genera with a “sharp incision in the lower border of the scutum” (Søli 2017)—including *Epicypta*, *Platurocypta*, *Aspidionia* and *Integricypta* Amorim & Oliveira, **gen. n.** —should form a clade. This is also the case of the katepisternum strongly compressed (in some keys a distinctive feature to separate *Epicypta* from *Mycetophila*), a largely developed mid coxa, the anterior end of the thorax modified to fit a compressed head, a largely reduced mediotergite and a compressed, reduced laterotergite. We are not sure about *Platyprosthiogyne* being monophyletic. The species of *Platyprosthiogyne* are plesiomorphic for all these features and the genus may be paraphyletic. *Platyprosthiogyne*— monophyletic or not as the sister genus of *Mycetophila* would make sense from the point of view of the head and thorax morphology.

With regard to the morphological nomenclature for the branches and lobes of the gonostylus, we here try to consistently apply the terms used by Magnussen et al. (2018, 2019) and Magnussen (2020) in mycetophilines. We used Kurina’s (1992) term “medioventral process” for a median extension at the posterior margin between the gonocoxites. This has been assumed to be an extension of sternite 9, even though a similar structure has originated independently more than once in the evolution of the family. It is worth clarifying that in the Mycetophilinae we call “vertex” the entire frontal area of the head capsule dorsally to the line connecting the lateral ocelli. In most flies, the entire head behind the eyes and the line of ocelli corresponds to the original “occiput” and faces posteriorly. The head of the mycetophilines is rather strongly compressed antero-posteriorly (especially in the Mycetophilini) and part of the head capsule posteriorly to the ocelli line extends dorsally and can be seen from an anterior view.

### Exechiini

Our Exechiini specimens included species of *Brachycampta*, *Rymosia*, and *Exechia*. The described species of these genera are mostly Holarctic, with relatively few known tropical areas. The Australasian and Oceanian fauna of the tribe also includes *Brevicornu* Marshall and *Synplasta* Skuse—maybe *Brevicornu* could eventually be found in Singapore.

There are over 100 described species in the genera *Allodia* and *Brachycampta* worldwide. They are predominantly Holarctic (Magnussen et al. 2019). *Allodia* and *Brachycampta* were originally proposed as separate genera, but have been treated as one genus by most authors (see a synthesis of the taxonomic history in Magnussen 2020). Kjærandsen (2007) and Magnussen (2020) discussed that they should be recognized again as separate genera (see Magnussen et al., 2022).

Part of the described species of this *Allodia-Brachycampta* complex needs reexamination to check their actual generic position. Nine species of the *Allodia-Brachycampta* complex have been described from the Oriental region—seven species from China (Wu et al. 2003), one from Sri Lanka (Senior-White 1922) and a single species from Gunong [=Mountain] Tahan, Malaya (Edwards 1928). Nine other species were added recently from the Himalayas (Nepal and Buthan) (Magnussen et al. 2019), apparently with Palaearctic affinities. There are seven species known from the Afrotropical region (Magnussen et al. 2019) and three from the Australasian region (Colless 1966).

There are clear differences in the color patterns between our species and, e.g., *Allodia micans* Edwards, from Malaya, and we consider all species of this group from Singapore new to science—three species of *Brachycampta*, besides one species of *Rymosia*, quite different from most Holarctic species of the genus. It is worth mentioning that some of the features present in the Palearctic species of *Brachycampta* are not present in the Singapore species and some taxonomic work is still needed to adjust the delimitation of the genus to include the non-Holarctic species. This also happens in some other cases, in which non-Holarctic species of mycetophilid genera do not strictly fit into the Holarctic-based diagnoses. A robust investment on the separation between *Allodia* and *Brachycampta* was recently made by Kjærandsen (2007), Magnussen et al. (2019, 2022), and Magnussen (2020). Earlier Tuomikoski (1966, p. 183) viewed *Allodia* s.l. as monophyletic, but he specifically mentioned that the reduction of the anal fold was possibly the only apomorphic in the diagnosis of the genus (p. 181). The male terminalia of *Allodia* and *Brachycampta* are particularly complex, with many different branches of the gonostylus, carefully studied by Magnussen (2020), but it is not easy to strictly establish homology of these branches or lobes. There are no conflicts between species delimitation methods in the exechiine genera (Fig. 89G).

### Brachycampta Winnertz

*Brachycampta* Winnertz 1864: 833. Type species: *Mycetophila alternans*, Winnertz, 1863, by designation of Coquillett 1910: 515. [Misidentification = *grata (*Meigen 1830)] (Further details about any possible uncertainties on the designated type species is given by Tuomikoski 1966).

**Diagnosis**: Well-developed discal bristles on the anterior part of scutum in addition to smaller flat-lying setae. Scutum and scutellum bristles with apex pointed, or somewhat splintery, never with splits of different lengths. Shape of gonostylus variable, often with a less intricate dorsal lobe and more elaborate medial and ventral lobes; hypandrial lobe prominent, with a complex outline of variable shape. Pale abdominal markings, when present, broader towards anterior margin of tergites. Base of posterior wing fork usually located basal to or opposite to base of r-m.

*Brachycampta glorialimae* Amorim & Oliveira, sp. n. (Figs. 89A–F, 90A–B)

https://singapore.biodiversity.online/species/A-Arth-Hexa-Diptera-000775

urn:lsid:zoobank.org:act:316D6A16-2556-433E-ADE4-019EE145F9F3

**Diagnosis**. Scutum light brown, yellowish along lateral margins, some few small bristles scattered on anterior half of scutum; pleural sclerites brownish, with lighter areas on anepisternum, katepisternum and mesepimeron. Sc short, fused to bR; r-m about as long as M_1+2_; R_5_ mostly straight on distal half; medial fork with a gentle constriction midway to apex; anal fold short, more or less straight, weakly sclerotized. Abdominal tergites 2–6 dark brown with cream-yellow areas on anterolateral corners, in more posterior segments extending inwards. Internal margin of ventral face of gonocoxites with a fringe of elongate setae; all gonostylus lobes slender, dorsal lobe with two branches, one digitiform directed posteriorly and one triangular directed inwards, widening towards tip; medial lobe digitiform, directed posteriorly; ventral lobe digitiform, curved, directed posteriorly on distal half; aedeagus strongly sclerotized, widening before apex.

***Brachycampta_glorialimae_ZRCBDP0049087_hapZRC BDP0049087_SMH_holotype* [55: A, 70: G, 85: T, 128: T, 154: C, 185: G, 199: T, 223: A]**

tctttcatctacaattgctcatgctggagcatcagttgatttagcaattttttcActtcatttagctggGatttcttcaattctTggagctgtaaattttattacaacaattattaatatacgagctTcaggaatttcatttgatcgaataccCttatttgtttgatcagttttaattactgctGttcttcttcttctTtctttaccagttttagctggagcAattactatattattaactgatcgaaatttaaatacatcattttttgatcctgcaggaggaggtgatccaattttatatcaacatttattt

#### Description

**Male** (Fig. 89A). Wing length, 1.92; width, 0.74. **Head** (Fig. 89B). Light brown, yellowish-brown at anterior margin of front, face greyish-brown. Head much higher than long, occiput flattened, head partially fit under anterior tip of scutum. Eyes densely covered with inter-ommatidial setulae. Antenna light brown, darker toward apex, pedicel longer than scape, both with setae on distal half, stronger setae along distal border, a strong seta dorsally; flagellomere 1 2× pedicel length, flagellomeres cylindrical, flagellomeres 2–13 about 1.4× longer than wide, covered only with setulae. Maxillary palpus light brown, whitish-yellow towards tip of distal palpomere, palpomere 1 not sclerotized, palpomere 2 bare, weakly sclerotized, palpomere 3 wider and longer, dorsal face bearing a band of sensillae, laterally and ventrally with small setae, palpomere 4 longer than palpomere 3, with scattered small setae on dorsal half, palpomere 5 inserted subapically on previous palpomere, more than twice length of palpomere 4, slender except at apex, with scattered setulae. Occiput with about eight dark setae around eye, vertex densely covered with brownish small setae. Mid ocellus minute, lateral ocelli surrounded by dark brown marking, displaced laterally, in contact with eye margin. Labella yellowish, long. **Thorax** (Fig. 89C). Scutum mostly brownish, light along margins and along dorsocentral line, scutellum brownish. Scutum clothed with short black setae, a line of stronger bristles along margin, two regular rows of strong dorsocentrals, including prescutellars. Scutellum large, with two strong scutellar bristles on posterior margin and a number of small setae covering scutellar disc. Antepronotum and proepisternum light whitish-brown, katepisternum and mesepimeron whitish-brown, darker on ventral half, anepisternum brown on anterior half, light brownish-yellow on posterior half, laterotergite light brown, darker along posterior margin, mediotergite dark brown, metepisternum whitish. A row of 4–5 antepronotal bristles and some additional small setae, proepisternum with two bristles and some small setae. Proepimeron small, rectangular, directed towards anterodorsal corner of anepisternum, bare. Anepisternum, katepisternum, mesepimeron and mediotergite bare. Laterotergite with four longer setae medially and about 20 additional small setae; metepisternum with 15–18 fine setae, one slightly stronger. Haltere light brown. **Legs**. Coxae whitish-yellow, femora yellowish-brown, tibiae and tarsi light brown, darker towards apex. Fore coxa covered with setae on anterior and internal face, setae larger towards tip; mid coxa entirely devoid of setae except on distal fifth, with some smaller and larger setae at distal margin; hind coxa with an isolate strong basal black bristle and a line of fine setae along its length, distally with setae on external and posterior faces. Femora evenly covered with small setae, some slightly longer along ventral margin close to tip. Tibiae and tarsi with regular rows of trichia. Fore tibia with few stronger short setae dorsally on distal fifth, a regular comb of setae on a small crest on external face at tip, anteroapical depressed area wide, densely lined with setulae. Mid tibia with stronger short setae on distal ¾ dorsally and laterally, external margin at tip also with a small crest bearing a regular comb of short setae; hind tibia with long irregular rows of dorsal, lateral and ventral short stronger setae, external margin at tip with a small crest bearing a regular comb of short setae. Fore leg tarsomere 1 slightly longer than tibia, some few stronger setae only at tip of tarsomeres, mid and hind tarsomeres with stronger setae ventrally and at tip. Tarsal claw with one long tooth coming out laterally from claw and two fine basal teeth. **Wing** (Fig. 89D). Membrane fumose brownish. C ending at tip of R_5_. Sc short, in contact with bR. R_1_ long, almost straight, ending at C at level of tip of M_4_. First sector of Rs short, oblique, R_5_ reaching margin before level of M_2_; r-m almost straight, oblique, slightly shorter than M_1+2_. M_1+2_ short, fork with constriction midway to apex, M_1_and M_2_ weak close to margin. M_4_ originating more basally than anterior end of M_1+2_. Cubital pseudovein not produced, CuP clearly extending beyond origin of M_4_; anal fold weakly sclerotized, only gently curved, extending to close posterior margin of anal lobe. Dorsal setae on bR, R_1_, second sector of Rs (R_5_); ventral setae on R_1_ and distal ¾ of second sector of Rs. **Abdomen**. Tergite 1 entirely brown, tergites 2–6 mostly brown, with cream-yellowish marks at anterior corners extending medially along anterior margin. Sternites whitish-yellow. **Terminalia** (Figs. 90A–B). Brownish-yellow. Gonocoxites well developed, in contact with each other medially but not fused, a row of conspicuous setae at along inner margin medially; no gonocoxite lobe dorsally or laterally projecting beyond base of gonostylus, syngonocoxite medial projection heavily sclerotized, slender, extending to level of base of gonostylus; at internal face of posterior margin ventrally, a short, curved digitiform projection with a fine seta medially. Gonostylus complex, composed of: (1) a main distal lobe with a basal setose lobe extending distally into a digitiform projection bearing setulae at distal third; (2) a median well sclerotized lobe, entirely devoid of seta, right gonostylus bearing an additional digitiform bare extension towards dorsal end of terminalia; (3) a ventral bifid lobe, more ventral branch straight with setulae along entire extension and a distal seta, dorsal branch curved, digitiform, bearing a long basal seta directed inwards and a pair of curved setae distally; and a striated internal part, extending from base of gonostylus towards dorsal face, reaching level of gonocoxal bridge. Tergite 9 small, medially divided, each part covered with microtrichia and with one strong apical bristle, besides scattered fine setae. Tergite 10 divided into a pair of elongated lobes covered with microtrichia and a long apical fine bristle. Cercus 1-segmented, small, elongate, close to each other at anterior end of terminalia, covered with fine trichia and scattered fine setae.

**Female**. As male, except for the following. **Wing l**ength, 1.95; width, 0.77. **Terminalia** (Figs. 90E–F). Sternite 8 slender, with median posterior incision reaching anterior margin, a pair of elongate lobes laterodistal lobes, fine setae restricted to lobes. Tergite 8 large, setose, with a pair of distal pointed lobose projections. Medial rhomboid extension beyond posterior margin of tergite 8. Cercus 2-segmented, cercomere 2 elongate.

#### Material examined

**Holotype**: male, ZRCBDP0049087, Nee Soon (NS1), 24.dec.14, MIP leg. (slide-mounted). **Paratype** (female): ZRCBDP0049093, Nee Soon (NS1), 24.dec.14, MIP leg. (slide-mounted).

**Etymology**. The species epithet of this species honors Gloria Lim (1930–), a mycology expert (author of >140 papers and book chapters) a first in many positions: first woman Dean of the Faculty of Science, National University of Singapore, in 1973; first woman appointee of the Public Service Commission in 1982, and first Foundation Director at the National Institute of Education. She was awarded the Public Service Star for her contributions to public service in 1993, and inducted into women’s Hall of Fame in 2014.

*Brachycampta murphyi* Amorim & Oliveira, sp. n. (Figs. 91A–G)

https://singapore.biodiversity.online/species/A-Arth-Hexa-Diptera-000785

urn:lsid:zoobank.org:act:FEF799BC-11C3-484C-B7DE-D91F089244FA

**Diagnosis**. Scutum light brown, ochre-yellowish along lateral margins, some few small bristles scattered on anterior half; pleural sclerites light ochre-yellowish, anepisternum light brown dorsoanteriorly. Sc short, fused to bR; r-m about as long as M_1+2_; R_5_ mostly straight on distal half; medial fork with a slight constriction midway to apex; anal fold short, straight, weakly sclerotized. Abdominal tergites 1–5 with a slender greyish-brown longitudinal mark medially, ochre-yellowish laterally, tergite 6 with a greyish-brown medial mark anteriorly, dark ochre-yellowish laterally and posteriorly. Gonocoxites strongly diverging, no stronger setae along inner margin at ventral face; gonostylus ventral lobe with a pointed branch directed inwards, medial lobe digitiform, directed posteriorly, dorsal lobe short, setose, well sclerotized; pair of aedeagus apodemes strongly sclerotized.

***Brachycampta_murphyi_ZRCBDP0048670_hapZRCBD P0048670_SMH_holotype* [22: C, 25: C, 28: T, 154: A, 1 99: T, 202: A, 208: A, 211: T, 220: T] --**

tatcatcaacaattgctcaCgcCggTgcatcagttgatttagctattttttctcttcatttagctggaatttcatcaattttaggagctgtaaattttattacaacaattattaatatacgagctcctggaattacttttgatcgaataccAttatttgtttgatcagtattaattacagctgttttattacttctTtcAttaccAgtTttagctggTgctattactatactattaactgatcgaaatttaaatacttcattttttgatcctgctggaggaggagatccaattttatatcaacatttattt

#### Description

**Male** (Fig. 91A). Wing length, 2.14; width, 0.80. **Head**. Yellowish-brown, occiput lighter towards ventral margin, face whitish-yellow. Occiput nearly flat, head much higher than long, partially fit under anterior end of scutum. Antennal scape and pedicel dirty whitish-yellow, flagellum light brown, darker towards apex, flagellomeres about as long as wide. Palpus light brown, labella whitish-yellow. **Thorax**. Scutum dark ochre-yellowish, with a pair of ochre-yellow bands along margins, scutellum brownish-ochre. Scutum clothed with short dark setae, some few strong bristles along margins, small bristles along weakly defined dorsocentral lines, one pair of prescutellar bristles; one pair of strong scutellar bristles and many additional small setae on disk. Scutum above wing with a small bulging area with marginal bristles. Antepronotum, proepisternum, proepimeron katepisternum, mesepimeron, and metepisternum whitish-yellow, anepisternum light brown on anterodorsal corner, dirty whitish-yellow on ventroposterior corner; laterotergite brownish-yellow, mediotergite light brown. Antepronotum with two strong bristles and many additional small setae; proepisternum with two bristles on ventral margin plus additional scattered small setae. Anepisternum, katepisternum, mesepimeron and mediotergite bare; laterotergite with about 20 longer and smaller setae. Metepisternum with 16–20 small, fine setae. Haltere yellow, with some few fine setae on pedicel and some setulae on knob. **Legs**. Coxae whitish-yellow, fore coxa yellowish on basal third; femora whitish-yellow, tibiae and tarsi light brown, darker towards tip. Hind coxa with one isolate strong basal black bristle. Mid tibia with long row of dorsal black setulae. Fore leg tarsomere 1 slightly longer than tibia. Tibial spurs over 4× longer than tibia width at apex. **Wing** (Fig. 91B). Membrane fumose brownish. C ending at tip of R_5_. Sc short, in contact with bR. R_1_ long, almost straight, ending at C at level of tip of M_4_. First sector of Rs short, oblique, R_5_ reaching margin before level of M_2_; r-m almost straight, oblique, slightly shorter than M_1+2_. M_1+2_ short, fork with constriction midway to apex, M_1_and M_2_ weak close to margin. M_4_ originating more basally than anterior end of M_1+2_. Cubital pseudovein ending at basal third of CuA, CuP inconspicuous; anal fold weakly sclerotized, curved only at distal fourth, extending to close to posterior margin of anal lobe. Dorsal setae on bR, R_1_, second sector of Rs (R_5_); ventral setae on R_1_ and distal ¾ of second sector of Rs. **Abdomen**. Tergite 1 light brown with yellowish laterals, tergites 2–5 cream-yellowish with a greyish-brown medial band, more dirty-yellowish at laterals, on more distal segments greyish-brown bands wider on posterior margin; tergite 6 greyish-brown medially on anterior margin, dirty-yellowish on posterior margin and along laterals. Sternites whitish-yellow. **Terminalia** (Figs. 91C–E). Ochre-yellowish with some darker sclerites. Gonocoxites well developed, setose, in contact only medially, not fused to each other; no projections of gonocoxite posterior margin beyond base of gonostylus dorsally or laterally, a trapezoid distal projection medially at ventral face; internal posterior corner of gonocoxite on dorsal face strongly sclerotized. Gonostylus complex, composed of: (1) an elongate, regularly-sclerotized setose dorsal lobe with lateral blade-like lateral extensions and a distal comb of setulae; (2) a medial lobe with a wider base bearing a pair of long setae and a digitiform extension inwards with two long subapical setae and a slightly falciform apex; (3) a digitiform ventral lobe, with a pair of long setae on internal face, some scattered small setae and a group of distal small and stronger setae; (4) a striated, wide internal part. Gonocoxal bridge wide. Aedeagal-parameral complex including a medial, well-sclerotized large sclerite with two pairs of short distal projections; an additional sclerite with an anterior medial area connecting a posterior membrane that ends slightly beyond insertion of gonostylus, posterior margin with a median shallow rounded incision. Tergite 9 present as a pair of elongate short lobes barely connecting anteriorly, ending with a strong apical bristle. Tergite 10 divided into a pair of elongate lobes covered with microtrichia and a long seta, ending close to level of base of gonostylus. Cerci small, elongate.

**Female**. As male, except for the following. **Wing l**ength, 2.24; width, 0.90. **Terminalia** (Figs. 91F–G). Sternite 8 slender, long, hypoginial valves long, separate medially by a deep incision, setulae restrict to valves. Tergite 8 short, wide, weakly sclerotized, entirely devoid of setae. Sternite 9 weakly sclerotized, oblong, with gonopore medially. Tergite 9, tergite 10 and cerci not recognizable as independent sclerites, a single trapezoid very elongate sclerite present, with a medial slender posterior incision, setulae on posterior half and a pair of long setae at posterior margin.

#### Material examined

**Holotype**: male, ZRCBDP0048670, Nee Soon (NS1), swamp forest, 04.apr. 12, MIP leg. (slide-mounted). **Paratypes** (2 females): ZRCBDP0048976, Nee Soon (NS1), 13.may.15, MIP leg.; ZRCBDP0048669, Nee Soon (NS1), swamp forest, 04.apr. 12, MIP leg. (slide-mounted).

**Etymology**. The species epithet honors Dennis Hugh “Paddy” Murphy (1931-2020). As a British citizen who moved to Singapore in the 1960s, his wealth of expertise in entomology made him an associate professor at then-University of Malaya in 1983, later National University of Singapore (NUS). Murphy was recognized as one of the most prominent entomologists in Southeast Asia, for his work on the Mangroves of Singapore, and has mentored a generation of zoologists in Singapore.

**Remarks**. We have two haplotypes for this species and no delimitation conflicts.

*Brachycampta limtzepengi* Amorim & Oliveira, sp. n. (Figs. 92A–E)

https://singapore.biodiversity.online/species/A-Arth-Hexa-Diptera-000754

urn:lsid:zoobank.org:act:41980F21-ED7D-43E1-84BD-ABF0C641D5DF

**Diagnosis**. Scutum with light brown longitudinal bands connecting posteriorly over a dark ochre-yellowish background, scattered small bristles on anterior half of scutum; pleural sclerites light ochre-yellowish, anepisternum light brown dorsoanteriorly. Sc short, fused to bR; r-m about as long as M_1+2_; R_5_ mostly straight on distal half; medial fork with a gentle constriction midway to apex; anal fold mostly straight, weakly sclerotized. Abdominal tergites 1–5 with a slender light brown longitudinal mark medially, ochre-yellowish laterally, tergite 6 light brown on anterior half, ochre-yellowish on posterior half. Gonocoxites strongly diverging, no stronger setae along inner margin at ventral face; gonostylus ventral lobe with a pointed branch directed inwards, medial lobe digitiform, directed posteriorly, dorsal lobe short, setose, well sclerotized; pair of aedeagus apodemes strongly sclerotized.

***Brachycampta_limtzepengi_ZRCBDP0047881_hapZRC BDP0047881_SMH_holotype* [16: C, 22: C, 88: T, 130: C, 199: C, 205: C, 220: T, 277: C]**

tctttcttcaacaatCgctcaCgcaggagcatcagttgatttagcaattttttctctacatttagcaggaatttcatctattttaggTgctgtaaattttattacaactattattaatatacgagccccCggaattaatttcgatcgaatacctttatttgtttgatcagttttaattactgctgttttacttcttctCtctctCccagttttagctggTgctattactatattattaactgatcgaaatttaaatacttcattttttgacccagcCggtggaggtgaccctattctttaccaacatttattt

#### Description

**Male** (Fig. 92A). Wing length, 2.05; width, 0.80. **Head**. Brownish-yellow, occiput lighter towards ventral margin, face whitish-yellow. Vertex elongate, head antero-posteriorly compressed. Antennal scape and pedicel dirty whitish-yellow, flagellum light brown, darker towards apex, flagellomeres 1.2 longer than wide. Palpus light brown, lighter towards apex. Labella yellowish. Six bristles on occiput dorsally around eye, vertex densely covered with brownish setulae, a row of dark setae along ventral margin of frons. **Thorax**. Scutum light ochre-brown, with a pair of slender yellowish bands over dorsocentral line connecting posteriorly and a band along lateral margins; scutellum light ochre-brown with a medial yellowish band. Scutum clothed with short dark setae, some few strong bristles along margins, irregular rows of dorsocentrals, and some longer prescutellars; scutellum with a pair of strong scutellar bristles on posterior border and scattered setae on disc. Scutum above wing with a small bulging area with marginal bristles. Antepronotum, proepisternum, proepimeron katepisternum, mesepimeron, and metepisternum whitish-yellow, anepisternum light brown on anterodorsal corner, dirty whitish-yellow on ventroposterior corner; laterotergite brownish-yellow, darker along posterior margin, mediotergite light brown. Antepronotum with two bristle and additional smaller setae; proepisternals with two bristles apart from ventral margin and some additional smaller setae. Anepisternum, katepisternum, and mesepimeron bare, laterotergite with four longer setae and 12 smaller setae; metepisternum with six setulae and only one longer seta. Haltere light yellowish-brown, setulae on distal half of pedicel and on anterior face of knob. **Legs**. Coxae light yellowish-brown; femora light brownish-yellow, tibiae and tarsi light brown, darker towards tip. Fore leg tarsomere 1 about as long as tibia. Tibial spurs over 4× longer than tibia width at apex. **Wing** (Fig. 92B). Membrane greyish-brown. **Abdomen**. Tergite 1 light brown, tergites 2–5 with a light brown medial band, dirty-yellowish at laterals; tergite 6 light brown on anterior margin, dirty-yellowish on posterior half. Sternites whitish-yellow, distal sternite slightly darker. **Terminalia** (Fig. 92C–E). Ochre-yellowish with some dark brown sclerites. Gonocoxites well developed, setose, only in contact medially not fused to each other; no projections of gonocoxite posterior margin beyond base of gonostylus dorsally or ventrally [medial projection of syngonocoxite not clear on slide-mounting]. Gonostylus complex, composed of: (1) an elongate, dorsal lobe with valve-shape main body, setose externally, with a wide blade-like internal extension bearing a comb of setae along entire distal margin; (2) a digitiform ventral lobe with a pair of basal long setae, a group of four long subapical setae directed inwards and a group of six smaller setae along distal margin; (3) a curved, digitiform median lobe with three setae at apex; (4) a large striated internal part that extends ventrally into a small digitiform projection with setulae at apex. Gonocoxal bridge wide. Aedeagal-parameral complex with a well-sclerotized sclerite, with an ejaculatory apodeme projected anteriorly, distally as a subtriangular membrane, rounded tip not reaching level of insertion of gonostylus. Tergite 9 present as a pair of elongate short lobes barely connecting anteriorly that end with a strong apical bristle. Tergite 10 divided into a pair of elongate lobes covered with microtrichia and a long seta, ending close to level of base of gonostylus. Cerci small elongate.

**Female**. Unknown.

#### Material examined

**Holotype**: male, ZRCBDP0047881, Nee Soon (NS1), swamp forest, 25.set.13, MIP leg. (slide-mounted). **Paratypes** (7 males): ZRCBDP0047943, Nee Soon (NS2), swamp forest, 20.nov.13, MIP leg.; ZRCBDP0048511, Nee Soon (NS1), swamp forest, 23.may. 12, MIP leg. (website photo specimen); ZRCBDP0048512, Nee Soon (NS1), swamp forest, 04.apr. 12, MIP leg.; ZRCBDP0048709, Nee Soon (NS2), 28.jan.15, MIP leg.; ZRCBDP0049080, Nee Soon (NS1), 24.dec.14, MIP leg.; ZRCBDP0049094, Nee Soon (NS1), 24.dec.14, MIP leg.; ZRCBDP0049097, Nee Soon (NS1), 24.dec.14, MIP leg.

**Etymology**. The species epithet honors Lim Tze Peng (1921–), one of the Singapore-born most significant artists and a living legend, renowned for Chinese ink drawings and paintings of post-independence Singapore, with masterpieces exhibited in the Singapore Art Museum and Nanyang Academy of Fine Arts. He has been bestowed several awards, including the Special Prize at the Commonwealth Art Exhibition in England (1977) and the prestigious Cultural Medallion in Singapore (2003).

### *Rymosia* Winnertz

*Rymosia* Winnertz 1863: 810. Type-species, *Mycetophila discoidea* Meigen (Johannsen, 1909: 102) = *Rymosia fasciata* (Meigen) (designated by Johannsen, 1909).

**Diagnosis** (modified from Tuomikoski, 1966). No conspicuous macrotrichia on flagellomeres, flagellomeres slightly longer than wide. Clypeus short, broadly truncate. Palpomere 3 with a sharply delimited rounded sensory pit. Proepisternum with one or two bristles; anepisternum bare or with a few minute setulae. Sc short, ending free; posterior fork typically long, basal half narrow, then abruptly wider, anal fold long and distinct. Medial fork, cubital fork, r-m, and bm bare of macrotrichia. Hind coxae with one or two basal bristles; pale markings of abdomen situated towards bases of tergites. Female cercus one-segmented.

*Rymosia teopohlengi* Amorim & Oliveira, sp. n. (Figs. 93A–E)

https://singapore.biodiversity.online/species/A-Arth-Hexa-Diptera-000833

urn:lsid:zoobank.org:act:76C14FF5-F8CB-47BE-A88A-1377C1E332DA

**Diagnosis**. Scutum brown, a whitish band along lateral margin, with a dark brown rounded mark above wing, no bristles over entire scutum; pleural sclerites mostly brownish, lighter areas on anepisternum, katepisternum, and mesepimeron, anepisternum mostly dark brown. Sc short, ending free; r-m 0.76× M_1+2_ length; R_5_ curved posteriorly on distal fourth of wing; no constriction on medial fork; anal fold long, gently curved, well-sclerotized. Abdominal tergites 2–5 with a slender yellowish-brown longitudinal band extending laterally along posterior margin, a large ochre-yellowish mark on anterolateral corners, tergite 6 yellowish-brown. Incision short between gonocoxites at ventral face. Gonostylus with a small, narrow ventral branch, a broad, bulbuous interior branch and row of 5 seta apically on anterior branch; aedeagus spoon-shaped, flat, blade-like.

***Rymosia_teopohlengi_ZRCBDP0049238_hapZRCBDP0 049238_SMH_holotype* [106: C, 142: C, 173: C, 187: C, 211: C, 223: A, 286: C]**

cctttcttcatcaattgctcatgctggggcatcagttgatttagctattttttctcttcatttagcaggaatctcttcaattttaggtgctgtaaattttattacCacaatt attaatatacgagctcctggaatttcattCgatcgaatacctttatttgtttgatctgttCtaattactgcagtCttattacttttatctttaccagtCctagctggagcAattacaatattattaacagatcgaaatttaaatacatctttcttcgatcctgcaggaggaggCgacccaatcctttatcaacacttattt

#### Description

**Male** (Fig. 93A). Wing length, 2.75; width, 0.90. **Head**. Vertex brownish, front darker, occiput light brown. Face and clypeus greyish-brown. Head quite compressed antero-posteriorly, much higher than long, partially fit under anterior tip of scutum. Maxillary palpus light brown, whitish-yellow towards tip of distal palpomere. Eyes densely covered with inter-ommatidial setulae. Antenna light brown, darker towards apex, pedicel longer than scape; pedicel with a group of concentrated small setae on dorsal half of internal face, one strong seta ventrally on posterior border; pedicel with two strong setae ventrally on posterior margin and one strong dorsal seta distally; flagellomeres cylindrical, flagellum 1 1.5× flagellomere 2 length, flagellomeres 2–13 about 1.4× longer than wide, covered only with setulae. Maxillary palpus light brown, whitish-yellow towards tip of distal palpomere, palpomere 2 bare, weakly sclerotized, palpomere 3 wider and longer, with a wide dorsal area lined with sensillae and small setae laterally and ventrally, palpomere 4 longer than palpomere 3, with scattered small setae, palpomere 5 inserted subapically on palpomere 4, more than twice its length, slender along most its length, widening at apex, with scattered setulae. Occiput with 8 dark setae posteriorly around eye, vertex densely covered with brownish small setae. Mid ocellus minute, lateral ocelli on a dark brown background, displaced laterally, in touch with eye margin. Labella yellowish, long. **Thorax** (Fig. 93B). Scutum light brown, with a whitish-yellow marginal band that extends from level of anterior spiracle to level of scuto-scutellar suture, a dark brown macula above wing and along scutum-scutellar suture, scutellum brown. Scutum densely clothed with short setae, stronger setae only along margins posteriorly to level of anepisternum, scutum with a sclerotized lobe above wing. Scutellum large, one pair of strong scutellar bristles at posterior margin, large number of small setae covering most of disk. Basisternum, antepronotum, and proepisternum light brown, proepimeron brown, anepisternum dark brown except for whitish ventral fifth, katepisternum brown on ventral half, whitish at dorsal half, mesepimeron whitish, laterotergite light brown, darker along posterior margin, mediotergite and metepisternum light brown. Basisternum bare, dorsoposterior arms largely developed; two strong antepronotal bristles and scattered small fine setae, proepisternum with one strong bristle and additional fine setae. Anepisternum, katepisternum, and mesepimeron bare; about 20 fine and 3 longer setae distributed along laterotergite length; metepisternum with a group of 16 to 18 small setae concentrated on posterior third. Mesepimeron reaching ventral margin of pleura. Haltere yellow, longer fine setae along pedicel, fine setae on basal half of knob. **Legs**. Coxae whitish-yellow, femora yellowish-brown, hind femur with a brownish tinge dorsally and ventrally, tibiae and tarsi light brown, progressively darker. Fore coxa covered with setae on anterior and internal faces, setae larger towards tip; mid coxa entirely devoid of setae except on distal fifth, with some smaller and larger setae at distal margin; hind coxa with a strong isolate basal black seta and a line of fine setae along its length, distally with setae on external and posterior faces. Femora evenly covered with small setae, some slightly longer setae only ventrally close to tip. Tibiae and tarsi with regular rows of trichia. Fore tibia with few stronger short setae dorsally on distal fifth, a regular comb of setae on a small crest on external face distally, anteroapical depressed area wide, densely lined with setulae. Mid tibia with stronger short setae on distal ¾ dorsally and laterally, with a regular comb of short setae on a small crest distally on external face; hind tibia with long irregular rows of dorsal, lateral and ventral short stronger setae, a small crest with a regular comb of short setae at external face distally. Tibiae and tarsi with regular rows of trichia. Fore tibia with short setae only at distal end, a regular comb of setae on a small distal crest on external face distally, anteroapical depressed area wide, densely lined with setulae. Mid tibia with stronger short setae on more or less irregular lines dorsally and laterally, a small crest with a regular comb of short setae at external margin distally; hind tibia with irregular rows of dorsal, lateral, and ventral short stronger setae along entire length, a small crest with regular comb of short setae at external margin distally. Fore leg tarsomere 1 longer than tibia. Tarsi with some few stronger setae on first and second tarsomeres; fore tarsomeres 3–5 modified for grasping or hanging, tarsomere 3 with regular rows of trichia, some stronger setae on both sides and a row of short, blunt spines ventrally on distal half, tarsomere 4 with a row of blunt spines along entire length, regular rows of trichia absent, with scattered short erected setae instead, tarsomere 5 only with short erect setae; mid and hind tarsomeres 3–5 only with rows of trichia and setae at distal end. Tibial spurs over 5× longer than tibia width at apex. Tarsal claw with one long tooth coming out dorsolaterally from claw and two fine basal teeth. **Wing** (Fig. 93C). Membrane fumose brownish. C ending at tip of R_5;_ Sc short, ending free, not close to R, humeral vein present. R_1_ long, reaching C almost at level of tip of M_2_, gently curved posterior close to margin. First sector of Rs short, transverse, R_5_ reaching C at level of M_1_, curved posteriorly at distal fifth; r-m almost straight, slightly oblique, 0.71× M_1+2_ length. M_1+2_ short, M_1_ gently curved anteriorly close to apex, M_1_and M_2_ weak distally. M_4_ originating way more basally than anterior end of M_1+2_. Cubital pseudovein clearly produced, long; CuP reaching over half of second sector of M_4_; anal fold well-developed, extending to close posterior margin of anal lobe. Dorsal setae on bR, R_1_, second sector of Rs (= R_5_), one single dorsal seta on Sc; ventral setae on distal half of R_1_ and entire length of second sector of Rs. No macrotrichia on wing membrane.

**Abdomen**. Tergite 1 entirely light brown, tergites 2–5 brown medially at anterior margin, expanding laterally towards posterior margin, yellowish at anterolateral corners; tergite 6almost entirely light brown. Sternites whitish-yellow. **Terminalia** (Fig. 93D–E). Light brownish-yellow, darker towards tip. Gonocoxites well-developed, setose, in contact with each other on anterior two-thirds, no median suture; no projections of gonocoxite posterior margin beyond base of gonostylus dorsally or ventrally; syngonocoxite medial projection with a basal neck, distally wider, blunt, barely reaching level of base of gonostylus. Gonostylus complex, composed of: (1) a short, setose, basal lobe; (2) a median, blade-like, flat lobe bifid distally, entirely devoid of setae; (3) a ventral, strongly curved lobe extending distally beyond rest of terminalia, with setae along inner face of basal half and with setae all over on distal half; (4) a complex, non-striated internal part, composed of a small lobose and setose branch, a blade-like wide falciform branch, and a wide, rather weakly sclerotized branch with a row of weak setae along margin bearing a digitiform distal projection with four strong, curved setae distally. Aedeagal-parameral complex with a pair of anterior apodemes diverging from each other, connected more distally, a medial elongate extension with a number of spinules on distal half. Tergite 9 wide, trapezoid, with a median shallow incision on posterior margin. Tergite 10 divided into a pair of elongate lobes covered with microtrichia and a long apical fine bristle. Cerci not detected.

**Female**. Unknown.

#### Material examined

**Holotype**: male, ZRCBDP0049238, Nee Soon (NS1), 10.dec.14, MIP leg. (website photo specimen, slide-mounted). **Additional sequenced specimens**: ZRCBDP0133914, Singapore; ZRCBDP0133932.

**Etymology**. The specific epithet honors the first notable Singaporean poet to publish in English, Teo Poh Leng (1912-1942). Malayan poet and teacher, he lived in the Crown Colony of Singapore and his modernist F. M. S. R. poem was published in 1937 in London, under the pseudonym of Francis P. Ng. He is accepted to be the first person from Singapore to have had a book-length publication in English.

**Remarks**. There are two haplotypes for *Rymosia teopohlengi* Amorim & Oliveira, **sp. n.** and all delimitation approaches keep them together. This *Rymosia* species is quite unique compared to the remaining Exechiini species and even compared to other species of *Rymosia*. Firstly, the thorax color pattern is unique. Secondly, R_5_ runs parallel to R_1_ on the basal two-thirds. Thirdly, r-m is quite more longitudinal. Fourthly, there is no constriction of the medial fork midway to apex and M_4_ only gradually diverges from CuA basally, with a slender anterior end of the posterior fork, in a rather typical *Rymosia* wing venation pattern. This species could be close to the European species *Rymosia signatipes* (Wulp). Like in *R. signatipes*, the male of *R. teohpolengi*, **sp.n.** has blunted, gripping spines on the foretarsi 3 and 4, and the female gonapophysis 9 is distinctly upcurved apically. *R. teohpolengi*, **sp.n.** differs from the European species in the exposed spoon-shaped hypandrial lobe (which has a narrow, triangular shape and more in-bent in *R. signatipes*), in the ventrocaudal margin of the gonocoxites being moderately excavated (convex in R. *signatipes*) and in details of the gonostylus, as the L-shaped dorsal branch and the sharply tipped ventral branch.

### *Exechia* Winnertz

*Exechia* Winnertz, 1863: 879. Type species: *Tipula fungorum* De Geer, 1776 [= *Mycetophila fusca* Meigen, 1804], des. Johannsen 1909: 106.

**Diagnosis** (modified from Tuomikoski 1966). Flagellomeres somewhat longer than wide, no conspicuous macrotrichia. Palpomere 3 short, with ovate sensory pit. Scutum and scutellum covered with pale, decumbent setae directed backwards; acrostichal and dorsocentral well developed rows of bristles, separated from marginal bristles by rather narrow areas (less often discal bristles reduced or absent); scutellum with two strong marginal bristles. Antepronotum and proepisternum each with two to four stronger bristles; anepisternum usually bare (in some larger species with minute setae). Wing membrane with microtrichia arranged regular rows; macrotrichia on anal lobe only in some species; medial fork and posterior fork bare. Sc short, ending free; R_1_ about as long as or slightly longer than bR; R_5_ almost straight, gently curved posteriorly close to tip; M_1_ clearly depressed on distal half, diverging from R_5_; r-m oblique, long, bare or with some macrotrichia at distal end, at least twice as long as M_1+2_. Posterior fork short, origin of M_4_ distinctly beyond that of medial fork; second sector of CuA straight. Fore leg tarsomere 1 slightly usually longer or slightly shorter than tibia; hind leg tarsomere 1 about twice as long as outer hind tibial spur; hind coxa with a single strong basal bristle; hind tibia with row of posterior bristles towards tip. Abdomen, if with pale markings, usually on anterior half of tergite. Male tergite 9 divided into two separate, somewhat pointed halves, each with one or two longer bristles at tip; sternite 9 usually inconspicuous, more or less fused with gonocoxites; gonostylus without a large blunt, dorsally striate inner lobe.

*Exechia* is a rather large genus with over 150 described species, over two thirds of which from the Holarctic regions (Lindemann et al. 2021) and there are no comprehensive revisions of the genus. The currently accepted limits of *Exechia* are based on Tuomikoski’s (1966) phylogenetic system for the Exechiini, describing *Pseudexechia* Tuomikoski and *Exechiopsis* Tuomikoski as separate genera. *Exechia* is characterized by a rather distal origin of M_4_, producing a short posterior fork. Tuomikoski (1966, p. 174) considered carefully the delimitation of *Exechia* and specifically mentions that “the possibility cannot be excluded that in *Exechia* the shortness of the posterior fork is to some extent a convergent rather than a true synapomorphic character, in which case the genus, in its traditional delimitation, cannot be maintained in a more natural and phylogenetic system.” In summary, the short posterior fork is an apomorphic condition, but this may have originated more than once in the Exechiini. Indeed, Edwards (1925) already recognized two separate groups in the genus. And *Exechia* may still be polyphyletic (Tuomikoski 1966). Only a study with worldwide scope of *Exechia* and other members of the tribe may properly answer these question.

There are 31 Oriental species of *Exechia*, 12 of which known from China and Nepal, and 16 from India and Sri Lanka. There are only three species described from Southeast Asia, two of them from the mountains in Gunong Tahan, Malaysia (Edwards 1928), and one from Mount Dulit, Borneo (Edwards 1926). Neither the descriptions nor the illustrations of the male terminalia of these species suggest conspecificity with the species we collected in Singapore.

The two species of *Exechia* described here differ significantly from each other. This includes the body color patterns and the male terminalia morphology, and there are also significant differences in wing venation. The homology or male terminalia sclerites in this clade of exechiine genera is detailed in Lindermann (2021). *Exechia tanswiehiani* Amorim & Oliveira, **sp. n.** has a longer posterior fork and M_1+2_ is nearly absent, while *E. alinewongae* Amorim & Oliveira, **sp. n.** has M_4_ originating more distally and M_1+2_is about half of the length of r-m. There are two haplotypes for *Exechia tanswiehiani* Amorim & Oliveira, **sp. n.** and only one haplotype for *Exechia alinewongae* Amorim & Oliveira, **sp. n.**; mPTP indicates a single species, while all other criteria clearly separate both species (Fig. 96G).

*Exechia tanswiehiani* Amorim & Oliveira, sp. n. (Figs. 94A–C, 95A–C)

https://singapore.biodiversity.online/species/A-Arth-Hexa-Diptera-000755

urn:lsid:zoobank.org:act:D751226D-E713-4560-9280-E394BC500BBE

**Diagnosis**. Head light brown, with ochreous-yellowish areas more ventrally. Scutum dark ochre-yellowish, lighter along lateral margins; a row of black elongate bristles along anterior margin of scutum; pleural sclerites light ochre-yellowish with lighter areas. R_5_ straight on basal third; M_1+2_ very short, medial fork with a constriction on distal third of wing; M_4_ originating almost at level of origin of Rs, posterior fork relatively long; anal fold well sclerotized. Abdominal tergites mostly ochre-yellow, a dark brown longitudinal mark medially on segments 1, 4–6; segment 7 elongate. Gonocoxites with a deep medial incision between them, no lobes projecting beyond insertion of gonostylus; gonostylus with a ventral short, pointed branch directed inwards and a medial, short, well-sclerotized branch directed medio-posteriorly, with a long, distinctive seta.

***Exechia_tanswiehiani_ZRCBDP0047947_hapZRCBDP0 047798_SMH_holotype* [7: C, 28: T, 40: C, 91: G, 149: C, 151: C, 154: A, 155: C, 157: T, 244: C]**

cctatcCtcatcaattgctcatgcaggTgcttctgttgaCttagctattttttctttacatttagctggtatttcttctattttaggagcGgttaattttattactacaattattaatatacgagcacctggaattacctttgatcgtCtCccACtTtttgtttgatcagtattaattacagctgtacttcttcttctttctttacctgttttagctggagctattactatactattaactgaCcgaaatttaaatacttccttttttgaccctgctggaggaggagatccaattttatatcaacatttattt

#### Description

**Male**. Wing length, 3.20; width, 1.06. **Head**. Vertex dark brown, ventral end of occiput and face brown, clypeus light brown. Vertex with seven small bristles posteriorly around eye, frons with a line of setae along anterior margin. Face wide, covered with small setae, clypeus bulging, covered with small setae. Antennal scape and pedicel light brownish-yellow, basal flagellomeres light brown, distal flagellomeres brown, basal two flagellomeres yellowish; scape about twice pedicel length; scape with setulae dorsally and ventrally along distal two-thirds, some stronger setae ventrally and one stronger seta dorsally on distal margin; pedicel with setulae all around distal two-thirds, a group of ventral strong setae and one dorsal strong seta on distal margin; flagellomere 1 about 1.4× flagellomere 2 length, flagellomeres 4 length 1.4× width. Palpus yellowish-brown, lighter towards tip, five palpomeres; palpomeres 1 and 2 weakly sclerotized, palpomere 3 well-developed, with a conspicuous subapical sensory pit, dorsally and laterally covered with small setae, palpomere 4 elongate, slightly longer than length of palpomere 3, covered with setulae, dorsal setae slightly longer, palpomere 5 2.5× palpomere 4 length, covered with setulae. Labellae brownish-yellow, largely developed backwards. **Thorax** (Fig. 95A). Scutum dark ochre-yellow, with a pair of light brown marks along dorsocentral lines, posterior end of scutum with dark markings, scutellum light brown. Scattered short setae over scutum, a regular row of stronger dorsocentrals and a line of bristles along lateral margin, a pair of prescutellar bristles medially and three prescutellar bristles at each lateroposterior corner; scutellum large, with a pair of strong bristles at posterior margin, two pairs of subterminal stronger setae, and small setae covering most of scutellum disc. Pleural sclerites light brownish-yellow except for a brownish mark on proepisternum and for brownish dorsal third of mediotergite. Antepronotum large, with two stronger bristles and many scattered setae; proepisternum with three strong bristles directed ventrally and many small setae; anepisternum, katepisternum, mesepimeron, and mediotergite bare; laterotergite with a large number of setulae on ventral and dorsal third, nine long setae medially; metepisternum with 15–17 setulae on posterior half and two stronger setae close to posterior margin. **Legs**. Coxae light whitish-yellow, mid and hind coxae more yellowish, femora light brownish-yellow; tibiae and tarsi light brown. Fore coxa covered with fine small setae on anterior and lateral faces, a row of strong setae along distal ⅔ of lateroposterior edge; mid coxa with small setae on distal third besides distal stronger setae anteriorly and one strong seta on posterior face at distal margin; hind coxa with one strong seta on lateral face dorsally, a row of small setae along almost entire length laterally, and some few long setae and one bristle laterally on distal end. Tibiae with dorsal and lateral regular rows of trichia. Tibiae with irregular dorsal and lateral rows of setae, stronger on mid and hind tibiae, distally with a small crest with regular comb of short setae at external margin. Anteroapical depressed area on inner face of fore tibia wide, lined with setulae. Tarsomeres 1 and 2 with irregular rows of dorsal and lateral setae, besides trichia and distal stronger setae ventrally, tarsomeres 3 and 4 with a single seta dorsally, besides trichia and distal stronger setae, tarsomere 5 only with trichia. Tarsal claw with a short blunt tooth and a long slender tooth almost as long as claw. Inner tibial spur over 5× tibia width at apex. **Wing** (Fig. 94B). Membrane light fumose, unmarked. C not extending beyond tip of R_5_, some few setulae dorsally on anal lobe membrane. Humeral vein present, oblique, Sc short, ending free. First sector of Rs short, slightly oblique; R_5_ gently curved posteriorly on distal fifth of extending slightly beyond base of gonostylus, aedeagus bifid distally, extending to level of tip of gonostylus. Tergite 9 present as a pair of lobes connected only at anterior end, microtrichia and fine setae along distal half, a strong seta at tip of each lobe. No sign of a separate tergite 10. Cercus long, extending beyond posterior margin of gonocoxite dorsally, covered with microtrichia and setae, especially along internal margin, a longer seta at tip. **Female** (Fig. 94A). As male, except for the following. **Wing l**ength, 3.30; width, 1.09. **Abdomen**. Tergites cream-yellowish, tergites 1–2 darker medially, tergites 4–5 with dark brown medial mark. **Terminalia** (Fig. 94C). Terminalia conical, elongate. Sternite 8 slender, very long, no posterior incision or lobes. Tergite 8 large, with median incision at posterior end medially, lateral borders extending towards ventral face of terminalia. Tergite 9 composing a conical membranous structure projecting beyond distal end of tergite 8, entirely bare. Distally on ventral face, sternite 10 membranous, with a pair of rows elongate fine setae and a pair of small distal dark setae; cerci 1-segmented with microtrichia, scattered setulae, an internal row of elongate fine setae and a pair of distal longer setae.

#### Material examined

**Holotype**: male, ZRCBDP0047947, Nee Soon (NS1), swamp forest, 03.jul.13, MIP leg. (slide-mounted). **Paratypes**: 2 males, 10 females. **Males**: ZRCBDP0048945, Nee Soon (NS2), 03.dec.14, MIP leg.; ZRCBDP0049174, Nee Soon (NS2), wing; r-m over 4× longer than M _1+2_, oblique; M1+2 very may.15, MIP leg. **Females**: ZRCBDP0047798, Nee Soon (NS2), swamp forest, 13.nov.13, MIP leg.; ZRCBDP0047898, short, medial fork long, M_1_ diverging from R_5_ on distal half, slightly divergent from M_2_ at apex; tip of M_1_, M_2_, and M_4_ weakly sclerotized. M_4_ originating slightly beyond level of origin of Rs. Cubital pseudovein barely extending beyond origin of M_4_; CuP nearly absent, anal fold clearly produced, gently curved. Distal third of R_1_ and distal ⅔ of Rs with ventral setae, bR, R_1_ and Rs with dorsal setae. **Abdomen**. Tergites 1–2 and tergite 4–7 ochre-yellowish with a dark brown medial mark, tergites 3 entirely ochre-yellowish. Sternites 1–7 cream-yellow. **Terminalia** (Figs. 95B–C). Yellowish, with dark brown marking ventrodistally. Gonocoxites large, fused at anterior fifth of terminalia, with a deep medial posterior incision, distal end of posterior margin internally slightly more projected, an acute oblique projection inwards and a short rounded lateral lobe extending slightly beyond base of gonostylus. Gonostylus complex, with: (1) a short basal lobe with one long fine seta; (2) a main distal lobe with a long seta on basal third, a subapical strong bristle and a sclerotized acute distal extension; (3) a ventral ovoid lobe pointed distally with a strong seta; (4) a medial, well sclerotized bifid lobe, ventral branch bare and secondarily bifurcated, dorsal branch short, digitiform, with elongate fine setae on distal half; (5) a long dorsal lobe with long setulae along distal half of internal margin and an apical long setae. A wide gonocoxal bridge without apodemes. An elongate aedeagal-parameral complex, with an anterior ejaculatory apodeme, and a pair of long parameral slender blades extending slightly beyond base of gonostylus, aedeagus bifid distally, extending to level of tip of gonostylus. Tergite 9 present as a pair of lobes connected only at anterior end, microtrichia and fine setae along distal half, a strong seta at tip of each lobe. No sign of a separate tergite 10. Cercus long, extending beyond posterior margin of gonocoxite dorsally, covered with microtrichia and setae, especially along internal margin, a longer seta at tip.

**Female** (Fig. 94A). As male, except for the following. Wing length, 3.30; width, 1.09. Abdomen. Tergites cream-yellowish, tergites 1-2 darker medially, tergites 4-5 with dark brown medial mark. Terminalia (Fig. 94C). Terminalia conical, elongate. Sternite 8 slender, very long, no posterior incision or lobes. Tergite 8 large, with median incision at posterior end medially, lateral borders extending towards ventral face of terminalia. Tergite 9 composing a conical membranous structure projecting beyond distal end of tergite 8, entirely bare. Distally on ventral face, sternite 10 membranous, with a pair of rows elongate fine setae and a pair of small distal dark setae; cerci 1-segmented with microtrichia, scattered setulae, an internal row of elongate fine setae and a pair of distal longer setae.

**Material examined**

Holotype: male, ZRCBDP0047947, Nee Soon (NS1), swamp forest, 03.jul.13, MIP leg. (slide-mounted). Paratypes: 2 males, 10 females. Males: ZRCBDP0048945, Nee Soon (NS2), 03.dec.14, MIP leg.; ZRCBDP0049174, Nee Soon (NS2), 13.may.15, MIP leg. Females: ZRCBDP0047798, Nee Soon (NS2), swamp forest, 13.nov.13, MIP leg.; ZRCBDP0047898, Nee Soon (NS2), swamp forest, 16.oct.13, MIP leg.; ZRCBDP0047899, Nee Soon (NS2), swamp forest, 16.oct.13, MIP leg.; ZRCBDP0048514, Nee Soon (NS1), swamp forest, 04.apr. 12, MIP leg. Female (website photo specimen); ZRCBDP0048668, Nee Soon (NS1), swamp forest, 23.may. 12, MIP leg. (slide-mounted); ZRCBDP0048855, Nee Soon (NS1), jan.15, MIP leg.; ZRCBDP0048946, Nee Soon (NS2), 03.dec.14, MIP leg.; ZRCBDP0048972, Nee Soon (NS1), 13.may.15, MIP leg.; ZRCBDP0048985, Nee Soon (NS2), 17.dec.14, MIP leg.; ZRCBDP0049179, Nee Soon (NS2), may.15, MIP leg.

**Etymology**. The species epithet of this species honors Tan Swie Hian (1943–), a Singaporean multidisciplinary artist known for his contemporary Chinese calligraphy, Chinese poetry and contemporary art sculptures, found in Singapore and many parts of the world.

**Remarks**. We have two haplotypes for this species, but no delimitation conflicts.

*Exechia alinewongae* Amorim & Oliveira, sp. n. (Figs. 96A–F)

urn:lsid:zoobank.org:act:934D9A59-2C49-4190-A152-7A4CF6DC3A3E

**Diagnosis**. Head dark brown, lighter more ventrally. Scutum dark ochre-yellowish, a pair of light brown marks along dorsocentral lines; a regular row of stronger dorsocentrals and a line of bristles along lateral margin; pleural sclerites light brownish-yellow, except for brownish mark on proepisternum and brownish dorsal third of mediotergite. R_5_ slightly sinuous on basal third; M_1+2_ half of r-m length, medial fork with a constriction on distal third of wing; M_4_ originating clearly beyond level of origin of Rs, posterior fork relatively short; anal fold weakly sclerotized. Tergite 1 ochre-yellowish with a light brown mark medially, tergites 2–3 entirely light brown, tergites 4–6 ochre-yellowish, dark brown medially, tergite 7 dark ochre-yellowish. Gonocoxites with a distinctive projection at posterior margin ventrally bearing long setae; gonostylus bifid, both branches elongate, slender, outer lobe with setae along entire length, inner lobe with only an elongate seta at tip. ***Exechia_alinewongae_ZRCBDP0140725_hapZRCBDP0 140725_SMH_holotype* [31: C, 151: C, 193: G, 199: T, 2 32: G, 235: T, 251: C]**

cctttcttcatcaattgcccacgctggagcCtctgtcgatttagctattttttcattacatttagcaggaatttcctcaattttaggggcagtaaactttattacaacaattattaatatgcgagccccaggtattacttttgatcgactCcctttatttgtatgatctgttttaattactgctgttttactGcttctTtctttaccagtattagctggagctattacaatGctTttaacagatcgaaatCtaaatacctcattttttgatcctgctggaggaggagatccaatcttataccaacatttattt

#### Description

**Male**. Wing length, 1.70; width, 0.67. **Head**. Vertex dark brown, face and clypeus light brown. Vertex with four small bristles around eye posteriorly to ocellus. Antenna light brownish-yellow; flagellomeres short, flagellomere 1 1.6× flagellomere 2 length, flagellomere 4 length 1.9× width (Fig. 96A). Palpus yellowish-brown. **Thorax**. Scutum ochre-yellowish, light brown towards posterior end, scutellum light brown. Pleural sclerites brownish. Scutum with two rows of dorsocentral bristles; scutellum with one pair of marginal bristles and a number of small setae spread over disc. Antepronotum with one strong bristle and many scattered setae; proepisternum with two strong bristles and some few setulae; anepisternum, katepisternum, mesepimeron, and mediotergite bare; laterotergite with six setulae and five long setae medially; metepisternum with nine setulae on posterior half and one stronger seta close to posterior margin. **Legs**. Coxae light whitish-yellow; femora light brownish-yellow, mid femur light brown on distal fifth of wing; tibiae and tarsi light brown. Inner tibial spurs about 3× tibia width at apex. **Wing** (Fig. 96B). Membrane light fumose, unmarked. R_5_ slightly sinuous on basal third; M_1+2_ half of r-m length, medial fork with a constriction on distal third of wing; M_4_ originating clearly beyond level of origin of Rs, posterior fork relatively short; anal fold weakly sclerotized. **Abdomen**. Tergites 1–2 brown medially, yellowish laterally, tergite 3 yellowish, tergites 4–6 blackish-brown medially; tergite 7 brownish-yellow. Sternites 1–6 light yellowish-brown. **Terminalia** (Figs. 96C–F). Light yellowish-brown. Gonocoxites large, close to each other on anterior third of terminalia but not fused, no medial syngonocoxite projection, each gonocoxite with a slender posterior extension on ventral face of terminalia reaching much beyond base of gonostylus, bearing four strong setae at tip; lateral borders extending dorsally, not projected beyond insertion of gonostylus. Gonostyli nearly bare, slightly asymmetric, each with three main lobes: (1) a ventral lobe blade-like, weakly sclerotized, with a sub-basal seta and subapically with three setulae and one longer seta; (2) left gonostylus with distal lobe bifid from nearly its base, dorsal branch digitiform with a single long seta at tip, ventral branch laminar, extending way beyond posterior extension of gonocoxite, falciform at apex, with a pair of subapical setulae, right gonostylus with two branches similarly to left gonostylus; and (3) an additional digitiform, more dorsal branch bearing setulae along its entire length and a distal seta, and a slender, digitiform basal lobe with a long seta at tip. Gonocoxal bridge wide, no apparent apodeme. Aedeagal-parameral complex with a pair of short lateral apodemes, a pair of pointed blade-like lateral arms extending beyond level of base of gonostylus and a medial plate ending distally on a fringe. Tergite 9 with a pair of lateral lobes connected medially at anterior end of terminalia, covered with microtrichia and one distal long seta, a pair of slightly smaller setae and some scattered small setae apically on each lobe. A pair of additional elongate lobes covered with microtrichia, setulae and some slightly longer setae at distal end.

**Female**. Unknown.

#### Material examined

**Holotype**: male, ZRCBDP0140725, Singapore, date range 2012-2018, MIP leg. (slide-mounted).

**Etymology**. Born in Singapore, Aline Wong (1941-) is a sociologist and has been Member of Parliament, as Minister of Education. She promoted early childhood, special needs, arts education programs, and introduced sex education to the national curriculum.

### Mycetophilini

The high species-diversity of Mycetophilini was one of the surprises of this study. This tribe has 13 valid genera, with almost 1,200 described species worldwide. The known diversity of *Mycetophila*, *Trichonta* Winnertz, *Phronia* Winnertz and *Epicypta* worldwide all exceed 100 described species. *Mycetophila* is especially diverse in temperate areas, while *Epicypta*, with about 120 described species, is species-rich in tropical areas of the world.

The samples from Singapore have only three species of *Mycetophila* but 30 species of *Epicypta*. Our samples as well include the first Oriental records for *Aspidionia*— originally described from Micronesia (Colless 1966) with an Afrotropical species added by Matile (1974b) from the Comores Island—and for *Platyprosthiogyne*, so far known from the Seychelles Islands, the Comores Islands and from continental Africa. Two species of *Platurocypta* Enderlein are described here from Singapore—a genus also known

of 20 species from South America, Europe, New Zealand, Africa and the Oriental region. Finally, a new genus of Mycetophilini with close affinities to *Aspidionia* is described here, with five species. The presence of five species of *Platyprosthiogyne*, three species of *Aspidionia*, two species of *Platurocypta*, and five species of the new genus connected to *Aspidionia* was indeed unexpected for Singapore.

### Mycetophila Meigen

*Mycetophila* Meigen, 1803: 263. Type-species, *Tipula agarici* Villers (Johannsen 1909: 116).

**Diagnosis**. Lateral ocelli touching eye margins, third palpomere not swollen. Border of scutum at anterior third only slightly curved; katepisternum at most slightly compressed; anepisternum with strong bristles at dorsal margin; mesepimeron with bristles; laterotergite haired, well developed. Wing membrane with microtrichia arranged in more or less regular longitudinal lines; Sc ending free or in bR; R_4_ absent; M_4_ present, complete at basal end, slightly divergent from M_2,_ parallel with or convergent towards CuA on distal half, tip of M_4_ closer to tip of CuA than to tip of M_2_.

*Mycetophila* has over 650 described species worldwide, most of which known from temperate areas in South America, Australia, New Zealand, and the Holarctic regions (Jürgenstein et al. 2015). There are some clades of the genus in tropical areas, but they are poor in number of species and not abundant.

A total of 40 species of *Mycetophila* have been described for the Oriental region. Of this total, 30 are from China, three are from Nepal, three from India, and four are from Southeast Asia—*M. borneana* Edwards, from Mt. Kinabalu, Malaysia (Edwards 1933), *M. lineicoxa* Edwards and *M. trimacula* Edwards from the mountains in Gunong Tahan, also Malaysia (Edwards 1928), and *M. reversa* Edwards, from Bukittinggi [= “Fort de Kock”], Sumatra, Indonesia (Edwards 1931). We found three species of *Mycetophila* in our Singapore samples, which might belong together to a small clade in the genus, different from the larger, showy groups in temperate areas in the southern hemisphere or in the northern hemisphere. All three species of *Mycetophila* already described from Southeast Asia are brownish—as *Mycetophila aishae* Amorim & Oliveira, **sp. n.** —, but they differ from the Singapore species in other details, including the male terminalia in the case of *M. trimacula*.

The haplotype network for the genus (Fig. 98D) does not depict any conflicts between different approaches, clearly recognizing three species. Most specimens of *Mycetophila* in our samples come from the swamp forest and the rain forest, but *M. aishae* Amorim & Oliveira, **sp. n.** was collected only in one of the urban forest.

*Mycetophila chngseoktinae* Amorim & Oliveira, sp. n. (Figs. 97A–E, 98A–C)

https://singapore.biodiversity.online/species/A-Arth-Hexa-Diptera-002129

urn:lsid:zoobank.org:act:8808A46C-BFE0-4FC2-9BC5-88F98C7EB385

**Diagnosis**. Head ochre-yellowish. Scutum with an ochre-yellowish background, a long light brown longitudinal band medially and a pair of shorter light brown bands more laterally; pleural sclerites basically ochre-yellowish, with a antepronotum, dorsal and dorsoposterior margins of anepisternum, dorsal end of mesepimeron, posterior margin of laterotergite and metepisternum brownish. C very short beyond tip of R_5_; bR devoid of setation at distal end. Tergite 1 brown, tergites 2–6 light yellowish with a medial brownish area of variable size between tergites, tergite 7 brownish-yellow. Male terminalia gonostylus with large posterior lobe, basal lobe short, not projected inwards.

***Mycetophila_chngseoktinae_ZRCBDP0048472_hapZRC BDP0047896_SMH_holotype* [1: A, 12: G, 13: A, 49: C, 139: C, 175: C, 251: C, 292: T]**

ActttcatctaGAattgcccatgctggagcatctgtagatttagcaatCttttctcttcatttagcaggtatttcttctattttaggagctattaattttattacaacaattattaatatacgagccccaggaataacCtttgatcgaatacctttatttgtatgagcagtactCattacagctattcttttattactttccttacctgttttagcaggggctattactatactactaactgatcgtaacCtaaatacttctttttttgaccctgctggaggaggtgacccTattctatatcaacatttattt

#### Description

**Male**. Wing length, 1.95; width, 0.83. **Head** (Fig. 97B). Ochre-yellowish, face greyish-yellow. Mid ocellus absent, lateral ocelli nearly touching eyes dark brown. Three bristles along margin of eye posteriorly to lateral ocellus, a row of short bristles along anterior margin of frons, scattered small setae covering vertex. Antennal scape and pedicel greyish-yellow, flagellomere 1 greyish-yellow on basal half, light brown on distal half, flagellomeres light brown, distal two flagellomeres lighter. Scape long, about twice longer than pedicel. Scape with setae on inner face and two dorsal small bristles. Pedicel with setation on lateral face and around posterior margin, in addition to a long dorsal seta. Flagellomeres almost twice as long as wide, except for flagellomere 1, about 3× longer than wide. Face short, light greyish-brown, clypeus slightly bulging, light brown, setose. Maxillary palpus light brown, labella brownish-yellow. Maxillary palpus light brown, lighter towards apex, with five palpomeres; palpomere 1 only with microtrichia, palpomere 2 short, with some setae along posterior margin dorsally, palpomere 3 with a conspicuous sensorial pit opening at inner face of basal half, and setae dorsally at inner face, palpomere 4 about as long as palpomere 3, with small setae dorsally at external face, palpomere 5 1.5× palpomere 4 length. Labella large, light brownish-yellow. **Thorax** (Fig. 97B). Scutum ochre-yellow, with a medial ochre-brown band on anterior half and a pair of light brown maculae on posterior half that merge together posteriorly and continues into scutellum. Scutum densely covered by short brownish small setae and a row of supra-alars, one marginal prescutellar bristle and one median prescutellar bristle. Scutellum large, with two pairs of strong bristles on posterior margin and additional small setae on posterior half. Antepronotum and proepisternum light brown, prosternum and proepimeron brown; anepisternum whitish ochre, light brown along anterior and dorsal margins, and brown along posterior margins; katepisternum and mesepimeron brownish-yellow; metepisternum light brown; laterotergite ochre-yellow, with ochre-brown posterior margin, mediotergite light brown with ochre-yellow laterals. Antepronotum with three bristles and additional larger and smaller setae; proepisternum with three bristles and additional small setae along dorsal margin; anepisternum with over 40 setae on dorsal ¾, and four bristles close to posterior margin; mesepimeron with three bristles along dorsal margin and some additional small setae; katepisternum and mediotergite bare; laterotergite with a group of three longer and six smaller setae; 5–6 metepisternal setae. Haltere light brown, setulae restricted to knob. **Legs**. Fore coxa light ochre-yellowish, mid and hind coxae whitish; fore femur light brownish-yellow, mid and hind femora whitish-yellow, yellowish-brown along dorsal edge; tibiae and tarsi light yellowish-brown. Hind femur considerably wide medially. Fore coxa densely covered with setae anteriorly and on internal face, some longer black setae anteriorly and close to tip; mid coxa with some fine small setae and some longer setae on distal fifth of wing; hind coxa with fine setae on basal 4/5 only on posterior face, a distinctive black subapical seta laterally, some few longer setae along distal end anteriorly. Femora densely covered with fine setae, mid femur with some longer setae only close to tip ventrally; hind femur with one long black seta ventrally midway to apex. Tibiae and tarsi with regular rows of trichia. Fore tibia with few stronger setae only on distal fifth. Mid tibia with five dorsal bristles, one bristle on inner face of basal half and and two on inner face on distal half, one stronger bristle on outer face of distal half; hind tibia with two rows of six posterodorsal bristles, area between rows flat and bare. Anteroapical depressed area on inner face of fore tibia wide, densely lined with setulae. Inner spur of hind tibia about 3× tibia width at apex. Fore leg tarsomeres with distal setae, only one ventral seta midway to apex; mid and hind tarsomeres 1–4 with two rows of ventrolateral setae in additional to those at apex. Tarsal claw with one proximal short tooth and one subproximal long tooth. **Wing** (Fig. 97C). Membrane light fumose, yellowish along anterior margin, no brown marks. Wing membrane microtrichia in regular rows, bare of macrotrichia except for four setulae dorsally on anal lobe posteriorly to anal fold. Sc short, ending free. C extending only shortly beyond tip of R_5_. R_1_ long, almost straight, reaching C at distal fourth of wing. First sector of Rs oblique, 1.4× r-m length; R_5_ gently curved along its entire length, reaching C beyond level of tip of M_2_. Vein r-m short, oblique, less than half extension of M_1+2_. M_1+2_ 3.3× r-m length. Medial fork long, M_1_ and M_2_ straight, only slightly divergent along their course. M_4_ very gently arched towards posterior margin, posterior fork slender, CuA straight. CuP not produced; anal fold long, arched. Dorsal setae present on Sc, bR, R_1_, second sector of Rs, and r-m; ventral setae present on distal half of bR, R_1_, Rs, and r-m. Wing posterior margin slightly emarginated at apex of CuA. **Abdomen**. Tergite 1 brown, tergites 2–5 light ochre-yellowish, with a medial brownish area, tergite 6–7 dark ochre-yellowish. Sternites 1–7 light yellowish-brown. **Terminalia** (Figs. 98A–B). Yellowish-brown. Gonocoxites fused along their entire length ventrally, no medioventral process, posterior margin of syngonocoxite ventrally with a pair of arms extending laterally towards base of gonostylus, no projection of gonocoxite distal border beyond base of gonostylus; gonocoxite lateral margin extended dorsally, partially encapsulating terminalia. Gonostylus large, composed of: (1) a large ventral lobe with fine setae and a short digitiform posterior extension with apical setulae; (2) a small digitiform ventral lobe with a subapical seta; (3) a large, elongate and setose dorsal lobe, projecting much beyond rest of terminalia; (4) a median lobe, with a pair of long setae directed inwards and two other smaller setae. Gonocoxal bridge large. Aedeagal-parameral complex with a pair of elongate parameral blades and a medial subquadrate aedeagal sclerite more distally. A pair of large ovoid lobes dorsally representing tergite 9+10, cercus weakly sclerotized.

**Female** (Fig. 97A). As male, except for the following. **Wing**. Membrane slightly darker. **Terminalia** (Figs. 97D– E). Sternite 8 elongate, no lobes produced, setae closer to posterior margin longer, reaching level of posterior ⅔ of cercomere 1. Sternite 9 elongate, weakly sclerotized, gonopore at level of posterior fourth of sternite 8, anterior end wide, reaching posterior third of segment 7. Tergite 8 wide, reaching level of mid of sternite 8. Tergite 9+10 short, with a short medial incision of posterior margin. Cercomeres elongate, setose, cercomere 1 2.8× length of cercomere 2.

#### Material examined

**Holotype**: male, ZRCBDP0048472, Nee Soon (NS1), swamp forest, 25.apr.2012, MIP leg. (slide-mounted). **Paratypes** (15 males, 6 females). **Males**: ZRCBDP0048719, Nee Soon (NS2), 28.jan. 2015, MIP leg.; ZRCBDP0048837, Nee Soon (NS1), jan. 2015, MIP leg.; ZRCBDP0048967, Nee Soon (NS1), apr. 2015, MIP leg.; ZRCBDP0048992, Nee Soon (NS2), 17.dec. 2014, MIP leg.; ZRCBDP0049001, Nee Soon (NS2), 17.dec. 2014, MIP leg.; ZRCBDP0049012, Nee Soon (NS2), 17.dec. 2014, MIP leg.; ZRCBDP0049014, Nee Soon (NS2), 17.dec. 2014, MIP leg.; ZRCBDP0049015, Nee Soon (NS2),17.dec. 2014, MIP leg.; ZRCBDP0049016, Nee Soon (NS2), 17.dec. 2014, MIP leg.; ZRCBDP0049017, Nee Soon (NS2), 17.dec. 2014, MIP leg.; ZRCBDP0049188, Nee Soon (NS2), 13.may. 2015, MIP leg.; ZRCBDP0049191, Nee Soon (NS2), 13.may. 2015, MIP leg. (slide-mounted) (MZUSP); ZRCBDP0049223, Nee Soon (NS1), 10.dec. 2014, MIP leg.; ZRCBDP0047921, Nee Soon (NS2), swamp forest, 09.oct. 2013, MIP leg.; ZRCBDP0154876, Singapore, date range 2012-2018, MIP leg. **Females**: ZRCBDP0047896, Nee Soon (NS2), swamp forest, 17.apr. 2013, MIP leg.; ZRCBDP0048447, Nee Soon (NS1), swamp forest, 04.apr. 12, MIP leg. (website photo specimen); ZRCBDP0048448, Nee Soon (NS1), swamp forest, 23.may. 12, MIP leg.; ZRCBDP0048674, Nee Soon (NS1), swamp forest, 11.apr. 2012, MIP leg. (slide-mounted); ZRCBDP0048703, Nee Soon (NS2), 28.jan. 2015, MIP leg.; ZRCBDP0049106, Nee Soon (NS1), 24.dec. 2014, MIP leg.

**Etymology**. This species epithet honors Chng Seok Tin, a visually impaired sculptor and artist, whose work was often inspired by the i-Ching and Buddhism. Her work has been shown internationally and she became the first Singaporean artist to exhibit her works at the Headquarters of the United Nations— Chng had over 26 solo shows and 100 group shows. She is a strong advocate for artists with disabilities and is a recipient of the Singapore Cultural Medallion. She was inducted into the Singapore Women’s Hall of Fame in 2014.

**Remarks**. We have two haplotypes and no delimitation conflicts.

*Mycetophila georgettechenae* Amorim & Oliveira, sp. n. (Fig. 99A–F)

https://singapore.biodiversity.online/species/A-Arth-Hexa-Diptera-002119-000803

urn:lsid:zoobank.org:act:22BC266B-23C1-48CA-8299-A1FAE38044BA

**Diagnosis**. Head dark ochre-yellowish. Scutum with an ochre-yellowish background and a long light brown longitudinal band medially and a pair of shorter light brown bands more laterally; pleural sclerites basically ochre-yellowish, antepronotum, dorsal and dorsoposterior margins of anepisternum, dorsal end of mesepimeron, posterior margin of laterotergite, and metepisternum brownish. C slightly projected beyond tip of R_5_; bR with some few dorsal setae at distal end. Tergite 1 brown, tergites 2–4 and 6 light yellowish with a medial brown longitudinal band, tergite 5 light brown, tergite 7 brownish-yellow. Male gonostylus with posterior lobe not too wide and a basal lobe with a short, triangular projection inwards.

***Mycetophila_georgettechenae_ZRCBDP0048061_hapZR CBDP0048061_SMH_holotype* [1: G, 4: G, 12: G, 34: C, 100: C, 167: G, 175: T]**

GctGtcatctaGtattgcccatgctggagcatcCgtagatttagcaattttttctcttcatttagcaggtatttcttcaattttaggagcaattaatttCattacaacaatcattaatatacgagcccctggaataacttttgatcgaatacccttatttgtttgaGcagttctTattacagctatcttactattactttcattacctgttctagcaggagctattactatactattaactgaccgtaatttaaatacttctttttttgacccagcaggaggaggggaccctatcttatatcaacatttattt

#### Description

**Male** (Fig. 99A). Wing length, 1.95; width, 0.77. **Head** (Fig. 99B). Ochre-yellowish, face greyish-yellow. Ocelli dark brown, lateral ocelli nearly touching eye margins. Antennal scape greyish-yellow, pedicel greyish-brown, flagellomere 1 whitish, flagellomeres light brown, lighter towards apex, flagellomeres more than twice longer than wide; scape and pedicel with a crown of setae around distal margin, but no dark, stronger setae. Maxillary palpus light brown, labella brownish-yellow. **Thorax** (Fig. 99C). Scutum ochre-yellow, with a medial ochre-brown band on anterior half and a pair of light brown maculae on posterior half. Scutellum light brown. Antepronotum ochre-brown, proepisternum brownish-yellow, prosternum light brown; proepimeron brownish-yellow; anepisternum ochre-yellow, light brown along anterior, dorsal and posterior margins, katepisternum and mesepimeron brownish-yellow; metepisternum light brown; laterotergite ochre-yellow, with ochre-brown posterior margin, mediotergite light brown with ochre-yellow laterals. **Legs**. Fore coxa light ochre-yellowish, mid and hind coxae whitish; fore femur light brownish-yellow, mid and hind femora whitish-yellow, mid femur with some yellowish tinge on basal half, hind femur brown on distal fourth along dorsal edge and a dark brown mark ventrally at tip; tibiae and tarsi light yellowish-brown. **Wing** (Fig. 99D). Membrane light brownish fumose. Microtrichia in regular rows, no macrotrichia on membrane. Sc short, ending free. C extending only shortly beyond tip of R_5_. Vein bR curved; R_1_ long, gently curved along its length, reaching C at distal fourth of wing. First sector of Rs oblique, 2.3× r-m length; R_5_ gently curved along its entire length, reaching C almost at level of tip of M_2_. Vein r-m very short, oblique, less than half extension of M_1+2_. M_1+2_ 3.2× r-m length. Medial fork long, M_1_ and M_2_ straight, only slightly divergent along their course. M_4_ gently arched towards posterior margin, posterior fork slender. Cubital pseudovein produced, short; CuP absent; anal fold long, arched. Dorsal setae present on bR, R_1_, second sector of Rs, r-m and distally on bM; ventral setae present on distal half of bR, R_1_, Rs and r-m. No macrotrichia on wing membrane. Wing posterior margin gently emarginated at apex of CuA. **Abdomen**. Tergite 1 brown, tergites 2–4 and 6 light yellowish with a medial brown longitudinal band, tergite 5 light brown, tergite 7 brownish-yellow. Sternites 1–6 light yellowish-brown. **Terminalia** (Figs. 99E–F). Yellowish-brown. Gonocoxites fused along their entire length ventrally, no suture of fusion, medioventrally syngonocoxite gently curved posteriorly, no ventral projection of gonocoxite posterior border beyond base of gonostylus, dorsally a short lobe extending slightly beyond base of gonostylus; gonocoxite lateral margins extended dorsally, partially encapsulating terminalia. Gonostylus weakly sclerotized, composed of: (1) a posterior lobe extending distally, weakly sclerotized, more or less digitiform, with some setae; (2) a basal short, slightly more sclerotized, subtriangular lobe projecting inwards; (3) a medial short lobe projecting inwards. Gonocoxal bridge wide. Aedeagal-parameral complex with a pair of elongate parameral blades and a medial subquadrate aedeagal sclerite more distally. A pair of elongate ovoid lobes dorsally, representing tergite 9+10, cerci weakly sclerotized, placed more distally.

**Female**. As male, except for the following. Wing length, 1.92; width, 0.70. **Terminalia**. Sternite 8 elongate, rounded distally, no lobes or medial incision, setae along posterior margin longer, distal margin of sternite 8 at level of distal half of cercomere 1. Sternite 9 elongate, weakly sclerotized, anterior end wide, reaching posterior third of segment 7. Tergite 8 wide, reaching level of mid of sternite 8. Tergite 9+10 short, with a short medial incision of posterior margin. Cercomeres elongate, setose, cercomere 1 2.5× length of cercomere 2.

#### Material examined

**Holotype**: male, ZRCBDP0048061, Nee Soon (NS2), swamp forest, 27.mar.13, MIP leg. (website photo specimen, slide-mounted). **Paratypes** (2 males, 1 female). **Males**: ZRCBDP0048745, Nee Soon (NS1), 25.feb.15, MIP leg.; ZRCBDP0154916, Singapore, date range 2012-2018, MIP leg.

**Female**: ZRCBDP0048450, Nee Soon (NS1), swamp forest, 11.apr. 12, MIP leg. (slide-mounted).

**Etymology**. The species epithet of this species honors Georgette Liying Chendana CHEN (1906-1993), an acclaimed first-generation Singaporean oil-painter. A key figure in the development modern art in Singapore, Chen is known not only for her oil paintings, but also for her contributions to art education as a teacher at the Nanyang Academy of Fine Arts (NAFA) from 1954 to 1981. She was one of a group of artists who established the Nanyang Style of painting, which combines Western technique with Asian themes. She was inducted into the Singapore Women’s Hall of Fame in 2014.

**Remarks**. We have two haplotypes and no delimitation conflicts.

*Mycetophila aishae* Amorim & Oliveira, sp. n. (Figs. 100A–F)

https://singapore.biodiversity.online/species/A-Arth-Hexa-Diptera-002078

urn:lsid:zoobank.org:act:94E54673-B4AF-4759-896C-61DD6A32927C

**Diagnosis**. Head light brown. Scutum mostly light brown, with dark ochre-yellowish marks on shoulders, anteriorly over scutum; pleural sclerites light brown. C not projected beyond tip of R_5_; bR entirely bare. Abdominal tergites entirely brown, lighter at lateral ends. Male terminalia gonostylus with posterior lobe projected, but quite slender distally, basal lobe wide, with a short, sclerotized beak midway to apex projecting inwards.

***Mycetophila_aishae_ZRCBDP0279129_holotype* [31: C, 44: A, 49: C, 118: C, 160: C, 187: C, 190: G, 220: T] -**

ctttcatcaactattgcccatgcaggggcCtcagtagatttaAcaatCttttctcttcatttagctggaatttcatcaattttaggagctatcaactttattacaactattattaaCatacgtgcccctggtataacttttgatcgattaccattattCgtttgatcagttttaatcacagccatCctGcttttactatcattacctgtcctagcaggTgctatcacaatacttttaacagaccgaaatttaaacacaacattttttgaccccgcaggaggaggagaccctattttataccaacatttattt

#### Description

**Male** (Fig. 100A). Wing length, 2.08; width, 0.77. **Head** (Fig. 100B). Brown, face brownish-yellow. Lateral ocelli nearly touching eyes. Antennal scape and pedicel greyish-yellow, flagellum light brown, scape 1.4× longer than pedicel, flagellomere 1 1.2× longer than flagellomere 2, flagellomere 2.3× longer than wide; scape and pedicel with a crown of setae around distal margin, pedicel with a stronger seta dorsally. Maxillary palpus light brown, palpomere 3 with a well-developed sensory pit, palpomere 4 1.1× length of palpomere 3, palpomere 5 1.9× palpomere 4 length. Labella brownish-yellow. **Thorax** (Fig. 100B). Scutum light brown, with brownish-yellow anterior corners, scutellum light brown. Some large supra-alar setae, two pairs of prescutellar bristles, scutellum with two pairs of marginal bristles and a number of small additional setae. Antepronotum, proepisternum, prosternum, katepisternum, and mesepimeron ochre-brown, anepisternum, metepisternum, laterotergite, and mediotergite light brown. **Legs**. Coxa ochre-yellowish, fore coxa slightly more brownish; femora light brownish-yellow, hind femur enlarged and flattened, with a brown mark apically on dorsal and ventral margins; tibiae and tarsi light yellowish-brown. **Wing** (Fig. 100C). Membrane light fumose brown. Wing membrane with microtrichia in regular rows. Sc ending free, barely visible. C not extending beyond tip of R_5_. R_1_ long, gently curved, reaching C at distal fourth of wing. Vein bR curved, first sector of Rs oblique, 1.6× r-m length, R_5_ gently curved, reaching C slightly beyond level of tip of M_2_. Vein r-m short, oblique. M_1+2_ short, 2.6× longer than first sector of Rs. Medial fork long, M_1_ and M_2_ slightly convergent at distal end. M_4_ gently arched towards posterior margin, posterior fork slender. CuA straight. Cubital pseudovein short, sclerotized to distal fourth of first sector of CuA, CuP not produced. Anal fold long, arched distally. Dorsal setae present on entire length of bR, R_1_, second sector of Rs, and r-m, posterior veins bare; ventral setae present on distal half of bR, R_1_, second sector of Rs, and r-m. Wing posterior margin gently emarginated at apex of CuA. **Abdomen**. Tergites 1–6 brown, slightly darker towards apex, sternites 1–6 light brown, sternites 5–6 darker; tergites and sternites 7–8 yellowish-brown. **Terminalia** (Figs. 100D–F). Small, yellowish-brown. Gonocoxites fused along their entire length ventrally, no medioventral posterior process. Gonostylus with an elongate posterior lobe with fine setae and a medial wide lobe with a short, sclerotized beak midway to apex. Aedeagal-parameral complex rectangular. Tergite 9+10 entirely divided into a pair of ovoid lobes dorsally.

**Female**. Unknown.

#### Material examined

**Holotype**: male, ZRCBDP0279129, Singapore, 31.may.18, MIP leg. (slide-mounted). **Paratype** (2 males): ZRCBDP0284233, Singapore, PU13, (date range 2012-2018), MIP leg; ZRCBDP0133534, National University of Singapore (PGP), 03.may.17.

**Etymology**. The species epithet honors Aisha Akbar (1930–2015), Malay music teacher, songwriter, author and broadcaster. A Singapore Women’s Hall of Fame inductee, she was concerned that traditional Malay folk songs would get lost in the wash of time and spent years researching and documenting important local songs such as ‘Rasa Sayang’ and ‘Dayung Sampan’, ensuring that Singapore’s musical heritage would be preserved.

### Platyprosthiogyne Enderlein

*Platyprosthiogyne* Enderlein 1910: 78. Type-species: *Platyprosthiogyne metameromelina* Enderlein 1910 (orig. design.).

**Diagnosis**. Small flies, wing length between 1.6 and 2.4 mm. Lateral ocelli touching eye margins, mid ocellus usually present (absent in *Platyprosthiogyne gohsookhimae* Amorim & Oliveira, **sp. n.**). Scutum with no incision at margin above level of antepronotum; anepisternum with scattered setae; mesepimeron and laterotergite setose. Wing membrane with microtrichia arranged in more or less regular longitudinal lines; Sc short, ending free; R_4_ absent; M_4_ absent, CuA running largely parallel to M_2_; anal fold well sclerotized, long, curved.

*Platyprosthiogyne* was erected by Enderlein (1910) for a species from the Seychelles Islands. It has been an obscure genus for a long while, until Matile (1974b, 1979a) described two Afrotropical species, one from Cameroun and one from the Comores Islands (see also Søli 2017). It was quite a surprise to find six additional species in Singapore, the first non-Afrotropical record of the genus.

*Platyprosthiogyne is* certainly not a genus sister to *Epicypta*, within the evolution of the tribe, and it may be closer to *Mycetophila*. All species described here from Singapore share the diagnostic features of *Platyprosthiogyne*. This includes the loss of M_4_ (also originated elsewhere in mycetophilines), the scutum border with no keel above antepronotum (a plesiomorphic condition), and an incision on the posterior wing margin at the tip of CuA (in some degree present in other genera). Most species also show a typically arched R_5_ running close to R_1_ and an extension of C beyond the tip of R_5_.

*Platyprosthiogyne snehalethaae* Amorim & Oliveira, **sp. n.** is relatively plesiomorphic for some of these features (especially the separation between R_5_ and R_1_, and the extension of C beyond the tip of R_5_). In the type-species of the genus (*P. metameromelina* Enderlein), the medial fork opens rather closer to the margin, as we see in *P. neilaae* Amorim & Oliveira, **sp. n.** and *P. snehalethaae*

Amorim & Oliveira, **sp. n.** Some strong setae along the posterior margin of the syngonocoxite are seen in *P. gohsookhimae* Amorim & Oliveira, **sp. n.** and *P. neilaae* Amorim & Oliveira, **sp. n.,** also present in *P. oresbia* Matile. Notwithstanding the position of *P. snehalethaae* Amorim & Oliveira, **sp. n.,** there are two consistent small clades in the genus: one including *P. gohsookhimae* Amorim & Oliveira, **sp. n.** and *P. neilaae* Amorim & Oliveira, **sp. n.,** and the other with *P. rahimahae* Amorim & Oliveira, **sp. n.,** *P. phanwaithongae* Amorim & Oliveira, **sp. n.** and *P. lynetteseahae* Amorim & Oliveira, **sp. n.**

There are three conflicts regarding the haplotype network for *Platyprosthiogyne* (Fig. 101). In one case, there are two very distinct species— *P. gohsookhimae* Amorim & Oliveira, **sp. n.** and *P. neilaae* Amorim & Oliveira, **sp. n.** —brought together by mPTP. *P. gohsookhimae* Amorim & Oliveira, **sp. n.** is split, on the other hand, in two separate species by ABGD, a divergence that may be in the grey-zone. Finally, *P. rahimahae* Amorim & Oliveira, **sp. n.,** *P. lynetteseahae* Amorim & Oliveira, **sp. n.** are brought together by mPTP, but there is molecular and morphological evidence that they are clearly separate species.

*Platyprosthiogyne phanwaithongae* Amorim & Oliveira, sp. n. (Figs. 102A–F)

https://singapore.biodiversity.online/species/A-Arth-Hexa-Diptera-000799

urn:lsid:zoobank.org:act:03FB165B-AF7A-4CDF-8268-71BFC6775CE0

**Diagnosis**. Vertex brown, ochre-yellow around eye and on face. Scutum dark brown, with ochre-yellow anterior fifth. Antepronotum ochre-yellow, proepisternum ochre-yellow with a brownish area anteriorly, other pleural sclerites brown. Legs mostly cream-whitish, hind coxa with brownish transverse band on basal end, femora entirely whitish. Wing with a darker macula over area around distal half of cell br; Sc nearly absent, not fused to C or to bR; C produced shortly beyond tip of R_5_; first section of Rs strictly transverse, R_5_ running very close to R_1_ and C; M_1+2_ slightly shorter than r-m; M_4_ absent, CuA running parallel to M_2_; anal fold long, well sclerotized, arched on distal third of wing; wing posterior margin with a gentle incision at level of tip of CuA. Abdominal tergites 1–5 brown, tergite 6 brown with a yellow band along posterior margin. Male syngonocoxite with no strong setae along posterior margin ventrally; gonostylus small, subquadrate, with a short digitiform basal projection directed inwards; paramere well sclerotized, present as an elongate, rectangular sclerite; tergite 9 at anterior half of terminalia wide, with rounded lateroposterior corners, setose and with microtrichia; cerci large.

***Platyprosthiogyne_phanwaithongae_ZRCBDP0048140_ hapZRCBDP0048140_SMH_holotype* [46: C, 103: C, 12 8: A, 133: G, 148: C, 151: G, 154: A, 212: C]**

actatcttcaactattgctcatgctggagcttcagttgatttagcCattttttctcttcatttagctggaatttcctcaattttaggagcaattaattttatCacaacaattattaatatacgagctAcaggGattacatttgatcgCatGccAttatttgtatggtctgtttttattacagctattttattacttttatcattaccagttCtagcaggagctattactatattattaacagaccgaaatttaaatacttcttttttcgaccctgctggaggaggagatcctattttataccaacatttattt

#### Description

**Male**. Wing length, 1.82; width, 0.70. **Head** (Fig. 102B). Brown medially on frons, brownish-yellow around eye, partially fit under anterior end of scutum. Face and clypeus dirty-yellowish. Lateral ocelli blackish-brown, nearly touching eye margin, no median ocellus, frontal furrow short. Antennal scape and pedicel light brown, flagellum brown. Maxillary palpus light brown, lighter toward apex. Labella light brownish-yellow. Dark brown small setae scattered over vertex, four slightly longer setae on occiput around dorsal margin of eye posteriorly to line of ocelli, a row of long setae close to anterior margin of frons. Antennal scape twice pedicel length, a crown of setae distally and on inner face of scape; setulae on both lateral faces and on distal margin of pedicel, in addition to one strong seta dorsally. Face slender, with a transverse line of setulae. Clypeus with scattered setulae. Mid ocellus present, at posterior end of frontal furrow. Flagellomere 1 twice flagellomere 2 length; flagellomere 2 1.7× longer then wide. Palpomere 1 twice as long as wide, covered only with microtrichia, palpomere 2 very short, with some setulae, palpomere 3 slightly longer than wide, sensorial pit conspicuous opening dorsally, setulae on external and dorsal faces, palpomere 4 almost twice flagellomere 3 length, with setulae on external and dorsal faces, palpomere 5 slender, about twice flagellomere 4 length, with scattered setulae. **Thorax** (Fig. 102C). Scutum shinning dark brown except for ochre-yellow anterior sixth, scutellum dark brown. Basisternum brown. Antepronotum ochre-yellow, proepisternum ochre-brown with ochre-yellow areas on distal half, anepisternum shining brown, katepisternum, mesepimeron, laterotergite, and metepisternum ochre-brown, mediotergite dark brown. Scutum densely covered with scattered fine setae, no shiny median keel anteriorly, five longer setae at small bulging area on margin behind level of wing, four pairs of prescutellar bristles along posterior margin of scutum; scutellum large, trapezoid, two pairs of strong bristles and some additional smaller setae along posterior margin. Basisternum dorsoposterior arms with some few setulae. Antepronotum only with short fine setae, proepisternum with three bristles on dorsal half and scattered small setae. Anepisternum entirely covered with scattered setae, a line of long setae along posterior margin; katepisternum small, slightly less than half height of anepisternum. Mesepimeron reaching ventral margin of thorax, with two strong bristles along dorsal margin and 11 setulae. Laterotergite bulging, quite flattened, with three long setae along posterior margin and nine scattered small setae; mediotergite small, inclined, bare; metepisternum with 12 fine setae along its length. **Legs**. Coxae whitish, mid coxa with orangish antero-basal corner, hind coxa with a light brown transverse band at basal end; femora whitish-yellow; tibiae light yellowish-brown with an orangish tinge, tarsi very light ochre-brown. Fore coxa entirely covered with setulae at anterior face, a row of brown bristles along posterior margin and around tip, and along margin on basal half of internal face; mid coxa largely developed, setation restricted to frontal face near distal end; hind coxa with some few small setae distally and one strong seta at external face on distal two-thirds. Femora covered with fine setae, a row 3–5 longer setae along ventral margin distally, stronger on hind femur. Tibiae and tarsi with regular rows of trichia. Fore tibia with a wide anteroapical depressed area lined with setulae, a pair of rows densely covered with setae dorsolaterally, in additional to setae at distal end. Mid and hind tibiae with two rows of setae dorsally and some strong setae on lateral faces, in additional to long setae at distal end. Fore leg tarsomere 1 shorter than tibia, twice tarsomere 2 length. Fore leg tarsomeres only with rows of trichia and a couple of distal setae; mid and hind tarsomeres 1–3 with some ventral setae besides rows of trichia. Tibial spurs light brown, subequal, spur of fore tibia about twice tibia width at apex, spurs of mid leg about 5× tibial apex, spurs of hind leg about 3× tibial apex. Tarsal claws with an inconspicuous basal tooth. **Wing** (Fig. 102D). Wing membrane light brown fumose, slightly darker along anterior margin, dark area extending to origin of M_1+2_. Membrane densely covered with regularly organized microtrichia on all cells, no macrotrichia on membrane; posterior margin gently emarginated at level of tip of CuP. Sc ending free, faint, barely recognizable. R_1_ relatively short, reaching C before distal third of wing; R_4_ absent; R_5_ short, reaching C before level of M_2_, running close to C beyond R_1_. C not extending beyond R_5_. First sector of Rs slightly oblique; r-m almost longitudinal, half M_1+2_ length. Vein bM over 7× r-m length; M_1+2_ short; M_1_ and M_2_ well sclerotized, running more or less parallel along most of their length; M_4_ absent; medial veins weakly sclerotized close to margin. CuA straight, long, reaching margin almost at level of tip of R_5_. Cubital pseudovein and CuP not produced. Anal fold long, gently curved on distal third, not reaching wing margin. Dorsal macrotrichia on bR, R_1_, and on second sector of Rs, ventral macrotrichia on distal fourth of bR, distal three fourth of R_1_, and on Rs. **Abdomen**. Tergite 1 white laterally with a medial brown mark, tergites 2–6 brown; sternite 1 white, sternite 2–6 light brown, darker towards distal segments; tergite and sternite 7 yellowish. **Terminalia** (Fig. 102E). Small, yellowish, weakly sclerotized. Gonocoxites fused together medially, no suture, no incision on posterior margin of syngonocoxite medially, a short medial keel extending medially, posterior margins of gonocoxite not extending beyond base of gonostylus, setae restricted to lateroposterior corners ventrally. Gonostylus complex, composed of: (1) ventral lobe, elongate towards mid of terminalia (medially almost touching tip of opposite gonocoxite ventral lobe), bearing distally a short digitiform projection with a distal seta, a short pointed lobe and a small sublobe with five fine setulae at tip; (2) a median lobe wider at base and with a distal digitiform projection bearing two spines and a fine seta at tip; (3) a dorsal wide digitiform lobe bearing fine setulae along distal half. Gonocoxal bridge present, weakly sclerotized, no apodemes visible. Aedeagal-parameral complex constituted of an elongate, subquadrate sclerite with a pair of rounded short lobes on laterodistal corners and a small distal keel. Tergite 9 trapezoid, tapering towards posterior margin. A pair of long lobes dorsally bearing microtrichia and fine setae, possibly fusion between tergite 10 and cerci (or loss of tergite 10).

**Female** (Fig. 102A). As males, except for the following. **Wing l**ength, 1.95; width, 0.83. Mesepimeron with one strong seta and eight smaller setae. Laterotergite with three longer setae and seven smaller ones; metepisternum with 17 small setae. **Legs**. Tarsomeres 1–2 of fore leg with a row of spines along entire length at external. **Abdomen**. Tergite 1 white, tergites 2–5 brown, tergite 6 brown with a cream-yellow transverse band at distal fourth; sternite 1 white, sternite 2–7 light brown, darker towards distal segments, sternite 7. **Terminalia** (Fig. 102F). Yellowish. Sternite 8 elongate, extending distally to reach distal third of cercomere 1, with a posterior incision between then. Sternite 9 with wide anterior end, distal medial end of sternite 9 reaching level of tip of lobes. Tergite 9 short, weakly sclerotized, with microtrichia and a row of setulae on posterior half, laterally extending towards ventral face of terminalia, partially overlapping sternite 8 anteriorly. Tergite 9+10 short, slightly more sclerotized, with microtrichia, but no setae, fused to sternite 9 laterally. Cercomere 1 long, densely covered by microtrichia and fine setae, apparently with a sensorial area on inner face. Cercomere 2 about as long as cercomere 1.

#### Material examined

**Holotype**: male, ZRCBDP0048140, Sungei Buloh (SB1), mangrove, 02.oct.2013, MIP leg. (slide-mounted). **Paratypes** (1 male, 3 females). **Male**: ZRCBDP0048149, Sungei Buloh (SB1), mangrove, 02.oct.13, MIP leg. **Females**: ZRCBDP0048144, Sungei Buloh (SB1), mangrove, 02.oct.13, MIP leg. (slide-mounted); ZRCBDP0048145, Sungei Buloh (SB1), mangrove, 02.oct.13, MIP leg. (website photo specimen); ZRCBDP0048148, Sungei Buloh (SB1), mangrove, 02.oct.13, MIP leg.

**Etymology**. The species epithet of this species honors Phan Wait Hong (1914–2016), called “grande dame of Beijing opera in Singapore”. Shanghai-born, she moved to Singapore at age 14 as part of an opera troupe. She performed regularly until the age of 82, with a limited number of performances for a decade afterwards. She received the Singapore Cultural Medallion for Chinese Opera in 1992 and was inducted to the Singapore Women’s Hall of Fame in 2014.

**Remarks**. There are two haplotypes and no conflicts between different delimitation approaches for this species.

*Platyprosthiogyne gohsookhimae* Amorim & Oliveira, sp. n. (Figs. 103A–I)

https://singapore.biodiversity.online/species/A-Arth-Hexa-Diptera-000806

urn:lsid:zoobank.org:act:B02F3C9B-AEF7-46E6-9DDB-58235FAEB080

**Diagnosis**. Head dark brown. Scutum dark brown, pleural sclerites dark brown, antepronotum slightly lighter. Coxae and femora mostly whitish, mid coxa with basal half brownish, mid coxa with basal ¾ brown, mid and hind femora with distal ⅔ brown. Wing mostly brownish, a lighter wide band along most of CuA; Sc basically absent; C produced well beyond tip of R_5_, over half way to tip of M_1_; first section of Rs almost transverse, R_5_ running very close to R_1_ and C; M_1+2_ slightly shorter than r-m; M_4_ absent, CuA running parallel to M_2_, slightly curved at distal end; anal fold long, well sclerotized, arched on distal third of wing; wing posterior margin with a deep incision at level of tip of CuA. Abdominal tergites 1–7 brown. Male syngonocoxite with a pair of short posterior projections at ventral face bearing a group of 2–3 strong setae slightly curved distally; gonostylus short, with a short ventral beak-like, bare projection directed inwards and a larger posterior, hairy projection; tergite 9 rectangular, wide; cerci regularly developed.

***Platyprosthiogyne_gohsookhimae_ZRCBDP0048727_ha pZRCBDP0048727_SMH_holotype* [58: C, 145: C, 146: A, 199: T, 205: T, 212: C, 235: T, 253: T]**

tttatcctcatctattgctcatacaggagcttctgttgatttaactattttttctctCcatttagctggtatttcctcaattttaggagctattaattttattacaactattattaatatacgagccccaggaatttattttgaCAaaatacctctatttgtttgatctgtttttattacagctattcttcttcttctTtctctTccagttCtagcaggagctattactatactTttaacagaccgaaatatTaatacatcattttttgaccccgcaggaggaggagacccaattctttatcaacatctattt

#### Description

**Male** (Fig. 103A). Wing length, 1.63; width, 0.75. **Head** (Fig. 103C). Dark brown, lighter on occiput along eye margin, face light brown, clypeus brown dorsally, lighter towards ventral margin. Antennal scape light brown, pedicel whitish-yellow, flagellomere 1 light brown, other flagellomeres brown. Palpus light brown, lighter towards apex. Labella light brownish-yellow. **Thorax** (Fig. 103E). Scutum and scutellum dark brown. Antepronotum ochre-brown, proepisternum brown. Anepisternum dark ochre-brown, katepisternum, mesepimeron, laterotergite, mediotergite, and metepisternum brown. Antepronotum only with long, fine setae, proepisternum elongate, with short setae and four bristles along ventral margin. Mesepimeron with three bristles and four small setae. Laterotergite with four bristles on dorsal half; metepisternum with seven long, fine setae. **Legs**. Fore coxa whitish with a yellowish-brown tinge posteriorly on basal half; mid and hind coxae brown at basal half, whitish distally. Fore femur light yellowish-brown, mid and hind femora whitish at basal third to fourth, dark brown at distal ⅔ or ¾. Tibiae and tarsi light brown, tibiae slightly darker basally, tarsi slightly darker towards tip. Haltere white. **Wing** (Fig. 103D). Wing membrane light brown fumose along axis of CuA, much darker along anterior margin, darker along radial and medial veins and posterodistally at anal lobe, a lighter wide band along most of CuA length. C extending beyond tip of R_5_ for about ⅔ of distance to M_1_. Sc extremely short, ending free. First sector of Rs nearly transverse; R_5_ running very close to C, curved along its length, reaching C before level of tip of CuA; M_1+2_ half of r-m length. Medial fork long, M_2_ diverging towards tip, M_1_ and M_2_ barely sclerotized at very tip; M_4_ absent, CuA basically parallel to M_2_. Cubital pseudovein short, restrict to basal fifth of CuA, CuP entirely absent; sclerotized anal fold present, long, gently curved on distal third. Dorsal setae present on bR, R_1_, R_5_, r-m and bM, ventral setae on R_1_ and distal ⅔ of R_5_. Posterior margin strongly emarginated at level of tip of CuA. **Abdomen**. Tergites 1– 6 brown, tergite 7 light brown; sternite 1–6 light greyish-brown, sternite 7 yellowish. **Terminalia** (Fig. 103F–H). Light yellowish-brown. Gonocoxites fused on anterior half of their extension, no suture, a short medial projection on syngonocoxite extending into aedeagal-parameral complex, some few long, fine setae close to posterior margin of gonocoxites ventrally, gonocoxite posterior margin not projected beyond base of gonostylus. Gonostylus complex, composed of: (1) a ventral lobe elongate towards mid of terminalia (almost touching medially tip of other gonocoxite ventral lobe), bearing close to base a long, fine setae and at distal end a digitiform projection with 2–3 long strong setae; (2) a distal lobe projecting inwards with a medial tooth and a short spine, distal end truncated; (3) a large dorsal subtriangular lobe with a long seta projected anteriorly and setulae on posterior face. Gonocoxal bridge not visible. Aedeagal-parameral complex present as a rhomboid sclerite wider at distal end, with three small pointed projection, and a medial short keel. Tergite 9 short, rectangular, weakly sclerotized, with microtrichia and some fine setae. A pair of large lobes possibly corresponding to fusion between tergite 10 and cerci (or loss of tergite 10).

**Female** (Fig. 103B). As male, except for the following.

**Wing l**ength, 1.76; width, 0.80. General coloration more reddish-brown, abdomen lighter. **Terminalia** (Fig. 103I). Sternite 8 trapezoid, elongate, extending distally to reach mid of cercomere 1, no posterior incision. Sternite 9 with slender anterior end, distal medial end of sternite 9 reaching level of tip of lobes of sternite 8, one seta at each side near tip coming out of a small digitiform projection. Tergite 8 trapezoid, with microtrichia and setulae close to anterior margin and fine setae along posterior margin, extending laterally towards ventral face of terminalia, partially overlapping sternite 8 anteriorly, posterior margin projecting beyond posterior margin of tergite 9+10. Tergite 9+10 short, slightly more sclerotized, a pair of rounded laterodistal projections, covered with microtrichia and three fine setae on each side close to posterior margin, fused to sternite 9 laterally. Cercomere 1 slender, very long, almost 10× longer than wide, covered with microtrichia, some fine setulae on distal half. Cercomere 2 minute at tip of cercomere 1.

#### Material examined

**Holotype**: male, ZRCBDP0048727, Nee Soon (NS2), 15.apr.2015, MIP leg. (website photo specimen, slide-mounted).

**Paratypes** (3 males): ZRCBDP0048764, Nee Soon (NS1), 25.feb.15, MIP leg.; ZRCBDP0048965, Nee Soon (NS1), 15.apr.15, MIP leg. (slide-mounted) (MZUSP); ZRCBDP0049008, Nee Soon (NS2), 17.dec.14, MIP leg.

**Possibly non-conspecific cluster** (1 female): ZRCBDP0047897, Nee Soon (NS2), swamp forest, 16.oct.13, MIP leg. (website photo specimen, slide-mounted).

**Etymology**. The species epithet of this species honors Goh Soo Khim (1944–), Singapore-born ballerina, instructor and principal dancer at the Singapore Ballet Academy. Credited for the development of ballet in Singapore and for nurturing many students who became successful dancers, soloists, and even choreographers. She served as a Co-director of the National Dance Company and co-founded the Singapore Dance Theatre. She was inducted into the Singapore Women’s Hall of Fame in 2014.

**Remarks**. Most delimitation approaches suggest two species because the distance between the subclusters of *Platyprosthiogyne gohsookhimae* Amorim & Oliveira, **sp. n.** is 4.18%. It is possible that the only specimen of one subcluster (a female: ZRCBDP0047897) may not be conspecific with the holotype and is thus not included as a paratype.

*Platyprosthiogyne rahimahae* Amorim & Oliveira, sp. n. (Figs. 104A–D)

https://singapore.biodiversity.online/species/A-Arth-Hexa-Diptera-000726,-002121

urn:lsid:zoobank.org:act:E543A704-4B5F-4FBE-803D-600AFACDB983

**Diagnosis**. Male vertex dark brown, dark ochre-yellowish around eye and on face, scutum dark brown; female head largely ochre-yellowish, light brown medially on vertex, a slender ochre-yellow band along anterior margin. Pleural sclerites brown, antepronotum slightly lighter. Legs whitish, mid coxa with slender brownish transverse band on basal end, mid coxa with basal third brownish, mid femur with brownish tip, hind femur with distal half brownish. Wing with a darker macula over area around distal half of cell br; Sc nearly absent, ending free; C produced beyond tip of R_5_ for a third of distance to tip of M_1_; first section of Rs almost transverse, R_5_ running very close to R_1_ and C; M_1+2_ about as long as r-m; M_4_ absent, CuA running parallel to M_2_; anal fold long, well sclerotized, curved on distal third of wing; wing posterior margin with a gentle incision at level of tip of CuA. Abdominal tergites 1–5 brown, tergite 6 brown with yellow band along posterior margin. Male syngonocoxite with no strong setae along posterior margin ventrally, a short medial projection; gonostylus with a short digitiform ventral projection with a distal slender spine and a short posterior projection; tergite 9 wide at anterior half of terminalia.

***Platyprosthiogyne_rahimahae_ZRCBDP0047098_hapZ RCBDP0047098_SMH_holotype* [25: T, 76: C, 94: C, 11 8: C, 128: A, 136: C, 277: C]**

tttatcttcaactattgctcatgcTggagcttcagtagatttagctattttttctcttcatttagcaggaatttcCtcaattttaggagcaatCaattttattacaacaattattaaCatacgagctAcaggaatCacatttgatcgaatacccttatttgtttgatctgtttttattacagctattttattacttttatcattaccagtattagctggagctattactatactattaactgatcgaaatttaaatacttcattttttgaccctgcCggaggaggagatcctattttatatcaacatttattt

#### Description

**Male** (Fig. 104A). Wing length, 1.54–1.63; width, 0.62–0.70. **Head**. Vertex mostly ochre-brown, greyish-brown medially; face and clypeus whitish-yellowish. A line of four longer setae on occiput around dorsal margin of eye posteriorly to ocelli line. Antennal scape, pedicel, and basal half of flagellomere 1 whitish-yellow, remaining of antenna brown. Maxillary palpus light yellowish-brown, lighter to apex, labella whitish-yellow. Flagellomere 4 length 1.9× width. **Thorax**. Scutum shinning dark brown, scutellum dark brown. All pleural sclerites dark brown, except for slightly lighter antepronotum. Proepisternum with three long setae close to dorsal margin, anepisternum with six long setae along posterior margin. Mesepimeron with two long setae and eight small setae, laterotergite with four long setae along posterior margin and nine small setae; metepisternum with 11 fine setae along its length. Haltere white. **Legs**. Fore coxa whitish with a light greyish tinge, mid and hind coxae whitish with a brown transverse band across basal end, wider brown area on hind coxa; fore femur whitish-yellow, light brown on dorsal and ventral borders; mid and hind femora whitish on anterior half, dark brown on distal half. Tibiae and tarsi light yellowish-brown. **Wing** (Fig. 104C). Wings fumose brown, darker along entire anterior margin, a dark mark on anterior third from margin to bM and along M_1+2_. C extending beyond tip of R_5_ to about a third of distance to M_1_. First sector of Rs slightly oblique; R_5_ curved along its entire length, running close to R_1_ and C; r-m short, M_1+2_ about 1.5× r-m length. Medial fork long, M_1_ and M_2_, mostly parallel, almost straight, tip of M_1_, M_2_ and CuA barely sclerotized; CuA straight, parallel to M_2_. Anal fold sclerotized, present, long, gently curved on distal third. Wing posterior margin emarginated at level of tip of CuA. **Abdomen**. Tergites 1–5 dark brown, tergite 6 brown with a yellowish transverse band along posterior margin, tergite 7 yellowish; sternite 1–7 light brown. **Terminalia** (Fig. 104D). Whitish-yellow. Gonocoxites fused medially along anterior half of terminalia, suture of fusion present, a short rhomboid medioventral process of syngonocoxite distally, distal margin of gonocoxite not extending beyond base of gonostylus, some few fine setae laterodistally. Gonostylus complex, composed of: (1) ventral lobe, with two sublobes, one more dorsal branch, digitiform and slightly arched, with a distal tooth, a spine and some setulae, and one more ventral branch, elongate, curved inwards, almost touching medially tip of opposed gonostylus, with one strong seta and some setulae along posterior margin; (2) a distal clavate lobe, with one spine and one setula on ventral face and two setae on other face; and (3) a dorsal slender lobe extended ventrodorsally, with four setulae at distal end. Gonocoxal bridge not visible. Aedeagal-parameral complex weakly sclerotized, not discernible. Tergite 9 rectangular, with lateroposterior corners slightly projected, covered with microtrichia and some fine setae. A pair of large lobes possibly corresponding to fusion between tergite 10 and cerci (or loss of tergite 10).

**Female** (Fig. 104B). As male, except for the following. **Wing l**ength, 1.63; width, 0.70. General coloration lighter, more yellowish, especial head, mostly ochre-yellowish, with a brown mark on top of vertex and anterior margin of scutum. **Terminalia**. Sternite 8 trapezoid, elongate, extending distally, no posterior incision, microtrichia and few scattered setulae. Sternite 9 with wide anterior end extending anteriorly to distal end of segment 7, distal medial end of sternite 9 reaching level of tip of lobes. Tergite 8 short, extending laterally towards ventral face of terminalia, partially overlapping sternite 8 anteriorly, with microtrichia and fine setae along posterior margin. Tergite 9+10 short, lateroposterior corners slightly projected, covered with microtrichia and fine setae, fused to sternite 9 laterally. Cercomere 1 slender, very long [cercomeres 2 probably broken].

#### Material examined

**Holotype**: male, ZRCBDP0047098, National University of Singapore (PGP), 08.jul.15, MIP leg. (slide-mounted). **Paratypes** (3 males, 2 females). **Males**: ZRCBDP0049342, National University of Singapore (PGP), 08.apr.15, MIP leg. (slide-mounted) (MZUSP); ZRCBDP0072463, Bukit Timah, primary forest (BT04), 08.dec. 16, MIP leg; ZRCBDP0155018, Nee Soon (NS2), 18.may.15. **Females**: ZRCBDP0048428, Nee Soon (NS2), swamp forest, 26.dec. 12-02.jan.13, MIP leg. (website photo specimen, slide-mounted); female, ZRCBDP0048789, National University of Singapore (PGP), 03.Jun.15, MIP leg. (slide-mounted).

**Additional sequenced specimens**: female, ZRCBDP0049075, National University of Singapore (PGP), 22.apr.15, MIP leg. (slide-mounted).

**Etymology**. The species epithet honors Rahimah Rahim (1955–), Singapore-born singer and actress, known as Singapore’s first Lady of Song. She released her first album, Mana Ibumu, in 1972, when she was 17 and her albums Gadis Dan Bunga and Bebas went gold. She was inducted into the Singapore Women’s Hall of Fame in 2017.

*Platyprosthiogyne lynetteseahae* Amorim & Oliveira, sp. n. (Figs. 105A–B)

urn:lsid:zoobank.org:act:500A65E6-A924-45E6-A1E1-6C245EC807BE

**Diagnosis**. Head and scutum dark brown, pleural sclerites mostly greyish-brown, some of sclerites darker. Mid coxa with light brown tinge on basal half. Wing light brown, with a brown mark at anterior margin at origin of Rs; C produced over 1/6 of distance between R_5_ and M_1_; first sector of Rs slightly oblique, R_5_ running close to R_1_ and C; M_1+2_ as long as r-m; anal fold long, well sclerotized, arched on distal third of wing.

***Platyprosthiogyne_lynetteseahae_ZRCBDP0048297_hap ZRCBDP0048297_SMH_holotype* [7: C, 19: A, 22: C, 2 5: C, 128: A, 148: T, 175: T, 256: C]**

tttatcCtcaactattgcAcaCgcCggagcttctgttgatttagcaattttttctcttcatttagcaggaatttcctcaattttaggagcaattaattttattacaacaattattaatatacgagctAcaggaattacatttgatcgTatacctctatttgtttgatctgttttTattacagctattttattacttttatcattaccagtattagctggagctattactatactattaactgatcgaaatttaaaCacttccttttttgaccctgctggaggaggagatcctattttatatcaacacttattt

#### Description

[Abdomen broken on holotype, only known specimen]. Wing length, 2.05; width, 0.82. **Head**. Strongly compressed, mostly dark ochre-brown, lighter towards ventral end of occiput; face and clypeus light brown. Mid ocellus present; four long setae on occiput around dorsal margin of eye. Antennal scape, pedicel, and basal half of flagellomere 1 brownish-yellow, remaining flagellomeres brown; flagellomere 4 length 1.4× width. Palpus light yellowish-brown, lighter towards apex; labella whitish-yellow. **Thorax** (Fig. 105A). Scutum shinning dark brown except for ochre-yellow transverse band along anterior margin that projects posteriorly at laterals to level of anterior spiracle. Scutellum dark brown. Antepronotum ochre-yellow, proepisternum greyish-brown, anepisternum dark brown, katepisternum greyish-brown, mesepimeron, laterotergite, and metepisternum brown, mediotergite dark brown. Antepronotum only covered with short setae, proepisternum with short setae and three bristles close to dorsal margin. Anepisternum with five long setae along posterior margin. Mesepimeron with two strong setae and five small setae, laterotergite with four long setae along posterior margin and 20 small setae; metepisternum with eight setulae along its length. **Legs**. Fore coxa whitish with a brownish tinge anteriorly, mid coxa whitish with a light brown tinge along basal end, hind coxa whitish with a brown transverse band across basal end [hind leg missing except for coxa]; fore and mid femur whitish-yellow with brownish distal end; tibiae and tarsi yellowish-brown, tarsi darker towards apex. Haltere white. **Wing** (Fig. 105B). Wings fumose light brown, with a brown mark at anterior margin at level of origin of Rs. C extending beyond tip of R_5_ for 1/6 of distance to tip of M_1_. First sector of Rs slightly oblique; R_5_ curved, running close to R_1_ and C; r-m about as long as M_1+2_. Medial fork long, M_1_ and M_2_ mostly parallel, tip of M_1_, M_2_, and CuA barely sclerotized at very tip, CuA straight. Cubital pseudovein short reaching level of anterior end of r-m; CuP entirely absent; sclerotized anal fold present, long, gently arched on distal half. Wing margin emarginated at level of tip of CuA.

#### Material examined

**Holotype**: ZRCBDP0048297, Pulau Semakau (SMN1), planted mangrove, 10.oct.13, MIP leg. (unknown gender, abdomen missing) (website photo specimen, slide-mounted).

**Etymology**. The species epithet honors violinist Lynnette Seah Mei Tsing (1957–), a Singapore-born Cultural Medallion-winner (the highest award for the arts in Singapore) for Music in 2006. She was a founding member of the Singapore Symphony Orchestra, which she co-leads, and has performed for Emperor Akihito and Empress Michiko of Japan. She was inducted into the Singapore Women’s Hall of Fame in 2014.

**Remarks**. We have a single specimen of this species, without abdomen. The scutum color pattern is similar to *P. phanwaithongae* Amorim & Oliveira, **sp. n.**, with yellowish shoulders, but diverges from the remaining four species of the genus in Singapore. All delimitation approaches confirm it as a separate species.

*Platyprosthiogyne neilaae* Amorim & Oliveira, sp. n. (Figs. 106A–E)

https://singapore.biodiversity.online/species/A-Arth-Hexa-Diptera-000767

urn:lsid:zoobank.org:act:B0E83FDB-2E55-4BED-A729-8EDE721A5844

**Diagnosis**. Head, scutum and pleural sclerites dark brown, some of pleural sclerites slightly lighter. Coxae and femora mostly whitish, mid coxa with basal half brownish, mid coxa with basal ¾ brown, mid and hind femora with distal ⅔ brown. Wing mostly brownish, a lighter elongate area around most of CuA length; Sc nearly absent; C produced over half way between R_5_ and M_1_; first section of Rs almost transverse, R_5_ running very close to R_1_ and C; M_1+2_ slightly shorter than r-m; M_4_ absent, CuA running parallel to M_2_; anal fold long, well sclerotized, arched on distal third of wing; wing posterior margin with clear incision at level of tip of CuA. Abdominal tergites 1–7 brown. Male syngonocoxite with a short medial posterior projection at ventral face bearing a group of 7–8 strong setae slightly curved distally; gonostylus small, with a short ventral digitiform projection directed inwards with a spine distally and a larger dorsal projection; tergite 9 rectangular, wide.

***Platyprosthiogyne_neilaae_ZRCBDP0048565_hapZRCB DP0048565_SMH_holotype* [12: A, 26: A, 52: C, 161: T, 162: G, 199: C, 202: C, 205: T]**

tctatcttcttAtattgctcatacaAgagcttctgttgatttaactattttCtctcttcatttagctggtatttcttcaatcttaggagcaattaatttcattacaactattattaatatacgagccccaggtatttattttgataaaatacctttatttTGctgatctgtatttattacagccattcttcttcttctCtcCctTccagttttagcaggagcaattactatattactaacagaccgaaatattaatacatcattttttgaccctgcaggaggaggagatcctattctttatcaacatttattt

#### Description

**Male** (Fig. 106A). Wing length, 1.63–1.92; width, 0.70–0.80. **Head**. Dark brown, occiput lighter around eye, face and clypeus brown. Antennal scape, pedicel and first third of flagellomere 1 whitish-yellow, other flagellomeres brown; flagellomere length over 3× width. Palpus light brown, lighter towards apex; labella light brownish-yellow. **Thorax**. Scutum and scutellum dark brown. Antepronotum ochre-brown, proepisternum brown; anepisternum dark ochre-brown, katepisternum, mesepimeron and metepisternum brown, laterotergite and mediotergite dark brown. Antepronotum only with short setae, proepisternum with short setae and three longer setae along ventral margin. Anepisternum with six long setae along posterior margin. Mesepimeron with eight setae, laterotergite with three setae; metepisternum with 3– 4 fine setae; other pleural sclerites bare. **Legs**. Fore coxa whitish-yellow, mid and hind coxae brown on basal half, hind coxa darker; fore femur whitish-yellow with brown tinge, darker towards apex, mid femur light ochre-yellow on basal half, brownish at distal half, hind femur whitish on basal fourth, dark brown on distal ¾; tibiae yellowish-brown, tarsi light brown, slightly darker towards tip. Haltere white. **Wing** (Fig. 106B). Membrane light greyish-brown fumose, lighter around length of CuA. R_5_ running very close to C, curved along its length, C extending beyond tip of R_5_ to about half distance to M_1_; r-m short, M_1+2_ short, as long as r-m. Medial fork long, M_1_ and M_2_ diverging on distal fifth, M_1_ and M_2_ barely sclerotized at very tip. CuA straight, reaching wing margin beyond level of tip of R_5_; only a stump of cubital pseudovein remaining; CuP entirely absent, sclerotized anal fold long, gently curved. Wing margin strongly emarginate at level of tip of CuA. **Abdomen**. Tergites 1–6 brown, sternites 1–6 light greyish-brown, tergite 7 light brown, sternite 7 yellowish-brown. **Terminalia** (Figs. 106C–E). Yellowish-brown. Gonocoxites fused medially along anterior half of terminalia, no evidence of suture, a short medioventral process of syngonocoxite distally bearing a row of seven long, strong setae on posterior margin, distal margin of gonocoxite not extended beyond base of gonostylus, some few fine setae laterodistally. Gonostylus relatively simple, composed of: (1) ventral short digitiform lobe, with a subapical setula and an apical spine; (2) a main distal or dorsal bifid lobe, with a short branch in a ventral position bearing three elongate setae a tip directed ventrally and a larger branch with a strong seta midway to apex directed inwards, bearing some fine setae on dorsoposterior end and a small laminar extension. Gonocoxal bridge present, no apodemes. Aedeagal-parameral complex, with a ventral subrectangular plate slightly widened towards apex, distally with a pair of short, rounded lobes on lateroposterior corners, and a large trapezoid plate extending posteriorly, with a pair of subapical long setae. Tergite 9 rectangular, lateroposterior corners slightly projected, with microtrichia and fine setae along posterior margin. A pair of large lobes possibly corresponding to fusion between tergite 10 and cerci.

**Female**. Unknown.

#### Material examined

**Holotype**: male, ZRCBDP0048565, Nee Soon (NS1), swamp forest, 04.apr. 2012, MIP leg. (website photo specimen). **Paratype**: male, ZRCBDP0140723, Singapore, date range 2012-2018, MIP leg. (slide-mounted).

**Etymology**. The species epithet of this species honors Neila Sathyalingam (1938-2017). Sri Lankan-born and with Singapore citizenship in 1994, she was a leading classical Indian dancer, teacher and choreographer in Singapore. She was appointed the dance instructor and choreographer for the Indian Dance Group of the People’s Association and was an artistic adviser to Singapore’s National Arts Council. For her contributions to dance, Neila was awarded the Singaporean Cultural Medallion in 1989. She was inducted into the Singapore Women’s Hall of Fame in 2014.

**Remarks**. *Platyprosthiogyne neilaae* Amorim & Oliveira, **sp. n.** and *P. gohsookhimae* Amorim & Oliveira, **sp. n.** seem to belong to a small clade within the genus, as the medio-posterior syngonocoxite projection with elongate, curved setae suggest. Their separation into distinct species is corroborated by all the species delimitation approaches.

*Platyprosthiogyne snehalethaae* Amorim & Oliveira, sp. n. (Figs. 107A–D)

https://singapore.biodiversity.online/species/A-Arth-Hexa-Diptera-000768

urn:lsid:zoobank.org:act:FDA6AF0E-48D2-4331-AAC6-6E757D1A7037

**Diagnosis**. Head light ochre-brown, frons dark brown. Scutum ochre-brown, darker on posterior half, scutellum dark brown. Pleural sclerite greyish-brown. Wing membrane light greyish fumose. Sc short, ending free. C extending for less than a ¼ of distance to M_1_. R_5_ gradually curved towards posterior margin; r-m short, oblique. M_4_ missing; CuA straight; sclerotized anal fold long. Wing posterior margin conspicuously emarginated at apex of CuA. Abdominal tergites 1–7 dark greyish-brown, tergites 1 with cream-yellow anterior mark medially, tergite 2 with cream-yellow mark on anterolateral corner. Gonocoxite elongate, no medial projection of posterior margin.

Gonostylus ovoid in lateral view, dorsal half folded over ventral half. Tergite 9 wide, short.

***Platyprosthiogyne_snehalethaae_ZRCBDP0049034_hap ZRCBDP0049034_SMH_holotype* [10: C, 106: C, 138: T, 149: C, 151: T, 269: A] -**

ctttcagcCtcaattgctcatgcaagagcttcagtagatttatctattttttctcttcacctagcaggtatttcatcaattttaggagctattaactttattacCactattattaacatacgaacaccaggaatatTttttgatcgtCtTcctttatttgtatgatccgttttaattactgccattttattattattatctttacctgtattagcaggagccatcacaatacttctaacagatcgtaatttaaatacttcattttttAacccagcagggggaggagaccctattttatatcaacatttattt

#### Description

**Male**. Wing length, 2.27; width, 1.02. **Head**. Light ochre-brown, frons dark brown. Lateral ocelli blackish-brown, nearly touching eye margin, mid ocellus absent, frontal furrow present. Dark brown setae scattered on vertex, a short line of four slightly longer setae on occiput around dorsal margin of eye posteriorly to ocelli line. Long interommatidial setulae over entire surface of eye. Antennal scape and pedicel light brown, flagellum brown. Scape long, about twice longer than pedicel. Scape with setae on inner face and dorsally. Pedicel with setation restricted to a crown of setae on posterior margin, in addition to a longer dorsal seta and some few additional smaller setae. Flagellomeres almost twice as long as wide, except for flagellomere 1, about 3× longer than wide. Face short, light greyish-brown, clypeus slightly bulging, light brown, setose. Maxillary palpus light brown, lighter toward apex, with five palpomeres; palpomere 1 only with microtrichia, palpomere 2 short, with some setae along distal margin dorsally, palpomere 3 with a conspicuous sensorial pit opening on basal half of internal face and setae on inner face, palpomere 4 almost twice palpomere 3 length, slender, with setulae dorsally and on internal face, palpomere 5 over twice palpomere 4 length. Labella large, light brownish-yellow. **Thorax**. Scutum ochre-brown, darker on posterior half, densely covered with scattered long setae, two small bristles above wing and a pair of prescutellar bristles medially. Scutellum dark brown, large, with four long subapical bristles along posterior margin and some scattered long setae. Pleural sclerite greyish-brown. Antepronotum with a number of shorter setae and some fine bristles, proepisternum with some few short setae and two bristles. Anepisternum covered with scattered setae, some slightly longer setae along posterior margin. Mesepimeron with 12 setae on dorsal third; laterotergite strongly bulging, with 18 setae; metepisternum with four small setae on posterior fourth. Katepisternum and mediotergite bare. **Legs**. Coxae ochre-yellowish, fore and hind coxae slightly darker; femora brownish-yellow, tip of hind femur brownish. Tibiae light yellowish-brown, tarsi light brown. Fore coxa covered with small fine setae on anterior and lateral faces, a row of strong setae along distal ⅔ of lateroposterior edge; mid coxa with small setae on distal third besides distal stronger setae anteriorly and one strong seta posteriorly on distal margin; hind coxa with one strong seta on lateral face dorsally, a row of small dark setae along posterior half of posterior face and one bristle and some setae laterally on distal end. Femora densely covered with fine setae, some longer setae on distal end ventrally. Fore tibia with scattered fine setae [mid tibia and tarsi missing]; hind tibia with scattered fine setae ventrally and laterally, and two laterodorsal rows of seven long setae. Anteroapical depressed area on inner face of fore tibia wide, densely lined with setulae. Tarsi with regular rows of trichia, setae only distally. Tarsal claw with a strong, large basal tooth. Tibial spurs about twice tibial width at apex. Haltere light brown. **Wing** (Fig. 107B). Wing membrane light greyish fumose, venation brownish, no markings, microtrichia exceptionally long. Sc short, ending free. C extending shortly beyond tip of R_5_, less than a fourth of distance to M_1_. R_1_ long, gently arched, reaching C at distal third of wing. First sector of Rs slightly oblique, R_5_ gradually curved towards posterior margin; r-m short, oblique. M_1+2_ short, about 2.5× r-m length. Medial fork long, M_1_ and M_2_ slightly divergent on distal fifth of wing; M_4_ missing. CuA straight; CuP produced, not reaching level of origin of M_1+2_; sclerotized anal fold present, long. Dorsal setae present on bR, R_1_, second sector of Rs, r-m, and bM; ventral setae present on distal half of bR, R_1_, second sector of Rs, r-m, and distal ⅔ of bM. Wing posterior margin conspicuously emarginated at apex of CuA. **Abdomen**. Tergites 1–7 dark greyish-brown, tergites 1 with cream-yellow anterior mark medially, tergite 2 with cream-yellow mark on anterolateral corner; sternite 1 light brown, sternites 2–6 cream-yellow, segment 7 light brown. **Terminalia** (Figs. 107C–D). Light brownish-yellow. Gonocoxite elongate, indistinguishably fused medially, no suture, no medial projection of posterior margin. Gonostylus ovoid-shape in lateral view, dorsal half folded over ventral half, marginal setae of both “valves” close to each other. Aedeagal-parameral complex long, more or less rectangular in ventral view, extending posteriorly beyond level of base of gonostylus. Tergite 9 wide, short, with long setae on posterior half. Tergite 10 present as an elongate trapezoid, weakly sclerotized plate, almost reaching level of tip of aedeagus. Cerci almost as long as tergite 10.

**Female** (Fig. 107A). As male, except for the following.

**Wing** length, 2.37; width, 1.12. Sternite 8 slender, with no posterior lobes, wide at base, extending distally almost to level of tip of first cercomere, covered with fine setulae. Tergite 8 fused to tergite 9 and tergite 10. Sternite 9 with wide base, extending distally to level of tip of sternite 8. Cercomere 1 very long, over 3× longer than cercomere 2, both covered with microtrichia and setulae.

#### Material examined

**Holotype**: male, ZRCBDP0049034, Nee Soon (NS2), 07.jan.15, MIP leg. (slide-mounted). **Paratype** (1 female): ZRCBDP0048566, Nee Soon (NS2), swamp forest, 09.may. 12, MIP leg. (website photo specimen, slide-mounted).

**Etymology**. The species epithet of this species honors Snehaletha Karunakaran. She was a student of D.H. Murphy and performed what was arguably the first extensive modern synthesis of the Chironomidae (Diptera) in Singapore. Her unpublished PhD thesis (1969) accumulated a wealth of information on taxonomy and bionomics of the group. She sadly died in a fire at home before she could be awarded her PhD.

**Remarks**. We have two haplotypes and no delimitation conflicts.

### Platurocypta Enderlein

*Platurocypta* Enderlein 1910: 76. Type-species, *P. limbatifemur* Enderlein (orig. des.).

**Diagnosis**. Lateral ocelli touching eye margins, third palpomere not swollen. Border of scutum at anterior third nearly straight or only slightly curved; katepisternum dorsoventrally compressed; laterotergite densely covered with small setae and a row of bristles along dorsoposterior margin; mesepimeron with bristles; laterotergite setose, compressed; mediotergite strongly compressed. Wing membrane with microtrichia arranged in more or less regular longitudinal lines; C clearly extending beyond tip of R_5_; Sc ending free; R_4_ absent; M_1+2_ about as long as r-m; M_4_ present, parallel to M_2_ on distal half, originating slightly beyond level of anterior end of medial fork, tip of M_4_ about equidistant between tip of M_2_ and tip of CuA.

*Platurocypta* is a rather poorly known genus with 20 described species: two from Europe, one from South America, two from New Zealand, 13 from the Comores and Seychelles Islands and continental Africa, and two from the Oriental region. The fact that the genus has species known from Europe and from South America may to some extent hidden its rather typical Afro-Oriental-Australasian distribution. We should expect more Oriental and Australasian species of the genus in the future.

*Platurocypta* may belong to a small clade with *Epicypta* but does not share all synapomorphies of the *Epicypta*. *Platurocypta* has a conspicuous incision on the margin of the scutum above the anterior spiracle, a long C beyond the tip of R_5_, and a reduced laterotergite and mediotergite. These features are shared by the Mycetophilini group of genera including *Platurocypta*, *Epicypta*, *Aspidionia*, and *Integricypta* Amorim & Oliveira, **gen. n.** .

In our Singapore samples, we found two species of *Platurocypta* but none of them fit the species description of Edwards (1929, 1931). The Oriental species of the genus—*P. intermedia* (Edwards 1929), *P. longiseta* (Edwards 1931)—were described respectively from the Philippines and from Sumatra. There are no conflicts on species delimitation (Fig. 110G), and the species have clearly divergent male terminalia morphology. Specimens of both species were collected in the mangrove and in the swamp forest.

*Platurocypta adeleneweeae* Amorim & Oliveira, sp. n. (Figs. 108A–E, 109A–C)

https://singapore.biodiversity.online/species/A-Arth-Hexa-Diptera-000723

urn:lsid:zoobank.org:act:CB0079D6-A0C2-4068-8334-6917E6914368

**Diagnosis**. Head dark brown; antenna yellow on basal half, greyish-brown on distal half, mouthparts brownish-yellow. Scutum blackish-brown; pleural sclerites dark brown, katepisternum, mesepimeron, and laterotergite lighter. Fore coxa light brownish-yellow, mid and hind coxae light brown; femora yellowish-brown medially with dark brown ventral and dorsal borders, tibiae greyish-brown; female fore tarsomeres 2-5 flattened and slightly swollen. Abdominal tergites 1–5 brown, tergite 6 brown, with yellowish transverse band along posterior margin. Male terminalia yellowish; gonocoxites with no medial suture, posterior border of syngonocoxite with no medial incision; gonostylus with a digitiform ventral projection and a main lobe with a row of short spines on small projections at distal border; tergite 9 present as a pair of large ovoid lobes medially. Female terminalia slightly elongate, weakly sclerotized.

***Platurocypta_adeleneweeae_ZRCBDP0048860_hapZRC BDP0048323_SMH_holotype* [10: C, 11: G, 12: T, 112: C, 118: C, 136: C, 181: G, 202: C]**

cctttcatcCGTtattgcccatgcaggtgcttctgttgatttagcaattttttctcttcatttagctggtatttcttcaattttaggagcaattaactttattacaactatCattaaCatacgagctccaggaatCtcttttgatcgattacctttatttgtatgatctgtattaatcacGgctattcttttacttttatcCttacctgttttagcaggagctattacaatattattaacagatcgaaatttaaatacatcattttttgaccccgcaggagggggagacccaattttataccaacatttattt

#### Description

**Male**. Wing length, 2.11 mm, width, 0.74 mm. **Head**. Vertex shinning brown, two longer setae dorsally to eye anteriorly to lateral ocellus, one seta dorsally to ocellus, 4– 5 setae posteriorly to ocellus. Occiput dark brown, light brown towards ventral end. Scape and pedicel ochre-yellow; basal flagellomeres ochre-yellow, gradually becoming greyish-brown. Face brown, clypeus light brown, labella whitish-yellow; four developed palpomeres, palpomeres 1+2 brownish-yellow, palpomeres 3–5 whitish-yellow. Entire eye surface with interommatidial short fine setae. Scape 2.0× length of pedicel, flagellomere 4 length 1.8× width. **Thorax**. Scutum shining blackish-brown, scutellum blackish-brown dirty-yellow at the sides anteriorly. Antepronotum and anepisternum shining blackish-brown, other pleural sclerites greyish-brown, mediotergite blackish-brown. Pleural membrane yellowish. Haltere with light brown pedicel and whitish knob. Scutum with seven supra-alar bristles and three pairs of prescutellar bristles, scutellum with two pairs of strong bristles. Antepronotum with three bristles at ventral margin directed ventrally. Anepisternum with six bristles along posterior margin. Mesepimeron with four bristles and 26 small setae and setulae, laterotergite with four bristles and 14 additional smaller setae. Metepisternum with three strong setae and seven additional small setae. **Legs**. Fore coxa whitish-yellow, mid and hind coxae light brown; trochanters brownish; femora brownish-yellow, with a brownish elongated mark along dorsal and ventral edges, hind femur with a wide brown area laterally on distal third of wing; tibiae and tarsi brownish-yellow. Forecoxa with a row of large brown setae laterodistally; mid coxa with some few strong setae on anterior face of distal margin; no setulae on proximal half of hind coxa, one lateral strong seta close to tip; femora only covered with setulae and very few apical larger setae ventrally. Tibiae and tarsi with trichia organized in regular rows. Fore tibia with one large seta on inner face laterally at distal third in addition to rows of trichia; mid tibia with a pair of laterodorsal rows of small bristles and a ventral row of bristles; hind tibia with two irregular laterodorsal rows of 3–5 strong bristles. Fore leg tarsomere 1 about as long as tibia, more than 2.5× tarsomere 2 length. Mid and hind tarsomeres with regular rows of setae ventrally in addition to regular rows of trichia. Tibial spurs 1:2:2, hind spur about 6× tibia width at apex, inner spur slightly shorter than outer spur. Tarsal claws with a small acute ventral tooth close to base and a lateral mid tooth directed outwards. **Wing**. Membrane fumose light brown, cell c and area around tip of R_1_, first sector of Rs and base medial fork slightly darker. Humeral vein present, oblique, Sc barely produced. R_1_ reaching C on distal fourth of wing; R_5_ reaching C before level of tip of M_2_; C extending beyond apex of R_5_ for about a third of distance to M_1_. First sector of Rs almost transverse, devoid of setae; r-m more or less oblique, well sclerotized, setose, slightly more than twice length of first sector of Rs. M_1+2_ short, about as long as r-m; bM about 5× r-m length; first sector of CuA about as long as second sector. Cubital pseudovein extending to slightly beyond origin of M_4_. Anal fold long, curved on posterior third. Dorsal microtrichia on radial veins, M_1_ on distal half, M_2_ on distal fifth, and M_4_ on distal end; anal lobe membrane with macrotrichia dorsally. **Abdomen**. Abdominal tergites 1-6 light brown, distal fourth of tergite 6 and tergite 7 cream-yellowish; sternites 1-7 whitish-yellow, no bristles ventrally on sternite 2. **Terminalia** (Figs. 109A–C). Yellowish. Gonocoxites large, fused along anterior half of terminalia, no suture of fusion, no medial incision on posterior border of syngonocoxite. Gonostylus composed of a digitiform ventral lobe with two short spines distally, a short, sclerotized flat inner lobe bare of setae and a main lobe with a row of small setae along posterior margin arising from short projections. Parameres composed of a pair of small plates connected anteriorly. Tergite 9 present as a pair of large ovoid lobes with microtrichia and no setae. Cerci weakly sclerotized. **Female** (Fig. 108A). As male, except for the following. Wing length, 2.27; width, 0.90. **Head** (Fig. 108B). Occiput with three longer setae dorsally to eye anteriorly to lateral ocellus, one seta dorsally to ocellus, 7 setae posteriorly to ocellus. **Thorax** (Fig. 108B). Eight supra-alar bristles. Proepisternum with one small and three strong bristles. Anepisternum with seven bristles along posterior margin. Mesepimeron with four bristles and 19 small setae and setulae, laterotergite with three bristles and four additional smaller setae. Metepisternum with four strong setae and six additional small setae. **Legs**. Fore leg tarsomere 1 about as long as tibia, more than 2.5× tarsomere 2 length; fore tarsomeres 2-5 slightly swollen. **Wing** (Fig. 108D). No dorsal setae on veins M_1_on distal half, M_2_ on distal fifth and M_4_, membrane on anal lobe with macrotrichia. **Abdomen**. Abdominal tergites 1-6 dark caramel-brown, tergite 6 with a yellowish band along posterior margin; tergite 7 brownish-yellow; sternites 1-7 brownish-yellowish, no bristles ventrally on sternite 2; sternites 3–6 with three longitudinal weakly sclerotized bands along sternite. **Terminalia** (Figs. 108D–E). Brownish-yellow. Sternite 8 rectangular, elongate, with rounded posterior margin, no medio-posterior incision, entirely covered with microtrichia and elongate setulae, no stronger setae along distal margin. Sternite 9 long and slender, anterior arm elongate, wide at tip, genital chamber slender, posterior end projected to level of tip of first cercomere. Tergite 8 short, with a medial weakly sclerotized area almost separating two lobes, extended lateroanteriorly at each side as apodemes. Tergite 9+10 reduced to a short, bare, sclerotized band. Cercomere 1 1.4× length of cercomere 2, both with microtrichia and setulae; cercomere 1 with a short projection distal to base of cercomere 2 ventrally, with a longer seta at tip; cercomere 2 with a longer seta at tip.

#### Material examined

**Holotype**: male, ZRCBDP0048860, Nee Soon (NS1), 14.jan.2015, MIP leg. (slide-mounted). **Paratypes** (12 females): ZRCBDP0048323, Nee Soon (NS2), swamp forest, 25.apr. 12, MIP leg. (website photo specimen); ZRCBDP0048327, Nee Soon (NS2), swamp forest, 16.may. 12, MIP leg. (website photo specimen); ZRCBDP0048424, Nee Soon (NS1), swamp forest, 09.jan.13, MIP leg. (slide-mounted); ZRCBDP0048705, Nee Soon (NS2), 28.jan.15, MIP leg.; ZRCBDP0048720, Nee Soon (NS2), 28.jan.15, MIP leg.; ZRCBDP0048821, Nee Soon (NS1), 14.jan.15, MIP leg.; ZRCBDP0048833, Nee Soon (NS1), 14.jan.15, MIP leg.; ZRCBDP0048841, Nee Soon (NS1), 14.jan.15, MIP leg.; ZRCBDP0048858, Nee Soon (NS1), 14.jan.15, MIP leg. (MZUSP); ZRCBDP0048866, Nee Soon (NS1), 10.dec.14, MIP leg.; ZRCBDP0048943, Nee Soon (NS2), 03.dec.14, MIP leg.; ZRCBDP0049109, Nee Soon (NS1), 24.dec.14, MIP leg. **Additional sequenced specimens**: male, ZRCBDP0134013, Singapore, data range 2012-2018, MIP leg.; ZRCBDP0134031, Singapore, data range 2012-2018, MIP leg.; ZRCBDP0078983, Singapore; ZRCBDP0134032, Singapore.

**Etymology**. The species epithet of this species honors bowling champion Adelene Wee (1965–). She was the youngest gold medal winner of the Philippines Women’s Open Masters in 1981, broke the world record for six-game singles in ten-pin bowling at the Sukhumvit Open in 1982, and won the Singapore International Bowling Championships. In June 1985, she won three gold medals at the Asian FIQ youth championships. She became the nation’s first and the world’s youngest World Bowling Champion, with the 1985 World Games title at the age 19. She was inducted into the Singapore Women’s Hall of Fame in 2014.

**Remarks**. We have three haplotypes and no delimitation conflicts.

*Platurocypta tanhoweliangi* Amorim & Oliveira, sp. n. (Figs. 110A–F)

https://singapore.biodiversity.online/species/A-Arth-Hexa-Diptera-000723,and-02073

urn:lsid:zoobank.org:act:CFA49724-36FA-4C4B-9BD8-00944BF904BA

**Diagnosis**. Head blackish-brown; antenna yellow on basal half, greyish-brown on distal half, mouthparts brownish. Scutum blackish-brown; pleural sclerites blackish-brown. Fore coxa light brownish-yellow, mid and hind coxae greyish brown, with a lighter medial area; femora greyish-brown, tibiae greyish-brown with yellowish basal and distal fifth; female fore tarsomeres 2-5 flattened and slightly swollen. Abdominal tergites 1–5 dark brown, tergite 6 brown on anterior ¾, yellowish on posterior fourth. Male terminalia brownish-yellow, gonocoxites with medial suture present, posterior border of syngonocoxite with a U-shaped medial incision; gonostylus with a short, sclerotized palm-like projection with a row of short blunt spines and a large median lobe; a pair of large ovoid lobes dorsally, cerci present as a pair of elongated lobes ventral to tergite 9. Female terminalia slightly elongate, cerci with a row of short spines along border.

***Platurocypta_tanhoweliangi_ZRCBDP0048063_hapZRC BDP0047827_SMH_holotype* [1: A, 11: T, 64: T, 67: T, 121: G, 133: C, 134: C, 187: A, 205: G]**

ActttcatctTctattgcccatgctggatcatcagttgatttagctattttttctcttcatctTgcTggtatttcttcaattttaggagcaattaattttattactacaattattaatatGcgagcccccggCCtatcttttgatcaattacctttatttgtatgatctgtattaattacagcagtActtttattattatccttGccagtattagctggagctattactatattattaacagatcgaaatttaaatacatctttctttgatcctgcaggaggaggagacccaatcctttaccaacatttattt

#### Description

**Male**. Wing length, 2.11 mm, width, 0.74 mm. **Head**. Head dark caramel-brown, scattered setulae over entire vertex, a row of six dark brown slightly longer setae dorsally to eyes on occiput. Scape and pedicel yellowish, flagellomeres 1–6 yellowish with a brownish tinge, flagellomeres 7–14 ochre-yellowish basally and greyish-brown distally, apical flagellomeres with more extensive greyish-brown area. Face and clypeus light brown. Maxillary palpus light brown. Labella whitish-yellow. **Thorax**. Scutum blackish-brown, yellowish-brown medially on anterior margin, scutellum blackish-brown. Pleural sclerites greyish-brown, proepisternum, most of anepisternum and dorsal half of mesepimeron darker. Pleural membrane dark ochre-yellow. Haltere whitish. Antepronotum with three long setae on ventral margin, proepisternum with three bristles. Mesepimeron with a row of six bristles and 13 smaller setae, laterotergite with four bristles and six additional smaller setae. Metepisternum with three long setae and ten setulae at posterior end. **Legs**. Fore coxa dirty yellowish-brown, mid and hind coxae mostly greyish-brown, mid coxa with a wide whitish-yellow area at ventral half. Femora greyish-brown, lighter on lateral face, darker along ventral and dorsal edges. Tibiae ochre-yellowish on both ends, darker medially, mid and hind tibiae darker. Tarsi light greyish-brown, hind tarsomere 1 with a basal dark mark. **Wing** (Fig. 110B). Membrane light fumose brown, slightly darker along anterior margin, some specimens with blackish-brown diffuse area around first sector of Rs and r-m. C extending beyond apex of R_5_ for a third distance to M_1_. First sector of Rs transverse. Setae on veins as in *Platurocypta adeleneweeae* Amorim & Oliveira, **sp. n.**, but bM with ventral and dorsal setae on distal half. Anal fold curved at distal third. **Abdomen**. Abdominal tergites 1-6 caramel-brown, tergite 1 darker, tergite 6 cream-yellow along posterior margin medially, tergite 7 cream-yellow. Sternites 1-7 light caramel-brown. **Terminalia** (Figs. 110C–E). Light ochre-yellow, cerci lighter. Gonocoxites large, fused along anterior half of terminalia, suture of fusion present, posterior border of syngonocoxite with a U-shaped medial incision, some few setulae along posterior third, a small sub-medial projection inwards. Gonostylus composed of a ventral lobe, with setulae along posterior margin, and a short, sclerotized palm-like projection with a row of five distal short blunt spines; a large, median lobe slightly widening distally, with a pointed tooth directed ventrally on inner distal end, some fine setulae on distal margin, and a group of small, fine setae on internal basal corner. Gonocoxal bridge weakly sclerotized, apodemes projected inwards, almost touching each other. Aedeagal-parameral encapsulated, a pair of weakly sclerotized pair of blades ventrally close to each other medially, aedeagal plate elongate, subquadrate, slightly more sclerotized on distal margin. A pair of large ovoid lobes dorsally, bearing microtrichia and fine setae, representing tergite 9. Cerci present as a pair of elongated lobes ventrad to tergite 9, with setae and microtrichia.

**Female** (Fig. 110A). As male, except for the following. **Wing l**ength, 2.08 mm; width, 0.77 mm. **Head**. Occiput with four long setae around eyes posteriorly to line of ocelli. **Thorax**. Mesepimeron with five bristles in line and 11 small setae, laterotergite with three bristles and eight additional smaller setae. Metepisternum with three longer setae on posterior end and 14 setulae along its length. **Abdomen**. Tergites 1–5 brown, tergite 6 mostly brown, with a medial yellowish mark medially along posterior margin, tergite 7 mostly yellowish. **Terminalia** (Fig. 110E–F). Yellowish-brown. Sternite 8 wide, with a pair of lateroposterior projections and a wide medial incision, sclerite covered with setulae and microtrichia. Sternite 9 well-sclerotized, a pair of posterior arms, gonopore medially, anterior medial furca well-sclerotized. Tergite 9+10 mostly bare, with a pair of lateroposterior projections. Cercomeres 1 and 2 partially fused, two strong spines on laterodistal end of cercomere 1, four strong spines on cercomere 2.

#### Material examined

**Holotype**: male, ZRCBDP0048063, Nee Soon (NS2), swamp forest, 27.mar.13, MIP leg. (slide-mounted). **Paratypes** (1 male, 9 females). **Male**: ZRCBDP0048423, Nee Soon (NS2), swamp forest, 16.may. 12, MIP leg. (MZUSP). **Females**: ZRCBDP0047827, Nee Soon (NS1), swamp forest, 06.mar.13, MIP leg.; ZRCBDP0047839, Nee Soon (NS1), swamp forest, 24.apr.13, MIP leg.; ZRCBDP0047891, Nee Soon (NS1), swamp forest, 20.mar.13, MIP leg.; ZRCBDP0047940, Nee Soon (NS1), swamp forest, 30.may-05.jun.13, MIP leg. (slide-mounted); ZRCBDP0047949, Nee Soon (NS1), swamp forest, 03.jul.13, MIP leg.; ZRCBDP0048324, Nee Soon (NS2), swamp forest, 18.jul. 12, MIP leg. (MZUSP); ZRCBDP0048325, Nee Soon (NS2), swamp forest, 09.may. 12, MIP leg. (website photo specimen); ZRCBDP0048326, Nee Soon (NS1), swamp forest, 25.apr. 12, MIP leg. (website photo specimen); ZRCBDP0048425, Nee Soon (NS1), swamp forest, 23.may. 12, MIP leg.

**Etymology**. The species epithet of this species honors Tan Howe Liang (1904–1999), Singapore’s first Olympic medalist, in weightlifting at the 1960 Rome games. Born in Swatow, China, Tan also broke the oldest-standing world record in the lightweight category in the clean and jerk in 1958. He was the only Singaporean Olympic medalist until the 2008 Summer Olympics.

**Remarks**. We have two haplotypes and no delimitation conflicts.

### Epicypta Winnertz

*Epicypta* Winnertz, 1863: 909. Type-species, *Mycetophila scatophora* Perris (Johannsen 1909: 110).

**Diagnosis**. Lateral ocelli touching eye margins, third palpomere not swollen. Border of scutum above anterior spiracle with a deep incision; katepisternum dorsoventrally compressed; separation between antepronotum and proepisternum barely recognizable, at most a short suture at anterior end; a row of bristles along dorsoposterior margin of anepisternum always present; mesepimeron with bristles; laterotergite and mediotergite strongly compressed. Mid coxa strongly developed. Wing membrane with microtrichia arranged in more or less regular longitudinal lines; C hardly extending beyond tip of R_5_ or not extending at all. Sc ending free, sometime barely recognizable; R_4_ absent; M_1+2_ about as long as r-m; M_4_ present, not interrupted at basal end, parallel to M_2_ on distal half, originating slightly at level of anterior end of medial fork or slightly beyond; tip of M_4_ about equidistant between tip of M_2_ and tip of CuA. Sternite 2 with a strong pair of bristles medially on distal half.

This is a large genus of Mycetophilidae, with almost 150 described species, but with a large number of undescribed species in almost any tropical forest of the planet. Indeed, the actual number of species of *Epicypta* may be much higher. In some localities there are 15-30 described species of the genus *Manota* in sympatry—as, e.g., in Costa Rica (Jaschhof & Hippa 2005), Peru (Hippa et al., 2017), southern Atlantic Forest (Kurina et al. 2018) or Malaysia (Hippa 2006)—, while we consistently find 30 to 40 species of *Epicypta* in sympatry in different tropical areas in the world, often with a complete turnover between localities. So far, the most species-rich region for *Epicypta* is the Neotropical, with 60 described species. There are certainly over one thousand undescribed species of *Epicypta* worldwide.

Lane (1954) created the subgenera *Epicypta* (*Boscmyia*) Lane and *Epicypta* (*Callimyia*) Lane in *Epicypta*, including only Neotropical species in these two subgenera. These names will probably be helpful for dividing *Epicypta* into several genera or subgenera. This will, however, require a study of the genus at a global scale.

A total of 45 species of *Epicypta* have been described from the Oriental region, of which nine are from India or Sri Lanka, 28 from China and eight from Southeast Asia. The Southeast Asian species are *Epicypta angusticollaris* (Edwards), *Epicypta boettcheri* (Edwards) and *Epicypta flavidula* (Edwards)—from the Philippines (see Edwards, 1929)—, *Epicypta pallida* (Edwards)—from Krakatau (see Edwards, 1927)—, *Epicypta sartrix* (Meijere), *Epicypta flavicauda* (Edwards) and *Epicypta leefmansi* (Edwards)— from Java (see, respectively, de Meijere, 1924 and Edwards, 1935)—, *Epicypta borneensis* (Edwards)—from Borneo (see Edwards, 1933)—, and *Epicypta* LJ*umatrensis* (Edwards), from Sumatra (see Edwards, 1931).

We carefully compared our species with the original descriptions of all Southeast Asian species. The species from Singapore do not fit any of them. We were particularly careful about the species described from Sumatra, Java and Borneo, but for all there are significant differences in the color of the coxa, thorax and abdomen.

The haplotype network for the genus (Figs. 111–112) is complex and the species delimitation approach for the species of the genus show conflicts for some of the species. There are four more complex cases—*Epicypta jennylauae* Amorim & Oliveira, **sp. N.,** *E. chezaharaae* Amorim & Oliveira, **sp. N.,** *E. janetyeeae* Amorim & Oliveira, **sp. N.,** and *E. purchoni* Amorim & Oliveira, **sp. N.**—, that deserve more attention. They are discussed under “remarks” after each species description. Some of the species with a large number of specimens would benefit from a population-level analysis, which is, however, beyond the scope of this paper.

*Epicypta constancesingamae* Amorim & Oliveira, sp. n. (Figs. 113A–G)

urn:lsid:zoobank.org:act:6A38734A-7B28-49AE-8EE0-81A38F530E4F

**Diagnosis**. Head ochre-yellowish. Scutum with three dark brown longitudinal bands connected to each other over ochre-yellow background, scutellum dark brown. Pleural sclerites ochre-yellowish except for a diffuse brown mark on antepronotum, dark brown paratergite, a dorsal brown mark on mesepimeron, brown laterotergite and dark brown mediotergite. Hind coxa with a brown band across proximal end of hind coxa. Wing membrane light brownish, darker on cells c and br. C barely produced beyond tip of R_5_; M_1+2_ 0.78× r-m length. Dorsal macrotrichia on posterior veins M_1_ and M_2_, M_4_, and on CuA close to tip; macrotrichia on anal lobe membrane. Abdominal tergites 1-2 dark-brown, tergite 3 light brown posteriorly, yellowish-brown at anterior fourth, tergite 4 yellowish anteriorly and laterally, a large medial brown mark posteriorly, tergite 5 dark brown with a slender yellowish band along anterior margin, tergite 6 dark brown, tergite 7 yellowish-brown. Male terminalia gonocoxite short, no lobes projecting beyond base of gonostylus; gonostylus digitiform, long; aedeagus with a pair of separate tubular extensions with independent gonopores; parameres with a pair of long lateral projections, each with a pair of distal setae; tergite 9 with a pair of long, digitiform separate extensions. Female terminalia sternite 8 posterior margin with no incision, distal part of cercus short.

***Epicypta_constancesingamae_ZRCBDP0137084_hapZR CBDP0137084_SMH_holotype* [25: T, 125: T, 128: T, 1 30: T, 181: T, 194: C, 280: G]**

tctttcttctactattgctcatgcTggctcttcagttgatttagctattttttctttacatttagctggtatttcttctattttaggagctattaattttattactacaattattaatatgcgaTctTcTggaattacttttgatcgaatacctttatttgtttgatcagttttaattacTgcagttttattaCttttatctttacctgtattagcaggagctattactatattattaacagatcgaaatttaaatacttctttttttgatccagcgggGggaggggatcctattttataccaacatttattt

#### Description

**Male**. Wing length, 2.18 mm, width, 0.73 mm. **Head**. Head ochre-yellowish. Scape and pedicel dark ochre-yellowish, flagellomeres light brown. Face and clypeus yellowish-brown. Basal three palpomeres dark yellowish-brown, distal two palpomeres lighter, labella light ochre-yellowish. Occiput with two longer setae dorsally to eye anteriorly to lateral ocellus, one seta dorsally, and six posteriorly. Scape 2.4× pedicel length; flagellomere 4 2.0× longer than wide. Maxillary palpomere 4 1.6× palpomere 3 length; palpomere 5 1.6× palpomere 4 length. **Thorax** (Fig. 113A). Scutum background color ochre-yellow, with three dark brown bands connected to each other; scutellum dark brown. Pleural sclerites ochre-yellowish except for a diffuse brown mark on antepronotum dorsally, dark brown paratergite, a dorsal brown mark on mesepimeron, brown laterotergite and dark brown metepisternum and mediotergite. Haltere with light brown pedicel, whitish knob with light brownish tinge. Pleural membrane yellowish. Scutum with no bristles except for 9+2 supra-alars and three pairs of prescutellar bristles; three pairs of scutellar bristles. Proepisternum with two long bristles, anepisternum with four bristles along posterior margin. Mesepimeron with two bristles and 38 fine small setae, laterotergite with four bristles, one long seta and eight fine setae. Metepisternum with 11-15 small setae. **Legs**. Fore coxa yellowish-brown, mid and hind coxae whitish with a brownish tinge, a brown band across proximal-posterior corner of hind coxa. Femora yellowish with a brownish tinge. Tibiae and tarsi brownish-yellow. Mid coxa with a band of fine setae across basal fifth, hind coxa with fine setae on basal fourth. Fore tibia with a single bristle medially on distal third of outer face, two dorsal and two ventral rows of dark brown trichia; mid tibia with two irregular dorsolateral rows of 3-5 bristles and three bristles along ventral edge, trichia along dorsal and ventral edges dark brown; hind tibia with two irregular rows of 5–6 bristles dorsolaterally, dorsal and dorsolateral rows of trichia dark brown. Fore leg tarsomere 1 0.9× tibia, 1.4× tarsomere 2 length. **Wing** (Fig. 113B). Membrane fumose light brown, slightly darker along anterior margin. C extending slightly beyond apex of R_5_; R_1_ reaching C on wing distal fourth; R_5_ reaching slightly beyond level of M_2_. First sector of Rs slightly oblique, 0.34× r-m length; r-m more or less oblique. M_1+2_ 0.78× of r-m length; bM 3.6× r-m length; first sector of CuA about 0.26× length of second sector of CuA. Cubital pseudovein absent, CuP extending to slightly beyond level of origin of M_4_. Anal fold gently curved along its length. Posterior wing veins M_1_and M_2_ with dorsal macrotrichia along most of their length, M_4_ on distal half, and CuA at distal fourth; anal lobe membrane with dorsal macrotrichia. **Abdomen**. Abdominal tergites 1-2 dark-brown, light brown laterally, tergite 3 light brown posteriorly, yellowish-brown at anterior fourth, tergite 4 yellowish anteriorly and laterally, with a large medial brown mark posteriorly, tergite 5 dark brown with a slender yellowish band along anterior margin, tergite 6 dark brown, tergite yellowish-brown; sternites 1-7 light yellowish-brown. Sternite 2 with a ventral pair of brown bristles, tergites 2-6 with long, darker setae medially. **Terminalia** (Figs. 113C–E). Yellowish. Gonocoxites fused medially, no suture present, bare ventrally, no dorsolateral lobes extending beyond insertion of gonostylus. Gonostylus long, inserted laterodistally on gonocoxite, digitiform, with fine setae on inner margin along basal third, setae on outer face along entire length, distal setae longer, extending beyond tip of tergite 9 digitiform projections. Aedeagus with a wide subquadrate plate articulating to gonocoxites laterally, bearing medially a pair of long digitiform laterodistal projections extending way beyond tip of aedeagus. Parameres triangular, in a more dorsal position, with a pair of sub-medial digitiform processes, with setulae and an elongate spine at tip, and a digitiform short medial projection with some fine setae at tip. Gonocoxal bridge with a pair of elongate apodemes directed inwards anteriorly. Tergite 9 present as a pair of long posterior digitiform extensions, wide, meeting each other at anterior half, covered with long setae, almost as long as gonostylus, with 4–5 long, curved setae distally. Cerci not visible.

**Female** (Fig. 113A). As male, except for the following. **Wing l**ength, 2.13–2.56; width, 0.82–0.93. **Terminalia** (Figs. 113F–G). Sternite 8 wide, rectangular, posterior margin straight, no medial incision, no lateroposterior projections, with scattered microtrichia and fine setae. Sternite 9 wide, anterior apodeme extending beyond anterior end of terminalia, genital chamber slender, well-sclerotized. Tergite 8 wide, covered with microtrichia and setulae. T9+10 bare, slender. Cercomeres 1 and 2 probably fused, no sign of suture, basal ⅔ large, ovoid, slender on distal third, covered with microtrichia and fine setae.

#### Material examined

**Holotype**: male, ZRCBDP0137084, Bukit Timah Forest (BT05, 29-Mar-17, MIP leg. (slide-mounted). **Paratypes** (3 females): ZRCBDP0074040, Pulau Ubin (PU18), mangrove, 10.may.18, MIP leg.; ZRCBDP0136995, Bukit Timah, maturing secondary forest (BT06), 12-Jul-17, MIP leg. (extracted, slide-mounted), ZRCBDP0143114, Nee Soon (NSM1), 21.jan.15, MIP leg.

**Sequence failure specimens**: ZRCBDP0278248, Pulau Ubin (PU18), mangrove, 31.may.18, MIP leg. (slide-mounted). ZRCBDP0040808; ZRCBDP0040813; ZRCBDP0040832; ZRCBDP0041086; ZRCBDP0041099; ZRCBDP0041101; ZRCBDP0041134; ZRCBDP0067260; ZRCBDP0067277; ZRCBDP0279215, Mandai Mangroves (MM05), 22.may.18, MIP leg. .

**Etymology**. The species epithet of this species honors Constance Singam (née D’Cruz 1936-). Born in Singapore, she is a writer and activist for women’s rights, migrant worker rights, and rape victims. She served as the president of Association of Women for Action and Research (AWARE) over three non-contiguous periods, and as president of the Singapore Council of Women’s Organizations (SCWO) for two years. In 2015, Constance was inducted into the Singapore Women’s Hall of Fame.

**Remarks**. There are four haplotypes for *Epicypta constancesingamae* Amorim & Oliveira, **sp. n.,** all kept as a single species by all delimitation approaches.

*Epicypta jennylauae* Amorim & Oliveira, sp. n. (Figs. 114A–D, 115A–B)

https://singapore.biodiversity.online/species/A-Arth-Hexa-Diptera-000728

urn:lsid:zoobank.org:act:A27ED5D4-732E-4AA0-BE3D-73F53052FF88

**Diagnosis**. Head yellowish, antennal scape and pedicel whitish-yellow, first flagellomere ochre-yellow, flagellum distally light greyish-brown. Scutum blackish-brown with a light ochre-yellowish collar along anterior fifth; antepronotum and proepisternum ochre-yellowish, katepisternum, anepisternum, and mesepimeron light ochre-brown; laterotergite, mediotergite, and metepisternum brown. Coxae and femora whitish, yellowish tinge on anterior leg, hind coxa with a brown mark at basal end. Wing membrane light brownish, area along anterior margin slightly darker. C ending at tip of R_5_; M_1+2_ shorter than r-m. Dorsal macrotrichia on posterior veins M_1_, M_2_, posterior half of M_4_, and distal end of CuA, anal lobe membrane with macrotrichia. Abdominal tergite 1 brown, tergites 2–3 caramel-brown medially, with ochre-yellowish margins, tergites 4–7 mostly caramel-yellowish, terminalia whitish-yellow. Female terminalia sternite 8 quite straight along posterior margin, cercus slightly reniform.

***Epicypta_jennylauae_ZRCBDP0278324_hapZRCBDP02 78161_SMH_holotype* [29: T, 125: T, 127: A, 128: T, 13 0: T, 149: T, 169: T, 208: A, 247: T] -**

ttatcttctacaattgctcatgcaggaTcatcagtagatttagctattttttctcttcatttagctggtatttcttctattttaggagctattaattttattacaacaattattaatatacgtTcATcTggaattacttttgatcgaTtaccattatttgtttgatcTgtattaattacagctattttattattattatcattaccAgtattagccggagctattaccatattattaacagatcgTaatttaaatacatctttttttgacccagcgggaggaggagacccgattttataccaacatttattt

#### Description

**Male**. Wing length, 1.98–2.18; width, 0.70–0.83 (n=2). **Head** (Fig. 114B). Head ochre-yellowish. Face and clypeus whitish-yellow. Scape and pedicel light ochre-yellowish, flagellomere 1 ochre-yellowish, remaining flagellomeres greyish-yellow. Basal palpomeres ochre-yellowish, distal two palpomeres whitish-yellow, labella light yellowish-brown. Occiput with three longer setae dorsally to eye anteriorly to lateral ocellus, two dorsally to ocellus, six posteriorly to ocellus. Scape 2.5× length of pedicel, flagellomere 4 1.4× longer than wide, covered with scattered setulae. Flagellomere 4 slightly longer than flagellomere 3, flagellomere 5 1.6× longer than flagellomere 4. **Thorax**. Scutum anterior fifth ochre-yellow, remaining dark brown, scutellum blackish-brown, with ochre-yellow anterolateral corners. Antepronotum, proepisternum, proepimeron, katepisternum, anterior ⅔ of anepisternum, and anterior half of mesepimeron ochre-yellow, other pleural sclerites greyish-brown, mediotergite dark greyish-brown, proepisternum with a diffuse greyish-brown area on dorsal half. Haltere whitish, no larger setae. Pleural membrane yellowish. Scutum with six long supra-alar setae, three pairs of prescutellar bristles; two pairs of scutellar bristles. Three bristles on proepisternum directed ventrally, anepisternum with four bristles along posterior margin. Mesepimeron with two bristles and seven small setae, laterotergite with two bristles and three smaller setae. Metepisternum with nine small setae. **Legs**. Coxae whitish, hind coxa with a brown band at dorsal sixth; femora, tibiae and tarsi light whitish-yellow, tarsi darker. Mid coxa with a band of small setae across basal fifth, hind coxae basal fourth with small setae. Fore tibia with a single dorsal strong seta medially; mid tibia with a pair of dorsolateral rows of 4–5 bristles along entire length, distal ones curved, and two ventral bristles, besides bristles at tip, hind tibia with two rows of dorsolateral bristles, no bristles ventrally. Mid and hind tarsomeres 1–3 with rows of ventrolateral longer setae, besides setae at tip. Fore leg tarsomere 1 1.2× tibia length, 1.6× tarsomere 2 length. Hind tibial outer spur almost 6× tibia width at apex. **Wing**. Membrane fumose light brown, slightly darker along anterior margin. Sc barely produced. C ending at tip of R_5_. R_1_ long, reaching C on distal fifth of wing; R_5_ reaching C before level of tip of M_1_. First sector of Rs slightly oblique, 0.92× r-m length; r-m almost longitudinal. M_1+2_ about as long as r-m; bM 7.0× r-m length. First sector of CuA about 0.27× length of second sector. Cubital pseudovein absent, CuP extending slightly beyond level of origin of M_4_. Anal fold almost reaching wing margin, bare. Dorsal macrotrichia on posterior veins M_1_ and M_2_ entire length, on M_4_ distal third, and tip of CuA. Dorsal macrotrichia on anal lobe membrane. **Abdomen**. Abdominal tergite 1 brownish, tergite 2 greyish-brown lighter laterally, tergites 3–7 dark ochre-yellow, tergites 3– 5 ochre-brownish medially; sternites 1-7 whitish-yellow, Sternite 2 with a pair of strong brown bristles. **Terminalia** (Figs. 115A–B). Yellowish. Gonocoxites fused medially, suture absent, bare ventrally, no dorsolateral gonocoxite lobes extending beyond insertion of gonostylus. Gonostylus inserted laterodistally on gonocoxite, digitiform, reaching level of tip of aedeagus, setae on outer face along entire length, distal setae slightly longer. Aedeagus with a wide subquadrate plate medially, opening at tip of a pair of long, digitiform laterodistal projections distally. Paramere with a triangular plate in a more dorsal position, medially with a short digitiform projection with some fine setae at tip and a pair of spines. Tergite 9 present as a pair of long posterior extensions, anterior half wider, close to each other at tip, longer than gonostylus, covered with long setae, distal setae longer. Cerci not visible.

**Female** (Fig. 114A). As male, except for the following.

**Head** (Fig. 114C). **Wing** (Fig. 114B). Length, 1.82; width, 0.67 (n=2). **Terminalia** (Fig. 114D). Whitish-yellow. Sternite 8 rectangular, elongate, posterior margin straight, no lobes or incision, microtrichia and fine setae covering entire sclerite, longer setae at posterior margin, lateral ones curved inwards. Sternite 9 with wide genital chamber, lined with microtrichia, gonopore connected to two gonoducts, anterior arm weakly sclerotized. Tergite 8 trapezoid, wide at base and more slender towards posterior margin, lateroanterior extensions sclerotized. T9+10 bare, with a pair of short lateral lobes connected medially by a slender, sclerotized band. Cercomeres 1 and 2 probably fused, no sign of suture, laterally compressed, wider midway to apex than at base, covered with microtrichia and elongate setae, distally with two stronger setae.

#### Material examined

**Holotype**: male, ZRCBDP0278324, Singapore, 03.may.18, (slide-mounted). **Paratypes** (1 male, 1 females). **Male**: ZRCBDP0284244, Singapore, date range 2012-2018, MIP leg. (slide-mounted) (MZUSP). **Females**: ZRCBDP0278161, Pulau Ubin (PU18), mangrove, 10.may.18, MIP leg. (slide-mounted). **Possibly non-conspecific cluster** (16 females): ZRCBDP0047842, Nee Soon (NS1), swamp forest, 24.apr.13, MIP leg. (slide-mounted); ZRCBDP0048438, Nee Soon (NS1), swamp forest, 23.may. 12, MIP leg. (website photo specimen); ZRCBDP0048439, Nee Soon (NS2), swamp forest, 16.may. 12, MIP leg. (website photo specimen); ZRCBDP0048702, Nee Soon (NS1), 13.may.15, MIP leg. (slide-mounted); ZRCBDP0048797, Nee Soon (NS1), 18.mar.15, MIP leg. (slide-mounted); ZRCBDP0048863, Nee Soon (NS1), 10.dec.14, MIP leg.; ZRCBDP0048865, Nee Soon (NS1), 10.dec.14, MIP leg. (slide-mounted); ZRCBDP0048918, Nee Soon (NS1), 31.dec.14, MIP leg.; ZRCBDP0072746, Bukit Timah, old secondary forest (BT03), 22.dec. 16, MIP leg.; ZRCBDP0072747, Bukit Timah, maturing secondary forest (BT06), 08.dec. 16, MIP leg.; ZRCBDP0133449, Singapore, date range 2012-2018, MIP leg. (slide-mounted); ZRCBDP0133905, Singapore, date range 2012-2018, MIP leg.; ZRCBDP0133954, Singapore, date range 2012-2018, MIP leg.; ZRCBDP0133976, Singapore, date range 2012-2018, MIP leg.; ZRCBDP0136997, maturing secondary forest (BT06),12.jul. 17, MIP leg. (imaged); ZRCBDP0048439.

**Etymology**. The species epithet honors Jenny Lau Buong Bee (1932-2013). Born in British Malaya, she became in 1960 the first woman to serve as a magistrate in Malaya and in 1966 and was the first woman to be appointed a district judge in Singapore. She was inducted into the Singapore Women’s Hall of Fame in 2014.

**Remarks**. Two clusters are separated at 4.49%—a species delimitation at this level finds support only with OC at 5% p-distance. Unfortunately, we only have females for the larger subcluster and the female-only cluster may eventually be revealed to be a different species. We do want to not recognize this subcluster as a separate species without examining males, the specimens of this subcluster are not included here as paratypes.

*Epicypta limchiumeiae* Amorim & Oliveira, sp. n. (Figs. 116A–F)

https://singapore.biodiversity.online/species/A-Arth-Hexa-Diptera-000721and-002099

urn:lsid:zoobank.org:act:7FEE70EC-3E88-4B5B-BB88-A00A6CCDE040

**Diagnosis**. Head yellowish, antennal scape, pedicel and first flagellomere ochre-yellowish, rest of flagellum light brownish. Anterior third of scutum yellowish, blackish-brown on posterior two-thirds, a slender yellowish transverse band at posterior end, scutellum mostly blackish brown; antepronotum and proepisternum yellowish, other pleural sclerites blackish-brown, katepisternum and mesepimeron more greyish. Fore coxa and femur cream-yellow, mid and hind coxae and femora whitish. Wing with a large brownish area along anterior margin. C extending shortly beyond tip of R_5_; M_1+2_ about as long as r-m. Dorsal macrotrichia on M_1_, M_2_, and distal half of M_4_, some few macrotrichia on anal lobe membrane. Abdominal tergite 1 dark brown, tergites 2–5 dark brown medially, yellowish laterally, extension of yellowish band variable, larger on tergite 4; tergite 6 with a dark brown anterior band, yellowish on most of tergite; tergite 7 yellowish. Female terminalia cercus short, widened subapically.

***Epicypta_limchiumeiae_ZRCBDP0047890_hapZRCBD P0047890_SMH_holotype* [46: G, 157: G, 163: C, 187: G, 194: T, 223: A]**

tctttcatctacaattgctcatgcaggttcttctgttgatttagcGattttttcccttcatttagcaggtatttcttcaattttaggagcaattaattttattacaactattattaatatgcgagcccccggcatttcttttgatcgaatacccttGtttgtCtgatcagtattaattacagctgtGttattaTtattgtctctcccagttttagcgggggcAattacaatattattaacagatcgaaatttaaatacatctttctttgatcccgcaggagggggagacccaattttataccaacatttattt

#### Description

**Female** (Figs. 116A–C). Wing length, 2.11; width, 0.83 mm. **Head** (Fig. 116D). Length 1.9× width in lateral view. Vertex ochre-yellowish, occiput light ochre-yellow. Fine, short interommatidial setae over entire eye surface. Scape and pedicel ochre, ochre-yellow flagellomeres, more brownish towards apex of flagellum. Frons, face and clypeus light ochre-yellow, palpomeres 1–3 yellowish-ochre, distal three flagellomeres whitish-yellow, labella whitish-yellow. Occiput dorsally to eye with a row of two stronger setae anteriorly to lateral ocellus, one above ocellus and eight posteriorly to ocellus. Scape 1.6× pedicel length, flagellomere 4 1.7× width. Palpomere 4 1.6× palpomere 3, palpomere 5 1.4× palpomere 4 length, palpomeres 3 and 4 gently projected distally beyond base of next palpomere. **Thorax**. Scutum yellowish on anterior third, dark brown on posterior two-third, light ochre-yellow laterally along posterior margin, scutellum blackish-brown medially, light ochre-yellow on anterior margin. Antepronotum, proepisternum, and proepimeron ochre-yellow, anepisternum and katepisternum greyish-brown, lighter along anterior margin, mesepimeron, laterotergite, and metepisternum blackish-brown. Mediotergite dark brown. Haltere whitish. Pleural membrane yellowish on anterior third, brown on posterior two-thirds. Seven longer supra-alar setae on area bulging beyond level of wing base, three pairs of prescutellar bristles. Antepronotum with two bristles directed ventrally. Anepisternum with two long setae medially and four brown bristles along posterior margin. Katepisternum strongly compressed, bare. Mesepimeron with two bristles and 17 fine setae along dorsal margin, laterotergite with four bristles and eight setulae or fine setae. Metepisternum with 12 fine, elongate setae. **Legs**. Coxae whitish, fore coxa with light brownish tinge. Fore femur and tibia light brownish-yellow, tarsomeres darker. Mid and hind femora and tibiae light brownish-yellow, and tarsi light ochre-yellow. Mid coxa with four long setulae near anterior end, hind coxa with a large band of long setulae on proximal third. Fore leg tarsomere 1 1.3× tibia length, 1.7× of tarsomere 2. Hind tibial spurs about 5× tibia width at apex. **Wing** (Fig. 116E). Membrane light brown fumose with a wide dark brown band along anterior margin from anterior third of cell br to distal fifth of R_1_, medially to M_2_, a brownish isolated mark between CuA and anal fold. C extending slightly beyond tip of R_5_; Sc barely produced; cell br slightly wider midway to apex. R_1_ reaching C on distal sixth of wing; R_5_ reaching C slightly before tip of M_1_. First sector of Rs slightly oblique, about half r-m length; r-m almost longitudinal. M_1+2_ slightly shorter than r-m; bM over 6× r-m length; first sector of CuA about 0.34× length of second sector of CuA; cubital pseudovein not even reaching level of origin of M_4_; CuP barely reaching origin of M_4_. Anal fold almost reaching wing margin. Dorsal macrotrichia on posterior veins M_1_, M_2_, and distal half of M_4_; anal lobe membrane with some few dorsal macrotrichia. **Abdomen**. Abdominal tergite 1 dark brown, tergites 2-6 blackish-brown, with different extensions of yellow-ochre marks laterally, tergite 4 mostly yellowish, tergite 5 blackish-brown mark almost reaching lateral end along posterior margin, tergite 6 brown only on anterior fourth, tergite 7 yellow; sternites 1-7 whitish, Sternite 2 with a pair of strong brown bristles. **Terminalia** (Fig. 116F). Yellowish. Sternite 8 large, with a pair of long lateroposterior lobes widely separated, rounded posterior incision deep, sclerite mostly covered with microtrichia and elongate setae, longer along posterior margin, anterior end ventral face less sclerotized and covered only with microtrichia. Sternite 9 with long anterior projection extending to anterior end of terminalia, genital chamber wide, laterodistal margin close to tip with some setae. Tergite 8 large, trapezoid, covered with setae and microtrichia, posterior margin straight. Tergite 9+10 short, slender, with sclerotized band along anterior margin extending anteriorly to lateral apodemes directed anteriorly, entirely bare of setae. Cercomeres 1 and 2 probably fused, cerci elongate, wider at base and slender towards apex.

**Male**. Unknown.

#### Material examined

**Holotype**: female, ZRCBDP0047890, Nee Soon (NS1), swamp forest, 20.mar.13, MIP leg. (slide-mounted). **Paratypes** (5 females): ZRCBDP0047958, Pulau Ubin (PU2), mangrove, 01.apr.13, MIP leg.; ZRCBDP0048320, Nee Soon (NS2), swamp forest, 28.nov. 12, MIP leg. (website photo specimen); ZRCBDP0048321, Nee Soon (NS2), swamp forest, 04.apr. 12, MIP leg. (website photo specimen); ZRCBDP0048701, Nee Soon (NS1), 13.may.15, MIP leg.; ZRCBDP0049010, Nee Soon (NS2), 17.dec.14, MIP leg.

**Etymology**. The species epithet honors Janet Lim Chiu Mei (1923-2014). Born in Hong Kong and having grew up in Guangdong, she became the first Asian hospital matron at the St Andrews Mission Hospital in 1954. Her autobiography, “Sold for Silver”, was the first English-language book by a Singaporean author, published in 1958. She was inducted into the Singapore Women’s Hall of Fame in 2014.

**Remarks**. This species is similar to some extent with three of the *Epicypta* species from Java, especially *Epicypta flavicauda* (Edwards, 1935). The similarities include, besides the wide anterior yellow band on the scutum, the transverse yellow band anteriorly to the scutellum and the wing dark mark along anterior margin medially. There are consistent differences, however, in the abdominal markings. *E. limchiumeiae* Amorim & Oliveira, **sp. n.** is also similar to *E. sartrix* (de Meijere), but does not present the wing markings. We have four haplotypes for this species and no delimitation conflicts.

*Epicypta janetyeeae* Amorim & Oliveira, sp. n. (Figs. 117A–F)

https://singapore.biodiversity.online/species/A-Arth-Hexa-Diptera-000816

urn:lsid:zoobank.org:act:7D437627-29FB-430F-8543-412324F6483E

**Diagnosis**. Head, thorax, and abdomen blackish-brown, antenna brownish, lighter on scape, pedicel and first flagellomere. Coxae whitish, except for brown band on proximal end of hind coxa; femora whitish, except for proximal and distal ends of mid femur, and for a brown line along dorsal edge of hind femur. Wing membrane light brownish, area along anterior margin slightly darker. C extending shortly beyond tip of R_5_; M_1+2_ shorter than r-m. Dorsal macrotrichia on veins M_1_, M_2_, and posterior half of M_4_, anal lobe membrane with macrotrichia. Gonocoxites without posterior projections beyond base of gonostylus; gonostylus flat, weakly sclerotized; paramere with a pair of strong spines; tergite 9 with a pair of long parallel projections. ***Epicypta_janetyeeae_ZRCBDP0048806_hapZRCBDP00 47948_SMH_holotype* [22: C, 67: T, 128: G, 129: T, 140 : C, 212: C, 214: T, 298: T]**

tctatcttcaactattgctcaCgcaggagcatctgttgatttagctattttttctcttcatttagcTggtatttcttctattttaggagctattaattttattacaactattattaatatacgatctGTaggaattactCttgatcgaatacctttatttgtttgatcagtattaattactgctattcttttattattatctttacctgttCtTgcaggagccattactatattattaacagatcgaaatttaaatacttctttctttgatcctgcagggggaggagaccctattctTtatcaacatttgttt

#### Description

**Male** (Fig. 117A). Wing length, 1.76–1.98; width, 0.67–0.72 (n=3). **Head**. Head compressed antero-posteriorly, 1,8× longer than wide, vertex large, facing anteriorly. Vertex dark brown. Occiput brown, lighter brown towards ventral margin. Frons, face, and clypeus light brown, maxillary palpus yellowish-brown, distal palpomeres lighter, labella whitish-yellow. Antennal scape and pedicel light brown, first two flagellomeres ochre-brown, remaining flagellomeres brown. Eye small, brownish. Mid ocellus absent, ocelli touching eye margin. Vertex and occiput densely covered of small setae ventrally to level of insertion of antenna, one bristle close to margin of foramen at level of dorsal end of eye, three strong setae at level of mid of eye. Scattered brownish setulae over entire vertex, two strong setae on occiput dorsally to eyes anteriorly to ocellus, two above ocellus and five posteriorly to ocellus. Eye entirely covered with conspicuous interommatidial setae. Frons with a long projection ventrally between antennae, bare along ventral margin, frontal furrow extending from ventral margin to level of ocelli. Face short, with a medial transverse depression, posterior margin slightly projected, fine setae on ventral half, clypeus short, slightly budging, with scattered short fine setae on ventral ¾. Scape and pedicel with dorsal a crown of darker setae at distal margin, one strong and some slightly smaller setae at dorsal end; scape 1.5× pedicel length; 14 flagellomeres, flagellomeres about 1.7× width, covered with scattered fine setulae. Maxillary palpomere 1 reduced to a small lobe with one seta directed posteriorly, palpomere 2 weakly sclerotized with some few setae, palpomere 3 with a conspicuous sensorial pit opening at internal face, no setae at internal face, palpomere 4 1.1× longer than palpomere 3, with a short distal projection beyond insertion of palpomere 5, no setae at internal face, palpomere 5 1.2× length of palpomere 4, setulae only at distal end. Labella developed backwards. **Thorax** (Fig. 117B). Scutum and scutellum dark brown. Antepronotum and anepisternum brown, other pleural sclerites dark brown except for dorsal third of laterotergite, brownish-yellow. Mediotergite dark brown on dorsal half, light brown on ventral half. Pleural membrane yellowish. Scutum with deep lateroanterior incision into which antepronotum fits, no sign of parapsidal or transverse sutures. Scutum entirely covered only with short light brown setae except for five strong setae placed on a short bulging area above wing and two pairs of strong prescutellar bristles. Scutellum large, with rounded posterior border, two pairs of scutellar bristles in addition to small setae directed backwards covering almost entire surface of disc. Antepronotum lobes not connected to each other medially. Antepronotum lobe and proepisternum large, entirely fused, sign of a suture only at anterior margin, basisternum dorsoposterior arm fused to proepisternum at ventral half, basisternum arm with a group of small setae ventrally at distal end. Antepronotum and proepisternum covered with small setae, proepisternum with no bristles directed ventrally. Anepisternum rectangular, entirely covered with light brown setulae, four strong bristles directed posteriorly on posterior margin, anapleural suture present only on distal fifth. Anterior and posterior basalare bare. Katepisternum strongly compressed, at most half of anepisternum height, entirely bare. Mesepimeron reaching ventral margin of pleura, dorsoventrally compressed, with two bristles and 26 small setae and setulae close to dorsal margin; laterotergite bulging, strongly compressed, with one isolated bristle. Metepisternum slender, with 10 small fine setae, metepimeron not discernible. Mediotergite short, “folded”, with ventral margin displaced anteriorly, entirely bare. Haltere whitish-yellow, two small setae on pedicel, knob with a number of small setae dorsally. **Legs**. Fore coxa whitish-yellow with a light brown tinge, mid and hind coxae whitish, mid coxa dorsal end with a band of brownish tinge, hind coxa with a dark brown band at dorsal third and distally along posterior margin; femora whitish-yellow with a light brown tinge, mid and hind femora whitish, with a dark brown mark along dorsal edge, hind femur with basal fourth brown; tibiae and tarsi brownish-yellow, darker towards tip. Coxae largely developed, fore coxa strong, mid coxa wide at base, hind coxa wide on basal third. Forecoxa laterally compressed, entirely covered with small setae on anterior and external face, a row of brown elongate strong setae along distal end of margin posteriorly; mid coxa with a band of fine setae across basal fifth, a group of fine setae along frontal face and a row of bristles along tip at anterior face; hind coxa with fine setae on anterior third, some fine setae near tip at anterior face, a row of strong setae along distal margin of anterior face, one isolated strong seta laterally near distal end. Femora covered with fine setae, a row of strong setae near apex along ventral edge. Tibiae and tarsi with regular rows of trichia, fore tibia with a lateral seta externally midway to apex; mid tibia with two irregular rows of 4-5 bristles dorsolaterally, one external seta medially, a row of three setae ventrally and two long bristles and some setae at distal margin; hind tibia with two irregular rows of five bristles dorsolaterally and some setae at distal margin, besides a comb of fine setae on internal face at distal end. Anteroapical depressed area of fore tibia covered with setulae. Tarsomere 1 of fore leg as long as tibia, 1.6× tarsomere 2 length. Hind tibial spurs about 5× tibia width at apex, inner spur subequal to outer spur, mid and hind leg tarsomeres 1 and 2 with ventral setae in addition to rows of trichia. Tarsal claws on basal half with a short tooth with wide base. **Wing** (Fig. 117C). Membrane fumose light brown, slightly darker along anterior margin. Membrane densely covered with regularly organized microtrichia on all cells, margin gently emarginated at level of tip of CuA. Humeral vein present, oblique, Sc barely produced. C extending beyond tip of R_5_ for about one fifth of distance to M_1_. R_1_ long, reaching C on distal fifth of wing, R_4_ absent, R_5_ reaching C slightly beyond level of tip of M_1_. First sector of Rs slightly oblique, about half of r-m length; r-m almost longitudinal, well sclerotized. M_1+2_ short, slightly over half of r-m length; M_1_ and M_2_ well sclerotized, mostly parallel to each other, faint close to tip; bM long, over 6× r-m length; M_4_ running parallel to M_2_ on distal two-thirds. First sector of CuA short, M_4_ originating more basally than origin of M_1+2_, M_4_ and CuA complete, well sclerotized, reaching wing margin. Cubital pseudovein barely produced, CuP very short, not reaching level of origin of M_4_, anal fold well sclerotized, long, gently curved along its entire length, not reaching margin. Macrotrichia dorsally on trunk of Sc, bR, R_1_, second sector of Rs, r-m, distal end of M_1_, M_2_, and M_4_; ventrally on distal end of bR, R_1_, R_5_, r-m, and distal half of bM; anal lobe membrane with macrotrichia. **Abdomen**. Abdominal tergites 1-7 dark brown; sternites 1-7 light yellowish-brown, sternite 2 with a pair of strong bristles medially on distal half. Tergites with some longer setae mixed with fine, smaller setae. **Terminalia** (Figs. 117D– F). Yellowish-brown. Gonocoxites relatively small, medially fused on anterior end of terminalia, suture of fusion conspicuous, no sign of sternite 9, lateroposteriorly with a weakly sclerotized extension rounded distally and a number of concentrated elongate setulae, medioventral process extending into the terminalia towards aedeagus. Gonostylus reduced to a wide flat blade covering ventrodistal end of gonocoxite, with a large number elongate fine setae and a digitiform process distally on external end bearing a long fine seta, a strong long seta on ventral margin medially and one long seta on external face distally. Parameres largely developed, a subquadrate plate extending ventrally into a pair of falciform short lobes medially, curved almost meeting along medial line and a pair of laterodistal digitiform projections with an elongate fine seta at tip, besides a pair of L-shaped sclerotized structures distally bearing a fine seta at dorsal end distally, one pointed strong subapical seta, and a strongly developed seta apically on ventrodistal arm, no tubular aedeagus evident. Gonocoxal apodeme not evident. Tergite 9 with a pair of long laterodistal projections, exceeding tip of parameres, external faces of lobes covered with long setae, connected medially only at anterior end, lobes slightly widening distally with curved posterior corners.

**Female**. Wing length, 2.02–2.05; width, 0.75–0.80 (n=2). **Head**. Antennal scape, pedicel and flagellomeres brown. **Thorax**. Mesepimeron with three bristles and 29 small setae and setulae close to dorsal margin. Metepisternum slender, with 13 fine setae. **Wing**. Membrane with 12 dorsal macrotrichia on anal lobe membrane, two dorsal setae on anal fold. **Abdomen**. Abdominal tergites 1–7 dark brown; sternites 1-7 light yellowish-brown, sternite 2 with a strong pair of bristles medially on distal half. **Terminalia**. Yellowish-brown. Sternite 8 wide at base, trapezoid, distal end with a shallow medial incision, fine setae with wide distribution along anterior margin, restricted medially close to posterior margin, setae along posterior margin stronger, lateroposterior setae gently curved inwards, short labia projected medio-posteriorly. Sternite 9 anterior end of vaginal furca extending beyond anterior margin of sternite 8, medial end of sternite 9 posteriorly, reaching level of posterior ¾ of cercomere 1, with short setae along margin. Tergite 8 with sclerotized anterior margin, laterally with a pair of long projections directed anteriorly almost as long as anterior extension of sternite 9, medial connection slender, setae restricted to lateroposterior end, some strong setae at lateroposterior corner curved inwards, ventrally to cercus. Tergite 9+10 reduced to a slender short band with microtrichia. Only one cercomere present, cercomeres possibly fused, wider at base and longer at apex, about 4× longer than wide midway to apex, covered with microtrichia and fine setae, slightly longer at apex.

#### Material examined

**Holotype**: male, ZRCBDP0048806, Nee Soon (NS1), 18.mar.2015, MIP leg. (slide-mounted). **Paratypes** (39 males, 30 females). **Males**: ZRCBDP0066804, Bukit Timah, maturing secondary forest (BT08), 16.aug. 16, MIP leg.; ZRCBDP0066810, Bukit Timah, maturing secondary forest (BT08), 16.aug. 16, MIP leg.; ZRCBDP0066813, Bukit Timah, maturing secondary forest (BT08), 16.aug. 16, MIP leg.; ZRCBDP0072666, Bukit Timah, primary forest (BT05), 28.dec. 16, MIP leg. (slide-mounted); ZRCBDP0072466, Bukit Timah, maturing secondary forest (BT08), 01.dec. 16, MIP leg.; ZRCBDP0047948, Nee Soon (NS1), swamp forest, 03.jul.13, MIP leg.; ZRCBDP0047787, Nee Soon (NS2), swamp forest, 05.feb.14, MIP leg.; ZRCBDP0048462, Nee Soon (NS1), swamp forest, 23.may. 12, MIP leg.; ZRCBDP0047892, Nee Soon (NS1), swamp forest, 20.mar.13, MIP leg.; ZRCBDP0047862, Nee Soon (NS2), swamp forest, 15.jan.14, MIP leg.; ZRCBDP0048950, Nee Soon (NS2), 03.dec.14, MIP leg.; ZRCBDP0049253, Nee Soon (NS1), 10.dec.14, MIP leg.; ZRCBDP0049261, Nee Soon (NS1), 10.dec.14, MIP leg.; ZRCBDP0049006, Nee Soon (NS2), 17.dec.14, MIP leg.; ZRCBDP0048909, Nee Soon (NS1), 31.dec.14, MIP leg. (website photo specimen); ZRCBDP0048912, Nee Soon (NS1), 31.dec.14, MIP leg.; ZRCBDP0048888, Nee Soon (NS1), 31.dec.14, MIP leg.; ZRCBDP0048908, Nee Soon (NS1), 31.dec.14, MIP leg.; ZRCBDP0049021, Nee Soon (NS2), 07.jan.15, MIP leg.; ZRCBDP0049037, Nee Soon (NS2), 07.jan.15, MIP leg.; ZRCBDP0048819, Nee Soon (NS1), 14.jan.15, MIP leg.; ZRCBDP0048827, Nee Soon (NS1), 14.jan.15, MIP leg.; ZRCBDP0048847, Nee Soon (NS1), 14.jan.15, MIP leg.; ZRCBDP0048759, Nee Soon (NS1), 25.feb.15, MIP leg.; ZRCBDP0048792, Nee Soon (NS1), 18.mar.15, MIP leg.; ZRCBDP0048795, Nee Soon (NS1), 18.mar.15, MIP leg.; ZRCBDP0049136, National University of Singapore (PGP), 22.apr.15, MIP leg.; ZRCBDP0048771, National University of Singapore (PGP), 20.may.15, MIP leg.; ZRCBDP0049332, National University of Singapore (PGP), 10.jun.15, MIP leg.; ZRCBDP0047063, National University of Singapore (PGP), 08.jul.15, MIP leg.; ZRCBDP0072727, Bukit Timah, primary forest (BT05), 14.dec. 16, MIP leg.; ZRCBDP0072740, Bukit Timah, primary forest (BT05), 14.dec. 16, MIP leg.; ZRCBDP0066780, Bukit Timah, primary forest (BT05), 16.aug. 16, MIP leg. (slide-mounted) (MZUSP); ZRCBDP0066786, Bukit Timah, primary forest (BT05), 16.aug. 16, MIP leg.; ZRCBDP0066788, Bukit Timah, primary forest (BT05), 16.aug. 16, MIP leg.; ZRCBDP0066791, Bukit Timah, primary forest (BT05), 16.aug. 16, MIP leg.; ZRCBDP0066797, Bukit Timah, primary forest (BT05), 23.aug. 16, MIP leg.; ZRCBDP0066798, Bukit Timah, primary forest (BT05), 23.aug. 16, MIP leg.; ZRCBDP0047066, National University of Singapore (Uhall), 15.jul.15, MIP leg. **Females**: ZRCBDP0048430, Nee Soon (NS1), swamp forest, 25.apr. 12, MIP leg. (website photo specimen); ZRCBDP0048877, Nee Soon (NS1), 31.dec.14, MIP leg.; ZRCBDP0049267, National University of Singapore (PGP), 06.may.15, MIP leg.; ZRCBDP0049269, National University of Singapore (PGP), 06.may.15, MIP leg.; ZRCBDP0049329, National University of Singapore (PGP), 10.jun.15, MIP leg.; ZRCBDP0066803, Bukit Timah, primary forest (BT05), 23.aug. 16, MIP leg.; ZRCBDP0072739, Bukit Timah, primary forest (BT05), 14.dec. 16, MIP leg.; ZRCBDP0072743, Bukit Timah, primary forest (BT05), 14.dec. 16, MIP leg.; ZRCBDP0047778, Nee Soon (NS1), swamp forest, 28.aug.13, MIP leg.; ZRCBDP0047783, Pulau Semakau (SMN2), planted mangrove, 16.may.13, MIP leg.; ZRCBDP0047861, Nee Soon (NS2), swamp forest, 31.dec.13, MIP leg.; ZRCBDP0047934, Nee Soon (NS1), swamp forest, 17.apr.13, MIP leg.; ZRCBDP0047950, Nee Soon (NS1), swamp forest, 03.jul.13, MIP leg.; ZRCBDP0048461, Nee Soon (NS1), swamp forest, 06.feb.13, MIP leg.; ZRCBDP0048464, Nee Soon (NS2), swamp forest, 26.dec. 12-02.jan.13, MIP leg.; ZRCBDP0048766, Nee Soon (NS1), 25.feb.15, MIP leg.; ZRCBDP0048791, Nee Soon (NS1), 18.mar.15, MIP leg.; ZRCBDP0048811, Nee Soon (NS1), 14.jan.15, MIP leg.; ZRCBDP0048814, Nee Soon (NS1), 14.jan.15, MIP leg. (slide-mounted); ZRCBDP0048831, Nee Soon (NS1), 14.jan.15, MIP leg.; ZRCBDP0048834, Nee Soon (NS1), 14.jan.15, MIP leg.; ZRCBDP0048856, Nee Soon (NS1), 14.jan.15, MIP leg.; ZRCBDP0048897, Nee Soon (NS1), 31.dec.14, MIP leg.; ZRCBDP0048905, Nee Soon (NS1), 31.dec.14, MIP leg.; ZRCBDP0048921, Nee Soon (NS1), 31.dec.14, MIP leg.; ZRCBDP0048922, Nee Soon (NS1), 31.dec.14, MIP leg.; ZRCBDP0048983, Nee Soon (NS2), 17.dec.14, MIP leg.; ZRCBDP0049100, Nee Soon (NS1), 24.dec.14, MIP leg.; ZRCBDP0049127, Nee Soon (NS1), 24.dec.14, MIP leg.; ZRCBDP0049220, Nee Soon (NS1), 10.dec.14, MIP leg.

**Additional sequenced specimens**: male, ZRCBDP0132892, Singapore, date range 2012-2018, MIP leg.; female, ZRCBDP0133471, Singapore, date range 2012-2018, MIP leg.; female, ZRCBDP0132812, Singapore, date range 2012-2018, MIP leg.; female, ZRCBDP0132845, Singapore, date range 2012-2018, MIP leg.; female, ZRCBDP0132872, Singapore, date range 2012-2018, MIP leg.; female, ZRCBDP0133153, Singapore, date range 2012-2018, MIP leg.; female, ZRCBDP0133479, Singapore, date range 2012-2018, MIP leg.; ZRCBDP0132861, Singapore, date range 2012-2018, MIP leg.; ZRCBDP0132867, Singapore, date range 2012-2018, MIP leg.; ZRCBDP0133102, Singapore, date range 2012-2018, MIP leg.; ZRCBDP0133403, Singapore, date range 2012-2018, MIP leg.; ZRCBDP0137042, Singapore, date range 2012-2018, MIP leg.; ZRCBDP0278335, Singapore, date range 2012-2018, MIP leg. ZRCBDP0078965, Singapore; ZRCBDP0078966; Singapore; ZRCBDP0278336, Singapore, 22.may.28; ZRCBDP0278338, Singapore, 22.may.18; ZRCBDP0278339, Singapore, 22.may.18; ZRCBDP0278340, Singapore, 22.may.18; ZRCBDP0279132, Singapore, 26.apr.18; ZRCBDP0279135, Singapore, 26.apr.18; ZRCBDP0279157, Singapore. 05.jun.18; ZRCBDP0284234, Sungei Buloh (SB12); ZRCBDP0284235, Sungei Buloh (SB12); ZRCBDP0284236, Sungei Buloh (SB12); ZRCBDP0284237, Sungei Buloh (SB12); ZRCBDP0284238, Sungei Buloh (SB12); ZRCBDP0284291, Singapore, 27.jun.18; ZRCBDP0284292, Singapore, 27.jun.18; ZRCBDP0284294, Singapore, 27.jun.18; ZRCBDP0284295, Singapore, 27.jun.18; ZRCBDP0284304, Singapore, 07.may.18.

**Etymology**. The species epithet of this species honors Janet Yee (1934–2019). Born in Singapore, she is recognized as a pioneering social worker who campaigned to ensure that abandoned babies would be considered citizens and thus able to receive social services. For her advocacy of children’s rights, she was one of 11 women inducted to Singapore Women’s Hall of Fame in 2015.

**Remarks**. This is one of the most abundant mycetophilid species in Singapore and contains 15 haplotypes. The sequenced specimens have two main subclusters, with 4.21% divergence between them. We do have males belonging to each of these subclusters. There are hardly any differences in the male terminalia between the males of both subclusters. This is one of the grey-zone cases above 3% divergence in which there is no evidence in morphology that would allow subclusters to be placed in separate species.

*Epicypta kohkhenglianae* Amorim & Oliveira, sp. n. (Figs. 118A–D, 119A–B)

https://singapore.biodiversity.online/species/A-Arth-Hexa-Diptera-000728

urn:lsid:zoobank.org:act:196BA363-5A48-4A25-9D97-6563D1547605

**Diagnosis**. Head ochre-yellow, antennal scape and pedicel ochre-yellowish, first flagellomere ochre-brown, flagellum distally grey-brown. Scutum blackish-brown with an ochre-brown collar along anterior fourth; most pleural sclerites dark brown, antepronotum, proepisternum, katepisternum, anteroventral corner of anepisternum, and ventral half of mesepimeron ochre-yellow. Coxae and femora whitish with yellowish tinge. Wing membrane light brownish, area along anterior margin slightly darker. C extending shortly beyond tip of R_5_; M_1+2_ shorter than r-m. Dorsal macrotrichia on posterior veins M_1_, M_2_, posterior half of M_4_ and distal end of CuA, anal lobe membrane with macrotrichia. Abdominal tergite 1 brown, tergites 2–5 brownish, with yellow margins larger (larger on tergites 4–5), tergite 6 mostly yellowish with some greyish-brown tinge, tergite 7 yellowish, terminalia yellowish. Female terminalia cercus reniform. ***Epicypta_kohkhenglianae_ZRCBDP0072736_hapZRCB DP0047893_SMH_holotype* [85: G, 91: C, 148: T, 187: A, 268: C, 274: C, 308: C]**

tctttcttctactattgctcatgcaggatcttctgtagatttagctattttttctcttcatttagcaggtatttcttcaattttGggagcCattaattttattactactattattaatatacgagctcctggaattacttttgatcgTatacctttatttgtttgatcagtcttaattactgctgtActtttactattatctttacctgttttagcaggtgctattactatattattaacagatcgaaatttaaatacatcttttttCgatccCgcaggaggaggagaccctattttatatcaacatCtattt

#### Description

**Male**. **Head**. Head ochre-yellowish. Scape and pedicel light ochre-yellowish, scape twice pedicel length; flagellomere 1 ochre-yellow, remaining flagellomeres greyish-yellow, face and clypeus light brown. Basal three palpomeres dark ochre-yellowish, distal two palpomeres whitish-yellow, labella slightly lighter than head color. Occiput with two longer setae dorsally to eye anteriorly to lateral ocellus, one seta dorsally to ocellus, six setae posteriorly to ocellus. Flagellomere 4 length 2.0× width. **Thorax**. Scutum anterior fifth ochre-yellow, remaining dark brown, scutellum blackish-brown with ochre-yellow anterolateral corners. Antepronotum, proepisternum, katepisternum, anteroventral corner of anepisternum, and anterior half of mesepimeron ochre-yellow, other pleural sclerites greyish-brown, mediotergite light brown dorsally, proepisternum with a diffuse greyish-brown mark along dorsal half. Pleural membrane yellowish. Scutum with no bristles except for some supra-alars and prescutellars, two pairs of scutellar bristles. Proepisternum with three long bristles directed ventrally, anepisternum with four bristles along posterior margin. Mesepimeron with one long seta and seven smaller fine setae, laterotergite with two bristles and seven fine setae. Metepisternum with seven small fine setae. Haltere with light brown pedicel, knob whitish with light brownish tinge. **Legs**. Fore coxa whitish-yellow, mid and hind coxae whitish, hind coxa with a brown band at dorsal sixth; femora, tibiae, and tarsi light brownish-yellow, tarsi darker. Mid coxa with a slender band of fine setae across basal fifth, hind coxa with fine setae on basal third. Fore tibia with a single dorsal strong seta dorsolaterally on distal third of wing; mid tibia with a row of three to five bristles dorsolaterally, three bristles ventrally and one bristle laterally midway to apex; hind tibia with two irregular rows of three to five small bristles dorsolaterally and two bristles on a lateroventral row. Fore leg tarsomere 1 as long as tibia, 1.8× tarsomere 2 length, mid and hind tarsomeres 1–3 with ventral rows of stronger short setae. Tibial spurs light brown, about 4× tibia width at apex, inner and outer spurs subequal. **Wing**. Membrane fumose light brown, slightly darker along anterior margin. C extending beyond apex of R_5_ for one third of distance to M_1_. First sector of Rs slightly oblique, less than half of r-m length; r-m oblique. M_1+2_ about as long as r-m; bM 5× r-m length; first sector of CuA about 0.38× length of second sector of CuA. Cubital pseudovein weakly sclerotized, CuP hardly extending beyond level of origin of M_4_. Among posterior veins, M_1_and M_2_ with microtrichia dorsally on most of their length, M_4_ on distal half, and CuA on distal fourth, anal fold bare; dorsal macrotrichia on anal lobe membrane. **Abdomen**. Abdominal tergites 1-2 greyish-brown, tergite 3 greyish-brown medially, yellow at laterals, tergites 4-5 with a greyish-brown medial mark and wide ochre-yellowish areas laterally, tergite 6 dark ochre-yellowish, tergite 7 ochre-yellowish; sternites 1-7 whitish, Sternite 2 with a pair of strong brown bristles. **Terminalia** (Figs. 119A–B). Gonocoxite small, slender, entirely bare, fused, with no medial suture, not projecting beyond insertion of gonostylus. Gonostylus dorsoventrally compressed, displaced laterally, slightly curved inwards at apex, with microtrichia and setulae on outer basal half, fine setae on distal half. Gonocoxal apodemes short, directed inwards. Aedeagus well-sclerotized distally; parameres projected beyond tip of aedeagus, with some setulae on a triangular distal end. Tergite 9 with a pair of large, subtriangular lateroposterior lobes bearing microtrichia and setae, longer setae on posterior margin internally. Cerci small, weakly sclerotized, separate from each other.

**Female** (Fig. 118A). As male, except for the following. **Wing** (Fig. 118B). Length, 2.08–2.14, width, 0.77–0.80 (n=2). **Terminalia** (Figs. 118C–D). Whitish-yellow.

Sternite 8 subquadrate, posterior end with a pair of short lobes, tips widely separated, with a wide, short posterior incision, microtrichia and some setae, longer setae along posterior margin. Sternite 9 wide, anterior apodeme wide at posterior end, triangular on distal third, setulae along margins subdistally, genital chamber and gonopore well sclerotized, two gonoducts reaching gonopore. T8 short and wide, no laterodistal lobe, covered with microtrichia and long setae. T9+10 present as a pair of bare lateral lobes with a short, slender medial connection. Cercomeres 1 and 2 apparently fused, no suture, basal half wider, distal half slender, projected ventroposteriorly.

#### Material examined

**Holotype**: male, ZRCBDP0072736, Bukit Timah, primary forest (BT05), 14.dec. 2016, MIP leg. (slide-mounted). **Paratypes** (13 males, 10 females). **Males**: ZRCBDP0047790, Pulau Ubin (PU4), mangrove, 20.apr. 2013, MIP leg.; ZRCBDP0047893, Nee Soon (NS1), swamp forest, 20.mar. 2013, MIP leg.; ZRCBDP0047959, Pulau Ubin (PU2), mangrove, 01.apr. 2013, MIP leg.; ZRCBDP0048432, Nee Soon (NS2), swamp forest, 18.jul. 2012, MIP leg.; ZRCBDP0048433, Nee Soon (NS1), swamp forest, 04.apr. 12, MIP leg. (website photo specimen); ZRCBDP0048434, Nee Soon (NS1), swamp forest, 23.may. 12, MIP leg.; ZRCBDP0048754, Nee Soon (NS1), 25.feb. 2015, MIP leg. (slide-mounted) (MZUSP); ZRCBDP0048763, Nee Soon (NS1), 25.feb. 2015, MIP leg.; ZRCBDP0049187, Nee Soon (NS2), 13.may. 2015, MIP leg.; ZRCBDP0049241, Nee Soon (NS1), 10.dec. 2014, MIP leg.; ZRCBDP0049242, Nee Soon (NS1), 10.dec. 2014, MIP leg.; ZRCBDP0072682, Bukit Timah, primary forest (BT05), 22.dec. 2016, MIP leg. (slide-mounted); ZRCBDP0072706, Bukit Timah, primary forest (BT05), 01.dec. 2016, MIP leg. **Females**: ZRCBDP0047782, Nee Soon (NS2), swamp forest, 20.feb. 2013, MIP leg. (slide-mounted); ZRCBDP0047951, Nee Soon (NS1), swamp forest, 03.jul. 2013, MIP leg.; ZRCBDP0048825, Nee Soon (NS1), 14.jan. 2015, MIP leg.; ZRCBDP0048994, Nee Soon (NS2), 17.dec. 2014, MIP leg.; ZRCBDP0049101, Nee Soon (NS1), 24.dec. 2014, MIP leg. (slide-mounted); ZRCBDP0049260, Nee Soon (NS1), 10.dec. 2014, MIP leg.; ZRCBDP0066775,

Bukit Timah, maturing secondary forest (BT08), 16.aug. 16, MIP leg.; ZRCBDP0066808, Bukit Timah, maturing secondary forest (BT08), 16.aug. 16, MIP leg.; ZRCBDP0066809, Bukit Timah, maturing secondary forest (BT08), 16.aug. 16, MIP leg.; ZRCBDP0072741, Bukit Timah, primary forest (BT05), 14.dec. 2016, MIP leg. **Additional sequenced specimens**: ZRCBDP0134055; ZRCBDP0143094 (missing terminalia); ZRCBDP0048435, Nee Soon (NS1), 30.may,12.

**Etymology**. The species epithet honors Koh Kheng Lian (1937–). She is an internationally recognized expert in Environmental law, who led the Asia-Pacific Centre for Environmental Law, at the National University of Singapore, to become a leading institution for the study of environmental law. In 2012, Kheng Lian was awarded Stockholm University’s Elizabeth Haub Prize for Environmental Law for her pioneering contributions to the development of the field in Singapore and the ASEAN region. She was inducted into the Singapore Women’s Hall of Fame in 2014.

**Remarks**. There are nine haplotypes for *Epicypta kohkhenglianae* Amorim & Oliveira, **sp.n.**. collected in different environments, that come together using any of the species delimitation approaches.

*Epicypta daintoni* Amorim & Oliveira, sp. n. (Figs. 120A–D)

https://singapore.biodiversity.online/species/A-Arth-Hexa-Diptera-000737

urn:lsid:zoobank.org:act:623FA481-44CE-4C4B-8074-636EE20FEC65

**Diagnosis**. Head and scutum bright yellowish, antennal scape, pedicel and first flagellomere more yellowish, other flagellomeres more brownish. Scutum with a brown medioposterior mark over an ochre-yellow background; scutellum brown; pleural sclerites mostly yellowish, with more brownish areas on proepisternum, katepisternum, mesepimeron, and laterotergite; metepisternum and mediotergite brown. Coxae and femora whitish, an orangish tinge along dorsal and ventral femora crests. Wing membrane light brownish, cells c and br darker. C produced slightly beyond tip of R_5_; M_1+2_ about half of r-m length. Dorsal macrotrichia on posterior veins M_1_ and M_2_, M_4_, and on CuA close to tip, macrotrichia on anal lobe membrane. Abdominal tergites 1 brown, tergites 2–6 dark brown medially, with cream-yellow laterals, brown area on tergite 6 wider, tergite 7 brownish-yellow, terminalia yellowish. Female terminalia sternite 8 with no medial incision along posterior margin; cercus slender, slightly reniform.

***Epicypta_daintoni_ZRCBDP0048460_hapZRCBDP0048 460_SMH_holotype* [62: C, 67: G, 106: C, 181: C, 220: T, 274: C, 304: G, 313: C]**

tctttcttctactattgctcatgcagggtcttctgtagatttagctattttttctcttcatCtagcGggaatttcttctattttaggggcaattaattttattacCactattattaatatacgagccccaggaattacatttgatcgaatacctttatttgtatgatctgttttaattacCgctattcttctacttttatctttacctgtattagcaggTgctattacaatattattaacagatcgaaatttaaatacatctttttttgatccCgcaggaggaggagaccctattttatatcaGcatttattC

#### Description

**Female** (Fig. 120A). Wing length, 3.07; width, 0.96. **Head** (Fig. 120B). Head bright ochre-yellow. Scape and pedicel light ochre-yellowish, flagellomeres light greyish-yellow, darker towards distal end. Face and clypeus ochre-yellow; palpomeres 1–3 brownish-yellow, palpomeres 4–5 lighter; labella whitish-yellow. Occiput with four longer setae dorsally to eye anteriorly to lateral ocellus, two dorsally to ocellus, five posteriorly to ocellus. Scape about twice pedicel length; flagellomere 4 1.7× longer than wide. Palpomere 4 1.2× palpomere 3 length, palpomere 5 1.6× palpomere 4 length. **Thorax**. Scutum ochre-yellowish, brownish on posterior half, scutellum brown with ochre-yellow anterolateral corners. Antepronotum, proepisternum and anepisternum bright ochre-yellow, katepisternum, mesepimeron, laterotergite, and metepisternum greyish ochre-brown; mediotergite brown medially, lighter laterally. Haltere pedicel light ochre, basal ⅔ of knob dark brown, whitish-yellow on apical third. Pleural membrane yellowish. Scutum with five long supra-alar bristles and three pairs of prescutellar bristles; two pairs of scutellar bristles. Proepisternum with four bristles; anepisternum with four bristles along posterior margin. Mesepimeron with two long setae and 27 small setae, laterotergite with three longer setae and two small setae. Metepisternum with 11 fine setae. **Legs**. Coxae whitish, fore coxa with light brownish tinge, hind coxa with a light brownish tinge on basal fourth; fore femur whitish with a reddish tinge, mid and hind femora more whitish, a brownish line along dorsal and ventral edges; tibiae and tarsi light ochre-brown, tips yellowish, tarsi darker. Mid coxa with some few setae across basal fifth, hind coxa with basal third covered with fine setae. Fore tibia with a single bristle dorsolaterally on distal third, mid tibia with a pair of dorsolateral rows of 4–5 bristles and four bristles along ventral edge, hind tibia with two rows of 5-7 bristles dorsolaterally; mid and hind tarsomeres 1–3 with rows of ventral setae. Fore tibia 1.4× tarsomere 1 length, 1.7× tarsomere 2 length. Hind tibial inner spur over 5× tibia width at apex. **Wing** (Fig. 120C). Membrane fumose yellowish-brown, slightly darker along anterior margin. C extending slightly beyond apex of R_5_. Sc short, ending free, R_1_ reaching C at apical fifth of wing; R_5_ reaching C at level of tip of M_1_. First sector of Rs slightly oblique, bare, 0.67× r-m length; r-m almost longitudinal. M_1+2_ 0.70× r-m length; bM 4.0× r-m length. First sector of CuA 0.37× length of second sector of CuA. Cubital pseudovein not reaching level of origin of M_4_. Anal fold only gently curved, almost reaching wing margin. Posterior veins M_1_and M_2_ with dorsal macrotrichia on almost entire length, M_4_ and CuA with macrotrichia on distal fourth of wing; dorsal macrotrichia on anal lobe membrane. **Abdomen**. Abdominal tergite 1 light brown, darker along posterior margin, tergites 2–5 light brown medially with a wide cream-yellow area laterally; tergite 6 mostly dark brown medially, yellowish-brown laterally, tergite 7 yellowish-brown; sternites 1-7 whitish-yellow, sternite 7 yellowish-brown. Sternite 2 with a pair of strong brown bristles. **Terminalia** (Fig. 120D). Whitish-yellow. Sternite 8 wide anteriorly, subrectangular, posterior margin straight, no medial incision, very short projection lateroposteriorly, covered with microtrichia and fine setae, setae on lateroposterior corners longer. Sternite 9 wide and well-sclerotized, notum wide anteriorly, gonopore at center of a subquadrate, well-sclerotized plate. Tergite 8 wide, very short medially, connecting a pair of lateral short lobes with setulae, partially overlapping laterally to sternite 8. Tergite 9+10 short, slender, bare. Cercomeres 1 and 2 apparently fused, no sign of suture, elongate, distal end curved ventrally, covered with microtrichia and setae, setae at tip longer.

**Male**. Unknown.

#### Material examined

**Holotype**: female, ZRCBDP0048460, Nee Soon (NS1), swamp forest, 23.may.2012, MIP leg. (website photo specimen, slide-mounted).

**Etymology**. The species epithet of this species honors Frederick Dainton, a British academic and university administrator. He was asked in 1979 by then Prime Minister Lee Kuan Yew to conduct a preliminary study of university education in Singapore. The report recommended the establishment of a single, strong university covering a wide range of academic disciplines. The recommendation was accepted in April 1980 and it was announced that the University of Singapore and Nanyang University would be merged to form the National University of Singapore, officially inaugurated on 8 August 1980 with about 9,000 students and 800 academic staff.

*Epicypta holltumi* Amorim & Oliveira, sp. N. (Figs. 121A–D)

urn:lsid:zoobank.org:act:EB680482-41B9-4D1B-8F61-19588F7219A2

**Diagnosis**. Head ochre-yellow, antennal scape and pedicel ochre-yellowish and basal flagellomeres ochre-brown, flagellum distally greyish-brown. Scutum blackish-brown with an ochre-yellowish mark medially on anterior end; pleural sclerites greyish-brown, dorso-posterior corner of anepisternum, laterotergite and mediotergite blackish-brown, some other sclerites with dark marks. Coxae and femora whitish, front leg with yellowish tinge, hind coxa with brown mark at basal end. Wing membrane light brownish, cells c and br slightly darker. C extending shortly beyond tip of R_5_; M_1+2_ about as long as r-m. Dorsal macrotrichia on all posterior veins, anal lobe with macrotrichia. Abdominal tergite 1 dark brown, tergites 2–5 dark brownish medially with yellow margins larger on posterior tergites, tergite 6 mostly yellowish with some greyish-brown tinge medially, tergite 7 yellowish, terminalia yellowish. Gonocoxites medially fused, no suture evident, a strongly sclerotized medial pair of projections, a long pair of blade-like dorsal lobes; gonostylus dorsoventrally compressed, setose, more or less rectangular; parameres trapezoid distally with some few fine setae at lateroposterior corners; tergite 9 with a pair of long digitiform, setose extensions.

***Epicypta_holltumi_ZRCBDP0048471_paratype* [67: G, 7 6: C, 79: T, 127: A, 133: G, 223: A, 259: C]**

tctttcttcaacaattgctcatgcagggtcttcagtagatttagctattttttctttacatttagcGggaatttcCtcTattttaggagctattaattttattacaacaattattaatatgcgagcAcctggGattacttttgaccgaatacctttatttgtatgatcagtattaattacagctgttttacttttattatctttaccagtattagcaggtgcAattactatattattaacagatcgaaatttaaatacCtcattttttgatcctgcaggagggggagatcctattttatatcaacatttattt

#### Description

**Female**. Wing length, 2.02; width, 0.74 mm. **Head**. Head ochre-yellowish. Scape and pedicel light ochre-yellowish, flagellomeres greyish-yellow. Face and clypeus ochre-yellowish, maxillary palpomeres 1–3 light brownish-yellow, palpomeres 4–5 lighter. Occiput with three longer setae dorsally to eye anteriorly to lateral ocellus, two dorsally to ocellus, six posteriorly to ocellus. Scape about 1.5× pedicel length; length of flagellomere 4 1.9× width.

Palpomere 4 1.1× palpomere 3 length, palpomere 5 1.9× palpomere 4 length. **Thorax**. Scutum anterior tip ochre-yellow, remaining dark brown, scutellum blackish-brown with ochre-yellow anterolateral corners. Antepronotum, proepisternum, katepisternum, anteroventral corner of anepisternum, and mid of mesepimeron greyish-brown, lighter areas on antepronotum dorsally, anepisternum anteroventrally, katepisternum and mesepimeron; mediotergite dark brown. Haltere light ochre-yellowish, no larger setae. Pleural membrane ochre-yellowish. Scutum with six supra-alars and three pairs of prescutellar bristles; two pairs of scutellar bristles. Proepisternum with three bristles, anepisternum with four bristles along posterior margin. Mesepimeron with two bristles and 31 small setae, laterotergite with two bristles and ten small setae. Metepisternum with 15 fine setae. **Legs**. Coxae whitish, Fore coxa with yellowish tinge, hind coxa with a small brown band along proximal end; femora concolor with coxae, slightly darker at tip, tibiae light brownish-yellow, tarsi slightly darker. Mid coxa with fine setae across basal fifth. Fore tibia with a single strong seta laterally; mid tibia with two dorsolateral rows of 4–5 bristles and a ventral row of three small bristles and one strong seta; hind tibia with a pair of dorsolateral rows of 5-6 bristles. Fore leg tarsomere 1 1.1× tibia length, 1.5× tarsomere 2 length. Hind tibial inner spur 5.4× tibia width at apex. **Wing** (Fig. 121B). Membrane fumose light brown, slightly darker along anterior margin. C extending slightly beyond apex of R_5_. Sc barely produced. R_1_ reaching C on distal fifth of wing; R_5_ reaching C at level of tip of M_1_. First sector of Rs slightly oblique, 0.76× r-m length. M_1+2_ 1.2× r-m length; bM about 8× r-m length; first sector of CuA short, about 0.37× length of second sector of CuA. Cubital pseudovein inconspicuous, CuP short, reaching level of origin of M_4_. Anal fold sclerotized, gently curved, nearly reaching wing margin. Posterior veins M_1_and M_2_ with dorsal macrotrichia on ¾ of length, M_4_, CuA, and anal fold with macrotrichia at distal fifth, anal lobe membrane with macrotrichia. **Abdomen**. Abdominal tergite 1 greyish-brown, tergites 2–5 brownish medially, with cream-yellow lateral bands, small band on tergite 2, wider on tergites 3–5, connecting each other along anterior margin on tergite 5, tergite 6 almost entirely cream-yellow, with only a brownish tinge medio-posteriorly, tergite 7 cream-yellow; sternites 1-7 whitish-yellow. Sternite 2 with a pair of strong brown bristles, sternites 3–6 medially with groups of short bristles. **Terminalia** (Fig. 121B). Whitish-yellow. Sternite 8 wide, subrectangular, posterior margin as wide as base, no lateroposterior projections, covered with microtrichia and fine setae, setae longer along distal margin. Gonopore plate on sternite 9 well-sclerotized. Tergite 8 wide, very short medially. Tergite 9+10 short, slender, bare. Cercomeres 1 and 2 apparently fused, no sign of suture, elongate, covered with microtrichia and setae, setae at tip longer.

#### Material examined

**Holotype**: female, ZRCBDP0048471, Nee Soon (NS1), 04.apr.12, MIP leg.

**Remarks**. Morphologically, this species seems close to but with relevant differences in the color of the head and of the abdomen with *Epicypta angusticollaris* (Edwards), from the Philippines.

*Epicypta* sp. B (Figs. 122A–D)

urn:lsid:zoobank.org:act:C467B833-6434-4C79-A645-A129C3FD86EF

**Diagnosis**. Head dark ochre-yellowish, occiput lighter, antennal scape and pedicel ochre-yellow, flagellomeres brownish-yellow. Scutum dark brown, anterior end ochre-yellow; scutellum blackish-brown; pleural sclerites mostly dark greyish-brown, katepisternum lighter, mediotergite dark brown on dorsal half. Coxae whitish, fore coxa with an orangish tinge, mid and hind coxae with a brownish band on proximal fifth. Wing membrane light brownish, more yellowish along cells c and br; C produced beyond tip of R_5_; M_1+2_ as long as r-m. Dorsal macrotrichia on posterior veins M_1_ and M_2_, M_4_, and distal half of CuA, anal lobe slender, macrotrichia on anal lobe membrane. Abdominal tergites 1–2 and 6 brownish, tergites 3–5 brown medially with cream-yellow area laterally, tergite 7 cream-yellow. Gonocoxites with no lobes projecting beyond base of gonostylus; gonostylus more or less dorsoventrally compressed; paramere with a row of curved short setae distally; tergite 9 with a pair of long parallel projections laterally on terminalia dorsal face.

#### Description

**Male**. Wing length, 2.18; width, 0.80. **Head**. Ochre-yellowish, occiput lighter towards ventral margin. Scape and pedicel greyish ochre-yellow, flagellomeres brownish-yellow, darker towards tip. Face and clypeus ochre-yellowish. Maxillary palpomeres 1–3 brownish-yellow, palpomeres 4–5 lighter; labella whitish-yellow. Occiput with two longer setae dorsally to eye anteriorly to lateral ocellus, one dorsally to ocellus, seven posteriorly to ocellus. Scape 1.7× pedicel length, flagellomere 4 1.9× longer than wide. Palpomere 4 1.2× palpomere 3 length, palpomere 5 twice palpomere 4 length. **Thorax** (Fig. 122A). Scutum dark brown, anterior end ochre-yellow; scutellum blackish-brown with lighter laterals. Antepronotum, proepisternum, anepisternum, katepisternum, mesepimeron, laterotergite, and metepisternum dark greyish-brown, katepisternum slightly lighter, mediotergite dark brown, lighter at ventral half. Haltere light ochre-yellowish, no larger setae. Pleural membrane ochre-yellow. Scutum with 4+1 supra-alars and three pairs of prescutellars. Proepisternum with three bristles, anepisternum with four bristles along posterior margin. Mesepimeron with two bristles and ten fine setae and setulae, laterotergite with 2–3 bristles and five fine setae. Metepisternum with 5–6 fine setae. **Legs**. Fore coxa whitish with an orangish tinge, mid and hind coxae whitish, a brownish band across proximal fifth of hind coxa [all three legs broken]. **Wing** (Fig. 122B). Membrane light fumose brown, slightly darker along anterior margin. C extending slightly beyond R_5_. Sc barely produced. R_1_ reaching C on distal fourth of wing; R_5_ reaching C slightly before level of tip of M_1_. First sector of Rs almost transverse, 0.68× r-m length; r-m oblique. M_1+2_ as long as r-m; bM 6.0× r-m length; first sector of CuA 0.39× length of second sector of CuA. Cubital pseudovein absent, CuP reaching slightly beyond level of origin of M_4_. Anal fold gently curved, almost reaching wing margin. Posterior veins M_1_ and M_2_ almost entirely with dorsal macrotrichia, M_4_ with macrotrichia on distal half, CuA with macrotrichia on distal third of wing; anal lobe membrane with macrotrichia. **Abdomen**. Abdominal tergites 1–2 and 6 brownish, tergites 3–5 brown medially with a cream-yellow band laterally, tergite 7 cream-yellow; sternites 1-7 light ochre-yellowish. Sternite 2 with a pair of strong brown bristles. **Terminalia** (Figs. 122C–D). Ochre-yellowish. Gonocoxites fused medially, no suture present, bare ventrally, no medioventral projection, a pair of short sub-medial projections, no laterodistal extensions. Gonostylus simple, elongate, dorsoventrally compressed, elongate setae on distal third and along inner margin, ventral face of proximal third densely covered by fine setulae. Aedeagus relatively small, anterior ejaculatory apodeme slender at tip, reaching level of insertion of gonostylus, widening medially, distally with a tubular projection with gonopore at level of tip of gonostylus; parameres present as a pair of elongate, weakly sclerotized sclerites laterally, connected to aedeagus. Gonocoxal bridge evident, without apodemes. Tergite 9 present as a pair of long laterodistal digitiform extensions, about as long as gonocoxite lateral extensions, setose at tip. Cerci weakly sclerotized between tergite 9 elongate lateral lobes. **Female**. Unknown.

**Material examined** (male): ZRCBDP0066714, Bukit Timah, maturing secondary forest (BT08), 22.sep.2016, MIP leg. (slide-mounted).

*Epicypta ridleyi* Amorim & Oliveira, sp. N. (Figs. 123A–D)

https://singapore.biodiversity.online/species/A-Arth-Hexa-Diptera-000821and-002079

urn:lsid:zoobank.org:act:37468856-0491-40A8-9B46-74A01AEF6D30

**Diagnosis**. Head light ochre-yellowish, antennal scape, pedicel and basal half of first flagellomere ochre-yellowish, other flagellomeres grey-brownish. Scutum with three connected blackish-brown longitudinal bands over a light ochre-yellowish background, scutellum mostly blackish-brown. Pleural sclerites light brownish, antepronotum and dorsoposterior end of anepisternum ochre-yellowish, mediotergite dark brown. Coxae and femora whitish, fore leg with an orangish tinge, a small brown mark across basal end of hind coxa. Wing membrane light brownish, darker along anterior margin. C barely produced beyond tip of R_5_; Sc incomplete, ending free; M_1+2_ 0.75× r-m length. Dorsal macrotrichia on posterior veins M_1_ and M_2_, M_4_, and on CuA close to tip, macrotrichia on anal lobe membrane. Abdominal tergite 1 brown, tergites 2–5 brown medially with light ochre-yellow laterals, wider on tergites 4–5, tergite 6 ochre-yellow with brownish tinge, tergite 7 and terminalia cream-yellow. Male terminalia gonocoxite with a lateroposterior lobe projecting beyond base of gonostylus, gonostylus wide, dorsoventrally compressed; aedeagus with a pair of separate tubular extensions with independent gonopores; parameres with a pair of long lateral projections; tergite 9 with a pair of long, digitiform separate extensions. Female terminalia with a wide and short medial incision on posterior margin of sternite 8.

***Epicypta_ridleyi_ZRCBDP0047083_hapZRCBDP00470 83_SMH_holotype* [10: A, 29: T, 148: T, 163: G, 187: A, 191: C, 229: C, 265: C, 286: T] -**

ctttcttcAacaattgctcatgcaggaTcttcagtagatttagctattttttctttacatttagcaggtatttcttctattttaggagctattaattttattactactattattaatatgcgagccccaggaattacttttgatcgTataccattatttgtGtgatctgtattaattacagctgtActtCttcttttatctttacctgttttagctggagctattacCatattattaacagatcgaaatttaaacacttcattCtttgatcctgcaggtgggggTgatcctattttataccaacatttattt

#### Description

**Male**. Wing length, 2.182. 26–2.56; width, 0.77–0.83 (n=2). **Head**. Light ochre-yellowish, face and clypeus whitish-yellow. Scape and pedicel light ochre-yellowish, flagellomeres 1–2 yellowish-brown, remaining flagellomeres greyish-brown. Palpomeres ochre-yellowish, palpomeres 3–5 whitish-yellow. Occiput with two longer setae dorsally to eye anteriorly to lateral ocellus, one dorsally to ocellus, five posteriorly to ocellus. Scape 1.4× pedicel length, length of flagellomere 4 1.1× width, tapering to the end. Palpomere 4 1.2× palpomere 3 length, palpomere 5 1.7× palpomere 4 length. **Thorax**. Scutum background ochre-yellow with three blackish-brown wide bands, scutellum blackish-brown with ochre-yellow anterolateral corners. Antepronotum and posterior half of anepisternum, proepisternum, anterior half of anepisternum, katepisternum, mesepimeron, laterotergite, and metepisternum greyish-yellow, mediotergite blackish-brown, lighter at laterals. Pleural membrane yellowish. Haltere pedicel and base of knob light brownish-yellowish, knob whitish distally. Scutum with 6+1 supra-alars and three pairs of prescutellars, scutellum with two pairs of bristles along posterior margin. Proepisternum with three bristles, anepisternum with four bristles along posterior margin. Mesepimeron with two bristles and nine small setae, laterotergite with three stronger setae and four small setae. Metepisternum with 11 fine setae. **Legs**. Coxae whitish, fore coxa with a rose tinge; fore femur concolor with coxa, mid and hind femora whitish, no brown proximal band on hind coxae; tibiae and tarsi light whitish-yellow, tarsi slightly darker, tibiae brownish-yellow at tip. Mid coxa with a band of fine setae across basal fifth, hind coxa with fine setae on proximal third. Fore tibia with one strong seta medially on distal third, hind tibia with two dorsolateral rows of 5-6 small bristles [mid tibiae and tarsi missing]. Fore leg tarsomere 1 1.2× tibia length, 1.7× tarsomere 2 length. Hind tibia inner spur 7.1× tibia width at apex. **Wing** (Fig. 122B). Membrane fumose ochre-yellowish, more yellowish along anterior margin. C produced only slightly beyond tip of R_5_. Sc barely produced. R_1_ reaching C on distal sixth of wing; R_5_ reaching C slightly beyond level of tip of M_2_. First sector of Rs slightly oblique, 0.75× r-m length; r-m almost longitudinal. M_1+2_ 1.0× r-m; bM 8.3× r-m length; first sector of CuA 0.40× length of second sector of CuA. Cubital pseudovein absent, CuP not reaching level of origin of M_4_. Anal fold gently curved, almost reaching wing margin. Posterior veins M_1_ and M_2_ with dorsal macrotrichia along most of their length, M_4_ and CuA with macrotrichia on distal fourth of wing; macrotrichia on anal lobe membrane. **Abdomen**. Abdominal tergite 1 dark brown, tergites 2–5 brown medially, with a wide cream-yellow band laterally, tergites 4–5 with yellow band extending towards mid along anterior margin, tergite 6 mostly dark cream-yellow, only with a medial brownish tinge, sternites 1-7 whitish-yellow; Sternite 2 with a pair of strong brown bristles. **Terminalia** (Fig. 123C). Whitish-yellow. Gonocoxites fused medially, no suture present, bare ventrally, a medioventral bare, flat, wide projection, a long digitiform laterodistal projection dorsally to insertion of gonostylus at each side, extending way beyond tip of gonostylus, with scattered microtrichia and fine setae, distal seta longer. Gonostylus simple, elongate, wide, dorsoventrally compressed, with fine setae on ventral face, longer setae along distal margin longer. Aedeagal-parameral complex almost as wide as terminalia, at each laterodistally corner with a long, digitiform, bare branch and a pair of short sclerotized lobes medially. Gonocoxal bridge wide, apodemes as a pair of long, slender sclerotized bands. Tergite 9 present as a pair of wide laterodistal digitiform extensions with microtrichia and setae. Cerci not recognizable.

**Female** (Fig. 123A). As male, except for the following. **Wing l**ength, 2.69; width, 0.96. **Terminalia** (Fig. 122D). Sternite 8 rectangular, elongate, posterior margin with a rather deep medial incision, with widespread microtrichia and small setae, posterior margin with longer setae. Sternite 9 with anterior apodeme extending beyond anterior end of terminalia widening close to tip, lateral arms wide, distal end triangular, acute, bare, genital chamber bell-shaped, well-sclerotized. Tergite 8 slender medially, wide, lateral lobes setose, projected posteriorly. Tergite 9+10 present as a pair of slender sclerotized inclined band, connected to sternite 9 lateroventrally. Sternite 10 triangular, with some setae close to tip directed dorsally. Cercus large, cercomeres 1 and 2 fused, no sign of suture, basal ⅔ wider, posterior end digitiform, distal setae longer.

#### Material examined

**Holotype**: male, ZRCBDP0047083, National University of Singapore (Icube), 17.jun.2015, MIP leg. (slide-mounted). **Paratypes** (13 males, 5 females). **Males**: ZRCBDP0047089, National University of Singapore (Icube), 15.jul.15, MIP leg.; ZRCBDP0048774, National University of Singapore (PGP), 20.may.15, MIP leg.; ZRCBDP0048776, National University of Singapore (PGP), 20.may.15, MIP leg.; ZRCBDP0048778, National University of Singapore (PGP), 20.may.15, MIP leg.; ZRCBDP0049063, National University of Singapore (PGP), 27.may.15, MIP leg.; ZRCBDP0049064, National University of Singapore (PGP), 27.may.15, MIP leg.; ZRCBDP0049065, National University of Singapore (PGP), 27.may.15, MIP leg.; ZRCBDP0049275, National University of Singapore (Uhall), 15.apr.15, MIP leg.; ZRCBDP0049287, National University of Singapore (Icube), 06.may.15, MIP leg.; ZRCBDP0049295, National University of Singapore (Icube), 20.may.15, MIP leg.; ZRCBDP0049328, National University of Singapore (PGP), 10.jun.15, MIP leg.; ZRCBDP0049337, National University of Singapore (PGP), 08.apr.15, MIP leg.; ZRCBDP0049339, National University of Singapore (PGP), 08.apr.15, MIP leg. **Females**: ZRCBDP0048072, Pulau Semakau (SMO2), old mangrove, 06.dec. 12, MIP leg.; ZRCBDP0048302, Sungei Buloh (SB1), mangrove, 16.oct.13, MIP leg.; ZRCBDP0049062, National University of Singapore (PGP), 27.may.15, MIP leg. (slide-mounted); ZRCBDP0049152, National University of Singapore (Uhall), 20.may.15, MIP leg. (website photo specimen); ZRCBDP0049341, National University of Singapore (PGP), 08.apr.15, MIP leg. (extracted). **Additional sequenced specimens**: ZRCBDP0132865, Singapore, date range 2012-2018, MIP leg.; ZRCBDP0133535, Singapore, date range 2012-2018, MIP leg.; ZRCBDP0136950, Singapore, date range 2012-2018, MIP leg.; ZRCBDP0278350, National of Singapore (CUGE), urban forest, 01.dec. 17, MIP leg.; ZRCBDP0278370, National of Singapore (CUGE), urban forest, 01.dec. 17, MIP leg.; ZRCBDP0278371, National of Singapore (CUGE), urban forest, 01.dec. 17, MIP leg.; ZRCBDP0278399, National of Singapore (CUGE), urban forest, 01.dec. 17, MIP leg.; ZRCBDP0278419, National University of Singapore (SDE), urban forest, 19.oct. 17, MIP leg.; ZRCBDP0278421, National University of Singapore (SDE), urban forest, 19.oct. 17, MIP leg.; ZRCBDP0278468, LCK02, mangrove, 26.apr.18, MIP leg. ZRCBDP0133517, Singapore; ZRCBDP0278357, Singapore Botanical Gardens (CUGE), 24.nov.17; ZRCBDP0278450, Singapore, 11.oct.17; ZRCBDP0278455, LCK02, mangrove, 02.may.18. **Sequence failure specimen**: ZRCBDP0278220, Pulau Ubin (PU18), costal forest, 31.may.18, MIP leg. (MZUSP).

**Etymology**. The species epithet honors Henry Nicholas Ridley (CMG) (FRS) (1855–1956), an English botanist, geologist and naturalist who served as the first Director of the Singapore Botanical Gardens. Also known as ‘Mad Ridley’ or ‘Rubber Ridley’, he had a fantastic amount of interest in furthering the cultivation of rubber trees in Malaya and is largely responsible for the bourgeoning rubber industry in the early 1900s. He lived much of his life in Singapore and was also an enthusiastic collector of insects during his tenure.

**Remarks**. We found six haplotypes among our specimens of *Epicypta ridleyi* Amorim & Oliveira, **sp. n.,** which cluster together as one species by all delimitation approaches except ABGD=0.001–0.005. The male terminalia of the clusters are identical and there is no evidence in morphology that they would belong to separate species.

*Epicypta chezaharaae* Amorim & Oliveira, sp. n. (Figs. 124A–E)

https://singapore.biodiversity.online/species/A-Arth-Hexa-Diptera-000731

urn:lsid:zoobank.org:act:C1BF367F-33C2-4233-AA63-2FC480179F3B

**Diagnosis**. Head light ochre-yellowish, antenna light ochre-yellowish. Scutum ochre-yellowish, anterior fifth lighter, scutellum brown. Antepronotum, proepisternum, katepisternum, anepisternum, and mesepimeron light ochre-yellowish with brownish diffuse marks; laterotergite, mediotergite, and metepisternum light brown. Coxae and femora whitish. Wing membrane light brownish, area along anterior margin slightly darker. C clearly extending beyond tip of R_5_; M_1+2_ almost as long as r-m. Dorsal macrotrichia on posterior veins M_1_, M_2_, posterior half of M_4_, and distal end of CuA, no macrotrichia on anal lobe membrane. Abdominal tergite 1 brown, tergites 2–5 cream-yellow, with a brown mark medially on posterior half, tergite 6 mostly brown, with cream-yellow mark laterally, terminalia whitish-yellow with brownish tinge. Male terminalia gonocoxite with no projection beyond base of gonostylus; gonostylus elongate, weakly sclerotized, with long setae; parameres with a pair of long, curved spines projecting distally; tergite 9 with a pair of long digitiform branches, each with a distal elongate spine. Female terminalia sternite 8 trapezoid, rounded along posterior border medially, sclerotization of vaginal furca cross-like, four short curved setae on ventral face of sternite 10, cercus digitiform.

***Epicypta_chezaharaae_ZRCBDP0048757_hapZRCBDP 0048444_SMH_holotype* [29: T, 31: T, 91: A, 163: A, 17 2: A, 181: T, 184: A, 187: A, 191: T, 199: A, 202: T, 205 : A]**

tctttcttctactattgctcatgctggaTcTtctgtagatttagcaattttttctttacatttagcaggtatttcttcaattttaggagcAattaattttattacaactattattaatatacgagctccaggtattacttttgatcgaatacctttatttgtAtgatcagtAttaattacTgcAgtActtTtattattAtcTttAccggtattagctggggctattactatattattaacagatcgaaacttaaatacttcattttttgacccagcaggaggtggggatcctattttataccaacatttattt

#### Description

**Male**. Wing length, 1.87–1.89; width, 0.75–0.77 (n=2). **Head**. Head entirely light ochre-yellowish. Scape and pedicel light ochre-yellowish, flagellomere 1 ochre-yellowish, remaining flagellomeres light greyish-yellow. Frons and clypeus light ochre-yellowish, maxillary palpomeres 1–3 ochre-yellowish, palpomeres 4–5 whitish-yellow, labella cream-yellow. Occiput with two longer setae dorsally to eye anteriorly to lateral ocellus, two dorsally to ocellus, six posteriorly to ocellus. Scape about 2.3× length of pedicel, flagellomere 4 1.4× longer than wide. Palpomere 4 1.4× palpomere 3 length, palpomere 5 1.4× palpomere 4 length. **Thorax**. Scutum anterior fifth light ochre-yellow, remaining ochre-yellow, scutellum brown with ochre-yellow anterolateral corners. Antepronotum anterior fifth and anepisternum posterior half, ventral ⅔ of mesepimeron ochre-yellow, anterior half of anepisternum, most of katepisternum, dorsal end of mesepimeron, laterotergite, and metepisternum ochre-brown, proepisternum ochre-yellowish with some diffuse ochre-brown marks; mediotergite dark brown medially, lighter laterally. Pleural membrane yellowish. Haltere light ochre-yellowish, no larger setae. Scutum with a row with four stronger and one smaller supra-alar setae and three pairs of prescutellars, two pairs of scutellar bristles. Proepisternum with three bristles, anepisternum with four bristles along posterior margin. Mesepimeron with two bristles and 14 small setae, laterotergite with four bristles and four smaller fine setae; metepisternum with six fine setae. **Legs**. Coxae whitish, fore coxa with yellowish-brown tinge; femora, tibiae, and tarsi light whitish-yellow, tarsi slightly darker, hind femur with a dark brown mark at tip. Fore coxa with four bristles along front distal margin, mid coxa with four fine setae across basal fifth, hind coxae covered with a number of fine setae at basal fourth. Fore tibia with a single laterodorsal strong seta medially; mid tibia with two rows of 2–5 small bristles, two ventral long setae; hind tibia with two rows of 4-5 bristles dorsolaterally and no long setae ventrally, a comb of long setae at outer face apically. Mid and hind tarsomeres 1–3 with rows of lateroventral longer setae. Fore leg tarsomere 1 1.2× tibia length, 1.6× tarsomere 2 length. Hind tibial inner spur 5× tibia width at apex. Tarsal claws with a large apical tooth and a smaller, more basal one. **Wing**. Membrane fumose brownish-yellow, darker on cells c, br and R_1_. C produced beyond tip of R_5_ for a ¼ of distance to M_1_. Sc barely produced. R_1_ long, reaching C on distal fifth of wing; R_5_ reaching C slightly before level of tip of M_1_. First sector of Rs slightly oblique, devoid of setae, 0.91× r-m length; r-m almost longitudinal. M_1+2_ short, 1.0× r-m length; bM slightly over 5× r-m length. First sector of CuA 0.27× length of second sector of CuA. Cubital pseudovein absent, CuP extending slightly beyond level of origin of M_4_. Anal fold gradually curved along its length, almost reaching wing margin. Posterior veins M_1_, M_2_, M_4_, and CuA with dorsal macrotrichia, on M_4_ restrict to distal half, on CuA restrict to tip. Anal lobe membrane with some few macrotrichia. **Abdomen**. Abdominal tergite 1 brownish, lighter on anterior third, tergites 2–5 brown medially, cream-yellow on anterior half and laterally, tergites 6–7 greyish-brown medially and cream-yellow laterally, sternites 1-7 whitish-yellow, Sternite 2 with a pair of strong brown bristles. **Terminalia** (Figs. 124C–D). Whitish-yellow. Gonocoxites fused together medially on their anterior third, no sign of suture, no medioventral process, syngonocoxite projected laterodistally, with a wide gap between arms, no setation on medioventral face of syngonocoxite, long setae on distal end of lateral projections, gonocoxites projecting dorsally, with lobes that contact each other medially with a complete suture between them, a slender V-shaped incision medially on posterior margin, slightly projected lateroposteriorly beyond level of insertion of gonostylus, long fine setae close to posterior margin laterally. Gonostylus laterally compressed, slender basal end gradually widening towards apex, no setae on inner face, setation on entire outer face, longer at distal margin, no branches. Aedeagus with a bulbous base at level of syngonocoxite, abruptly slendering distally, a pair of lateral parameral blades across terminalia connected together dorsally to aedeagus, extending posteriorly into a medial single arm with two digitiform short extensions close to each other medially beyond level of insertion of gonostylus, bearing two long spines. Gonocoxal bridge wide, with a short sclerotized peak medially, apodemes almost touching each other. Tergite 9 divided into a pair of well sclerotized large projections at distal margin of gonocoxal medio-dorsal lobes, extending way beyond distal end of gonostylus, close to each other at anterior end, gradually diverging towards apex, long setae dorsally and laterally, a comb of spines at tip.

**Female** (Fig. 124A). Wing (Fig. 124B) length, 1.79–2.08; width, 0.69–0.78 (n=3). **Head**. Occiput with two longer setae dorsally to eye anteriorly to lateral ocellus, two dorsally to ocellus, four posteriorly to ocellus. **Thorax**. Scutum with a row with six long supra-alar bristles and three pairs of prescutellars; two pairs of scutellar bristles. Proepisternum with three bristles, anepisternum with four bristles along posterior margin. Mesepimeron with two bristles and 18 small setae, laterotergite with two bristles and four small fine setae; metepisternum with six longer setae at posterior end, six small setae at anterior end. **Terminalia** (Fig. 124E). Sternite 8 rectangular, elongate, posterior margin straight medially, with short lateroposterior extensions, microtrichia and fine setae covering entire sclerite, longer setae at posterior margin, lateral ones curved inwards, labia beneath distal margin extending to close to tip of cerci, with four long fine setae. Sternite 9 with wide genital chamber, lined with microtrichia, gonopore connected to two gonoducts, sclerotized part sword-like, with a pair of arms extending laterally towards T9+10, an anterior apodeme widening towards apex and a posterior median sclerotized band reaching genital chamber. Tergite 8 short and wide, with microtrichia and short and long setae along posterior margin. T9+10 bare, with a pair of short lateral lobes connected medially by a slender, sclerotized band, lateral ends extending posteriorly, fused to sternite 9 laterally. Cercomeres 1 and 2 probably fused, no sign of suture, digitiform, covered with microtrichia and elongate setae, one stronger seta at apex, apparently an ovoid sensorial area at inner face on anterior third.

#### Material examined

**Holotype**: male, ZRCBDP0048757, Nee Soon (NS1), 25.feb. 2015, MIP leg. (slide-mounted). **Paratypes** (7 males, 28 females). **Males**: ZRCBDP0048796, Nee Soon (NS1), 18.mar. 2015, MIP leg.; ZRCBDP0048803, Nee Soon (NS1), 18.mar. 2015, MIP leg. (slide-mounted); ZRCBDP0048823, Nee Soon (NS1), 14.jan. 2015, MIP leg.; ZRCBDP0048842, Nee Soon (NS1), 14.jan. 2015, MIP leg.; ZRCBDP0048862, Nee Soon (NS1), 14.jan. 2015, MIP leg.; ZRCBDP0049213, Nee Soon (NS1), 10.dec. 2014, MIP leg. (extracted); ZRCBDP0066746, Bukit Timah, maturing secondary forest (BT06), 28.sep.2016, MIP leg. (slide-mounted) (MZUSP). **Females**: ZRCBDP0047799, Nee Soon (NS2), swamp forest, 13.nov. 2013, MIP leg. (slide-mounted); ZRCBDP0047908, Nee Soon (NS2), swamp forest, 30.oct. 2013, MIP leg.; ZRCBDP0048442, Singapore, NS01, 23.may. 12, MIP leg. (website imaged specimen, extracted); ZRCBDP0048444, Nee Soon (NS1), swamp forest, 06.jun. 2012, MIP leg. (extracted, slide-mounted); ZRCBDP0048445, Nee Soon (NS2), swamp forest, 23.jan.13, MIP leg.; ZRCBDP0048722, Nee Soon (NS2), 28.jan. 2015, MIP leg.; ZRCBDP0048741, Nee Soon (NS1), 25.feb. 2015, MIP leg.; ZRCBDP0048802, Nee Soon (NS1), 18.mar. 2015, MIP leg.; ZRCBDP0048871, Nee Soon (NS1), 31.dec. 2014, MIP leg.; ZRCBDP0049085, Nee Soon (NS1), 24.dec. 2014, MIP leg. (slide-mounted) (MZUSP); ZRCBDP0049122, Nee Soon (NS1), 24.dec. 2014, MIP leg.; ZRCBDP0066819, Bukit Timah, maturing secondary forest (BT09), 05.oct. 2016, MIP leg.; ZRCBDP0072458, Bukit Timah, old secondary forest (BT07), 08.dec. 2016, MIP leg.; ZRCBDP0072688, Bukit Timah, old secondary forest (BT07), 01.dec. 2016, MIP leg.; ZRCBDP0072691, Singapore, date range 2012-2018, MIP leg.; ZRCBDP0072713, Bukit Timah, primary forest (BT05), 01.dec. 2016, MIP leg.; ZRCBDP0072717, Bukit Timah, primary forest (BT05), 01.dec. 2016, MIP leg.; ZRCBDP0074026, Bukit Timah, maturing secondary forest (BT08), 14.dec. 2016, MIP leg.; ZRCBDP0074029, Bukit Timah, maturing secondary forest (BT08), 05.oct. 2016, MIP leg.; ZRCBDP0078975, Singapore, date range 2012-2018, MIP leg.; ZRCBDP0133430, Singapore, (date range 2012-2018), MIP leg. (slide-mounted); ZRCBDP0133443, Singapore, (date range 2012-2018), MIP leg. (slide-mounted) (MZUSP); ZRCBDP0136948, Singapore, (date range 2012-2018), MIP leg.; ZRCBDP0137218, Bukit Timah Forest (BT06), 26.apr. 17, MIP leg. (extracted, slide-mounted); ZRCBDP0154808, Singapore, (date range 2012-2018), MIP leg.; ZRCBDP0154856, Singapore, (date range 2012-2018), MIP leg. (slide-mounted) (MZUSP); ZRCBDP0154860, Singapore, (date range 2012-2018), MIP leg.; ZRCBDP0154872, Singapore, (date range 2012-2018), MIP leg.

**Additional sequenced specimens:** ZRCBDP0048913, Nee Soon (NS1), 31.dec.14; ZRCBDP0066805, Bukit Timah, maturing secondary forest, 16.aug.16. **Sequence failure specimens**: ZRCBDP0041130; ZRCBDP0067192; ZRCBDP0067324.

**Possibly non-conspecific cluster** (10 males, 5 females): ZRCBDP0278180, Pulau Ubin (PU18), mangrove, 10.may. 2018, MIP leg.; ZRCBDP0278191, Pulau Ubin (PU18), mangrove, 31.may. 2018, MIP leg.; ZRCBDP0278218, Pulau Ubin (PU18), mangrove, 31.may. 2018, MIP leg.; ZRCBDP0278237, Pulau Ubin (PU18), mangrove, 31.may. 2018, MIP leg.; ZRCBDP0278258, Singapore, 07.Jun. 2018, MIP leg. (slide-mounted); ZRCBDP0278260, Singapore, 07.Jun. 2018, MIP leg.; ZRCBDP0279113, Singapore, 31.may. 2018, MIP leg.; ZRCBDP0279121, Singapore, 31.may. 2018, MIP leg.; ZRCBDP0279198, Singapore, 07.Jun. 2018, MIP leg.; ZRCBDP0284178, Pulau Ubin (PU18), mangrove, no date, MIP leg.; ZRCBDP0284248, Singapore, date range 2012-2018, MIP leg.; ZRCBDP0048094, Pulau Semakau (SMO2), old mangrove, 31.October-6.nov. 2013, MIP leg. (slide-mounted); ZRCBDP0048120, Pulau Semakau (SMO2), old mangrove, 18.jul. 2013, MIP leg. (slide-mounted); ZRCBDP0048237, Pulau Semakau (SMO2), old mangrove, 14.nov. 2013, MIP leg. (website photo specimen, slide-mounted); ZRCBDP0278259, Singapore, 07.Jun. 2018, MIP leg. (slide-mounted).

**Etymology**. The species epithet of this species honors Che Zahara binte Noor Mohamed (1907-1962). Born in Singapore, she was one of the first Malay women to fight for modern women’s rights in Singapore. She founded the Malay Women’s Welfare Association, the first Muslim women’s organization in Singapore, and was instrumental in the passage of the Women’s Charter, a women’s rights act. Che Zahara was inducted into the Singapore Women’s Hall of Fame in 2014.

**Remarks**. This is the second most abundant species of *Epicypta* in Singapore. There are three subclusters (one of them with a single specimen), diverging from each other by 3.53% and 3.19%—only OC at p-distance 4-5% and ABGD P=0.1 would bring all 20 haplotypes together. We assume this as another grey-zone case. There are no males of two of the three subclusters. The specimens are thus not designated paratypes and it is not possible to decide about conspecificity of these subclusters.

*Epicypta tanjiakkimi* Amorim & Oliveira, sp. n. (Figs. 125A–D)

https://singapore.biodiversity.online/species/A-Arth-Hexa-Diptera-000735

urn:lsid:zoobank.org:act:70001E9C-2EE6-4BD9-8D1A-895E3DBD9A61

**Diagnosis**. Head ochre-yellow, antennal scape, pedicel and basal half of flagellum ochre-yellowish, distal half greyish-yellow. Scutum ochre-yellow, scutellum brown; most pleural sclerites ochre-yellow, anepisternum, katepisternum, mesepimeron, mediotergite, and metepisternum with brownish marks. Fore coxa and femur cream-yellowish, mid and hind coxae and femora whitish, hind femur with brownish tip. Wing membrane light brownish, area along anterior margin on basal half more yellowish. C extending beyond tip of R_5_; M_1+2_ much shorter than r-m, r-m almost longitudinal. Dorsal macrotrichia on posterior veins M_1_ and M_2_, and on M_4_ close to tip, no macrotrichia on anal lobe membrane. Abdominal tergite 1 yellowish-brown, tergites 2–6 dark ochre-yellowish with a dark brown medial mark, tergite 7 cream-yellow, terminalia whitish-yellow. Gonocoxites fused medially and extending dorsally, displacing tergite 9 distally; gonostylus simple, elongate, displaced medially. Aedeagus tubular, simple. Parameres with a pair of strongly sclerotized distal flaps. Tergite 9 with a pair of large, independent lobes with a row of short spines at distal margin.

***Epicypta_tanjiakkimi_ZRCBDP0048748_hapZRCBDP0 048748_SMH_holotype* [67: A, 70: T, 109: A, 185: G, 18 8: T, 194: T, 203: C, 208: T, 247: T]**

tctttcttctacaattgctcatgcaggttcatctgttgatttagcaattttttctttacatttggcAggTatttcttctattttgggggctattaattttattactacAattattaatatacgagctccagggattacttttgatcgaatacctttatttgtttgatctgttttaattactgctGttTtattgTtattatctCttccTgttttagcaggggctattacaatattattaacagatcgTaatttaaatacttcattttttgatccagcgggaggaggagatcctattttatatcaacatttattt

#### Description

**Male**. Wing length, 2.08–2.14; width, 0.78–0.80 (n=2). **Head**. Head light ochre-yellowish. Scape and pedicel ochre-yellowish, flagellomere 1 ochre-yellowish, remaining flagellomeres light greyish-yellow. Face and clypeus ochre-yellowish; palpomeres 1–3 light brown, palpomeres 4–5 lighter; labella ochre-yellowish. Occiput with three longer setae dorsally to eye anteriorly to lateral ocellus, two dorsally to ocellus, seven posteriorly to ocellus. Scape about 1.7× pedicel length; flagellomere 4 2.0× longer than wide. Palpomere 4 as long as palpomere 3, palpomere 5 2.0× palpomere 4 length. **Thorax**. Scutum light ochre-yellow, scutellum brown with ochre-yellow anterolateral corners. Most antepronotum, dorsal half of proepisternum, posterior half of anepisternum, and most katepisternum ochre-yellowish, posterior half of antepronotum, anterior half of anepisternum, dorsal half of katepisternum, dorsal end of mesepimeron, laterotergite, and metepisternum greyish ochre-brown; mediotergite brown, lighter lateroventrally. Haltere light ochre-yellowish, no larger setae. Pleural membrane yellowish. Scutum with seven supra-alar and two post-alar bristles, and three pairs of prescutellar bristles; scutellum with two pairs of marginal bristles. Proepisternum with three bristles, anepisternum with five bristles along posterior margin. Mesepimeron with two bristles and 13 small setae and setulae, laterotergite with one bristle and two small setae. Metepisternum with 15 fine setae. **Legs**. Fore coxa light ochre-yellowish, mid and hind coxae whitish; fore femur light ochre-yellowish, mid and hind femora whitish-yellowish, mid femur with a basal brown mark ventrally and a small brown mark distally, hind femur with a basal mark, tibiae ventrally with an ochre-brownish mark distally; tarsi light ochre-yellowish. Mid coxa with a band of fine setae at basal fifth, hind coxa covered with fine setae on basal third. Fore tibia with a single dorsal bristle medially, mid tibia with a row of 4-5 brown bristles dorsolaterally and three fine bristles ventrally, besides distal strong bristles, hind tibia with two rows of 4–6 dorsolateral bristles. Mid and hind tarsomeres with rows of ventrolateral setae. Fore leg tarsomere 1 1.2× tibia length, 1.6× tarsomere 2 length. Hind inner tibial spur almost 4.4× tibia width at apex. **Wing** (Fig. 125B). Membrane fumose light brown, yellowish along anterior margin. Sc barely produced. C extending beyond apex of R_5_ for one third of distance to M_1_. R_1_ reaching C on distal 1/6 of wing, R_5_ reaching C beyond level of tip of M_2_. First sector of Rs slightly oblique, 1.1× length of r-m; r-m quite longitudinal. M_1+2_ very short, 0.50× length of r-m; bM slightly over 7.4× r-m length; first sector of CuA 0.33× length of second sector of CuA. Cubital pseudovein entirely absent, CuP ending slightly beyond level of origin of M_4_. Anal fold almost reaching wing margin. Dorsal macrotrichia on posterior veins M_1_ and M_2_, and on M_4_ close to tip, no macrotrichia on anal lobe membrane. **Abdomen**. Abdominal tergite 1 light brown, tergites 2–6 brown medially, cream-yellow on anterior half and laterally, tergites 2 and 5 with larger darker areas, tergite 6 ochre-yellowish with a brown band anteriorly and medially, tergite 7 ochre-yellow; sternites 1-7 whitish-yellow, Sternite 2 with a pair of strong brown bristles. **Terminalia** (Fig. 125C). Whitish-yellow, with dark brown tip of parameres and brown cerci. Gonocoxites fused together medially on ventral face, no suture, entirely devoid of setae ventrally, syngonocoxite with a pair of posterolateral extensions with setae distally, long straight setae along margin, no medioventral process, dorsomedial lobes of gonocoxites fused to each other on dorsal face of terminalia, no sign of fusion. Gonostylus elongate, more or less compressed dorsoventrally, with setae on both faces, setae along distal end longer. Parameral sclerites large, semicircular at base, with strongly sclerotized medio-posterior projections bearing a beak directed dorsally at distal end, a weakly sclerotized triangular, weakly sclerotized connection between parameres with a row of small setae on distal margin. Aedeagus weakly sclerotized, present between parameres. Tergite 9 present as a pair of entirely separated long lobes extending way beyond tip of parameres and gonostylus, in contact at anterior end, each lobe with a subdistal winglet on inner edge and a sclerotized, slightly curved tip with a comb of short spines. Cerci present as a pair of short stripes ventrad to base of tergite 9 lobes.

**Female** (Fig. 125A). As male, except for the following. **Wing l**ength, 2.02–2.03; width, 0.77–0.80 (n=2). **Head**. Occiput with two longer setae dorsally to eye anteriorly to lateral ocellus, two dorsally to ocellus, seven posteriorly to ocellus. **Thorax**. Scutum with seven supra-alar long setae and three pairs of prescutellars, two pairs of scutellar bristles. Proepisternum with three bristles directed ventrally, anepisternum with four bristles along posterior margin. Mesepimeron with two bristles and 17 small setae and setulae, laterotergite with two long setae, no smaller setae or setulae. Metepisternum with 17 small fine setae and setulae. **Legs**. Mid tibia with two dorsolateral rows of 3–5 bristles and 3 bristles along ventral edge, hind tibia with a row of 5–6 bristles dorsally, a row of three long setae ventrally. **Wing**. Posterior veins M_1_ and M_2_ with over half of their length with dorsal macrotrichia, M_4_ with macrotrichia at distal fourth, one macrotrichium on CuA close to tip, anal fold with four macrotrichia on distal fourth, no macrotrichia on anal lobe membrane. **Terminalia** (Fig. 125D). Sternite 8 wide, a pair of lateroposterior projections and a medial, acute sclerotized posterior projection separated by a pair of curved medio-lateral incisions, few microtrichia and fine setae scattered on anterior half, some few fine setae and a pair of subapical strong setae on medial projections, setulae along incisions and long setae posteriorly on lateroposterior projections. Sternite 9 with a pair of lateral wide arms and an anterior apodeme with a sclerotized axis extending to anterior end of terminalia, genital chamber sclerotized, displaced distally. Tergite 8 wide, short medially and with a pair of short lateral lobes, no microtrichia medially, long setae restricted to lateroventral lobes. T9+10 bare, slender medially, with a pair of wide lateral lobes, fused to sternite 9. Cercomeres 1 and 2 probably fused, no sign of suture, basal half ovoid, large, slender on distal half, covered with microtrichia and elongate setae, distal setae longer and curved.

#### Material examined

**Holotype**: male, ZRCBDP0048748, Nee Soon (NS1), 25.feb.15, MIP leg. (slide-mounted). **Paratypes**: 6 males, 6 females. **Males**: ZRCBDP0133427, Singapore, Nee Soon, swamp forest, (date range 2012-2018), MIP leg.; ZRCBDP0133432, Singapore, Nee Soon, swamp forest, (date range 2012-2018), MIP leg.; ZRCBDP0133437, Singapore, Nee Soon Swamp Forest, (date range 2012-2018), MIP leg.; ZRCBDP0133447, Singapore, Nee Soon, swamp forest, (date range 2012-2018), MIP leg.; ZRCBDP0133468, Singapore, (date range 2012-2018), MIP leg. (slide-mounted) (MZUSP); ZRCBDP0133561, Singapore, (date range 2012-2018), MIP leg. **Females**: ZRCBDP0047866, Nee Soon (NS2), swamp forest, 02.oct.13, MIP leg. (slide-mounted) (MZUSP); ZRCBDP0048929, Nee Soon (NS1), 31.dec.14, MIP leg.; ZRCBDP0133458, Singapore, (date range 2012-2018), MIP leg.; ZRCBDP0154874, Singapore, (date range 2012-2018), MIP leg.; ZRCBDP0154967, Singapore, Nee Soon, swamp forest, (date range 2012-2018), MIP leg; ZRCBDP0048452, Singapore. **Additional sequenced specimens:** ZRCBDP0048853, Nee Soon (NS1), 14.jun.15; ZRCBDP0072721, Bukit Timah (BT01), 14.dec.16; ZRCBDP0072737, Bukit Timah (BT01), 14.dec.16; ZRCBDP0072751, Bukit Timah (BT08), 14.dec.16; ZRCBDP0137017, Singapore; ZRCBDP0154873, Singapore.

**Etymology**. The species epithet of this species honors Tan Jiak Kim (1859–1917). A Straits-born Chinese, he was businessman and philanthropist, President of the Straits Chinese British Association. He was a leading force to establish a medical school in Singapore, petitioning the colonial Governor of the Straits Settlements, Sir John Anderson, in name of a group of representatives of the Chinese and other non-European communities. He disbursed medical scholarships to poorer students, giving them an opportunity to study abroad while pursuing their academic career.

**Remarks**. *Epicypta tanjiakkimi* Amorim & Oliveira, **sp.n.** has eight haplotypes which fall into three 1% subclusters with no delimitation conflicts. We have specimens from the swamp forest in Singapore in our samples.

*Epicypta gehminae* Amorim & Oliveira, sp. n. (Figs. 126A–D)

urn:lsid:zoobank.org:act:8603180E-B187-4649-9914-A819E4374275

**Diagnosis**. Head brown, most of antenna light brown. Scutum mostly brown, anterior end and laterals more yellowish, scutellum dark brown; most pleural sclerites light brown, anepimeron, proepisternum, and katepisternum light brownish-yellow. Fore coxa ochre-yellowish, mid and hind coxae whitish, mid femur with brownish tip. Wing vein C not extending beyond tip of R_5_; M_1+2_ as long as r-m, r-m almost longitudinal. Dorsal macrotrichia on posterior veins M_1_, M_2_, M_4_, and CuA, no macrotrichia on anal lobe membrane. Abdominal tergites brown. Female terminalia with no posterior lobes on sternite 8, a medial sclerotized longitudinal line on sternite 9.

***Epicypta_gehminae_ZRCBDP0284297_hapZRCBDP028 4297_SMH_holotype* [34: C, 121: G, 139: A, 187: A, 188 : T, 190: A, 211: G, 241: T, 262: C]**

tctatcttctactattgctcatgcaggatcttcCgtagatttagcaattttttctttacatttagcaggaatttcttctattttaggagctattaattttattacaacaattattaatatGcgagccccaggaatttcAtttgatcgaatacctttatttgtttgatcagttttaattactgctgtATtAttattattatctttaccagtGttagctggagctattactatattattaacTgatcgaaatttaaatacttcCttttttgatccagcaggaggaggggatccaattttataccaacatttattc

#### Description

**Female**. Wing length, 2.08; width, 0.78. **Head** (Fig. 126A). Head brown. Scape light brown, pedicel ochre-yellowish, flagellomeres light brown. Face and clypeus light brown; palpomeres 1–3 dark ochre-yellowish [both palpomeres 4–5 broken in the holotype]; labella brownish-yellow. Occiput with two longer setae dorsally to eye anteriorly to lateral ocellus, one dorsally to ocellus, four posteriorly to ocellus. Scape about 1.8× pedicel length; flagellomere 4 2.3× longer than wide. **Thorax** (Fig. 126B). Scutum brown, anterior end and lateral margins brownish-yellow, scutellum dark brown. Antepronotum, proepisternum, and katepisternum light brownish-yellow, anepisternum, mesepimeron, and metepisternum light brown, laterotergite and mediotergite dark brown. Pleural membrane light yellowish-brown. Scutum with five supra-alars and three pairs of prescutellar bristles; scutellum with two pairs of marginal bristles. Proepisternum with three bristles, anepisternum with four bristles along posterior margin. Mesepimeron with two bristles and six setulae, laterotergite with two bristles and some small setae. **Legs**. Fore coxa ochre-yellowish, mid and hind coxae whitish; fore femur light ochre-yellowish, mid and hind femora whitish-yellow, mid femur with a small brown mark distally; tarsi light ochre-yellowish. Mid coxa with a band of fine setae at basal fifth, hind coxa covered with fine setae on basal third. Fore tibia with a single dorsal bristle medially, mid tibia with two rows of 4–5 brown bristles dorsolaterally and three fine bristles ventrally, besides strong distal bristles, hind tibia with two rows of 3–5 dorsolateral bristles. Mid and hind tarsomeres with rows of ventrolateral setae. Fore leg tarsomere 1 1.0× tibia length, 1.6× tarsomere 2 length. Hind inner tibial spur almost 6.3× tibia width at apex. **Wing** (Fig. 126C). Membrane fumose light brown, yellowish along anterior margin. Sc barely produced. C not produced beyond apex of R_5_. R_1_ reaching C on distal fifth of wing, R_5_ reaching C beyond level of tip of M_2_. First sector of Rs slightly oblique, 0.8× length of r-m; r-m almost longitudinal. M_1+2_ very short, 1.0× length of r-m; bM slightly over 6.0× r-m length; first sector of CuA 0.40× length of second sector of CuA. Cubital pseudovein entirely absent, CuP not even reaching level of origin of M_4_. Anal fold almost reaching wing margin. Dorsal macrotrichia on posterior veins M_1_ and M_2_, distal half of M_4_, and distal third of CuA, no macrotrichia on anal lobe membrane. **Abdomen**. Abdominal tergites brown, sternites light brownish-yellow, sternite 2 with a pair of strong brown bristles. **Terminalia** (Fig. 126D). Sternite 8 subtrapezoid, no lateroposterior projections. Sternite 9 with a pair of medial sclerotized line, anterior arm of genital fork extending to anterior end of terminalia, slightly wider at tip. Tergite 8 much wider than sternite 8, short medially, lateroanterior arms extending anteriorly, some setae along posterior margin. T9+10 bare, short medially, with a pair of long lateral projections. Cercomeres 1 and 2 probably fused, no sign of suture, basal half ovoid, slender on distal half, covered with microtrichia and elongate setae, distal setae longer and curved.

#### Material examined

**Holotype**: female, ZRCBDP0284297, Singapore, 16.may.18, MIP leg. (slide-mounted).

**Etymology**. The species epithet of this species honors Geh Min (1950-), an eye surgeon by trade, best known for her work as a conservationist, who served as the head of the Nature Society (Singapore) from 2000 to 2008. Her efforts were important in ensuring the preservation of the biodiverse Chek Jawa wetlands at Pulau Ubin. For her contributions to environmental sustainability, Geh Min was awarded the inaugural President’s Award for the Environment as well as the Stellar Award from the United Nations Development Fund for Women in 2006. She was inducted into the Singapore Women’s Hall of Fame in 2014.

*Epicypta jackieyingae* Amorim & Oliveira, sp. n. (Figs. 127A–E)

urn:lsid:zoobank.org:act:AAF6D8B4-8DB6-4F90-A06C-8D593A64301F

**Diagnosis**. Head ochre-yellow, antennal scape, pedicel ochre-yellowish, flagellum yellowish-brown. Scutum ochre-yellow, with a dark brown mark above wing; scutellum dark brown; pleural sclerites ochre-yellowish, with brown marks on dorsal half of mesepimeron and paratergite, mediotergite dark brown, yellowish laterally. Fore coxa ochre-yellowish with brownish tinge, mid and hind coxae whitish with a brown mark at tip, mid and hind femora yellowish-brown with a brown mark at proximal end, hind femur with a brown mark also at distal end. Wing membrane light brownish, cells c and br more yellowish. C extending way beyond tip of R_5_; M_1+2_ 1.0× r-m length. Dorsal macrotrichia on posterior veins M_1_, M_2_, M_4_, CuA, and anal fold, no macrotrichia on anal lobe membrane. Abdominal tergite 1 whitish-yellow laterally, a brown medial mark, tergites 2–3 and 5 brown medially with a cream-yellow lateral, tergite 4 yellowish-brown with lighter laterals, tergite 6 yellowish with a pair of large separate brownish areas on anterior two-thirds, tergites 7– 8 yellowish. Gonocoxites with a pair of short dorsolateral lobes extending slightly beyond base of gonostylus, dorsal borders of gonocoxites fused to each other dorsally. Gonostylus with a flat sclerite at level of laterodistal extension of gonocoxite, extending medio-posteriorly as a bare, slightly curved blade and an elongate digitiform lateral lobe. Aedeagus tubular, no clear opening medially. Tergite 9 with a pair of long posterior digitiform setose extensions, extending way beyond tip of parameres, with a distal spine. Female sternum 8 with no incision medially along posterior border, genital furca slender at anterior end.

***Epicypta_jackieyingae_ZRCBDP0278332_hapZRCBDP 0278332_SMH_holotype* [28: T, 58: G, 127: A, 163: G, 1 66: G, 169: T, 241: T, 253: T] -**

ttatcttctacaattgctcatgcaggTtcatctgttgatttagcaattttttctttGcatttagcaggtatttcttctattttaggagctattaattttattacaactattattaatatacgagcAccaggaattacttttgatcgaatacctttatttgtGtgGtcTgttttaattacagctgttttattattattatcattacctgttttagcaggtgctattacaatattattaacTgatcgaaatttTaatacttctttttttgacccagctggtggnggagatccaattttatatcaacatttattt

#### Description

**Male**. Wing length, 2.43; width, 0.91. **Head** (Fig. 127A). Ochre-yellowish. Antennal scape and pedicel ochre-yellowish, flagellomeres yellowish-brown. Face and clypeus light brown. Palpomeres light yellowish-brown. Labella cream-yellowish. Occiput with two longer setae dorsally to eye anteriorly to lateral ocellus, two dorsally to ocellus, seven posteriorly to ocellus. Scape about 1.6× pedicel length; length of flagellomere 4 2.2× width. Palpomere 4 1.3× palpomere 3 length, palpomere 5 1.6× palpomere 4 length. **Thorax** (Fig. 127A). Scutum ochre-yellow, with a dark brown mark laterally above wing, scutellum dark brown, ochre-yellow along anterior and posterior margins. Pleural sclerites ochre-yellowish, with brown marks on dorsal half of mesepimeron and anterior basalare, mediotergite dark brown with yellowish laterals and ventral end. Haltere whitish-yellow. Pleural membrane yellowish. Scutum with 5+1 supra-alars, three pairs of prescutellar bristles and two additional smaller pairs in a slightly more anterior line, scutellum with two pairs of marginal bristles. Proepisternum with three bristles, anepisternum with four bristles along posterior margin. Mesepimeron with two bristles and 16 small setae and setulae, laterotergite with one large bristle and one small bristle, and two smaller setae; metepisternum with 12 fine setae. **Legs**. Fore coxa ochre-yellowish with a brownish tinge, mid coxa whitish with a brown mark at tip on ventral face, hind coxa whitish, with a brown mark at tip on external face. Mid and hind femora yellowish-brown, mid femur with a brown mark at proximal end ventrally, hind femur with a brown mark at proximal and distal ends ventrally. Tibiae and tarsi yellowish-brown, hind leg tarsomere 1 proximal fifth with a dark brown mark. Mid coxa with some few fine setae across basal fifth, hind coxa with fine setae on basal third extending more distally along anterior margin. Fore tibia with one bristle medially on inner face, mid tibia with two irregular rows of 3–5 bristles dorsolaterally, three bristles along ventral edge, distalmost one stronger, hind tibia with two irregular dorsolateral rows of 5–6 bristles. Hind tibia inner spur over 5.7× tibia width at apex. **Wing** (Fig. 127B). Membrane fumose brown, darker along anterior margin, a dark brown mark at wing base. Sc faint (visible only in phase contrast), with two dorsal macrotrichia. C produced beyond tip of R_5_ for over a third of distance to M_1_. R_1_ reaching C on distal fifth of wing; R_5_ reaching C before level of tip of M_1_. First sector of Rs clearly oblique, 1.0× r-m length; r-m almost longitudinal. M_1+2_ 0.63× r-m length; bM slightly over 7.6× r-m length; first sector of CuA 0.34× length of second sector of CuA. Cubital pseudovein present only as a fold, CuP extending slightly beyond level of origin of M_4_. Anal fold gradually curved, almost reaching wing margin. Posterior veins M_1_ and M_2_ with dorsal macrotrichia along most of their length, M_4_ with macrotrichia on distal half, CuA with setae on distal fifth, no macrotrichia on anal lobe membrane. **Abdomen**. Abdominal tergite 1 whitish-yellow with a brown medial mark, tergites 2–3 and 5 brown medially with cream-yellow lateral bands, tergite 4 yellowish-brown with lighter laterals, tergite 6 yellowish with a pair of large separate brownish areas on anterior two-thirds, tergites 7– 8 yellowish; sternites 1-7 whitish-yellow. Sternite 2 with a pair of strong brown bristles, tergites 2-6 with long, darker setae medially. **Terminalia** (Figs. 127C–D). Whitish-yellow. Gonocoxites fused medially, no suture present, bare ventrally, and with a pair of short dorsolateral lobes extending only slightly beyond insertion of gonostylus, gonocoxites fused to each other dorsally with no suture, displacing tergite 9 to a posterior position. Gonostylus as a flat sclerite at level of laterodistal extension of gonocoxite, extending medio-posteriorly as a slightly curved bare blade extending almost to level of tip of paramere spines, and an elongate digitiform lateral lobe, with setae at distal half on both faces, at tip a longer seta. Aedeagus tubular, without a clear anterior medial ejaculatory apodeme, opening medially between distal arms of parameres on a sclerotized distal structure. Parameres connected together medially, each side with a pair of projections, one more ventral, blade-like branch distally falciform and one more dorsal, strongly sclerotized bearing a pair of digitiform distal sublobes, each ending with a long spine, weakly sclerotized medially, a pair of long sub-medial setae on posterior margin. Gonocoxal bridge not detected. Tergite 9 present as a pair of long posterior digitiform extensions close to each other at anterior end, covered with long setae, extending way beyond tip of parameres, with a spine at tip. Cerci small, setose, dorsad to posterior margin of parameres.

**Female**. As male, except for the following. **Wing l**ength, 2.37; width, 0.90. **Head**. Occiput with two longer setae dorsally to eye anteriorly to lateral ocellus, one dorsally to ocellus, six posteriorly to ocellus. **Thorax**. Scutum with 4+1 supra-alar long setae and three pairs of prescutellars, lateral pair smaller; two pairs of scutellar bristles. Proepisternum with three bristles directed ventrally, anepisternum with four bristles along posterior margin. Mesepimeron with two bristles and 14 small setae, laterotergite with two bristles (one of them smaller) and two small setae. Metepisternum with 14 small fine setae. **Abdomen**. Tergite 1 cream-yellow, tergites 2–3 cream-yellow with a pair of separated large brown marks, tergite 4 cream-yellow with a small brownish mark medially, tergite 5 cream-yellow with a median brown mark and a pair of additional, larger brown marks more laterally, tergite 6 yellowish-brown. **Terminalia** (Fig. 127E). Sternite 8 wide, rectangular, posterior margin straight, no medial incision, no lateroposterior projections, covered with microtrichia and fine setae, longer setae on posterior margin, setae on lateroposterior corners strong and curved. Sternite 9 wide, anterior apodeme extending beyond anterior end of terminalia, with a sclerotized axis and slightly widening at tip, genital chamber slender. Tergite 8 short, wide, covered with microtrichia and with setulae along posterior margin, long setae restrict to lateroventral lobes. T9+10 bare, U-shaped, projecting laterodistally at sides of cerci, rounded at tip. Cercomeres 1 and 2 probably fused, no sign of suture, basal half large, ovoid, slender on distal half, covered with microtrichia and elongate setae, distal setae longer and curved.

#### Material examined

**Holotype**: male, ZRCBDP0278332, Singapore, 03.may.18, MIP leg. (extracted, slide-mounted). **Paratype** (female): ZRCBDP0284204, Singapore, PU20, date range 2012-2018, MIP leg. (slide-mounted).

**Etymology**. The species epithet honors Jackie Yi-Ru Ying (1966-). Taipei-born, she is a leading researcher in nanotechnology who left a professorship (as one of the youngest full Professors at 35) at MIT to found the Institute of Bioengineering and Nanotechnology in Singapore to advance biomedical research. She was inducted into the Singapore Women’s Hall of Fame in 2014.

**Remarks**. The pair of long, curved projections of tergite 9 with a short blunt spine at the apex is an unique features of the male terminalia of *Epicypta jackieyingae* Amorim & Oliveira, **sp. n.** The tergite 9 of *E. tanjiakkimi* Amorim & Oliveira, **sp. n.** is also long (though not curved) and the structure of the male terminalia is in some extension similar to that of *Epicypta jackieyingae* Amorim & Oliveira, **sp. n.** They may correspond to a small clade in the genus.

*Epicypta khatijunae* Amorim & Oliveira, sp. n. (Figs. 128A–C)

https://singapore.biodiversity.online/species/A-Arth-Hexa-Diptera-000722and-002091

urn:lsid:zoobank.org:act:14F60D4D-C64B-45E3-A867-7576747C2863

**Diagnosis**. Head dark brown, antennal scape and pedicel ochre-yellowish, flagellum light brown. Scutum and scutellum blackish-brown, scutum with a slender, ochre-brown collar along anterior margin; pleural sclerites dark brown. Coxae whitish, with a brown band at basal end. Wing membrane light brownish, a large brownish area along anterior margin. C not extending beyond tip of R_5_; M_1+2_ shorter than r-m. Dorsal macrotrichia on posterior veins M_1_, M_2_, posterior half of M_4_, and distal end of CuA. Abdominal tergites 1–6 dark brown, tergite 6 with yellowish posterior margin, tergite 7 and terminalia yellowish. Female terminalia cercus digitiform, elongate. ***Epicypta_khatijunae_ZRCBDP0048068_hapZRCBDP00 48068_SMH_holotype* [1: C, 16: C, 23: T, 124: T, 127: G, 196: C, 199: T, 203: C, 205: C]**

CctttcatctacaatCgcccatTcaggatcatctgtagatttagctattttttctttacatttagctggtatttcttctattttgggggcaattaattttattactacaattattaatatacgTtcGccaggtatttcatttgatcgaatacctttatttgtatgatcagtattaattacagctgttttattactCctTtctCtCccagttttagccggagctattactatattattaacagatcgaaatttaaatacatctttttttgatcctgctggagggggtgatccaattttataccaacatttattt

#### Description

**Female** (Fig. 128A). Wing length, 2.58; width, 0.93 mm. **Head**. Vertex dark brown, two longer setae dorsally to eye anteriorly to lateral ocellus, one seta dorsally to ocellus, eight setae posteriorly to ocellus. Occiput dark brown, yellowish-brown towards ventral margin, a yellowish slender gena with dark brown ventral margin. Fine interommatidial setae over entire eye surface. Scape and pedicel light ochre, first two flagellomeres ochre-yellowish, remaining greyish-brownish. Frons, face, and clypeus brown; maxillary palpus with first two palpomeres yellowish-brown, distal three flagellomeres whitish; labella whitish-yellow. Scape twice length of pedicel, flagellomere 4 length 2.0× width. Maxillary palpomere 4 × palpomere 3, palpomere 5 1.5× palpomere 4 length. **Thorax**. Scutum shining blackish-brown, lighter on anterior margin. Pleural sclerites blackish-brown except for light brown mediotergite and metepisternum. Pleural membrane yellowish. Antepronotum with three bristles directed ventrally. Anepisternum with five bristles at posterior margin directed posteriorly. Mesepimeron with two bristles and 16 smaller setae; laterotergite with four bristles and eight smaller setae. Metepisternum with 12 setulae. Haltere whitish. **Legs**. Coxae and femora light ochre-yellowish, with a brown band basally, femora slightly darker with a brown tinge along dorsal edge, tibiae and tarsomeres dark ochre-yellowish, mid and hind tibiae greyish-brown at basal end. A group of four bristles and some long setae along frontal distal margin of fore coxa; a band of small fine setae across basal end of mid coxa, hind coxa with fine setae on basal half; femora with a short row of 4-5 brownish setae ventrally on distal end. Hind tibial spur over 5× tibia width at apex. Tarsal claws with an elongate pointed tooth coming out on inner margin medially. **Wing** (Fig. 128B). Membrane fumose light brown, darker cells c, br and r1. Humeral vein oblique, Sc virtually not produced. C barely produced beyond tip of R_5_. R_1_ long, reaching C on wing distal fifth; R_5_ reaching C slightly before level of tip of M_1_. First sector of Rs slightly oblique, devoid of setae; r-m more or less longitudinal, over 3× longer than first sector of Rs. M_1+2_ short, 0.67× r-m length; bM about 5× r-m length; first sector of CuA about 0.30× length of second sector of CuA. Cubital pseudovein absent, CuP weakly sclerotized, barely extending beyond level of origin of M_4_. Anal fold long, nearly reaching wing margin. Dorsal setae on posterior veins M_1_, M_2_, M_4_, and distal fourth of CuA. Some few dorsal macrotrichia on anal lobe membrane. **Abdomen**. Abdominal tergites 1-6 shining brown, posterior margin of tergite 6 and tergite 7 yellow; sternites 1-7 whitish, Sternite 2 with a pair of strong brown bristles. **Terminalia** (Fig. 128C). Dirty-yellow. Sternite 8 large, with a pair of long lateroposterior lobes close together, with a medial posterior incision, some stronger setae along posterior margin longer. Sternite 9 well-defined, tapered at anterior end, genital chamber slender, laterodistal margin close to tip with some fine setae, distal end acute. Tergite 8 large, subquadrate, covered with setae and microtrichia, posterior margin straight. Tergite 9+10 short, slender, with sclerotized band along anterior margin extending anteriorly to lateral apodemes directed anteriorly, entirely bare of setae. Cercomeres 1 and 2 probably fused, cerci very long, about 7× longer than wide.

**Male**. Unknown.

#### Material examined

**Holotype**: female, ZRCBDP0048068, Nee Soon (NS1), swamp forest, 12.jun.2013, MIP leg. (slide-mounted). **Paratypes** (5 females): ZRCBDP0048322, Nee Soon (NS2), swamp forest,

20.jun. 12, MIP leg. (website photo specimen); ZRCBDP0048426, Nee Soon (NS1), swamp forest, 09.may. 12, MIP leg. (website photo specimen); ZRCBDP0048431, Nee Soon (NS2), swamp forest, 18.jul. 12, MIP leg. (website photo specimen); ZRCBDP0048749, Nee Soon (NS1), 25.feb.15, MIP leg.; ZRCBDP0048861, Nee Soon (NS1), 14.jan.15, MIP leg.

**Etymology**. The species epithet of this species honors Khatijun Nissa Siraj (1925-), a Singaporean women’s rights activist and the co-founder of the Young Women’s Muslim Association (PPIS) and the Muslim Women’s Welfare Council in 1964. In response to an epidemic of women in the Singaporean Muslim community being abandoned through inexpensive and easy divorces, she pressed for the formation of a Syariah Court, and served as its first caseworker in the 1960s. She was inducted into the Singapore Women’s Hall of Fame in 2014.

**Remarks**. There are three haplotypes for *Epicypta khatijunae* Amorim & Oliveira, **sp. n.** and all delimitation approaches point to a single species.

*Epicypta purchoni* Amorim & Oliveira, sp. N. (Figs. 129A–D)

https://singapore.biodiversity.online/species/A-Arth-Hexa-Diptera-000786

urn:lsid:zoobank.org:act:F0E2CF0F-8234-41E0-862B-053388899FA5

**Diagnosis**. Head, scutum, scutellum and thoracic pleural sclerites blackish-brown, antenna light brown, scape, pedicel, and first flagellomere lighter. Coxae whitish, except for brown band on proximal fourth of hind coxa; femora whitish, except for proximal and distal ends of mid femur and for a brown line along dorsal edge of hind femur. Wing membrane light brownish, area along anterior margin slightly darker. C extending shortly beyond tip of R_5_. First section of Rs short, 0.46× r-m length; M_1+2_ short, 0.27× of r-m length. Dorsal macrotrichia on posterior veins M_1_, M_2_, and posterior half of M_4_, anal lobe membrane with macrotrichia. Abdominal tergites 1–2 dark brown, tergites 3–5 greyish-brown medially, with wide lateral cream-yellow bands, tergites 6–7 greyish-brown. Gonocoxites with a long lateral digitiform lobe and a long, blade-like dorsal lobe; gonostylus flat, small, displaced medially; paramere with a pair of strong spines on a triangular projection distally; tergite 9 with a pair of long parallel projections.

***Epicypta_purchoni_ZRCBDP0047803_hapZRCBDP004 7803_SMH_holotype* [10: A, 61: C, 118: C, 130: C, 199: T, 251: C, 253: T] -**

ctatcttcAactattgctcatgcaggagcatcagtagatttagctattttttctcttcaCttagctggtatttcttcaattttaggagctattaattttattactacaattattaaCatgcgatctccCgggatttcttttgatcgaatacctttatttgtttgatctgttttaattacagcaattcttttattactTtcattaccagtattagctggagct attactatattattaacagatcgtaatCtTaatacatctttttttgatcctgcaggagggggagacccaattttatatcaacatttattt

#### Description

**Male**. Wing length, 2.37; width, 0.86. **Head**. Vertex dark brown, occiput dark caramel-brown. Scape and pedicel ochre-yellow, flagellomere 1 dark yellow-brown, remaining greyish-brown. Face and clypeus light caramel-brown; palpomeres 1–3 light brownish, palpomeres 4–5 lighter; labella whitish-yellow. Occiput with three longer setae dorsally to eye anteriorly to lateral ocellus, two dorsally to ocellus, six posteriorly to ocellus. Scape about × pedicel length; flagellomere 4 1.6× longer than wide.

Palpomere 4 1.5× palpomere 3 length, palpomere 5 1.6× palpomere 4 length. **Thorax**. Scutum blackish-brown. Pleural sclerites dark brown. Pleural membrane ochre-yellow. Haltere whitish-yellow. Scutum with seven supra-alar and three pairs of prescutellar bristles; two pairs of scutellar bristles and one additional outer pair of long setae. Proepisternum with three bristles, anepisternum with five bristles along posterior margin. Mesepimeron with two bristles and 23 small setae, laterotergite with three bristles and 16 small setae. Metepisternum with 16 fine setae. **Legs**. Coxae whitish-yellow with orangish tinge, hind coxa with a brown band on basal fifth; femora, tibiae and tarsi whitish-yellow with orangish tinge, tibiae, and tarsi darker. Mid tibia with a band of fine setae across proximal fourth, hind coxa covered with setulae on basal third, extending posteriorly close to anterior margin. Fore tibia with two dorsolateral bristles; mid tibia with two dorsolateral rows of four short bristles, a pair of lateral bristles and a row with four long setae along ventral edge; hind tibiae with a pair of dorsolateral rows of six bristles and four long setae on inner face. Fore leg tarsomere 1 1.1× tibia length, 1.5× tarsomere 2 length. Hind leg inner tibial spur 5.0× tibia width at apex. **Wing** (Fig. 129B). Membrane light brown fumose, darker along anterior margin. Sc faint, with dorsal macrotrichia (visible only with phase contrast). C extending beyond apex of R_5_ for about a ¼ of distance to M_1_. R_1_ reaching C at wing distal fourth; R_5_ reaching C slightly before level of M_1_. First sector of Rs transverse, bare, 0.46× r-m length; r-m almost longitudinal. M_1+2_ almost absent, 0.27× r-m length; bM 3.1× r-m length. First sector of CuA 0.35× length of second sector of CuA. Cubital pseudovein inconspicuous, sclerotized, CuP reaching level of origin of M_4_. Anal fold almost reaching wing margin, curved along most of its length. Posterior veins M_1_ and M_2_ with of dorsal macrotrichia on almost entire length, M_4_ on distal half and CuA on distal fifth of wing; no macrotrichia on anal lobe membrane. **Abdomen**. Abdominal tergite 1 greyish-brown with a dark brown slender band across posterior end, tergites 2-5 brown medially with yellowish-brown to cream-yellow lateral bands, tergites 6–7 light brown; sternites 1-7 light brownish-yellow. Sternite 2 with a pair of long, slightly curved bristles. **Terminalia** (Fig. 129C). Light brown, cerci lighter. Gonocoxites fused medially, no suture present, bare ventrally, a medioventral digitiform projection with setulae at tip, reaching level of tip of gonostylus, sided by a pair of shorter digitiform projections, each with an elongate seta at tip, a pair of long digitiform laterodistal extensions reaching way beyond tip of gonostylus, a large lobe just external to base of gonostylus, with setae on both faces, setae on dorsoposterior margin longer, curved, extending to level of inner arms of aedeagus, gonocoxites fused medially on dorsal face, no suture left. Gonostylus simple, elongate, displaced to a more medial position, dorsoventrally compressed, with setae on both faces, ventral setae on basal half longer than distal setae. Aedeagus with an anterior ejaculatory apodeme, widening midway to apex, then divided distally into a pair of lobes in contact medially and extending distally beyond tip of gonostylus, gonopore apparently short, medially; parameres present as a triangular sclerite extending distally, with a medial short beak at tip, two pairs of small setae close to each other subapically and a pair of strong curved spines close to tip, laterally with a pair of long, slender projections extending beyond tip of gonocoxite lateral lobes, each with five setulae on distal end. Gonocoxal bridge not evident. Tergite 9 present as a pair of long laterodistal digitiform extensions, entirely disconnected from each other, about as long as gonocoxite lateral extensions, setose at tip. Cerci weakly sclerotized between tergite 9 elongate lateral lobes. **Female** (Fig. 129A). As male, except for the following. **Wing l**ength, 2.21–2.69; width, 0.82–0.99. **Head**. Occiput with three longer setae dorsally to eye anteriorly to lateral ocellus, one seta dorsally to ocellus, ten setae posteriorly to ocellus. **Thorax**. Scutum with seven supra-alar bristles and three pairs of prescutellars; three pairs of scutellar bristles, outer pair smaller. Proepisternum with three bristles directed ventrally, anepisternum with four bristles along posterior margin. Mesepimeron with two bristles and 30 small setae and setulae, laterotergite with three bristles and 18 small setae. Metepisternum with 16 small fine setae. **Legs**. Fore tibia with two strong setae at outer face, mid tibia with two dorsolateral rows of 5–6 bristles, one outer bristle and three bristles along ventral edge, hind tibia with a row of four dorsal bristles, one lateral subdistal strong seta and three setae, one bristle along ventral edge. **Terminalia** (Fig. 129D). Light brownish-yellow. Sternite 8 elongate, trapezoid, posterior margin nearly straight, microtrichia and setae evenly distributed, long setae at posterior margin. Sternite 9 slender, elongate, anterior apodeme extending to slightly beyond anterior end of terminalia, widening at apex, a pair of sclerotized bands extending into lateral arms, genital chamber elongate. Tergite 8 with a pair of separate large lateral lobes touching medially, microtrichia and setae on lateral lobes. Tergite 9+10 slender, a pair of sclerotized lateral bands, lateroventrally fused to sternite 9. Cercomeres 1 and 2 fused, no sign of suture, elongate, wider midway to apex posterior end digitiform, distal setae longer.

#### Material examined

**Holotype**: male, ZRCBDP0047803, Nee Soon (NS2), swamp forest, 13.nov.13, MIP leg. (slide-mounted). **Paratype** (1 male): ZRCBDP0047857, Nee Soon (NS1), swamp forest, 10.jul.13, MIP leg. (website photo specimen, slide-mounted) (MZUSP).

**Possibly non-conspecific cluster**: female, ZRCBDP0047813, Nee Soon (NS2), swamp forest, 26.mar.14, MIP leg. (slide-mounted).

**Etymology**. The species epithet of this species honors Professor Richard Denison Purchon (1916–1992), first Head of the Department of Zoology, later merged with the Department of Botany to constitute the present Department of Biological Sciences, National University of Singapore. A specialist in marine biology, he was a recognized authority in malacology, with a famous book *The Biology of the Mollusca*.

**Remarks**. *Epicypta purchoni* Amorim & Oliveira, **sp. n.** has three haplotypes, one of which with a single male and another one with a single female. They are split into two mOTUs by OC p-distance 2% and ABGD (p=0.001-0.005), but kept together by all other delimitation approaches. This as another grey-zone case. We keep two of the haplotypes together in a single species. The third haplotype subcluster only includes only a female, not included as a paratype. *Epicypta purchoni* Amorim & Oliveira, **sp. n.** has overall similar male terminalia as *Epicypta wallacei* Amorim & Oliveira, **sp. n.** and *Epicypta peterngi* Amorim & Oliveira, **sp. n.**

*Epicypta foomaoshengi* Amorim & Oliveira, sp. n. (Figs. 130A–D, 131A-C)

https://singapore.biodiversity.online/species/A-Arth-Hexa-Diptera-000807and-002087

urn:lsid:zoobank.org:act:FEB0EE0C-65F3-41A8-9C52-3A299765980D

**Diagnosis**. Head and antenna ochre-yellowish; scutum brownish, with a dark ochre-yellowish diffuse area along anterior third, scutellum dark brown; pleural sclerites dark brown, with ochre-yellowish area on antepronotum and on ventroposterior end of anepisternum. Coxae and femora whitish, on fore leg with brownish light tinge, a dark brown line along posterior border of anterior coxa, fore femur with a blackish brown mark at proximal end and along ventral crest. Wing membrane light brownish, basal third of cell c and cell br with a brown mark. C slightly produced beyond tip of R_5_; M_1+2_ nearly absent. Dorsal macrotrichia on posterior veins M_1_ and M_2_, M_4_, and on CuA distal fourth, microtrichia present on anal lobe membrane. Abdominal tergite 1 ochre-yellow with a brown transverse band close to posterior margin, tergites 2–5 mostly light brown with a yellowish-brown band on anterior third; tergite 6 bright cream-yellow, terminalia bright ochre-yellow. No medioventral male syngonocoxite projection; gonocoxite with a long lateroposterior lobe, bifid midway to apex; gonostylus with a more ventral rhomboid branch and a more dorsal digitiform projection; parameres with a pair of long falciform blades; tergite 9 present as a pair of long digitiform extensions entirely disconnected from each other. Female terminalia with a wide, short incision on posterior margin of sternite 8.

***Epicypta_foomaoshengi_ZRCBDP0284196_hapZRCBD P0049068_holotype* [29: T, 56: C, 148: T, 149: T, 196: T, 199: T, 220: T, 223: A, 277: A] -**

ctttcatctactattgctcatgcaggaTcttctgttgatttagctattttttctCttcatttagctggtatttcttcaattttaggagctattaattttattactacaattattaatatacgatctcctggaatttcttttgatcgTTtacctttatttgtatgatcagtattaattactgctattcttttactTctTtctttacctgttttagcaggTgcAattactatactattaacagatcgaaatttaaatacttctttttttgatcctgcGggaggaggggatcctattttatatcaacatttattt

#### Description

**Male** (Fig. 130A). Wing length, 2.82; width, 0.98. **Head**. Caramel-brown (Fig. 131A). Scape and pedicel light ochre-yellowish, flagellomeres light brownish-yellow. Face and clypeus light caramel-brown. Maxillary palpomeres 1–3 light brownish-yellow, palpomeres 4–5 lighter; labella whitish-yellow. Occiput with three longer setae dorsally to eye anteriorly to lateral ocellus, two dorsally to ocellus, seven posteriorly to ocellus. Scape 1.7× pedicel length, flagellomere 4 1.5× longer than wide.

Palpomere 4 1.5× palpomere 3 length, palpomere 5 1.6× palpomere 4 length. **Thorax**. Scutum caramel-brown, anterior third ochre-yellowish; scutellum dark brown with ochre-yellow anterolateral corners. Antepronotum ochre-yellow on dorsal end, greyish-brown ventrally, proepisternum greyish-brown, anepisternum dark greyish-brown with an ochre-yellow area medially along posterior margin, katepisternum, mesepimeron, laterotergite, and metepisternum dark greyish-brown, mediotergite dark brown. Pleural membrane yellowish. Scutum with 4+2 supra-alars and three pairs of prescutellar bristles. Proepisternum with four bristles (ventroposterior one smaller), anepisternum with six bristles along posterior margin (two dorsal ones smaller). Mesepimeron with two bristles and 12 fine setae, laterotergite with four bristles and four setae. Metepisternum bare. Haltere pedicel and most knob light whitish-ochre, base of knob brownish, no larger setae. **Legs**. Fore coxa whitish with a brownish tinge, brownish along posterior margin and a brown mark at tip; mid and hind coxae whitish, both with a light brown band basally, dark on hind coxa. Femora whitish with a grey-brownish tinge, brownish at proximal end, darker along dorsal and ventral margin; tibiae and tarsi slightly darker, tips of mid and hind tibiae darker. Mid coxa with some fine setae across proximal fifth, hind coxa with a band of setulae at basal fourth. Fore tibia, besides regular rows of fine trichia, with two dorsolateral small bristles medially; mid tibia with three irregular dorsal rows of 3–5 bristles, one small bristle on distal third of inner face, and one bristle and four strong setae along ventral margin; hind tibia, besides regular rows of fine trichia, with a regular line of small bristles dorsally along entire length, a row of brown trichia at outer face, and three irregular dorsal rows of bristles and six small bristles on inner face.

Fore tibia tarsomere 1 1.1× tibia length, 1.9× tarsomere 2 length. Hind leg tibial inner spur 6.0× tibia width at apex. **Wing** (Fig. 130B). Membrane fumose brown, darker along anterior margin, an elongate brownish mark on cell c to origin of Rs, on cell br and around base of M_1+2_. C produced slightly beyond tip of R_5_. Sc barely produced. R_1_ reaching C on distal fifth of wing; R_5_ reaching C before level of tip of M_1_. First sector of Rs transverse, 0.36× r-m length; r-m almost longitudinal. M_1+2_ nearly absent, M_1_ forking from M_2_ almost directly from r-m; bM 3.3× r-m length; first sector of CuA short, 0.23× length of second sector of CuA. Cubital pseudovein absent, CuP extending to slightly beyond level of origin of M_4_. Anal fold long, almost reaching wing margin. Posterior veins M_1_ and M_2_ almost entirely with dorsal macrotrichia, M_4_ with macrotrichia on distal half, macrotrichia on anal lobe membrane. **Abdomen**. Abdominal tergite 1 ochre-yellow, with a brown transverse band close to posterior margin, tergites 2–5 light brown on posterior half, yellowish-brown on anterior half, lateral margins more yellowish; tergite 6 bright cream-yellow; sternites 1-7 whitish-yellow. Tergites 1–5 with dark brown elongate setae on dorsal part of sclerites. Sternite 2 with a pair of strong brown bristles. **Terminalia** (Figs. 131B–C). Light brown. Gonocoxites connected medially at anterior end of terminalia, no medioventral projection; a long lateroposterior lobe bifid midway to apex, ventral branch triangular, blade-like, posterior branch falciform, acuminating towards apex, extending to slightly before level of tip of parameres, gonocoxites fused medially on dorsal face through a slender connection. Gonostylus with two arms, a more ventral rhomboid branch and a more dorsal digitiform projection, ventral arm more setose, distal arm with setae on distal fifth. Aedeagus with an anterior inverted T-shape anterior process extending posteriorly into a triangular medial process extending to level of tip of gonostylus branches, anterior end rounded; parameres present as a pair of long, slender falciform blades extending distally to level of tip of gonocoxite laterodistal process. Gonocoxal bridge not evident. Tergite 9 present as a pair of long digitiform extensions, slightly wider at base, entirely disconnected from each other, longer than gonocoxite lateral extensions, setose at entire length. Cerci weakly sclerotized between tergite 9 elongate lateral lobes.

**Female**. As male, except for the following. **Terminalia** (Figs. 130C–D). Cream-yellow. Sternite 8 rectangular, elongate, lateral ends of posterior margin slightly more projected than medially, microtrichia and small setae widespread, posterior margin with three pairs of longer setae, those on lateroposterior corners stronger. Sternite 9 elongate, anterior apodeme widening close to tip, genital chamber elongate, distal end acute. Tergite 8 wide, lateral lobes projected posteriorly, medial connection slender, entirely bare. Tergite 9+10 present as a pair of slender sclerotized inclined band, connected to sternite 9 lateroventrally. Cercus large, cercomeres 1 and 2 fused, no sign of suture, basal ⅔ wider, posterior end digitiform, distal setae longer.

#### Material examined

**Holotype**: male, ZRCBDP0284196, Kranji Marshes (KM03), freshwater swamp, date range 2018-2019, MIP leg. **Paratypes** (9 females): ZRCBDP0047057, National University of Singapore (PGP), 17.jun. 2015, MIP leg. (slide-mounted); ZRCBDP0047058, National University of Singapore (PGP), 17.jun. 2015, MIP leg.; ZRCBDP0047075, National University of Singapore (PGP), 15.jul. 2015, MIP leg.; ZRCBDP0047076, National University of Singapore (PGP), 15.jul. 2015, MIP leg.; ZRCBDP0048770, National University of Singapore (PGP), 20.may. 2015, MIP leg.; ZRCBDP0049066, National University of Singapore (PGP), 27.may. 2015, MIP leg.; ZRCBDP0049068, National University of Singapore (PGP), 27.may. 2015, MIP leg. (website photo specimen); ZRCBDP0049326, National University of Singapore (PGP), 10.jun. 2015, MIP leg.; ZRCBDP0049330, National University of Singapore (PGP), 10.jun. 2015, MIP leg. **Additional sequenced specimens:** ZRCBDP0078989, Singapore; ZRCBDP0279153, Singapore, 05.jun.18; ZRCBDP0279154, Singapore, 05.jun.18; ZRCBDP0279163, Singapore, 26.apr.18; ZRCBDP0279170, Singapore, 26.apr.18; ZRCBDP0284190, Singapore; ZRCBDP0284299, Singapore, 26.apr.18; ZRCBDP0284302, Singapore, 26.apr.18.

**Etymology**. The species epithet of this species honors Foo Maosheng (1988-), a curator at the Lee Kong Chian Natural History Museum, specializing in Blattodea. He is one of the researchers associated with the Singapore Mangrove Insect Project and helped the development of this study of the Mycetophilidae in different occasions.

**Remarks**. All specimens of the type-series are from the National University of Singapore campus, which has patches of secondary forest. *Epicypta foomaoshengi* Amorim & Oliveira, **sp. n.** has five haplotypes that are clustered into one mOTU by all delimitation approaches. There is also no support from morphology to split the cluster into separate species.

*Epicypta ganengsengi* Amorim & Oliveira, sp. n. (Figs. 132A–C, 133A–B)

https://singapore.biodiversity.online/species/A-Arth-Hexa-Diptera-000795

urn:lsid:zoobank.org:act:B9A73545-2327-495E-8937-10C764228B43

**Diagnosis**. Head brown, antennal scape and pedicel ochre-yellowish, flagellum greyish-brown; scutum dark brown, a diffuse dark ochre-yellowish area along anterior margin, scutellum blackish-brown; pleural sclerites dark brown, antepronotum and proepisternum dark ochre-yellowish. Coxae and femora whitish, on fore leg with brownish light tinge, mid and hind coxae with a brown band on proximal end, tip brownish. Wing membrane light brownish, cells c and br darker. C barely produced beyond tip of R_5_; M_1+2_ 0.58× r-m length. Dorsal macrotrichia on posterior veins M_1_ and M_2_, M_4_, and on CuA distal fourth; macrotrichia on anal lobe membrane. Abdominal tergites 1–6 dark brown, tergite 7 brownish-yellow. Gonocoxites fused medially, with a short and wide medioventral projection, a pair of laterodistal projections over five times longer than length of syngonocoxite medially, with scattered fine setae and a couple of subdistal strong setae and some longer setae at tip. Gonostylus simple, digitiform. Aedeagus rhomboid, bare, well sclerotized, parameres weakly sclerotized, with a pair of strong, straight spines close together medially. Tergite 9 present as a pair of wide posterior lobes covered with microtrichia and setae, straight on outer margins, wider at base. Female terminalia with short incision on posterior margin of sternite 8.

***Epicypta_ganengsengi_ZRCBDP0133402_hapZRCBDP 0132824_SMH_holotype* [28: T, 85: G, 133: T, 146: A, 1 72: C, 193: T, 277: G]**

tctttcatctactattgctcacgcaggTtcttctgtagatttagctattttttctcttcatttagcaggaatttcttcaattttGggagctattaattttattactactattatcaatatacgagcccctggTatttcttttgatAaaatacctttatttgtttgatcagtCttaattacagctattttactTcttttatctttacctgttttagctggagctattactatattattaacagatcgaaatttaaatacttctttttttgaccctgcGggagggggagaccctattttatatcaacatttattt

#### Description

**Male**. Wing length, 2.38; width, 0.83. **Head**. Vertex dark caramel-brown, occiput dark caramel-brown. Scape and pedicel ochre-yellowish, first two flagellomeres light brown, remaining yellowish-brown at basal third, light brown at distal two-thirds. Face and clypeus light caramel-brown. Maxillary palpomeres light brownish. Scape 1.6× pedicel length; flagellomere 4 1.5× longer than wide, covered with scattered whitish setae. Face and clypeus light caramel-brown; face covered with short darker setae, clypeus short, densely covered with setulae. Palpomeres light brownish; labella whitish-yellow. Occiput with three longer setae dorsally to eye anteriorly to lateral ocellus, one dorsally to ocellus, seven posteriorly to ocellus. Scape 1.6× pedicel length, flagellomere 4 1.5× width. Palpomere 4 1.1× palpomere 3 length, palpomere 5 2.3× palpomere 4 length. **Thorax**. Scutum dark caramel-brown, scutellum blackish-brown. Antepronotum and proepisternum light caramel-brown, anepisternum dark caramel-brown, light along anterior margin, other sclerites greyish-brown, mediotergite dark brown. Pleural membrane ochre-yellow. Haltere ochre-yellowish. Scutum with 5+1 supra-alars (first small) and three pairs of prescutellar bristles, scutellum with two pairs of bristles along posterior margin. Proepisternum with three bristles, anepisternum with five bristles along posterior margin. Mesepimeron with two bristles and 16 small setae, laterotergite with two long setae and seven small setae. Metepisternum with three fine setae. **Legs**. Coxae light ochre-yellowish, mid and hind coxae with a brown band basally, hind coxa with a brown posterior mark distally; femora light ochre-yellowish with a brown tinge along dorsal and ventral edges; tibiae ochre-yellowish, mid and hind tibiae greyish-brown at basal end, yellowish-brown at tip, tarsi light greyish-brown. Fore tibia with one strong seta on distal third of inner face; mid tibia with three irregular dorsolateral rows of 3–5 small bristles, one strong seta on distal third of inner face; hind tibia with three irregular rows of 3–5 dorsolateral small bristles and five larger setae on distal third of inner face. Fore leg tarsomere 1 1.2× tibia length [tarsomeres 2–5 broken]. Hind tibial inner spur over 4.6× tibia width at apex. **Wing** (Fig. 132B). Membrane light brown fumose. C not extending beyond apex of R_5_. Sc barely produced. R_1_ reaching C on distal fifth of wing; R_5_ reaching C slightly before level of tip of M_1_. First sector of Rs nearly transverse, 0.52× r-m length; r-m almost longitudinal. M_1+2_ 0.58× r-m length; bM about 4.7× r-m length; first sector of CuA about 0.18× length of second sector of CuA. Cubital pseudovein absent, CuP extending slightly beyond level of origin of M_4_. Anal fold gradually curved, nearly reaching wing margin. Posterior veins M_1_ and M_2_ with dorsal macrotrichia on most of their length, M_4_ with macrotrichia on distal ¾, CuA with macrotrichia on distal fourth, macrotrichia on anal lobe membrane. **Abdomen**. Abdominal tergites 1-6 brown, posterior margin of tergite 6 yellowish, tergite 7 yellowish; sternites 1-7 whitish. Sternite 2 with a pair of strong brown bristles. **Terminalia** (Figs. 133A–B). Gonocoxites fused medially, no suture present, bare ventrally, a large and short medioventral projection, a laterodistal projection dorsally to insertion of gonostylus at each side that is over five times longer than length of syngonocoxite medially, with scattered fine setae along entire length, a couple of subdistal strong setae and some longer setae at tip. Gonostylus simple, digitiform, with fine setae, slightly stronger distally. Aedeagus rhomboid, bare, well sclerotized, parameres weakly sclerotized, projecting posteriorly beyond aedeagus, with a pair of strong, straight spines close together medially. Gonocoxal bridge wide, apodemes as a pair of long slender sclerotized bands. Tergite 9 present as a pair of wide posterior lobes covered with microtrichia and setae, straight on outer margins, wider at base. Cerci not recognizable.

**Female** (Fig. 132A). As male, except for the following. **Terminalia** (Fig. 132C). Light brownish-yellow, cerci lighter. Sternite 8 trapezoid, elongate, posterior margin straight, with a rather deep medial incision, widespread microtrichia and small setae, posterior margin with longer setae. Sternite 9 with anterior apodeme extending beyond anterior end of terminalia, widening close to tip, genital chamber bell-shaped, lateral arms wide, distal end triangular, acute, bare. Tergite 8 wide, lateral lobes projected posteriorly with setae, with a slender medial connection. Tergite 9+10 present as a pair of slender sclerotized inclined band, connected to sternite 9 lateroventrally. Sternite 10 triangular, with some setae close to tip directed dorsally. Cercus large, cercomeres 1 and 2 fused, no sign of suture, basal ¾ wider, posterior end digitiform, distal setae longer.

#### Material examined

**Holotype**: male, ZRCBDP0133402, National University of Singapore (Uhall), 17.may.2017, MIP leg. (slide-mounted). **Paratypes** (5 females): ZRCBDP0049044, Nee Soon (NS2), 07.jan.15, MIP leg.; ZRCBDP0049300, National University of Singapore (Icube), 01.apr.15, MIP leg. (website photo specimen, slide-mounted); ZRCBDP0072662, Bukit Timah, primary forest (BT05), 28.dec. 16, MIP leg.; ZRCBDP0072722, Bukit Timah, old secondary forest (BT01), 14.dec. 16, MIP leg. (slide-mounted); ZRCBDP0279158, Singapore, 7.jun.18, MIP leg. (slide-mounted). **Additional sequenced specimens**: ZRCBDP0132824, National University of Singapore (PGP), urban forest, 24.may. 17; ZRCBDP0133115, National University of Singapore (PGP), urban forest, 17.may. 17; ZRCBDP0133129, National University of Singapore (PGP), urban forest, 17.may. 17; ZRCBDP0133169, National University of Singapore (Uhall), urban forest, 26.apr. 17, MIP leg.; ZRCBDP0133378, National University of Singapore (PGP), urban forest, 14.may. 17; ZRCBDP0133417, National University of Singapore (PGP), urban forest, 21.may. 17; ZRCBDP0133490, National University of Singapore (PGP), urban forest, 31.may. 17; ZRCBDP0133504, Singapore, date range 2012-2018, MIP leg.; ZRCBDP0133552; ZRCBDP0279142, Singapore, date range 2012-2018, MIP leg.; ZRCBDP0279159, Singapore, date range 2012-2018, MIP leg.; ZRCBDP0284194, Kranji Marshes (KM03), freshwater swamp, date range 2018-2019, MIP leg; ZRCBDP0133158, National University of Singapore (RVR), 03.may.17; ZRCBDP0133945, Singapore.

**Etymology**. The species epithet of this species honors Gan Eng Seng (1844–1899), businessman and philanthropist, an early pioneer in Singapore. He is known for his generosity to many charitable causes in Malaya and Singapore during the British colonial era. He founded in 1885 the first school established by overseas Chinese in Singapore, which is one of the oldest educational institutions in the nation-state.

**Remarks**. There are five haplotypes for *Epicypta ganengsengi* Amorim & Oliveira, **sp. n.,** clustering together into one mOTU by all species delimitation methods. Specimens were collected in the urban forests and the swamp forests.

*Epicypta* sp. A (Figs. 134A-B)

#### Description

**Male**. Wing length, 2.66; width, 0.96. **Head**. Dark brown, antenna brown, scape and pedicel slightly lighter. **Thorax** (Fig. 134A). Scutum and scutellum dark brown; pleural sclerites dark brown. Haltere with light brown pedicel, knob whitish with light brownish tinge. Pleural membrane ochre-yellowish. Scutum with setae, but no bristles except for nine supra-alars and three pairs of prescutellar bristles; three pairs of scutellar bristles and an additional pair of longer setae on margin outer end. Proepisternum with four long bristles, anepisternum with five bristles along posterior margin. Mesepimeron with two bristles and 11 fine small setae, laterotergite with five bristles and some few additional fine setae. **Legs**. Coxae yellowish, fore coxa with a brownish tinge, all three hind coxae with a brown proximal band; femora yellowish with a brownish tinge, fore femur with a light brown band along dorsal and ventral edges, mid and hind femora with a slender brown band along ventral edge and a brown area at proximal and distal ends, mid femur with a row of small, conspicuous setae along entire dorsal edge. **Wing** (Fig. 134B). Membrane light brownish, darker on cells c and br. C not produced beyond tip of R_5_; Sc short, incomplete. R_1_ reaching C on distal fifth of wing; R_5_ reaching beyond level of tip of M_1_. First sector of Rs transverse, 0.30× r-m length; r-m almost longitudinal. M_1+2_ 0.35× r-m; bM 2.9× r-m length; first sector of CuA about 0.34× length of second sector of CuA. Cubital pseudovein absent, CuP not reaching level of origin of M_4_. Anal fold gently curved along basal three-fourth. Veins bR, R_1_, R_5_, distal half of r-m, M_1_, M_2_, M_4_, and distal third of CuA with dorsal macrotrichia; anal lobe membrane with dorsal macrotrichia. **Abdomen**. Abdominal tergites 1-7 dark-brown, sternite 1–7 light brown. **Terminalia** (Figs. 134C– D). Gonocoxite short, a pair of long lateral setose lobes projecting beyond tip of gonostylus, dorsal lobe digitiform, long, mostly bare, a modified distal short spine. Gonostylus dorsoventrally compressed, capitate, with small setae distally. Aedeagus with a pair of parallel blades close to each other; parameres with a pair of long, distal spines close to each other. Tergite 9 with a pair of long, digitiform extensions, not reaching level of tip of gonocoxite lateral lobes.

**Material examined** (male): ZRCBDP0132893, Singapore, date range 2012-2018, MIP leg. (slide-mounted, sequence failure).

**Remarks**. This is one of the species of *Epicypta* recognized based only on morphological data—the only known specimen corresponds to a specimen that failed to sequence. We describe the species, but do not formally name it. The male terminalia morphology shows obvious differences from all remaining species of *Epicypta*, particularly with regard to the pair of elongate distal spines of the parameres, the flat, capitate gonostyle and the typical shape of the aedeagus, with sigmoid laterals.

*Epicypta nanyangu* Amorim & Oliveira, sp. n. (Figs. 135A–D, 136A–B)

https://singapore.biodiversity.online/species/A-Arth-Hexa-Diptera-000736

urn:lsid:zoobank.org:act:EABE5AC7-C89C-4C8A-904C-658433DE36FB

**Diagnosis**. Head and scutum dark ochre-yellowish, scutellum brown; pleural sclerites greyish brown. Coxae and femora whitish, anterior leg with orangish tinge, hind coxa with a large brown area basally, hind femur conspicuously brownish along dorsal and ventral crest. Membrane fumose light brown, slightly darker on cells c, br and r1. C extending beyond tip of R_5_; M_1+2_ 0.71× r-m length, r-m almost longitudinal. Dorsal macrotrichia on posterior veins M_1_ and M_2_, and on M_4_ close to tip; no macrotrichia on anal lobe membrane. Abdominal tergites 1–2 light brown, tergites 3–5 light brown with cream-yellow laterals, wider on tergites 4–5, tergite 6 light brown on anterior half, cream-yellow on posterior half, tergite 7 cream-yellow, terminalia whitish-yellow. Female terminalia sternite 8 with no medial incision along posterior margin.

***Epicypta_nanyangu_ZRCBDP0048844_hapZRCBDP00 47903_SMH_holotype* [19: A, 37: C, 70: G, 130: T, 155: C, 190: C, 202: C]**

cctttcttcaacaattgcAcacgcaggagcttctgtCgatttagcaattttttctcttcatttagctggGatttcttctattttaggggctattaattttattactacaattattaatatacgagctccTggaatttcttttgatcgaatacctCtttttgtttgatcagttttaattacagctattctCttattattatcCttacctgtattagcaggtgctattacaatattattaacagatcgaaatttaaatacatcattttttgaccctgcaggagggggagaccctattttatatcaacatttattt

#### Description

**Female** (Fig. 135A). Wing length, 2.02; width, 0.74. **Head**. Head dark ochre-yellowish. Scape and pedicel ochre-yellowish, flagellomeres light greyish-yellow. Face and clypeus ochre-yellow, labella whitish-yellow. Maxillary palpomeres 1–3 light brownish-yellow, palpomeres 4–5 lighter. Occiput with three longer setae dorsally to eye anteriorly to lateral ocellus, two dorsally to ocellus, seven posteriorly to ocellus. Scape about 1.5× pedicel length; flagellomere 4 2.0× longer than wide. Palpomere 4 1.2× palpomere 3 length, palpomere 5 1.5× palpomere 4 length. **Thorax**. Scutum dark ochre-yellow; scutellum brown with ochre-yellow anterolateral corners. Scutum with five supra-alars and three pairs of prescutellar bristles; two pairs of scutellar bristles. Most antepronotum, proepisternum, anepisternum, katepisternum mesepimeron, laterotergite, and metepisternum greyish ochre-brown, dorsoposterior corner ochre-yellow; mediotergite greyish ochre-brown dorsally, lighter ventrally. Haltere light ochre-yellowish, dark at base of knob, no larger setae. Pleural membrane yellowish. Proepisternum with three bristles, anepisternum with five bristles along posterior margin. Mesepimeron with two bristles and 13 smaller setae, laterotergite with one bristle and six smaller setae. Metepisternum 24 small fine setae. **Legs**. Fore coxa whitish-yellow with brownish tinge, mid and hind coxae whitish, hind coxa with dark brown basal fourth. Femora whitish-yellow with brownish tinge, a brown line along dorsal and ventral edges, hind femur with light brown distal fifth; tibiae and tarsi light ochre-brown, tarsi darker. Fore tibia with two dorsal strong setae, mid tibia with two dorsolateral rows with 4–5 bristles, five bristles ventrally and one laterally on inner face, hind tibia with two dorsolateral rows with four bristles, three bristles on distal half of lateral external face. Mid and hind tarsomeres with rows of lateral and some few ventral setae. Fore tibia 1.3× length of tarsomere 1, 1.7× tarsomere 2 length. Hind tibia inner spur over 4× tibia width at apex. **Wing** (Fig. 135B) Membrane fumose light brown, darker on cells c, br and r1. R_1_ reaching C on distal fourth. C not extending beyond tip of R_5_; R_5_ reaching C at level of tip of M_1_. First sector of Rs slightly oblique, 0.77× r-m length; r-m slightly oblique. M_1+2_ 0.71× r-m; bM slightly over 7× r-m length. First sector of CuA about 0.35× length of second sector of CuA. Cubital pseudovein absent, CuP ending at about level of origin of M_4_. Anal fold gently curved, almost reaching wing margin. Dorsal macrotrichia on posterior veins M_1_ and M_2_ on most of their length, M_4_ on distal half, CuA close to apex, some few macrotrichia on anal lobe membrane. **Abdomen**. Abdominal tergites 1–2 light brown, tergite 3 mostly light brown with a small cream-yellow area laterodistally; tergites 4–5 light brown medially with wide cream-yellow area laterally, tergite 6 light brown on anterior half, cream-yellow on posterior half, tergite 7 cream-yellow; sternites 1-7 whitish-yellow, sternite 2 with a pair of strong brown bristles. **Terminalia** (Figs. 135C– D). Whitish-yellow. Sternite 8 subquadrate, posterior end slightly more slender than anterior end, no lobes or incision along posterior margin, covered with microtrichia and fine setae, setae longer towards posterior end, four large setae along posterior margin. Sternite 9 slender, elongate, genital chamber slender, anterior arm long, reaching anterior end of terminalia, slightly widening at tip, lateral arms fused lateroposteriorly to tergite 9+10, three pairs of elongate setae at medio-distal margins. Tergite 8 straight at posterior margin, lateroanterior ends extending ventrally, bare medially, microtrichia and setae only at laterals, a wide and deep U-shape medial incision. Tergite 9+10 slender medially, laterals projected posteriorly, entirely bare. Cercomeres 1 and 2 probably fused, no sign of suture, digitiform, slightly curved inwards towards apex, covered with microtrichia and setae, four stronger distal setae.

**Male**. As male, except for the following. **Wing l**ength, 2.08; width, 0.75. **Terminalia** (Figs. 136A–B). Gonocoxites elongate, fused medially with no suture, a long lateral digitiform lobe projecting beyond tip of gonostylus. Gonostylus slender basally, much wider towards apex, densely setose on distal half. Gonocoxal apodemes parallel, close to each other. Parameres elongate, blade-like, with a triangular distal end with a medial short and slender incision distally. Tergite 9 with a pair of setose digitiform lobes laterally connected anteriorly.

#### Material examined

**Holotype**: male, ZRCBDP0048844, Nee Soon (NS1), 14.jan.15, MIP leg. (slide-mounted). **Paratypes** (4 males, 16 females). **Males**: ZRCBDP0048456, Nee Soon (NS1), swamp forest, 06.feb. 13MIP leg. (slide-mounted); ZRCBDP0048864, Nee Soon (NS1), 10.dec.14, MIP leg. (slide-mounted) (MZUSP). ZRCBDP0049088, Nee Soon (NS1), 24.dec.14, MIP leg. (slide-mounted); ZRCBDP0049254, Nee Soon (NS1), 10.dec.14, MIP leg. **Females**: ZRCBDP0047843, Nee Soon (NS1), swamp forest, 24.apr.13, MIP leg.; ZRCBDP0047903, Nee Soon (NS2), swamp forest, 30.oct.13, MIP leg.; ZRCBDP0048454, Nee Soon (NS1), swamp forest, 31.oct. 12, MIP leg.; ZRCBDP0048457, Nee Soon (NS2), swamp forest, 11.apr. 12, MIP leg.; ZRCBDP0048458, Nee Soon (NS2), swamp forest, 13.jun. 12, MIP leg. (website photo specimen, slide-mounted); ZRCBDP0048463, Nee Soon (NS1), swamp forest, 07.nov. 12, MIP leg.; ZRCBDP0048682, Nee Soon (NS2), swamp forest, 05.Sep.12, MIP leg.; ZRCBDP0048910, Nee Soon (NS1), 31.dec.14, MIP leg.; ZRCBDP0048926, Nee Soon (NS1), 31.dec.14, MIP leg. (slide-mounted); ZRCBDP0049002, Nee Soon (NS2), 17.dec.14, MIP leg. (slide-mounted); ZRCBDP0049004, Nee Soon (NS2), 17.dec.14, MIP leg.; ZRCBDP0049083, Nee Soon (NS1), 24.dec.14, MIP leg.; ZRCBDP0049102, Nee Soon (NS1), 24.dec.14, MIP leg.; ZRCBDP0049110, Nee Soon (NS1), 24.dec.14, MIP leg.; ZRCBDP0049224, Nee Soon (NS1), 10.dec.14, MIP leg; ZRCBDP0048455, Singapore, 06.jun.2012. **Additional sequenced specimens**: ZRCBDP0133452, Singapore, date range 2012-2018, MIP leg.; female, ZRCBDP0133913, Singapore, date range 2012-2018, MIP leg.; ZRCBDP0133926, Singapore, date range 2012-2018, MIP leg. (abdomen missing);female, ZRCBDP0155116, Nee Soon (NSM2a), swamp forest, 01.apr.15, MIP leg. (website photo specimen), ZRCBDP0048470, Mandai Mangroves (MM05), 02.jan.13; ZRCBDP0048846, Nee Soon (NS1), 14.jan.15; ZRCBDP0048990, Nee Soon (NS2),

17.dec.14; ZRCBDP0049046, Nee Soon (NS1), 24.dec.14; ZRCBDP0049103, Nee Soon (NS1), 24.dec.14; ZRCBDP0049116, Nee Soon (NS1), 24.dec.14, ZRCBDP0049126, Nee Soon (NS1), 24.dec.14.

**Etymology**. The species epithet refers to The Nanyang University (Nan Yang meaning Southern Ocean, the sinocentric term for Southeast Asia), merged in 1980 with the University of Singapore to constitute de present National University of Singapore. The Nanyang University, which existed from 1956 to 1980, has been until 2005 the only private university in Singapore in the Chinese language.

**Remarks**. There are 12 haplotypes in this species and all delimitation approaches bring them together into a single mOTU consisting of two 1% subclusters.

*Epicypta nus* Amorim & Oliveira, sp. n. (Figs. 137A–D, 138A–B)

https://singapore.biodiversity.online/species/A-Arth-Hexa-Diptera-000734,and-002127

urn:lsid:zoobank.org:act:A6AF43FF-73C0-4203-8560-4ECF5081CEC8

**Diagnosis**. Head bright ochre-yellow, antennal scape, pedicel, and first two flagellomeres ochre-yellowish, other flagellomeres grey-brownish. Scutum ochre-yellow, scutellum light brown; pleural sclerites light greyish-brown, antepronotum, proepisternum, anterior, and posterior ends of anepisternum more ochre-yellow; laterotergite and mediotergite brown. Fore coxa and femur whitish with orange tinge, mid and hind coxae whitish with a brown mark at basal end, mid and hind femora light whitish-yellow with a brownish line along ventral and dorsal crests. Wing membrane light brownish, darker along cells c and br. C produced beyond tip of R_5_; M_1+2_ almost as long as r-m. Dorsal macrotrichia on posterior veins M_1_ and M_2_, M_4_, and on CuA close to tip, macrotrichia on anal lobe membrane. Abdominal tergite 1 ochre-yellowish, tergites 2–5 ochre-yellowish with a small brown mark lateroposteriorly, tergite 6 brownish on anterior half, ochre-yellow on posterior half, tergite 7 ochre-yellow, terminalia yellowish. Syngonocoxite medial suture evident, gonocoxite with a long lateral lobe widening towards apex and an even longer digitiform dorsal lobe; gonostylus small, digitiform, slightly wider subapically; parameres with a pair of subapical spines and a pair of lateral lobes similar to can openers; tergite 9 with a pair of elongate digitiform extensions.

***Epicypta_nus_ZRCBDP0047864_hapZRCBDP0047864_ SMH_holotype* [109: T, 127: A, 166: G, 197: C, 199: T, 241: T, 262: C, 307: C]**

tctttcatcaacaattgcccatgcaggagcatctgttgatttagcaattttttctcttcatttagcaggaatttcttcaattttaggagctattaattttattactacTattattaatatgcgagcAccaggaatttcttttgatcgaatacctttatttgtttgGtcagttttaattactgctattcttttattaCtTtctttacctgtattagcaggcgctattactatattattaacTgatcgaaatttaaatacttcCttttttgaccctgcgggaggaggagatccaattttatatcaacaCttattt

#### Description

**Female** (Fig. 137A). Wing length, 2.11; width, 0.77. **Head** (Fig. 137B). Vertex ochre-yellowish. Scape and pedicel whitish-yellow, flagellomeres ochre-yellow with a whitish band basally. Face and clypeus light ochre-yellow. Palpomeres yellowish-ochre; labella whitish. Occiput with three longer setae dorsally to eye anteriorly to lateral ocellus, two dorsally to ocellus, seven posteriorly to ocellus. Scape about 1.5× pedicel length; flagellomere 4 1.5× longer than wide. Palpomere 4 1.3× palpomere 3 length, palpomere 5 1.4× palpomere 4 length. **Thorax**. Scutum ochre-yellowish, with a pair of dark brown marks medially at posterior margin, scutellum light brown. Scutum with seven long supra-alar setae and three pairs of prescutellars bristles; two pairs of scutellar bristles and one additional outer pair of long setae. Antepronotum yellowish-ochre on dorsal half, brownish-ochre on ventral half, anepisternum light brown medially, light yellowish-ochre on anteroventral corner, yellowish-ochre on posterior fourth, katepisternum, anepimeron, laterotergite, and metepisternum light brown. Pleural membrane yellowish. Anepisternum with five bristles along posterior margin. Mesepimeron with two bristles and 15 small setae, laterotergite with one bristle and 8 small setae; metepisternum with 18 small, fine setae. Haltere whitish with some ochre areas, no larger setae. **Legs**. Coxae whitish, fore coxa with brownish tinge, mid coxa with brownish tinge on basal half, hind coxa light brown on basal fifth. Femora whitish-yellow with brownish tinge, tibiae and tarsi light ochre-yellow. Mid coxa with a small band of fine setae across proximal fifth, hind coxa covered with fine setae on proximal fifth. Fore tibia with one bristle dorsoventrally, mid tibia with two dorsolateral rows of 4–5 bristles, a row with three bristles on outer face and three bristles ventrally, hind tibia with two dorsolateral rows with five bristles and a lateral row with two longer setae. Fore leg tarsomere 1 0.9× tibial length, 1.4× tarsomere 2 length. Hind tibial inner spur 5.0× tibia width at apex. **Wing** (Fig. 137C). Membrane light yellowish fumose, darker along anterior margin. Sc very short (visible with phase contrast), ending free. C extending beyond tip of R_5_ for one third of distance to M_1_. R_1_ reaching C on wing distal fourth; R_5_ reaching C before level of tip of M_1_. First sector of Rs slightly oblique, bare, **Holotype**: male, ZRCBDP0047864, Nee Soon (NS2), swamp forest, 19.mar.2014, MIP leg. (imaged, slide-mounted). **Paratypes** (13 males, 13 females). **Males**: ZRCBDP0048469, Nee Soon (NS1), swamp forest, 07.nov. 12, MIP leg. (website photo specimen, slide-mounted); ZRCBDP0048747, Nee Soon (NS1), 25.feb.15, MIP leg.; ZRCBDP0048799, Nee Soon (NS1), 18.mar.15, MIP leg.; ZRCBDP0049244, Nee Soon (NS1), 10.dec.14, MIP leg.; ZRCBDP0066812, Bukit Timah, maturing secondary forest (BT08), 16.aug. 16, MIP leg.; ZRCBDP0072663, Bukit Timah, primary forest (BT05), 28.dec. 16, MIP leg. (slide-mounted); ZRCBDP0072667, Bukit Timah, primary forest (BT05), 28.dec. 16, MIP leg.; ZRCBDP0072718, Bukit Timah, primary forest (BT05), 01.dec. 16, MIP leg.; ZRCBDP0072723, Bukit Timah, maturing secondary forest (BT06), 01.dec. 16, MIP leg.; ZRCBDP0072726, Bukit Timah, 0.52× r-m length; r-m oblique. M1+2 0.89 r-m length; bM primary forest (BT05), 14.dec. 16, MIP leg.; ZRCBDP0072734, Bukit Timah, primary forest (BT05), 14.dec. 16, MIP leg.; 4.3× longer than r-m; first sector of CuA about 0.45 length of second sector of CuA. CuP well produced to level of origin of M_4_. Anal fold only gently curved, nearly reaching wing margin. Posterior veins M_1_ and M_2_ with dorsal macrotrichia on about distal third, M_4_ on distal half, CuA on distal end; anal lobe membrane with dorsal macrotrichia. **Abdomen**. Abdominal tergites dark ochre-yellowish with light brown marks, tergites 2-5 light brown on posterior half or third, ochre-yellowish on anterior half or two-thirds, tergite 6 light brown on anterior half, ochre-yellowish on posterior half; sternites 1-7 light ochre-yellowish. Sternite 2 with a pair of strong brown bristles. **Terminalia** (Fig. 137D). Light brownish-yellow. Sternite 8 large, trapezoid, distal end more slender than anterior end, posterior margin with a short medial incision, microtrichia and setae concentrated mainly on distal half, two pairs of long setae subapically. Sternite 9 with wide medial arms, anterior apodeme (notum) long, extending beyond anterior end of terminalia, genital chamber elongate. Sternite 10 with distal end acute, with some subapical setulae laterally. Tergite 8 large, a complete suture dividing tergite into a pair of plates barely in contact medially, a pair of apodemes extending anteriorly from lateral corners, microtrichia spread on posterior half, setae restrict to posterior margins, longer laterally. Tergite 9+10 also separate into a pair of separate sclerites with a slender medial connection, bare. Cercomeres 1 and 2 fused, no sign of suture, basal half wider, posterior end digitiform, distal setae longer.

**Male**. As male, except for the following. **Wing l**ength, 1.92; width, 0.69 mm. **Terminalia** (Figs. 138A–B). Yellowish. Gonocoxites medially fused at ventral face, suture evident, bare ventrally, with a long lateral, setose lobe widening towards apex, much longer than gonostylus. Gonostylus small, simple, digitiform, slightly wider subapically. Parameres with a pair of strong subapical spines on a triangular projection, laterally with a pair of strong bifid lobes similar to can openers. Tergite 9 present as a pair of long digitiform, setose extensions. No sign of cerci.

#### Material examined

Holotype: male, ZRCBDP0047864, Nee Soon (NS2), swamp forest, 19.mar.2014, MIP leg. (imaged, slide-mounted). Paratypes (13 males, 13 females). Males: ZRCBDP0048469, Nee Soon (NS1), swamp forest, 07.nov. 12, MIP leg. (website photo specimen, slide-mounted); ZRCBDP0048747, Nee Soon (NS1), 25.feb.15, MIP leg.; ZRCBDP0048799, Nee Soon (NS1), 18.mar.15, MIP leg.; ZRCBDP0049244, Nee Soon (NS1), 10.dec.14, MIP leg.; ZRCBDP0066812, Bukit Timah, maturing secondary forest (BT08), 16.aug. 16, MIP leg.; ZRCBDP0072663, Bukit Timah, primary forest (BT05), 28.dec. 16, MIP leg. (slide-mounted); ZRCBDP0072667, Bukit Timah, primary forest (BT05), 28.dec. 16, MIP leg.; ZRCBDP0072718, Bukit Timah, primary forest (BT05), 01.dec. 16, MIP leg.; ZRCBDP0072723, Bukit Timah, maturing secondary forest (BT06), 01.dec. 16, MIP leg.; ZRCBDP0072726, Bukit Timah, primary forest (BT05), 14.dec. 16, MIP leg.; ZRCBDP0072734, Bukit Timah, primary forest (BT05), 14.dec. 16, MIP leg.; ZRCBDP0072742, Bukit Timah, primary forest (BT05), 14.dec. 16, MIP leg.; ZRCBDP0074042, Bukit Timah, primary forest (BT05), 08.dec. 16, MIP leg. **Females**: ZRCBDP0048451, Nee Soon (NS2), swamp forest, 12.dec. 12, MIP leg. (extracted); ZRCBDP0048798, Nee Soon (NS1), 18.mar.15, MIP leg.; ZRCBDP0048805, Nee Soon (NS1), 18.mar.15, MIP leg.; ZRCBDP0048881, Nee Soon (NS1), 31.dec.14, MIP leg.; ZRCBDP0048900, Nee Soon (NS1), 31.dec.14, MIP leg. (slide-mounted); ZRCBDP0048916, Nee Soon (NS1), 31.dec.14, MIP leg.; ZRCBDP0048920, Nee Soon (NS1), 31.dec.14, MIP leg.; ZRCBDP0049059, Nee Soon (NS2), 07.jan.15, MIP leg.; ZRCBDP0049178, Nee Soon (NS2), 13.may.15, MIP leg.; ZRCBDP0072690, Bukit Timah, maturing secondary forest (BT06), 28.dec. 16, MIP leg.; ZRCBDP0072708, Bukit Timah, primary forest (BT05), 01.dec. 16, MIP leg.; ZRCBDP0072728, Bukit Timah, primary forest (BT05), 14.dec. 16, MIP leg.; ZRCBDP0074036, Bukit Timah, primary forest (BT05), 08.dec. 16, MIP leg. **Additional sequenced specimens**: ZRCBDP0066820; ZRCBDP0078947; ZRCBDP0078970; ZRCBDP0078974; ZRCBDP0133970; ZRCBDP0279165; ZRCBDP0284198; ZRCBDP0049119, Nee Soon (NS1), dec.14; ZRCBDP0049125, Nee Soon (NS1), 24.dec.14; ZRCBDP0066767, Bukit Timah (BT05), 25.jan.17, MIP leg. (MZUSP). **Sequence failure specimen**: ZRCBDP0067855.

**Etymology**. The species epithet of this species refers to the National University of Singapore, NUS, the national research university of Singapore. Founded in 1905 as the Straits Settlements and Federated Malay States Government Medical School, NUS is the oldest higher education institution in Singapore. It is consistently ranked within the top 20 universities in the world and is considered to be the best university in the Asia-Pacific by the QS ranking. NUS offers a wide range of disciplines, including sciences, medicine and dentistry, design and environment, law, arts and social sciences, engineering, business, computing and music at both the undergraduate and postgraduate levels. The name is used in aposition.

**Remarks**. There are four haplotypes in our samples for *Epicypta nus* Amorim & Oliveira, **sp. n.,** which cluster together into a single mOTU according to all delimitation approaches.

*Epicypta peterngi* Amorim & Oliveira, sp. n. (Figs. 139A–E)

https://singapore.biodiversity.online/species/A-Arth-Hexa-Diptera-000794

urn:lsid:zoobank.org:act:6F1F1B69-94C1-421B-A172-BA3756255029

**Diagnosis**. Head ochre-yellowish with large brownish areas, antennal scape, pedicel, and first flagellomere yellowish-brown, remaining flagellomeres light greyish-brown. Scutum dark brown, with ochre-yellowish band at anterior end; scutellum blackish-brown; thoracic pleural sclerites dark greyish-brown. Coxae whitish, hind coxa with a light brown basal tinge, tip of mid and hind femora with a brown mark. Wing membrane light brownish fumose, area along anterior margin on basal half of wing more yellowish. C not extending beyond tip of R_5_; M_1+2_ about half of r-m length. Dorsal macrotrichia on posterior veins M_1_, M_2_, and posterior half of M_4_, no macrotrichia on anal lobe membrane. Abdominal tergite 1 dark brown, tergites 2–5 greyish-brown medially with a cream-yellow lateral band, larger on tergites 3–4, connecting each other medially on tergite 5, tergites 6–7 greyish-brown. Gonocoxites with a long lateral lobe and a long digitiform dorsal lobe; gonostylus bifid at base, with a short, digitiform internal lobe and a longer outer lobe; aedeagus with a pair of parallel projections distally; paramere with a pair of strong spines on a triangular projection distally; tergite 9 with a pair of long projections slender towards apex.

***Epicypta_peterngi_ZRCBDP0047888_hapZRCBDP0047 888_SMH_holotype* [19: A, 149: G, 151: C, 226: C, 229: T, 238: G]**

cctttcttctactattgcAcatgcaggagcatcagtagatttagctattttttctcttcatttagcaggaatttcttctattttaggggctattaactttattacaacaattattaatatacgagctccaggaattacatttgatcgaGtCcctttatttgtttgatctgttttaattactgccgttttattattactatctttaccagtattagcaggggctatCacTatattattGacagatcgaaatttaaatacatcattttttgatccagcaggaggaggagaccctattttatatcaacatttattt

#### Description

**Male**. Wing length, 2.56; width, 0.93. **Head**. Light brownish-yellow medially, ochre-yellowish laterally. Scape and pedicel greyish ochre-yellow, flagellomeres ochre-yellow. Face and clypeus light brownish-yellow, palpomeres 1–3 light brownish-yellow, palpomeres 4–5 lighter, labella whitish-yellow. Occiput with four longer setae dorsally to eye anteriorly to lateral ocellus, two dorsally to ocellus, ten posteriorly to ocellus. Scape about 2.0× pedicel length; flagellomere 4 1.5× longer than wide.

Palpomere 4 1.5× palpomere 3 length, palpomere 5 1.9× palpomere 4 length. **Thorax**. Scutum dark brown, anterior end ochre-yellow, scutellum blackish-brown with ochre-yellow anterolateral corners. Antepronotum, proepisternum, anepisternum, katepisternum, mesepimeron, laterotergite, and metepisternum dark greyish-brown, katepisternum slightly lighter, dark brown. Haltere light ochre-yellowish, no larger setae. Pleural membrane ochre-yellow. Scutum with nine supra-alars and three pairs of prescutellar bristles, two pairs of scutellar bristles and one outer pair of long setae. Proepisternum with three bristles, anepisternum with five bristles along posterior margin. Mesepimeron with two small bristles and 18 small setae, laterotergite with three bristles, one long seta and 18 small setae. Metepisternum with 18 fine setae. **Legs**. Fore coxa whitish with an orangish tinge, mid and hind coxae whitish, hind coxa with a light brownish tinge at basal end; femora whitish with a light brown tinge, mid and hind femora darker at tip; tibiae and tarsi light brownish-yellow, tibia and each tarsomere with a yellowish tinge at tip. Fore tibia with two lateral bristles medially; mid tibia with 2–3 irregular rows of 4–6 small bristles dorsally, one bristle on inner face medio-distally and three long setae along ventral edge; hind tibiae with two irregular rows of six small bristles dorsally and four longer setae at inner face medio-distally. Fore leg tarsomere 1 as long as tibia, 1.4× tarsomere 2 length. Hind tibial inner spur 4.4× tibia width at apex. **Wing** (Fig. 139B). Membrane fumose light brown, darker along anterior margin. Sc barely produced. C not produced beyond tip of R_5_. R_1_ reaching C on distal sixth of wing; R_5_ reaching C before level of tip of M_1_. First sector of Rs oblique, 0.63× r-m length; r-m oblique. M_1+2_ 0.61× r-m length; bM 4.6× r-m length; first sector of CuA 0.29× length of second sector of CuA. Cubital pseudovein not produced, CuP extending slightly beyond level of origin of M_4_. Anal fold almost reaching wing margin. Posterior veins M_1_ and M_2_ almost entirely with dorsal macrotrichia, M_4_ with macrotrichia on distal half, CuA with macrotrichia on distal fourth of wing; some few macrotrichia on anal lobe membrane. **Abdomen**. Abdominal tergite 1 greyish-brown medially, ochre-yellowish laterally, tergites 2–5 light brown medially, with cream-yellow areas laterally and anteriorly, wider on posterior segments, tergite 6 almost entirely brownish, an ochre-yellow tinge lateroanteriorly, tergite 7 cream-yellow; sternites 1-7 light ochre-yellowish. Sternite 2 with a pair of strong brown bristles. **Terminalia** (Figs. 139C– D). Ochre-yellowish. Gonocoxites fused medially, no suture present, bare ventrally, a medioventral projection slightly wider at base than at tip, about half of gonostylus length, with four setulae at distal end, sided by a pair of shorter digitiform projections with three distal small setae, a pair of long laterodistal extensions slightly shorter than gonostylus. Gonostylus simple, elongate, slightly clavate, dorsoventrally compressed, with fine setae along inner margin. Aedeagus large, oblong, ejaculatory apodeme slender at anterior end, wide midway to apex, then divided distally into a pair of lobes extending posteriorly and pointed at tip, extending beyond tip of laterodistal projections of gonocoxite, gonopore dorsally to base, between these lobes; parameres present as a triangular sclerite extending distally, medial short beak at tip with a pair of subapical small setae close to each other and a pair of strong curved spines close to tip, laterally with a pair of long, digitiform projections extending beyond tip of gonocoxite lateral lobes, five setae along most of their length. Gonocoxal bridge not evident. Tergite 9 present as a pair of long laterodistal digitiform extensions, about as long as gonocoxite lateral extensions, setose at tip. Cerci weakly sclerotized between tergite 9 elongate lateral lobes. **Female** (Fig. 139A). As male, except for the following. **Wing l**ength, 2.75; width, 1.01 mm. **Head**. Occiput with four longer setae dorsally to eye anteriorly to lateral ocellus, one dorsally to ocellus, 8 posteriorly to ocellus. **Thorax**. Scutum with eight supra-alar bristles and three pairs of prescutellars; three pairs of scutellar bristles, outer pair smaller. Proepisternum with four bristles directed ventrally, anepisternum with 5–6 bristles along posterior margin. Mesepimeron with three bristles and 19 small setae and setulae, laterotergite with three bristles and 17 smaller setae or setulae. Metepisternum with 20 small fine setae. **Legs**. Fore tibia with two bristles at outer face, mid tibia with two dorsolateral rows of 4–5 small bristles, one bristle on outer face medio-distally and four strong setae along ventral margin; hind tibia with three irregular rows of 4–6 small bristles. **Terminalia** (Fig. 139E). Light brownish-yellow. Sternite 8 elongate, posterior margin with a short median incision, microtrichia and setae evenly distributed, long setae at posterior margin. Sternite 9 slender, anterior apodeme extending to slightly beyond anterior end of terminalia, lateral arms ovoid, elongate, genital chamber elongate. Tergite 8 with a pair of separate large lateral lobes with microtrichia and setae. Tergite 9+10 slender, a pair of sclerotized bands connected medially. Cercomeres 1 and 2 fused, no sign of suture, elongate, wider midway to apex posterior end digitiform, distal setae longer.

#### Material examined

**Holotype**: male, ZRCBDP0047888, Nee Soon (NS2), swamp forest, 23.oct.2013, MIP leg. (slide-mounted). **Paratypes** (1 male, 1 female). **Male**: ZRCBDP0047907, Nee Soon (NS2), swamp forest, 30.oct.13, MIP leg. **Female**: ZRCBDP0047927, Nee Soon (NS1), swamp forest, 17.jul.13, MIP leg. (website photo specimen, slide-mounted).

**Etymology**. The species epithet of this species honors Peter Ng Kee Lin (1960-), prominent crustacean and fish systematist, with contributions to conservation biology and aquatic ecology. He has been the Director of the Raffles Museum of Biodiversity Research since 1998 and published over 1,000 scientific papers.

**Remarks**. *Epicypta peterngi* Amorim & Oliveira, **sp. n.,** with *E. wallacei* Amorim & Oliveira, **sp. n.** and *E. purchoni* Amorim & Oliveira, **sp. n.,** seem to be part of a small clade within the genus, as suggested by the shape of the aedeagus, besides similarities shared by a larger group of species, as the spines distally on the parameres.

*Epicypta maggielimae* Amorim & Oliveira, sp. n. (Figs. 140A–D)

https://singapore.biodiversity.online/species/A-Arth-Hexa-Diptera-000828

urn:lsid:zoobank.org:act:CE9AF0A3-0FAF-4257-8C14-7E245A9C133D

**Diagnosis**. Head dark brown, antennal scape, pedicel, and first flagellomere ochre-yellowish, other flagellomeres grey-brownish. Scutum dark ochre-yellow with a blackish-brown mark on posterior fifth, scutellum blackish-brown; pleural sclerites with a dark ochre-yellow background, antepronotum, proepisternum, katepisternum, and mesepimeron more brownish, mediotergite blackish-brown on posterior half, paratergite blackish-brown. Fore coxa and femur whitish with an orange tinge, mid coxa and femur whitish, femur with a faint brownish mark dorsally close to tip. Wing membrane light brownish, more yellowish along cells c and br; C clearly produced beyond tip of R_5_; M_1+2_ 0.80× r-m. Dorsal macrotrichia on posterior veins M_1_ and M_2_, M_4_ and on CuA distal half, anal lobe slender, no macrotrichia on membrane. Abdominal tergites ochre-yellowish. Female terminalia vaginal furca with wide, rounded anterior half.

***Epicypta_maggielimae_ZRCBDP0048259_hapZRCBDP 0048259_SMH_holotype* [22: C, 49: C, 97: C, 133: G, 13 9: A, 193: G, 250: C, 310: T]**

tctttcttctactattgctcaCgcaggttcatctgtagatttagcaatCttttctcttcatttagcaggtatttcttcaattttaggggctattaaCtttattacaacaattattaatatacgagcccctggGatttcAtttgatcgaatacctttatttgtttgatcagtattaattacagctattcttttGttattatctttacctgtattagcaggagctattacaatactattaactgatcgtaaCttaaatacatctttttttgaccccgctggagggggagacccaattttatatcaacatctTttt

#### Description

**Female** (Fig. 140A). Wing length, 2.00; width, 0.75. **Head**. Dark brown. Scape and pedicel greyish ochre-yellow, flagellomere 1 light brownish-yellow, flagellomeres 2–14 light greyish-brown. Face and clypeus brown. Palpomeres 1–3 brownish-yellow, palpomeres 4–5 lighter. Labella whitish-yellow. Occiput with two longer setae dorsally to eye anteriorly to lateral ocellus, one dorsally to ocellus, six posteriorly to ocellus. Scape 1.8× pedicel length, flagellomere 4 1.8× longer than wide. Palpomere 4 1.5× palpomere 3 length, palpomere 5 1.7× palpomere 4 length. Scutum with 4+3 supra-alar bristles, three pairs of prescutellars; two pairs of scutellar bristles and one additional pair of long external setae. Proepisternum with three bristles, anepisternum with five bristles along posterior margin. Mesepimeron with two bristles and 12 small setae and setulae, laterotergite with two bristles and 13 small setae. Metepisternum with 23 fine setae. **Thorax** (Fig. 140B). Scutum caramel-yellow, a blackish-brown medial band at posterior end; scutellum blackish-brown with ochre-yellow laterals. Antepronotum, proepisternum, katepisternum, dorsal half of mesepimeron and anterior end of laterotergite brownish, ventral half of mesepimeron, and most of laterotergite caramel-brown, anepisternum caramel-brown with dark areas dorsoanteriorly, mediotergite blackish-brown medially, caramel-yellow laterally. Pleural membrane ochre-yellow. Scutum with 4+3 long supra-alar bristles, three pairs of prescutellar bristles; two pairs of bristles and one additional external pair of long setae. Proepisternum with three bristles, anepisternum with five bristles along posterior margin. Mesepimeron with two longer setae and 17 small setae and setulae, laterotergite with no bristles, only four small setae [hind legs and posterior part of thorax missing in holotype]. **Legs**. Fore coxa light yellowish-brown with an orangish tinge, mid coxa whitish. Fore and mid femora concolor with fore coxa, mid femur with light brown tip. Fore and mid tibiae and tarsi ochre-yellow, tibiae with brownish-yellow tip. Mid coxa with a small group of fine setae across proximal fifth. Fore tibia with one strong seta medially on external face, mid tibia with two irregular dorsolateral rows of 3–5 brownish bristles laterally, one lateral seta on external face on distal fourth, three bristles along ventral edge. Hind tibia inner spur 6.7× tibia width at apex. **Wing** (Fig. 140C). Membrane fumose light brown, slightly darker along anterior margin. C extending beyond apex of R_5_ for a third of distance to M_1_. Sc short, weakly sclerotized beyond humeral vein. R_1_ reaching C on distal fourth of wing; R_5_ reaching C before level of tip of M_1_. First sector of Rs slightly oblique, 0.82× r-m length; r-m almost longitudinal. M_1+2_ 1.8× r-m length; bM 9× r-m length; first sector of CuA 0.25× length of second sector of CuA. Cubital pseudovein absent, CuP reaching slightly beyond level of origin of M_4_. Anal fold only gently curved, almost reaching wing margin. Posterior veins M_1_ and M_2_ with dorsal macrotrichia along most of their length, M_4_ with macrotrichia along distal half, CuA with macrotrichia on distal third, anal lobe with no macrotrichia. **Abdomen**. Abdominal tergite 1 ochre-yellowish, tergites 2–5 light brownish-yellow medially, with a wide light ochre-yellowish band laterally, tergites 6–7 light ochre-yellowish; sternites 1-7 light ochre-yellowish. Sternite 2 with a pair of strong brown bristles. **Terminalia** (Fig. 140D). Ochre-yellowish. Sternite 8 trapezoid, posterior margin with no medial incision, microtrichia and setae concentrated mostly on distal half. Sternite 9 with lateral arms slender, anterior apodeme wide, rounded anteriorly, genital chamber elongate. Sternite 10 with distal end acute, with some subapical setulae laterally. Tergite 8 short and bare medially, lateral lobes slightly projected posteriorly and entirely separated medially, microtrichia spread on posterior half, setae restrict to posterior margins, longer laterally. Tergite 9+10 with a slender, bare sclerotized medial band connecting a pair of lobes fused to sternite 9 lateroventrally. Cercomeres 1 and 2 fused, no sign of suture, basal ⅔ wider, posterior end digitiform, distal setae longer.

**Male**. Unknown.

#### Material examined

**Holotype**: female, ZRCBDP0048259, Sungei Buloh (SB1), mangrove, 09.oct.13, MIP leg. (website photo specimen, slide-mounted).

**Etymology**. The species epithet honors Maggie Lim (née Tan; 1913–1995), a physician, public health official and family planning and reproductive rights advocate. She was the first girl in Singapore to win the Queen’s Scholarship in 1930. During World War II, Lim was a camp medical doctor at Endau Settlement in Johor. After the war, she worked as an obstetrician and public health official in Singapore. She was president of the Family Planning and Population Board, and an advisor to the Midwives’ Council. Later in her career, Lim became a professor of epidemiology and public health at the University of Hawai’i’s East–West Center. She was inducted into the Singapore Women’s Hall of Fame in 2014.

*Epicypta yupeigaoae* Amorim & Oliveira, sp. n. (Figs. 141A–C)

urn:lsid:zoobank.org:act:D2514A4D-613B-431A-8131-D1BFCFB63892

**Diagnosis**. Light brownish-yellow, lighter laterally, antennal scape and pedicel ochre-yellowish, flagellomeres light brown. Scutum ochre-yellow with a transverse dark brown mark medially at posterior end, scutellum ochre-yellow with a brown medial mark. Pleural sclerites light brown, antepronotum and proepisternum yellowish-brown. Wing membrane light brown, cells c and br darker. C produced way beyond tip of R_5_; M_1+2_ 1.0× r-m length; dorsal macrotrichia on posterior veins M_1_ and M_2_, M_4_ and on distal end of CuA, macrotrichia on anal lobe membrane. Abdominal tergites 1–6 light yellowish-brown on anterior half, light brown on posterior half, tergite 7 ochre-yellowish. Gonocoxites medially fused, a medioposterior projection rounded distally, a long lateral lobe widening towards apex and an even longer digitiform lobe dorsally; gonostylus small; parameres with a pair of subapical spines and a pair of lateral lobes similar to can openers; tergite 9 with a pair of long, setose digitiform extensions.

***Epicypta_yupeigaoae_ZRCBDP0066740_hapZRCBDP0 066740_SMH_holotype* [10: C, 55: C, 148: C, 149: T, 17 5: T, 181: T, 196: G, 251: A, 253: T]**

tctttcctcCtcaattgcccatgctggggcttctgttgatttagctattttttcCcttcatttagcaggtatttcctctattttaggggctgtaaattttattacaacaattattaatatacgagcccccggaattacttttgatcgCTtacctttattcgtttgatcagttctTattacTgcagtactattattGctatcattacctgtattagccggagctattacaatacttctaacagatcgaaatAtTaatacttcattttttgatcctgccggaggaggagaccctattttatatcaacatttattt

#### Description

**Male**. Wing length, 1.86; width, 0.70. **Head** (Fig. 141A). Light brownish-yellow, lighter laterally on occiput. Scape and pedicel ochre-yellowish, flagellomeres light brown. Face and clypeus light brownish-yellow, palpomeres light brown, labella whitish-yellow. Occiput with three longer setae dorsally to eye anteriorly to lateral ocellus, three dorsally to ocellus, eight posteriorly to ocellus. Scape about 1.4× pedicel length; length of flagellomere 4 1.9× width. Palpomere 4 1.2× palpomere 3 length, palpomere 5 2.3× palpomere 4 length. **Thorax**. Scutum ochre-yellow with a transverse dark brown mark medially at posterior end; scutellum ochre-yellow with a brown transverse mark medially at anterior end. Antepronotum and proepisternum yellowish-brown, anepisternum, katepisternum, mesepimeron, and laterotergite light brown, metepisternum and mediotergite light brown. Scutum with 5+2 supra-alars and three pairs of prescutellar bristles, two pairs of scutellar bristles and one outer pair of long setae. Proepisternum with three bristles, anepisternum with five bristles along posterior margin. Mesepimeron with two small bristles and eight small setae, laterotergite with one bristle and ten small setae. Metepisternum with 12 fine setae and setulae. Haltere ochre-yellowish. Pleural membrane ochre-yellow. **Legs**. Fore coxa whitish with an orangish tinge, mid and hind coxae whitish, mid and hind coxae with a brownish band at proximal end, mid coxa brownish at anterior face distally, hind coxa with a brown mark at distal third on posterior face; Fore femur light yellowish-brown, mid and hind femora light brown; fore tibia and tarsus light yellowish-brown, hind tibia light brown, tarsus light yellowish-brown [mid tibiae and tarsi missing]. Fore tibia with two lateral bristles medially; hind tibia with 2–3 irregular rows of 5 small bristles dorsally, 2–3 bristles on inner face medio-distally. Fore leg tarsomere 1 about as long as tibia, 1.4× tarsomere 2 length. Hind tibial inner spur 5.3× tibia width at apex. **Wing** (Fig. 141B). Membrane fumose light brown, darker along anterior margin. Sc barely produced. C produced beyond tip of R_5_ for about a fourth of distance to M_1_. R_1_ reaching C on distal fourth of wing; R_5_ reaching C close to level of tip of M_2_. First sector of Rs almost transverse, 0.60× r-m length; r-m almost longitudinal. M_1+2_ about 1.0× r-m length; bM 4.6× r-m length; first sector of CuA short, 0.46× length of second sector of CuA. Cubital pseudovein absent, CuP not reaching level of origin of M_4_. Anal fold only gently curved, almost reaching wing margin. Posterior veins M_1_ and M_2_ almost entirely with dorsal macrotrichia, M_4_ with macrotrichia on distal ¾, CuA with macrotrichia on distal fifth of wing; macrotrichia on anal lobe membrane. **Abdomen**. Abdominal tergites 1–6 light yellowish-brown on anterior half, light brown on posterior half, tergite 7 ochre-yellowish; sternites 1-7 light ochre-yellowish. Sternite 2 with a pair of strong brown bristles. **Terminalia** (Fig. 141C). Ochre-yellowish. Gonocoxites fused medially, no suture present, bare ventrally, a large medioventral projection of syngonocoxite, wide at base, with two pairs of setulae subapically, extending to about ¾ of gonostylus length, a pair of long laterodistal extensions dorsally to insertion of gonostylus and extending much beyond tip of gonostylus, with fine setae on dorsal face on basal half and with five fine setae on inner margin on distal half, a setula at tip. Gonostylus composed of two lobes connected at base, inner lobe smaller, with a pair of longer setae at base of lobe on dorsal face and fine setae on ventral face at distal half; outer lobe larger, extending to level of tip of aedeagal distal projections, also compressed, more densely setose, with setae on both faces, more distal setae on curved inner face, larger setae along posterior end of outer face. Aedeagal-parameral complex with a pair of sclerites connected together more ventrally and projected posteriorly as a strongly sclerotized bottle-opener extending slightly beyond tip of gonostylus, and a medial short distal triangular projection more dorsally at tip, with two pairs of subapical small setae close to each other and a pair of strong spines close to tip. Gonocoxal bridge evident, incomplete medially, a pair of oblique apodemes directed anteriorly. Tergite 9 present as a pair of long laterodistal digitiform extensions, about as long as gonocoxite lateral extensions, setose at tip. Cerci not visible.

**Female**. Unknown.

#### Material examined

**Holotype**: male, ZRCBDP0066740, Bukit Timah, primary forest (BT05), 30.aug.2016, MIP leg. (slide-mounted).

**Etymology**. The species epithet honors Madame Yu Pei Gao, inaugural principal of the Singapore Nanyang Girls’ School – the first Chinese educational institution for girls in Singapore. Yu was an independent-minded principal who challenged obsolete traditions, such as allowing only female teachers for a girl’s school, and hired teachers based on their capabilities instead of gender.

**Remarks**. *Epicypta yupeigaoae* Amorim & Oliveira, **sp. n.** is obviously close to *E. nus* Amorim & Oliveira, **sp. n.,** as can be inferred by the uniquely derived can-opener shape of the parameres. The haplotype network leaves no question that they are separate species.

*Epicypta annwee* Amorim & Oliveira, sp. n. (Figs. 142A–D)

https://singapore.biodiversity.online/species/A-Arth-Hexa-Diptera-000732,-002110

urn:lsid:zoobank.org:act:99F26EFA-3D7F-4548-BBB5-876D9C82AEDA

**Diagnosis**. Head ochre-yellow, antennal scape, pedicel, and flagellomeres 1 and 2 ochre-yellowish, remaining flagellomeres brown. Scutum ochre-yellow, scutellum dark brown; most pleural sclerites dark ochre-yellow, some sclerites with diffuse brownish marks; mediotergite and metepisternum brown. Coxae and femora whitish, fore coxa with brownish tinge, femora with dorsal and ventral orange-brown crest, hind femur with brownish tip. Wing membrane light brownish, area along anterior margin slightly darker. C not extending beyond tip of R_5_; M_1+2_ clearly shorter than r-m. Dorsal macrotrichia on posterior veins M_1_, M_2_, posterior half of M_4_, and distal end of CuA, macrotrichia on anal lobe membrane. Abdominal tergite 1 brown, tergites 2–6 dark caramel-brown, whitish laterally, tergite 7 cream-yellow, terminalia whitish-yellow with brownish tinge. Gonocoxites fused medially, no suture, a pair of long, digitiform laterodistal lobes slightly widening towards apex. Gonostylus simple, elongate, directed inwards. Aedeagus wider midway to apex, with a pair of separate distal gonopores at tip of tubular extensions. Parameres with a pair of strong setae at tip. Tergite 9 with a pair of entirely separated, elongate and compressed lobes, as long as gonocoxite laterodistal lobes.

***Epicypta_annwee_ZRCBDP0047808_hapZRCBDP0047 808_SMH_holotype* [10: G, 109: C, 128: G, 129: T, 148: G, 208: A]**

tctttcatcGactattgcccatgctgggtcttcagttgatttagctattttttctcttcatttagcgggaatttcttctattttaggagctattaattttattactacCattattaatatacgttctGTaggaattacttttgaccgGatacctttatttgtatgatcagtattaattacagctattcttcttcttctttcattaccAgttttagcgggagctattacaatacttttaacagatcgtaatttaaatacttctttttttgatcctgctgggggaggagaccctatcttatatcaacatttattt

#### Description

**Male**. Wing length, 2.27; width, 0.83 mm. **Head**. Head ochre-yellowish. Scape and pedicel light ochre-yellowish, first two flagellomeres ochre-yellowish, remaining flagellomeres brown. Face and clypeus light brown, palpomeres 1–3 ochre-yellowish, 4–5 whitish-yellow, labella cream-yellow. Occiput with two longer setae dorsally to eye anteriorly to lateral ocellus, two dorsally to ocellus, five posteriorly to ocellus. Scape 1.6× pedicel length; flagellomere 4 twice as long as wide. Palpomere 4 1.3× palpomere 3 length, palpomere 5 2.0× palpomere 4 length. **Thorax**. Scutum ochre-yellow, scutellum blackish-brown, with ochre-yellow anterolateral corners. Antepronotum and ventral half of katepisternum light ochre, proepisternum, proepimeron, most of anepisternum, dorsal half of katepisternum, mesepimeron, and laterotergite light greyish-brown, with some diffuse lighter areas, metepisternum dark greyish-brown, mediotergite brown, lighter lateroventrally. Haltere pedicel whitish, knob brownish basally, whitish distally. Pleural membrane yellowish. Scutum with six supra-alar long setae and three pairs of prescutellars, two pairs of scutellar bristles and one outer pair of longer setae. Proepisternum with three bristles directed ventrally, anepisternum with five bristles along posterior margin. Mesepimeron with three long setae and 29 small setae, laterotergite with two long setae and five smaller setae. Metepisternum with eight small setae. **Legs**. Fore coxa light whitish-yellow, mid and hind coxae whitish; femora, tibiae, and tarsi light whitish-yellow, tarsi darker, femora with a yellowish-brown line along dorsal edge, hind femur with brownish tip. Fore tibia with two dorsolateral strong setae medially, mid tibia with two dorsolateral rows of 4–5 bristles and two strong setae along ventral edge, hind tibia with a row of six bristles dorsally, a line of four long setae on outer lateral face, and five bristles on inner lateral face. Fore leg tarsomere 1 1.2× tibia length, 1.9× tarsomere 2 length, mid and hind tarsomeres 1–3 with rows of ventral setae besides trichia. Hind tibia inner spur 5× tibia width at apex. **Wing** (Fig. 142B). Membrane fumose light brown, darker on cells c, br and r1. Sc barely produced. C not extending beyond apex of R_5_. R_1_ reaching C on distal sixth, R_5_ reaching C slightly beyond level of tip of M_1_. First sector of Rs oblique, slightly over half of r-m length; r-m almost longitudinal. M_1+2_ short, 0.68× r-m length; bM about 5× r-m length; tip of M_1_ slightly curved posteriorly at tip. First sector of CuA 0.34× length of second sector of CuA. Cubital pseudovein not produced, CuP extending slightly beyond level of origin of M_4_. Anal fold long, gently curved, almost reaching wing margin. Posterior veins M_1_ and M_2_ with dorsal macrotrichia along most of their length, M_4_ and CuA restricted to distal end; anal lobe with some macrotrichia. **Abdomen**. Abdominal tergite 1 greyish-brown, with dark brown mark along posterior margin, tergite 2 brown, tergites 3–5 light-brown medially with cream-yellow marks laterally and along anterior margin, tergite 7 cream-yellow; sternites 1-7 whitish-yellow. Sternite 2 with a pair of strong ventral brown bristles. **Terminalia** (Fig. 142C). Whitish-yellow. Gonocoxites fused medially, no suture, bare ventrally, a pair of laterodistal long digitiform lobes with long setae on ventral, outer and dorsal faces, dorsal face of terminalia without clear dorsomedial borders, a pair of digitiform projections medio-posteriorly with 4–5 fine setae at tip. Gonostylus simple, elongate, directed inwards, basally about as wide as base of laterodistal projection of gonocoxite, slender towards tip, slightly tapered distally, two fine sub-basal setae and some few setulae at tip. Aedeagus widening midway to apex, divided into a pair of separate distal gonopores at tip of tubular extensions. Parameres present as a triangular sclerite with rounded tip dorsally to aedeagus, a pair of strong setae at tip. Gonocoxal apodemes elongate, extending from inner base of gonocoxite laterodistal lobe, directed inwards. Tergite 9 present dorsally as a pair of entirely separate, elongate and compressed lobes extending as distally as gonocoxite laterodistal lobes, covered with microtrichia and setae, distal setae longer. Cerci not discernible.

**Female** (Fig. 142A). As male, except for the following. **Wing l**ength, 2.59–2.66; width, 0.91–0.94 (n=2). **Head**. Occiput with four longer setae dorsally to eye anteriorly to lateral ocellus, two dorsally to ocellus, eight posteriorly to ocellus. **Thorax**. Scutum with seven long supra-alar setae and three pairs of prescutellars; two pairs of scutellar bristles. Proepisternum with four bristles directed ventrally, anepisternum with five bristles along posterior margin. Mesepimeron with two long setae and 31 small setae and setulae, laterotergite with two bristles and 13 smaller setae. Metepisternum with 63 small fine setae and setulae. **Legs**. Fore tibia with two dorsolateral strong setae medially, mid tibia with two dorsolateral rows of 3–5 bristles and three strong setae along ventral edge, hind tibia with a row of 5–6 bristles dorsally, a line of seven long setae on inner lateral face. Fore leg tarsomere 1 1.2× tibia length, 1.7× tarsomere 2 length, mid and hind tarsomeres 1–3 with rows of ventral setae besides trichia. Hind tibia inner spur 5× tibia width at apex. **Wing**.

Membrane fumose light brown, darker on cells c, br and r1. Posterior veins M_1_, M_2_, and M_4_ with dorsal macrotrichia along most of their length, on CuA restricted to distal end; anal lobe membrane with macrotrichia. **Abdomen**. Abdominal tergite 1 greyish-brown with dark brown mark along posterior margin, tergite 2 brown, tergites 3–5 light-brown medially, with cream-yellow laterally and along anterior margin, tergite 7 cream-yellow; sternites 1-7 whitish-yellow. Sternite 2 with a pair of strong brown bristles. Tergite 2 with a pair of concentrated setae medially. **Terminalia** (Fig. 142D). Sternite 8 trapezoid, posterior end more slender than anterior end, posterior margin straight, no lateroposterior extensions, microtrichia and fine setae covering entire sclerite, some long setae at posterior margin, labia beneath distal margin with four long fine setae. Sternite 9 with slender genital chamber, lined with microtrichia, gonopore connected to two gonoducts, sclerotized area with a pair of arms extending laterally, an anterior apodeme extending into segment 7. Tergite 8 short medially and wide, a pair of lobes laterally, with microtrichia and some few elongate setae on lateroposterior corners. T9+10 bare, short medially, a pair of short lobes extending lateroposteriorly fused to sternite 9. Cercomeres 1 and 2 probably fused, no sign of suture, ovoid, laterally compressed, covered with microtrichia and elongate setae, distal setae longer and curved.

#### Material examined

**Holotype**: male, ZRCBDP0047808, Nee Soon (NS1), swamp forest, 01.may. 2013, MIP leg. (slide-mounted). **Paratypes** (14 males, 10 females). **Males**: ZRCBDP0047797, Nee Soon (NS2), swamp forest, 13.nov. 2013, MIP leg.; ZRCBDP0048697, Nee Soon (NS1), 13.may. 2015, MIP leg.; ZRCBDP0048793, Nee Soon (NS1), 18.mar. 2015, MIP leg.; ZRCBDP0048794, Nee Soon (NS1), 18.mar. 2015, MIP leg.; ZRCBDP0048815, Nee Soon (NS1), 14.jan. 2015, MIP leg.; ZRCBDP0048816, Nee Soon (NS1), 14.jan. 2015, MIP leg.; ZRCBDP0048903, Nee Soon (NS1), 31.dec. 2014, MIP leg.; ZRCBDP0048928, Nee Soon (NS1), 31.dec. 2014, MIP leg.; ZRCBDP0048935, Nee Soon (NS1), 31.dec. 2014, MIP leg.; ZRCBDP0048969, Nee Soon (NS1), 15.apr. 2015, MIP leg.; ZRCBDP0048970, Nee Soon (NS1), 15.apr. 2015, MIP leg.; ZRCBDP0049234, Nee Soon (NS1), 10.dec. 2014, MIP leg.; ZRCBDP0072716, Bukit Timah, primary forest (BT05), 01.dec. 2016, MIP leg.; ZRCBDP0072725, Bukit Timah, old secondary forest (BT01), 08.dec. 2016, MIP leg. **Females**: ZRCBDP0047811, Nee Soon (NS1), swamp forest, 01.may. 2013, MIP leg.; ZRCBDP0047917, Nee Soon (NS2), swamp forest, 27.nov. 2013, MIP leg. (slide-mounted) (MZUSP); ZRCBDP0048059, Pulau Ubin (PU2), mangrove, 01.apr. 2013, MIP leg.; ZRCBDP0048804, Nee Soon (NS1), 18.mar. 2015, MIP leg.; ZRCBDP0049084, Nee Soon (NS1), 24.dec. 2014, MIP leg.; ZRCBDP0049098, Nee Soon (NS1), 24.dec. 2014, MIP leg.; ZRCBDP0049232, Nee Soon (NS1), 10.dec. 2014, MIP leg.; ZRCBDP0049236, Nee Soon (NS1), 10.dec. 2014, MIP leg.; ZRCBDP0072661, Bukit Timah, primary forest (BT05), 28.dec. 2016, MIP leg.; ZRCBDP0074037, Bukit Timah, primary forest (BT05), 08.dec. 2016, MIP leg; ZRCBDP0048446, Singapore, apr.12. **Additional sequenced specimens**: female, ZRCBDP0078955, Singapore, date range 2012-2018, MIP leg.; female, ZRCBDP0078960, Singapore, date range 2012-2018, MIP leg.; female, ZRCBDP0078981, Singapore, date range 2012-2018, MIP leg.; female, ZRCBDP0078982, Singapore, date range 2012-2018, MIP leg.; ZRCBDP0128600, Bukit Timah (BT07), old secondary forest, 25.jan-01.feb. 2017, MIP leg.; female, ZRCBDP0133921, Singapore, date range 2012-2018, MIP leg.; female, ZRCBDP0133952, Singapore, date range 2012-2018, MIP leg.; female, ZRCBDP0134005, Singapore, date range 2012-2018, MIP leg.; male, ZRCBDP0134023, Singapore, date range 2012-2018, MIP leg; ZRCBDP0049216, Nee Soon (NS1), 10.dec.14; ZRCBDP0049233, Nee Soon (NS1), 10.dec.14; ZRCBDP0078956, Singapore; ZRCBDP0133935, Singapore. **Sequence failure specimens**: ZRCBDP0137195, Bukit Timah (BT06), maturing secondary forest (BT06), 22.feb-08.mar.2017, MIP leg. (sequence failure); ZRCBDP0137199, Bukit Timah (BT06), maturing secondary forest (BT06), 22.feb-08.mar.2017, MIP leg.

**Etymology**. The species epithet of this species honors Ann Elizabeth Wee (1926-2019; née Wilcox), a British-born academic and social worker. Considered the “founding mother of social work in Singapore”, she worked with the abused and abandoned, before joining the staff of the then-University of Malaya, pushing for the development of a four-year degree program to train social workers. She was the inaugural recipient of the lifetime volunteer achievement award of the Ministry of Community Development, Youth and Sports in 2009, was honored with the Meritorious Service Medal in 2010 and was inducted into the Singapore Women’s Hall of Fame in 2014. The name is used in apposition.

**Remarks**. *Epicypta annwee* Amorim & Oliveira, **sp. n.** has six different haplotypes clustered together by the species delimitation approaches.

*Epicypta wallacei* Amorim & Oliveira, sp. n. (Figs. 143A–D, 144A–B)

https://singapore.biodiversity.online/species/A-Arth-Hexa-Diptera-000738

urn:lsid:zoobank.org:act:724B5B7C-E6EC-4285-9027-260BCDA8A75A

**Diagnosis**. Head and scutum ochre-yellowish, antenna brownish; scutellum yellowish-brown; pleural sclerites dark ochre-yellowish, antepronotum and proepisternum lighter, laterotergite, metepisternum, and mediotergite brown. Coxae and femora whitish, a small brown mark at basal tip of hind coxa and at tip of hind femur. Wing membrane light brownish, cell c and cell br darker. C not produced beyond tip of R_5_; Sc long; M_1+2_ about half of r-m length. Dorsal macrotrichia on posterior veins M_1_ and M_2_, M_4_, and on CuA close to tip, macrotrichia on anal lobe membrane. Abdominal tergites 1–6 greyish-brown with light ochre-yellow anterolateral corners, wider on tergites 4–5, tergite 7 ochre-yellow, terminalia yellowish. Male terminalia gonocoxite with a medioventral digitiform projection and a digitiform laterodistal extension reaching way beyond tip of gonostylus; gonostylus simple, elongate, dorsoventrally compressed; aedeagus with a pair of separate tubular extensions each with a gonopore; parameres triangular with a pair of strong, curved spines close to tip; tergite 9 present as a pair of long laterodistal separate extensions, much longer than gonostylus or gonocoxite lobes.

***Epicypta_wallacei_ZRCBDP0048465_hapZRCBDP0048 465_SMH_holotype* [1: C, 7: C, 19: C, 29: T, 67: C, 106: C, 178: C, 196: T, 199: T]**

CctatcCtcaactattgcCcatgctggaTcttctgtagatttagctattttttctttacatttagcCggaatctcctcaattttaggggcaattaattttattacCacaattattaatatacgatccccaggaattacttttgatcgaatacctctatttgtttgatcagtattaatCacagctattttacttctTctTtctttacctgttttagcgggagctattacaatattattaacagatcgaaatttaaatacttctttttttgatccagcgggaggaggggatcctattttatatcaacatttattt

#### Description

**Male** (Fig. 143A). Wing length, 2.21; width, 0.80. **Head**. Head ochre-yellowish. Scape greyish-yellow, pedicel ochre-yellowish, flagellomeres light brown. Face and clypeus ochre-yellowish, palpomeres 1–3 ochre-yellowish, palpomeres 4–5 whitish-yellow, labella cream-yellow. Occiput with four longer setae dorsally to eye anteriorly to lateral ocellus, two dorsally to ocellus, eight posteriorly to ocellus. Scape about 1.3× pedicel length; flagellomere 4 1.7× as long as wide. Palpomere 4 1.1× palpomere 3 length, palpomere 5 2.0× palpomere 4 length. **Thorax**. Scutum ochre-yellow, scutellum yellowish-brown with lighter corners. Pleural sclerites light ochre-yellowish, laterotergite and metepisternum greyish-brown, mediotergite yellowish-brown. Haltere pedicel whitish, knob brownish basally, whitish distally. Pleural membrane yellowish. Scutum with seven supra-alar bristles and three pairs of prescutellar bristles, plus a pair of long setae between inner two, three pairs of scutellar bristles and one outer pair of longer setae. Proepisternum with three bristles, anepisternum with five bristles along posterior margin. Mesepimeron with three bristles and 20 small setae, laterotergite with three bristles and 18 smaller setae. Metepisternum with 29 small setae. **Legs**. Fore coxa whitish-yellow with a light brownish tinge, mid and hind coxae whitish, hind coxa with a greyish-brown mark at basal end; femora, tibiae, and tarsi light whitish-yellow, tarsi darker, hind femur with brownish mark at distal end. Fore tibia with two dorsolateral bristles, mid tibia with two dorsolateral rows of 3–5 bristles and three bristles along ventral edge, hind tibia with a row of 5–6 bristles dorsolaterally, a line of five long setae on outer lateral face. Fore leg tarsomere 1 1.1× tibia length, 1.7× tarsomere 2 length, mid and hind tarsomeres 1–3 with rows of ventral setae besides trichia. Hind tibia inner spur 5.0× tibia width at apex. **Wing** (Fig. 143B). Membrane fumose light brown, darker on cells c, br and r1. Sc barely produced. C barely extending beyond apex of R_5_. R_1_ reaching C on distal sixth, R_5_ reaching C slightly before level of tip of M_1_. First sector of Rs oblique, slightly over half of r-m length; r-m almost longitudinal. M_1+2_ 0.8× r-m length; bM 5.0× r-m length. First sector of CuA 0.45× length of second sector of CuA. Cubital pseudovein barely reaching level of origin of M_4_. Anal fold long, only gently curved, almost reaching wing margin. Posterior veins M_1_, M_2_, and M_4_ with dorsal macrotrichia along more than half their length, CuA with macrotrichia on distal fourth of wing; macrotrichia present on anal lobe membrane. **Abdomen**. Abdominal tergite 1 greyish-brown with dark brown mark along posterior margin, tergites 2–6 greyish-brown medially, cream-yellow laterally and along anterior margin, cream-yellow marks larger on segments 4–5, tergite 7 cream-yellow; sternites 1-7 whitish-yellow. Sternite 2 with a pair of strong brown bristles. **Terminalia** (Figs. 143C–D). Whitish-yellow. Gonocoxites fused medially, suture present, bare ventrally, a medioventral digitiform projection with setulae at tip sided by a pair of short pointed lobes, each with a small seta at tip, a pair of long, digitiform laterodistal extensions reaching way beyond tip of gonostylus. Gonostylus simple, elongate, dorsoventrally compressed, with setae along inner margin and at distal half on both faces. Aedeagus with an anterior medial ejaculatory apodeme, widening midway to apex, then divided distally into a pair of separate tubular extensions with independent gonopores, laterally with a pair of long, slender distal projections, slightly capitate at tip. Parameres as a triangular sclerite dorsal to aedeagus extending distally, with a medial short beak at tip, a pair of small setae close to each other subapically, a pair of strong, curved spines close to tip. Gonocoxal bridge present, slender medially, no apodemes visible. Tergite 9 present as a pair of long laterodistal extensions, much longer than gonostylus or gonocoxite extensions, setose at tip. Cerci not visible.

**Female**. As male, except for the following. **Wing l**ength, 2.27; width, 0.86. **Terminalia** (Figs. 144A–B). Brownish-yellow. Sternite 8 wide anteriorly, subquadrate, posterior margin slightly more slender than base, a very short medial incision, covered with microtrichia and fine setae, longer setae medially. Sternite 9 wide medially, anterior end slender, extending beyond anterior end of sternite 8, gonopore at center of a well-sclerotized plate. Tergite 8 wide, short medially, connecting a pair of lateral short lobes with fine setae, partially overlapping to sternite 8. Tergite 9+10 short, slender, bare. Cercomeres 1 and 2 apparently fused, no sign of suture, elongate, distal end curved ventrally, covered with microtrichia and setae, setae at tip longer.

#### Material examined

**Holotype**: male, ZRCBDP0048465, Nee Soon (NS1), swamp forest, 04.apr.2012, MIP leg. (website photo specimen, slide-mounted). **Paratypes** (4 females): ZRCBDP0048832, Nee Soon (NS1), 14.jan.15, MIP leg. (slide-mounted); ZRCBDP0048868, Nee Soon (NS1), 31.dec.14, MIP leg. (slide-mounted); ZRCBDP0048884, Nee Soon (NS1), 31.dec.14, MIP leg.; ZRCBDP0133462, Singapore, (date range 2012-2018), MIP leg.

**Etymology**. The species epithet of this species honors Alfred Russel Wallace (1823–1913), British naturalist, biogeographer, anthropologist, evolutionist and illustrator.

He is best known for independently conceiving the very idea of phylogeny and the theory of evolution through natural selection. Aside from scientific work, he was a social activist, critical of an unjust social and economic system in 19^th^-century Britain. He had extensive field work in the Amazon and in the Malay Archipelago, including Singapore. His 1858 paper on the subject was one of the triggers for Charles Darwin to publish his own writings on evolution.

**Remarks**. There are three haplotypes of *Epicypta wallacei* Amorim & Oliveira, **sp. n.** and they cluster together based on all species delimitation methods.

*Epicypta lamtoongjini* Amorim & Oliveira, sp. N. (Figs. 145A–E)

https://biodiversity.Online/species/A-Arth-Hexa-Diptera-000729

urn:lsid:zoobank.org:act:B25D0E92-84B8-4BB5-B695-CE21755E6C99

**Diagnosis**. Head dark ochre-orangish, scutum ochre-orangish anteriorly, caramel-brown on posterior two-thirds. Antepronotum and proepisternum dark ochre-yellow, proepisternum with a small greyish-brown mark; other sclerites dark greyish-brown, mediotergite dark brown. Coxae and femora whitish, fore coxa and femora with an orangish tinge, a small brown mark at distal end of hind femur. Wing membrane light brownish, darker along anterior margin. C barely produced beyond tip of R_5_; Sc faint, with a row of dorsal macrotrichia; M_1+2_ about half of r-m length. Dorsal macrotrichia on posterior veins M_1_ and M_2_, M_4_ and on CuA close to tip, macrotrichia on anal lobe membrane. Abdominal tergite 1 dark brown, tergites 2–5 brown with light ochre-yellow laterals extending inwards along anterior margin, wider on tergites 4–5, tergite 6 mostly ochre-yellow with brownish tinge. Male terminalia syngonocoxite with a long medioventral projection, a long digitiform laterodistal extension, and a bare dorsal projection; gonostylus simple, elongate, dorsoventrally compressed; aedeagus with a pair of separate extensions with independent gonopores; parameres triangular with a pair of strong, curved spines close to tip; tergite 9 present as a pair of long, setose separate extensions.

***Epicypta_lamtoongjini_ZRCBDP0047900_hapZRCBDP 0047816_SMH_holotype* [28: C, 29: T, 97: C, 106: A, 13 0: A, 172: A, 193: T, 202: A, 208: T, 217: G, 241: T]**

tctttcttctacaattgctcatgcaggCTcttctgtagatttagctattttttctcttcatttagcaggaatttcttctattttaggggcaattaaCtttattacAacaattattaatatacgttctccAggaattacttttgatcgaatacctttatttgtatgatcagtAttaattacagcagttttactTcttttatcAttaccTgttttagcGggagctattacaatattattaacTgatcgaaatttaaatacttctttttttgaccctgccggaggaggagaccctattttatatcaacatctattt

#### Description

**Male**. Wing length, 2.59; width, 0.90. **Head**. Head light caramel-brown. Scape and pedicel light ochre-yellowish, flagellomeres light greyish-yellow, basal two flagellomeres lighter. Face and clypeus light caramel-brown. Palpomeres 1–3 light brownish-yellow, palpomeres 4–5 lighter. Labella cream-yellowish. Occiput with four longer setae dorsally to eye anteriorly to lateral ocellus, two dorsally to ocellus, eight posteriorly to ocellus. Scape about 2.0× pedicel length; flagellomere 4 1.7× longer than wide. Palpomere 4 1.1× palpomere 3 length, palpomere 5 1.5× palpomere 4 length. **Thorax**. Scutum compressed, caramel-brown on posterior two-thirds, ochre-orangish on anterior third and laterally to level of wing base, scutellum dark brown with ochre-yellow anterolateral corners. Antepronotum and proepisternum light ochre-yellow, proepisternum with a small ventral greyish-brown mark; anepisternum mostly ochre-brown, light on anteroventral corner; katepisternum light greyish-brown, mesepimeron, laterotergite, and metepisternum dark ochre-brown; mediotergite dark brown. Haltere pedicel light whitish-ochre. Pleural membrane yellowish. Scutum with eight supra-alar bristles, three pairs of prescutellar bristles, two additional smaller pairs in a slightly more anterior line, scutellum with three pairs of marginal bristles, outer pair smaller. Proepisternum with four bristles directed ventrally. Anepisternum with six bristles along posterior margin. Mesepimeron with four bristles and 35 small setae, laterotergite with three longer setae and 11 smaller setae. Metepisternum with 18 fine setae. **Legs**. Coxae and femora whitish with a brownish tinge, mid and hind femora with a brownish mark at tip; tibiae and tarsi ochre-yellowish, tarsi darker. Mid coxa with some few fine setae across basal fifth, hind coxa with fine setae on basal fourth. Fore tibia with two strong setae laterally on distal half, mid tibia with two dorsolateral rows with 4–6 bristles, ventrally a row of five strong setae. Mid and hind tarsomeres 1–3 with rows of ventral longer setae. Fore leg tarsomere 1 as long as tibia, 1.6× tarsomere 2 length. Hind tibia inner spur over 5× tibia width at apex. **Wing** (Fig. 145C). Membrane fumose brown, darker along anterior margin. Sc faint, with a row of dorsal macrotrichia. C produced slightly beyond tip of R_5_. R_1_ reaching C on distal sixth of wing; R_5_ reaching C slightly before level of tip of M_1_. First sector of Rs almost transverse, 0.54× r-m length; r-m slightly oblique. M_1+2_ short, 0.52× r-m length; bM slightly over 6× r-m length; first sector of CuA 0.34× length of second sector of CuA. Cubital pseudovein absent, CuP reaching level of origin of M_4_. Anal fold gradually curved, almost reaching wing margin. Posterior veins M_1_, M_2_, and M_4_ with dorsal macrotrichia along more than half their length, CuA with setae on distal fourth, macrotrichia on anal lobe membrane. **Abdomen**. Abdominal tergite 1 greyish-brown, tergites 2–6 brownish medially with cream-yellow area laterally, bands larger from segments 3–5, tergite 6 almost entirely cream-yellow, with only a brownish tinge medio-posteriorly, tergite 7 cream-yellow; sternites 1-7 whitish-yellow. Sternite 2 with a pair of strong brown bristles. **Terminalia** (Figs. 145D–E). Whitish-yellow. Gonocoxites fused medially, no suture present, bare ventrally, with two dorsolateral lobes: laterally a long, digitiform lobe with setae on outer face and a strong seta at tip, extending beyond tip of gonostylus; a blade-like projection dorsally as long as other lobe, slender and curved inwards towards tip, almost entirely bare except for three setulae apically. Gonostylus simple, elongate, dorsoventrally compressed, with setae at distal half on both faces, longer setae along inner margin. Aedeagus with an anterior medial ejaculatory apodeme, widening towards apex and then divided distally into a pair of separate tubular extensions. Parameres trapezoid, extending distally, some setulae apically and a pair of strong subapical straight blunt spines. Gonocoxal bridge weakly sclerotized, slender medially, no apodemes visible. Tergite 9 with a pair of long laterodistal extensions, longer than gonostylus or gonocoxite extensions, covered with microtrichia and setae. Cerci not visible.

**Female** (Figs. 145A–B). As male, except for the following. **Wing l**ength, 2.48; width, 0.94. **Head**. Occiput with three longer setae dorsally to eye anteriorly to lateral ocellus, two dorsally to ocellus, seven posteriorly to ocellus. **Thorax**. Scutum with six supra-alar bristles and three pairs of prescutellar bristles, three pairs of scutellar bristles, outer pair smaller. Proepisternum with three bristles directed ventrally, anepisternum with five bristles along posterior margin. Mesepimeron with two bristles and 39 small setae and setulae, laterotergite with one bristle, one long seta and 10 smaller setae. Metepisternum with five small fine setae. **Legs**. Fore tibia with two bristles at outer face, mid tibia with two dorsolateral rows of 4–5 bristles and three bristles along ventral edge, hind tibia two dorsolateral rows of six bristles and a row of six long setae on outer face. **Wing**. C not produced beyond tip of R_5_. Posterior veins M_1_ and M_2_ with over half of their length with dorsal macrotrichia, M_4_ and CuA with macrotrichia at distal fourth, macrotrichia on anal lobe membrane. **Terminalia**. Sternite 8 wide, no lateroposterior projections, posterior margin straight, covered with microtrichia and fine setae, longer setae on posterior margin. Sternite 9 wide medially, notum extending to anterior end of terminalia, blunt at tip, genital chamber slender, two gonoducts. Tergite 8 short medially, wide, a pair of wide ventrolateral lobes, no microtrichia or setae medially, setae restricted to lateral lobes. T9+10 bare, medially slender, lateral lobes fused to sternite 9. Cercomeres 1 and 2 probably fused, no sign of suture, basal half ovoid, distal half slender, covered with microtrichia and elongate setae, distal setae longer and curved.

#### Material examined

**Holotype**: male, ZRCBDP0047900, Nee Soon (NS2), swamp forest, 16.oct.2013, MIP leg. (slide-mounted). **Paratypes** (5 males, 12 females). **Males**: ZRCBDP0047919, Nee Soon (NS2), swamp forest, 27.nov.13, MIP leg.; ZRCBDP0048466, Nee Soon (NS1), swamp forest, 06.jun. 12, MIP leg. (slide-mounted); ZRCBDP0048852, Nee Soon (NS1), 14.jan.15, MIP leg.; ZRCBDP0048982, Nee Soon (NS2), 17.dec.14, MIP leg.; ZRCBDP0049123, Nee Soon (NS1), 24.dec.14, MIP leg.

**Females**: ZRCBDP0047816, Nee Soon (NS1), swamp forest, 18.dec.13, MIP leg.; ZRCBDP0048071, Nee Soon (NS1), swamp forest, 12.jun.13, MIP leg.; ZRCBDP0048441, Nee Soon (NS2), swamp forest, 30.may. 12, MIP leg. (website photo specimen, slide-mounted); ZRCBDP0048824, Nee Soon (NS1), 14.jan.15, MIP leg.; ZRCBDP0048851, Nee Soon (NS1), 14.jan.15, MIP leg.; ZRCBDP0048998, Nee Soon (NS2), 17.dec.14, MIP leg.; ZRCBDP0049013, Nee Soon (NS2), 17.dec.14, MIP leg.; ZRCBDP0049096, Nee Soon (NS1), 24.dec.14, MIP leg.; ZRCBDP0049115, Nee Soon (NS1), 24.dec.14, MIP leg.; ZRCBDP0049117, Nee Soon (NS1), 24.dec.14, MIP leg.; ZRCBDP0049248, Nee Soon (NS1), 10.dec.14, MIP leg; ; ZRCBDP0048440, Singapore, 04.apr.12.. **Additional sequenced specimens**: male, ZRCBDP0133442, Singapore, date range 2012-2018, MIP leg.; female, ZRCBDP0133915, Singapore, date range 2012-2018, MIP leg.; female, ZRCBDP0134054, Singapore, date range 2012-2018, MIP leg.; female, ZRCBDP0134063, Singapore, date range 2012-2018, MIP leg; ZRCBDP0048436, Singapore, 04.apr.12, ZRCBDP0048467, Nee Soon (NS1), 30.may.12.

**Etymology**. The species epithet of this species honors Professor Lam Toong Jin, Head of Zoology, National University of Singapore (1981–1996), then Director of School of Biological Sciences (1996–1998) and eventually Head of Department of Biological Sciences (1998–1999).

**Remarks**. There is a single haplotype for *Epicypta lamtoongjini* Amorim & Oliveira, **sp. n.**

*Epicypta catherinelimae* Amorim & Oliveira, sp. n. (Figs. 146A–D)

https://singapore.biodiversity.online/species/A-Arth-Hexa-Diptera-000778

urn:lsid:zoobank.org:act:8201362A-0752-4984-A1DB-84649F8D5913

**Diagnosis**. Head and scutum ochre-yellow, posterior end of scutum slightly darker, scutellum brown; antennal scape and pedicel ochre-yellowish, flagellum light ochre-brown; pleural sclerites ochre-brown, posterior sclerites darker. Coxae and femora whitish, on anterior leg with an orangish tinge, tip of mid and hind coxae and tip of hind femur with a light brown small mark. Wing membrane light brownish, area along cells c and br slightly darker. C barely extending beyond tip of R_5_; M_1+2_ 0.89× r-m length, r-m almost longitudinal. Dorsal macrotrichia on posterior veins M_1_ and M_2_, M_4_ and CuA close to tip, macrotrichia on anal lobe membrane. Abdominal tergites 1–2 light brown, tergites 3–5 light brown with slender cream-yellow laterals, slightly larger on tergites 4–5, tergite 6 mostly cream-yellow, light brown on lateroanterior corners, tergite 7 cream-yellow, terminalia whitish-yellow. Female terminalia sternite 8 trapezoid, elongate, no medio-posterior incision, genital furca long, slightly wider at anterior end, cercus slender, digitiform.

***Epicypta_catherinelimae_ZRCBDP0048468_hapZRCBD P0048468_SMH_holotype* [4: C, 7: G, 34: C, 118: C, 12 8: G, 129: T, 193: C, 196: G, 205: G] --**

tCtcGtcaactattgcccatgctgggtcttcCgttgatttagctattttttctcttcatttagcagggatttcttctattttaggagctattaattttattactacaattattaaCatacgttctGTaggaattacttttgatcgaatacccttatttgtatgatcagtattaattacagctattcttctCttGctttcattGcctgttttagcaggggctatcactatacttttaacagatcgtaatttaaatacctccttttttgaccctgcggggggaggagaccctattttatatcaacatttattt

#### Description

**Female** (Fig. 146A). Wing length, 2.14; width, 0.78. **Head**. Dark ochre-yellowish. Scape and pedicel greyish ochre-yellow, with a crown of darker, longer setae at distal margin, flagellomeres greyish-yellow. Face and clypeus ochre-yellowish. Palpomeres 1–3 light brownish-yellow, palpomeres 4–5 lighter. Labella whitish-yellow. Occiput with four longer setae dorsally to eye anteriorly to lateral ocellus, two dorsally to ocellus, ten posteriorly to ocellus. Scape 1.6× pedicel length, flagellomere 4 1.8× longer than wide. Palpomere 4 1.2× palpomere 3 length, palpomere 5 2.1× palpomere 4 length. **Thorax** (Fig. 146B). Scutum caramel-yellow, darker towards posterior end, scutellum blackish-brown with ochre-yellow anterolateral corners. Pleural sclerites caramel-yellow, proepimeron and katepisternum slightly lighter, mediotergite dark brown, lighter laterally. Haltere light ochre-yellowish with brown base of knob. Pleural membrane ochre-yellow. Scutum with 5+2 supra-alar bristles and long setae, three pairs of prescutellars; two pairs of scutellar bristles and one additional external pair of long setae. Proepisternum with three bristles, anepisternum with five bristles along posterior margin. Mesepimeron with two bristles and 17 small setae, laterotergite with two bristles and 15 small setae. Metepisternum with six fine setae. **Legs**. Coxae whitish, Fore coxa with a light orangish tinge; femora whitish with a light orangish tinge, mid and hind femora with brownish tips, tibiae and tarsi ochre-yellow with brownish tips, tarsi darker. Some few fine setae across basal fifth of mid coxa, hind coxa with fine setae along posterior half. Fore tibia with a single lateral bristle medially, mid tibia with a pair of irregular rows of 3-5 brownish bristles dorsolaterally, one bristle laterally, four bristles along ventral edge; hind tibia with a pair of irregular dorsolateral rows of 4-6 brownish bristles and three bristles laterally. Hind tibia inner spurs 4.1× tibia width at apex. **Wing** (Fig. 146C). Membrane fumose light brown, slightly darker along anterior margin. Sc short (visible only under phase contrast). C ending at apex of R_5_. R_1_ reaching C on distal fifth of wing; R_5_ reaching C at level of tip of M_1_. First sector of Rs slightly oblique, 0.72× r-m length; r-m oblique. M_1+2_ 0.89× r-m length; bM 6.1× r-m length; first sector of CuA 0.31× length of second sector of CuA. Cubital pseudovein not produced, CuP extending to slightly beyond level of origin of M_4_. Anal fold almost reaching wing margin. Posterior veins M_1_and M_2_ with dorsal macrotrichia along most of their length, M_4_ with macrotrichia along distal ¾, CuA with macrotrichia along distal fourth, anal lobe membrane with macrotrichia. **Abdomen**. Abdominal tergite 1 dark greyish-brown with a dark brown transverse band along posterior margin, tergite 2 greyish-brown, tergites 3–5 light brown, with slender cream-yellow area laterally, lateral band of tergite 5 wider, tergite 6 largely cream-yellow with light brown lateral band, tergite 7 cream-yellow; sternites 1-7 light ochre-yellowish. Sternite 2 with a pair of strong brown bristles. **Terminalia** (Fig. 146D). Ochre-yellowish. Sternite 8 subquadrate, lateroposterior ends slightly more projected than posterior margin medially, microtrichia and setae concentrated mainly on distal half, two pairs of long setae subapically. Sternite 9 anterior apodeme not extending beyond anterior end of terminalia, widening at anterior end, lateral arms sclerotized as a pair of diverging slender bands, genital chamber elongate. Sternite 10 with distal end acute, with some subapical setulae laterally. Tergite 8 short and bare medially, lateral lobes slightly projected posteriorly, entirely separated medially, microtrichia spread on posterior half, setae restrict to posterior margins. Tergite 9+10 with a slender medial bare sclerotized band connecting a pair of lobes fused to sternite 9 ventrolaterally. Cercomeres 1 and 2 fused, no sign of suture, basal ⅔ wider, posterior end digitiform, distal setae longer.

**Male**. Unknown.

#### Material examined

**Holotype**: female, ZRCBDP0048468, Nee Soon (NS2), swamp forest, 25.apr. 12, MIP leg. (slide-mounted). **Paratype**: female, ZRCBDP0048062, Nee Soon (NS2), swamp forest, 27.mar.13, MIP leg. (website photo specimen).

**Etymology**. The species epithet honors Catherine Lim Poh Imm (1942–), known as the “doyenne of Singapore writers”. Singaporean fiction author, she writes about the Singapore society and themes of traditional Chinese culture. She has published many collections of short stories, novels, and poetry collections. She was inducted into the Singapore Women’s Hall of Fame in 2014.

**Remarks**. All delimitation approaches point to separate species.

*Epicypta grootaerti* Amorim & Oliveira, sp. n. (Figs. 147A–C)

https://singapore.biodiversity.online/species/A-Arth-Hexa-Diptera-000779

urn:lsid:zoobank.org:act:77370B6D-9D79-401E-B74A-218F18A0F980

**Diagnosis**. Head brown, dark ochre-yellow posteriorly, antennal scape, pedicel, and first three flagellomeres ochre-yellowish, other flagellomeres grey-brownish, fourth flagellomere with a dark brown ring. Scutum dark ochre-yellow, with a blackish-brown mark on posterior fifth, scutellum blackish-brown; pleural sclerites dark ochre-yellow, darker on proepisternum, paratergite dark brown, laterotergite and mediotergite blackish-brown.

Fore coxa and femur whitish with orange tinge, mid and hind coxae whitish, mid and hind femora light whitish-yellow with a brownish line along ventral and dorsal edges. Wing membrane light brownish, darker along cells c and br; C not produced beyond tip of R_5_; M_1+2_ about as long as r-m. Dorsal macrotrichia on posterior veins M_1_ and M_2_, M_4_, and CuA close to tip, macrotrichia on anal lobe membrane. Abdominal tergite 1 brownish, tergites 2–5 ochre-yellowish with a slender brownish mark medially, tergites 6–7 and terminalia ochre-yellowish. Female terminalia sternite 8 trapezoid without medial incision posteriorly, genital furca long, not wider at anterior end.

***Epicypta_grootaerti_ZRCBDP0048128_hapZRCBDP004 8128_SMH_holotype* [28: T, 29: T, 121: G, 125: T, 130: C, 136: C, 137: T, 139: A, 193: T, 197: C, 199: T] --**

tttcttcaacaattgctcatgctggTTcttcagttgatttagctattttttctcttcatttagctggaatttcttctattttaggagcaattaattttattactacaattattaatatGcgaTctccCggaatCTcAtttgatcgaatacctttatttgtatgatctgtattaattacagctattttactTcttCtTtctttacctgttttagcaggagctattactatattattaacagatcgaaatttaaatacttctttttttgacccagcaggagggggggatcctattttataccaacatttattt

#### Description

**Female** (Fig. 147A). Wing length, 2.16 mm, width, 0.80 mm. **Head**. Dark ochre-yellowish, occiput lighter towards ventral margin. Scape and pedicel greyish ochre-yellow, flagellomeres 1–4 light ochre-yellow, flagellomere 4 light brown with a dark brown band on basal half, flagellomeres 5–14 light brown. Face and clypeus dark ochre-yellowish. Palpomeres 1–3 brownish-yellow, palpomeres 4–5 lighter; labella whitish-yellow. Occiput with four longer setae dorsally to eye anteriorly to lateral ocellus, two dorsally to ocellus, ten posteriorly to ocellus. Scape 1.6× pedicel length, flagellomere 4 1.5× as long as wide. Palpomere 4 1.5× palpomere 3 length, palpomere 5 1.7× palpomere 4 length. **Thorax**. Scutum caramel-yellow, a dark brown band at posterior sixth; scutellum blackish-brown with ochre-yellow anterolateral corners. Pleural sclerites caramel-yellow, except for proepisternum and mesepimeron blackish-brown, ochre-yellow laterally. Pleural membrane ochre-yellow. Scutum with 4+3 supra-alar bristles, three pairs of prescutellars; two pairs of scutellar bristles and one additional external pair of long setae. Proepisternum with three bristles, anepisternum with five bristles along posterior margin. Mesepimeron with two bristles and 12 small setae, laterotergite with two bristles and 13 small setae. Metepisternum with 23 fine setae. Haltere light ochre-yellowish with brown knob base. **Legs**. Coxae whitish, fore coxa with a light orangish tinge, basal sixth of hind coxa with a light brown band; femora whitish with an orangish tinge, mid and hind femora with brownish distal tips, tibiae and tarsi ochre-yellow with brownish distal tips, tarsi darker. Mid coxa with a band of fine setae across proximal fifth, hind coxa with setae covering basal fourth. Fore tibia with two bristles dorsolaterally, mid tibia with a pair of irregular dorsolateral rows of 4-5 bristles, one bristle laterally on outer face, and three long setae along ventral edge. Fore leg tarsomere 1 1.1× tibia length, 1.3× tarsomere 2 length. Hind tibial inner spur 4.9× tibia width at apex. **Wing** (Fig. 147B). Membrane fumose light brown, slightly darker along anterior margin. C not produced beyond tip of R_5_. Sc barely produced. R_1_ reaching C on distal fourth of wing; R_5_ reaching C at level of tip of M_1_. First sector of Rs slightly oblique, 0.7× r-m length; r-m oblique. M_1+2_ 0.76× r-m length; bM 5.9× r-m length; first sector of CuA 0.34× length of second sector. Cubital pseudovein absent, CuP extending to slightly beyond level of origin of M_4_. Anal fold almost reaching wing margin. Posterior veins M_1_ and M_2_ with dorsal macrotrichia along almost entire length, M_4_ along distal three-fourth, CuA on distal fourth, anal lobe membrane with macrotrichia. **Abdomen**. Abdominal tergite 1 brown, with a dark brown transverse band close to posterior margin, lighter laterally, tergites 2–5 light brown medially, with a wide cream-yellow area laterally, tergites 6–7 light brownish-yellow; sternites 1-7 light ochre-yellowish. Sternite 2 with a pair of strong brown bristles. **Terminalia** (Fig. 147C). Ochre-yellowish. Sternite 8 trapezoid, slender, posterior margin with no incision medio-posteriorly, scattered microtrichia and setae, setae along posterior margin longer. Sternite 9 slender, anterior apodeme extending slightly beyond anterior end of terminalia, pointed at apex, genital chamber elongate, distal end acute, with some subapical setulae laterally. Tergite 8 short, lateral lobes slightly projected posteriorly, not in contact medially, microtrichia spread on posterior half, setae restrict to posterior margins. Tergite 9+10 with a slender medial bare sclerotized band connecting a pair of lobes fused to sternite 9 lateroventrally. Cercomeres 1 and 2 fused, no sign of suture, basal ⅔ wider, posterior end digitiform, distal setae longer.

**Male**. Unknown.

#### Material examined

**Holotype**: female, ZRCBDP0048128, Pulau Semakau (SMO2), old mangrove, 11.jul.13, MIP leg. (website photo specimen, slide-mounted).

**Etymology**. The species epithet of this species honors Patrick Grootaert, member of the Royal Belgian Institute of Natural Sciences, Belgium. Specialist in Dolichopodidae, the long-legged flies, he is one of the researchers associated to the Singapore Mangrove Insect Project.

**Remarks**. This species seems close to *Epicypta pallida* (Edwards), from Krakatau, with minor differences in the color of the antenna, mesepimeron and metepisternum, and in the extension of C beyond the tip of R_5_.

*Epicypta joaquimae* Amorim & Oliveira, sp. n. (Figs. 148A–D)

urn:lsid:zoobank.org:act:BAB60260-2ED4-4368-A227-F3D8D252F39A

**Diagnosis**. Head ochre-yellow, antennal scape, pedicel and flagellomeres 1–3 ochre, ochre basally with a brownish ring distally. Scutum ochre-yellow, scutellum light yellowish-brown; pleural sclerites dark ochre-yellow, laterotergite with a dark brown mark. Coxae and femora whitish, fore coxa with brownish tinge, femora dorsal crest greyish-brown. Wing membrane light brownish, cells c and br more yellowish. C not extending beyond tip of R_5_; M_1+2_ 0.88× r-m. Dorsal macrotrichia on posterior veins M_1_, M_2_, posterior half of M_4_ and distal end of CuA; macrotrichia on anal lobe membrane. Abdominal tergite 1 ochre-yellow, tergites 2–4 ochre-yellowish with a brownish diffuse mark medially, tergite 5 and 7 entirely ochre-yellowish, tergite 6 ochre-yellowish with two pairs of brownish marks laterally, one at anterior margin and another one at posterior margin; terminalia whitish-yellow. Female sternum 8 with a shallow incision medially along posterior border, genital furca wide at anterior end.

***Epicypta_joaquimae_ZRCBDP0072680_hapZRCBDP00 72680_SMH_holotype* [28: G, 76: C, 79: T, 121: G, 125: T, 127: T, 133: T, 137: T, 155: C]**

tctttcttctactattgctcatgcaggGgcatctgttgatttagctattttttctttacatttagcaggaatttcCtcTattttaggagctattaattttattacaacaattattaatatGcgaTcTcctggTattTcttttgatcgaatacctCtatttgtatgatctgttttaattacagctgttcttttacttttatctttacctgtattagcaggagctattactatattattaacagatcgaaacttaaatacttcattttttgaccctgcaggagggggagatcctattttatatcaacatttattt

#### Description

**Female** (Fig. 148A). Wing length, 2.30; width, 0.83. **Head**. Vertex dark yellowish-brown, occiput lighter laterally. Scape and pedicel ochre-yellow, flagellomeres light-brown. Face and clypeus light brown; palpomeres 1–3 light yellowish-brownish, palpomeres 4–5 lighter; labella whitish-yellow. Occiput with two longer setae dorsally to eye anteriorly to lateral ocellus, one dorsally to ocellus, nine posteriorly to ocellus. Scape about 1.9× pedicel length; flagellomere 4 1.9× longer than wide. Palpomere 4 1.3× palpomere 3 length, palpomere 5 2.0× palpomere 4 length. **Thorax** (Fig. 148B). Scutum yellowish-brown, scutellum light yellowish-brown. Antepronotum and proepisternum light brown, anepisternum, laterotergite, and mediotergite yellowish-brown, katepisternum ochre-brown, mesepimeron brown on dorsal half, ochre-brown on ventral half, metepisternum brown. Pleural membrane ochre-yellow. Haltere with whitish-yellow pedicel, knob brownish on proximal half, whitish-yellow on distal half. Scutum with 6+3 supra-alars and three pairs of prescutellar bristles; three pairs of scutellar bristles, outer pair smaller. Proepisternum with three bristles, anepisternum with five bristles along posterior margin. Mesepimeron with one bristle and 17 small setae, laterotergite with four bristles and 6 small setae, posterior bristle smaller. Metepisternum with seven fine setae. **Legs**. Coxae whitish-yellow with orangish tinge, hind coxa with a brown band on basal fifth; fore femur yellowish-brown, mid and hind femora lighter, tibiae and tarsi light ochre-yellow. Fore tibia with dorsal rows of strong setae, two dorsolateral small brown bristles medially; mid tibia with three irregular dorsolateral rows of 3–5 bristles, one strong lateral seta on inner face and a row with four bristles along ventral edge; hind tibia with two dorsolateral rows of 4–5 bristles and four long setae on inner face. Fore leg tarsomere 1 1.0× tibia length, 1.6× tarsomere 2 length. Hind tibia inner spur over 5× tibia width at apex. **Wing** (Fig. 148C). Membrane light brown fumose, darker along anterior margin. Sc faint, incomplete (visible in phase contrast) with dorsal macrotrichia. C ending at tip of R_5_. R_1_ reaching C at wing distal fourth; R_5_ reaching C slightly before level of M_1_. First sector of Rs oblique, bare, 0.40× r-m length; r-m almost longitudinal. M_1+2_ 0.88× r-m length; bM 4.7× r-m length. First sector of CuA 0.26× length of second sector of CuA. Cubital pseudovein absent, CuP barely reaching level of origin of M_4_. Anal fold gently curved along basal half, almost reaching wing margin. Posterior veins M_1_ and M_2_ with of dorsal macrotrichia on almost entire length, M_4_ on distal two-thirds, CuA on distal fourth, macrotrichia on anal lobe membrane. **Abdomen**. Abdominal tergite 1 ochre-yellow, tergites 2–4 ochre-yellowish, with a diffuse brownish mark medially, tergite 5 and 7 entirely ochre-yellowish, tergite 6 ochre-yellowish with two pairs of brownish marks laterally, one at anterior margin and another one at posterior margin; sternite 2 with a pair of long, slightly curved ventral bristles. **Terminalia** (Fig. 148D). Light brownish-yellow. Sternite 8 elongate, trapezoid, posterior margin nearly straight, with a short medial incision, microtrichia and setae evenly distributed, long setae at posterior margin. Sternite 9 elongate, anterior apodeme extending to anterior end of terminalia, wide, genital chamber elongate. Tergite 8 with a pair of separate large lateral lobes with microtrichia and setae. Tergite 9+10 slender, a pair of sclerotized bands connected medially, lateroposteriorly fused to sternite 9. Cerci long, cercomeres 1 and 2 fused, no sign of suture, elongate, wider midway to apex posterior end digitiform, distal setae longer.

#### Material examined

**Holotype**: female, ZRCBDP0072680, Bukit Timah Forest (BT05), 22.dec. 16, MIP leg. (slide-mounted). **Paratypes** (1 female): ZRCBDP0137314, Bukit Timah (BT05), 17.may.17.

**Etymology**. The species epithet of this species honors Agnes Joaquim (or Ashkhen Hovakimian) (1854-1899). A Singaporean Armenian horticulturalist, who bred the first hybrid orchid, the Vanda ‘Miss Joaquim’, now the national flower of Singapore.

### Aspidionia Colless

*Aspidionia* Colless 1966: 664. Type-species, *Aspidionia palauensis* Colless (orig. des.).

**Diagnosis**. Head anteroposteriorly compressed, partially under anterior end of scutum, vertex displaced to a frontal position. Scutum lateral margin with sharp incision at above level of anterior spiracle; shining medial keel on anterior third of scutum due to modification of setulae sockets. Antepronotum lateral lobes completely separated, katepisternum strongly compressed; laterotergite and mediotergite small, strongly compressed. M_1+2_ long, over 3× r-m length, aligned with M_2_; M_4_ missing; CuA straight, distance from tip of CuA to tip of M_2_ as long as from tip of M_2_ to tip of M_1_. Anal fold long, straight, only gently curved close to posterior margin.

This is one of the least known mycetophilid genera worldwide. Originally described from the Caroline Island, Palau, in Micronesia based on two specimens (Colless 1966), it later had a second species formally described based on six specimens from the Comores Island (Matile 1974b)—between Madagascar and continental Africa. With this paper, the number of described species in the genus moves from two to five. Colless (1966) also referred to an additional undescribed species from Australia, making this another case of Indo-Pacific distribution, quite similar to that of *Platyprosthiogyne*. *Aspidionia palauensis* Colless was described based on males and *A. balachowski* Matile was described based on males and females. The genus is rare in our Singapore samples, with only four females known of three species altogether in our samples.

Two of the Singapore species—*Aspidionia cheesweeleeae* Amorim & Oliveira, **sp. n.** and *Aspidionia fatimahae* Amorim & Oliveira, **sp. n.**—are brought together by mPTP, but all other criteria show the three of them as separate species, what is corroborated by morphological data. As well, different aspects of the morphology show clear differences between these two species. *A. fatimahae* Amorim & Oliveira, **sp. n.** has two haplotypes and the other two species have only one (Fig. 150B). All three species are described based only on females, since there is enough information to discriminate between them and we have a diagnoses based on the color patterns and other details of the general morphology.

*Aspidionia cheesweeleeae* Amorim & Oliveira, sp. n. (Figs. 149A–D, 150A)

https://singapore.biodiversity.online/species/A-Arth-Hexa-Diptera-000818

urn:lsid:zoobank.org:act:5C7605D5-3BB4-4428-8FB1-3C242CB80EF8

**Diagnosis**. Scutum mostly ochre-yellowish, with a dark brown band along anterior margin, a dark brown mark above wing and a dark brown band along posterior end. Thoracic pleura mostly ochre-brown with a light brown mark on posterior half of anepisternum, mid and hind coxae entirely whitish, hind coxa with a brown basal band. Wing vein r-m present. Abdominal tergites ochre-yellowish with a brown medial longitudinal band, sternites ochre-yellowish.

***Aspidionia_cheesweeleeae_ZRCBDP0049018_hapZRCB DP0049018_SMH_holotype* [13: C, 16: A, 76: A, 214: T, 223: A, 241: T, 283: T]**

cttatcttcatcCatAgctcacgcaggagcatcagtagatttagcaattttctctttacatttagccggtatttcAtcaattttaggagcaattaattttattactacaattattaatatacgatctcctgggatatattttgatcgtttacctctatttgtatgatctgttttaattacagcaattctcttattactatctttacctgttctTgctggagcAattactatacttttaacTgaccgaaatttaaatacttcattttttgatcctgctggtggTggagacccaattttataccaacatttattt

#### Description

**Female** (Fig. 149A). Wing length, 1.66; width, 0.67. **Head** (Figs. 149B–C). Brown medially on frons, brownish-yellow around eye, head partially fit under anterior end of scutum. Face and clypeus dirty-yellowish. Lateral ocelli blackish-brown, nearly touching eye margin, mid ocellus small, at dorsal end of short frontal furrow. Antennal scape and pedicel light brown, flagellum brown. Maxillary palpus light brown, lighter towards apex. Labella light brownish-yellow. Dark brown small setae scattered over vertex, four slightly longer setae on occiput around dorsal margin of eye posteriorly to ocellus, a line of long setae close to anterior margin of frons. Face slender, with a transverse line of setulae. Clypeus with scattered setulae. Antennal scape twice pedicel length, a crown of setae distally and on inner face of scape; setulae on both lateral faces and on distal margin of pedicel, in addition to one strong dorsal seta. Flagellomere 1 twice flagellomere 2 length; flagellomere 4 1.5× longer than wide. Palpomere 1 twice length of palpomere 2, covered only with microtrichia, palpomere 2 short, with some setulae, palpomere 3 slightly longer than wide, sensorial pit opening conspicuous, dorsal, setulae on external and dorsal faces, palpomere 4 almost twice palpomere 3 length, with setulae on external and dorsal faces, palpomere 5 slender, about twice palpomere 4 length, with scattered setulae. **Thorax**. Scutum ochre-yellow, with an elongate yellowish-brown mark along anterior margin, a brown mark along margin above anepisternum and a transverse band at posterior end of scutum; scutellum dark brown. Pleural sclerites mostly ochre-brown, basisternum brown, proepisternum with an ochre-yellow area on distal half, anepisternum with a brown mark on posterior half, mediotergite dark brown. Scutum densely covered with scattered fine setae, shiny median keel anteriorly (composed of modified setal sockets), five longer setae at small bulging area on margin above level of wing, four pairs of prescutellar bristles along posterior margin of scutum; a deep incision at margin of scutum above level of anterior spiracle. Scutellum large, trapezoid, two pairs of strong bristles aligned at distal end, additional setae along posterior margin. Basisternum dorsoposterior arms with some few setulae. Antepronotum lateral lobes completely separated, medial connection not sclerotized, only with short fine setae, proepisternum with three longer setae and two additional slightly smaller setae along ventral margin, scattered small setae on sclerite. Proepimeron reduced to a slender stripe at ventrodistal end of proepisternum, reaching anterodorsal end of katepisternum. Anepisternum covered with scattered setulae, a line of four long setae along posterior margin; katepisternum strongly compressed dorsoventrally, about a third of height of anepisternum. Mesepimeron with two bristles along dorsoposterior end, not reaching ventral margin of thorax, a small connection between katepisternum dorsoposterior end and laterotergite anterior end. Laterotergite bulging, with two long setae and seven small setae; metepisternum with 14 fine setae along its length; mediotergite small, strongly folded, bare. **Legs**. Fore coxa ochreish with an orangish tinge, mid and hind coxae whitish, hind coxa with a brown transverse band at basal end; femora whitish-yellow, fore and mid femora with brownish-yellow tinge, mid femur brownish along dorsal edge close to basal end, hind femur whitish, with brown dorsal edge; tibiae and tarsi light yellowish-brown with an orangish tinge. Fore coxa entirely covered with setulae at anterior face, a row of brown bristles along posterior margin and distal margins; mid coxa largely developed, anterior face covered with setae, a line of bristles along distal margin; hind coxa with a line of small setae on distal half of external face, including one subapical strong seta, a few small setae and one longer seta at anterior face distally. Femora covered with fine setae, a row 3–5 longer setae along ventral margin distally, stronger on hind femur. Tibiae and tarsi with regular rows of trichia. Fore tibia with a wide anteroapical depressed area lined with setulae and some few strong setae at distal end; mid tibia with two rows of bristles dorsally and two bristles on internal face, in additional to a bristle at distal end; hind tibia with two rows of laterodorsal setae along entire length separated by a flat, bare area, a comb of long setae at internal face of distal end of tibia. First fore tarsomere shorter than tibia, about twice tarsomere 2 length. Fore leg tarsomere only with rows of trichia and a couple of distal setae; mid and hind tarsomeres 1–4 with rows of ventral setae besides rows of trichia. Tibial spurs yellowish-brown, mid tibia spur almost twice length of inner spur, outer spur of mid tibia about 5× tibial apex, outer spur of hind leg about 3× tibial apex. Tarsal claws with a long basal tooth. **Wing** (Fig. 150A). Wing membrane light yellowish-brown fumose, slightly darker along anterior margin. Membrane densely covered with regularly organized microtrichia on all cells, no macrotrichia on membrane; posterior margin emarginated at level of tip of CuP. Sc faint; R_1_ reaching C at distal third of wing; R_4_ absent; R_5_ short, reaching C at level of M_2_, running almost parallel to R_1_, gently curved on distal third. C extending beyond R_5_ for half distance to M_1_. First sector of Rs slightly oblique, 1.1× r-m length; r-m short, oblique. M_1+2_ 5.3× r-m length; M_1_ and M_2_ gently diverging on distal half; M_4_ absent; bM over 7× r-m length; posterior veins weak at tip. CuA straight, long, reaching posterior wing margin slightly beyond level of tip of R_1_. Cubital pseudovein absent, CuP produced to level of basal end of M_1+2_. Anal fold long, almost straight, not reaching wing margin. Dorsal macrotrichia on bR, R_1_, and Rs, ventral macrotrichia on distal half of bR, R_1_, and on second sector Rs. **Abdomen**. Tergite 1 whitish laterally with brown medial mark, tergites 2–6 light yellowish-brown laterally and brownish medially; sternite 1 whitish, sternite 2–6 light yellowish-brown, darker towards distal segments; tergite and sternite 7 yellowish. **Terminalia** (Fig. 149D). Yellowish, small, weakly sclerotized. Sternite 8 trapezoid, a pair of distal lobes reaching level of tip of cercomere 2, with a median incision at posterior margin, microtrichia and elongate setae on ventral face, laterally bare; sternite 9 extending distally to level of tip of lobes sternite 8, anterior end extending to distal third of segment Tergite 8 rectangular, longer than tergite 9+10. Tergite 9+10 short. Cercus short, cercomere 1 over twice length of cercomere 2.

**Male**. Unknown.

#### Material examined

**Holotype**: female, ZRCBDP0049018, Nee Soon (NS2), 17. dec. 2014, MIP leg. (website photo specimen, slide-mounted).

**Etymology**. The species epithet of this species honors Chee Swee Lee (1955–), Singapore-born sprinter who won medals at the Southeast Asian Games (SEAG) in 1969, 1971, and 1973, before becoming the first woman from Singapore to win a gold medal and break a record at the Asian Games, in the 1974. She won a second gold medal the subsequent year before retiring from sports. She was inducted into the Singapore Women’s Hall of Fame in 2014.

**Remarks**. An ochre-yellow background color of *Aspidionia cheesweeleeae* Amorim & Oliveira, **sp. n.** is shared with *A. fatimahae* Amorim & Oliveira, **sp. n.,** and these two species are probably closer to each other than any of them with *A. janetjesudasonae* Amorim & Oliveira, **sp. N.**

*Aspidionia janetjesudasonae* Amorim & Oliveira, sp. n. (Figs. 151A–F)

https://singapore.biodiversity.online/species/A-Arth-Hexa-Diptera-000831

urn:lsid:zoobank.org:act:4BBBFB35-C6FC-4657-8D9D-A44F27E53A30

**Diagnosis**. Scutum mostly blackish-brown, with yellowish band along on anterior fifth. Thoracic pleura dark brown, mid coxa whitish with slender brown band at base, hind coxa whitish with brown mark at anterior third. Wing vein r-m absent. Abdomen with tergites and sternites brown.

***Aspidionia_janetjesudasonae_ZRCBDP0048147_hapZR CBDP0048147_SMH_holotype* [22: C, 55: A, 64: T, 76: A, 125: A, 126: G, 127: A, 238: T]**

tctttcttcaacaattgctcaCgcaggagcatcagtagatttagcaattttttcAttacatctTgcaggtatttcAtcaattttaggagcaattaattttattactacaattattaatatacgaAGAccaggaatatattttgatcgaataccattattt gtttgatctgttttaattactgctattctattattattatctttaccagtattagcaggagctattactatattactTacagatcgaaatttaaatacttcattttttgatccagctggaggaggggatcctatcttatatcaacatttattt

#### Description

**Female** (Fig. 151A). Wing length, 1.47; width, 0.54. **Head**. Dark brown, face and clypeus dirty-yellowish. Antennal scape and pedicel ochre-yellowish, flagellum light brown; flagellomere 4 1.3× longer than wide. Palpus and labella light yellowish-brown. **Thorax** (Figs. 151B– C). Scutum shinning dark brown except for ochre-yellow transverse band along anterior margin, two pairs of long prescutellar bristles and an additional pair of smaller bristles. Scutellum dark brown, with two pairs of strong bristles and two pairs of small additional small setae. All pleural sclerites dark brown except for brown antepronotum, proepisternum, and metepisternum. Antepronotum and proepisternum only with short setae. Anepisternum entirely covered with scattered setae, five longer setae along posterior margin. Mesepimeron with two strong bristles at dorsal margin. Laterotergite with five elongate setae; metepisternum with nine elongate setae. **Legs**. Fore coxa whitish with orangish tinge, mid and hind coxae whitish with a brown transverse band across basal end; fore and mid femora whitish-yellow, hind femur mostly whitish-yellow, a dark brown mark along dorsal edge and distal third of femur. Tibiae light yellowish-brown, tarsi light brown. Fore tibia with regular rows of setulae, but no distinctive setae. Haltere white. **Wing** (Fig. 151D). Wing membrane fumose light brown, slightly darker along anterior margin. C extending beyond tip of R_5_ over half of distance to M_1_. Sc short, ending free. First sector of Rs nearly transverse; first half of R_5_ straight, gently curved on distal half, not too close to R_1_ and C; r-m basically absent, origin of M_1+2_ at level of distal end of bM; M_1+2_ relatively long, about 4× length of first sector of Rs. Medial fork long, M_1_ and M_2_ gently diverging towards apex; tip of posterior veins weakly sclerotized; CuA straight, reaching margin before level of tip of R_5_. Sclerotized anal fold present, long, mostly straight. Dorsal setae present on bR, R_1_, R_5_, tip of bM, and distal ⅔ of M_1_, ventral setae on bR, R_1_ and R_5_. **Abdomen**. Tergites 1–7 brown, sternite 1 white, sternites 2–7 light brown. **Terminalia** (Figs. 151E–F). Yellowish. Sternite 8 subtriangular with rounded distal end, reaching level of tip of cercomere 2, extending laterally, microtrichia and elongate setae on ventral face, laterally bare; sternite 9 extending medially to level of posterior margin of sternite 8, anterior end with two rounded short lateral lobes, reaching distal third of segment 7. Tergite 8 rectangular, longer than tergite 9+10. Tergite 9+10 short. Cercomere 1 almost 3× cercomere 2 length.

#### Material examined

**Holotype**: female, ZRCBDP0048147, Sungei Buloh (SB1), mangrove, 02.oct.2013, MIP leg. (website photo specimen, slide-mounted). **Additional sequenced specimen**: ZRCBDP0314085.

**Etymology**. The species epithet of this species honors Janet “Speedy Gonzales” Jesudason (1936–), Singapore-born pioneer women who represented Singapore in the 1956 Olympics in the 100 meters sprint. She was inducted into the Singapore Women’s Hall of Fame in 2016.

*Aspidionia fatimahae* Amorim & Oliveira, sp. n. (Figs. 152A–E)

https://singapore.biodiversity.online/species/A-Arth-Hexa-Diptera-000822

urn:lsid:zoobank.org:act:7C794F02-79E5-4FF8-A679-124E0B53B737

**Diagnosis**. Scutum ochre-yellowish, with a dark brown band along anterior margin, and a curved brown mark above anepisternum connecting to brown area on posterior fourth of scutum; scutellum dark brown. Anterior half of anepisternum and dorsoposterior corner of katepisternum greyish-ochre, antepronotum, proepisternum, anteroventral corner of anepisternum, katepisternum and mesepimeron brownish, a brown mark on dorsoposterior corner of anepisternum, mediotergite dark brown. Mid and hind coxae whitish, hind coxa with a greyish-brown mark at basal fourth. Wing vein r-m present. Abdomen with ochre-brownish tergites and ochre-yellowish sternites.

***Aspidionia_fatimahae_ZRCBDP0047810_hapZRCBDP0 047810_SMH_holotype* [41: C, 97: C, 124: C, 133: C, 15 2: T, 280: C]**

actttcttcttctattgctcatgcaggtgcatcagttgatCtagcaattttttcccttcacttagcaggtatttcttcaattttaggagcaattaaCtttattactacaattatcaatatacgCtcccccggCatatattttgatcgattaTctttatttgtatgatctgttttaatcacagcagttcttttattattatcattaccagtattagctggagctattacaatattattaactgatcgaaatttaaatacatctttttttgacccagcaggCgggggagacccaattttatatcaacatttattt

#### Description

**Female** (Fig. 152A). Wing length, 1.84; width, 0.67. **Head** (Fig. 152B). Brown, face and clypeus light brown. A line of five longer setae on occiput around dorsal margin of eye posteriorly to ocellus. Antennal scape, pedicel and basal half of flagellomere 1 whitish-yellow, distal half of flagellomere 1 and other flagellomeres brown; flagellomere 4 1.8× longer than wide. Palpus whitish, labella whitish-yellow. **Thorax**. Scutum background light ochre-yellow, a dark brown transverse band along anterior margin, a pair of curved elongate brown maculae above anepisternum connecting medially to a brown macula across posterior fourth of scutum. Scutellum dark brown. Antepronotum brown, lighter on posterior half, proepisternum brownish. Ventroanterior corner and posterior third of anepisternum dark brown, ochre medially, as well as posterior fourth of katepisternum; proepisternum, most katepisternum, mesepimeron, laterotergite, and metepisternum brownish, mediotergite dark brown. Antepronotum with some few short setae, proepisternum with short setae and a line with four long setae. Anepisternum with two long setae on ventroanterior corner and four long setae along posterior margin. Mesepimeron with two strong setae close to dorsal margin. Laterotergite with 16 setae; metepisternum with 14 fine setae. Haltere pedicel light brown, knob mostly cream brown. **Legs**. Fore coxa light yellowish-brown, mid and hind coxae whitish, hind coxa with a brown transverse band along basal end; mid and hind femora light yellowish-brown, mid femur with a short brown band at dorsal edge close to anterior end, hind femur with a dark brown longitudinal band along dorsal edge; tibiae and tarsi yellowish-brown, tarsi slightly darker towards tip [fore femora, tibiae and tarsi, and mid tibiae and tarsi missing]. **Wing** (Fig. 152C). Wing membrane light brown fumose. C extending beyond tip of R_5_ to almost half distance to M_1_. First sector of Rs almost transverse; R_5_ running not too close to R_1_ and to C, straight on basal two-thirds, gently curved distally. First sector of Rs 1.0× r-m length, r-m short but produced, M_1+2_ 4.0× r-m length; base of M_1_ weakly sclerotized; M_1_ and M_2_ only gently diverging, tip of M_1_, M_2_, and CuA barely sclerotized at very tip; CuA straight. Cubital pseudovein not produced, CuP present to slightly beyond basal end of M_1+2_; sclerotized anal fold long, straight. Setae present dorsally on bR, R_1_, R_5_, r-m, and distal end of bM, entirely absent on medial and cubital veins, setae ventrally on bR, R_1_ and R_5_. **Abdomen**. Tergite 1 brown, tergites 2–6 brown medially, ochre-yellowish laterally, tergite 7 yellowish-brown. Sternites 1–7 yellowish-brown. **Terminalia** (Figs. 152D–E). Sternite 8 subtriangular, with rounded distal end, reaching level of tip of cercomere 2, extending laterally, microtrichia and elongate setae on ventral face, laterally bare; sternite 9 posterior end extending to level of posterior margin of sternite 8, anterior end wide. Tergite 8 rectangular, longer than tergite 9+10, mostly covered with microtrichia, three elongate setae at slightly projected lateroposterior corner, a short rounded incision present medially. Tergite 9+10 short, lateroposterior corners slightly extended posteriorly, with one seta at each lateroposterior corner. Cercomere 1 about 2× cercomere 2 length.

#### Material examined

**Holotype**: female, ZRCBDP0047810, Nee Soon (NS1), swamp forest, 01.may. 2013, MIP leg. (slide-mounted) (website photo specimen). **Additional sequenced specimens:** ZRCBDP0133550, National University of Singapore.

**Etymology**. The species epithet of this species honors Hajjah Fatimah binte Sulaiman (1754?–1852?). Born in Malacca, she was a Singaporean merchant and philanthropist. She donated money and land for the establishment of the mosque Masjid Hajjah Fatimah, which has her name, as well as funded homes for the poor adjacent to it. Fatimah was inducted into the Singapore Women’s Hall of Fame in 2014.

**Remarks**. We have two haplotypes and no delimitation conflicts.

*Integricypta* Amorim & Oliveira, gen. n. *Integricypta*Amorim & Oliveira. Type-species, *Integricypta shirinae* Amorim & Oliveira

urn:lsid:zoobank.org:act:B6AC9706-A13A-4562-A88B-BB739D807755

**Diagnosis**. Head partially fit under anterior end of scutum, antero-posteriorly compressed, vertex largely displaced to a frontal position; long interommatidial setae. Mid ocellus absent. Scutum lateral border with sharp incision above antepronotum; no shining medial keel on scutum. Katepisternum strongly compressed; laterotergite and mediotergite small, strongly compressed. Wing membrane with a dark brown mark over area around origin of Rs; M_1+2_ only slightly longer than r-m length; M_2_ not aligned basally to M_1+2_; M_4_ present, arched, anterior end disconnected from CuA, slightly converging towards CuA on distal third; CuA long, straight, tip before level of tip of R_5_; anal fold long, almost straight.

Some of the parallel evolution of features often used in identification keys for genera of Mycetophilini leads to a puzzling situation. The species of *Integricypta* Amorim & Oliveira, **gen. n.** have M_4_ (as *Mycetophila*, *Epicypta* and *Platurocypta*), differently from several Mycetophilini genera and mycetophilid genera in other subfamilies, that have the base of M_4_ disconnected from CuA. The origin of M_4_ in the wing in *Integricypta* Amorim & Oliveira, **gen. n.,** however, is beyond the level of the base of the medial fork and with its base disconnected from CuA. The slightly curved M_4_ towards CuA at its distal fourth, diverging from M_2_is also seen in *Mycetophila*. The wing membrane has a dark mark medially along the anterior margin of the wing, as some few species of *Epicypta*. It has a long C beyond the tip of R_5_, as some of the species of *Platyprosthiogyne*, diverging from what is seen in *Epicypta*.

Helpful in this discussion is the fact that the incision at the margin of the scutum above the anterior spiracle is a uniquely derived feature of the clade including *Integricypta* Amorim & Oliveira, **gen. n.,** *Epicypta*, *Platurocypta* and *Aspidionia*. This feature alone places this new genus away from the remaining genera of Mycetophilini. The dorsoventral compression of the katepisternum, the reduction in size of the laterotergite and the mediotergite, the largely developed mid coxa, and the size and position of the head are additional derived features shared by this small mycetophiline clade. The set of species of *Integricypta* Amorim & Oliveira, **gen. n.** seem to be sister to *Aspidionia*. These species could be included in *Aspidionia*, but this would require significant changes to the diagnosis of the genus. M_4_ is entirely missing in *Aspidionia* and the modified sockets on the scutum anteromedial “keel” are unique for this genus, absent in *Integricypta* Amorim & Oliveira, **gen. n.**

This new genus raises the number of genera in the Mycetophilini to 14. It is interesting to note that this entire clade of mycetophilines have largely an Indo-Pacific distribution: *Aspidionia*, *Platyprosthiogyne* and *Integricypta* Amorim & Oliveira, **gen. n.** are exclusively distributed in this area, where *Platurocypta* has most of its diversification.

A new genus with five new species described here, two of which quite abundant, clearly depicts our deficit of understanding of the biodiversity of dark taxa in tropical forests.

There are no conflicts between the different delimitation approaches of the species of *Integricypta* Amorim & Oliveira, **gen. n.** (Fig. 154E). Differently from *Aspidionia*, where all three species are rare in our samples, *Integricypta shirinae* Amorim & Oliveira, **sp. n.** is quite abundant, present in freshwater swamp, mangrove and urban forest traps.

**Etymology**. The name of the genus is feminine and brings together the Latin word *integrum*, for whole, complete, and the Greek word κύπτω—borrowed from the related genus *Epicypta*—, that means “bending the head forward”. The name is a reference to integrative taxonomy, with reciprocal illumination between morphology and molecular data as sources of information.

*Integricypta fergusondavie* Amorim & Oliveira, sp. n. (Figs. 153A–G, 154A–D)

https://singapore.biodiversity.online/species/A-Arth-Hexa-Diptera-000827

urn:lsid:zoobank.org:act:D98967C1-1DEF-40A7-BEA0-49E89784721A

**Diagnosis**. Head blackish-brown, flagellum dark brown with lighter scape and pedicel. Scutum and pleural sclerites blackish-brown. Coxae and femora mostly whitish, mid and hind coxae with brown transverse band at proximal end, hind femur brown at distal third, anterior end of hind tibia with a brown mark. Wing membrane with dark brown mark along area of origin of Rs; C produced beyond tip of R_5_ on ⅔ of distance to M_1_; r-m oblique; M_1+2_1.4× r-m length; wing margin emarginated at level of tip of CuA. Abdominal tergites brown. Gonocoxites large, no medioventral process, no gonocoxite distal projection beyond base of gonostylus; gonostylus with an elongate ventral lobe with macrosetae on ventral end, a small digitiform median lobe with an apical seta and a weakly sclerotized dorsal lobe with a dorsal and a ventral sublobe with a megaseta apically.

***Integricypta_fergusondavie_ZRCBDP0049297_hapZRC BDP0049297_SMH_holotype* [29: T, 67: T, 122: A, 123: A, 125: T, 127: C, 146: A, 155: C, 181: T, 229: C]**

cctatcatcttcattagctcatgcaggaTcatcagttgatttagctattttttctttacatttagcTggaatttcttcaattttaggagctattaattttattacaactattattaatataAAaTcCccaggaatatctttagatAaaatacctCtatttgtttgatctgttttaattacTgctattcttttattattatctttaccagttttagcaggagctattacCatattattaacagatcgaaatttaaatacatctttttttgaccctgcaggaggaggggacccaattttatatcaacatttattt

#### Description

**Male** (Fig. 153A). Wing length, 1.79; width, 0.74. **Head** (Figs. 153C–D). Vertex and frons dark brown, face and clypeus light brown. Eyes small, blackish, with long interommatidial setae. Antennal scape and pedicel whitish-yellow, flagellum light brown, except for lighter flagellomere 1 and distal four flagellomeres. Maxillary palpus whitish-yellow. Labella small, whitish-yellow. Head dorsoventrally compressed, scattered setulae over entire vertex, a row of five longer setae on occiput around eyes posteriorly to lateral ocellus, a row of setae along anterior margin of frons. Mid ocellus absent, lateral ocelli over a blackish background, touching eye margin. Eyes densely covered with long interommatidial setulae except along dorsal margin. Face covered with short darker setae, clypeus slightly bulging, short, densely covered with setulae. Antennal scape 1.5× pedicel length; flagellomere 1 barely longer than second flagellomere, covered with scattered light setae. Maxillary palpus with five palpomeres, palpomere 1 barely produced, palpomere 2 short, with small dorsal setae, palpomere 3 about twice as long as wide, sensorial pit conspicuous, opening at inner face on basal half, palpomere 4 slightly longer than palpomere 3, with dorsal and lateral setulae, palpomere 5 slender, over twice longer than palpomere 4. **Thorax** (Figs. 153E–F). Scutum compressed, dark brown, scutellum dark brown. Scutum entirely covered only with short light brown setae except for some supra-alar longer setae and three pairs of prescutellar bristles, second one at each side smaller. Small bulging area on scutum margin posteriorly to level of insertion of wing. Scutellum large, two pairs of bristles along posterior margin, some small scattered setae also along posterior margin, most of scutellum bare. Pleural sclerites dark brown. Pleural membrane ochre-yellow. Deep incision at scutum lateral margin above level of anterior spiracle. Basisternum extending dorsolateral arms to base of fore coxa, not fused to proepisternum, bare. Antepronotum triangular, fused to proepisternum at posterior margin, suture separating from proepisternum complete, covered only with small setae. Proepisternum largely developed, with small setae and three bristles along ventral margin. Anepisternum large, rectangular, entirely covered with brown setulae, a row of five long setae at posterior margin. Katepisternum strongly compressed, bare, anapleural suture restrict to posterior half. Mesepimeron and laterotergite strongly compressed, mesepimeron with two long setae and three small fine setae, laterotergite with two stronger setae and one small setae ventrally. Metepisternum long and slender, with a row of five fine setae; metepimeron not discernible. Mediotergite short, strongly curved, bare. Haltere pedicel whitish, knob brown, very few setulae on knob. **Legs**. Coxae strongly developed, especially mid and hind coxae. Fore coxa whitish with very light orangish tinge, mid and hind coxae whitish, all coxae with a light brown band basally; Fore femur concolor with coxa, mid and hind femora whitish on basal two-thirds, dark brown on distal third of wing; tibiae ochre-yellowish, mid and hind tibiae brown at basal end, yellowish-brown at tip; tarsi light greyish-brown. Fore coxa entirely covered with setulae at anterior face, a row of longer brown setae along margin on distal half of external face and at tip, and along margin on basal half of internal face; mid coxa nearly bare; hind coxa with some few small setae and one strong seta distally at external face. Femora covered with fine setae, a row of longer setae along ventral margin, hind femur with stronger setae. Tibiae and tarsi with regular rows of trichia. Fore tibia with a wide anteroapical depressed area lined with setulae. Mid and hind tibiae with a regular row of long, fine setae externally on apex. Fore tibia with one stronger dorsal seta at distal half; mid tibia with a row of six bristles dorsally, two bristles laterally on anterior face and one strong ventral bristle midway to apex besides distal strong setae; hind tibia with four dorsal bristles and seven stronger lateral bristles, besides distal strong setae. Fore leg tarsomeres only with rows of trichia and a couple of distal stronger setae; mid leg tarsomeres 1–3 with rows of stronger setae besides rows of trichia; hind leg tarsomeres 1–5 with stronger or longer setae, besides regular rows of trichia. Tarsomere 1 of fore leg 0.7× tibia length and 2.0× tarsomere 2 length. Tibial spurs light brown, subequal, less than 3× tibia width at apex. Tarsal claws with an inconspicuous basal tooth. **Wing** (Fig. 153G). Membrane light brown fumose, a small light brown band on wing base and a dark brown macula medially on wing from anterior margin to base of medial fork; membrane densely covered with regularly organized microtrichia on all cells, no macrotrichia on membrane. Membrane emarginated at tip of CuA. Sc faint, present as a fold towards C distally. R_1_ relatively short, reaching C at distal third of wing; R_4_ absent; R_5_ short, reaching C before level of tip of M_4_ running very close to C; C extending for ⅔ of distance to M_1_. First sector of Rs oblique, devoid of setae, 0.82× r-m length; r-m oblique, short. Posterior wing veins weakly sclerotized close to wing margin. M_1+2_ short, .4× r-m length; M_1_ and M_2_ well sclerotized, running more or less parallel along most of their length; bM over 7× r-m length; M_4_ short, arched, well sclerotized, origin beyond medial fork, base of M_4_ detached from CuA. CuA straight, long, reaching margin beyond level of tip of R_1_; first sector of CuA 1.3× length of second sector of CuA. Cubital pseudovein absent, CuP barely recognizable, reduced to basal fourth of length of CuA. Anal fold long, almost straight, not reaching wing margin. Dorsal macrotrichia on bR, R_1_, second sector of Rs, and r-m; ventral macrotrichia on distal fourth of bR, R_1_ and second sector of Rs. Wing margin emarginated at level of tip of CuA. **Abdomen**. Abdominal tergites 1–7 brown. Sternites 1-6 very slender, light brown. **Terminalia** (Figs. 154A– B). Light brown, cerci lighter. Gonocoxites large, close together medially, no medioventral process, no projection of gonocoxite distal border beyond base of gonostylus. Gonostylus composed of: (1) an elongate ventral lobe with strong setae and macrosetae on ventral end; (2) a small digitiform median lobe with an apical seta; (3) a dorsal, weakly sclerotized lobe with a dorsal branch with setae externally and a ventral branch, each branch with a megaseta apically. Gonocoxal bridge very weakly sclerotized. Aedeagal-parameral complex subquadrate, weakly sclerotized, with a pair of short, pointed extension dorsolaterally. Tergite 9 and cerci not clear.

**Female** (Fig. 153B). As male, except for the following. **Wing l**ength, 1.95; width, 0.75. **Head**. Head with four long setae on occiput around eyes posteriorly to lateral ocellus. **Thorax**. Laterotergite with four setae. Metepisternum with seven longer and four smaller setae along its length. **Terminalia** (Figs. 154C–D). Whitish-yellow. Sternite 8 elongate, no lobes posteriorly, setulae on posterior half, sternite reaching level of mid of cercomere 1. Sternite 9 elongate, Y-shaped, weakly sclerotized, anterior end slender, reaching proximal end of sternite 8. Tergite 8 wide anteriorly, tapering at posterior end, reaching level of base of cercomere 1. Tergite 9+10 elongate, wider midway to apex, weakly sclerotized. Cercomeres 1 and 2 produced, elongate, setose, subequal in length.

#### Material examined

**Holotype**: male, ZRCBDP0049297, National University of Singapore (Icube), 01.apr. 2015, MIP leg. (slide-mounted). **Paratypes**: 12 males, 5 females. **Males**: ZRCBDP0047053, National University of Singapore (PGP), 17.jun. 2015, MIP leg.; ZRCBDP0047056, National University of Singapore (PGP), 17.jun. 2015, MIP leg.; ZRCBDP0047102, National University of Singapore (PGP), 08.jul. 2015, MIP leg.; ZRCBDP0048133, Pulau Semakau (SMO2), old mangrove, 11.jul. 2013, MIP leg.; ZRCBDP0048728, Nee Soon (NS2), 15.apr. 2015, MIP leg.; ZRCBDP0049298, National University of Singapore (Icube), 01.apr. 2015, MIP leg.; ZRCBDP0049313, National University of Singapore (Icube), 08.apr. 2015, MIP leg.; ZRCBDP0049316, National University of Singapore (Icube), 08.apr. 2015, MIP leg.; ZRCBDP0066817, Bukit Timah, maturing secondary forest (BT09), 05.oct. 2016, MIP leg.; ZRCBDP0133553, Singapore, date range 2012-2018, MIP leg.; ZRCBDP0155002, Singapore, date range 2012-2018, MIP leg; ZRCBDP0155020, Nee Soon (NS2), 18.mar.15. **Females**: ZRCBDP0047092, National University of Singapore (PGP), 24.jun. 2015, MIP leg.; ZRCBDP0049314, National University of Singapore (Icube), 08.apr. 2015, MIP leg. (slide-mounted); ZRCBDP0049318, National University of Singapore (Icube), 27.may. 2015, MIP leg.; ZRCBDP0049325, National University of Singapore (Icube), 29.apr. 2015, MIP leg; ZRCBDP0049135, National University of Singapore (PGP), 22.apr.2015. **Additional sequenced specimens**: ZRCBDP0132881, National University of Singapore (PGP), 19.jul. 2017, MIP leg.; ZRCBDP0133420, National University of Singapore (PGP), 21.jun. 2017, MIP leg.; ZRCBDP0133513, National University of Singapore (PGP), 03.may. 2017, MIP leg.; ZRCBDP0133529, National University of Singapore (PGP), 03.may. 2017, MIP leg.; ZRCBDP0278325, Singapore, date range 2012-2108, MIP leg.; ZRCBDP0278457, Pulau Ubin (PU12), coastal forest, 17.may. 2018, MIP leg.; ZRCBDP0279137, Singapore, date range 2012-2108, MIP leg.; ZRCBDP0279177, Singapore, date range 2012-2108, MIP leg.; ZRCBDP0279178, Singapore, date range 2012-2108, MIP leg.; ZRCBDP0279182, Singapore, date range 2012-2108, MIP leg.; ZRCBDP0279186, Singapore, date range 2012-2108, MIP leg.; ZRCBDP0284177, Pulau Ubin (PU18), coastal forest, 2018, MIP leg.; ZRCBDP0284180, Singapore, freshwater swamp (KM03), date range, 2018-2019, MIP leg.; ZRCBDP0284182, Singapore, freshwater swamp (KM03), date range, 2018-2019, MIP leg.; ZRCBDP0284183, Singapore, freshwater swamp (KM03), date range, 2018-2019, MIP leg.; ZRCBDP0284186, Singapore, freshwater swamp (KM03), date range, 2018-2019, MIP leg.; ZRCBDP0284187, Singapore, freshwater swamp (KM03), date range, 2018-2019, MIP leg.; ZRCBDP0284191, Singapore, freshwater swamp (KM03), date range, 2018-2019, MIP leg.; ZRCBDP0284197, Singapore, freshwater swamp (KM03), date range, 2018-2019, MIP leg.; ZRCBDP0284199, Singapore, freshwater swamp (KM03), date range, 2018-2019, MIP leg.; ZRCBDP0284202, Singapore, freshwater swamp (KM03), date range, 2018-2019, MIP leg.; ZRCBDP0284255, Singapore, date range 2012-2108, MIP leg.; ZRCBDP0284268, Singapore, date range 2012-2108, MIP leg.; ZRCBDP0284284, Singapore, date range 2012-2108, MIP leg.; ZRCBDP0284300, Singapore, date range 2012-2108, MIP leg.; ZRCBDP0314075, Singapore, freshwater swamp (KM03), 04.apr. 2018, MIP leg.; ZRCBDP0314078, Singapore, freshwater swamp (KM03), 04.apr. 2018, MIP leg.; ZRCBDP0314090, Singapore, freshwater swamp (KM03), 04.apr. 2018, MIP leg.; ZRCBDP0314091, Singapore, freshwater swamp (KM03), 04.apr. 2018, MIP leg.; ZRCBDP0314096, Pulau Ubin (PU14), coastal forest, 05.apr. 2018, MIP leg.; ZRCBDP0314133, Pulau Ubin (PU13), mangrove, 05.apr. 2018, MIP leg.; ZRCBDP0314135, Pulau Ubin (PU13), mangrove, 05.apr. 2018, MIP leg.; ZRCBDP0314147, Singapore, freshwater swamp (KM03), 19.apr. 2018, MIP leg.; ZRCBDP0314149, Singapore, freshwater swamp (KM03), 19.apr. 2018, MIP leg.; ZRCBDP0314151, Singapore, freshwater swamp (KM03), 19.apr. 2018, MIP leg.; ZRCBDP0314153, Singapore, freshwater swamp (KM03), 19.apr. 2018, MIP leg.; ZRCBDP0314181, Singapore, freshwater swamp (KM03), 25.mar. 2018, MIP leg.; ZRCBDP0314184, Sungei Buloh (SB08), mangrove, 19.apr. 2018, MIP leg.; ZRCBDP0314195, Pulau Ubin (PU13), mangrove, 12.apr. 2018, MIP leg.; ZRC_MIS0000053, National University of Singapore (L06-MT), urban forest, 02.may. 2019, MIP leg; ZRCBDP0047084, National University of Singapore (Icube), 17.jun.15; ZRCBDP0048101, Sungei Buloh (SB01), 02.oct.13; ZRCBDP0049340, National University of Singapore (PGP), 08.apr.15; ZRCBDP0278330, Singapore, 03.may.18; ZRCBDP0279141, Singapore, 16.may.18; ZRCBDP0284228, Singapore; ZRCBDP0314081, Kranji Marshes (KM03), 04.apr.18; ZRCBDP0314132, Pulau Ubin (PU13), 05.apr.18; ZRCBDP0314155, Kranji Marshes (KM03), 19.apr.18; ZRCBDP0314162, Kranji Marshes (KM03), 19.apr.18; ZRCBDP0314164, Kranji Marshes (KM03), 19.apr.18.

**Etymology**. The species epithet of this species honors Charlotte Elizabeth Ferguson-Davie (1880–1943). Born in Essex and moving to Singapore in 1909 already as a physician, she founded St. Andrew’s Mission Hospital, the first women’s and children’s clinic in Singapore, and oversaw some of Singapore’s first programs to train female midwives and nurses. She was inducted into the Singapore Women’s Hall of Fame in 2014. The name is used in apposition.

**Remarks**. This is a pretty abundant species in Singapore, with nine different haplotypes, collected in different environments.

*Integricypta teosoonkimae* Amorim & Oliveira, sp. n. (Figs. 155A–D)

https://singapore.biodiversity.online/species/A-Arth-Hexa-Diptera-001482

urn:lsid:zoobank.org:act:66C8A0B6-2E65-4052-AB54-AA7722DCBFFE

**Diagnosis**. Head blackish-brown, flagellum dark brown, with lighter scape and pedicel. Scutum and pleural sclerites blackish-brown. Coxae and femora mostly whitish, mid and hind coxae with brown transverse band at proximal end, hind femur brown at distal fourth, proximal end of mid and hind tibia brown. Wing membrane with dark brown mark along over origin of Rs; C well produced beyond tip of R_5_; vein r-m present, M_1+2_ 0.88× r-m length; wing margin emarginated at level of tip of CuA. Abdominal tergites brown. Gonocoxites fused to each other along anterior half of syngonocoxite, no medioventral process; gonostylus with a flat lobe with short denticules covering inner face, a ventral digitiform lobe with some setae, and a falciform dorsal lobe.

***Integricypta_teosoonkimae_ZRCBDP0047054_hapZRC BDP0047054_SMH_holotype* [10: C, 14: T, 16: A, 24: G, 25: T, 55: A, 106: T, 122: A, 133: T, 217: C, 286: T]**

cctttcttcCacaTtAgctcatgGTggatcatctgttgatttagctattttttcAttacatttagcaggaatttcttcaattttaggagctattaattttattacTactattattaatataAaatctccaggTatatcttttgataaaatacctttatttgtttgatctgtattaattacagctattttattattattatctttaccagtattagcCggagctattactatattattaacagatcgaaatttaaatacttcattttttgatcctgcaggaggaggTgatccaattttataccaacatttattt

#### Description

**Male** (Fig. 155A). Wing length, 1.76; width, 0.67 mm. **Head**. Vertex and frons brown, face and clypeus light brown. Antennal scape and pedicel whitish-yellow [flagelli of both antennae missing in the holotype]. Maxillary palpus whitish-yellow. Labella small, whitish-yellow. A row of six long setae on occiput around eyes posteriorly to lateral ocellus. **Thorax** (Fig. 155B). Scutum brown, scutellum light brown. Mesepimeron with two long setae and three small fine setae, laterotergite with two long setae. Metepisternum with six long setae along sclerite and three setulae on anterior end. **Legs**. Coxae whitish, fore coxa with a slender light brown band basally, mid and hind coxae with larger brown band basally. **Wing** (Fig. 155C). Membrane light brown fumose, a small light brown band on wing base and a dark brown macula medially on wing from anterior margin to base of medial fork, more or less fading beyond tip of costal cell beyond R_1_. Sc faint, present as a fold towards C distally. R_1_ relatively short, reaching C at distal third of wing; R_4_ absent; R_5_ short, running close to C, reaching margin at level of tip of M_4_; C extending for over half of distance to M_1_. First sector of Rs oblique, devoid of setae; r-m oblique, short. M_1+2_ short, 1.5× r-m length; M_1_ and M_2_ well sclerotized, slightly diverging from each other close to wing margin. M_4_ origin beyond level of medial fork, base of M_4_ detached from CuA, reaching wing margin at level of tip of R_5_, arched along most of its length; bM over 9.1× r-m length. Posterior wing veins weakly sclerotized close to wing margin. CuA straight, long, reaching margin beyond level of tip of R_1_; first sector of CuA 1.2 length of second sector of CuA. Cubital pseudovein not produced, CuP weak, short, not reaching level of origin of M_4_. Dorsal macrotrichia on bR, R_1_, second sector of Rs, and r-m; ventral macrotrichia on distal fourth of bR, R_1_, and second sector of Rs. Wing margin emarginated at level of tip of CuA. **Abdomen**. Abdominal tergites 1–7 light brown. Sternites 1-7 yellowish-brown. **Terminalia** (Figs. 155D). Light yellowish-brown. Gonocoxites fused to each other along anterior half of syngonocoxite, no medioventral process. Gonostylus composed of: (1) a large, flat lobe with short denticules covering inner face; (2) a ventral, digitiform lobe with some few setae; and (3) a dorsal, falciform bare lobe. Tergite 9 well developed, placed at anterior end of terminalia dorsally.

#### Material examined

**Holotype**: male, ZRCBDP0047054, National University of Singapore (PGP), 17.jun.2015, MIP leg. (website photo specimen, slide-mounted).

**Etymology**. The species epithet of this species honors Teo Soon Kim (1904-1978). She was the third Malayan Chinese woman to be admitted to the bar of England and Wales (1927), the first woman admitted to the Straits Settlement bar (1928) after she returned to Singapore, and the first woman barrister in Hong Kong (1932). She was the first woman to argue a case in front of the Supreme Court and drew a crowd in the public gallery. She was inducted into the Singapore Women’s Hall of Fame in 2014.

*Integricypta shirinae* Amorim & Oliveira, sp. n. (Figs. 156A–H)

https://singapore.biodiversity.online/species/A-Arth-Hexa-Diptera-000725

urn:lsid:zoobank.org:act:030362EC-3000-409D-927A-408CACCAE650

**Diagnosis**. Head blackish-brown, flagellum dark brown with lighter scape and pedicel. Scutum, scutellum and pleural sclerites blackish-brown. Coxae and femora mostly whitish, mid and hind coxae with brown transverse band at proximal end; tip of mid femur and distal third of hind femur dark brown, proximal end of mid and hind tibiae brown. Wing membrane with dark brown mark along of origin of Rs; C well produced beyond tip of R_5_; r-m present, 0.77× M_1+2_ length; wing margin strongly emarginated at level of tip of CuA. Abdominal tergites 1–brown. Gonocoxites fused to each other along anterior half of syngonocoxite, no medioventral posterior process; gonostylus large, ventral lobe triangular, with a row of slender spines along inner margin, a digitiform median lobe and a large, digitiform dorsal lobe.

***Integricypta_shirinae_ZRCBDP0047077_hapZRCBDP0 040965_SMH_holotype* [14: T, 24: G, 37: A, 73: C, 88: T, 123: A, 256: C, 259: C]**

cctttcttctacaTtagctcatgGaggatcatctgtAgatttagcaattttttctttacatttagcaggaatCtcttcaattttaggTgctattaattttattacaactattattaatataaAatctccaggaatatcttttgataaaatacctttatttgtttgatctgttttaattacagctattttattattattatctttacctgtattagcaggagctattacaatattattaacagatcgaaatttaaaCacCtcattttttgaccccgccggaggaggagatccaattttatatcaacatttattt

#### Description

**Male** (Fig. 156A). Wing length, 1.73–1.76 mm; width, 0.67–0.74 mm. **Head**. Dark caramel-brown, a row of five dark brown longer setae dorsally to eyes on occiput. Scape and pedicel yellowish, scape twice pedicel length; flagellomeres greyish-brown, flagellomere 1 much lighter. Face and clypeus light brown. Maxillary palpus whitish; labella whitish-yellow. **Thorax** (Fig. 156C). Scutum caramel-brown, scutellum blackish-brown. Pleural sclerites caramel-brown, antepronotum, proepisternum, and mesepimeron darker, laterotergite ochre-yellow. Pleural membrane ochre-yellow. Two pairs of scutellar bristles. Antepronotum only with small setae, proepisternum with three bristles along ventral margin. Mesepimeron with two strong setae and two small setae, laterotergite with three strong setae. Metepisternum with a row of five long setae and five setulae close to anterior end. Haltere pedicel whitish, knob brownish. **Legs**. Coxae strongly developed, especially mid and hind coxae. Fore coxa whitish with very light orangish tinge, mid and hind coxae whitish, all coxae with a light brown band basally; fore femur concolor with coxa, mid and hind femora whitish on basal two-thirds, dark brown on distal fourth; tibiae ochre-yellowish, mid and hind tibiae brown at basal end, yellowish-brown at tip; tarsi light greyish-brown. **Wing** (Fig. 156D). Membrane light brown fumose, a dark brown macula from anterior margin to base of medial fork, extending distally along cell c, a faint brownish mark on membrane at very base of wing anteriorly. Sc faint, present as a fold. R_1_ relatively short, reaching C at distal third of wing; R_4_ absent; R_5_ short, reaching C before level of tip of M_4_; C extending for over half distance to M_1_. First sector of Rs oblique, devoid of setae; r-m oblique, short, slightly longer than first sector of Rs. M_1+2_ short, 2.0× r-m length; M_1_ and M_2_ well sclerotized, slightly diverging close to wing margin. M_4_ origin beyond level of base of medial fork, base of M_4_ detached from CuA, reaching wing margin beyond level of tip of R_5_,arched along most of its length; bM over 8.9× r-m length. Posterior wing veins weakly sclerotized close to wing margin. CuA straight, long, reaching margin almost at level of tip of R_5_; first sector of CuA 1.1× length of (projected) second sector of CuA. Cubital pseudovein not produced, CuP barely produced, not reaching level of origin of M_4_. Dorsal macrotrichia on bR, R_1_, second sector of Rs, and r-m; ventral macrotrichia on distal fourth of bR, R_1_, and second sector of Rs. Wing margin strongly emarginated at level of tip of CuA. **Abdomen**. Abdominal tergite 1 dark greyish-brown, tergites 1-6 light greyish-brown, tergite 6 cream-yellow along posterior margin, tergite 7 cream-yellow. Sternites 1-7 cream-yellow. **Terminalia** (Figs. 156E–F). Light brown, cerci lighter. Gonocoxites short, fused medially along anterior half of syngonocoxite, suture of fusion present only at anterior end, posterior border of syngonocoxite without a medioventral process, posterior border of gonocoxite with short rounded lobes laterally, anteriorly and posteriorly to insertion of gonostylus, few long setae close to posterior margins of syngonocoxite ventrally and some few elongate fine setae on posterior half laterally. Gonostylus composed of: (1) a triangular ventral lobe (more a ventral extension of distal lobe) with a long row of long, curved setae on posterior margin bearing a short, digitiform subterminal projection on anterior margin with two long distal setae; (2) a long digitiform, weakly sclerotized distal lobe densely covered with long fine setae; (3) a large dorsal lobe with a dense group of stronger setae distally and a strong seta directed inwards; a short digitiform median lobe with a single fine apical seta. Gonocoxal bridge very weakly sclerotized. Aedeagal-parameral complex present as a ventral elongate capsule extending into a distal aedeagal plate more sclerotized on distal margin, with a pair of sub-medial setae and a pair of laterodistal short pointed projections. Tergite 9 fused to tergite 10 and cerci, present as a pair of large ovoid lobes dorsally, bearing microtrichia and fine setae.

**Female** (Fig. 156B). As male, except for the following. **Wing l**ength, 1.70 mm; width, 0.67 mm. **Head**. Four long setae on occiput around eyes posteriorly to lateral ocellus. **Thorax**. Mesepimeron with five longer setae in a line and 11 small setae, laterotergite with three long setae and eight additional small setae. Metepisternum with three longer setae on posterior end and 14 setulae along its length. **Abdomen**. Tergites 1–5 brown, tergite 6 mostly brown, with a medial yellowish mark medially along posterior margin, tergite 7 mostly yellowish. **Terminalia** (Figs. 156G–H). Whitish-yellow. Sternite 8 elongate, trapezoid, weakly sclerotized, with no posterior lobes, covered with microtrichia and fine setae. Sternite 9 long and weakly sclerotized, anterior arm reaching level of mid of segment 7, posteriorly extending to level of tip of cerci. Tergite 9+10 wide, covered with microtrichia over entire sclerite and some setulae on posterior half. Cercomeres 1 and 2 partially fused, with setae, setulae and microtrichia, cercomere 2 with a long apical seta.

#### Material examined

**Holotype**: male, ZRCBDP0047077, National University of Singapore (PGP), 15.jul.15, MIP leg. (slide-mounted). **Paratypes** (14 males, 5 females). **Males**: ZRCBDP0047093, National University of Singapore (PGP), 24.jun.15, MIP leg.; ZRCBDP0047097, National University of Singapore (PGP), 08.jul.15, MIP leg. (MZUSP); ZRCBDP0048260, Sungei Buloh (SB1), mangrove, 09.oct.13, MIP leg.; ZRCBDP0048261, Sungei Buloh (SB1), mangrove, 09.oct.13, MIP leg.; ZRCBDP0048767, National University of Singapore (PGP), 20.may.15, MIP leg.; ZRCBDP0048780, National University of Singapore (PGP), 20.may.15, MIP leg.; ZRCBDP0048782, National University of Singapore (PGP), 20.may.15, MIP leg.; ZRCBDP0048787, National University of Singapore (PGP), 03.jun.15, MIP leg.; ZRCBDP0048788, National University of Singapore (PGP), 03.jun.15, MIP leg.; ZRCBDP0049069, National University of Singapore (PGP), 27.may.15, MIP leg.; ZRCBDP0049321, National University of Singapore (Uhall), 01.apr.15, MIP leg.; ZRCBDP0049346, National University of Singapore (PGP), 29.apr.15, MIP leg.; ZRCBDP0049347, National University of Singapore (PGP), 29.apr.15, MIP leg, ZRCBDP0133375, National University of Singapore (PGP), 14.apr.17. **Females**: ZRCBDP0040965, National University of (date range 2012-2018), MIP leg. ZRCBDP0047085, National University of Singapore (Icube), 17.jun.15, MIP leg.; ZRCBDP0049331, National University of Singapore (PGP), 10.jun.15, MIP leg.; ZRCBDP0049345, National University of Singapore (PGP), 29.apr.15, MIP leg. (slide-mounted), ZRCBDP0048427, Singapore, 25.apr.12. **Additional sequenced specimens**: ZRCBDP0078962, Singapore, date range 2012-2018; ZRCBDP0078972, Singapore, date range 2012-2018; ZRCBDP0132829, Singapore, date range 2012-2018; ZRCBDP0132829, Singapore, date range 2012-2018; ZRCBDP0132832, Singapore, date range 2012-2018; ZRCBDP0132833, Singapore, date range 2012-2018; ZRCBDP0132851, Singapore, date range 2012-2018; ZRCBDP0132855, Singapore, date range 2012-2018; ZRCBDP0132856, Singapore, date range 2012-2018; ZRCBDP0132878, Singapore, date range 2012-2018; ZRCBDP0132880, Singapore, date range 2012-2018; ZRCBDP0132894, Singapore, date range 2012-2018; ZRCBDP0133136, Singapore, date range 2012-2018; ZRCBDP0133161, Singapore, date range 2012-2018; ZRCBDP0133484, Singapore, date range 2012-2018; ZRCBDP0133512, Singapore, date range 2012-2018; ZRCBDP0133520, Singapore, date range 2012-2018; ZRCBDP0133521, Singapore, date range 2012-2018; ZRCBDP0133538, Singapore, date range 2012-2018; ZRCBDP0278282, Singapore, 17.may.18, MIP leg.; ZRCBDP0278291, Singapore, 17.may.18, MIP leg.; ZRCBDP0278294, Singapore, 17.may.18, MIP leg.; ZRCBDP0279136, Singapore, 16.may.18, MIP leg.; ZRCBDP0279140, Singapore, 16.may.18, MIP leg.; ZRCBDP0279145, Singapore, 16.may.18, MIP leg.; ZRCBDP0279146, Singapore, 16.may.18, MIP leg.; ZRCBDP0279152, Singapore, 30.may-05.jun.18, MIP leg.; ZRCBDP0279155, Singapore, 13.jun.18, MIP leg.; ZRCBDP0279161, Singapore, 26.apr.18, MIP leg.; ZRCBDP0279190, Singapore, 25.apr.18, MIP leg.; ZRCBDP0284189, Kranji Marshes (KM03), freshwater swamp, date range 2018-2019, MIP leg.; ZRCBDP0284193, Kranji Marshes (KM03), freshwater swamp, date range 2018-2019, MIP leg.; ZRCBDP0284201, Kranji Marshes (KM03), freshwater swamp, date range 2018-2019, MIP leg.; ZRCBDP0284242, Singapore, date range 2012-2018, ZRCBDP0284253, Singapore, date range 2012-2018, ZRCBDP0284258, Singapore, 03.may.18, MIP leg.; ZRCBDP0284260, Singapore, 03.may.18, MIP leg; ZRCBDP0047096, National University of Singapore (PGP), 08.jul.15; ZRCBDP0048768, National University of Singapore (PGP), 20.may.15; ZRCBDP0048783, National University of Singapore (PGP), 20.may.15; ZRCBDP0049268, National University of Singapore (PGP), 06.may.15; ZRCBDP0049333, National University of Singapore (PGP), 10.jun.15;. ZRCBDP0049334, National University of Singapore (PGP), 10.jun.15; ZRCBDP0049343, National University of Singapore (PGP), 08.apr.15; ZRCBDP0049344, National University of Singapore (PGP), 08.apr.15; ZRCBDP0133170, Singapore; ZRCBDP0133397, Singapore; ZRCBDP0278304. Singapore, 03.may.18; ZRCBDP0279144, Singapore, 16.may.18; ZRCBDP0279196, Singapore, 07.jun.18; ZRCBDP0284259, Singapore, 03.may.18; ZRCBDP0284261, Singapore, 03.may.18; ZRCBDP0284263, Singapore, 03.may.18. **Sequence failure specimens**: male, ZRCBDP0279115, Singapore, 31.may. 2018, MIP leg. (slide-mounted).

**Etymology**. The species epithet of this species honors Shirin Fozdar (1905-1992). Mumbai-born, she worked on women’s rights and welfare issues in India in the 1930s and 1940s, moving to Singapore in 1950, where she and her husband helped spreading the BaháDí. In Singapore, she championed against marriage inequality and polygamy, and was instrumental in the founding of the Singapore Council of Women and of the nation’s Syariah Court. Shirin was a leader in the advocacy effort that saw the Singapore Council of Women become law and played a major role in the creation of the Syariah Court.

**Remarks**. There are three haplotypes, *Integricypta shirinae* Amorim & Oliveira, **sp. n.** and all species delimitation approaches indicate a single species. Two sequence failure males have a terminalia identical to the holotype.

*Integricypta hoyuenhoeae* Amorim & Oliveira, sp. n. (Figs. 157A–E)

https://singapore.biodiversity.online/species/A-Arth-Hexa-Diptera-001545

urn:lsid:zoobank.org:act:5B515576-4D43-4947-B4FA-EDD34D7CCB5D

**Diagnosis**. Head blackish-brown, flagellum dark brown, scape and pedicel lighter. Scutum, scutellum, and pleural sclerites shining blackish-brown. Coxae and femora mostly whitish, mid and hind coxae with brown transverse band at proximal end; mid femur with no brown mark at basal end. Wing membrane light yellowish-brown fumose, with a light brown mark at wing base anteriorly, a dark brown mark anteriorly from C to M_2_ between origin of Rs and tip of R_1_; C produced beyond tip of R_5_ for over half of distance to tip of M_1_; r-m oblique, 0.79× of M_1+2_ length; wing margin strongly emarginated at level of tip of CuA. Abdominal tergites 1–5 brown, tergite 6 yellowish-brown. ***Integricypta_hoyuenhoeae_ZRCBDP0103977_holotype* [ 14: T, 24: G, 123: A, 125: T, 129: C, 133: G, 136: A, 146: A]**

cctttcttctacaTtagctcatgGagggtcatctgttgatttagcaattttttctctacatttagcaggaatttcttcaattttaggagctattaattttattacaactattattaatataaAaTctcCaggGatAtcttttgacAaaatacctttatttgtt tgatctgtattaattacagctattttattattattatctttaccagtattagctggagctattactatattattaacagatcgaaatttaaatacttcattttttgaccctgc tgggggaggagatccaattttatatcaacatctattt

#### Description

**Female** (Figs. 157A–C). Wing length, 1.98 mm; width, 0.75 mm. **Head**. Dark greyish-brown, a row of five dark brown longer setae dorsally to eyes on occiput. Scape and pedicel yellowish, flagellomeres light brown. Face and clypeus light brown. Maxillary palpus whitish. Labella whitish-yellow. **Thorax** (Fig. 157C). Scutum caramel-brown, scutellum light brown. Pleural sclerites blackish-brown. Pleural membrane ochre-yellow. Antepronotum only with small setae, proepisternum with three bristles along ventral margin. Mesepimeron with two longer setae and two smaller setae, laterotergite with two long setae. Metepisternum with a row of four long setae and three setulae close to anterior end. Haltere pedicel whitish, knob brownish. **Legs**. Coxae whitish with light orangish tinge, all coxae with a light brown band at basal end, less extensive in fore coxa. Fore femur concolor with coxa except for darker tip, mid and hind femora whitish with a greyish tinge; tibiae and tarsi light greyish-brown, at fore leg darker. **Wing** (Fig. 157D). Membrane light brown fumose, a dark brown macula from anterior margin to base of medial fork, extending distally along cell c, a faint brownish mark on membrane at very base of wing anteriorly. Sc faint, a fold towards C. R_1_ relatively short, reaching C at distal third of wing; R_4_ absent; R_5_ short, reaching C slightly before level of tip of M_4_; C extending for over half of distance to M_1_. First sector of Rs oblique, devoid of setae; r-m oblique, short, 0.82× first sector of Rs. M_1+2_ short, 1.7× r-m length; M_1_ and M_2_ well sclerotized, M_1_ quite parallel to M_2_ close to wing margin. M_4_ origin beyond level of base of medial fork, base of M_4_ detached from CuA, reaching wing margin before level of tip of R_5_,gently curved along distal half; bM over 9.7× r-m length. Posterior wing veins weakly sclerotized close to wing margin. CuA straight, long, reaching margin beyond level of tip of R_1_; first sector of CuA 1.2× length of (projected) second sector of CuA. Cubital pseudovein absent, CuP restrict to first third of CuA. Dorsal macrotrichia on bR, R_1_, second sector of Rs, and r-m; ventral macrotrichia on distal fourth of bR, R_1_, and second sector of Rs. **Abdomen**. Abdominal tergite 1 dark greyish-brown, tergites 2-5 dark greyish-brown with margins grey-yellowish, tergite 6 darker only medially, large lighter portions laterally, tergite 7 cream-yellow. Sternites 1-7 cream-yellow. **Terminalia** (Fig. 157E). Light brown, cerci lighter. Sternite 8 elongate, trapezoid, weakly sclerotized, with no lobes on posterior margin, covered with microtrichia and fine setae. Sternite 9 weakly sclerotized, very long, anterior end reaching level of mid of segment 7, extending posteriorly to level of tip of cerci. Tergite 9+10 wide, covered with microtrichia, two pairs of setae along posterior margin. Cercomeres 1 and 2 partially fused, with setae, setulae and microtrichia, cercomere 2 with a long apical seta.

#### Material examined

**Holotype**: female, ZRCBDP103977, Sungei Buloh (SB1), 06.nov. 2013, MIP leg. (website photo specimen, slide-mounted).

**Etymology**. The species epithet honors Abbess Ho Yuen Hoe (1908-2006), who had as Dharma name Venerable Jing Run. Born in Guangzhou, she was a Buddhist nun who was affectionately known as Singapore’s “grand dame of charity” for her lifelong devotion in helping the old and needy. She was the founder and abbess of the Lin Chee Cheng Sia Temple and founded in 1969 the Man Fu Tong Nursing Home, the first Buddhist nursing home. She received the Public Service Award from the President of Singapore in 2001 in recognition of her contribution to the country. She was inducted into the Singapore Women’s Hall of Fame in 2014.

**Remarks**. The holotype of *Integricypta hoyuenhoeae* Amorim & Oliveira, **sp. n.** is a female and can be distinguished from the remaining four species of *Integricypta* Amorim & Oliveira, **gen. n.** based on the color of the abdomen and details of the wing venation: the strongly emarginated wing margins, M_4_ not arched along the distal half, M_1_ and M_2_ not divergent at distal end etc.

## Discussion

### Singapore Mycetophilidae are a dark taxon

This study of the fauna of Mycetophilidae of Singapore indicates the size of the taxonomic impediment for the family, which is clearly a dark taxon based on the large number of new species. After over 250 years of biodiversity studies, it is still possible to find 115 new species within a relatively small set of specimens from one small area of study. There is no reason to believe that this is due to an unusually large number of species of the family in Singapore. To the contrary, Singapore has lost most of its natural habitats and the original species diversity would undoubtedly be much higher (Brook et al. 2003). For example, Singapore only retains half of its original documented butterfly fauna (Theng et al. 2020). We therefore predict that the fauna of countries with more undisturbed forests will yield many more new species than what has been found in Singapore. It is also unlikely that this applies only to Southeast Asia. We predict that many tropical forests worldwide will have a similar number of new species of the family, as can be seen from the diversity of mycetophilids in tropical forests elsewhere (e.g., Brown et al. 2018; Amorim et al. 2022).

### Revising integrative taxonomy protocols

The kind of results obtained with this study raises the central question of how the species diversity of dark taxa should be tackled. We argue for a shift towards more faunistic treatments based on freshly collected specimens that can be readily processed and described based on multiple sources of data. This process will become even more efficient as specimen handling is automated (e.g., Wührl et al., 2022). We would also like to mention that this study exemplifies why a minimalist approach, based on barcode-only data is damaging to taxonomy. Barcode data can be analyzed with various algorithms and parameters: the haplotype networks demonstrate that there is a significant number of cases where the same barcode data yield different number of MOTUs. Without additional evidence, it becomes impossible to choose among conflicting hypotheses. Indeed, there are a good number of cases where the morphological evidence is only congruent with some MOTU delimitations. In other words, the use of morphology greatly helps with clearing up the confusion caused by ambiguous signal in the barcode data. Using only barcodes for species delimitation method would lead likely to mistakes that would damage the taxonomy of the family. Overall, we would argue that having 15–20% of the species with delimitation issues is already a compelling reason to retain morphology in species delimitation and descriptions.

A minimalist approach only using DNA barcodes would have generated even more problems because it impedes with clarifying the meaning of already-available names and interferes with the identification of older material that is not suitable for large-scale barcoding. Thirdly, it renders the species unavailable to those labs that do not have access to barcoding facilities (see also Zamani et al. 2021, 2022; Engel et al. 2021). This is a particularly serious problem, because DNA barcode data are essentially unavailable in many tropical countries (see Figure 3 in Meier et al., 2024) Lastly, barcode-alone approaches ignore that *COI* evolution is largely clockwise and unlikely to capture particularly slow or fast speciation events.

Barcode information, however, was essential in this study for effectively guiding and simplifying the morphological work. Barcodes highlighted, for example, which haplotypes should have specimens checked to confirm species limits. In some cases, the final decisions were made only after dissecting male terminalia for specimens of all haplotypes. This is direct evidence of the power of a truly integrative approach for taxonomy. We also argue that morphology-alone approaches are undesirable because it is difficult to scale them sufficiently for dark taxa and it struggles with resolving “grey-zone” cases. Morphology-only approaches also struggle with recognizing species that are only available from females and thus considerably underestimate diversity.

Lastly, we would like to emphasize the need to track all specimens with unique identifiers. All specimens are important, and revisions of dark taxa will have thousands of them. It is imperative that each specimen has a unique code that is used to track information at all steps of the analysis—for example, in large spreadsheets including label data, details of the sequencing process, extraction, identification, slide-mounting, imaging etc. It is similarly important that an efficient archival system for all specimens is established.

Indeed, in many biodiversity treatments, not all specimens are given equal attention. Often molecular studies do not even keep vouchers. The different stages of mass species discovery like this—sequencing, imaging, slide-mounting, pinning, and storying—are frequently done in different spaces or labs. Tracking specimens is easy in small studies, but it is a challenge when dealing with thousands of vials each with a single specimen representing hundreds of haplotypes and species. Careful and strict procedures are needed not only for tracking specimens, but also to match illustrations.

Work on this project started several years ago, what explains why the descriptions are still fairly detailed. In future large-scale revisions, we will shorten the descriptions further and the diagnoses will be even more focused. However, we will not cut down on imaging, which will remain indispensable, while descriptions can be reduced to the habitus and diagnostic features (often male terminalia structures with more detailed illustrations). As mentioned above, more comprehensive descriptions would still be useful in some contexts (e.g., production of identification keys for genera). Similarly, drawings will also remain important for complex male terminalia, but overall the length of species descriptions can be reduced without major loss to the taxonomy of the group.

### Singapore mycetophilids and habitat diversity

The sampling effort in this project was uneven across the different habitats, mangroves having had more sampling weeks than the remaining habitats. Despite limited sampling, the swamp forest was surprisingly the habitat with the highest number of mycetophilid species (89 species in the first batch). Rainforests were next, with 45 species, while the mangroves only have 26 species. The smallest number of species was found in the freshwater swamp and a coastal forest (both had 9 species based on very limited sampling).

The diversity of mycetophilids in the mangrove samples was surprisingly high given that this environment has never been associated with mycetophilid diversity. This is in line with the recent finding that mangroves have a surprisingly large and distinct insect fauna (Yeo et al. 2021). Eight species (one species of *Manota*, two species of *Platyprosthiogyne*, three species of *Epicypta*, one species of *Aspidionia* and one species of *Integricypta* Amorim & Oliveira, **gen.n.**) were recorded only in mangrove samples. This is an interesting example of how massive biodiversity biomonitoring can unveil relevant and unexpected information of microhabitat faunal composition and species-richness, guiding conservation priorities.

### Mycetophilidae species-richness in tropical environments

The most species-rich genera in our samples are the mycomyine genus *Neoempheria*, with 31 species, the mycetophiline genus *Epicypta*, with 29 species, and the leiine genus *Manota*, with 14 species.

Many of the genera collected in Singapore have a worldwide distribution. In the Mycetophilidae, some genera are species-rich in temperate zones, but have small clades that extend into tropical areas. Examples from Singapore are *Brachycampta*, *Exechia*, *Mycetophila*, and maybe *Ectrepesthoneura*. However, genera that are diverse in tropical areas of the world (occasionally with species in temperate areas) correspond to the bulk of the species diversity and the abundance of the family in Singapore. Examples are the species of *Tetragoneura*, *Allactoneura*, *Eumanota*, *Manota*, *Mohelia*, *Neoempheria*, *Parempheriella*, *Clastobasis*, *Metanepsia*, *Chalastonepsia*, *Platyprosthiogyne*, *Platurocypta*, *Epicypta*, *Aspidionia*, and *Integricypta*. They comprise 106 of the 120 species found in our samples. In this context, the number of species of *Clastobasis* in Singapore—only three—was surprisingly low. The most abundant species found in our samples is *Clastobasis oranglaut* Amorim & Oliveira, **sp. n.,** with 357 collected specimens, but the species of *Epicypta* overall are responsible for almost half of the mycetophilid specimens collected in Singapore.

This pattern of generic composition and species richness across mycetophilids is also seen in tropical areas of other parts of the globe. In samples of the Amazon forest, including the canopy, the most species-rich genera were *Epicypta*, with 29 species, *Neoempheria*, with 21 species, *Cluzobra*, with eight species, and *Dziedzickia* and *Leia*, with five species each (Amorim et al. 2022). A cloud forest in Costa Rica, with an overlap of Nearctic and Neotropical elements, had 136 species of *Mycetophila*, 23 species of *Epicypta*, 18 species of *Zygomyia*, 16 species of *Mycomya*, nine species of *Sciophila*, eight species of *Exechia*, and seven species of *Neoempheria (*Brown et al. 2018).

Some elements of the Singaporean fauna are clearly part of a more restricted distribution pattern—Afro-Oriental or Afro-Oriental-Australasian. This is the case of *Allactoneura*, *Eumanota*and the (*Aspidionia*+*Integricypta*) clade. *Platurocypta* and *Platyprosthiogyne* are predominantly Afro-Oriental, but also have some few Palearctic and Neotropical species (one undescribed Neotropical species of *Platyprosthiogyne* was mentioned in Brown et al. 2018). The clade including the genera *Chalastonepsia* and *Metanepsia* indeed has Neotropical representatives, but they are still nominally kept within *Dziedzickia*.

It is worth mentioning that the great similarity of the Singaporean fauna of Mycetophilidae to that of South American tropical forests suggests common evolutionary origins between the transtropical disjunct elements in the Oriental and Neotropical regions (many of which also in Africa). This is interesting because most mycetophilid clades originated more recently than the beginning of the separation between the Gondwanam continents (Oliveira & Amorim 2021). A better explanation than a Gondwanan origins for this Neotropical-Oriental pattern seems to be a connection through the paleotropical environments in Laurasian terranes in the first half of the Cenozoic (Amorim et al. 2018). These elements of a tropical fauna seem to have expanded southwards in South America, Africa and Australasia, followed by an extensive extinction in Europe and North America. The resulting pattern is the typical pantropical distribution (i.e., Oriental + Afrotropical + Australasian + Neotropical) with a post-Gondwanan origin. This is a case of pseudo-congruence and the pattern can be referred to as pseudo-Gondwanan (Amorim et al. 2018).

As we further address entire faunas of different regions, we will be able to understand the biogeographical evolution of the Mycetophilidae in more detail. With the study of the phylogeny, evolution and associated fossils of Leiinae (Oliveira & Amorim 2021), for example, it was possible not only to track the origin of the subfamily to the southern part of Gondwana, but also to spot clades of the subfamily that extended their distribution northwards, secondarily occupying other parts of the world. Understanding the patterns of mycetophilid faunal composition in different parts of the world, with studies of complete faunas, will provide helpful information not only on the evolution of fungus-gnats, but also of the evolution of forests and of the fungi associated to these forests along the last 150 million years.

## Acknowledgements

All work described here was carried out as part of a comprehensive insect survey of Singapore developed in collaboration and with support from the National Parks Board of Singapore (NParks). Special thanks also go to Dr. Patrick Grootaert, Foo Maosheng, Jayanthi Puniamoorthy, and the team from the National Biodiversity Centre of NParks for their assistance in fieldwork (Permits: NP/RP12-022-4, NP/RP12-022-5, NP/RP12-022-6). In addition, we would like to also thank many research staff, lab technicians, undergraduate students, and interns of the Evolutionary Biology Laboratory at the National University of Singapore for their help and assistance with molecular work. This project would have been impossible without their hard work and support by Peter K. L. Ng. Financial support was provided by Tier 2 funding from the Ministry of Education (Singapore).

## Legend for Plates

**Figs. 1A-B.**
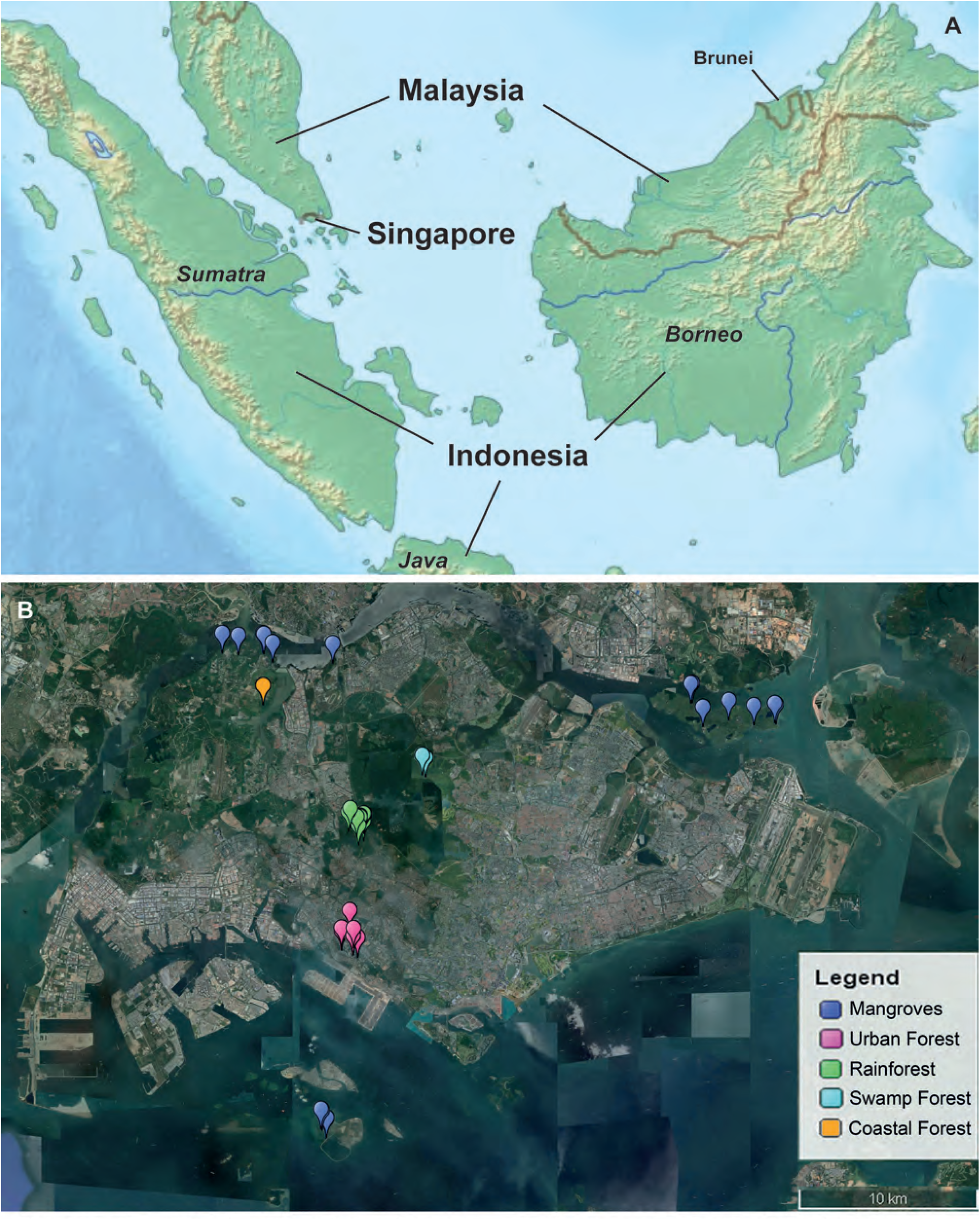
Geographic distribution of traps of the Singapore Mangrove Insect Project (MIP) (modified from www.freeworldmaps.net). **A.** Southeast Asia, with relative position of Singapore. **B.** Collecting sites in different Singapore natural environments.

**Figs. 2A-E.**
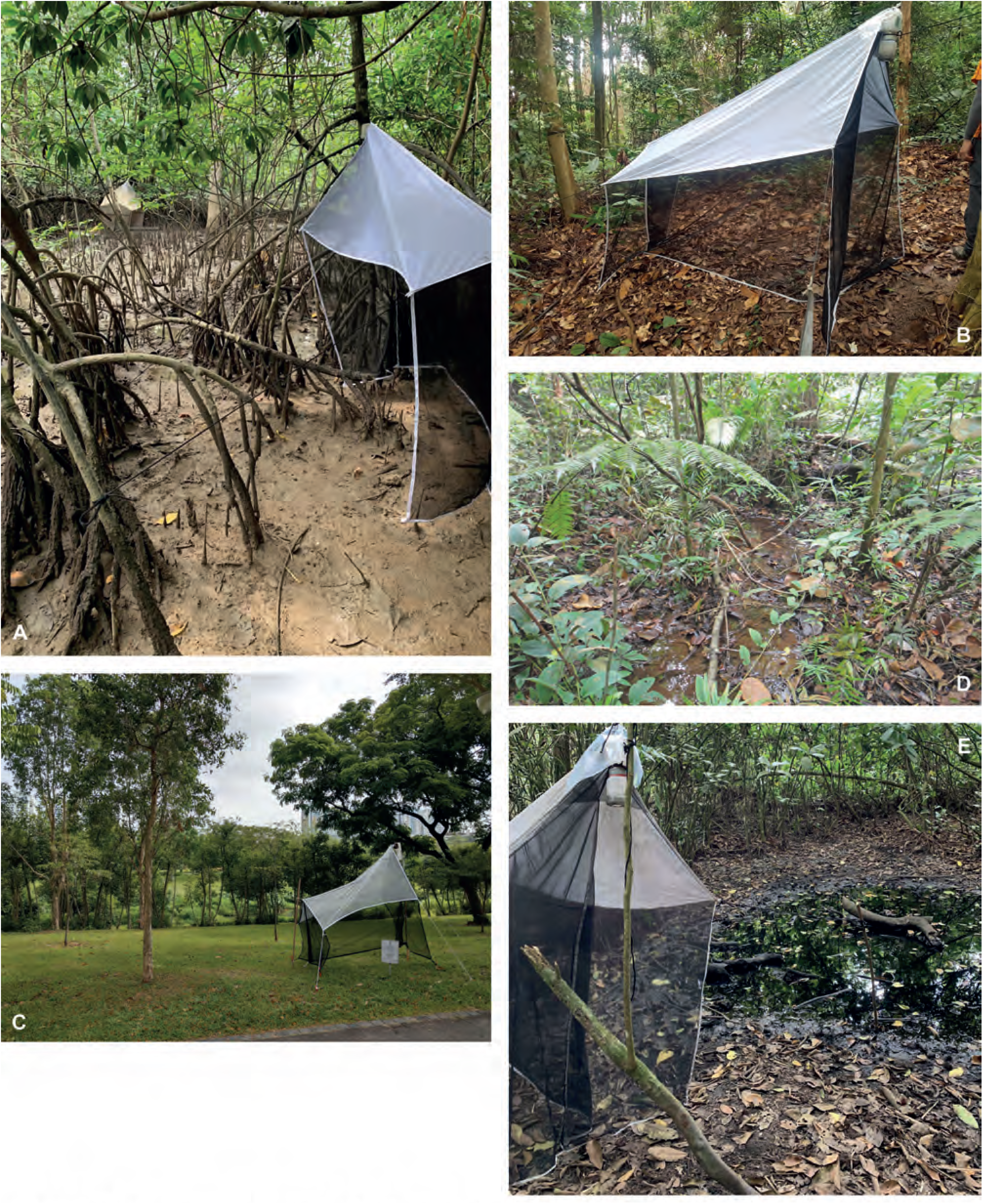
Environments sampled in the Singapore Mangrove Insect Project. **A.** Mangrove. **B.** Rainforest. **C.** Urban forest. **D.** Swamp forest. **E.** Freshwater swamp.

**Figs. 3A-G.**
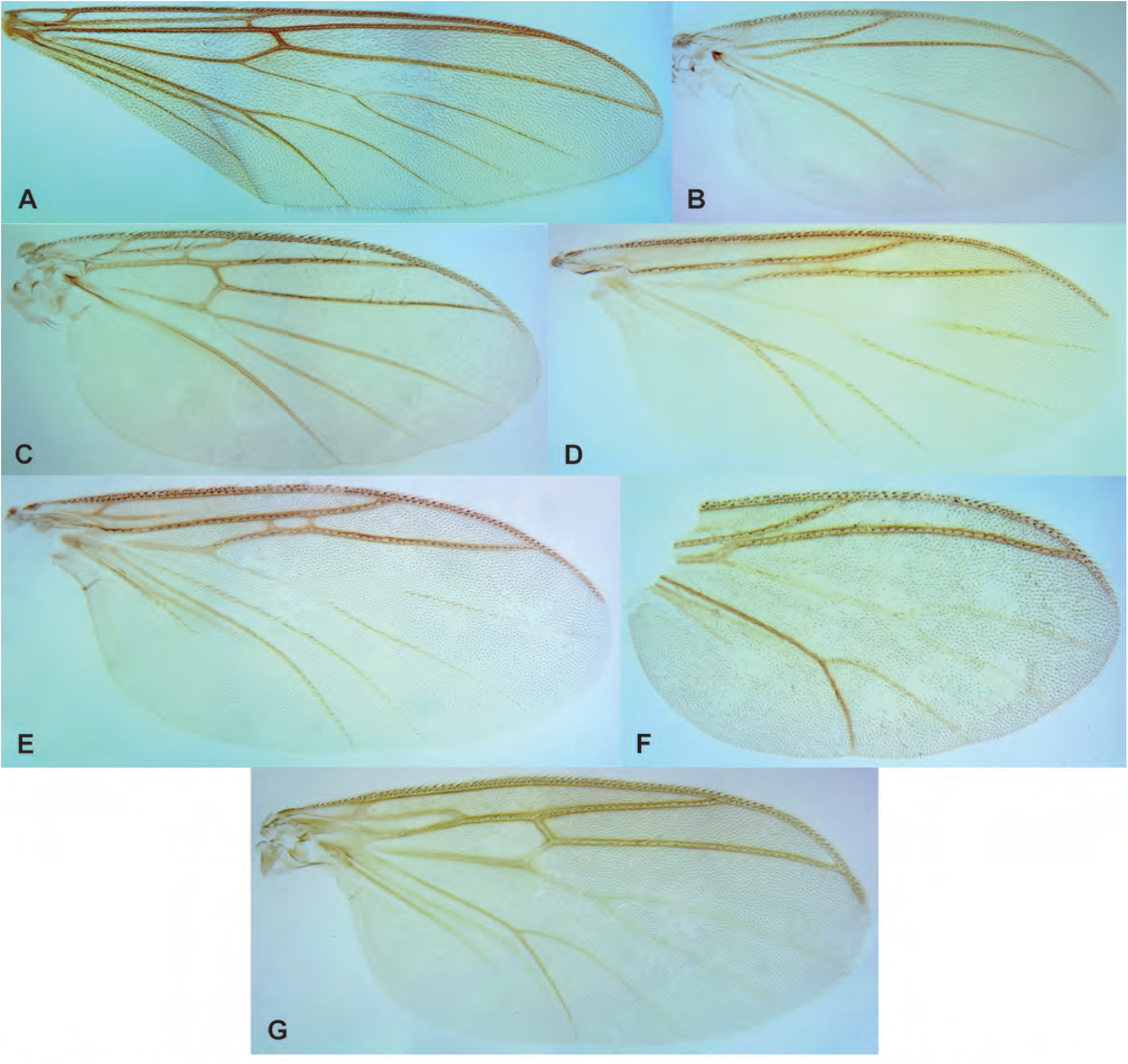
Wings of Mycetophilidae genera in the Singapore Mangrove Insect Project. Sciophilinae: **A.** *Leptomorphus rafflesi* Amorim & Oliveira, **sp. n. B.** *Azana leekongchiani* Amorim & Oliveira, **sp. n. C.** *Monoclona simhapura* Amorim & Oliveira, **sp. n.** Tetragoneurinae: **D.** *Tetragoneura crawfurdi* Amorim & Oliveira, **sp. n. E.** *Ectrepesthoneura johor* Amorim & Oliveira, **sp. n.** Gnoristinae: **F.** *Metanepsia malaysiana* Kallweit. **G.** *Chalastonepsia* sp.

**Figs. 4A-J.**
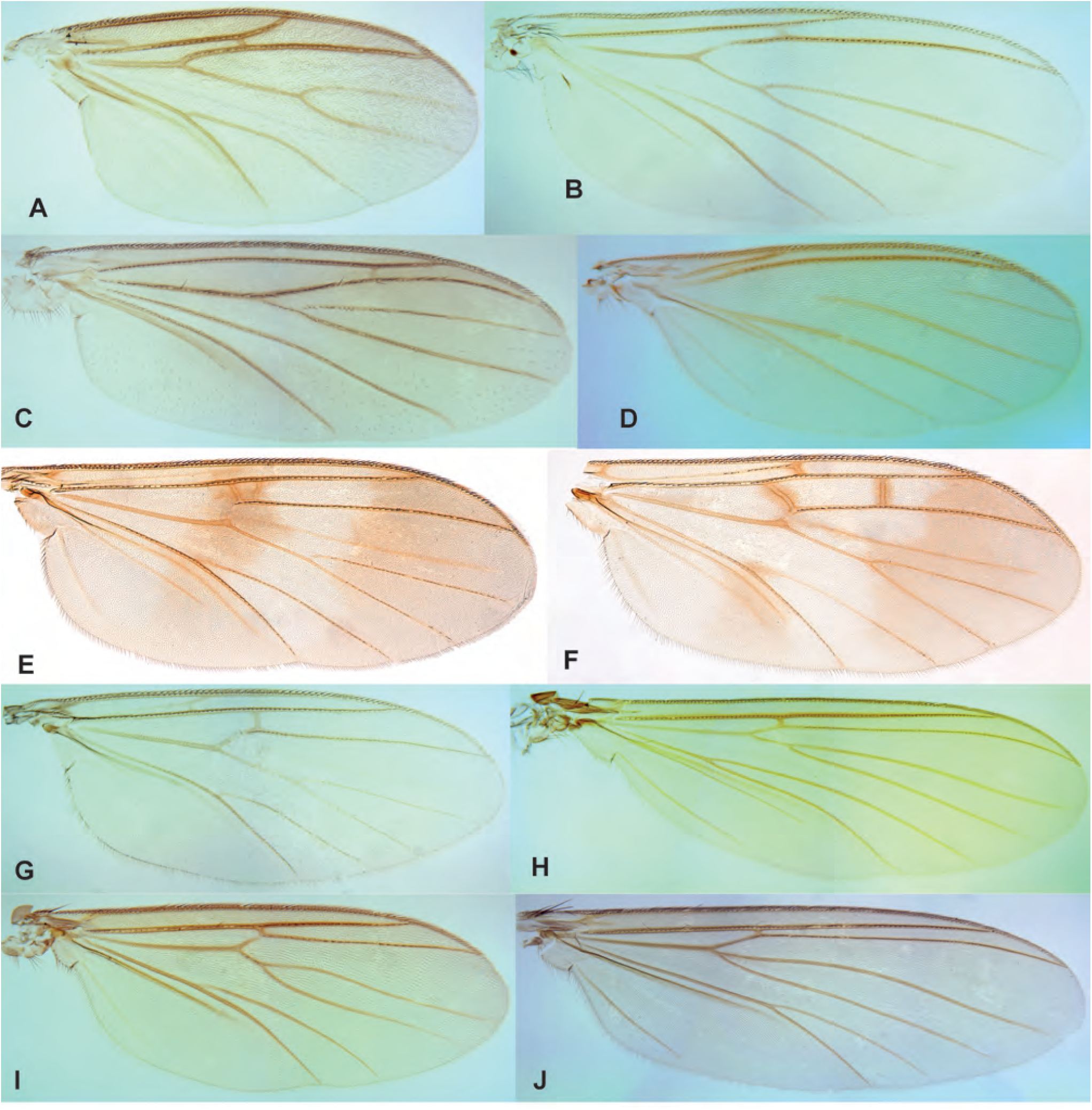
Wings of Mycetophilidae genera in the Singapore Mangrove Insect Project. Leiinae: **A.** *Mohelia zubirsaidi* Amorim & Oliveira, **sp. n. B.** *Clastobasis oranglaut* Amorim & Oliveira, **sp. n. C.** *Eumanota rakola* Søli. **D.** *Manota temenggong* Amorim & Oliveira, **sp. n.** Mycomyinae. **E.** *Neoempheria dizonalis* Edwards. **F.** *Neoempheria* sp. **G.** *Parempheriella defectiva* Edwards. Mycetophilinae Exechiini: **H.** *Allodia teopohlengi* Amorim & Oliveira, **sp. n. I.** *Brachycampta glorialimae* Amorim & Oliveira, **sp. n. J.** *Exechia alinewongae* Amorim & Oliveira, **sp. n**.

**Figs. 5A-H.**
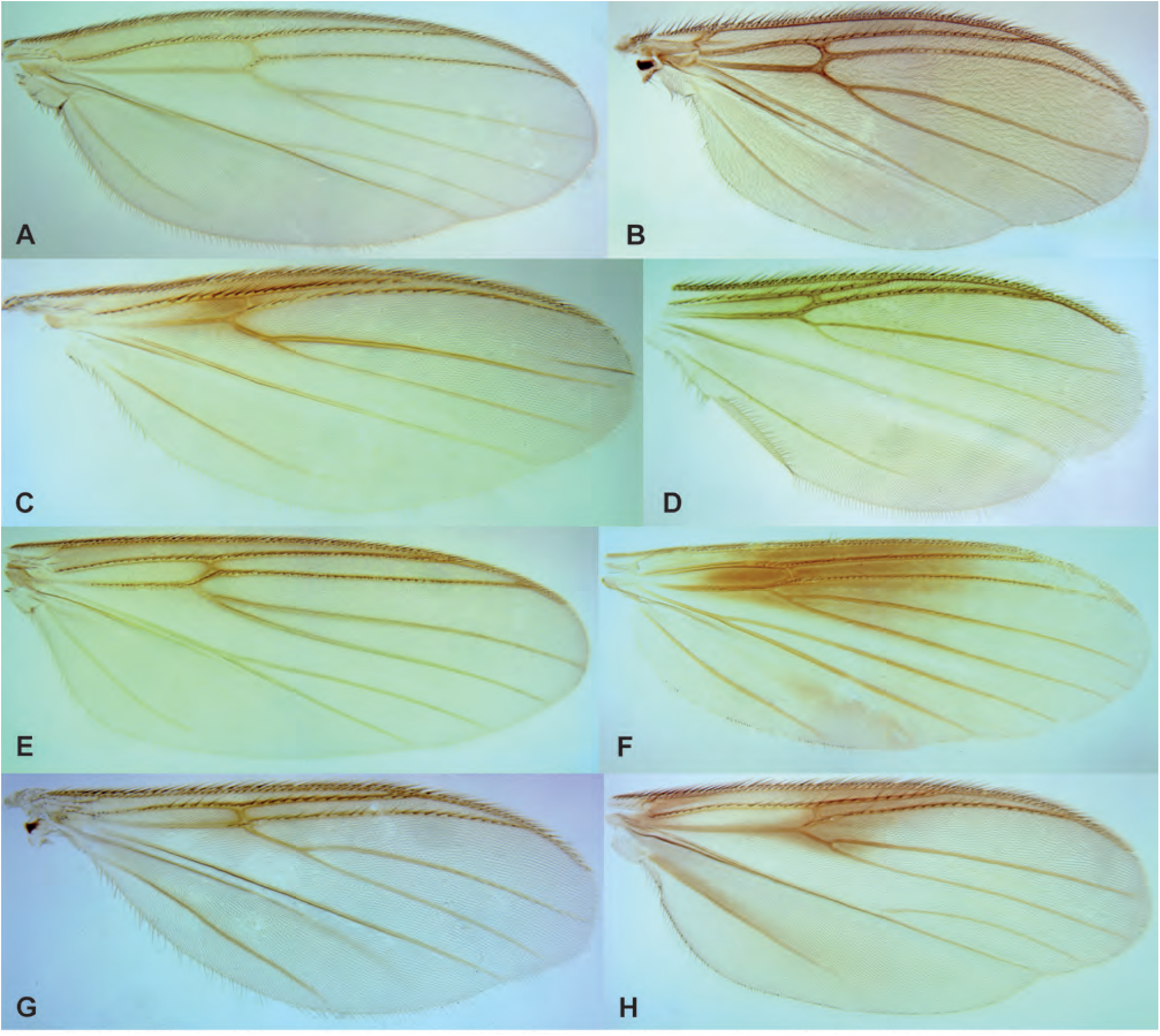
Wings of Mycetophilidae genera in the Singapore Mangrove Insect Project. Mycetophilinae Mycetophilini: **A.** *Mycetophila chngseoktinae* Amorim & Oliveira, **sp. n. B.** *Platyprosthiogyne snehalethaae* Amorim & Oliveira, **sp. n. C.** *Platyprosthiogyne phanwaithongae* Amorim & Oliveira, **sp. n. D.** *Platyprosthiogyne gohsookhimae* Amorim & Oliveira, **sp. n. E.** *Platurocypta adeleneweeae* Amorim & Oliveira, **sp. n. F.** *Epicypta limchiumeiae* Amorim & Oliveira, **sp. n. G.** *Aspidionia janetjesudasonae* Amorim & Oliveira, **sp. n. H.** *Integricypta shirinae* Amorim & Oliveira, **gen. n. sp. n. Sciophilinae**

## Sciophilinae

**Figs. 6A-D.**
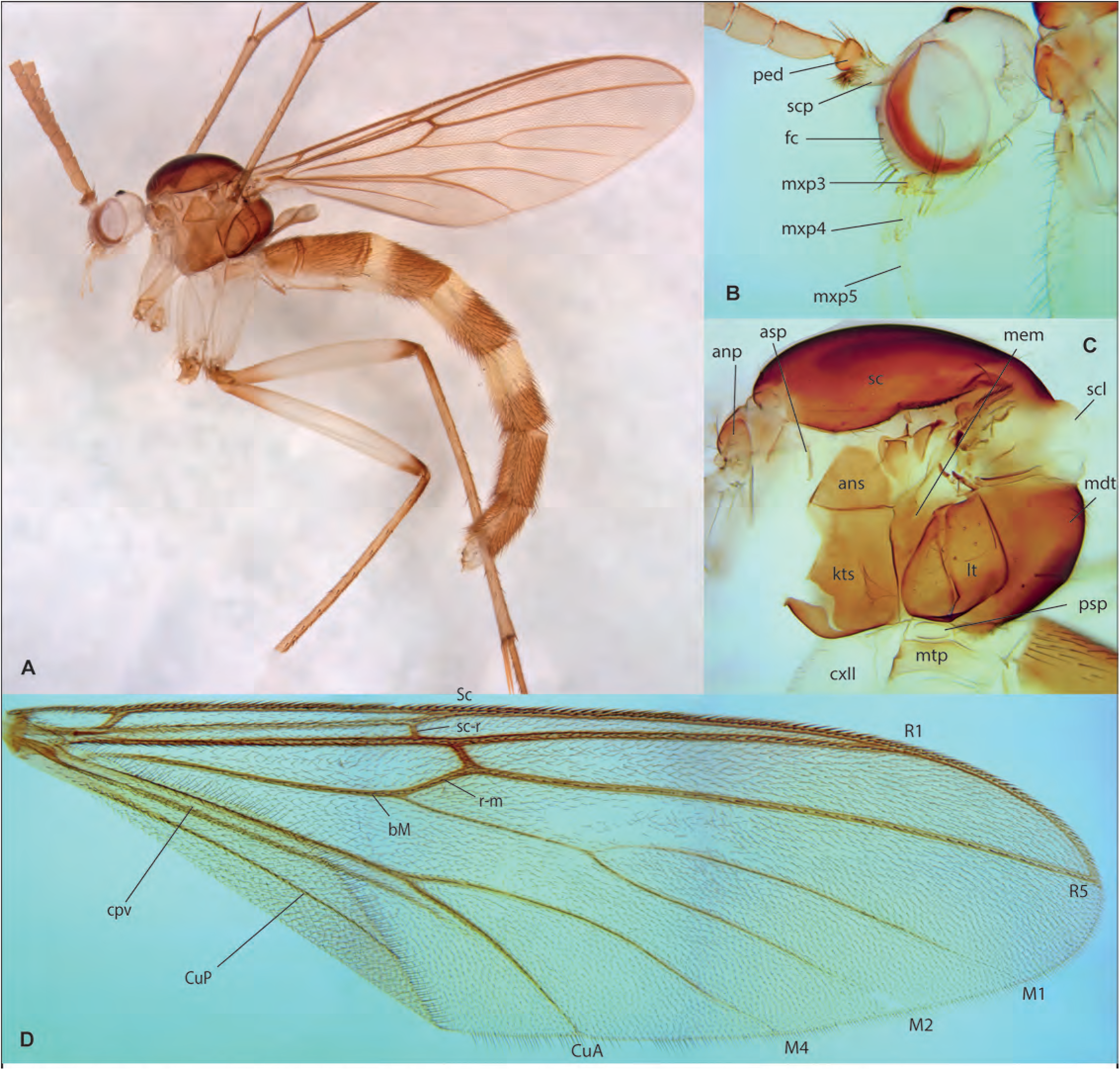
*Leptomorphus rafflesi* Amorim & Oliveira, **sp. n.**, male holotype, ZRCBDP0066807. **A.** Habitus, lateral view. **B.** Head, lateral view. **C.** Thorax, lateral view. **D.** Wing.

**Figs. 7A-D.**
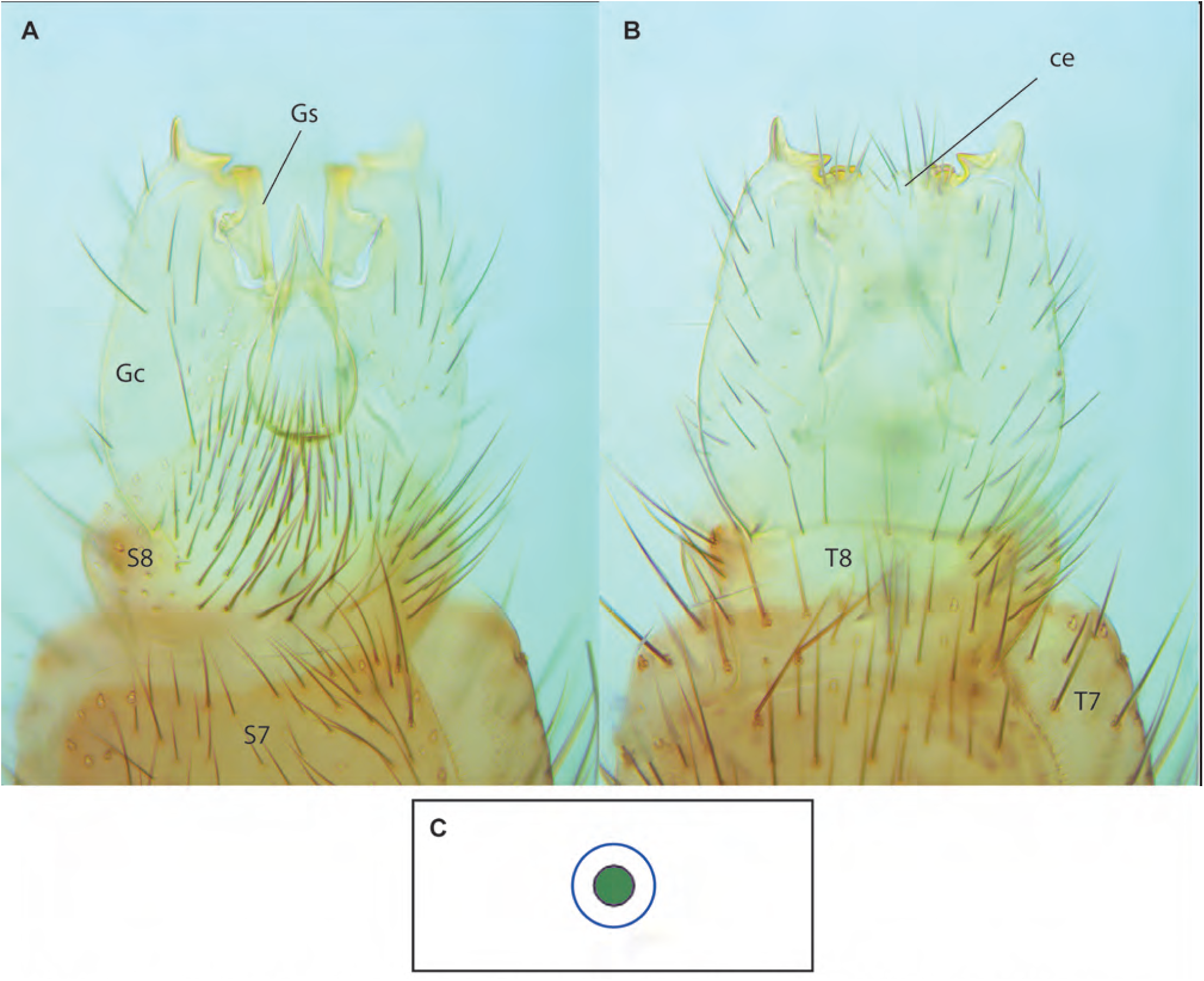
*Leptomorphus rafflesi* Amorim & Oliveira, sp. n., male holotype, ZRCBDP0066807. A. Terminalia, ventral view. B. Terminalia, dorsal view. C. Haplotype network for *Leptomorphus*.

**Figs. 8A-F.**
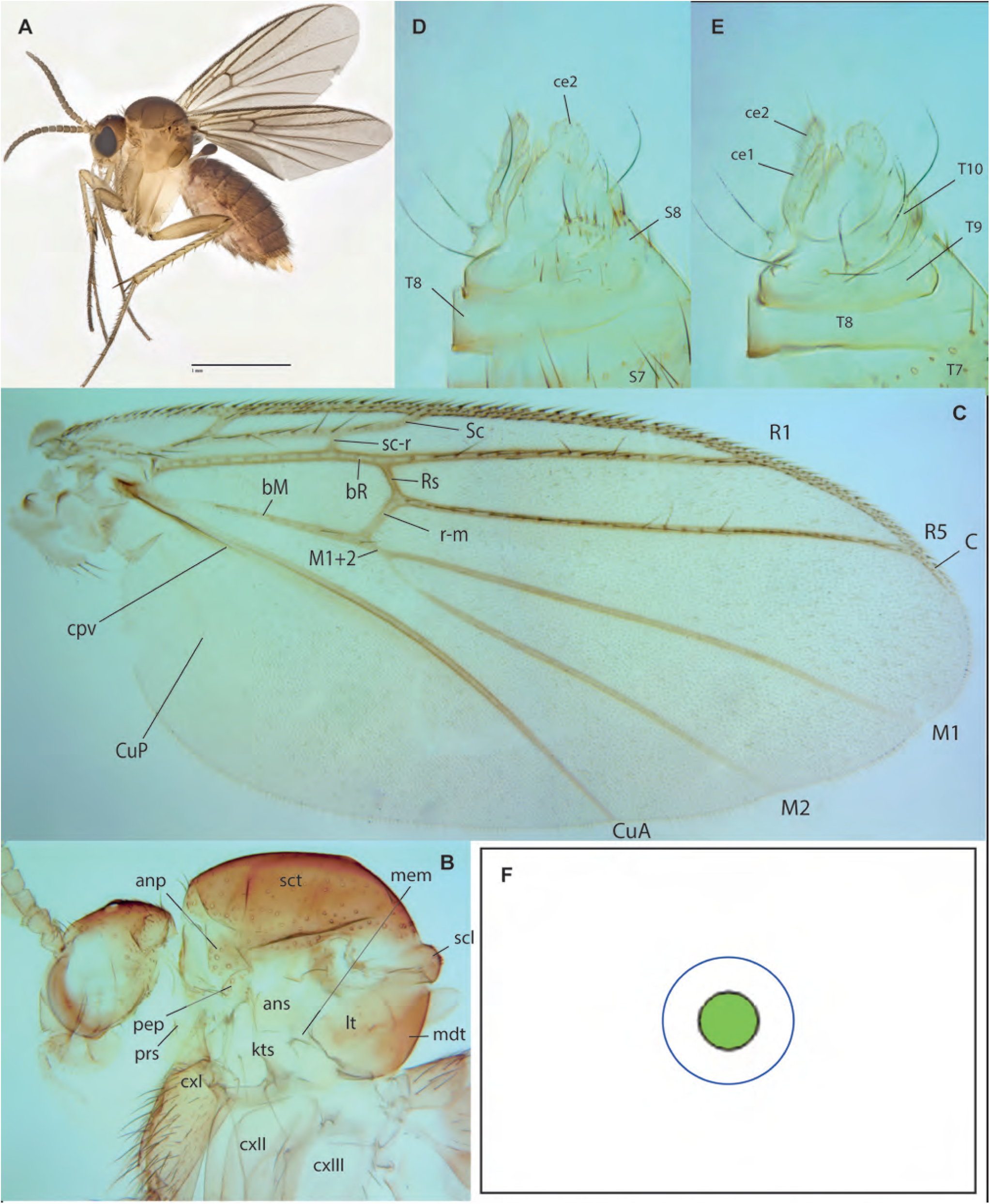
*Monoclona simhapura* Amorim & Oliveira, sp. n. A. Habitus, lateral view, female paratype, ZRCBDP_0048568. B. Head and thorax, lateral view, female holotype, ZRCBDP0048567. C. Wing, same. D. Female terminalia, ventral view, same. E. Female terminalia, dorsal view, same. F. Haplotype network for *Monoclona*.

**Figs. 9A-I.**
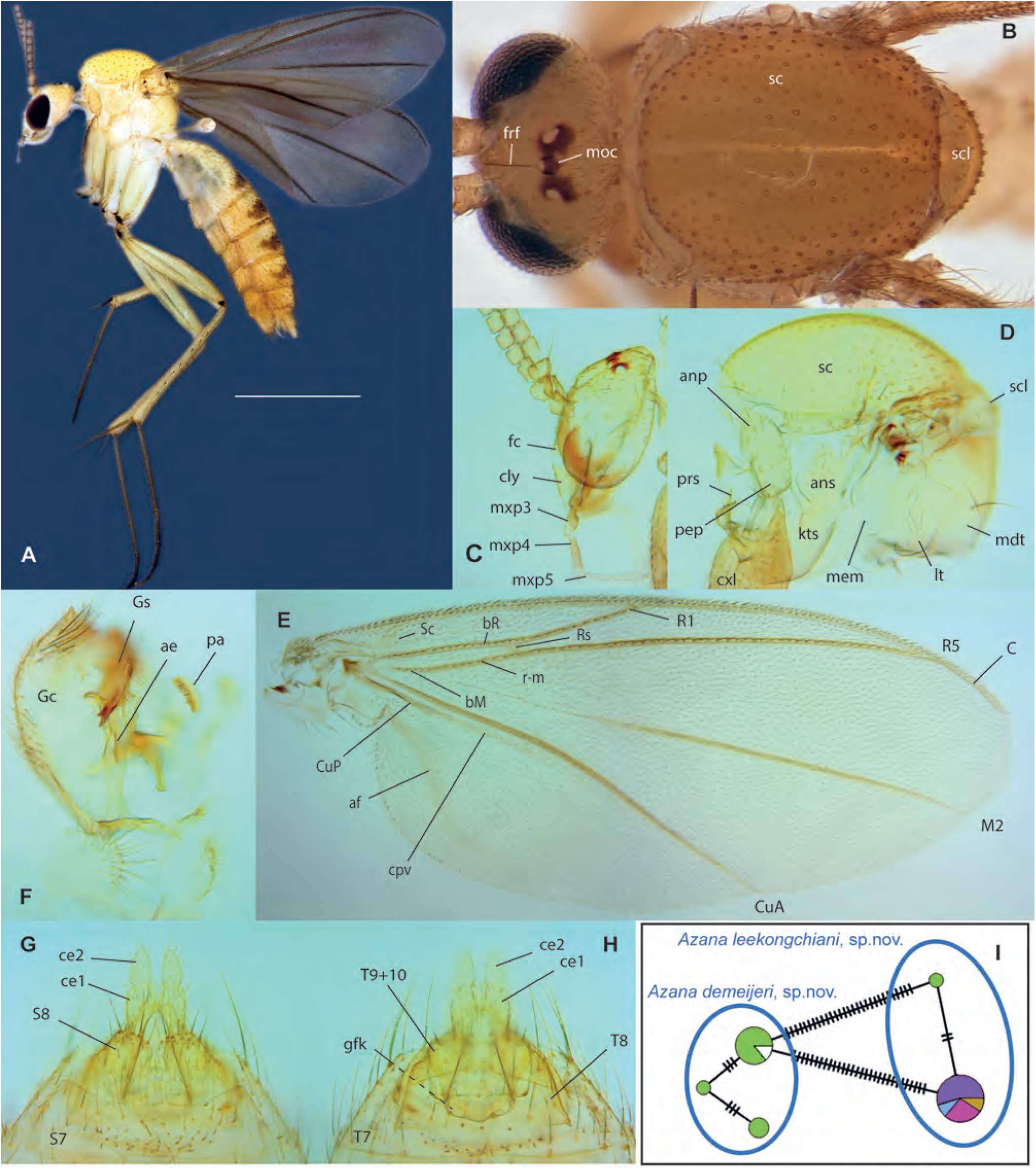
*Azana demeijeri* Amorim & Oliveira, **sp. n. A.** Habitus, lateral view, male holotype. **B.** Dorsal view of head and thorax, male holotype. **C.** Head, lateral view, female paratype, ZRCBDP0047941. **D.** Thorax, lateral view, same. **E.** Wing, female paratype, ZRCBDP0047941. **F.** Terminalia, Internal view, male holotype. **G.** Terminalia, ventral view, same. **H.** Terminalia, dorsal view, same. **I.** Haplotype network for *Azana*.

**Figs. 10A-F.**
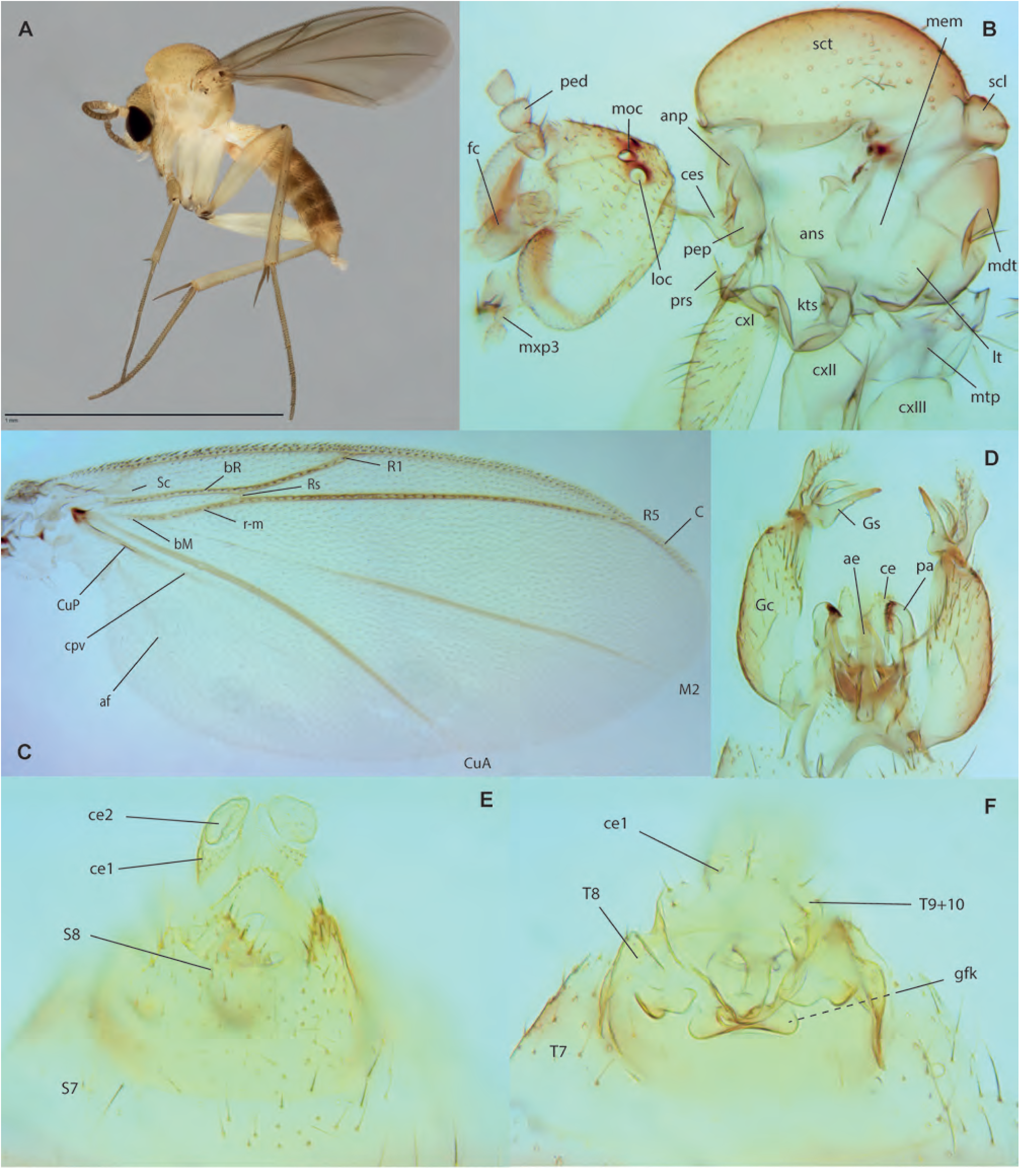
*Azana leekongchiani* Amorim & Oliveira, sp. n. A. Habitus, lateral view, female paratype ZRCBDP0049320. B. Head and thorax, lateral view, male holotype. C. Wing, male holotype. D. Male terminalia, ventral view, holotype. E. Female terminalia, ventral view, paratype ZRCBDP0049121. F. Female terminalia, dorsal view, same. Tetragoneurinae

## Tetragoneurinae

**Fig. 11.**
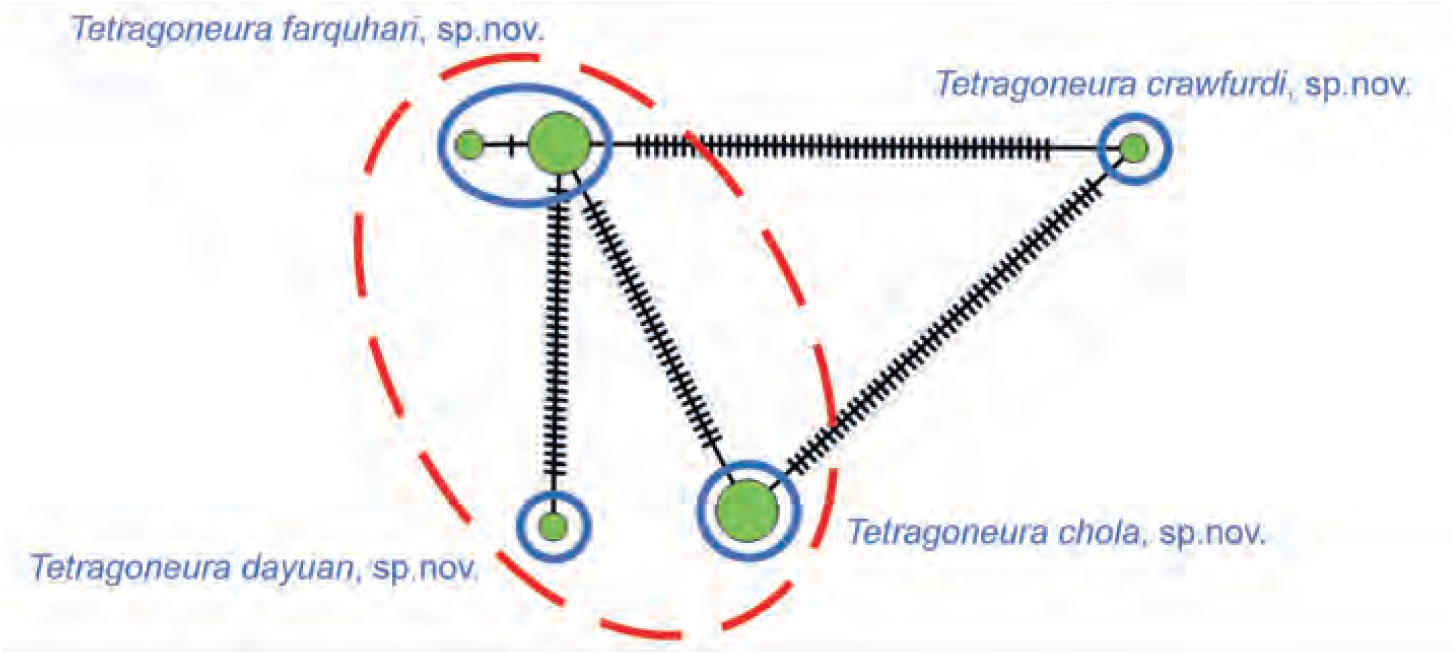
Haplotype network for *Tetragoneura*.

**Figs. 12A-D.**
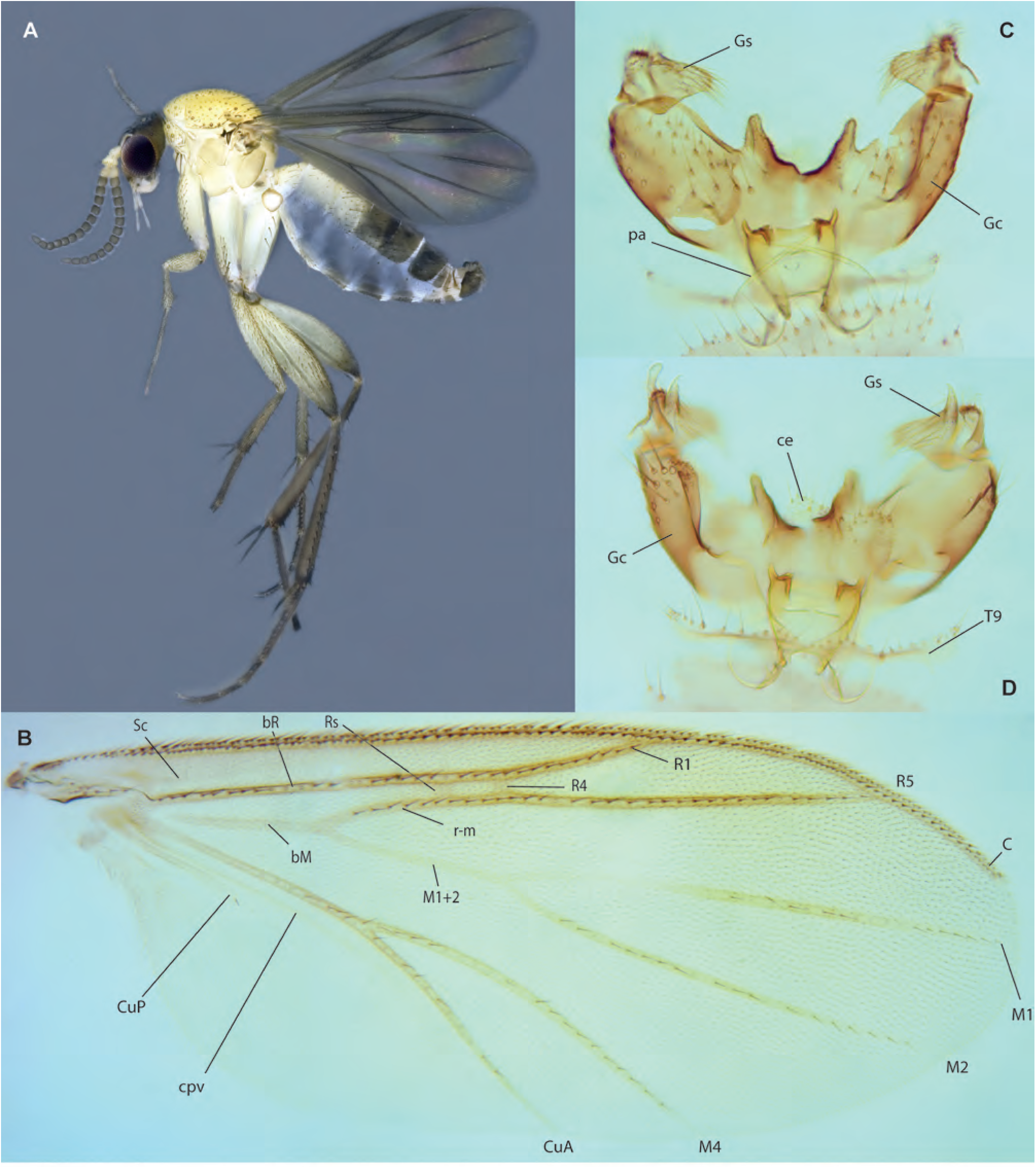
*Tetragoneura crawfurdi* Amorim & Oliveira, **sp. n.,** male holotype. **A.** Habitus, lateral view. **B.** Wing. **C.** Terminalia, ventral view. **D.** Terminalia, dorsal view.

**Figs. 13A-E.**
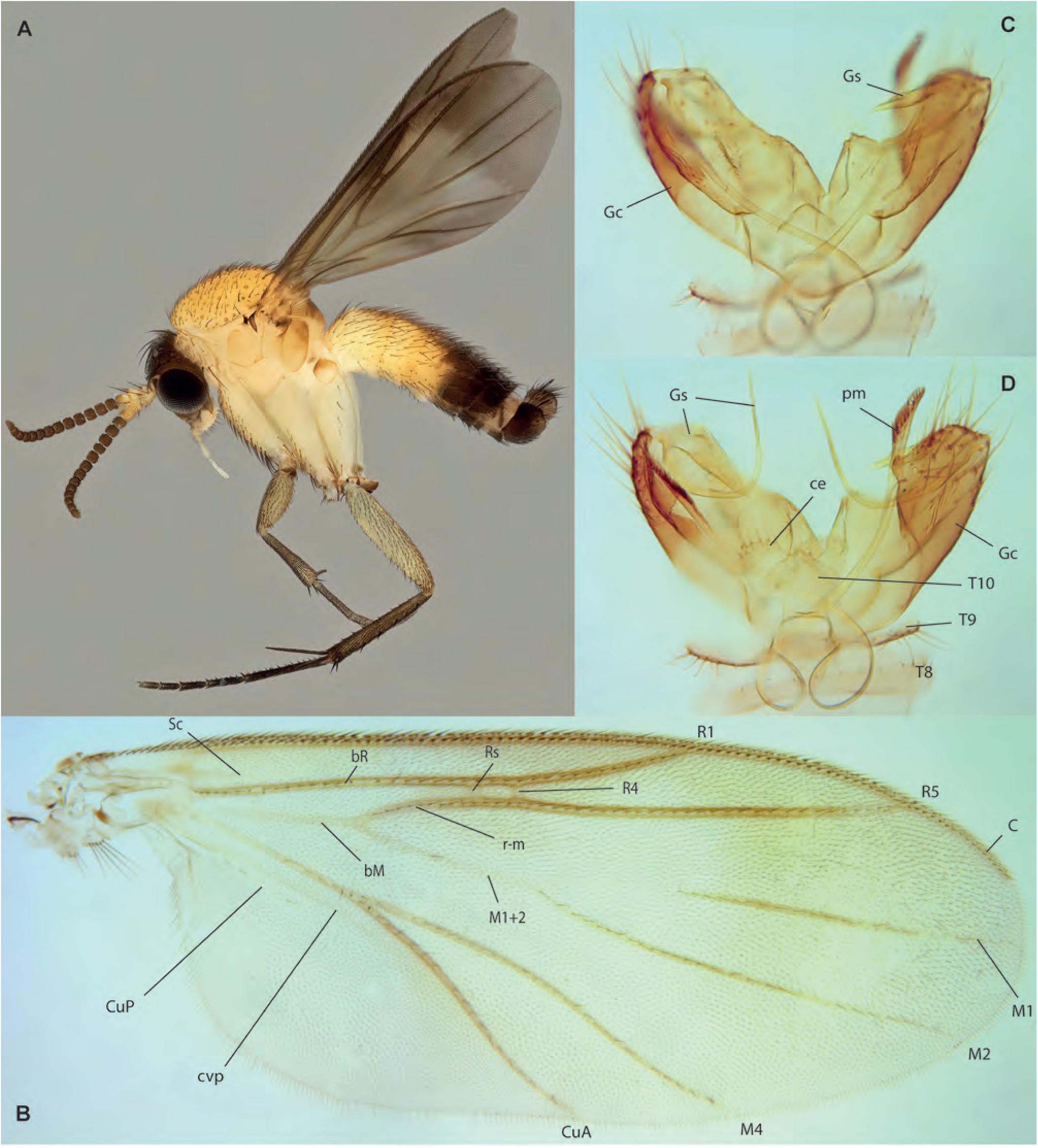
*Tetragoneura chola* Amorim & Oliveira, **sp. n. A.** Male, habitus, lateral view, paratype ZRCBDP0048501. **B.** Wing, male holotype. **C.** Male terminalia, ventral view, same. **D.** Male terminalia, ventral view, same.

**Figs. 14A-F.**
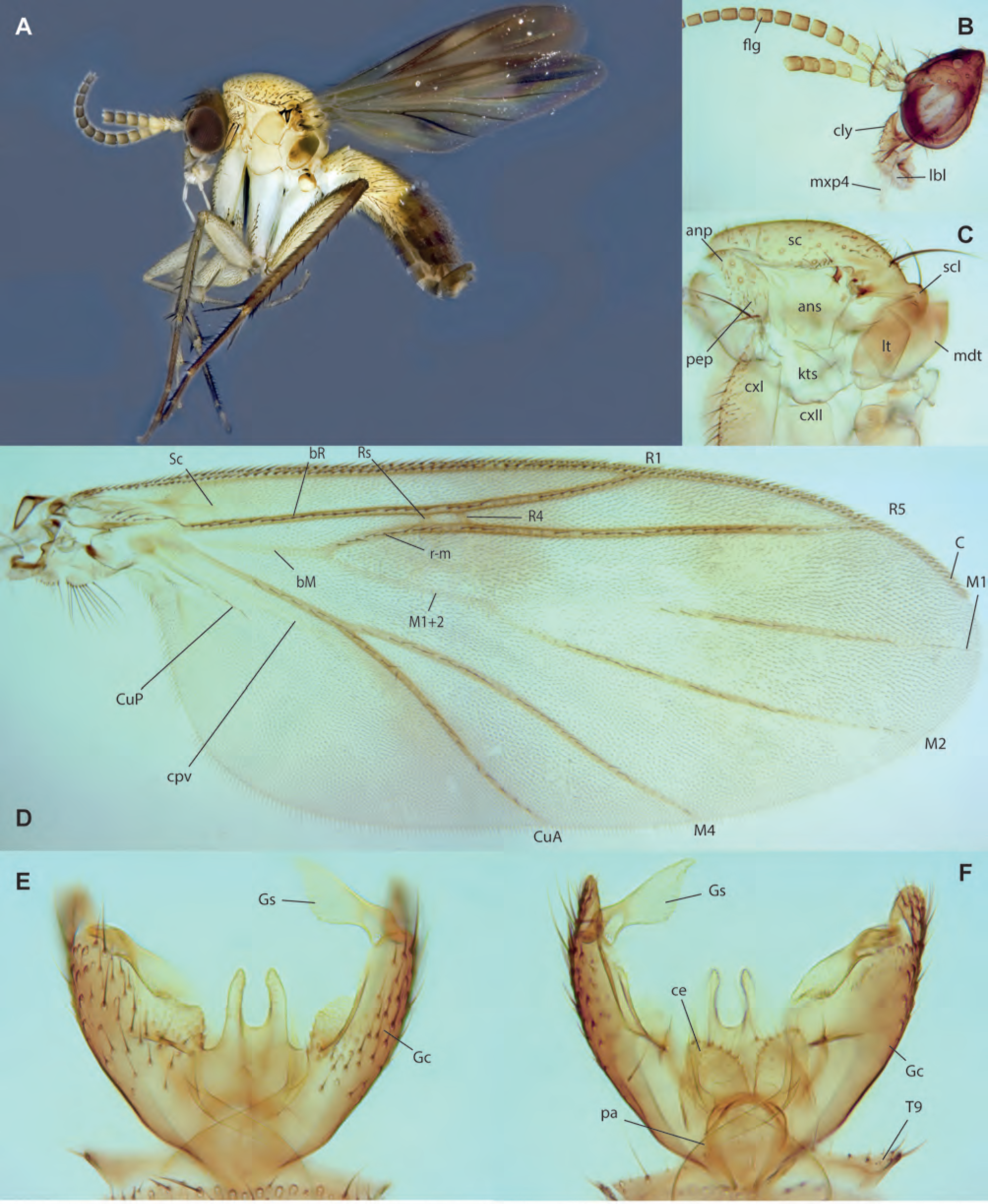
*Tetragoneura dayuan* Amorim & Oliveira, **sp. n.,** male holotype. **A.** Habitus, lateral view. **B.** Head, lateral view. **C.** Thorax, lateral view. **D.** Wing. **E.** Male terminalia, ventral view. **F.** Male terminalia, dorsal view.

**Figs. 15A-D.**
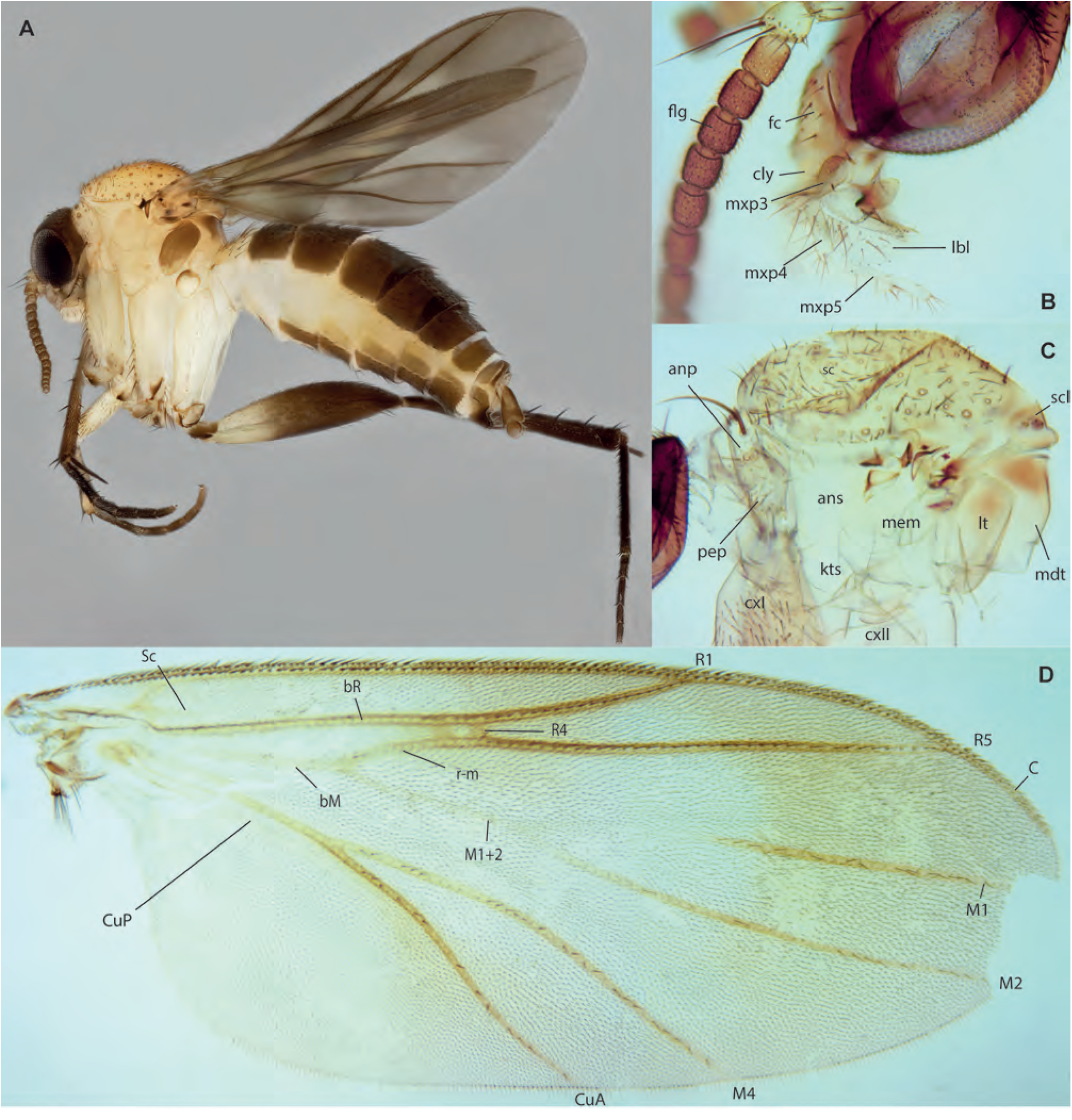
*Tetragoneura farquhari* Amorim & Oliveira, **sp. n. A.** Habitus, lateral view, female paratype ZRCBDP0048503. **B.** Head, lateral view, male holotype. **C.** Thorax, lateral view, same. **D.** Wing, same.

**Figs. 16A-D.**
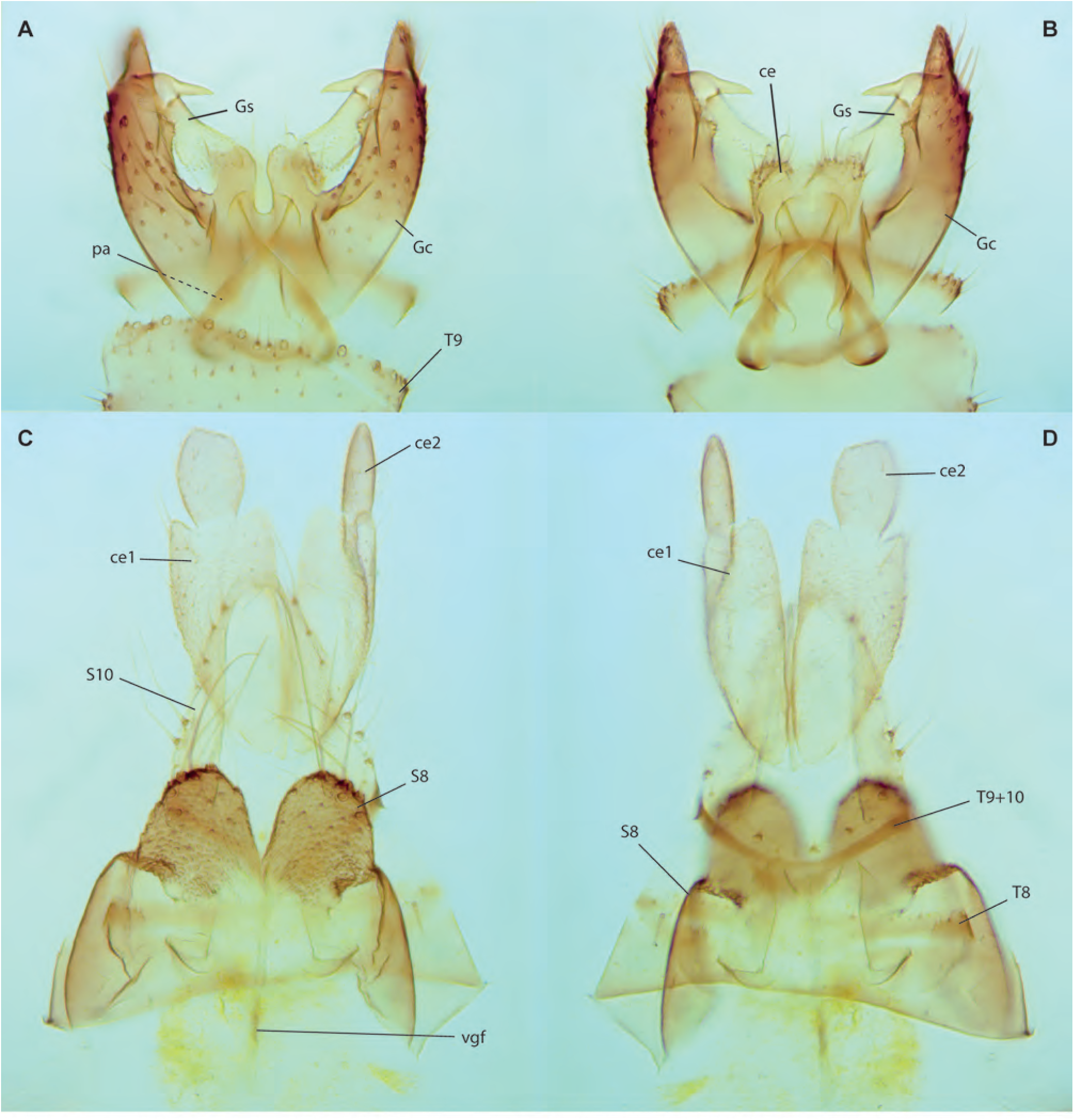
*Tetragoneura farquhari* Amorim & Oliveira, **sp. n. A.** Male terminalia, ventral view, holotype. **B.** Male terminalia, dorsal view, same. **C.** Female terminalia, ventral view, paratype ZRCBDP0048503. **D.** Female terminalia, dorsal view, same.

**Figs. 17A-J.**
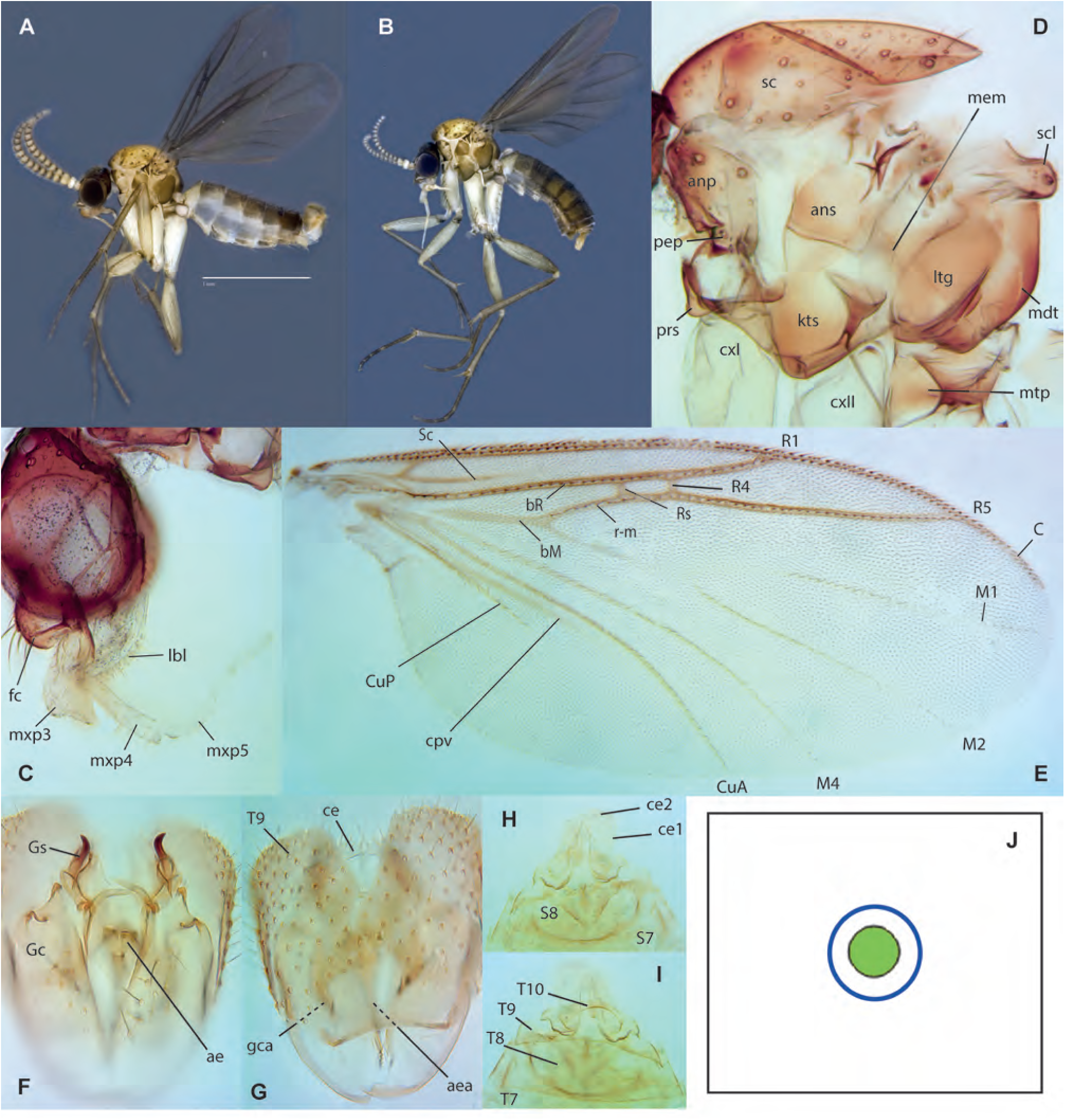
*Ectrepesthoneura johor* Amorim & Oliveira, **sp. n. A.** Habitus, lateral view, male holotype. **B.** Habitus, lateral view, female paratype ZRCBDP0048506. **C.** Head, lateral view, same. **D.** Thorax, lateral view, male holotype. **E.** Wing, same. **F.** Male terminalia, ventral, same. **G.** Male terminalia, dorsal, same. **H.** Female terminalia, ventral view, paratype ZRCBDP0048506. **I.** Female terminalia, dorsal view, same. **J.** Haplotype network for *Ectrepesthoneura*. **Leiinae**

## Leiinae

**Figs. 18A-G.**
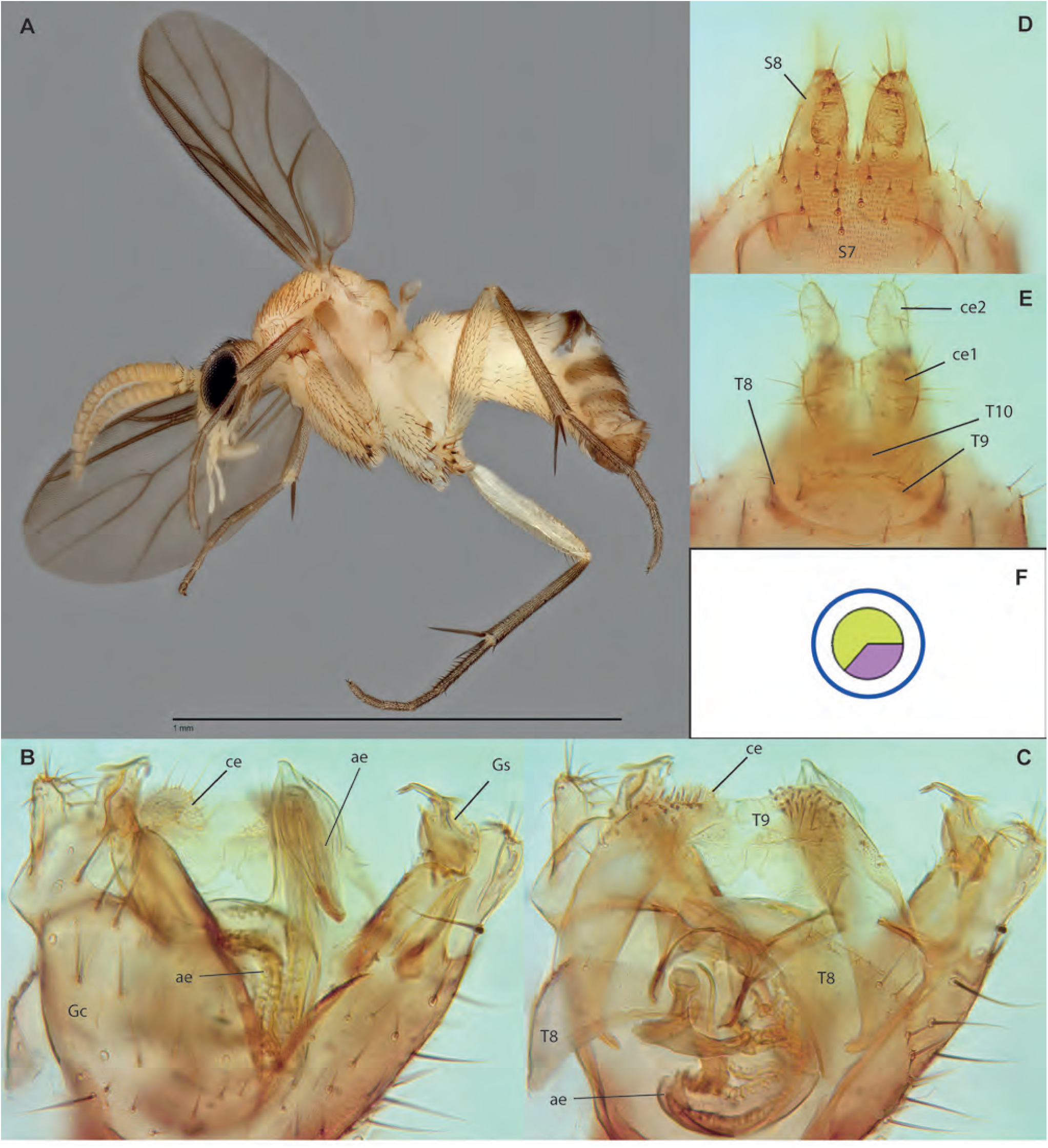
*Mohelia zubirsaidi* Amorim & Oliveira, **sp. n. A.** Male, habitus, lateroventral view, paratype ZRCBDP0048984. **B.** Male terminalia, ventral view, same. **C.** Male terminalia, internal view, same. **D.** Male terminalia, dorsal view, same. **E.** Female terminalia, ventral, paratype ZRCBDP0048999. **F.** Female terminalia, dorsal view, same. **G.** Haplotype network for *Mohelia*.

**Fig. 19.**
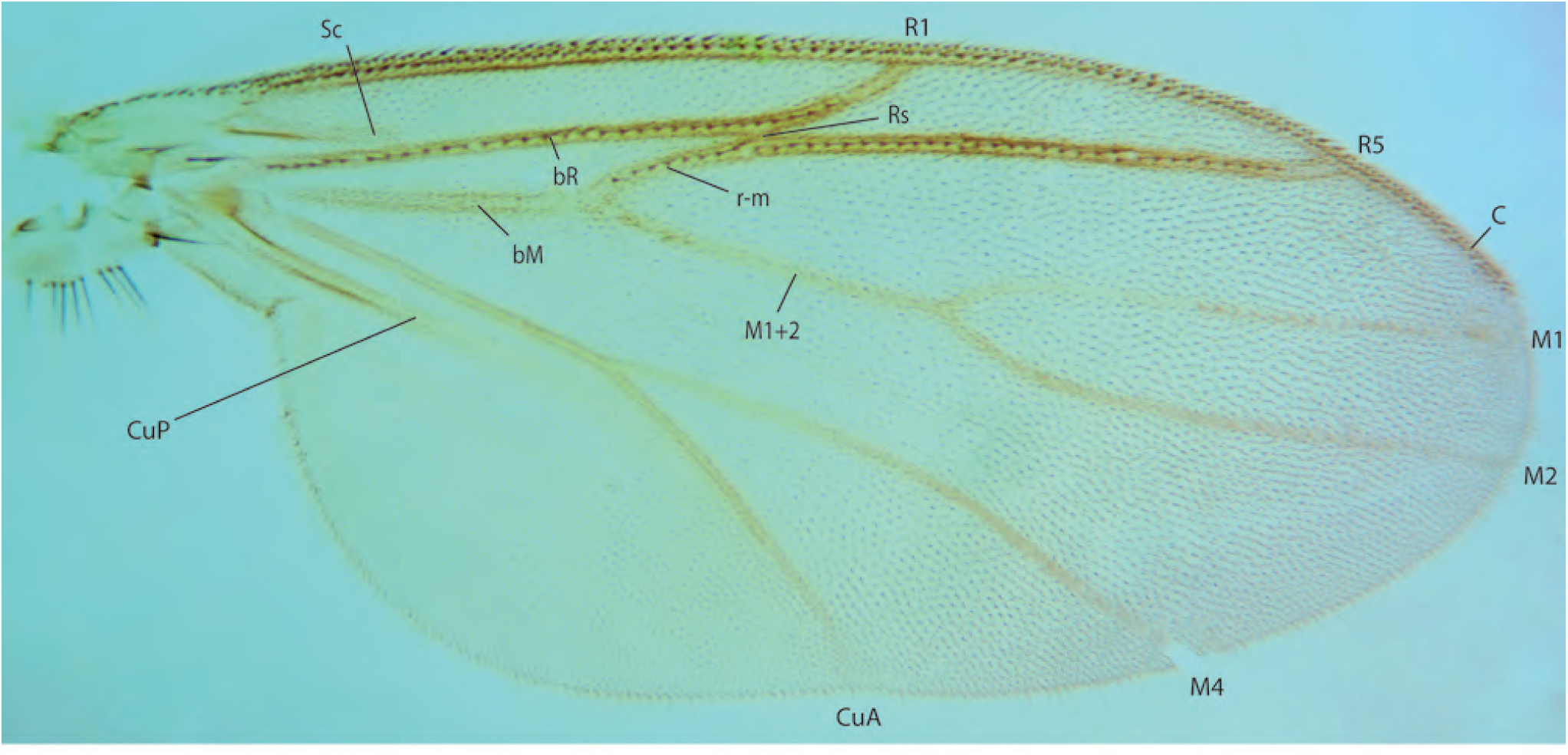
*Mohelia zubirsaidi* Amorim & Oliveira, **sp. n.,** wing, male holotype.

**Figs. 20A-B.**
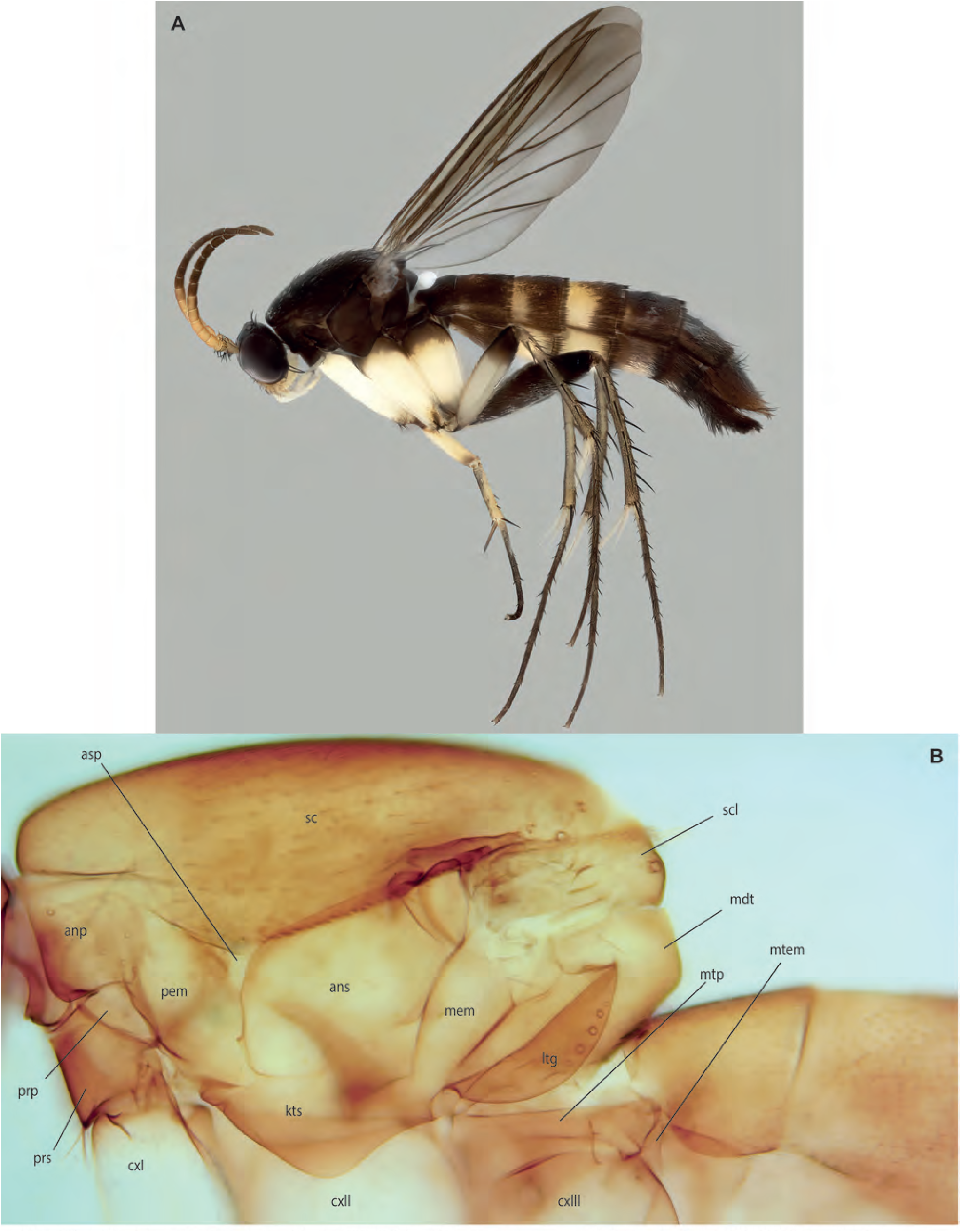
*Allactoneura tumasik* Amorim & Oliveira, **sp. n.,** male holotype, ZRCBDP0048240. **A.** Habitus, lateral view. **B.** Thorax, lateral view.

**Figs. 21A-F.**
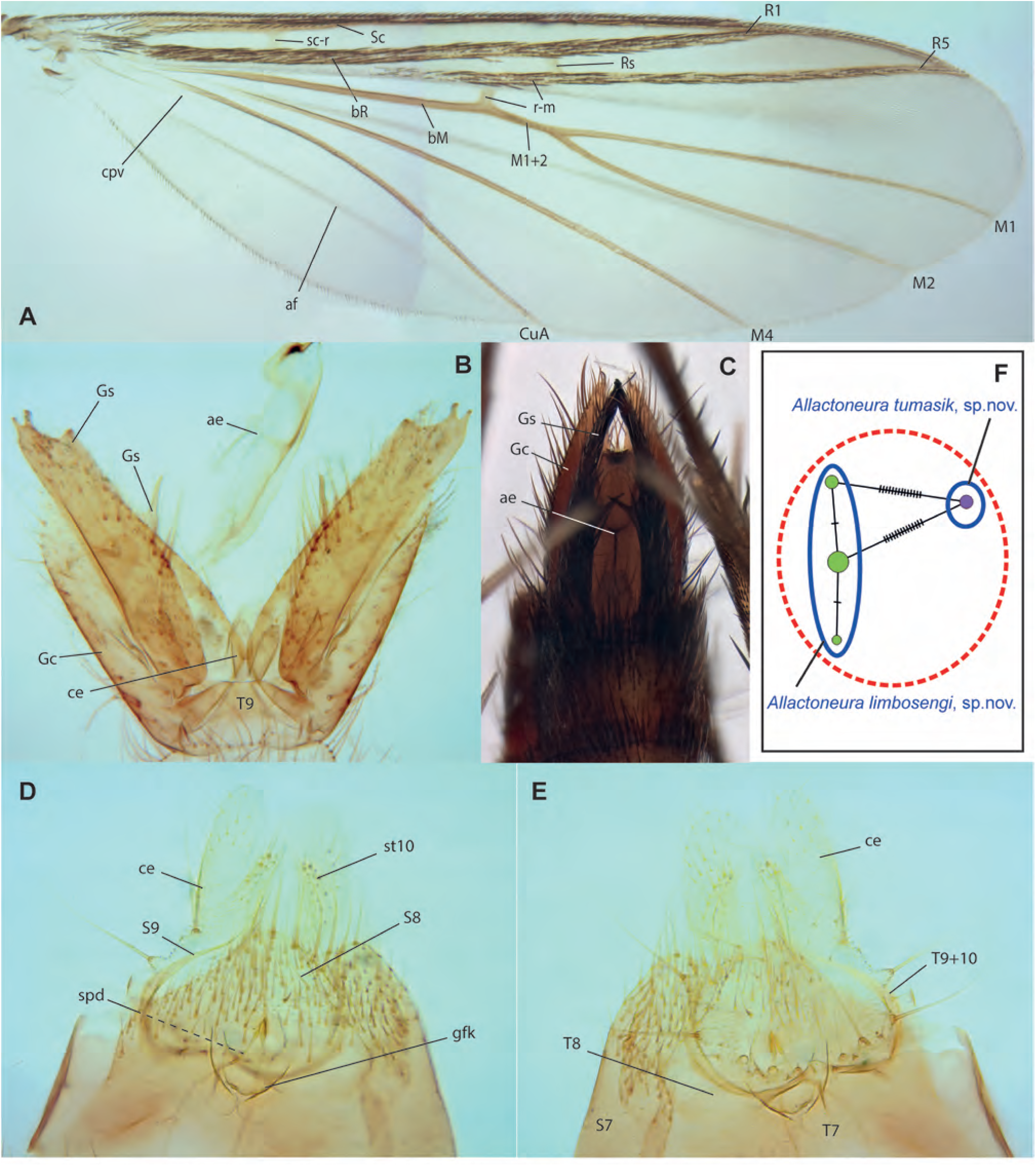
*Allactoneura tumasik*, Amorim & Oliveira, **sp. n. A.** Wing, holotype. **B.** Male terminalia, ventral view, same. **C.** Detail of distal end of gonostylus, same. **D.** Female terminalia, ventral view, paratype ZRCBDP0048284. **E.** Same, dorsal view. **F.** Haplotype network for *Allactoneura*.

**Figs. 22.**
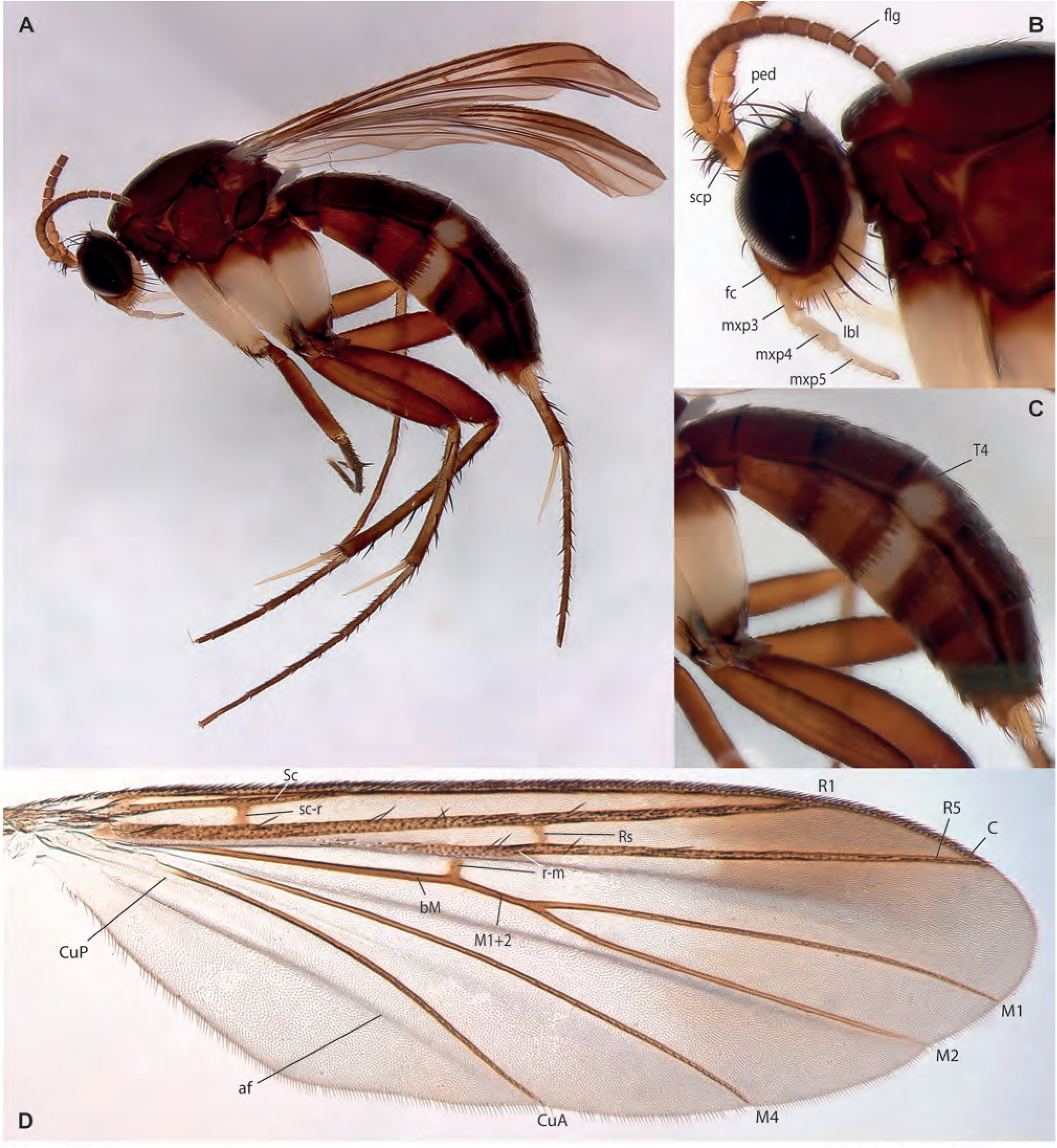
*Allactoneura limbosengi* Amorim & Oliveira, **sp. n. A.** Habitus, lateral view, female paratype, ZRCBDP0047786. **B.** Head, lateral view, same. **C.** Abdomen, lateral view, same. **D.** Wing, male holotype ZRCBDP0278244.

**Figs. 23.**
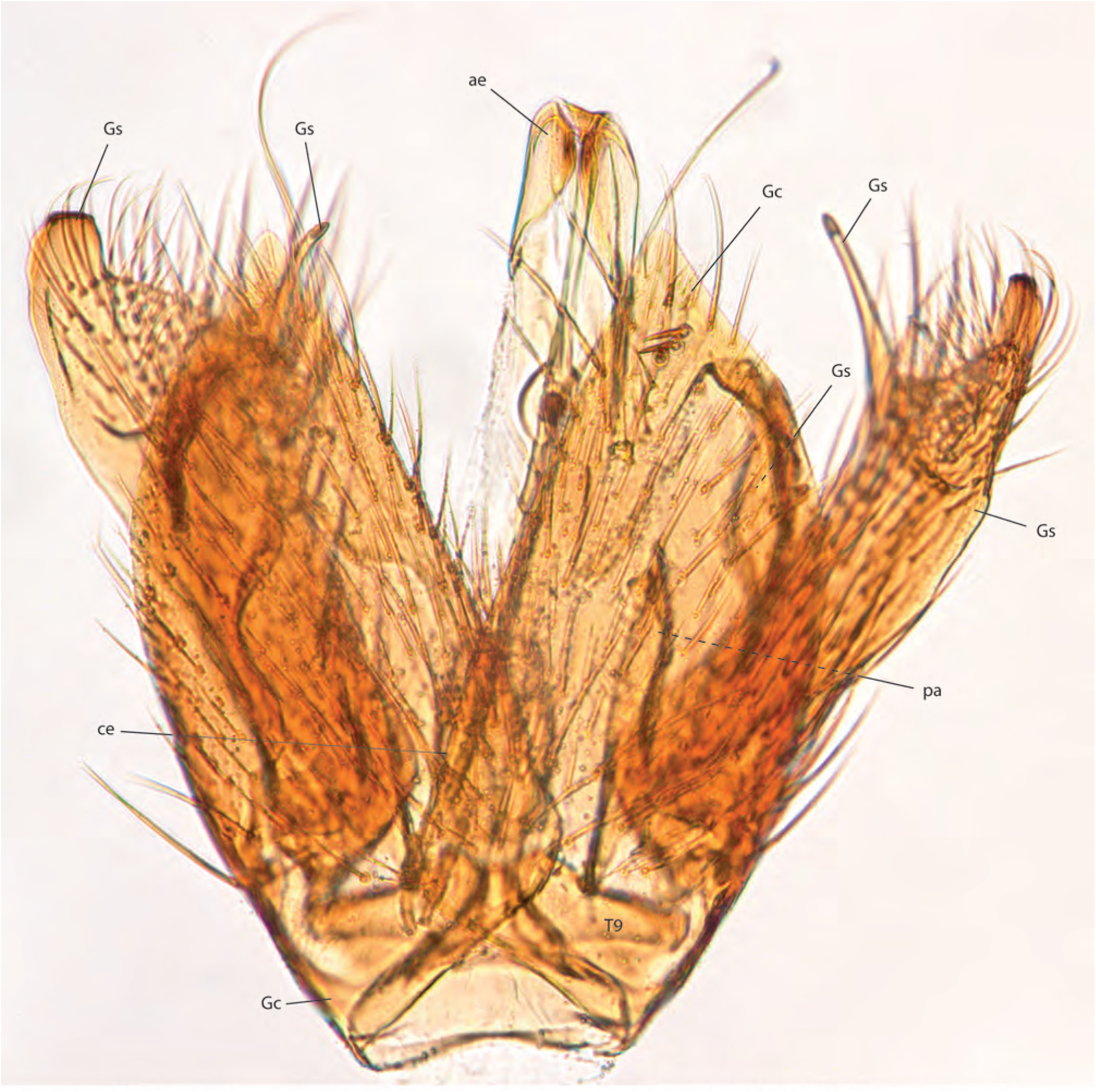
*Allactoneura limbosengi* Amorim & Oliveira, **sp. n.,** male holotype ZRCBDP0278244, terminalia, dorsal view.

**Figs. 24A-H.**
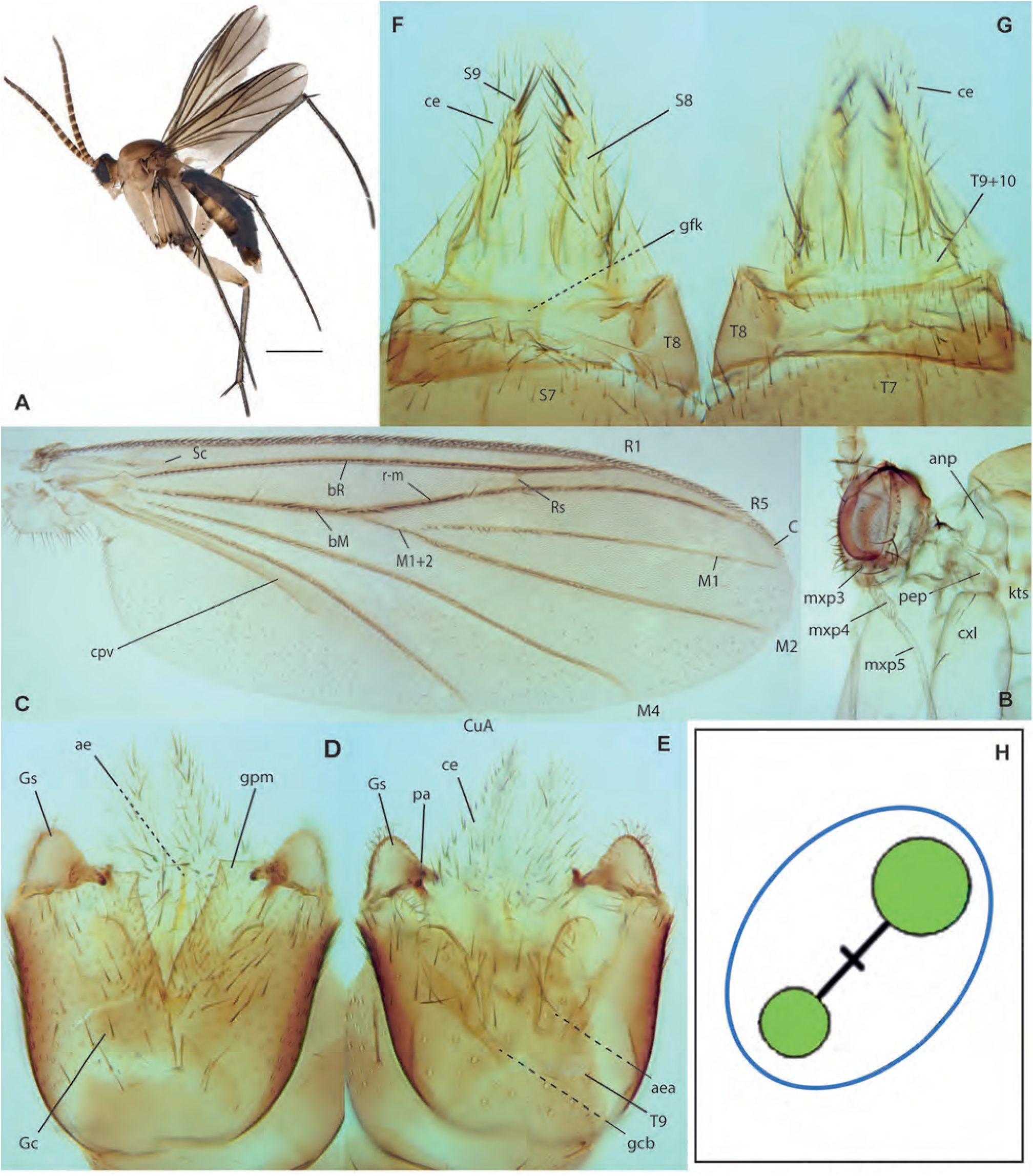
*Eumanota racola* Søli. **A.** Habitus, lateral view, female ZRCBDP0048561. **B.** Head, lateral view, male ZRCBDP0048560. **C.** Wing, same. **D.** Male terminalia, ventral view, same. **E.** Male terminalia, dorsal view, same. **F.** Female terminalia, ventral view, ZRCBDP0048927. **G.** Terminalia, dorsal view, same. **H.** Haplotype network for *Eumanota*.

**Figs. 25A-C.**
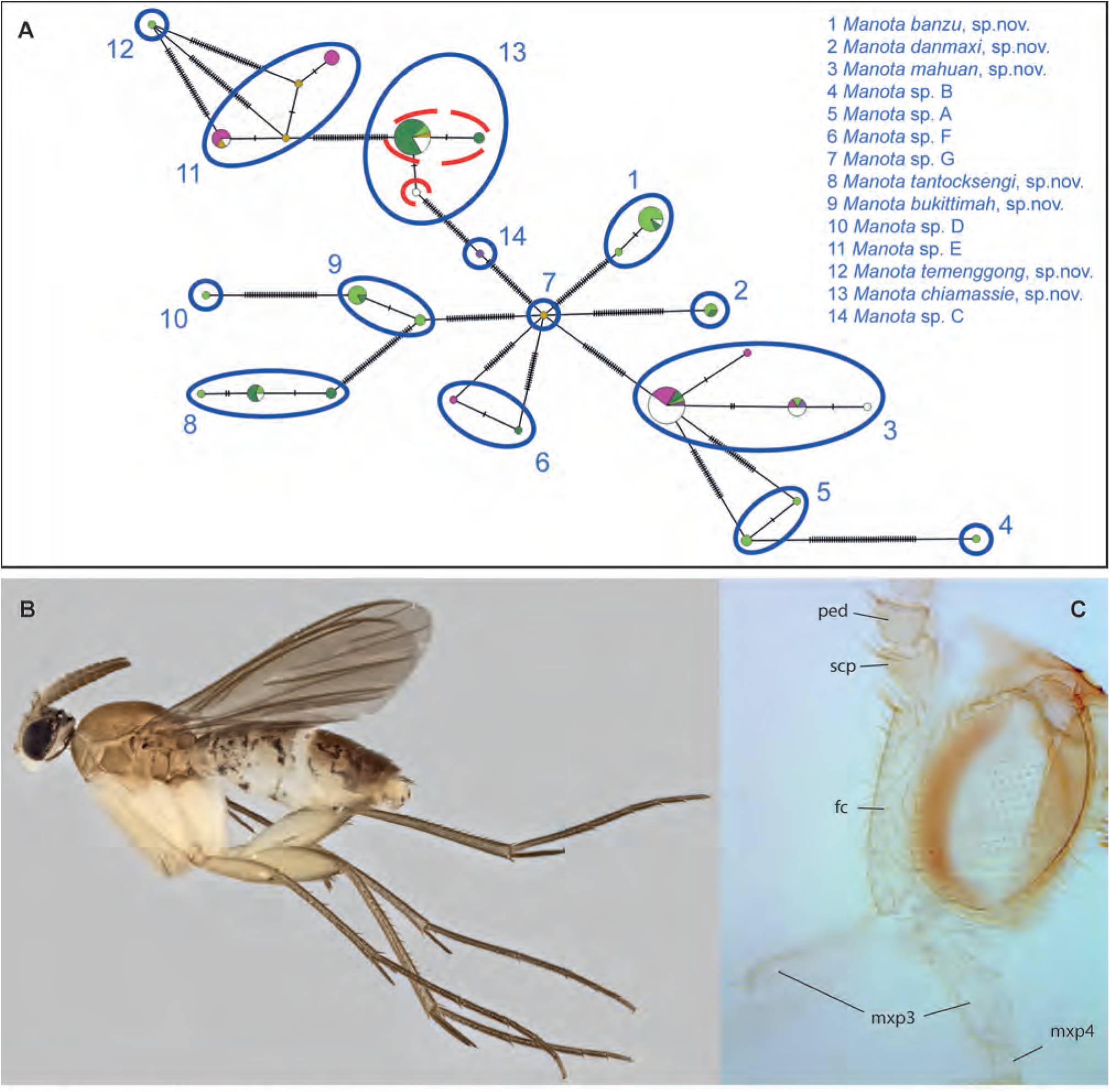
A. Haplotype network for *Manota*. **B-C.** *Manota banzu* Amorim & Oliveira, **sp. n. B.** Habitus, lateral view, female paratype ZRCBDP0048517. **C.** Head, lateral view, male paratype ZRCBDP0137281.

**Figs. 26A-D.**
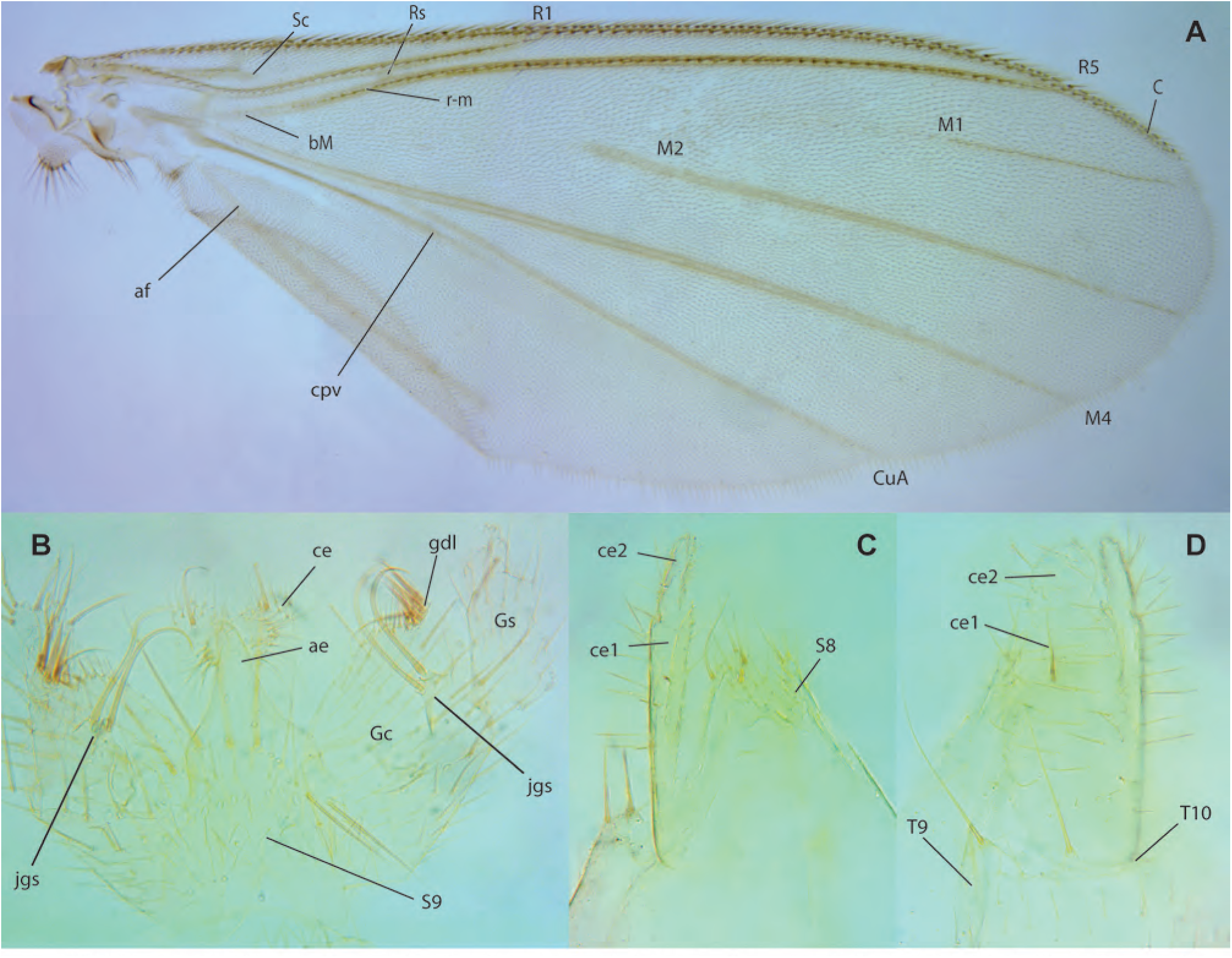
*Manota banzu* Amorim & Oliveira, **sp. n. A.** Wing, male paratype ZRCBDP0047877. **B.** Terminalia, dorsal view, same. **C.** Female terminalia, ventral view, paratype ZRCBDP0048677. **D.** Female terminalia, dorsal view, same.

**Figs. 27A-E.**
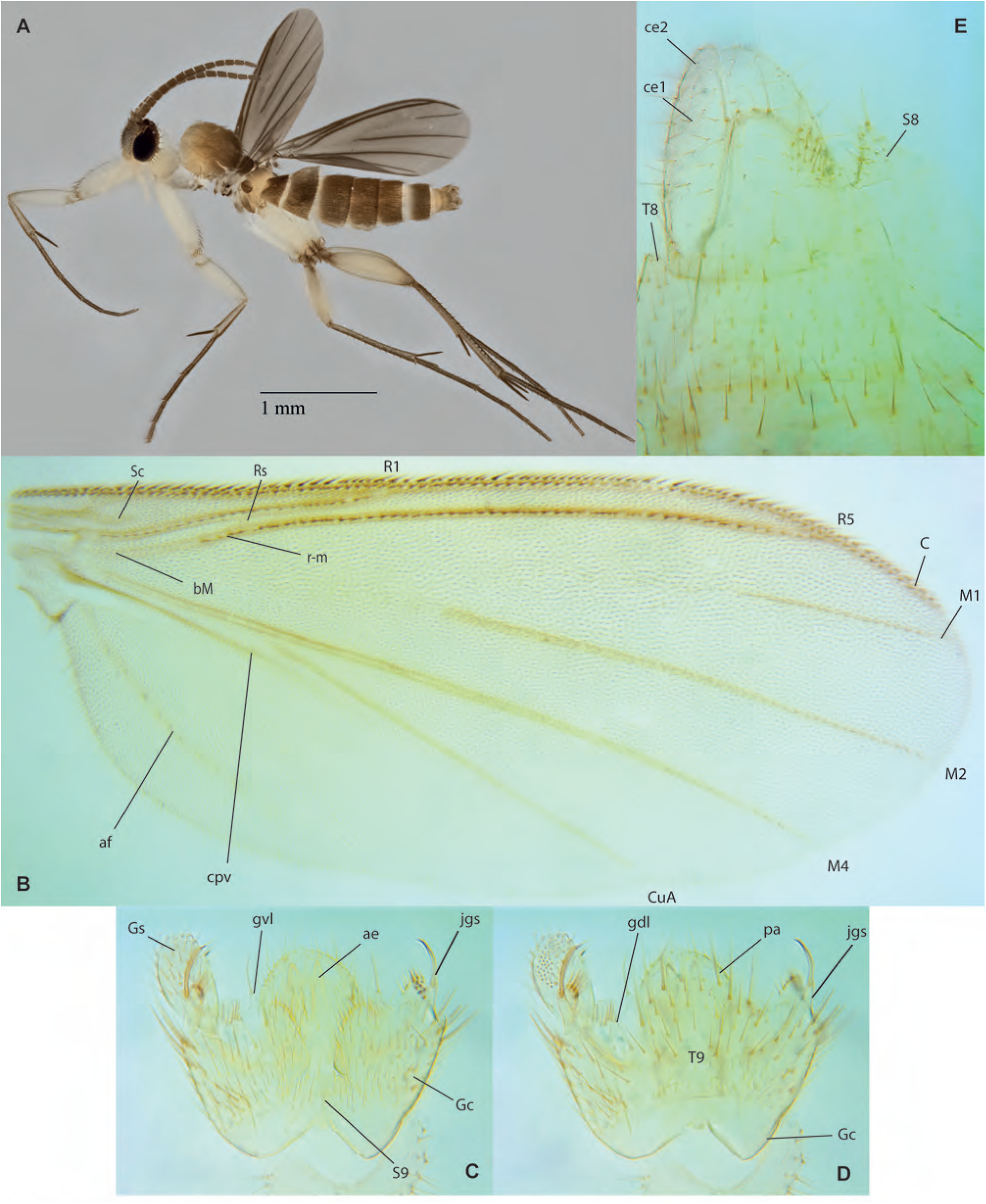
*Manota tantocksengi* Amorim & Oliveira, **sp. n. A.** Habitus, lateral view, male paratype ZRCBDP0137247. **B.** Wing, male holotype. **C.** Male terminalia, ventral view, same. **D.** Male terminalia, dorsal view, same. **E.** Female terminalia, ventrolateral view, paratype ZRCBDP0072744.

**Figs. 28A-G.**
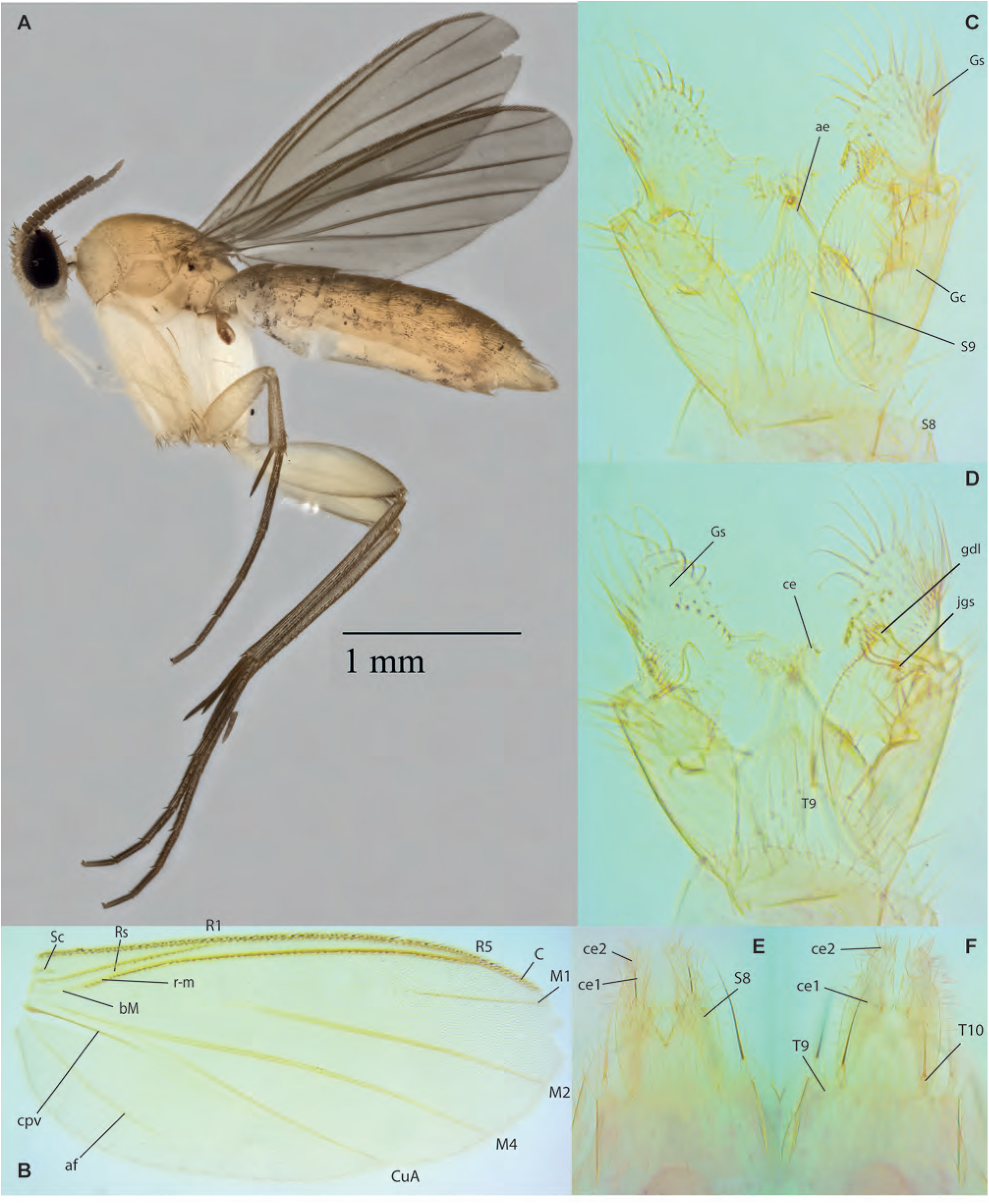
*Manota bukittimah* Amorim & Oliveira, **sp. n. A.** Habitus, lateral view, female paratype, ZRCBDP0137035. **B.** Wing, male holotype. **C.** Terminalia, ventral view, same. **D.** Terminalia, dorsal view, same. **E.** Female terminalia, ventral view, paratype ZRCBDP0048527. **F.** Female terminalia, dorsal view, same.

**Figs. 29A-E.**
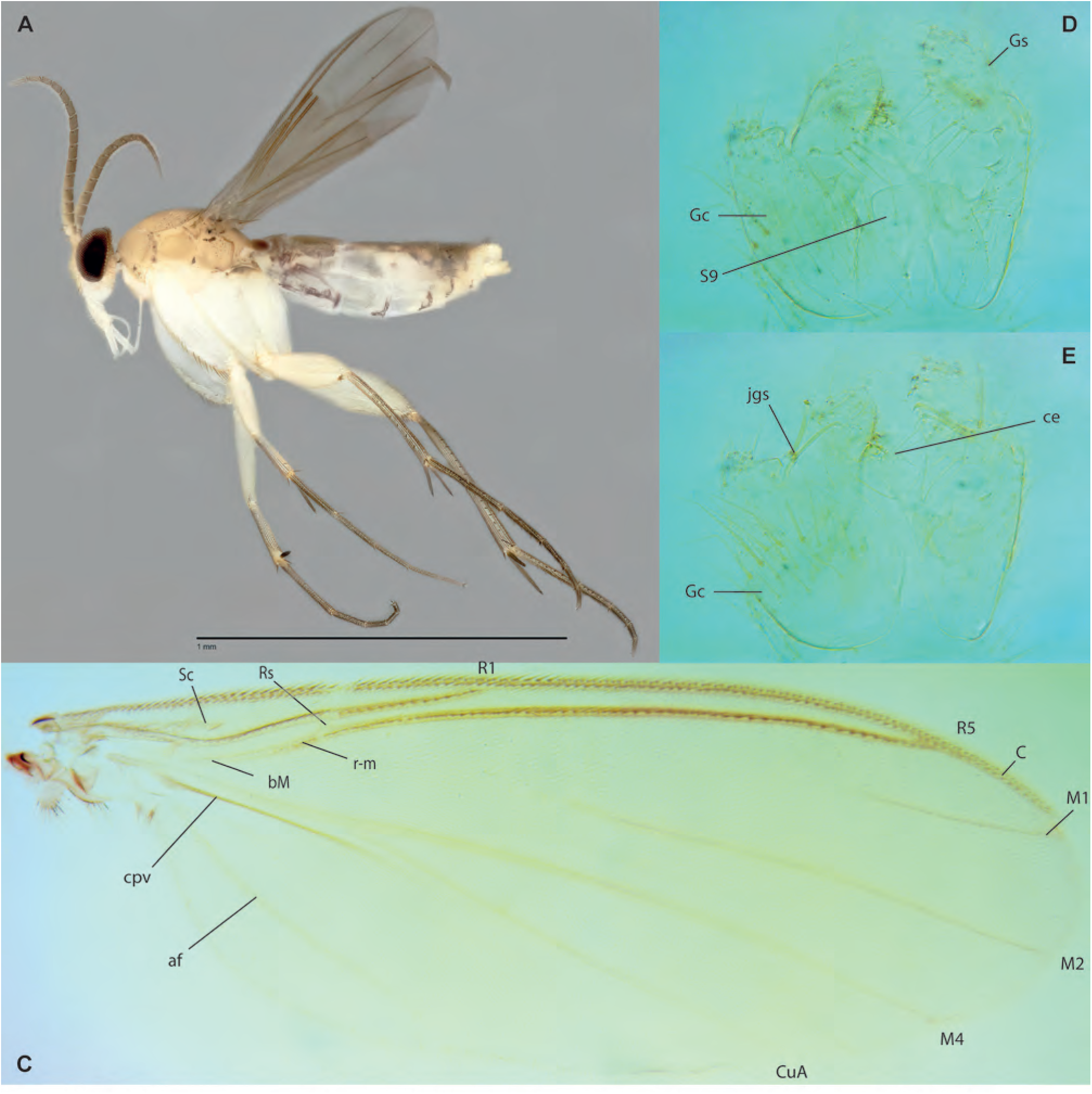
*Manota chiamassie* Amorim & Oliveira, **sp. n.,** male holotype. **A.** Habitus, lateral view. **B.** Head, lateral view. **C.** Wing. **D.** Terminalia, ventral view. **E.** Terminalia, dorsal view, same.

**Figs. 30A-H.**
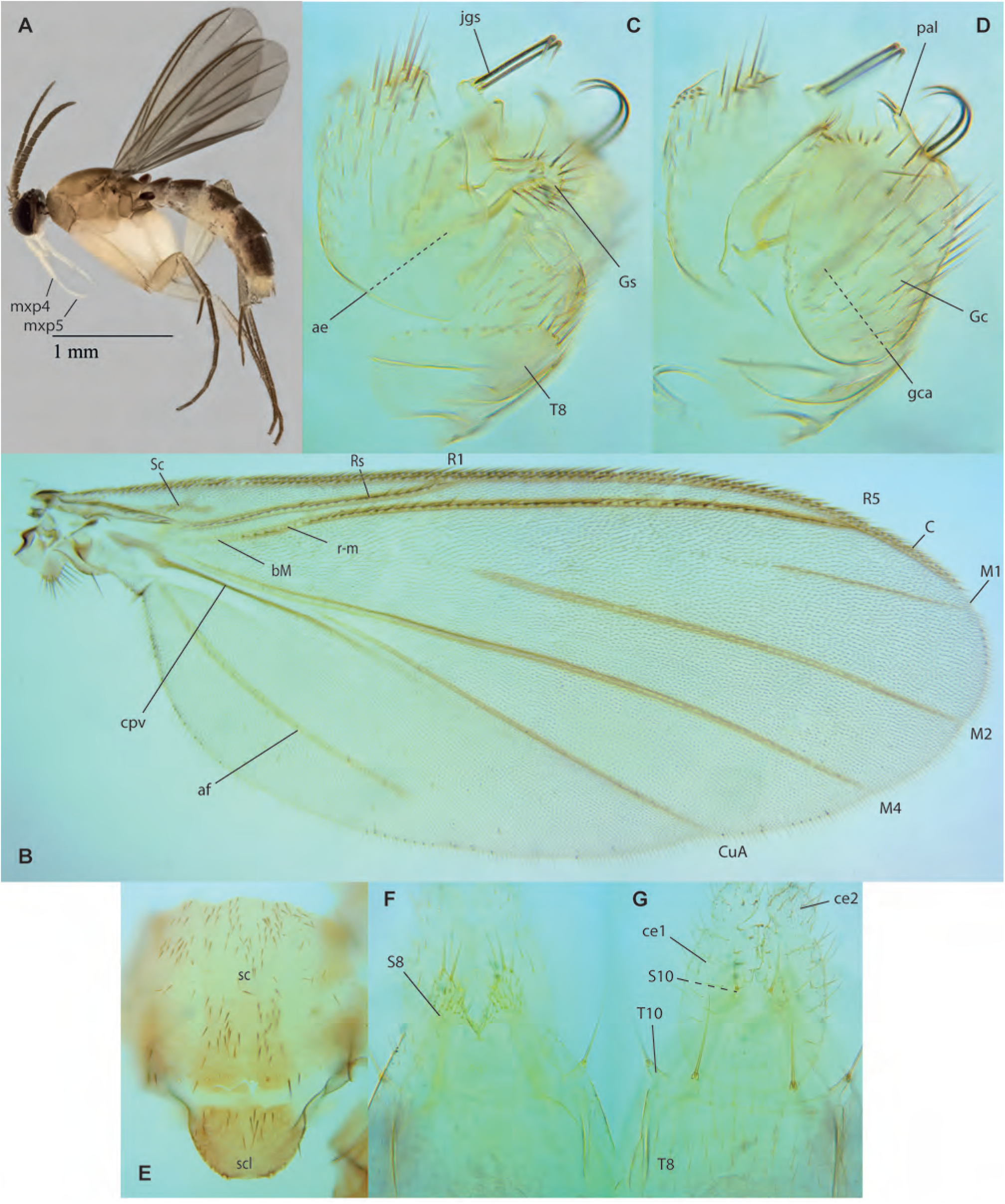
*Manota danmaxi* Amorim & Oliveira, **sp. n. A.** Habitus, lateral view, male paratype, ZRCBDP0137040. B. Wing, male holotype. C. Mesonotum, dorsal view, same. D-E Terminalia, lateral view, same. F. Female terminalia, ventral view, paratype ZRCBDP0074033. G. Female terminalia, dorsal view, same.

**Figs. 31A-E.**
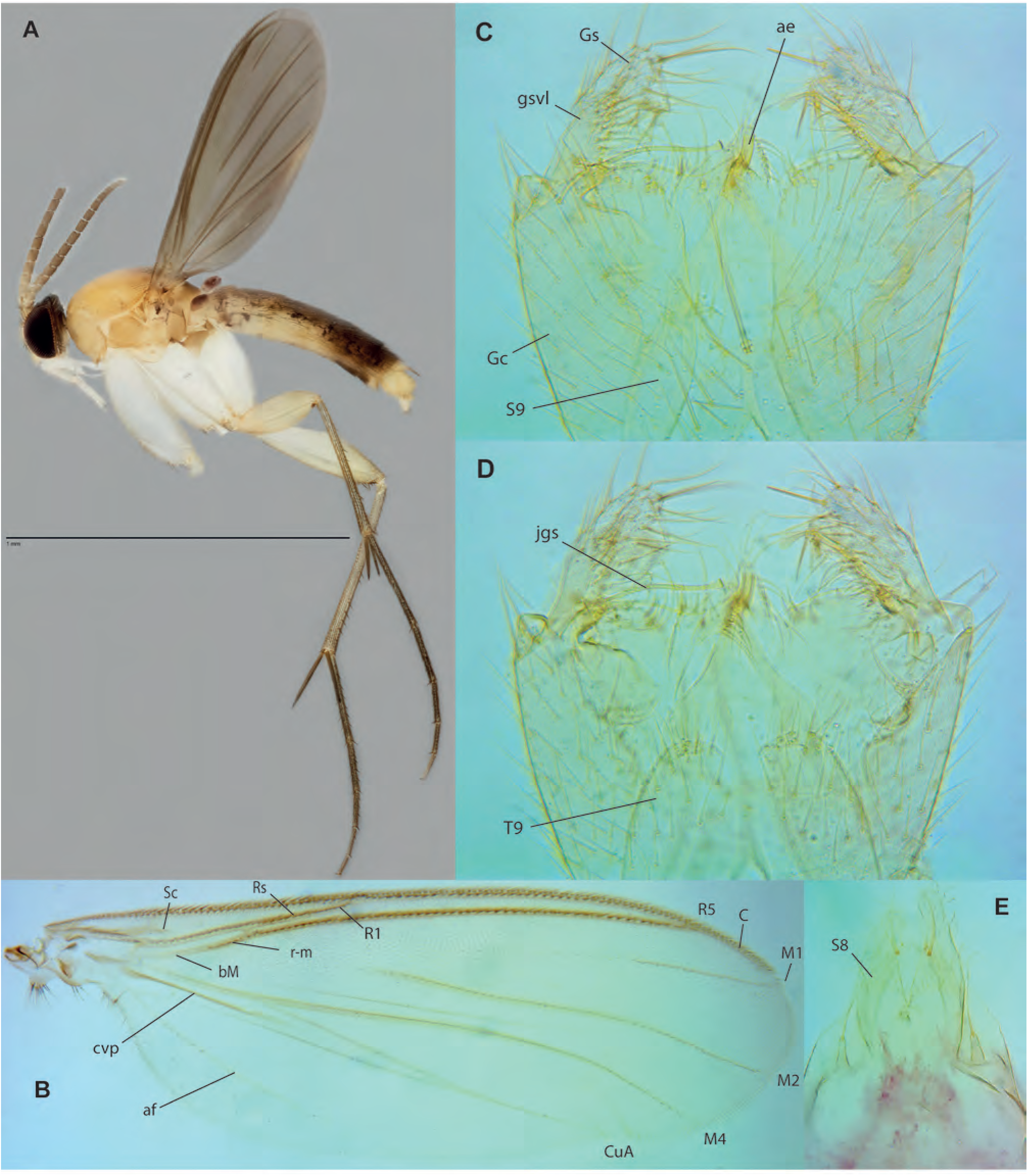
*Manota mahuan* Amorim & Oliveira, sp. n. A. Habitus, lateral view, male paratype, ZRCBDP0049338. B. Wing, male holotype (wing partially folded). C. Male terminalia, ventral view, same. F. Male terminalia, dorsal view. E. Female terminalia, ventral view, paratype ZRCBDP0049137

**Figs. 32A-E.**
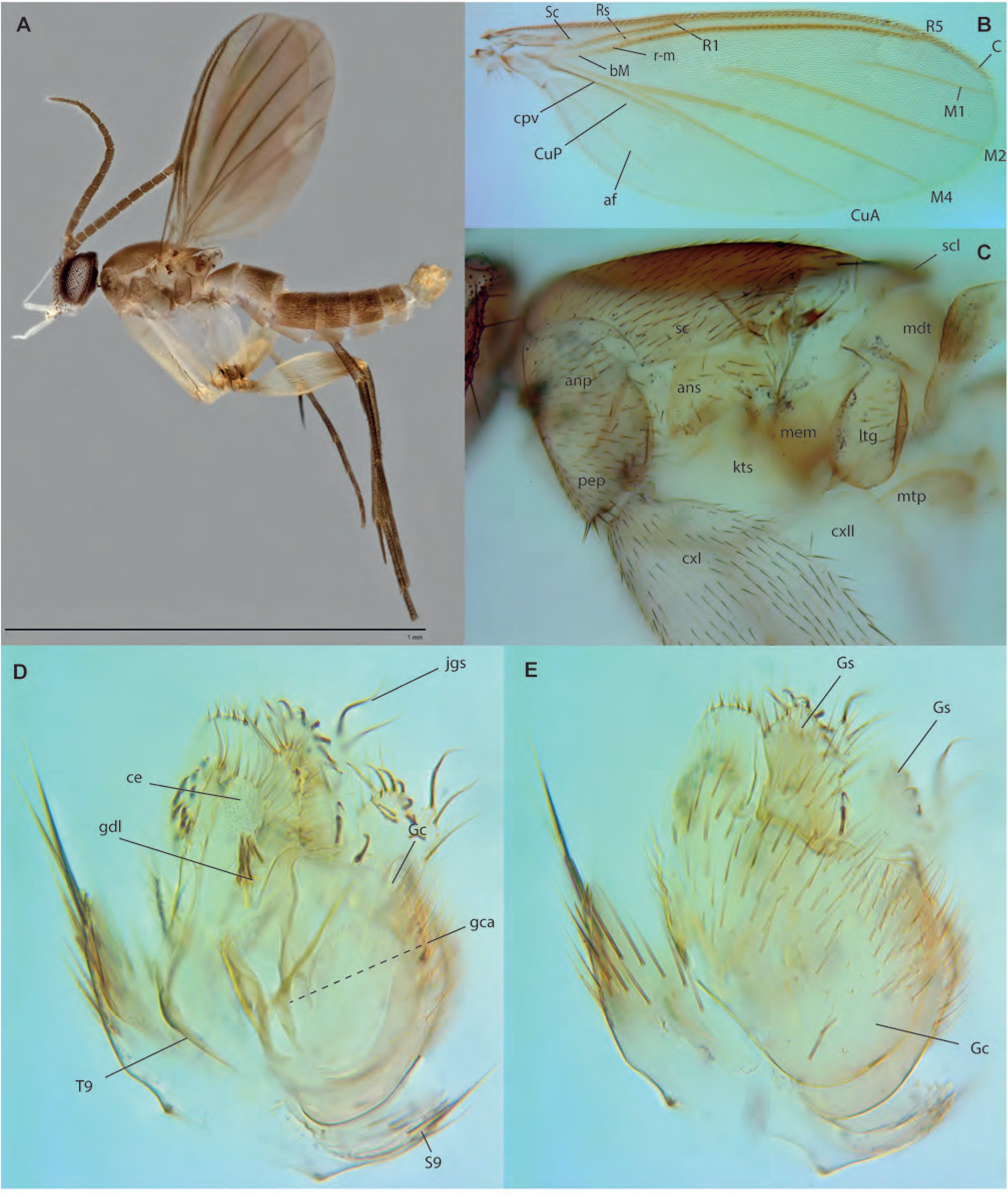
*Manota temenggong* Amorim & Oliveira, **sp. n.** 11, male holotype. **A.** Habitus, lateral view. **B.** Wing. **C.** Thorax, lateral view. **D.** Terminalia, lateral view, mid-section. **E.** Terminalia, lateral view, external section.

**Figs. 33A-B.**
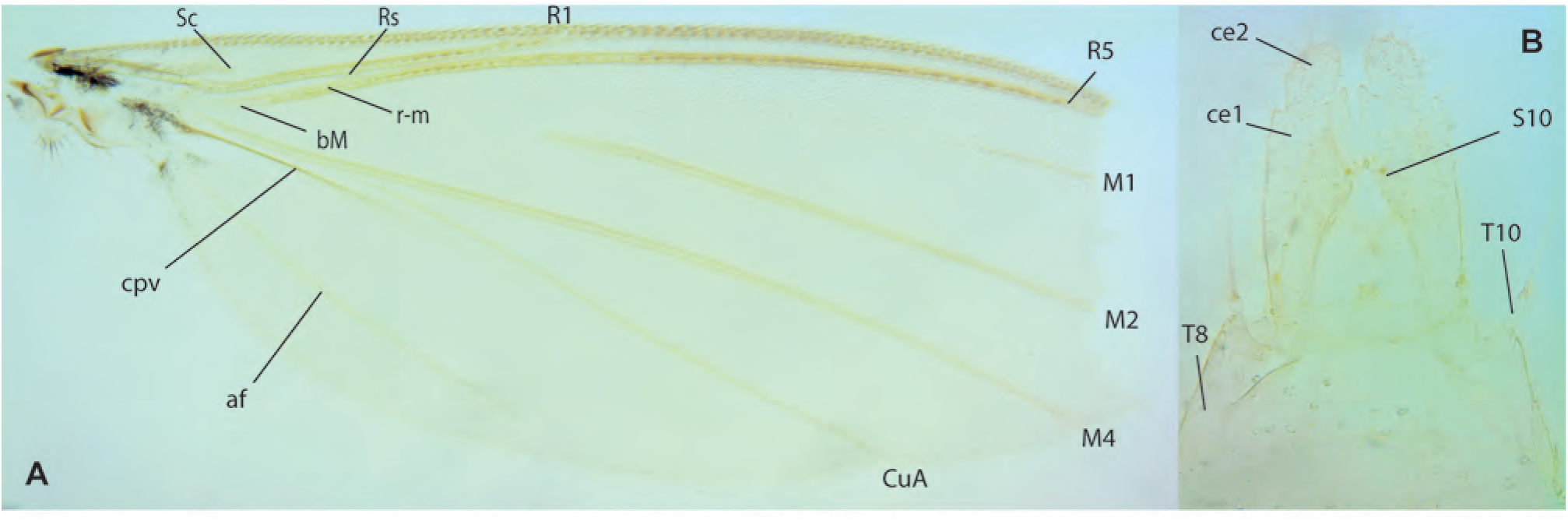
*Manota* sp. A, female, ZRCBDP0047870. **A.** Wing. **B.** Terminalia, dorsal view.

**Figs. 34A-C.**
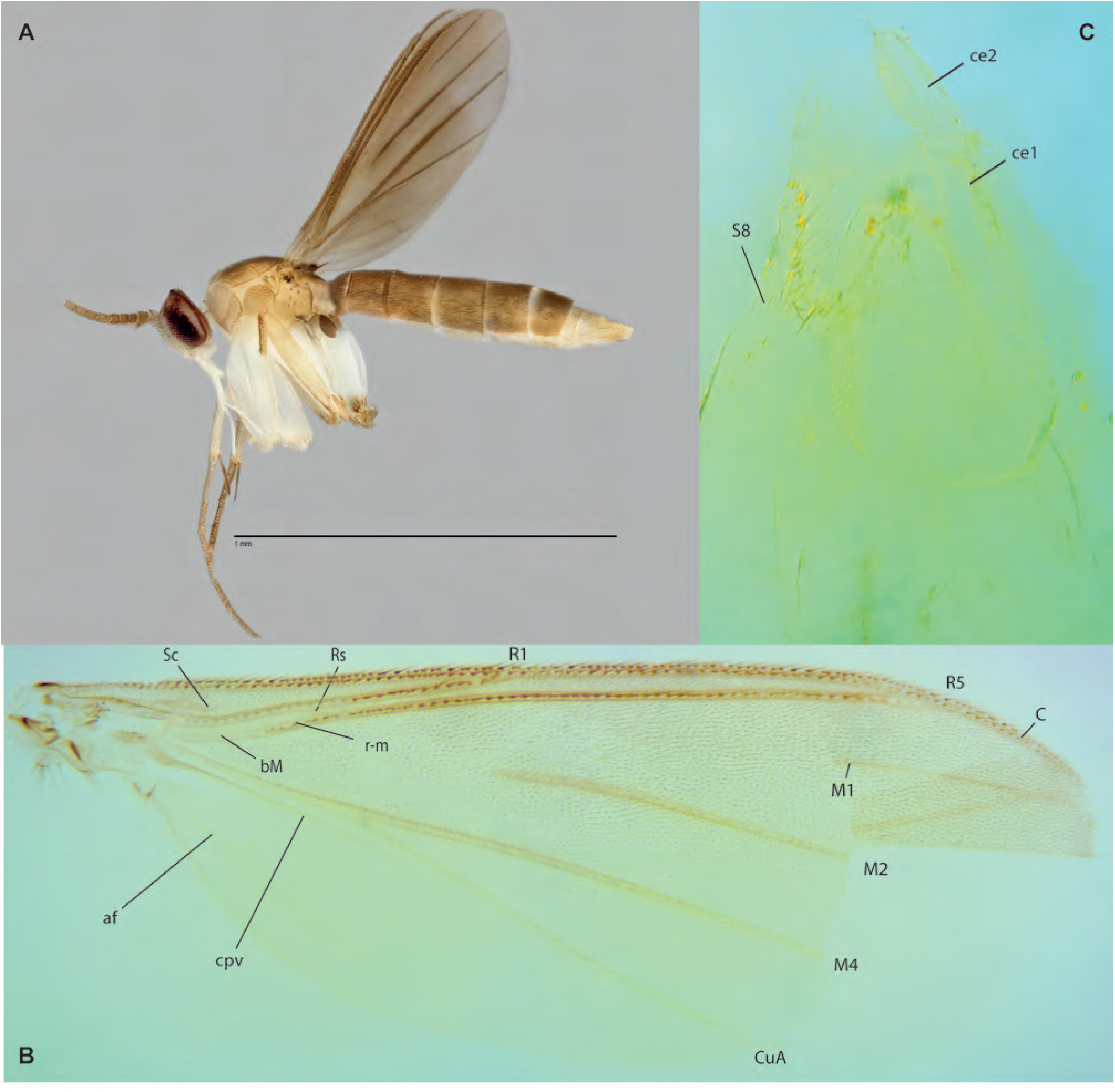
*Manota* sp. B, female ZRCBDP0047826. **A.** Habitus, lateral view. **B.** Wing. **C.** Terminalia, ventrolateral view.

**Figs. 35A-F.**
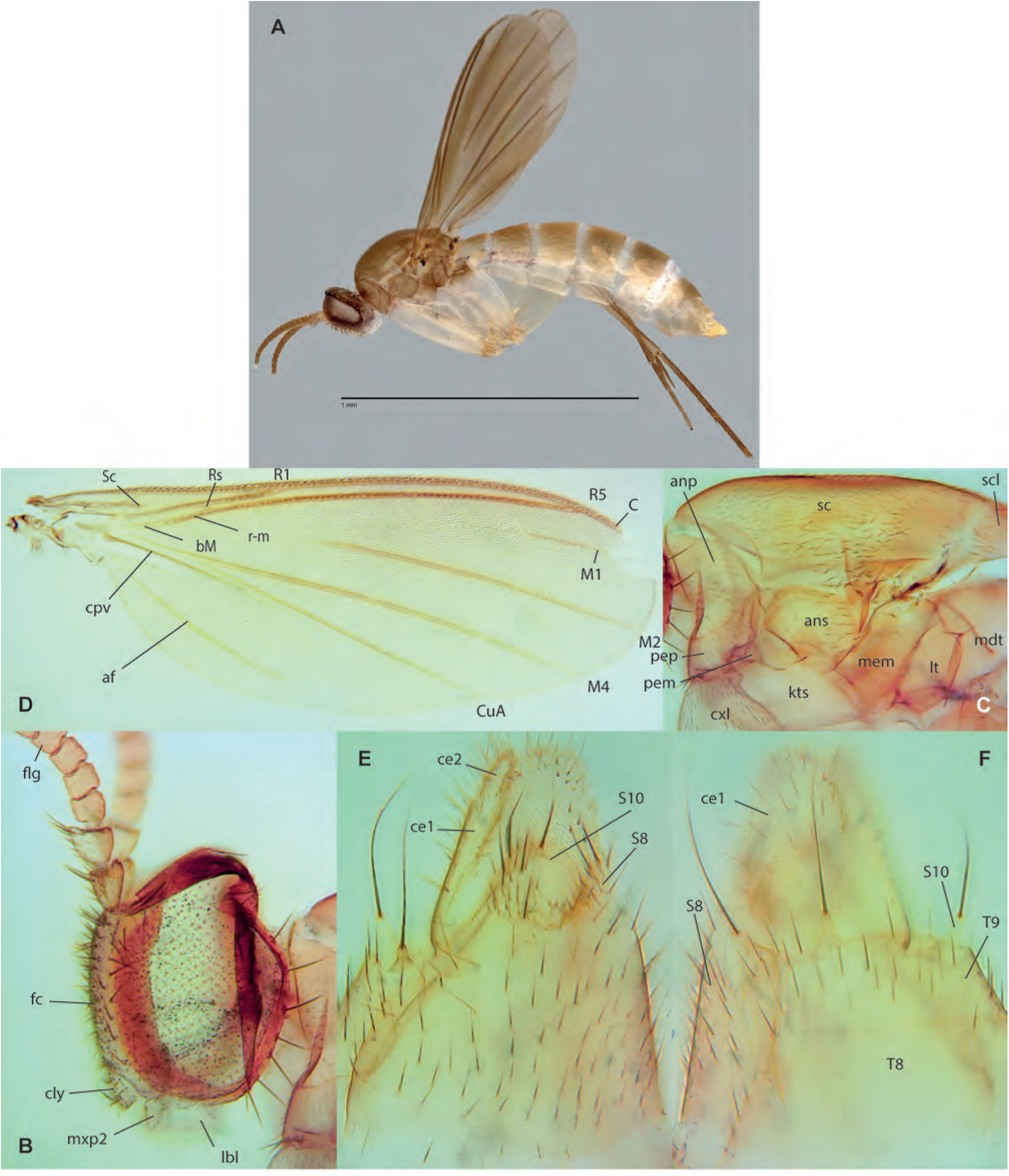
*Manota* sp. C, female ZRCBDP0048300. **A.** Habitus, lateral view. **B.** Head, lateral view. **C.** Thorax, lateral view. **D.** Wing. **E.** Female terminalia, ventral view. **F.** Female terminalia, dorsal view, same.

**Figs. 36A-D.**
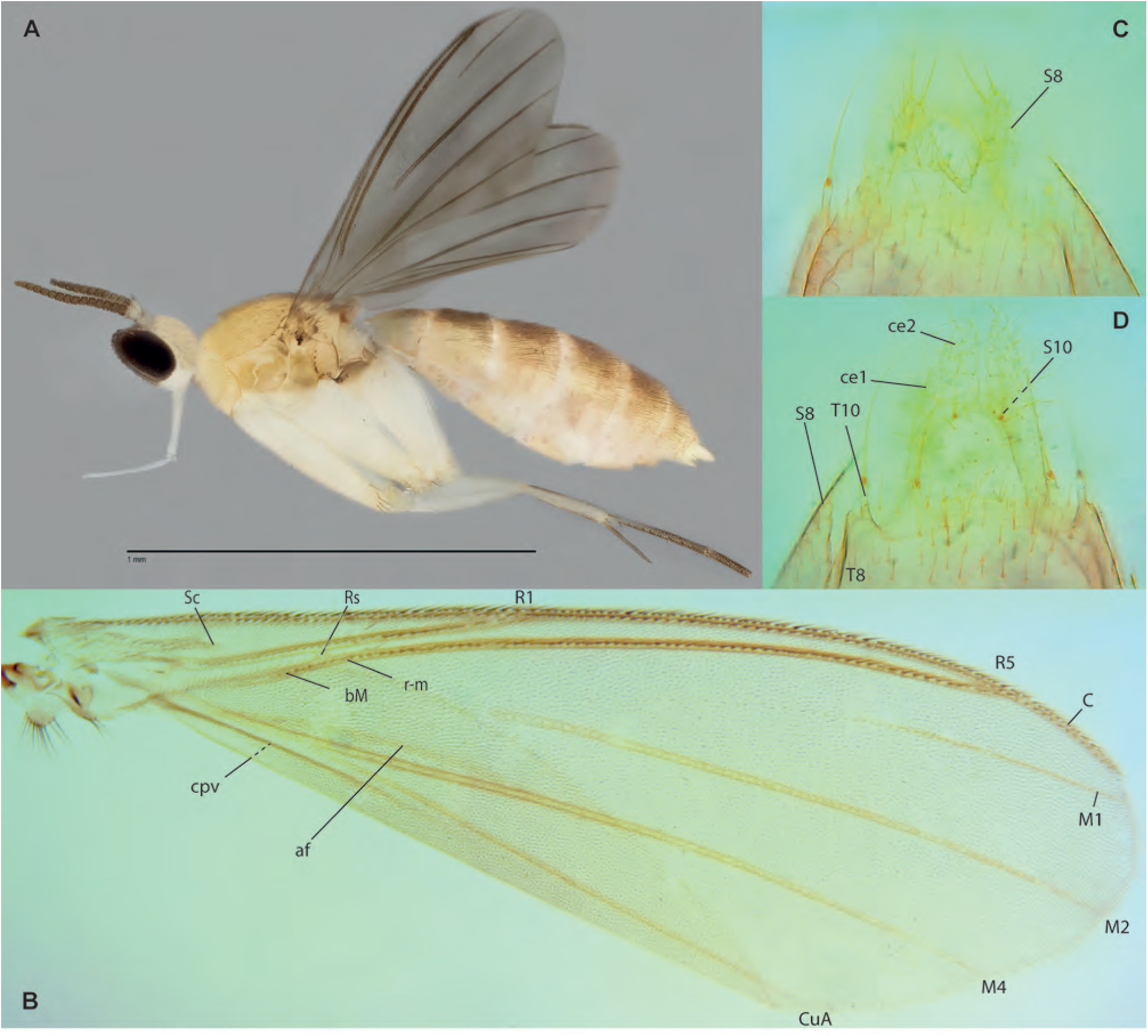
*Manota* sp. D, female ZRCBDP0048676. **A.** Habitus, lateral view. **B.** Wing. **C.** Female terminalia, ventral view. **D.** Female terminalia, dorsal view, same.

**Figs. 37A-C.**
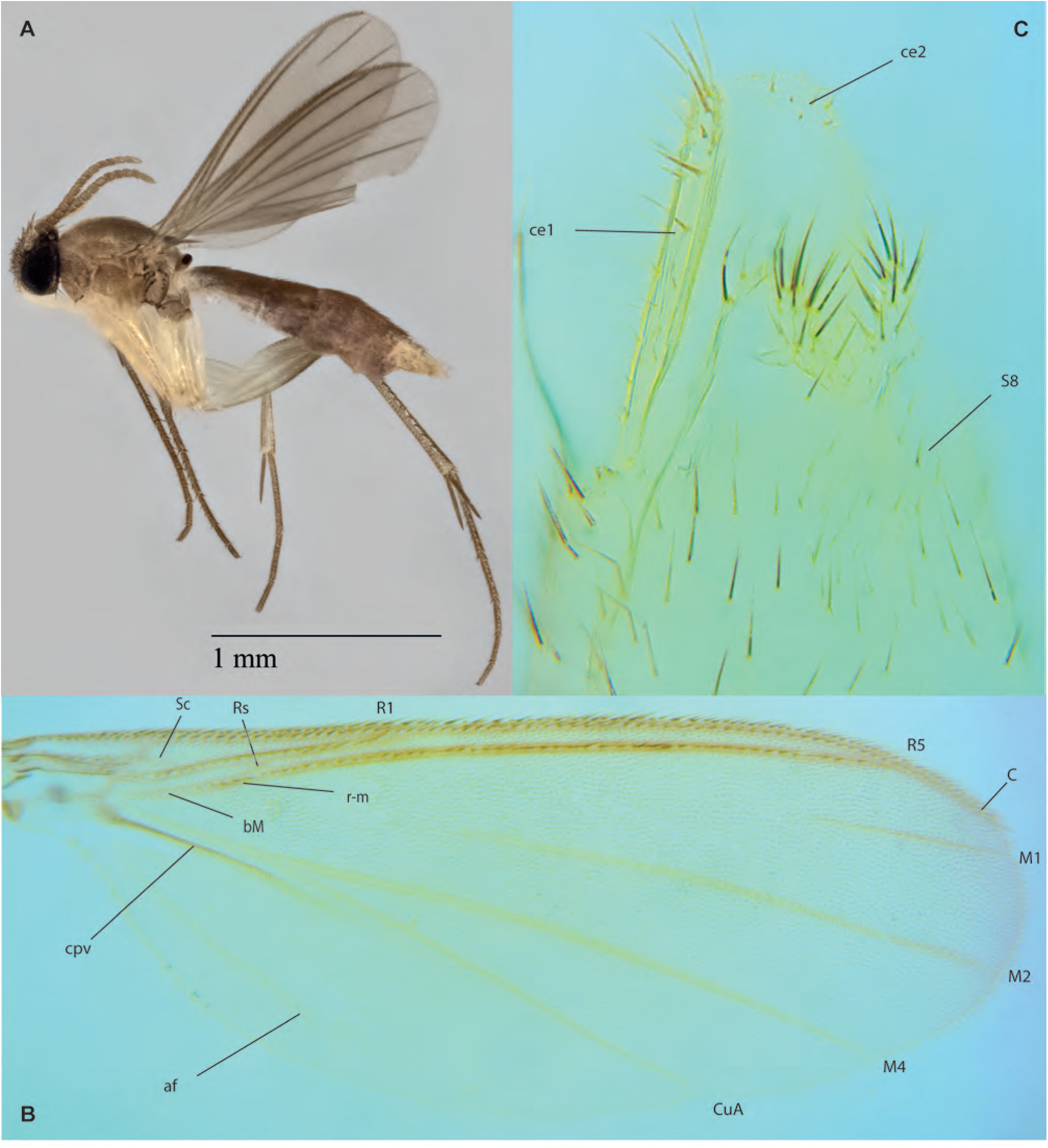
*Manota* sp. E, female. **A.** Habitus, ZRCBDP0133494. **B.** Wing, ZRCBDP0047060. **C.** Terminalia, ventrolateral view, same.

**Figs. 38A-E.**
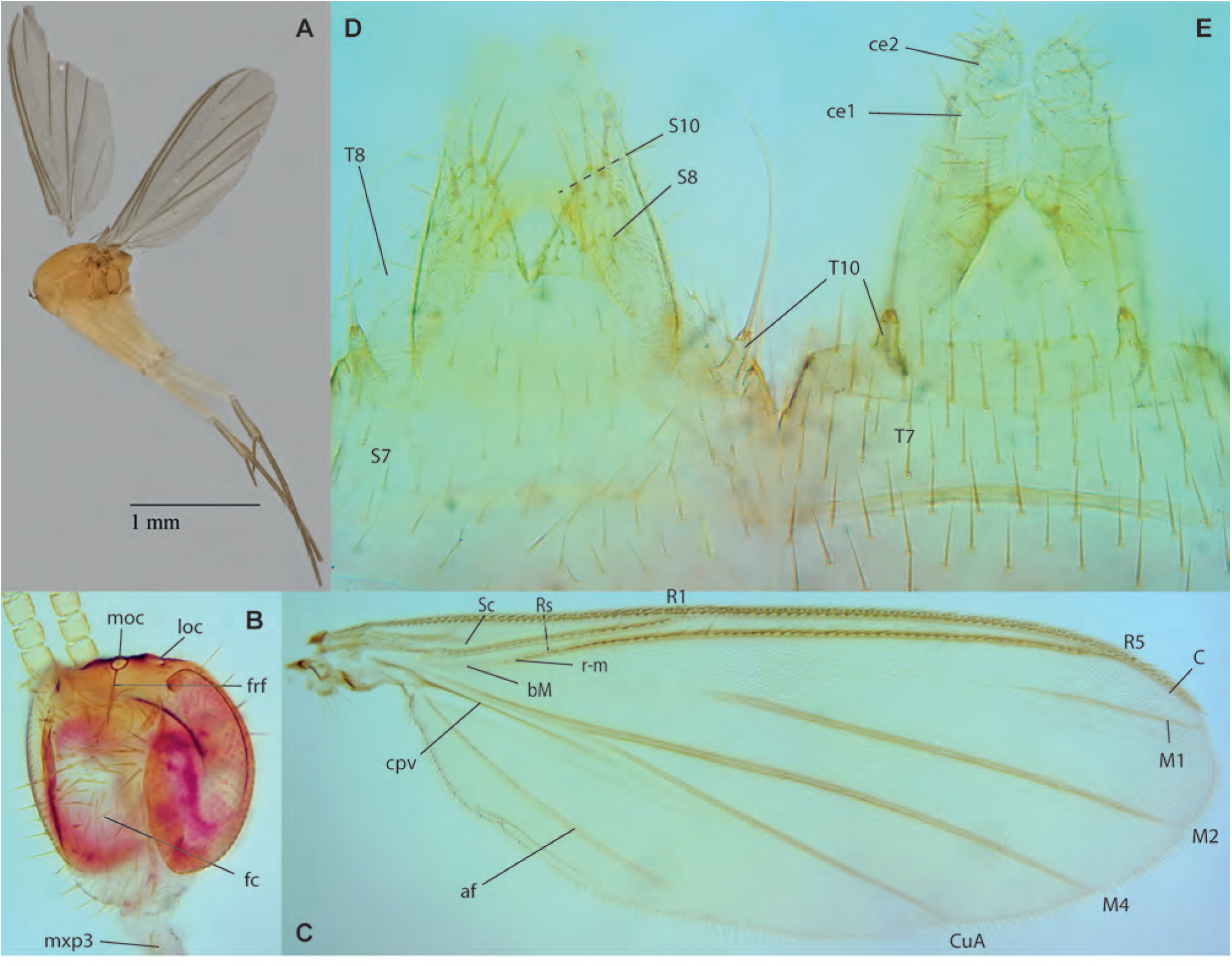
*Manota* sp. F, female. **A.** Habitus, lateral view, ZRCBDP0133440. **B.** Head, frontal view, same. **C.** Wing, ZRCBDP0072692. **D.** Terminalia, ventral view, same. **E.** Terminalia, dorsal view, same.

**Figs. 39A-F.**
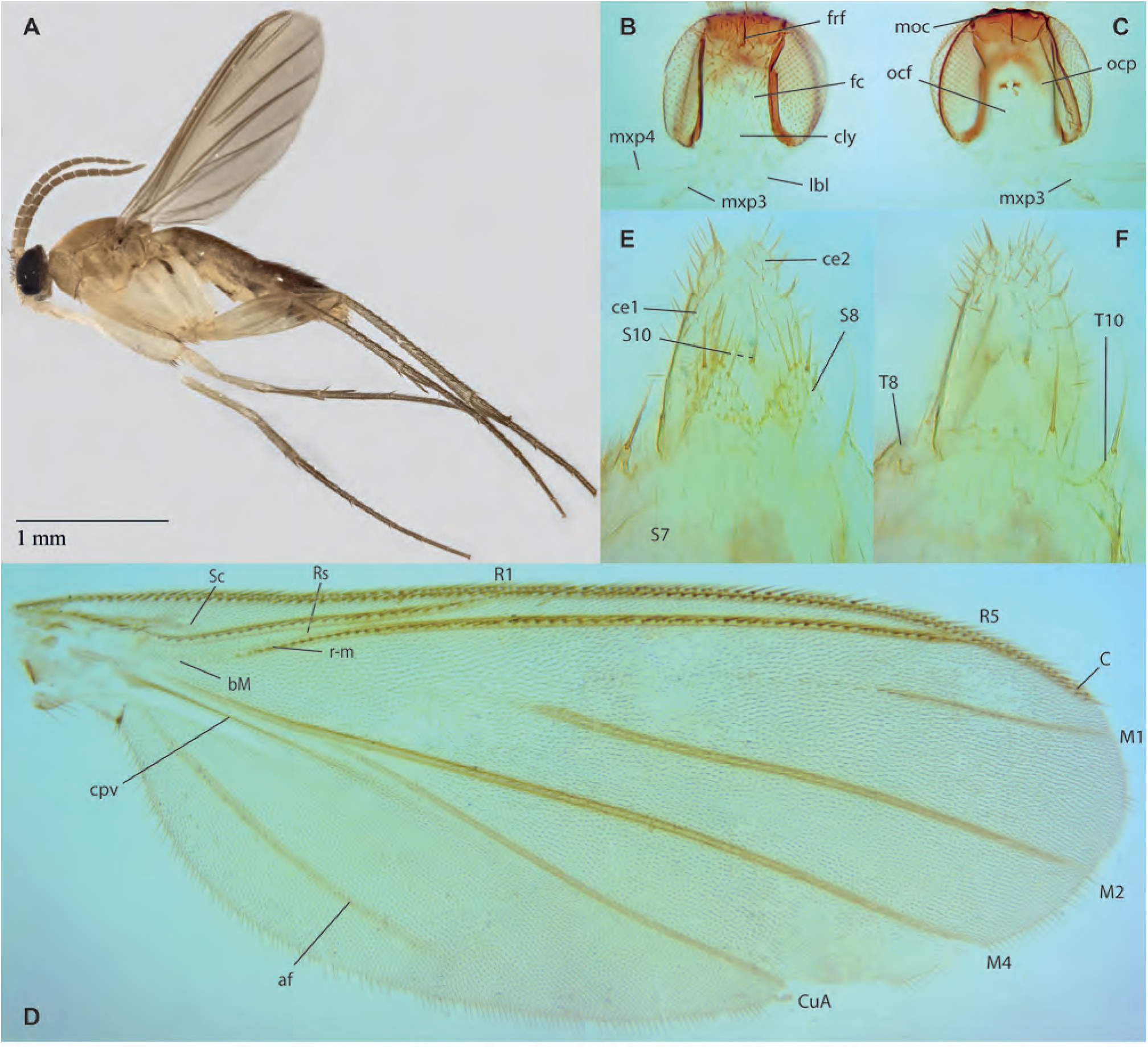
*Manota* sp. G, female. **A.** Habitus, lateral view, ZRCBDP0132831. **B.** Head, frontal view, ZRCBDP0278331. **C.** Head, posterior view, same. **D.** Wing, same. **E.** Terminalia, ventral view, same. **F.** Terminalia, dorsal view, same.

**Figs. 40A-E.**
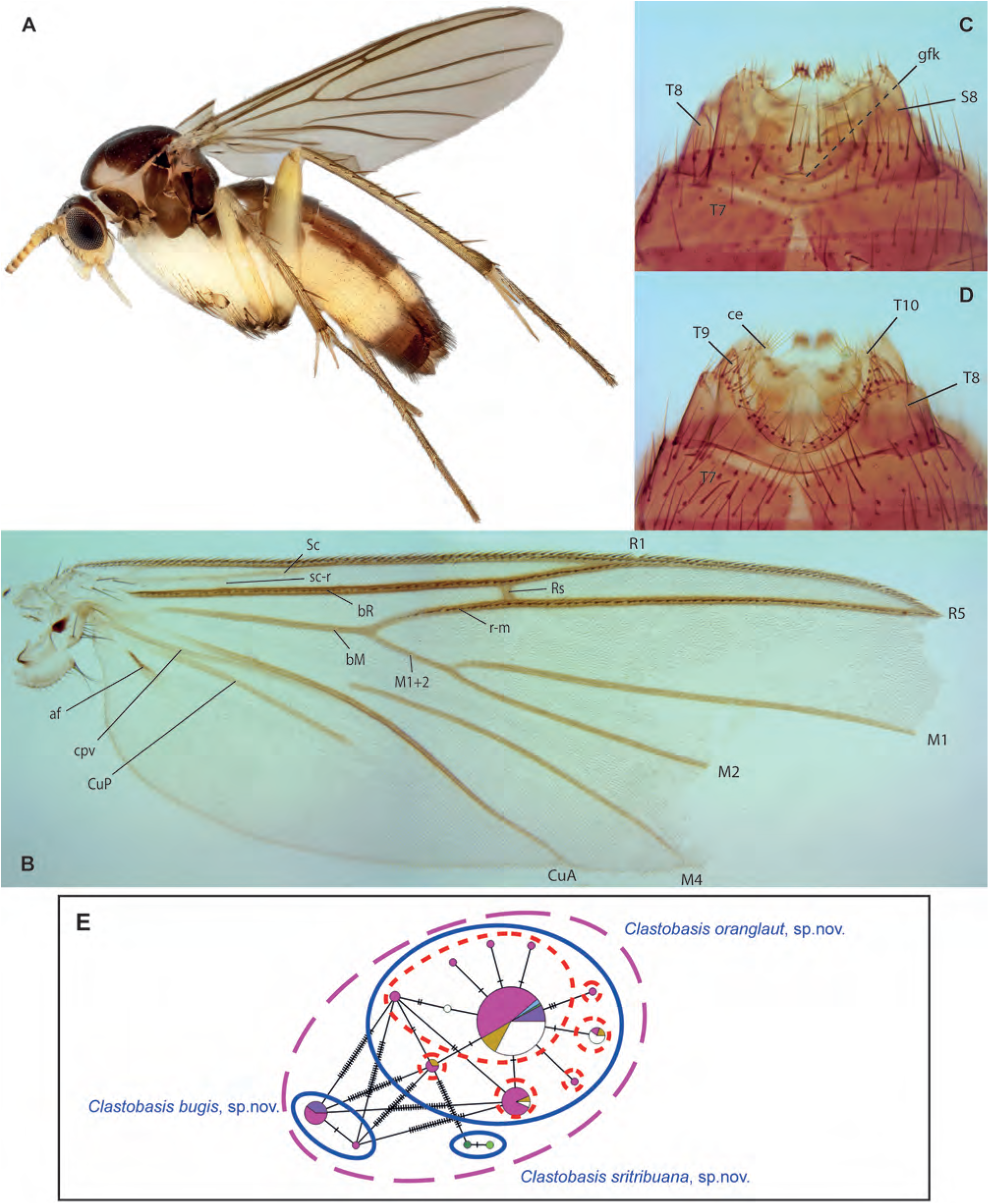
*Clastobasis sritribuana* Amorim & Oliveira, **sp. n.,** female holotype. **A.** Habitus, lateral view. **B.** Wing. **C.** Terminalia, ventral view. **D.** Terminalia, dorsal view. E Haplotype network for *Clastobasis*.

**Figs. 41A-F.**
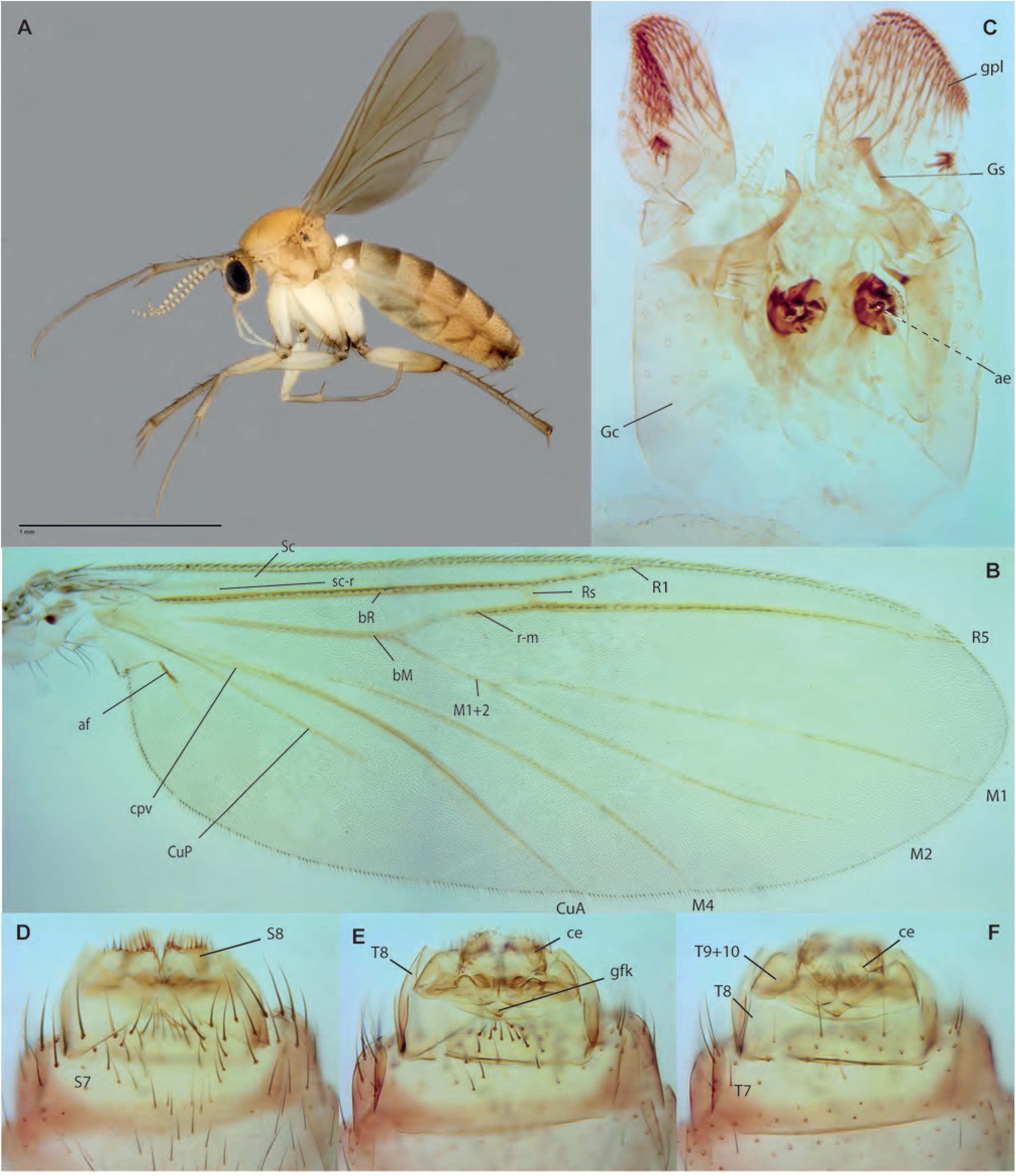
*Clastobasis bugis* Amorim & Oliveira, **sp. n. A.** Habitus, lateral view, female paratype ZRCBDP0048245. **B.** Wing. **C.** Male terminalia, ventral view, holotype. **D.** Female terminalia, ventral view, paratype ZRCBDP0048242. **E.** Female terminalia, mid-section, same. **F.** Female terminalia, dorsal view, same.

**Figs. 42A-F.**
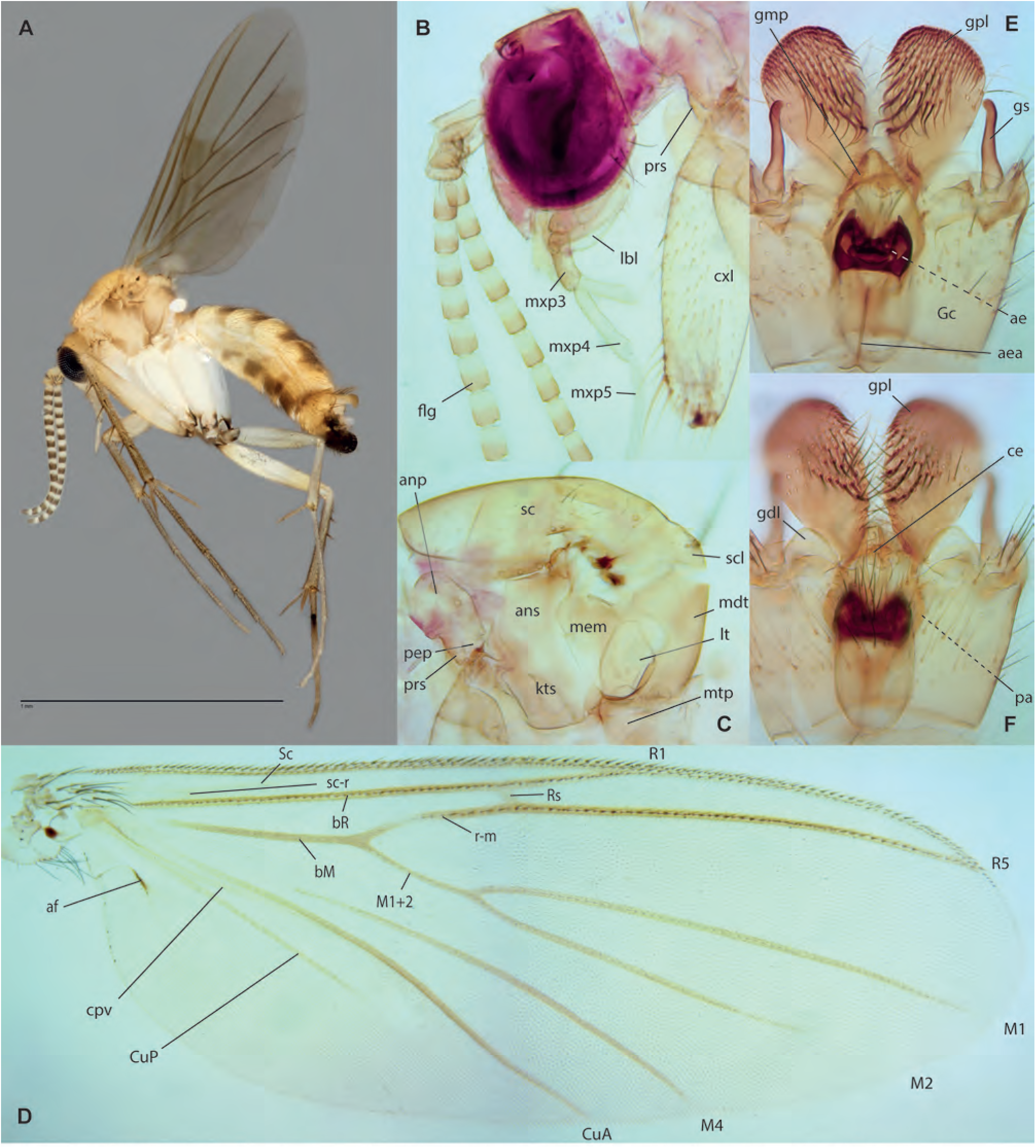
*Clastobasis oranglaut* Amorim & Oliveira, **sp. n. A.** Habitus, lateral view, male paratype ZRCBDP0049312. **B.** Head, lateral view, male holotype. **C.** Thorax, lateral view, same. **D.** Wing, same. **E.** Male terminalia, ventral view, same. **F.** Male terminalia, dorsal view, same.

**Figs. 43A-B.**
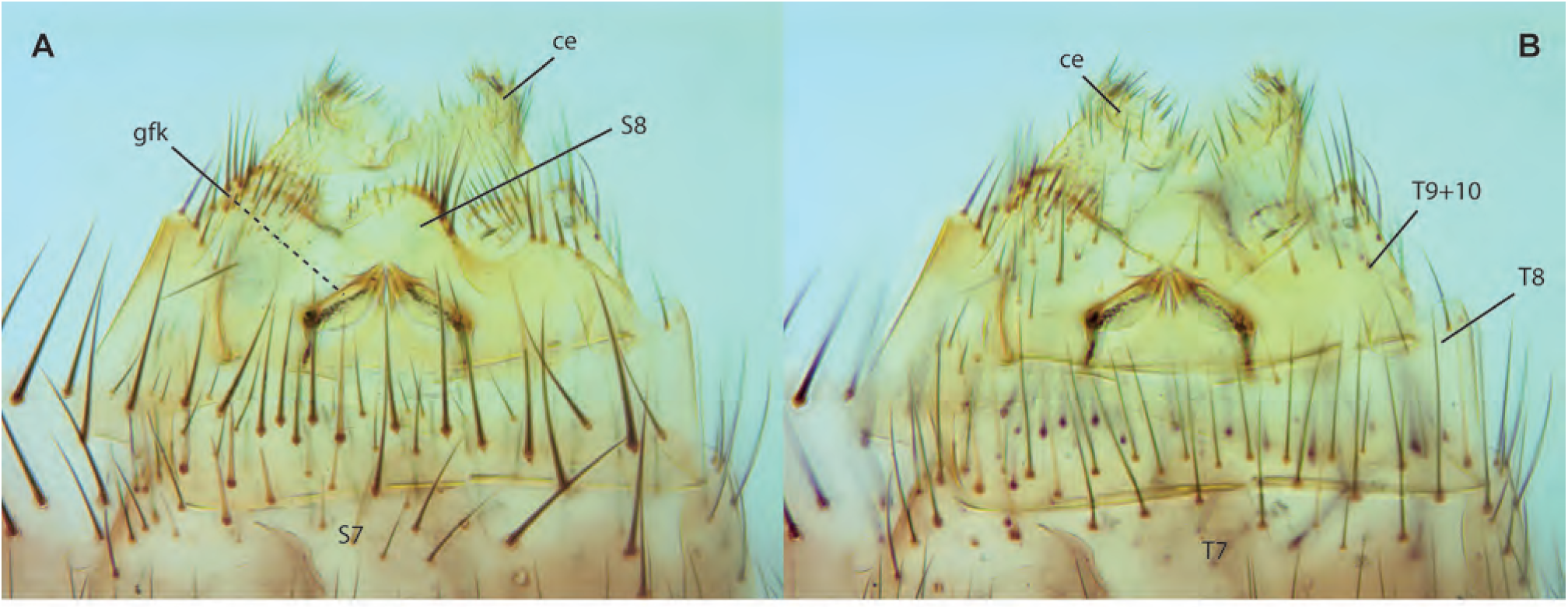
*Clastobasis oranglaut* Amorim & Oliveira, **sp. n.**, female terminalia, paratype ZRCBDP0049336. **A.** Ventral view, same. **B.** Dorsal view. **Gnoristinae**

## Gnoristinae

**Figs. 44A-D.**
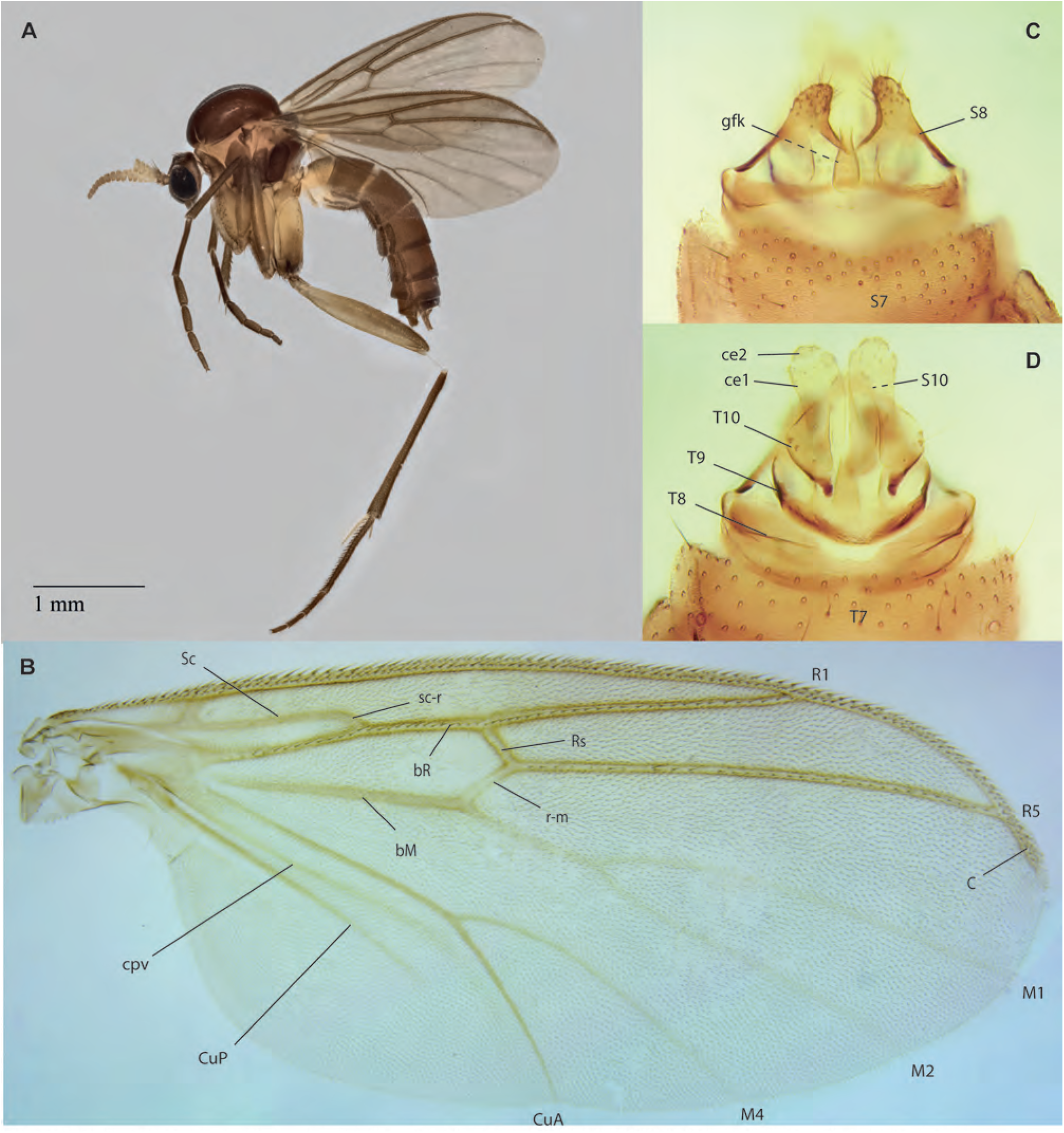
*Chalastonepsia* sp. **A.** Habitus, lateral view, female ZRCBDP0143086. **B.** Wing, female. **C.** Terminalia, ventral view, same. **D.** Terminalia, dorsal view, same.

**Figs. 45A-C.**
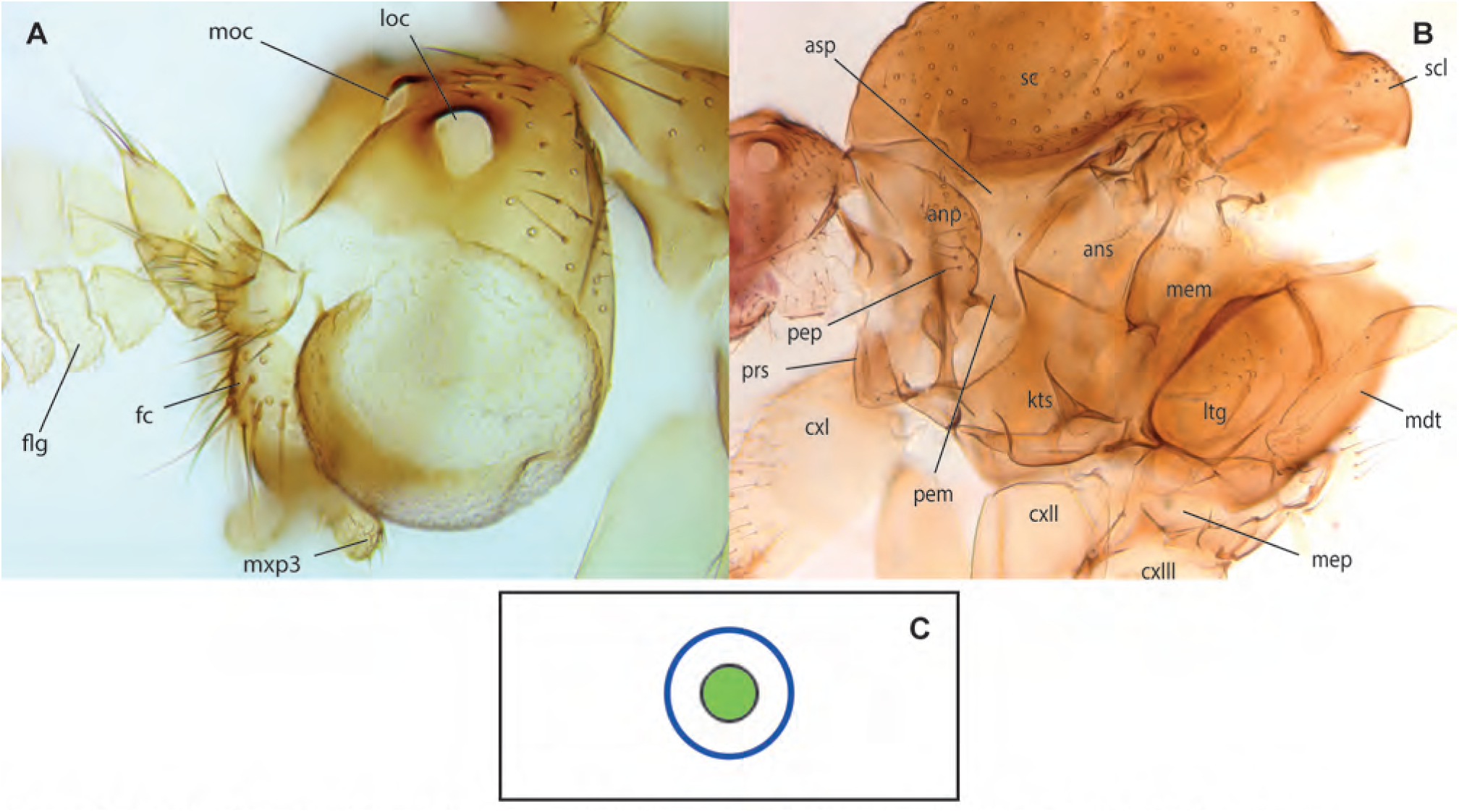
*Chalastonepsia* sp. **A.** Head, laterodorsal view, female. **B.** Thorax, female ZRCBDP0047804. **C.** Haplotype network for *Chalastonepsia*.

**Figs. 46A-D.**
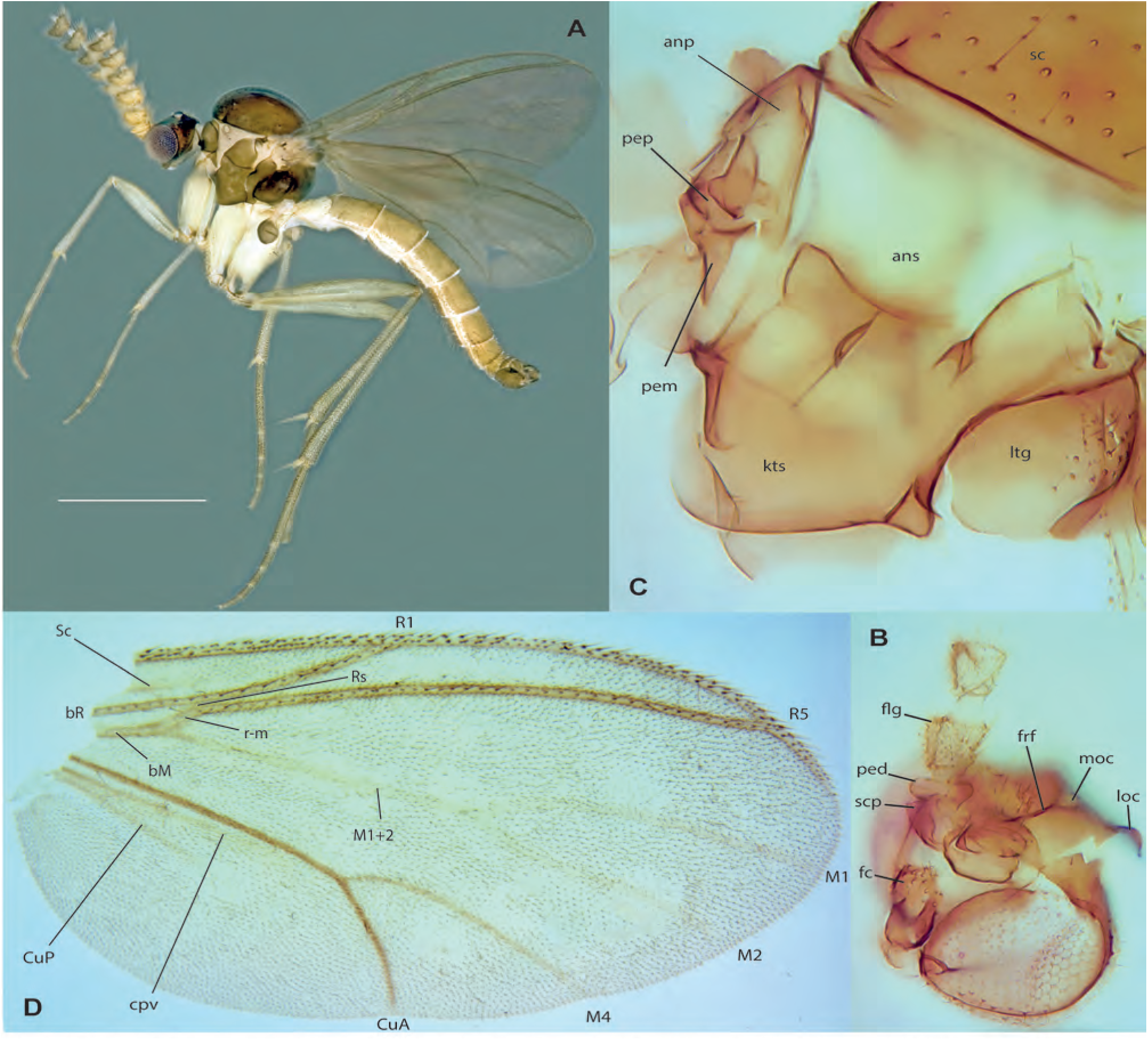
*Metanepsia malaysiana* Kallweit. **A.** Habitus, lateral view, male, ZRCBDP0048531. **B.** Head, dorsolateral view, male, ZRCBDP0048680. **C.** Thorax, lateral view, same. **D.** Wing, same.

**Figs. 47A-F.**
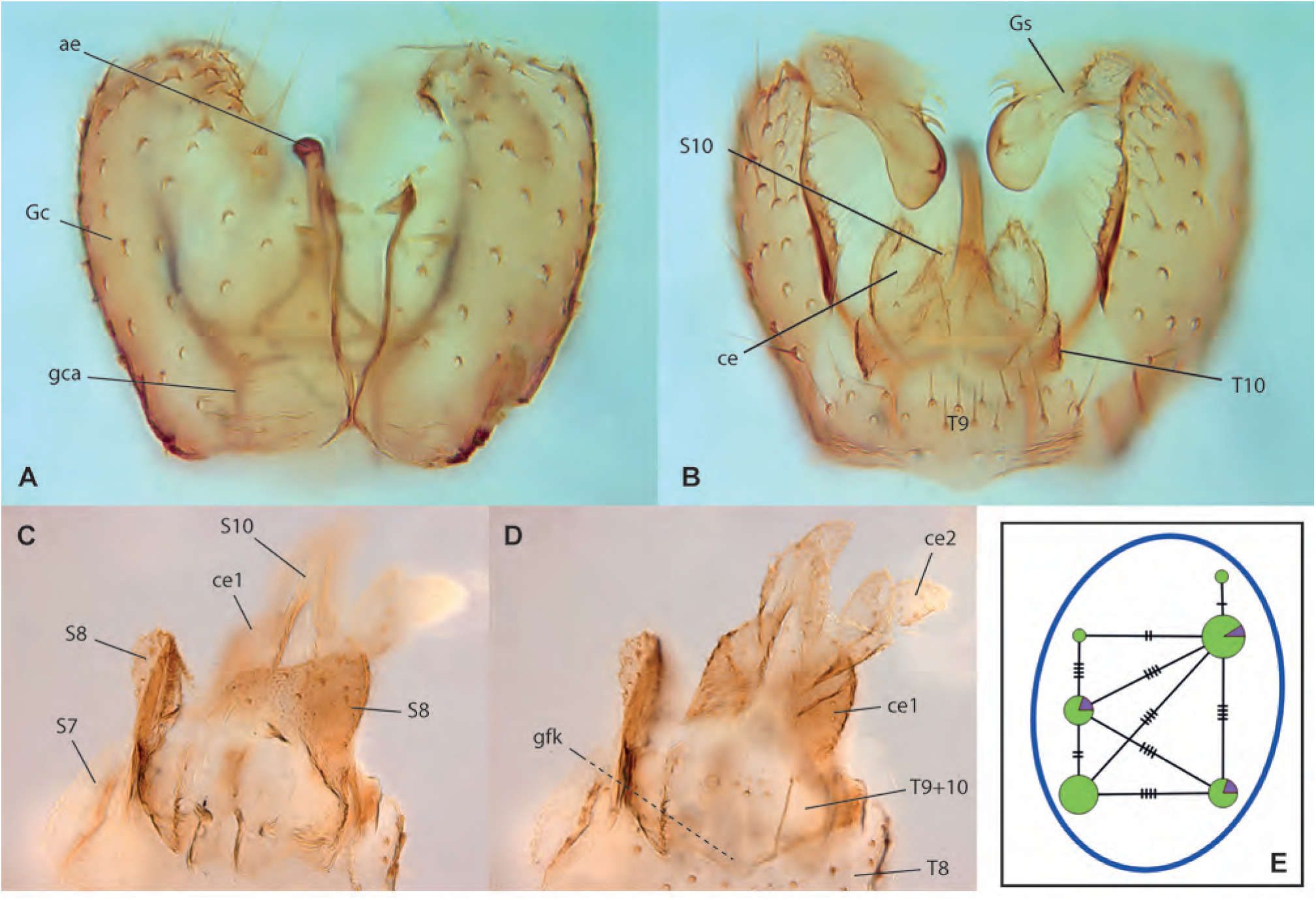
*Metanepsia malaysiana* Kallweit. **A.** Male terminalia, ventral view, ZRCBDP0048680. **C.** Male terminalia, dorsal view, same. **D.** Female terminalia, ventral view, ZRCBDP0048869. **E.** Female terminalia, dorsal view, same. **F.** Haplotype network for *Metanepsia*. **Mycomyinae**

## Mycomyinae

**Figs. 48A-I.**
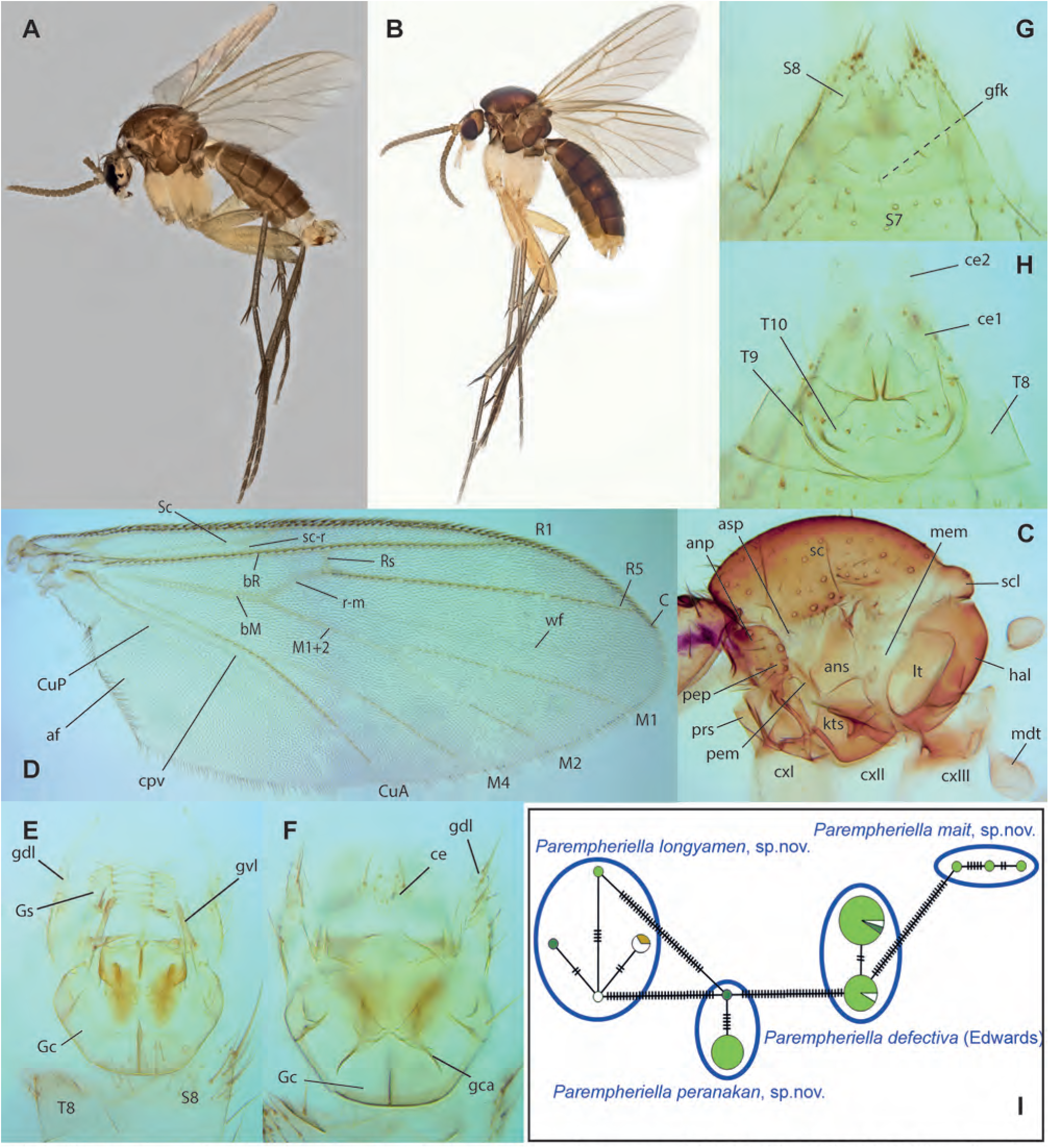
*Parempheriella defectiva* Edwards. **A.** Habitus, lateral view, male, ZRCBDP0155005. **B.** Habitus, lateral view, female, ZRCBDP0048556. **C.** Thorax, lateral view, male, ZRCBDP0048828. **D.** Wing, same. **E.** Male terminalia, ventral view, same. **F.** Male terminalia, dorsal view, same. **G.** Female terminalia, ventral view, ZRCBDP0048556. **H.** Female terminalia, dorsal view, same. **I.** Haplotype network for *Parempheriella*.

**Figs. 49A-D.**
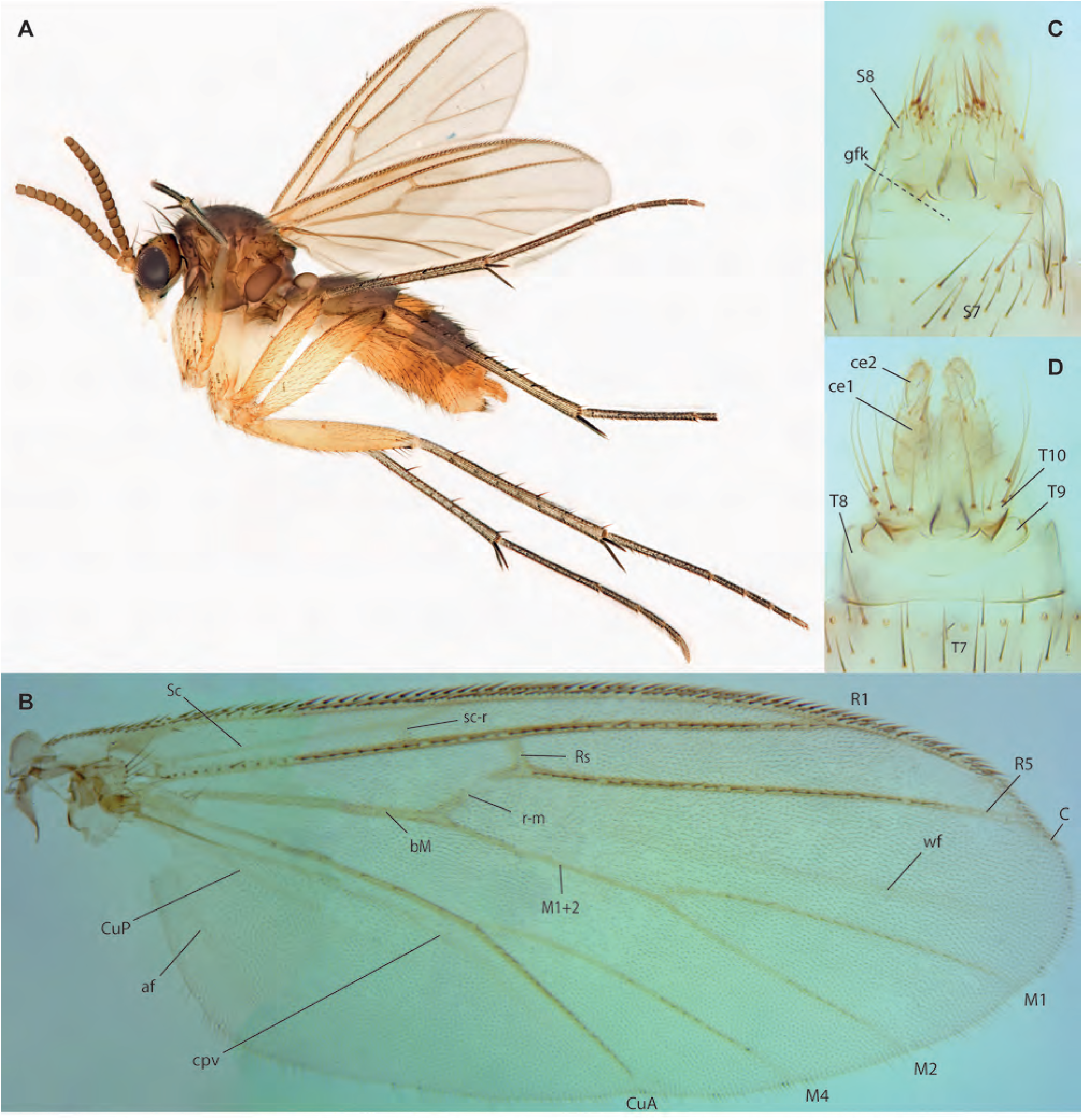
*Parempheriella mait* Amorim & Oliveira, **sp. n.,** female paratype ZRCBDP0048559. **A.** Habitus, lateral view. **B.** Wing. **C.** Terminalia, ventral view. **D.** Terminalia, dorsal view.

**Figs. 50A-D.**
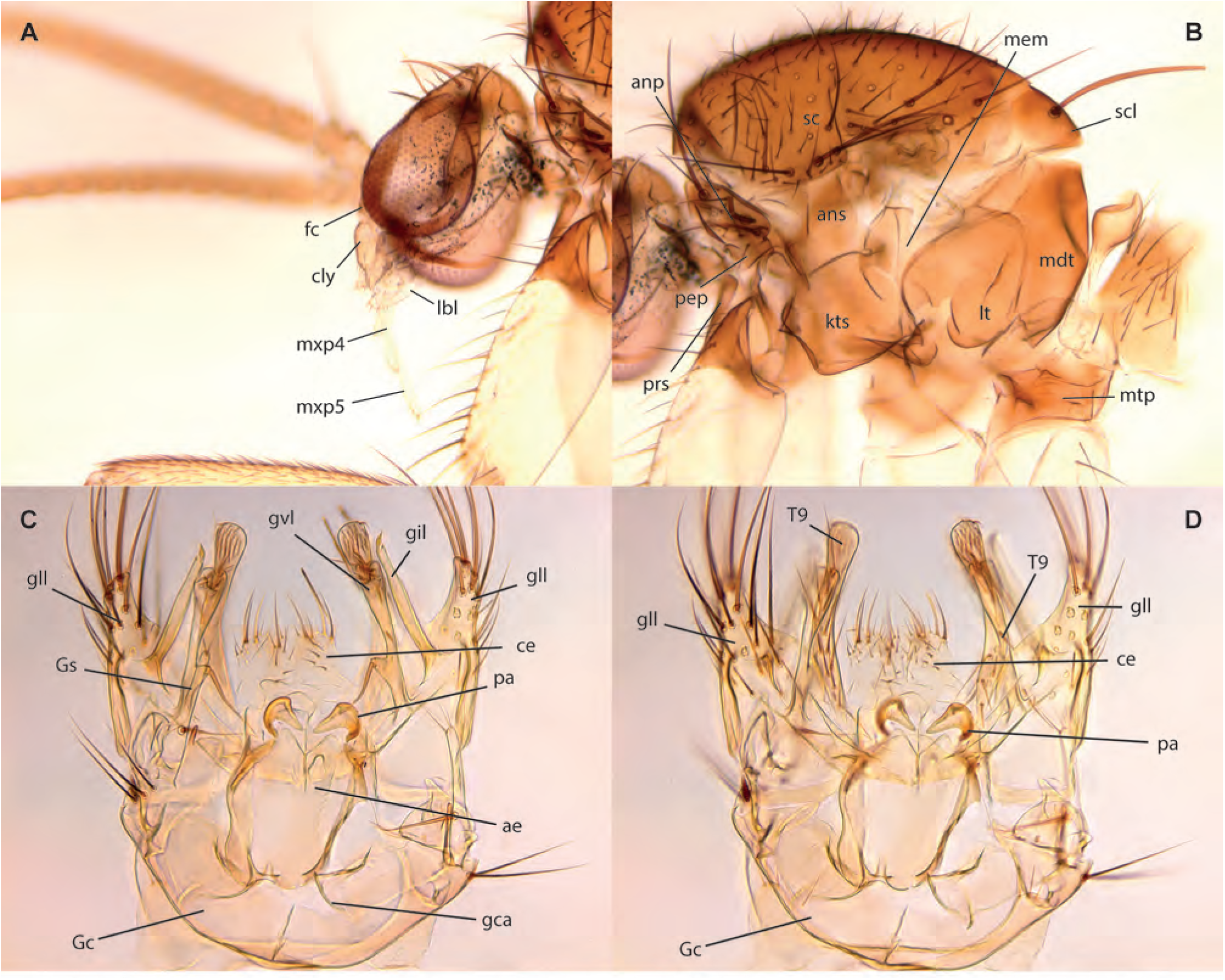
*Parempheriella mait* Amorim & Oliveira, **sp. n.,** male holotype. **A.** Head, lateral view. **B.** Thorax, lateral view. **C.** Terminalia, ventral view. **D.** Terminalia, dorsal view.

**Figs. 51A-G.**
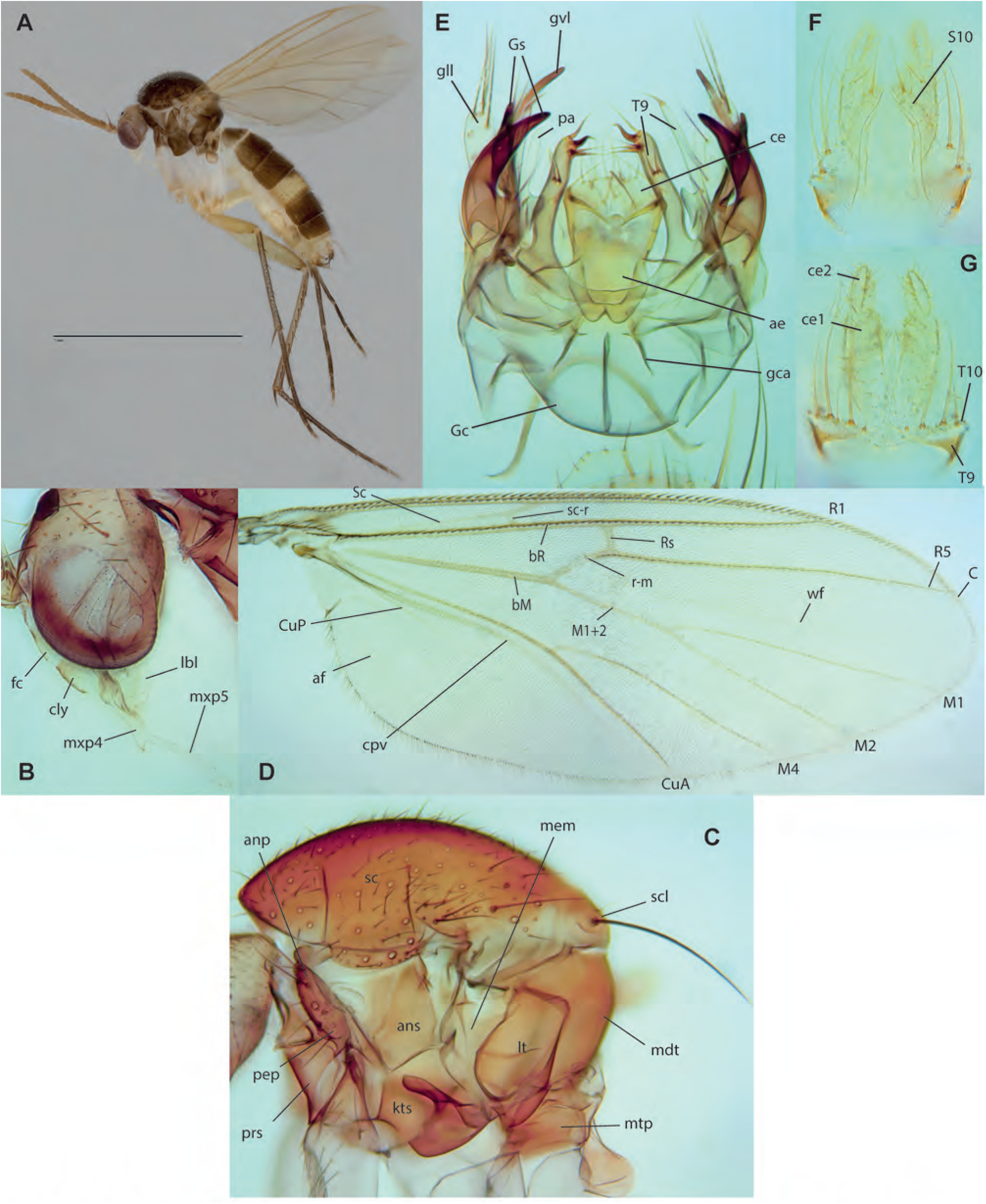
*Parempheriella longyamen* Amorim & Oliveira, **sp. n. A.** Habitus, lateral view, male holotype. **B.** Head, lateral view, same. . **C.** Thorax, lateral view, same. **D.** Wing, same. **E.** Male terminalia, ventral view, same. **F.** Female terminalia, dorsal view, paratype ZRCBDP0072675. **G.** Female terminalia, ventral view, same.

**Figs. 52A-E.**
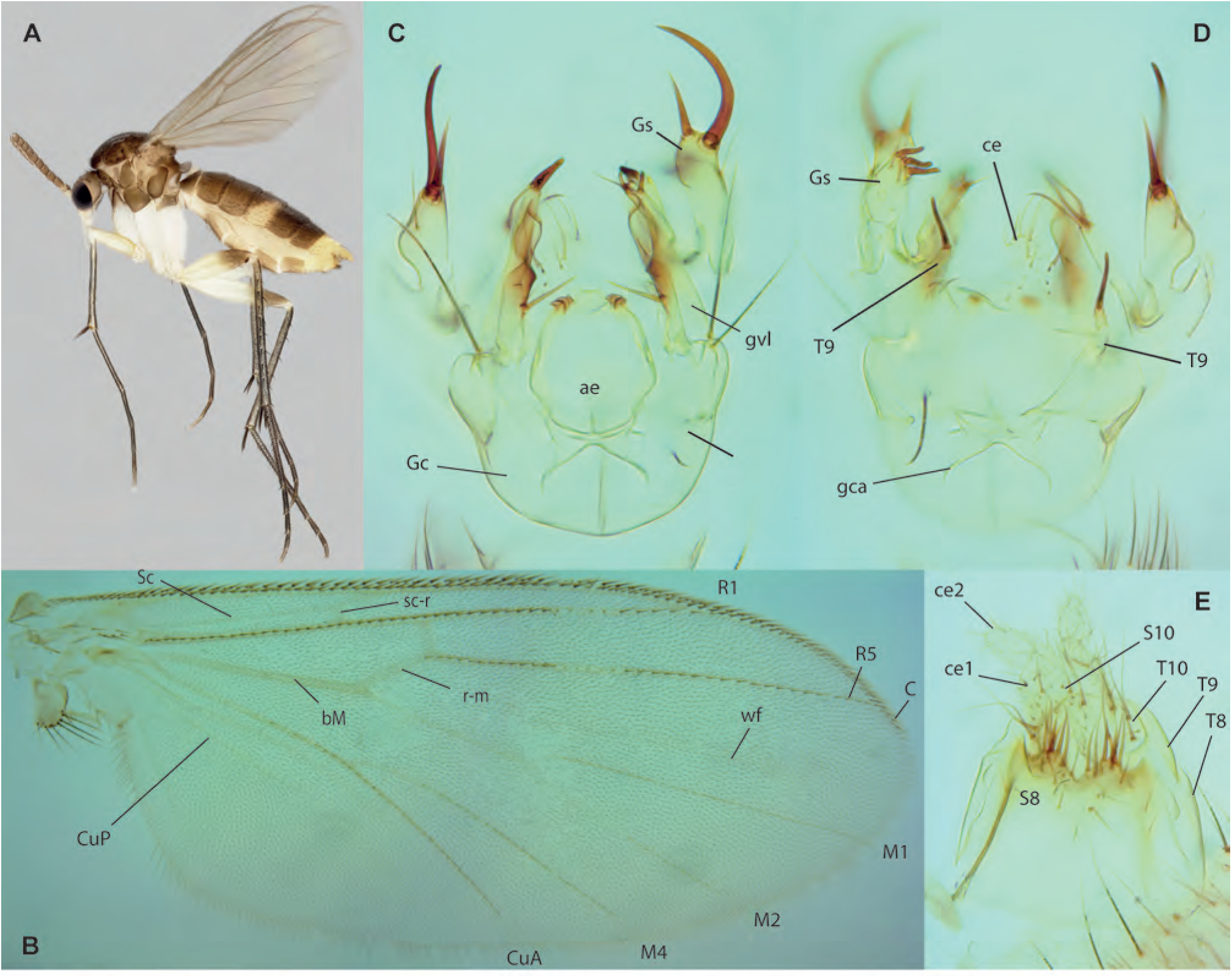
*Parempheriella peranakan* Amorim & Oliveira, **sp. n. A.** Habitus, lateral view, female, paratype ZRCBDP0049227. **B.** Wing, male, paratype ZRCBDP0049222. **C.** Male terminalia, ventral view, same. **D.** Male terminalia, dorsal view, same. **E.** Female terminalia, paratype ZRCBDP0133974.

**Fig. 53.**
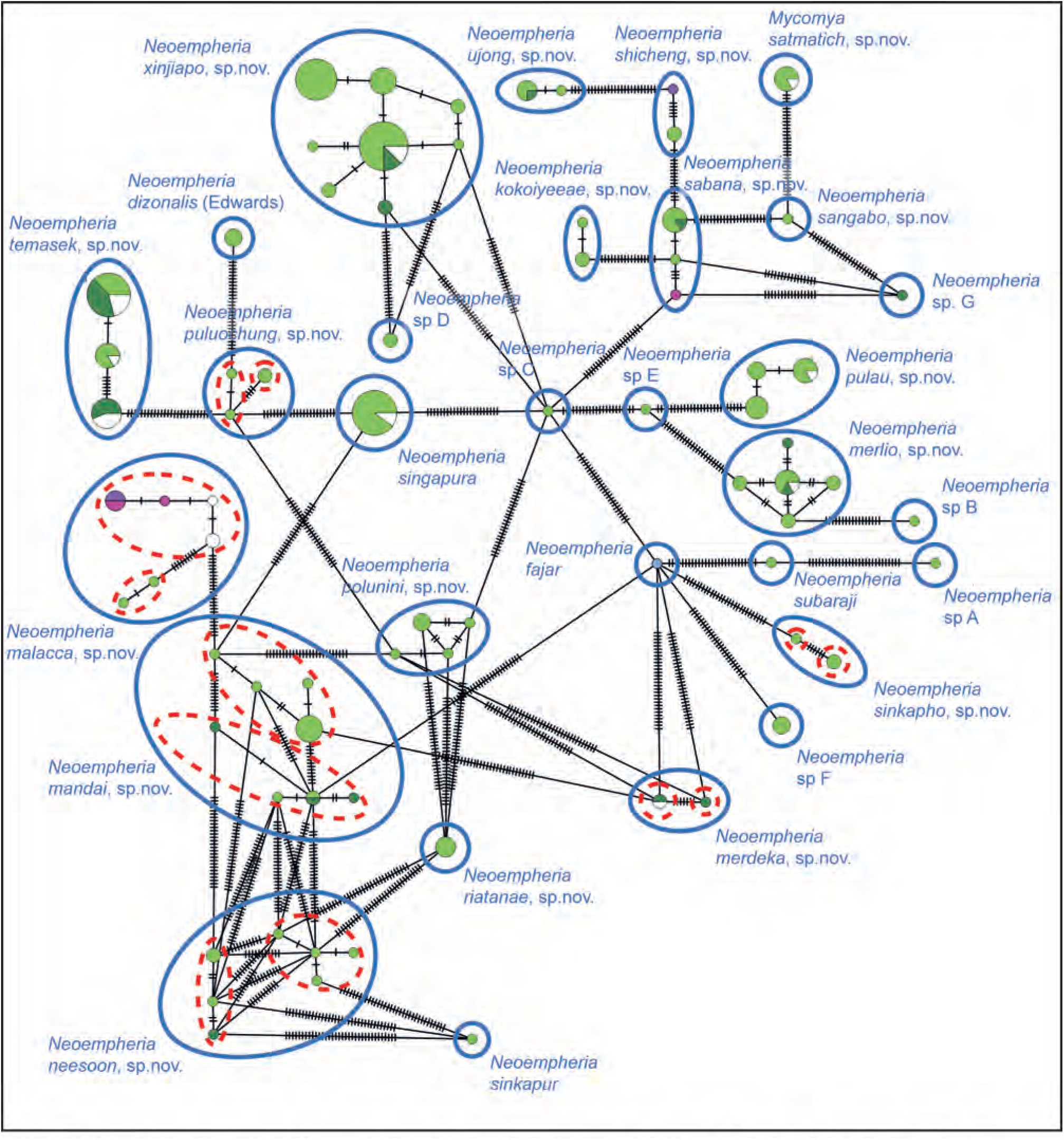
Haplotype network for *Neoempheria*. Species delimitation methods used: methods congruent with morphology unless conflict specified OC: 2-5% ABGD: P=0.001-P=0.06 PTP.

**Figs. 54A-D.**
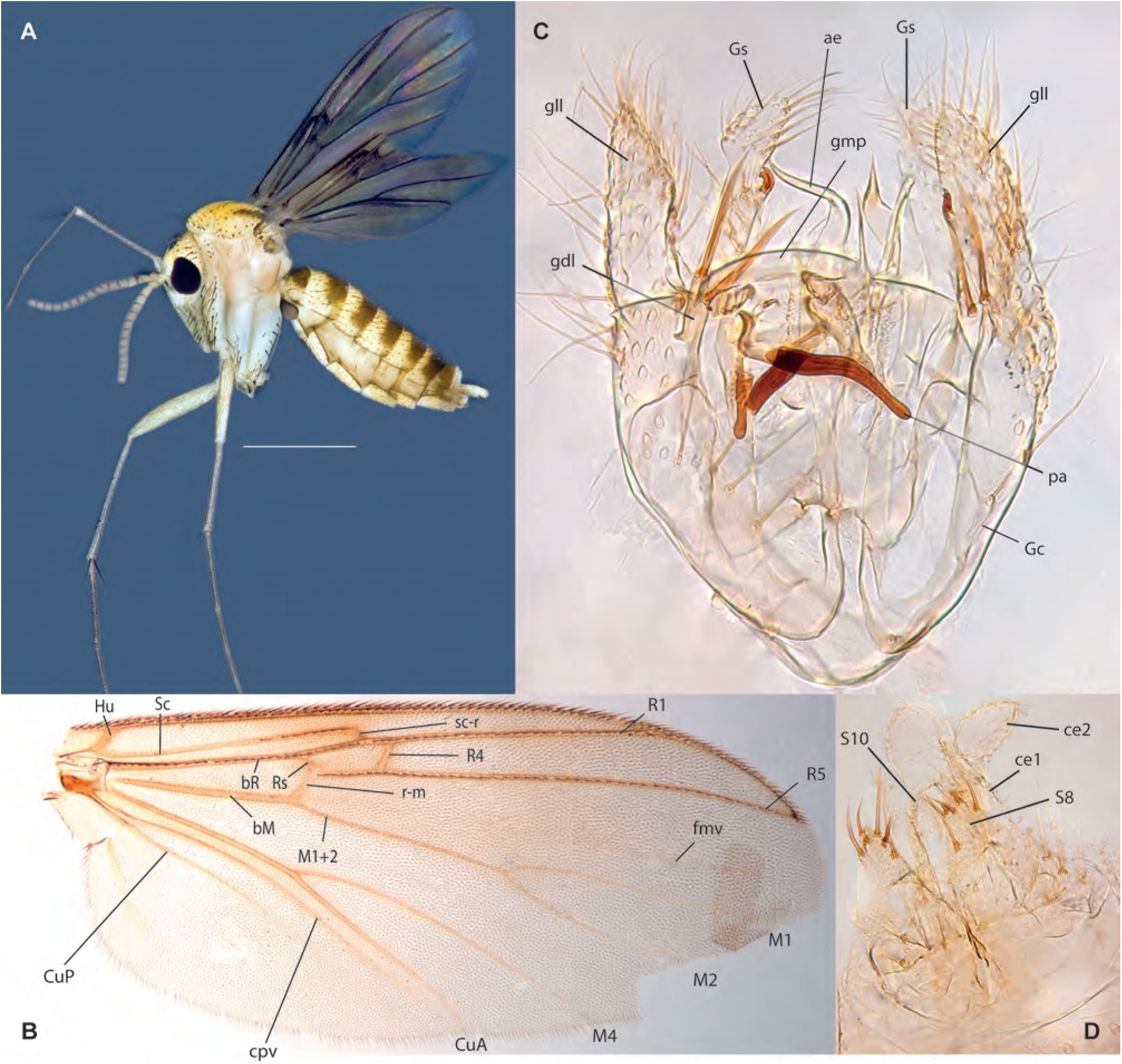
*Mycomya sachmatich* Amorim & Oliveira, **sp. n. A.** Habitus, lateral view, female, ZRCBDP0048498. **B.** Wing, male holotype. **C.** Male terminalia, ventral view, same. **D.** Female terminalia, ventral view, ZRCBDP48498.

**Figs. 55A-C.**
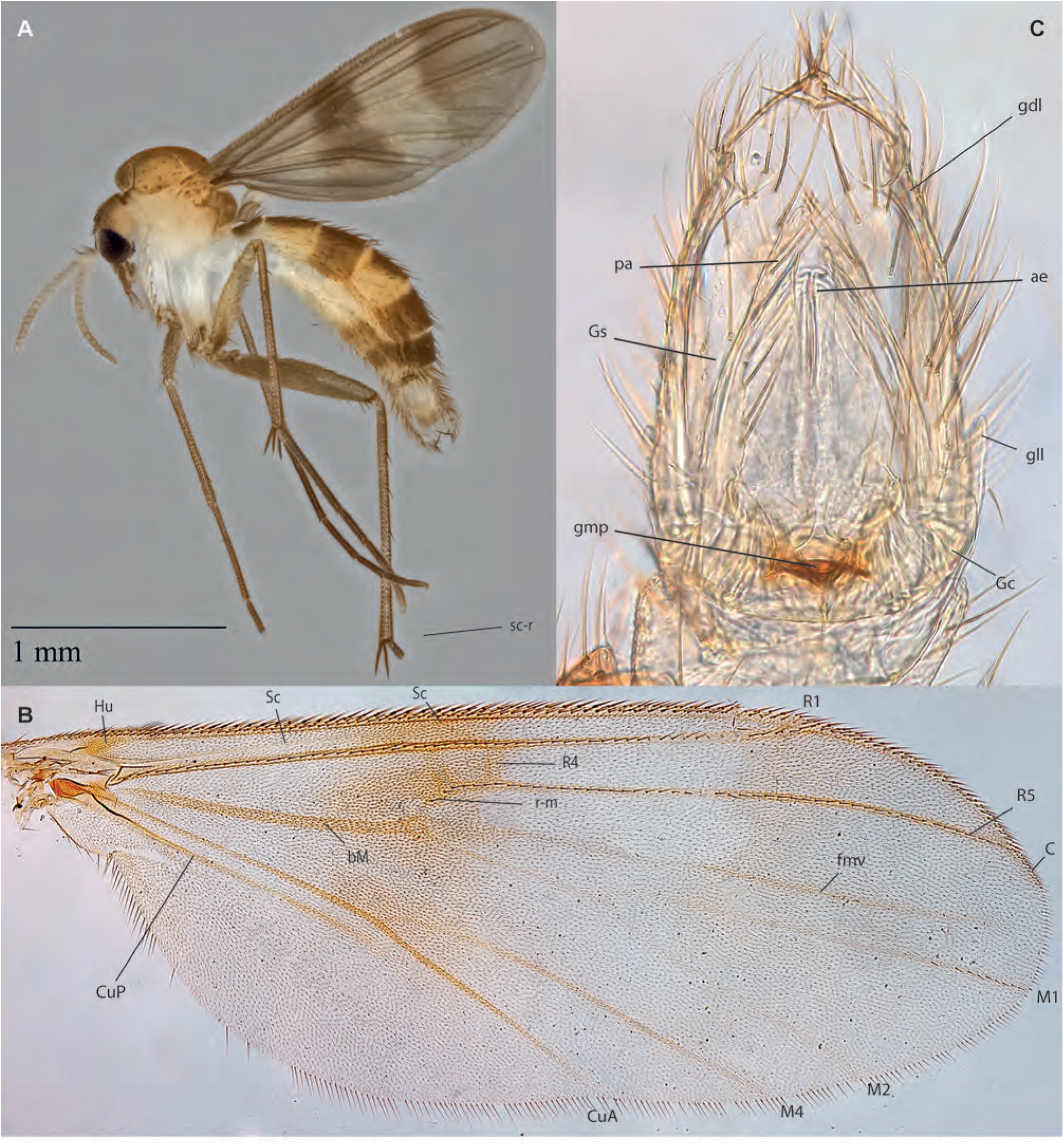
*Neoempheria merlio* Amorim & Oliveira, **sp. n. A.** Habitus, lateral view, male, ZRCBDP0155067. **B.** Wing, female, ZRCBDP0049192. **C.** Male terminalia, ventral view, holotype.

**Figs. 56A-D.**
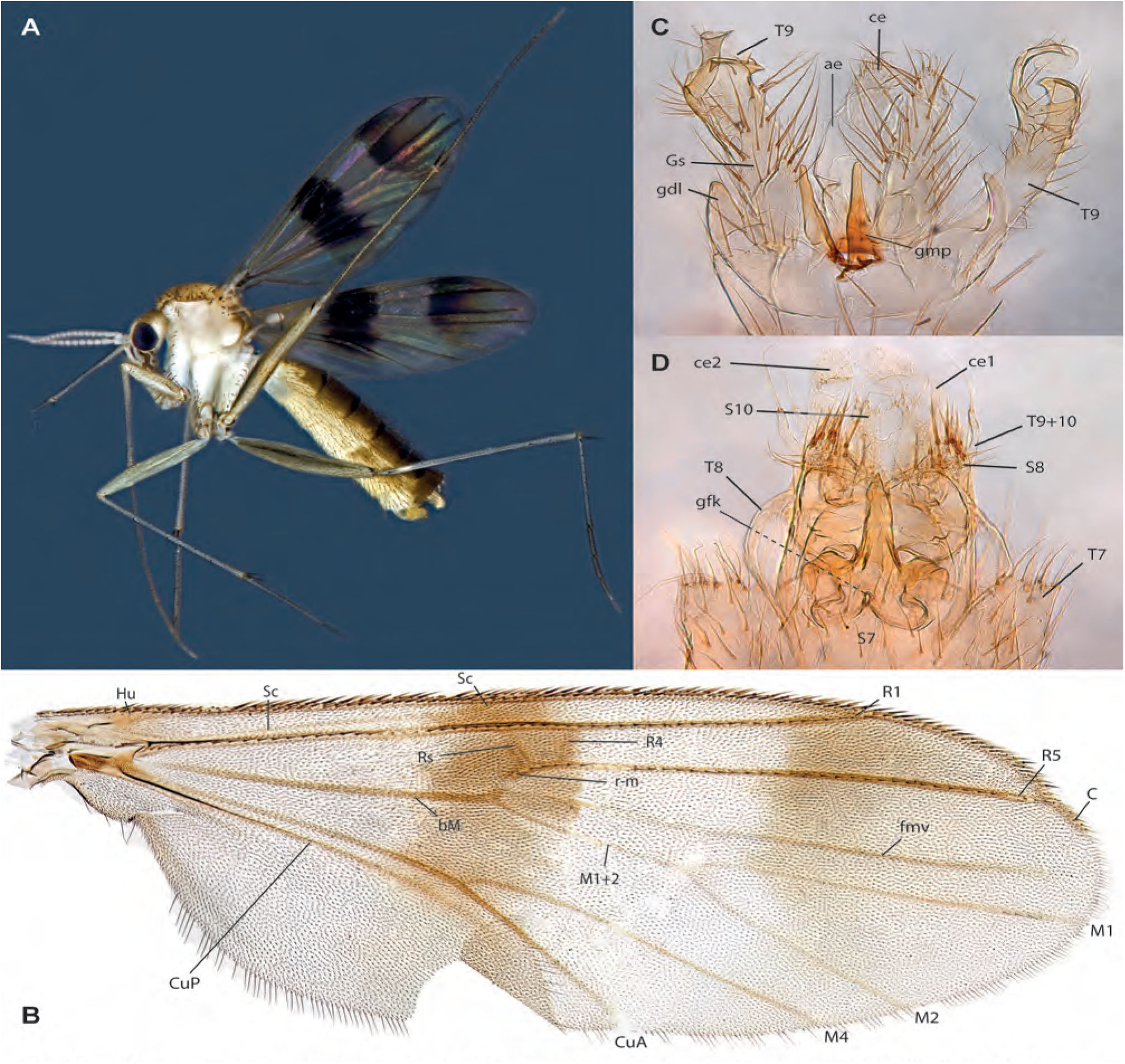
*Neoempheria sabana* Amorim & Oliveira, **sp. n. A.** Habitus, lateral view, female, ZRCBDP0048495. **B.** Wing, male holotype. **C.** Male terminalia, ventral view, same. **D.** Female terminalia, ventral view, female ZRCBDP0047068.

**Figs. 57A-D.**
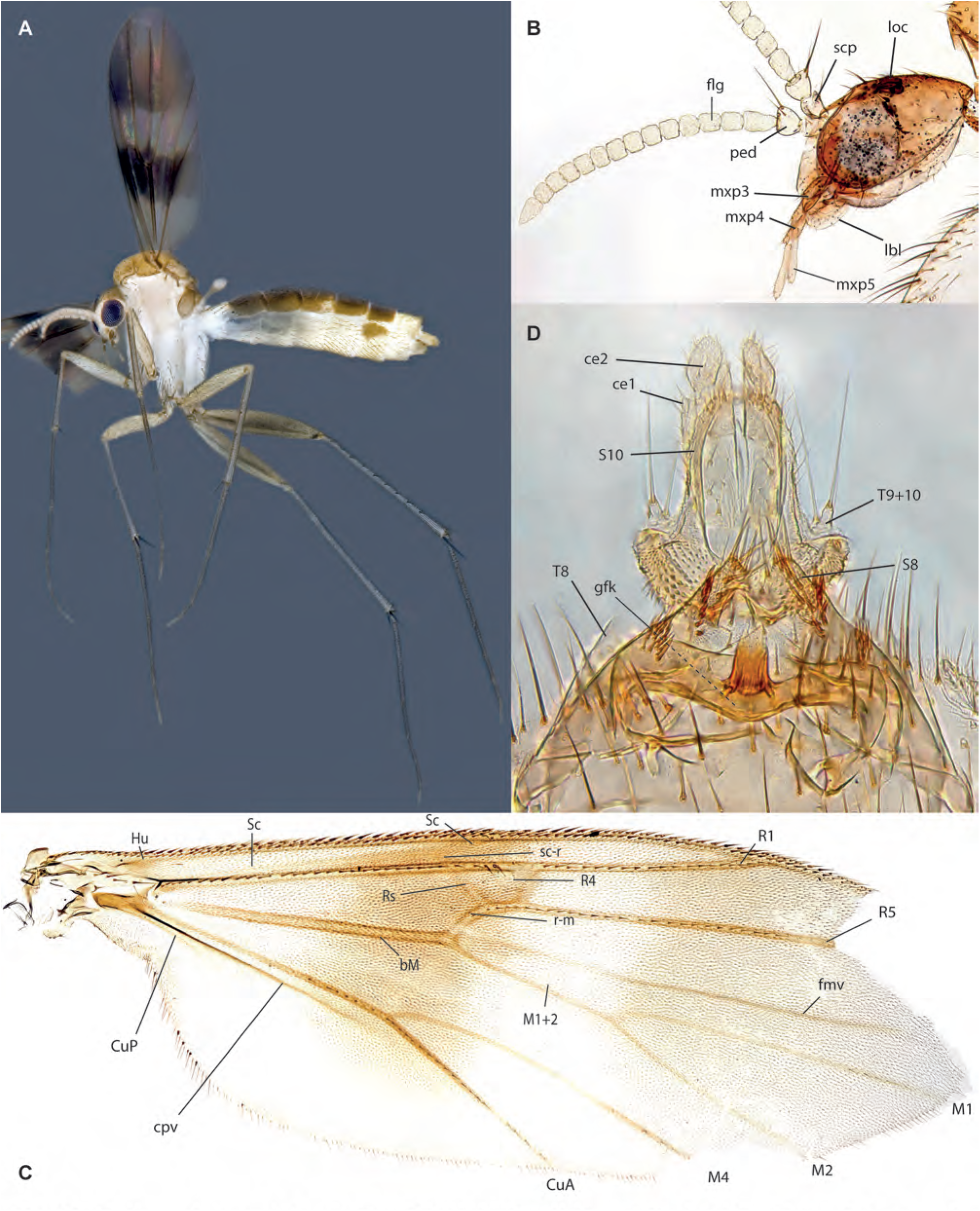
*Neoempheria* sp. A, female, ZRCBDP0048497. **A.** Habitus, lateral view. **B.** Head, lateral view. **C.** Wing. **D.** Terminalia, ventral view.

**Figs. 58A-D.**
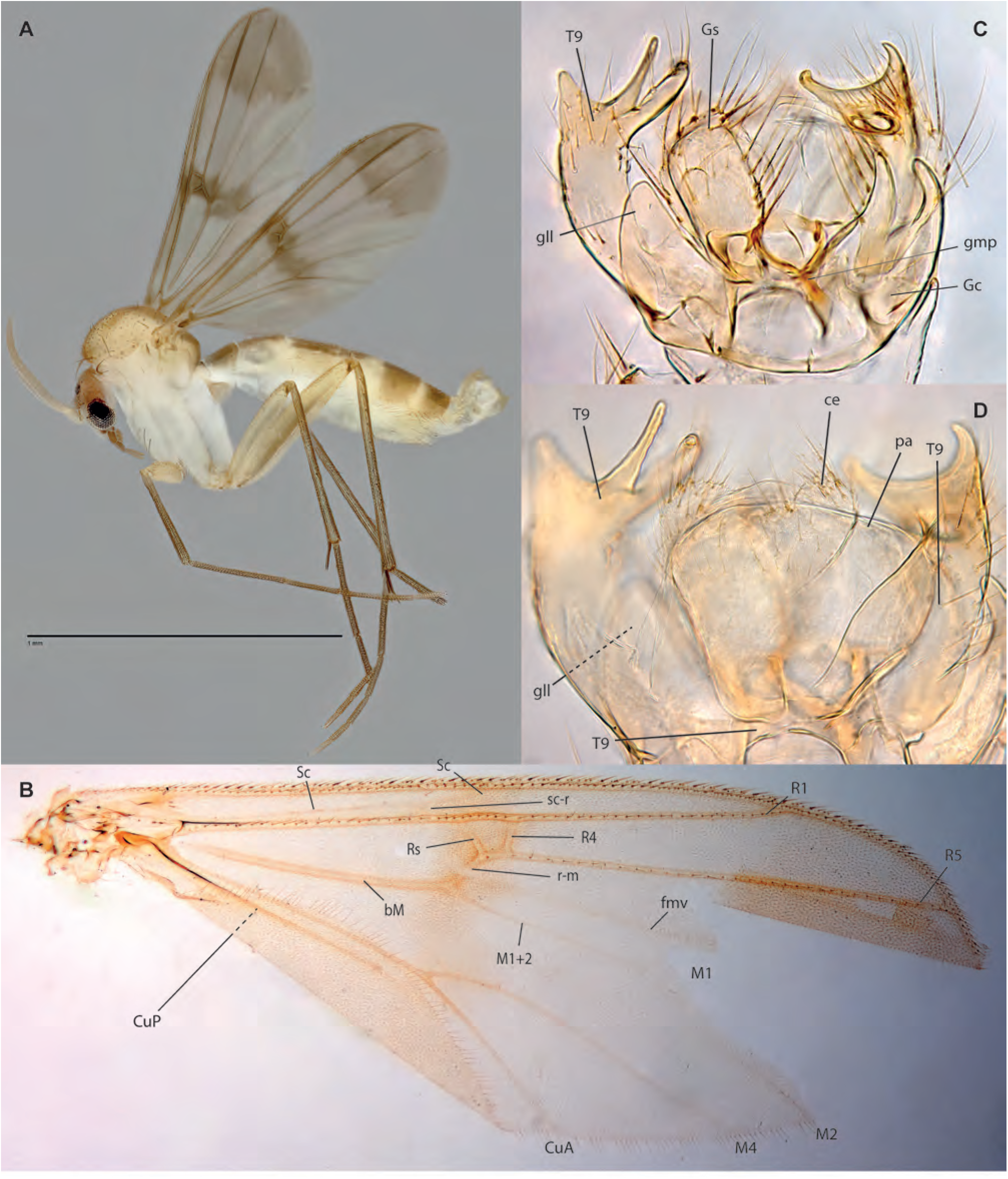
*Neoempheria sangabo* Amorim & Oliveira, **sp. n.** male holotype. **A.** Habitus, lateral view. **B.** Wing. **C.** Terminalia, ventral view. **D.** Detail of terminalia, dorsal view.

**Figs. 59A-D.**
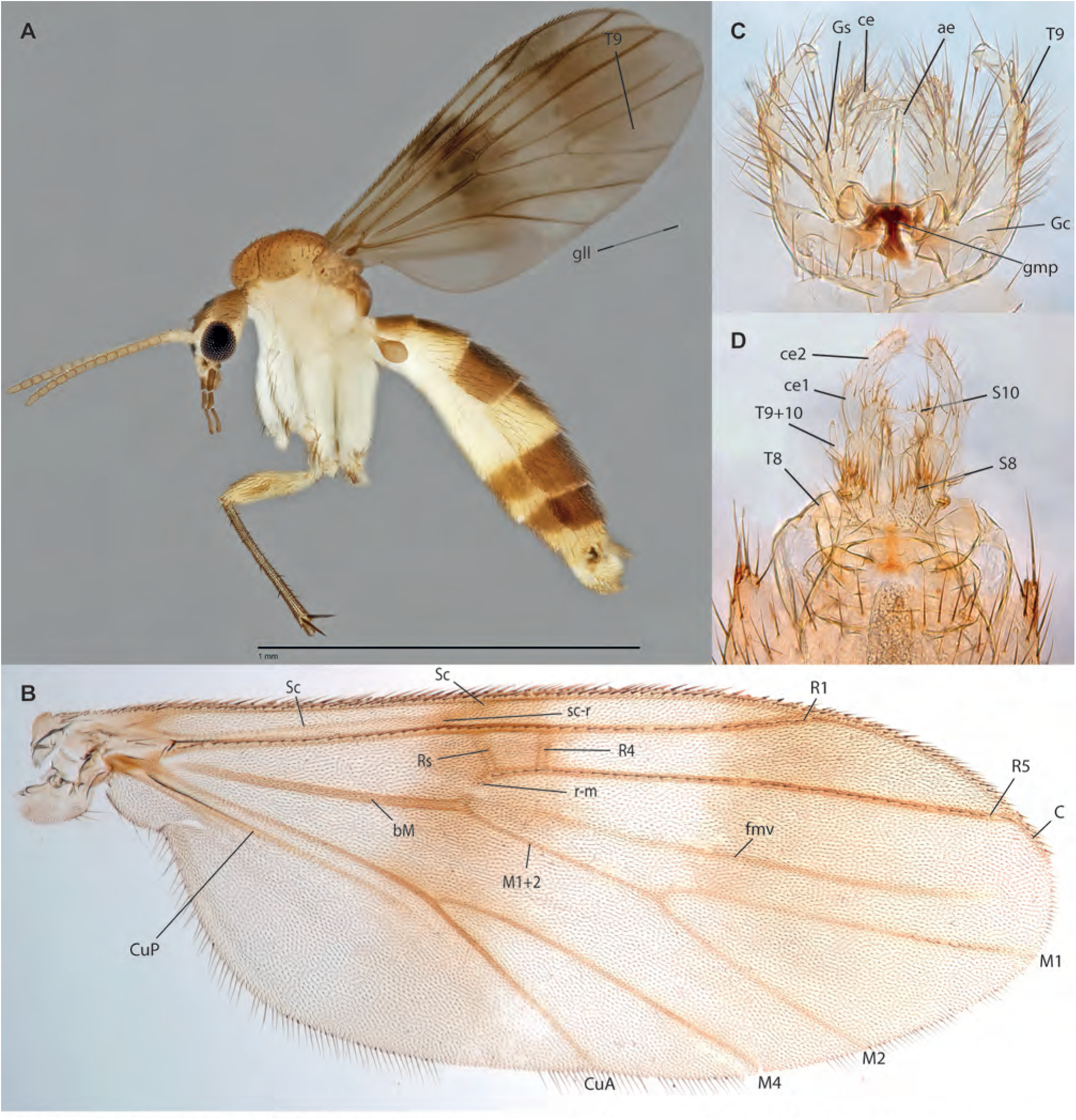
*Neoempheria shicheng* Amorim & Oliveira, **sp. n.** . **A.** Habitus, lateral view, male holotype. **B.** Wing, same. **C.** Male terminalia, ventral view, same. **D.** Female terminalia, ventral view, ZRCBDP0047906.

**Figs. 60A-D.**
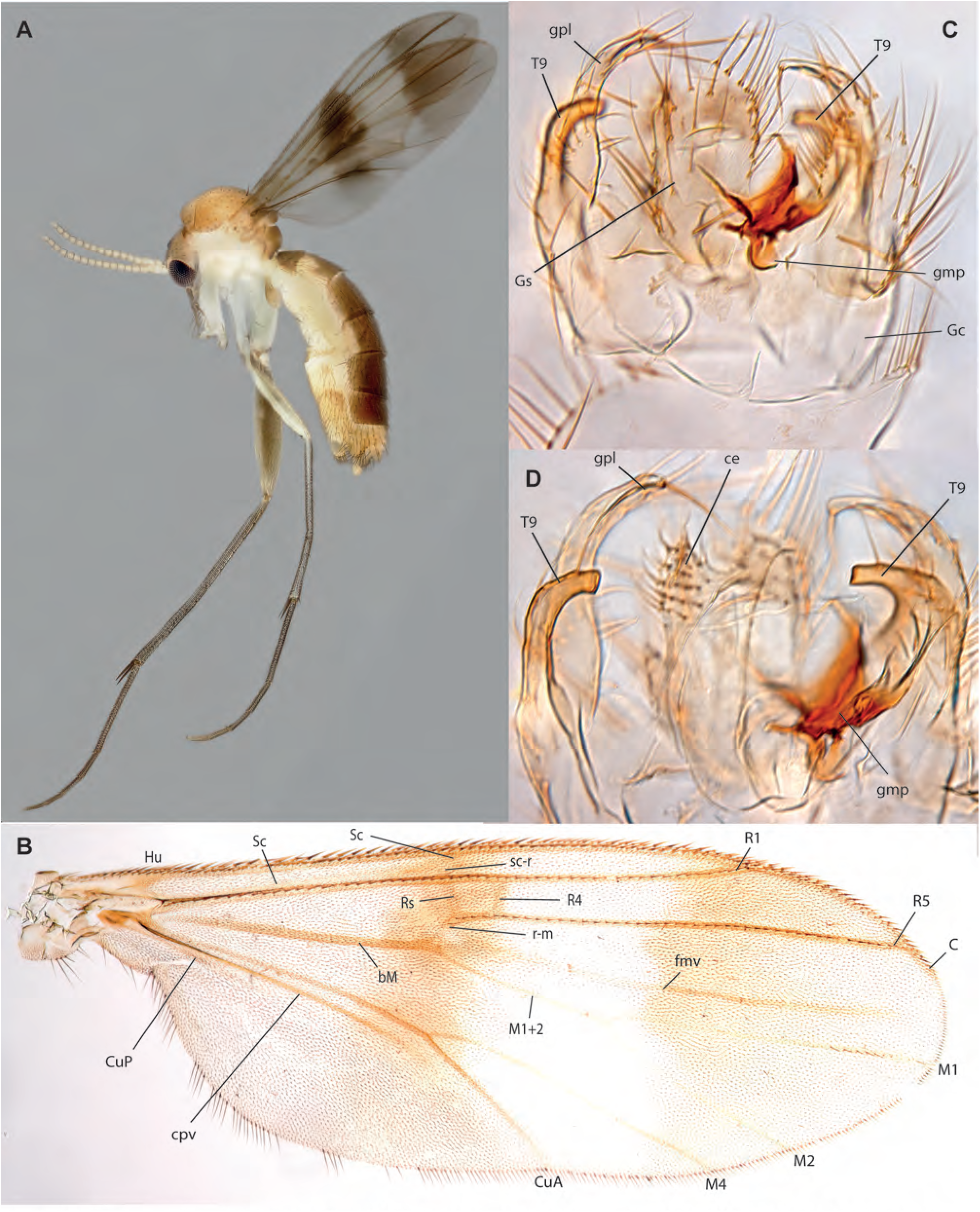
*Neoempheria ujong* Amorim & Oliveira, **sp. n. A.** Habitus, lateral view, male ZRCBDP0048978. **B.** Wing, male holotype. **C.** Male terminalia, ventral view, same. **D.** Male terminalia, mid-section, same.

**Figs. 61A-D.**
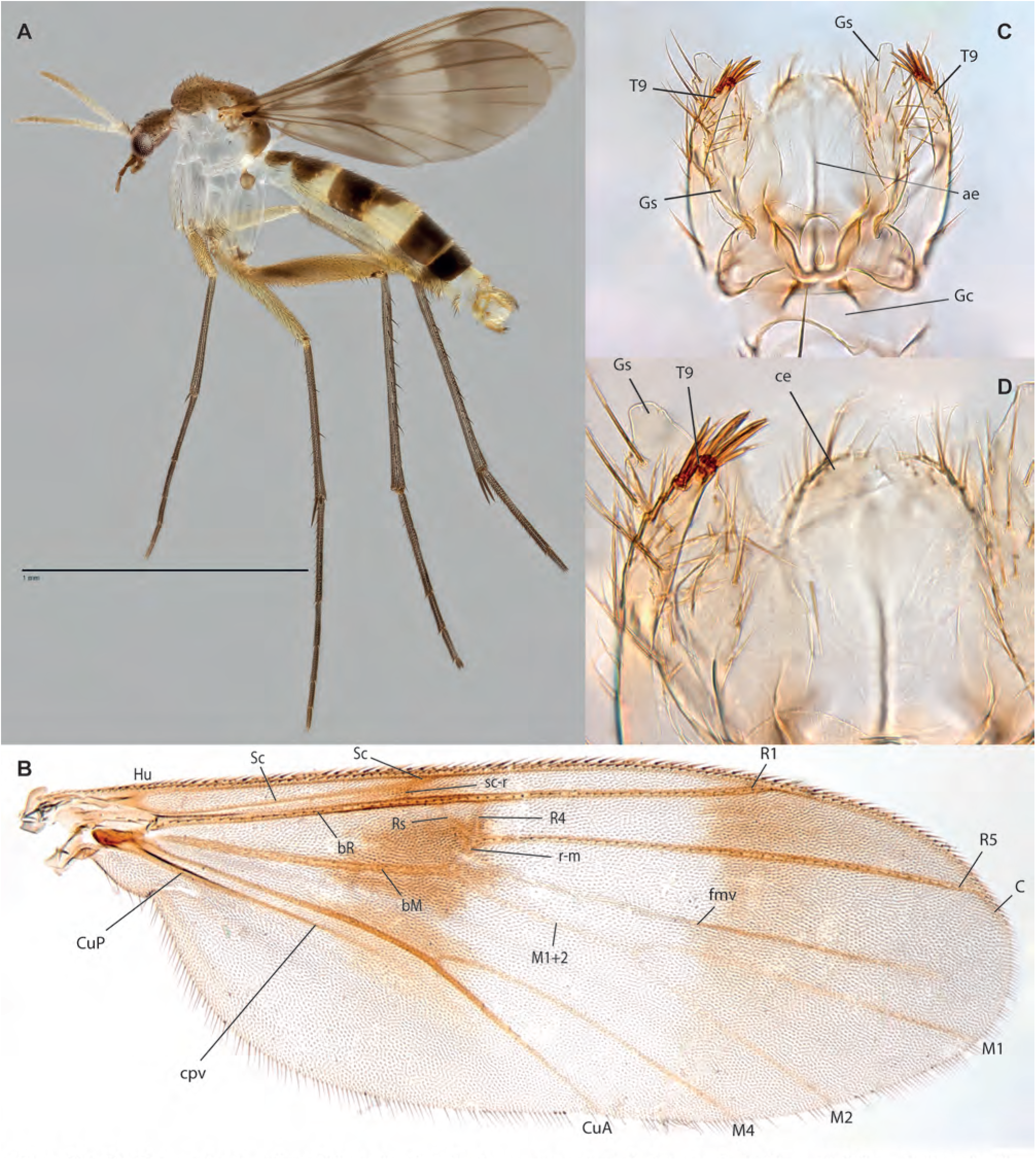
*Neoempheria subaraji* Amorim & Oliveira, **sp. n.,** male holotype. **A.** Habitus, lateral view. **B.** Wing. **C.** Terminalia, ventral view. **D.** Details of terminalia, dorsal view.

**Figs. 62A-D.**
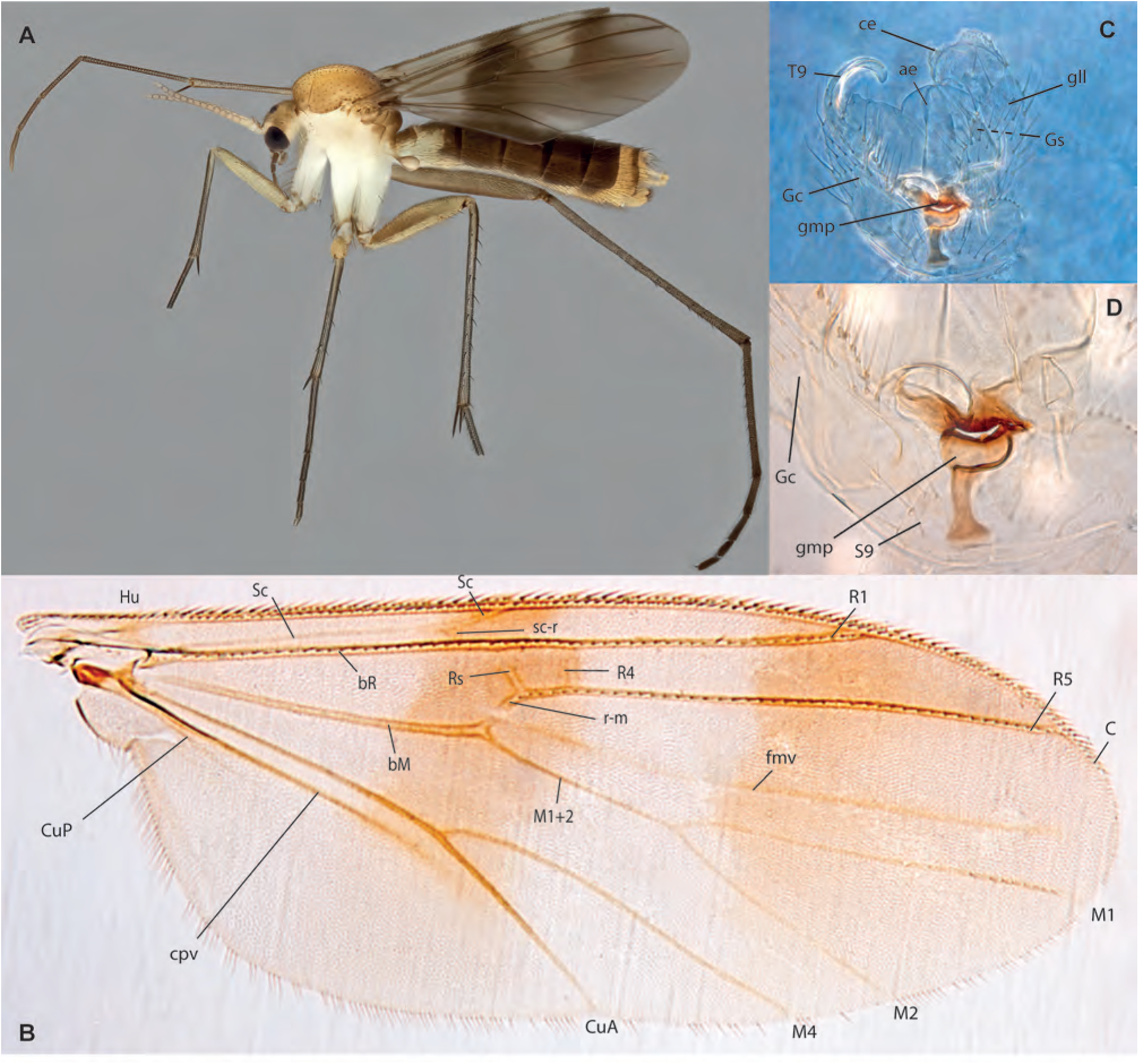
*Neoempheria kokoiyeeae* Amorim & Oliveira, **sp. n. A.** Habitus, lateral view, female, ZRCBDP0049205. **B.** Wing, male holotype. **C.** Male terminalia, ventral view, same. **D.** Detail of male terminalia, ventral view, same.

**Figs. 63A-E.**
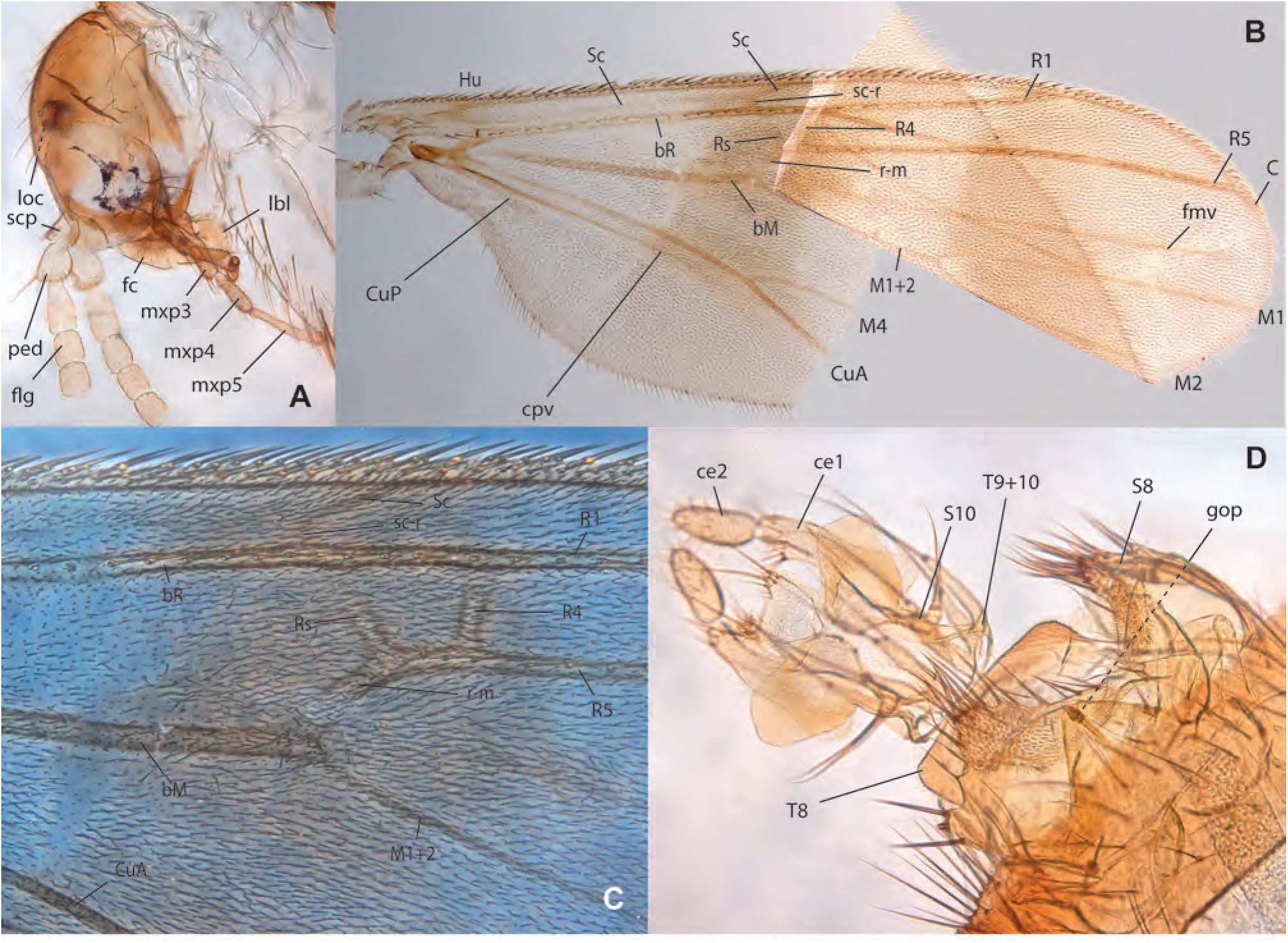
*Neoempheria* sp. G, female, ZRCBDP0137178. **A.** Head, lateral view. **B.** Wing. **C.** Detail of wing under phase contrast **D.** Terminalia, ventral view. **E.** Detail of terminalia, dorsal view.

**Figs. 64A-D.**
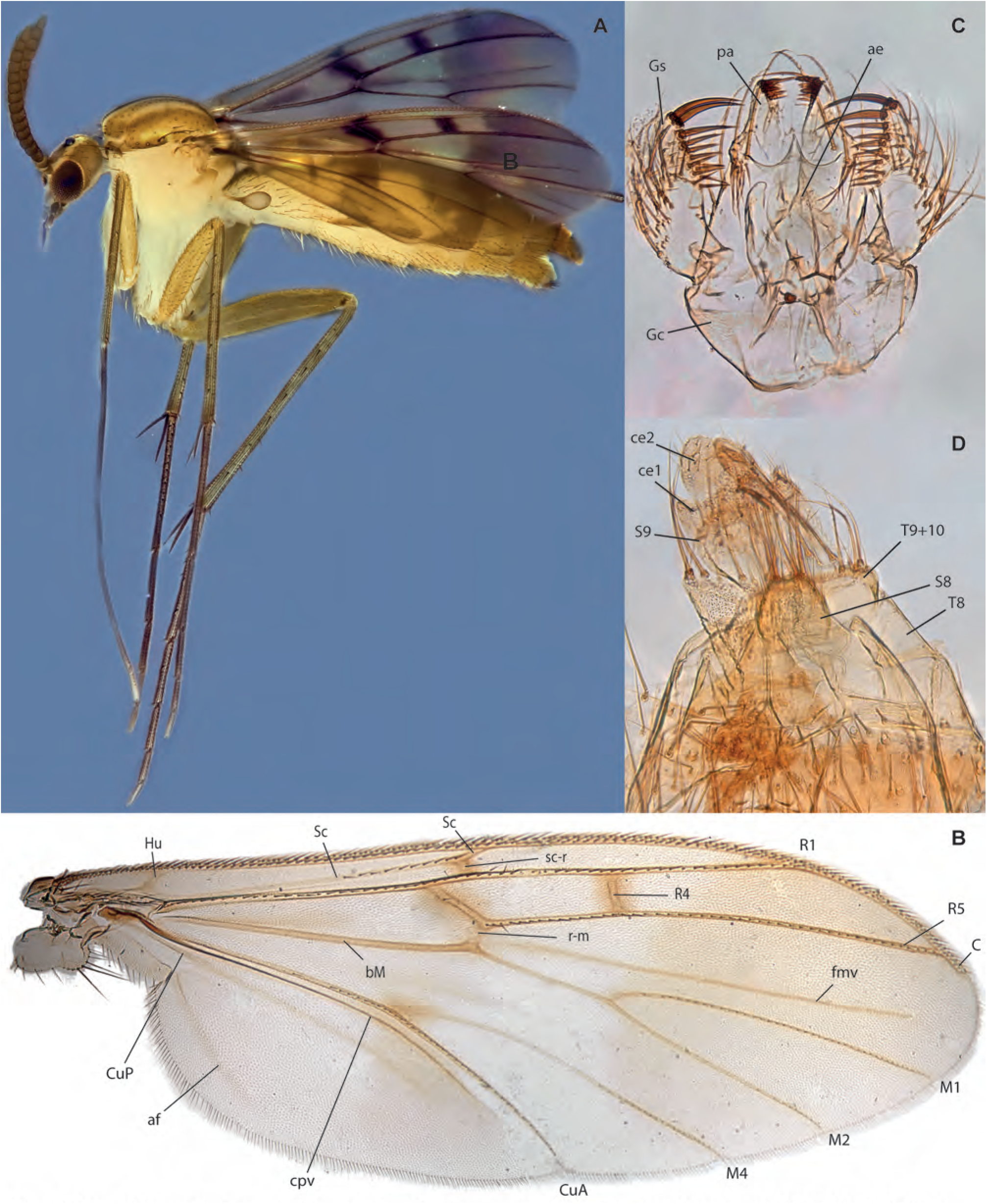
*Neoempheria mandai* Amorim & Oliveira, **sp. n. A.** Habitus, lateral view, female paratype ZRCBDP0048478. **B.** Wing, holotype. **C.** Male terminalia, ventral view, same. **D.** Female terminalia, ventral view, paratype ZRCBDP0048962.

**Figs. 65A-C.**
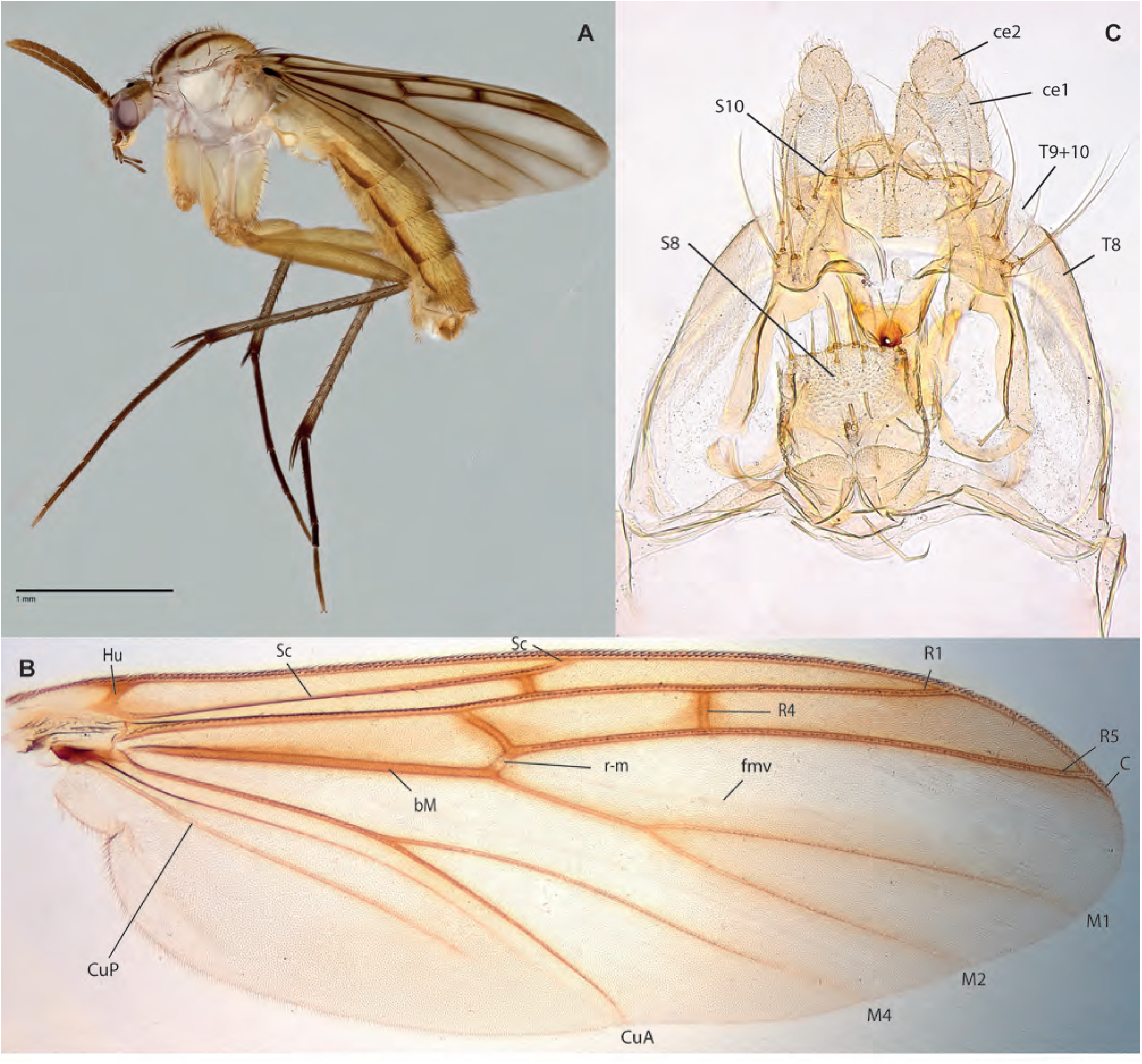
*Neoempheria malacca* Amorim & Oliveira, **sp. n. A.** Habitus, lateral view, male holotype. **B.** Female terminalia, ventral view, paratype ZRCBDP0048482. **C.** Wing, same.

**Figs. 66A-D.**
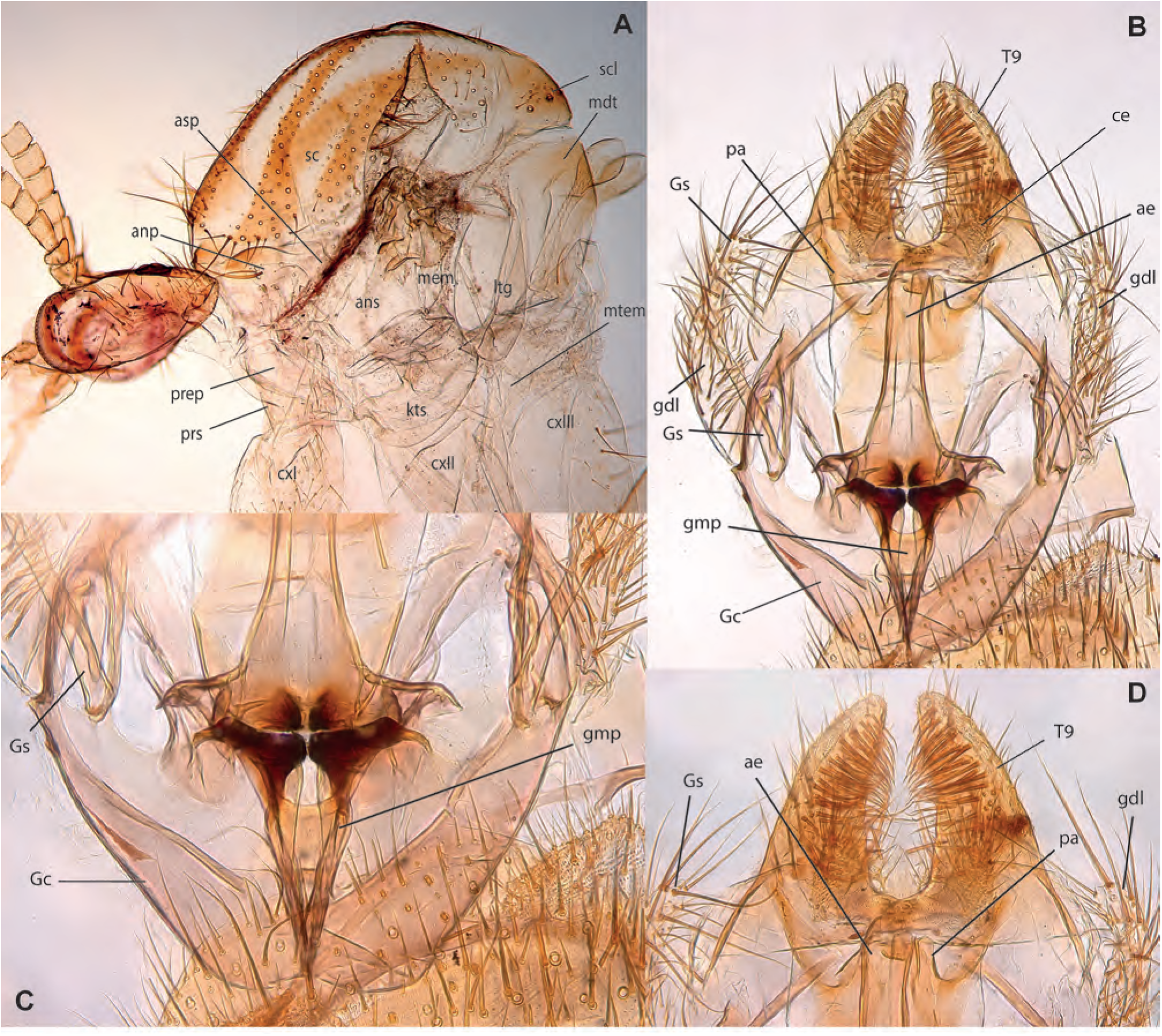
*Neoempheria malacca* Amorim & Oliveira, **sp. n. A.** Thorax, lateral view, female paratype ZRCBDP0048482. **B.** Male holotype, terminalia, ventral view. **C.** Same, detail of anterior end. **D.** Same, detail of posterior end.

**Figs. 67A-D.**
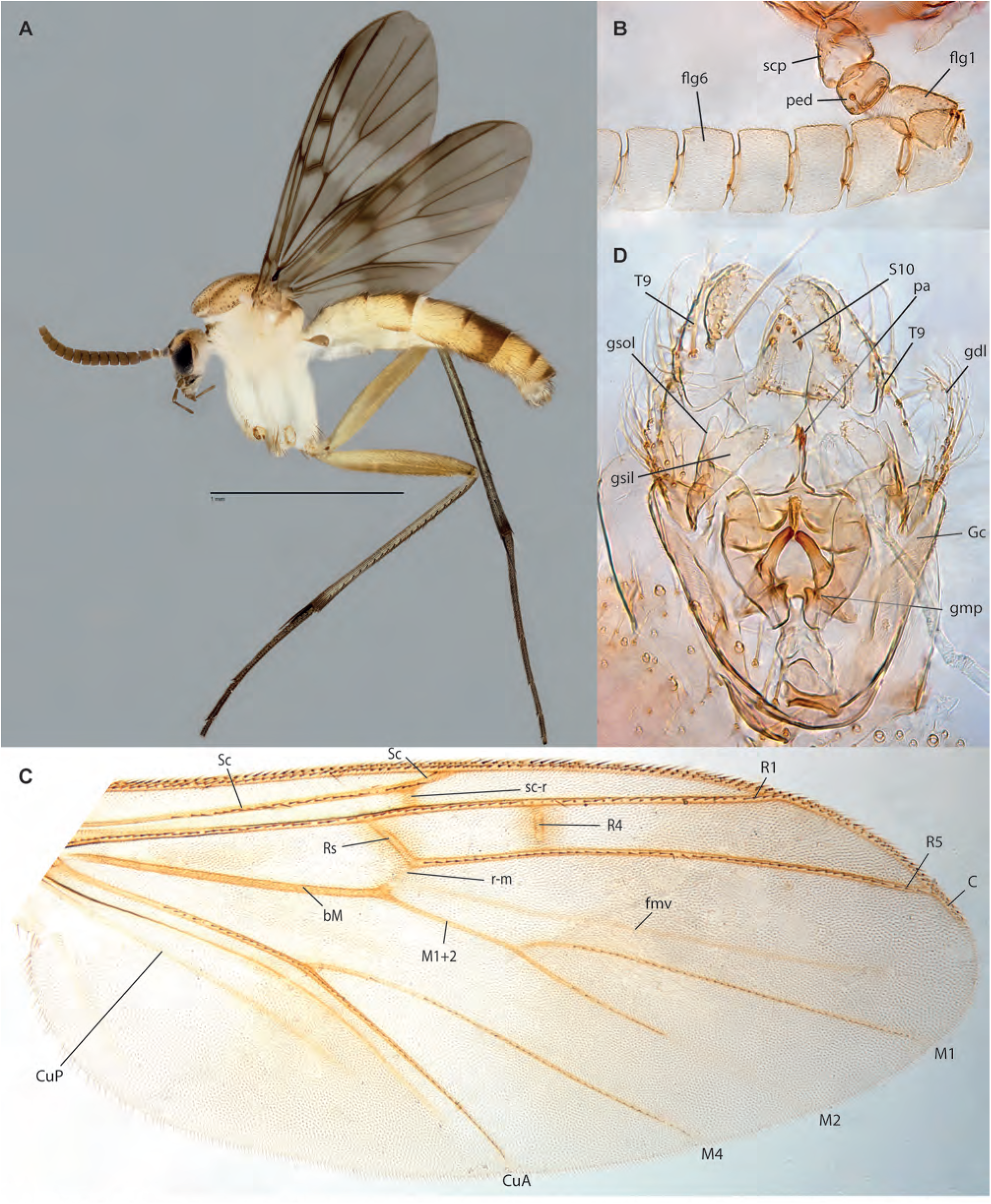
*Neoempheria sinkapho Amorim* & Oliveira, **sp. n.,** male holotype. **A.** Habitus, lateral view. **B.** Detail of antenna. **C.** Wing. **D.** Male terminalia, ventral view.

**Figs. 68A-F.**
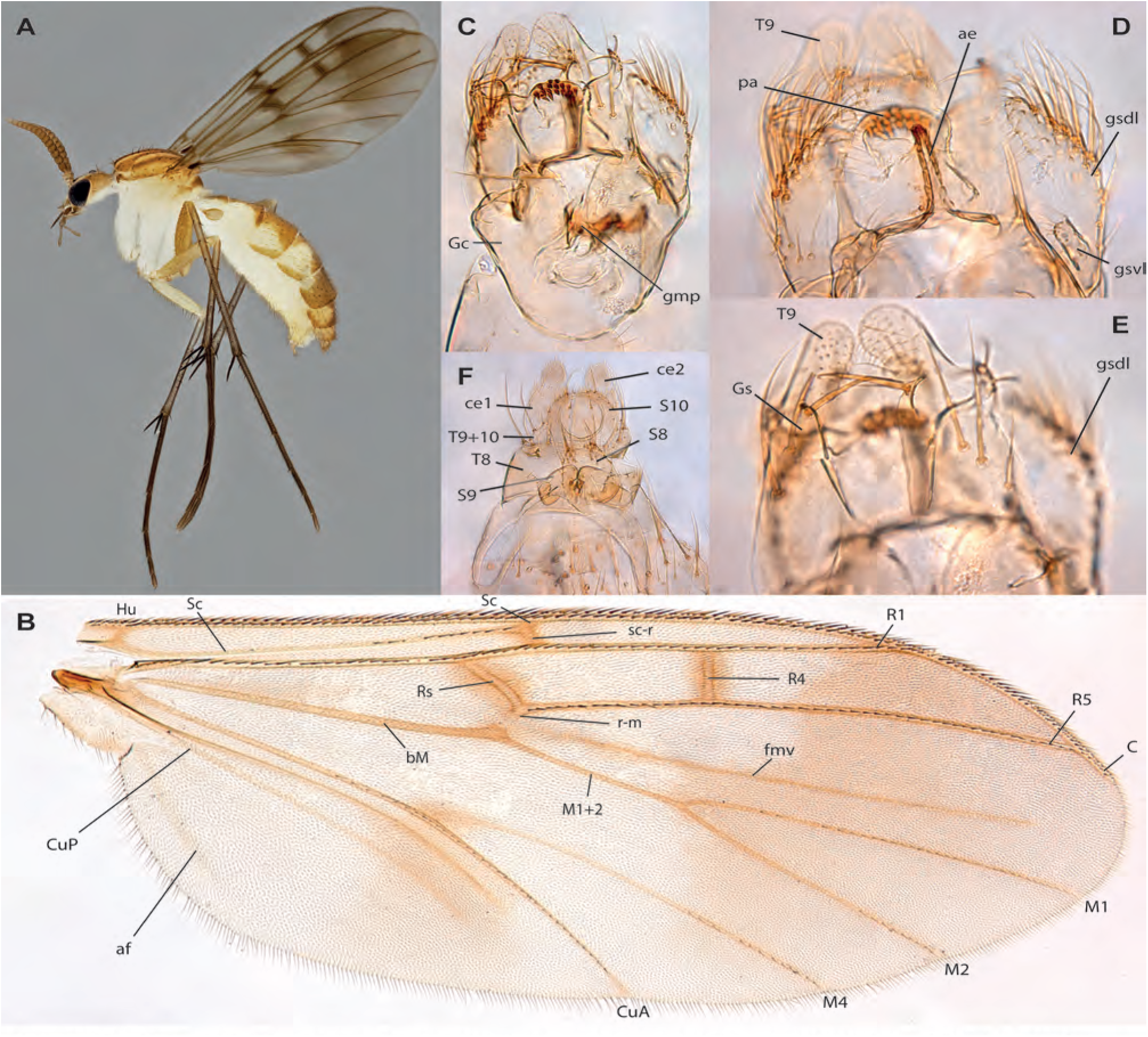
*Neoempheria singapura Amorim* & Oliveira, **sp. n. A.** Habitus, lateral view, female ZRCBDP0047930. **B.** Wing, male holotype. **C.** Male terminalia, ventral view, same. **D.** Detail of male terminalia, ventral view, same. **E.** Detail of male terminalia, dorsal view, same. **F.** Female terminalia, ventral view, ZRCBDP0047796.

**Figs. 69A-C.**
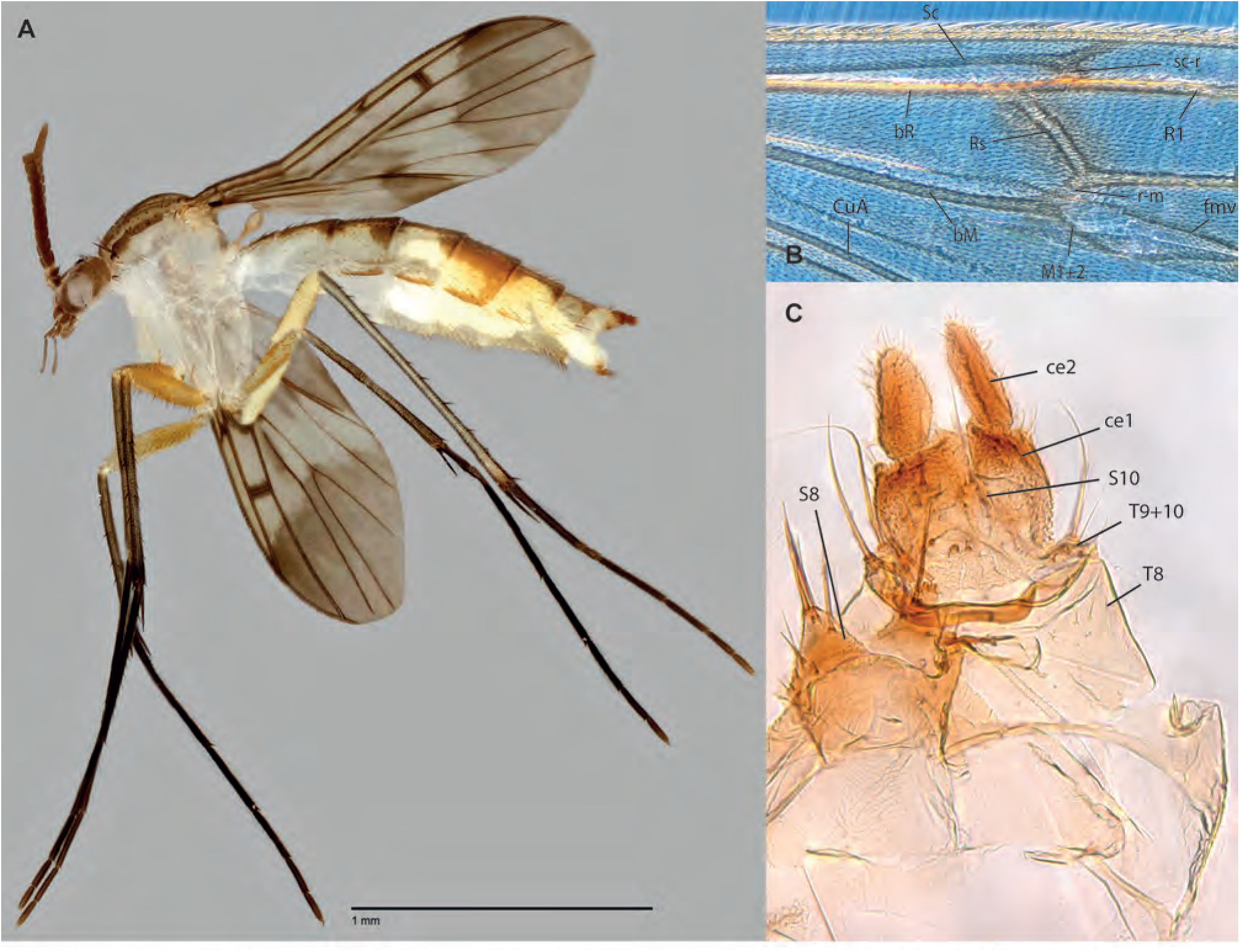
*Neoempheria* sp. C, female, ZRCBDP0048477. **A.** Habitus, lateral view. **B.** Detail of wing under phase contrast. **C.** Terminalia, ventral view.

**Figs. 70A-D.**
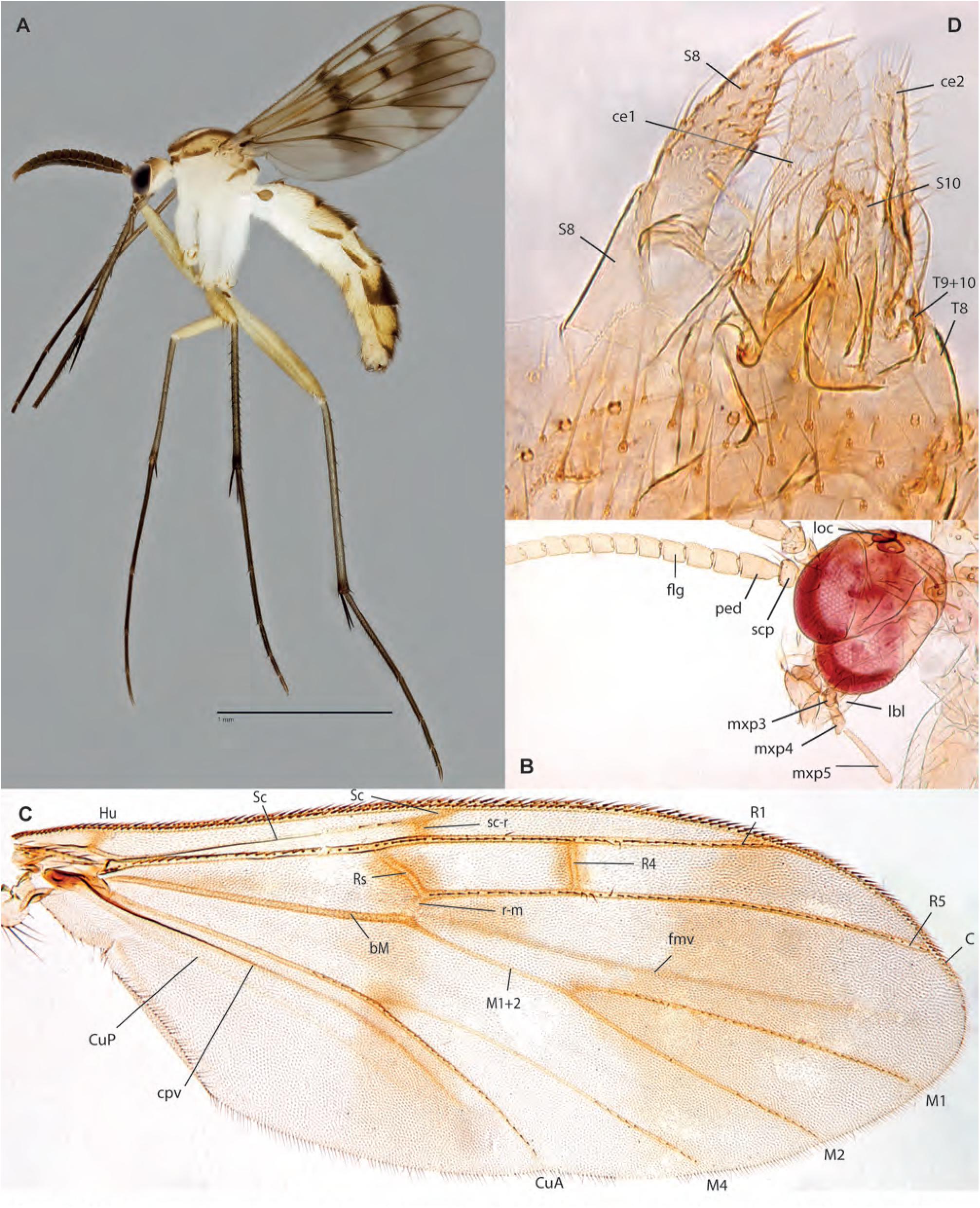
*Neoempheria xinjiapo* Amorim & Oliveira, **sp. n. A.** Habitus, lateral view, male ZRCBDP0048892. **B.** Head, lateral view, female paratype ZRCBDP0047795. **C.** Wing, same. **D.** Female terminalia, ventral view, same.

**Figs. 71A-B.**
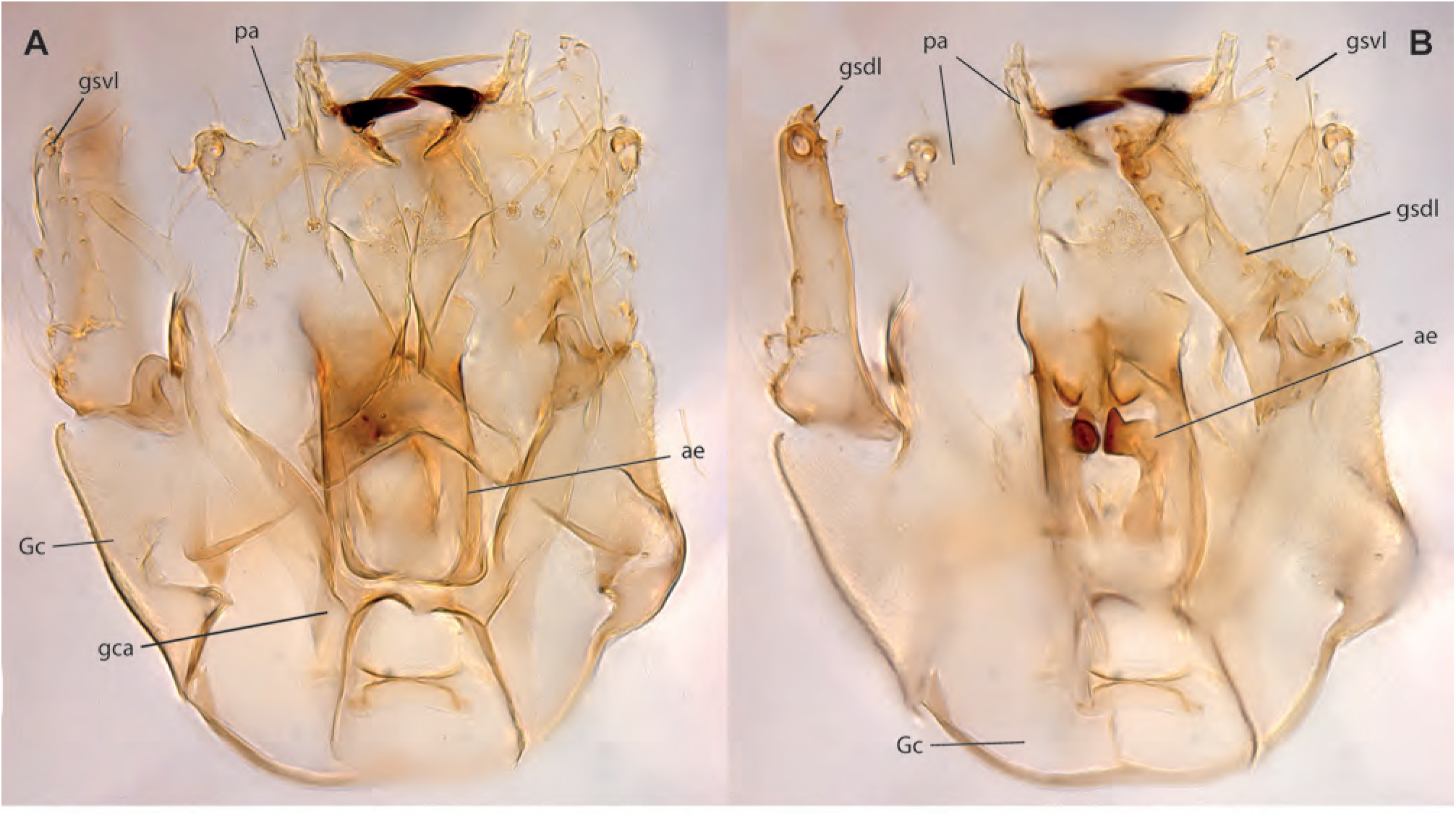
*Neoempheria xinjiapo* Amorim & Oliveira, **sp. n.,** male holotype. **A.** Terminalia, ventral view. **B.** Terminalia, dorsal view.

**Figs. 72A-D.**
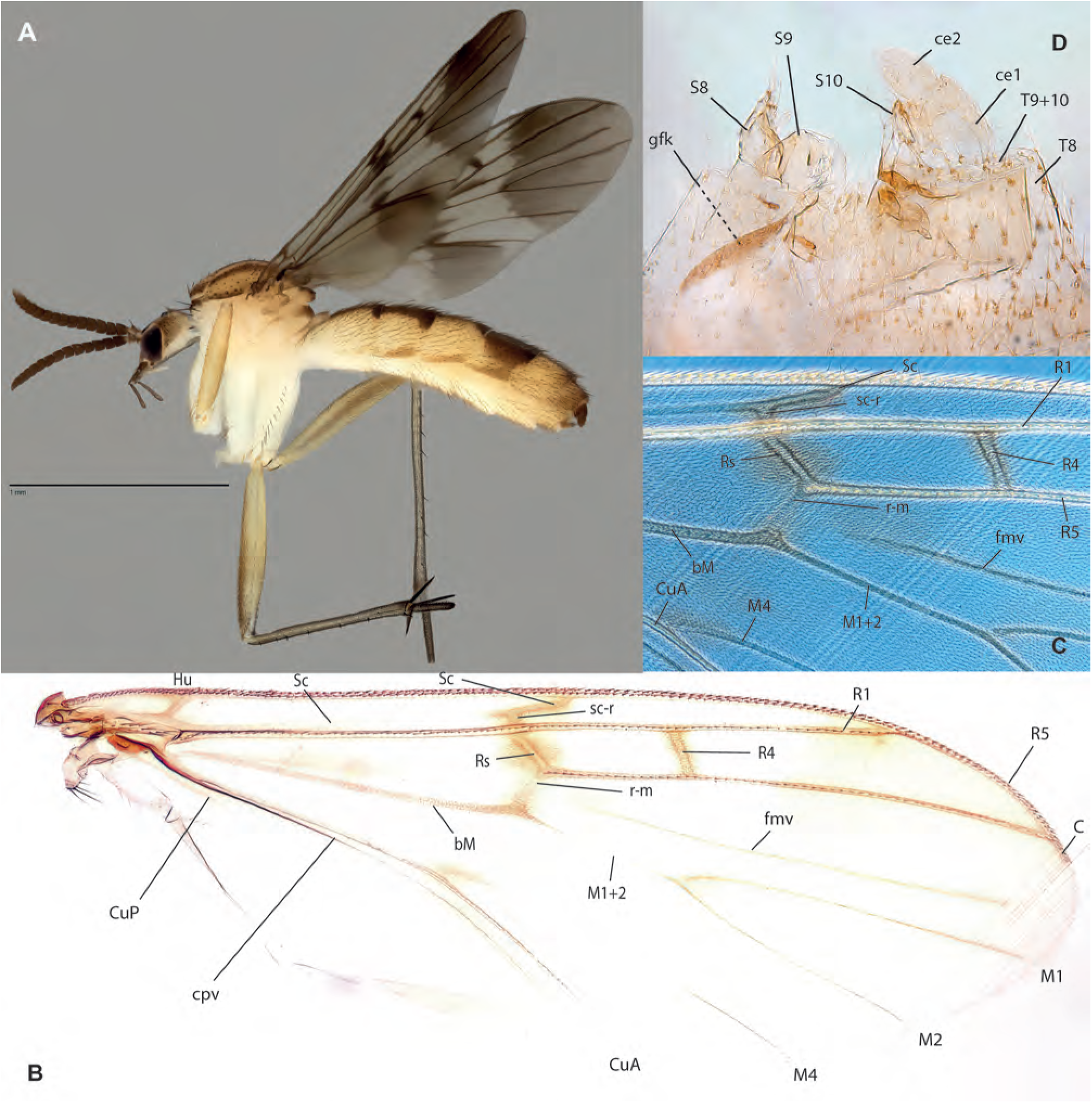
*Neoempheria* sp. D, female ZRCBDP0049022. **A.** Habitus, lateral view. **B.** Wing. **C.** Detail of wing under phase contrast. **D.** Female terminalia, lateral view.

**Figs. 73A-C.**
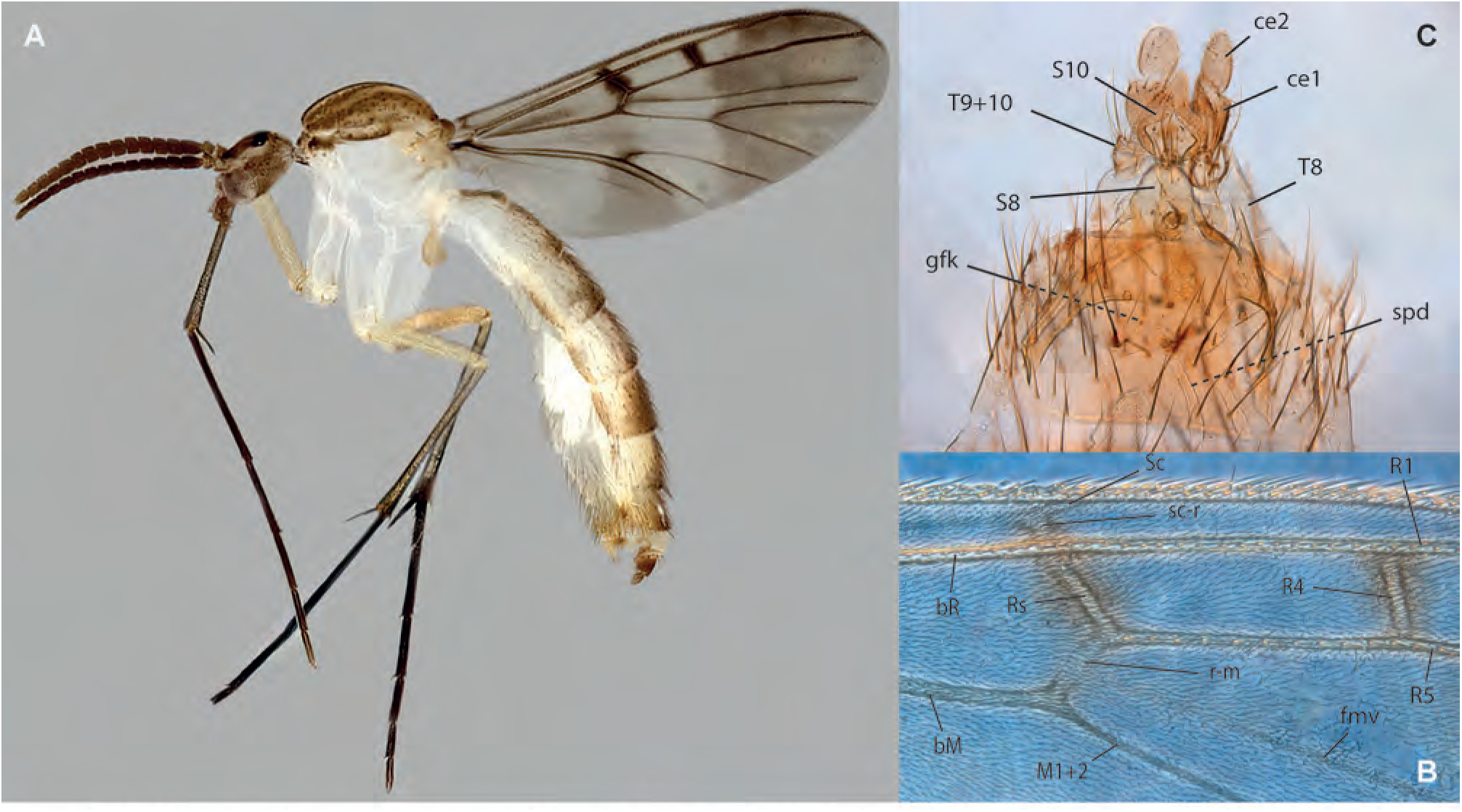
*Neoempheria* sp. E, female ZRCBDP00049180. **A.** Habitus, lateral view. **B.** Detail of wing under phase contrast. **C.** Female terminalia, ventral view.

**Figs. 74A-D.**
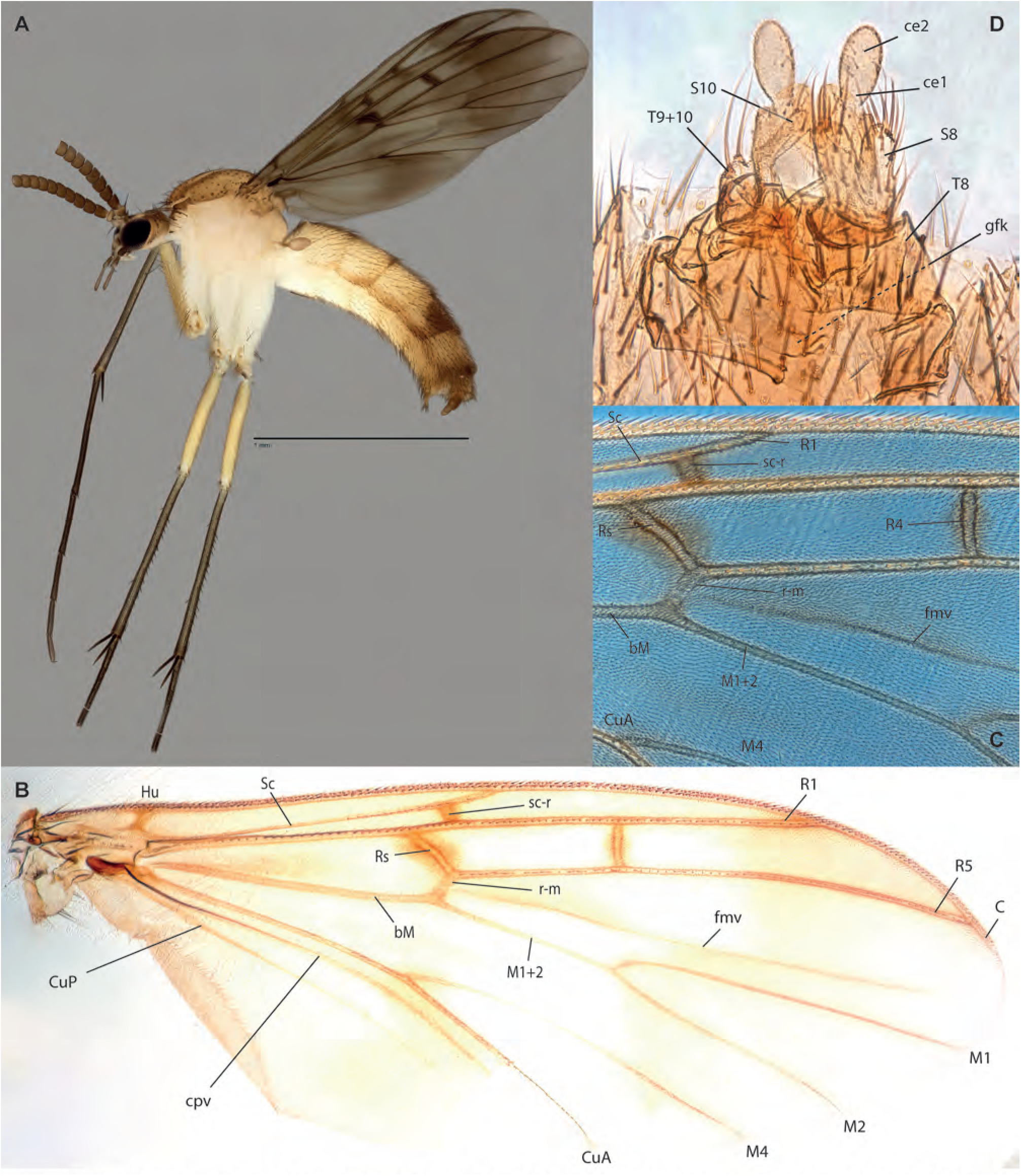
*Neoempheria* sp. F, female. **A.** Habitus, lateral view, ZRCBDP0047902. **B.** Wing, ZRCBDP0047836. **C.** Detail of wing under phase contrast, same. **D.** Female terminalia, ventral view, same.

**Figs. 75A-E.**
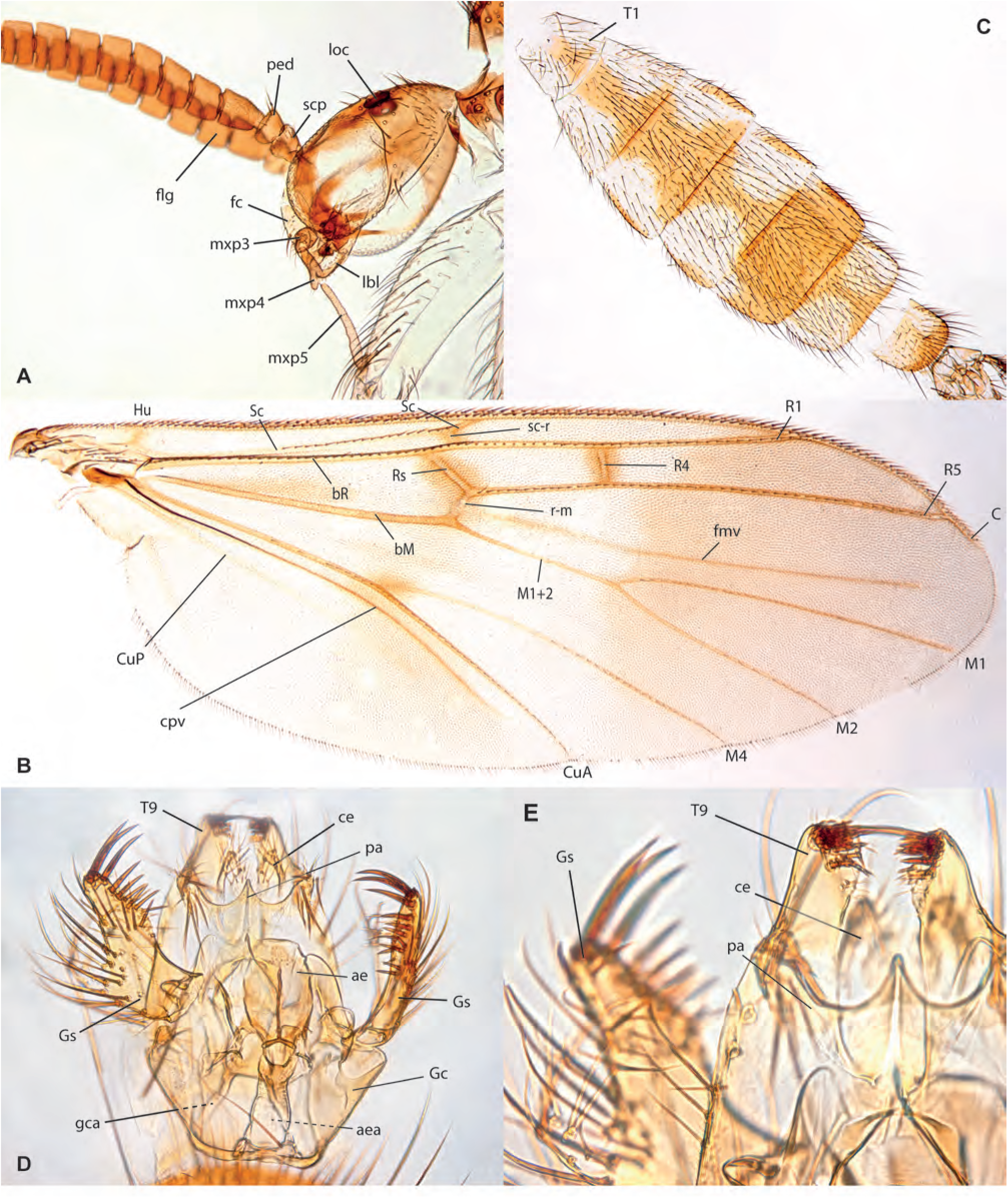
*Neoempheria* **sp. H,** male, ZRCBDP0066766. **A.** Head, lateral view. **B.** Wing. **C.** Abdomen, dorsal view. **D.** Terminalia, ventral view. **E.** Detail of terminalia, dorsal view.

**Figs. 76A-D.**
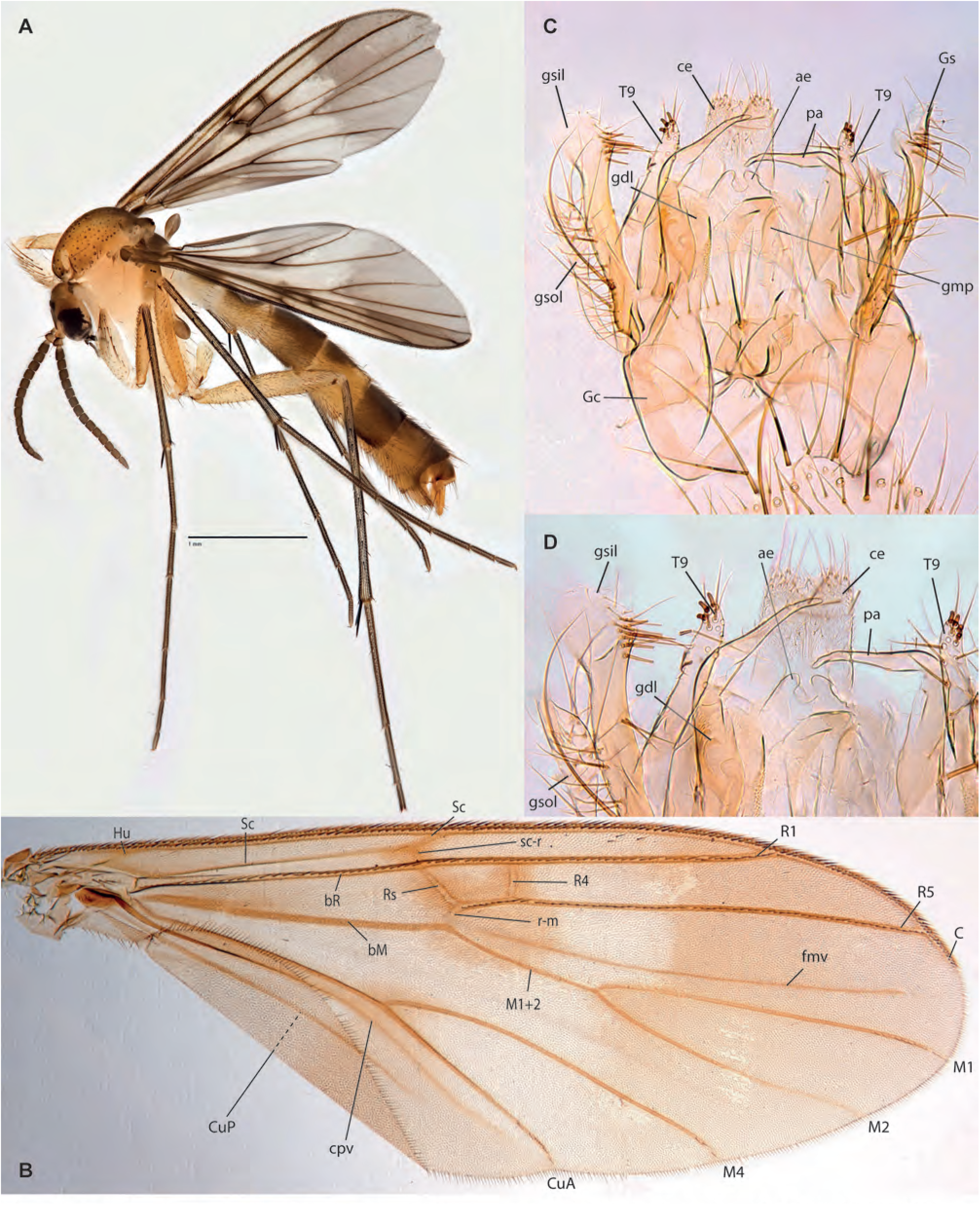
*Neoempheria puluochung* Amorim & Oliveira, **sp. n. A.** Habitus, lateral view, female, ZRCBDP0048494. **B.** Wing, male holotype. **C.** Male terminalia, ventral view, same. **D.** Detail of male terminalia, ventral view, same.

**Figs. 77A-B.**
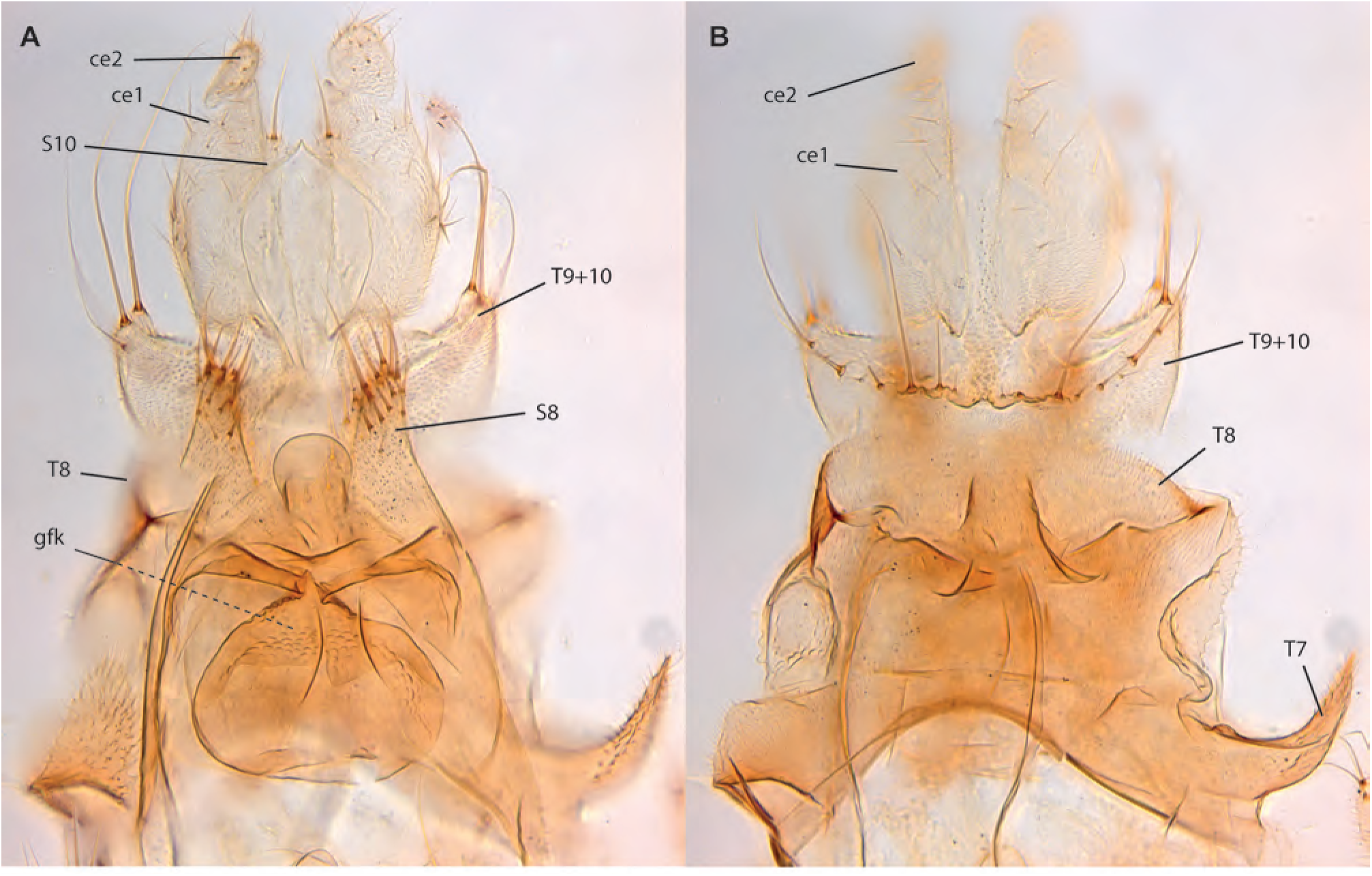
*Neoempheria puluochung* Amorim & Oliveira, **sp. n. A.** Female terminalia, paratype ZRCBDP0048493, ventral view. **B.** Female terminalia, dorsal view, same.

**Figs. 78A-D.**
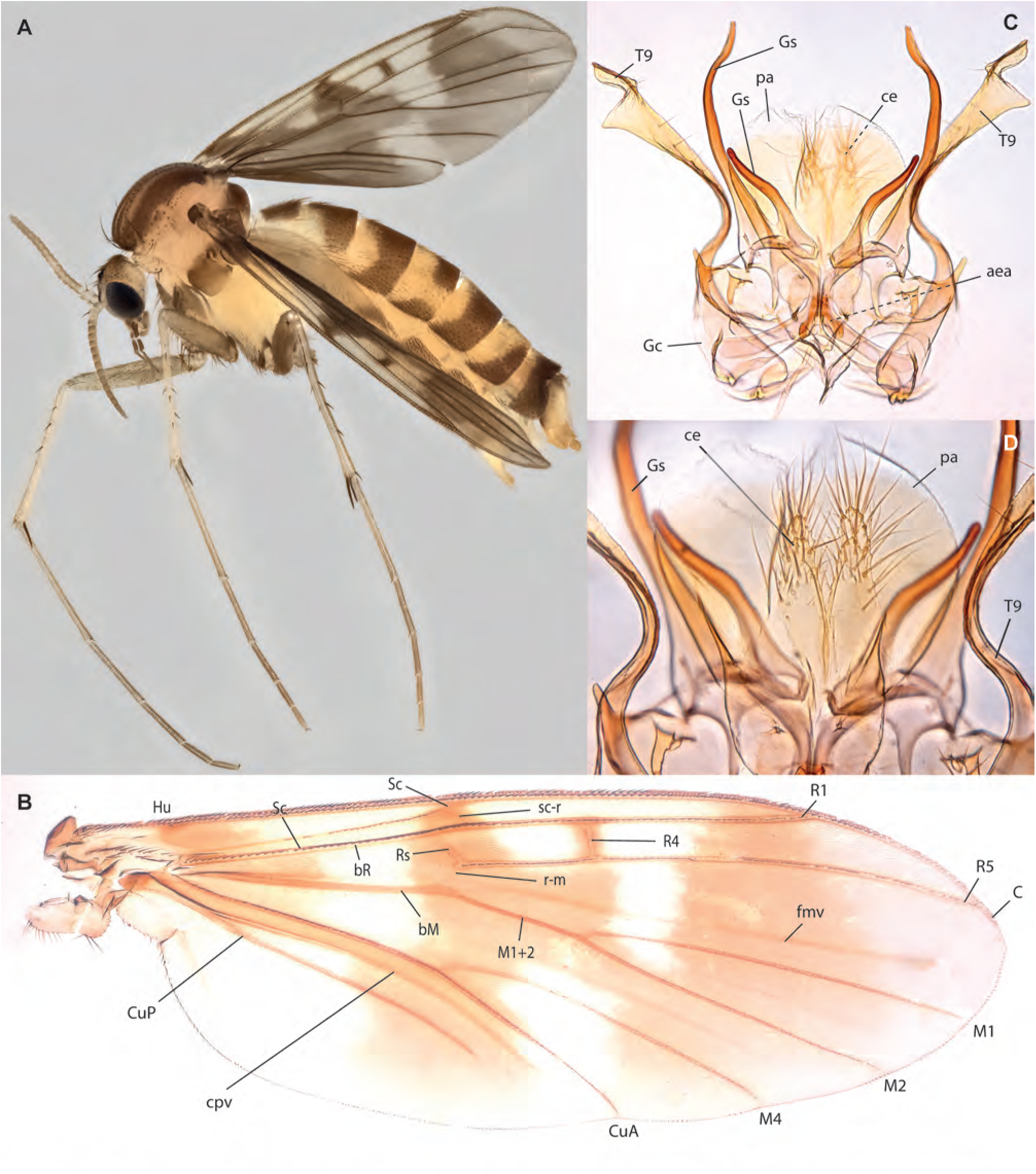
*Neoempheria merdeka* Amorim & Oliveira, **sp. n. A.** Habitus, lateral view, female paratype ZRCBDP0155081. **B.** Wing, male holotype. **C.** Male terminalia, ventral view, same. **D.** Detail of male terminalia, dorsal view, same.

**Figs. 79A-B.**
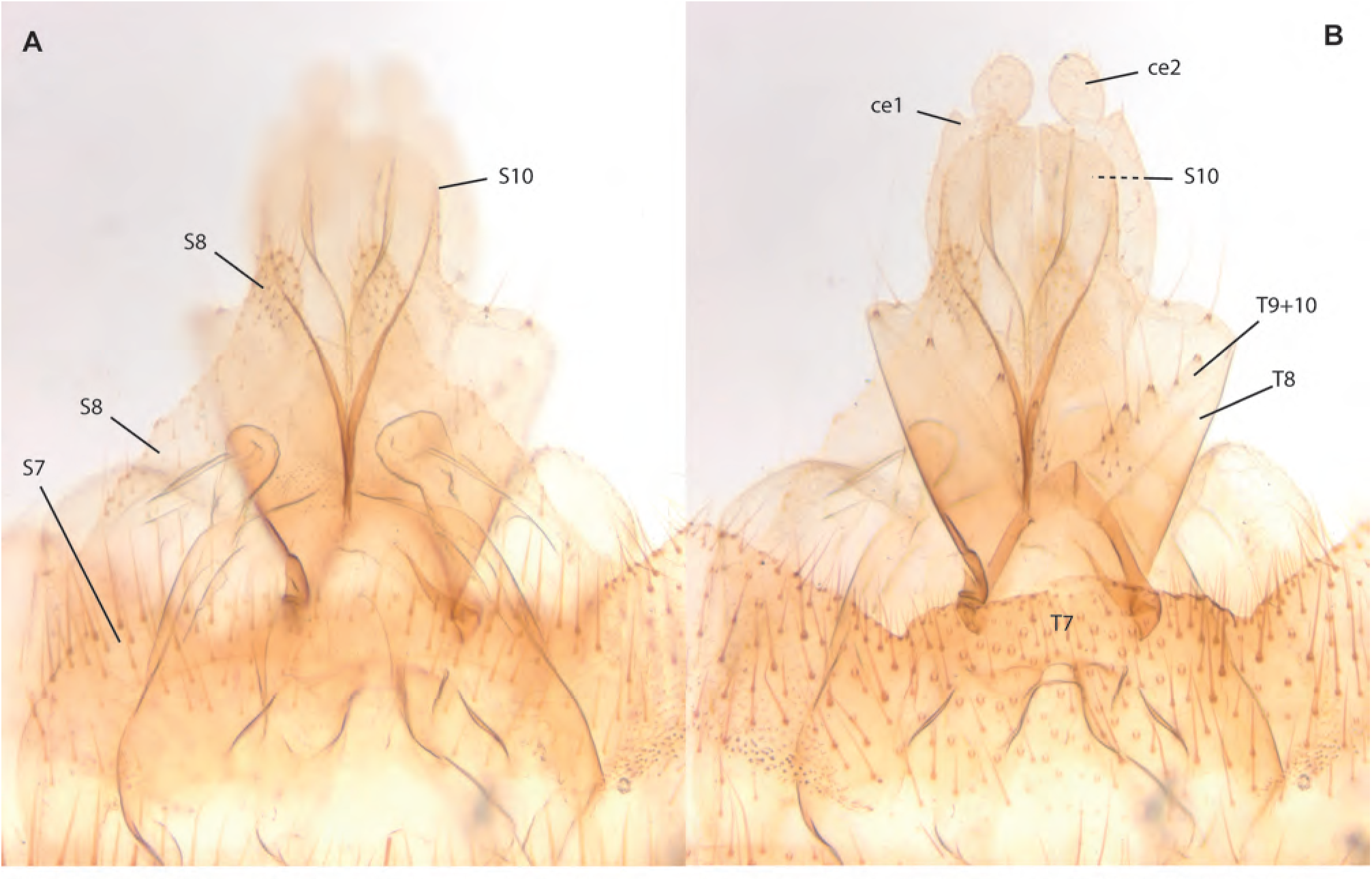
*Neoempheria merdeka* Amorim & Oliveira, **sp. n.,** female terminalia, paratype ZRCBDP0074035. **A.** Ventral view. **B.** Dorsal view.

**Figs. 80A-D.**
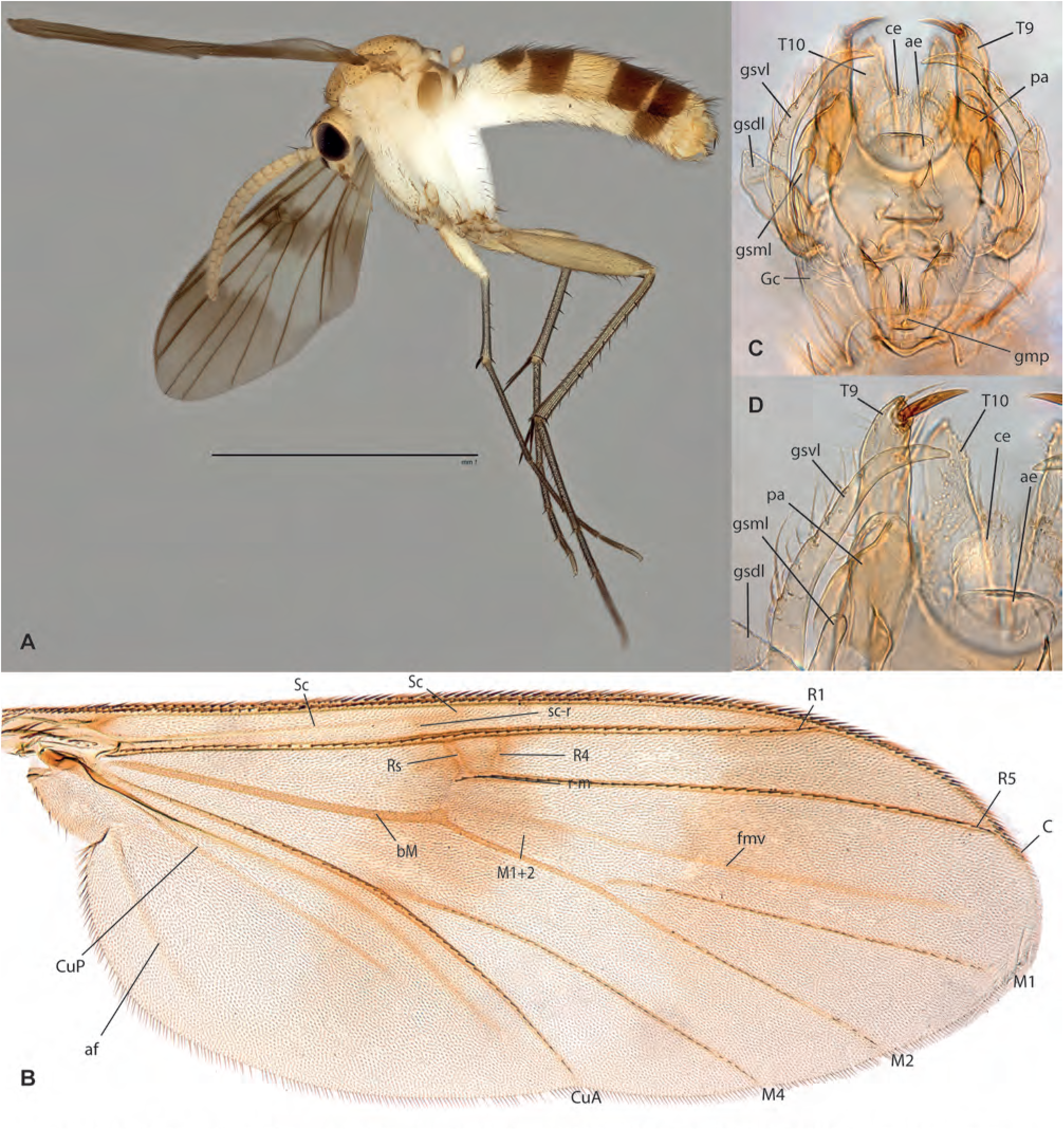
*Neoempheria dizonalis* (Edwards). **A.** Habitus, lateral view, male ZRCBDP0047918. **B.** Wing, male holotype. **C.** Male terminalia, ventral view, same. **D.** Detail of male terminalia, dorsal view, same.

**Figs. 81A-C.**
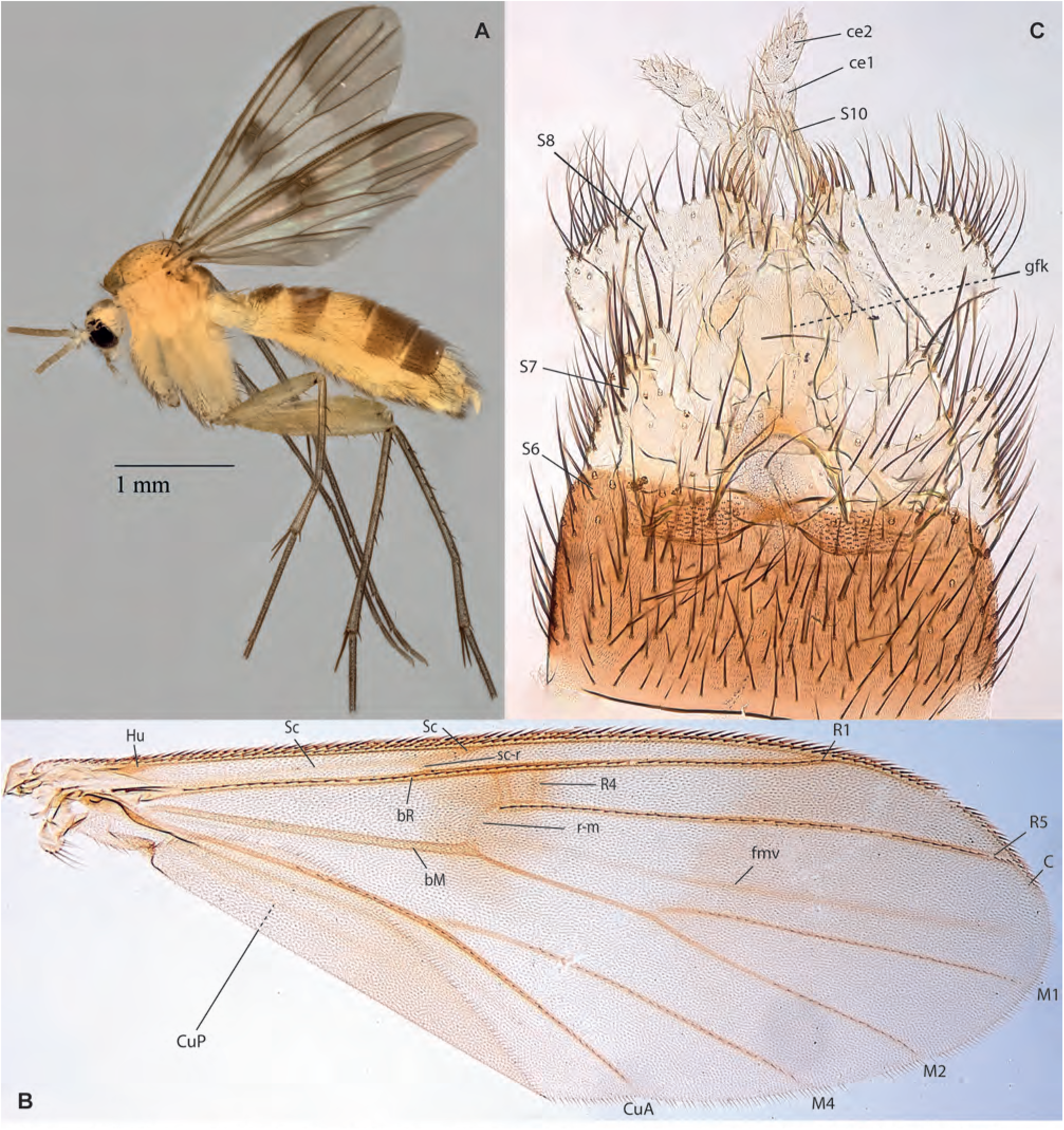
*Neoempheria neesoon* Amorim & Oliveira, **sp. n. A.** Habitus, lateral view, female, ZRCBDP0154986. **B.** Wing, female holotype. **C.** Female terminalia, ventral view, ZRCBDP0049243.

**Figs. 82A-H.**
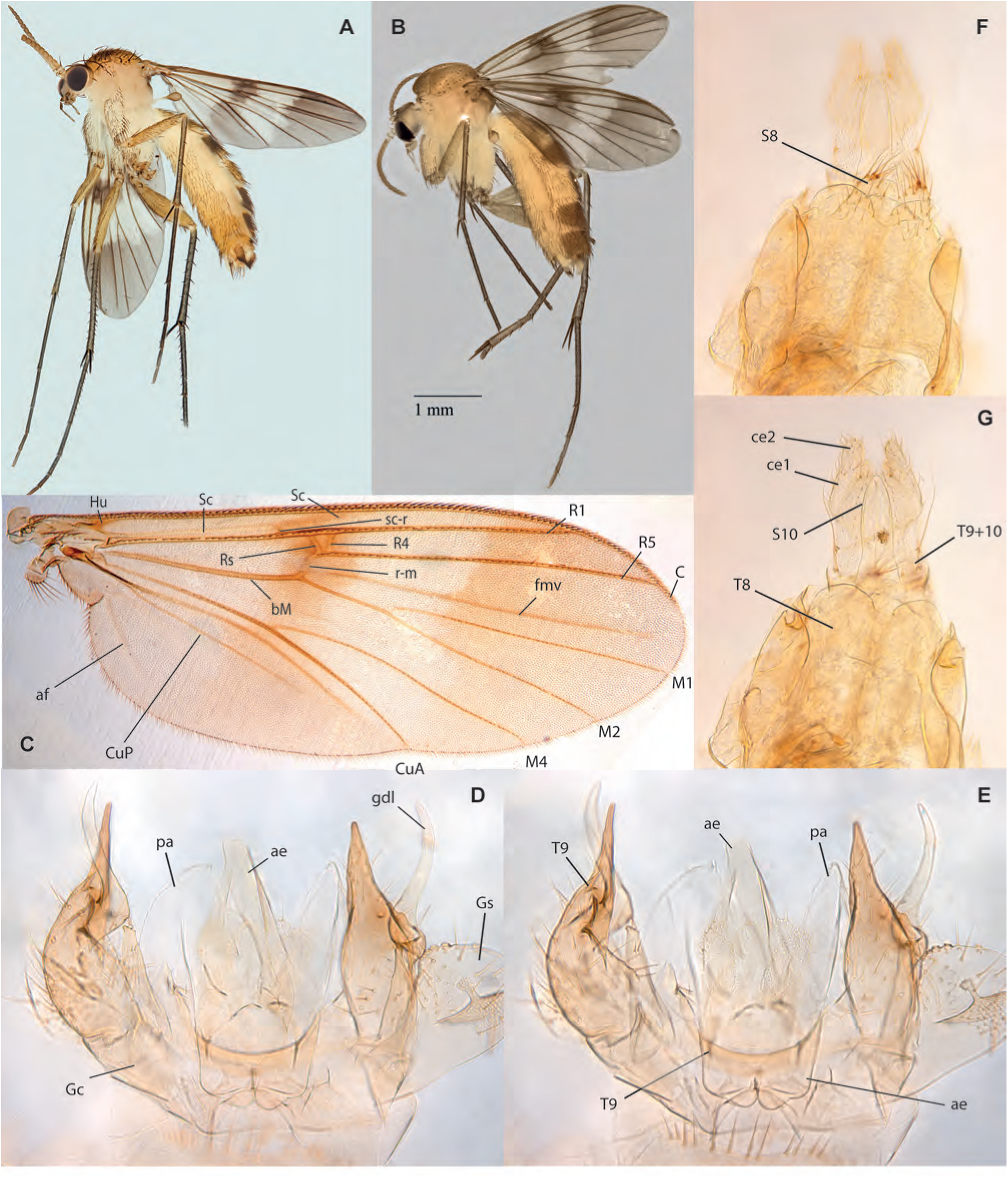
*Neoempheria pulau* Amorim & Oliveira, **sp. n. A.** Habitus, lateral view, male paratype, ZRCBDP0048491. **B.** Habitus, lateral view, female paratype, ZRCBDP0155089. **C.** Head, lateral view, male holotype. **D.** Wing, same. **E.** Male terminalia, ventral view, same. **F.** Male terminalia, dorsal view, same. **G.** Female terminalia, ventral view, paratype, ZRCBDP0048485. **H.** Female terminalia, dorsal view, same.

**Figs. 83A-D.**
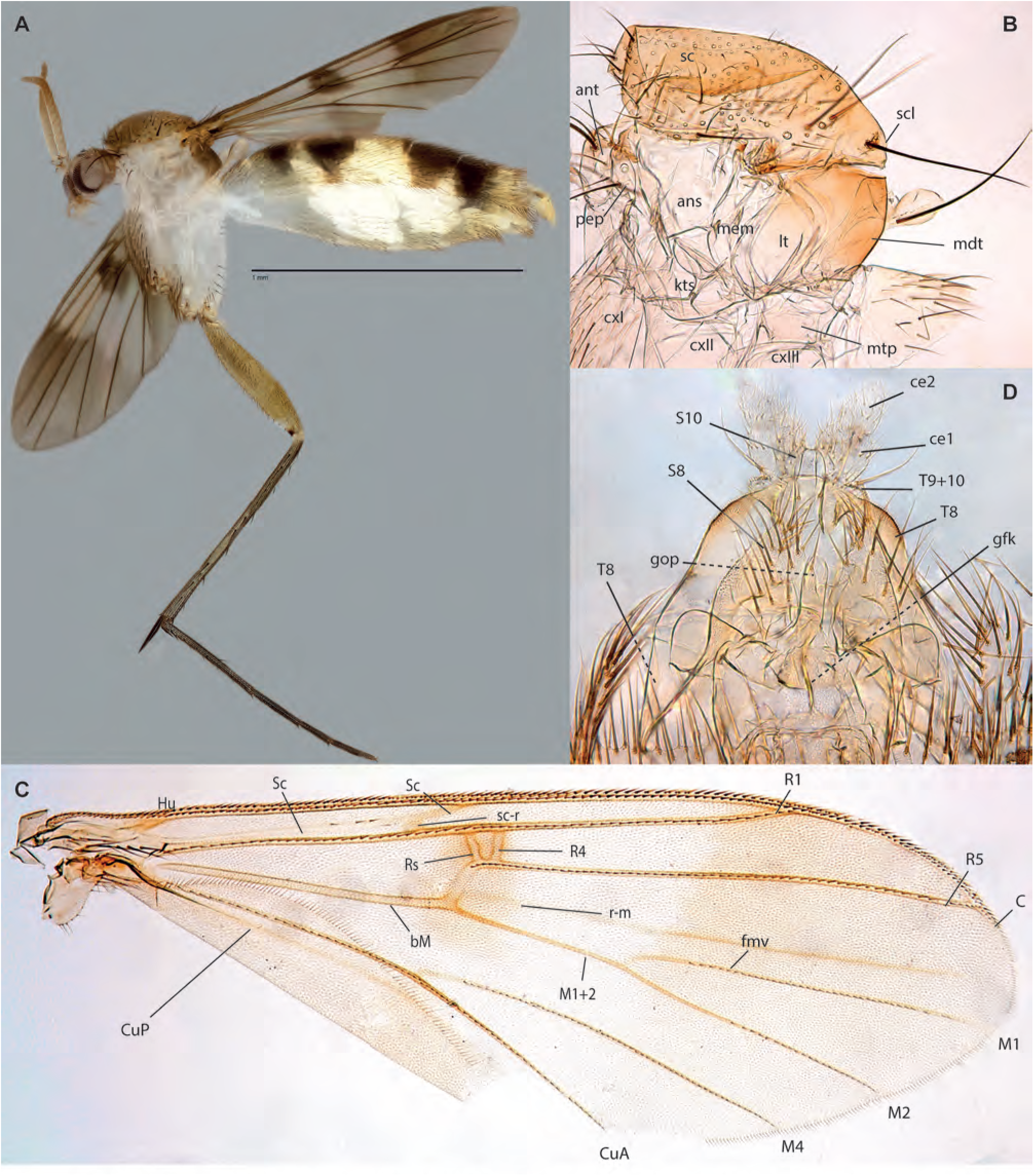
*Neoempheria cinkappur* Amorim & Oliveira, **sp. n.,** female holotype. **A.** Habitus, lateral view. **B.** Thorax, lateral view. **C.** Wing. **D.** Terminalia, ventral view.

**Figs. 84A-D.**
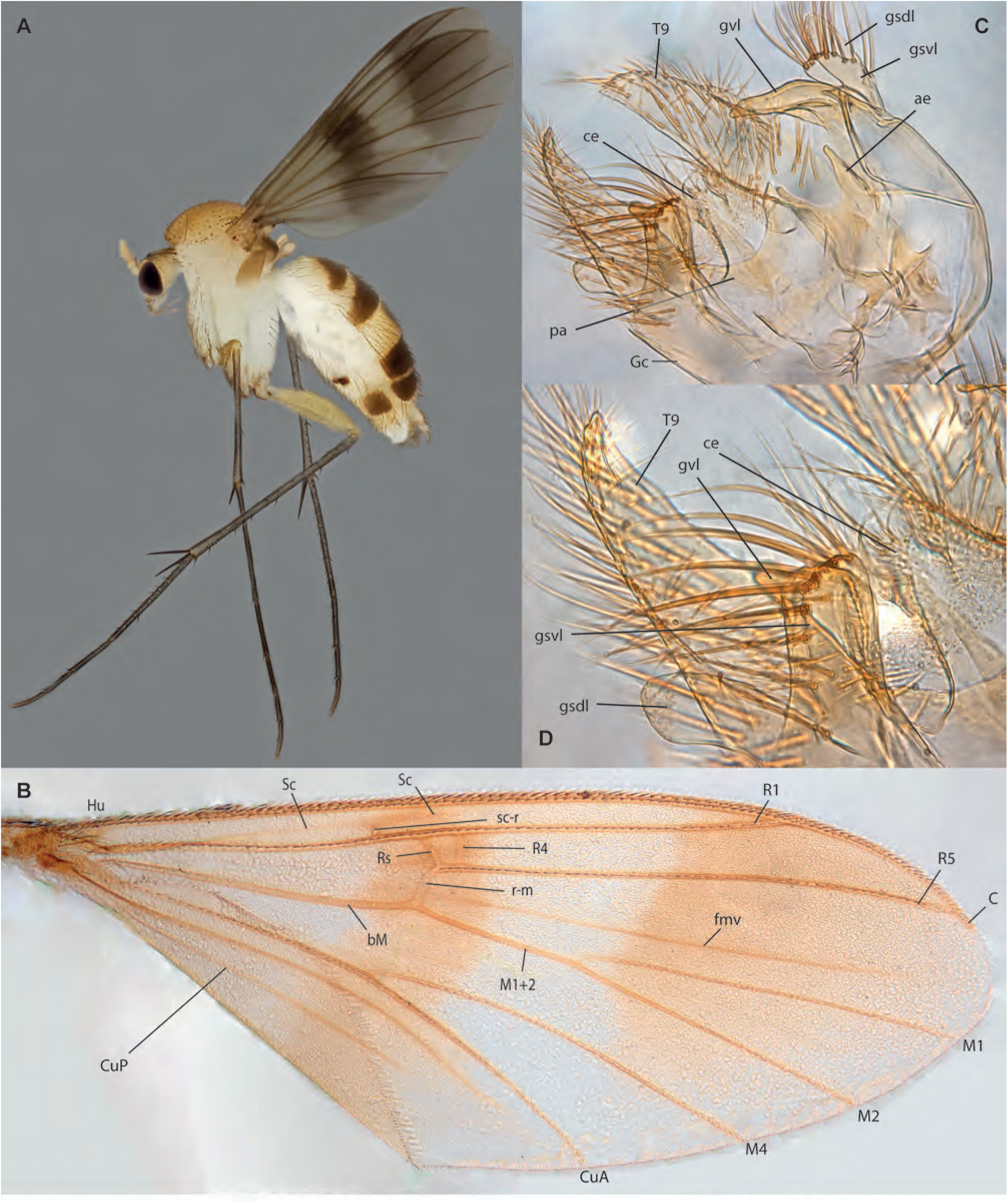
*Neoempheria temasek* Amorim & Oliveira, **sp. n. A.** Habitus, lateral view, male ZRCBDP0048695. **B.** Wing, male holotype. **C.** Male terminalia, ventral view, same. **D.** Detail of male terminalia, same.

**Figs. 85A-D.**
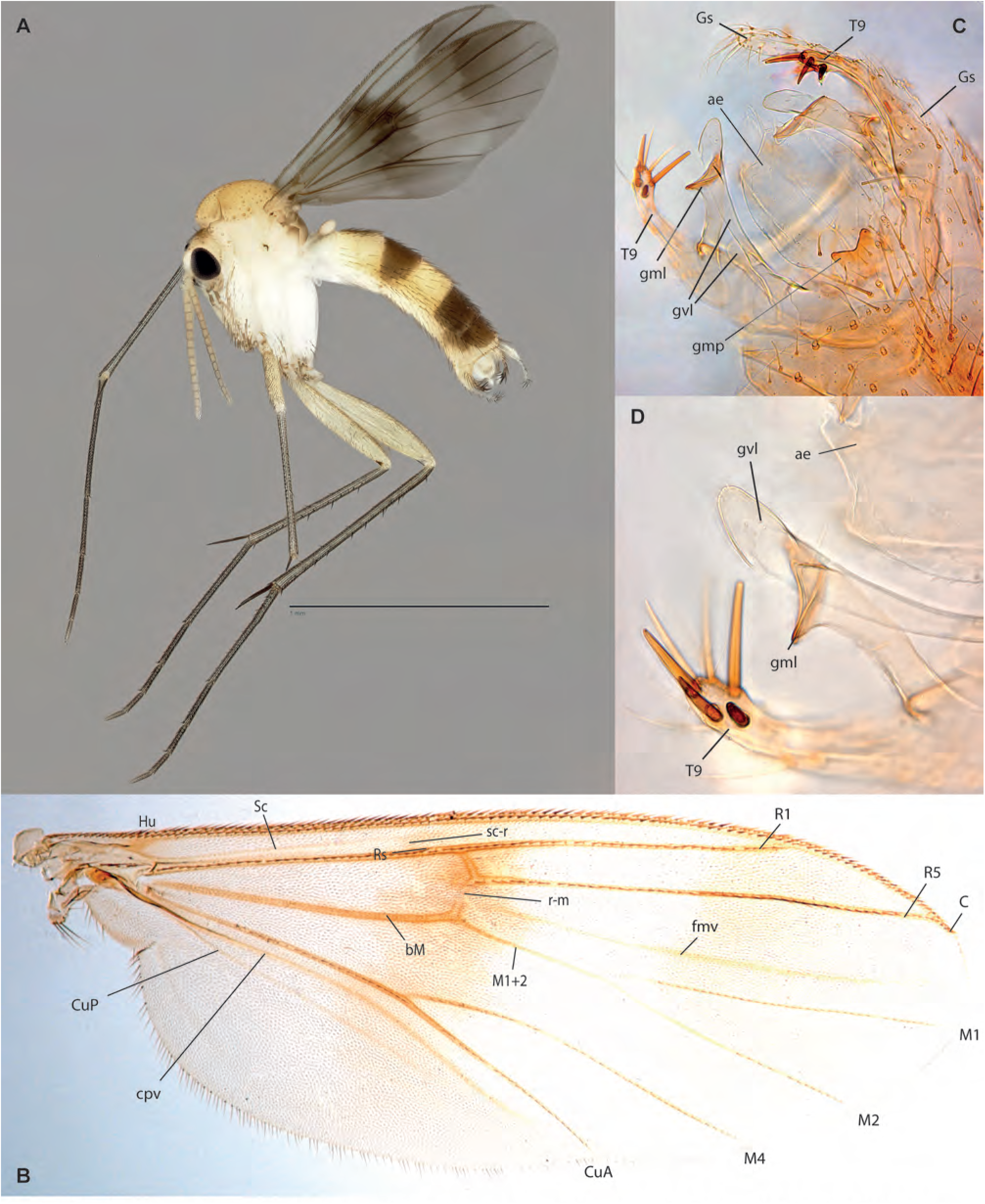
*Neoempheria polunini* Amorim & Oliveira, **sp. n. A.** Habitus, lateral view, male, ZRCBDP0049204. **B.** Wing, male holotype. **C.** Male terminalia, ventral view, same. **D.** Detail of terminalia, ventral view, same.

**Figs. 86A-C.**
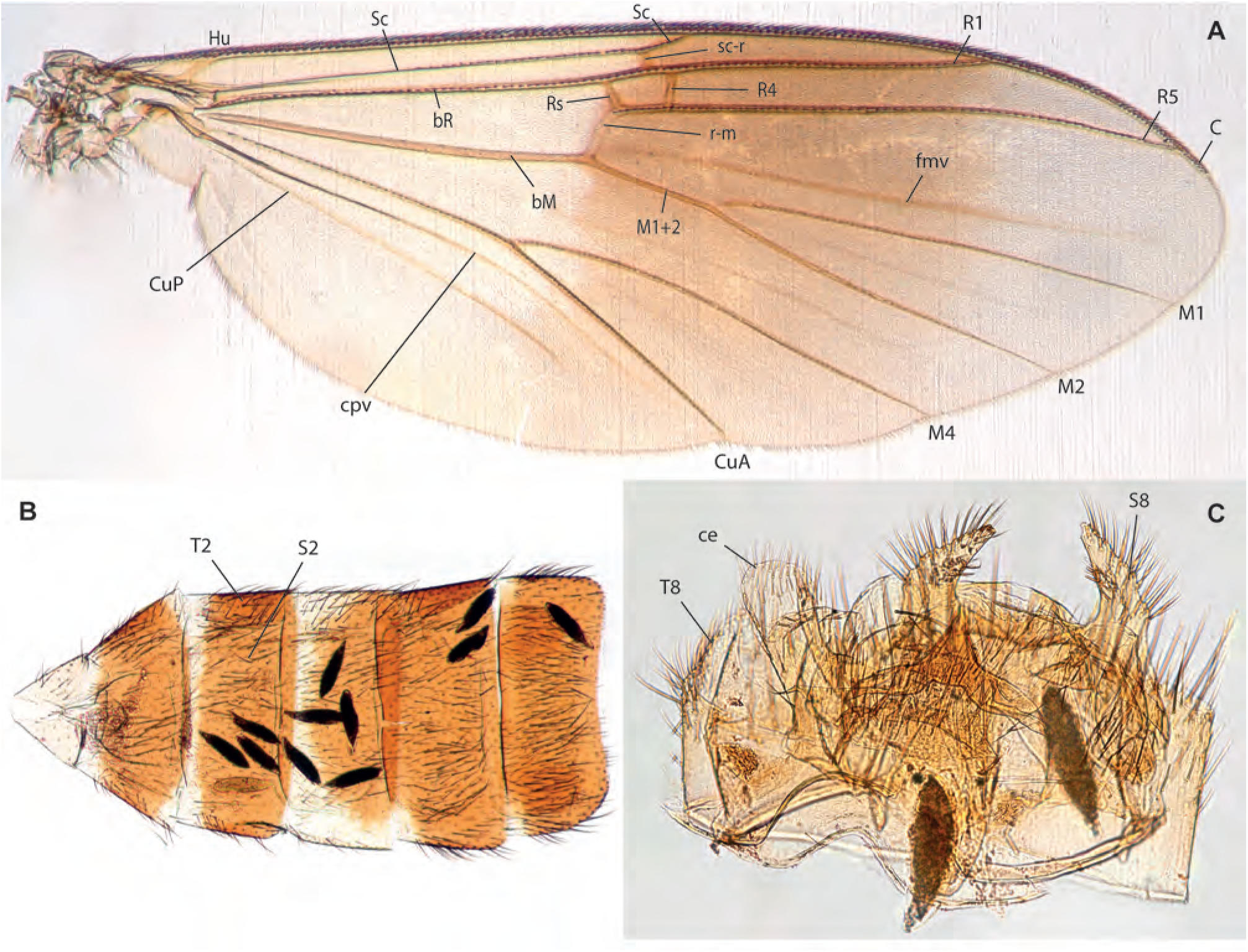
*Neoempheria fajar* Amorim & Oliveira, **sp. n.,** female holotype. **A.** Wing. **B.** Abdomen, ventral view. **C.** Terminalia, ventral view.

**Figs. 87A-D.**
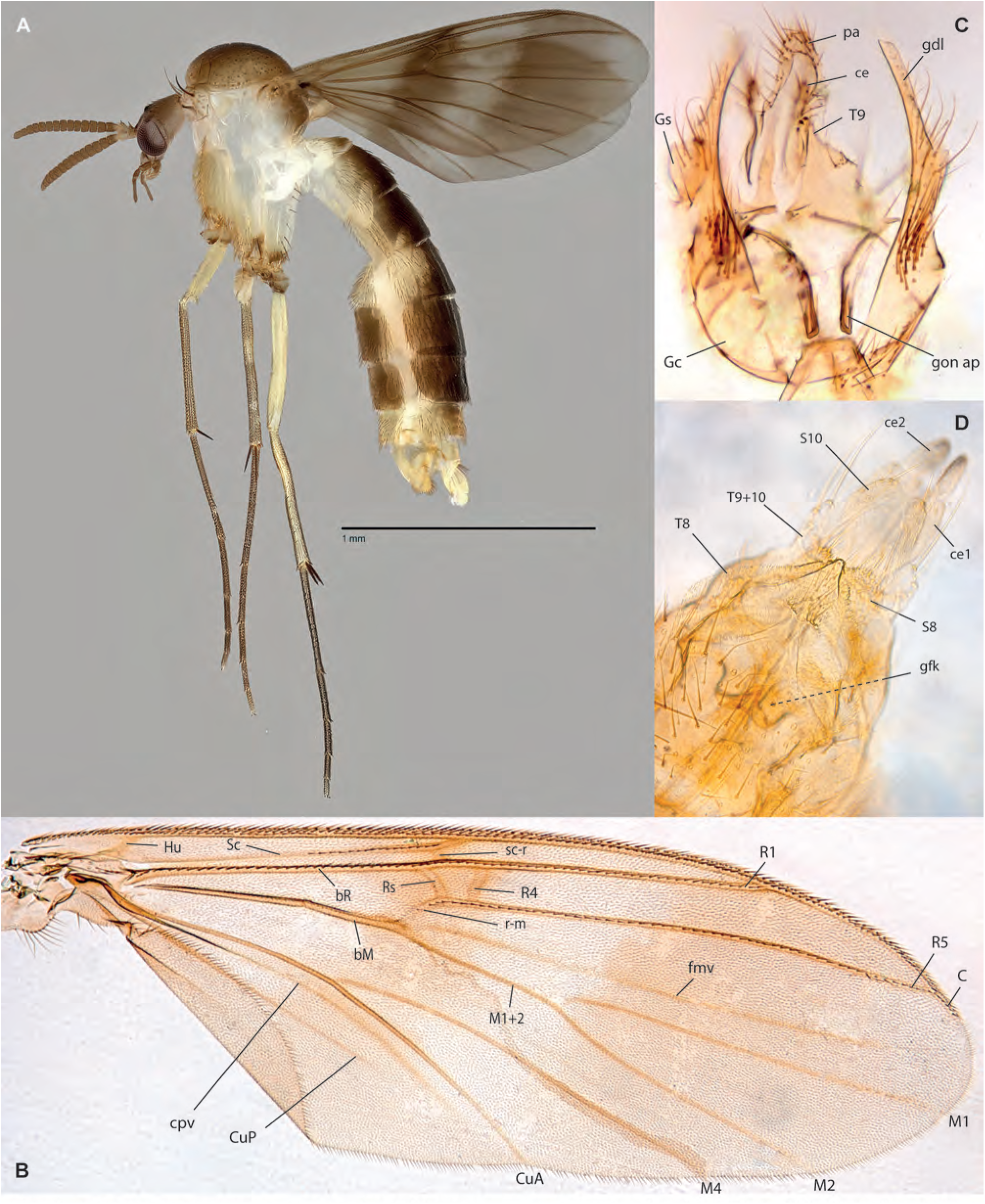
*Neoempheria riatanae* Amorim & Oliveira, **sp. n. A.** Habitus, lateral view, female, ZRCBDP0049247. **B.** Wing, female paratype ZRCBDP0047840. **C.** Male terminalia, dorsal view, holotype. **D.** Female terminalia, ventral view, female paratype ZRCBDP0047840.

**Figs. 88A-D.**
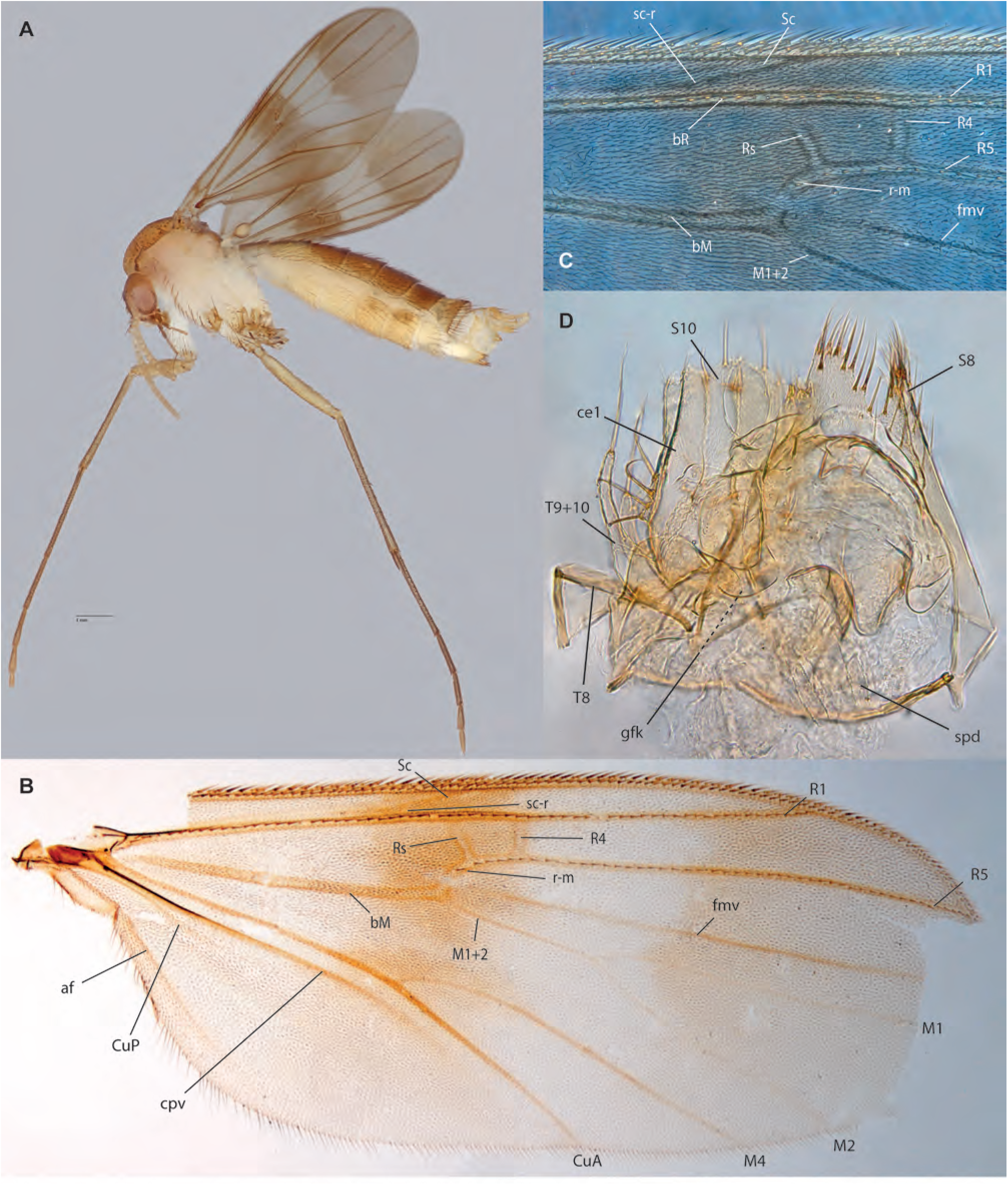
*Neoempheria* sp. B, female, ZRCBDP0047779. **A.** Habitus, lateral view. **B.** Wing. **C.** Detail of anterior wing margin under phase contrast. **D.** Female terminalia, ventral view. **Mycetophilinae Exechiini**

## Mycetophilinae Exechiini

**Figs. 89A-G.**
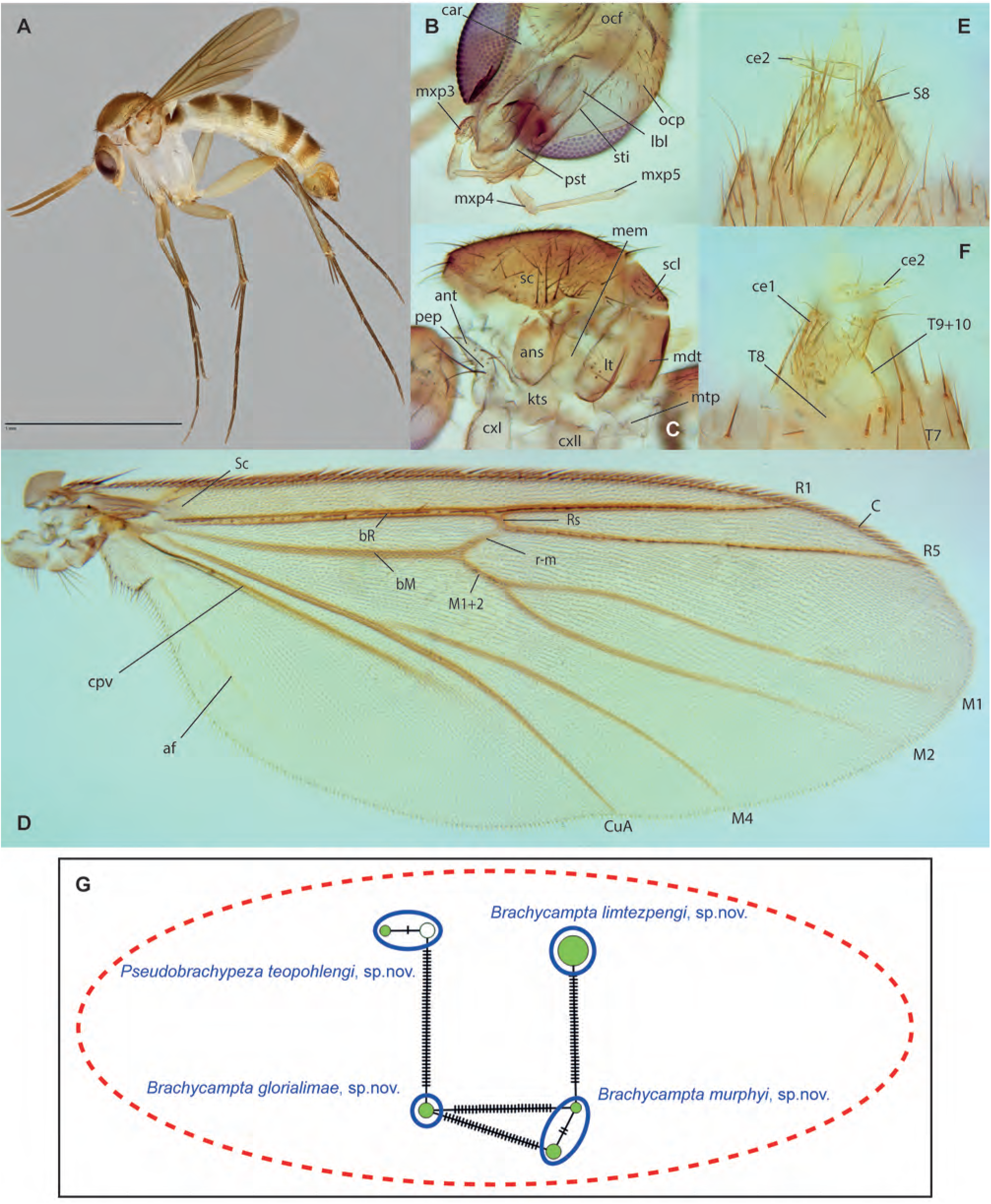
*Brachycampta glorialimae* Amorim & Oliveira, **sp. n. A.** Habitus, lateral view, male holotype. **B.** Head, ventral view, same. **C.** Thorax, lateral view, same. **D.** Wing, same. **E.** Female terminalia, dorsal view, paratype ZRCBDP0049093. **F.** Female terminalia, dorsal view, same. **G.** Haplotype network for *Brachycampta*+*Rymosia*.

**Figs. 90A-B.**
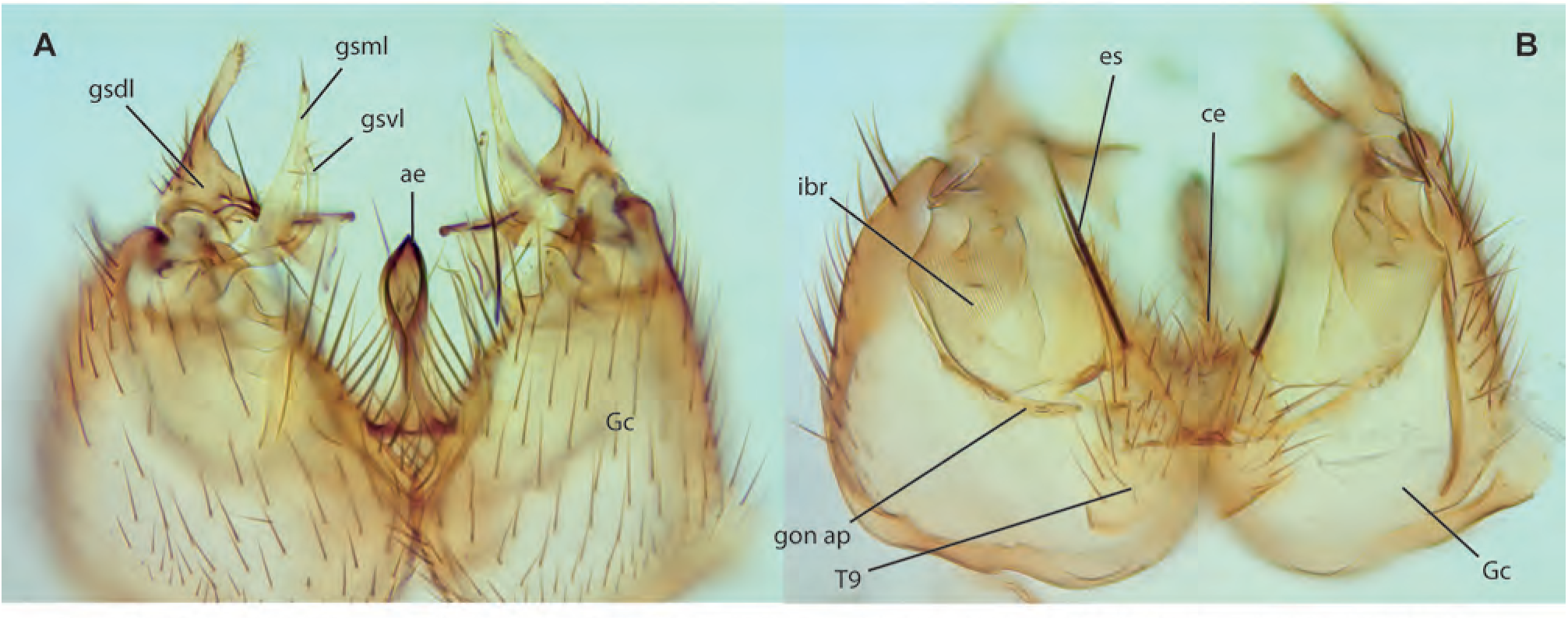
*Brachycampta glorialimae* Amorim & Oliveira, **sp. n.** holotype. **A.** Male terminalia, dorsal view. **B.** Male terminalia, dorsal view.

**Figs. 91A-G.**
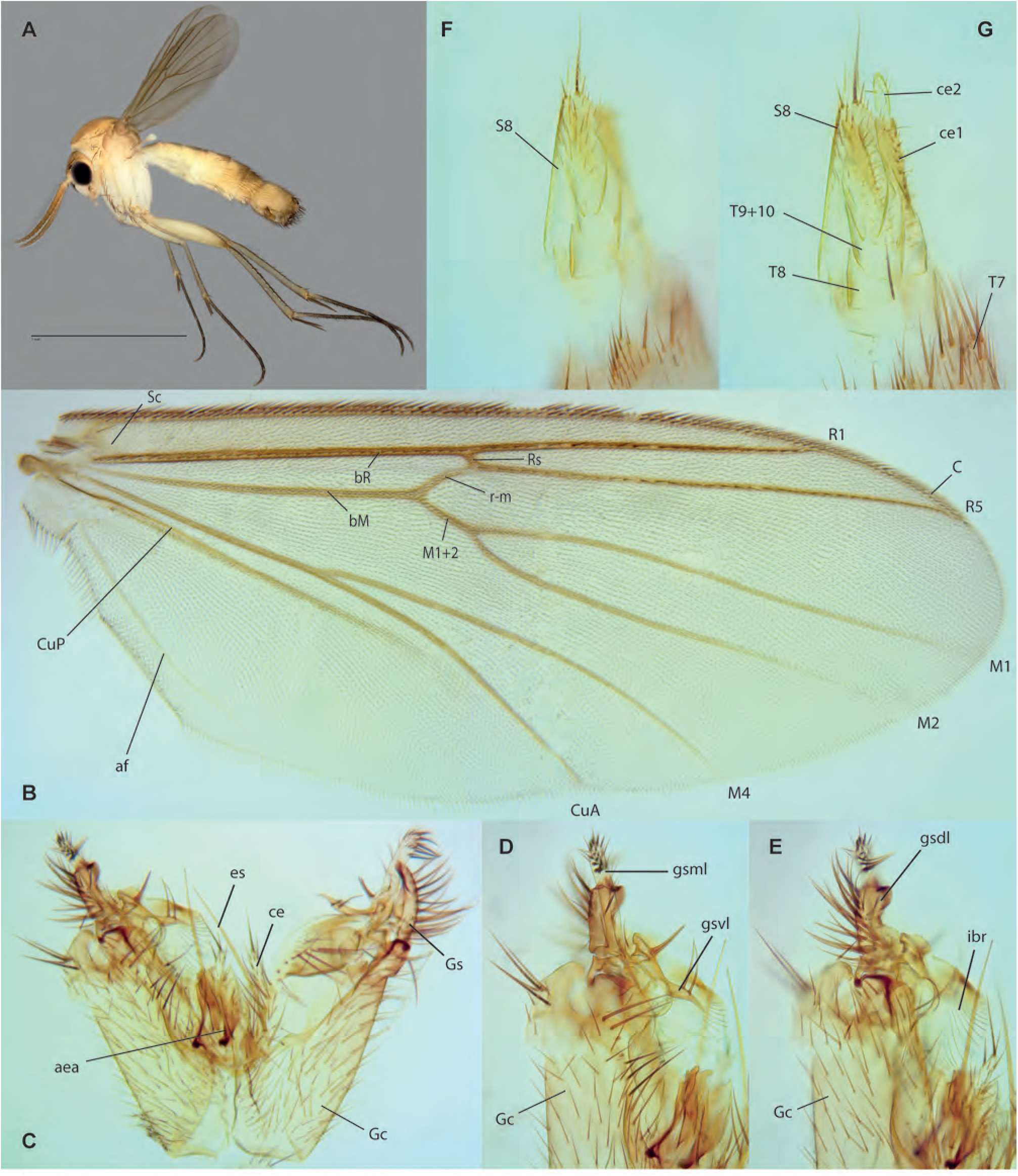
*Brachycampta murphyi* Amorim & Oliveira, **sp. n. A.** Habitus, lateral view, male paratype ZRCBDP0048976. **B.** Wing, male holotype. **C.** Male terminalia, ventral view, same. **D.** Detail of gonostyle, ventral view, same. **E.** Detail of gonostylus, dorsal view, same. **F.** Female terminalia, ventral view, paratype ZRCBDP0048669. **G.** Female terminalia, ventral view, same.

**Figs. 92A-E.**
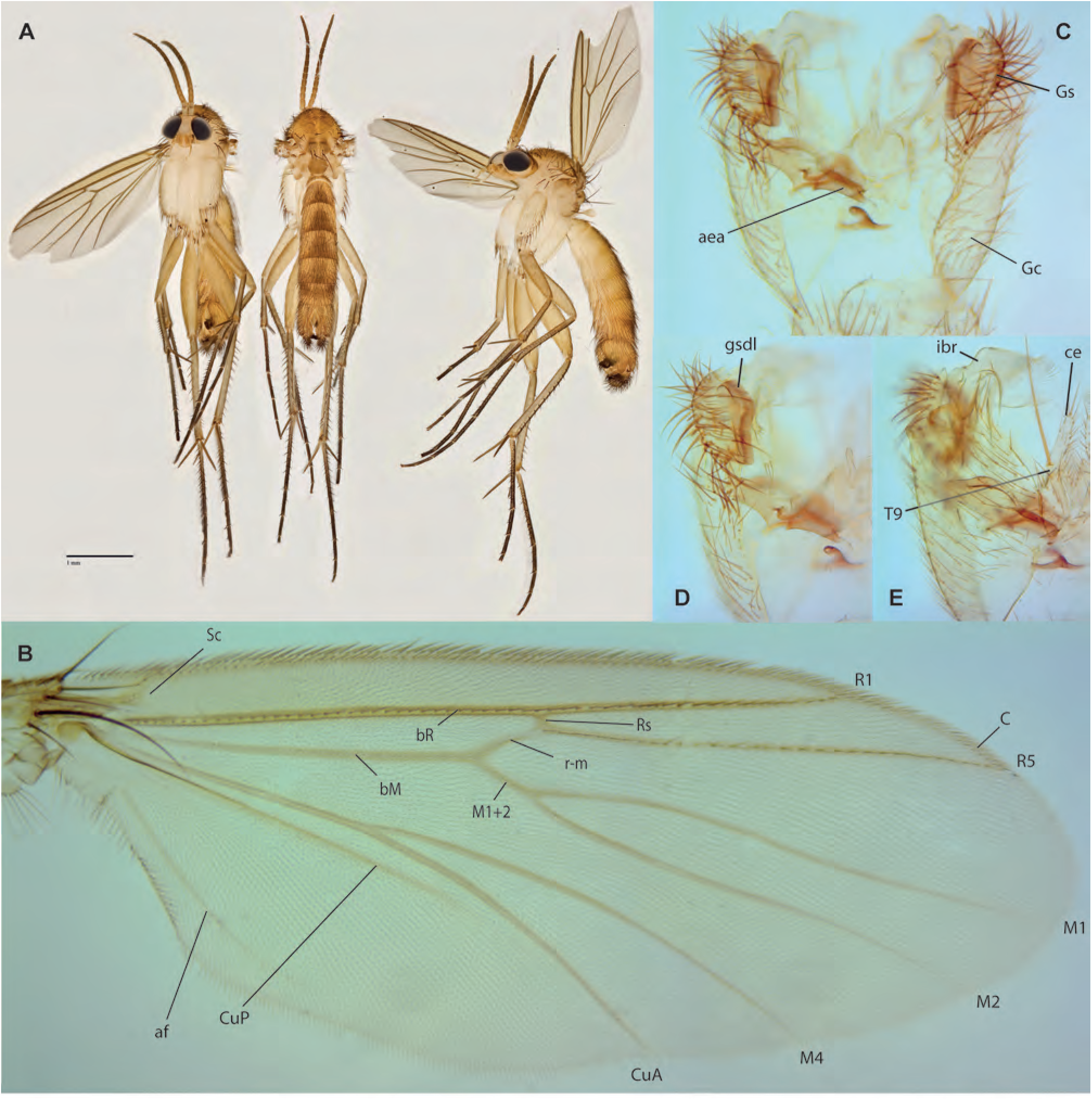
*Brachycampta limtzepengi* Amorim & Oliveira, **sp. n. A.** Habitus, lateral view, male paratype ZRCBDP0048511. **B.** Wing, male holotype. **C.** Male terminalia, ventral view, same. **D.** Detail of gonostyle, ventral view, same. **E.** Detail of gonostyle, dorsal view, same.

**Figs. 93A-E.**
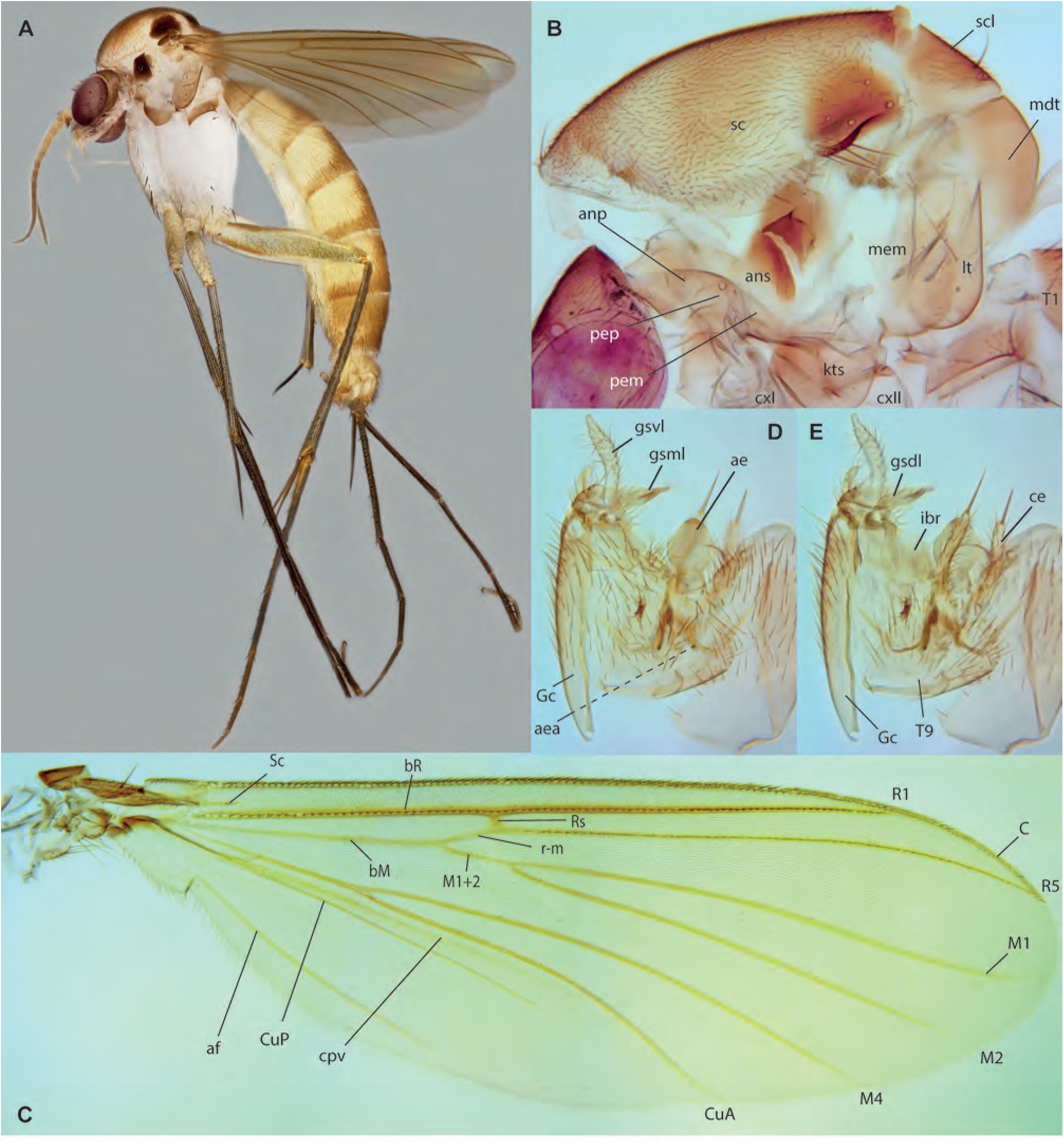
*Rymosia teopohlengi* Amorim & Oliveira, **sp. n.,** male holotype. **A.** Habitus, lateral view. **B.** Thorax, lateral view. **C.** Wing. **D.** Terminalia, ventral view. **E.** Terminalia, dorsal view.

**Figs. 94A-C.**
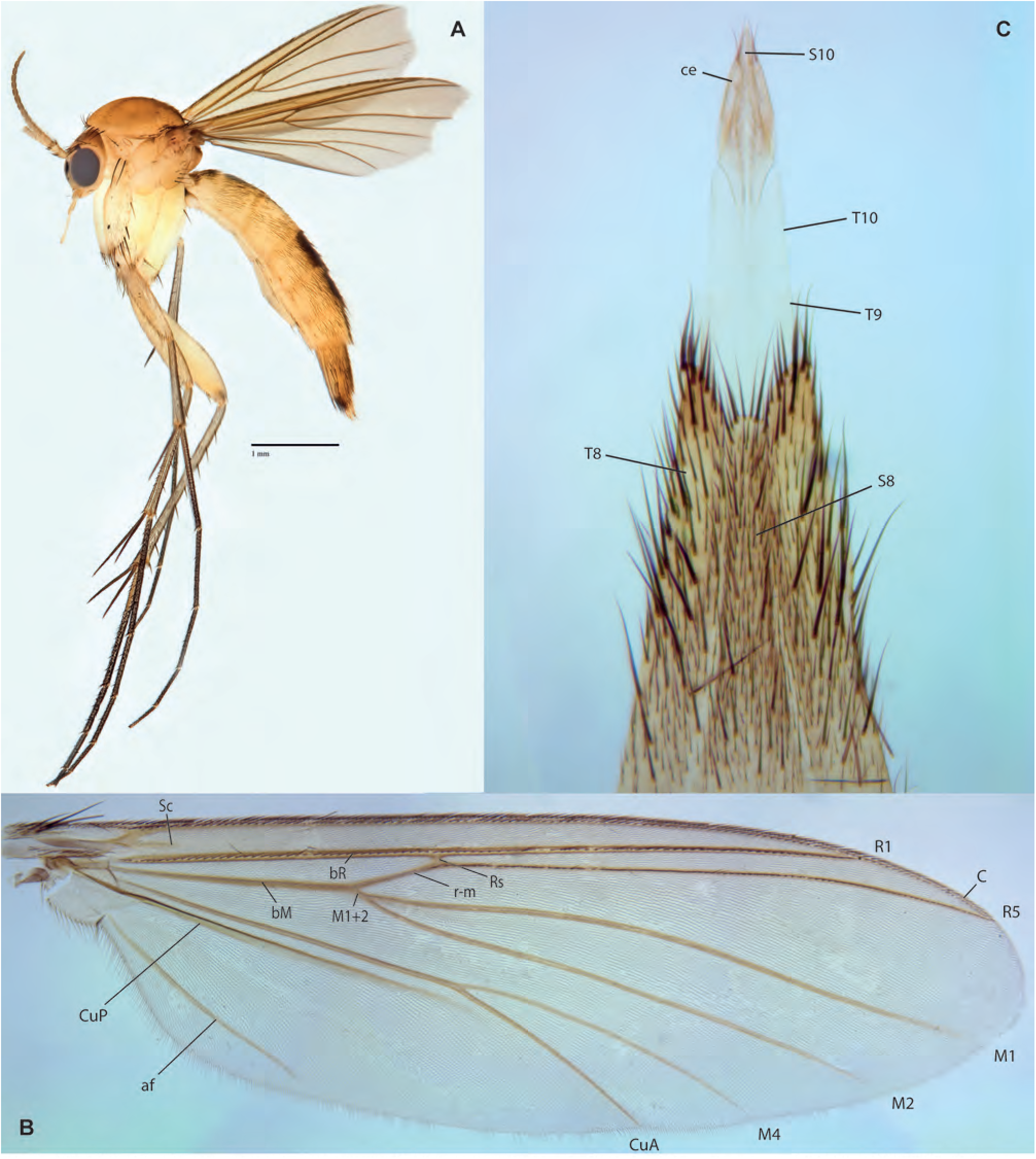
*Exechia tanswiehiani* Amorim & Oliveira, **sp. n. A.** Habitus, lateral view, female paratype ZRCBDP0048514. **B.** Wing, male holotype. **C.** Female terminalia, ventral view, paratype ZRCBDP0048668.

**Figs. 95A-C.**
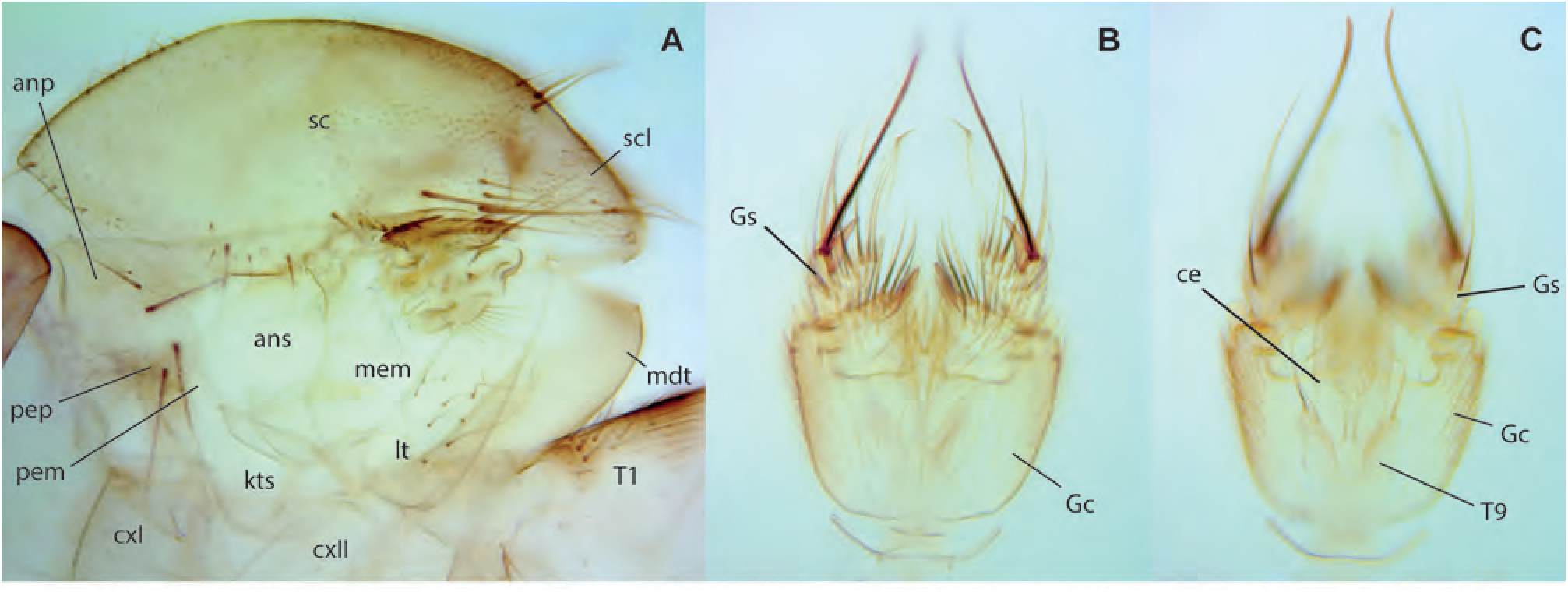
*Exechia tanswiehiani* Amorim & Oliveira, **sp. n.,** male holotype. **A.** Thorax, lateral view. **B.** Terminalia, ventral view. **C.** Terminalia, dorsal view.

**Figs. 96A-G.**
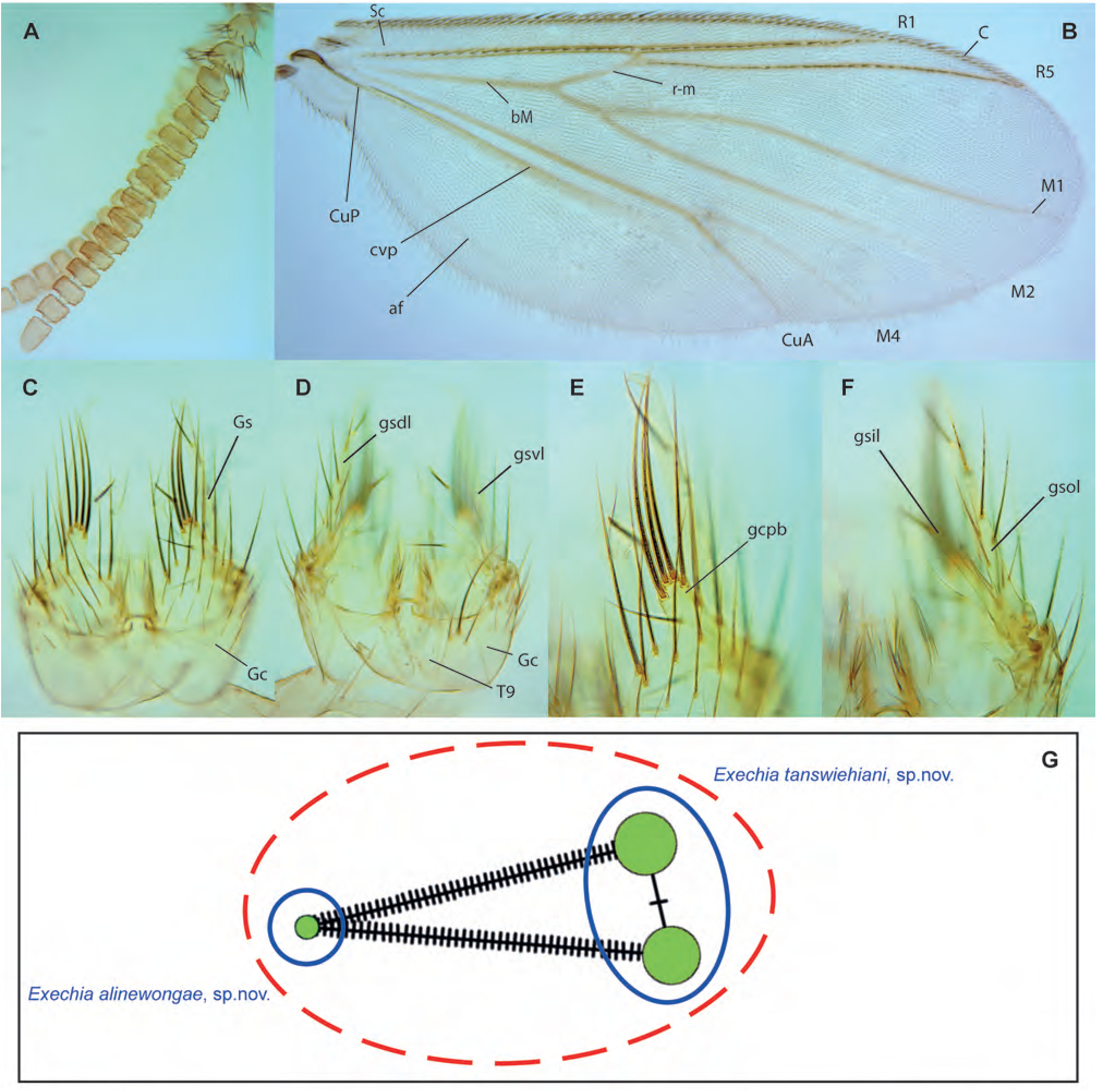
*Exechia alinewongae* Amorim & Oliveira, **sp. n.,** male holotype. **A.** Antenna, lateral view. **B.** Wing. **C.** Male terminalia, ventral view. **D.** Gonocoxite posterior margin, ventral view. **E.** Gonostylus, dorsal view. **G.** Haplotype network for *Exechia*. **Mycetophilini**

## Mycetophilini

**Figs. 97A-H.**
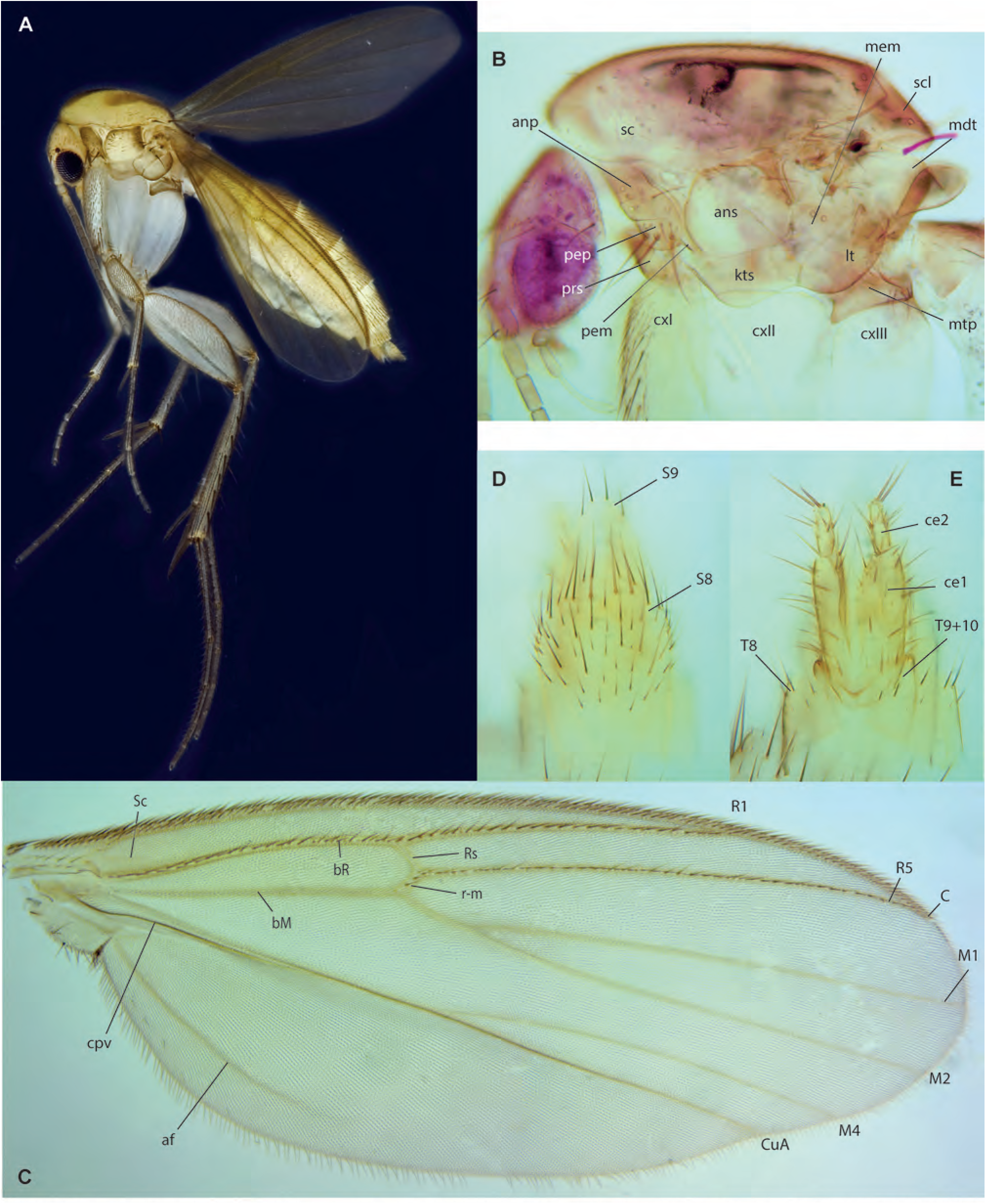
*Mycetophila chngseoktinae* Amorim & Oliveira, **sp. n. A.** Habitus, lateral view, female paratype ZRCBDP0048447. **B.** Head and thorax, lateral view, male paratype ZRCBDP0049191. **C.** Wing, same. **D.** Female terminalia, ventral view, paratype ZRCBDP0048674. **E.** Female terminalia, dorsal view, same.

**Figs. 98A-D.**
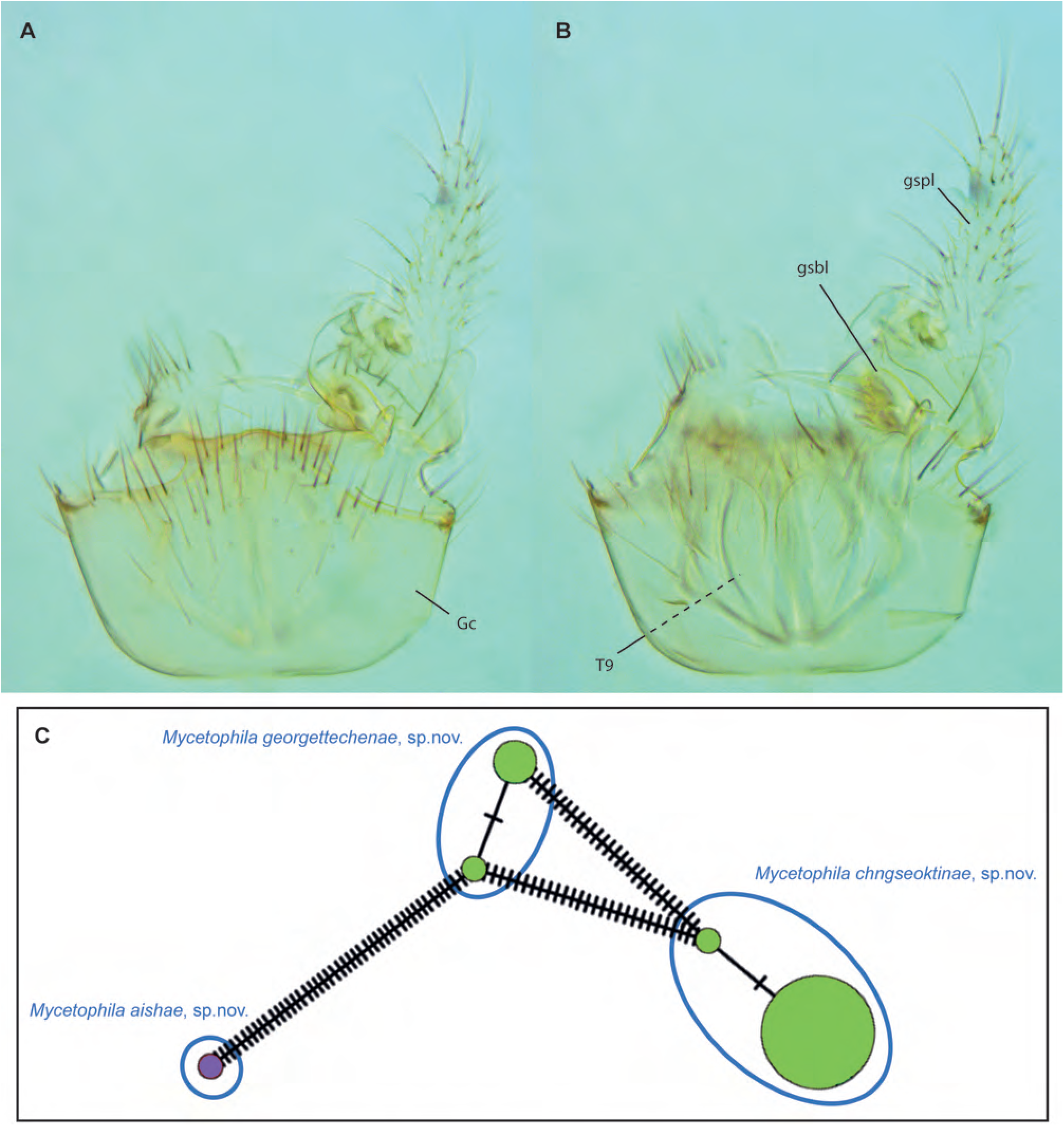
*Mycetophila chngseoktinae* Amorim & Oliveira, **sp. n. A.** Male terminalia, ventral view, male paratype ZRCBDP0049191. **B.** Male terminalia, dorsal view, same. **C.** Haplotype network for *Mycetophila*.

**Figs. 99A-F.**
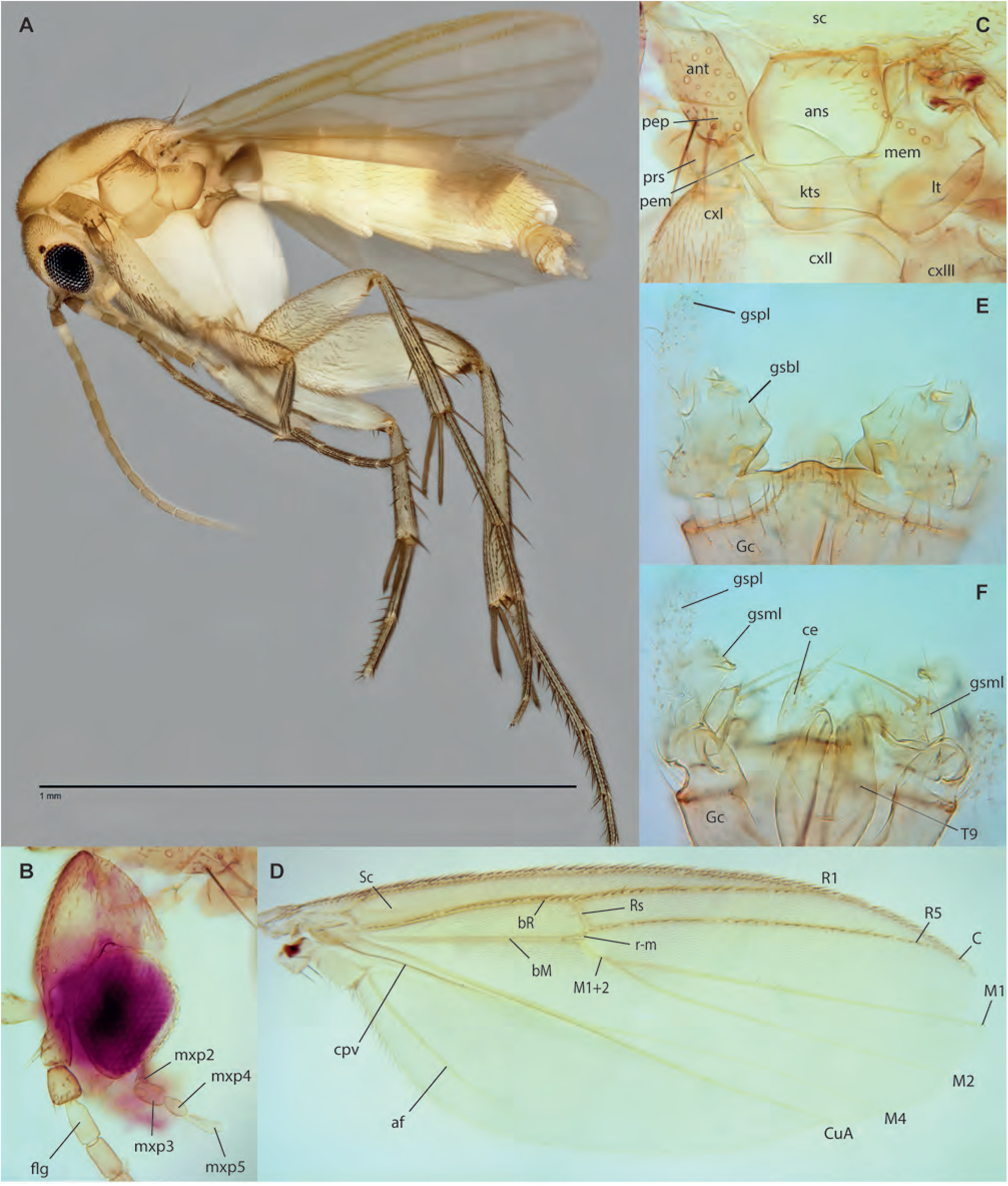
*Mycetophila georgettechenae* Amorim & Oliveira, **sp. n.,** male holotype. **A.** Habitus, lateral view. **B.** Head, lateral view. **C.** Thorax, lateral view. **D.** Wing. **E.** Male terminalia, ventral view. **F.** Male terminalia, dorsal view.

**Figs. 100A-F.**
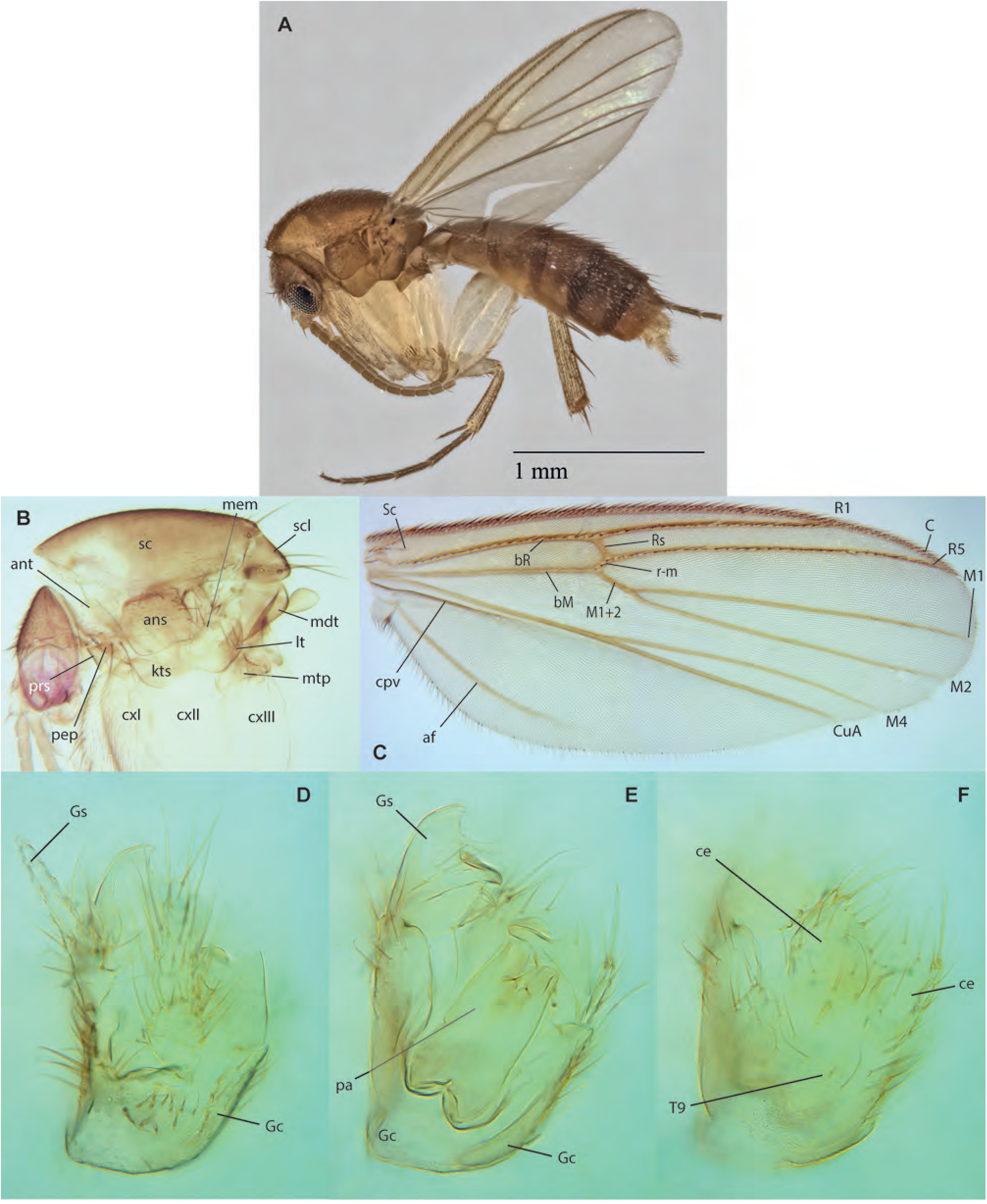
*Mycetophila aishae* Amorim & Oliveira, **sp. n. A.** Habitus, lateral view, male paratype ZRCBDP0133534. **B.** Thorax, lateral view, male holotype. **C.** Wing, same. **D.** Male terminalia, mid-section. **E.** Male terminalia, ventral view. **F.** Male terminalia, dorsal view.

**Fig. 101.**
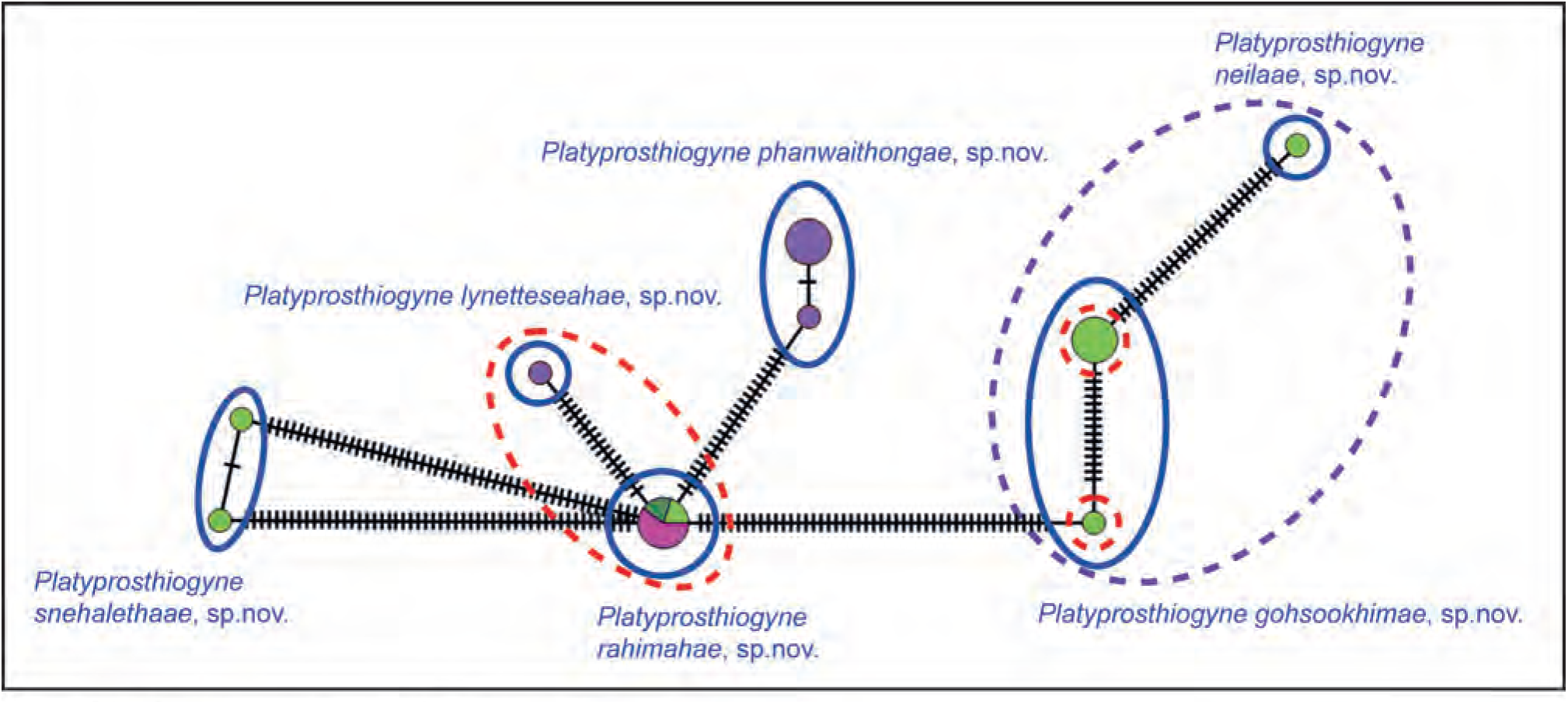
Haplotype network for *Platyprosthiogyne*.

**Figs. 102A-F.**
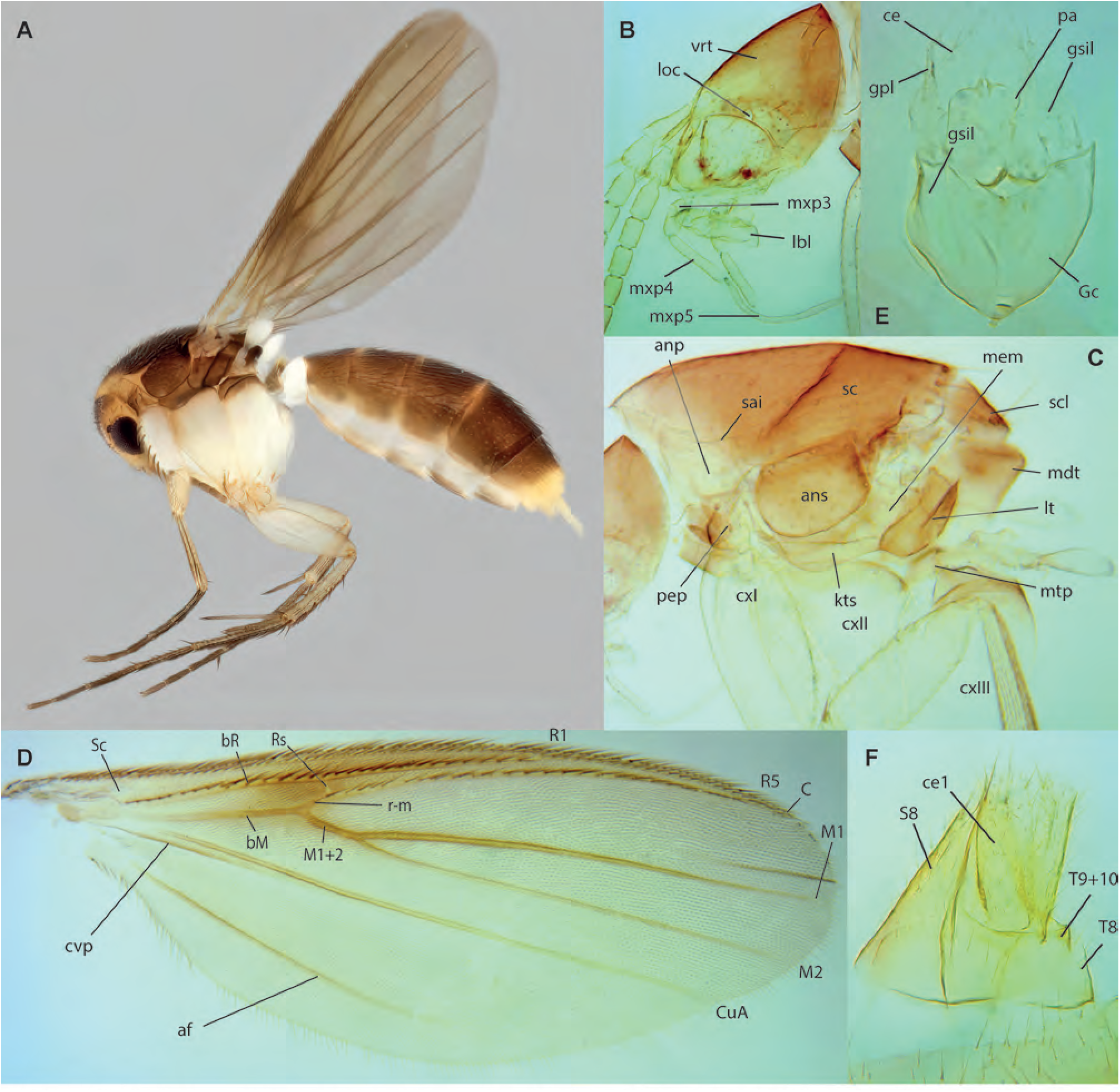
*Platyprosthiogyne phanwaithongae* Amorim & Oliveira, **sp. n. A.** Habitus, lateral view, female paratype ZRCBDP0048145. **B.** Head, lateral view, male holotype. **C.** Thorax, lateral view, same. **D.** Wing, same (wing folded close to margin anteriorly to M_1_). **E.** Male terminalia, ventral view, same. **F.** Female terminalia, laterodorsal view, female paratype, ZRCBDP0048144.

**Figs. 103A-I.**
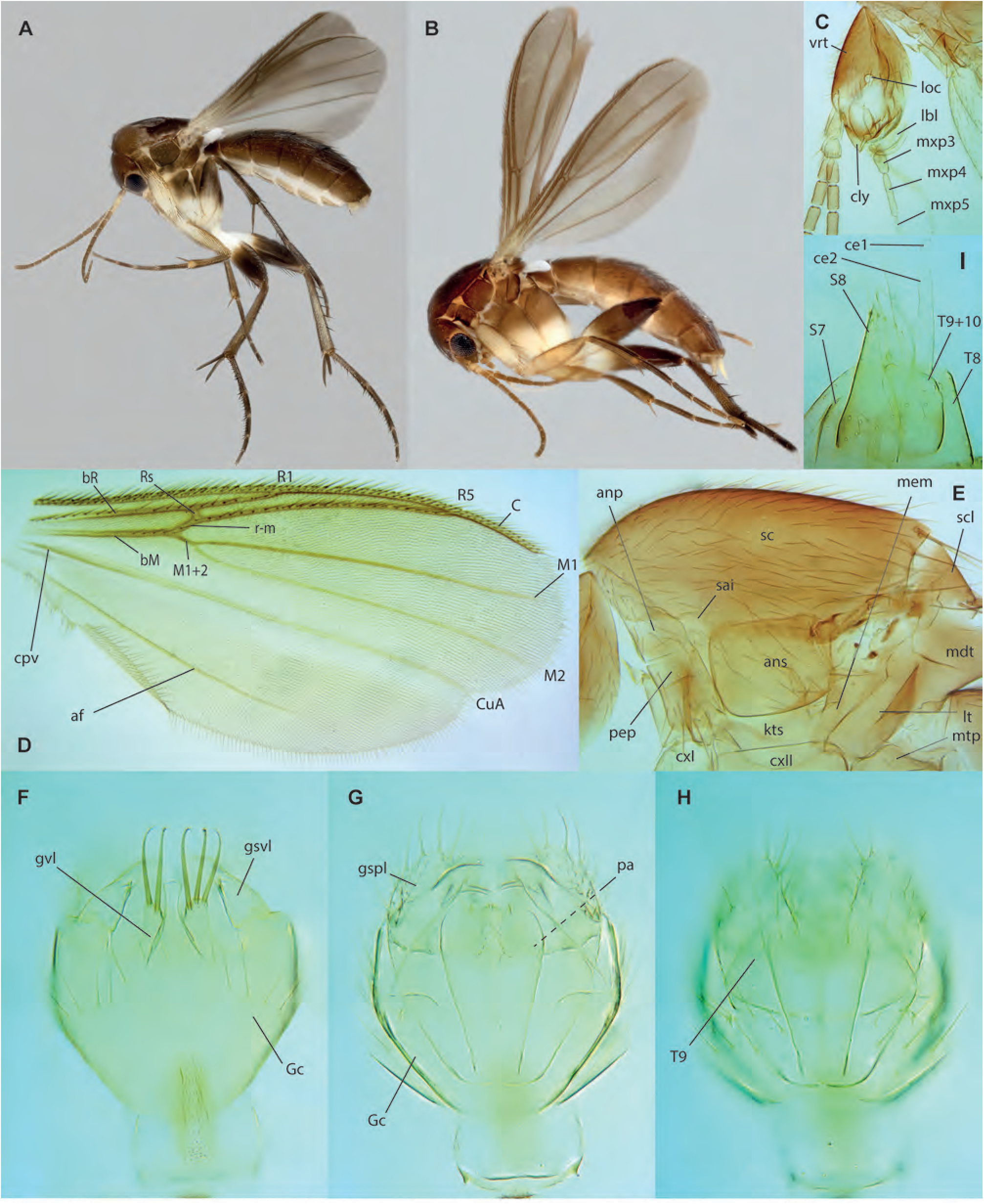
*Platyprosthiogyne gohsookhimae* Amorim & Oliveira, **sp. n. A.** Habitus, lateral view, male holotype ZRCBDP0048727. **B.** Habitus, lateral view, female paratype, ZRCBDP0047897. **C.** Head, lateral view, male paratype, ZRCBDP0048965. **D.** Wing, same. **E.** Thorax, lateral view, same. **F.** Male terminalia, ventral view, same. **G.** Male terminalia, mid-section, same. **H.** Male terminalia, dorsal view, same. **I.** Female terminalia, lateral view, paratype, ZRCBDP0047897.

**Figs. 104A-D.**
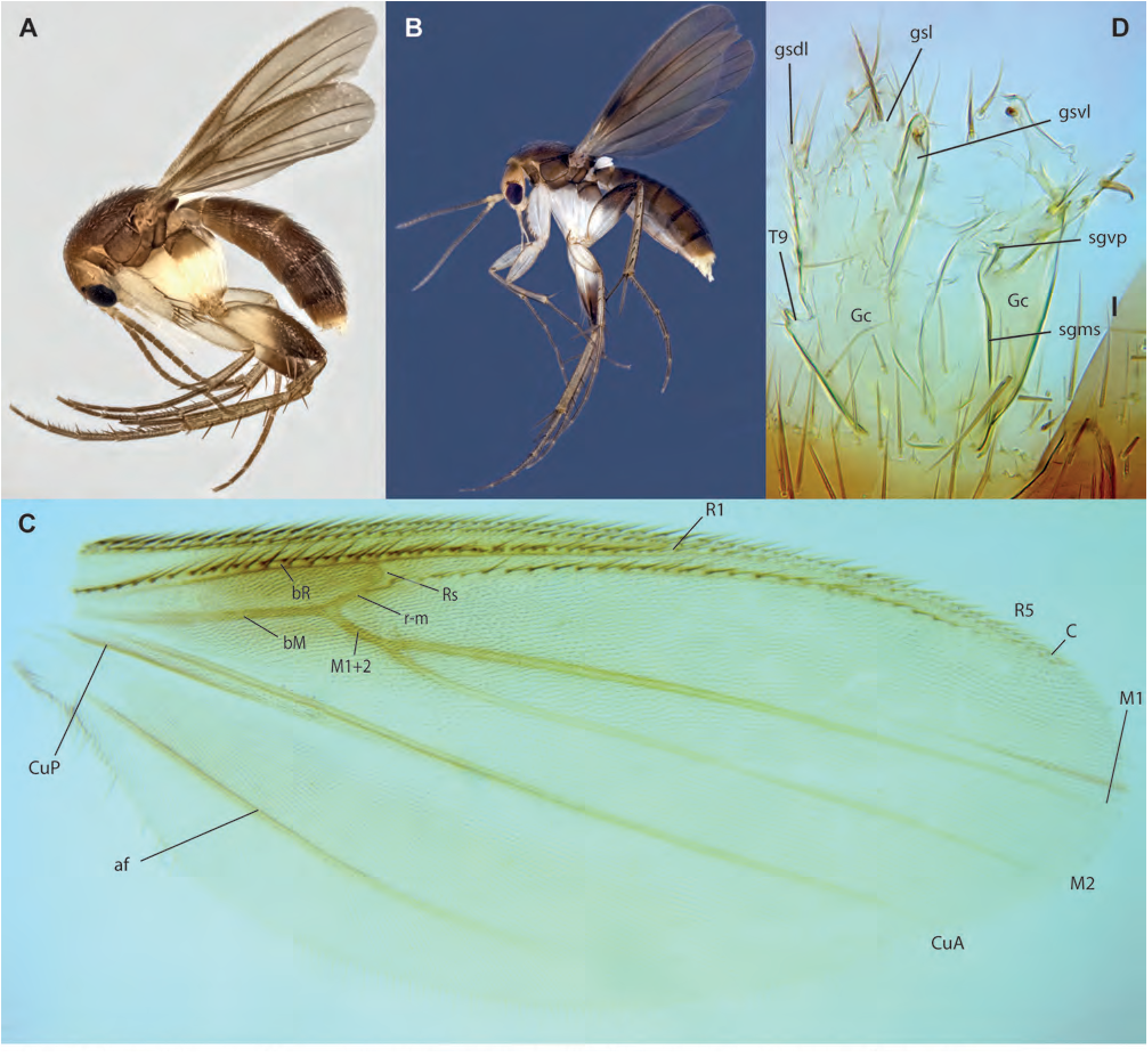
*Platyprosthiogyne rahimahae* Amorim & Oliveira, **sp. n.** . **A.** Habitus, lateral view, male, paratype ZRCBDP0155018. **B.** Habitus, lateral view, female, ZRCBDP0048428. **C.** Wing, male holotype. **D.** Male terminalia, ventral view, paratype ZRCBDP0049342.

**Figs. 105A-B.**
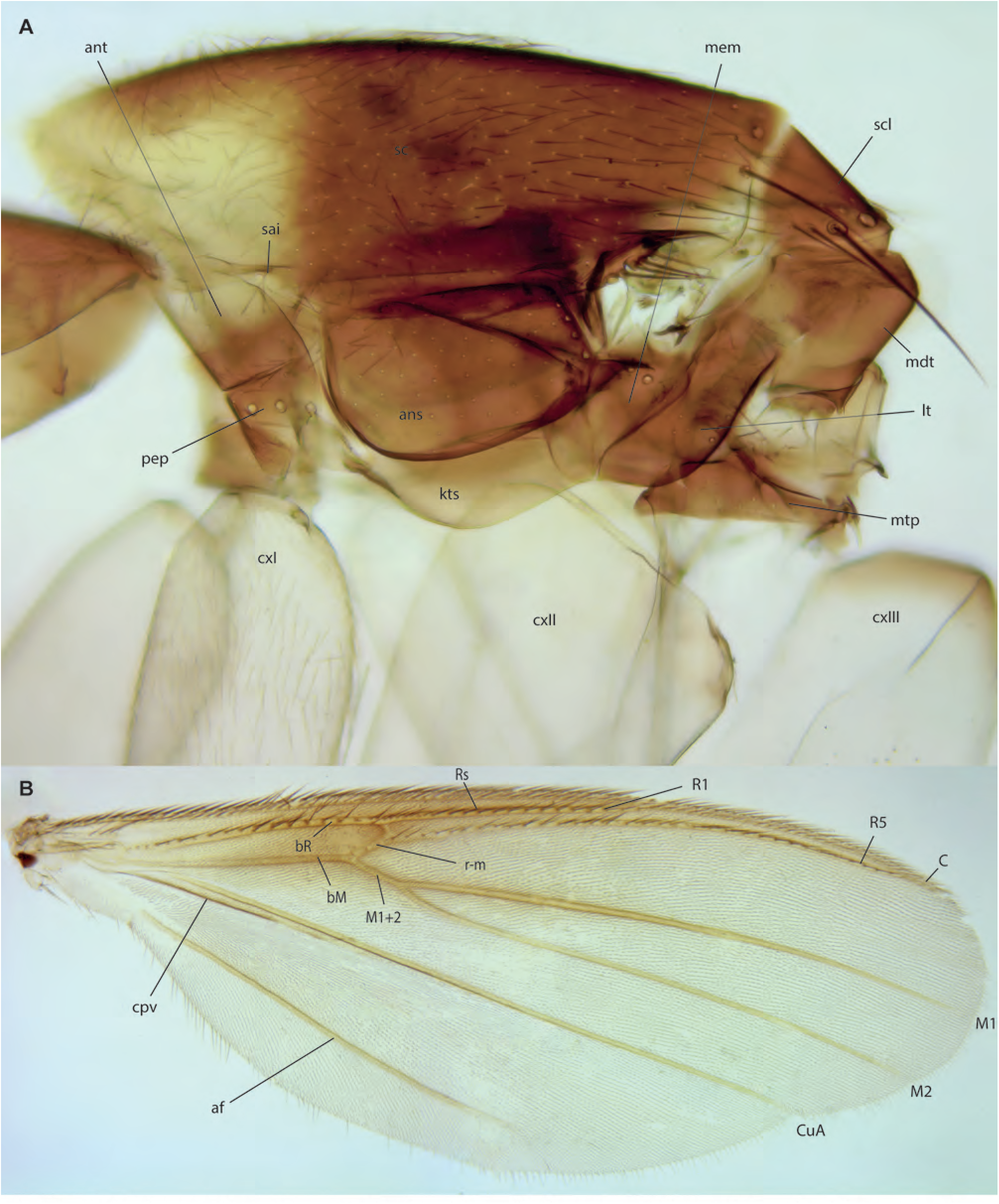
*Platyprosthiogyne lynetteseahae* Amorim & Oliveira, **sp. n.,** holotype. **A.** Head and thorax, lateral view. **B.** Wing.

**Figs. 106A-E.**
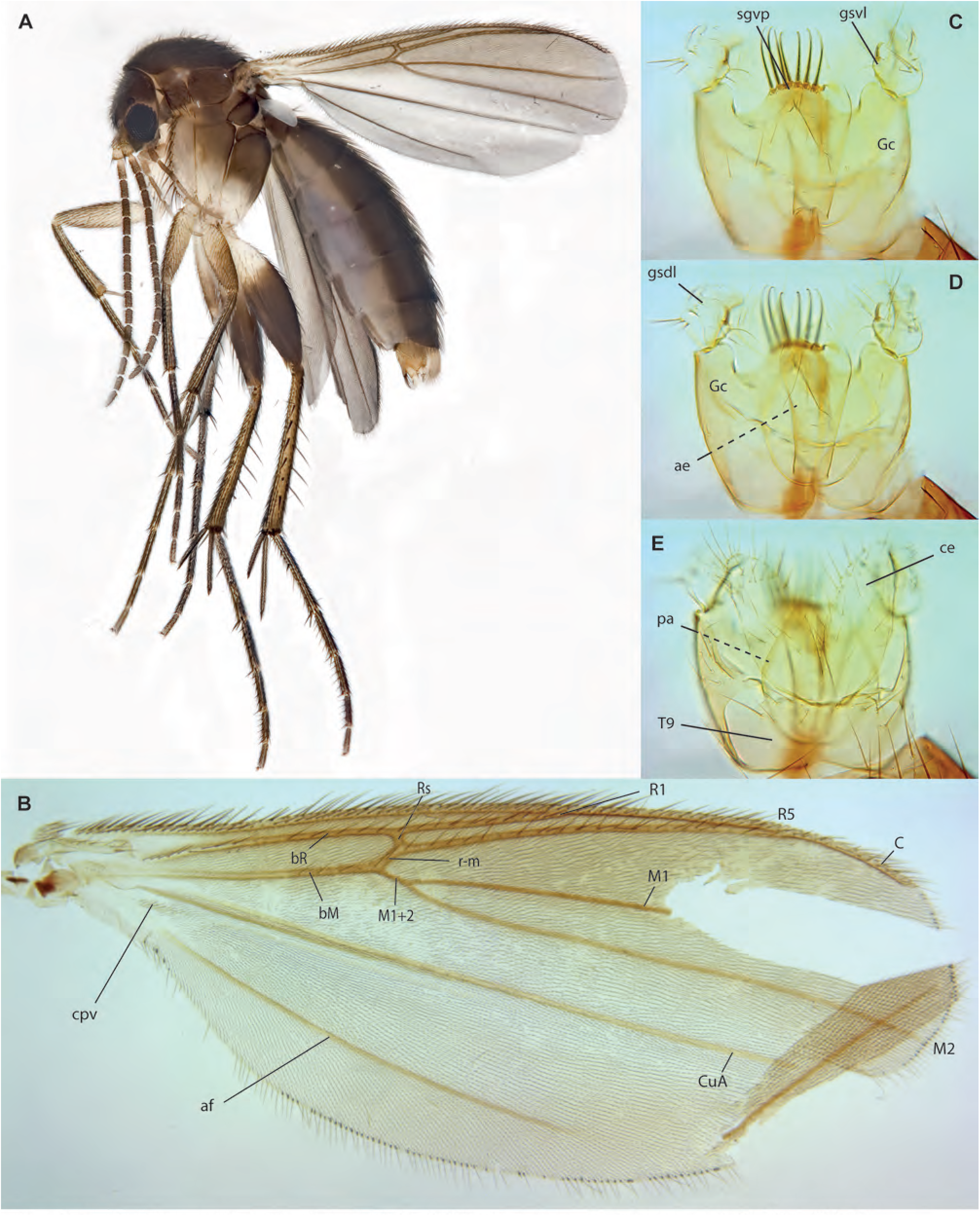
*Platyprosthiogyne neilaae* Amorim & Oliveira, **sp. n.,** male holotype. **A.** Habitus, lateral view. **B.** Wing. **C.** Terminalia, ventral view. **D.** Terminalia, dorsal view. **E.** Terminalia, mid-section.

**Figs. 107A-D.**
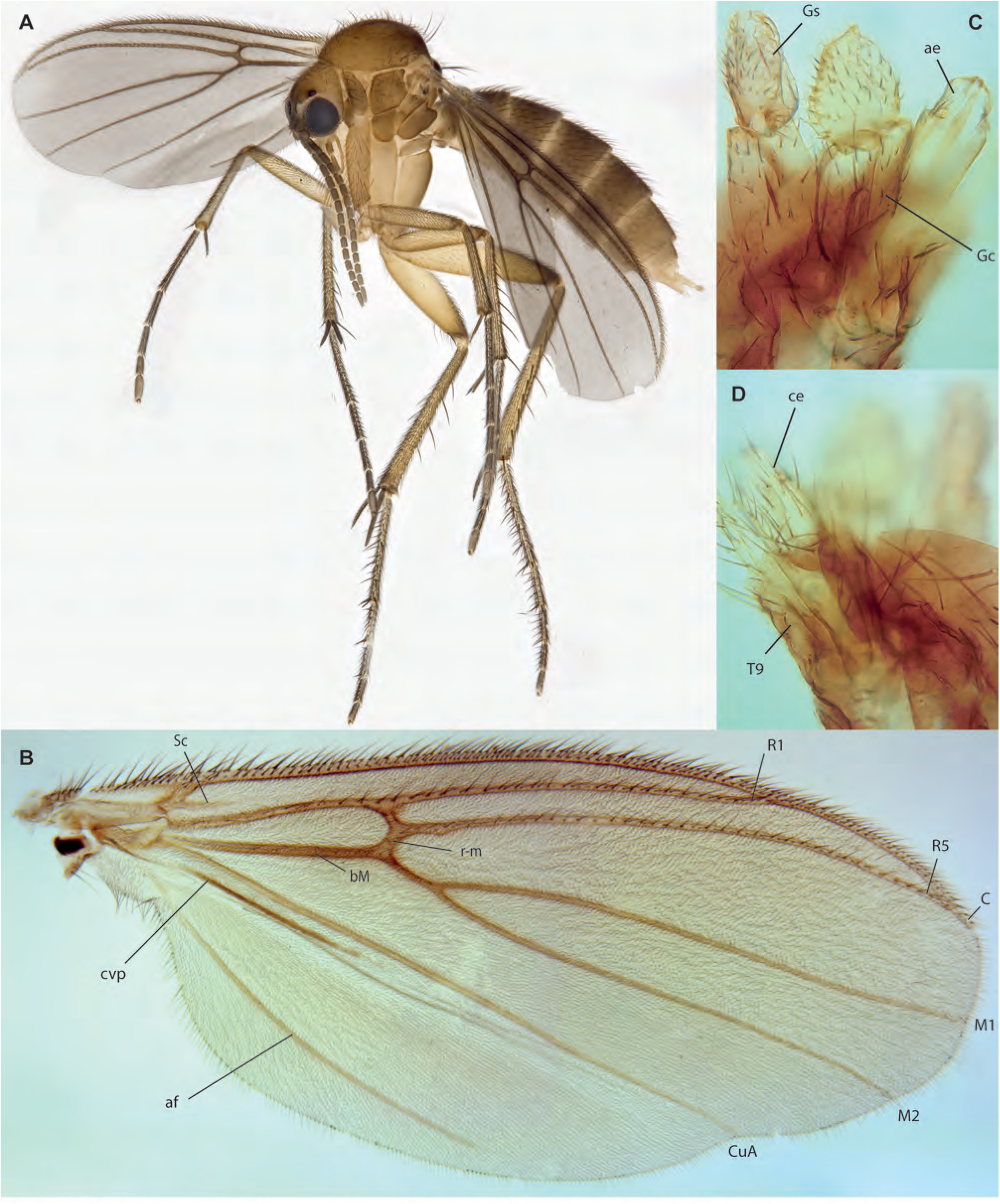
*Platyprosthiogyne snehalethaae* Amorim & Oliveira, **sp. n.** . **A.** Habitus, lateral view, female paratype ZRCBDP0048566. **B.** Wing, male holotype. **C.** Male terminalia, ventral view, same. **D.** Male terminalia, dorsal view, same.

**Figs. 108A-E.**
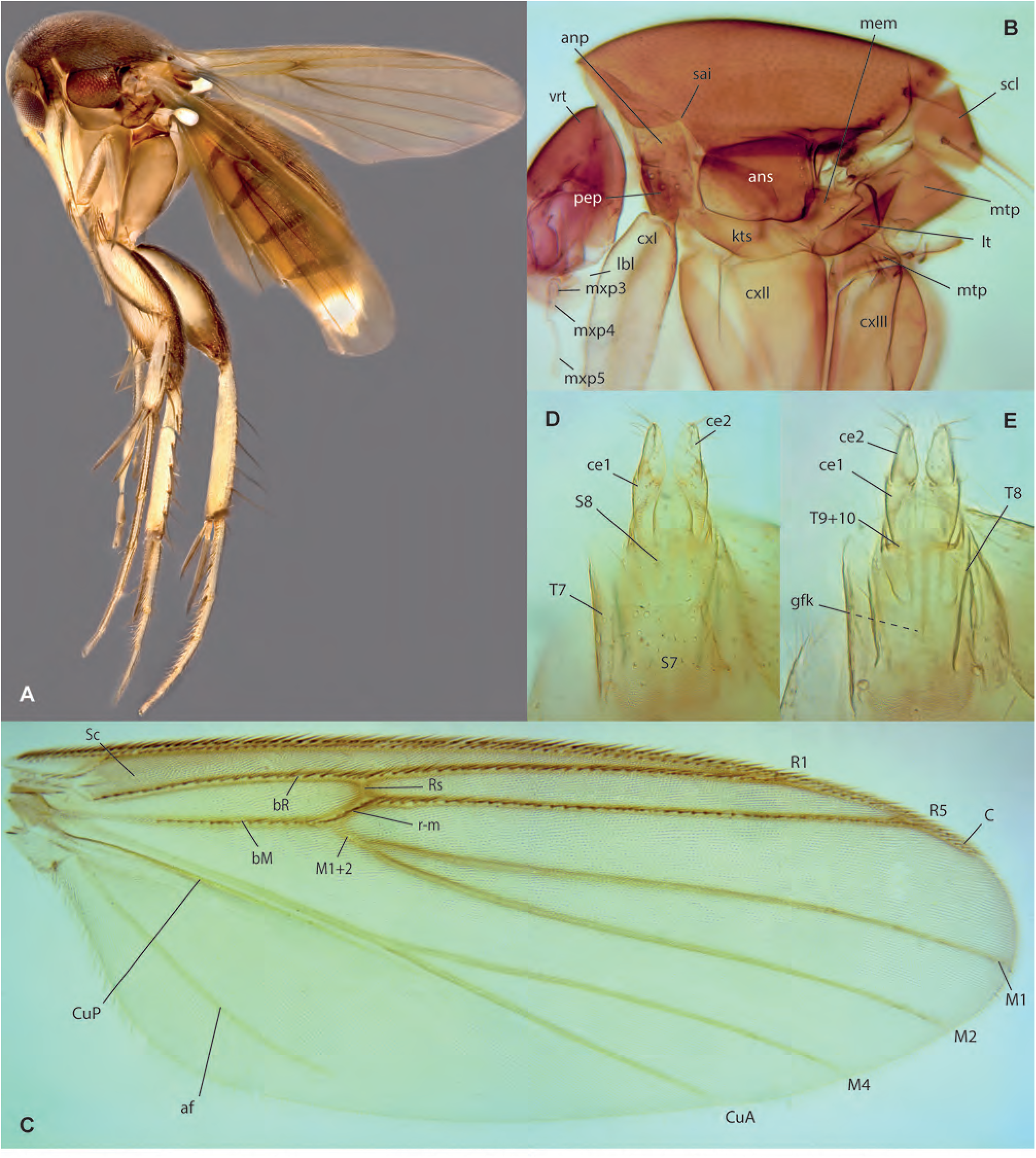
*Platurocypta adeleneweeae* Amorim & Oliveira, **sp. n.,** female. **A.** Habitus, lateral view, paratype ZRCBDP0048323. **B.** Thorax, lateral view, female paratype ZRCBDP0048424. **C.** Wing, paratype ZRCBDP0048323. **D.** Terminalia, ventral view, paratype ZRCBDP0048424. **E.** Terminalia, dorsal view, same.

**Figs. 109A-C.**
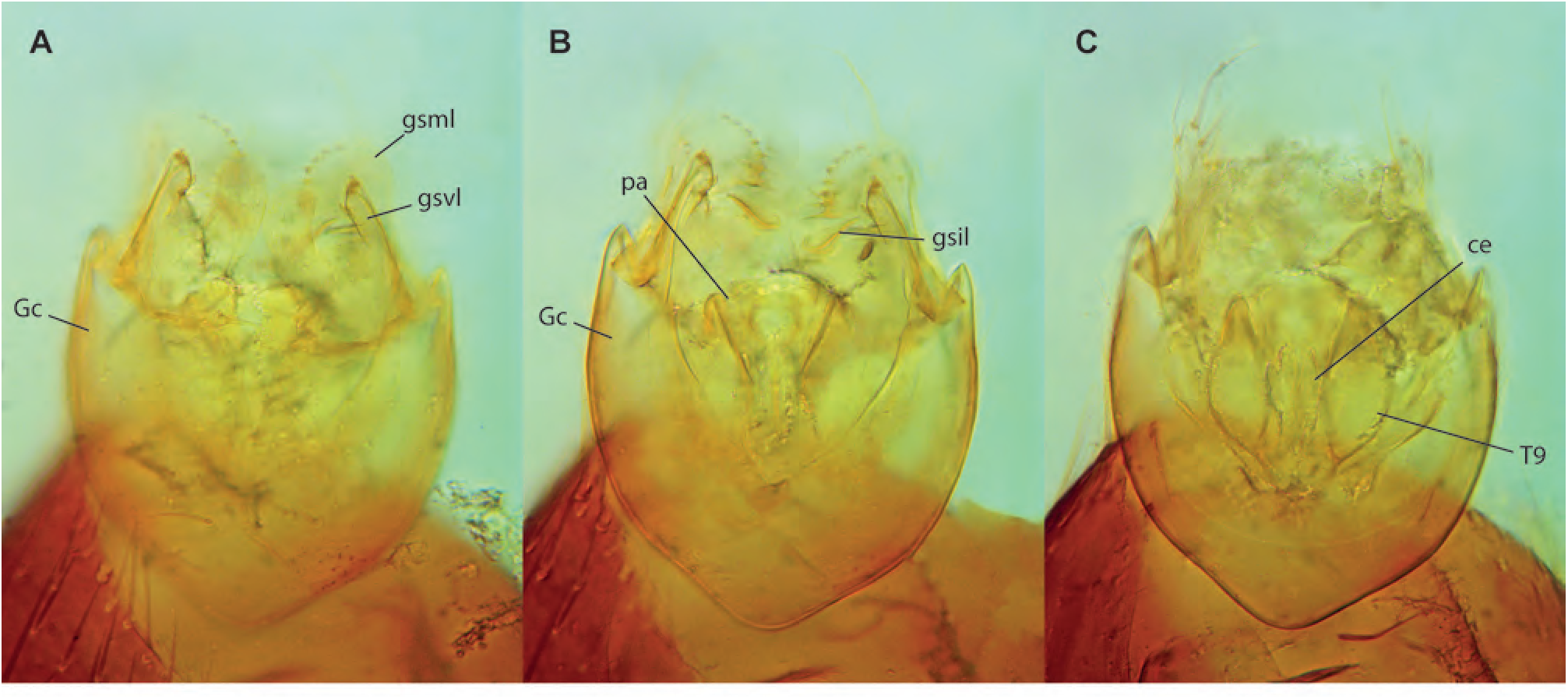
*Platurocypta adeleneweeae* Amorim & Oliveira, **sp. n.,** male holotype, terminalia. **A.** Ventral view. **B.** Mid-section. **C.** Dorsal view.

**Figs. 110A-G.**
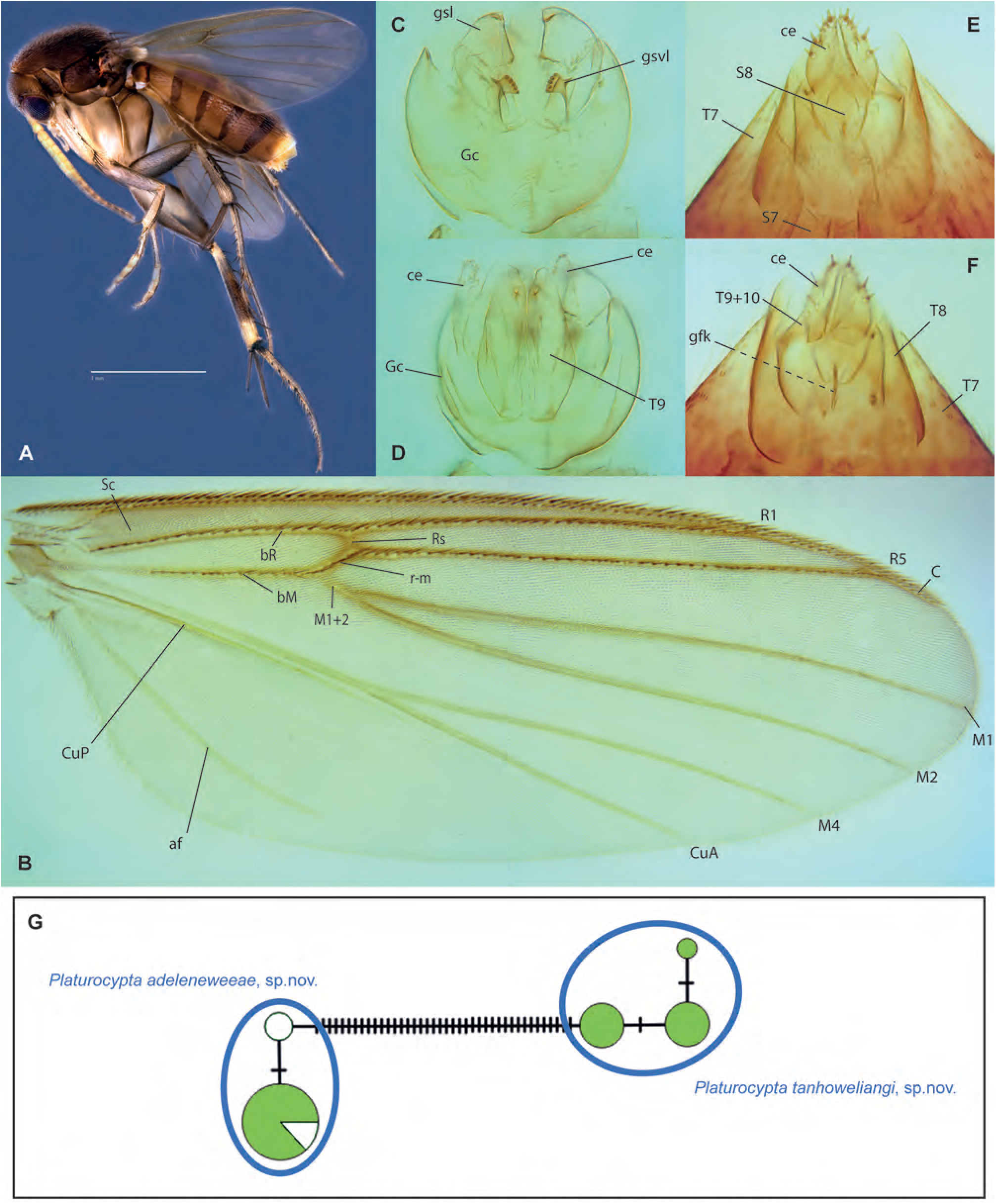
*Platurocypta tanhoweliangi* Amorim & Oliveira, **sp. n. A.** Habitus, lateral view, female paratype ZRCBDP0048325. **B.** Wing, male holotype. **C.** Male terminalia, ventral view, same. **D.** Male terminalia, dorsal view, same. **E.** Female terminalia, ventral view, paratype ZRCBDP0047940. **F.** Female terminalia, ventral view, same. **G.** Haplotype network for *Platurocypta*.

**Fig. 111.**
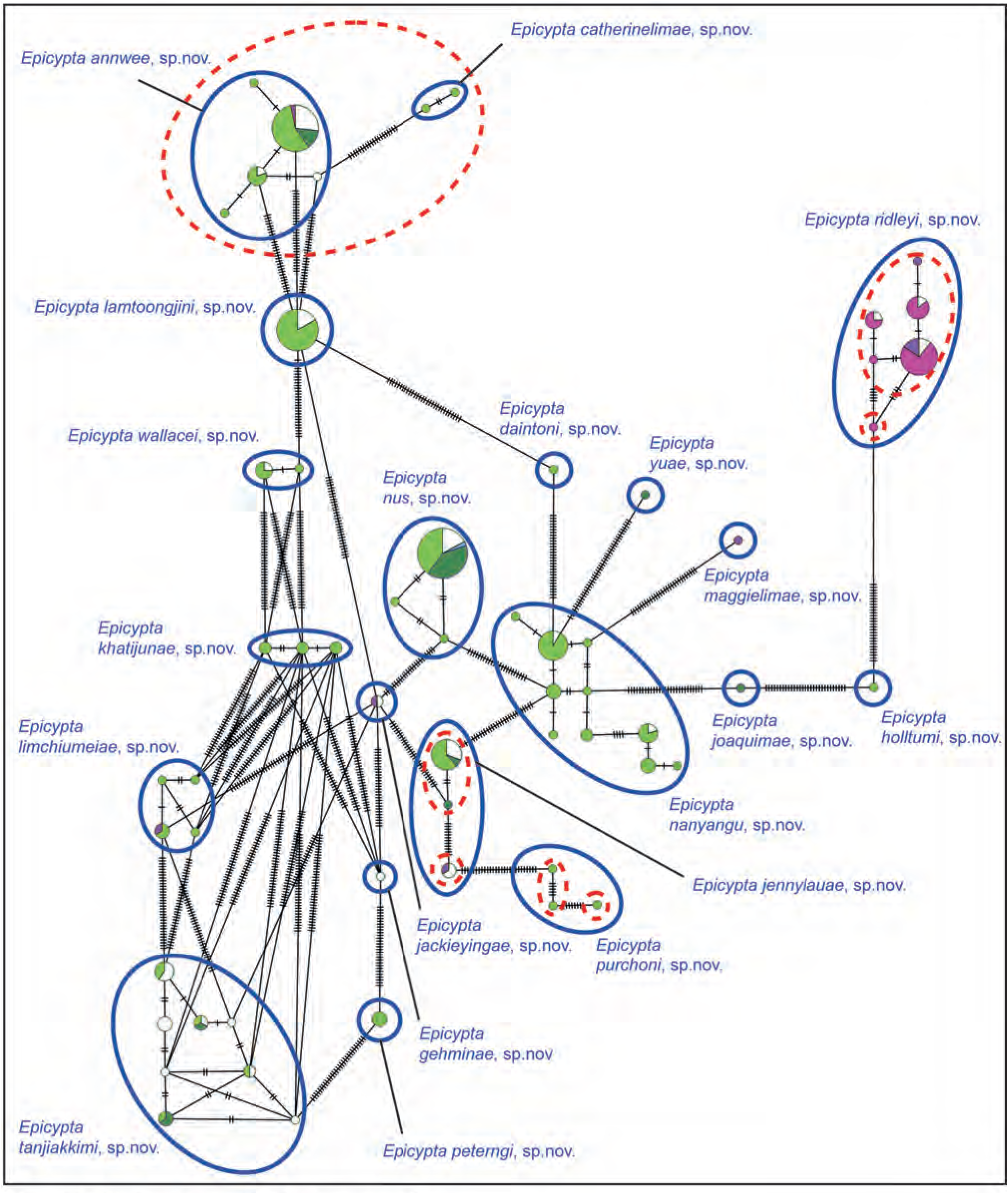
Haplotype network of *Epicypta* (part 1).

**Fig. 112.**
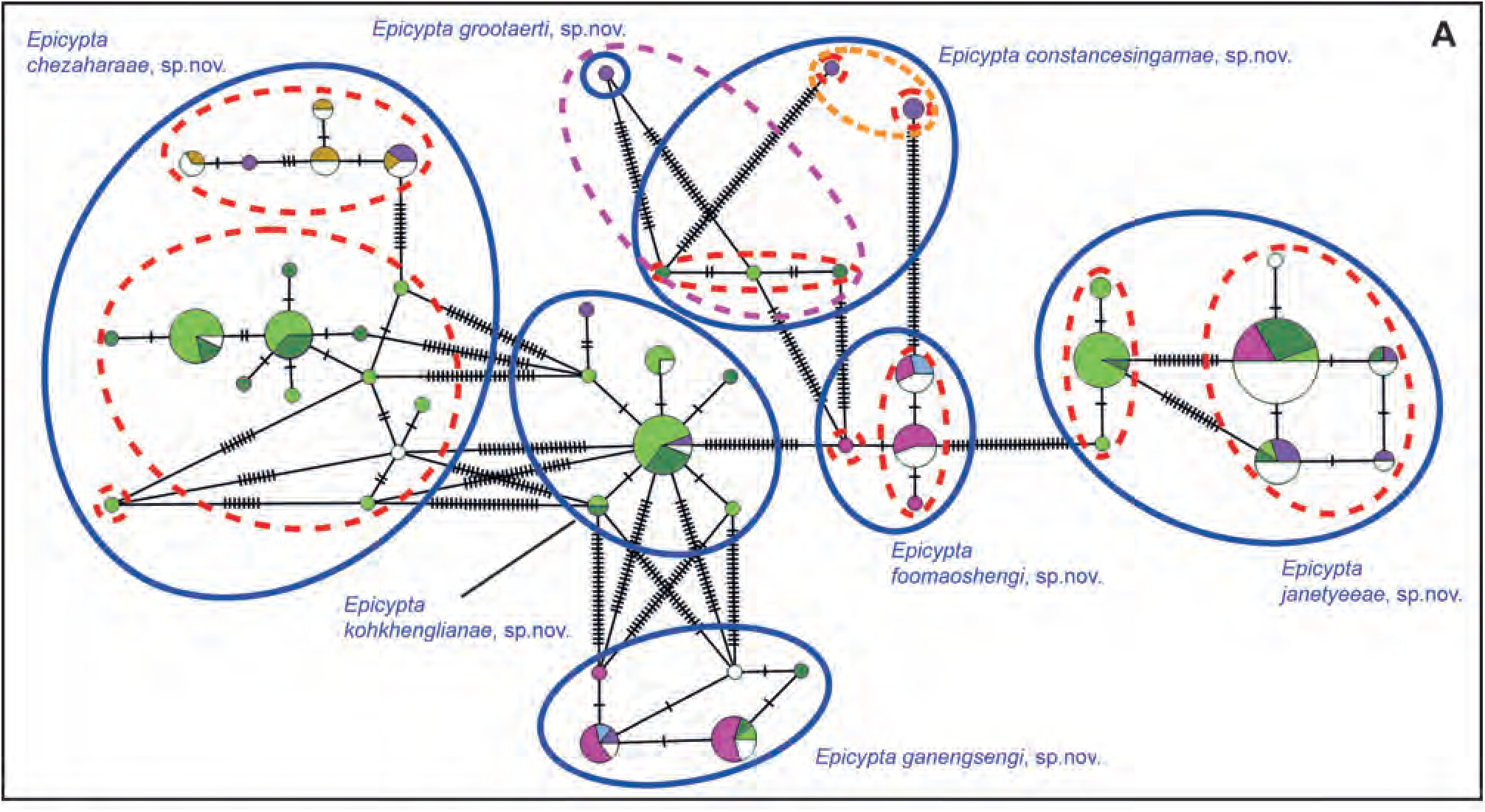
Haplotype network of *Epicypta* (part 2).

**Figs. 113A-G.**
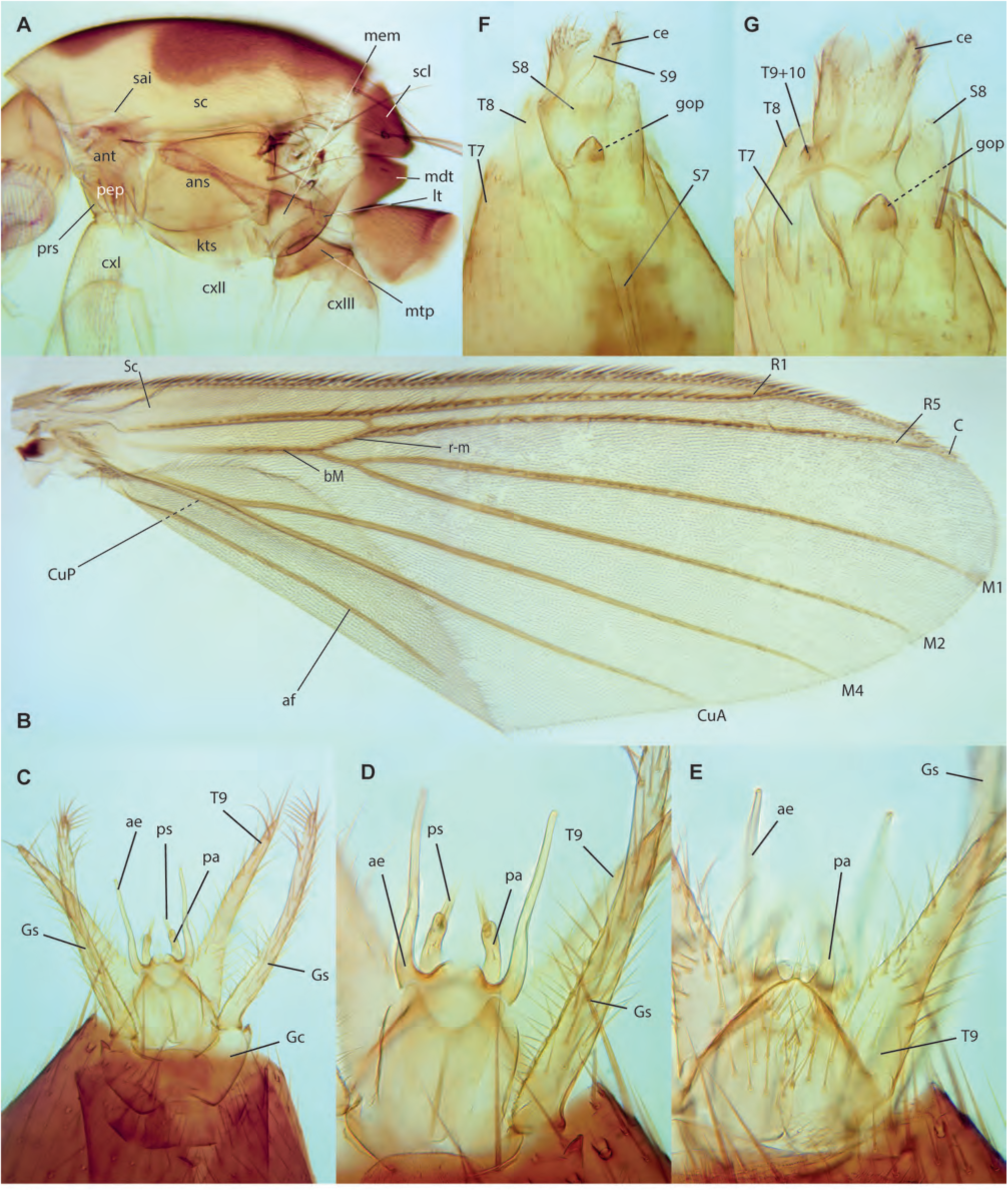
*Epicypta constancesingamae* Amorim & Oliveira, **sp. n. A.** Thorax, lateral view, female paratype, ZRCBDP0278248. **B.** Wing, male holotype. **C.** Male terminalia, ventral view, same. **D.** Detail of male terminalia, ventral view, same. **E.** Detail of male terminalia, dorsal view, same. **F.** Female terminalia, ventral view, paratype ZRCBDP0278248. **G.** Female terminalia, dorsal view, same.

**Figs. 114A-D.**
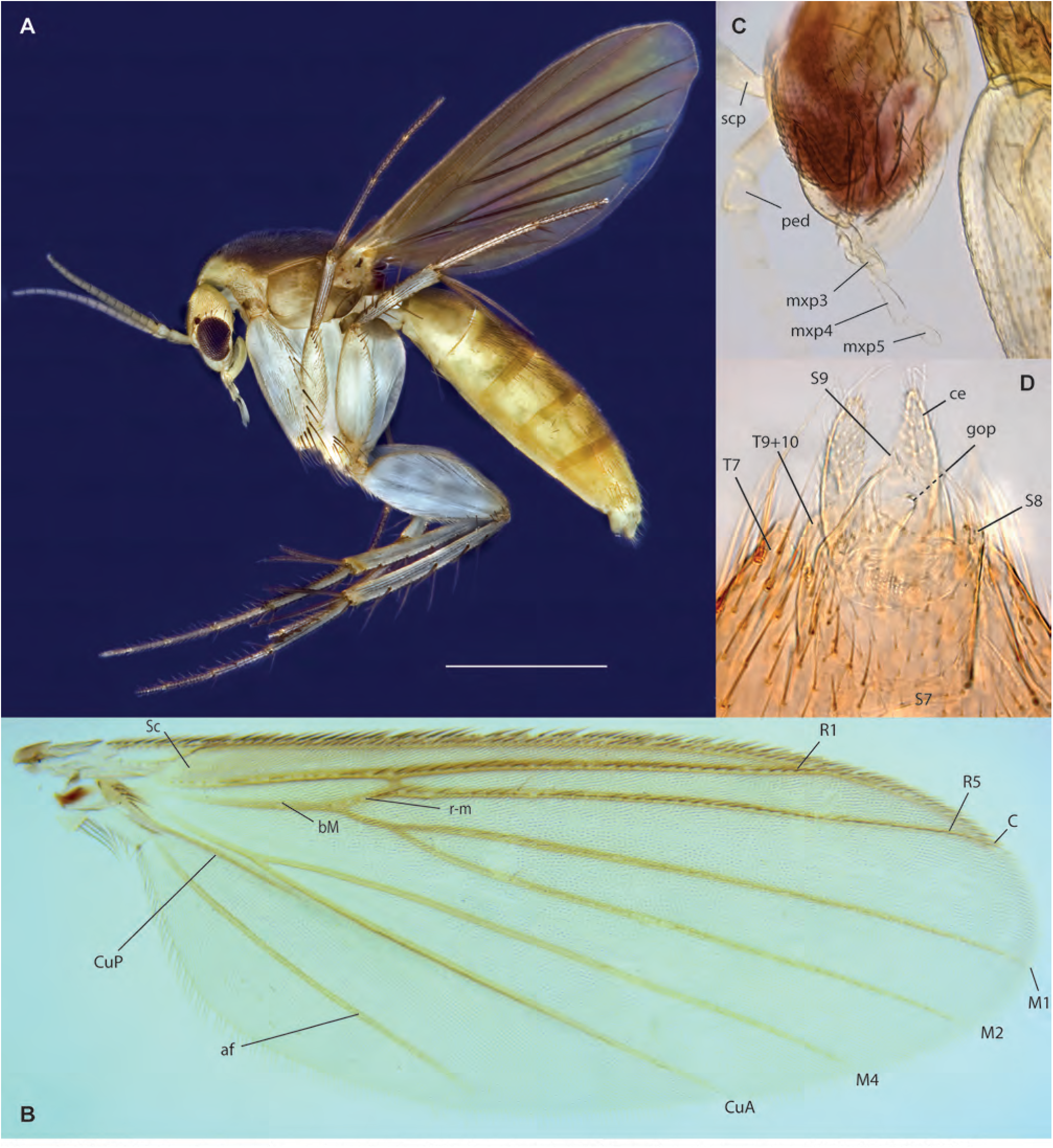
*Epicypta jennylauae* Amorim & Oliveira, **sp. n. A.** Habitus, lateral view, female paratype, ZRCBDP0048438. **B.** Wing, female paratype, ZRCBDP0048797. **C.** Head, lateral view, female holotype. **D.** Female terminalia, ventral view, holotype.

**Figs. 115A-B.**
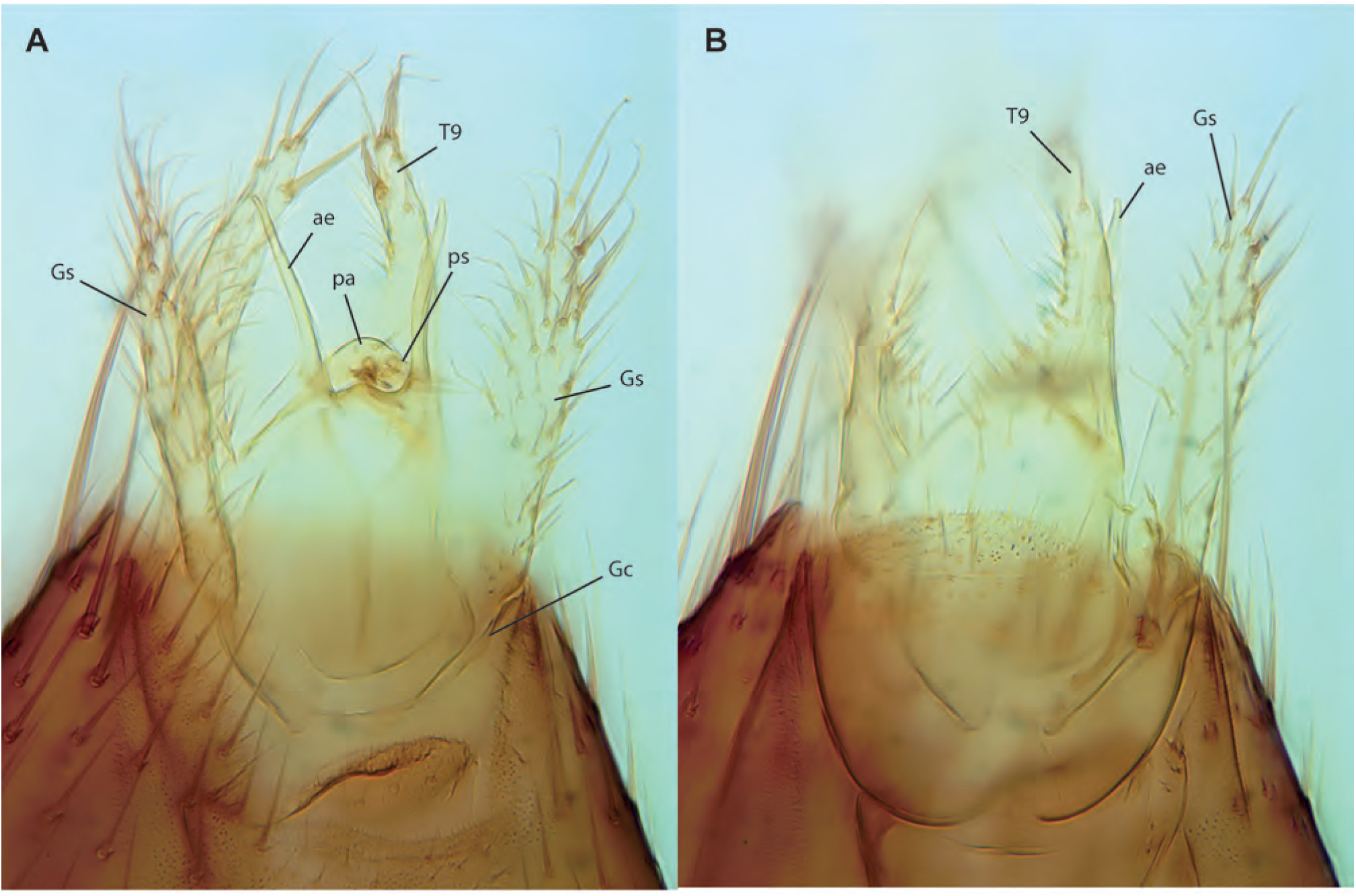
*Epicypta jennylauae* Amorim & Oliveira, **sp. n.,** male ZRCBDP0278324, terminalia. **A.** Ventral view, holotype. **B.** Dorsal view.

**Figs. 116A-E.**
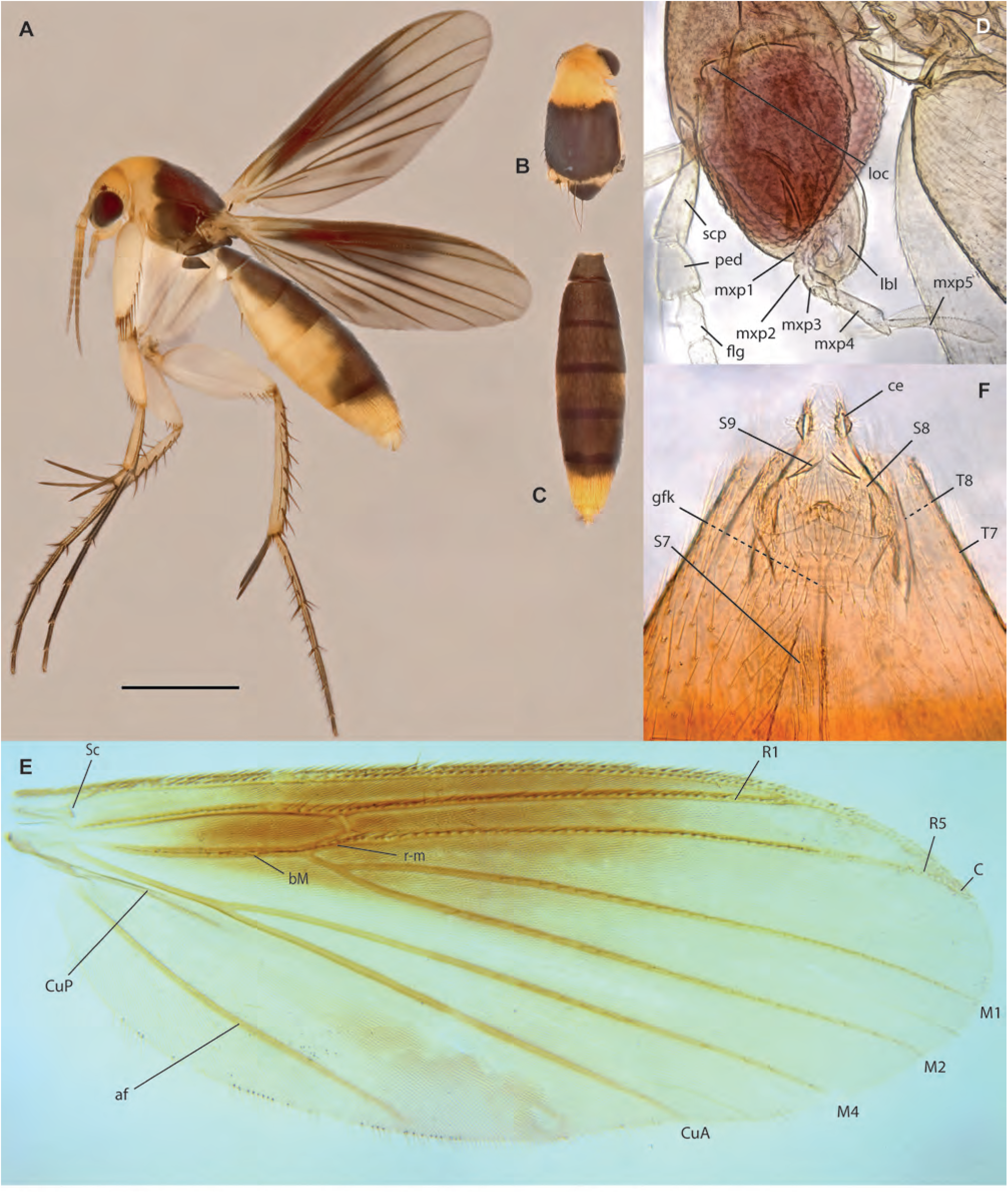
*Epicypta limchiumeiae* Amorim & Oliveira, **sp. n. A.** Habitus, lateral view, female paratype, ZRCBDP0048320. **B.** Thorax, dorsal view, same. **C.** Abdomen, dorsal view, same. **D.** Head, lateral view, female holotype. **E.** Wing, same. **F.** Terminalia, ventral view, same.

**Figs. 117A-F.**
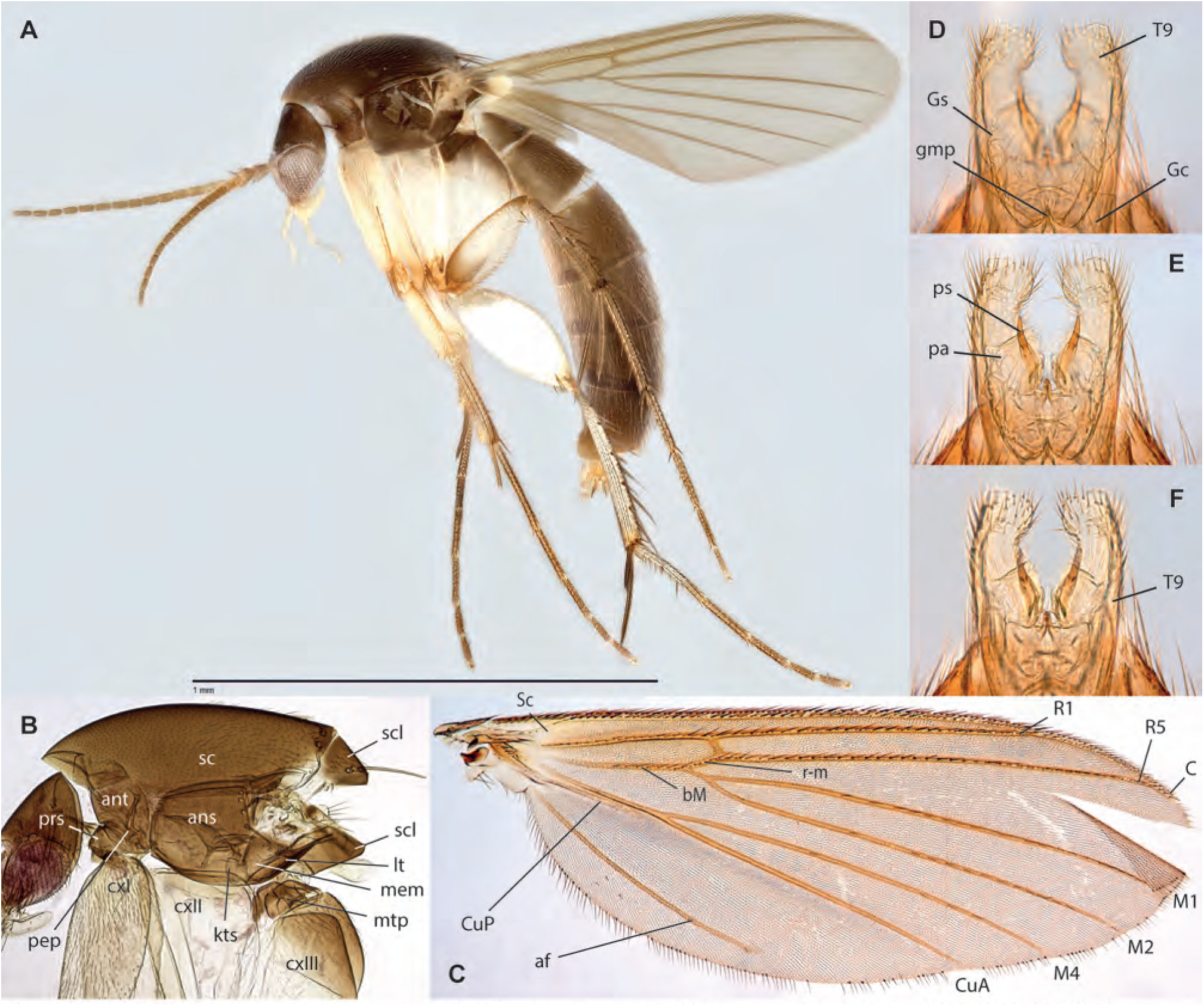
*Epicypta janetyeeae* Amorim & Oliveira, **sp. n. A.** Habitus, lateral view, male paratype, ZRCBDP0048909. **B.** Thorax, lateral view, same. **C.** Wing, male holotype. **D.** Male terminalia, ventral view, paratype ZRCBDP0072666. **E.** Same, mid-section. **F.** Same, dorsal view.

**Figs. 118A-D.**
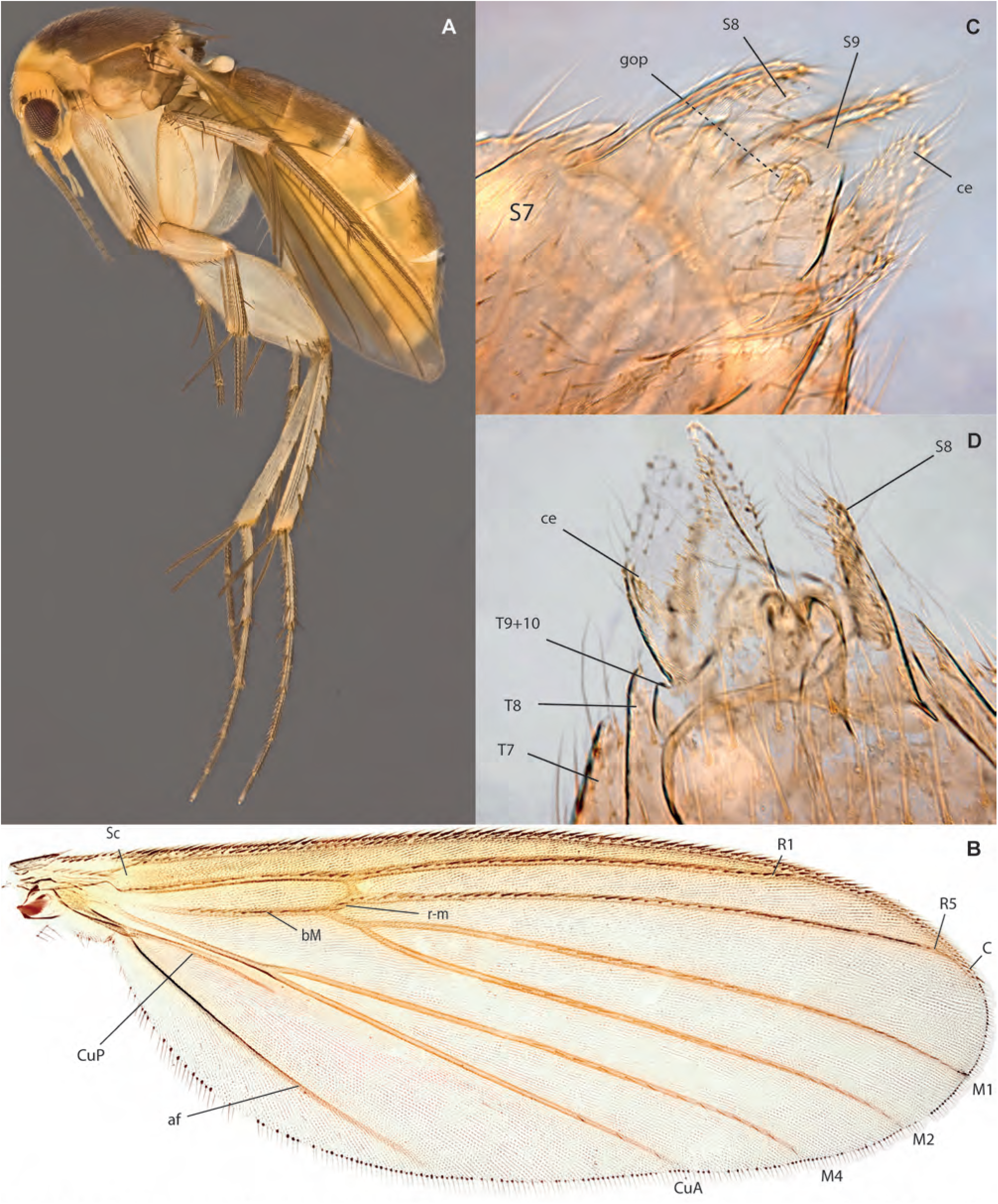
*Epicypta kohkhenglianae* Amorim & Oliveira, **sp. n. A.** Habitus, lateral view, female paratype, ZRCBDP0048433. **B.** Wing, same. **C.** Female terminalia, ventral view, paratype ZRCBDP0049101. **D.** Female terminalia, dorsal view, paratype ZRCBDP_0047782.

**Figs. 119A-B.**
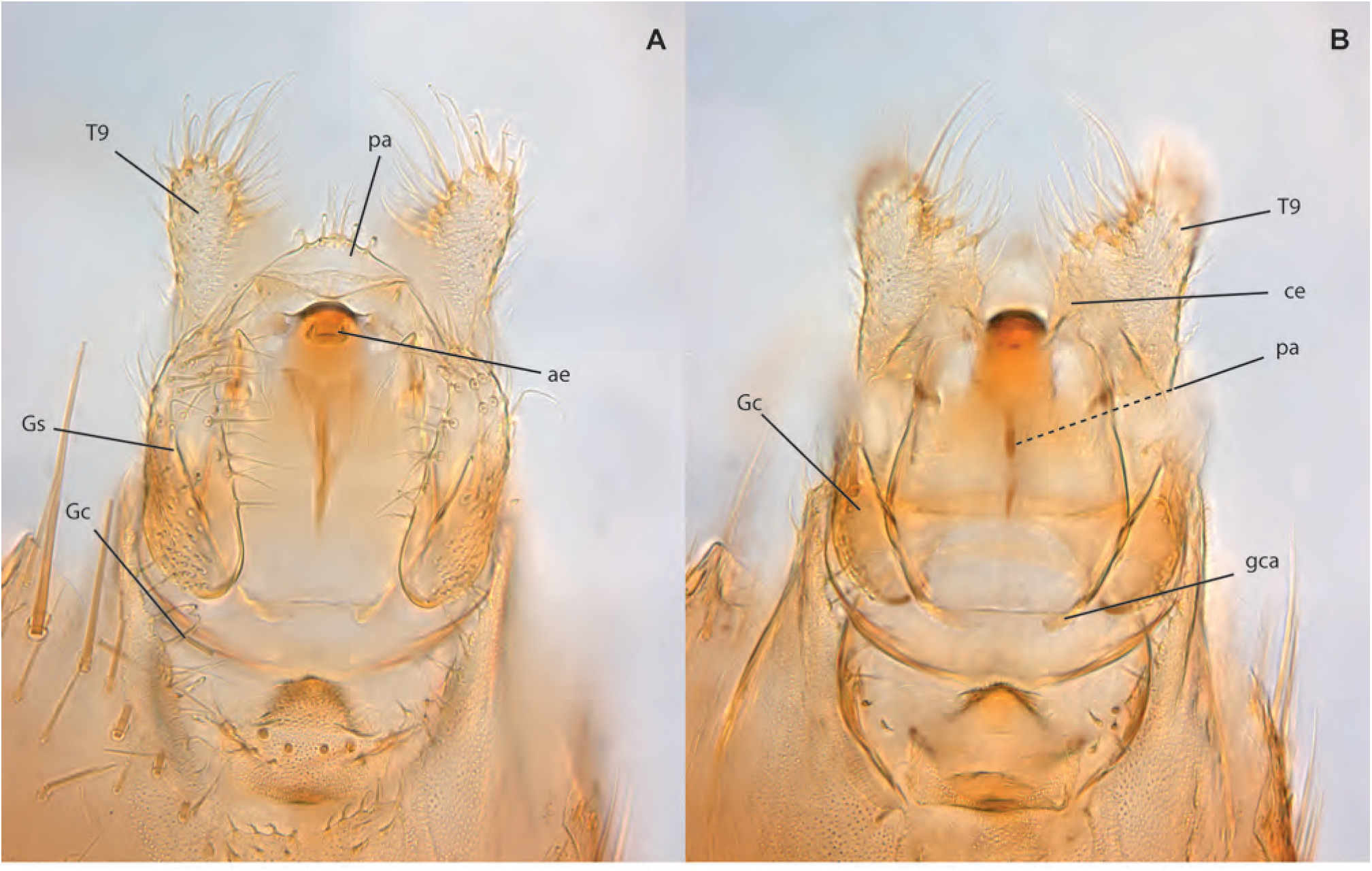
*Epicypta kohkhenglianae* Amorim & Oliveira, **sp. n.,** male holotype, terminalia. **A.** Ventral view. **B.** Dorsal view.

**Figs. 120A-D.**
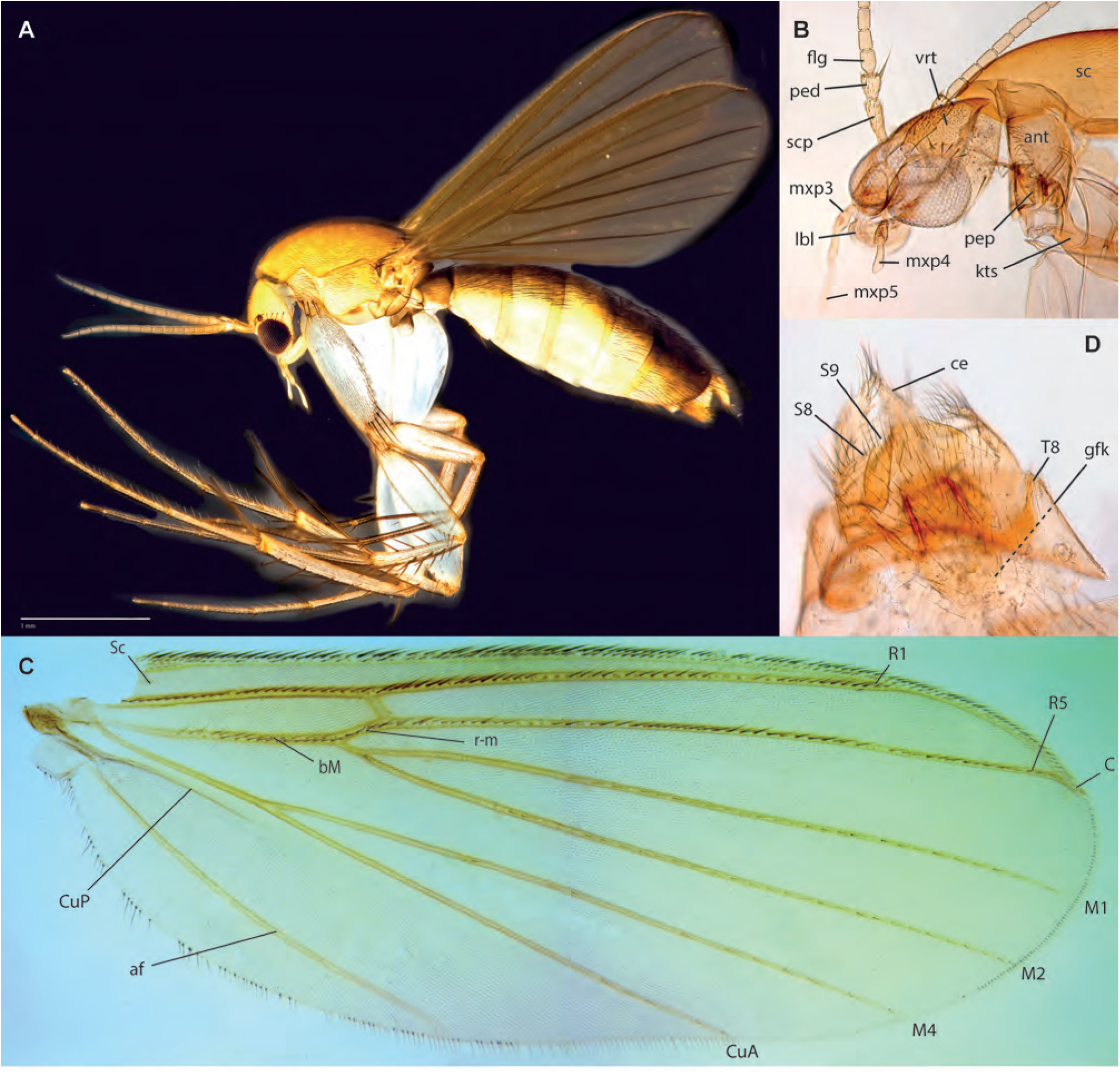
*Epicypta daintoni* Amorim & Oliveira, **sp. n.,** female holotype. **A.** Habitus, lateral view. **B.** Head, lateral view. **C.** Wing. **D.** Female terminalia, ventral view.

**Figs. 121A-D.**
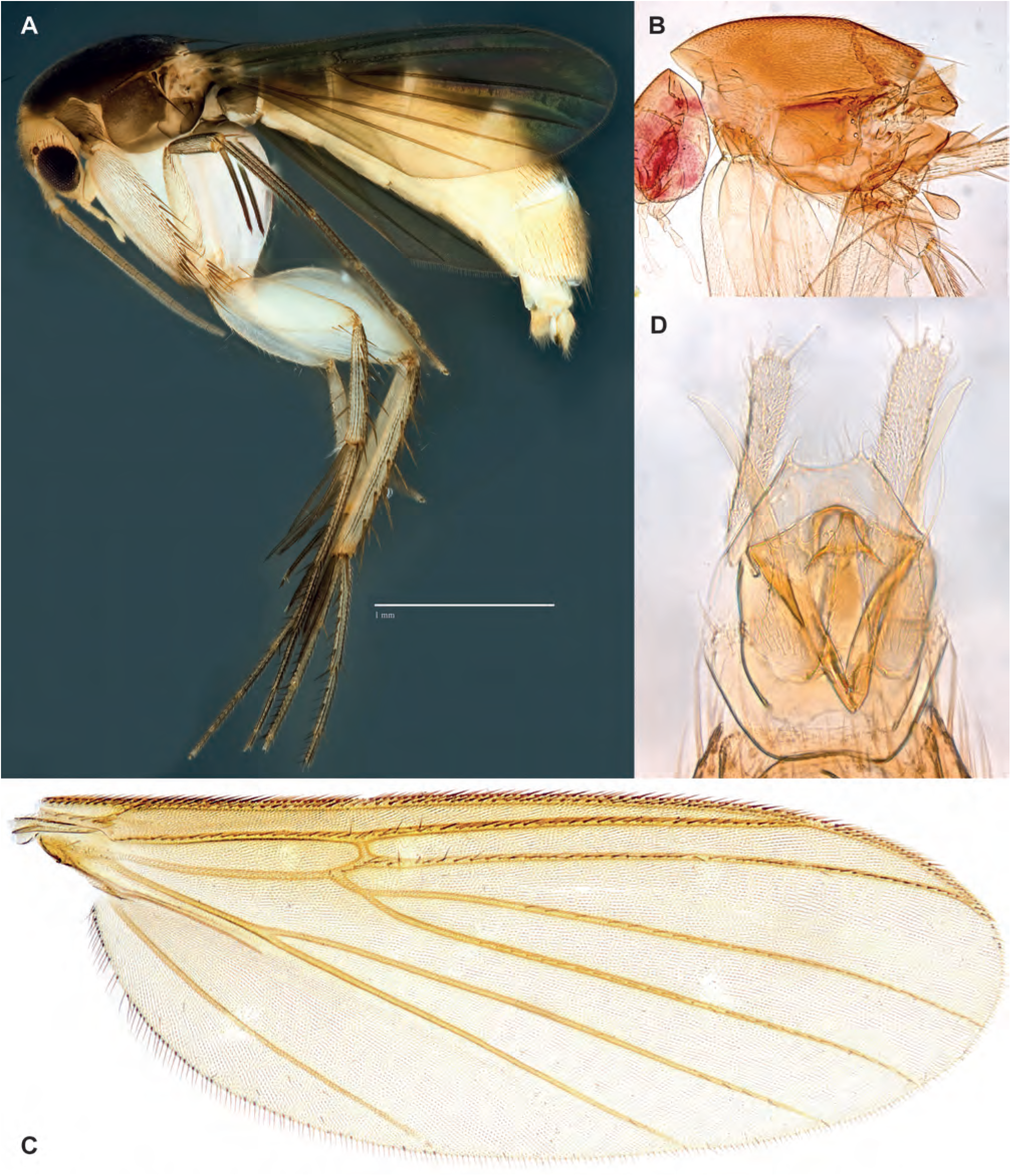
*Epicypta holltumi*. Habitus, lateral view, female ZRCBDP0048471.

**Figs. 122A-D.**
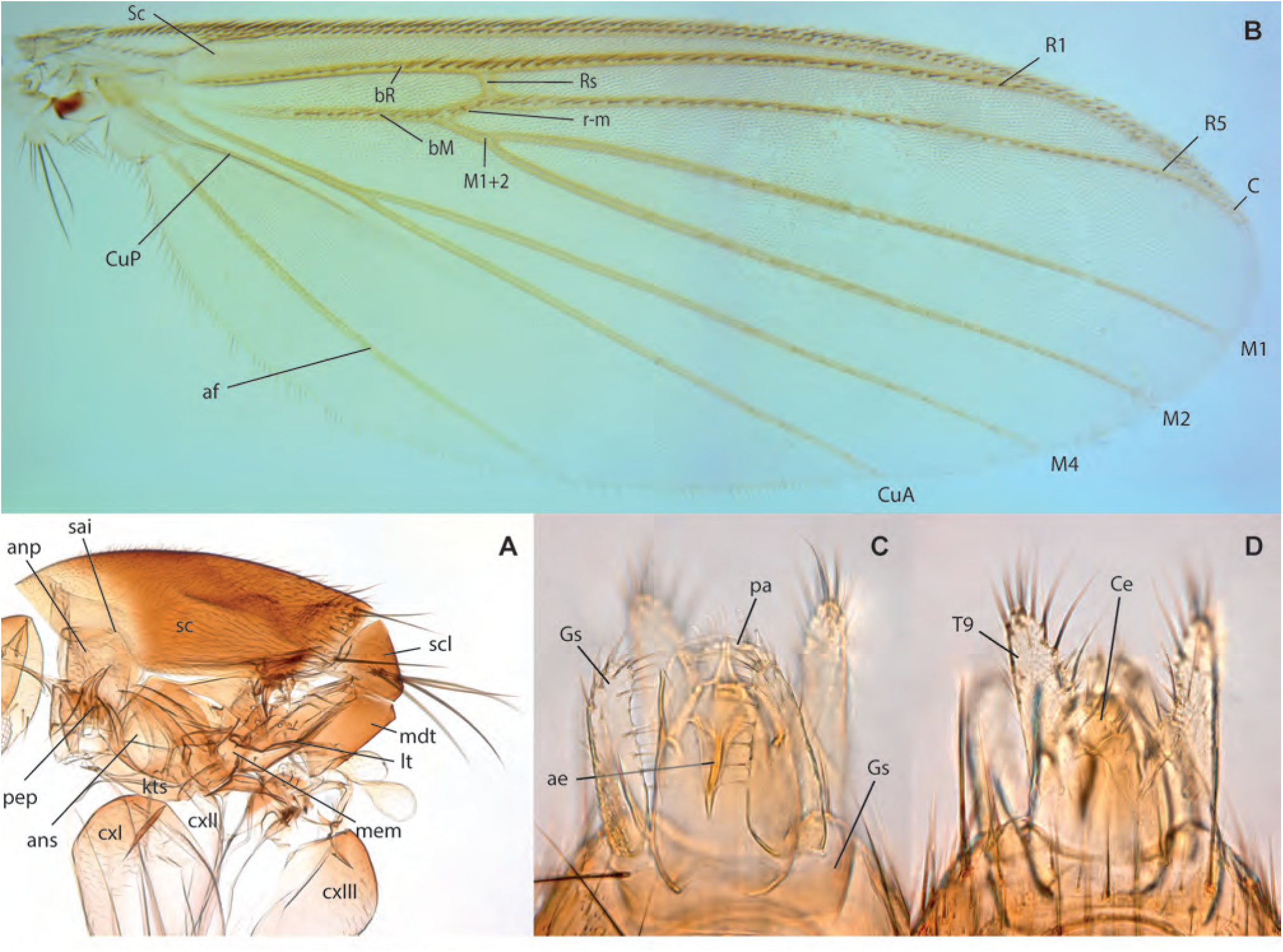
*Epicypta* sp. B. **A.** Thorax, lateral view. **B.** Wing. **C.** Terminalia, ventral view. **D.** Terminalia, dorsal view.

**Figs. 123A-D.**
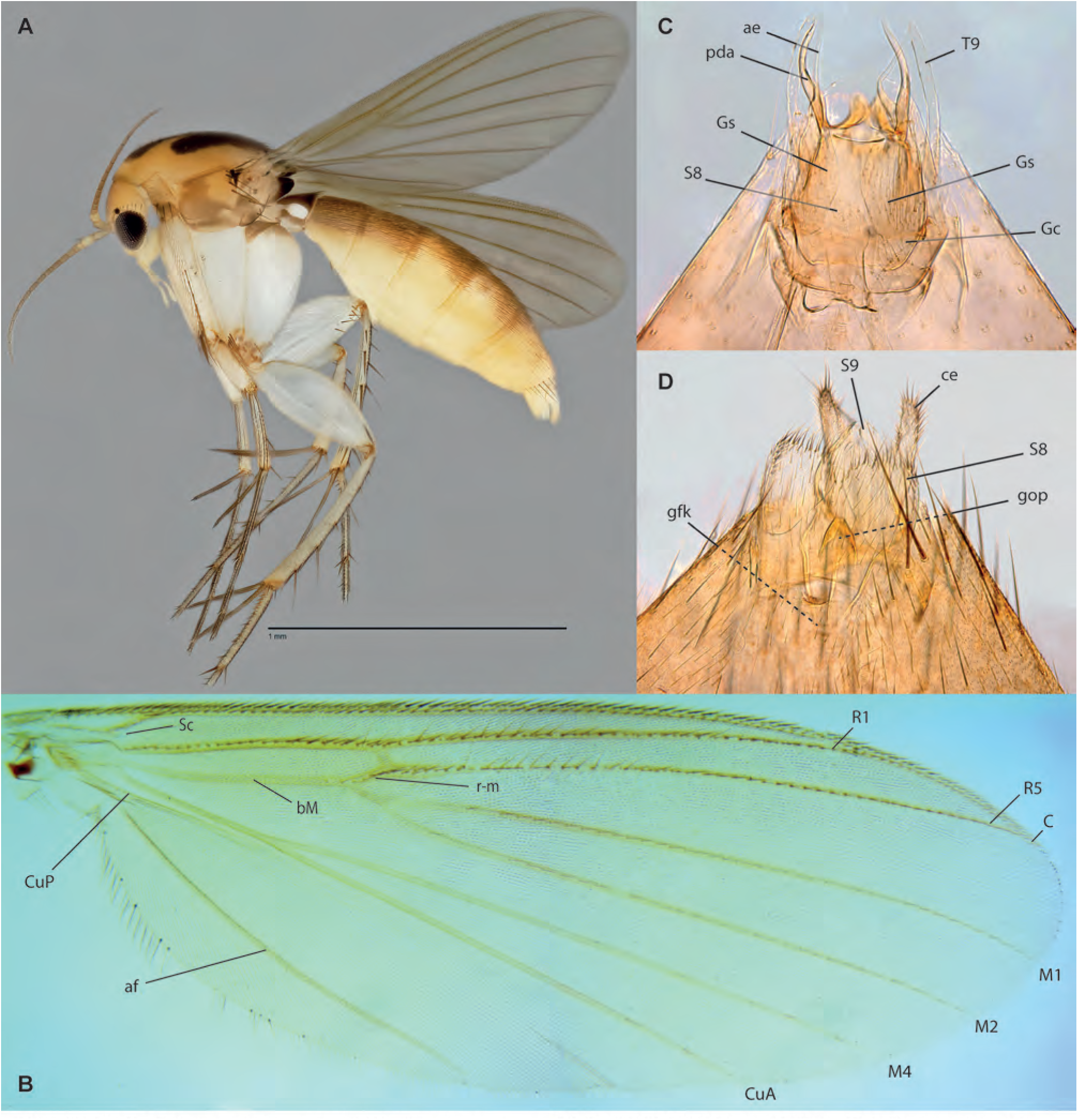
*Epicypta ridleyi* Amorim & Oliveira, **sp. n.**, female paratype, ZRCBDP0049152. **A.** Habitus, lateral view. **B.** Wing, male holotype. **C.** Male terminalia, ventral view, same. **D.** Female terminalia, ventral view, paratype ZRCBDP0049062.

**Figs. 124A-E.**
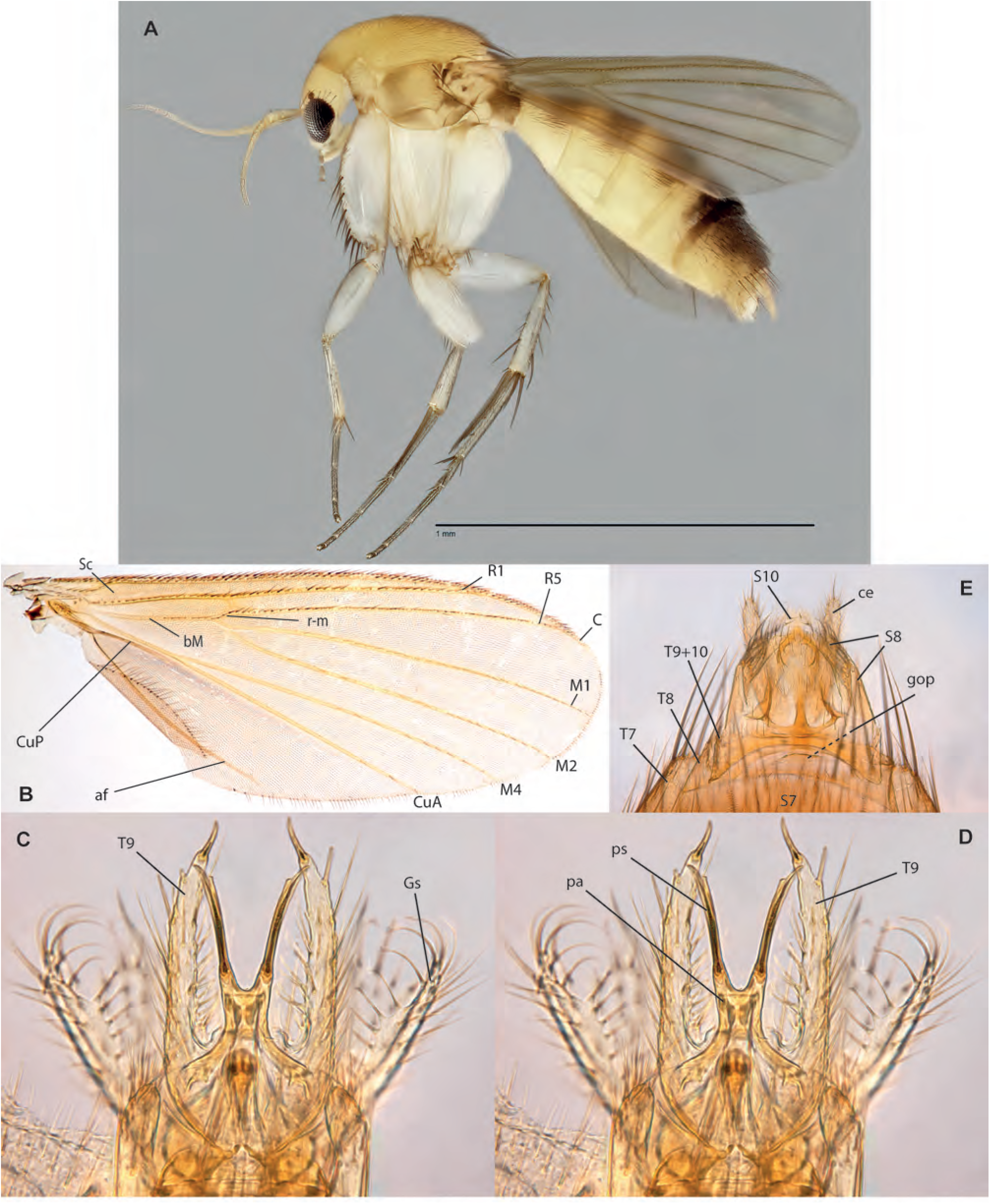
*Epicypta chezaharaae* Amorim & Oliveira, **sp. n. A.** Habitus, lateral view, female paratype, ZRCBDP0048442. **B.** Wing, female paratype ZRCBDP0154856. **C.** Male terminalia, ventral view, holotype. **D.** Male terminalia, dorsal view, same. **E.** Female terminalia, ventral view, paratype ZRCBDP0048120.

**Figs. 125A-D.**
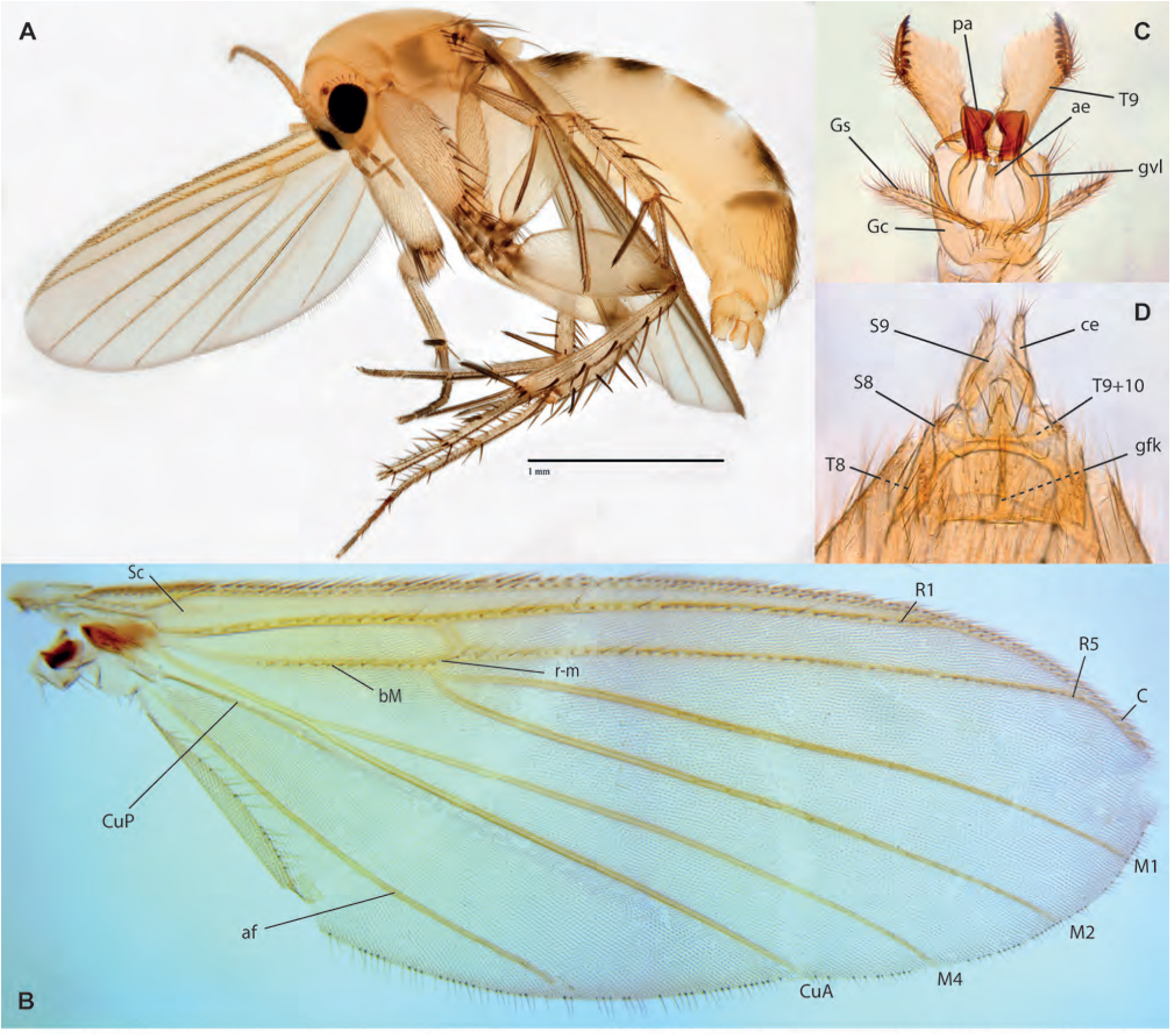
*Epicypta tanjiakkimi* Amorim & Oliveira, **sp. n. A.** Habitus, lateral view, female paratype, ZRCBDP0048452. **B.** Wing, male holotype. **C.** Male terminalia, ventral view, same. **D.** Female terminalia, ventral view, paratype ZRCBDP0047866.

**Figs. 126A-D.**
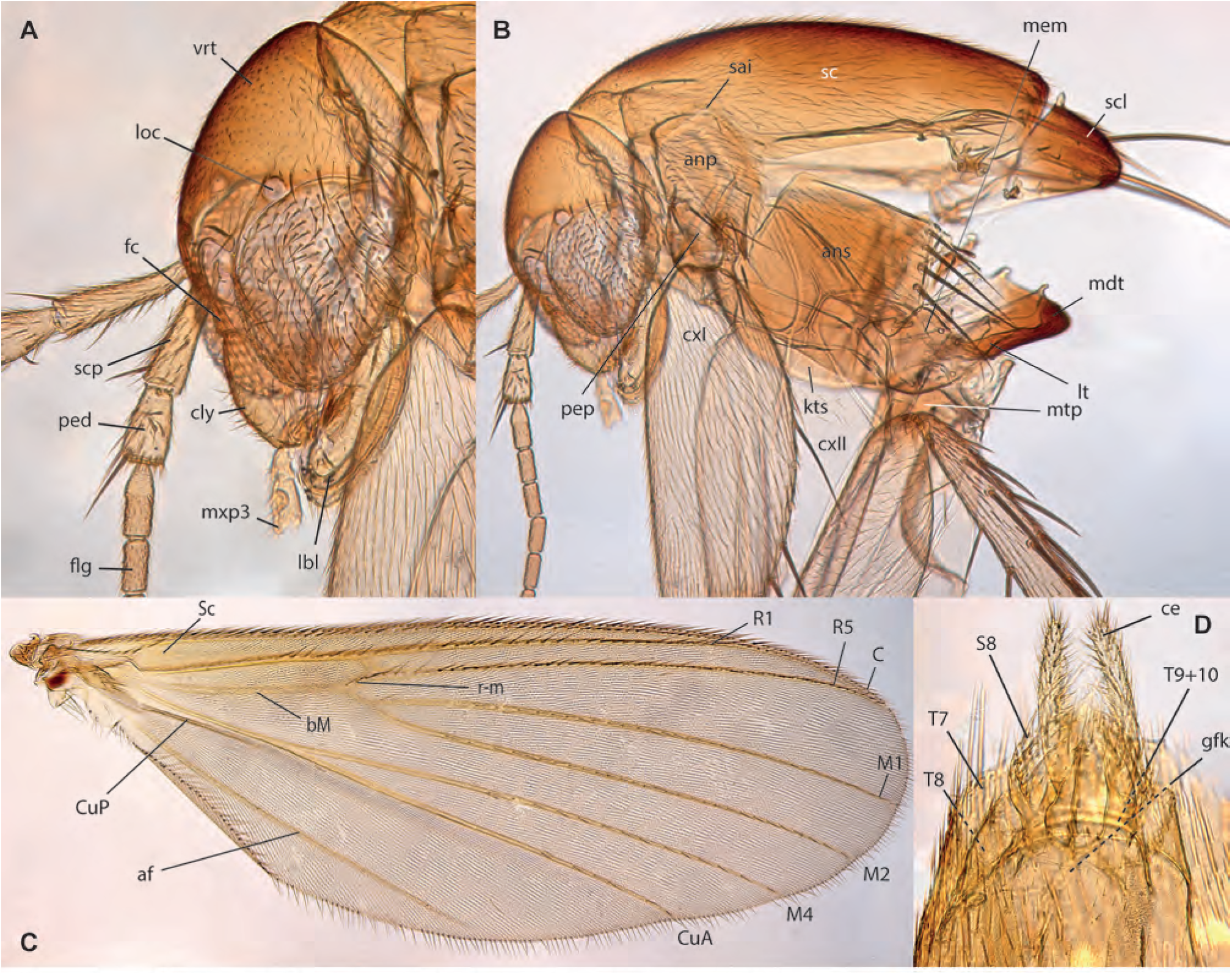
*Epicypta gehminae* Amorim & Oliveira, **sp. n.,** female holotype. **A.** Head, lateral view. **B.** Thorax, lateral view. **C.** Wing. **D.** Female terminalia, ventral view.

**Figs. 127A-E.**
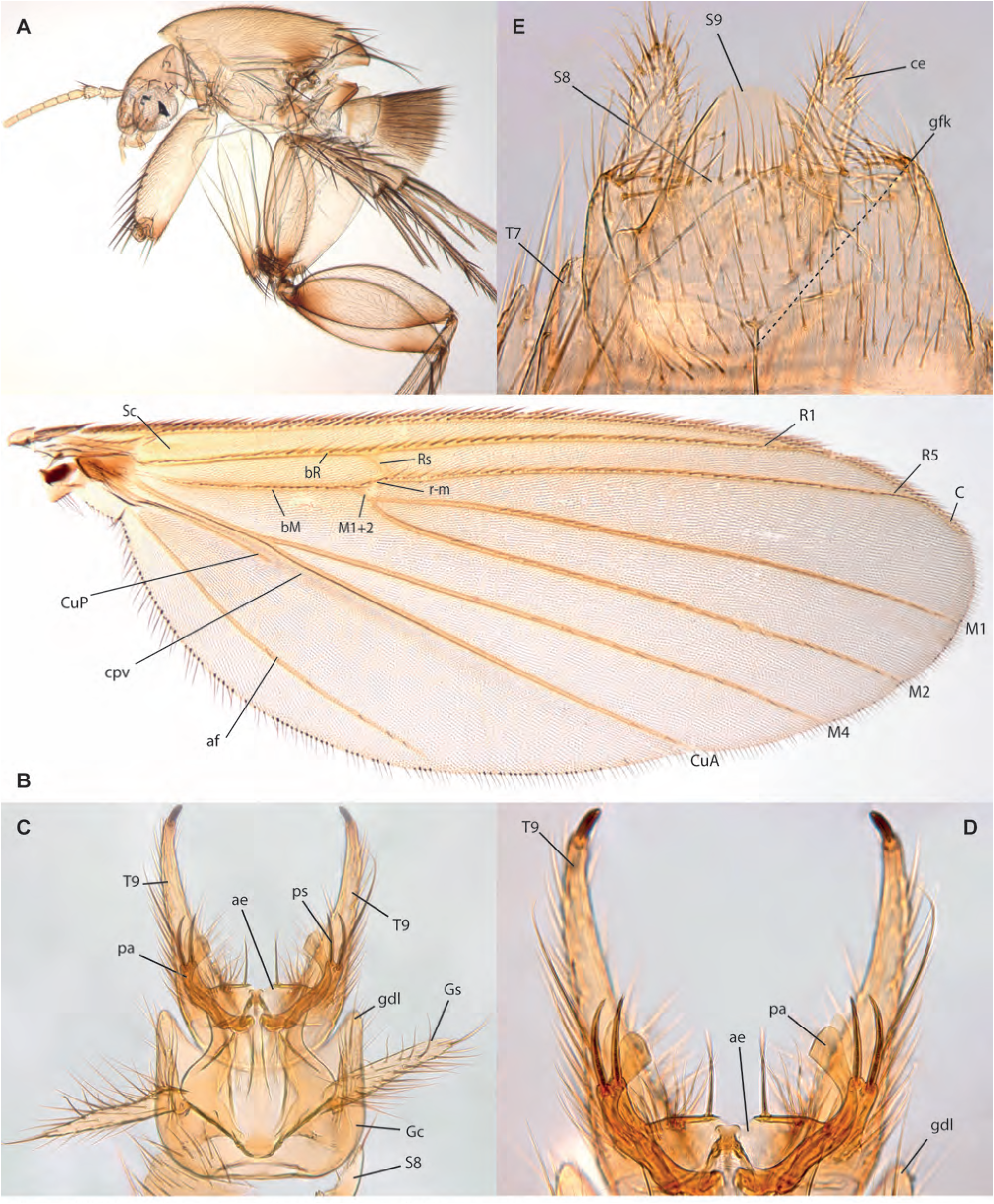
*Epicypta jackieyingae* Amorim & Oliveira, **sp. n. A.** Thorax and head, lateral view, male holotype. **B.** Wing, same. **C.** Male terminalia, ventral view, same. **D.** Male terminalia, distal end, ventral view, same. **E.** Female terminalia, paratype ZRCBDP0284204.

**Figs. 128A-C.**
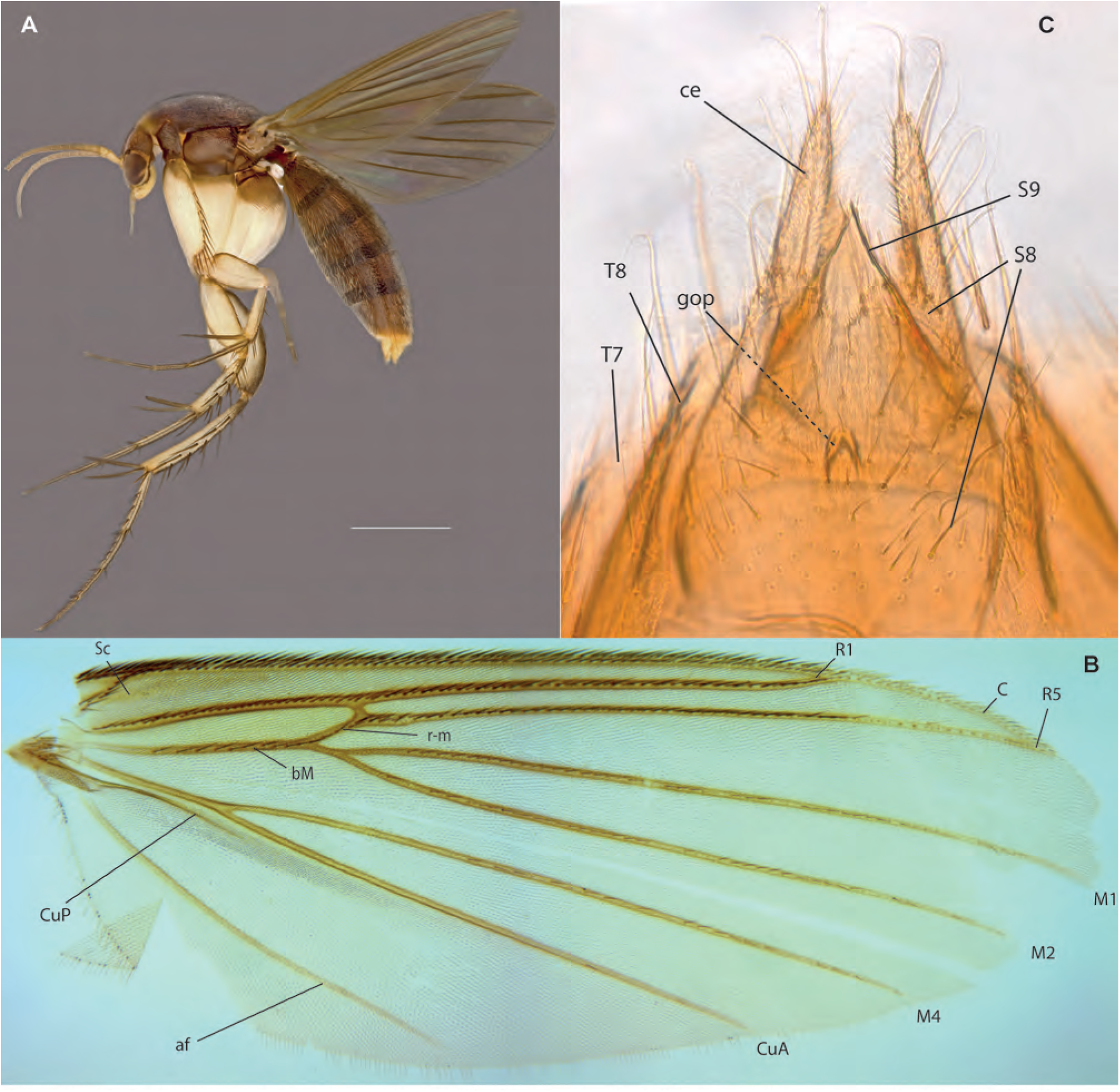
*Epicypta khatijunae* Amorim & Oliveira, **sp. n. A.** Habitus, lateral view, female paratype, ZRCBDP0048431. **B.** Wing, male holotype. **C.** Female terminalia, ventral view.

**Figs. 129A-D.**
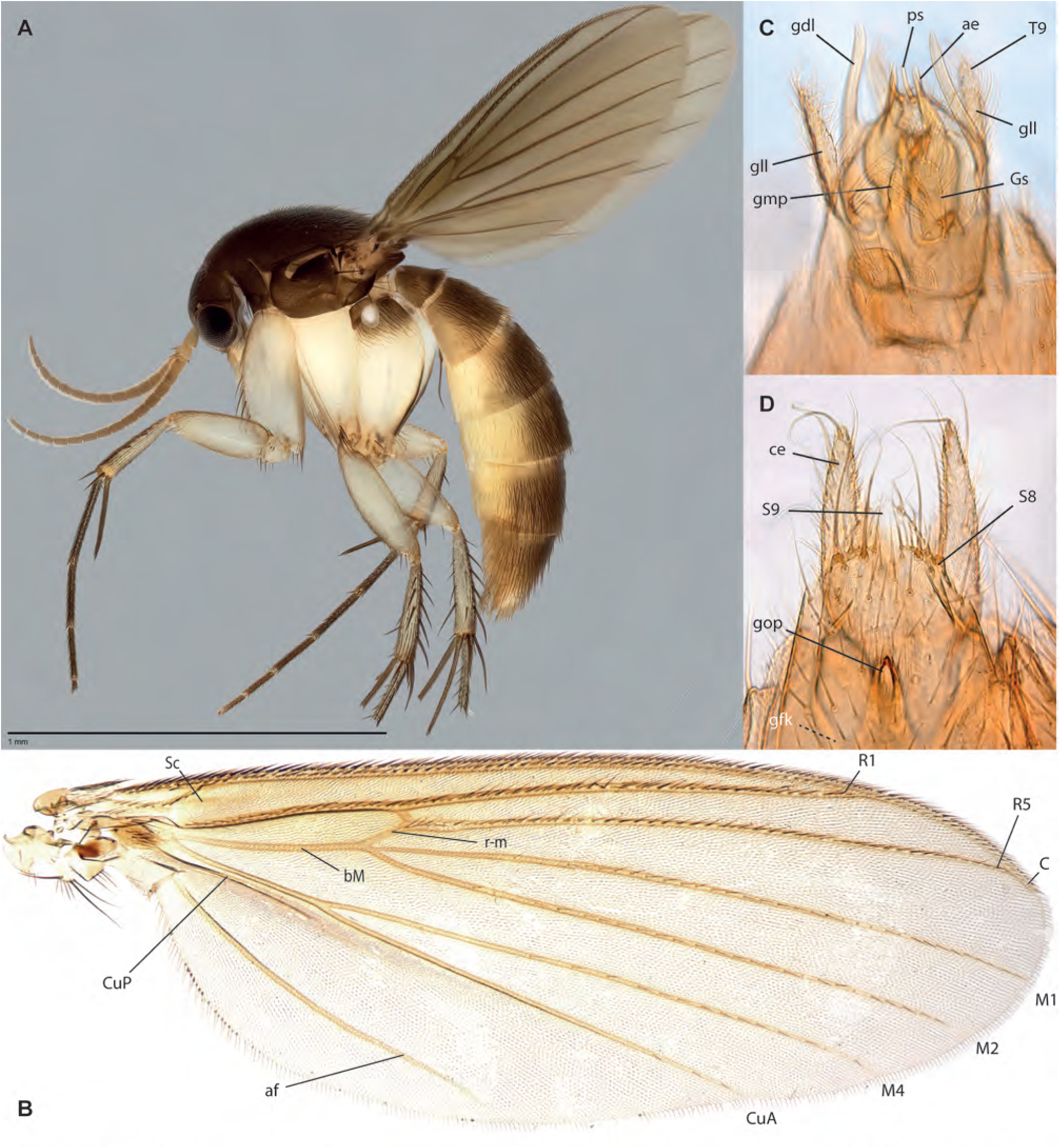
*Epicypta purchoni* Amorim & Oliveira, **sp. n. A.** Habitus, lateral view, female paratype ZRCBDP0047857. **B.** Wing, male holotype. **C.** Male terminalia, ventral view, same. **D.** Female terminalia, ventral view, ZRCBDP0047813.

**Figs. 130A-D.**
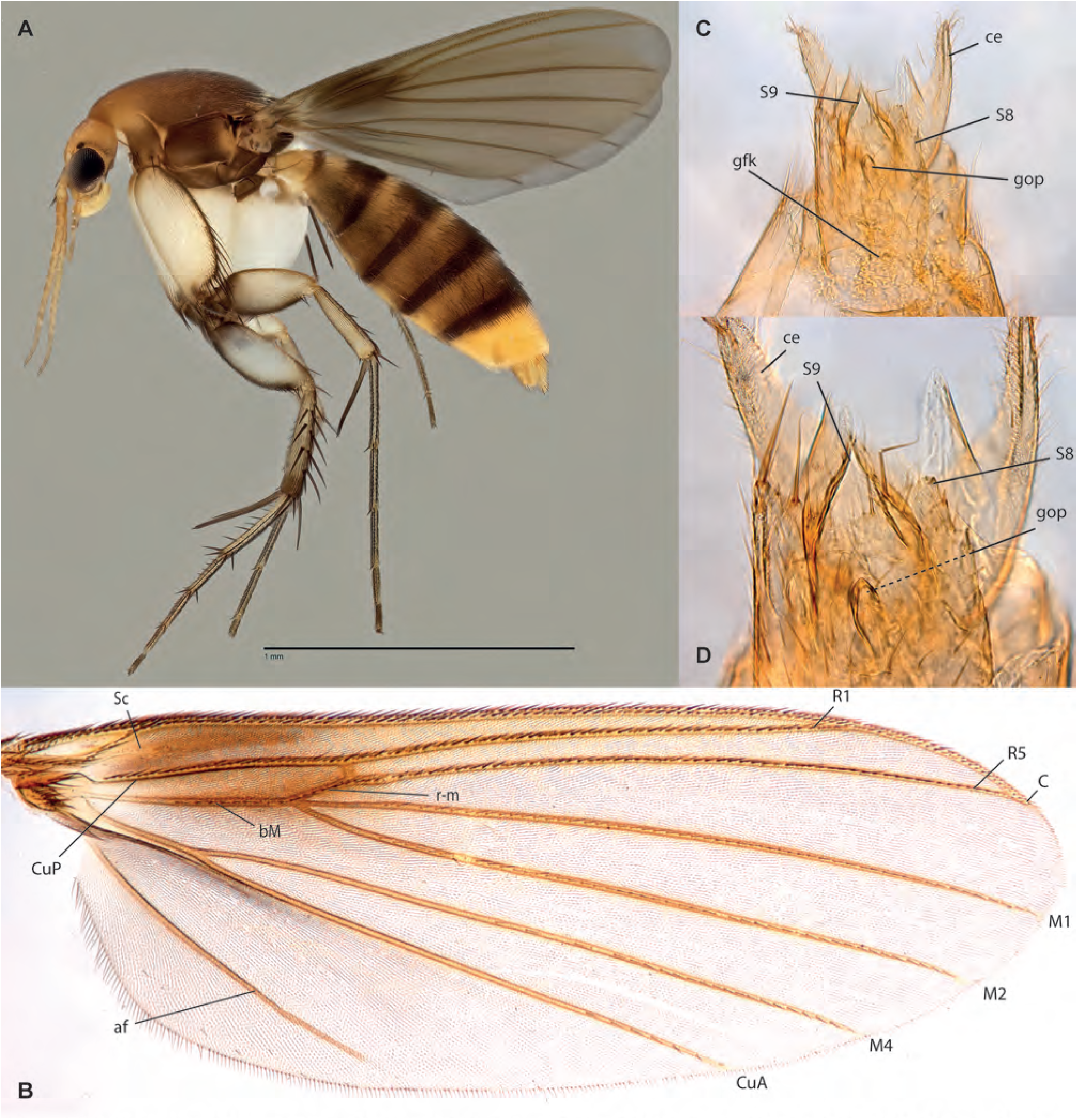
*Epicypta foomaoshengi Amorim* & Oliveira, **sp. n. A.** Habitus, lateral view, female paratype, ZRCBDP0049068. **B.** Wing, male holotype. . **C.** Detail of female terminalia, same. **D.** Terminalia, ventral view, same.

**Figs. 131A-C.**
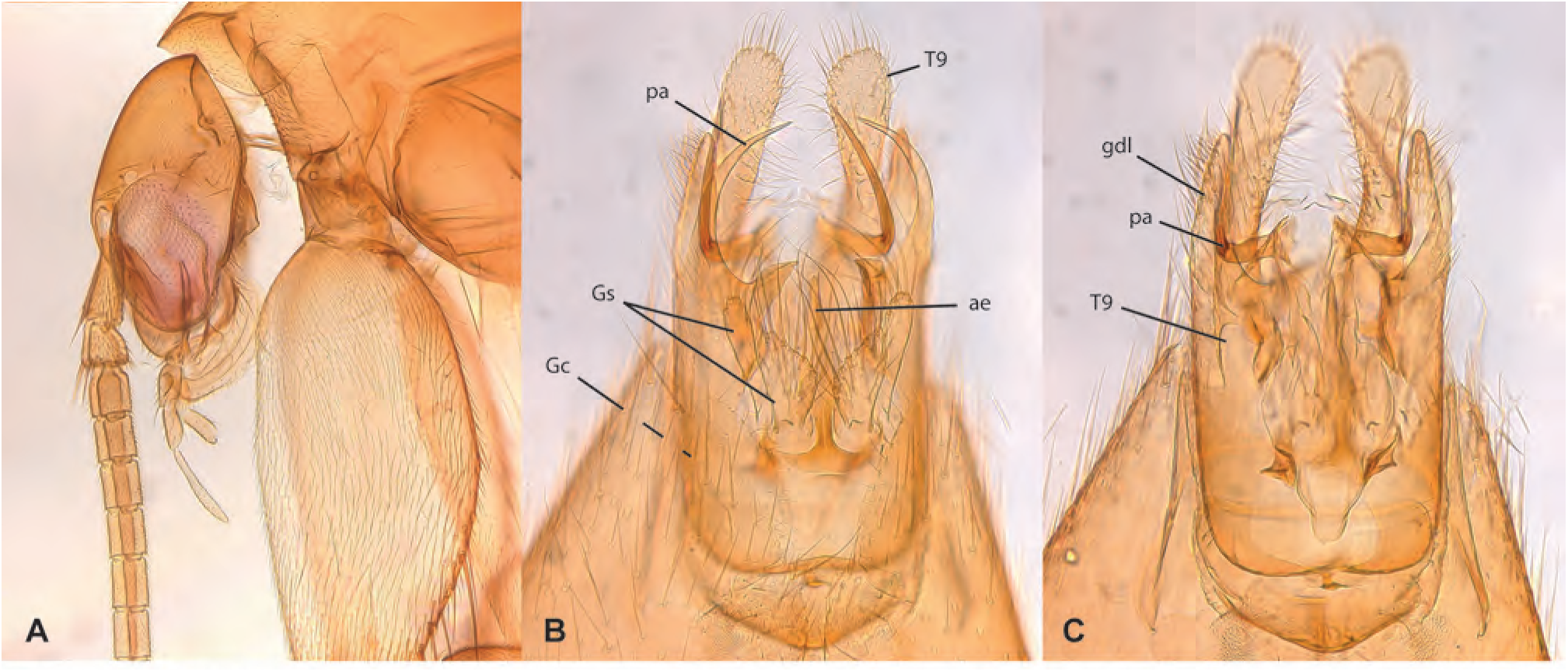
*Epicypta foomaoshengi Amorim* & Oliveira, **sp. n.,** male holotype, ZRCBDP0284196. **A.** Head, lateral view. **B.** Terminalia, ventral view. **C.** Terminalia, dorsal view.

**Figs. 132A-C.**
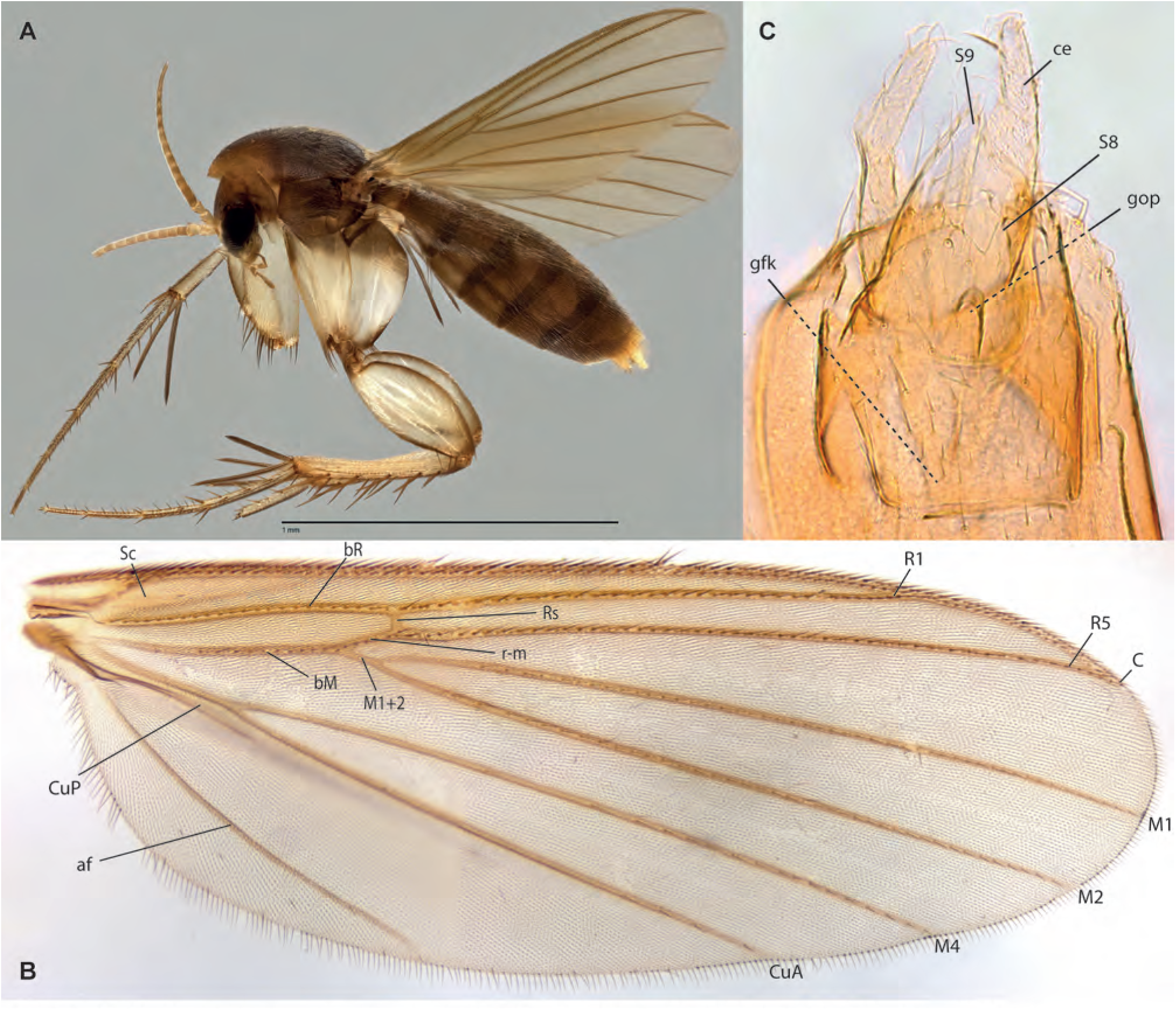
*Epicypta ganengsengi* Amorim & Oliveira, **sp. n.,** female holotype. **A.** Habitus, lateral view. **B.** Wing. **C.** Terminalia, ventral view.

**Figs. 133A-B.**
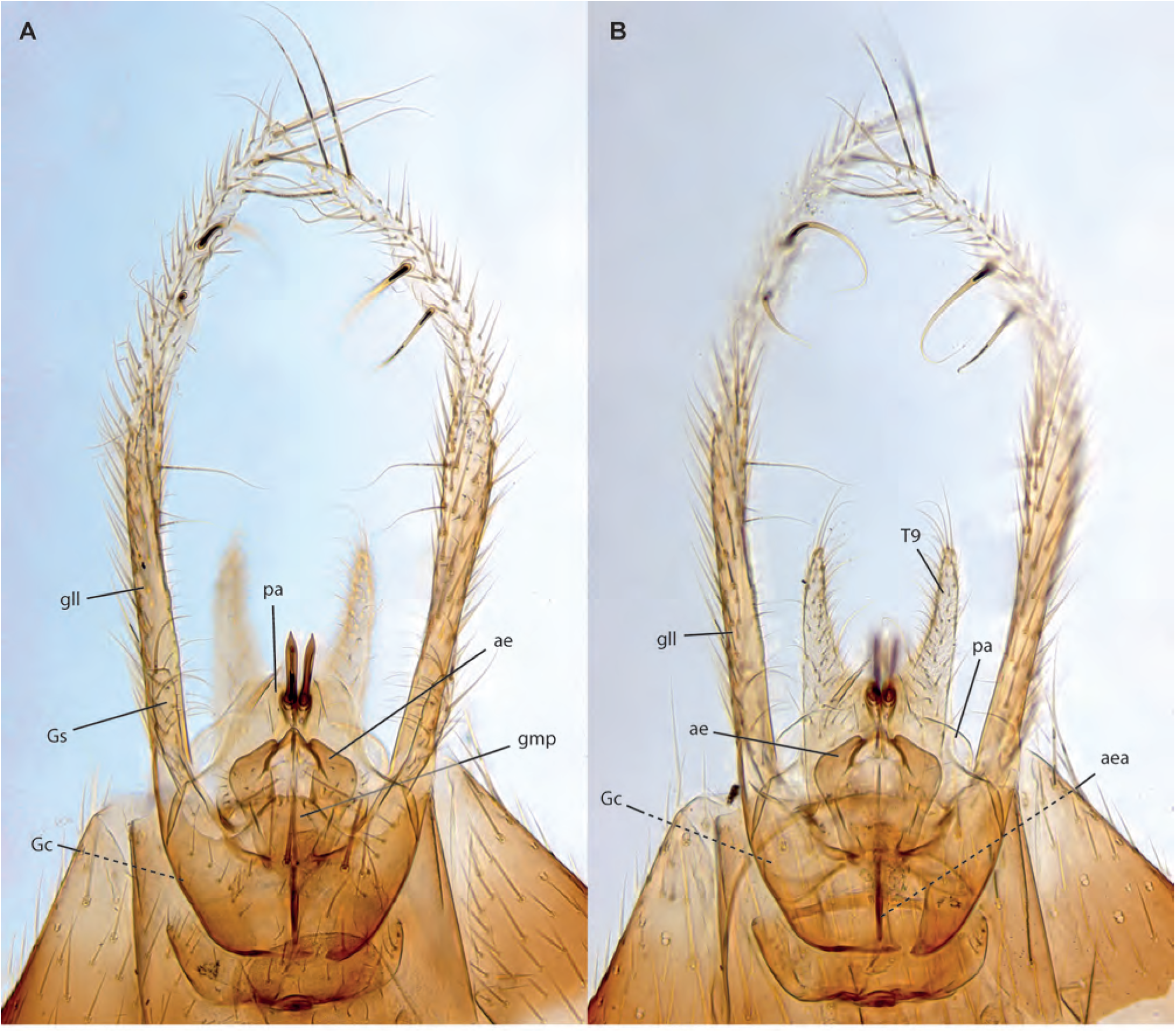
*Epicypta ganengsengi* Amorim & Oliveira, **sp. n.** male holotype, terminalia. **A.** Ventral view. **B.** Dorsal view.

**Figs. 134A-D.**
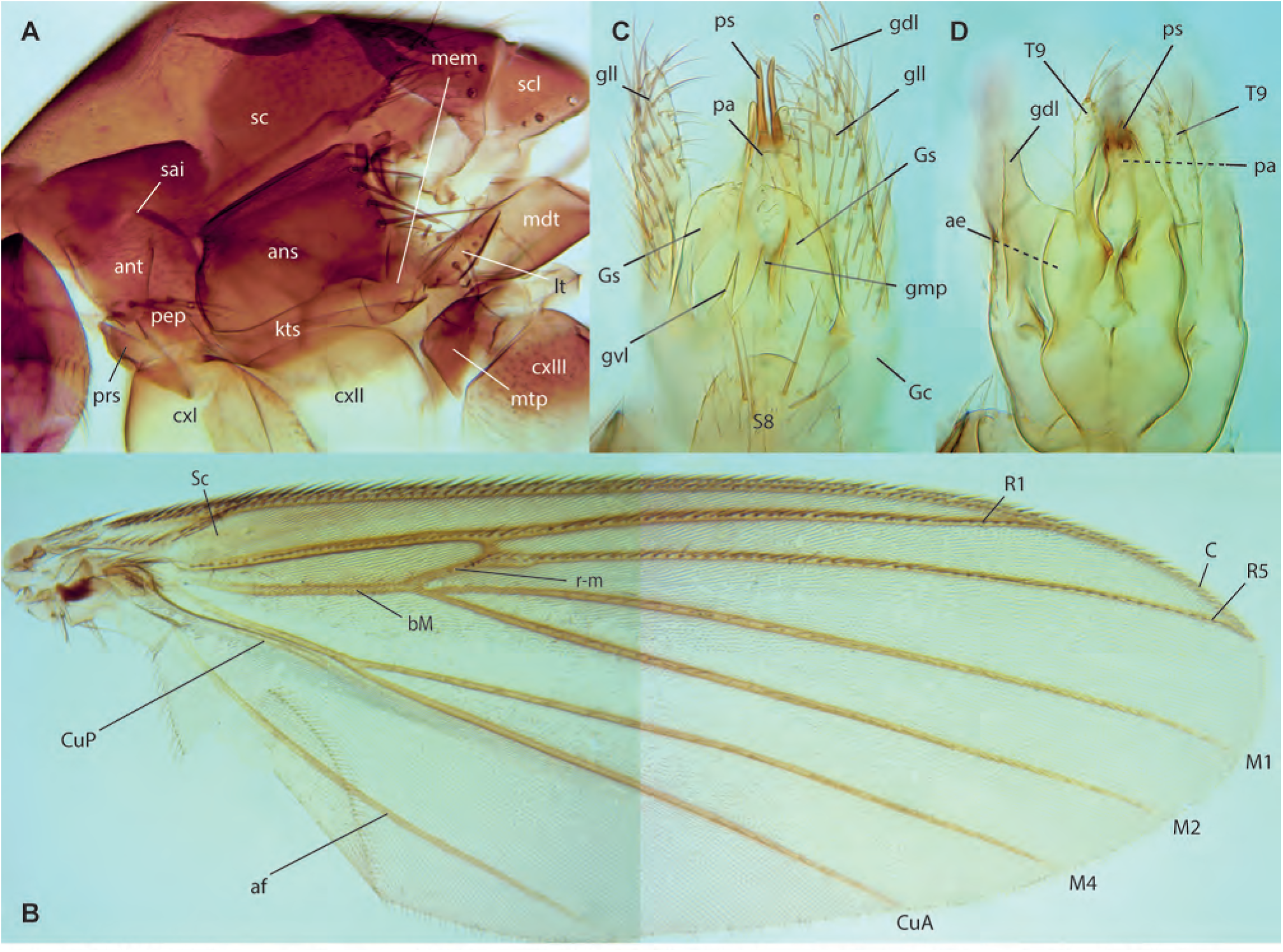
*Epicypta* sp. A, male. **A.** Thorax, lateral view. **B.** Wing. **C.** Terminalia, ventral view. **D.** Terminalia, dorsal view.

**Figs. 135A-D.**
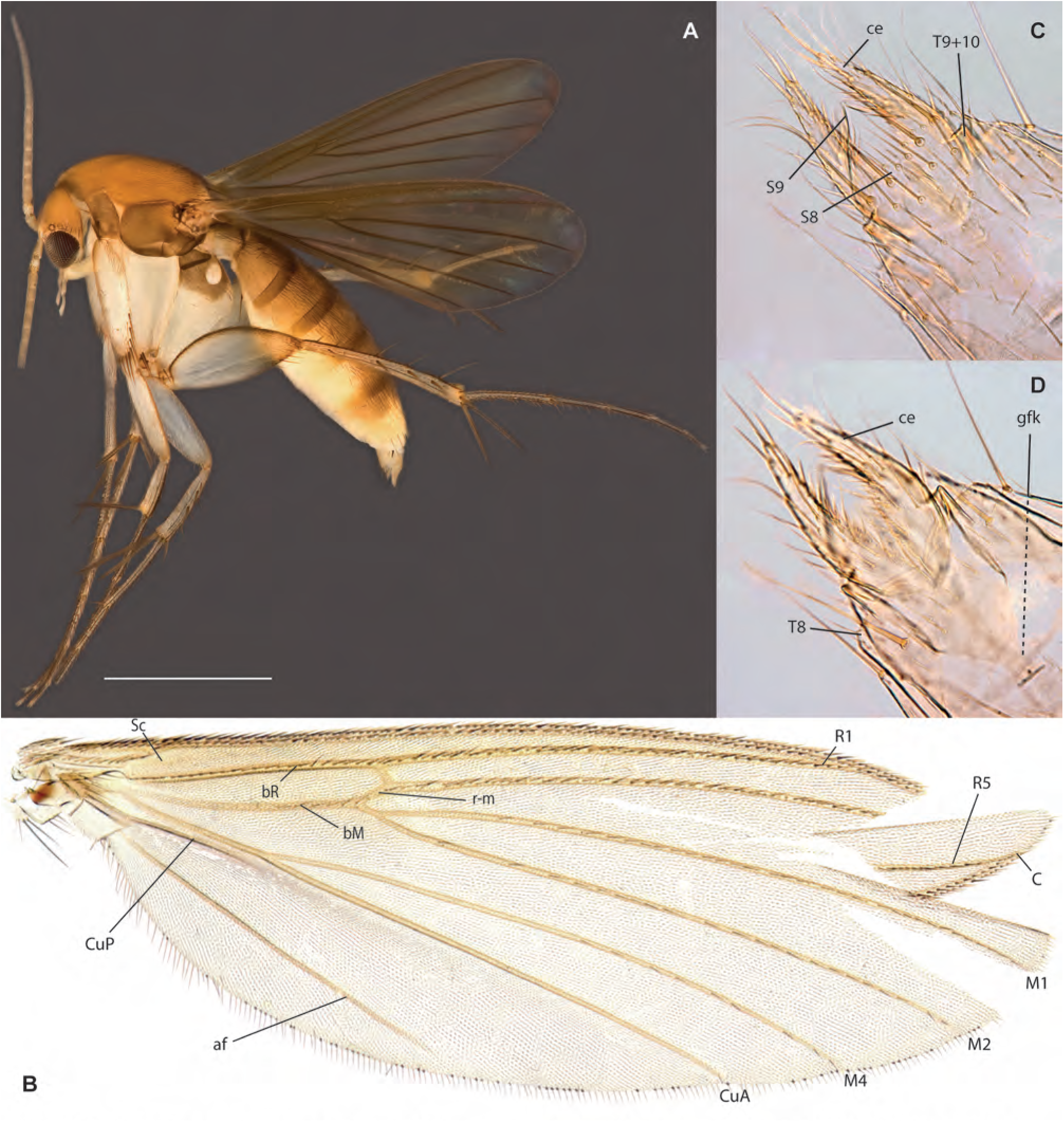
*Epicypta nanyangu* Amorim & Oliveira, **sp. n. A.** Habitus, lateral view, female paratype, ZRCBDP0048455. **B.** Wing, female paratype ZRCBDP0048926. **C.** Female terminalia, ventral view, same. **D.** Female terminalia, dorsal view, same.

**Figs. 136A-B.**
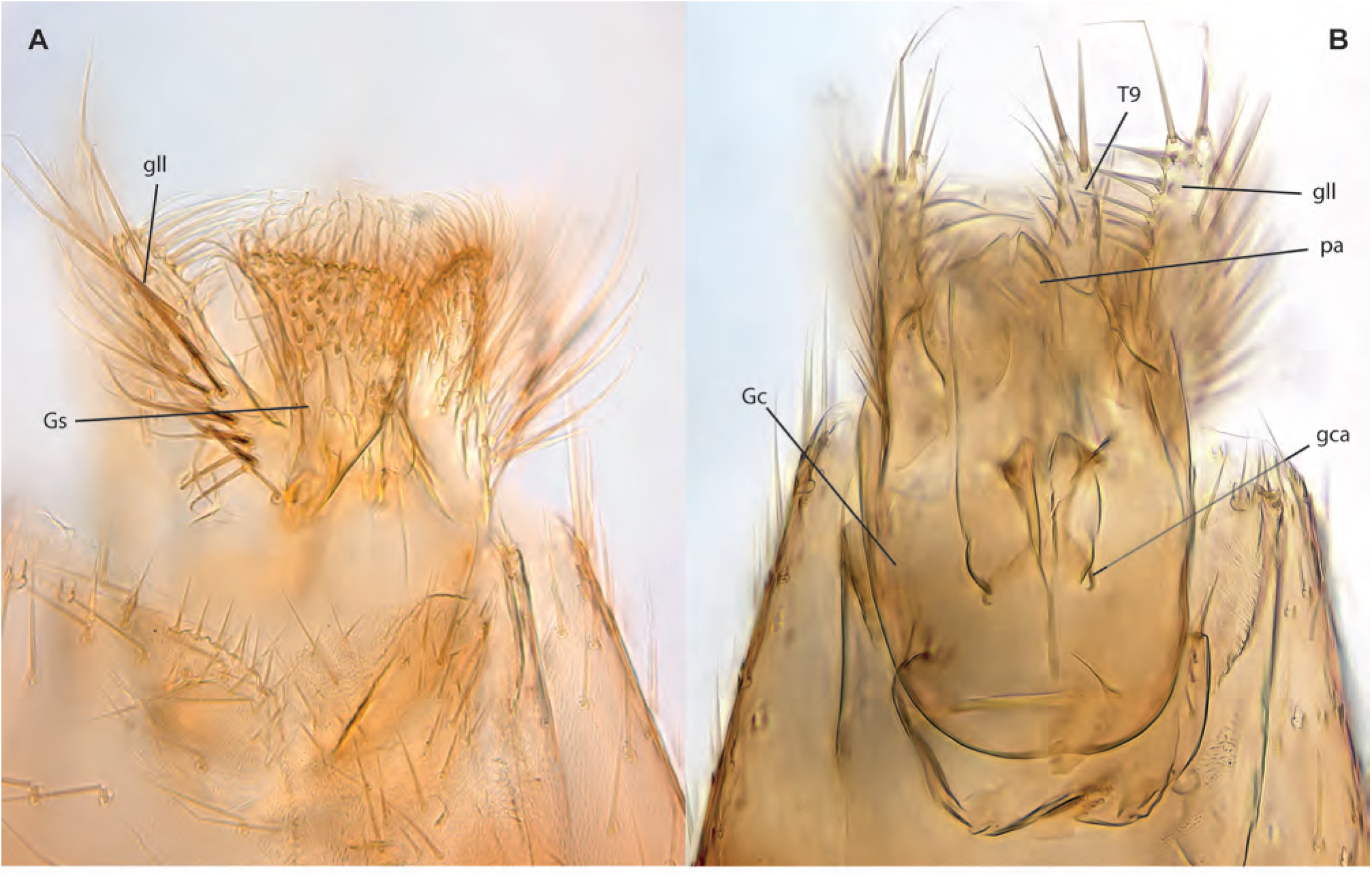
*Epicypta nanyangu* Amorim & Oliveira, **sp. n.,** male terminalia. **A.** Ventral view, paratype ZRCBDP0048864. **B.** Dorsal view, paratype ZRCBDP0049088.

**Figs. 137A-D.**
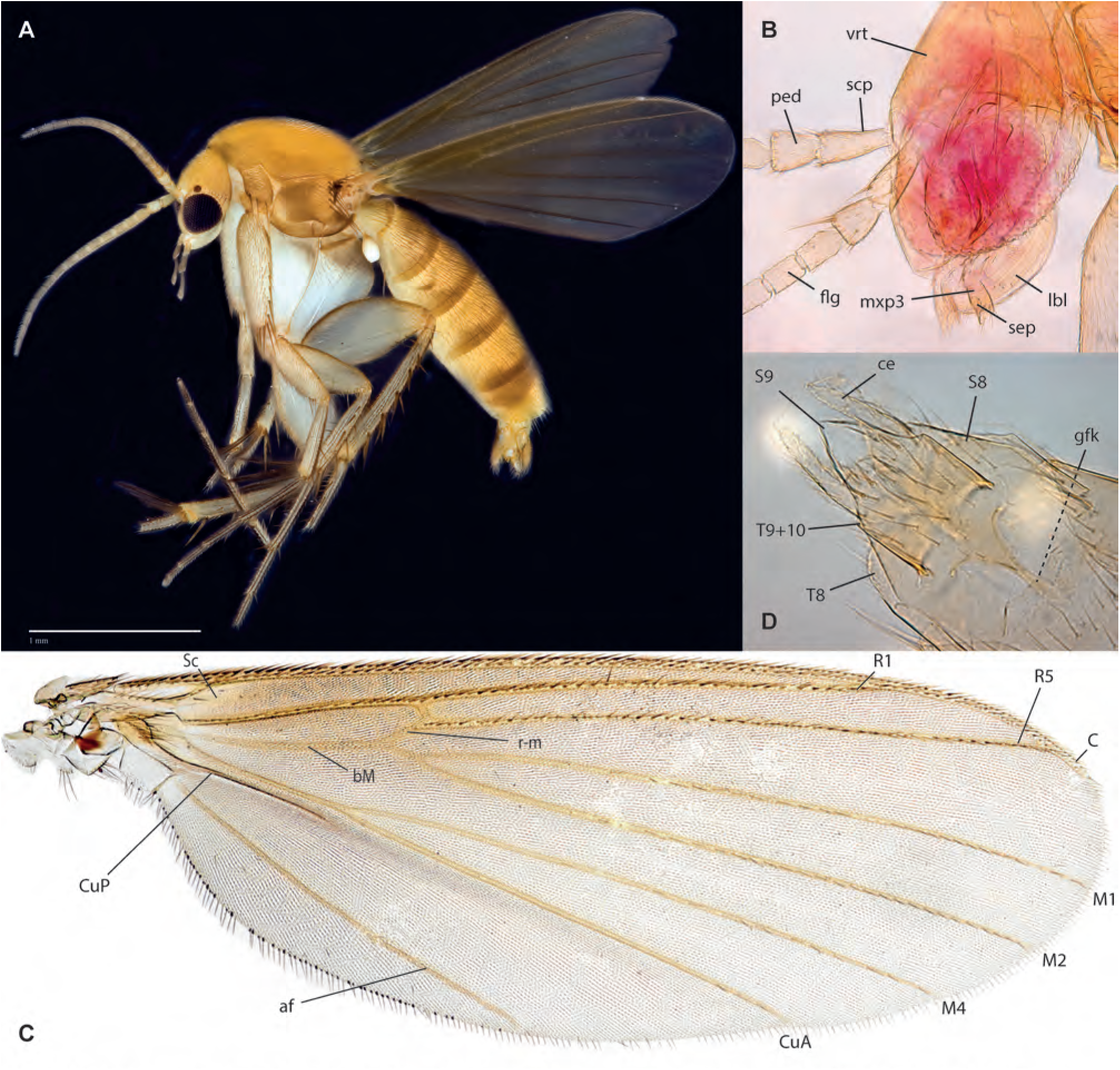
*Epicypta nus* Amorim & Oliveira, **sp. n. A.** Habitus, lateral view, female paratype ZRCBDP0048469. **B.** Head, female paratype ZRCBDP0048900 (maxillary palpomeres 4–5 missing). **C.** Wing, same. **D.** Female terminalia, ventral view, same.

**Figs. 138A-B.**
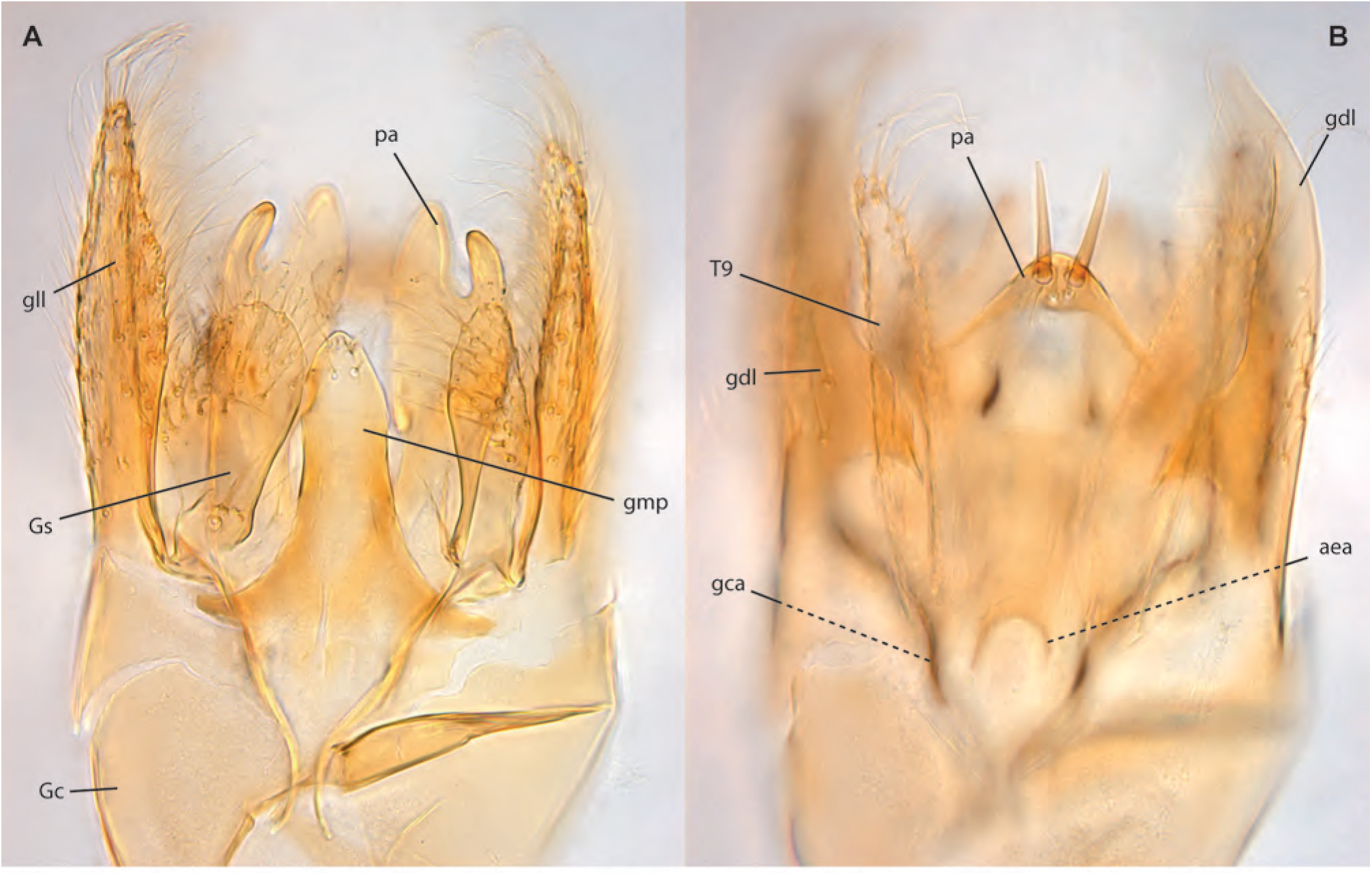
*Epicypta nus* Amorim & Oliveira, **sp. n.,** male terminalia, holotype. **A.** Ventral view (gonocoxite lateral lobe broken and slightly displaced). **B.** Dorsal view.

**Figs. 139A-E.**
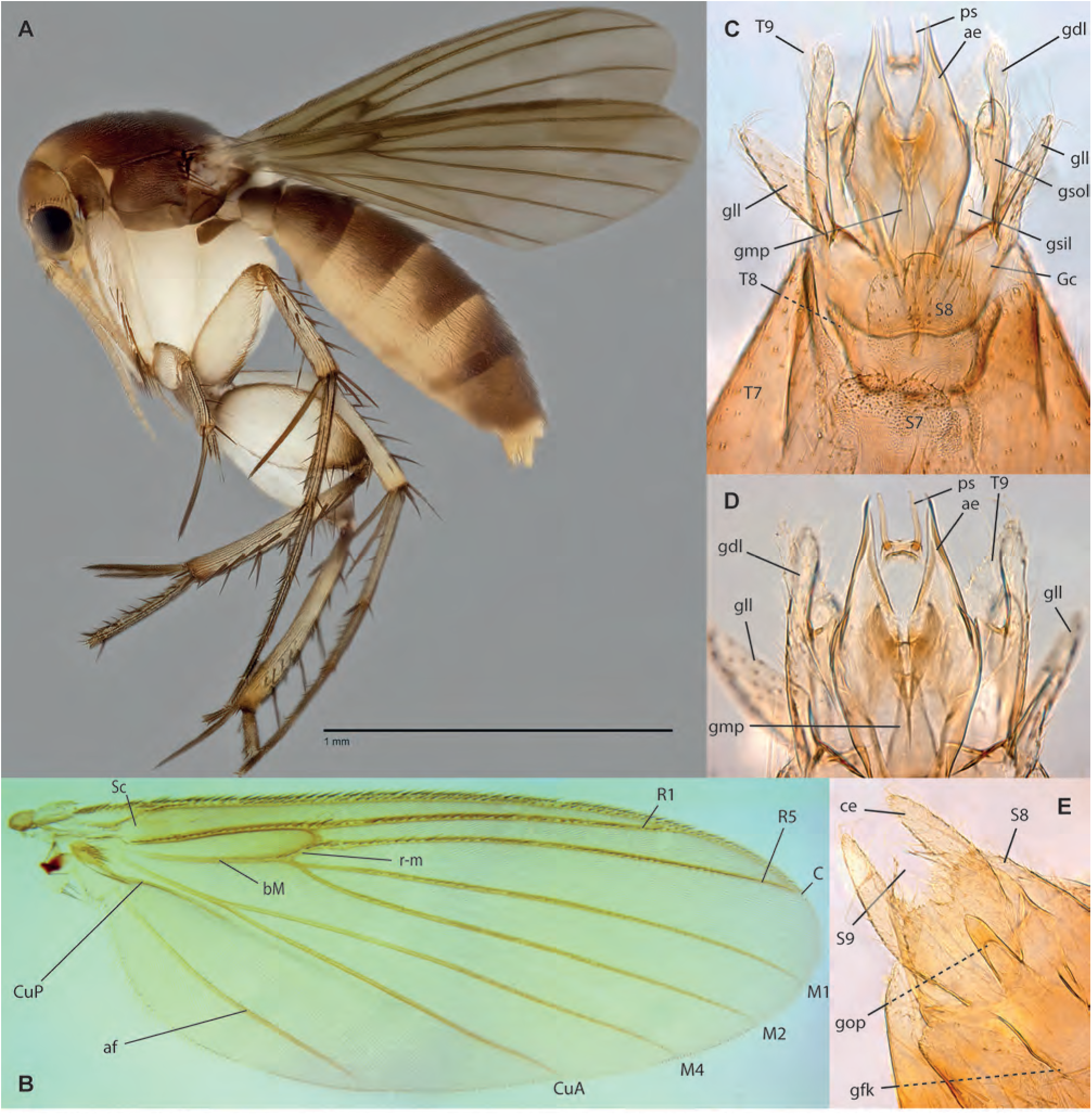
*Epicypta peterngi* Amorim & Oliveira, **sp. n. A.** Habitus, lateral view, female paratype ZRCBDP0047927. **B.** Wing, male holotype. **C.** Male terminalia, ventral view, same. **D.** Detail of distal end of male terminalia, same. **E.** Female terminalia, ventral view, ZRCBDP0047927.

**Figs. 140A-D.**
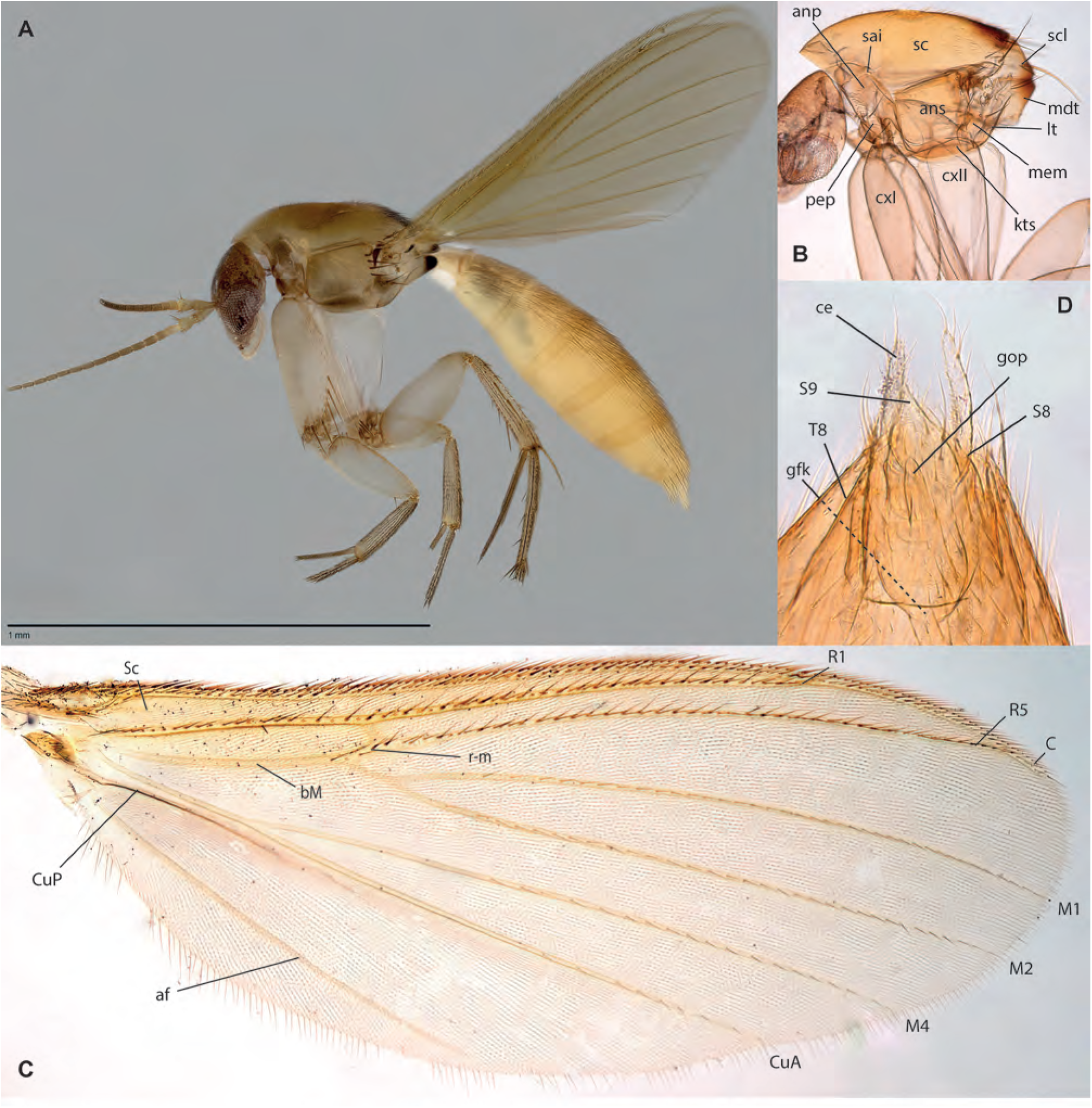
*Epicypta maggielimae* Amorim & Oliveira, **sp. n.,** female holotype. **A.** Habitus, lateral view. **B.** Thorax, lateral view. **C.** Wing. **D.** Terminalia, ventral view.

**Figs. 141A-C.**
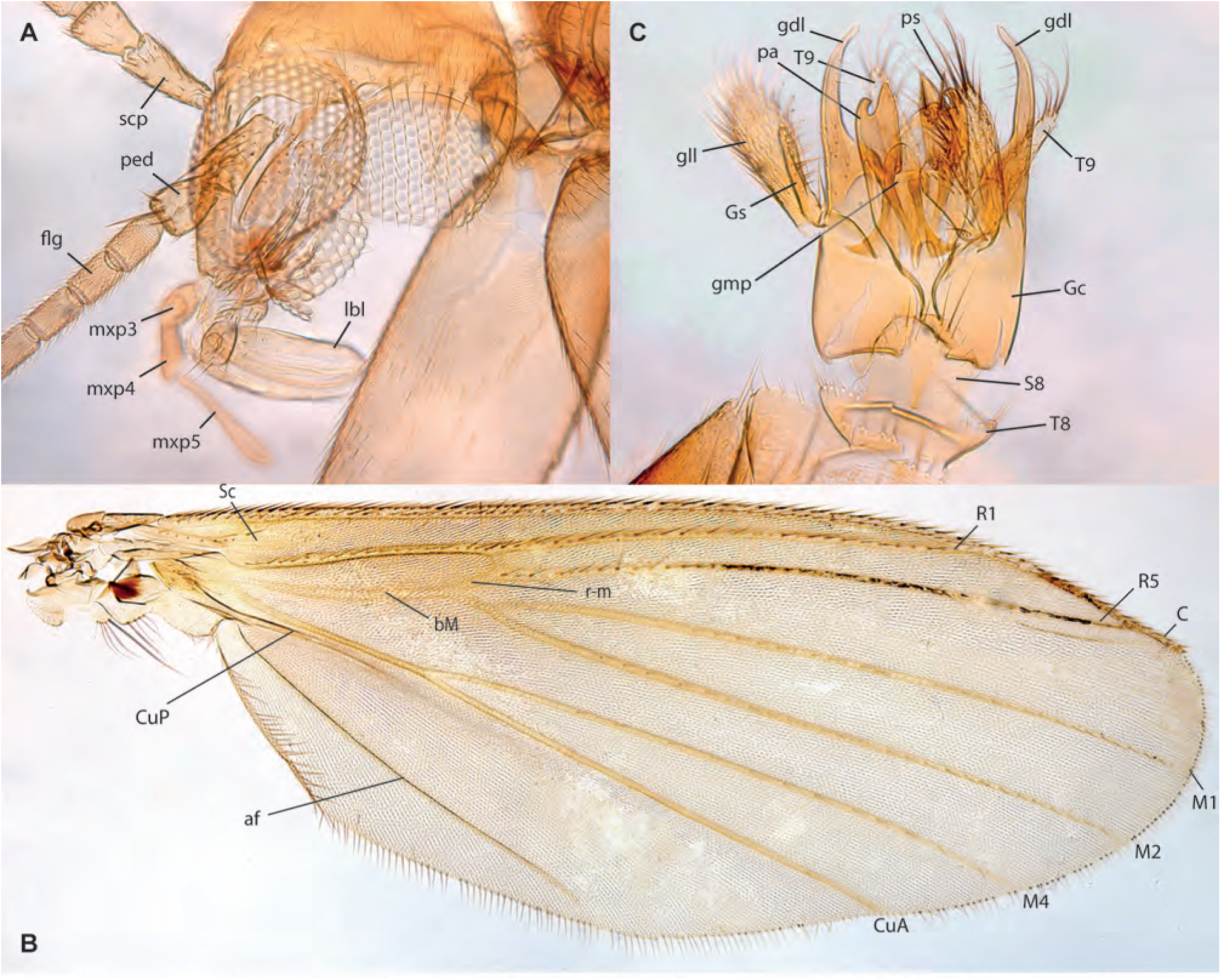
*Epicypta yupeigaoae* Amorim & Oliveira, **sp. n.**, male holotype. **A.** Head, lateral view. **B.** Wing. **C.** Male terminalia, ventral view.

**Figs. 142A-D.**
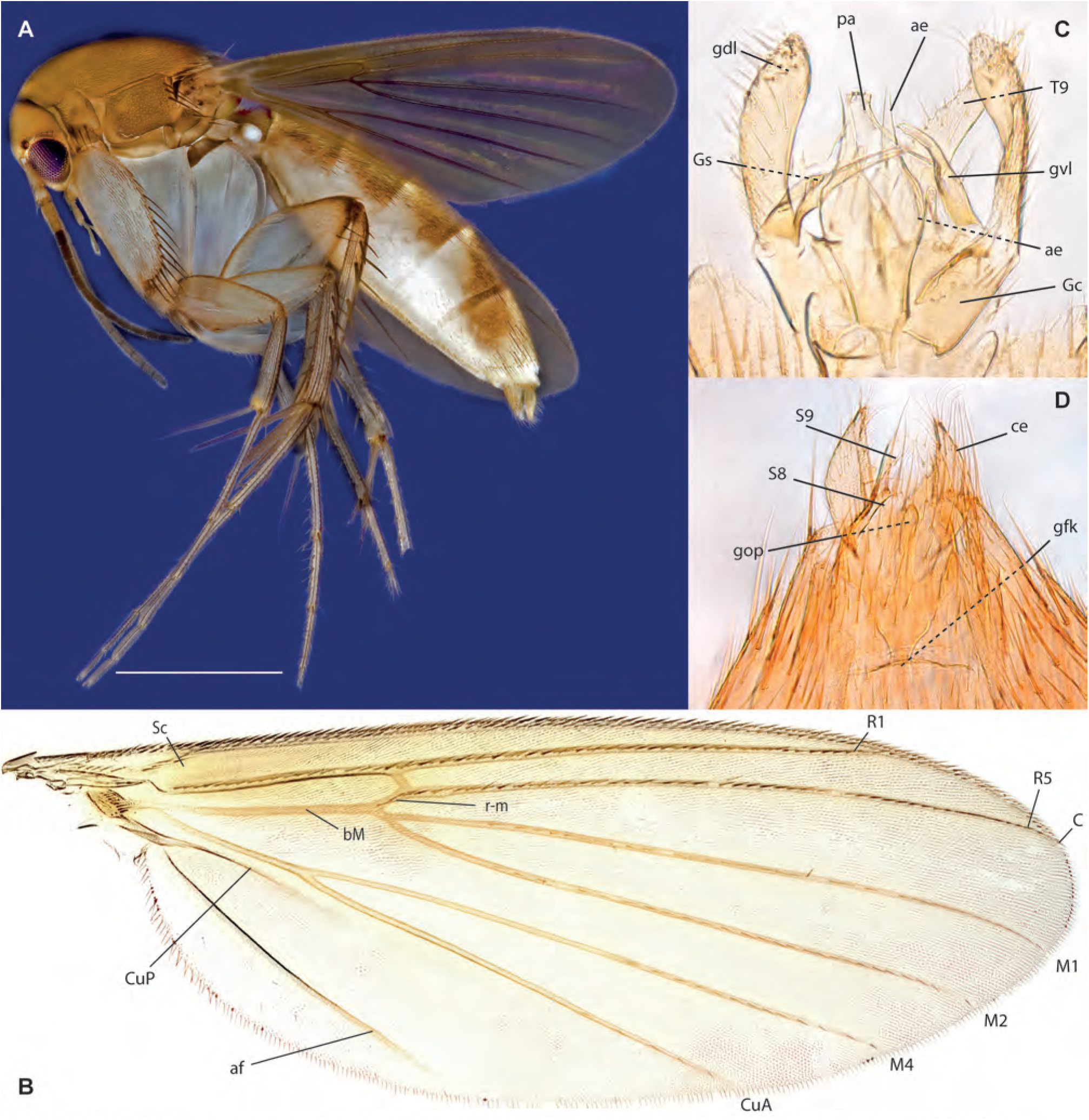
*Epicypta annwee* Amorim & Oliveira, **sp. n. A.** Habitus, lateral view, female paratype, ZRCBDP0048446. **B.** Wing, male holotype. **C.** Male terminalia, ventral view, same. **D.** Female terminalia, ventral view, paratype ZRCBDP0047917.

**Figs. 143A-D.**
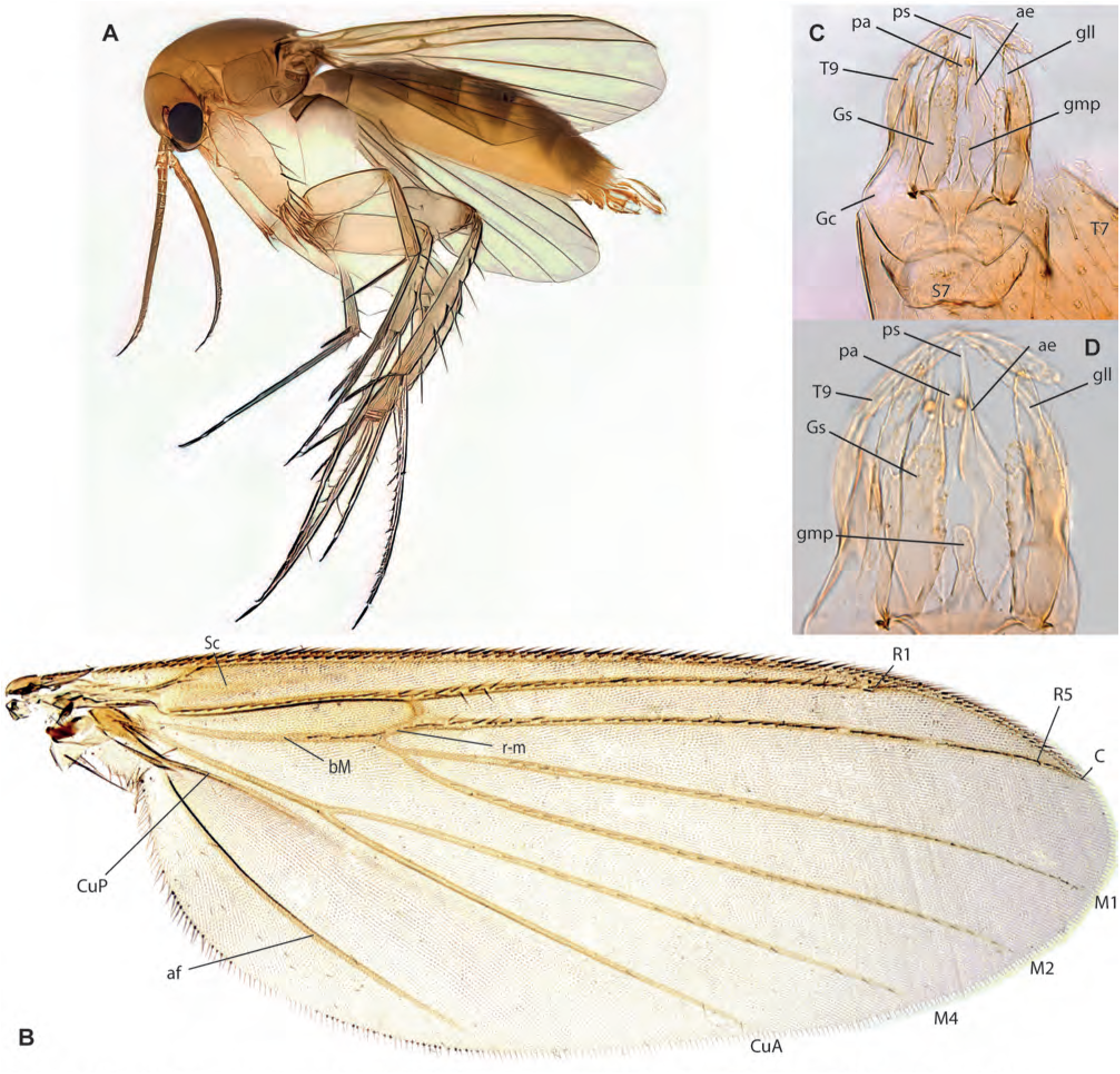
*Epicypta wallacei* Amorim & Oliveira, **sp. n. A.** Habitus, lateral view, male paratype ZRCBDP0048465. **B.** Wing, male holotype. **C.** Male terminalia, ventral view, same. **D.** Detail of distal end of male terminalia, same.

**Figs. 144A-B.**
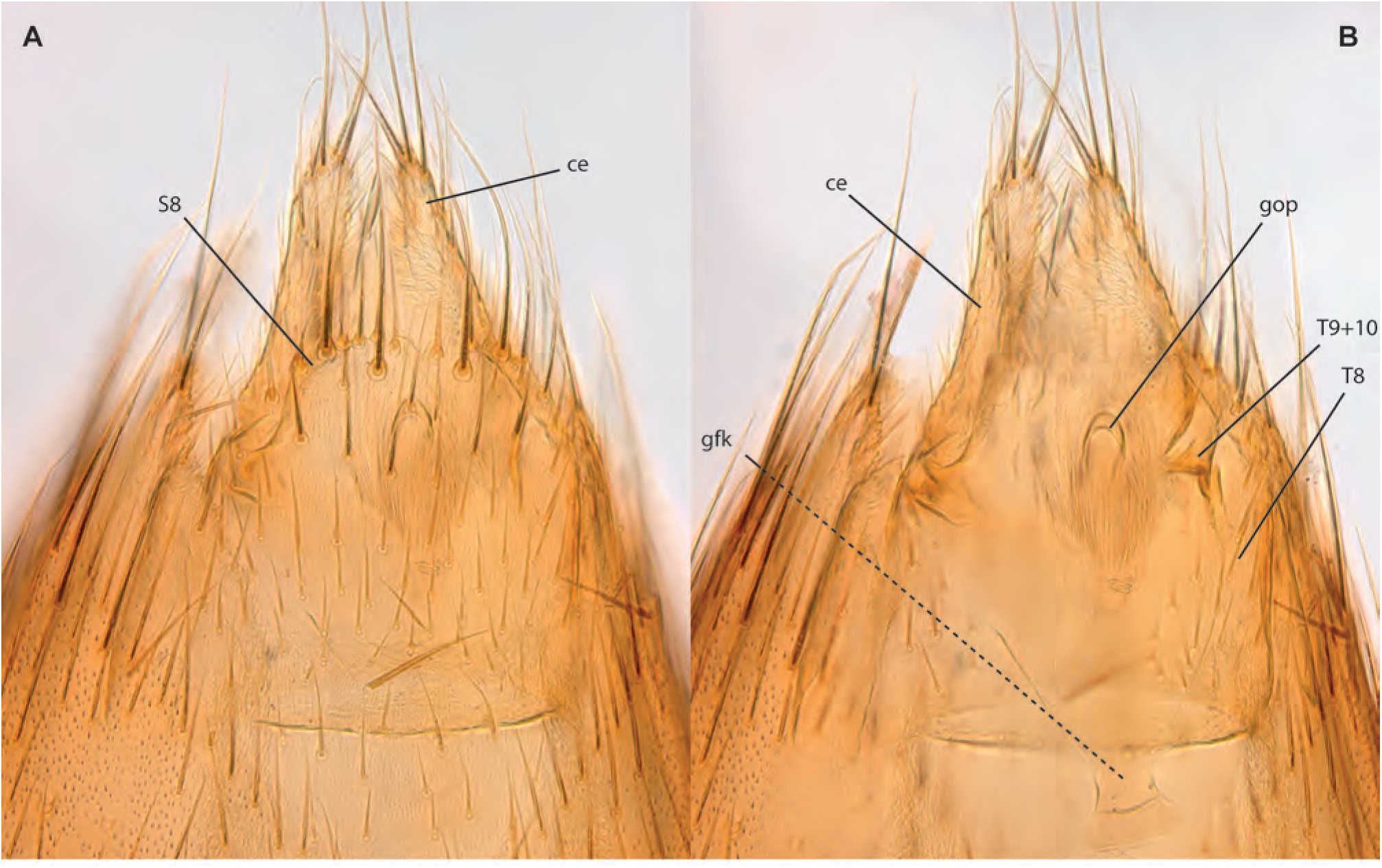
*Epicypta wallacei* Amorim & Oliveira, **sp. n.,** female terminalia, paratype ZRCBDP0048868. **A.** Ventral view. **B.** Dorsal view.

**Figs. 145A-E.**
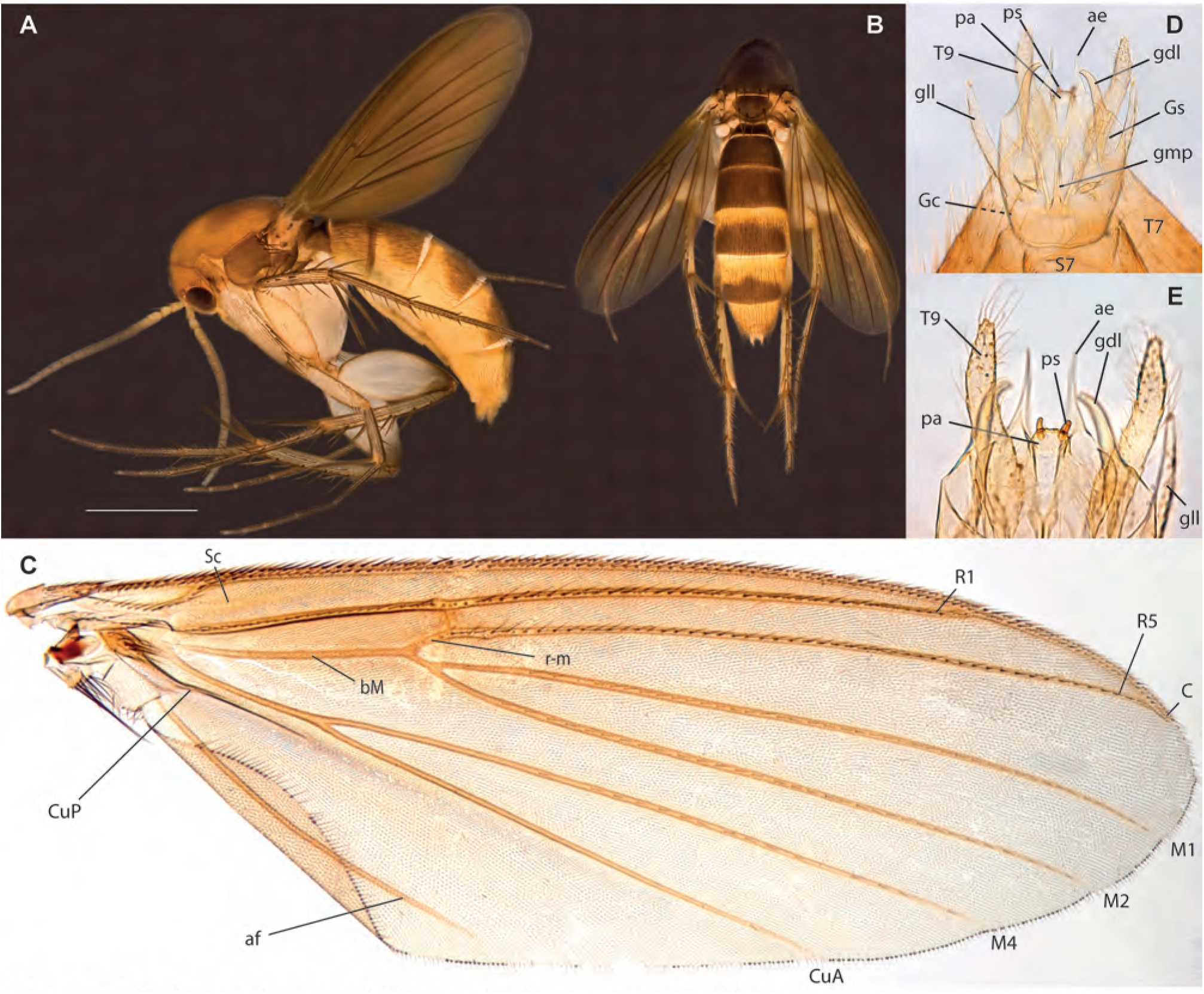
*Epicypta lamtoongjini* Amorim & Oliveira, **sp. n. A.** Habitus, lateral view, lateral view, female paratype ZRCBDP0048440. **B.** Same, dorsal view. **C.** Wing, male holotype. **D.** Male terminalia, ventral view, same. **E.** Detail of distal end of male terminalia, same.

**Figs. 146A-D.**
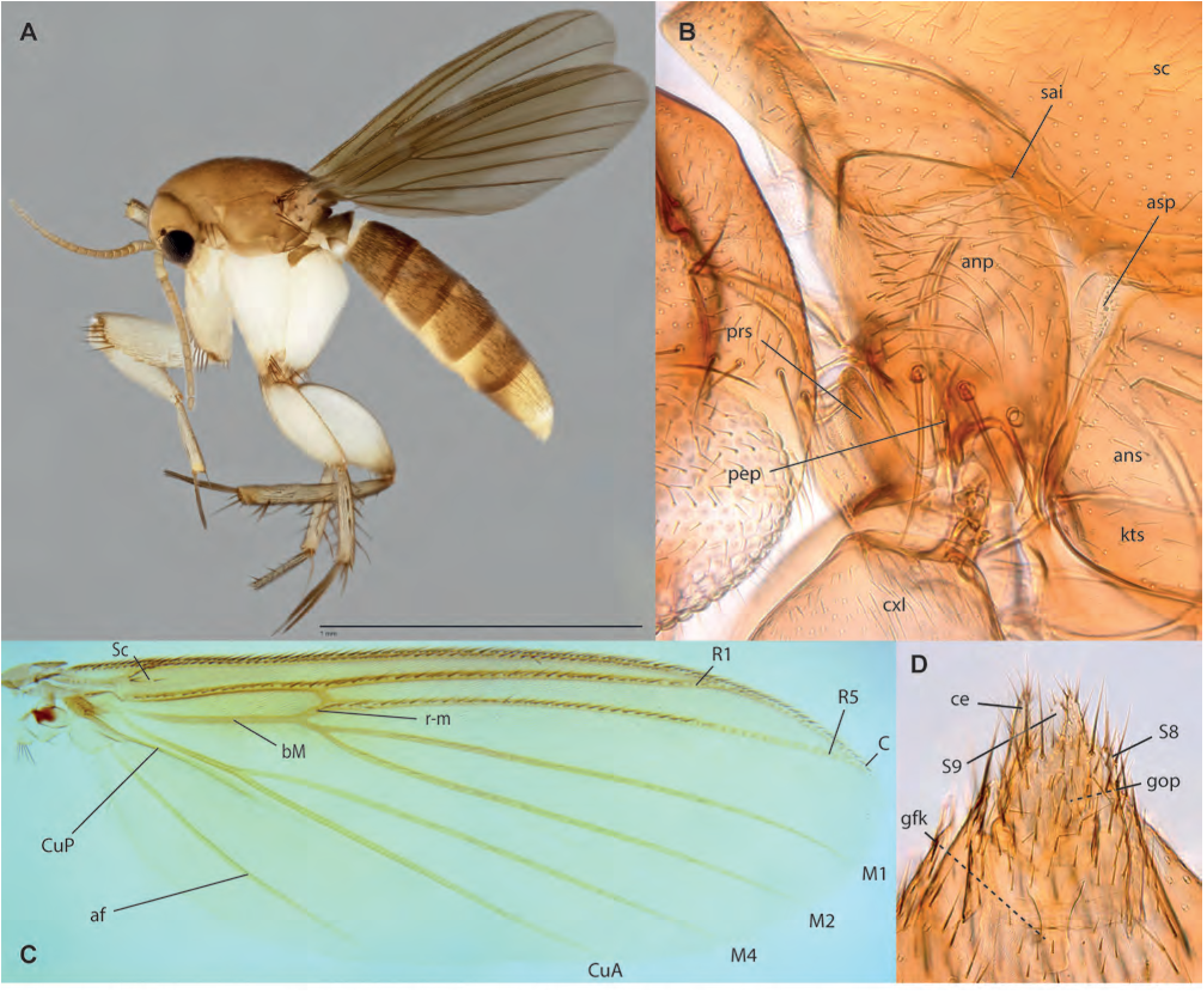
*Epicypta catherinelimae* Amorim & Oliveira, **sp. n. A.** Habitus, lateral view, female paratype ZRCBDP0048062. **B.** Anterior end of thoracic pleura, lateral view, female holotype. **C.** Wing, same. **D.** Female terminalia, ventral view, same.

**Figs. 147A-C.**
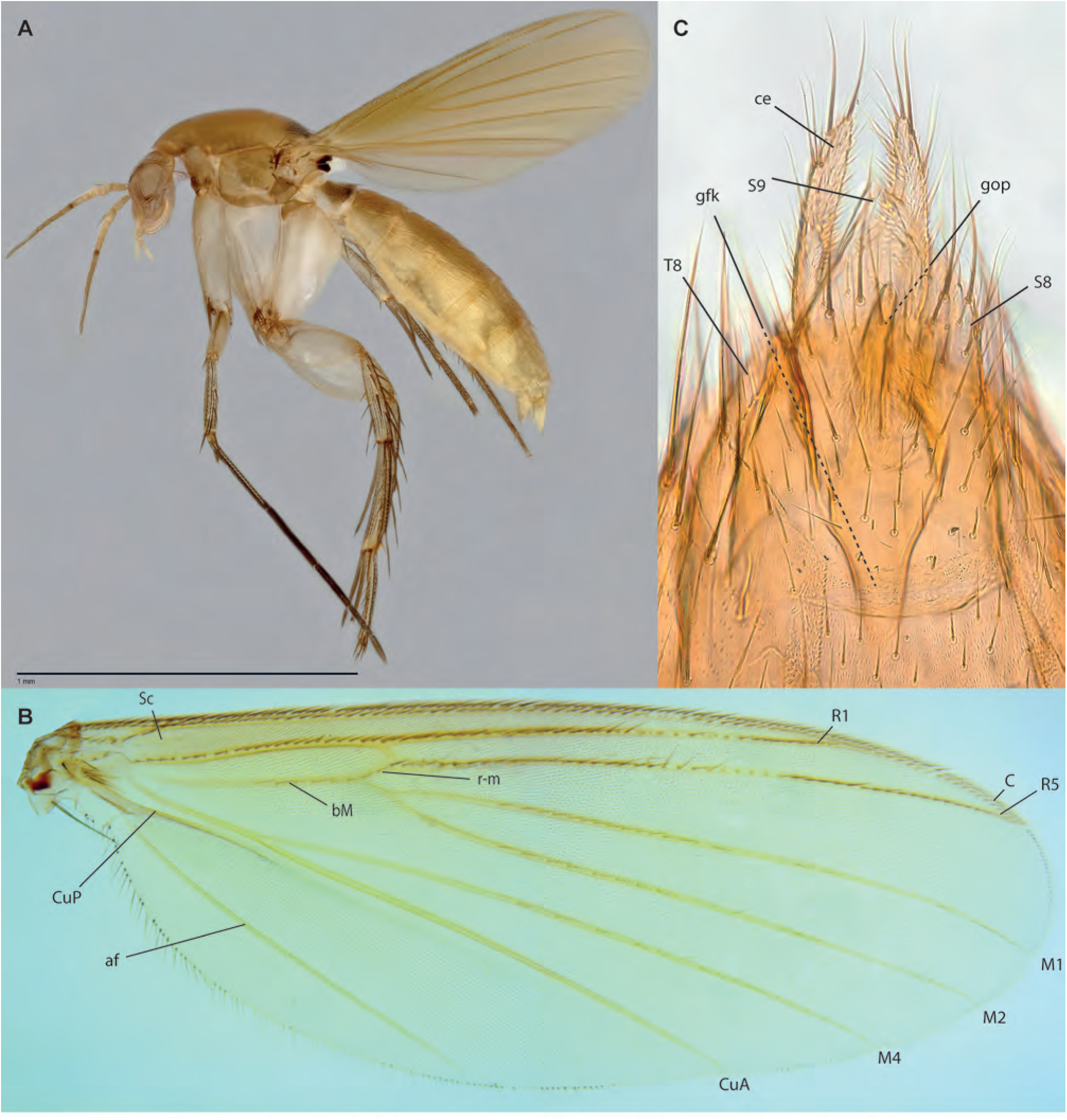
*Epicypta grootaerti* Amorim & Oliveira, **sp. n. A.** Habitus, lateral view, female holotype. **B.** Wing, same. **C.** Female terminalia, ventral view, same.

**Figs. 148A-D.**
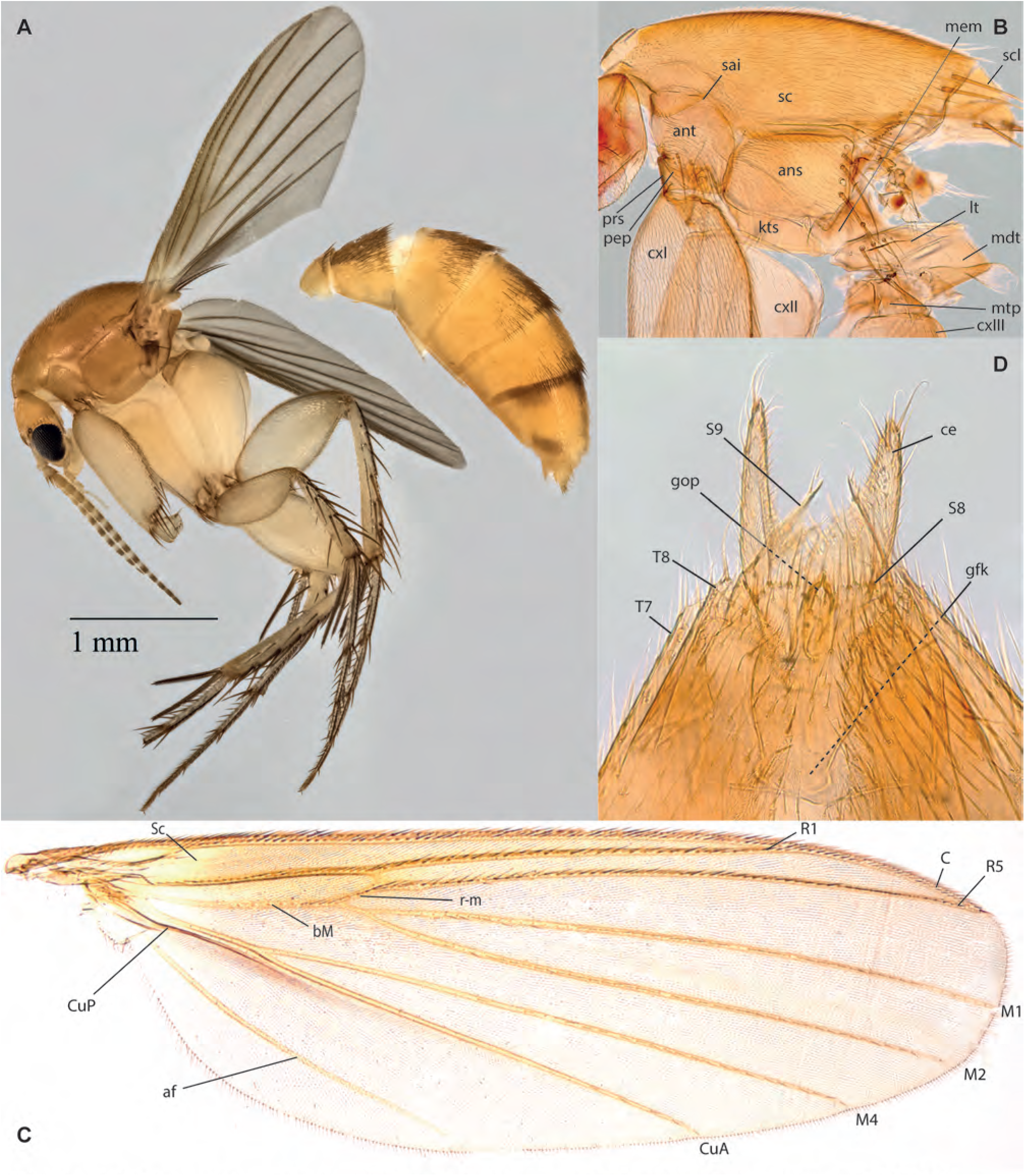
*Epicypta joaquimae* Amorim & Oliveira, **sp. n. A.** Habitus, lateral view, female paratype ZRCBDP0137314. **B.** Thorax, lateral view, female holotype. **C.** Wing, same. **D.** Female terminalia, ventral view, same.

**Figs. 149A-D.**
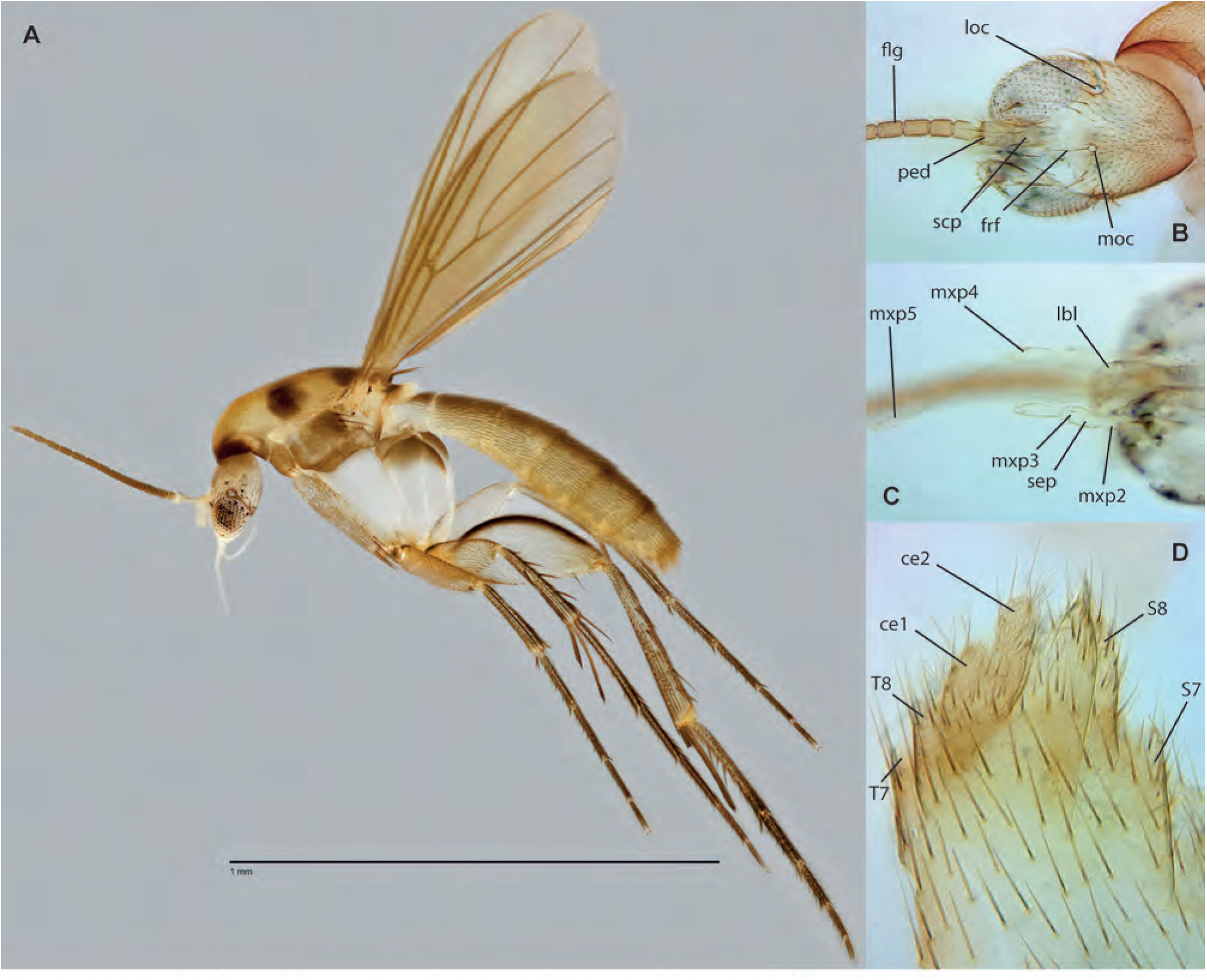
*Aspidionia cheesweeleeae* Amorim & Oliveira, **sp. n.,** female holotype. **A.** Habitus, lateral view. **B.** Head, dorsal view. **C.** Head, ventral view. **D.** Terminalia, lateral view.

**Figs. 150A-B.**
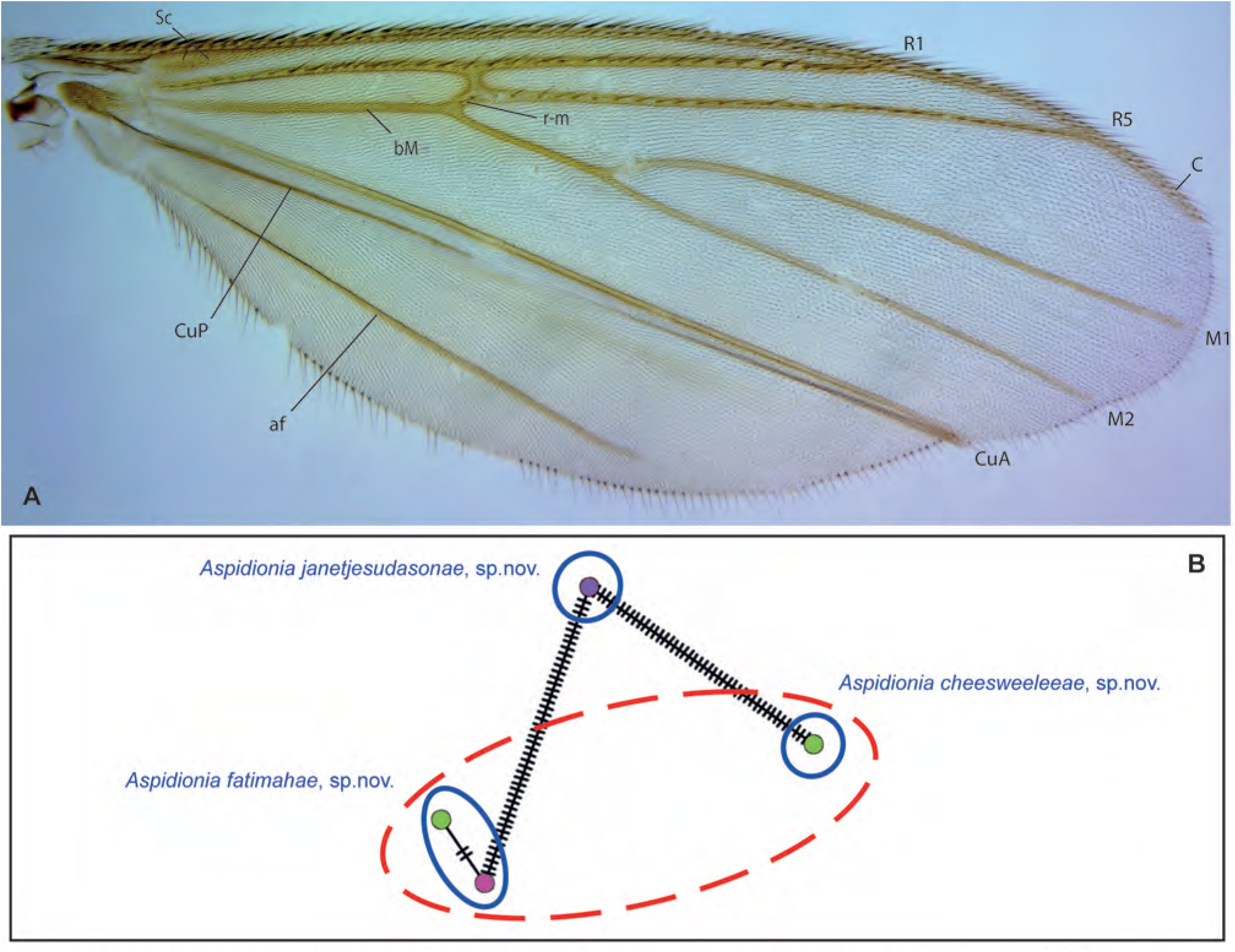
A. *Aspidionia cheesweeleeae* Amorim & Oliveira, **sp. n.,** wing, female holotype. **B.** Haplotype network for *Aspidionia*.

**Figs. 151A-F.**
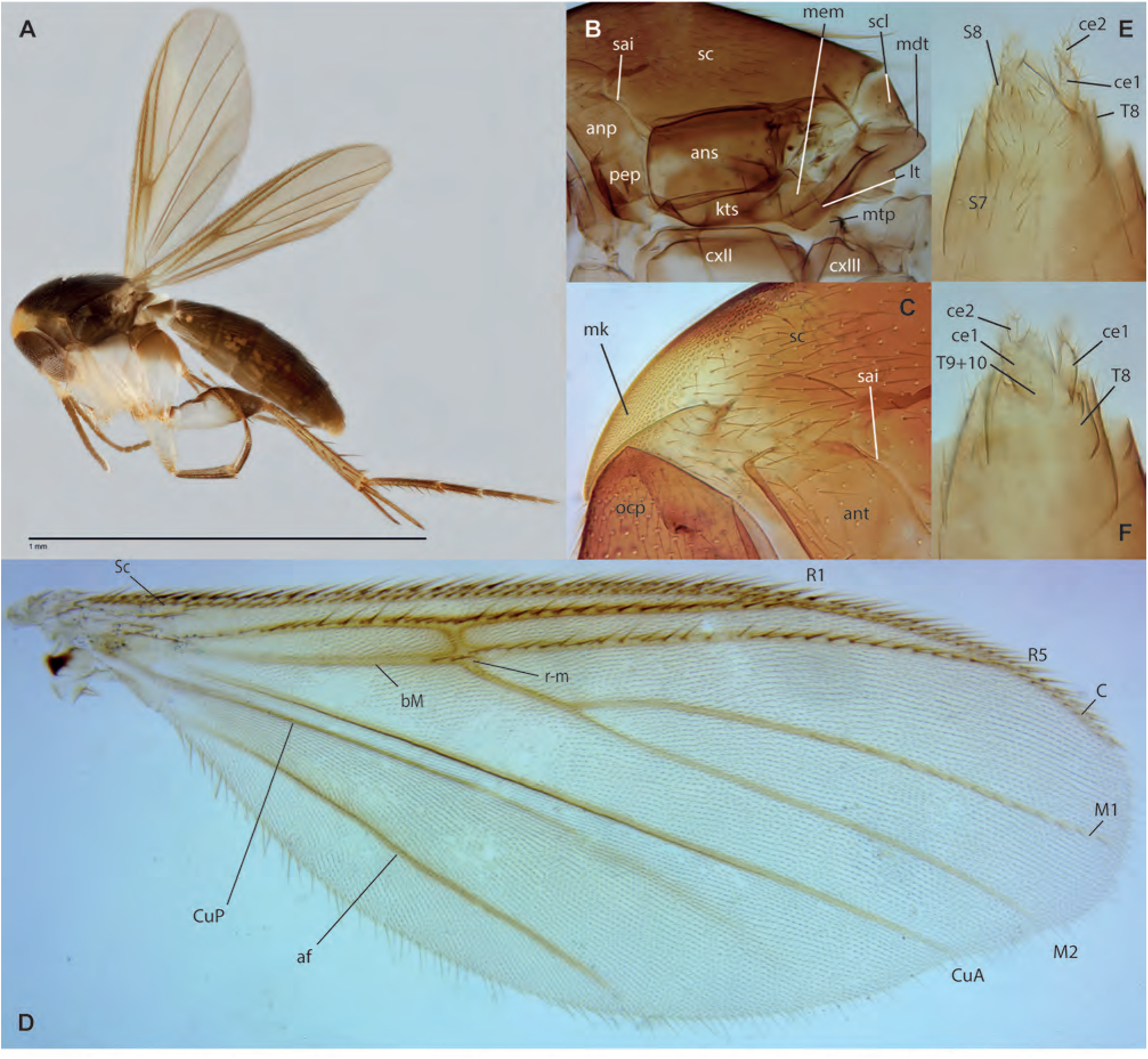
*Aspidionia janetjesudasonae* Amorim & Oliveira, **sp. n.**, female holotype. **A.** Habitus, lateral view. **B.** Thorax, lateral view. **C.** Anterior end of scutum, lateral view. **D.** Wing. **E.** Terminalia, ventral view. **F.** Terminalia, dorsal view.

**Figs. 152A-E.**
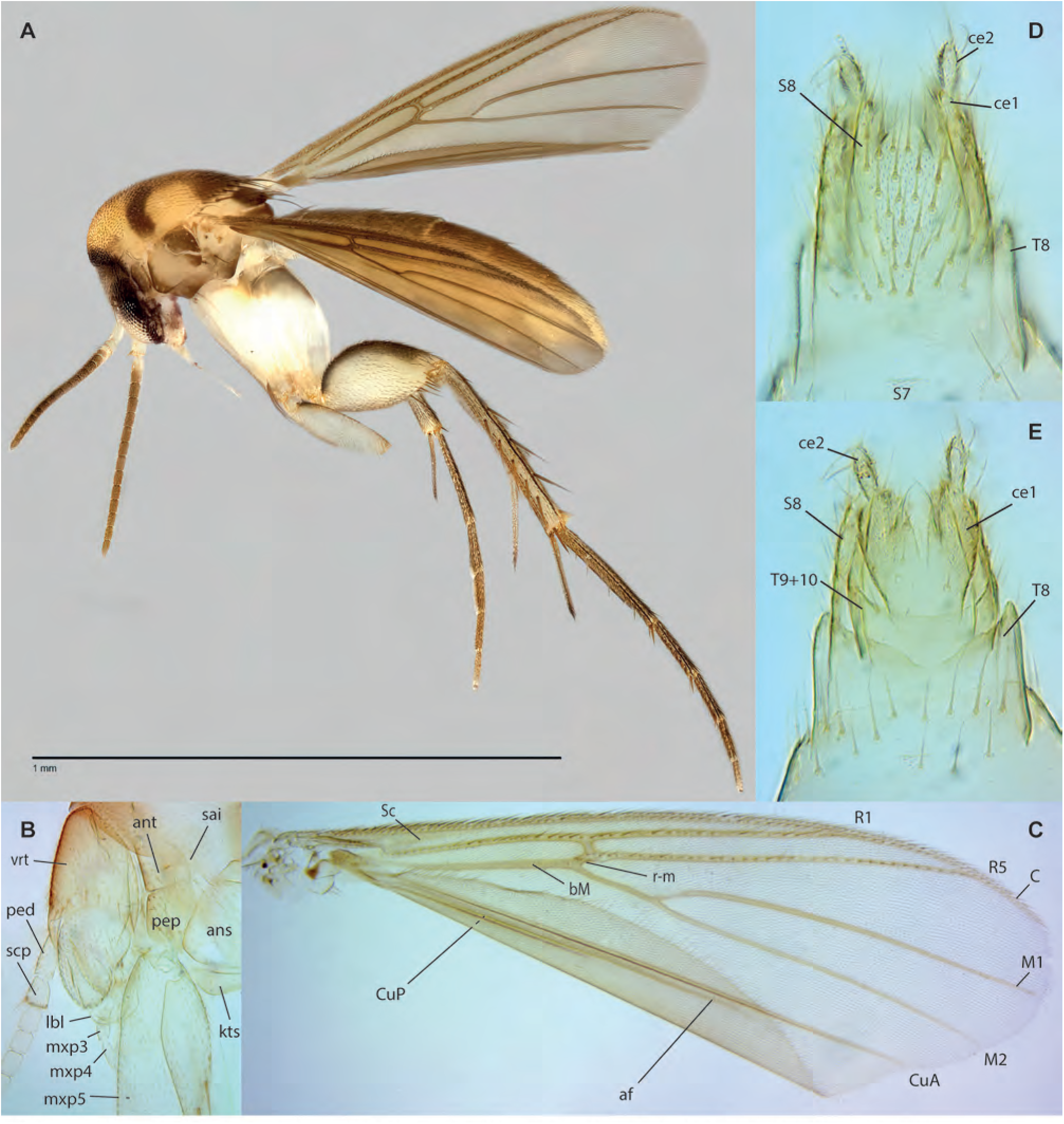
*Aspidionia fatimahae* Amorim & Oliveira, **sp. n.,** female holotype. **A.** Habitus, lateral view. **B.** Head, lateral view. **C.** Wing. **D.** Terminalia, ventral view. **E.** Terminalia, dorsal view.

**Figs. 153A-G.**
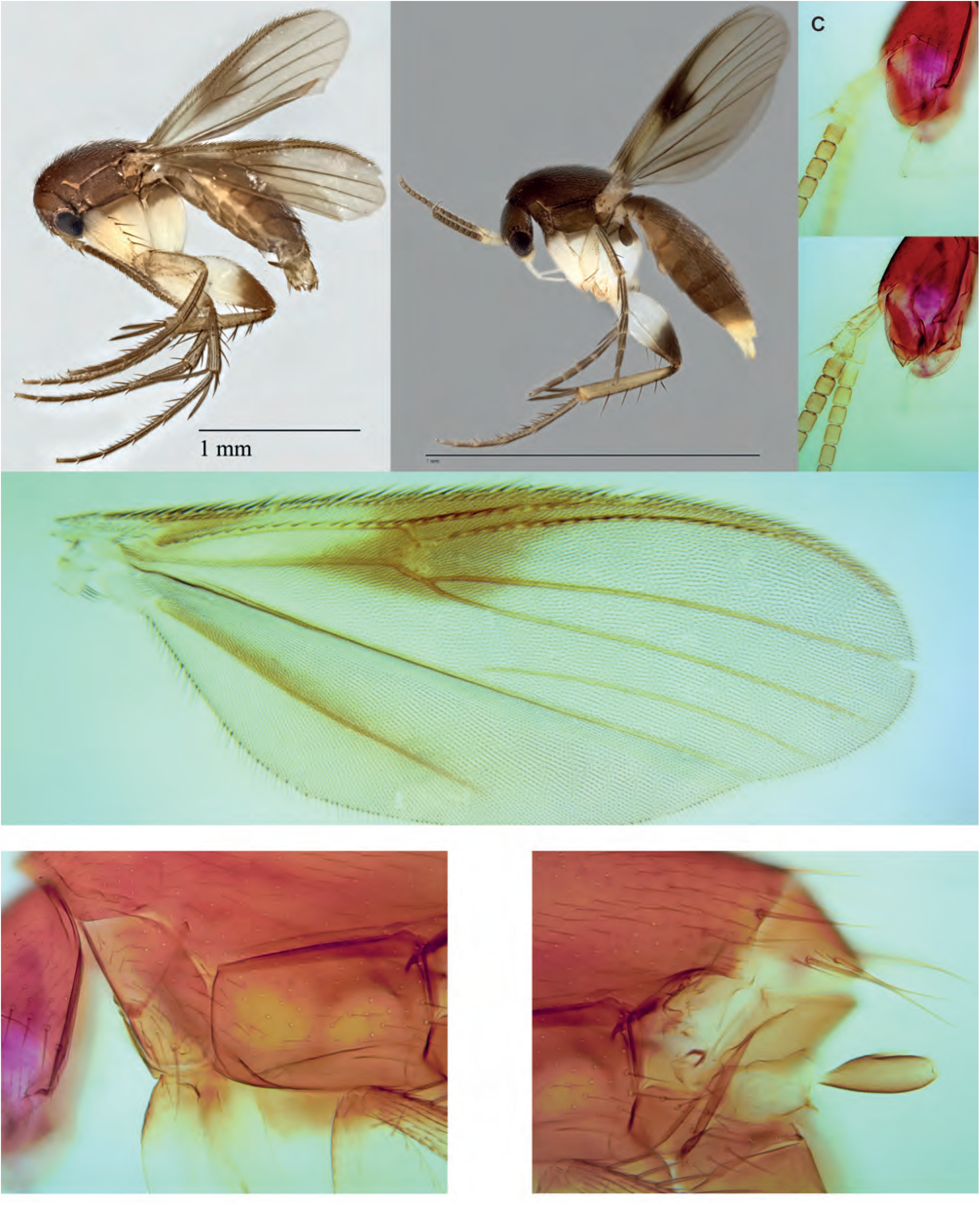
*Integricypta fergusondavie* Amorim & Oliveira, **sp. n. A.** Habitus, lateral view, male, paratype ZRCBDP0155020. **B.** Habitus, lateral view, female, paratype ZRCBDP0049135. **C.** Maxillary palpus, lateral view, male holotype. **D.** Antennae and labella, lateral view, same. **E.** Thorax, lateral view, anterior half, same. **F.** Thorax, posterior half, lateral view, same. **G.** Wing, same.

**Figs. 154A-E.**
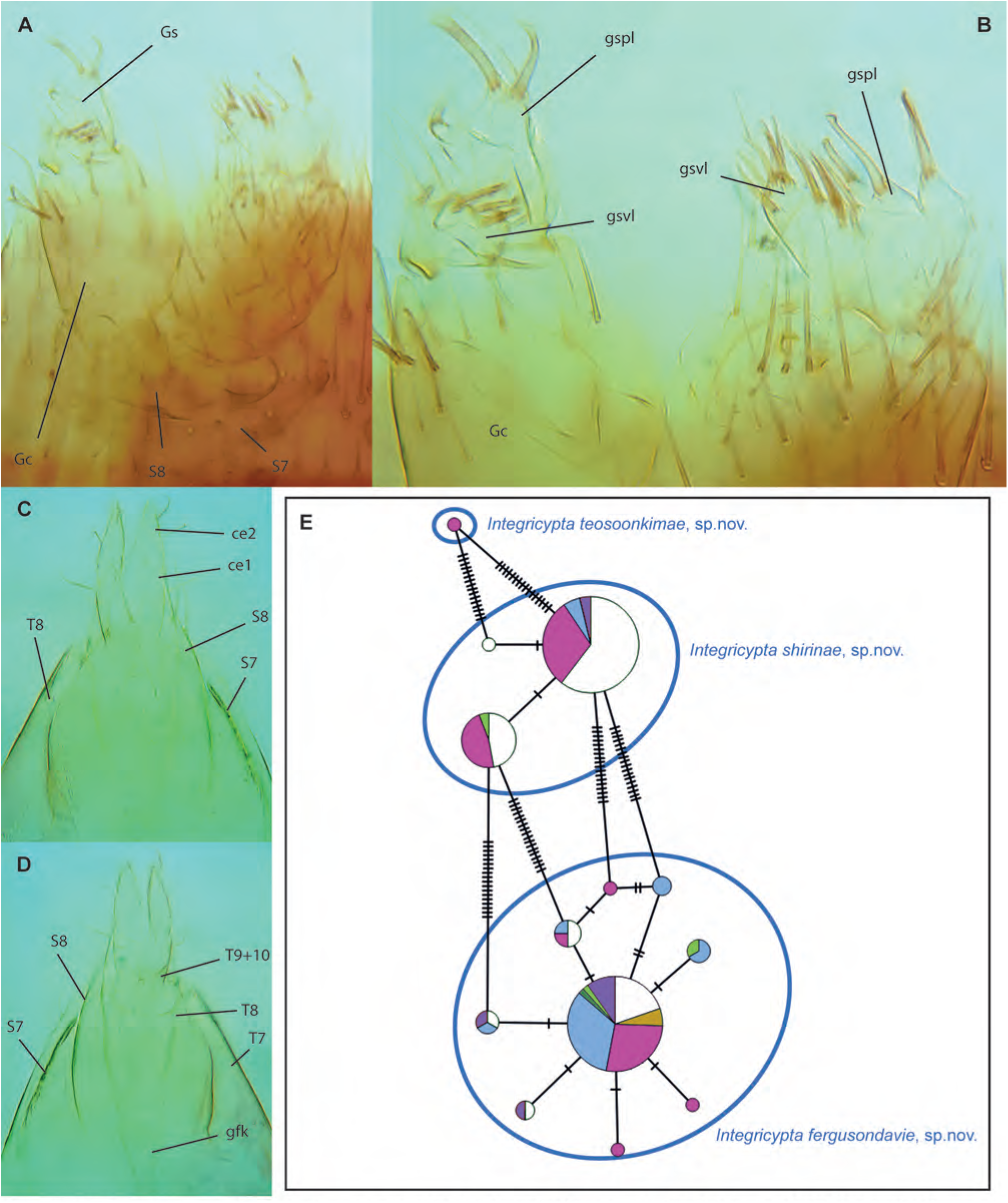
*Integricypta fergusondavie* **sp. n. A.** Male terminalia, ventral view, holotype. **B.** Male terminalia, dorsal view, same. **C.** Female terminalia, ventral view, paratype ZRCBDP0049314. **D.** Female terminalia, ventral view, same. **E.** Haplotype network of *Integricypta Amorim* & Oliveira, **gen. n.** —*I. hoyuenhoeae*, **sp. n.** is not included in the network.

**Figs. 155A-E.**
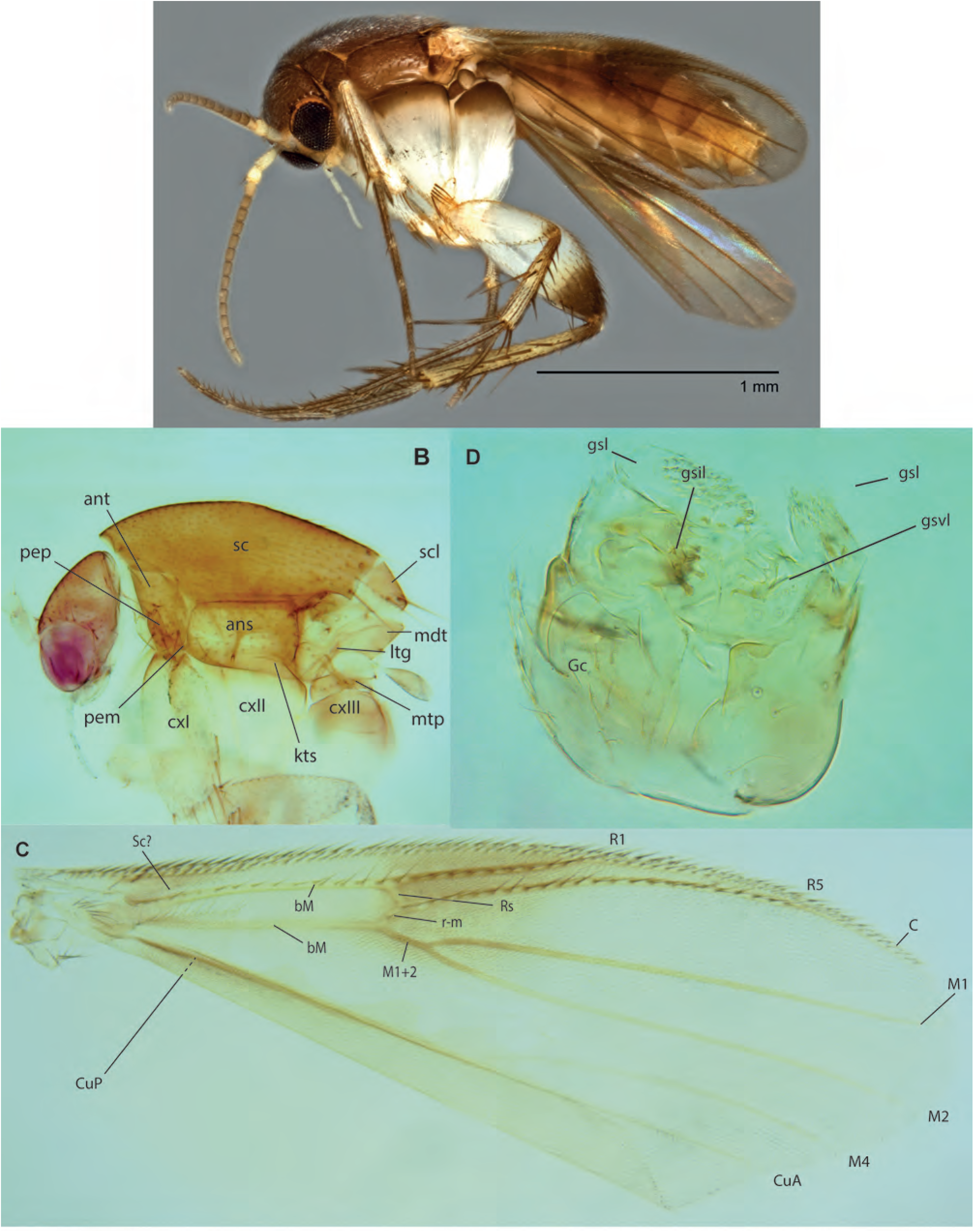
*Integricypta teosoonkimae* Amorim & Oliveira, **sp. n.,** male holotype. **A.** Habitus, lateral view. **B.** Thorax, lateral view. **C.** Wing. **D.** Terminalia, ventral view. **E.** Terminalia, dorsal view.

**Figs. 156A-H.**
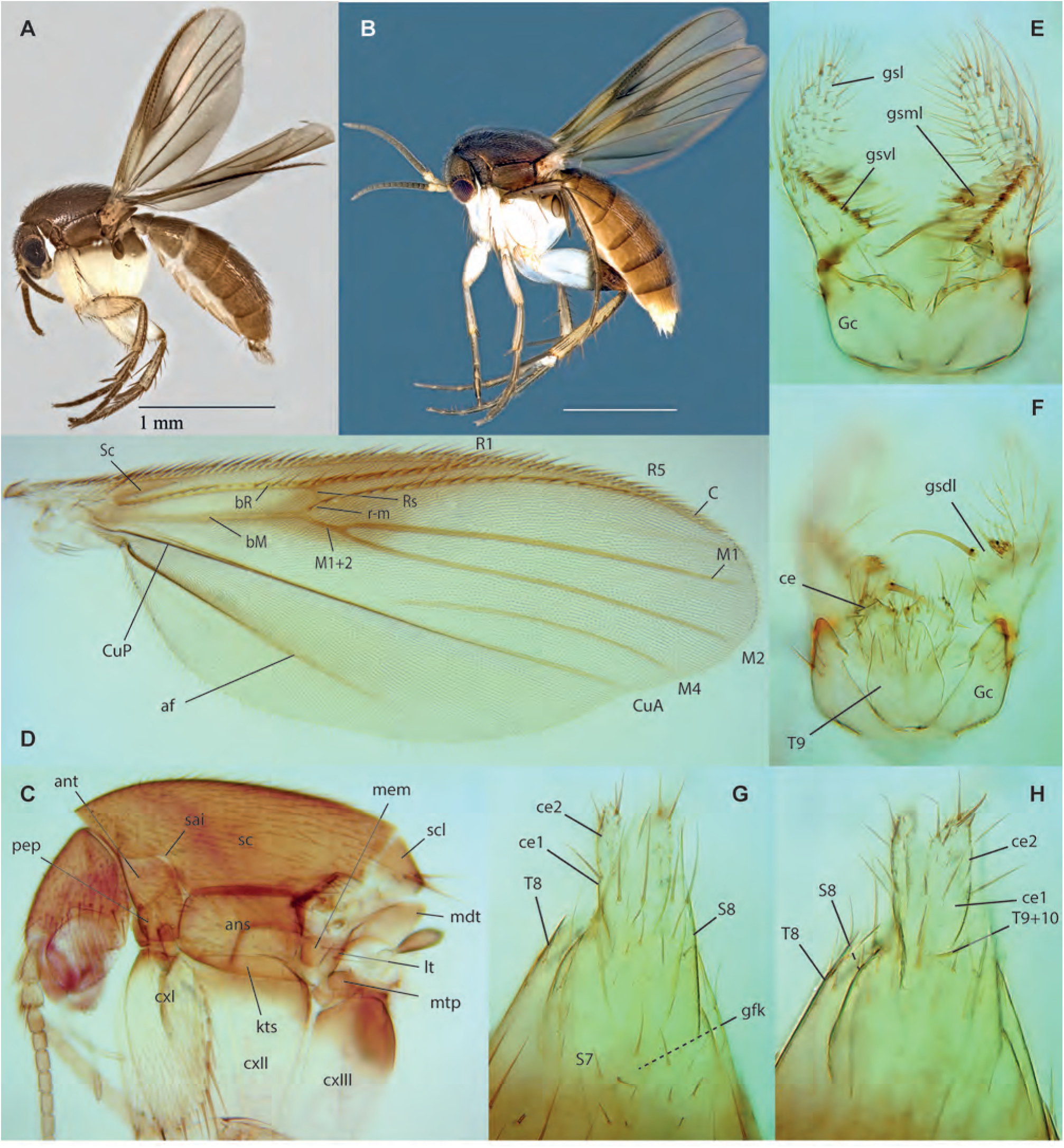
*Integricypta shirinae* Amorim & Oliveira, **sp. n. A.** Habitus, lateral view, male, paratype ZRCBDP0133375. **B.** Habitus, lateral view, female, paratype ZRCBDP0048427. **C.** Thorax, lateral view, male holotype. **D.** Wing, same. **E.** Male terminalia, ventral view, same. **F.** Male terminalia, dorsal view, same. **G.** Female terminalia, ventral view, paratype ZRCBDP0049345. **H.** Female terminalia, dorsal view, same.

**Figs. 157A-E.**
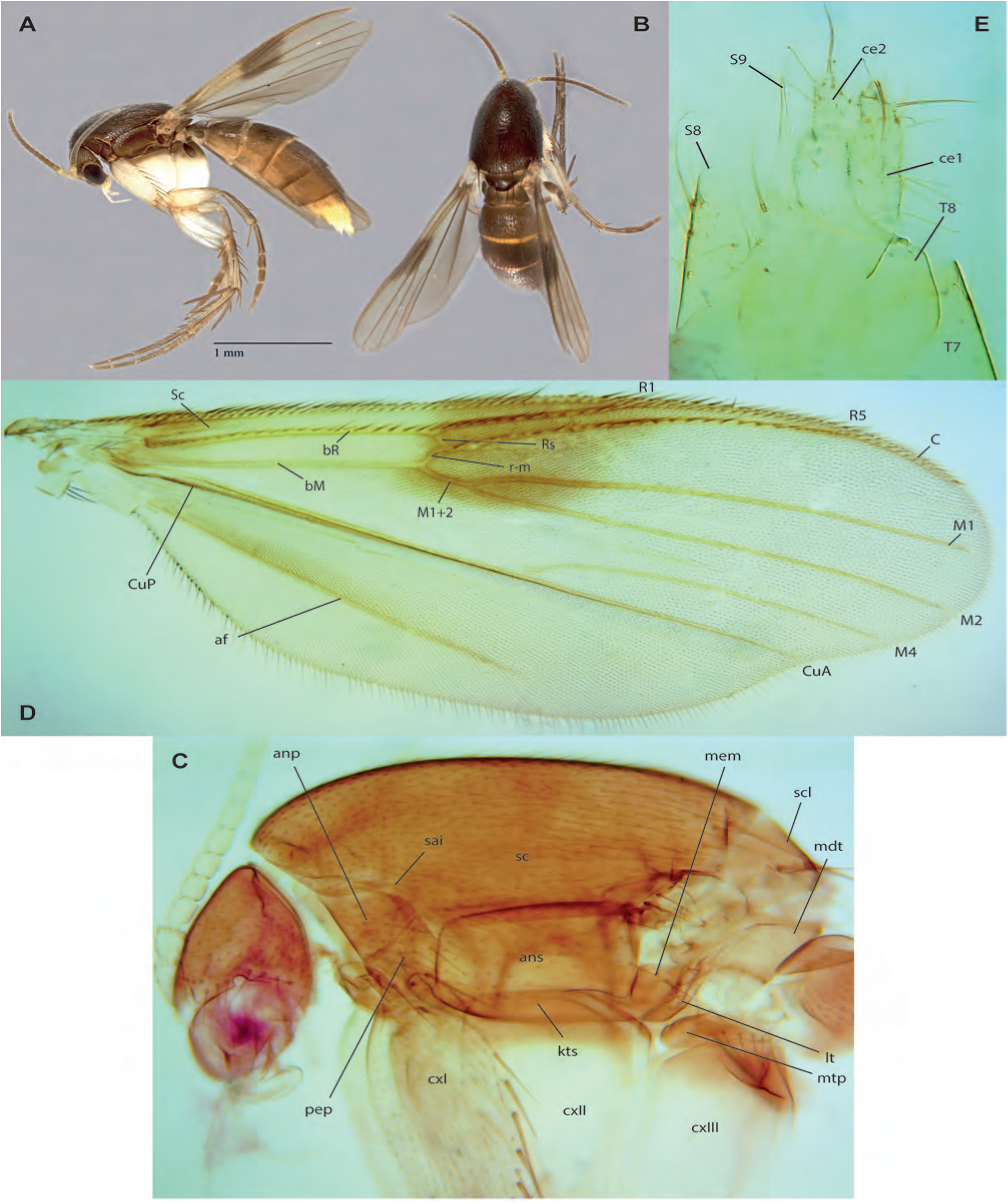
*Integricypta hoyuenhoeae* Amorim & Oliveira, **sp. n.**, female holotype, ZRCBDP0103977. **A.** Habitus, lateral view. **B.** Same, dorsal view. **C.** Head and thorax, lateral view. **D.** Wing. **E.** Terminalia, ventral view.

## References

1. Amaral, E. M., Oliveira, S. S. & Falaschi, R. L. (2022): An unknown world in the Neotropical region: a complete life cycle of a new species of *Monoclona* Mik, 1886 (Diptera: Mycetophilidae: Sciophilinae). – Zootaxa 5091: 107–130.

2. Amorim, D. S. (1993): A phylogenetic analysis of the basal groups of Bibionomorpha, with a critical reanalysis of the wing vein homology. – Revista Brasileira de Biologia 52: 379–399.

3. Amorim, D. S., Oliveira, S. S. & Henao–Sepulveda, A. C. (2018): A new species of *Eumanota* Edwards (Diptera: Mycetophilidae: Manotine) from Colombia: evidence for a pseudogondwanan pattern. – American Museum Novitates 3915: 1–19.

4. Amorim, D. S., Oliveira, S. S. & Balbi, M. I. P. A. (2008): *Azana atlantica*, n. sp., with reduced mouthparts and two ocelli: first record of Azana for the Neotropical region (Diptera: Mycetophilidae: Sciophilinae). – Zootaxa 1789: 57–65.

5. Amorim, D. S., Oliveira, S. S. & Balbi, M. I. P. A. (2008b): First Neotropical species of genus *Azana* (Diptera: Mycetophilidae: Sciophilinae). – Zootaxa (1937): 67–68.

6. Amorim, D. S. & Rindal, E. (2007): A phylogenetic study of the Mycetophiliformia, with creation of the subfamilies Heterotrichinae, Ohakuneinae, and Chiletrichinae for the Rangomaramidae (Diptera, Bibionomorpha). – Zootaxa 1535: 1–92.

7. Amorim, D. S. & Tozoni, S. H. S. (1995): Phylogenetic and biogeographic analysis of the Anisopodoidea (Diptera, Bibionomorpha), with an area cladogram for intercontinental relationships. – Revista Brasileira de Entomologia 38: 517–543.

8. Amorim, D. S. & Yeates, D.K. (2006): Pesky Gnats: Ridding Dipteran Classification of the “Nematocera”. – Studia Dipterologica 13, 3–9.

9. Amorim, D. S., Brown, B. V., Boscolo, D., Ale-Rocha, R., Alvarez-Garcia, D. M., Balbi, M. I. P. A., Barbosa, A. M., Capellari, R. S., Carvalho, C. J. B. de, Couri, M. S., Perez-Dios, R. V., Fachin, D. A., Ferro, G. B., Flores, H. F., Frare, L. M., Gudin, F. M., Hauser, M., Lamas, C. J. E., Lindsay, K., Marinho, M. A. T., Marques, D. W. A., Marshall, S. A., Melo-Patiu, C., Menezes, M. A., Morales, M. N., Nihei, S. S., Oliveira, S. S., Pirani, G., Ribeiro, G. C., Riccardi, P. R., Santis, M. D., Santos, D., Santos, J. R. dos, Silva, V. C. & Rafael, J. A. (2022): Vertical stratification of Diptera abundance and species richness in an Amazonian tropical forest. – Scientific Reports (2022) **12**:1734.

10. Ang, Y., Wong, L.J. & Meier, R. (2013): Using seemingly unnecessary illustrations to improve the diagnostic usefulness of descriptions in taxonomy–a case study on *Perochaeta orientalis* (Diptera, Sepsidae). – ZooKeys (355): 9–27.

11. Bechev, D. (1995): *Allactoneura papuensis* spec. nov. from New Guinea (Insecta: Diptera: Mycetophilidae). – Reichenbachia 31: 81–82.

12. Bechev, D. (1999): The zoogeographic classification of the Palaearctic genera of fungus gnats (Diptera: Sciaroidea, excluding Sciaridae). – Studia dipterologica 6: 321–326.

13. Bechev, D. & Kazandzhieva, S. (2020): A new species of Stenophragma Skuse from Borneo (Diptera: Mycetophilidae: Sciophilinae). – Zootaxa 4819(1): 195–200.

14. Bickel, D. (2009): Why Hilara is not amusing: the problem of open-ended taxa and the limits of taxonomic knowledge. – In: Diptera diversity: status, challenges, and tools. Pape, T., Bickel, D.J.; Meier, R. (eds.): Diptera diversity: status, challenges and tools, pp. 279–301; Leiden (Brill).

15. Blagoderov, V. A. & Grimaldi, D. A. (2004): Fossil Sciaroidea (Diptera) in Cretaceous ambers, exclusive of Cecidomyiidae, Sciaridae, and Keroplatidae. – American Museum Novitates 3433: 1–76.

16. Borkent, C. J. & Wheeler, T. A. (2012): Systematics and Phylogeny of *Leptomorphus* Curtis (Diptera: Mycetophilidae). – Zootaxa 3549: 1–117.

17. Borkent, C. J. & Wheeler, T. A. (2013): Phylogeny of the tribe Sciophilini (Diptera: Mycetophilidae: Sciophilinae). – Systematic Entomology 38: 407–427.

18. Borkent. A., Brown, B. V., Adler, P. H., Amorim, D. S., Barber, K., Bickel, D., Boucher, S., Brooks, S. E., Burger, J., Burington, Z. L., Capellari, R. S., Costa, D. N. R., Cumming, J. M., Curler, G., Dick, C. W., Epler, J. H., Fisher, E., Gaimari, S. D., Gelhaus, J., Grimaldi, D. A., Hash, J., Hauser, M., Hippa, H., Ibáñez-Bernal, S., Jaschhof, M., Kameneva, E. P., Kerr, P. H., Korneyev, V., Korytkowski, C. A., Kung, G. -A., Kvifte, G. M., Lonsdale, O., Marshall, S. A., Mathis, W., Michelsen, V., Naglis, S., Norrbom, A. A., Paiero, S., Pape, T., Pereira-Colavite, A., Pollet, M., Rochefort, S., Rung, A., Runyon, J. B., Savage, J., Silva, V. C., Sinclair, B. J., Skevington, J. H., Stireman, J. O. III, Swann, J., Vilkamaa, P., Wheeler, T., Withworth, T., Wong, M., Wood, D. M., Woodley, N., Yau, T., Zavortink, T. J., Zumbado, M. A. 2018. Remarkable fly (Diptera) diversity in a patch of Costa Rican cloud forest: why inventory is a vital science. – Zootaxa 4402(1): 53–90.

19. Brook, B. W., Sodhi, N. S. and Ng, P. K. (2003): Catastrophic extinctions follow deforestation in Singapore. – Nature 424(6947): 420–423.

20. Brown, B. V., Borkent, A., Adler, P. H., De Souza, A. D., Barber, K., Bickel, D., Boucher, S., Brooks, S. E., Burger, J., Burington, Z. L., Capellari, R. S., Costa, D. N. R., Cumming, J. M., Curler, G., Dick, C. W., Epler, J. H., Fisher, E., Gaimari, S. D., Gelhaus, J., Grimaldi, D. A., Hash, J., Hauser, M., Hippa, H., Ibáñez-Bernal, S., Jaschhof, M., Kameneva, E. P., Kerr, P. H., Korneyev, V., Korytkowski, C. A., Kung, G. -A., Kvifte, G. M., Lonsdale, O., Marshall, S. A., Mathis, W., Michelsen, V., Naglis, S., Norrbom, A. A., Paiero, S., Pape, T., Pereira-Colavite, A., Pollet, M., Rochefort, S., Rung, A., Runyon, J. B., Savage, J., Silva, V. C., Sinclair, B. J., Skevington, J. H., Stireman III, J. O., Swann, J., Thopson, F. C., Vilkamaa, P., Wheeler, T., Withworth, T., Wong, M., Wood, D. M., Woodley, N., Yau, T., Zavortink, T. J., Zumbado, M. A. (2018): Comprehensive inventory of true flies (Diptera) at a tropical site. – Communications Biology 1(21): 8.

21. Brunetti, E. (1912) The fauna of British India, including Ceylon and Burma. Diptera Nematocera (excluding Chironomidae and Culicidae), 581 pp., XII pls.; London (Taylor and Francis).

22. Burdíková, N., Kjærandsen, J., Lindemann, J. P., Kaspřák, D., Tóthová, A. & Ševčík, J. (2019): Molecular phylogeny of the Paleogene fungus gnat tribe Exechiini (Diptera: Mycetophilidae) revisited: Monophyly of genera established and rapid radiation confirmed. – Journal of Zoological Systematics and Evolutionary Research 57(4): 806–821.

23. Chandler, P. J., Bechev, D. N. & Caspers, N. (2005): The Fungus Gnats (Diptera: Bolitophilidae, Diadocidiidae, Ditomyiidae, Keroplatidae and Mycetophilidae) of Greece, its islands and Cyprus. – Studia Dipterologica 12: 255–314.

24. Coher, E.I. (1959): A synopsis of american Mycomyiini with description of new species (Diptera: Mycetophilidae). – Entomologica Americana, n.s. 37: 1–155.

25. Colless, D. H. (1966): Insects of Micronesia. Diptera: Mycetophilidae. – Insects of Micronesia 12(6): 637– 666.

26. Colless, D. H. & Liepa, Z. (1973): Superfamily Mycetophiloidea. Family Mycetophilidae (Fungivoridae). – In: Delfinado, M. D. & Hardy, D. E. (eds.): A catalogue of the Diptera of the Oriental Region. Volume I. Suborder Nematocera, pp. 442–463; Honolulu (University of Hawaii Press).

27. Coquillett, D. W. (1910): The type-species of the North American genera of Diptera. – Proceedings of the United States National Museum 37: 499–647.

28. Cumming, J. W. & Wood, D. M. (2017): Adult morphology and terminology. – In: Kirk–Spriggs, A. H. & Sinclair, B. J. (eds.): Manual of Afrotropical Diptera, vol. 1. Introductory chapters and keys to Diptera families, pp. 89–133; Suricata 4. Pretoria (South African National Biodiversity Institute).

29. Curtis, J. (1831): Plate 365, Leptomorphus walkeri. – In: Curtis, J. British entomology; Being illustrations and descriptions of the genera of insects found in Great Britain and Ireland. Vol. 8, 439 pp.; London (Privately published).

30. Curtis, J. (1837): British Entomology: Being illustrations and descriptions of the genera of insects found in Great Britain and Ireland 13, pls. 626–673; London (Privately published).

31. De Geer, D. (1776): Memoires pour servir a l’historie des Insects. Volume 6, 361pp., 22 pl.; Stockholm (Hesselberg)

32. De Meijere, J. C. H. (1924): Studien über südostasiatische Dipteren XVI. – Tijdschrift Voor Entomologie 67: 197–224.

33. De Meijere, J. C. H. (1907): Studien uber sudostasiatische Dipteren. I. – Tijdschrift voor entomologie 50: 196–264, pls. 5–6.

34. Duff, A., Lott, D. A. & Buckland, P. C. (2012): Checklist of beetles of the British Isles, 174 pp.; Iver (Pemberley Books).

35. Edwards, F. W. (1925): Diptera Nematocera from the Dutch East Indies. – Treubia 6: 154–172.

36. Edwards, F. W. (1926): XIV – Diptera Nematocera from them mountains of Borneo. – Sarawale Museum Journal 3: 243–278.

37. Edwards, F. W. (1927): Diptera Nematocera from the Dutch East Indies. – Treubia 9: 352–370.

38. Edwards, F. W. (1928): Diptera Nematocera from the Federated Malay States Museum. – Journal of the Federated Malay States Museuns 14: 1–139.

39. Edwards, F. W. (1929): Philipinne Nematocerous Diptera III. – Notulae Entomologicae 9: 70–81.

40. Edwards, F. W. (1931): Fauna Sumatrensis. Mycetophilidae, Diptera. – Beijdrage 69: 262–278.

41. Edwards, F. W. (1933): Diptera Nematocera from Mount Kinabalu. – Journal of the Federated Malay States Museuns 17: 223–296.

42. Edwards, F. W. (1935): Two new *Delopsis* collected by Dr. Leefmans in Java. Diptera, Mycetophilidae. – Tijdschrift Voor Entomologie 78: 186–187.

43. Edwards, F. W. (1940): Neotropical Neoempheria (Diptera, Mycetophilidae). – Novitates Zoologicae 42: 107–129.

44. Enderlein, G. (1910): Diptera, Mycetophilidae [of the Seychelles]. Proceedings of the Linnean Society of London, (2) 14: 59–81.

45. Enderlein, G. (1911): Neue Gattungen und Arten aussereuropaischer Fliegen. – Stettiner Entomologische Zeitung 72: 135–209.

46. Engel, M. S., Ceríaco, L. M., Daniel, G. M., Dellapé, P. M., Löbl, I., Marinov, M., Reis, R. E., Young, M. T., Dubois, A., Agarwal, I. & Lehmann A, P. (2021): The taxonomic impediment: a shortage of taxonomists, not the lack of technical approaches. – Zoological Journal of the Linnean Society, 193(2): 381–387.

47. Evenhuis, N. L. & T. Pape. (2022): Systema Dipterorum. http://www.diptera.org/ (accessed January 16, 2023).

48. Fitzgerald, S. J. (2017): *Phthinia* Winnertz (Diptera: Mycetophilidae): new species and records from the Neotropical and Oriental regions. Zootaxa 4231: 107– 113.

49. Freeman, P. (1951): Diptera of Patagonia and South Chile based mainly on material in the British Museum (Natural History). Part III–Mycetophilidae, 133 pp + 49 pls.; London (British Museum (Natural History).

50. Gammelmo, O. (2004): A review of the Afrotropical genus *Mycomyiella* Matile, 1973 (Diptera, Mycetophilidae, Mycomyini), with the description of six new species. – Zootaxa 625: 1-30.

51. Grootaert, P. (2018): Revision of the genus *Thinophilus* Wahlberg (Diptera: Dolichopodidae) from Singapore and adjacent regions: A long term study with a prudent reconciliation of a genetic to a classic morphological approach. – Raffles Bulletin of Zoology 66: 413–473.

52. Grootaert, P. & Puniamoorthy, J. (2014): Revision of Ngirhaphium (Insecta: Diptera: Dolichopodidae), with the description of two new species from Singapore’s mangroves. – Raffles Bulletin of Zoology 62:146–160.

53. Hartop, E., Srivathsan, A., Ronquist, F. & Meier, R. (2022): Towards Large-scale Integrative Taxonomy (LIT): resolving the data conundrum for dark taxa. – Systematic Biology 71(6): 1404–1422.

54. Hausmann A., Krogmann L., Peters R., Rduch V., Schmidt, S. (2020): GBOL III: Dark Taxa. Available from https://ibol.org/barcodebulletin/research/gbol-iii-dark-taxa/.

55. Hebert, P. D., Ratnasingham, S., Zakharov, E. V., Telfer, A. C., Levesque-Beaudin, V., Milton, M. A., Pedersen, S., Jannetta, P., de Waard, J. R. (2016): Counting animal species with DNA barcodes: Canadian insects. – Philosophical Transactions of the Royal Society of London, Series B, Biological Sciences 371(1702): 20150333.

56. Hippa, H. (2006): Diversity of *Manota* Williston (Diptera: Mycetophilidae) in a Malaysian rainforest: description of twenty-seven new sympatric species. – Zootaxa 1161: 1–49.

57. Hippa, H. (2007): The genus *Manota* Williston (Diptera: Mycetophilidae) in Melanesia and Oceania. – Zootaxa 1502: 1–44.

58. Hippa, H. (2008): New species and new records of Manota Williston (Diptera, Mycetophilidae) from the Oriental region. – Zootaxa 1723: 1–41.

59. Hippa, H. (2009): New species and new records of *Manota* Williston (Diptera, Mycetophilidae) from Thailand. – Zootaxa 2017: 1–33.

60. Hippa, H. (2011): New species and new records of *Manota* Williston (Diptera, Mycetophilidae) from Thailand, with a key to the Oriental and Palaearctic species. – Zootaxa 2763: 39–60.

61. Hippa, H., Jaschhof, M. & P. Vilkamaa. (2005): Phylogeny of the Manotinae, with a review of *Eumanota* Edwards, *Paramanota* Tuomikoski and *Promanota* Tuomikoski (Diptera, Mycetophilidae). – Studia Dipterologica 11: 405–428.

62. Hippa, H. & Kurina, O. (2018): Four new species and new records of *Manota* (Diptera: Mycetophilidae) from Sulawesi, Indonesia. – Acta Entomologica Musei Nationalis Pragae 58: 249–256.

63. Hippa, H., Kurina, O. & Sääksjärvi, I. (2017): The genus *Manota* Williston (Diptera: Mycetophilidae) in Peruvian Amazonia, with description of sixteen new species. – Zootaxa 4236: 1–40.

64. Hippa, H. & Ševčík, J. (2010): Notes on Oriental and Australasian Manotinae (Diptera, Mycetophilidae), with the description of thirteen new species. – Zootaxa 2333: 1–25.

65. Hippa, H. & Ševčík, J. (2013): Five new species and a new record of *Manota* (Diptera: Mycetophilidae) from Sulawesi. – Acta Entomologica Musei Nationalis Pragae 53: 763–775.

66. Hippa, H. & Saigusa, T. (2016): Notes on Oriental and East Palaearctic *Manota* Williston (Diptera, Mycetophilidae), with the description of seven new species. – Zootaxa 4084: 377–390.

67. Hippa, H., Søli, G. & Kurina, O. (2019): New data on the genus *Manota* Williston (Diptera: Mycetophilidae) from Africa, with an updated key to the species. – Zootaxa 4652: 401–441.

68. Høye, T.T., Ärje, J., Bjerge, J.K., Hansen, O.L., Iosifidis, A., Leese, F., Mann, H.M., Meissner, K., Melvad, C., Raitoharju, J. (2021): Deep learning and computer vision will transform entomology. – Proceedings of the National Academy of Sciences USA 118(2), e2002545117. Doi: 10.1073/pnas.2002545117

69. Jaschhof, M. & Hippa, H. (2005): The genus *Manota* in Costa Rica (Diptera: Mycetophilidae). – Zootaxa 1011: 1–54.

70. Jaschhof, M. & Kallweit, U. (2004): The genus *Aphrastomyia* Coher and Lane, 1949 in Costa Rica (Insecta: Diptera: Mycetophilidae). – Faunistische Abhandlungen 25: 107–123.

71. Jaschhof, M., Jaschhof, C., Rulik, B. & Kjærandsen, J. (2011): New records of *Manota* Williston (Diptera: Mycetophilidae) in Europe and North America, including a redescription of *Manota unifurcata* Lundström and pointers towards the interrelationships among Palaearctic species. – Studia dipterologica 17: 55–66.

72. Johannsen, O. A. (1909): Diptera. Fam. Mycetophilidae. – In: Wytsman, P. (ed.). Genera Insectorum 93, 141 pp. + 8 pls; Bruxelles.

73. Jürgenstein, S., Kurina, O., & Põldmaa, K. (2015): The *Mycetophila ruficollis* Meigen (Diptera, Mycetophilidae) group in Europe: elucidating species delimitation with COI and ITS2 sequence data. – ZooKeys 508, 15.

74. Kallweit, U. (1998): Notes on the genus Metanepsia Edwards and its relatives from East Asia (Insecta: Diptera: Mycetophilidae). – Reichenbachia Staatliches Museum für Tierkund Dresden 53: 341–353.

75. Kaspřák, D., Borkent, C. & Wahab, R.H.A. (2017): *Leptomorphus sevciki* sp. Nov., a remarkable new wasp-mimicking fungus gnat from Brunei (Diptera: Mycetophilidae). – Acta Entomologica Musei Nationalis Pragae 57(1): 195–203.

76. Kaspřák, D., Kerr, P., Sýkora, V., Tóthová, A. & Ševčík, J. (2019): Molecular phylogeny of the fungus gnat subfamilies Gnoristinae and Mycomyinae, and their position within Mycetophilidae (Diptera). – Systematic Entomology 44: 128–138.

77. Kerr, P. H. (2010): New *Azana* species from Western North America (Diptera: Mycetophilidae). – Zootaxa 2397, 1–14.

78. Kerr, P. H. (2014): The *Megophthalmidia* (Diptera, Mycetophilidae) of North America including eight new species. – ZooKeys 386: 29–83.

79. Kjærandsen, J. (2005): A review of fungus gnats in the tribe Exechiini (Diptera, Mycetophilidae) from the J. W. Zetterstedt collection at the Museum of Zoology in Lund, Sweden. – Zootaxa 856: 1–35.

80. Kjærandsen, J. (2007): Two new species of *Allodia* subgenus *Brachycampta* Winnertz from Norway and Sweden (Diptera: Mycetophilidae). – Entomologica Fennica 18: 17–23

81. Kjaerandsen, J. (2009): The genus *Pseudexechia* Tuomikoski re-characterized, with a review of European species (Diptera: Mycetophilidae). – Zootaxa 2056(1): 1–45.

82. Kurina, O. (2020): Three new species and new records of *Clastobasis* Skuse (Diptera: Mycetophilidae) from Japan and the Kuril Islands. – Zootaxa 4810(3): 589– 600.

83. Kurina, O. (1992): A new species of the genus *Mycetophila* Meigen (Diptera, Mycetophilidae) found in Estonia. – Proceedings of the Estonian Academy of Sciences, Biologia 41(3): 127–130.

84. Kurina, O. & Hippa, H. (2014): The genus *Manota* Williston (Diptera: Mycetophilidae) in the Congo basin with description of five new species. – Zootaxa 3827: 214–230.

85. Kurina, O. & Hippa, H. (2015): A review of the South Pacific *Manota* Williston (Diptera, Mycetophilidae), with description of thirteen new species. – Zootaxa 4020: 257–288.

86. Kurina, O., Hippa, H. & Amorim, D. S. (2018): A contribution to the systematics of the genus *Manota* Williston (Diptera: Mycetophilidae) in Brazil. – Zootaxa 4472: 1–59.

87. Kurina, O., Hippa, H. & Amorim, D. S. (2019): Notes on *Manota* Williston (Diptera: Mycetophilidae) from Australia and Papua New Guinea, with description of two new species. – Zootaxa 4555: 385–395.

88. Kurina, O. & Hippa, H. (2021): Additions to the knowledge of *Manota* Williston (Diptera: Mycetophilidae) from the Neotropical region, with description of four new species. – Zootaxa 4938: 85– 100.

89. Lane, J. (1954) Revision of the genus *Epicypta* Winnertz, 1863 in the neotropical region. – Revista Brasileira de Entomologia 2: 111-138.

90. Magnussen, T. (2020): – Integrative taxonomy and systematics of *Allodia* Winnertz (Diptera, Mycetophilidae). PhD Thesis, 208 pp.; Oslo (University of Oslo).

91. Magnussen, T., Kjærandsen J., Johnsen A., Søli G. E. E. (2018): Six new species of Afrotropical Allodia (Diptera: Mycetophilidae): DNA barcodes indicate recent diversification with a single origin. – Zootaxa 4407(3): 301–320.

92. Magnussen, T., Søli, G. E. E. & Kjærandsen, J. (2019): *Allodia* Winnertz from the Himalayas, with nine species new to science (Diptera, Mycetophilidae). – ZooKeys 820: 119–138.

93. Magnussen, T., Johnsen, A., Kjærandsen, J., Struck, T. H., & Søli, G. E. (2022): Molecular phylogeny of *Allodia* (Diptera: Mycetophilidae) constructed using genome skimming. – Systematic Entomology 47(1), 267–281.

94. Matile, L. (1971a): Une nouvelle tribu de Mycetophilidae: les Metanepsiini (Dipt.). – Bulletin de la Société entomologique de France 76: 91–97.

95. Matile. L. (1971b): Un *Metanepsia* nouveau du Kenya (Dipt. Mycetophilidae). – Bulletin de la Société entomologique de France 76: 271–272.

96. Matile, L. (1973a): Diptères Mycetophilidae de Fernando-Póo. – Bulletin du Muséum National d’Historie Naturelle 111: 189–213.

97. Matile, L. (1973b): Diptéres Mycetophilidae de l’Afrique Orientale. – Stuttgarter Beiträge zur Naturkunde 250: 1–6.

98. Matile, L. (1974a): Diptères Mycetophilidae du Cameroun et de République Centrafricaine. 3. Sciophilinae, genre *Parempheriella*. – Bulletin de l’Institut Fondamental d’Afrique Noire 35: 609–664.

99. Matile, L. (1974b): Note sur les genres *Aspidionia* et *Platyprosthiogyne* en Région Éthiopienne (Dipt. Mycetophilidae). – Annales de la Société Entomologique de France, 10 (3): 389–392.

100. Matile, L. (1976a): Diptères Mycetophilidae du Cameroun et de République Centrafricaine. IV. Sciophilinae, genre *Syndocosia*. – Bulletin de l’Institut Fondamental d’Afrique Noire 37: 687–701.

101. Matile, L. (1976b): Un genre nouveau de Mycomyini à nervation alaire réduite; diagnose preliminaire. Diptera, Mycetophilidae, Sciophilinae. – Bulletin de la Société entomologique de France 81: 139–140.

102. Matile, L. (1979a): Diptères Mycetophilidae de ĺArchipel des Comores. – Mémoires Museum National d’Histoire naturelle 109: 247–306.

103. Matile, L. (1979b): Um nouveau genre Afrotropical de Mycomyini [Diptera Mycetophilidae]. – Revue française d’Entomologie 1 (3): 106–116.

104. Matile, L. (1980a): Nouvelles Données sur les *Metanepsia* Afrotropicaux (Diptera, Mycetophilidae). – Revue française d’Entomologie 2(3): 119–122.

105. Matile, L. (1980b): Superfamily Mycetophiloidea. 15. Family Mycetophilidae. – In: Crosskey, R. W. (ed.): Catalogue of the Diptera of the Afrotropical Region, pp. 216–230;. London (British Museum Natural History)

106. Matile, L. (1989): 10. Family Mycetophilidae. – In: Evenhuis, N. L. (ed.): Catalog of the Diptera of the Australasian and Oceanic Regions, pp. 135–145; Honolulu (Bishop Museum Press).

107. Matile, L. (1999): Première citation du genre *Parempheriella* Matile dans la Région Paléarctique (Diptera, Mycetophilidae. – Revue française d’Entomologie 21(3):115–118.

108. Meier, R. (2017): Citation of taxonomic publications: the why, when, what and what not. – Systematic Entomology 42: 301–304.

109. Meier, R., Lawniczak, M.K. Srivathsan, A. (2024). Illuminating entomological dark matter with DNA barcodes in an era of insect decline, deep learning, and genomics. Annual Review of Entomology 70 10.1146/annurev-ento-040124-014001.

110. Meier, R., Srivathsan, A., Yeo, D., Oliveira, S. S., Balbi, M. I. P. A., Ang, Y., Amorim, D. S. (2023): “Dark taxonomy”: a new protocol for overcoming the taxonomic impediments for dark taxa and broadening the taxon base for biodiversity assessment. – bioRxiv 2023.08.31.555664.

111. Meigen, J. W. (1803): Versuch einer neuen Gattungseintheilung der europaischen zweiflugeligen Insekten. – Magazin fur Insektenkunde Herausgegeben von Karl Illiger 2: 259–281.

112. Meigen, J. W. (1804): Klassifikazion und Beschreibung der europäischen zweiflügeligen Insecten (Dipteren Linn.) **1**(1): xxvii + 142 pp., 8 pls.; (2), vi + pp. 153–314, pls. 9–15; Braunschweig.

113. Meigen, J. W. (1818): Systematische Beschreibung der bekannten europaischen zweifl ugeligen Insekten 1: xxxvi + 332 + [1] pp., 11 pls.; Aachen.

114. Meigen, J.W. (1830): Systematische Beschreibung der bekannten europäischen zweiflügeligen Insekten **6**: xi + 401 pp.; Hamm (Schulz).

115. Mik, J. (1886): Dipterologische Miscellen. II. – Wiener Entomologische Zeitung 5: 276–279.

116. Økland, B. (1994): Mycetophilidae (Diptera), an insect group vulnerable to forestry? A comparison of clearcut, managed and seminatural spruce forests in southern Norway. – Biodiversity and Conservation 3: 68–85.

117. Økland, B. (1996): Unlogged forests: Important sites for preserving the diversity of mycetophilids (Diptera: Sciaroidea). – Biological Conservation 76: 297–310

118. Økland, B., Götmark, F., Nordén, B., Franc, N., Kurina, O. & Polevoi, A. (2005): Regional diversity of mycetophilids (Diptera: Sciaroidea) in Scandinavian oak-dominated forests. – Biological Conservation 121: 9–20.

119. Oliveira, S. S. (2015): On Afrotropical *Mohelia* Matile (Diptera, Mycetophilidae): new species and phylogenetic comments. – Zootaxa 3947: 251–263.

120. Oliveira, S. S. & Amorim. D. S. (2021): Phylogeny, classification, Mesozoic fossils and biogeography of the Leiinae (Diptera: Mycetophilidae). – Bulletin of the American Museum of Natural History 446: 1–108.

121. Oliveira, S. S. & Amorim, D. S. (2014): Catalogue of the Neotropical Diptera. Mycetophilidae. – Neotropical Diptera 25: 1–87.

122. Osten-Sacken, C. R. (1878): Catalogue of the described Diptera of North America (2nd Edition). – Smithsonian Miscellaneous Collections 16: 1–276.

123. Papp, L. (2004): Seven new species of Manotinae (Diptera: Mycetophilidae) from Asia and Papua New Guinea. – Acta Zoologica Academiae Scientiarum Hungaricae 50: 227–244.

124. Papp, L. & Ševčík, J. (2011): Eight new Oriental and Australasian species of *Leptomorphus* (Diptera: Mycetophilidae). – Acta Zoologica Academiae Scientiarum Hungaricae 57: 139–159.

125. Ratnasingham, S. & Hebert, P.D. (2007) Bold: The Barcode of Life Data System (http://www.barcodinglife.org). Molecular Ecology Notes 7(3): 355–364.

126. Rheindt, F.E., Bouchard, P., Pyle, R.L., Welter-Schultes, F., Aescht, E., Ahyong, S.T., Ballerio, A., Bourgoin, T., Ceríaco, L.M., Dmitriev, D. & Evenhuis, N. (2023): Tightening the requirements for species diagnoses would help integrate DNA-based descriptions in taxonomic practice. – PLoS Biology 21(8): p.e3002251.

127. Rindal, E. & Søli, G. E. E. (2006): Phylogeny of the subfamily Mycetophilinae (Diptera: Mycetophilidae). – Zootaxa 1302: 43–59.

128. Rindal, E., Søli, G. E. E. & Bachmann, L. (2009): On the systematics of the fungus gnat subfamily Mycetophilinae (Diptera): a combined morphological and molecular approach. – Journal of Zoological Systematics and Evolutionary Research 47: 227–233.

129. Rindal, E., Søli, G. E. E., Kjærandsen, J. & Bachmann, L. (2007): Molecular phylogeny of the fungus gnat tribe Exechiini (Mycetophilidae, Diptera). – Zoologica Scripta 36: 327–335.

130. Rondani, C. (1856): Dipterologiae Italicae Prodromus. Vol. 1: Genera Italica ordinis dipterorum ordinatim disposita et distincta et in familias et stirpes aggregata, 228 pp.; Parma.

131. Savage J., Borkent, A., Brodo, F., Cumming, J. M., Curler, G., Currie, D.C., deWaard, J.R., Gibson, J.F., Hauser, M., Laplante, L., Lonsdale, O., Marshall, S.A., O’Hara, J.E., Sinclair, B.J., Skevington, J.H. (2019): Diptera of Canada. – In: Langor, D. W. & Sheffield, C. S. (eds.): The Biota of Canada. A Biodiversity Assessment. Part 1: The Terrestrial Arthropods. – ZooKeys 819: 397-450.

132. Senior-White, R. A. (1922): New Ceylon Diptera (Part II). – Spolia Zeylanica 12: 195–206.

133. Ševčík, J. (2001): First record of *Cordyla* Meigen and Monoclona Mik from the Oriental Region (Diptera: Mycetophilidae). – Beiträge zur Entomologie 51(1): 155–160.

134. Ševčík, J., Bechev, D. & Hippa, H. (2011): New Oriental species of Gnoristinae with pectinate antennae (Diptera: Mycetophilidae). – Acta Entomologica Musei Nationalis Pragae 51(2): 687–696.

135. Ševčík, J. & Hippa, H. (2010): New species of *Chalastonepsia* and *Pectinepsia* gen. nov. (Diptera: Mycetophilidae) from the Oriental and Australasian Regions. – Acta Entomologica Musei Nationalis Pragae 50(2): 595–608.

136. Ševčík, J., Kaspřák, D. & Tóthová, A. (2013): Molecular phylogeny of fungus gnats (Diptera: Mycetophilidae) revisited: position of Manotinae, Metanepsiini, and other enigmatic taxa as inferred from multigene analysis. – Systematic Entomology 38: 654–660.

137. Ševčík, J. & Kjaerandsen, J. (2012): *Brachyradia*, a new genus of the tribe Exechiini (Diptera: Mycetophilidae) from the Oriental and Australasian regions. – Raffles Bulletin of Zoology 60(1): 117–127.

138. Sivec, I. & Plassmann, E. (1982): Sechs neue Pilzmücken aus Sri Lanka (Diptera, Nematocera, Mycetophilidae). – Spixiana 5(1): 7–13.

139. Skuse, F. A. (1890): Diptera of Australia. Nematocera. Supplement II. – Proceedings of the Linnean Society of New South Wales (2)**5**: 595–640.

140. Søli, G. E. E. (1996): *Chalastonepsia orientalis* gen. n., **sp. n.**, a second genus in the tribe Metanepsiini (Diptera, Mycetophilidae). – Tijdschrift voor Entomologie 139: 79–83.

141. Søli, G. E. E. (1997): The adult morphology of Mycetophilidae (s. str.), with a tentative phylogeny of the family (Diptera, Sciaroidea). – Entomologica Scandinavica Supplement 50: 5–55.

142. Søli, G. E. E. (2002): New species of *Eumanota* Edwards, 1933 (Diptera: Mycetophilidae). – Annales de la Societé entomologique de France 38: 45–53.

143. Søli, G. E. E. (2017): 20. Mycetophilidae (Fungus Gnats). – In: Kirk-Springgs, A. H. & Sinclair, B. J. (eds.): Manual of Afrotropical Diptera. Volume 2. Nematocerous Diptera and lower Brachycera. Suricata 5, pp. 533–555; Pretoria (South African National Biodiversity Institute).

144. Søli, G. E. E., Vockeroth, J. R. & Matile, L. (2000): Families of Sciaroidea. – In: Papp L. & Darvas B. (eds.), Contributions to a Manual of Palaearctic Diptera Appendix, pp. 49–92; Budapest (Science Herald).

145. Srivathsan, A., Ang, Y., Heraty, J., Hwang, W. S., Wan, J. F. A., Kutty, S. N., Puniamoorthy, J., Yeo, D., Roslin, T. & Meier, R. (2023): Convergence of dominance and neglect in flying insect diversity. – Nature Ecology and Evolution 7: 1012–1021.

146. Staeger, C. (1840): Systematisk Fortegnelse over de i Danmark hidtil funde Diptera. Tipularia Fungicolae. – Naturhistorisk Tidsskrift 3: 228–288.

147. Sueyoshi, M. (2014): Taxonomy of fungus gnats allied to *Neoempheria ferruginea* (Brunetti, 1912) (Diptera: Mycetophilidae), with descriptions of 11 new species from Japan and adjacent areas. – Zootaxa 3790(1): 139–164.

148. Sueyoshi, M., Mukai, H., Kitajima, H. & Huang, J. (2019) *Allactoneura akasakana* Sasakawa, 2005 (Diptera, Mycetophilidae), occurred in cultivation facilities of shiitake mushroom. Bulletin of the Forestry and Forest Products Research Institute, 18, 319-324.

149. Theng, M., Jusoh, W. F., Jain, A., Huertas, B., Tan, D. J., Tan, H. Z., Kristensen, N. P., Meier, R. & Chisholm, R. A. (2020): A comprehensive assessment of diversity loss in a well-documented tropical insect fauna: Almost half of Singapore’s butterfly species extirpated in 160 years. – Biological Conservation 242: 108401.

150. Torres, A., Lee, L., Srivathsan, A., & Meier, R. (2024). UITOTO: User Interface for Optimal Molecular Diagnoses in High-Throughput Taxonomy (v1.0.0). Zenodo. 10.5281/zenodo.11236953

151. Tuomikoski, R. (1966): Generic taxonomy of the Exechiini (Dipt., Mycetophillidae). – Annales entomologici Fennici 32: 159–194.

152. Ulimah, F. A., Suheriyanto, D. & Ahmad, M. (2021): *Keanekaragaman Serangga Aerial di Perkebunan Jeruk Semi Organik dan Anorganik Dusun Kasin Desa Sepanjang Kecamatan Gondanglegi Kabupaten Malang*, 212 p.; Malang (Program Studi Biologi, Fakultas Sains dan Teknologi, Universitas Islam Negeri Maulana Malik Ibrahim). [In Malay, Diversity of Aerial Insects in Semi-Organic and Inorganic Citrus Plantations, Kasin Hamlet, Sepanjang Village, Gondanglegi District, Malang Regency].

153. Väisänen, R. (1984): A monograph of the genus *Mycomyia* Rondani in the Holarctic region (Diptera, Mycetophilidae). – Acta Zoologica Fennica 177: 1– 346.

154. Väisänen, R. (1986): The delimitation of the Gnoristinae: criteria for the classification of recent European genera (Diptera, Mycetophilidae). – Annales Zoologici Fennici 23: 197–206.

155. Väisänen, R. (1996): New *Mycomya* species from the Himalayas (Diptera, Mycetophilidae): 1. Subgenus *Mycomya* s. str. – Entomologica Fennica **7**(3), 99–132. Van der Wulp, F.M. (1876): Verslag van de dertigste zomervergadering der Nederlandsche Entomologische Vereeniging gehouden te Amsterdam op Zaturdag 24 Julij 1875. – Tijdschrift voor Entomologie **19**: i-liv, 2 figs.

156. Walker, F. (1848): List of the specimens of dipterous insects in the collection of the British Museum. Part 1, 229 pp.; London (British Museum).

157. Walker, F. (1856): Insecta Britannica, Diptera. Volume 3, xxiv + 352 p.; London (L. Reeve).

158. Wang, W. Y., Srivathsan, A., Foo, M., Yamane, S. K. & Meier, R. (2018): Sorting specimenDrich invertebrate samples with costDeffective NGS barcodes: Validating a reverse workflow for specimen processing. – Molecular ecology resources 18(3): 490–501.

159. Wiedemann, C.R.W. (1817): Neue Zweiflügler (Diptera Linn.) aus der Gegend um Kiel. – Zoologisches Magazin 1: 61–86.

160. Winnertz, J. (1846): Beschreibung einiger neuen Gattungen aus der Ordnung der Zweiflügler. – Stettiner Entolomologishe Zeitung 7: 11–20.

161. Winnertz, J. (1864): Beitrag zu einer Monographie der Pilzmücken (Mycetophilidae). – Verhandlungen der Zoologisch-Botanischen Gesellschaft in Wien 13[1863]: 637–964.

162. Wu, H., Zheng, L-Y. & Xu, H.-C. (2003). First record of Allodia Winnertz (Diptera, Mycetophilidae) in China with descriptions of seven new species. – Acta Zootaxonomica Sinica 28(2): 349–355.

163. Wu, H.-C., & Yang, C. (1996). A new genus and a new species of Mycomyiinae from Fujian Province, China (Diptera: Mycetophilidae). – Acta Entomologica Sinica 39: 86–89.

164. Wührl, L., Pylatiuk, C., Giersch, M., Lapp, F., von Rintelen, T., Balke, M. et al. (2022): DiversityScanner: robotic handling discovery of small invertebrates with machine learning methods. – Molecular Ecology Resources 22(4), 1626– 1638.

165. Xu, H.-C. & Wu, H. (2002): Two new species of the genus *Azana* from China (Diptera: Mycetophilidae). – Acta Zootaxonomica Sinica 27(3): 621–623.

166. Yeo, D., Puniamoorthy, J., Ngiam, R. W. J., & Meier, R. (2018): Towards holomorphology in entomology: rapid and costDeffective adult–larva matching using NGS barcodes. – Systematic entomology 43(4): 678– 691.

167. Yeo, D., Srivathsan, A. & Meier, R. (2020): Longer is not always better: Optimizing barcode length for large-scale species discovery and identification. – Systematic Biology 69(5): 999–1015.

168. Yeo, D., Srivathsan, A., Puniamoorthy, J. et al. (2021): Mangroves are an overlooked hotspot of insect diversity despite low plant diversity. – BMC Biology 19: 202.

169. Zaitzev, A. I. (1982): Composition and systematic position of the genus *Allactoneura* de Meijere (Diptera, Mycetophilidae). – Entomologicheskoye Obozreniye 60, 901–913.

170. Zaitzev, A. I. (2001): New species of fungus gnats from Russia and Italy (Diptera: Mycetophilidae). – Zoosystematica Rossica 9(2): 453–458.

171. Zamani, A., Vahtera, V., Sääksjärvi, I. E., & Scherz, M. D. (2021): The omission of critical data in the pursuit of ‘revolutionary’methods to accelerate the description of species. – Systematic Entomology 46(1): 1–4.

172. Zamani, A., Dal Pos, D., Fric, Z. F., Orfinger, A. B., Scherz, M. D., Bartoňová, A. S., & Gante, H. F. (2022): The future of zoological taxonomy is integrative, not minimalist. – Systematics and biodiversity 20(1): 1–14.

