## Supplementary material for "An integrative taxonomic treatment of the Mycetophilidae (Diptera: Bibionomorpha) from Singapore reveals 115 new species on 730km^2^": ALnID2.r

```
## Title: ALnID2
## Version: 0.1-1
## Date: 2024-05-14
## Author: Ambrosio Torres (Researcher [Ctr. Integr. Biodivers. Discov. - Museum für Naturkunde, Berlin, Germany)
## Maintainer: Ambrosio Torres <> <Ambrosio.TorresGalvis@mfn.berlin>
## Depends: R version (>= 4.2.2). Packages   seqinr (4.2-23); Biostrings (2.66.0); DECIPHER (2.30.0)
## Description: identifies the specimens of the unaligned sequences given in a fasta file, based on a pool of DMCs (e.g., those included in an Output-File from OpDMC).
## License: GPL (3)
## Usage: ALnID("OpDMC_output.csv", "DatasetAlignedTarget.fas", "DatasetUnalignedQuery.fas", mismatches = 1, against= "First", perfectMatch=5, misMatch=0, gapOpening = -14, gapExtension = -2, gapPower = -1, terminalGap = 0, OutName = "IdentificationOutput.csv", MissLogFile= "LogMissing.csv", AligName = "ResultingAlignment.fasta")

require("Biostrings")
require("seqinr")
require("DECIPHER")

ALnID2 <- function(DMC_Output, dataTarget, dataQuery, 
				  mismatches = 1,
				  against = c("First","All"),
				  perfectMatch=5, 
				  misMatch=0, 
				  gapOpening = -14,
				  gapExtension = -2,
				  gapPower = -1,
				  terminalGap = 0,
				  OutName = "IdentificationOutput.csv",
				  MissLogFile= "LogMissing.csv",
				  AligName = "ResultingAlignment.fasta"				  
				  ) { 
	
	datos <- read.fasta(dataQuery, whole.header = TRUE, forceDNAtolower = FALSE)
	nombres <- names(datos)
	raw_records <- read.fasta(dataTarget, whole.header = TRUE, forceDNAtolower = FALSE)
	datiles <- read.csv(DMC_Output, header = TRUE)
	
	datiles$DMC <- gsub(":", ",", datiles$DMC) 
	datiles$DMC <- gsub("'", "", datiles$DMC)
	datiles$DMC <- gsub(",", "", datiles$DMC) 
	datiles$DMC <- gsub("\\[", "", datiles$DMC)
	datiles$DMC <- gsub("\\]", "", datiles$DMC)
	datiles2 <- gsub('[ACGT]','',datiles$DMC); datiles2 <- gsub("  "," ", datiles2)
	datiles2 <- paste(datiles2, collapse = "")
	datiles2 <- strsplit(datiles2, " ")[[1]]
	datiles2 <- sort(unique(as.numeric(datiles2)))	
	
	cadena <- NULL
	identi <- NULL
	for (i in 1:length(raw_records)) {
		cuerda <- as.character(raw_records[[i]]); cuerda <- paste(cuerda, collapse= "")
		if (is.element(cuerda, cadena) == TRUE) {
			next
		} else {
			cadena <- c(cadena, cuerda)
			identi <- c(identi, i)
			nombrecito <- attributes(raw_records)$names[i]
		}
	}
	raw_records <- raw_records[identi]	
	
	FinalTable <- NULL
	options(width = 50)
	texto <- paste("Identifying the specimens", "of the", dataQuery, "file.")
	logMissing <- NULL
	liniers <- paste("Sequence","SpeciesDMC","N_Missing", sep=",")
	logMissing <- rbind(logMissing, liniers)
	for (i in 1:length(datos)) {
		extra <- nchar('||100%')
		width <- options()$width
		step <- round(i / length(datos) * (width - extra))
		text <- sprintf('|%s%s|% 3s%%', strrep('=', step),
						strrep(' ', width - step - extra), round(i / length(datos) * 100))
		cat(paste(texto,'\n',text))
		count <- 0
		resultado <- NULL
		
		query <- paste(as.vector(datos[[i]]), collapse = "")
		query <- toupper(query)
		query <- DNAString(gsub("-", "N", query))
		query2 <- as.character(complement(query))
		query3 <- as.character(reverseComplement(query))
		query4 <- as.character(rev(query))
		query <- as.character(query)
		queries <- c(query, query2, query3, query4)
		
		if (against == "First") {			
			patron <- raw_records[[1]]
			patron <- paste(patron, collapse= "")
			
			prueba <- AlignProfiles(DNAStringSet(patron), DNAStringSet(queries))
			indigo <- which.min(DistanceMatrix(prueba, verbose = F)[2:5,1])
			prueba2 <- prueba[1:(indigo+1)]
			
			resul2 <- compareStrings(as.character(prueba2[2]), as.character(prueba2[1]))
			if ( grepl("\\+", resul2) ) {
				paeliminar <- strsplit(resul2, split="")[[1]]
				funar <- NULL
				for (mas in 1:length(paeliminar)){
					if(paeliminar[mas] == "+") {
						funar <- c(funar, mas)
					}	
				}
				alineado <- strsplit( as.character(prueba2[2]), split="")[[1]]
				datos[[i]] <- alineado[-c(funar)]
			} else {
				datos[[i]] <- strsplit(as.character(prueba2[2]), split="")[[1]]
			}
			nomina <- names(raw_records)[1]
		}
		
		if (against == "All") {			
			FUN = function(x)
			paste(getSequence(x), collapse = "")
			raw_records2 <- as(vapply(raw_records, FUN, character(1)), "DNAStringSet")
			 
			prueba <- AlignProfiles(raw_records2, DNAStringSet(queries))
			indigo <- which.min(DistanceMatrix(prueba, verbose = F)[(length(prueba)-3):length(prueba),1])
			indigo2 <- which.min(DistanceMatrix(prueba, verbose = F)[1:length(raw_records2),(indigo+length(raw_records2))])
			prueba2 <- prueba[c(indigo2,(indigo+length(raw_records2)))]
		
			resul2 <- compareStrings(as.character(prueba2[2]), as.character(prueba2[1]))
			if ( grepl("\\+", resul2) ) {
				paeliminar <- strsplit(resul2, split="")[[1]]
				funar <- NULL
				for (mas in 1:length(paeliminar)){
					if(paeliminar[mas] == "+") {
						funar <- c(funar, mas)
					}	
				}
				alineado <- strsplit( as.character(prueba2[2]), split="")[[1]]
				datos[[i]] <- alineado[-c(funar)]
			} else {
				datos[[i]] <- strsplit(as.character(prueba2[2]), split="")[[1]]
			}
			nomina <- names(prueba2)[1]
		}
		
		simil <- 1 - DistanceMatrix(prueba2, verbose = F)[1,2]
		
		chequeo <- 0
		if (MissLogFile == "none" | MissLogFile == "None" | MissLogFile == "NONE") {
		} else {
			if (grepl(paste(datiles$Species, collapse='|'), attributes(datos)$names[i] )) {
				MissEntries <- which(as.character(datos[[i]]) == "-")
				if (length(MissEntries) > 0) {
					ImporMiss <- c(MissEntries, datiles2)[which(duplicated(c(MissEntries, datiles2)))]
					if (length(ImporMiss) > 0) {
						chequeo <- 1
					}		
				}
			}	
		}
		
		for (j in 1:dim(datiles)[1]) {
			DMC <- datiles$DMC[j]
			DMC <- strsplit(DMC, " ")[[1]]
			
			if (chequeo == 1) {
				#if (grepl(datiles$Species[j], attributes(datos)$names[i] )) {
					DMCtemp <- gsub('[ACGT]','', DMC); DMCtemp <- gsub("  "," ", DMCtemp)
					DMCtemp <- as.numeric(na.exclude(as.numeric(DMCtemp)))
					cuantos <- c(ImporMiss, DMCtemp)[which(duplicated(c(ImporMiss, DMCtemp)))]
					if (length(cuantos) > mismatches) {
						liniers <- paste(attributes(datos)$names[i],
									datiles$Species[j], 
									paste("[", cuantos, "]", collapse=" "), sep=",")
						logMissing <- rbind(logMissing, liniers)
						write(logMissing, file= MissLogFile)
						next
					} 
				#}
			}
			
			key <- NULL
			totest <- NULL
			for (k in 1:(length(DMC)/2)){
				key <- c(key, DMC[2*k])
				totest <- c(totest, datos[[i]][(as.numeric(DMC[k+(k-1)]))])
			}
			matchi <- (sum((key == toupper(totest)), na.rm = TRUE))
			if (identical(toupper(key),toupper(totest))) {
				pares <- paste("-", datiles$Species[j], "[", matchi, "/", (length(DMC)/2), " {", format(simil, digits = 3), " similar to ", nomina, "}",  "]", sep = "")
				resultado <- paste(resultado, pares, sep = "")
				count <- count + 1
			} else if ( matchi < (length(DMC)/2)    &    matchi >= ((length(DMC)/2)-mismatches) ) {
				pares <- paste("-", datiles$Species[j], "[", matchi, "/", (length(DMC)/2), " {", format(simil, digits = 3), " similar to ", nomina, "}",  "]", sep = "")
				resultado <- paste(resultado, pares, sep = "")
				count <- count + 1
			}	
		}
		if (count == 0) {
			resultado <- "Unidentified"
		}
		palata <- cbind(attributes(datos)$names[i], resultado)
		FinalTable <- rbind(FinalTable, palata)
		colnames(FinalTable) <- c("SpecimenName", "MatchesWith")
		write.csv(FinalTable, file= OutName)
		if (AligName == "none") {
		} else {
			write.fasta(datos, names= nombres, file.out = AligName)	
		}
		cat(if (i == length(datos)) '\n' else '\014')
		#print(as.character(prueba2[1]))
		#print(as.character(prueba2[2]))	
	}
}
```
