## Supplementary material for "An integrative taxonomic treatment of the Mycetophilidae (Diptera: Bibionomorpha) from Singapore reveals 115 new species on 730km^2^": ConfuClass.R

```
## Title: ConfuClass
## Version: 0.1-1
## Date: 2024-05-16
## Author: Ambrosio Torres (Researcher [Ctr. Integr. Biodivers. Discov. - Museum für Naturkunde, Berlin, Germany)
## Depends: R version (>= 4.2.2 )
## Description: function used in the paper to analyze the performance of the DMCs in terms of identification/Discrimination.
## License: GPL (3)
## Usage: ConfuClass("SpeciesListEuroButterflies.csv", "Id_NPM_MolD_EuroButterfliesIter50K_MxLen7.csv", "MissMolD_NPM_EuroButterfliesIter50K_MxLen7.csv", OutConfuFile = "ConfuMatrixMolD_NPM_EuroButterfliesIter50K_MxLen7.csv", OutClassFile= "ClassStatsMolD_NPM_EuroButterfliesIter50K_MxLen7.csv")

ConfuClass <- function(SpList, IdFile, MissFile, OutConfuFile, OutClassFile) {

	especies <- read.csv(SpList, header=F)
	identificando <- read.csv(IdFile, header=T)
	misifu <- read.csv(MissFile, header=T)
	
	identificando$MatchesWith <- gsub("\\{.*?\\}", "", identificando$MatchesWith)
	
	PaRevisar <- NULL
	for (i in 1:dim(misifu)[1]) {
		cat('\014')
		avance <- (i/(dim(misifu)[1]))*100
		print (paste("Evaluating the Missing information. ", format(round(avance, 2), nsmall = 2), "% of progress."))
		for (j in 1:dim(especies)[1]) {
			if (grepl(as.character(especies[j,1]), as.character(misifu$Sequence[i]))) {
				if ((grepl(as.character(especies[j,1]), as.character(misifu$SpeciesDMC[i])))) {
					PaRevisar <- c(PaRevisar, i)
				}
			}
		}				
	}
	if(length(PaRevisar)> 0) {
		Paila <- misifu[PaRevisar,]
		identificando$SpecimenName <- as.character(identificando$SpecimenName)
		lugares <- NULL
		for (i in 1:dim(Paila)[1]) {
			cual <- which(identificando$SpecimenName == as.character(Paila$Sequence[i]))
			identificando$SpecimenName[cual] <- gsub(Paila$SpeciesDMC[i], "", identificando$SpecimenName[cual])
			identificando$MatchesWith[cual] <- gsub(Paila$SpeciesDMC[i], "", identificando$MatchesWith[cual])
			lugares <- c(lugares, cual)
		}
	}
	
	matera <- NULL
	superindice <- NULL
	for (i in 1: (dim(especies)[1] + 1) ) {
		if (i == 1) {
			cat('\014')
			print ("Building the Confusion Matrix. 0 % of progress.")
		}
		
		indice <- grep(especies[i,1], identificando$SpecimenName)
		if (i < (dim(especies)[1] + 1) ) {
			superindice <- c(superindice, indice)
			if (length(indice) < 1) {
				linea <- rep(0, (dim(especies)[1] + 1) )
				matera <- rbind(matera, linea)
				next
			} else {
				submatriz <- identificando[indice,]
				linea <- rep(0, (dim(especies)[1] + 1) )
				for (j in 1:dim(submatriz)[1] ) {
					ide <- 0
					for (k in 1:dim(especies)[1] ) {
						if ( grepl(especies[k,1], submatriz$MatchesWith[j]) ) {
							linea[k] <- linea[k] + 1
							ide <- ide + 1
						} 
					}
					if (ide < 1) {
						linea[length(linea)] <- linea[length(linea)] + 1
					}	
				}
				matera <- rbind(matera, linea)
			}
		} else {
			superindice <- unique(superindice)
			if ( length(superindice) < 1) {
				linea <- rep(0, (dim(especies)[1] + 1) )
				matera <- rbind(matera, linea)
				next
			} else {
				submatriz <- identificando[-(superindice),]
				linea <- rep(0, (dim(especies)[1] + 1) )
				for ( j in 1:dim(submatriz)[1] ) {
					ide <- 0
					for (k in 1:dim(especies)[1] ) {
						if ( grepl(especies[k,1], submatriz$MatchesWith[j]) ) {
							linea[k] <- linea[k] + 1
							ide <- ide + 1
						}
					}
					if (ide < 1) {
						linea[length(linea)] <- linea[length(linea)] + 1
					}
				}
				matera <- rbind(matera, linea)
			}
		}
		cat('\014')
		avance <- (i/(dim(especies)[1] + 1))*100
		print (paste("Building the Confusion Matrix. ", avance, "% of progress."))
	}
	matera <- data.frame(matera)
	colnames(matera) <- c(as.character(especies[,1]), "Unidentified")
	rownames(matera) <- c(as.character(especies[,1]), "Unidentified")
	write.csv(matera, file= OutConfuFile)
	
	matera <- read.csv(OutConfuFile, header=T)	
	matera <- matera[,-c(1)] 
	Tabletilla <- NULL
	Nombrados <- 0
	for (i in 1:dim(matera)[1] ) {
		TruePos <- matera[i,i]
		
		if (i == dim(matera)[1]) {
			FalseNeg <- (dim(identificando)[1] - Nombrados) - TruePos
		} else {
			EnLaEspecie <- length(grep(colnames(matera)[i], identificando$SpecimenName))
			Nombrados <- Nombrados + EnLaEspecie
			FalseNeg <- EnLaEspecie - TruePos
		}
		FalsePos <- sum(matera[-c(i),i])
		TrueNeg <- dim(identificando)[1] - (TruePos + FalsePos + FalseNeg)
	
	
		Precision <- (TruePos) / (TruePos + FalsePos)
		Precision <- signif(Precision, digits =5)
		if (is.nan(Precision)) {Precision <- 0}
		if (TruePos == 0 & FalseNeg == 0) {
			Precision <- NA
		}
		
		Recall <- (TruePos) / (TruePos + FalseNeg)
		Recall <- signif(Recall, digits =5)
		if (TruePos == 0 & FalseNeg == 0) {
			Recall <- NA
		}
		
		Specificity <- (TrueNeg) / (TrueNeg + FalsePos)
		Specificity <- signif(Specificity, digits =5)
		
		if (is.na(Recall)) {
			Accuracy <- NA
		} else {
			Accuracy <- (Recall + Specificity) / 2
		}
		Accuracy <- signif(Accuracy, digits =5)	
		
		if (is.na(Recall) & is.na(Precision)) {
			F1Score <- NA
		} else {
			F1Score <- 2 * ((Precision*Recall) / (Precision+Recall))
		}
		if (is.nan(F1Score)) {F1Score <- 0}
		F1Score <- signif(F1Score, digits =5)	
	
		Linea <- c(Precision, Recall, Specificity, Accuracy, F1Score)
		Tabletilla <- rbind(Tabletilla, Linea)
	}
	Tabletilla <- data.frame(Tabletilla)
	rownames(Tabletilla) <- colnames(matera)
	colnames(Tabletilla) <- c("Precision", "Recall", "Specificity", "Accuracy", "F1Score")	
	Tabletilla <- rbind(Tabletilla, Average= colMeans(Tabletilla, na.rm=T))
	write.csv(Tabletilla, OutClassFile)	
	
	if(length(PaRevisar)> 0) {
		if (length(lugares) > 0) {
			print(paste("## The following sequences contain missing data conflicting with the positions of the DMCs.",
						"For that reason they were not taking into account in the calculation of the performance of their corresponding DMCs:"))
			print(Paila)
		}
	}		
}
```
