## Supplementary material for "An integrative taxonomic treatment of the Mycetophilidae (Diptera: Bibionomorpha) from Singapore reveals 115 new species on 730km^2^": ScriptMolecularDiagnosis.R

```
## Description: this is the script used to find and evaluate Diagnostic Molecular Combinations (DMCs) for a set of Mycetophilidae new species.
## Date: 2024-05-22
## Author: Ambrosio Torres (Researcher [Ctr. Integr. Biodivers. Discov. - Museum für Naturkunde, Berlin, Germany).
## Environment used: R version (4.3.2); RStudio (2023.12.0+369); Package UITOTO (1.0.0) and its dependencies.
## License: GPL (3)
## Citation: if you use this script please cite the related study (Amorim et al. 2024 [Integrative Systematics, Stuttgart Contributions to Natural History]), 
##			 as well as the UITOTO package and its dependencies (https://zenodo.org/records/11236953 / https://github.com/atorresgalvis/UITOTO). 

##Load the required resources
	library(UITOTO) #Load the UITOTO package (download and installation instructions in: https://github.com/atorresgalvis/UITOTO)
	source("ALnID2.r") #Load the function "ALnID2" (a modified version of the "ALnID" function from UITOTO for this study).
	source("ConfuClass.R") #Load the function "ConfuClass"

##Obtention of DMCs
	OpDMC("Mycetos_dataset1_27mar24.fasta", "MycetoSpeciesList.csv", iter = 50000, MnLen = 3, exclusive = 3, RefStrength = 0.25, OutName = "OpDMC_MycetoExclu3.csv", GapsNew = FALSE)
	OpDMC("Mycetos_dataset1_27mar24.fasta", "MycetoSpeciesList.csv", iter = 50000, MnLen = 4, exclusive = 4, RefStrength = 0.33, OutName = "OpDMC_MycetoExclu4.csv", GapsNew = FALSE)
	OpDMC("Mycetos_dataset1_27mar24.fasta", "MycetoSpeciesList.csv", iter = 50000, MnLen = 5, exclusive = 5, RefStrength = 0.50, OutName = "OpDMC_MycetoExclu5.csv", GapsNew = FALSE)
	OpDMC("Mycetos_dataset1_27mar24.fasta", "MycetoSpeciesList.csv", iter = 50000, MnLen = 10, exclusive = 4, RefStrength = 0.33, OutName = "OpDMC_MycetoExclu4Min10B.csv", GapsNew = FALSE)

	##We obtained a fifth set of DMCs using the most frequent sites presented in the DMCs from the previous four searches																														
	##For this, we followed this set of commands for each Species (we included the results in a file named "DMCsMycetoMostFrecuentSites.csv"):
		#  a <- "138: T, 149: C, 151: T, 169: C, 223: C, 269: A" #DMC from the set of parameters A
		#  b <- "10: C, 26: A, 62: C, 106: C, 138: T, 149: C, 151: T, 236: C, 247: T, 269: A"  #DMC from the set of parameters B
		#  c <- "10: C, 106: C, 149: C, 151: T, 269: A"  #DMC from the set of parameters C
		#  d <- "106: C, 149: C, 151: T, 269: A"  #DMC from the set of parameters D
		#  
		#  a <- strsplit(a, ', ')[[1]]
		#  b <- strsplit(b, ', ')[[1]]
		#  c <- strsplit(c, ', ')[[1]]
		#  d <- strsplit(d, ', ')[[1]]
		#  e <- c(a,b,c,d)
		#  final<- unique(e[duplicated(e)])
		#  paste("[", paste(final, collapse= ", "), "]", sep= "")
		#  	[1] "[10: C, 106: C, 138: T, 149: C, 151: T, 269: A]" #Resultant DMC based on the most frequent sites in the DMCs A, B, C and D
	
##Identification of unknown sequences (testing dataset) using the obtained DMCs. This step is aimed for testing the performance of the DMCs.
	ALnID2("OpDMC_MycetoExclu3.csv", "Mycetos_dataset1_27mar24.fasta", "MycetophilidaeTesting.fasta", mismatches = 1, against= "First", perfectMatch=5, misMatch=0, gapOpening = -14, gapExtension = -2, gapPower = -1, terminalGap = 0, OutName = "Id_UITOTO_NPM_Mycetophilidae3Free.csv", MissLogFile= "MissUITOTO_NPM_Mycetophilidae3Free.csv", AligName = "none")
	ALnID2("OpDMC_MycetoExclu4.csv", "Mycetos_dataset1_27mar24.fasta", "MycetophilidaeTesting.fasta", mismatches = 1, against= "First", perfectMatch=5, misMatch=0, gapOpening = -14, gapExtension = -2, gapPower = -1, terminalGap = 0, OutName = "Id_UITOTO_NPM_Mycetophilidae4Free.csv", MissLogFile= "MissUITOTO_NPM_Mycetophilidae4Free.csv", AligName = "none")
	ALnID2("OpDMC_MycetoExclu5.csv", "Mycetos_dataset1_27mar24.fasta", "MycetophilidaeTesting.fasta", mismatches = 1, against= "First", perfectMatch=5, misMatch=0, gapOpening = -14, gapExtension = -2, gapPower = -1, terminalGap = 0, OutName = "Id_UITOTO_NPM_Mycetophilidae5Free.csv", MissLogFile= "MissUITOTO_NPM_Mycetophilidae5Free.csv", AligName = "none")
	ALnID2("OpDMC_MycetoExclu4Min10B.csv", "Mycetos_dataset1_27mar24.fasta", "MycetophilidaeTesting.fasta", mismatches = 1, against= "First", perfectMatch=5, misMatch=0, gapOpening = -14, gapExtension = -2, gapPower = -1, terminalGap = 0, OutName = "Id_UITOTO_NPM_Mycetophilidae4Min10.csv", MissLogFile= "MissUITOTO_NPM_Mycetophilidae4Min10.csv", AligName = "none")
	ALnID2("DMCsMycetoMostFrecuentSites.csv", "Mycetos_dataset1_27mar24.fasta", "MycetophilidaeTesting.fasta", mismatches = 1, against= "First", perfectMatch=5, misMatch=0, gapOpening = -14, gapExtension = -2, gapPower = -1, terminalGap = 0, OutName = "Id_UITOTO_NPM_MycetophilidaeMostFrecuentSites.csv", MissLogFile= "MissUITOTO_NPM_MycetophilidaeMostFrecuentSites.csv", AligName = "none")	
	
##CrossValidation. Generation of confusion matrices and Clasification Statistics.
	ConfuClass("MycetoSpeciesList.csv", "Id_UITOTO_NPM_Mycetophilidae3Free.csv", "MissUITOTO_NPM_Mycetophilidae3Free.csv", OutConfuFile = "ConfuMatrix_NPM_Mycetophilidae3Free.csv", OutClassFile= "ClassStats_NPM_Mycetophilidae3Free.csv")
	ConfuClass("MycetoSpeciesList.csv", "Id_UITOTO_NPM_Mycetophilidae4Free.csv", "MissUITOTO_NPM_Mycetophilidae4Free.csv", OutConfuFile = "ConfuMatrix_NPM_Mycetophilidae4Free.csv", OutClassFile= "ClassStats_NPM_Mycetophilidae4Free.csv")
	ConfuClass("MycetoSpeciesList.csv", "Id_UITOTO_NPM_Mycetophilidae5Free.csv", "MissUITOTO_NPM_Mycetophilidae5Free.csv", OutConfuFile = "ConfuMatrix_NPM_Mycetophilidae5Free.csv", OutClassFile= "ClassStats_NPM_Mycetophilidae5Free.csv")
	ConfuClass("MycetoSpeciesList.csv", "Id_UITOTO_NPM_Mycetophilidae4Min10.csv", "MissUITOTO_NPM_Mycetophilidae4Min10.csv", OutConfuFile = "ConfuMatrix_NPM_Mycetophilidae4Min10.csv", OutClassFile= "ClassStats_NPM_Mycetophilidae4Min10.csv")
	ConfuClass("MycetoSpeciesList.csv", "Id_UITOTO_NPM_MycetophilidaeMostFrecuentSites.csv", "MissUITOTO_NPM_MycetophilidaeMostFrecuentSites.csv", OutConfuFile = "ConfuMatrix_NPM_MycetophilidaeMostFrecuentSites.csv", OutClassFile= "ClassStats_NPM_MycetophilidaeMostFrecuentSites.csv")

##Evaluation of the CrossValidation results.
		especies <- read.csv("MycetoSpeciesList.csv", header=F)
		especies <- as.character(especies[c(order(especies[,1])),])
	
		DMCs <- c("OpDMC_MycetoExclu3.csv",
			"OpDMC_MycetoExclu4.csv",
			"OpDMC_MycetoExclu5.csv",
			"OpDMC_MycetoExclu4Min10B.csv",
			"DMCsMycetoMostFrecuentSites.csv"
		)
		Archivote <- data.frame(Species = especies)
		for (i in 1:length(DMCs)) {
			archi <- read.csv(DMCs[i], header=T)
			longitudes <- NULL
			for (j in 1:(dim(Archivote)[1])) {
				combi <- as.character(archi$DMC[archi$Species == Archivote$Species[j]])
				len <- lengths(regmatches(combi, gregexpr(":", combi)))
				longitudes <- c(longitudes, len)
			}
			Archivote <- cbind(Archivote, longitudes)
		}
		colnames(Archivote)[2:6] <- c("LengthDMC_Exclu3","LengthDMC_Exclu4","LengthDMC_Exclu5","LengthDMC_Exclu4Min10","LengthDMC_MFS")
	
		performances <- c("ClassStats_NPM_Mycetophilidae3Free.csv",
			"ClassStats_NPM_Mycetophilidae4Free.csv",
			"ClassStats_NPM_Mycetophilidae5Free.csv",
			"ClassStats_NPM_Mycetophilidae4Min10.csv",
			"ClassStats_NPM_MycetophilidaeMostFrecuentSites.csv"
		)
		
		for (med in 2:6) {
			for (i in 1:length(performances)) {
				archi <- read.csv(performances[i], header=T)
				valores <- NULL
				for (j in 1:(dim(Archivote)[1])) {
					submatriz <- data.frame(especie= archi$X, medida = archi[,med])
					valor <- as.numeric(submatriz$medida[as.character(submatriz$especie) == as.character(Archivote$Species[j])])
					valores <- c(valores, valor)
				}
				Archivote <- cbind(Archivote, valores)
			}
		}
		colnames(Archivote)[7:11] <- c("Precision_Exclu3","Precision_Exclu4","Precision_Exclu5","Precision_Exclu4Min10","Precision_MFS")
		colnames(Archivote)[12:16] <- c("Recall_Exclu3","Recall_Exclu4","Recall_Exclu5","Recall_Exclu4Min10","Recall_MFS")
		colnames(Archivote)[17:21] <- c("Specificity_Exclu3","Specificity_Exclu4","Specificity_Exclu5","Specificity_Exclu4Min10","Specificity_MFS")
		colnames(Archivote)[22:26] <- c("Accuracy_Exclu3","Accuracy_Exclu4","Accuracy_Exclu5","Accuracy_Exclu4Min10","Accuracy_MFS")
		colnames(Archivote)[27:31] <- c("F1Score_Exclu3","F1Score_Exclu4","F1Score_Exclu5","F1Score_Exclu4Min10","F1Score_MFS")
	
		write.csv(Archivote, file= "TotalClassificationStatisticsMycetophilidae.csv")
		datos<- read.csv("TotalClassificationStatisticsMycetophilidae.csv", header=T)
		datos <- datos[,-1]
		sumatorias <- NULL
		for (i in 1:dim(datos)[1]) {
			a <- sum(datos[i,5:19], na.rm = T)
			sumatorias <- c(sumatorias, a)
		}
		sumatorias <- order(sumatorias, decreasing = T)
		
		datos <- datos[sumatorias,]
		write.csv(datos, file= "TotalClassificationStatisticsMycetophilidae.csv")	
		
##Evaluate the performance of the Final selected DMCs
	ALnID2("DMCsMycetoFinal.csv", "Mycetos_dataset1_27mar24.fasta", "MycetophilidaeTesting.fasta", mismatches = 1, against= "First", perfectMatch=5, misMatch=0, gapOpening = -14, gapExtension = -2, gapPower = -1, terminalGap = 0, OutName = "Id_FinalNPM_First.csv", MissLogFile= "MissFinalNPM_First.csv", AligName = "none")
	ConfuClass("MycetoSpeciesList.csv", "Id_FinalNPM_First.csv", "MissFinalNPM_First.csv", OutConfuFile = "ConfuMatrix_NPM_MycetophilidaeFinalNPM_First.csv", OutClassFile= "ClassStats_NPM_MycetophilidaeFinalNPM_First.csv")
```
